# A Predictive Energy Landscape Model of Metamorphic Protein Conformational Specificity

**DOI:** 10.1101/2021.11.16.468851

**Authors:** James O. Wrabl, Keila Voortman-Sheetz, Vincent J. Hilser

## Abstract

“Metamorphic” proteins challenge state-of-the-art structure prediction methods reliant on amino acid similarity. Unfortunately, this obviates a more effective thermodynamic approach necessary to properly evaluate the impact of amino acid changes on the stability of two different folds. A vital capability of such a thermodynamic approach would be the quantification of the free energy differences between 1) the energy landscape minima of each native fold, and 2) each fold and the denatured state. Here we develop an energetic framework for conformational specificity, based on an ensemble description of protein thermodynamics. This energetic framework was able to successfully recapitulate the structures of high-identity enginerered sequences experimentally shown to adopt either *Streptococcus* protein G_A_ or G_B_ folds, demonstrating that this approach indeed reflected the energetic determinants of fold. Residue-level decomposition of the conformational specificity suggested several testable hypotheses, notably among them that fold-switching could be affected by local de-stabilization of the populated fold at positions sensitive to equilibrium perturbation. Since this ensemble-based compatibility framework is applicable to any structure and any sequence, it may be practically useful for the future targeted design, or large-scale proteomic detection, of novel metamorphic proteins.

**Impact Statement:** Metamorphic proteins are single amino acid sequences capable of adopting more than one structure at equilibrium. Detection and design of these molecules hold great promise for biological understanding and materials engineering, but to do so requires a thermodynamic framework capable of estimating the free energy differences between the two structures and the denatured state. We present such a framework, show it to be effective for the well-studied metamorphic protein G_A_/G_B_ system, and suggest testable hypotheses for engineering novel fold-switch proteins.

## Introduction

Metamorphic proteins [1–3], capable of more than one distinct tertiary structure, are among the most interesting examples of biological rule-bending. Traditionally, the central dogma asserted that one linear sequence of amino acids was only capable of forming one final, functional structure. Recent exploration and characterization of the natural proteomes has revealed an alternative scheme: a small population of structural dissentients capable of more than one complete and functional fold. Examples that have been described thus far include XCL1 (lymphotactin) [4], CLIC1 [5], RfaH-CTD [6], KaiB/C [7], and MAD2 [8]. Altogether, it has been estimated that up to 4% of the PDB (nearly 6500 proteins) may display this paradigm-changing behavior [9].

Metamorphic proteins are of particular interest in the field of protein folding because they provide an exceptional window into the evolutionary, chemical, and physical forces responsible for the relationship between primary sequence and tertiary structure. Distinct from proteins that undergo minor conformational change, a metamorphic protein populates two completely different tertiary structures under different temperature, pH, or oxidative conditions and may provide insight into the determinants of the physical and chemical interactions within the primary sequence that drive folding and binding activity. Additionally, investigation into the genetic history of these fold-switching proteins has contributed to the understanding of natural gain-of-function events in protein evolution [10–12].

Metamorphic protein design efforts have been ongoing for more than thirty years [13, 14]. Among the few engineered metamorphic proteins that exist [15–18], one of the most useful examples for benchmarking predictive approaches has been the design and structural characterization of an extensive series of small, high sequence identity proteins based on domains isolated from *Streptococcus* protein G [19]. This process culminated in several sequences where as little as one amino acid change resulted in a complete fold and function switch between two tertiary folds, G_A_ and G_B_. Including intermediate mutations, the entire sequence space separating the two terminal folds was reduced to variability in only 13 out of the proteins’ 56 positions, which was further restricted by choosing between one of two possibilities at each position. Thus, a “binary sequence space” was experimentally defined [19] that localized the conformational specificity to 2^13^ = 8,192 sequence permutations (as compared to a theoretical 20^56^ + 19^56^ ~ 10^73^, expected if the two folds had no sequence identity).

Such a massive reduction in sequence space for this G_A_/G_B_ system has motivated efforts from many laboratories to understand the biophysical determinants of metamorphic proteins, to engineer novel fold-switching proteins, and to search for additional naturally occurring examples. Predictive applications, with varying degrees of success, have ranged from primary sequence analysis [20], secondary structure prediction [21, 22], and all-atom molecular dynamics simulation [23–25].

Still elusive, however, is a unified description of metamorphic protein behavior from an energy landscape perspective. This knowledge gap is not trivial, even if heuristic prediction algorithms were error-free. The energy landscape description is crucial for a complete understanding because metamorphic behavior is a chemical equilibrium between the denatured state and multiple folded states. To date, the small number of reported metamorphic proteins are only able to be documented because they exhibit simultaneous populations of both folds at equilibrium.

Unfortunately, such coupled equilibria, at least for molecules as large as globular proteins, are at or beyond the limits of state-of-the-art all-atom simulations [26–29]. Moreover, physico-chemical processes are simply ignored in heuristic approaches based on sequence or secondary structure, and recent extraordinary deep-learning success in protein folding does not transfer to metamorphic protein sequences [30]. To our knowledge, current predictive methods applied to metamorphic proteins must make unfavorable compromises between efficacy, applicability, computational efficiency, and degree of insight.

To address these shortcomings and gain predictive understanding into conformational specificity, we introduce an ensemble-based thermodynamic description of proteins and apply it to the energy landscape of a metamorphic protein, specifically the G_A_/G_B_ system. Although this ensemble-based description was previously benchmarked against folded proteins with a single energy minimum, we demonstrate here that it is also accurate when applied to the double-well G_A_/G_B_ system. In particular, a scoring framework is developed to relate the difference in free energies of one sequence adopting either of two distinct folds (*i.e.* the free energy of conformational specificity) to the thermodynamic propensities of sequence for structure obtained from a large database of natural proteins. We further show that our scoring framework is singularly effective at identifying which fold is populated for a given sequence amongst the plethora of high-identity sequences within the binary space. Regional analysis of the determinants of conformational specificity reveals that not all parts of the folded proteins contribute equally to fold-switching behavior and suggests testable design approaches to influence the equilibrium populations.

## Results

### An energy landscape view of conformational specificity

Decades of investigation into protein folding and stability have converged on the energy landscape theory, summarized in Figure 1 [31–33]. In this descriptive formalism, a protein folding reaction is treated as a chemical equilibrium between native and denatured states, where the native state of the protein is populated at a thermodynamic minimum of free energy (Figure 1, left bottom). Attention has been primarily focused on the energy well (Fold A), but also necessary to complete the energy landscape are the fully unfolded state (U) and the higher energy conformations of vanishingly populated, but nonetheless posssible, alternatively folded states (*e.g.* Fold B). Possible to locate in the energy landscape, but difficult to experimentally measure, is the conformational specificity [34] of amino acid sequence for structure, defined as the free energy difference between two alternatively folded states given a single sequence (*i.e.* ΔG_AB_).

**Figure 1.**
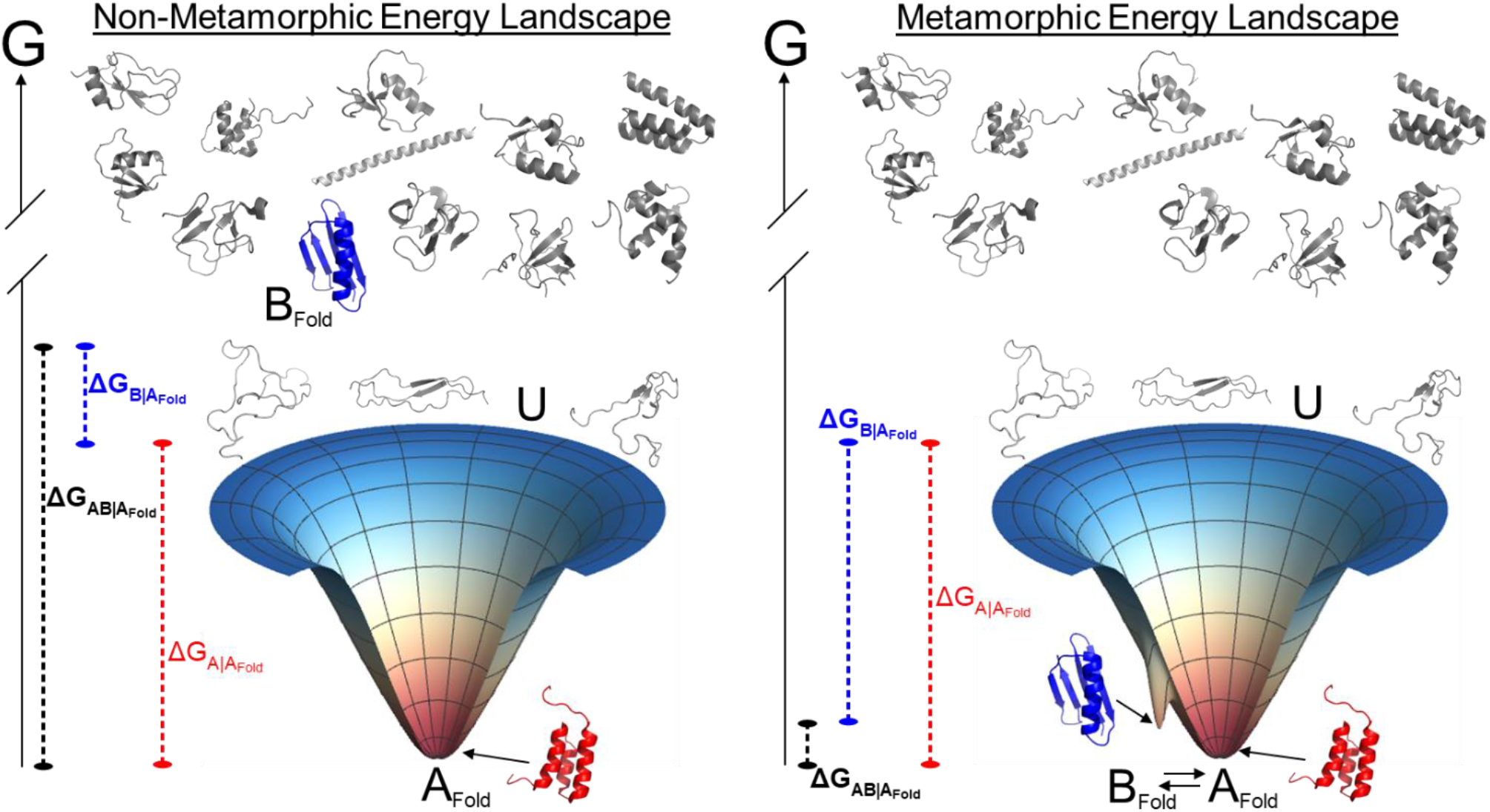
The free energy landscapes of non-metamorphic *versus* metamorphic proteins. The funnel on the left side depicts an idealized three-dimensional reaction coordinate of a globular alpha-helical protein, A, at a single energy minimum. The funnel on the right side depicts protein A as the predominant species with a competing beta-sheet protein, B, populated to a smaller extent at equilibrium. U stands for the unfolded state, occupying an intermediate free energy level (G) as compared with a variety of higher energy competing folds (grayscale cartoons). Given the energy landscape of fold A, the free energy difference between amino acid sequence A adopting fold A and sequence A adopting the unfolded state is ΔG_A|A_. ΔG_AB|A_ is the free energy difference between sequence A adopting fold A or fold B, given the energy landscape of fold A. If fold A is stable, then knowledge of ΔG_AB|A_ is useful in predicting whether sequence A is a metamorphic protein or not. Thus, ΔG_A|A_ is a *stability* measurement, while ΔG_AB|A_ is a *conformational specificity* measurement.

The energy landscape of a metamorphic protein introduces complexity due to the population of a second fold (Figure 1, right). In particular, the conformational specificity between Folds A and B has decreased to a point where both folds have become stabilized relative to the unfolded state. In this situation, because both folds are located within a common energy landscape, knowledge of the free energies of stability of both folds relative to the unfolded state is sufficient to determine the conformational specificity of sequence A for Fold A over Fold B. Thus, we set out to develop an approach that will provide a measure of the energy of each fold relative to the denatured state, providing implicitly a means of comparing the energy difference between the native folds.

We employed *COREX,* an established statistical thermodynamic framework developed in our lab over the last two decades, which is based on an ensemble description of protein structural thermodynamics [35–37]. As we have already shown, this approximation to the energy landscape, when applied to a database of diverse proteins, has resulted in an effective fold recognition method that matches one sequence to one structure [38–41], given the energy landscape of the single structure in question.

Within this framework, a testable hypothesis arises as to whether a metamorphic amino acid sequence could be simultaneously compatible with more than one energy landscape (Figure 2). In this scenario, multiple extremely high identity sequences, capable of simultaneously populating both structures, would exhibit high fold recognition scores for both folds. When scored against only one of the two folds (Figure 2, right side) these special, nearly identical, sequences would nonetheless achieve high scores to the exclusion of sequences adopting alternative folds (Figure 2, right bottom). Since the alternative folds are not measurably populated under experimental conditions, their free energies must be less than or equal to that of the unfolded state (Figure 2, left top). Note that this scoring system need not possess a zero point coincident with the unfolded state to be able to reflect a conformational specificity (Figure 2, ΔG|_αβ_).

**Figure 2.**
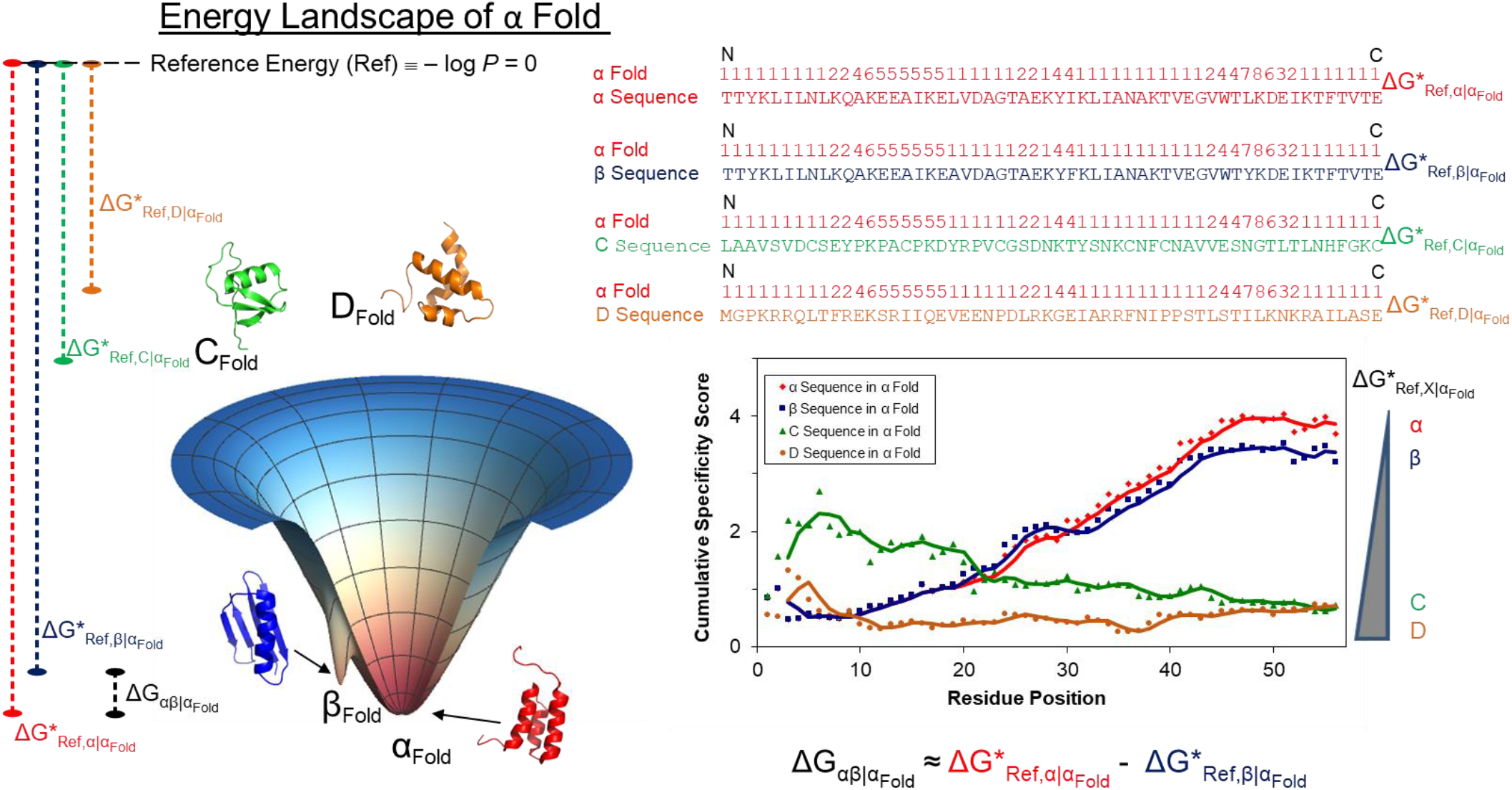
Ensemble-based energetic framework for conformational specificity. A *COREX* ensemble generated for α fold (red) results in a simulated energy landscape (funnel) and a set of Thermodynamic Environments for each residue in α fold (red numbers 1-8, top right). As described in Methods, mapping amino acid sequences for folds α, β, C, D onto the aligned environments results in sequence:environment scores accumulating from the N to C termini of each protein (red, blue, green, orange sequences), with the total score approximating a ΔG (ΔG*)with respect to a Reference Energy (Ref). Since the total score is highest for the correct sequence in the correct fold (chart), it is assumed that the total score can generally represent a conformational specificity of an arbitrary sequence for an arbitrary fold. Thus, the difference in scores between two sequences for the same environments (equation) could approximate the free energy difference between two folds, if the two sequences in question were known to adopt two different folds.

Interrogating this hypothesis using our ensemble-based scoring framework, we find that metamorphic, or nearly metamorphic, amino acid sequences are compatible with both structures of the G_A_95/G_B_95 metamorphic pair (Figure 3). Out of a library of high-identity engineered sequences, where the sequence identity between any two pairs is at least 77%, sequences experimentally determined to adopt the G_A_95 structure cluster in a distinct region (red circle), as compared to sequences adopting the G_B_95 fold (blue circle). The cluster separation between the red and blue populations is statistically significant (Supplementary Material). Adding confidence to the interpretation that the scoring is indeed reflecting conformational specificity of metamorphic proteins, amino acid sequences of natural homologs to Protein G (the G_B_95 fold) score highly when threaded onto the G_B_95 fold but score poorly when threaded onto the G_A_95 fold (Figure 3A, black dashed circle).

**Figure 3.**
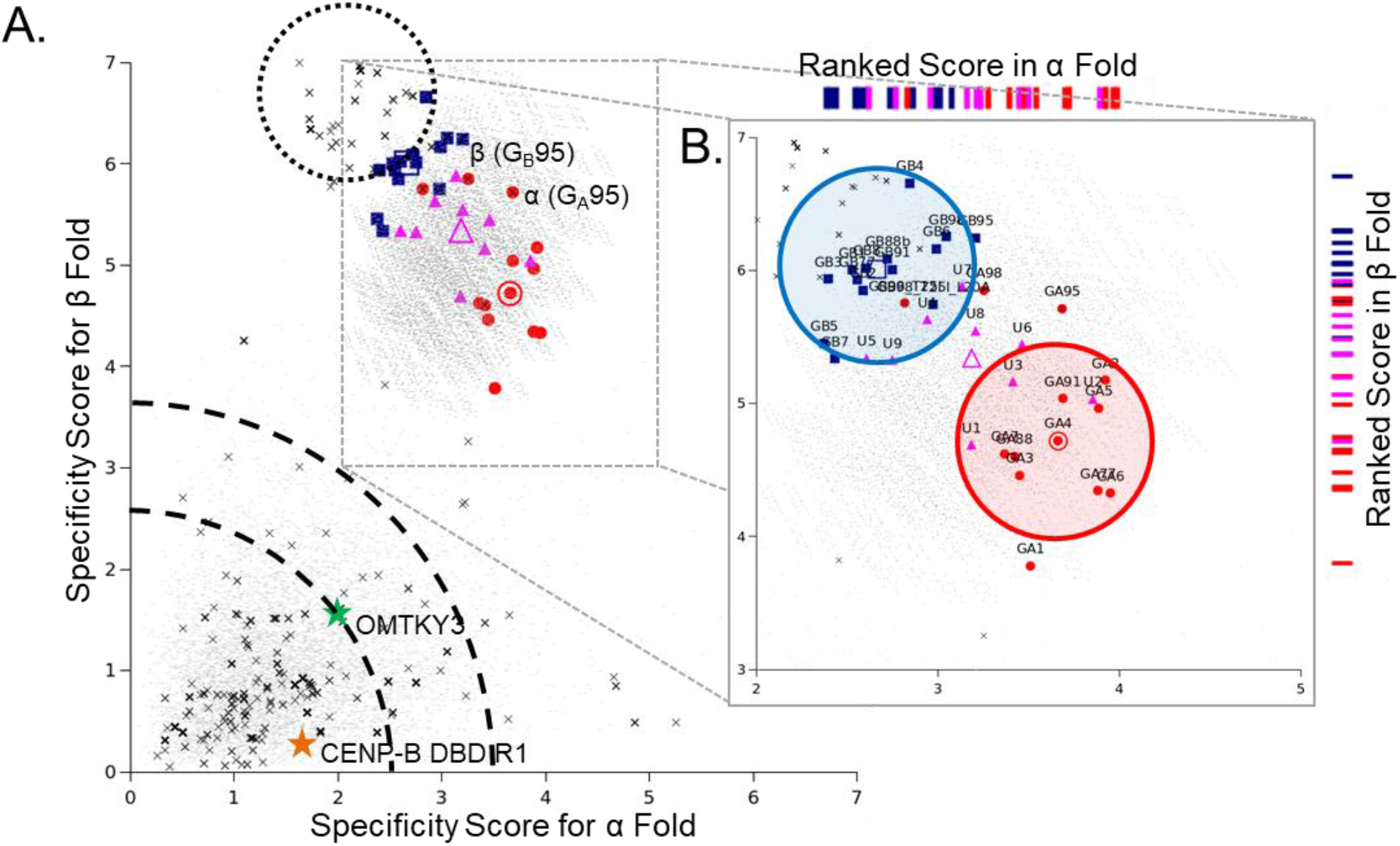
Ensemble-based energetic framework recapitulates experimental fold-switch behavior of high-identity amino acid sequences. **A.** Each point on the plot represents a total score between one amino acid sequence and either the Thermodynamic Environments of α fold (G_A_95, 2kdl; x-axis) or the Thermodynamic Environments of β fold (G_B_95, 2kdm; y-axis). Red points are engineered sequences experimentally reported to adopt the α fold, Blue points are sequences reported to adopt the β fold, and Pink points are reported to misfold. Dark Gray dashes represent 8192 sequences of high-identity space (Table S8), as described [19]; all heretofore mentioned sequences on the plot exhibit 77% - 99% identity with both G_A_95 and G_B_95. Black X points represent structured sequences of 56 residues found in the PDB (Table S7). Green and Orange stars represent specific folds (1ovo and 1bw6) that are neither G_A_95 nor G_B_95, discussed in the text. Light gray dashes represent random amino acid sequences (Table S9). Median scores for experimental α and β sequences are denoted by large open symbols. The dashed circle encompasses structured sequences adopting the β fold that have less than 77% identity to G_B_95. The probability of obtaining such clusters at random is approximately *p* < 0.02 (Supplementary Material). Dashed arcs represent an empirical gap between random sequences and high-identity metamorphic sequence space. **B.** Enlarged area (inset) focusing on high-identity sequence space as described [19]. Individual proteins from that work are labeled. Open symbols represent median scores for each fold, clearly showing that experimental alpha fold sequences exhibit a higher conformational specificity for the α fold than do β sequences, and vice-versa. Red and blue circles represent arbitrary areas centered on median scores for experimental α and β sequences. Conformational specificity scores are projected onto each axis to demonstrate the individual rankings of each sequence in each fold.

### The relationship between specificity score and the energy landscape

Several lines of evidence suggest that the scoring of sequences in each fold (Fig. 3) is related to its location in the energy landscape (Fig. 1). Primarily, the locations of non-homologous structured and random amino acid sequences in Figure 3A are significantly different. Both groups generally exhibit significances of greater than 0.01 (*i.e.* – log P-value < 2). However, a scoring gap exists on both axes between this value and the lowest scoring experimentally folded sequences – although variable, depending on the fold, significances between 2.5 and 3.5 (dashed arcs Figure 3A) exhibit comparatively fewer points as compared to significances between 1 and 2. This observation suggests an inverse correspondence between the folding funnel energy landscape and the ensemble-based scoring framework: alternative non-populated folds achieve scores of approximately 1-2, populated folds achieve scores greater than approximately 4, and the intervening region could indicate the location of the unfolded state. Thus, Figure 3A can be interpreted as a visualization of features of two energy landscapes that are normally invisible to experimental measurement, highlighting a special region of sequence space that simultaneously exhibits increased specificity to more than one fold.

Additonally, focusing on the region of Figure 3A where experimentally determined sequences are located (Figure 3B), several observations are noteworthy. First, it is clear that each compatibility axis can be independently considered, since the projected rankings from each axis comport with experimental data in isolation. Second, considering the average positions of experimental alpha and beta fold proteins (large open points), a population of points (pink) lies intermediate to the red and blue clouds; these are classified as “unfolded” [19]. Third, two red points, G_A_98 and G_B_98_T25I [10], are conspicuously separated from the preponderance of experimental alpha fold proteins and appear to be located more within the cloud of beta points.

### Sequence determinants of metamorphic conformational specificity

Why do the sequences cluster as they do? One advantage of the COREX statistics is that the total scores can be decomposed into residue-level contributions. When this is done for the parent proteins G_A_95 and G_B_95 (Figure 4), several insights can be gleaned into the determinants of conformational specificity. First, almost all structured regions for both folds make positive contributions to the scoring, contrasting with the negative scores of the unstructured termini in the G_A_95 protein (Figure 4A, top). Second, the three amino acid sequence changes between the two proteins (positions 20, 30 and 45), exhibit scoring differences with respect to the alternative fold that extend beyond the residue position of the change (Figure 4, top, differences between red / blue lines, and bottom cartoons). For two out of the three positions (30 and 45), the difference is enough to drive the cumulative score in the correct direction of the observed conformational specificity (Figure 4, middle panels). Position 20 represents an interesting exception, as its score is neutral between the two folds in G_B_95 and actually scores unfavorably in the G_A_95 sequence:environments pair.

**Figure 4.**
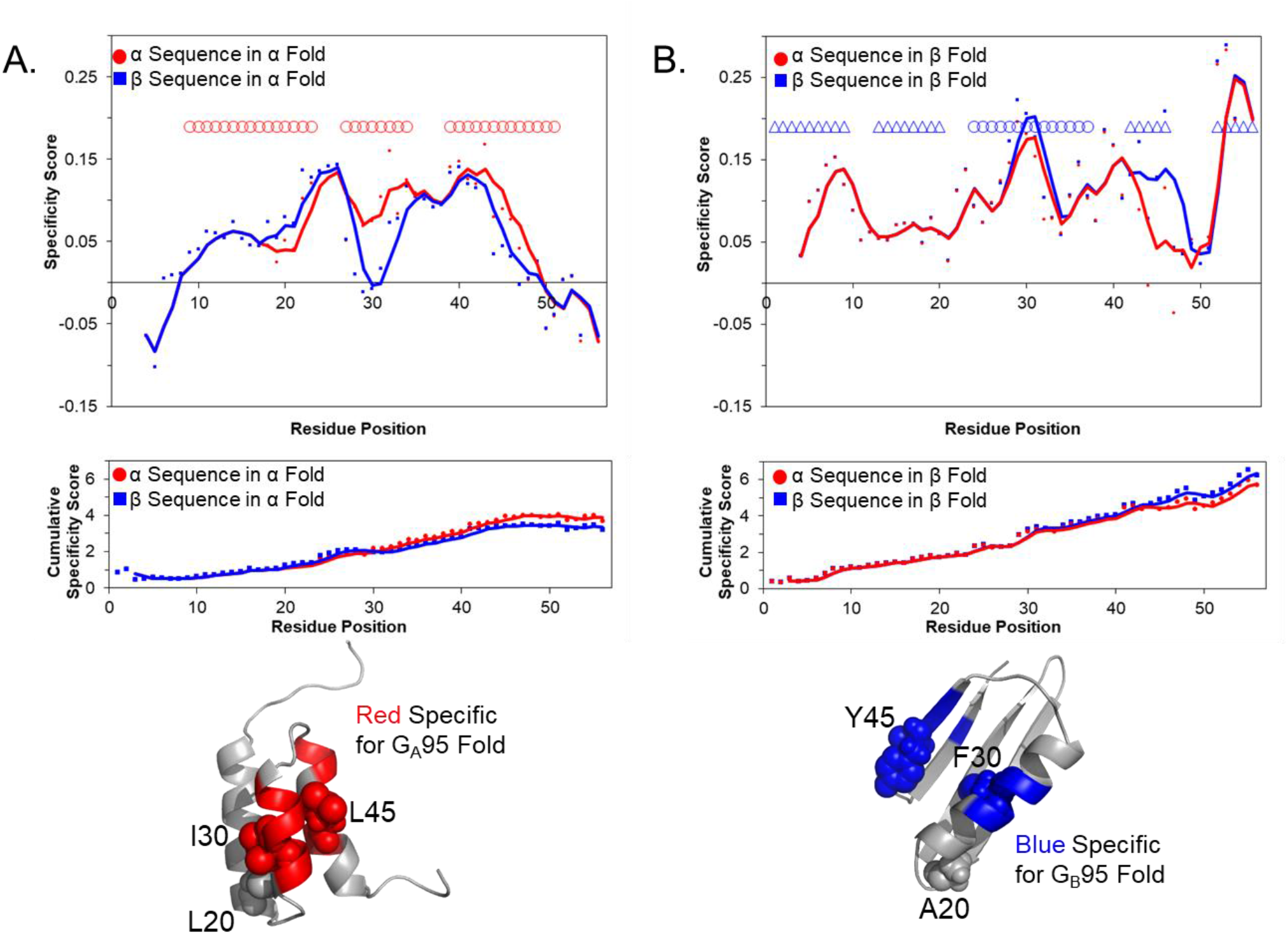
Regional contributions (top) to cumulative conformational specificity (bottom) emphasize non-uniform effects and the importance of the denatured state. The lower plots were obtained by cumulatively evaluating Equations 1 and 2 at every residue position, with a guiding line drawn by averaging every five residues. The upper plots were obtained by decomposing the data in the lower plots, subtracting from every residue the previous, averaging over a centered five residue window, with a guiding line drawn by further averaging nearest neighbors. **A.** Scoring in the G_A_95 fold (2kdl) environments reveals that the C-terminal half of the protein contributes more to the score, and to the specificity, than does the N-terminal half. Positions in the neighborhood of amino acid substitutions 30 and 45 drive the specificity between G_A_95 and G_B_95, with position 20 contributing a lesser amount. **B**. Scoring in the G_B_95 fold (2kdm) environments reveals that the C-terminal half of the protein contributes more to the score than does the N-terminal half, similarly with positions 30 and 45 contributing more than position 20. **C.** Cartoon figures for G_A_95 fold (top) and G_B_95 fold (bottom) demonstrate the propagation of conformational specificity in the neighborhood of mutation sites 20, 30 and 45 (labeled). Colored regions indicate residue positions exhibiting increased scoring in one fold over the other, conferring specificity for the populated fold.

When the scoring changes are averaged across all experimentally investigated sequence variants in this system, the determinants in Figure 4 are generally present among all variants (Supplementary Material, Figure S1). Thus, these calculations suggest that, despite the background of high sequence identity, the entire protein contributes to the conformational specificity of both folds simultaneously, and the engineered positions serve simply to tip the energetic balance one way or another.

This observation is particularly important as it could inform protein design efforts. We hypothesize that the positions that tip the energetic balance can be deduced from the energy landscape of the protein. Specifically, the case of G_A_95/G_B_95 suggests that the positions influencing the conformational specificity scoring are located in aligned regions of the two folds’ stability profiles exhibiting stability mismatch (Figure 5). Two of the three substitutions determining the fold switch involve a high-stability region of one fold aligned with a low-stability region of the other (Figure 5, green boxes). Mapping all such regions in the G_A_95/G_B_95 system (Figure 5 cartoons) suggests that large portions of the proteins are amenable to designed fold-switching, notably the N and C-termini, previously identified in the original report. If true, what the substitution does is to de-populate the high-stability fold by de-stabilization at that position. Since the alternative fold is presumably close in energy, if the de-stabilization is large enough, the alternative fold will be populated. The great advantage of the ensemble-based scoring is that estimation of the degree of destabilization is possible, providing position-specific control of mutation effects.

**Figure 5.**
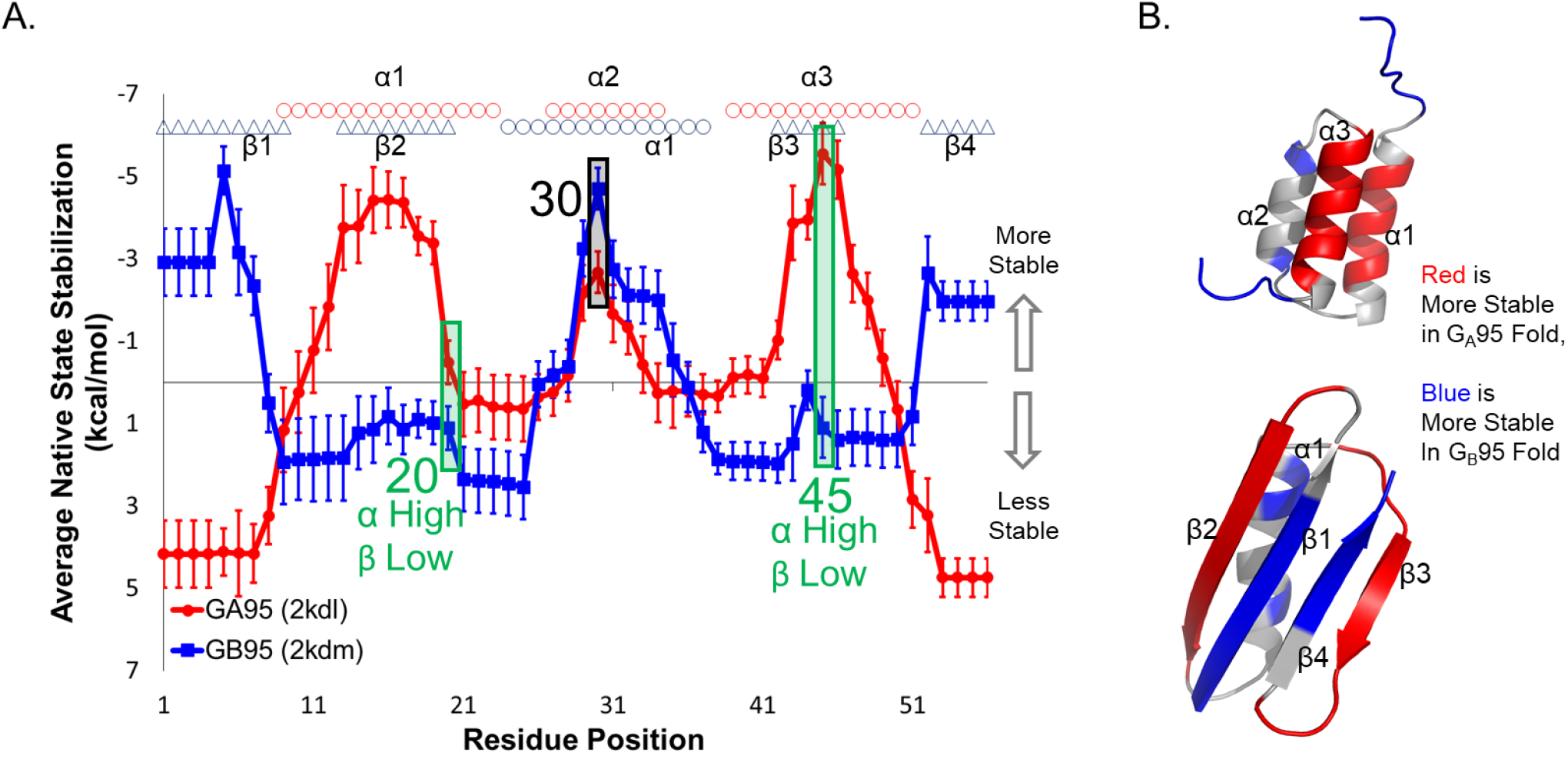
Ensemble-based modeling predicts sensitive regions for fold-switch perturbation, targeting conformational specificity. **A.** *COREX* stability profiles for G_A_95 (red) and G_B_95 (blue) suggest that two out the three amino acid substitutions occur at positions where high stability (“High”) in one fold is juxtaposed with low stability (“Low”) in the other (*i.e.* positions 20 and 45, green). In this plot, the average stability of each profile is set to zero. **B.** Red and blue colors in the molecular cartoons indicate design positions in both folds, where high stability in the colored fold is juxtaposed with low stability in the alternative fold. Red color denotes regions of high stability in the G_A_95 fold juxtaposed with low stability in the G_B_95 fold, and blue denotes regions of high stability in the G_B_95 fold juxtaposed with low stability in the G_A_95 fold. Note that the N- and C-termini of the G_B_95 fold (β1 and β4) are particularly amenable to fold-switching.

## Discussion

How can a single amino acid chain adopt two different structures? The ultimate answer to this puzzle lies in the shape and the depth of the two local minima in the free energy landscape of the protein. Although possible, all-atom modeling of complex landscapes push the limits of state-of-the-art molecular simulation, and few solutions have been achieved. Here we explore a simpler approach to mapping the energy landscape of an engineered metamorphic protein, based on thermodynamic environments derived from simplified protein ensembles of the two structures involved in the equilibrium.

Our approach uses a scoring algorithm that is based on the frequency of finding certain amino acids in different thermodynamic environments within folded proteins, as sampled across a database of protein ensembles, much in the same way that structure prediction algorithms’ scores are based on how often amino acids are found in particular structural environments. Although this approach may at first appear too simplistic, precedence may be found in our earlier work [42] as well as more recently in the impactful work of Stormo and colleagues [43, 44] who find that the energetics of protein-DNA interactions are well captured by similar log-odds probabilistic models. In the case of DNA, the linear, non-cooperative nature of transcription factor binding sites naturally permit an additive, independent-site quantification of free energy of the system. In the case of proteins, clustering of thermodynamic environments by relative stability removes most of the cooperativity, converting sequence-structure specificity into a linear, additive, independent-site system (Tables S4 and S5).

Another advantage of the thermodynamic characterization is that energy is a continuous function, while structural characterics of proteins (*i.e.* helix, sheet, turn, coil) share no objective quantifiable metrics [38, 39]. Indeed, as the structure is ignored in our approach, the resulting frequencies can be universally applied providing quantitative differences both within and between fold families and classes. If this is the case, our approach provides a unique opportunity to dissect the fold switch mechanism in energetic terms and understand more completely how this phenomenon is possible.

The main findings from this simpler energetic approach are presented in Figure 3. Several prominent features of an inverted textbook energy landscape, mapped to ranked conformational specificity scores along each axis, are reflected in these data. First, considering each structure separately, correctly folded sequences are ranked most highly along each axis, while other sequences corresponding to incorrect specific conformations are ranked low. Second, an obvious gap buffers these two categories of sequence, possibly indicating the location of nonspecific denatured conformations. Third, correctly folded sequences lie within a cloud of experimentally determined “binary sequence space”, which contains all permutations of the possible residue pairs at the positions most responsible for conformational specificilty within this metamorphic protein system. Most importantly, this space is unique, relative to randomly generated sequences and alternative folds, because it is ranked highly *along both axes simultaneously*. Thus, we agree with the original authors’ interpretation [19], that this subset of 13 residue positions contains key conformational specificity information; meaning whether an amino acid sequence adopts fold A or fold B (*i.e.* positive design), but also that it does not adopt fold C or fold D (*i.e.* negative design). If true, this logic could be used prospectively with other structures to potentially identify novel fold-switch proteins, searching for one sequence’s scores against any two structures that are substantially higher than random.

### Relationship between specificity scores and global stability

Because the rankings in the current system appear to map to an inverted energy landscape, it may be the case that the conformational specificity scores are quantitatively related to the experimental stabilities of the individual proteins. To test this hypothesis, we plotted the Figure 3 scores against the experimental free energies of folding as reported previously (Figure 6). Remarkably, positive correlations were observed between scores and stabilities for each fold (Figure 6A); a higher score implies a higher stability. The intriguing implication is that stability and specificity, rather than being separate phenomena [34], are actually both manifestations of Gibbs free energy of stability. Thus, we see no mystery in the specific native protein conformation existing once in a large population of molecules in 6M urea to the exclusion of all other folds. Rather, this is a simple consequence of the native conformation being relatively more stable, and relatively more populated, than all other folds under denaturing conditions.

**Figure 6.**
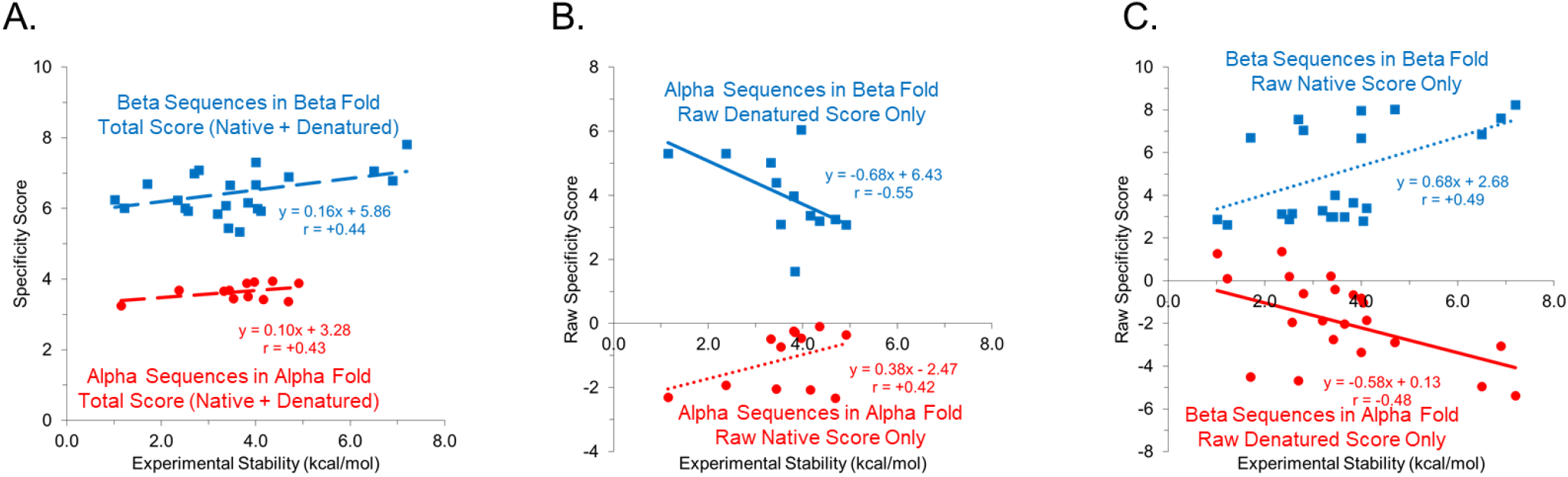
Correlation between conformational specificity scores and experimental stabilities of high-identity proteins. In all panels, red circles represent alpha sequences scored in alpha fold and blue squares represent beta sequences scored in beta fold. Lines represent Pearson correlations with indicated coefficients, *r*. Experimental Stability values are given in Table S6. **A. Conformational Specificity Score (Equation 3) *vs.* Experimental Stability (Equation 4).** Both correlations are positive but not significantly different from zero (*p* < 0.10). Positive slopes suggest that a higher score is related to a higher experimental stability. **B. Native or Denatured Raw Score contributions to Equation 3 involving alpha sequences.** Alpha sequences scored in the beta fold exhibit a negative and significant (*p* < 0.01) correlation between raw denatured score and alpha fold stability. This suggests that the alternative (*i.e.* beta) fold is designed out of the energy landscape for sequences that ultimately adopt the alpha fold. **C. Native or Denatured Raw Score contributions to Equation 2 involving beta sequences.** Likewise, beta sequences scored in the alpha fold exhibit a negative and significant (*p* < 0.01) correlation between raw denatured score and beta fold stability. This suggests that the alternative (*i.e.* alpha) fold is designed out of the energy landscape for sequences that ultimately adopt the beta fold. In both panels B and C, positive and significant (*p* < 0.05) correlations involving raw native state scores suggest that the correct fold is stabilized by the correct sequence.

### Negative design and the denatured state

Perhaps one of the most interesting findings in this study involves the denatured state. As we showed previously [40, 41], just as it is possible to score a sequence in the thermodynamic profile of the native state of a protein, it is also possible to score a sequence in the denatured state, thus providing a vehicle for evaluating the impact of mutations on the native versus the denatured states of each fold. These impacts are shown in Figure 6A for the series of high-identity proteins in the G_A_/G_B_ system.

We find that for both folds in this metamorphic system, native contributions are positively correlated with the stability of the fold while denatured contributions are negatively correlated with the stability of the *alternative* fold (Figures 6B-C). This result is consistent with “designing-in” the native fold (*i.e.* positive design) and “designing-out” incorrect folds (*i.e.* negative design) [41, 45–48]. Thus, the approach outlined here reveals both positive and negative design. In this case, the native state scores are largely responsible for the energy gap between most other folds, but that designing out the alternative fold using the denatured state becomes important for cases where two folds are closely matched. Is this a general phenomenon? If true, this would explain why computational force-fields that approximate free energy, but do not explicitly take the denatured state into account, are nonetheless relatively useful for the study of many proteins. But they also suggest that using these approaches to capture fold-switching may be more problematic as the denatured state is seldom considered.

Although remarkable, the clustering of experimental alpha and beta proteins in Figure 3 is not perfect, nor do we expect that it should be. As our approach scores sequences based on the average amino acid use across a database of proteins, it will undoubtedly be the case that certain amino acids, when placed in particular structural contexts, will cause a steric clash at that position, even if the statistics for the thermodynamic environment in which that structure resides is generally accommodating for the amino acid in question. This will result in cases where the score of a sequence in a particular fold is predicted high, but that substitution in actuality produces a counterindicating experimental outcome. Our success at discriminating the folds of high identity sequences suggests that such cases do not significantly impede the overall performance of the algorithm, but that their influence may nonetheless be surmised from the results.

With this in mind, there are at least three observations suggesting imperfect performance of the conformational specificity scoring. First, some experimentally unfolded sequences appear inseparable from the alpha and beta clouds. Second, the sequence corresponding to the G_A_95 fold lies uncharacteristically far from the visual center of the alpha cluster. Third, two alpha points are located within the beta cloud, these sequences are G_A_98 and G_B_98_T25I. These two sequences are particularly interesting as they are both single amino acid changes away from adopting the alternative fold. Indeed, evidence has been reported that both proteins populate the alternative fold to 5% or less in solution [10, 19]. These two ostensibly metamorphic proteins approach a middle ground in Figure 3, located between the alpha and beta point clusters. We speculate that there is a region of sequence space enriched in potentially metamorphic proteins, and that this region can be quantified by specificity scores that are moderately high for both folds but not exceedingly so. Experimental testing of this hypothesis is currently in progress.

Because this potentially metamorphic region lies between two clusters of stable, “monomorphic” proteins, it is tempting to propose that relatively small changes in the ensemble scoring could shift a protein originally in the metamorphic region to one or the other of the monomorphic regions. In favorable cases, it may be that a protein can move directly from one monomorphic cloud to the other due to targeted engineering, as suggested by Figures 4 and 5. Examination of scoring tables S1 and S2 reveals that, of the 20 amino acids, Glycine has the largest potential effects on the specificity score that could drive re-positioning between clouds. If an amino acid substitution were to involve Glycine, this could be a special case of controlled local unfolding [49], imparting an entropic energy penalty of around 1 kcal/mol [50], already demonstrated to be important in populating alternative states in adenylate kinase [51]. Experimental testing of this local unfolding hypothesis with regards to fold-switch behavior is currently in progress.

## Conclusions and Outlook

Ultimately, we find that our established statistical thermodynamic framework is able to recapitulate the energy landscape of potentially “metamorphic” proteins with access to two energy minima relative to the unfolded state. This framework, which is based on an ensemble description of protein structural thermodynamics, has been previously validated for fold recognition of traditional, “monomorphic” proteins. This body of evidence indicates that such an approach is applicable and valuable when applied to the complex case of fold-switching proteins and may be more broadly applied to both the discovery and design of these unique molecules.

In evaluating the implications of the *COREX* algorithm’s success in addressing metamorphic proteins, it is important to look back and take stock of the other important insights uncovered over the last two decades using this ensemble-based model of protein stability. These include deciphering how the ensemble affects regional stability [35], folding kinetics [36], electrostatics [52], and allostery [53]. Now added to this list are the origins and determinants of conformational specificity, fold switching, and global stability. Looking forward, this model thus provides an unprecedented opportunity to simultaneously and quantitatively; 1) investigate the effects of mutations on function and stability, 2) explore the evolution of new functions, and 3) to probe for distant relationships between structurally distant proteins. Indeed, taken together this body of work demonstrates, for the first time to our knowledge, that the entirety of the energetic, functional and structural landscape can be reconciled within a unified framework that is based simply on local and global unfolding equilibria.

Finally, although it has been the subject of speculation for years, perhaps the most important implication of the success of an energetic, rather than a structural, framework is that it provides strong evidence that the thermodynamic hypothesis, first described by Anfinsen more than 50 years ago [54] to explain how a single protein sequence determines its final structure, is in fact a foundational organizing principle that operates at the molecular level. This foundational principle is not only responsible for governing the folds and functions of the myriad of proteins, but also the ability of the proteome to expand and evolve.

## Materials and Methods

Metamorphic protein sequences and G_A_95/G_B_95 structures 2kdl/2kdm were obtained from Figure 2 and Table S1 of Alexander, *et al.* [19], and the Protein Data Bank [55]. Because 2kdl and 2kdm are NMR structural ensembles, *COREX* [35, 56, 57] was run separately on each model, using parameters Window Size = 5, Minimum Window Size = 4, Temperature of 25.0 °C, and Entropy Weight of either 0.5 (Native State) or 1.5 (Denatured State). Thermodynamic Descriptors of { ΔG, ΔH_ap_, ΔH_pol_, TΔS_conf_ }, obtained from *COREX* Native State [39, 41] and Denatured State [40] results, were mapped to Thermodynamic Environments [38] using Equations S1 and S2. These procedures resulted in two 4-vectors of Thermodynamic Environments for each residue (56 total), in each conformer (20 total), of each protein structure (2 total). For specificity scoring purposes, the most frequent thermodynamic environment (*i.e.* the mode of 20 conformers) was selected as the single most accurate descriptor at each residue position (Table S6).

Given a set of native state thermodynamic environments and an amino acid sequence of length *L* residues, a raw native score was computed as previously described [41], using a published lookup table (Table S4). From the raw score, a probability value for the native state, P_native_, was computed as previously described [41], according to Equation 1:

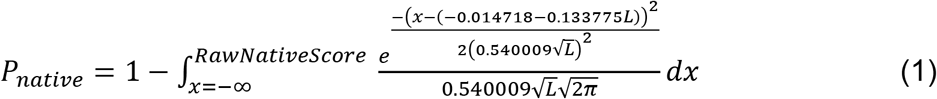

P-values for the denatured state were computed in a similar way, except that denatured environments (mapped using Equation S2) were substituted for native environments (mapped using Equation S1), a denatured log-odds scoring table (Table S5) was substituted for the native log-odds table (Table S4), and P-values were computed from raw scores using a Gaussian background distribution specific to the *COREX* denatured state (Equation 2):

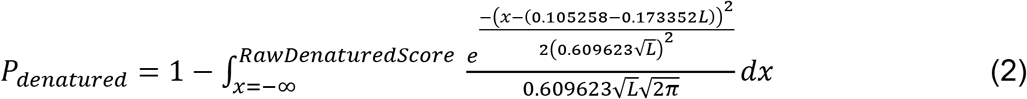

Thus, specificity scores displayed in Figures 3 and 4 are, under the assumption of statistical independence, the product of native and denatured P-values, *i.e.* the sum of the negated logs of native and denatured P-values:

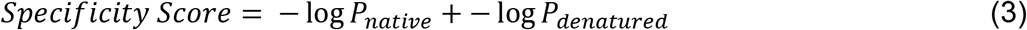

Using Equations 1–3, any sequence of thermodynamic environments can be scored against any amino acid sequence, regardless of length differences between the sequences [41] or indels, adopting appropriate gap penalties [39, 40]. For simplicity and for application to protein design, in this work we ignore indels and restrict possible amino acid sequences to be of equal length to those of the G_A_95/G_B_95 structures, *i.e.* 56 residues. For comparison, random sequences of this length (Table S9) were generated using natural background frequencies [58], and all 56 residue proteins of known structure (Table S7) were obtained from the Protein Data Bank (rcsb.org, November 5, 2020).

Experimental stabilities for individual proteins at T = 25 °C were inferred from published Tm values [19] using a modified Gibbs-Helmholtz equation [59]:

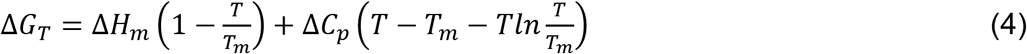

In Equation 4, ΔC_p_ for the G_A_ and G_B_ folds were 260 cal mol^-1^ K^-1^ and 830 cal mol^-1^ K^-1^ respectively [19]. For variants where ΔH_m_ was unavailable, empirical relationships based on linear correlation between ΔH_m_ and T_m_ were used to estimate ΔH_m_ (Equations 5 and 6). Equation 5 was applied to variants folding to G_A_ fold and Equation 6 was applied to variants folding to G_B_ fold.

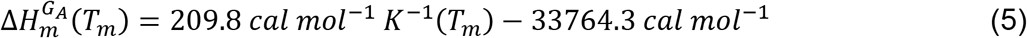

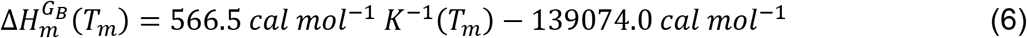

Stabilities for other Protein G Protein Data Bank entries used in Figure 6 were taken from [60–64]; all stability values used in Figure 6 are tabulated in Table S6.

## Acknowledgements

The authors wish to thank Dr. Jordan Hoffmann for insightful discussions regarding the fold-switching potential of Glycine. This work was supported by National Institutes of Health grants GM63747 and GM126130 to VJH and T32-GM080189.

One Supplementary PDF file *Supplementary_Material.pdf,* containing random and engineered protein sequences, scoring tables, and thermodynamic environments.

## References

1. Murzin, A.G., Biochemistry. Metamorphic proteins. Science, 2008. 320(5884): p. 1725–6.

2. Lella, M. and R. Mahalakshmi, Metamorphic Proteins: Emergence of Dual Protein Folds from One Primary Sequence. Biochemistry, 2017. 56(24): p. 2971–2984.

3. Das, M., et al., Identification and characterization of metamorphic proteins: Current and future perspectives. Biopolymers, 2021: p. e23473.

4. Dishman, A.F., F.C. Peterson, and B.F. Volkman, Specific binding-induced modulation of the XCL1 metamorphic equilibrium. Biopolymers, 2020: p. e23402.

5. Goodchild, S.C., et al., Metamorphic response of the CLIC1 chloride intracellular ion channel protein upon membrane interaction. Biochemistry, 2010. 49(25): p. 5278–89.

6. Galaz-Davison, P., et al., Differential Local Stability Governs the Metamorphic Fold Switch of Bacterial Virulence Factor RfaH. Biophys J, 2020. 118(1): p. 96–104.

7. Chang, Y.G., et al., Circadian rhythms. A protein fold switch joins the circadian oscillator to clock output in cyanobacteria. Science, 2015. 349(6245): p. 324–8.

8. Luo, X. and H. Yu, Protein metamorphosis: the two-state behavior of Mad2. Structure (London, England: 1993), 2008. 16(11): p. 1616–1625.

9. Porter, L.L. and L.L. Looger, Extant fold-switching proteins are widespread. Proc Natl Acad Sci U S A, 2018. 115(23): p. 5968–5973.

10. He, Y., et al., Mutational tipping points for switching protein folds and functions. Structure, 2012. 20(2): p. 283–91.

11. Kumirov, V.K., et al., Multistep mutational transformation of a protein fold through structural intermediates. Protein Sci, 2018. 27(10): p. 1767–1779.

12. Dishman, A.F., et al., Evolution of fold switching in a metamorphic protein. Science, 2021. 371(6524): p. 86–90.

13. Rose, G.D. and T.P. Creamer, Protein folding: predicting predicting. Proteins, 1994. 19(1): p. 1–3.

14. Dalal, S. and L. Regan, Understanding the sequence determinants of conformational switching using protein design. Protein Sci, 2000. 9(9): p. 1651–9.

15. Ambroggio, X.I. and B. Kuhlman, Computational design of a single amino acid sequence that can switch between two distinct protein folds. J Am Chem Soc, 2006. 128(4): p. 1154–61.

16. Porter, L.L., et al., Subdomain interactions foster the design of two protein pairs with ~80% sequence identity but different folds. Biophys J, 2015. 108(1): p. 154–62.

17. Wei, K.Y., et al., Computational design of closely related proteins that adopt two well-defined but structurally divergent folds. Proc Natl Acad Sci U S A, 2020. 117(13): p. 7208–7215.

18. Dawson, W.M., et al., Structural resolution of switchable states of a de novo peptide assembly. Nat Commun, 2021. 12(1): p. 1530.

19. Alexander, P.A., et al., A minimal sequence code for switching protein structure and function. Proc Natl Acad Sci U S A, 2009. 106(50): p. 21149–54.

20. Mezei, M., Foldability and chameleon propensity of fold-switching protein sequences. Proteins, 2020.

21. Chen, N., et al., Sequence-Based Prediction of Metamorphic Behavior in Proteins. Biophys J, 2020. 119(7): p. 1380–1390.

22. Kim, A.K., L.L. Looger, and L.L. Porter, A high-throughput predictive method for sequence-similar fold switchers. Biopolymers, 2021: p. e23416.

23. Allison, J.R., et al., Current computer modeling cannot explain why two highly similar sequences fold into different structures. Biochemistry, 2011. 50(50): p. 10965–73.

24. Tian, P. and R.B. Best, Exploring the sequence fitness landscape of a bridge between protein folds. PLoS Comput Biol, 2020. 16(10): p. e1008285.

25. Seifi, B. and S. Wallin, The C-terminal domain of transcription factor RfaH: Folding, fold switching and energy landscape. Biopolymers, 2021: p. e23420.

26. Robustelli, P., S. Piana, and D.E. Shaw, Developing a molecular dynamics force field for both folded and disordered protein states. Proc Natl Acad Sci U S A, 2018. 115(21): p. E4758–e4766.

27. Dauber-Osguthorpe, P. and A.T. Hagler, Biomolecular force fields: where have we been, where are we now, where do we need to go and how do we get there? J Comput Aided Mol Des, 2019. 33(2): p. 133–203.

28. Hagler, A.T., Force field development phase II: Relaxation of physics-based criteria … or inclusion of more rigorous physics into the representation of molecular energetics. J Comput Aided Mol Des, 2019. 33(2): p. 205–264.

29. Appadurai, R., J. Nagesh, and A. Srivastava, High resolution ensemble description of metamorphic and intrinsically disordered proteins using an efficient hybrid parallel tempering scheme. Nat Commun, 2021. 12(1): p. 958.

30. Jumper, J., et al., Highly accurate protein structure prediction with AlphaFold. Nature, 2021. 596(7873): p. 583–589.

31. Wolynes, P.G., Evolution, energy landscapes and the paradoxes of protein folding. Biochimie, 2015. 119: p. 218–30.

32. Rose, G.D., Protein folding - seeing is deceiving. Protein Sci, 2021. 30(8): p. 1606–1616.

33. Nassar, R., et al., The Protein Folding Problem: The Role of Theory. J Mol Biol, 2021: p. 167126.

34. Lattman, E.E. and G.D. Rose, Protein folding--what’s the question? Proc Natl Acad Sci U S A, 1993. 90(2): p. 439–41.

35. Hilser, V.J. and E. Freire, Structure-based calculation of the equilibrium folding pathway of proteins. Correlation with hydrogen exchange protection factors. J Mol Biol, 1996. 262(5): p. 756–72.

36. Hilser, V.J., et al., A statistical thermodynamic model of the protein ensemble. Chem Rev, 2006. 106(5): p. 1545–58.

37. Liu, T., et al., Quantitative assessment of protein structural models by comparison of H/D exchange MS data with exchange behavior accurately predicted by DXCOREX. J Am Soc Mass Spectrom, 2012. 23(1): p. 43–56.

38. Wrabl, J.O., S.A. Larson, and V.J. Hilser, Thermodynamic environments in proteins: fundamental determinants of fold specificity. Protein Sci, 2002. 11(8): p. 1945–57.

39. Larson, S.A. and V.J. Hilser, Analysis of the “thermodynamic information content” of a Homo sapiens structural database reveals hierarchical thermodynamic organization. Protein Sci, 2004. 13(7): p. 1787–801.

40. Wang, S., et al., Denatured-state energy landscapes of a protein structural database reveal the energetic determinants of a framework model for folding. J Mol Biol, 2008. 381(5): p. 1184–201.

41. Hoffmann, J., J.O. Wrabl, and V.J. Hilser, The role of negative selection in protein evolution revealed through the energetics of the native state ensemble. Proteins, 2016. 84(4): p. 435–47.

42. Wrabl, J.O., S.A. Larson, and V.J. Hilser, Thermodynamic propensities of amino acids in the native state ensemble: implications for fold recognition. Protein Sci, 2001. 10(5): p. 1032–45.

43. Stormo, G.D., Modeling the specificity of protein-DNA interactions. Quant Biol, 2013. 1(2): p. 115–130.

44. Ruan, S. and G.D. Stormo, Inherent limitations of probabilistic models for protein-DNA binding specificity. PLoS Comput Biol, 2017. 13(7): p. e1005638.

45. Bolon, D.N., et al., Specificity versus stability in computational protein design. Proc Natl Acad Sci U S A, 2005. 102(36): p. 12724–9.

46. Anderson, T.A., M.H. Cordes, and R.T. Sauer, Sequence determinants of a conformational switch in a protein structure. Proc Natl Acad Sci U S A, 2005. 102(51): p. 18344–9.

47. Berezovsky, I.N., K.B. Zeldovich, and E.I. Shakhnovich, Positive and negative design in stability and thermal adaptation of natural proteins. PLoS Comput Biol, 2007. 3(3): p. e52.

48. Xu, F., et al., De novo self-assembling collagen heterotrimers using explicit positive and negative design. Biochemistry, 2010. 49(11): p. 2307–16.

49. Schrank, T.P., et al., Strategies for the thermodynamic characterization of linked binding/local folding reactions within the native state application to the LID domain of adenylate kinase from Escherichia coli. Methods Enzymol, 2011. 492: p. 253–82.

50. Matthews, B.W., H. Nicholson, and W.J. Becktel, Enhanced protein thermostability from site-directed mutations that decrease the entropy of unfolding. Proc Natl Acad Sci U S A, 1987. 84(19): p. 6663–7.

51. Saavedra, H.G., et al., Dynamic allostery can drive cold adaptation in enzymes. Nature, 2018. 558(7709): p. 324–328.

52. Whitten, S.T., E.B. Garcia-Moreno, and V.J. Hilser, Local conformational fluctuations can modulate the coupling between proton binding and global structural transitions in proteins. Proc Natl Acad Sci U S A, 2005. 102(12): p. 4282–7.

53. Motlagh, H.N., et al., The ensemble nature of allostery. Nature, 2014. 508(7496): p. 331–9.

54. Anfinsen, C.B., Principles that govern the folding of protein chains. Science, 1973. 181(4096): p. 223–30.

55. Berman, H.M., et al., The Protein Data Bank. Nucleic Acids Res, 2000. 28(1): p. 235–42.

56. Vertrees, J., et al., COREX/BEST server: a web browser-based program that calculates regional stability variations within protein structures. Bioinformatics, 2005. 21(15): p. 3318–9.

57. Hilser, V.J. and S.T. Whitten, Using the COREX/BEST server to model the native-state ensemble. Methods Mol Biol, 2014. 1084: p. 255–69.

58. Robinson, A.B. and L.R. Robinson, Distribution of glutamine and asparagine residues and their near neighbors in peptides and proteins. Proc Natl Acad Sci U S A, 1991. 88(20): p. 8880–4.

59. Robertson, A.D. and K.P. Murphy, Protein Structure and the Energetics of Protein Stability. Chem Rev, 1997. 97(5): p. 1251–1268.

60. Strop, P., A.M. Marinescu, and S.L. Mayo, Structure of a protein G helix variant suggests the importance of helix propensity and helix dipole interactions in protein design. Protein Sci, 2000. 9(7): p. 1391–4.

61. Maxwell, K.L., et al., Protein folding: defining a “standard” set of experimental conditions and a preliminary kinetic data set of two-state proteins. Protein Sci, 2005. 14(3): p. 602–16.

62. Wunderlich, M., et al., Optimization of the gbeta1 domain by computational design and by in vitro evolution: structural and energetic basis of stabilization. J Mol Biol, 2007. 373(3): p. 775–84.

63. Davey, J.A., et al., Rational design of proteins that exchange on functional timescales. Nat Chem Biol, 2017. 13(12): p. 1280–1285.

64. Maniaci, B., et al., Design of High-Affinity Metal-Controlled Protein Dimers. Biochemistry, 2019. 58(17): p. 2199–2207.

