## Supplemental Methods, Tables, Figures for "A Predictive Energy Landscape Model of Metamorphic Protein Conformational Specificity"

**Supplementary Material for "A Predictive Energy Landscape Model for Metamorphic Protein Conformational Specificity" by James O. Wrabl, Keila Voortman-Sheetz, and Vincent J. Hilser.**

#### *Mapping of COREX output to Native and Denatured State Thermodynamic Environments*

In detail, the mapping of each protein in terms of Thermodynamic Environments (TEs) was accomplished as follows. The *COREX* output for every amino acid *j* in the protein was treated as two four-dimensional vectors, one each for the native (N) and denatured (D) states, *i.e.* $\{\Delta G_j^N, \Delta H_{apolar,j}^N, \Delta H_{polar,j}^N, T\Delta S_{conf,j}^N\}$ and $\{\Delta G_j^D, \Delta H_{apolar,j}^D, \Delta H_{polar,j}^D, T\Delta S_{conf,j}^D\}$. The native and denatured TEs corresponding to such vectors were defined as the TEs whose cluster centers in high-dimensional space, over a large database of proteins, were closest in Manhattan distance according to Equations (S1) and (S2).

$$TE_j^N \equiv \min_k\left[abs\left(\overline{\Delta G_k^N} - \Delta G_j^N\right) + abs\left(\overline{\Delta H_{apolar,k}^N} - \Delta H_{apolar,j}^N\right) + abs\left(\overline{\Delta H_{polar,k}^N} - \Delta H_{polar,j}^N\right) + 3\, abs\left(\overline{T\Delta S_{conf,k}^N} - T\Delta S_{conf,j}^N\right)\right] \quad \text{(S1)}$$

$$TE_j^D \equiv \min_k\left[abs\left(\overline{\Delta G_k^D} - \Delta G_j^D\right) + abs\left(\overline{\Delta H_{apolar,k}^D} - \Delta H_{apolar,j}^D\right) + abs\left(\overline{\Delta H_{polar,k}^D} - \Delta H_{polar,j}^D\right) + abs\left(\overline{T\Delta S_{conf,k}^D} - T\Delta S_{conf,j}^D\right)\right] \quad \text{(S2)}$$

The index *k* runs over the eight native state TEs for Equation (S1), and the index *k* runs over the eight denatured state environments in Equation (S2).

Average values $\left\{\overline{\Delta G_k^N}, \overline{\Delta H_{apolar,k}^N}, \overline{\Delta H_{polar,k}^N}, \overline{T\Delta S_{conf,k}^N}\right\}$ and $\left\{\overline{\Delta G_k^D}, \overline{\Delta H_{apolar,k}^D}, \overline{\Delta H_{polar,k}^D}, \overline{T\Delta S_{conf,k}^D}\right\}$ for each environment have been published (Wang, *et al.* 2008, Hoffmann, *et al.* 2016) and are given in Supplementary Tables S1 and S2.

#### *Empirical Procedure to Estimate the Clustering Significance of Figure 3.*

Although more sophisticated models are possible, we took a simple approach to estimate the significance of the Figure 3 clusters, using random sampling of the binary sequence space (Table S8, 8192 sequences). We assumed that this space was a complete enumeration of the degrees of freedom of the system. 1000 randomly sampled clusters of (x, y) points from the 8192 points in the Figure 3 two-dimensional plot represented possible clusters of experimental alpha, beta, or unfolded proteins in binary space. Each randomly sampled cluster contained a number of (x, y) points, equal to the number of experimentally observed alpha, beta, and unfolded proteins; these numbers were 13, 14, and 9 respectively. For each group of experimentally observed alpha, beta, and unfolded proteins, we then assumed that the observed standard deviations along the x and y axes of Figure 3 were good measures of the tightness of the cluster relative to the binary space.

Comparing the standard deviations of the randomly selected clusters with the observed clusters, 21/1000 random clusters exhibited smaller standard deviations along both x and y axes than the observed alpha cluster. Thus, we estimated that the tightness of the observed alpha cluster had a 21/1000 = 2.1% probability of chance occurrence. Similarly, the beta cluster had a 0/1000 = 0.0% probability of chance occurrence, and the unfolded cluster had a 5/1000 = 0.5% probability of chance occurrence. Thus, we concluded that observed clusters in Figure were unlikely to have arisen by chance.

This simple approach had a number of limitations, including tacit assumptions that all clusters were perfect rectangles and all cluster axes were perfectly aligned to the x and y axes of Figure 3. However, we believe these probabilities are underestimates of the true significance, since the random clusters were considered in isolation, instead of altogether, and the lack of overlap seen between the true clusters was not enforced. Conservatively, then, we report that the significance of the clustering in Figure 3 is less than 0.02, the least significant of the groups.

**Figure S1. Average regional contributions (top) to cumulative conformational specificity (bottom) for all experimentally measured sequence variants. (Compare to Main Text Figure 4.)** The lower plots were obtained by cumulatively evaluating Main Text Equations 1 and 2 at every residue position in every sequence, with a guiding line drawn by averaging every five residues. The upper plots were obtained by decomposing the data in the lower plots, subtracting from every residue the previous, averaging over a centered five residue window, with a guiding line drawn by further averaging nearest neighbors. **A.** Scoring in the $G_A$ fold (α fold, 2kdl) environments reveals that the C-terminal half of the protein contributes more to the score, and to the specificity, than does the N-terminal half for all variants in binary space. Positions in the neighborhood of amino acid substitutions 30 and 45 drive the specificity between alpha and beta, with position 20 contributing a lesser amount. **B**. Scoring in the $G_B$ fold (β fold, 2kdm) environments reveals that the C-terminal half of the protein contributes more to the score than does the N-terminal half, similarly with positions 30 and 45 contributing more than position 20.

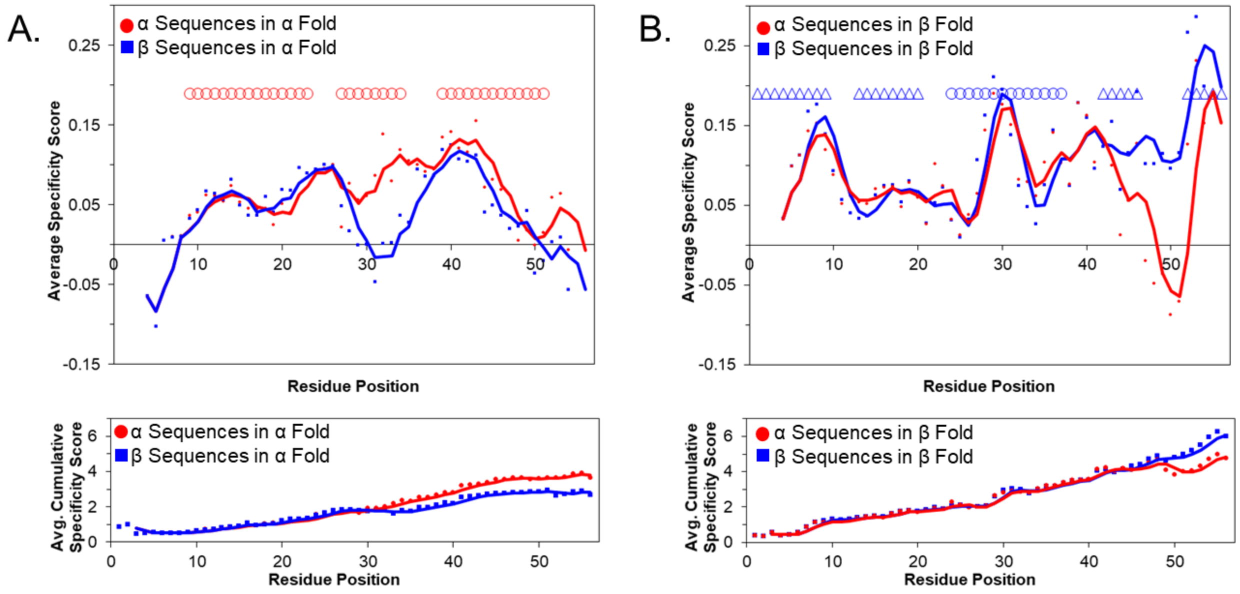


**Table S1. Average Native State Thermodynamic Environments (Cluster Centers).** Native State Thermodynamic Environments (NTE) are organized such that NTE1 is least stable on average and NTE8 is most stable.

| Native | NTE1 | NTE2 | NTE3 | NTE4 | NTE5 | NTE6 | NTE7 | NTE8 |
|---|---|---|---|---|---|---|---|---|
| N $\Delta G$ | -3.5 * | -4.4 | -6.4 | -7.5 | -8.5 | -9.8 | -10.5 | -12.4 |
| N $\Delta H_{apolar}$ | 4.9 | 6.3 | 6.5 | 10.9 | 8.8 | 9.1 | 14.0 | 14.2 |
| N $\Delta H_{polar}$ | -6.1 | -8.6 | -11.5 | -9.4 | -12.4 | -15.1 | -12.5 | -16.3 |
| N $T\Delta S_{conf}$ | -3.1 | -4.1 | -4.9 | -4.1 | -4.4 | -5.2 | -4.6 | -5.6 |

* Values are in kcal/mol at 25.0 C.

**Table S2. Average Denatured State Thermodynamic Environments (Cluster Centers).** Denatured State Thermodynamic Environments (DTE) are organized such that DTE1 is least stable on average and DTE8 is most stable.

| Denatured | DTE1 | DTE2 | DTE3 | DTE4 | DTE5 | DTE6 | DTE7 | DTE8 |
|---|---|---|---|---|---|---|---|---|
| D $\Delta G$ | 10.7 * | 10.2 | 9.8 | 9.1 | 9.1 | 8.4 | 7.6 | 7.5 |
| D $\Delta H_{apolar}$ | -1.5 | -1.5 | -1.0 | 0.7 | -1.0 | -1.1 | -0.3 | -1.1 |
| D $\Delta H_{polar}$ | 1.1 | 2.4 | -1.1 | -0.4 | 0.0 | 1.1 | -2.2 | -0.9 |
| D $T\Delta S_{conf}$ | -9.9 | -8.4 | -10.9 | -10.3 | -9.3 | -7.7 | -9.8 | -8.3 |

* Values are in kcal/mol at 25.0 C.

**Table S3. Native and Denatured Thermodynamic Environments (NTE and DTE) for $G_A$95 (2kdl.pdb) and $G_B$95 (2kdm.pdb) Folds.**

| Residue Position 2kdl.pdb | Most Frequent NTE | Number of Conformers (20 max.) | | Residue Position 2kdm.pdb | Most Frequent NTE | Number of Conformers (20 max.) |
|---|---|---|---|---|---|---|
| 1 | 1 | 19 | | 1 | 7 | 15 |
| 2 | 1 | 19 | | 2 | 7 | 15 |
| 3 | 1 | 19 | | 3 | 7 | 15 |
| 4 | 1 | 19 | | 4 | 7 | 15 |
| 5 | 1 | 17 | | 5 | 8 | 17 |
| 6 | 1 | 16 | | 6 | 8 | 18 |
| 7 | 1 | 16 | | 7 | 7 | 9 |
| 8 | 1 | 17 | | 8 | 3 | 14 |
| 9 | 1 | 10 | | 9 | 2 | 17 |
| 10 | 2 | 10 | | 10 | 2 | 16 |
| 11 | 2 | 8 | | 11 | 2 | 16 |
| 12 | 4 | 10 | | 12 | 2 | 16 |
| 13 | 6 | 6 | | 13 | 2 | 17 |
| 14 | 5 | 11 | | 14 | 2 | 11 |
| 15 | 5 | 10 | | 15 | 2 | 11 |
| 16 | 5 | 12 | | 16 | 4 | 14 |
| 17 | 6 | 10 | | 17 | 2 | 13 |
| 18 | 5 | 15 | | 18 | 2 | 15 |
| 19 | 5 | 11 | | 19 | 2 | 17 |
| 20 | 1 | 18 | | 20 | 2 | 19 |
| 21 | 1 | 19 | | 21 | 2 | 18 |
| 22 | 1 | 18 | | 22 | 2 | 18 |
| 23 | 1 | 14 | | 23 | 2 | 18 |
| 24 | 1 | 15 | | 24 | 2 | 18 |
| 25 | 1 | 15 | | 25 | 2 | 18 |
| 26 | 2 | 12 | | 26 | 3 | 10 |
| 27 | 2 | 15 | | 27 | 4 | 8 |
| 28 | 1 | 14 | | 28 | 5 | 8 |
| 29 | 4 | 18 | | 29 | 7 | 19 |
| 30 | 4 | 18 | | 30 | 7 | 17 |
| 31 | 2 | 8 | | 31 | 6 | 12 |
| 32 | 1 | 11 | | 32 | 6 | 11 |
| 33 | 2 | 11 | | 33 | 5 | 10 |
| 34 | 1 | 17 | | 34 | 5 | 9 |
| 35 | 1 | 17 | | 35 | 3 | 13 |
| 36 | 1 | 16 | | 36 | 3 | 18 |
| 37 | 1 | 17 | | 37 | 3 | 14 |
| 38 | 1 | 16 | | 38 | 2 | 18 |
| 39 | 1 | 12 | | 39 | 2 | 19 |
| 40 | 1 | 10 | | 40 | 2 | 19 |

| | | | | | | |
|---|---|---|---|---|---|---|
| 41 | 1 | 9 | | 41 | 2 | 19 |
| 42 | 2 | 10 | | 42 | 2 | 19 |
| 43 | 4 | 11 | | 43 | 2 | 13 |
| 44 | 4 | 12 | | 44 | 3 | 12 |
| 45 | 7 | 16 | | 45 | 2 | 11 |
| 46 | 8 | 16 | | 46 | 2 | 14 |
| 47 | 5 | 8 | | 47 | 2 | 12 |
| 48 | 4 | 6 | | 48 | 2 | 12 |
| 49 | 2 | 9 | | 49 | 2 | 13 |
| 50 | 1 | 11 | | 50 | 3 | 12 |
| 51 | 1 | 18 | | 51 | 3 | 18 |
| 52 | 1 | 20 | | 52 | 7 | 20 |
| 53 | 1 | 20 | | 53 | 4 | 17 |
| 54 | 1 | 20 | | 54 | 4 | 17 |
| 55 | 1 | 20 | | 55 | 4 | 17 |
| 56 | 1 | 20 | | 56 | 4 | 17 |

| Residue Position 2kdl.pdb | Most Frequent DTE | Number of Conformers (20 max.) | | Residue Position 2kdm.pdb | Most Frequent DTE | Number of Conformers (20 max.) |
|---|---|---|---|---|---|---|
| 1 | 6 | 11 | | 1 | 8 | 10 |
| 2 | 6 | 11 | | 2 | 8 | 10 |
| 3 | 6 | 11 | | 3 | 8 | 10 |
| 4 | 6 | 11 | | 4 | 8 | 10 |
| 5 | 5 | 14 | | 5 | 5 | 9 |
| 6 | 1 | 10 | | 6 | 2 | 12 |
| 7 | 1 | 14 | | 7 | 2 | 16 |
| 8 | 5 | 11 | | 8 | 5 | 13 |
| 9 | 5 | 10 | | 9 | 5 | 18 |
| 10 | 4 | 9 | | 10 | 5 | 18 |
| 11 | 4 | 9 | | 11 | 5 | 18 |
| 12 | 4 | 9 | | 12 | 5 | 18 |
| 13 | 5 | 9 | | 13 | 5 | 18 |
| 14 | 5 | 14 | | 14 | 1 | 19 |
| 15 | 5 | 16 | | 15 | 1 | 18 |
| 16 | 5 | 19 | | 16 | 1 | 12 |
| 17 | 5 | 16 | | 17 | 2 | 10 |
| 18 | 5 | 18 | | 18 | 1 | 12 |
| 19 | 5 | 19 | | 19 | 1 | 16 |
| 20 | 5 | 17 | | 20 | 1 | 19 |
| 21 | 5 | 18 | | 21 | 1 | 19 |
| 22 | 5 | 11 | | 22 | 1 | 12 |
| 23 | 1 | 12 | | 23 | 1 | 12 |
| 24 | 1 | 19 | | 24 | 3 | 11 |
| 25 | 1 | 17 | | 25 | 4 | 6 |
| 26 | 5 | 14 | | 26 | 4 | 20 |
| 27 | 5 | 18 | | 27 | 4 | 20 |

| | | | | | | |
|---|---|---|---|---|---|---|
| 28 | 5 | 18 | | 28 | 4 | 20 |
| 29 | 5 | 10 | | 29 | 4 | 18 |
| 30 | 6 | 16 | | 30 | 5 | 17 |
| 31 | 6 | 15 | | 31 | 5 | 15 |
| 32 | 6 | 17 | | 32 | 5 | 15 |
| 33 | 6 | 20 | | 33 | 5 | 12 |
| 34 | 6 | 20 | | 34 | 5 | 11 |
| 35 | 5 | 18 | | 35 | 5 | 14 |
| 36 | 5 | 18 | | 36 | 5 | 14 |
| 37 | 2 | 10 | | 37 | 5 | 13 |
| 38 | 1 | 19 | | 38 | 5 | 9 |
| 39 | 1 | 19 | | 39 | 5 | 12 |
| 40 | 1 | 19 | | 40 | 1 | 16 |
| 41 | 1 | 19 | | 41 | 1 | 20 |
| 42 | 5 | 15 | | 42 | 1 | 18 |
| 43 | 5 | 14 | | 43 | 1 | 20 |
| 44 | 5 | 10 | | 44 | 1 | 17 |
| 45 | 5 | 12 | | 45 | 4 | 10 |
| 46 | 5 | 11 | | 46 | 3 | 16 |
| 47 | 5 | 11 | | 47 | 3 | 14 |
| 48 | 5 | 16 | | 48 | 3 | 12 |
| 49 | 5 | 18 | | 49 | 3 | 12 |
| 50 | 5 | 18 | | 50 | 3 | 10 |
| 51 | 5 | 14 | | 51 | 1 | 16 |
| 52 | 5 | 16 | | 52 | 1 | 16 |
| 53 | 6 | 10 | | 53 | 2 | 19 |
| 54 | 6 | 10 | | 54 | 2 | 19 |
| 55 | 6 | 10 | | 55 | 2 | 19 |
| 56 | 6 | 10 | | 56 | 2 | 19 |

**Table S4. Native State Log-Odds Score Matrix (Hoffmann, *et al.* 2016).** Native State Thermodynamic Environments (NTE) are organized such that NTE1 is least stable on average and NTE8 is most stable, according to Table S1.

| | W | F | Y | L | I | V | M | A | G | P | C | T | S | Q | N | E | D | H | K | R |
|---|---|---|---|---|---|---|---|---|---|---|---|---|---|---|---|---|---|---|---|---|
| NTE1 | -0.90 | -1.53 | -0.93 | -0.23 | 0.12 | 0.20 | -0.28 | 0.45 | 0.37 | 0.99 | -0.29 | 0.15 | -0.03 | -0.19 | -0.27 | -0.37 | -0.22 | -0.18 | -0.11 | -0.53 |
| NTE2 | -0.82 | -1.05 | -1.09 | -0.20 | -0.24 | -0.03 | -0.39 | 0.05 | 0.66 | 0.63 | -0.18 | 0.17 | 0.05 | -0.25 | 0.10 | -0.07 | 0.11 | -0.24 | 0.05 | -0.42 |
| NTE3 | -0.83 | -1.05 | -0.95 | -0.85 | -0.77 | -0.60 | -0.11 | -0.29 | 0.65 | -0.42 | 0.36 | 0.19 | 0.04 | 0.15 | 0.54 | 0.41 | 0.47 | -0.26 | 0.39 | -0.55 |
| NTE4 | -0.03 | 0.18 | 0.05 | 0.39 | 0.56 | 0.51 | -0.57 | 0.02 | -0.25 | 0.38 | -1.62 | 0.24 | -0.39 | -0.89 | -0.47 | -0.33 | -0.42 | -0.19 | -0.12 | -0.29 |
| NTE5 | -1.04 | -0.50 | -0.55 | -0.03 | -0.29 | -0.10 | 0.07 | 0.39 | -0.27 | -0.14 | -0.09 | -0.16 | 0.37 | 0.22 | 0.07 | -0.12 | 0.15 | 0.04 | 0.01 | 0.28 |
| NTE6 | -0.79 | -0.66 | -0.09 | -0.44 | -0.75 | -0.63 | 0.18 | -0.45 | -0.37 | -1.36 | 0.66 | -0.11 | 0.40 | 0.44 | 0.45 | 0.39 | 0.32 | 0.08 | 0.20 | 0.40 |
| NTE7 | 0.72 | 0.98 | 0.42 | 0.55 | 0.57 | 0.38 | 0.12 | 0.15 | -1.08 | -0.49 | -0.56 | -0.16 | -0.58 | -0.59 | -1.11 | -0.47 | -0.61 | 0.10 | -0.43 | -0.44 |
| NTE8 | 1.09 | 0.58 | 1.06 | -0.09 | -0.21 | -0.44 | 0.57 | -0.93 | -1.28 | -2.00 | 0.39 | -0.51 | -0.37 | 0.44 | -0.26 | 0.13 | -0.41 | 0.45 | -0.25 | 0.66 |

**Table S5. Denatured State Log-Odds Score Matrix (Wang, *et al.* 2008).** Denatured State Thermodynamic Environments (DTE) are organized such that DTE1 is least stable on average and DTE8 is most stable.

| | W | F | Y | L | I | V | M | A | G | P | C | T | S | Q | N | E | D | H | K | R |
|---|---|---|---|---|---|---|---|---|---|---|---|---|---|---|---|---|---|---|---|---|
| DTE1 | 0.09 | -0.29 | -0.59 | -0.22 | -0.65 | 0.21 | -0.19 | -0.46 | 0.98 | -0.12 | 0.38 | 0.25 | -1.36 | -0.38 | 0.36 | 0.10 | 0.18 | 0.10 | 0.13 | -1.86 |
| DTE2 | -0.39 | -0.55 | -0.79 | 0.20 | 0.96 | 0.80 | -0.04 | -0.08 | -0.80 | -0.12 | 0.75 | 0.53 | -1.28 | -1.11 | -0.60 | -0.51 | -0.79 | -0.00 | -0.05 | -0.03 |
| DTE3 | -0.03 | -0.11 | -0.58 | -0.72 | -1.62 | -0.87 | -0.18 | -0.55 | 0.90 | -0.56 | -0.58 | -0.23 | -0.49 | 0.34 | 0.85 | 0.48 | 0.73 | -0.18 | 0.18 | -1.83 |
| DTE4 | 0.78 | 0.96 | 1.48 | -0.46 | -1.14 | -1.16 | 0.07 | -0.30 | -1.18 | -0.16 | -0.56 | -0.11 | -0.51 | 0.33 | -0.00 | 0.29 | -0.11 | 0.10 | 0.04 | -0.47 |
| DTE5 | 0.03 | -0.09 | -0.52 | 0.16 | -0.10 | 0.02 | 0.24 | 0.26 | -0.49 | 0.14 | -0.06 | 0.09 | 0.05 | -0.04 | -0.06 | 0.06 | 0.04 | 0.25 | 0.07 | -0.53 |

```
DTE6 -0.52 -0.59 -0.51  0.31  0.67  0.31 -0.07  0.19 -1.52  0.49  0.19 -0.05 -0.03 -0.53 -1.20 -0.85 -1.14  0.12 -0.15  0.94
DTE7  0.15  0.42  0.26 -0.08 -0.82 -1.02  0.10  0.14 -0.98 -0.27 -1.07 -0.70  0.89  0.55 -0.29  0.12  0.16 -0.61 -0.25  0.35
DTE8 -1.06 -0.70 -1.20  0.38  0.13 -0.15 -0.01  0.46 -1.76  0.27 -0.66 -0.57  0.86  0.11 -1.29 -0.54 -0.55 -0.18 -0.18  0.80
```

**Table S6. Experimental Stabilities, Specificity Scores, and Raw Native and Denatured Scores for High-Identity Proteins.**

| Protein | Experimental Stability kcal/mol @ 25 °C | Specificity Score | Raw Native Score in $G_A$ Fold | Raw Denatured Score in $G_B$ Fold |
|---|---|---|---|---|
| GA77 | 4.9 | 3.88 | -0.35 | 3.08 |
| GA1 | 3.8 | 3.51 | -0.27 | 1.63 |
| GA2 | 4.0 | 3.92 | -0.45 | 6.05 |
| GA3 | 3.5 | 3.45 | -0.73 | 3.09 |
| GA4 | 3.3 | 3.66 | -0.48 | 5.02 |
| GA5 | 3.8 | 3.89 | -0.23 | 3.98 |
| GA6 | 4.4 | 3.95 | -0.09 | 3.20 |
| GA7 | 4.7 | 3.37 | -2.33 | 3.25 |
| GA88 | 4.2 | 3.42 | -2.07 | 3.37 |
| GA91 | 3.4 | 3.69 | -2.04 | 4.40 |
| GA95 | 2.4 | 3.69 | -1.92 | 5.30 |
| GA98 | 1.2 | 3.25 | -2.30 | 5.31 |

| Protein | Experimental Stability kcal/mol @ 25 °C | Specificity Score | Raw Native Score in $G_B$ Fold | Raw Denatured Score in $G_A$ Fold |
|---|---|---|---|---|
| GB77 | 4.1 | 5.93 | 3.40 | -1.84 |
| GB1 | 4.0 | 6.00 | 2.79 | -1.03 |
| GB2 | 3.2 | 5.85 | 3.28 | -1.86 |
| GB3 | 2.6 | 5.93 | 3.15 | -1.94 |
| GB4 | 3.5 | 6.65 | 4.01 | -0.40 |
| GB5 | 3.4 | 5.45 | 2.99 | -2.74 |
| GB6 | 3.8 | 6.16 | 3.66 | -0.66 |
| GB7 | 3.7 | 5.33 | 2.99 | -2.03 |
| GB88b | 3.4 | 6.09 | 2.99 | 0.22 |
| GB91 | 2.5 | 6.01 | 2.87 | 0.20 |
| GB95 | 2.4 | 6.24 | 3.13 | 1.38 |
| GB8 | 1.2 | 6.02 | 2.62 | 0.10 |
| GB98 | 1.0 | 6.25 | 2.88 | 1.28 |
| 2onq_A | 7.2 | 7.82 | 8.23 | -5.37 |
| 1em7_A | 6.9 | 6.79 | 7.61 | -3.06 |
| 2on8_A | 6.5 | 7.06 | 6.86 | -4.95 |
| 1gb1_A | 4.7 | 6.90 | 8.02 | -2.88 |
| 2gb1_A | 4.7 | 6.90 | 8.02 | -2.88 |
| 6nlb_A | 4.0 | 6.67 | 6.68 | -0.82 |
| 5ub0_A | 4.0 | 7.31 | 7.96 | -3.35 |

| | | | | |
|---|---|---|---|---|
| 6nl8_A | 2.8 | 7.08 | 7.05 | -0.59 |
| 5ubs_A | 2.7 | 6.99 | 7.55 | -4.67 |
| 5uce_A | 1.7 | 6.70 | 6.70 | -4.50 |
| 5ucf_A | 1.7 | 6.70 | 6.70 | -4.50 |

**Table S7. 56-Residue Chains Contained in the Protein Data Bank.**

```
1a02_F M K R R I R R E R N K M A A A K S R N R R R E L T D T L Q A E T D Q L E D E K S A L Q T E I A N L L K E K E K L
1a02_J M K A E R K R M R N R I A A S K S R K R K L E R I A R L E E K V K T L K A Q N S E L A S T A N M L R E Q V A Q L
1aze_A M E A I A K V D F K A T A D D E L S F K R G D I L K V L N E E S D Q N W Y K A E L N G K D G F I P K N Y I E M K
1be3_K M L T R F L G P R Y R Q L A R N W V P T A Q L W G A V G A V G L V S A T D S R L I L D W V P Y I N G K F K K D D
1bgy_K M L T R F L G P R Y R Q L A R N W V P T A Q L W G A V G A V G L V S A T D S R L I L D W V P Y I N G K F K K D D
1bgy_W M L T R F L G P R Y R Q L A R N W V P T A Q L W G A V G A V G L V S A T D S R L I L D W V P Y I N G K F K K D D
1bor_A E E E F Q F L R C Q Q C Q A E A K C P K L L P C L H T L C S G C L E A S G M Q C P I C Q A P W P L G A D T P A L
1brc_I V R E V C S E Q A E T G P C R A M I S R W Y F D V T E G K C A P F F Y G G C G G N R N N F D T E E Y C M A V C G
1bw6_A M G P K R R Q L T F R E K S R I I Q E V E E N P D L R K G E I A R R F N I P P S T L S T I L K N K R A I L A S E
1ccv_A E E C G P N E V F N T C G S A C A P T C A Q P K T R I C T M Q C R I G C Q C Q E G F L R N G E G A C V L P E N C
1cgi_I D S L G R E A K C Y N E L N G C T Y E Y R P V C G T D G D T Y P N E C V L C F E N R K R Q T S I L I Q K S G P C
1cgj_I D S L G R E A K C Y N E L N G C T L E Y R P V C G T D G D T Y P N E C V L C F E N R K R Q T S I L I Q K S G P C
1cho_I L A A V S V D C S E Y P K P A C T L E Y R P L C G S D N K T Y G N K C N F C N A V V E S N G T L T L S H F G K C
1cqu_A M K V I F L K D V K G K G K K G E I K N V A D G Y A N N F L F K Q G L A I E A T P A N L K A L E A Q K Q K E Q R
1edi_A A Q H D E A Q Q N A F Y Q V L N M P N L N A D Q R N G F I Q S L K D D P S Q S A N V L G E A Q K L N D S Q A P K
1edj_A A Q H D E A Q Q N A F Y Q V L N M P N L N A D Q R N G F I Q S L K D D P S Q S A N V L G E A Q K L N D S Q A P K
1edk_A A Q H D E A Q Q N A F Y Q V L N M P N L N A D Q R N G F I Q S L K D D P S Q S A N V L G E A Q K L N D S Q A P K
1edl_A A Q H D E A Q Q N A F Y Q V L N M P N L N A D Q R N G F I Q S L K D D P S Q S A N V L G E A Q K L N D S Q A P K
1em7_A T T Y K L I L N G K T L K G E T T T E A V D A E T A E R V F K E Y A K K N G V D G E W T Y D D A T K T F T V T E
1eq7_A S S N A K I D Q L S S D V Q T L N A K V D Q L S N D V N A M R S D V Q A A K D D A A R A N Q R L D N M A T K Y R
1f6u_A M Q K G N F R N Q R K T V K C F N C G K E G H I A K N C R A P R K K G C W K C G K E G H Q M K D C T E R Q A N X
1fcc_C T T Y K L V I N G K T L K G E T T T E A V D A A T A E K V F K Q Y A N D N G V D G E W T Y D D A T K T F T V T E
1fcc_D T T Y K L V I N G K T L K G E T T T E A V D A A T A E K V F K Q Y A N D N G V D G E W T Y D D A T K T F T V T E
1fcl_A T T F K L I I N G K T L K G E T T T E A V D A A T A E K V L K Q Y I N D N G I D G E W T Y D D A T K T W T V T E
1ffk_X T G A G T P S Q G K K N T T T H T K C R R C G E K S Y H T K K K V C S S C G F G K S A K R R D Y E W Q S K A G E
1gb1_A M T Y K L I L N G K T L K G E T T T E A V D A A T A E K V F K Q Y A N D N G V D G E W T Y D D A T K T F T V T E
1h9e_A P E F L E D P S V L T K D K L K S E L V A N N V T L P A G E Q R K D V Y V Q L Y L Q H L T A R N R P P L P A G T
1hpt_A D S L G R E A K C Y N E L N G C T Y E Y R P V C G T D G D T Y P N E C V L C F E N R K R Q T S I L I Q K S G P C
1jcc_A S S N A K I D Q L S S D A Q T A N A K A D Q A S N D A N A A R S D A Q A A K D D A A R A N Q R L D N M A T K Y R
1jcc_B S S N A K I D Q L S S D A Q T A N A K A D Q A S N D A N A A R S D A Q A A K D D A A R A N Q R L D N M A T K Y R
1jcc_C S S N A K I D Q L S S D A Q T A N A K A D Q A S N D A N A A R S D A Q A A K D D A A R A N Q R L D N M A T K Y R
1jj2_Z T G A G T P S Q G K K N T T T H T K C R R C G E K S Y H T K K K V C S S C G F G K S A K R R D Y E W Q S K A G E
1jy2_O K V E R K P P D A D G C L H A D P D L G V L C P T G C K L Q D T L V R Q E R P I R K S I E D L R N T V D S V S R
1jy2_R K V E R K P P D A D G C L H A D P D L G V L C P T G C K L Q D T L V R Q E R P I R K S I E D L R N T V D S V S R
1jy3_O K V E R K P P D A D G C L H A D P D L G V L C P T G C K L Q D T L V R Q E R P I R K S I E D L R N T V D S V S R
1jy3_R K V E R K P P D A D G C L H A D P D L G V L C P T G C K L Q D T L V R Q E R P I R K S I E D L R N T V D S V S R
1k73_2 T G A G T P S Q G K K N T T T H T K C R R C G E K S Y H T K K K V C S S C G F G K S A K R R D Y E W Q S K A G E
1k8a_2 T G A G T P S Q G K K N T T T H T K C R R C G E K S Y H T K K K V C S S C G F G K S A K R R D Y E W Q S K A G E
1k9m_2 T G A G T P S Q G K K N T T T H T K C R R C G E K S Y H T K K K V C S S C G F G K S A K R R D Y E W Q S K A G E
1kc8_2 T G A G T P S Q G K K N T T T H T K C R R C G E K S Y H T K K K V C S S C G F G K S A K R R D Y E W Q S K A G E
1kd1_2 T G A G T P S Q G K K N T T T H T K C R R C G E K S Y H T K K K V C S S C G F G K S A K R R D Y E W Q S K A G E
1kfm_A S S N A K I D Q L S S D V Q T L N A K V D Q L S N D V N A A R S D A Q A A K D D A A R A N Q R L D N M A T K Y R
1kfn_A S S N A K I D Q L S S D V Q T L N A K V D Q A S N D A N A A R S D A Q A A K D D A A R A N Q R L D N M A T K Y R
1kql_A M D K V E E L L S K N Y H L E N E V A R L K K L V D D L E D E L Y A Q K L K Y K A I S E E L D H A L N D M T S I
1kql_B M D K V E E L L S K N Y H L E N E V A R L K K L V D D L E D E L Y A Q K L K Y K A I S E E L D H A L N D M T S I
1kqs_Z T G A G T P S Q G K K N T T T H T K C R R C G E K S Y H T K K K V C S S C G F G K S A K R R D Y E W Q S K A G E
1l0l_K M L T R F L G P R Y R Q L A R N W V P T A Q L W G A V G A V G L V W A T D W R L I L D W V P Y I N G K F K K D D
1l0n_K M L T R F L G P R Y R Q L A R N W V P T A Q L W G A V G A V G L V S A T D S R L I L D W V P T I N G K F K K D D
1lgh_A S N P K D D Y K I W L V I N P S T W L P V I W I V A T V V A I A V H A A V L A A P G F N W I A L G A A K S A A K
1lgh_D S N P K D D Y K I W L V I N P S T W L P V I W I V A T V V A I A V H A A V L A A P G F N W I A L G A A K S A A K
1lgh_G S N P K D D Y K I W L V I N P S T W L P V I W I V A T V V A I A V H A A V L A A P G F N W I A L G A A K S A A K
1lgh_J S N P K D D Y K I W L V I N P S T W L P V I W I V A T V V A I A V H A A V L A A P G F N W I A L G A A K S A A K
1lqc_A M K P V T L Y D V A E Y A G V S Y Q T V S R V V N Q A S H V S A K T R E K V E A A M A E L N Y I P N R V A Q Q L
1m1k_2 T G A G T P S Q G K K N T T T H T K C R R C G E K S Y H T K K K V C S S C G F G K S A K R R D Y E W Q S K A G E
1m8b_A L A A V S V D C S E Y P K D A C T L E Y R P L C G S D N K T Y G N K C N F C N A V V E S N G T L T L S H F G K C
1m8c_A L A A V S V D C S E Y P K D A C T L E Y R P L C G S D N K T Y G N K C N F C N A V V E S N G T L T L S H F G K C
1m90_2 T G A G T P S Q G K K N T T T H T K C R R C G E K S Y H T K K K V C S S C G F G K S A K R R D Y E W Q S K A G E
1mpe_A M Q Y K V I L N G K T L K G E T T T E A V D A A T F E K V V K Q F F N D N G V D G E W T Y D D A T K T F T V T E
1mpe_B M Q Y K V I L N G K T L K G E T T T E A V D A A T F E K V V K Q F F N D N G V D G E W T Y D D A T K T F T V T E
1mpe_C M Q Y K V I L N G K T L K G E T T T E A V D A A T F E K V V K Q F F N D N G V D G E W T Y D D A T K T F T V T E
1mpe_D M Q Y K V I L N G K T L K G E T T T E A V D A A T F E K V V K Q F F N D N G V D G E W T Y D D A T K T F T V T E
1mvk_A M Q Y K V I L N G K T L K G E T T T E A V D A A T F E K V V K Q F F N D N G V D G E W T Y D D A T K T F T V T E
1mvk_B M Q Y K V I L N G K T L K G E T T T E A V D A A T F E K V V K Q F F N D N G V D G E W T Y D D A T K T F T V T E
1mvk_C M Q Y K V I L N G K T L K G E T T T E A V D A A T F E K V V K Q F F N D N G V D G E W T Y D D A T K T F T V T E
1mvk_D M Q Y K V I L N G K T L K G E T T T E A V D A A T F E K V V K Q F F N D N G V D G E W T Y D D A T K T F T V T E
1mvk_E M Q Y K V I L N G K T L K G E T T T E A V D A A T F E K V V K Q F F N D N G V D G E W T Y D D A T K T F T V T E
1mvk_F M Q Y K V I L N G K T L K G E T T T E A V D A A T F E K V V K Q F F N D N G V D G E W T Y D D A T K T F T V T E
1mvk_G M Q Y K V I L N G K T L K G E T T T E A V D A A T F E K V V K Q F F N D N G V D G E W T Y D D A T K T F T V T E
1mvk_H M Q Y K V I L N G K T L K G E T T T E A V D A A T F E K V V K Q F F N D N G V D G E W T Y D D A T K T F T V T E
1mvk_I M Q Y K V I L N G K T L K G E T T T E A V D A A T F E K V V K Q F F N D N G V D G E W T Y D D A T K T F T V T E
1mvk_J M Q Y K V I L N G K T L K G E T T T E A V D A A T F E K V V K Q F F N D N G V D G E W T Y D D A T K T F T V T E
1mvk_K M Q Y K V I L N G K T L K G E T T T E A V D A A T F E K V V K Q F F N D N G V D G E W T Y D D A T K T F T V T E
1mvk_L M Q Y K V I L N G K T L K G E T T T E A V D A A T F E K V V K Q F F N D N G V D G E W T Y D D A T K T F T V T E
1n87_A M L C A I S G K V P R R P V L S P K S R T I F E K S L L E Q Y V K D T G N D P I T N E P L S I E E I V E I V P S
1n8r_2 T G A G T P S Q G K K N T T T H T K C R R C G E K S Y H T K K K V C S S C G F G K S A K R R D Y E W Q S K A G E
```

```
1nay_A G P P G P P G P P G P P G P P G P P G P P G P P G P P G S G Y I P E A P R D G Q A Y V R K D G E W V L L S T F L
1nay_B G P P G P P G P P G P P G P P G P P G P P G P P G P P G S G Y I P E A P R D G Q A Y V R K D G E W V L L S T F L
1nay_C G P P G P P G P P G P P G P P G P P G P P G P P G P P G S G Y I P E A P R D G Q A Y V R K D G E W V L L S T F L
1nji_2 T G A G T P S Q G K K N T T T H T K C R R C G E K S Y H T K K K V C S S C G F G K S A K R R D Y E W Q S K A G E
1ntk_K M L T R F L G P R Y R Q L A R N W V P T A Q L W G A V G A V G L V W A T D W R L I L D W V P Y I N G K F K K D D
1ntm_K M L T R F L G P R Y R Q L A R N W V P T A Q L W G A V G A V G L V W A T D W R L I L D W V P Y I N G K F K K D D
1ntz_K M L T R F L G P R Y R Q L A R N W V P T A Q L W G A V G A V G L V W A T D W R L I L D W V P Y I N G K F K K D D
1nu1_K M L T R F L G P R Y R Q L A R N W V P T A Q L W G A V G A V G L V W A T D W R L I L D W V P Y I N G K F K K D D
1occ_K I H Q K R A P D F H D K Y G N A V L A S G A T F C V A V W V Y M A T Q I G I E W N P S P V G R V T P K E W R E Q
1occ_X I H Q K R A P D F H D K Y G N A V L A S G A T F C V A V W V Y M A T Q I G I E W N P S P V G R V T P K E W R E Q
1oco_K I H Q K R A P D F H D K Y G N A V L A S G A T F C V A V W V Y M A T Q I G I E W N P S P V G R V T P K E W R E Q
1oco_X I H Q K R A P D F H D K Y G N A V L A S G A T F C V A V W V Y M A T Q I G I E W N P S P V G R V T P K E W R E Q
1ocr_K I H Q K R A P D F H D K Y G N A V L A S G A T F C V A V W V Y M A T Q I G I E W N P S P V G R V T P K E W R E Q
1ocr_X I H Q K R A P D F H D K Y G N A V L A S G A T F C V A V W V Y M A T Q I G I E W N P S P V G R V T P K E W R E Q
1ocz_K I H Q K R A P D F H D K Y G N A V L A S G A T F C V A V W V Y M A T Q I G I E W N P S P V G R V T P K E W R E Q
1ocz_X I H Q K R A P D F H D K Y G N A V L A S G A T F C V A V W V Y M A T Q I G I E W N P S P V G R V T P K E W R E Q
1oks_A M A S R S V I R S I I K S S R L E E D R K R Y L M T L L D D I K G A N D L A K F H Q M L V K I I M K H H H H H H
1omt_A L A A V S V D C S E Y P K P A C T L E Y R P L C G S D N K T Y G N K C N F C N A V V E S N G T L T L S H F G K C
1omu_A L A A V S V D C S E Y P K P A C T L E Y R P L C G S D N K T Y G N K C N F C N A V V E S N G T L T L S H F G K C
1ovo_A L A A V S V D C S E Y P K P A C P K D Y R P V C G S D N K T Y S N K C N F C N A V V E S N G T L T L N H F G K C
1ovo_B L A A V S V D C S E Y P K P A C P K D Y R P V C G S D N K T Y S N K C N F C N A V V E S N G T L T L N H F G K C
1ovo_C L A A V S V D C S E Y P K P A C P K D Y R P V C G S D N K T Y S N K C N F C N A V V E S N G T L T L N H F G K C
1ovo_D L A A V S V D C S E Y P K P A C P K D Y R P V C G S D N K T Y S N K C N F C N A V V E S N G T L T L N H F G K C
1p7e_A M Q Y K L V I N G K T L K G E T T T K A V D A E T A E K A F K Q Y A N D N G V D G V W T Y D D A T K T F T V T E
1p7f_A M Q Y K L V I N G K T L K G E T T T K A V D A E T A E K A F K Q Y A N D N G V D G V W T Y D D A T K T F T V T E
1pga_A M T Y K L I L N G K T L K G E T T T E A V D A A T A E K V F K Q Y A N D N G V D G E W T Y D D A T K T F T V T E
1pgb_A M T Y K L I L N G K T L K G E T T T E A V D A A T A E K V F K Q Y A N D N G V D G E W T Y D D A T K T F T V T E
1ppf_I L A A V S V D C S E Y P K P A C T L E Y R P L C G S D N K T Y G N K C N F C N A V V E S N G T L T L S H F G K C
1pru_A M A T I K D V A K R A N V S T T T V S H V I N K T R F V A E E T R N A V W A A I K E L H Y S P S A V A R S L K V
1prv_A M A T I K D V A K R A N V S T T T V S H V I N K T R F V A E E T R N A V W A A I K E L H Y S P S A V A R S L K V
1q10_A M Q Y K V I L N G K T L K G E T T T E A V D A A T A E K V V K Q F F N D N G V D G E W T Y D D A T K T F T V T E
1q10_B M Q Y K V I L N G K T L K G E T T T E A V D A A T A E K V V K Q F F N D N G V D G E W T Y D D A T K T F T V T E
1q7y_2 T G A G T P S Q G K K N T T T H T K C R R C G E K S Y H T K K K V C S S C G F G K S A K R R D Y E W Q S K A G E
1q81_2 T G A G T P S Q G K K N T T T H T K C R R C G E K S Y H T K K K V C S S C G F G K S A K R R D Y E W Q S K A G E
1q82_2 T G A G T P S Q G K K N T T T H T K C R R C G E K S Y H T K K K V C S S C G F G K S A K R R D Y E W Q S K A G E
1q86_2 T G A G T P S Q G K K N T T T H T K C R R C G E K S Y H T K K K V C S S C G F G K S A K R R D Y E W Q S K A G E
1qvf_Z T G A G T P S Q G K K N T T T H T K C R R C G E K S Y H T K K K V C S S C G F G K S A K R R D Y E W Q S K A G E
1qvg_Z T G A G T P S Q G K K N T T T H T K C R R C G E K S Y H T K K K V C S S C G F G K S A K R R D Y E W Q S K A G E
1rso_B G L L A A E R A V S Q V L D S L E E I H A L T D S S E K D L D F L H S V F Q D Q H L H T L L D L Y D K I N T K S
1rso_D G L L A A E R A V S Q V L D S L E E I H A L T D S S E K D L D F L H S V F Q D Q H L H T L L D L Y D K I N T K S
1sqb_K M L T R F L G P R Y R Q L A R N W V P T A Q L W G A V G A V G L V W A T D S R L I L D W V P Y I N G K F K K D D
1sqp_K M L T R F L G P R Y R Q L A R N W V P T A G L W G A V G A V G L V W A T D W R L I L D W V P Y I N G K F K K D D
1sqq_K M L T R F L G P R Y R Q L A R N W V P T A S L W G A V G A V G L V W A T D W R L I L D W V P Y I N G K F K K D D
1sqv_K M L T R F L G P R Y R Q L A R N W V P T A S L W G A V G A V G L V W A T D W R L I L D W V P Y I N G K F K K D D
1sqx_K M L T R F L G P R Y R Q L A R N W V P T A S L W G A V G A V G L V W A T D W R L I L D W V P Y I N G K F K K D D
1tgs_I T S P Q R E A T C T S E V S G C P K I Y N P V C G T D G I T Y S N E C V L C S E N K K R Q T P V L I Q K S G P C
1ttw_B M T L Q Q V S T Q A I T S D E R R F A Y A V L E H A K N T I L N R Q D V A K L L P R A S N F E L S Q H H H H H H
1tur_A L A A V S V D C S E Y P K P A C T L E Y R P L C G S D N K T Y G N K C N F C N A V V E S N G T L T L S H F G K C
1tus_A L A A V S V D C S E Y P K P A C T L E Y R P L C G S D N K T Y G N K C N F C N A V V E S N G T L T L S H F G K C
1u6p_A A T V V S G Q K Q D R Q G G E R R R S Q L D R D Q C A Y C K E K G H W A K D C P K K P R G P R G P R P Q T S L L
1uua_A D F C L E P P Y T G P C K A R I I R Y F Y N A K A G L C Q T F V Y G G C R A K R N N F K S A E D C M R T C G G A
1uub_A D F C L E P P Y T G P C R A A I I R Y F Y N A K A G L C Q T F V Y G G C R A K S N N F K S A E D C M R T C G G A
1v54_K I H Q K R A P D F H D K Y G N A V L A S G A T F C V A V W V Y M A T Q I G I E W N P S P V G R V T P K E W R E Q
1v54_X I H Q K R A P D F H D K Y G N A V L A S G A T F C V A V W V Y M A T Q I G I E W N P S P V G R V T P K E W R E Q
1v55_K I H Q K R A P D F H D K Y G N A V L A S G A T F C V A V W V Y M A T Q I G I E W N P S P V G R V T P K E W R E Q
1v55_X I H Q K R A P D F H D K Y G N A V L A S G A T F C V A V W V Y M A T Q I G I E W N P S P V G R V T P K E W R E Q
1vex_A G S I P C L L S P W S E W S D C S V T C G K G M R T R Q R M L K S L A E L G D C N E D L E Q A E K C M L P E C P
1w2b_Z T G A G T P S Q G K K N T T T H T K C R R C G E K S Y H T K K K V C S S C G F G K S A K R R D Y E W Q S K A G E
1wwd_A A T V V S G Q K Q D R Q G G E R R R S Q L D R D Q C A Y C K E K G H W A K D C P K K P R G P R G P R P Q T S L L
1wwe_A A T V V S G Q K Q D R Q G G E R R R S Q L D R D Q C A Y C K E K G H W A K D C P K K P R G P R G P R P Q T S L L
1wwf_A A T V V S G Q K Q D R Q G G E R R R S Q L D R D Q C A Y C K E K G H W A K D C P K K P R G P R G P R P Q T S L L
1wwg_A A T V V S G Q K Q D R Q G G E R R R S Q L D R D Q C A Y C K E K G H W A K D C P K K P R G P R G P R P Q T S L L
1ykt_B R P D F C L E P P Y T G P C K A R I I R Y F Y N A K A G L C Q T F V Y G G C R A K R N N F K S A E D C M R T C G
1ylc_B R P D F A L E P P Y T G P C K A R I I R Y F Y N A K A G L A Q T F V Y G G C R A K R N N F K S A E D A M R T A G
1yld_B R P D F A L E P P Y T G P C K A R I I R Y F Y N A P D G L A Q T F V Y G G C R A K R N N F K S A E D A M R T A G
1zxh_A M Y Y L V V N K G Q N A F Y E T L T K A V D A E T A R N A F I Q S L K D D G V Q G V W T Y D D A T K T F T V Q A
2ayj_A M P L T D P A K L Q I V Q Q R V F L K K V C R K C G A L N P I R A T K C R R C H S T N L R L K K K E L P T K K G
2bap_D M R R A K L A K E K A E K E R L E K Q Q K R E Q L I D M N A E G D E T G V M D S L L E A L Q S G A A F R R K R G
2bap_C M R R A K L A K E K A E K E R L E K Q Q K R E Q L I D M N A E G D E T G V M D S L L E A L Q S G A A F R R K R G
2dyr_K I H Q K R A P D F H D K Y G N A V L A S G A T F C V A V W V Y M A T Q I G I E W N P S P V G R V T P K E W R E Q
2dyr_X I H Q K R A P D F H D K Y G N A V L A S G A T F C V A V W V Y M A T Q I G I E W N P S P V G R V T P K E W R E Q
2dys_K I H Q K R A P D F H D K Y G N A V L A S G A T F C V A V W V Y M A T Q I G I E W N P S P V G R V T P K E W R E Q
2dys_X I H Q K R A P D F H D K Y G N A V L A S G A T F C V A V W V Y M A T Q I G I E W N P S P V G R V T P K E W R E Q
2eij_K I H Q K R A P D F H D K Y G N A V L A S G A T F C V A V W V Y M A T Q I G I E W N P S P V G R V T P K E W R E Q
2eij_X I H Q K R A P D F H D K Y G N A V L A S G A T F C V A V W V Y M A T Q I G I E W N P S P V G R V T P K E W R E Q
2eik_K I H Q K R A P D F H D K Y G N A V L A S G A T F C V A V W V Y M A T Q I G I E W N P S P V G R V T P K E W R E Q
2eik_X I H Q K R A P D F H D K Y G N A V L A S G A T F C V A V W V Y M A T Q I G I E W N P S P V G R V T P K E W R E Q
2eil_K I H Q K R A P D F H D K Y G N A V L A S G A T F C V A V W V Y M A T Q I G I E W N P S P V G R V T P K E W R E Q
2eil_X I H Q K R A P D F H D K Y G N A V L A S G A T F C V A V W V Y M A T Q I G I E W N P S P V G R V T P K E W R E Q
2eim_K I H Q K R A P D F H D K Y G N A V L A S G A T F C V A V W V Y M A T Q I G I E W N P S P V G R V T P K E W R E Q
2eim_X I H Q K R A P D F H D K Y G N A V L A S G A T F C V A V W V Y M A T Q I G I E W N P S P V G R V T P K E W R E Q
2ein_K I H Q K R A P D F H D K Y G N A V L A S G A T F C V A V W V Y M A T Q I G I E W N P S P V G R V T P K E W R E Q
2ein_X I H Q K R A P D F H D K Y G N A V L A S G A T F C V A V W V Y M A T Q I G I E W N P S P V G R V T P K E W R E Q
2erw_A E V R N P C A C F R N Y V P V C G S D G K T Y G N P C M L N C A A Q T K V P G L K L V H E G R C Q R S N V E Q F
2ewl_A G S H M A E P Q R H K I L C V C C K C D G R I E L T V E S S A E D L R T L Q Q L F L S T L S F V C P W C A T N Q
2f8b_A G S H M A E P Q R H K I L C V C C K C D G R I E L T V E S S A E D L R T L Q Q L F L S T L S F V C P W C A T N Q
2f8b_B G S H M A E P Q R H K I L C V C C K C D G R I E L T V E S S A E D L R T L Q Q L F L S T L S F V C P W C A T N Q
2fs1_A M E A V D A N S L A Q A K E A A I K E L K Q Y G I G D Y Y I K L I N N A K T V E G V E S L K N E I L K A L P T E
2fyu_K M L T R F L G P R Y R Q L A R N W V P T A S L W G A V G A V G L V W A T D W R L I L D W V P Y I N G K F K K D D
2gb1_A M T Y K L I L N G K T L K G E T T T E A V D A A T A E K V F K Q Y A N D N G V D G E W T Y D D A T K T F T V T E
2ggv_A M S T D M W I E R T A D I S W E S D A E I T G S S E R V D V R L D D D G N F Q L M N D P G A G G G G S G G G R R
2gi9_A M Q Y K L I L N G K T L K G E T T T E A V D A A T A E K V F K Q Y A N D N G V D G E W T Y D D A T K T F T V T E
2gus_A S S N A K F D Q F S S D F Q T F N A K F D Q F S N D M N A F R S D F Q A F K D D F A R F N Q R F D N F A T K Y R
2guv_A S S N A K F D Q F S S D F Q T F N A K F D Q F S N D F N A F R S D F Q A F K D D F A R F N Q R F D N F A T K Y R
2guv_B S S N A K F D Q F S S D F Q T F N A K F D Q F S N D F N A F R S D F Q A F K D D F A R F N Q R F D N F A T K Y R
```

```
2guv_C S S N A K F D Q F S S D F Q T F N A K F D Q F S N D F N A F R S D F Q A F K D D F A R F N Q R F D N F A T K Y R
2guv_D S S N A K F D Q F S S D F Q T F N A K F D Q F S N D F N A F R S D F Q A F K D D F A R F N Q R F D N F A T K Y R
2guv_E S S N A K F D Q F S S D F Q T F N A K F D Q F S N D F N A F R S D F Q A F K D D F A R F N Q R F D N F A T K Y R
2j28_0 A V Q Q N K P T R S K R G M R R S H D A L T A V T S L S V D K T S G E K H L R H H I T A D G Y Y R G R K V I A K
2j52_A M T Y K L I L N G K T L K G E T T T E A V D A A T A E K V F K Q Y A N D N G V D G E W T Y D A A T K T F T V T E
2j53_A M T Y K L I L N G K T L K G E T T T E A V D A A T A E K V F K Q Y A N D N G V D G E W T Y D A A T K T F T V T E
2jsv_X M Q Y K L I L N G K T L K G E T T T E A V D A A T A E K V F K Q Y A N D N G V D G E W T Y D D A T K T F T V T E
2ju6_X M Q Y K L I L N G K T L K G E T T T E A V D A A T A E K V F K Q Y A N D N G V D G E W T Y D D A T K T F T V T E
2juj_A A T A S P Q L S S E I E N L M S Q G Y S Y Q D I Q K A L V I A Q N N I E M A K N I L R E F V S I S S P A H V A T
2jws_A T T Y K L I L N L K Q A K E E A I K E L V D A G I A E K Y I K L I A N A K T V E G V W T L K D E I L T F T V T E
2jwu_A T T Y K L I L N L K Q A K E E A I K E L V D A A T A E K Y F K L Y A N A K T V E G V W T Y K D E T K T F T V T E
2k0p_A M Q Y K L I L N G K T L K G E T T T E A V D A A T A E K V F K Q Y A N D N G V D G E W T Y D D A T K T F T V T E
2k6i_A M K H H H H H H P M G V S V M E A I K E V K L A E E Q A V K E I E E A K N R A E Q I K A E A I E E A K K L I A C
2kdl_A T T Y K L I L N L K Q A K E E A I K E L V D A G T A E K Y I K L I A N A K T V E G V W T L K D E I K T F T V T E
2kdm_A T T Y K L I L N L K Q A K E E A I K E A V D A G T A E K Y F K L I A N A K T V E G V W T Y K D E I K T F T V T E
2kel_A M Q T Q E Q S Q K K K Q K A V F G I Y M D K D L K T R L K V Y C A K N N L Q L T Q A I E E A I K E Y L Q K R N G
2kel_B M Q T Q E Q S Q K K K Q K A V F G I Y M D K D L K T R L K V Y C A K N N L Q L T Q A I E E A I K E Y L Q K R N G
2kft_A G S K N E D E C A V C R D G G E L I C C D G C P R A F H L A C L S P P L R E I P S G T W R C S S C L Q A T V Q E
2kk9_A K G L R R D L D A S R E A K K Q L E A E Q Q K L E E Q N K I S E A S R K G L R R D L D A S R E A K K Q V E K A X
2klk_A M Q Y K L I L N G K T L K G E T T T E A V D A A T A E K V F K Q Y F N D N G V D G E W T Y D D A T K T F T V T E
2klk_B M Q Y K L I L N G K T L K G E T T T E A V D A A T A E K V F K Q Y F N D N G V D G E W T Y D D A T K T F T V T E
2kmu_A G S M E R L R D V R E R L Q A W E R A F R R Q R G R R P S Q D D V E A A P E E T R A L Y R E Y R T L K R T T G Q
2kpi_A M P L E A G L L E I L A C P A C H A P L E E R D A E L I C T G Q D C G L A Y P V R D G I P V L L V D E A R R P E
2kq4_X M Q Y K L I L N G K T L K G E T T T E A V D A A T A E K V F K Q Y A N D N G V D G E W T Y D D A T K T F T V T E
2kwd_A M Q Y K L I L N G K T L K G E T T T E A V D A A T A E K V F K Q Y A N D N G V D G E W T Y D D A T K T F T V T E
2kwd_B M Q Y K L I L N G K T L K G E T T T E A V D A A T A E K V F K Q Y A N D N G V D G E W T Y D D A T K T F T V T E
2kwd_C M Q Y K L I L N G K T L K G E T T T E A V D A A T A E K V F K Q Y A N D N G V D G E W T Y D D A T K T F T V T E
2kwd_D M Q Y K L I L N G K T L K G E T T T E A V D A A T A E K V F K Q Y A N D N G V D G E W T Y D D A T K T F T V T E
2kwd_E M Q Y K L I L N G K T L K G E T T T E A V D A A T A E K V F K Q Y A N D N G V D G E W T Y D D A T K T F T V T E
2kwh_A G S E T Q A G I K E E I R R Q E F L L N S L H R D L Q G G I K D L S K E E R L W E V Q R I L T A L K R K L R E A
2kwi_B G S E T Q A G I K E E I R R Q E F L L N S L H R D L Q G G I K D L S K E E R L W E V Q R I L T A L K R K L R E A
2l12_A G E Q V F A V E S I R K K R V R K G K V E Y L V K W K G W P P K Y S T W E P E E H I L D P R L V M A Y E E K E E
2l1b_A G E Q V F A V E S I R K K R V R K G K V E Y L V K W K G W P P K Y S T W E P E E H I L D P R L V M A Y E E K E E
2le2_A M V Q N D F V D S Y D V T M L L Q D D D G K Q Y Y E Y H K G L S L S D F E V L Y G N T A D E I I K L R L D K V L
2le2_B M V Q N D F V D S Y D V T M L L Q D D D G K Q Y Y E Y H K G L S L S D F E V L Y G N T A D E I I K L R L D K V L
2lgi_A M Q Y K L I L N G K T L K G E T T T E A V D A A T A E K V F K Q Y A N D N G V D G E W T Y D D A T K T F T V T E
2lhc_A T T Y K L I L N L K Q A K E E A I K E L V D A G T A E K Y F K L I A N A K T V E G V W T L K D E I K T F T V T E
2lhd_A T T Y K L I L N L K Q A K E E A I K E L V D A G T A E K Y F K L I A N A K T V E G V W T Y K D E I K T F T V T E
2lhe_A T T Y K L I L N L K Q A K E E A I K E A V D A G I A E K Y F K L I A N A K T V E G V W T Y K D E I K T F T V T E
2lhg_A T T Y K L I L N L K Q A K E E A I K E L V D A G I A E K Y F K L I A N A K T V E G V W T Y K D E I K T F T V T E
2lum_A M Q Y K L V I N G K T L K G E T T T K A V D A E T A E K A F K Q Y A N D N G V D G V W T Y D D A T K T F T V T E
2m4y_A M T A Y R C P V C D Y T Y D E G K G D P R E G F P A G T R W D Q I P D D W C C P D C S V R E K V D F E R M G G K
2md7_B G A M G M R N L D E C E V C R D G G E L F C C D T C S R V F H E D C H I P P V E A E R T P W N C I F C R M K E S
2md8_C G A M G M R N L D E C E V C R D G G E L F C C D T C S R V F H E D C H I P P V E A E R T P W N C I F C R M K E S
2mdi_A G S D L P P P S P P A A A T I V L P P N W K T A R D P E G K I Y Y Y H V I T R Q T Q W D P P T W E S P G D D A S
2mdj_A G S D L P P P S P P K P K T I V L P P N W K T A R D P E G K I Y Y Y H V I T R Q T Q W D P P T W E S P G D D A S
2mh8_A N G D K G Y N G L A E A K E K A I K D L K I Y G I G E H Y I K L I E K A K Q V A A V E D L K D E I L K A H D R F
2mqv_A A T V V S G Q K Q D R Q G G E R R R S Q L D R D Q C A Y C K E K G H W A K D C P K K P R G P R G P R P Q T S L L
2ms0_A A T V V S G Q K Q D R Q G G E R R R S Q L D R D Q C A Y C K E K G H W A K D C P K K P R G P R G P R P Q T S L L
2ms0_C A T V V S G Q K Q D R Q G G E R R R S Q L D R D Q C A Y C K E K G H W A K D C P K K P R G P R G P R P Q T S L L
2ms1_A A T V V S G Q K Q D R Q G G E R R R S Q L D R D Q C A Y C K E K G H W A K D C P K K P R G P R G P R P Q T S L L
2mwx_A E E E F Q F L R C Q Q C Q A E A K C P K L L P C L H T L C S G C L E A S G M Q C P I C Q A P W P L G A D T P A L
2n69_A Q D K C K K V Y E N Y P V S K C Q L R I A N Q C N Y D C K L D K H A R S G E C F Y D E K R N L Q C I C D Y C E Y
2n7j_A M Q Y K L V I N G K T L K G E T T T K A V D A E T A E K A F K Q Y A N D N G V D G V W T Y D D A T K T F T V T E
2nc2_A K F G G E C S L K H N T C T Y L K G G K N H V V N C G S A A N K K C K S D R H H C E Y D E H H K R V D C Q T P V
2occ_K I H Q K R A P D F H D K Y G N A V L A S G A T F C V A V W V Y M A T Q I G I E W N P S P V G R V T P K E W R E Q
2occ_X I H Q K R A P D F H D K Y G N A V L A S G A T F C V A V W V Y M A T Q I G I E W N P S P V G R V T P K E W R E Q
2oed_A M Q Y K L V I N G K T L K G E T T T K A V D A E T A E K A F K Q Y A N D N G V D G V W T Y D D A T K T F T V T E
2on8_A M Q F K L I I N G K T L K G E I T L E A V D A A E A E K K F K Q Y A N D N G I D G E W T Y D D A T K T F T V T E
2onq_A M Q F K L I I N G K T L K G E I T I E A V D A A E A E K F F K Q Y A N D N G I D G E W T Y D D A T K T F T V T E
2ovo_A L A A V S V D C S E Y P K P A C T M E Y R P L C G S D N K T Y G N K C N F C N A V V E S N G T L T L S H F G K C
2p63_A G A M G S P E Y L S D E I F S A I N N N L P H A Y F K N L L F R L V A N M D R S E L S D L G T L I K D N L K R D
2p63_B G A M G S P E Y L S D E I F S A I N N N L P H A Y F K N L L F R L V A N M D R S E L S D L G T L I K D N L K R D
2p63_C G A M G S P E Y L S D E I F S A I N N N L P H A Y F K N L L F R L V A N M D R S E L S D L G T L I K D N L K R D
2p63_D G A M G S P E Y L S D E I F S A I N N N L P H A Y F K N L L F R L V A N M D R S E L S D L G T L I K D N L K R D
2qmt_A M Q Y K L I L N G K T L K G E T T T E A V D A A T A E K V F K Q Y A N D N G V D G E W T Y D D A T K T F T V T E
2rdo_0 A V Q Q N K P T R S K R G M R R S H D A L T A V T S L S V D K T S G E K H L R H H I T A D G Y Y R G R K V I A K
2rmm_A M Q Y K L I L N G K T L K G E T T T E A V D A A T A E K V F K Q Y F N D N G V D G E W T Y D D A T K T F T V T E
2rmm_B M Q Y K L I L N G K T L K G E T T T E A V D A A T A E K V F K Q Y F N D N G V D G E W T Y D D A T K T F T V T E
2vvk_A G S V Q A L F D F D P Q E D G E L G F R R G D F I H V M D N S D P N W W K G A C H G Q T G M F P R N Y V T P V N
2w6b_A G P L G S K S L V D T V Y A L K D E V Q E L R Q D N K K M K K S L E E E Q R A R K D L E K L V R K V L K N M N D
2x7n_D M K V E I D S F S G A K I Y P G R G T L F V R G D S K I F R F Q N S K S A S L F K Q R K N P R R I A W T V L F R
2y0n_E G A M G I Q E S E P E V T S F F P E P D D V E S L L I T P F L P V V A F G R P L P K L A P Q N F E L P W L D E R
2y0n_F G A M G I Q E S E P E V T S F F P E P D D V E S L L I T P F L P V V A F G R P L P K L A P Q N F E L P W L D E R
2y0n_G G A M G I Q E S E P E V T S F F P E P D D V E S L L I T P F L P V V A F G R P L P K L A P Q N F E L P W L D E R
2y0n_H G A M G I Q E S E P E V T S F F P E P D D V E S L L I T P F L P V V A F G R P L P K L A P Q N F E L P W L D E R
2y21_A H M S T I T R P I I E L S N T V D K I A E G N L E A E V P H Q N R A D E I G I L A K S I E R L R R S L K V A M E
2y21_B H M S T I T R P I I E L S N T V D K I A E G N L E A E V P H Q N R A D E I G I L A K S I E R L R R S L K V A M E
2y21_C H M S T I T R P I I E L S N T V D K I A E G N L E A E V P H Q N R A D E I G I L A K S I E R L R R S L K V A M E
2y21_D H M S T I T R P I I E L S N T V D K I A E G N L E A E V P H Q N R A D E I G I L A K S I E R L R R S L K V A M E
2y21_E H M S T I T R P I I E L S N T V D K I A E G N L E A E V P H Q N R A D E I G I L A K S I E R L R R S L K V A M E
2y21_F H M S T I T R P I I E L S N T V D K I A E G N L E A E V P H Q N R A D E I G I L A K S I E R L R R S L K V A M E
2y21_G H M S T I T R P I I E L S N T V D K I A E G N L E A E V P H Q N R A D E I G I L A K S I E R L R R S L K V A M E
2y21_H H M S T I T R P I I E L S N T V D K I A E G N L E A E V P H Q N R A D E I G I L A K S I E R L R R S L K V A M E
2y21_I H M S T I T R P I I E L S N T V D K I A E G N L E A E V P H Q N R A D E I G I L A K S I E R L R R S L K V A M E
2y21_J H M S T I T R P I I E L S N T V D K I A E G N L E A E V P H Q N R A D E I G I L A K S I E R L R R S L K V A M E
2y21_K H M S T I T R P I I E L S N T V D K I A E G N L E A E V P H Q N R A D E I G I L A K S I E R L R R S L K V A M E
2y21_L H M S T I T R P I I E L S N T V D K I A E G N L E A E V P H Q N R A D E I G I L A K S I E R L R R S L K V A M E
2ybb_K M L T R F L G P R Y R Q L A R N W V P T A Q L W G A V G A V G L V S A T D S R L I L D W V P Y I N G K F K K D D
2ybb_k M L T R F L G P R Y R Q L A R N W V P T A Q L W G A V G A V G L V S A T D S R L I L D W V P Y I N G K F K K D D
2ybb_V I H Q K R A P D F H D K Y G N A V L A S G A T F C V A V W V Y M A T Q I G I E W N P S P V G R V T P K E W R E Q
2yql_A G S S G S S G H E D F C S V C R K S G Q L L M C D T C S R V Y H L D C L D P P L K T I P K G M W I C P R C Q D Q
2yqq_A G S S G S S G L K C S T V V C V I C L E K P K Y R C P A C R V P Y C S V V C F R K H K E Q C N P E T S G P S S G
2ys0_A G S S G S S G W T C N K F R C G E K R L T R S L C A C S D D C K D Q G D C C I N Y S S V C Q G E K S S G P S S G
2zxw_K I H Q K R A P D F H D K Y G N A V L A S G A T F C V A V W V Y M A T Q I G I E W N P S P V G R V T P K E W R E Q
```

2zxw_X I H Q K R A P D F H D K Y G N A V L A S G A T F C V A V W V Y M A T Q I G I E W N P S P V G R V T P K E W R E Q
3a03_A M T S F S R S Q V L E L E R R F L R Q K Y L A S A E R A A L A K A L R M T D A Q V K T W F Q N R R T K W R R Q T
3abk_K I H Q K R A P D F H D K Y G N A V L A S G A T F C V A V W V Y M A T Q I G I E W N P S P V G R V T P K E W R E Q
3abk_X I H Q K R A P D F H D K Y G N A V L A S G A T F C V A V W V Y M A T Q I G I E W N P S P V G R V T P K E W R E Q
3abl_K I H Q K R A P D F H D K Y G N A V L A S G A T F C V A V W V Y M A T Q I G I E W N P S P V G R V T P K E W R E Q
3abl_X I H Q K R A P D F H D K Y G N A V L A S G A T F C V A V W V Y M A T Q I G I E W N P S P V G R V T P K E W R E Q
3abm_K I H Q K R A P D F H D K Y G N A V L A S G A T F C V A V W V Y M A T Q I G I E W N P S P V G R V T P K E W R E Q
3abm_X I H Q K R A P D F H D K Y G N A V L A S G A T F C V A V W V Y M A T Q I G I E W N P S P V G R V T P K E W R E Q
3ag1_K I H Q K R A P D F H D K Y G N A V L A S G A T F C V A V W V Y M A T Q I G I E W N P S P V G R V T P K E W R E Q
3ag1_X I H Q K R A P D F H D K Y G N A V L A S G A T F C V A V W V Y M A T Q I G I E W N P S P V G R V T P K E W R E Q
3ag2_K I H Q K R A P D F H D K Y G N A V L A S G A T F C V A V W V Y M A T Q I G I E W N P S P V G R V T P K E W R E Q
3ag2_X I H Q K R A P D F H D K Y G N A V L A S G A T F C V A V W V Y M A T Q I G I E W N P S P V G R V T P K E W R E Q
3ag3_K I H Q K R A P D F H D K Y G N A V L A S G A T F C V A V W V Y M A T Q I G I E W N P S P V G R V T P K E W R E Q
3ag3_X I H Q K R A P D F H D K Y G N A V L A S G A T F C V A V W V Y M A T Q I G I E W N P S P V G R V T P K E W R E Q
3ag4_K I H Q K R A P D F H D K Y G N A V L A S G A T F C V A V W V Y M A T Q I G I E W N P S P V G R V T P K E W R E Q
3ag4_X I H Q K R A P D F H D K Y G N A V L A S G A T F C V A V W V Y M A T Q I G I E W N P S P V G R V T P K E W R E Q
3asn_K I H Q K R A P D F H D K Y G N A V L A S G A T F C V A V W V Y M A T Q I G I E W N P S P V G R V T P K E W R E Q
3asn_X I H Q K R A P D F H D K Y G N A V L A S G A T F C V A V W V Y M A T Q I G I E W N P S P V G R V T P K E W R E Q
3aso_K I H Q K R A P D F H D K Y G N A V L A S G A T F C V A V W V Y M A T Q I G I E W N P S P V G R V T P K E W R E Q
3aso_X I H Q K R A P D F H D K Y G N A V L A S G A T F C V A V W V Y M A T Q I G I E W N P S P V G R V T P K E W R E Q
3bbx_0 A V Q Q N K P T R S K R G M R R S H D A L T A V T S L S V D K T S G E K H L R H H I T A D G Y Y R G R K V I A K
3cxc_Z T G A G T P S Q G K K N T T T H T K C R R C G E K S Y H T K K K V C S S C G F G K S A K R R D Y E W Q S K A G E
3e7k_A G A G S R V T F E R V E Q M S I Q I K E V G D R V N Y I K R S L Q S L D S Q I G H L Q D L S A L T V D T L K T L
3e7k_B G A G S R V T F E R V E Q M S I Q I K E V G D R V N Y I K R S L Q S L D S Q I G H L Q D L S A L T V D T L K T L
3e7k_C G A G S R V T F E R V E Q M S I Q I K E V G D R V N Y I K R S L Q S L D S Q I G H L Q D L S A L T V D T L K T L
3e7k_D G A G S R V T F E R V E Q M S I Q I K E V G D R V N Y I K R S L Q S L D S Q I G H L Q D L S A L T V D T L K T L
3e7k_E G A G S R V T F E R V E Q M S I Q I K E V G D R V N Y I K R S L Q S L D S Q I G H L Q D L S A L T V D T L K T L
3e7k_F G A G S R V T F E R V E Q M S I Q I K E V G D R V N Y I K R S L Q S L D S Q I G H L Q D L S A L T V D T L K T L
3e7k_G G A G S R V T F E R V E Q M S I Q I K E V G D R V N Y I K R S L Q S L D S Q I G H L Q D L S A L T V D T L K T L
3e7k_H G A G S R V T F E R V E Q M S I Q I K E V G D R V N Y I K R S L Q S L D S Q I G H L Q D L S A L T V D T L K T L
3fil_A M Q Y K L I L N G K T L K G V L T I E A V D A A T A E K V F K Q Y A N D L G V D G E W T Y D D A T K T F T V T E
3fil_B M Q Y K L I L N G K T L K G V L T I E A V D A A T A E K V F K Q Y A N D L G V D G E W T Y D D A T K T F T V T E
3g4s_1 T G A G T P S Q G K K N T T T H T K C R R C G E K S Y H T K K K V C S S C G F G K S A K R R D Y E W Q S K A G E
3g6e_1 T G A G T P S Q G K K N T T T H T K C R R C G E K S Y H T K K K V C S S C G F G K S A K R R D Y E W Q S K A G E
3g71_1 T G A G T P S Q G K K N T T T H T K C R R C G E K S Y H T K K K V C S S C G F G K S A K R R D Y E W Q S K A G E
3gb1_A M T Y K L I L N G K T L K G E T T T E A V D A A T A E K V F K Q Y A N D N G V D G E W T Y D D A T K T F T V T E
3he4_A G S Q K V A Q L K N R V A Y K L K E N A K L E N I V A R L E N D N A N L E K D I A N L E K D I A N L E R D V A R
3he4_C G S Q K V A Q L K N R V A Y K L K E N A K L E N I V A R L E N D N A N L E K D I A N L E K D I A N L E R D V A R
3he4_E G S Q K V A Q L K N R V A Y K L K E N A K L E N I V A R L E N D N A N L E K D I A N L E K D I A N L E R D V A R
3he4_G G S Q K V A Q L K N R V A Y K L K E N A K L E N I V A R L E N D N A N L E K D I A N L E K D I A N L E R D V A R
3hon_A G S S G V R L W A T R Q A M L G Q V H E V P E G W L I F V A E Q E E L Y V R V Q N G F R K V Q L E A R T P L P R
3hsh_A G S S G V R L W A T R Q A M L G Q V H E V P E G W L I F V A E Q E E L Y V R V Q N G F R K V Q L E A R T P L P R
3hsh_B G S S G V R L W A T R Q A M L G Q V H E V P E G W L I F V A E Q E E L Y V R V Q N G F R K V Q L E A R T P L P R
3hsh_C G S S G V R L W A T R Q A M L G Q V H E V P E G W L I F V A E Q E E L Y V R V Q N G F R K V Q L E A R T P L P R
3hsh_D G S S G V R L W A T R Q A M L G Q V H E V P E G W L I F V A E Q E E L Y V R V Q N G F R K V Q L E A R T P L P R
3hsh_E G S S G V R L W A T R Q A M L G Q V H E V P E G W L I F V A E Q E E L Y V R V Q N G F R K V Q L E A R T P L P R
3hsh_F G S S G V R L W A T R Q A M L G Q V H E V P E G W L I F V A E Q E E L Y V R V Q N G F R K V Q L E A R T P L P R
3j45_E X E A R T E V R K V I W P T R Q E T L H T T L I V A A V T A V M S L I L W G L D G I L V R L V S F I T G L R F X
3j46_E X E A R T E V R K V I W P T R Q E T L H T T L I V A A V T A V M S L I L W G L D G I L V R L V S F I T G L R F X
3j5l_0 A V Q Q N K P T R S K R G M R R S H D A L T A V T S L S V D K T S G E K H L R H H I T A D G Y Y R G R K V I A K
3j5l_H M Q V I L L D K V A N L G S L G D Q V N V K A G Y A R N F L V P Q G K A V P A T K K N I E F F E A R R A E L E A
3j6x_29 M A H E N V W F S H P R R Y G K G S R Q C R V C S S H T G L I R K Y G L N I C R Q C F R E K A N D I G F N K F R
3j6y_29 M A H E N V W F S H P R R Y G K G S R Q C R V C S S H T G L I R K Y G L N I C R Q C F R E K A N D I G F N K F R
3j77_29 M A H E N V W F S H P R R Y G K G S R Q C R V C S S H T G L I R K Y G L N I C R Q C F R E K A N D I G F N K F R
3j78_29 M A H E N V W F S H P R R Y G K G S R Q C R V C S S H T G L I R K Y G L N I C R Q C F R E K A N D I G F N K F R
3j7p_Sd M G P Q G F Y W S H P R K F G Q G S R S C R V C S N R H G L I R K Y G L N M C R Q C F R Q Y A K D I G F I K L D
3j7r_Sd M G P Q G F Y W S H P R K F G Q G S R S C R V C S N R H G L I R K Y G L N M C R Q C F R Q Y A K D I G F I K L D
3j80_d M A H E N V W Y S H P R K F G K G S R Q C R I S G S H S G L I R K Y G L N I D R Q S F R E K A N D I G F Y K Y R
3j81_d M A H E N V W Y S H P R K F G K G S R Q C R I S G S H S G L I R K Y G L N I D R Q S F R E K A N D I G F Y K Y R
3j9z_L1 A V Q Q N K P T R S K R G M R R S H D A L T A V T S L S V D K T S G E K H L R H H I T A D G Y Y R G R K V I A K
3ja1_L3 A V Q Q N K P T R S K R G M R R S H D A L T A V T S L S V D K T S G E K H L R H H I T A D G Y Y R G R K V I A K
3jam_d M A H E N V W Y S H P R K F G K G S R Q C R I S G S H S G L I R K Y G L N I D R Q S F R E K A N D I G F Y K Y R
3jap_d M A H E N V W Y S H P R K F G K G S R Q C R I S G S H S G L I R K Y G L N I D R Q S F R E K A N D I G F Y K Y R
3jaq_d M A H E N V W Y S H P R K F G K G S R Q C R I S G S H S G L I R K Y G L N I D R Q S F R E K A N D I G F Y K Y R
3kxt_A M K P V K V K T P A G K E A E L V P E K V W A L A P K G R K G V K I G L F K D P E T G K Y F R H K L P D D Y P I
3mmy_B T G T T I K F N P P T G T D T M V K A G V S T N I S T K H Q C I T A M K E Y E S K S L E E L R L E D Y Q A N R K
3mmy_D T G T T I K F N P P T G T D T M V K A G V S T N I S T K H Q C I T A M K E Y E S K S L E E L R L E D Y Q A N R K
3mmy_F T G T T I K F N P P T G T D T M V K A G V S T N I S T K H Q C I T A M K E Y E S K S L E E L R L E D Y Q A N R K
3mmy_H T G T T I K F N P P T G T D T M V K A G V S T N I S T K H Q C I T A M K E Y E S K S L E E L R L E D Y Q A N R K
3o6q_B M E R A F Q N R C E P R A A K P F K I L K K R S T T S V A S Y Q V S P H T A R I F K E N E R L I D E Y K R K K A
3o6q_D M E R A F Q N R C E P R A A K P F K I L K K R S T T S V A S Y Q V S P H T A R I F K E N E R L I D E Y K R K K A
3ovo_A L A A V S V D C S E Y P K P A C P K D Y R P V C G S D N K T Y S N K C N F C N A V V E S N G T L T L N H F G K C
3ow2_Z T G A G T P S Q G K K N T T T H T K C R R C G E K S Y H T K K K V C S S C G F G K S A K R R D Y E W Q S K A G E
3sgb_I L A A V S V D C S E Y P K P A C T L E Y R P L C G S D N K T Y G N K C N F C N A V V E S N G T L T L S H F G K C
3v3x_A M Q Y K L I L C G K T L K G E T T T E A V D A A T A E C V F K Q Y A N D N G V D G E W T Y D D A T K T F T V T E
3v3x_B M Q Y K L I L C G K T L K G E T T T E A V D A A T A E C V F K Q Y A N D N G V D G E W T Y D D A T K T F T V T E
3v3x_C M Q Y K L I L C G K T L K G E T T T E A V D A A T A E C V F K Q Y A N D N G V D G E W T Y D D A T K T F T V T E
3v3x_D M Q Y K L I L C G K T L K G E T T T E A V D A A T A E C V F K Q Y A N D N G V D G E W T Y D D A T K T F T V T E
3vpb_E M V V L K C P V C N G D V N V P D D A L P G E I V E H E C G A Q L E V Y N D H G R L A L R L A E Q V G E D W G E
3vpb_F M V V L K C P V C N G D V N V P D D A L P G E I V E H E C G A Q L E V Y N D H G R L A L R L A E Q V G E D W G E
3wg7_K I H Q K R A P D F H D K Y G N A V L A S G A T F C V A V W V Y M A T Q I G I E W N P S P V G R V T P K E W R E Q
3wg7_X I H Q K R A P D F H D K Y G N A V L A S G A T F C V A V W V Y M A T Q I G I E W N P S P V G R V T P K E W R E Q
3x2q_K I H Q K R A P D F H D K Y G N A V L A S G A T F C V A V W V Y M A T Q I G I E W N P S P V G R V T P K E W R E Q
3x2q_X I H Q K R A P D F H D K Y G N A V L A S G A T F C V A V W V Y M A T Q I G I E W N P S P V G R V T P K E W R E Q
3zoq_B M V Q N D F V D S Y D V T M L L Q D D D G K Q Y Y E Y H K G L S L S D F E V L Y G N T A D E I I K L R L D K V L
3zoq_C M V Q N D F V D S Y D V T M L L Q D D D G K Q Y Y E Y H K G L S L S D F E V L Y G N T A D E I I K L R L D K V L
4ce4_o X X X X X X X X X X X X X X X X X X X X X X X X X X X X X X X X X X X X X X X X X X X X X X X X X X X X X X X X
4csu_3 A V Q Q N K P T R S K R G M R R S H D A L T A V T S L S V D K T S G E K H L R H H I T A D G Y Y R G R K V I A K
4d5l_d M G H Q Q L Y W S H P R K F G Q G S R S C R V C S N R H G L I R K Y G L N M C R Q C F R Q Y A K D I G F I K L D
4d61_d M G H Q Q L Y W S H P R K F G Q G S R S C R V C S N R H G L I R K Y G L N M C R Q C F R Q Y A K D I G F I K L D
4dnn_A G S T P D Y L M Q L M N D K K L M S S L P N F S G I F N H L E R L L D E E I S R V R K D M Y N D T L N G S T E K
4dnn_B G S T P D Y L M Q L M N D K K L M S S L P N F S G I F N H L E R L L D E E I S R V R K D M Y N D T L N G S T E K
4e5r_A P P T E P L P D G W I M T F H N S G L P V Y L H R E T R V V T W S R P Y F L G T G S I R K H D P P I S S I P C L
4gln_D T Y K L I L N G K T L K G E T T T E A V D V F D A F D V F F V Y A A S N F S D F D D W T Y D D A T K T F T V T E
4gln_H T Y K L I L N G K T L K G E T T T E A V D V F D A F D V F F V Y A A S N F S D F D D W T Y D D A T K T F T V T E

4gls_C T Y K L I L N G K T L K G E T T T E A V D V F D A F D V F F V Y A A S N F S D F D D W T Y D D A T K T F T V T E
4gls_G T Y K L I L N G K T L K G E T T T E A V D V F D A F D V F F V Y A A S N F S D F D D W T Y D D A T K T F T V T E
4gls_D T Y K L I L N G K T L K G E T T T E A V D V F D A F D V F F V Y A A S N F S D F D D W T Y D D A T K T F T V T E
4gls_H T Y K L I L N G K T L K G E T T T E A V D V F D A F D V F F V Y A A S N F S D F D D W T Y D D A T K T F T V T E
4kzx_d M G H Q Q L Y W S H P R K F G Q G S R S C R V C S N R H G L I R K Y G L N M C R Q C F R Q Y A K D I G F I K L D
4kzy_d M G H Q Q L Y W S H P R K F G Q G S R S C R V C S N R H G L I R K Y G L N M C R Q C F R Q Y A K D I G F I K L D
4kzz_d M G H Q Q L Y W S H P R K F G Q G S R S C R V C S N R H G L I R K Y G L N M C R Q C F R Q Y A K D I G F I K L D
4l9u_A S E L R H L R L P T Y Q E L E Q E I N T L K A D N D A L K I Q L K Y A Q K K I E S L Q L E K S N H V L A Q M E Q
4l9u_B S E L R H L R L P T Y Q E L E Q E I N T L K A D N D A L K I Q L K Y A Q K K I E S L Q L E K S N H V L A Q M E Q
4mn3_A G E Q V F A V E S I R K K R V R K G K V E Y L V K W K G W P P K Y S T W E P E E H I L D P R L V M A Y E E K E E
4ovo_A L A A V S V D C S E Y P K P A C T M E Y R P L C G S D N K T Y G N K C N F C N A V V E S N G T L T L S H F G K C
4riq_A G K V D L Q S L P T R A Y L D Q T V V P I L L Q G L A V L A K E R P P N P I E F L A S Y L L K N K A Q F E D R N
4riq_B G K V D L Q S L P T R A Y L D Q T V V P I L L Q G L A V L A K E R P P N P I E F L A S Y L L K N K A Q F E D R N
4riq_D G K V D L Q S L P T R A Y L D Q T V V P I L L Q G L A V L A K E R P P N P I E F L A S Y L L K N K A Q F E D R N
4riq_E G K V D L Q S L P T R A Y L D Q T V V P I L L Q G L A V L A K E R P P N P I E F L A S Y L L K N K A Q F E D R N
4riq_G G K V D L Q S L P T R A Y L D Q T V V P I L L Q G L A V L A K E R P P N P I E F L A S Y L L K N K A Q F E D R N
4riq_H G K V D L Q S L P T R A Y L D Q T V V P I L L Q G L A V L A K E R P P N P I E F L A S Y L L K N K A Q F E D R N
4riq_J G K V D L Q S L P T R A Y L D Q T V V P I L L Q G L A V L A K E R P P N P I E F L A S Y L L K N K A Q F E D R N
4riq_K G K V D L Q S L P T R A Y L D Q T V V P I L L Q G L A V L A K E R P P N P I E F L A S Y L L K N K A Q F E D R N
4riq_M G K V D L Q S L P T R A Y L D Q T V V P I L L Q G L A V L A K E R P P N P I E F L A S Y L L K N K A Q F E D R N
4riq_N G K V D L Q S L P T R A Y L D Q T V V P I L L Q G L A V L A K E R P P N P I E F L A S Y L L K N K A Q F E D R N
4riq_P G K V D L Q S L P T R A Y L D Q T V V P I L L Q G L A V L A K E R P P N P I E F L A S Y L L K N K A Q F E D R N
4riq_Q G K V D L Q S L P T R A Y L D Q T V V P I L L Q G L A V L A K E R P P N P I E F L A S Y L L K N K A Q F E D R N
4riq_S G K V D L Q S L P T R A Y L D Q T V V P I L L Q G L A V L A K E R P P N P I E F L A S Y L L K N K A Q F E D R N
4riq_T G K V D L Q S L P T R A Y L D Q T V V P I L L Q G L A V L A K E R P P N P I E F L A S Y L L K N K A Q F E D R N
4riq_V G K V D L Q S L P T R A Y L D Q T V V P I L L Q G L A V L A K E R P P N P I E F L A S Y L L K N K A Q F E D R N
4riq_Z G K V D L Q S L P T R A Y L D Q T V V P I L L Q G L A V L A K E R P P N P I E F L A S Y L L K N K A Q F E D R N
4u1u_B0 A V Q Q N K P T R S K R G M R R S H D A L T A V T S L S V D K T S G E K H L R H H I T A D G Y Y R G R K V I A K
4u1u_D0 A V Q Q N K P T R S K R G M R R S H D A L T A V T S L S V D K T S G E K H L R H H I T A D G Y Y R G R K V I A K
4u1v_B0 A V Q Q N K P T R S K R G M R R S H D A L T A V T S L S V D K T S G E K H L R H H I T A D G Y Y R G R K V I A K
4u1v_D0 A V Q Q N K P T R S K R G M R R S H D A L T A V T S L S V D K T S G E K H L R H H I T A D G Y Y R G R K V I A K
4u20_B0 A V Q Q N K P T R S K R G M R R S H D A L T A V T S L S V D K T S G E K H L R H H I T A D G Y Y R G R K V I A K
4u20_D0 A V Q Q N K P T R S K R G M R R S H D A L T A V T S L S V D K T S G E K H L R H H I T A D G Y Y R G R K V I A K
4u24_B0 A V Q Q N K P T R S K R G M R R S H D A L T A V T S L S V D K T S G E K H L R H H I T A D G Y Y R G R K V I A K
4u24_D0 A V Q Q N K P T R S K R G M R R S H D A L T A V T S L S V D K T S G E K H L R H H I T A D G Y Y R G R K V I A K
4u25_B0 A V Q Q N K P T R S K R G M R R S H D A L T A V T S L S V D K T S G E K H L R H H I T A D G Y Y R G R K V I A K
4u25_D0 A V Q Q N K P T R S K R G M R R S H D A L T A V T S L S V D K T S G E K H L R H H I T A D G Y Y R G R K V I A K
4u26_B0 A V Q Q N K P T R S K R G M R R S H D A L T A V T S L S V D K T S G E K H L R H H I T A D G Y Y R G R K V I A K
4u26_D0 A V Q Q N K P T R S K R G M R R S H D A L T A V T S L S V D K T S G E K H L R H H I T A D G Y Y R G R K V I A K
4u27_B0 A V Q Q N K P T R S K R G M R R S H D A L T A V T S L S V D K T S G E K H L R H H I T A D G Y Y R G R K V I A K
4u27_D0 A V Q Q N K P T R S K R G M R R S H D A L T A V T S L S V D K T S G E K H L R H H I T A D G Y Y R G R K V I A K
4ug0_Sd M G H Q Q L Y W S H P R K F G Q G S R S C R V C S N R H G L I R K Y G L N M C R Q C F R Q Y A K D I G F I K L D
4ujc_Cd M G H Q Q L Y W S H P R K F G Q G S R S C R V C S N R H G L I R K Y G L N M C R Q C F R Q Y A K D I G F I K L D
4ujd_Cd M G H Q Q L Y W S H P R K F G Q G S R S C R V C S N R H G L I R K Y G L N M C R Q C F R Q Y A K D I G F I K L D
4uje_Bd M G H Q Q L Y W S H P R K F G Q G S R S C R V C S N R H G L I R K Y G L N M C R Q C F R Q Y A K D I G F I K L D
4uy8_0 A V Q Q N K P T R S K R G M R R S H D A L T A V T S L S V D K T S G E K H L R H H I T A D G Y Y R G R K V I A K
4v47_AZ A V Q Q N K P T R S K R G M R R S H D A L T A V T S L S V D K T S G E K H L R H H I T A D G Y Y R G R K V I A K
4v48_AZ A V Q Q N K P T R S K R G M R R S H D A L T A V T S L S V D K T S G E K H L R H H I T A D G Y Y R G R K V I A K
4v4b_BS M K V E I D S F S G A K I Y P G R G T L F V R G D S K I F R F Q N S K S A S L F K Q R K N P R R I A W T V L F R
4v4n_BP M A K A D Y N K R K P R K F G K G A R R C I R C G Q Y G P I I R I Q G L M L C R H C F R E V A P K L G F R K Y E
4v4q_B0 A V Q Q N K P T R S K R G M R R S H D A L T A V T S L S V D K T S G E K H L R H H I T A D G Y Y R G R K V I A K
4v4q_D0 A V Q Q N K P T R S K R G M R R S H D A L T A V T S L S V D K T S G E K H L R H H I T A D G Y Y R G R K V I A K
4v4v_BX T I K I T Q T R S A I G R L P K H K A T L L G L G L R R I G H T V E R E D T P A I R G M I N A V S F M V K V E E
4v4w_BX T I K I T Q T R S A I G R L P K H K A T L L G L G L R R I G H T V E R E D T P A I R G M I N A V S F M V K V E E
4v50_B0 A V Q Q N K P T R S K R G M R R S H D A L T A V T S L S V D K T S G E K H L R H H I T A D G Y Y R G R K V I A K
4v50_D0 A V Q Q N K P T R S K R G M R R S H D A L T A V T S L S V D K T S G E K H L R H H I T A D G Y Y R G R K V I A K
4v52_B0 A V Q Q N K P T R S K R G M R R S H D A L T A V T S L S V D K T S G E K H L R H H I T A D G Y Y R G R K V I A K
4v52_D0 A V Q Q N K P T R S K R G M R R S H D A L T A V T S L S V D K T S G E K H L R H H I T A D G Y Y R G R K V I A K
4v53_B0 A V Q Q N K P T R S K R G M R R S H D A L T A V T S L S V D K T S G E K H L R H H I T A D G Y Y R G R K V I A K
4v53_D0 A V Q Q N K P T R S K R G M R R S H D A L T A V T S L S V D K T S G E K H L R H H I T A D G Y Y R G R K V I A K
4v54_B0 A V Q Q N K P T R S K R G M R R S H D A L T A V T S L S V D K T S G E K H L R H H I T A D G Y Y R G R K V I A K
4v54_D0 A V Q Q N K P T R S K R G M R R S H D A L T A V T S L S V D K T S G E K H L R H H I T A D G Y Y R G R K V I A K
4v55_B0 A V Q Q N K P T R S K R G M R R S H D A L T A V T S L S V D K T S G E K H L R H H I T A D G Y Y R G R K V I A K
4v55_D0 A V Q Q N K P T R S K R G M R R S H D A L T A V T S L S V D K T S G E K H L R H H I T A D G Y Y R G R K V I A K
4v56_B0 A V Q Q N K P T R S K R G M R R S H D A L T A V T S L S V D K T S G E K H L R H H I T A D G Y Y R G R K V I A K
4v56_D0 A V Q Q N K P T R S K R G M R R S H D A L T A V T S L S V D K T S G E K H L R H H I T A D G Y Y R G R K V I A K
4v57_B0 A V Q Q N K P T R S K R G M R R S H D A L T A V T S L S V D K T S G E K H L R H H I T A D G Y Y R G R K V I A K
4v57_D0 A V Q Q N K P T R S K R G M R R S H D A L T A V T S L S V D K T S G E K H L R H H I T A D G Y Y R G R K V I A K
4v5b_A0 A V Q Q N K P T R S K R G M R R S H D A L T A V T S L S V D K T S G E K H L R H H I T A D G Y Y R G R K V I A K
4v5b_C0 A V Q Q N K P T R S K R G M R R S H D A L T A V T S L S V D K T S G E K H L R H H I T A D G Y Y R G R K V I A K
4v5h_B3 A V Q Q N K P T R S K R G M R R S H D A L T A V T S L S V D K T S G E K H L R H H I T A D G Y Y R G R K V I A K
4v5y_B0 A V Q Q N K P T R S K R G M R R S H D A L T A V T S L S V D K T S G E K H L R H H I T A D G Y Y R G R K V I A K
4v5y_D0 A V Q Q N K P T R S K R G M R R S H D A L T A V T S L S V D K T S G E K H L R H H I T A D G Y Y R G R K V I A K
4v5z_An M G H Q Q L Y W S H P R K F G Q G S R S C R V C S N R H G L I R K Y G L N M C R Q C F R Q Y A K D I G F I K L D
4v64_B0 A V Q Q N K P T R S K R G M R R S H D A L T A V T S L S V D K T S G E K H L R H H I T A D G Y Y R G R K V I A K
4v64_D0 A V Q Q N K P T R S K R G M R R S H D A L T A V T S L S V D K T S G E K H L R H H I T A D G Y Y R G R K V I A K
4v69_B0 A V Q Q N K P T R S K R G M R R S H D A L T A V T S L S V D K T S G E K H L R H H I T A D G Y Y R G R K V I A K
4v6i_AN M A H E N V W F S H P R R Y G K G S R Q C R V C S S H T G L I R K Y G L N I C R Q C F R E K A N D I G F N K F R
4v6m_B0 A V Q Q N K P T R S K R G M R R S H D A L T A V T S L S V D K T S G E K H L R H H I T A D G Y Y R G R K V I A K
4v6n_A3 A V Q Q N K P T R S K R G M R R S H D A L T A V T S L S V D K T S G E K H L R H H I T A D G Y Y R G R K V I A K
4v6o_B3 A V Q Q N K P T R S K R G M R R S H D A L T A V T S L S V D K T S G E K H L R H H I T A D G Y Y R G R K V I A K
4v6p_B3 A V Q Q N K P T R S K R G M R R S H D A L T A V T S L S V D K T S G E K H L R H H I T A D G Y Y R G R K V I A K
4v6q_B3 A V Q Q N K P T R S K R G M R R S H D A L T A V T S L S V D K T S G E K H L R H H I T A D G Y Y R G R K V I A K
4v6r_B3 A V Q Q N K P T R S K R G M R R S H D A L T A V T S L S V D K T S G E K H L R H H I T A D G Y Y R G R K V I A K
4v6s_A3 A V Q Q N K P T R S K R G M R R S H D A L T A V T S L S V D K T S G E K H L R H H I T A D G Y Y R G R K V I A K
4v6t_B0 A V Q Q N K P T R S K R G M R R S H D A L T A V T S L S V D K T S G E K H L R H H I T A D G Y Y R G R K V I A K
4v6u_AP M A K A D Y N K R K P R K F G K G A R R C I R C G Q Y G P I I R I Q G L M L C R H C F R E V A P K L G F R K Y E
4v6v_B5 A V Q Q N K P T R S K R G M R R S H D A L T A V T S L S V D K T S G E K H L R H H I T A D G Y Y R G R K V I A K
4v6w_Ad M G F A T L W Y S H P R K Y G Q G S R C C R A C S N R H G L I R K Y G L N I C R Q C F R E Y A N D I G F K K L D
4v6x_Ad M G H Q Q L Y W S H P R K F G Q G S R S C R V C S N R H G L I R K Y G L N M C R Q C F R Q Y A K D I G F I K L D
4v7c_B3 A V Q Q N K P T R S K R G M R R S H D A L T A V T S L S V D K T S G E K H L R H H I T A D G Y Y R G R K V I A K
4v7d_A4 A V Q Q N K P T R S K R G M R R S H D A L T A V T S L S V D K T S G E K H L R H H I T A D G Y Y R G R K V I A K
4v7e_Bd M G H S N V W N S H P K N Y G A G S R V C R V C G N S H G L I R K Y G L M C C R Q C F R S D A K D I G F I K Y R
4v7r_AS M A H E N V W F S H P R R Y G K G S R Q C R V C S S H T G L I R K Y G L N I C R Q C F R E K A N D I G F N K F R
4v7r_CS M A H E N V W F S H P R R Y G K G S R Q C R V C S S H T G L I R K Y G L N I C R Q C F R E K A N D I G F N K F R

4v7s_B0 A V Q Q N K P T R S K R G M R R S H D A L T A V T S L S V D K T S G E K H L R H H I T A D G Y Y R G R K V I A K
4v7s_D0 A V Q Q N K P T R S K R G M R R S H D A L T A V T S L S V D K T S G E K H L R H H I T A D G Y Y R G R K V I A K
4v7t_B0 A V Q Q N K P T R S K R G M R R S H D A L T A V T S L S V D K T S G E K H L R H H I T A D G Y Y R G R K V I A K
4v7t_D0 A V Q Q N K P T R S K R G M R R S H D A L T A V T S L S V D K T S G E K H L R H H I T A D G Y Y R G R K V I A K
4v7u_B0 A V Q Q N K P T R S K R G M R R S H D A L T A V T S L S V D K T S G E K H L R H H I T A D G Y Y R G R K V I A K
4v7u_D0 A V Q Q N K P T R S K R G M R R S H D A L T A V T S L S V D K T S G E K H L R H H I T A D G Y Y R G R K V I A K
4v7v_B0 A V Q Q N K P T R S K R G M R R S H D A L T A V T S L S V D K T S G E K H L R H H I T A D G Y Y R G R K V I A K
4v7v_D0 A V Q Q N K P T R S K R G M R R S H D A L T A V T S L S V D K T S G E K H L R H H I T A D G Y Y R G R K V I A K
4v88_Ad M A H E N V W F S H P R R Y G K G S R Q C R V C S S H T G L I R K Y G L N I C R Q C F R E K A N D I G F N K F R
4v88_Cd M A H E N V W F S H P R R Y G K G S R Q C R V C S S H T G L I R K Y G L N I C R Q C F R E K A N D I G F N K F R
4v8y_A3 M A H E N V W F S H P R R Y G K G S R Q C R V C S S H T G L I R K Y G L N I C R Q C F R E K A N D I G F N K F R
4v8z_A3 M A H E N V W F S H P R R Y G K G S R Q C R V C S S H T G L I R K Y G L N I C R Q C F R E K A N D I G F N K F R
4v9d_C0 A V Q Q N K P T R S K R G M R R S H D A L T A V T S L S V D K T S G E K H L R H H I T A D G Y Y R G R K V I A K
4v9d_D0 A V Q Q N K P T R S K R G M R R S H D A L T A V T S L S V D K T S G E K H L R H H I T A D G Y Y R G R K V I A K
4v9f_6 M S T Y T V R G S F P A R D G P Q Q F E K E V E A P N E N V A E E R V Y S D F G S Q H N L K R T Q I T I E E V A
4wf1_B0 A V Q Q N K P T R S K R G M R R S H D A L T A V T S L S V D K T S G E K H L R H H I T A D G Y Y R G R K V I A K
4wf1_D0 A V Q Q N K P T R S K R G M R R S H D A L T A V T S L S V D K T S G E K H L R H H I T A D G Y Y R G R K V I A K
4wh4_A M Q Y K L H L H G K T L K G E T T T E A V D A A T A E H V F K H Y A N D N G V D G E W T Y D D A T K T F T V T E
4wh4_B M Q Y K L H L H G K T L K G E T T T E A V D A A T A E H V F K H Y A N D N G V D G E W T Y D D A T K T F T V T E
4www_R0 A V Q Q N K P T R S K R G M R R S H D A L T A V T S L S V D K T S G E K H L R H H I T A D G Y Y R G R K V I A K
4www_Y0 A V Q Q N K P T R S K R G M R R S H D A L T A V T S L S V D K T S G E K H L R H H I T A D G Y Y R G R K V I A K
4xrs_G K P R T I Y S S Y Q L A A L Q R R F Q K A Q Y L A L P E R A E L A A Q L G L T Q T Q V K I W F Q N R R S K F K K
4xrs_I K P R T I Y S S Y Q L A A L Q R R F Q K A Q Y L A L P E R A E L A A Q L G L T Q T Q V K I W F Q N R R S K F K K
4ybb_AU P V I K V R E N E P F D V A L R R F K R S C E K A G V L A E V R R R E F Y E K P T T E R K R A K A S A V K R H A
4ybb_BU P V I K V R E N E P F D V A L R R F K R S C E K A G V L A E V R R R E F Y E K P T T E R K R A K A S A V K R H A
4ybb_C1 A V Q Q N K P T R S K R G M R R S H D A L T A V T S L S V D K T S G E K H L R H H I T A D G Y Y R G R K V I A K
4ybb_D1 A V Q Q N K P T R S K R G M R R S H D A L T A V T S L S V D K T S G E K H L R H H I T A D G Y Y R G R K V I A K
4z6y_C R G Q L Q A A E S R Y E A Q K R I T Q V F E L E I L D L Y G R L E K D G L L K K L E E E K A E A A A E A A E E R L
4z6y_H R G Q L Q A A E S R Y E A Q K R I T Q V F E L E I L D L Y G R L E K D G L L K K L E E E K A E A A A E A A E E R L
4z6y_F R G Q L Q A A E S R Y E A Q K R I T Q V F E L E I L D L Y G R L E K D G L L K K L E E E K A E A A A E A A E E R L
4z6y_D R G Q L Q A A E S R Y E A Q K R I T Q V F E L E I L D L Y G R L E K D G L L K K L E E E K A E A A A E A A E E R L
5a2q_e G S L A R A G K V R G Q T P K V A K Q E K K K K K T G R A K R R M Q Y N R R F V N V V P T F G K K K G P N A N S
5aj0_Bd M G H Q Q L Y W S H P R K F G Q G S R S C R V C S N R H G L I R K Y G L N M C R Q C F R Q Y A K D I G F I K L D
5aka_0 A V Q Q N K P T R S K R G M R R S H D A L T A V T S L S V D K T S G E K H L R H H I T A D G Y Y R G R K V I A K
5ayw_C R Q V S G D E A Y L E A A P L A E L H A P A G M I L P V T S G D Y A I P V T N G S G A V G K A L D I R P P A Q P
5b1a_K I H Q K R A P D F H D K Y G N A V L A S G A T F C V A V W V Y M A T Q I G I E W N P S P V G R V T P K E W R E Q
5b1a_X I H Q K R A P D F H D K Y G N A V L A S G A T F C V A V W V Y M A T Q I G I E W N P S P V G R V T P K E W R E Q
5b1b_K I H Q K R A P D F H D K Y G N A V L A S G A T F C V A V W V Y M A T Q I G I E W N P S P V G R V T P K E W R E Q
5b1b_X I H Q K R A P D F H D K Y G N A V L A S G A T F C V A V W V Y M A T Q I G I E W N P S P V G R V T P K E W R E Q
5b3s_K I H Q K R A P D F H D K Y G N A V L A S G A T F C V A V W V Y M A T Q I G I E W N P S P V G R V T P K E W R E Q
5b3s_X I H Q K R A P D F H D K Y G N A V L A S G A T F C V A V W V Y M A T Q I G I E W N P S P V G R V T P K E W R E Q
5bmg_A M Q Y K L I L N G K T L K G C T T T E A V D A A T A E K V F K Q Y A N D N G V D G E W T Y D D A T K T F T V T E
5bmg_B M Q Y K L I L N G K T L K G C T T T E A V D A A T A E K V F K Q Y A N D N G V D G E W T Y D D A T K T F T V T E
5bmg_C M Q Y K L I L N G K T L K G C T T T E A V D A A T A E K V F K Q Y A N D N G V D G E W T Y D D A T K T F T V T E
5bmg_D M Q Y K L I L N G K T L K G C T T T E A V D A A T A E K V F K Q Y A N D N G V D G E W T Y D D A T K T F T V T E
5bmg_E M Q Y K L I L N G K T L K G C T T T E A V D A A T A E K V F K Q Y A N D N G V D G E W T Y D D A T K T F T V T E
5bmg_F M Q Y K L I L N G K T L K G C T T T E A V D A A T A E K V F K Q Y A N D N G V D G E W T Y D D A T K T F T V T E
5bmg_G M Q Y K L I L N G K T L K G C T T T E A V D A A T A E K V F K Q Y A N D N G V D G E W T Y D D A T K T F T V T E
5bmg_H M Q Y K L I L N G K T L K G C T T T E A V D A A T A E K V F K Q Y A N D N G V D G E W T Y D D A T K T F T V T E
5bmh_A M Q Y K L I L N G K T L K G E T T T E A V D A A T A E K V F K Q Y A N D N G V D G E W C Y D D A T K T F T V T E
5bmi_A M Q Y K L I L N G K T L K G E T T T E A V D A A T A E K V F K Q Y A N D N G V D G E W C Y D D A T K T F T V T E
5bwk_F M S T S A S G P E H E F V S K F L T L A T L T E P K L P K S Y T K P L K D V T N L G V P L P T L K Y K Y K Q N R
5bwk_H M S T S A S G P E H E F V S K F L T L A T L T E P K L P K S Y T K P L K D V T N L G V P L P T L K Y K Y K Q N R
5bwk_J M S T S A S G P E H E F V S K F L T L A T L T E P K L P K S Y T K P L K D V T N L G V P L P T L K Y K Y K Q N R
5bwk_L M S T S A S G P E H E F V S K F L T L A T L T E P K L P K S Y T K P L K D V T N L G V P L P T L K Y K Y K Q N R
5bwk_R M S T S A S G P E H E F V S K F L T L A T L T E P K L P K S Y T K P L K D V T N L G V P L P T L K Y K Y K Q N R
5bwk_T M S T S A S G P E H E F V S K F L T L A T L T E P K L P K S Y T K P L K D V T N L G V P L P T L K Y K Y K Q N R
5bwk_V M S T S A S G P E H E F V S K F L T L A T L T E P K L P K S Y T K P L K D V T N L G V P L P T L K Y K Y K Q N R
5bwk_X M S T S A S G P E H E F V S K F L T L A T L T E P K L P K S Y T K P L K D V T N L G V P L P T L K Y K Y K Q N R
5cai_A G C S S N Y V M H T N D G R T I V T D G K P Q T D N D T G M I S Y K D A W G N K Q Q I N R S D V K Q L G E L D E
5cai_B G C S S N Y V M H T N D G R T I V T D G K P Q T D N D T G M I S Y K D A W G N K Q Q I N R S D V K Q L G E L D E
5cai_C G C S S N Y V M H T N D G R T I V T D G K P Q T D N D T G M I S Y K D A W G N K Q Q I N R S D V K Q L G E L D E
5cai_D G C S S N Y V M H T N D G R T I V T D G K P Q T D N D T G M I S Y K D A W G N K Q Q I N R S D V K Q L G E L D E
5cai_E G C S S N Y V M H T N D G R T I V T D G K P Q T D N D T G M I S Y K D A W G N K Q Q I N R S D V K Q L G E L D E
5cai_F G C S S N Y V M H T N D G R T I V T D G K P Q T D N D T G M I S Y K D A W G N K Q Q I N R S D V K Q L G E L D E
5dma_A P G G G W K V G D R A N H R K W G I G T V V S V R G G G D D Q E L D I A F P S P I G I K R L L A K F A P I E K V
5epj_A G E Q V F A V E S I R K K R V R K G K V E Y L V K W K G W P P K Y S T W E P E E H I L D P R L V M A Y E E K E E
5flx_d M G H Q Q L Y W S H P R K F G Q G S R S C R V C S N R H G L I R K Y G L N M C R Q C F R Q Y A K D I G F I K L D
5gaf_b A V Q Q N K P T R S K R G M R R S H D A L T A V T S L S V D K T S G E K H L R H H I T A D G Y Y R G R K V I A K
5gpn_K M L T R F L G P R Y R Q L A R N W V P T A S L W G A V G A V G L V W A T D W R L I L D W V P Y I N G K F K K D D
5gpn_V M L T R F L G P R Y R Q L A R N W V P T A S L W G A V G A V G L V W A T D W R L I L D W V P Y I N G K F K K D D
5gpn_8 I H Q K R A P D F H D K Y G N A V L A S G A T F C V A V W V Y M A T Q I G I E W N P S P V G R V T P K E W R E Q
5gup_3 M L S R F L G P R Y R E L A R N W I P T A S M W G A V G A V G L V W A T D W R L I L D W V P Y I N G K F K K E D
5gup_Ad M L S R F L G P R Y R E L A R N W I P T A S M W G A V G A V G L V W A T D W R L I L D W V P Y I N G K F K K E D
5gup_Ai I H Q K R A P D F H D K Y G N A V L A S G A T F C V A V W V Y M A T Q I G I E W N P S P V G R V T P K E W R E Q
5h1s_b A V P K K R T S I Y K K R I R K N I W K K K G Y W A A L K A F S L A K S L S T G N S K S F F V R K I S N Q T L E
5h4p_z T I T A A D R R M V S G V T E K Q Y K K G M K K M Q Q L E K N A I N T Q I R R E I K R V N F Q T H Y R D E L L Q
5h5u_b A V Q Q N K P T R S K R G M R R S H D A L T A V T S L S V D K T S G E K H L R H H I T A D G Y Y R G R K V I A K
5he9_E S N A M M V T K E F L K T K L E C S D M Y A Q K L I D E A Q G D E N R L Y D L F I Q K L A E R H T R P A I V E Y
5it8_AU P V I K V R E N E P F D V A L R R F K R S C E K A G V L A E V R R R E F Y E K P T T E R K R A K A S A V K R H A
5it8_BU P V I K V R E N E P F D V A L R R F K R S C E K A G V L A E V R R R E F Y E K P T T E R K R A K A S A V K R H A
5it8_C1 A V Q Q N K P T R S K R G M R R S H D A L T A V T S L S V D K T S G E K H L R H H I T A D G Y Y R G R K V I A K
5it8_D1 A V Q Q N K P T R S K R G M R R S H D A L T A V T S L S V D K T S G E K H L R H H I T A D G Y Y R G R K V I A K
5j5b_AU P V I K V R E N E P F D V A L R R F K R S C E K A G V L A E V R R R E F Y E K P T T E R K R A K A S A V K R H A
5j5b_BU P V I K V R E N E P F D V A L R R F K R S C E K A G V L A E V R R R E F Y E K P T T E R K R A K A S A V K R H A
5j5b_C1 A V Q Q N K P T R S K R G M R R S H D A L T A V T S L S V D K T S G E K H L R H H I T A D G Y Y R G R K V I A K
5j5b_D1 A V Q Q N K P T R S K R G M R R S H D A L T A V T S L S V D K T S G E K H L R H H I T A D G Y Y R G R K V I A K
5j7l_AU P V I K V R E N E P F D V A L R R F K R S C E K A G V L A E V R R R E F Y E K P T T E R K R A K A S A V K R H A
5j7l_BU P V I K V R E N E P F D V A L R R F K R S C E K A G V L A E V R R R E F Y E K P T T E R K R A K A S A V K R H A
5j7l_C1 A V Q Q N K P T R S K R G M R R S H D A L T A V T S L S V D K T S G E K H L R H H I T A D G Y Y R G R K V I A K
5j7l_D1 A V Q Q N K P T R S K R G M R R S H D A L T A V T S L S V D K T S G E K H L R H H I T A D G Y Y R G R K V I A K
5j88_AU P V I K V R E N E P F D V A L R R F K R S C E K A G V L A E V R R R E F Y E K P T T E R K R A K A S A V K R H A
5j88_BU P V I K V R E N E P F D V A L R R F K R S C E K A G V L A E V R R R E F Y E K P T T E R K R A K A S A V K R H A
5j88_C1 A V Q Q N K P T R S K R G M R R S H D A L T A V T S L S V D K T S G E K H L R H H I T A D G Y Y R G R K V I A K
5j88_D1 A V Q Q N K P T R S K R G M R R S H D A L T A V T S L S V D K T S G E K H L R H H I T A D G Y Y R G R K V I A K

5j8a_AU P V I K V R E N E P F D V A L R R F K R S C E K A G V L A E V R R R E F Y E K P T T E R K R A K A S A V K R H A
5j8a_BU P V I K V R E N E P F D V A L R R F K R S C E K A G V L A E V R R R E F Y E K P T T E R K R A K A S A V K R H A
5j8a_C1 A V Q Q N K P T R S K R G M R R S H D A L T A V T S L S V D K T S G E K H L R H H I T A D G Y Y R G R K V I A K
5j8a_D1 A V Q Q N K P T R S K R G M R R S H D A L T A V T S L S V D K T S G E K H L R H H I T A D G Y Y R G R K V I A K
5j91_AU P V I K V R E N E P F D V A L R R F K R S C E K A G V L A E V R R R E F Y E K P T T E R K R A K A S A V K R H A
5j91_BU P V I K V R E N E P F D V A L R R F K R S C E K A G V L A E V R R R E F Y E K P T T E R K R A K A S A V K R H A
5j91_C1 A V Q Q N K P T R S K R G M R R S H D A L T A V T S L S V D K T S G E K H L R H H I T A D G Y Y R G R K V I A K
5j91_D1 A V Q Q N K P T R S K R G M R R S H D A L T A V T S L S V D K T S G E K H L R H H I T A D G Y Y R G R K V I A K
5jb3_P M A K A D Y N K R K P R K F G K G A R R C I R C G Q Y G P I I R I Q G L M L C R H C F R E V A P K L G F R K Y E
5jbh_P M A K A D Y N K R K P R K F G K G A R R C I R C G Q Y G P I I R I Q G L M L C R H C F R E V A P K L G F R K Y E
5jc9_AU P V I K V R E N E P F D V A L R R F K R S C E K A G V L A E V R R R E F Y E K P T T E R K R A K A S A V K R H A
5jc9_BU P V I K V R E N E P F D V A L R R F K R S C E K A G V L A E V R R R E F Y E K P T T E R K R A K A S A V K R H A
5jc9_C1 A V Q Q N K P T R S K R G M R R S H D A L T A V T S L S V D K T S G E K H L R H H I T A D G Y Y R G R K V I A K
5jc9_D1 A V Q Q N K P T R S K R G M R R S H D A L T A V T S L S V D K T S G E K H L R H H I T A D G Y Y R G R K V I A K
5juo_AC M A H E N V W F S H P R R Y G K G S R Q C R V C S S H T G L I R K Y G L N I C R Q C F R E K A N D I G F N K F R
5jup_AC M A H E N V W F S H P R R Y G K G S R Q C R V C S S H T G L I R K Y G L N I C R Q C F R E K A N D I G F N K F R
5jus_AC M A H E N V W F S H P R R Y G K G S R Q C R V C S S H T G L I R K Y G L N I C R Q C F R E K A N D I G F N K F R
5jut_AC M A H E N V W F S H P R R Y G K G S R Q C R V C S S H T G L I R K Y G L N I C R Q C F R E K A N D I G F N K F R
5juu_AC M A H E N V W F S H P R R Y G K G S R Q C R V C S S H T G L I R K Y G L N I C R Q C F R E K A N D I G F N K F R
5jxv_A M Q Y K L I L N G K T L K G E T T T E A V D A A T A E K V F K Q Y A N D N G V D G E W T Y D D A T K T F T V T E
5li0_4 A V P K R R T S K T R K N K R R T H F K I S V P G M T E C P N C G R E Y K L S H R V C K N C G S Y N G E E V A A
5lks_Sd M G H Q Q L Y W S H P R K F G Q G S R S C R V C S N R H G L I R K Y G L N M C R Q C F R Q Y A K D I G F I K L D
5luf_k M L T R F L G P R Y R Q L A R N W V P T A Q L W G A V G A V G L V S A T D S R L I L D W V P Y I N G K F K K D D
5luf_w M L T R F L G P R Y R Q L A R N W V P T A Q L W G A V G A V G L V S A T D S R L I L D W V P Y I N G K F K K D D
5luf_8 I H Q K R A P D F H D K Y G N A V L A S G A T F C V A V W V Y M A T Q I G I E W N P S P V G R V T P K E W R E Q
5lza_0 A V Q Q N K P T R S K R G M R R S H D A L T A V T S L S V D K T S G E K H L R H H I T A D G Y Y R G R K V I A K
5lzb_0 A V Q Q N K P T R S K R G M R R S H D A L T A V T S L S V D K T S G E K H L R H H I T A D G Y Y R G R K V I A K
5lzc_0 A V Q Q N K P T R S K R G M R R S H D A L T A V T S L S V D K T S G E K H L R H H I T A D G Y Y R G R K V I A K
5lzd_0 A V Q Q N K P T R S K R G M R R S H D A L T A V T S L S V D K T S G E K H L R H H I T A D G Y Y R G R K V I A K
5lze_0 A V Q Q N K P T R S K R G M R R S H D A L T A V T S L S V D K T S G E K H L R H H I T A D G Y Y R G R K V I A K
5lzf_0 A V Q Q N K P T R S K R G M R R S H D A L T A V T S L S V D K T S G E K H L R H H I T A D G Y Y R G R K V I A K
5lzs_dd M G H Q Q L Y W S H P R K F G Q G S R S C R V C S N R H G L I R K Y G L N M C R Q C F R Q Y A K D I G F I K L D
5lzt_dd M G H Q Q L Y W S H P R K F G Q G S R S C R V C S N R H G L I R K Y G L N M C R Q C F R Q Y A K D I G F I K L D
5lzu_dd M G H Q Q L Y W S H P R K F G Q G S R S C R V C S N R H G L I R K Y G L N M C R Q C F R Q Y A K D I G F I K L D
5lzv_dd M G H Q Q L Y W S H P R K F G Q G S R S C R V C S N R H G L I R K Y G L N M C R Q C F R Q Y A K D I G F I K L D
5lzw_dd M G H Q Q L Y W S H P R K F G Q G S R S C R V C S N R H G L I R K Y G L N M C R Q C F R Q Y A K D I G F I K L D
5lzx_dd M G H Q Q L Y W S H P R K F G Q G S R S C R V C S N R H G L I R K Y G L N M C R Q C F R Q Y A K D I G F I K L D
5lzy_dd M G H Q Q L Y W S H P R K F G Q G S R S C R V C S N R H G L I R K Y G L N M C R Q C F R Q Y A K D I G F I K L D
5lzz_dd M G H Q Q L Y W S H P R K F G Q G S R S C R V C S N R H G L I R K Y G L N M C R Q C F R Q Y A K D I G F I K L D
5m1s_F Q T E M D K V N V D L A A A G V A F K E R Y N M P V I A E A V E R E Q P E H L R S W F R E R L I A H R L A S V N
5mc6_M M A H E N V W F S H P R R Y G K G S R Q C R V C S S H T G L I R K Y G L N I C R Q C F R E K A N D I G F N K F R
5mc9_B G A L A S K L Q L L E D L E R K Y E D N Q K Y L E D K A Q E L V R L E G E V R S L L K D I S E K V A V Y S T C L
5mgp_0 A V Q Q N K P T R S K R G M R R S H D A L T A V T S L S V D K T S G E K H L R H H I T A D G Y Y R G R K V I A K
5nco_b A V Q Q N K P T R S K R G M R R S H D A L T A V T S L S V D K T S G E K H L R H H I T A D G Y Y R G R K V I A K
5nco_h A T V A F A R E A R T E V R K V I W P T R Q E T L H T T L I V A A V T A V M S L I L W G L D G I L V R L V S F I
5np6_y A V Q Q N K P T R S K R G M R R S H D A L T A V T S L S V D K T S G E K H L R H H I T A D G Y Y R G R K V I A K
5nx1_D A M I S R W Y F D V T E G K C A P F F Y G G C G G N R N N F D T E E Y C M A V C G S A I P R H H H H H H A A A N
5nx3_D A L F F R W Y F D V T E G K C A P F V Y G G C G G N R N N F D T E E Y C M A V C G S A I P R H H H H H H A A A N
5o2r_0 A V Q Q N K P T R S K R G M R R S H D A L T A V T S L S V D K T S G E K H L R H H I T A D G Y Y R G R K V I A K
5o94_A M Q F K L I L N G K T L K G V I T I E A V D H A E A E K F F K Q Y A N D N G V D G E W T Y D E A T H T F T V T E
5oa3_e G S L A R A G K V R G Q T P K V A K Q E K K K K K T G R A K R R M Q Y N R R F V N V V P T F G K K K G P N A N S
5ofs_A M Q F K L I L N G K T L K G V I T I E A V D H A E A E K F F K Q Y A N D N G V D G E W T Y D E A T H T F T V T E
5ofs_B M Q F K L I L N G K T L K G V I T I E A V D H A E A E K F F K Q Y A N D N G V D G E W T Y D E A T H T F T V T E
5ofs_C M Q F K L I L N G K T L K G V I T I E A V D H A E A E K F F K Q Y A N D N G V D G E W T Y D E A T H T F T V T E
5ofs_D M Q F K L I L N G K T L K G V I T I E A V D H A E A E K F F K Q Y A N D N G V D G E W T Y D E A T H T F T V T E
5t2c_AG M G H Q Q L Y W S H P R K F G Q G S R S C R V C S N R H G L I R K Y G L N M C R Q C F R Q Y A K D I G F I K L D
5tc2_A G K I Q N F R V Y Y R D S R N S L W K G P A K L L W K G E G A V V I Q D N S D I K V V P R R K A K I I R D Y G K
5tc2_B G K I Q N F R V Y Y R D S R N S L W K G P A K L L W K G E G A V V I Q D N S D I K V V P R R K A K I I R D Y G K
5tcu_S9 G I T H I D Y K D T E L L K R F I S E R G K I L P R R V T G T S A K Y Q R M L T T A I K R S R H M A L L P Y V K
5ub0_A M T F K L I I N G K T L K G E T T T E A V D A A T A E K V L K Q Y A N D N G I D G E W T Y D D A T K T F T V T E
5ubs_A M T F K L I I N G K T L K G E T T T E A V D A A T A E K V F K Q Y F N D N G I D G E W T Y D D A T K T F T I T E
5uce_A M T F K A I I N G K T L K G E T T T E A V D A A T A E K V F K Q Y F N D N G L D G E W T Y D D A T K T F T V T E
5ucf_A M T F K A I I N G K T L K G E T T T E A V D A A T A E K V F K Q Y F N D N G L D G E W T Y D D A T K T F T V T E
5uyk_30 A V Q Q N K P T R S K R G M R R S H D A L T A V T S L S V D K T S G E K H L R H H I T A D G Y Y R G R K V I A K
5uyl_30 A V Q Q N K P T R S K R G M R R S H D A L T A V T S L S V D K T S G E K H L R H H I T A D G Y Y R G R K V I A K
5uym_30 A V Q Q N K P T R S K R G M R R S H D A L T A V T S L S V D K T S G E K H L R H H I T A D G Y Y R G R K V I A K
5uyn_30 A V Q Q N K P T R S K R G M R R S H D A L T A V T S L S V D K T S G E K H L R H H I T A D G Y Y R G R K V I A K
5uyp_30 A V Q Q N K P T R S K R G M R R S H D A L T A V T S L S V D K T S G E K H L R H H I T A D G Y Y R G R K V I A K
5uyq_30 A V Q Q N K P T R S K R G M R R S H D A L T A V T S L S V D K T S G E K H L R H H I T A D G Y Y R G R K V I A K
5vyc_d1 M G H Q Q L Y W S H P R K F G Q G S R S C R V C S N R H G L I R K Y G L N M C R Q C F R Q Y A K D I G F I K L D
5vyc_d2 M G H Q Q L Y W S H P R K F G Q G S R S C R V C S N R H G L I R K Y G L N M C R Q C F R Q Y A K D I G F I K L D
5vyc_d3 M G H Q Q L Y W S H P R K F G Q G S R S C R V C S N R H G L I R K Y G L N M C R Q C F R Q Y A K D I G F I K L D
5vyc_d4 M G H Q Q L Y W S H P R K F G Q G S R S C R V C S N R H G L I R K Y G L N M C R Q C F R Q Y A K D I G F I K L D
5vyc_d5 M G H Q Q L Y W S H P R K F G Q G S R S C R V C S N R H G L I R K Y G L N M C R Q C F R Q Y A K D I G F I K L D
5vyc_d6 M G H Q Q L Y W S H P R K F G Q G S R S C R V C S N R H G L I R K Y G L N M C R Q C F R Q Y A K D I G F I K L D
5w97_K I H Q K R A P D F H D K Y G N A V L A S G A T F C V A V W V Y M A T Q I G I E W N P S P V G R V T P K E W R E Q
5w97_k I H Q K R A P D F H D K Y G N A V L A S G A T F C V A V W V Y M A T Q I G I E W N P S P V G R V T P K E W R E Q
5wau_K I H Q K R A P D F H D K Y G N A V L A S G A T F C V A V W V Y M A T Q I G I E W N P S P V G R V T P K E W R E Q
5wau_k I H Q K R A P D F H D K Y G N A V L A S G A T F C V A V W V Y M A T Q I G I E W N P S P V G R V T P K E W R E Q
5wdt_Z K T I K I T Q T R S A I G R L P K H K A T L L G L G L R R I G H T V E R E D T P A I R G M I N A V S F M V K V E
5we4_Z K T I K I T Q T R S A I G R L P K H K A T L L G L G L R R I G H T V E R E D T P A I R G M I N A V S F M V K V E
5we6_Z K T I K I T Q T R S A I G R L P K H K A T L L G L G L R R I G H T V E R E D T P A I R G M I N A V S F M V K V E
5wf0_Z K T I K I T Q T R S A I G R L P K H K A T L L G L G L R R I G H T V E R E D T P A I R G M I N A V S F M V K V E
5wfk_Z K T I K I T Q T R S A I G R L P K H K A T L L G L G L R R I G H T V E R E D T P A I R G M I N A V S F M V K V E
5wfs_Z K T I K I T Q T R S A I G R L P K H K A T L L G L G L R R I G H T V E R E D T P A I R G M I N A V S F M V K V E
5x19_K I H Q K R A P D F H D K Y G N A V L A S G A T F C V A V W V Y M A T Q I G I E W N P S P V G R V T P K E W R E Q
5x19_X I H Q K R A P D F H D K Y G N A V L A S G A T F C V A V W V Y M A T Q I G I E W N P S P V G R V T P K E W R E Q
5x1b_K I H Q K R A P D F H D K Y G N A V L A S G A T F C V A V W V Y M A T Q I G I E W N P S P V G R V T P K E W R E Q
5x1b_X I H Q K R A P D F H D K Y G N A V L A S G A T F C V A V W V Y M A T Q I G I E W N P S P V G R V T P K E W R E Q
5x1f_K I H Q K R A P D F H D K Y G N A V L A S G A T F C V A V W V Y M A T Q I G I E W N P S P V G R V T P K E W R E Q
5x1f_X I H Q K R A P D F H D K Y G N A V L A S G A T F C V A V W V Y M A T Q I G I E W N P S P V G R V T P K E W R E Q
5x8p_1 A V P K K R T S I Y K K R I R K N I W K K K G Y W A A L K A F S L A K S L S T G N S K S F F V R K I S N Q T L E
5x8t_1 A V P K K R T S I Y K K R I R K N I W K K K G Y W A A L K A F S L A K S L S T G N S K S F F V R K I S N Q T L E
5xdq_K I H Q K R A P D F H D K Y G N A V L A S G A T F C V A V W V Y M A T Q I G I E W N P S P V G R V T P K E W R E Q
5xdq_X I H Q K R A P D F H D K Y G N A V L A S G A T F C V A V W V Y M A T Q I G I E W N P S P V G R V T P K E W R E Q

```
5xdx_K I H Q K R A P D F H D K Y G N A V L A S G A T F C V A V W V Y M A T Q I G I E W N P S P V G R V T P K E W R E Q
5xdx_X I H Q K R A P D F H D K Y G N A V L A S G A T F C V A V W V Y M A T Q I G I E W N P S P V G R V T P K E W R E Q
5xtc_n N L L Q I V R D H W V H V L V P M G F V I G C Y L D R K S D E R L T A F R N K S M L F K R E L Q P S E E V T W K
5xtd_n N L L Q I V R D H W V H V L V P M G F V I G C Y L D R K S D E R L T A F R N K S M L F K R E L Q P S E E V T W K
5xth_n N L L Q I V R D H W V H V L V P M G F V I G C Y L D R K S D E R L T A F R N K S M L F K R E L Q P S E E V T W K
5xth_6 F E N R V A E K Q K L F Q E D N G L P V H L K G G A T D N I L Y R V T M T L C L G G T L Y S L Y C L G W A S F P
5xti_n N L L Q I V R D H W V H V L V P M G F V I G C Y L D R K S D E R L T A F R N K S M L F K R E L Q P S E E V T W K
5xti_Bn N L L Q I V R D H W V H V L V P M G F V I G C Y L D R K S D E R L T A F R N K S M L F K R E L Q P S E E V T W K
5xti_6 F E N R V A E K Q K L F Q E D N G L P V H L K G G A T D N I L Y R V T M T L C L G G T L Y S L Y C L G W A S F P
5xti_B6 F E N R V A E K Q K L F Q E D N G L P V H L K G G A T D N I L Y R V T M T L C L G G T L Y S L Y C L G W A S F P
5yvv_A A D L S L E K A A N V Q W D E M A D I T G S S P I I E V K Q D E D G S F S I R D V E E T N M I G G G G S G G G G
5z62_J K N K V P E K Q K L F Q E D D E I P L Y L K G G V A D A L L Y R A T M I L T V G G T A Y A I Y E L A V A S F P K
5z84_K I H Q K R A P D F H D K Y G N A V L A S G A T F C V A V W V Y M A T Q I G I E W N P S P V G R V T P K E W R E Q
5z84_X I H Q K R A P D F H D K Y G N A V L A S G A T F C V A V W V Y M A T Q I G I E W N P S P V G R V T P K E W R E Q
5z85_K I H Q K R A P D F H D K Y G N A V L A S G A T F C V A V W V Y M A T Q I G I E W N P S P V G R V T P K E W R E Q
5z85_X I H Q K R A P D F H D K Y G N A V L A S G A T F C V A V W V Y M A T Q I G I E W N P S P V G R V T P K E W R E Q
5z86_K I H Q K R A P D F H D K Y G N A V L A S G A T F C V A V W V Y M A T Q I G I E W N P S P V G R V T P K E W R E Q
5z86_X I H Q K R A P D F H D K Y G N A V L A S G A T F C V A V W V Y M A T Q I G I E W N P S P V G R V T P K E W R E Q
5zco_K I H Q K R A P D F H D K Y G N A V L A S G A T F C V A V W V Y M A T Q I G I E W N P S P V G R V T P K E W R E Q
5zco_X I H Q K R A P D F H D K Y G N A V L A S G A T F C V A V W V Y M A T Q I G I E W N P S P V G R V T P K E W R E Q
5zcp_K I H Q K R A P D F H D K Y G N A V L A S G A T F C V A V W V Y M A T Q I G I E W N P S P V G R V T P K E W R E Q
5zcp_X I H Q K R A P D F H D K Y G N A V L A S G A T F C V A V W V Y M A T Q I G I E W N P S P V G R V T P K E W R E Q
5zcq_K I H Q K R A P D F H D K Y G N A V L A S G A T F C V A V W V Y M A T Q I G I E W N P S P V G R V T P K E W R E Q
5zcq_X I H Q K R A P D F H D K Y G N A V L A S G A T F C V A V W V Y M A T Q I G I E W N P S P V G R V T P K E W R E Q
6a3z_A A T P E E L Q A M D N V C I I C R E E M V T G A K R L P C N H I F H T S C L R S W F Q R Q Q T C P T C R M D V L
6bu8_30 A V Q Q N K P T R S K R G M R R S H D A L T A V T S L S V D K T S G E K H L R H H I T A D G Y Y R G R K V I A K
6by1_C1 A V Q Q N K P T R S K R G M R R S H D A L T A V T S L S V D K T S G E K H L R H H I T A D G Y Y R G R K V I A K
6by1_D1 A V Q Q N K P T R S K R G M R R S H D A L T A V T S L S V D K T S G E K H L R H H I T A D G Y Y R G R K V I A K
6c9o_A G Q Y K L I L N G K T L K G E T T T E A V D A A T A E K M F K Q Y A N D N G V D G E W T Y D D A T K T F T V T E
6c9o_B G Q Y K L I L N G K T L K G E T T T E A V D A A T A E K M F K Q Y A N D N G V D G E W T Y D D A T K T F T V T E
6cfz_H M N A F E P A F A E L A D A V A D L E A N M M H F Q L M H E S L A R F S E S F A S F L Y G L N M N A F C V D F P
6che_A G Q Y K L I L N G K T L K G E T T T E A V D A A T A E K V F K Q Y M N D N G V D G E W T Y D D A T K T F T V T E
6cku_A D V Q C G G G F S C H D G E T C C P T S Q T T W G C C P S P K A V C C D D M Q H C C P A G Y K C G P G G T C I S
6cne_A G Q Y K M I L N G K T L K G E T T T E A V D A A T A E K V F K Q Y A N D N G V D G E W T Y D D A T K T F T V T E
6cne_B G Q Y K M I L N G K T L K G E T T T E A V D A A T A E K V F K Q Y A N D N G V D G E W T Y D D A T K T F T V T E
6cpz_A G Q Y K L M L N G K T L K G E T T T E A V D A A T A E K V F K Q Y A N D N G V D G E W T Y D D A T K T F T V T E
6cpz_B G Q Y K L M L N G K T L K G E T T T E A V D A A T A E K V F K Q Y A N D N G V D G E W T Y D D A T K T F T V T E
6cte_A G Q Y K L I L N G K T L K G E T T T E A V D A A T A E K V F K Q Y A N D N G M D G E W T Y D D A T K T F T V T E
6cte_B G Q Y K L I L N G K T L K G E T T T E A V D A A T A E K V F K Q Y A N D N G M D G E W T Y D D A T K T F T V T E
6d90_ee M G H Q Q L Y W S H P R K F G Q G S R S C R V C S N R H G L I R K Y G L N M C R Q C F R Q Y A K D I G F I K L D
6d9j_ee M G H Q Q L Y W S H P R K F G Q G S R S C R V C S N R H G L I R K Y G L N M C R Q C F R Q Y A K D I G F I K L D
6e6q_A M Y V C L C Q G V T D N Q I R D A I Y E G C C S Y R E V R E A T G V G T Q C G K C A S L A K Q V V R E T L N D L
6e6q_B M Y V C L C Q G V T D N Q I R D A I Y E G C C S Y R E V R E A T G V G T Q C G K C A S L A K Q V V R E T L N D L
6e6r_A M Y V C L C Q G V T D N Q I R D A I Y E G C C S Y E E V R E A T G V G T Q C G K C A S L A K Q V V R E T L N D L
6e6s_A M Y V C L C Q G V T D N Q I R D A I Y E G C C S Y E E V R E A T G V G T Q C G K C A S L A Y Q V V R E T L N D L
6e6s_B M Y V C L C Q G V T D N Q I R D A I Y E G C C S Y E E V R E A T G V G T Q C G K C A S L A Y Q V V R E T L N D L
6ek0_Sd M G H Q Q L Y W S H P R K F G Q G S R S C R V C S N R H G L I R K Y G L N M C R Q C F R Q Y A K D I G F I K L D
6enf_0 A V Q Q N K P T R S K R G M R R S H D A L T A V T S L S V D K T S G E K H L R H H I T A D G Y Y R G R K V I A K
6enj_0 A V Q Q N K P T R S K R G M R R S H D A L T A V T S L S V D K T S G E K H L R H H I T A D G Y Y R G R K V I A K
6enu_0 A V Q Q N K P T R S K R G M R R S H D A L T A V T S L S V D K T S G E K H L R H H I T A D G Y Y R G R K V I A K
6et5_1 M A D L K P S L T G L T E E E A K E F H G I F V T S T V L Y L A T A V I V H Y L V W T A R P W I A P I P K G W V
6et5_G M A D L K P S L T G L T E E E A K E F H G I F V T S T V L Y L A T A V I V H Y L V W T A R P W I A P I P K G W V
6et5_N M A D L K P S L T G L T E E E A K E F H G I F V T S T V L Y L A T A V I V H Y L V W T A R P W I A P I P K G W V
6et5_Q M A D L K P S L T G L T E E E A K E F H G I F V T S T V L Y L A T A V I V H Y L V W T A R P W I A P I P K G W V
6et5_T M A D L K P S L T G L T E E E A K E F H G I F V T S T V L Y L A T A V I V H Y L V W T A R P W I A P I P K G W V
6et5_W M A D L K P S L T G L T E E E A K E F H G I F V T S T V L Y L A T A V I V H Y L V W T A R P W I A P I P K G W V
6et5_Z M A D L K P S L T G L T E E E A K E F H G I F V T S T V L Y L A T A V I V H Y L V W T A R P W I A P I P K G W V
6et5_c M A D L K P S L T G L T E E E A K E F H G I F V T S T V L Y L A T A V I V H Y L V W T A R P W I A P I P K G W V
6et5_f M A D L K P S L T G L T E E E A K E F H G I F V T S T V L Y L A T A V I V H Y L V W T A R P W I A P I P K G W V
6et5_i M A D L K P S L T G L T E E E A K E F H G I F V T S T V L Y L A T A V I V H Y L V W T A R P W I A P I P K G W V
6et5_l M A D L K P S L T G L T E E E A K E F H G I F V T S T V L Y L A T A V I V H Y L V W T A R P W I A P I P K G W V
6et5_o M A D L K P S L T G L T E E E A K E F H G I F V T S T V L Y L A T A V I V H Y L V W T A R P W I A P I P K G W V
6et5_r M A D L K P S L T G L T E E E A K E F H G I F V T S T V L Y L A T A V I V H Y L V W T A R P W I A P I P K G W V
6et5_u M A D L K P S L T G L T E E E A K E F H G I F V T S T V L Y L A T A V I V H Y L V W T A R P W I A P I P K G W V
6et5_x M A D L K P S L T G L T E E E A K E F H G I F V T S T V L Y L A T A V I V H Y L V W T A R P W I A P I P K G W V
6et5_4 M A D L K P S L T G L T E E E A K E F H G I F V T S T V L Y L A T A V I V H Y L V W T A R P W I A P I P K G W V
6et5_7 M A D L K P S L T G L T E E E A K E F H G I F V T S T V L Y L A T A V I V H Y L V W T A R P W I A P I P K G W V
6f5n_A M Q F K L I L N G K T L K G V I T I E A V D H A E A E K F F K Q Y A N D N G V D G E W T Y D E A T H T F T V T E
6f5n_B M Q F K L I L N G K T L K G V I T I E A V D H A E A E K F F K Q Y A N D N G V D G E W T Y D E A T H T F T V T E
6fai_d M A H E N V W F S H P R R Y G K G S R Q C R V C S S H T G L I R K Y G L N I C R Q C F R E K A N D I G F N K F R
6fyx_d M A H E N V W Y S H P R K F G K G S R Q C R I S G S H S G L I R K Y G L N I D R Q S F R E K A N D I G F Y K Y R
6fyy_d M A H E N V W Y S H P R K F G K G S R Q C R I S G S H S G L I R K Y G L N I D R Q S F R E K A N D I G F Y K Y R
6g51_d M G H Q Q L Y W S H P R K F G Q G S R S C R V C S N R H G L I R K Y G L N M C R Q C F R Q Y A K D I G F I K L D
6g53_d M G H Q Q L Y W S H P R K F G Q G S R S C R V C S N R H G L I R K Y G L N M C R Q C F R Q Y A K D I G F I K L D
6g5h_d M G H Q Q L Y W S H P R K F G Q G S R S C R V C S N R H G L I R K Y G L N M C R Q C F R Q Y A K D I G F I K L D
6g5i_d M G H Q Q L Y W S H P R K F G Q G S R S C R V C S N R H G L I R K Y G L N M C R Q C F R Q Y A K D I G F I K L D
6gbz_0 A V Q Q N K P T R S K R G M R R S H D A L T A V T S L S V D K T S G E K H L R H H I T A D G Y Y R G R K V I A K
6gc0_0 A V Q Q N K P T R S K R G M R R S H D A L T A V T S L S V D K T S G E K H L R H H I T A D G Y Y R G R K V I A K
6gc4_0 A V Q Q N K P T R S K R G M R R S H D A L T A V T S L S V D K T S G E K H L R H H I T A D G Y Y R G R K V I A K
6gc6_0 A V Q Q N K P T R S K R G M R R S H D A L T A V T S L S V D K T S G E K H L R H H I T A D G Y Y R G R K V I A K
6gc7_0 A V Q Q N K P T R S K R G M R R S H D A L T A V T S L S V D K T S G E K H L R H H I T A D G Y Y R G R K V I A K
6gc8_0 A V Q Q N K P T R S K R G M R R S H D A L T A V T S L S V D K T S G E K H L R H H I T A D G Y Y R G R K V I A K
6gwt_0 A V Q Q N K P T R S K R G M R R S H D A L T A V T S L S V D K T S G E K H L R H H I T A D G Y Y R G R K V I A K
6gxm_0 A V Q Q N K P T R S K R G M R R S H D A L T A V T S L S V D K T S G E K H L R H H I T A D G Y Y R G R K V I A K
6gxn_0 A V Q Q N K P T R S K R G M R R S H D A L T A V T S L S V D K T S G E K H L R H H I T A D G Y Y R G R K V I A K
6gxo_0 A V Q Q N K P T R S K R G M R R S H D A L T A V T S L S V D K T S G E K H L R H H I T A D G Y Y R G R K V I A K
6gxp_0 A V Q Q N K P T R S K R G M R R S H D A L T A V T S L S V D K T S G E K H L R H H I T A D G Y Y R G R K V I A K
6h4n_0 A V Q Q N K P T R S K R G M R R S H D A L T A V T S L S V D K T S G E K H L R H H I T A D G Y Y R G R K V I A K
6h58_0 A V Q Q N K P T R S K R G M R R S H D A L T A V T S L S V D K T S G E K H L R H H I T A D G Y Y R G R K V I A K
6h58_00 A V Q Q N K P T R S K R G M R R S H D A L T A V T S L S V D K T S G E K H L R H H I T A D G Y Y R G R K V I A K
6hcf_e1 M G H Q Q L Y W S H P R K F G Q G S R S C R V C S N R H G L I R K Y G L N M C R Q C F R Q Y A K D I G F I K L D
6hcj_e2 M G H Q Q L Y W S H P R K F G Q G S R S C R V C S N R H G L I R K Y G L N M C R Q C F R Q Y A K D I G F I K L D
6hcm_e1 M G H Q Q L Y W S H P R K F G Q G S R S C R V C S N R H G L I R K Y G L N M C R Q C F R Q Y A K D I G F I K L D
6hcq_e2 M G H Q Q L Y W S H P R K F G Q G S R S C R V C S N R H G L I R K Y G L N M C R Q C F R Q Y A K D I G F I K L D
6hhq_e M A H E N V W F S H P R R Y G K G S R Q C R V C S S H T G L I R K Y G L N I C R Q C F R E K A N D I G F N K F R
```

6hhq_d9 M A H E N V W F S H P R R Y G K G S R Q C R V C S S H T G L I R K Y G L N I C R Q C F R E K A N D I G F N K F R
6hiv_AB A A A A A A A A A A A A A A A A A A A A A A A A A A A A A A A A A A A A A A A A A A A A A A A A A A A A A A A A
6hix_AB A A A A A A A A A A A A A A A A A A A A A A A A A A A A A A A A A A A A A A A A A A A A A A A A A A A A A A A A
6hrm_b A V Q Q N K P T R S K R G M R R S H D A L T A V T S L S V D K T S G E K H L R H H I T A D G Y Y R G R K V I A K
6i7v_C1 A V Q Q N K P T R S K R G M R R S H D A L T A V T S L S V D K T S G E K H L R H H I T A D G Y Y R G R K V I A K
6i7v_D1 A V Q Q N K P T R S K R G M R R S H D A L T A V T S L S V D K T S G E K H L R H H I T A D G Y Y R G R K V I A K
6ip5_3E M G H Q Q L Y W S H P R K F G Q G S R S C R V C S N R H G L I R K Y G L N M C R Q C F R Q Y A K D I G F I K L D
6ip6_3E M G H Q Q L Y W S H P R K F G Q G S R S C R V C S N R H G L I R K Y G L N M C R Q C F R Q Y A K D I G F I K L D
6ip8_3E M G H Q Q L Y W S H P R K F G Q G S R S C R V C S N R H G L I R K Y G L N M C R Q C F R Q Y A K D I G F I K L D
6j8m_K I H Q K R A P D F H D K Y G N A V L A S G A T F C V A V W V Y M A T Q I G I E W N P S P V G R V T P K E W R E Q
6j8m_X I H Q K R A P D F H D K Y G N A V L A S G A T F C V A V W V Y M A T Q I G I E W N P S P V G R V T P K E W R E Q
6jy3_K I H Q K R A P D F H D K Y G N A V L A S G A T F C V A V W V Y M A T Q I G I E W N P S P V G R V T P K E W R E Q
6jy4_K I H Q K R A P D F H D K Y G N A V L A S G A T F C V A V W V Y M A T Q I G I E W N P S P V G R V T P K E W R E Q
6kza_C M K N V G D L M Q R L Q K M M P A H I K P A F K T G E E L L A W Q K E Q G A I R S A A L E R E N R A M K M Q R T
6kza_D M K N V G D L M Q R L Q K M M P A H I K P A F K T G E E L L A W Q K E Q G A I R S A A L E R E N R A M K M Q R T
6l91_A D T Y K L I L N G V T L K G E T T T E A V D A A T A E K V F K Q Y A N D N G V D G E W T Y D D A T K T F T V T E
6l9b_A D T Y K L I L N G V A L K G E T T T E A V D A A T A E K V F K Q Y A N D N G V D G E W T Y D D A T K T F T V T E
6l9d_A D T Y K L I L N G V S L K G E T T T E A V D A A T A E K V F K Q Y A N D N G V D G E W T Y D D A T K T F T V T E
6lji_A D T Y K L I L N G V V L K G E T T T E A V D A A T A E K V F K Q Y A N D N G V D G E W T Y D D A T K T F T V T E
6lji_B D T Y K L I L N G V V L K G E T T T E A V D A A T A E K V F K Q Y A N D N G V D G E W T Y D D A T K T F T V T E
6lkq_i A V Q Q N K P T R S K R G M R R S H D A L T A V T S L S V D K T S G E K H L R H H I T A D G Y Y R G R K V I A K
6nkn_K I H Q K R A P D F H D K Y G N A V L A S G A T F C V A V W V Y M A T Q I G I E W N P S P V G R V T P K E W R E Q
6nkn_X I H Q K R A P D F H D K Y G N A V L A S G A T F C V A V W V Y M A T Q I G I E W N P S P V G R V T P K E W R E Q
6nl6_A M T Y K L I L N G K T L K G E F T A E A E D A A L A E Y I F R H L A K H Q G V D G E W T Y D D A T K T F T V T E
6nl6_B M T Y K L I L N G K T L K G E F T A E A E D A A L A E Y I F R H L A K H Q G V D G E W T Y D D A T K T F T V T E
6nl6_C M T Y K L I L N G K T L K G E F T A E A E D A A L A E Y I F R H L A K H Q G V D G E W T Y D D A T K T F T V T E
6nl6_D M T Y K L I L N G K T L K G E F T A E A E D A A L A E Y I F R H L A K H Q G V D G E W T Y D D A T K T F T V T E
6nl7_A M T Y K L I L N G K T L K G E F T A E A H D A A H A E Y I F R A L A K A Q G V D G E W T Y D D A T K T F T V T E
6nl7_B M T Y K L I L N G K T L K G E F T A E A H D A A H A E Y I F R A L A K A Q G V D G E W T Y D D A T K T F T V T E
6nl7_C M T Y K L I L N G K T L K G E F T A E A H D A A H A E Y I F R A L A K A Q G V D G E W T Y D D A T K T F T V T E
6nl7_D M T Y K L I L N G K T L K G E F T A E A H D A A H A E Y I F R A L A K A Q G V D G E W T Y D D A T K T F T V T E
6nl8_A M T Y K L I L N G K T H K G E L T T E A V D A A T A E K H F K Q H A N D L G V D G E W T Y D D A T K T F T V T E
6nl9_A M T Y K L I L N G K T H K G E L T T E A V D A A T A E K H F K Q H A N D L G V D G E W T Y D D A T K T F T V T E
6nl9_B M T Y K L I L N G K T H K G E L T T E A V D A A T A E K H F K Q H A N D L G V D G E W T Y D D A T K T F T V T E
6nl9_C M T Y K L I L N G K T H K G E L T T E A V D A A T A E K H F K Q H A N D L G V D G E W T Y D D A T K T F T V T E
6nl9_D M T Y K L I L N G K T H K G E L T T E A V D A A T A E K H F K Q H A N D L G V D G E W T Y D D A T K T F T V T E
6nla_A M T Y K L I L N G K T H K G V L T I E A V D A A T A E K H F K Q H A N D L G V D G E W T Y D D A T K T F T V T E
6nlb_A M T Y K L I L N G K T H K G V L T I E A V D A A T A E K H F K Q H A N D L G V D G E W T Y D D A T K T F T V T E
6nlb_B M T Y K L I L N G K T H K G V L T I E A V D A A T A E K H F K Q H A N D L G V D G E W T Y D D A T K T F T V T E
6nlb_C M T Y K L I L N G K T H K G V L T I E A V D A A T A E K H F K Q H A N D L G V D G E W T Y D D A T K T F T V T E
6nlb_D M T Y K L I L N G K T H K G V L T I E A V D A A T A E K H F K Q H A N D L G V D G E W T Y D D A T K T F T V T E
6nmf_K I H Q K R A P D F H D K Y G N A V L A S G A T F C V A V W V Y M A T Q I G I E W N P S P V G R V T P K E W R E Q
6nmf_X I H Q K R A P D F H D K Y G N A V L A S G A T F C V A V W V Y M A T Q I G I E W N P S P V G R V T P K E W R E Q
6nmp_K I H Q K R A P D F H D K Y G N A V L A S G A T F C V A V W V Y M A T Q I G I E W N P S P V G R V T P K E W R E Q
6nmp_X I H Q K R A P D F H D K Y G N A V L A S G A T F C V A V W V Y M A T Q I G I E W N P S P V G R V T P K E W R E Q
6nu4_A E A S V R Y I T Y P A I D R G D H A V H C D K A H P N T C K K K Q A N P Y R R G C G V L E G C H R E T G P K P T
6o8z_2 A V P A R R T S K A K K A K R R T H Y K L T I K G L N A C S N C G E M K K S H H V C P A C G H Y D G K D V M S K
6o9j_1 A V Q Q N K P T R S K R G M R R S H D A L T A V T S L S V D K T S G E K H L R H H I T A D G Y Y R G R K V I A K
6o9k_5 A V Q Q N K P T R S K R G M R R S H D A L T A V T S L S V D K T S G E K H L R H H I T A D G Y Y R G R K V I A K
6ofx_B A V Q Q N K P T R S K R G M R R S H D A L T A V T S L S V D K T S G E K H L R H H I T A D G Y Y R G R K V I A K
6og7_B A V Q Q N K P T R S K R G M R R S H D A L T A V T S L S V D K T S G E K H L R H H I T A D G Y Y R G R K V I A K
6ore_b A V Q Q N K P T R S K R G M R R S H D A L T A V T S L S V D K T S G E K H L R H H I T A D G Y Y R G R K V I A K
6orl_b A V Q Q N K P T R S K R G M R R S H D A L T A V T S L S V D K T S G E K H L R H H I T A D G Y Y R G R K V I A K
6osk_b A V Q Q N K P T R S K R G M R R S H D A L T A V T S L S V D K T S G E K H L R H H I T A D G Y Y R G R K V I A K
6osq_b A V Q Q N K P T R S K R G M R R S H D A L T A V T S L S V D K T S G E K H L R H H I T A D G Y Y R G R K V I A K
6ost_b A V Q Q N K P T R S K R G M R R S H D A L T A V T S L S V D K T S G E K H L R H H I T A D G Y Y R G R K V I A K
6osw_B G A Q A G A A N R S L T E G F V L D T M N D S L S K I L V D I S F S G L E D E D L G M G N I S W S Q F I P E A K
6ot3_b A V Q Q N K P T R S K R G M R R S H D A L T A V T S L S V D K T S G E K H L R H H I T A D G Y Y R G R K V I A K
6ouo_b A V Q Q N K P T R S K R G M R R S H D A L T A V T S L S V D K T S G E K H L R H H I T A D G Y Y R G R K V I A K
6p4g_e M G H Q Q L Y W S H P R K F G Q G S R S C R V C S N R H G L I R K Y G L N M C R Q C F R Q Y A K D I G F I K L D
6p4h_e M G H Q Q L Y W S H P R K F G Q G S R S C R V C S N R H G L I R K Y G L N M C R Q C F R Q Y A K D I G F I K L D
6p5i_e M G H Q Q L Y W S H P R K F G Q G S R S C R V C S N R H G L I R K Y G L N M C R Q C F R Q Y A K D I G F I K L D
6p5j_e M G H Q Q L Y W S H P R K F G Q G S R S C R V C S N R H G L I R K Y G L N M C R Q C F R Q Y A K D I G F I K L D
6p5k_e M G H Q Q L Y W S H P R K F G Q G S R S C R V C S N R H G L I R K Y G L N M C R Q C F R Q Y A K D I G F I K L D
6p5n_e M G H Q Q L Y W S H P R K F G Q G S R S C R V C S N R H G L I R K Y G L N M C R Q C F R Q Y A K D I G F I K L D
6pj6_i A V Q Q N K P T R S K R G M R R S H D A L T A V T S L S V D K T S G E K H L R H H I T A D G Y Y R G R K V I A K
6q97_Z T I K I T Q T R S A I G R L P K H K A T L L G L G L R R I G H T V E R E D T P A I R G M I N A V S F M V K V E E
6q9a_Z K T I K I T Q T R S A I G R L P K H K A T L L G L G L R R I G H T V E R E D T P A I R G M I N A V S F M V K V E
6qwn_F M G R R Q L A R G R R Y I G Y D A L K K N N V P C S R R G R S Y Y D C K K R R R N N P Y R R G C S A I T H C Y R
6qwn_G M G R R Q L A R G R R Y I G Y D A L K K N N V P C S R R G R S Y Y D C K K R R R N N P Y R R G C S A I T H C Y R
6qwn_H M G R R Q L A R G R R Y I G Y D A L K K N N V P C S R R G R S Y Y D C K K R R R N N P Y R R G C S A I T H C Y R
6qwn_I M G R R Q L A R G R R Y I G Y D A L K K N N V P C S R R G R S Y Y D C K K R R R N N P Y R R G C S A I T H C Y R
6qwn_J M G R R Q L A R G R R Y I G Y D A L K K N N V P C S R R G R S Y Y D C K K R R R N N P Y R R G C S A I T H C Y R
6qxp_I M G R R Q L A R G R R Y I G Y D A L K K N N V P C S R R G R S Y Y D C K K R R R N N P Y R R G C S A I T H C Y R
6qxp_J M G R R Q L A R G R R Y I G Y D A L K K N N V P C S R R G R S Y Y D C K K R R R N N P Y R R G C S A I T H C Y R
6qxp_K M G R R Q L A R G R R Y I G Y D A L K K N N V P C S R R G R S Y Y D C K K R R R N N P Y R R G C S A I T H C Y R
6qxp_L M G R R Q L A R G R R Y I G Y D A L K K N N V P C S R R G R S Y Y D C K K R R R N N P Y R R G C S A I T H C Y R
6qxp_M M G R R Q L A R G R R Y I G Y D A L K K N N V P C S R R G R S Y Y D C K K R R R N N P Y R R G C S A I T H C Y R
6qxp_N M G R R Q L A R G R R Y I G Y D A L K K N N V P C S R R G R S Y Y D C K K R R R N N P Y R R G C S A I T H C Y R
6qxp_O M G R R Q L A R G R R Y I G Y D A L K K N N V P C S R R G R S Y Y D C K K R R R N N P Y R R G C S A I T H C Y R
6qxp_P M G R R Q L A R G R R Y I G Y D A L K K N N V P C S R R G R S Y Y D C K K R R R N N P Y R R G C S A I T H C Y R
6rbd_d M A H E N V W F S H P R R Y G K G S R Q C R V C S S H T G L I R K Y G L N I C R Q C F R E K A N D I G F N K F R
6rbe_d M A H E N V W F S H P R R Y G K G S R Q C R V C S S H T G L I R K Y G L N I C R Q C F R E K A N D I G F N K F R
6sgc_e1 M G H Q Q L Y W S H P R K F G Q G S R S C R V C S N R H G L I R K Y G L N M C R Q C F R Q Y A K D I G F I K L D
6skf_Ar M A K A D Y N K R K P R K F G K G A R R C M R C G Q Y G P I I R I H G L M L C R H C F R E V A P K L G F K K Y E
6skg_Ar M A K A D Y N K R K P R K F G K G A R R C M R C G Q Y G P I I R I H G L M L C R H C F R E V A P K L G F K K Y E
6snt_d M A H E N V W F S H P R R Y G K G S R Q C R V C S S H T G L I R K Y G L N I C R Q C F R E K A N D I G F N K F R
6spc_r K E I D Y K D L N T L K A Y V S E T G K I V P S R I T G T K A K Y Q R Q L A T A I K R A R Y L A L L P Y T D S H
6spf_r K E I D Y K D L N T L K A Y V S E T G K I V P S R I T G T K A K Y Q R Q L A T A I K R A R Y L A L L P Y T D S H
6sv4_M M A H E N V W F S H P R R Y G K G S R Q C R V C S S H T G L I R K Y G L N I C R Q C F R E K A N D I G F N K F R
6sv4_Mb M A H E N V W F S H P R R Y G K G S R Q C R V C S S H T G L I R K Y G L N I C R Q C F R E K A N D I G F N K F R
6sv4_Mc M A H E N V W F S H P R R Y G K G S R Q C R V C S S H T G L I R K Y G L N I C R Q C F R E K A N D I G F N K F R
6sw9_P M A K A D Y N K R K P R K F G K G A R R C I R C G Q Y G P I I R I H G L M L C R H C F R E V A P K L G F R K Y E
6swc_P M A K A D Y N K R K P R K F G K G A R R C I R C G Q Y G P I I R I H G L M L C R H C F R E V A P K L G F R K Y E
6swe_P M A K A D Y N K R K P R K F G K G A R R C I R C G Q Y G P I I R I H G L M L C R H C F R E V A P K L G F R K Y E

```
6t7i_Sd M A H E N V W F S H P R R Y G K G S R Q C R V C S S H T G L I R K Y G L N I C R Q C F R E K A N D I G F N K F R
6t83_db M A H E N V W F S H P R R Y G K G S R Q C R V C S S H T G L I R K Y G L N I C R Q C F R E K A N D I G F N K F R
6t83_4 M A H E N V W F S H P R R Y G K G S R Q C R V C S S H T G L I R K Y G L N I C R Q C F R E K A N D I G F N K F R
6th6_Ar M A K A D Y N K R K P R K F G K G A R R C M R C G Q Y G P I I R I H G L M L C R H C F R E V A P K L G F K K Y E
6tmf_Q M A K A D Y N K R K P R K F G K G A R R C V R C G Q Y G P V I R V H G L M L C R H C F R E I A P K L G F K K Y E
6tmf_W K F E V P V C T S C G K E I T P R E H A T H F V C P N C G E A I I W R C E S C R V L S V P Y K C P K C G W E G P
6u48_C1 A V Q Q N K P T R S K R G M R R S H D A L T A V T S L S V D K T S G E K H L R H H I T A D G Y Y R G R K V I A K
6uz7_d M A H E N V W Y S H P R K F G K G S R Q C R I S G S H S G L I R K Y G L N I D R Q S F R E K A N D I G F Y K Y R
6v2r_A G E Q V F A A E S I R K K R V R K G K V E Y L V K W K G W P P K Y S T W E P E E H I L D P R L V M A Y E E K E E
6vu3_i A V Q Q N K P T R S K R G M R R S H D A L T A V T S L S V D K T S G E K H L R H H I T A D G Y Y R G R K V I A K
6vz2_i A V Q Q N K P T R S K R G M R R S H D A L T A V T S L S V D K T S G E K H L R H H I T A D G Y Y R G R K V I A K
6vz3_i A V Q Q N K P T R S K R G M R R S H D A L T A V T S L S V D K T S G E K H L R H H I T A D G Y Y R G R K V I A K
6vz7_i A V Q Q N K P T R S K R G M R R S H D A L T A V T S L S V D K T S G E K H L R H H I T A D G Y Y R G R K V I A K
6vzj_i A V Q Q N K P T R S K R G M R R S H D A L T A V T S L S V D K T S G E K H L R H H I T A D G Y Y R G R K V I A K
6w2s_e M G H Q Q L Y W S H P R K F G Q G S R S C R V C S N R H G L I R K Y G L N M C R Q C F R Q Y A K D I G F I K L D
6w2t_e M G H Q Q L Y W S H P R K F G Q G S R S C R V C S N R H G L I R K Y G L N M C R Q C F R Q Y A K D I G F I K L D
6wd0_B A V Q Q N K P T R S K R G M R R S H D A L T A V T S L S V D K T S G E K H L R H H I T A D G Y Y R G R K V I A K
6wd1_B A V Q Q N K P T R S K R G M R R S H D A L T A V T S L S V D K T S G E K H L R H H I T A D G Y Y R G R K V I A K
6wd2_B A V Q Q N K P T R S K R G M R R S H D A L T A V T S L S V D K T S G E K H L R H H I T A D G Y Y R G R K V I A K
6wd3_B A V Q Q N K P T R S K R G M R R S H D A L T A V T S L S V D K T S G E K H L R H H I T A D G Y Y R G R K V I A K
6wd4_B A V Q Q N K P T R S K R G M R R S H D A L T A V T S L S V D K T S G E K H L R H H I T A D G Y Y R G R K V I A K
6wd5_B A V Q Q N K P T R S K R G M R R S H D A L T A V T S L S V D K T S G E K H L R H H I T A D G Y Y R G R K V I A K
6wd6_B A V Q Q N K P T R S K R G M R R S H D A L T A V T S L S V D K T S G E K H L R H H I T A D G Y Y R G R K V I A K
6wd7_B A V Q Q N K P T R S K R G M R R S H D A L T A V T S L S V D K T S G E K H L R H H I T A D G Y Y R G R K V I A K
6wd8_B A V Q Q N K P T R S K R G M R R S H D A L T A V T S L S V D K T S G E K H L R H H I T A D G Y Y R G R K V I A K
6wd9_B A V Q Q N K P T R S K R G M R R S H D A L T A V T S L S V D K T S G E K H L R H H I T A D G Y Y R G R K V I A K
6wda_B A V Q Q N K P T R S K R G M R R S H D A L T A V T S L S V D K T S G E K H L R H H I T A D G Y Y R G R K V I A K
6wdb_B A V Q Q N K P T R S K R G M R R S H D A L T A V T S L S V D K T S G E K H L R H H I T A D G Y Y R G R K V I A K
6wdc_B A V Q Q N K P T R S K R G M R R S H D A L T A V T S L S V D K T S G E K H L R H H I T A D G Y Y R G R K V I A K
6wdd_B A V Q Q N K P T R S K R G M R R S H D A L T A V T S L S V D K T S G E K H L R H H I T A D G Y Y R G R K V I A K
6wde_B A V Q Q N K P T R S K R G M R R S H D A L T A V T S L S V D K T S G E K H L R H H I T A D G Y Y R G R K V I A K
6wdf_B A V Q Q N K P T R S K R G M R R S H D A L T A V T S L S V D K T S G E K H L R H H I T A D G Y Y R G R K V I A K
6wdg_B A V Q Q N K P T R S K R G M R R S H D A L T A V T S L S V D K T S G E K H L R H H I T A D G Y Y R G R K V I A K
6wdh_B A V Q Q N K P T R S K R G M R R S H D A L T A V T S L S V D K T S G E K H L R H H I T A D G Y Y R G R K V I A K
6wdi_B A V Q Q N K P T R S K R G M R R S H D A L T A V T S L S V D K T S G E K H L R H H I T A D G Y Y R G R K V I A K
6wdj_B A V Q Q N K P T R S K R G M R R S H D A L T A V T S L S V D K T S G E K H L R H H I T A D G Y Y R G R K V I A K
6wdk_B A V Q Q N K P T R S K R G M R R S H D A L T A V T S L S V D K T S G E K H L R H H I T A D G Y Y R G R K V I A K
6wdl_B A V Q Q N K P T R S K R G M R R S H D A L T A V T S L S V D K T S G E K H L R H H I T A D G Y Y R G R K V I A K
6wdm_B A V Q Q N K P T R S K R G M R R S H D A L T A V T S L S V D K T S G E K H L R H H I T A D G Y Y R G R K V I A K
6wnt_B A V Q Q N K P T R S K R G M R R S H D A L T A V T S L S V D K T S G E K H L R H H I T A D G Y Y R G R K V I A K
6wnv_B A V Q Q N K P T R S K R G M R R S H D A L T A V T S L S V D K T S G E K H L R H H I T A D G Y Y R G R K V I A K
6wnw_B A V Q Q N K P T R S K R G M R R S H D A L T A V T S L S V D K T S G E K H L R H H I T A D G Y Y R G R K V I A K
6ww4_B A R T K Q T I H P G V T C D G C Q M F P I N G S R F K C R N C D D F D F C E T C F K T K K H N T R H T F G R I N
6ww4_A A R T K Q T I H P G V T C D G C Q M F P I N G S R F K C R N C D D F D F C E T C F K T K K H N T R H T F G R I N
6xiq_AT M A H E N V W F S H P R R Y G K G S R Q C R V C S S H T G L I R K Y G L N I C R Q C F R E K A N D I G F N K F R
6xir_AT M A H E N V W F S H P R R Y G K G S R Q C R V C S S H T G L I R K Y G L N I C R Q C F R E K A N D I G F N K F R
6xz7_a A V Q Q N K P T R S K R G M R R S H D A L T A V T S L S V D K T S G E K H L R H H I T A D G Y Y R G R K V I A K
6y0g_Sd M G H Q Q L Y W S H P R K F G Q G S R S C R V C S N R H G L I R K Y G L N M C R Q C F R Q Y A K D I G F I K L D
6y2l_Sd M G H Q Q L Y W S H P R K F G Q G S R S C R V C S N R H G L I R K Y G L N M C R Q C F R Q Y A K D I G F I K L D
6y57_Sd M G H Q Q L Y W S H P R K F G Q G S R S C R V C S N R H G L I R K Y G L N M C R Q C F R Q Y A K D I G F I K L D
6y69_0 A V Q Q N K P T R S K R G M R R S H D A L T A V T S L S V D K T S G E K H L R H H I T A D G Y Y R G R K V I A K
6y7c_d M A H E N V W F S H P R R Y G K G S R Q C R V C S S H T G L I R K Y G L N I C R Q C F R E K A N D I G F N K F R
6yal_e M G H Q Q L Y W S H P R K F G Q G S R S C R V C S N R H G L I R K Y G L N M C R Q C F R Q Y A K D I G F I K L D
6ybs_i M G H Q Q L Y W S H P R K F G Q G S R S C R V C S N R H G L I R K Y G L N M C R Q C F R Q Y A K D I G F I K L D
6z6j_Sd M A H E N V W F S H P R R Y G K G S R Q C R V C S S H T G L I R K Y G L N I C R Q C F R E K A N D I G F N K F R
6z6k_Sd M A H E N V W F S H P R R Y G K G S R Q C R V C S S H T G L I R K Y G L N I C R Q C F R E K A N D I G F N K F R
6z6l_Sd M G H Q Q L Y W S H P R K F G Q G S R S C R V C S N R H G L I R K Y G L N M C R Q C F R Q Y A K D I G F I K L D
6z6m_Sd M G H Q Q L Y W S H P R K F G Q G S R S C R V C S N R H G L I R K Y G L N M C R Q C F R Q Y A K D I G F I K L D
6z6n_Sd M G H Q Q L Y W S H P R K F G Q G S R S C R V C S N R H G L I R K Y G L N M C R Q C F R Q Y A K D I G F I K L D
6zlw_f M G H Q Q L Y W S H P R K F G Q G S R S C R V C S N R H G L I R K Y G L N M C R Q C F R Q Y A K D I G F I K L D
6zm7_Sd M G H Q Q L Y W S H P R K F G Q G S R S C R V C S N R H G L I R K Y G L N M C R Q C F R Q Y A K D I G F I K L D
6zme_Sd M G H Q Q L Y W S H P R K F G Q G S R S C R V C S N R H G L I R K Y G L N M C R Q C F R Q Y A K D I G F I K L D
6zmi_Sd M G H Q Q L Y W S H P R K F G Q G S R S C R V C S N R H G L I R K Y G L N M C R Q C F R Q Y A K D I G F I K L D
6zmo_Sd M G H Q Q L Y W S H P R K F G Q G S R S C R V C S N R H G L I R K Y G L N M C R Q C F R Q Y A K D I G F I K L D
6zmt_f M G H Q Q L Y W S H P R K F G Q G S R S C R V C S N R H G L I R K Y G L N M C R Q C F R Q Y A K D I G F I K L D
6zn5_e G S L A R A G K V R G Q T P K V A K Q E K K K K K T G R A K R R M Q Y N R R F V N V V P T F G K K K G P N A N S
6zoj_e G S L A R A G K V R G Q T P K V A K Q E K K K K K T G R A K R R M Q Y N R R F V N V V P T F G K K K G P N A N S
6zok_e G S L A R A G K V R G Q T P K V A K Q E K K K K K T G R A K R R M Q Y N R R F V N V V P T F G K K K G P N A N S
6zon_l M G H Q Q L Y W S H P R K F G Q G S R S C R V C S N R H G L I R K Y G L N M C R Q C F R Q Y A K D I G F I K L D
6zp4_l M G H Q Q L Y W S H P R K F G Q G S R S C R V C S N R H G L I R K Y G L N M C R Q C F R Q Y A K D I G F I K L D
```

### Table S8. 56-Residue Chains Contained in Binary Sequence Space.

```
BINARY_1 T T Y K L I L N L K Q A K E E A I K E L V D A G I A E K Y I K L I A N A K T V E G V W T L K D E I L K A T V T E
BINARY_2 T T Y K L I L N L K Q A K E E A I K E L V D A G I A E K Y I K L I A N A K T V E G V W T L K D E I L K F T V T E
BINARY_3 T T Y K L I L N L K Q A K E E A I K E L V D A G I A E K Y I K L I A N A K T V E G V W T L K D E I L T A T V T E
BINARY_4 T T Y K L I L N L K Q A K E E A I K E L V D A G I A E K Y I K L I A N A K T V E G V W T L K D E I L T F T V T E
BINARY_5 T T Y K L I L N L K Q A K E E A I K E L V D A G I A E K Y I K L I A N A K T V E G V W T L K D E I K K A T V T E
BINARY_6 T T Y K L I L N L K Q A K E E A I K E L V D A G I A E K Y I K L I A N A K T V E G V W T L K D E I K K F T V T E
BINARY_7 T T Y K L I L N L K Q A K E E A I K E L V D A G I A E K Y I K L I A N A K T V E G V W T L K D E I K T A T V T E
BINARY_8 T T Y K L I L N L K Q A K E E A I K E L V D A G I A E K Y I K L I A N A K T V E G V W T L K D E I K T F T V T E
BINARY_9 T T Y K L I L N L K Q A K E E A I K E L V D A G I A E K Y I K L I A N A K T V E G V W T L K D E T L K A T V T E
BINARY_10 T T Y K L I L N L K Q A K E E A I K E L V D A G I A E K Y I K L I A N A K T V E G V W T L K D E T L K F T V T E
BINARY_11 T T Y K L I L N L K Q A K E E A I K E L V D A G I A E K Y I K L I A N A K T V E G V W T L K D E T L T A T V T E
BINARY_12 T T Y K L I L N L K Q A K E E A I K E L V D A G I A E K Y I K L I A N A K T V E G V W T L K D E T L T F T V T E
BINARY_13 T T Y K L I L N L K Q A K E E A I K E L V D A G I A E K Y I K L I A N A K T V E G V W T L K D E T K K A T V T E
BINARY_14 T T Y K L I L N L K Q A K E E A I K E L V D A G I A E K Y I K L I A N A K T V E G V W T L K D E T K K F T V T E
BINARY_15 T T Y K L I L N L K Q A K E E A I K E L V D A G I A E K Y I K L I A N A K T V E G V W T L K D E T K T A T V T E
BINARY_16 T T Y K L I L N L K Q A K E E A I K E L V D A G I A E K Y I K L I A N A K T V E G V W T L K D E T K T F T V T E
```

BINARY_17 T T Y K L I L N L K Q A K E E A I K E L V D A G I A E K Y I K L I A N A K T V E G V W T Y K D E I L K A T V T E
BINARY_18 T T Y K L I L N L K Q A K E E A I K E L V D A G I A E K Y I K L I A N A K T V E G V W T Y K D E I L K F T V T E
BINARY_19 T T Y K L I L N L K Q A K E E A I K E L V D A G I A E K Y I K L I A N A K T V E G V W T Y K D E I L T A T V T E
BINARY_20 T T Y K L I L N L K Q A K E E A I K E L V D A G I A E K Y I K L I A N A K T V E G V W T Y K D E I L T F T V T E
BINARY_21 T T Y K L I L N L K Q A K E E A I K E L V D A G I A E K Y I K L I A N A K T V E G V W T Y K D E I K K A T V T E
BINARY_22 T T Y K L I L N L K Q A K E E A I K E L V D A G I A E K Y I K L I A N A K T V E G V W T Y K D E I K K F T V T E
BINARY_23 T T Y K L I L N L K Q A K E E A I K E L V D A G I A E K Y I K L I A N A K T V E G V W T Y K D E I K T A T V T E
BINARY_24 T T Y K L I L N L K Q A K E E A I K E L V D A G I A E K Y I K L I A N A K T V E G V W T Y K D E I K T F T V T E
BINARY_25 T T Y K L I L N L K Q A K E E A I K E L V D A G I A E K Y I K L I A N A K T V E G V W T Y K D E T L K A T V T E
BINARY_26 T T Y K L I L N L K Q A K E E A I K E L V D A G I A E K Y I K L I A N A K T V E G V W T Y K D E T L K F T V T E
BINARY_27 T T Y K L I L N L K Q A K E E A I K E L V D A G I A E K Y I K L I A N A K T V E G V W T Y K D E T L T A T V T E
BINARY_28 T T Y K L I L N L K Q A K E E A I K E L V D A G I A E K Y I K L I A N A K T V E G V W T Y K D E T L T F T V T E
BINARY_29 T T Y K L I L N L K Q A K E E A I K E L V D A G I A E K Y I K L I A N A K T V E G V W T Y K D E T K K A T V T E
BINARY_30 T T Y K L I L N L K Q A K E E A I K E L V D A G I A E K Y I K L I A N A K T V E G V W T Y K D E T K K F T V T E
BINARY_31 T T Y K L I L N L K Q A K E E A I K E L V D A G I A E K Y I K L I A N A K T V E G V W T Y K D E T K T A T V T E
BINARY_32 T T Y K L I L N L K Q A K E E A I K E L V D A G I A E K Y I K L I A N A K T V E G V W T Y K D E T K T F T V T E
BINARY_33 T T Y K L I L N L K Q A K E E A I K E L V D A G I A E K Y I K L Y A N A K T V E G V W T L K D E I L K A T V T E
BINARY_34 T T Y K L I L N L K Q A K E E A I K E L V D A G I A E K Y I K L Y A N A K T V E G V W T L K D E I L K F T V T E
BINARY_35 T T Y K L I L N L K Q A K E E A I K E L V D A G I A E K Y I K L Y A N A K T V E G V W T L K D E I L T A T V T E
BINARY_36 T T Y K L I L N L K Q A K E E A I K E L V D A G I A E K Y I K L Y A N A K T V E G V W T L K D E I L T F T V T E
BINARY_37 T T Y K L I L N L K Q A K E E A I K E L V D A G I A E K Y I K L Y A N A K T V E G V W T L K D E I K K A T V T E
BINARY_38 T T Y K L I L N L K Q A K E E A I K E L V D A G I A E K Y I K L Y A N A K T V E G V W T L K D E I K K F T V T E
BINARY_39 T T Y K L I L N L K Q A K E E A I K E L V D A G I A E K Y I K L Y A N A K T V E G V W T L K D E I K T A T V T E
BINARY_40 T T Y K L I L N L K Q A K E E A I K E L V D A G I A E K Y I K L Y A N A K T V E G V W T L K D E I K T F T V T E
BINARY_41 T T Y K L I L N L K Q A K E E A I K E L V D A G I A E K Y I K L Y A N A K T V E G V W T L K D E T L K A T V T E
BINARY_42 T T Y K L I L N L K Q A K E E A I K E L V D A G I A E K Y I K L Y A N A K T V E G V W T L K D E T L K F T V T E
BINARY_43 T T Y K L I L N L K Q A K E E A I K E L V D A G I A E K Y I K L Y A N A K T V E G V W T L K D E T L T A T V T E
BINARY_44 T T Y K L I L N L K Q A K E E A I K E L V D A G I A E K Y I K L Y A N A K T V E G V W T L K D E T L T F T V T E
BINARY_45 T T Y K L I L N L K Q A K E E A I K E L V D A G I A E K Y I K L Y A N A K T V E G V W T L K D E T K K A T V T E
BINARY_46 T T Y K L I L N L K Q A K E E A I K E L V D A G I A E K Y I K L Y A N A K T V E G V W T L K D E T K K F T V T E
BINARY_47 T T Y K L I L N L K Q A K E E A I K E L V D A G I A E K Y I K L Y A N A K T V E G V W T L K D E T K T A T V T E
BINARY_48 T T Y K L I L N L K Q A K E E A I K E L V D A G I A E K Y I K L Y A N A K T V E G V W T L K D E T K T F T V T E
BINARY_49 T T Y K L I L N L K Q A K E E A I K E L V D A G I A E K Y I K L Y A N A K T V E G V W T Y K D E I L K A T V T E
BINARY_50 T T Y K L I L N L K Q A K E E A I K E L V D A G I A E K Y I K L Y A N A K T V E G V W T Y K D E I L K F T V T E
BINARY_51 T T Y K L I L N L K Q A K E E A I K E L V D A G I A E K Y I K L Y A N A K T V E G V W T Y K D E I L T A T V T E
BINARY_52 T T Y K L I L N L K Q A K E E A I K E L V D A G I A E K Y I K L Y A N A K T V E G V W T Y K D E I L T F T V T E
BINARY_53 T T Y K L I L N L K Q A K E E A I K E L V D A G I A E K Y I K L Y A N A K T V E G V W T Y K D E I K K A T V T E
BINARY_54 T T Y K L I L N L K Q A K E E A I K E L V D A G I A E K Y I K L Y A N A K T V E G V W T Y K D E I K K F T V T E
BINARY_55 T T Y K L I L N L K Q A K E E A I K E L V D A G I A E K Y I K L Y A N A K T V E G V W T Y K D E I K T A T V T E
BINARY_56 T T Y K L I L N L K Q A K E E A I K E L V D A G I A E K Y I K L Y A N A K T V E G V W T Y K D E I K T F T V T E
BINARY_57 T T Y K L I L N L K Q A K E E A I K E L V D A G I A E K Y I K L Y A N A K T V E G V W T Y K D E T L K A T V T E
BINARY_58 T T Y K L I L N L K Q A K E E A I K E L V D A G I A E K Y I K L Y A N A K T V E G V W T Y K D E T L K F T V T E
BINARY_59 T T Y K L I L N L K Q A K E E A I K E L V D A G I A E K Y I K L Y A N A K T V E G V W T Y K D E T L T A T V T E
BINARY_60 T T Y K L I L N L K Q A K E E A I K E L V D A G I A E K Y I K L Y A N A K T V E G V W T Y K D E T L T F T V T E
BINARY_61 T T Y K L I L N L K Q A K E E A I K E L V D A G I A E K Y I K L Y A N A K T V E G V W T Y K D E T K K A T V T E
BINARY_62 T T Y K L I L N L K Q A K E E A I K E L V D A G I A E K Y I K L Y A N A K T V E G V W T Y K D E T K K F T V T E
BINARY_63 T T Y K L I L N L K Q A K E E A I K E L V D A G I A E K Y I K L Y A N A K T V E G V W T Y K D E T K T A T V T E
BINARY_64 T T Y K L I L N L K Q A K E E A I K E L V D A G I A E K Y I K L Y A N A K T V E G V W T Y K D E T K T F T V T E
BINARY_65 T T Y K L I L N L K Q A K E E A I K E L V D A G I A E K Y F K L I A N A K T V E G V W T L K D E I L K A T V T E
BINARY_66 T T Y K L I L N L K Q A K E E A I K E L V D A G I A E K Y F K L I A N A K T V E G V W T L K D E I L K F T V T E
BINARY_67 T T Y K L I L N L K Q A K E E A I K E L V D A G I A E K Y F K L I A N A K T V E G V W T L K D E I L T A T V T E
BINARY_68 T T Y K L I L N L K Q A K E E A I K E L V D A G I A E K Y F K L I A N A K T V E G V W T L K D E I L T F T V T E
BINARY_69 T T Y K L I L N L K Q A K E E A I K E L V D A G I A E K Y F K L I A N A K T V E G V W T L K D E I K K A T V T E
BINARY_70 T T Y K L I L N L K Q A K E E A I K E L V D A G I A E K Y F K L I A N A K T V E G V W T L K D E I K K F T V T E
BINARY_71 T T Y K L I L N L K Q A K E E A I K E L V D A G I A E K Y F K L I A N A K T V E G V W T L K D E I K T A T V T E
BINARY_72 T T Y K L I L N L K Q A K E E A I K E L V D A G I A E K Y F K L I A N A K T V E G V W T L K D E I K T F T V T E
BINARY_73 T T Y K L I L N L K Q A K E E A I K E L V D A G I A E K Y F K L I A N A K T V E G V W T L K D E T L K A T V T E
BINARY_74 T T Y K L I L N L K Q A K E E A I K E L V D A G I A E K Y F K L I A N A K T V E G V W T L K D E T L K F T V T E
BINARY_75 T T Y K L I L N L K Q A K E E A I K E L V D A G I A E K Y F K L I A N A K T V E G V W T L K D E T L T A T V T E
BINARY_76 T T Y K L I L N L K Q A K E E A I K E L V D A G I A E K Y F K L I A N A K T V E G V W T L K D E T L T F T V T E
BINARY_77 T T Y K L I L N L K Q A K E E A I K E L V D A G I A E K Y F K L I A N A K T V E G V W T L K D E T K K A T V T E
BINARY_78 T T Y K L I L N L K Q A K E E A I K E L V D A G I A E K Y F K L I A N A K T V E G V W T L K D E T K K F T V T E
BINARY_79 T T Y K L I L N L K Q A K E E A I K E L V D A G I A E K Y F K L I A N A K T V E G V W T L K D E T K T A T V T E
BINARY_80 T T Y K L I L N L K Q A K E E A I K E L V D A G I A E K Y F K L I A N A K T V E G V W T L K D E T K T F T V T E
BINARY_81 T T Y K L I L N L K Q A K E E A I K E L V D A G I A E K Y F K L I A N A K T V E G V W T Y K D E I L K A T V T E
BINARY_82 T T Y K L I L N L K Q A K E E A I K E L V D A G I A E K Y F K L I A N A K T V E G V W T Y K D E I L K F T V T E
BINARY_83 T T Y K L I L N L K Q A K E E A I K E L V D A G I A E K Y F K L I A N A K T V E G V W T Y K D E I L T A T V T E
BINARY_84 T T Y K L I L N L K Q A K E E A I K E L V D A G I A E K Y F K L I A N A K T V E G V W T Y K D E I L T F T V T E
BINARY_85 T T Y K L I L N L K Q A K E E A I K E L V D A G I A E K Y F K L I A N A K T V E G V W T Y K D E I K K A T V T E
BINARY_86 T T Y K L I L N L K Q A K E E A I K E L V D A G I A E K Y F K L I A N A K T V E G V W T Y K D E I K K F T V T E
BINARY_87 T T Y K L I L N L K Q A K E E A I K E L V D A G I A E K Y F K L I A N A K T V E G V W T Y K D E I K T A T V T E
BINARY_88 T T Y K L I L N L K Q A K E E A I K E L V D A G I A E K Y F K L I A N A K T V E G V W T Y K D E I K T F T V T E
BINARY_89 T T Y K L I L N L K Q A K E E A I K E L V D A G I A E K Y F K L I A N A K T V E G V W T Y K D E T L K A T V T E
BINARY_90 T T Y K L I L N L K Q A K E E A I K E L V D A G I A E K Y F K L I A N A K T V E G V W T Y K D E T L K F T V T E
BINARY_91 T T Y K L I L N L K Q A K E E A I K E L V D A G I A E K Y F K L I A N A K T V E G V W T Y K D E T L T A T V T E
BINARY_92 T T Y K L I L N L K Q A K E E A I K E L V D A G I A E K Y F K L I A N A K T V E G V W T Y K D E T L T F T V T E
BINARY_93 T T Y K L I L N L K Q A K E E A I K E L V D A G I A E K Y F K L I A N A K T V E G V W T Y K D E T K K A T V T E
BINARY_94 T T Y K L I L N L K Q A K E E A I K E L V D A G I A E K Y F K L I A N A K T V E G V W T Y K D E T K K F T V T E
BINARY_95 T T Y K L I L N L K Q A K E E A I K E L V D A G I A E K Y F K L I A N A K T V E G V W T Y K D E T K T A T V T E
BINARY_96 T T Y K L I L N L K Q A K E E A I K E L V D A G I A E K Y F K L I A N A K T V E G V W T Y K D E T K T F T V T E
BINARY_97 T T Y K L I L N L K Q A K E E A I K E L V D A G I A E K Y F K L Y A N A K T V E G V W T L K D E I L K A T V T E
BINARY_98 T T Y K L I L N L K Q A K E E A I K E L V D A G I A E K Y F K L Y A N A K T V E G V W T L K D E I L K F T V T E
BINARY_99 T T Y K L I L N L K Q A K E E A I K E L V D A G I A E K Y F K L Y A N A K T V E G V W T L K D E I L T A T V T E
BINARY_100 T T Y K L I L N L K Q A K E E A I K E L V D A G I A E K Y F K L Y A N A K T V E G V W T L K D E I L T F T V T E
BINARY_101 T T Y K L I L N L K Q A K E E A I K E L V D A G I A E K Y F K L Y A N A K T V E G V W T L K D E I K K A T V T E
BINARY_102 T T Y K L I L N L K Q A K E E A I K E L V D A G I A E K Y F K L Y A N A K T V E G V W T L K D E I K K F T V T E
BINARY_103 T T Y K L I L N L K Q A K E E A I K E L V D A G I A E K Y F K L Y A N A K T V E G V W T L K D E I K T A T V T E
BINARY_104 T T Y K L I L N L K Q A K E E A I K E L V D A G I A E K Y F K L Y A N A K T V E G V W T L K D E I K T F T V T E
BINARY_105 T T Y K L I L N L K Q A K E E A I K E L V D A G I A E K Y F K L Y A N A K T V E G V W T L K D E T L K A T V T E
BINARY_106 T T Y K L I L N L K Q A K E E A I K E L V D A G I A E K Y F K L Y A N A K T V E G V W T L K D E T L K F T V T E
BINARY_107 T T Y K L I L N L K Q A K E E A I K E L V D A G I A E K Y F K L Y A N A K T V E G V W T L K D E T L T A T V T E
BINARY_108 T T Y K L I L N L K Q A K E E A I K E L V D A G I A E K Y F K L Y A N A K T V E G V W T L K D E T L T F T V T E
BINARY_109 T T Y K L I L N L K Q A K E E A I K E L V D A G I A E K Y F K L Y A N A K T V E G V W T L K D E T K K A T V T E
BINARY_110 T T Y K L I L N L K Q A K E E A I K E L V D A G I A E K Y F K L Y A N A K T V E G V W T L K D E T K K F T V T E
BINARY_111 T T Y K L I L N L K Q A K E E A I K E L V D A G I A E K Y F K L Y A N A K T V E G V W T L K D E T K T A T V T E

BINARY_112 T T Y K L I L N L K Q A K E E A I K E L V D A G I A E K Y F K L Y A N A K T V E G V W T L K D E T K T F T V T E
BINARY_113 T T Y K L I L N L K Q A K E E A I K E L V D A G I A E K Y F K L Y A N A K T V E G V W T Y K D E I L K A T V T E
BINARY_114 T T Y K L I L N L K Q A K E E A I K E L V D A G I A E K Y F K L Y A N A K T V E G V W T Y K D E I L K F T V T E
BINARY_115 T T Y K L I L N L K Q A K E E A I K E L V D A G I A E K Y F K L Y A N A K T V E G V W T Y K D E I L T A T V T E
BINARY_116 T T Y K L I L N L K Q A K E E A I K E L V D A G I A E K Y F K L Y A N A K T V E G V W T Y K D E I L T F T V T E
BINARY_117 T T Y K L I L N L K Q A K E E A I K E L V D A G I A E K Y F K L Y A N A K T V E G V W T Y K D E I K K A T V T E
BINARY_118 T T Y K L I L N L K Q A K E E A I K E L V D A G I A E K Y F K L Y A N A K T V E G V W T Y K D E I K K F T V T E
BINARY_119 T T Y K L I L N L K Q A K E E A I K E L V D A G I A E K Y F K L Y A N A K T V E G V W T Y K D E I K T A T V T E
BINARY_120 T T Y K L I L N L K Q A K E E A I K E L V D A G I A E K Y F K L Y A N A K T V E G V W T Y K D E I K T F T V T E
BINARY_121 T T Y K L I L N L K Q A K E E A I K E L V D A G I A E K Y F K L Y A N A K T V E G V W T Y K D E T L K A T V T E
BINARY_122 T T Y K L I L N L K Q A K E E A I K E L V D A G I A E K Y F K L Y A N A K T V E G V W T Y K D E T L K F T V T E
BINARY_123 T T Y K L I L N L K Q A K E E A I K E L V D A G I A E K Y F K L Y A N A K T V E G V W T Y K D E T L T A T V T E
BINARY_124 T T Y K L I L N L K Q A K E E A I K E L V D A G I A E K Y F K L Y A N A K T V E G V W T Y K D E T L T F T V T E
BINARY_125 T T Y K L I L N L K Q A K E E A I K E L V D A G I A E K Y F K L Y A N A K T V E G V W T Y K D E T K K A T V T E
BINARY_126 T T Y K L I L N L K Q A K E E A I K E L V D A G I A E K Y F K L Y A N A K T V E G V W T Y K D E T K K F T V T E
BINARY_127 T T Y K L I L N L K Q A K E E A I K E L V D A G I A E K Y F K L Y A N A K T V E G V W T Y K D E T K T A T V T E
BINARY_128 T T Y K L I L N L K Q A K E E A I K E L V D A G I A E K Y F K L Y A N A K T V E G V W T Y K D E T K T F T V T E
BINARY_129 T T Y K L I L N L K Q A K E E A I K E L V D A G T A E K Y I K L I A N A K T V E G V W T L K D E I L K A T V T E
BINARY_130 T T Y K L I L N L K Q A K E E A I K E L V D A G T A E K Y I K L I A N A K T V E G V W T L K D E I L K F T V T E
BINARY_131 T T Y K L I L N L K Q A K E E A I K E L V D A G T A E K Y I K L I A N A K T V E G V W T L K D E I L T A T V T E
BINARY_132 T T Y K L I L N L K Q A K E E A I K E L V D A G T A E K Y I K L I A N A K T V E G V W T L K D E I L T F T V T E
BINARY_133 T T Y K L I L N L K Q A K E E A I K E L V D A G T A E K Y I K L I A N A K T V E G V W T L K D E I K K A T V T E
BINARY_134 T T Y K L I L N L K Q A K E E A I K E L V D A G T A E K Y I K L I A N A K T V E G V W T L K D E I K K F T V T E
BINARY_135 T T Y K L I L N L K Q A K E E A I K E L V D A G T A E K Y I K L I A N A K T V E G V W T L K D E I K T A T V T E
BINARY_136 T T Y K L I L N L K Q A K E E A I K E L V D A G T A E K Y I K L I A N A K T V E G V W T L K D E I K T F T V T E
BINARY_137 T T Y K L I L N L K Q A K E E A I K E L V D A G T A E K Y I K L I A N A K T V E G V W T L K D E T L K A T V T E
BINARY_138 T T Y K L I L N L K Q A K E E A I K E L V D A G T A E K Y I K L I A N A K T V E G V W T L K D E T L K F T V T E
BINARY_139 T T Y K L I L N L K Q A K E E A I K E L V D A G T A E K Y I K L I A N A K T V E G V W T L K D E T L T A T V T E
BINARY_140 T T Y K L I L N L K Q A K E E A I K E L V D A G T A E K Y I K L I A N A K T V E G V W T L K D E T L T F T V T E
BINARY_141 T T Y K L I L N L K Q A K E E A I K E L V D A G T A E K Y I K L I A N A K T V E G V W T L K D E T K K A T V T E
BINARY_142 T T Y K L I L N L K Q A K E E A I K E L V D A G T A E K Y I K L I A N A K T V E G V W T L K D E T K K F T V T E
BINARY_143 T T Y K L I L N L K Q A K E E A I K E L V D A G T A E K Y I K L I A N A K T V E G V W T L K D E T K T A T V T E
BINARY_144 T T Y K L I L N L K Q A K E E A I K E L V D A G T A E K Y I K L I A N A K T V E G V W T L K D E T K T F T V T E
BINARY_145 T T Y K L I L N L K Q A K E E A I K E L V D A G T A E K Y I K L I A N A K T V E G V W T Y K D E I L K A T V T E
BINARY_146 T T Y K L I L N L K Q A K E E A I K E L V D A G T A E K Y I K L I A N A K T V E G V W T Y K D E I L K F T V T E
BINARY_147 T T Y K L I L N L K Q A K E E A I K E L V D A G T A E K Y I K L I A N A K T V E G V W T Y K D E I L T A T V T E
BINARY_148 T T Y K L I L N L K Q A K E E A I K E L V D A G T A E K Y I K L I A N A K T V E G V W T Y K D E I L T F T V T E
BINARY_149 T T Y K L I L N L K Q A K E E A I K E L V D A G T A E K Y I K L I A N A K T V E G V W T Y K D E I K K A T V T E
BINARY_150 T T Y K L I L N L K Q A K E E A I K E L V D A G T A E K Y I K L I A N A K T V E G V W T Y K D E I K K F T V T E
BINARY_151 T T Y K L I L N L K Q A K E E A I K E L V D A G T A E K Y I K L I A N A K T V E G V W T Y K D E I K T A T V T E
BINARY_152 T T Y K L I L N L K Q A K E E A I K E L V D A G T A E K Y I K L I A N A K T V E G V W T Y K D E I K T F T V T E
BINARY_153 T T Y K L I L N L K Q A K E E A I K E L V D A G T A E K Y I K L I A N A K T V E G V W T Y K D E T L K A T V T E
BINARY_154 T T Y K L I L N L K Q A K E E A I K E L V D A G T A E K Y I K L I A N A K T V E G V W T Y K D E T L K F T V T E
BINARY_155 T T Y K L I L N L K Q A K E E A I K E L V D A G T A E K Y I K L I A N A K T V E G V W T Y K D E T L T A T V T E
BINARY_156 T T Y K L I L N L K Q A K E E A I K E L V D A G T A E K Y I K L I A N A K T V E G V W T Y K D E T L T F T V T E
BINARY_157 T T Y K L I L N L K Q A K E E A I K E L V D A G T A E K Y I K L I A N A K T V E G V W T Y K D E T K K A T V T E
BINARY_158 T T Y K L I L N L K Q A K E E A I K E L V D A G T A E K Y I K L I A N A K T V E G V W T Y K D E T K K F T V T E
BINARY_159 T T Y K L I L N L K Q A K E E A I K E L V D A G T A E K Y I K L I A N A K T V E G V W T Y K D E T K T A T V T E
BINARY_160 T T Y K L I L N L K Q A K E E A I K E L V D A G T A E K Y I K L I A N A K T V E G V W T Y K D E T K T F T V T E
BINARY_161 T T Y K L I L N L K Q A K E E A I K E L V D A G T A E K Y I K L Y A N A K T V E G V W T L K D E I L K A T V T E
BINARY_162 T T Y K L I L N L K Q A K E E A I K E L V D A G T A E K Y I K L Y A N A K T V E G V W T L K D E I L K F T V T E
BINARY_163 T T Y K L I L N L K Q A K E E A I K E L V D A G T A E K Y I K L Y A N A K T V E G V W T L K D E I L T A T V T E
BINARY_164 T T Y K L I L N L K Q A K E E A I K E L V D A G T A E K Y I K L Y A N A K T V E G V W T L K D E I L T F T V T E
BINARY_165 T T Y K L I L N L K Q A K E E A I K E L V D A G T A E K Y I K L Y A N A K T V E G V W T L K D E I K K A T V T E
BINARY_166 T T Y K L I L N L K Q A K E E A I K E L V D A G T A E K Y I K L Y A N A K T V E G V W T L K D E I K K F T V T E
BINARY_167 T T Y K L I L N L K Q A K E E A I K E L V D A G T A E K Y I K L Y A N A K T V E G V W T L K D E I K T A T V T E
BINARY_168 T T Y K L I L N L K Q A K E E A I K E L V D A G T A E K Y I K L Y A N A K T V E G V W T L K D E I K T F T V T E
BINARY_169 T T Y K L I L N L K Q A K E E A I K E L V D A G T A E K Y I K L Y A N A K T V E G V W T L K D E T L K A T V T E
BINARY_170 T T Y K L I L N L K Q A K E E A I K E L V D A G T A E K Y I K L Y A N A K T V E G V W T L K D E T L K F T V T E
BINARY_171 T T Y K L I L N L K Q A K E E A I K E L V D A G T A E K Y I K L Y A N A K T V E G V W T L K D E T L T A T V T E
BINARY_172 T T Y K L I L N L K Q A K E E A I K E L V D A G T A E K Y I K L Y A N A K T V E G V W T L K D E T L T F T V T E
BINARY_173 T T Y K L I L N L K Q A K E E A I K E L V D A G T A E K Y I K L Y A N A K T V E G V W T L K D E T K K A T V T E
BINARY_174 T T Y K L I L N L K Q A K E E A I K E L V D A G T A E K Y I K L Y A N A K T V E G V W T L K D E T K K F T V T E
BINARY_175 T T Y K L I L N L K Q A K E E A I K E L V D A G T A E K Y I K L Y A N A K T V E G V W T L K D E T K T A T V T E
BINARY_176 T T Y K L I L N L K Q A K E E A I K E L V D A G T A E K Y I K L Y A N A K T V E G V W T L K D E T K T F T V T E
BINARY_177 T T Y K L I L N L K Q A K E E A I K E L V D A G T A E K Y I K L Y A N A K T V E G V W T Y K D E I L K A T V T E
BINARY_178 T T Y K L I L N L K Q A K E E A I K E L V D A G T A E K Y I K L Y A N A K T V E G V W T Y K D E I L K F T V T E
BINARY_179 T T Y K L I L N L K Q A K E E A I K E L V D A G T A E K Y I K L Y A N A K T V E G V W T Y K D E I L T A T V T E
BINARY_180 T T Y K L I L N L K Q A K E E A I K E L V D A G T A E K Y I K L Y A N A K T V E G V W T Y K D E I L T F T V T E
BINARY_181 T T Y K L I L N L K Q A K E E A I K E L V D A G T A E K Y I K L Y A N A K T V E G V W T Y K D E I K K A T V T E
BINARY_182 T T Y K L I L N L K Q A K E E A I K E L V D A G T A E K Y I K L Y A N A K T V E G V W T Y K D E I K K F T V T E
BINARY_183 T T Y K L I L N L K Q A K E E A I K E L V D A G T A E K Y I K L Y A N A K T V E G V W T Y K D E I K T A T V T E
BINARY_184 T T Y K L I L N L K Q A K E E A I K E L V D A G T A E K Y I K L Y A N A K T V E G V W T Y K D E I K T F T V T E
BINARY_185 T T Y K L I L N L K Q A K E E A I K E L V D A G T A E K Y I K L Y A N A K T V E G V W T Y K D E T L K A T V T E
BINARY_186 T T Y K L I L N L K Q A K E E A I K E L V D A G T A E K Y I K L Y A N A K T V E G V W T Y K D E T L K F T V T E
BINARY_187 T T Y K L I L N L K Q A K E E A I K E L V D A G T A E K Y I K L Y A N A K T V E G V W T Y K D E T L T A T V T E
BINARY_188 T T Y K L I L N L K Q A K E E A I K E L V D A G T A E K Y I K L Y A N A K T V E G V W T Y K D E T L T F T V T E
BINARY_189 T T Y K L I L N L K Q A K E E A I K E L V D A G T A E K Y I K L Y A N A K T V E G V W T Y K D E T K K A T V T E
BINARY_190 T T Y K L I L N L K Q A K E E A I K E L V D A G T A E K Y I K L Y A N A K T V E G V W T Y K D E T K K F T V T E
BINARY_191 T T Y K L I L N L K Q A K E E A I K E L V D A G T A E K Y I K L Y A N A K T V E G V W T Y K D E T K T A T V T E
BINARY_192 T T Y K L I L N L K Q A K E E A I K E L V D A G T A E K Y I K L Y A N A K T V E G V W T Y K D E T K T F T V T E
BINARY_193 T T Y K L I L N L K Q A K E E A I K E L V D A G T A E K Y F K L I A N A K T V E G V W T L K D E I L K A T V T E
BINARY_194 T T Y K L I L N L K Q A K E E A I K E L V D A G T A E K Y F K L I A N A K T V E G V W T L K D E I L K F T V T E
BINARY_195 T T Y K L I L N L K Q A K E E A I K E L V D A G T A E K Y F K L I A N A K T V E G V W T L K D E I L T A T V T E
BINARY_196 T T Y K L I L N L K Q A K E E A I K E L V D A G T A E K Y F K L I A N A K T V E G V W T L K D E I L T F T V T E
BINARY_197 T T Y K L I L N L K Q A K E E A I K E L V D A G T A E K Y F K L I A N A K T V E G V W T L K D E I K K A T V T E
BINARY_198 T T Y K L I L N L K Q A K E E A I K E L V D A G T A E K Y F K L I A N A K T V E G V W T L K D E I K K F T V T E
BINARY_199 T T Y K L I L N L K Q A K E E A I K E L V D A G T A E K Y F K L I A N A K T V E G V W T L K D E I K T A T V T E
BINARY_200 T T Y K L I L N L K Q A K E E A I K E L V D A G T A E K Y F K L I A N A K T V E G V W T L K D E I K T F T V T E
BINARY_201 T T Y K L I L N L K Q A K E E A I K E L V D A G T A E K Y F K L I A N A K T V E G V W T L K D E T L K A T V T E
BINARY_202 T T Y K L I L N L K Q A K E E A I K E L V D A G T A E K Y F K L I A N A K T V E G V W T L K D E T L K F T V T E
BINARY_203 T T Y K L I L N L K Q A K E E A I K E L V D A G T A E K Y F K L I A N A K T V E G V W T L K D E T L T A T V T E
BINARY_204 T T Y K L I L N L K Q A K E E A I K E L V D A G T A E K Y F K L I A N A K T V E G V W T L K D E T L T F T V T E
BINARY_205 T T Y K L I L N L K Q A K E E A I K E L V D A G T A E K Y F K L I A N A K T V E G V W T L K D E T K K A T V T E
BINARY_206 T T Y K L I L N L K Q A K E E A I K E L V D A G T A E K Y F K L I A N A K T V E G V W T L K D E T K K F T V T E

BINARY_207 T T Y K L I L N L K Q A K E E A I K E L V D A G T A E K Y F K L I A N A K T V E G V W T L K D E T K T A T V T E
BINARY_208 T T Y K L I L N L K Q A K E E A I K E L V D A G T A E K Y F K L I A N A K T V E G V W T L K D E T K T F T V T E
BINARY_209 T T Y K L I L N L K Q A K E E A I K E L V D A G T A E K Y F K L I A N A K T V E G V W T Y K D E I L K A T V T E
BINARY_210 T T Y K L I L N L K Q A K E E A I K E L V D A G T A E K Y F K L I A N A K T V E G V W T Y K D E I L K F T V T E
BINARY_211 T T Y K L I L N L K Q A K E E A I K E L V D A G T A E K Y F K L I A N A K T V E G V W T Y K D E I L T A T V T E
BINARY_212 T T Y K L I L N L K Q A K E E A I K E L V D A G T A E K Y F K L I A N A K T V E G V W T Y K D E I L T F T V T E
BINARY_213 T T Y K L I L N L K Q A K E E A I K E L V D A G T A E K Y F K L I A N A K T V E G V W T Y K D E I K K A T V T E
BINARY_214 T T Y K L I L N L K Q A K E E A I K E L V D A G T A E K Y F K L I A N A K T V E G V W T Y K D E I K K F T V T E
BINARY_215 T T Y K L I L N L K Q A K E E A I K E L V D A G T A E K Y F K L I A N A K T V E G V W T Y K D E I K T A T V T E
BINARY_216 T T Y K L I L N L K Q A K E E A I K E L V D A G T A E K Y F K L I A N A K T V E G V W T Y K D E I K T F T V T E
BINARY_217 T T Y K L I L N L K Q A K E E A I K E L V D A G T A E K Y F K L I A N A K T V E G V W T Y K D E T L K A T V T E
BINARY_218 T T Y K L I L N L K Q A K E E A I K E L V D A G T A E K Y F K L I A N A K T V E G V W T Y K D E T L K F T V T E
BINARY_219 T T Y K L I L N L K Q A K E E A I K E L V D A G T A E K Y F K L I A N A K T V E G V W T Y K D E T L T A T V T E
BINARY_220 T T Y K L I L N L K Q A K E E A I K E L V D A G T A E K Y F K L I A N A K T V E G V W T Y K D E T L T F T V T E
BINARY_221 T T Y K L I L N L K Q A K E E A I K E L V D A G T A E K Y F K L I A N A K T V E G V W T Y K D E T K K A T V T E
BINARY_222 T T Y K L I L N L K Q A K E E A I K E L V D A G T A E K Y F K L I A N A K T V E G V W T Y K D E T K K F T V T E
BINARY_223 T T Y K L I L N L K Q A K E E A I K E L V D A G T A E K Y F K L I A N A K T V E G V W T Y K D E T K T A T V T E
BINARY_224 T T Y K L I L N L K Q A K E E A I K E L V D A G T A E K Y F K L I A N A K T V E G V W T Y K D E T K T F T V T E
BINARY_225 T T Y K L I L N L K Q A K E E A I K E L V D A G T A E K Y F K L Y A N A K T V E G V W T L K D E I L K A T V T E
BINARY_226 T T Y K L I L N L K Q A K E E A I K E L V D A G T A E K Y F K L Y A N A K T V E G V W T L K D E I L K F T V T E
BINARY_227 T T Y K L I L N L K Q A K E E A I K E L V D A G T A E K Y F K L Y A N A K T V E G V W T L K D E I L T A T V T E
BINARY_228 T T Y K L I L N L K Q A K E E A I K E L V D A G T A E K Y F K L Y A N A K T V E G V W T L K D E I L T F T V T E
BINARY_229 T T Y K L I L N L K Q A K E E A I K E L V D A G T A E K Y F K L Y A N A K T V E G V W T L K D E I K K A T V T E
BINARY_230 T T Y K L I L N L K Q A K E E A I K E L V D A G T A E K Y F K L Y A N A K T V E G V W T L K D E I K K F T V T E
BINARY_231 T T Y K L I L N L K Q A K E E A I K E L V D A G T A E K Y F K L Y A N A K T V E G V W T L K D E I K T A T V T E
BINARY_232 T T Y K L I L N L K Q A K E E A I K E L V D A G T A E K Y F K L Y A N A K T V E G V W T L K D E I K T F T V T E
BINARY_233 T T Y K L I L N L K Q A K E E A I K E L V D A G T A E K Y F K L Y A N A K T V E G V W T L K D E T L K A T V T E
BINARY_234 T T Y K L I L N L K Q A K E E A I K E L V D A G T A E K Y F K L Y A N A K T V E G V W T L K D E T L K F T V T E
BINARY_235 T T Y K L I L N L K Q A K E E A I K E L V D A G T A E K Y F K L Y A N A K T V E G V W T L K D E T L T A T V T E
BINARY_236 T T Y K L I L N L K Q A K E E A I K E L V D A G T A E K Y F K L Y A N A K T V E G V W T L K D E T L T F T V T E
BINARY_237 T T Y K L I L N L K Q A K E E A I K E L V D A G T A E K Y F K L Y A N A K T V E G V W T L K D E T K K A T V T E
BINARY_238 T T Y K L I L N L K Q A K E E A I K E L V D A G T A E K Y F K L Y A N A K T V E G V W T L K D E T K K F T V T E
BINARY_239 T T Y K L I L N L K Q A K E E A I K E L V D A G T A E K Y F K L Y A N A K T V E G V W T L K D E T K T A T V T E
BINARY_240 T T Y K L I L N L K Q A K E E A I K E L V D A G T A E K Y F K L Y A N A K T V E G V W T L K D E T K T F T V T E
BINARY_241 T T Y K L I L N L K Q A K E E A I K E L V D A G T A E K Y F K L Y A N A K T V E G V W T Y K D E I L K A T V T E
BINARY_242 T T Y K L I L N L K Q A K E E A I K E L V D A G T A E K Y F K L Y A N A K T V E G V W T Y K D E I L K F T V T E
BINARY_243 T T Y K L I L N L K Q A K E E A I K E L V D A G T A E K Y F K L Y A N A K T V E G V W T Y K D E I L T A T V T E
BINARY_244 T T Y K L I L N L K Q A K E E A I K E L V D A G T A E K Y F K L Y A N A K T V E G V W T Y K D E I L T F T V T E
BINARY_245 T T Y K L I L N L K Q A K E E A I K E L V D A G T A E K Y F K L Y A N A K T V E G V W T Y K D E I K K A T V T E
BINARY_246 T T Y K L I L N L K Q A K E E A I K E L V D A G T A E K Y F K L Y A N A K T V E G V W T Y K D E I K K F T V T E
BINARY_247 T T Y K L I L N L K Q A K E E A I K E L V D A G T A E K Y F K L Y A N A K T V E G V W T Y K D E I K T A T V T E
BINARY_248 T T Y K L I L N L K Q A K E E A I K E L V D A G T A E K Y F K L Y A N A K T V E G V W T Y K D E I K T F T V T E
BINARY_249 T T Y K L I L N L K Q A K E E A I K E L V D A G T A E K Y F K L Y A N A K T V E G V W T Y K D E T L K A T V T E
BINARY_250 T T Y K L I L N L K Q A K E E A I K E L V D A G T A E K Y F K L Y A N A K T V E G V W T Y K D E T L K F T V T E
BINARY_251 T T Y K L I L N L K Q A K E E A I K E L V D A G T A E K Y F K L Y A N A K T V E G V W T Y K D E T L T A T V T E
BINARY_252 T T Y K L I L N L K Q A K E E A I K E L V D A G T A E K Y F K L Y A N A K T V E G V W T Y K D E T L T F T V T E
BINARY_253 T T Y K L I L N L K Q A K E E A I K E L V D A G T A E K Y F K L Y A N A K T V E G V W T Y K D E T K K A T V T E
BINARY_254 T T Y K L I L N L K Q A K E E A I K E L V D A G T A E K Y F K L Y A N A K T V E G V W T Y K D E T K K F T V T E
BINARY_255 T T Y K L I L N L K Q A K E E A I K E L V D A G T A E K Y F K L Y A N A K T V E G V W T Y K D E T K T A T V T E
BINARY_256 T T Y K L I L N L K Q A K E E A I K E L V D A G T A E K Y F K L Y A N A K T V E G V W T Y K D E T K T F T V T E
BINARY_257 T T Y K L I L N L K Q A K E E A I K E L V D A A I A E K Y I K L I A N A K T V E G V W T L K D E I L K A T V T E
BINARY_258 T T Y K L I L N L K Q A K E E A I K E L V D A A I A E K Y I K L I A N A K T V E G V W T L K D E I L K F T V T E
BINARY_259 T T Y K L I L N L K Q A K E E A I K E L V D A A I A E K Y I K L I A N A K T V E G V W T L K D E I L T A T V T E
BINARY_260 T T Y K L I L N L K Q A K E E A I K E L V D A A I A E K Y I K L I A N A K T V E G V W T L K D E I L T F T V T E
BINARY_261 T T Y K L I L N L K Q A K E E A I K E L V D A A I A E K Y I K L I A N A K T V E G V W T L K D E I K K A T V T E
BINARY_262 T T Y K L I L N L K Q A K E E A I K E L V D A A I A E K Y I K L I A N A K T V E G V W T L K D E I K K F T V T E
BINARY_263 T T Y K L I L N L K Q A K E E A I K E L V D A A I A E K Y I K L I A N A K T V E G V W T L K D E I K T A T V T E
BINARY_264 T T Y K L I L N L K Q A K E E A I K E L V D A A I A E K Y I K L I A N A K T V E G V W T L K D E I K T F T V T E
BINARY_265 T T Y K L I L N L K Q A K E E A I K E L V D A A I A E K Y I K L I A N A K T V E G V W T L K D E T L K A T V T E
BINARY_266 T T Y K L I L N L K Q A K E E A I K E L V D A A I A E K Y I K L I A N A K T V E G V W T L K D E T L K F T V T E
BINARY_267 T T Y K L I L N L K Q A K E E A I K E L V D A A I A E K Y I K L I A N A K T V E G V W T L K D E T L T A T V T E
BINARY_268 T T Y K L I L N L K Q A K E E A I K E L V D A A I A E K Y I K L I A N A K T V E G V W T L K D E T L T F T V T E
BINARY_269 T T Y K L I L N L K Q A K E E A I K E L V D A A I A E K Y I K L I A N A K T V E G V W T L K D E T K K A T V T E
BINARY_270 T T Y K L I L N L K Q A K E E A I K E L V D A A I A E K Y I K L I A N A K T V E G V W T L K D E T K K F T V T E
BINARY_271 T T Y K L I L N L K Q A K E E A I K E L V D A A I A E K Y I K L I A N A K T V E G V W T L K D E T K T A T V T E
BINARY_272 T T Y K L I L N L K Q A K E E A I K E L V D A A I A E K Y I K L I A N A K T V E G V W T L K D E T K T F T V T E
BINARY_273 T T Y K L I L N L K Q A K E E A I K E L V D A A I A E K Y I K L I A N A K T V E G V W T Y K D E I L K A T V T E
BINARY_274 T T Y K L I L N L K Q A K E E A I K E L V D A A I A E K Y I K L I A N A K T V E G V W T Y K D E I L K F T V T E
BINARY_275 T T Y K L I L N L K Q A K E E A I K E L V D A A I A E K Y I K L I A N A K T V E G V W T Y K D E I L T A T V T E
BINARY_276 T T Y K L I L N L K Q A K E E A I K E L V D A A I A E K Y I K L I A N A K T V E G V W T Y K D E I L T F T V T E
BINARY_277 T T Y K L I L N L K Q A K E E A I K E L V D A A I A E K Y I K L I A N A K T V E G V W T Y K D E I K K A T V T E
BINARY_278 T T Y K L I L N L K Q A K E E A I K E L V D A A I A E K Y I K L I A N A K T V E G V W T Y K D E I K K F T V T E
BINARY_279 T T Y K L I L N L K Q A K E E A I K E L V D A A I A E K Y I K L I A N A K T V E G V W T Y K D E I K T A T V T E
BINARY_280 T T Y K L I L N L K Q A K E E A I K E L V D A A I A E K Y I K L I A N A K T V E G V W T Y K D E I K T F T V T E
BINARY_281 T T Y K L I L N L K Q A K E E A I K E L V D A A I A E K Y I K L I A N A K T V E G V W T Y K D E T L K A T V T E
BINARY_282 T T Y K L I L N L K Q A K E E A I K E L V D A A I A E K Y I K L I A N A K T V E G V W T Y K D E T L K F T V T E
BINARY_283 T T Y K L I L N L K Q A K E E A I K E L V D A A I A E K Y I K L I A N A K T V E G V W T Y K D E T L T A T V T E
BINARY_284 T T Y K L I L N L K Q A K E E A I K E L V D A A I A E K Y I K L I A N A K T V E G V W T Y K D E T L T F T V T E
BINARY_285 T T Y K L I L N L K Q A K E E A I K E L V D A A I A E K Y I K L I A N A K T V E G V W T Y K D E T K K A T V T E
BINARY_286 T T Y K L I L N L K Q A K E E A I K E L V D A A I A E K Y I K L I A N A K T V E G V W T Y K D E T K K F T V T E
BINARY_287 T T Y K L I L N L K Q A K E E A I K E L V D A A I A E K Y I K L I A N A K T V E G V W T Y K D E T K T A T V T E
BINARY_288 T T Y K L I L N L K Q A K E E A I K E L V D A A I A E K Y I K L I A N A K T V E G V W T Y K D E T K T F T V T E
BINARY_289 T T Y K L I L N L K Q A K E E A I K E L V D A A I A E K Y I K L Y A N A K T V E G V W T L K D E I L K A T V T E
BINARY_290 T T Y K L I L N L K Q A K E E A I K E L V D A A I A E K Y I K L Y A N A K T V E G V W T L K D E I L K F T V T E
BINARY_291 T T Y K L I L N L K Q A K E E A I K E L V D A A I A E K Y I K L Y A N A K T V E G V W T L K D E I L T A T V T E
BINARY_292 T T Y K L I L N L K Q A K E E A I K E L V D A A I A E K Y I K L Y A N A K T V E G V W T L K D E I L T F T V T E
BINARY_293 T T Y K L I L N L K Q A K E E A I K E L V D A A I A E K Y I K L Y A N A K T V E G V W T L K D E I K K A T V T E
BINARY_294 T T Y K L I L N L K Q A K E E A I K E L V D A A I A E K Y I K L Y A N A K T V E G V W T L K D E I K K F T V T E
BINARY_295 T T Y K L I L N L K Q A K E E A I K E L V D A A I A E K Y I K L Y A N A K T V E G V W T L K D E I K T A T V T E
BINARY_296 T T Y K L I L N L K Q A K E E A I K E L V D A A I A E K Y I K L Y A N A K T V E G V W T L K D E I K T F T V T E
BINARY_297 T T Y K L I L N L K Q A K E E A I K E L V D A A I A E K Y I K L Y A N A K T V E G V W T L K D E T L K A T V T E
BINARY_298 T T Y K L I L N L K Q A K E E A I K E L V D A A I A E K Y I K L Y A N A K T V E G V W T L K D E T L K F T V T E
BINARY_299 T T Y K L I L N L K Q A K E E A I K E L V D A A I A E K Y I K L Y A N A K T V E G V W T L K D E T L T A T V T E
BINARY_300 T T Y K L I L N L K Q A K E E A I K E L V D A A I A E K Y I K L Y A N A K T V E G V W T L K D E T L T F T V T E
BINARY_301 T T Y K L I L N L K Q A K E E A I K E L V D A A I A E K Y I K L Y A N A K T V E G V W T L K D E T K K A T V T E

BINARY_302 T T Y K L I L N L K Q A K E E A I K E L V D A A I A E K Y I K L Y A N A K T V E G V W T L K D E T K K F T V T E
BINARY_303 T T Y K L I L N L K Q A K E E A I K E L V D A A I A E K Y I K L Y A N A K T V E G V W T L K D E T K T A T V T E
BINARY_304 T T Y K L I L N L K Q A K E E A I K E L V D A A I A E K Y I K L Y A N A K T V E G V W T L K D E T K T F T V T E
BINARY_305 T T Y K L I L N L K Q A K E E A I K E L V D A A I A E K Y I K L Y A N A K T V E G V W T Y K D E I L K A T V T E
BINARY_306 T T Y K L I L N L K Q A K E E A I K E L V D A A I A E K Y I K L Y A N A K T V E G V W T Y K D E I L K F T V T E
BINARY_307 T T Y K L I L N L K Q A K E E A I K E L V D A A I A E K Y I K L Y A N A K T V E G V W T Y K D E I L T A T V T E
BINARY_308 T T Y K L I L N L K Q A K E E A I K E L V D A A I A E K Y I K L Y A N A K T V E G V W T Y K D E I L T F T V T E
BINARY_309 T T Y K L I L N L K Q A K E E A I K E L V D A A I A E K Y I K L Y A N A K T V E G V W T Y K D E I K K A T V T E
BINARY_310 T T Y K L I L N L K Q A K E E A I K E L V D A A I A E K Y I K L Y A N A K T V E G V W T Y K D E I K K F T V T E
BINARY_311 T T Y K L I L N L K Q A K E E A I K E L V D A A I A E K Y I K L Y A N A K T V E G V W T Y K D E I K T A T V T E
BINARY_312 T T Y K L I L N L K Q A K E E A I K E L V D A A I A E K Y I K L Y A N A K T V E G V W T Y K D E I K T F T V T E
BINARY_313 T T Y K L I L N L K Q A K E E A I K E L V D A A I A E K Y I K L Y A N A K T V E G V W T Y K D E T L K A T V T E
BINARY_314 T T Y K L I L N L K Q A K E E A I K E L V D A A I A E K Y I K L Y A N A K T V E G V W T Y K D E T L K F T V T E
BINARY_315 T T Y K L I L N L K Q A K E E A I K E L V D A A I A E K Y I K L Y A N A K T V E G V W T Y K D E T L T A T V T E
BINARY_316 T T Y K L I L N L K Q A K E E A I K E L V D A A I A E K Y I K L Y A N A K T V E G V W T Y K D E T L T F T V T E
BINARY_317 T T Y K L I L N L K Q A K E E A I K E L V D A A I A E K Y I K L Y A N A K T V E G V W T Y K D E T K K A T V T E
BINARY_318 T T Y K L I L N L K Q A K E E A I K E L V D A A I A E K Y I K L Y A N A K T V E G V W T Y K D E T K K F T V T E
BINARY_319 T T Y K L I L N L K Q A K E E A I K E L V D A A I A E K Y I K L Y A N A K T V E G V W T Y K D E T K T A T V T E
BINARY_320 T T Y K L I L N L K Q A K E E A I K E L V D A A I A E K Y I K L Y A N A K T V E G V W T Y K D E T K T F T V T E
BINARY_321 T T Y K L I L N L K Q A K E E A I K E L V D A A I A E K Y F K L I A N A K T V E G V W T L K D E I L K A T V T E
BINARY_322 T T Y K L I L N L K Q A K E E A I K E L V D A A I A E K Y F K L I A N A K T V E G V W T L K D E I L K F T V T E
BINARY_323 T T Y K L I L N L K Q A K E E A I K E L V D A A I A E K Y F K L I A N A K T V E G V W T L K D E I L T A T V T E
BINARY_324 T T Y K L I L N L K Q A K E E A I K E L V D A A I A E K Y F K L I A N A K T V E G V W T L K D E I L T F T V T E
BINARY_325 T T Y K L I L N L K Q A K E E A I K E L V D A A I A E K Y F K L I A N A K T V E G V W T L K D E I K K A T V T E
BINARY_326 T T Y K L I L N L K Q A K E E A I K E L V D A A I A E K Y F K L I A N A K T V E G V W T L K D E I K K F T V T E
BINARY_327 T T Y K L I L N L K Q A K E E A I K E L V D A A I A E K Y F K L I A N A K T V E G V W T L K D E I K T A T V T E
BINARY_328 T T Y K L I L N L K Q A K E E A I K E L V D A A I A E K Y F K L I A N A K T V E G V W T L K D E I K T F T V T E
BINARY_329 T T Y K L I L N L K Q A K E E A I K E L V D A A I A E K Y F K L I A N A K T V E G V W T L K D E T L K A T V T E
BINARY_330 T T Y K L I L N L K Q A K E E A I K E L V D A A I A E K Y F K L I A N A K T V E G V W T L K D E T L K F T V T E
BINARY_331 T T Y K L I L N L K Q A K E E A I K E L V D A A I A E K Y F K L I A N A K T V E G V W T L K D E T L T A T V T E
BINARY_332 T T Y K L I L N L K Q A K E E A I K E L V D A A I A E K Y F K L I A N A K T V E G V W T L K D E T L T F T V T E
BINARY_333 T T Y K L I L N L K Q A K E E A I K E L V D A A I A E K Y F K L I A N A K T V E G V W T L K D E T K K A T V T E
BINARY_334 T T Y K L I L N L K Q A K E E A I K E L V D A A I A E K Y F K L I A N A K T V E G V W T L K D E T K K F T V T E
BINARY_335 T T Y K L I L N L K Q A K E E A I K E L V D A A I A E K Y F K L I A N A K T V E G V W T L K D E T K T A T V T E
BINARY_336 T T Y K L I L N L K Q A K E E A I K E L V D A A I A E K Y F K L I A N A K T V E G V W T L K D E T K T F T V T E
BINARY_337 T T Y K L I L N L K Q A K E E A I K E L V D A A I A E K Y F K L I A N A K T V E G V W T Y K D E I L K A T V T E
BINARY_338 T T Y K L I L N L K Q A K E E A I K E L V D A A I A E K Y F K L I A N A K T V E G V W T Y K D E I L K F T V T E
BINARY_339 T T Y K L I L N L K Q A K E E A I K E L V D A A I A E K Y F K L I A N A K T V E G V W T Y K D E I L T A T V T E
BINARY_340 T T Y K L I L N L K Q A K E E A I K E L V D A A I A E K Y F K L I A N A K T V E G V W T Y K D E I L T F T V T E
BINARY_341 T T Y K L I L N L K Q A K E E A I K E L V D A A I A E K Y F K L I A N A K T V E G V W T Y K D E I K K A T V T E
BINARY_342 T T Y K L I L N L K Q A K E E A I K E L V D A A I A E K Y F K L I A N A K T V E G V W T Y K D E I K K F T V T E
BINARY_343 T T Y K L I L N L K Q A K E E A I K E L V D A A I A E K Y F K L I A N A K T V E G V W T Y K D E I K T A T V T E
BINARY_344 T T Y K L I L N L K Q A K E E A I K E L V D A A I A E K Y F K L I A N A K T V E G V W T Y K D E I K T F T V T E
BINARY_345 T T Y K L I L N L K Q A K E E A I K E L V D A A I A E K Y F K L I A N A K T V E G V W T Y K D E T L K A T V T E
BINARY_346 T T Y K L I L N L K Q A K E E A I K E L V D A A I A E K Y F K L I A N A K T V E G V W T Y K D E T L K F T V T E
BINARY_347 T T Y K L I L N L K Q A K E E A I K E L V D A A I A E K Y F K L I A N A K T V E G V W T Y K D E T L T A T V T E
BINARY_348 T T Y K L I L N L K Q A K E E A I K E L V D A A I A E K Y F K L I A N A K T V E G V W T Y K D E T L T F T V T E
BINARY_349 T T Y K L I L N L K Q A K E E A I K E L V D A A I A E K Y F K L I A N A K T V E G V W T Y K D E T K K A T V T E
BINARY_350 T T Y K L I L N L K Q A K E E A I K E L V D A A I A E K Y F K L I A N A K T V E G V W T Y K D E T K K F T V T E
BINARY_351 T T Y K L I L N L K Q A K E E A I K E L V D A A I A E K Y F K L I A N A K T V E G V W T Y K D E T K T A T V T E
BINARY_352 T T Y K L I L N L K Q A K E E A I K E L V D A A I A E K Y F K L I A N A K T V E G V W T Y K D E T K T F T V T E
BINARY_353 T T Y K L I L N L K Q A K E E A I K E L V D A A I A E K Y F K L Y A N A K T V E G V W T L K D E I L K A T V T E
BINARY_354 T T Y K L I L N L K Q A K E E A I K E L V D A A I A E K Y F K L Y A N A K T V E G V W T L K D E I L K F T V T E
BINARY_355 T T Y K L I L N L K Q A K E E A I K E L V D A A I A E K Y F K L Y A N A K T V E G V W T L K D E I L T A T V T E
BINARY_356 T T Y K L I L N L K Q A K E E A I K E L V D A A I A E K Y F K L Y A N A K T V E G V W T L K D E I L T F T V T E
BINARY_357 T T Y K L I L N L K Q A K E E A I K E L V D A A I A E K Y F K L Y A N A K T V E G V W T L K D E I K K A T V T E
BINARY_358 T T Y K L I L N L K Q A K E E A I K E L V D A A I A E K Y F K L Y A N A K T V E G V W T L K D E I K K F T V T E
BINARY_359 T T Y K L I L N L K Q A K E E A I K E L V D A A I A E K Y F K L Y A N A K T V E G V W T L K D E I K T A T V T E
BINARY_360 T T Y K L I L N L K Q A K E E A I K E L V D A A I A E K Y F K L Y A N A K T V E G V W T L K D E I K T F T V T E
BINARY_361 T T Y K L I L N L K Q A K E E A I K E L V D A A I A E K Y F K L Y A N A K T V E G V W T L K D E T L K A T V T E
BINARY_362 T T Y K L I L N L K Q A K E E A I K E L V D A A I A E K Y F K L Y A N A K T V E G V W T L K D E T L K F T V T E
BINARY_363 T T Y K L I L N L K Q A K E E A I K E L V D A A I A E K Y F K L Y A N A K T V E G V W T L K D E T L T A T V T E
BINARY_364 T T Y K L I L N L K Q A K E E A I K E L V D A A I A E K Y F K L Y A N A K T V E G V W T L K D E T L T F T V T E
BINARY_365 T T Y K L I L N L K Q A K E E A I K E L V D A A I A E K Y F K L Y A N A K T V E G V W T L K D E T K K A T V T E
BINARY_366 T T Y K L I L N L K Q A K E E A I K E L V D A A I A E K Y F K L Y A N A K T V E G V W T L K D E T K K F T V T E
BINARY_367 T T Y K L I L N L K Q A K E E A I K E L V D A A I A E K Y F K L Y A N A K T V E G V W T L K D E T K T A T V T E
BINARY_368 T T Y K L I L N L K Q A K E E A I K E L V D A A I A E K Y F K L Y A N A K T V E G V W T L K D E T K T F T V T E
BINARY_369 T T Y K L I L N L K Q A K E E A I K E L V D A A I A E K Y F K L Y A N A K T V E G V W T Y K D E I L K A T V T E
BINARY_370 T T Y K L I L N L K Q A K E E A I K E L V D A A I A E K Y F K L Y A N A K T V E G V W T Y K D E I L K F T V T E
BINARY_371 T T Y K L I L N L K Q A K E E A I K E L V D A A I A E K Y F K L Y A N A K T V E G V W T Y K D E I L T A T V T E
BINARY_372 T T Y K L I L N L K Q A K E E A I K E L V D A A I A E K Y F K L Y A N A K T V E G V W T Y K D E I L T F T V T E
BINARY_373 T T Y K L I L N L K Q A K E E A I K E L V D A A I A E K Y F K L Y A N A K T V E G V W T Y K D E I K K A T V T E
BINARY_374 T T Y K L I L N L K Q A K E E A I K E L V D A A I A E K Y F K L Y A N A K T V E G V W T Y K D E I K K F T V T E
BINARY_375 T T Y K L I L N L K Q A K E E A I K E L V D A A I A E K Y F K L Y A N A K T V E G V W T Y K D E I K T A T V T E
BINARY_376 T T Y K L I L N L K Q A K E E A I K E L V D A A I A E K Y F K L Y A N A K T V E G V W T Y K D E I K T F T V T E
BINARY_377 T T Y K L I L N L K Q A K E E A I K E L V D A A I A E K Y F K L Y A N A K T V E G V W T Y K D E T L K A T V T E
BINARY_378 T T Y K L I L N L K Q A K E E A I K E L V D A A I A E K Y F K L Y A N A K T V E G V W T Y K D E T L K F T V T E
BINARY_379 T T Y K L I L N L K Q A K E E A I K E L V D A A I A E K Y F K L Y A N A K T V E G V W T Y K D E T L T A T V T E
BINARY_380 T T Y K L I L N L K Q A K E E A I K E L V D A A I A E K Y F K L Y A N A K T V E G V W T Y K D E T L T F T V T E
BINARY_381 T T Y K L I L N L K Q A K E E A I K E L V D A A I A E K Y F K L Y A N A K T V E G V W T Y K D E T K K A T V T E
BINARY_382 T T Y K L I L N L K Q A K E E A I K E L V D A A I A E K Y F K L Y A N A K T V E G V W T Y K D E T K K F T V T E
BINARY_383 T T Y K L I L N L K Q A K E E A I K E L V D A A I A E K Y F K L Y A N A K T V E G V W T Y K D E T K T A T V T E
BINARY_384 T T Y K L I L N L K Q A K E E A I K E L V D A A I A E K Y F K L Y A N A K T V E G V W T Y K D E T K T F T V T E
BINARY_385 T T Y K L I L N L K Q A K E E A I K E L V D A A T A E K Y I K L I A N A K T V E G V W T L K D E I L K A T V T E
BINARY_386 T T Y K L I L N L K Q A K E E A I K E L V D A A T A E K Y I K L I A N A K T V E G V W T L K D E I L K F T V T E
BINARY_387 T T Y K L I L N L K Q A K E E A I K E L V D A A T A E K Y I K L I A N A K T V E G V W T L K D E I L T A T V T E
BINARY_388 T T Y K L I L N L K Q A K E E A I K E L V D A A T A E K Y I K L I A N A K T V E G V W T L K D E I L T F T V T E
BINARY_389 T T Y K L I L N L K Q A K E E A I K E L V D A A T A E K Y I K L I A N A K T V E G V W T L K D E I K K A T V T E
BINARY_390 T T Y K L I L N L K Q A K E E A I K E L V D A A T A E K Y I K L I A N A K T V E G V W T L K D E I K K F T V T E
BINARY_391 T T Y K L I L N L K Q A K E E A I K E L V D A A T A E K Y I K L I A N A K T V E G V W T L K D E I K T A T V T E
BINARY_392 T T Y K L I L N L K Q A K E E A I K E L V D A A T A E K Y I K L I A N A K T V E G V W T L K D E I K T F T V T E
BINARY_393 T T Y K L I L N L K Q A K E E A I K E L V D A A T A E K Y I K L I A N A K T V E G V W T L K D E T L K A T V T E
BINARY_394 T T Y K L I L N L K Q A K E E A I K E L V D A A T A E K Y I K L I A N A K T V E G V W T L K D E T L K F T V T E
BINARY_395 T T Y K L I L N L K Q A K E E A I K E L V D A A T A E K Y I K L I A N A K T V E G V W T L K D E T L T A T V T E
BINARY_396 T T Y K L I L N L K Q A K E E A I K E L V D A A T A E K Y I K L I A N A K T V E G V W T L K D E T L T F T V T E

BINARY_397 T T Y K L I L N L K Q A K E E A I K E L V D A A T A E K Y I K L I A N A K T V E G V W T L K D E T K K A T V T E
BINARY_398 T T Y K L I L N L K Q A K E E A I K E L V D A A T A E K Y I K L I A N A K T V E G V W T L K D E T K K F T V T E
BINARY_399 T T Y K L I L N L K Q A K E E A I K E L V D A A T A E K Y I K L I A N A K T V E G V W T L K D E T K T A T V T E
BINARY_400 T T Y K L I L N L K Q A K E E A I K E L V D A A T A E K Y I K L I A N A K T V E G V W T L K D E T K T F T V T E
BINARY_401 T T Y K L I L N L K Q A K E E A I K E L V D A A T A E K Y I K L I A N A K T V E G V W T Y K D E I L K A T V T E
BINARY_402 T T Y K L I L N L K Q A K E E A I K E L V D A A T A E K Y I K L I A N A K T V E G V W T Y K D E I L K F T V T E
BINARY_403 T T Y K L I L N L K Q A K E E A I K E L V D A A T A E K Y I K L I A N A K T V E G V W T Y K D E I L T A T V T E
BINARY_404 T T Y K L I L N L K Q A K E E A I K E L V D A A T A E K Y I K L I A N A K T V E G V W T Y K D E I L T F T V T E
BINARY_405 T T Y K L I L N L K Q A K E E A I K E L V D A A T A E K Y I K L I A N A K T V E G V W T Y K D E I K K A T V T E
BINARY_406 T T Y K L I L N L K Q A K E E A I K E L V D A A T A E K Y I K L I A N A K T V E G V W T Y K D E I K K F T V T E
BINARY_407 T T Y K L I L N L K Q A K E E A I K E L V D A A T A E K Y I K L I A N A K T V E G V W T Y K D E I K T A T V T E
BINARY_408 T T Y K L I L N L K Q A K E E A I K E L V D A A T A E K Y I K L I A N A K T V E G V W T Y K D E I K T F T V T E
BINARY_409 T T Y K L I L N L K Q A K E E A I K E L V D A A T A E K Y I K L I A N A K T V E G V W T Y K D E T L K A T V T E
BINARY_410 T T Y K L I L N L K Q A K E E A I K E L V D A A T A E K Y I K L I A N A K T V E G V W T Y K D E T L K F T V T E
BINARY_411 T T Y K L I L N L K Q A K E E A I K E L V D A A T A E K Y I K L I A N A K T V E G V W T Y K D E T L T A T V T E
BINARY_412 T T Y K L I L N L K Q A K E E A I K E L V D A A T A E K Y I K L I A N A K T V E G V W T Y K D E T L T F T V T E
BINARY_413 T T Y K L I L N L K Q A K E E A I K E L V D A A T A E K Y I K L I A N A K T V E G V W T Y K D E T K K A T V T E
BINARY_414 T T Y K L I L N L K Q A K E E A I K E L V D A A T A E K Y I K L I A N A K T V E G V W T Y K D E T K K F T V T E
BINARY_415 T T Y K L I L N L K Q A K E E A I K E L V D A A T A E K Y I K L I A N A K T V E G V W T Y K D E T K T A T V T E
BINARY_416 T T Y K L I L N L K Q A K E E A I K E L V D A A T A E K Y I K L I A N A K T V E G V W T Y K D E T K T F T V T E
BINARY_417 T T Y K L I L N L K Q A K E E A I K E L V D A A T A E K Y I K L Y A N A K T V E G V W T L K D E I L K A T V T E
BINARY_418 T T Y K L I L N L K Q A K E E A I K E L V D A A T A E K Y I K L Y A N A K T V E G V W T L K D E I L K F T V T E
BINARY_419 T T Y K L I L N L K Q A K E E A I K E L V D A A T A E K Y I K L Y A N A K T V E G V W T L K D E I L T A T V T E
BINARY_420 T T Y K L I L N L K Q A K E E A I K E L V D A A T A E K Y I K L Y A N A K T V E G V W T L K D E I L T F T V T E
BINARY_421 T T Y K L I L N L K Q A K E E A I K E L V D A A T A E K Y I K L Y A N A K T V E G V W T L K D E I K K A T V T E
BINARY_422 T T Y K L I L N L K Q A K E E A I K E L V D A A T A E K Y I K L Y A N A K T V E G V W T L K D E I K K F T V T E
BINARY_423 T T Y K L I L N L K Q A K E E A I K E L V D A A T A E K Y I K L Y A N A K T V E G V W T L K D E I K T A T V T E
BINARY_424 T T Y K L I L N L K Q A K E E A I K E L V D A A T A E K Y I K L Y A N A K T V E G V W T L K D E I K T F T V T E
BINARY_425 T T Y K L I L N L K Q A K E E A I K E L V D A A T A E K Y I K L Y A N A K T V E G V W T L K D E T L K A T V T E
BINARY_426 T T Y K L I L N L K Q A K E E A I K E L V D A A T A E K Y I K L Y A N A K T V E G V W T L K D E T L K F T V T E
BINARY_427 T T Y K L I L N L K Q A K E E A I K E L V D A A T A E K Y I K L Y A N A K T V E G V W T L K D E T L T A T V T E
BINARY_428 T T Y K L I L N L K Q A K E E A I K E L V D A A T A E K Y I K L Y A N A K T V E G V W T L K D E T L T F T V T E
BINARY_429 T T Y K L I L N L K Q A K E E A I K E L V D A A T A E K Y I K L Y A N A K T V E G V W T L K D E T K K A T V T E
BINARY_430 T T Y K L I L N L K Q A K E E A I K E L V D A A T A E K Y I K L Y A N A K T V E G V W T L K D E T K K F T V T E
BINARY_431 T T Y K L I L N L K Q A K E E A I K E L V D A A T A E K Y I K L Y A N A K T V E G V W T L K D E T K T A T V T E
BINARY_432 T T Y K L I L N L K Q A K E E A I K E L V D A A T A E K Y I K L Y A N A K T V E G V W T L K D E T K T F T V T E
BINARY_433 T T Y K L I L N L K Q A K E E A I K E L V D A A T A E K Y I K L Y A N A K T V E G V W T Y K D E I L K A T V T E
BINARY_434 T T Y K L I L N L K Q A K E E A I K E L V D A A T A E K Y I K L Y A N A K T V E G V W T Y K D E I L K F T V T E
BINARY_435 T T Y K L I L N L K Q A K E E A I K E L V D A A T A E K Y I K L Y A N A K T V E G V W T Y K D E I L T A T V T E
BINARY_436 T T Y K L I L N L K Q A K E E A I K E L V D A A T A E K Y I K L Y A N A K T V E G V W T Y K D E I L T F T V T E
BINARY_437 T T Y K L I L N L K Q A K E E A I K E L V D A A T A E K Y I K L Y A N A K T V E G V W T Y K D E I K K A T V T E
BINARY_438 T T Y K L I L N L K Q A K E E A I K E L V D A A T A E K Y I K L Y A N A K T V E G V W T Y K D E I K K F T V T E
BINARY_439 T T Y K L I L N L K Q A K E E A I K E L V D A A T A E K Y I K L Y A N A K T V E G V W T Y K D E I K T A T V T E
BINARY_440 T T Y K L I L N L K Q A K E E A I K E L V D A A T A E K Y I K L Y A N A K T V E G V W T Y K D E I K T F T V T E
BINARY_441 T T Y K L I L N L K Q A K E E A I K E L V D A A T A E K Y I K L Y A N A K T V E G V W T Y K D E T L K A T V T E
BINARY_442 T T Y K L I L N L K Q A K E E A I K E L V D A A T A E K Y I K L Y A N A K T V E G V W T Y K D E T L K F T V T E
BINARY_443 T T Y K L I L N L K Q A K E E A I K E L V D A A T A E K Y I K L Y A N A K T V E G V W T Y K D E T L T A T V T E
BINARY_444 T T Y K L I L N L K Q A K E E A I K E L V D A A T A E K Y I K L Y A N A K T V E G V W T Y K D E T L T F T V T E
BINARY_445 T T Y K L I L N L K Q A K E E A I K E L V D A A T A E K Y I K L Y A N A K T V E G V W T Y K D E T K K A T V T E
BINARY_446 T T Y K L I L N L K Q A K E E A I K E L V D A A T A E K Y I K L Y A N A K T V E G V W T Y K D E T K K F T V T E
BINARY_447 T T Y K L I L N L K Q A K E E A I K E L V D A A T A E K Y I K L Y A N A K T V E G V W T Y K D E T K T A T V T E
BINARY_448 T T Y K L I L N L K Q A K E E A I K E L V D A A T A E K Y I K L Y A N A K T V E G V W T Y K D E T K T F T V T E
BINARY_449 T T Y K L I L N L K Q A K E E A I K E L V D A A T A E K Y F K L I A N A K T V E G V W T L K D E I L K A T V T E
BINARY_450 T T Y K L I L N L K Q A K E E A I K E L V D A A T A E K Y F K L I A N A K T V E G V W T L K D E I L K F T V T E
BINARY_451 T T Y K L I L N L K Q A K E E A I K E L V D A A T A E K Y F K L I A N A K T V E G V W T L K D E I L T A T V T E
BINARY_452 T T Y K L I L N L K Q A K E E A I K E L V D A A T A E K Y F K L I A N A K T V E G V W T L K D E I L T F T V T E
BINARY_453 T T Y K L I L N L K Q A K E E A I K E L V D A A T A E K Y F K L I A N A K T V E G V W T L K D E I K K A T V T E
BINARY_454 T T Y K L I L N L K Q A K E E A I K E L V D A A T A E K Y F K L I A N A K T V E G V W T L K D E I K K F T V T E
BINARY_455 T T Y K L I L N L K Q A K E E A I K E L V D A A T A E K Y F K L I A N A K T V E G V W T L K D E I K T A T V T E
BINARY_456 T T Y K L I L N L K Q A K E E A I K E L V D A A T A E K Y F K L I A N A K T V E G V W T L K D E I K T F T V T E
BINARY_457 T T Y K L I L N L K Q A K E E A I K E L V D A A T A E K Y F K L I A N A K T V E G V W T L K D E T L K A T V T E
BINARY_458 T T Y K L I L N L K Q A K E E A I K E L V D A A T A E K Y F K L I A N A K T V E G V W T L K D E T L K F T V T E
BINARY_459 T T Y K L I L N L K Q A K E E A I K E L V D A A T A E K Y F K L I A N A K T V E G V W T L K D E T L T A T V T E
BINARY_460 T T Y K L I L N L K Q A K E E A I K E L V D A A T A E K Y F K L I A N A K T V E G V W T L K D E T L T F T V T E
BINARY_461 T T Y K L I L N L K Q A K E E A I K E L V D A A T A E K Y F K L I A N A K T V E G V W T L K D E T K K A T V T E
BINARY_462 T T Y K L I L N L K Q A K E E A I K E L V D A A T A E K Y F K L I A N A K T V E G V W T L K D E T K K F T V T E
BINARY_463 T T Y K L I L N L K Q A K E E A I K E L V D A A T A E K Y F K L I A N A K T V E G V W T L K D E T K T A T V T E
BINARY_464 T T Y K L I L N L K Q A K E E A I K E L V D A A T A E K Y F K L I A N A K T V E G V W T L K D E T K T F T V T E
BINARY_465 T T Y K L I L N L K Q A K E E A I K E L V D A A T A E K Y F K L I A N A K T V E G V W T Y K D E I L K A T V T E
BINARY_466 T T Y K L I L N L K Q A K E E A I K E L V D A A T A E K Y F K L I A N A K T V E G V W T Y K D E I L K F T V T E
BINARY_467 T T Y K L I L N L K Q A K E E A I K E L V D A A T A E K Y F K L I A N A K T V E G V W T Y K D E I L T A T V T E
BINARY_468 T T Y K L I L N L K Q A K E E A I K E L V D A A T A E K Y F K L I A N A K T V E G V W T Y K D E I L T F T V T E
BINARY_469 T T Y K L I L N L K Q A K E E A I K E L V D A A T A E K Y F K L I A N A K T V E G V W T Y K D E I K K A T V T E
BINARY_470 T T Y K L I L N L K Q A K E E A I K E L V D A A T A E K Y F K L I A N A K T V E G V W T Y K D E I K K F T V T E
BINARY_471 T T Y K L I L N L K Q A K E E A I K E L V D A A T A E K Y F K L I A N A K T V E G V W T Y K D E I K T A T V T E
BINARY_472 T T Y K L I L N L K Q A K E E A I K E L V D A A T A E K Y F K L I A N A K T V E G V W T Y K D E I K T F T V T E
BINARY_473 T T Y K L I L N L K Q A K E E A I K E L V D A A T A E K Y F K L I A N A K T V E G V W T Y K D E T L K A T V T E
BINARY_474 T T Y K L I L N L K Q A K E E A I K E L V D A A T A E K Y F K L I A N A K T V E G V W T Y K D E T L K F T V T E
BINARY_475 T T Y K L I L N L K Q A K E E A I K E L V D A A T A E K Y F K L I A N A K T V E G V W T Y K D E T L T A T V T E
BINARY_476 T T Y K L I L N L K Q A K E E A I K E L V D A A T A E K Y F K L I A N A K T V E G V W T Y K D E T L T F T V T E
BINARY_477 T T Y K L I L N L K Q A K E E A I K E L V D A A T A E K Y F K L I A N A K T V E G V W T Y K D E T K K A T V T E
BINARY_478 T T Y K L I L N L K Q A K E E A I K E L V D A A T A E K Y F K L I A N A K T V E G V W T Y K D E T K K F T V T E
BINARY_479 T T Y K L I L N L K Q A K E E A I K E L V D A A T A E K Y F K L I A N A K T V E G V W T Y K D E T K T A T V T E
BINARY_480 T T Y K L I L N L K Q A K E E A I K E L V D A A T A E K Y F K L I A N A K T V E G V W T Y K D E T K T F T V T E
BINARY_481 T T Y K L I L N L K Q A K E E A I K E L V D A A T A E K Y F K L Y A N A K T V E G V W T L K D E I L K A T V T E
BINARY_482 T T Y K L I L N L K Q A K E E A I K E L V D A A T A E K Y F K L Y A N A K T V E G V W T L K D E I L K F T V T E
BINARY_483 T T Y K L I L N L K Q A K E E A I K E L V D A A T A E K Y F K L Y A N A K T V E G V W T L K D E I L T A T V T E
BINARY_484 T T Y K L I L N L K Q A K E E A I K E L V D A A T A E K Y F K L Y A N A K T V E G V W T L K D E I L T F T V T E
BINARY_485 T T Y K L I L N L K Q A K E E A I K E L V D A A T A E K Y F K L Y A N A K T V E G V W T L K D E I K K A T V T E
BINARY_486 T T Y K L I L N L K Q A K E E A I K E L V D A A T A E K Y F K L Y A N A K T V E G V W T L K D E I K K F T V T E
BINARY_487 T T Y K L I L N L K Q A K E E A I K E L V D A A T A E K Y F K L Y A N A K T V E G V W T L K D E I K T A T V T E
BINARY_488 T T Y K L I L N L K Q A K E E A I K E L V D A A T A E K Y F K L Y A N A K T V E G V W T L K D E I K T F T V T E
BINARY_489 T T Y K L I L N L K Q A K E E A I K E L V D A A T A E K Y F K L Y A N A K T V E G V W T L K D E T L K A T V T E
BINARY_490 T T Y K L I L N L K Q A K E E A I K E L V D A A T A E K Y F K L Y A N A K T V E G V W T L K D E T L K F T V T E
BINARY_491 T T Y K L I L N L K Q A K E E A I K E L V D A A T A E K Y F K L Y A N A K T V E G V W T L K D E T L T A T V T E

BINARY_492 T T Y K L I L N L K Q A K E E A I K E L V D A A T A E K Y F K L Y A N A K T V E G V W T L K D E T L T F T V T E
BINARY_493 T T Y K L I L N L K Q A K E E A I K E L V D A A T A E K Y F K L Y A N A K T V E G V W T L K D E T K K A T V T E
BINARY_494 T T Y K L I L N L K Q A K E E A I K E L V D A A T A E K Y F K L Y A N A K T V E G V W T L K D E T K K F T V T E
BINARY_495 T T Y K L I L N L K Q A K E E A I K E L V D A A T A E K Y F K L Y A N A K T V E G V W T L K D E T K T A T V T E
BINARY_496 T T Y K L I L N L K Q A K E E A I K E L V D A A T A E K Y F K L Y A N A K T V E G V W T L K D E T K T F T V T E
BINARY_497 T T Y K L I L N L K Q A K E E A I K E L V D A A T A E K Y F K L Y A N A K T V E G V W T Y K D E I L K A T V T E
BINARY_498 T T Y K L I L N L K Q A K E E A I K E L V D A A T A E K Y F K L Y A N A K T V E G V W T Y K D E I L K F T V T E
BINARY_499 T T Y K L I L N L K Q A K E E A I K E L V D A A T A E K Y F K L Y A N A K T V E G V W T Y K D E I L T A T V T E
BINARY_500 T T Y K L I L N L K Q A K E E A I K E L V D A A T A E K Y F K L Y A N A K T V E G V W T Y K D E I L T F T V T E
BINARY_501 T T Y K L I L N L K Q A K E E A I K E L V D A A T A E K Y F K L Y A N A K T V E G V W T Y K D E I K K A T V T E
BINARY_502 T T Y K L I L N L K Q A K E E A I K E L V D A A T A E K Y F K L Y A N A K T V E G V W T Y K D E I K K F T V T E
BINARY_503 T T Y K L I L N L K Q A K E E A I K E L V D A A T A E K Y F K L Y A N A K T V E G V W T Y K D E I K T A T V T E
BINARY_504 T T Y K L I L N L K Q A K E E A I K E L V D A A T A E K Y F K L Y A N A K T V E G V W T Y K D E I K T F T V T E
BINARY_505 T T Y K L I L N L K Q A K E E A I K E L V D A A T A E K Y F K L Y A N A K T V E G V W T Y K D E T L K A T V T E
BINARY_506 T T Y K L I L N L K Q A K E E A I K E L V D A A T A E K Y F K L Y A N A K T V E G V W T Y K D E T L K F T V T E
BINARY_507 T T Y K L I L N L K Q A K E E A I K E L V D A A T A E K Y F K L Y A N A K T V E G V W T Y K D E T L T A T V T E
BINARY_508 T T Y K L I L N L K Q A K E E A I K E L V D A A T A E K Y F K L Y A N A K T V E G V W T Y K D E T L T F T V T E
BINARY_509 T T Y K L I L N L K Q A K E E A I K E L V D A A T A E K Y F K L Y A N A K T V E G V W T Y K D E T K K A T V T E
BINARY_510 T T Y K L I L N L K Q A K E E A I K E L V D A A T A E K Y F K L Y A N A K T V E G V W T Y K D E T K K F T V T E
BINARY_511 T T Y K L I L N L K Q A K E E A I K E L V D A A T A E K Y F K L Y A N A K T V E G V W T Y K D E T K T A T V T E
BINARY_512 T T Y K L I L N L K Q A K E E A I K E L V D A A T A E K Y F K L Y A N A K T V E G V W T Y K D E T K T F T V T E
BINARY_513 T T Y K L I L N L K Q A K E E A I K E A V D A G I A E K Y I K L I A N A K T V E G V W T L K D E I L K A T V T E
BINARY_514 T T Y K L I L N L K Q A K E E A I K E A V D A G I A E K Y I K L I A N A K T V E G V W T L K D E I L K F T V T E
BINARY_515 T T Y K L I L N L K Q A K E E A I K E A V D A G I A E K Y I K L I A N A K T V E G V W T L K D E I L T A T V T E
BINARY_516 T T Y K L I L N L K Q A K E E A I K E A V D A G I A E K Y I K L I A N A K T V E G V W T L K D E I L T F T V T E
BINARY_517 T T Y K L I L N L K Q A K E E A I K E A V D A G I A E K Y I K L I A N A K T V E G V W T L K D E I K K A T V T E
BINARY_518 T T Y K L I L N L K Q A K E E A I K E A V D A G I A E K Y I K L I A N A K T V E G V W T L K D E I K K F T V T E
BINARY_519 T T Y K L I L N L K Q A K E E A I K E A V D A G I A E K Y I K L I A N A K T V E G V W T L K D E I K T A T V T E
BINARY_520 T T Y K L I L N L K Q A K E E A I K E A V D A G I A E K Y I K L I A N A K T V E G V W T L K D E I K T F T V T E
BINARY_521 T T Y K L I L N L K Q A K E E A I K E A V D A G I A E K Y I K L I A N A K T V E G V W T L K D E T L K A T V T E
BINARY_522 T T Y K L I L N L K Q A K E E A I K E A V D A G I A E K Y I K L I A N A K T V E G V W T L K D E T L K F T V T E
BINARY_523 T T Y K L I L N L K Q A K E E A I K E A V D A G I A E K Y I K L I A N A K T V E G V W T L K D E T L T A T V T E
BINARY_524 T T Y K L I L N L K Q A K E E A I K E A V D A G I A E K Y I K L I A N A K T V E G V W T L K D E T L T F T V T E
BINARY_525 T T Y K L I L N L K Q A K E E A I K E A V D A G I A E K Y I K L I A N A K T V E G V W T L K D E T K K A T V T E
BINARY_526 T T Y K L I L N L K Q A K E E A I K E A V D A G I A E K Y I K L I A N A K T V E G V W T L K D E T K K F T V T E
BINARY_527 T T Y K L I L N L K Q A K E E A I K E A V D A G I A E K Y I K L I A N A K T V E G V W T L K D E T K T A T V T E
BINARY_528 T T Y K L I L N L K Q A K E E A I K E A V D A G I A E K Y I K L I A N A K T V E G V W T L K D E T K T F T V T E
BINARY_529 T T Y K L I L N L K Q A K E E A I K E A V D A G I A E K Y I K L I A N A K T V E G V W T Y K D E I L K A T V T E
BINARY_530 T T Y K L I L N L K Q A K E E A I K E A V D A G I A E K Y I K L I A N A K T V E G V W T Y K D E I L K F T V T E
BINARY_531 T T Y K L I L N L K Q A K E E A I K E A V D A G I A E K Y I K L I A N A K T V E G V W T Y K D E I L T A T V T E
BINARY_532 T T Y K L I L N L K Q A K E E A I K E A V D A G I A E K Y I K L I A N A K T V E G V W T Y K D E I L T F T V T E
BINARY_533 T T Y K L I L N L K Q A K E E A I K E A V D A G I A E K Y I K L I A N A K T V E G V W T Y K D E I K K A T V T E
BINARY_534 T T Y K L I L N L K Q A K E E A I K E A V D A G I A E K Y I K L I A N A K T V E G V W T Y K D E I K K F T V T E
BINARY_535 T T Y K L I L N L K Q A K E E A I K E A V D A G I A E K Y I K L I A N A K T V E G V W T Y K D E I K T A T V T E
BINARY_536 T T Y K L I L N L K Q A K E E A I K E A V D A G I A E K Y I K L I A N A K T V E G V W T Y K D E I K T F T V T E
BINARY_537 T T Y K L I L N L K Q A K E E A I K E A V D A G I A E K Y I K L I A N A K T V E G V W T Y K D E T L K A T V T E
BINARY_538 T T Y K L I L N L K Q A K E E A I K E A V D A G I A E K Y I K L I A N A K T V E G V W T Y K D E T L K F T V T E
BINARY_539 T T Y K L I L N L K Q A K E E A I K E A V D A G I A E K Y I K L I A N A K T V E G V W T Y K D E T L T A T V T E
BINARY_540 T T Y K L I L N L K Q A K E E A I K E A V D A G I A E K Y I K L I A N A K T V E G V W T Y K D E T L T F T V T E
BINARY_541 T T Y K L I L N L K Q A K E E A I K E A V D A G I A E K Y I K L I A N A K T V E G V W T Y K D E T K K A T V T E
BINARY_542 T T Y K L I L N L K Q A K E E A I K E A V D A G I A E K Y I K L I A N A K T V E G V W T Y K D E T K K F T V T E
BINARY_543 T T Y K L I L N L K Q A K E E A I K E A V D A G I A E K Y I K L I A N A K T V E G V W T Y K D E T K T A T V T E
BINARY_544 T T Y K L I L N L K Q A K E E A I K E A V D A G I A E K Y I K L I A N A K T V E G V W T Y K D E T K T F T V T E
BINARY_545 T T Y K L I L N L K Q A K E E A I K E A V D A G I A E K Y I K L Y A N A K T V E G V W T L K D E I L K A T V T E
BINARY_546 T T Y K L I L N L K Q A K E E A I K E A V D A G I A E K Y I K L Y A N A K T V E G V W T L K D E I L K F T V T E
BINARY_547 T T Y K L I L N L K Q A K E E A I K E A V D A G I A E K Y I K L Y A N A K T V E G V W T L K D E I L T A T V T E
BINARY_548 T T Y K L I L N L K Q A K E E A I K E A V D A G I A E K Y I K L Y A N A K T V E G V W T L K D E I L T F T V T E
BINARY_549 T T Y K L I L N L K Q A K E E A I K E A V D A G I A E K Y I K L Y A N A K T V E G V W T L K D E I K K A T V T E
BINARY_550 T T Y K L I L N L K Q A K E E A I K E A V D A G I A E K Y I K L Y A N A K T V E G V W T L K D E I K K F T V T E
BINARY_551 T T Y K L I L N L K Q A K E E A I K E A V D A G I A E K Y I K L Y A N A K T V E G V W T L K D E I K T A T V T E
BINARY_552 T T Y K L I L N L K Q A K E E A I K E A V D A G I A E K Y I K L Y A N A K T V E G V W T L K D E I K T F T V T E
BINARY_553 T T Y K L I L N L K Q A K E E A I K E A V D A G I A E K Y I K L Y A N A K T V E G V W T L K D E T L K A T V T E
BINARY_554 T T Y K L I L N L K Q A K E E A I K E A V D A G I A E K Y I K L Y A N A K T V E G V W T L K D E T L K F T V T E
BINARY_555 T T Y K L I L N L K Q A K E E A I K E A V D A G I A E K Y I K L Y A N A K T V E G V W T L K D E T L T A T V T E
BINARY_556 T T Y K L I L N L K Q A K E E A I K E A V D A G I A E K Y I K L Y A N A K T V E G V W T L K D E T L T F T V T E
BINARY_557 T T Y K L I L N L K Q A K E E A I K E A V D A G I A E K Y I K L Y A N A K T V E G V W T L K D E T K K A T V T E
BINARY_558 T T Y K L I L N L K Q A K E E A I K E A V D A G I A E K Y I K L Y A N A K T V E G V W T L K D E T K K F T V T E
BINARY_559 T T Y K L I L N L K Q A K E E A I K E A V D A G I A E K Y I K L Y A N A K T V E G V W T L K D E T K T A T V T E
BINARY_560 T T Y K L I L N L K Q A K E E A I K E A V D A G I A E K Y I K L Y A N A K T V E G V W T L K D E T K T F T V T E
BINARY_561 T T Y K L I L N L K Q A K E E A I K E A V D A G I A E K Y I K L Y A N A K T V E G V W T Y K D E I L K A T V T E
BINARY_562 T T Y K L I L N L K Q A K E E A I K E A V D A G I A E K Y I K L Y A N A K T V E G V W T Y K D E I L K F T V T E
BINARY_563 T T Y K L I L N L K Q A K E E A I K E A V D A G I A E K Y I K L Y A N A K T V E G V W T Y K D E I L T A T V T E
BINARY_564 T T Y K L I L N L K Q A K E E A I K E A V D A G I A E K Y I K L Y A N A K T V E G V W T Y K D E I L T F T V T E
BINARY_565 T T Y K L I L N L K Q A K E E A I K E A V D A G I A E K Y I K L Y A N A K T V E G V W T Y K D E I K K A T V T E
BINARY_566 T T Y K L I L N L K Q A K E E A I K E A V D A G I A E K Y I K L Y A N A K T V E G V W T Y K D E I K K F T V T E
BINARY_567 T T Y K L I L N L K Q A K E E A I K E A V D A G I A E K Y I K L Y A N A K T V E G V W T Y K D E I K T A T V T E
BINARY_568 T T Y K L I L N L K Q A K E E A I K E A V D A G I A E K Y I K L Y A N A K T V E G V W T Y K D E I K T F T V T E
BINARY_569 T T Y K L I L N L K Q A K E E A I K E A V D A G I A E K Y I K L Y A N A K T V E G V W T Y K D E T L K A T V T E
BINARY_570 T T Y K L I L N L K Q A K E E A I K E A V D A G I A E K Y I K L Y A N A K T V E G V W T Y K D E T L K F T V T E
BINARY_571 T T Y K L I L N L K Q A K E E A I K E A V D A G I A E K Y I K L Y A N A K T V E G V W T Y K D E T L T A T V T E
BINARY_572 T T Y K L I L N L K Q A K E E A I K E A V D A G I A E K Y I K L Y A N A K T V E G V W T Y K D E T L T F T V T E
BINARY_573 T T Y K L I L N L K Q A K E E A I K E A V D A G I A E K Y I K L Y A N A K T V E G V W T Y K D E T K K A T V T E
BINARY_574 T T Y K L I L N L K Q A K E E A I K E A V D A G I A E K Y I K L Y A N A K T V E G V W T Y K D E T K K F T V T E
BINARY_575 T T Y K L I L N L K Q A K E E A I K E A V D A G I A E K Y I K L Y A N A K T V E G V W T Y K D E T K T A T V T E
BINARY_576 T T Y K L I L N L K Q A K E E A I K E A V D A G I A E K Y I K L Y A N A K T V E G V W T Y K D E T K T F T V T E
BINARY_577 T T Y K L I L N L K Q A K E E A I K E A V D A G I A E K Y F K L I A N A K T V E G V W T L K D E I L K A T V T E
BINARY_578 T T Y K L I L N L K Q A K E E A I K E A V D A G I A E K Y F K L I A N A K T V E G V W T L K D E I L K F T V T E
BINARY_579 T T Y K L I L N L K Q A K E E A I K E A V D A G I A E K Y F K L I A N A K T V E G V W T L K D E I L T A T V T E
BINARY_580 T T Y K L I L N L K Q A K E E A I K E A V D A G I A E K Y F K L I A N A K T V E G V W T L K D E I L T F T V T E
BINARY_581 T T Y K L I L N L K Q A K E E A I K E A V D A G I A E K Y F K L I A N A K T V E G V W T L K D E I K K A T V T E
BINARY_582 T T Y K L I L N L K Q A K E E A I K E A V D A G I A E K Y F K L I A N A K T V E G V W T L K D E I K K F T V T E
BINARY_583 T T Y K L I L N L K Q A K E E A I K E A V D A G I A E K Y F K L I A N A K T V E G V W T L K D E I K T A T V T E
BINARY_584 T T Y K L I L N L K Q A K E E A I K E A V D A G I A E K Y F K L I A N A K T V E G V W T L K D E I K T F T V T E
BINARY_585 T T Y K L I L N L K Q A K E E A I K E A V D A G I A E K Y F K L I A N A K T V E G V W T L K D E T L K A T V T E
BINARY_586 T T Y K L I L N L K Q A K E E A I K E A V D A G I A E K Y F K L I A N A K T V E G V W T L K D E T L K F T V T E

BINARY_587 T T Y K L I L N L K Q A K E E A I K E A V D A G I A E K Y F K L I A N A K T V E G V W T L K D E T L T A T V T E
BINARY_588 T T Y K L I L N L K Q A K E E A I K E A V D A G I A E K Y F K L I A N A K T V E G V W T L K D E T L T F T V T E
BINARY_589 T T Y K L I L N L K Q A K E E A I K E A V D A G I A E K Y F K L I A N A K T V E G V W T L K D E T K K A T V T E
BINARY_590 T T Y K L I L N L K Q A K E E A I K E A V D A G I A E K Y F K L I A N A K T V E G V W T L K D E T K K F T V T E
BINARY_591 T T Y K L I L N L K Q A K E E A I K E A V D A G I A E K Y F K L I A N A K T V E G V W T L K D E T K T A T V T E
BINARY_592 T T Y K L I L N L K Q A K E E A I K E A V D A G I A E K Y F K L I A N A K T V E G V W T L K D E T K T F T V T E
BINARY_593 T T Y K L I L N L K Q A K E E A I K E A V D A G I A E K Y F K L I A N A K T V E G V W T Y K D E I L K A T V T E
BINARY_594 T T Y K L I L N L K Q A K E E A I K E A V D A G I A E K Y F K L I A N A K T V E G V W T Y K D E I L K F T V T E
BINARY_595 T T Y K L I L N L K Q A K E E A I K E A V D A G I A E K Y F K L I A N A K T V E G V W T Y K D E I L T A T V T E
BINARY_596 T T Y K L I L N L K Q A K E E A I K E A V D A G I A E K Y F K L I A N A K T V E G V W T Y K D E I L T F T V T E
BINARY_597 T T Y K L I L N L K Q A K E E A I K E A V D A G I A E K Y F K L I A N A K T V E G V W T Y K D E I K K A T V T E
BINARY_598 T T Y K L I L N L K Q A K E E A I K E A V D A G I A E K Y F K L I A N A K T V E G V W T Y K D E I K K F T V T E
BINARY_599 T T Y K L I L N L K Q A K E E A I K E A V D A G I A E K Y F K L I A N A K T V E G V W T Y K D E I K T A T V T E
BINARY_600 T T Y K L I L N L K Q A K E E A I K E A V D A G I A E K Y F K L I A N A K T V E G V W T Y K D E I K T F T V T E
BINARY_601 T T Y K L I L N L K Q A K E E A I K E A V D A G I A E K Y F K L I A N A K T V E G V W T Y K D E T L K A T V T E
BINARY_602 T T Y K L I L N L K Q A K E E A I K E A V D A G I A E K Y F K L I A N A K T V E G V W T Y K D E T L K F T V T E
BINARY_603 T T Y K L I L N L K Q A K E E A I K E A V D A G I A E K Y F K L I A N A K T V E G V W T Y K D E T L T A T V T E
BINARY_604 T T Y K L I L N L K Q A K E E A I K E A V D A G I A E K Y F K L I A N A K T V E G V W T Y K D E T L T F T V T E
BINARY_605 T T Y K L I L N L K Q A K E E A I K E A V D A G I A E K Y F K L I A N A K T V E G V W T Y K D E T K K A T V T E
BINARY_606 T T Y K L I L N L K Q A K E E A I K E A V D A G I A E K Y F K L I A N A K T V E G V W T Y K D E T K K F T V T E
BINARY_607 T T Y K L I L N L K Q A K E E A I K E A V D A G I A E K Y F K L I A N A K T V E G V W T Y K D E T K T A T V T E
BINARY_608 T T Y K L I L N L K Q A K E E A I K E A V D A G I A E K Y F K L I A N A K T V E G V W T Y K D E T K T F T V T E
BINARY_609 T T Y K L I L N L K Q A K E E A I K E A V D A G I A E K Y F K L Y A N A K T V E G V W T L K D E I L K A T V T E
BINARY_610 T T Y K L I L N L K Q A K E E A I K E A V D A G I A E K Y F K L Y A N A K T V E G V W T L K D E I L K F T V T E
BINARY_611 T T Y K L I L N L K Q A K E E A I K E A V D A G I A E K Y F K L Y A N A K T V E G V W T L K D E I L T A T V T E
BINARY_612 T T Y K L I L N L K Q A K E E A I K E A V D A G I A E K Y F K L Y A N A K T V E G V W T L K D E I L T F T V T E
BINARY_613 T T Y K L I L N L K Q A K E E A I K E A V D A G I A E K Y F K L Y A N A K T V E G V W T L K D E I K K A T V T E
BINARY_614 T T Y K L I L N L K Q A K E E A I K E A V D A G I A E K Y F K L Y A N A K T V E G V W T L K D E I K K F T V T E
BINARY_615 T T Y K L I L N L K Q A K E E A I K E A V D A G I A E K Y F K L Y A N A K T V E G V W T L K D E I K T A T V T E
BINARY_616 T T Y K L I L N L K Q A K E E A I K E A V D A G I A E K Y F K L Y A N A K T V E G V W T L K D E I K T F T V T E
BINARY_617 T T Y K L I L N L K Q A K E E A I K E A V D A G I A E K Y F K L Y A N A K T V E G V W T L K D E T L K A T V T E
BINARY_618 T T Y K L I L N L K Q A K E E A I K E A V D A G I A E K Y F K L Y A N A K T V E G V W T L K D E T L K F T V T E
BINARY_619 T T Y K L I L N L K Q A K E E A I K E A V D A G I A E K Y F K L Y A N A K T V E G V W T L K D E T L T A T V T E
BINARY_620 T T Y K L I L N L K Q A K E E A I K E A V D A G I A E K Y F K L Y A N A K T V E G V W T L K D E T L T F T V T E
BINARY_621 T T Y K L I L N L K Q A K E E A I K E A V D A G I A E K Y F K L Y A N A K T V E G V W T L K D E T K K A T V T E
BINARY_622 T T Y K L I L N L K Q A K E E A I K E A V D A G I A E K Y F K L Y A N A K T V E G V W T L K D E T K K F T V T E
BINARY_623 T T Y K L I L N L K Q A K E E A I K E A V D A G I A E K Y F K L Y A N A K T V E G V W T L K D E T K T A T V T E
BINARY_624 T T Y K L I L N L K Q A K E E A I K E A V D A G I A E K Y F K L Y A N A K T V E G V W T L K D E T K T F T V T E
BINARY_625 T T Y K L I L N L K Q A K E E A I K E A V D A G I A E K Y F K L Y A N A K T V E G V W T Y K D E I L K A T V T E
BINARY_626 T T Y K L I L N L K Q A K E E A I K E A V D A G I A E K Y F K L Y A N A K T V E G V W T Y K D E I L K F T V T E
BINARY_627 T T Y K L I L N L K Q A K E E A I K E A V D A G I A E K Y F K L Y A N A K T V E G V W T Y K D E I L T A T V T E
BINARY_628 T T Y K L I L N L K Q A K E E A I K E A V D A G I A E K Y F K L Y A N A K T V E G V W T Y K D E I L T F T V T E
BINARY_629 T T Y K L I L N L K Q A K E E A I K E A V D A G I A E K Y F K L Y A N A K T V E G V W T Y K D E I K K A T V T E
BINARY_630 T T Y K L I L N L K Q A K E E A I K E A V D A G I A E K Y F K L Y A N A K T V E G V W T Y K D E I K K F T V T E
BINARY_631 T T Y K L I L N L K Q A K E E A I K E A V D A G I A E K Y F K L Y A N A K T V E G V W T Y K D E I K T A T V T E
BINARY_632 T T Y K L I L N L K Q A K E E A I K E A V D A G I A E K Y F K L Y A N A K T V E G V W T Y K D E I K T F T V T E
BINARY_633 T T Y K L I L N L K Q A K E E A I K E A V D A G I A E K Y F K L Y A N A K T V E G V W T Y K D E T L K A T V T E
BINARY_634 T T Y K L I L N L K Q A K E E A I K E A V D A G I A E K Y F K L Y A N A K T V E G V W T Y K D E T L K F T V T E
BINARY_635 T T Y K L I L N L K Q A K E E A I K E A V D A G I A E K Y F K L Y A N A K T V E G V W T Y K D E T L T A T V T E
BINARY_636 T T Y K L I L N L K Q A K E E A I K E A V D A G I A E K Y F K L Y A N A K T V E G V W T Y K D E T L T F T V T E
BINARY_637 T T Y K L I L N L K Q A K E E A I K E A V D A G I A E K Y F K L Y A N A K T V E G V W T Y K D E T K K A T V T E
BINARY_638 T T Y K L I L N L K Q A K E E A I K E A V D A G I A E K Y F K L Y A N A K T V E G V W T Y K D E T K K F T V T E
BINARY_639 T T Y K L I L N L K Q A K E E A I K E A V D A G I A E K Y F K L Y A N A K T V E G V W T Y K D E T K T A T V T E
BINARY_640 T T Y K L I L N L K Q A K E E A I K E A V D A G I A E K Y F K L Y A N A K T V E G V W T Y K D E T K T F T V T E
BINARY_641 T T Y K L I L N L K Q A K E E A I K E A V D A G T A E K Y I K L I A N A K T V E G V W T L K D E I L K A T V T E
BINARY_642 T T Y K L I L N L K Q A K E E A I K E A V D A G T A E K Y I K L I A N A K T V E G V W T L K D E I L K F T V T E
BINARY_643 T T Y K L I L N L K Q A K E E A I K E A V D A G T A E K Y I K L I A N A K T V E G V W T L K D E I L T A T V T E
BINARY_644 T T Y K L I L N L K Q A K E E A I K E A V D A G T A E K Y I K L I A N A K T V E G V W T L K D E I L T F T V T E
BINARY_645 T T Y K L I L N L K Q A K E E A I K E A V D A G T A E K Y I K L I A N A K T V E G V W T L K D E I K K A T V T E
BINARY_646 T T Y K L I L N L K Q A K E E A I K E A V D A G T A E K Y I K L I A N A K T V E G V W T L K D E I K K F T V T E
BINARY_647 T T Y K L I L N L K Q A K E E A I K E A V D A G T A E K Y I K L I A N A K T V E G V W T L K D E I K T A T V T E
BINARY_648 T T Y K L I L N L K Q A K E E A I K E A V D A G T A E K Y I K L I A N A K T V E G V W T L K D E I K T F T V T E
BINARY_649 T T Y K L I L N L K Q A K E E A I K E A V D A G T A E K Y I K L I A N A K T V E G V W T L K D E T L K A T V T E
BINARY_650 T T Y K L I L N L K Q A K E E A I K E A V D A G T A E K Y I K L I A N A K T V E G V W T L K D E T L K F T V T E
BINARY_651 T T Y K L I L N L K Q A K E E A I K E A V D A G T A E K Y I K L I A N A K T V E G V W T L K D E T L T A T V T E
BINARY_652 T T Y K L I L N L K Q A K E E A I K E A V D A G T A E K Y I K L I A N A K T V E G V W T L K D E T L T F T V T E
BINARY_653 T T Y K L I L N L K Q A K E E A I K E A V D A G T A E K Y I K L I A N A K T V E G V W T L K D E T K K A T V T E
BINARY_654 T T Y K L I L N L K Q A K E E A I K E A V D A G T A E K Y I K L I A N A K T V E G V W T L K D E T K K F T V T E
BINARY_655 T T Y K L I L N L K Q A K E E A I K E A V D A G T A E K Y I K L I A N A K T V E G V W T L K D E T K T A T V T E
BINARY_656 T T Y K L I L N L K Q A K E E A I K E A V D A G T A E K Y I K L I A N A K T V E G V W T L K D E T K T F T V T E
BINARY_657 T T Y K L I L N L K Q A K E E A I K E A V D A G T A E K Y I K L I A N A K T V E G V W T Y K D E I L K A T V T E
BINARY_658 T T Y K L I L N L K Q A K E E A I K E A V D A G T A E K Y I K L I A N A K T V E G V W T Y K D E I L K F T V T E
BINARY_659 T T Y K L I L N L K Q A K E E A I K E A V D A G T A E K Y I K L I A N A K T V E G V W T Y K D E I L T A T V T E
BINARY_660 T T Y K L I L N L K Q A K E E A I K E A V D A G T A E K Y I K L I A N A K T V E G V W T Y K D E I L T F T V T E
BINARY_661 T T Y K L I L N L K Q A K E E A I K E A V D A G T A E K Y I K L I A N A K T V E G V W T Y K D E I K K A T V T E
BINARY_662 T T Y K L I L N L K Q A K E E A I K E A V D A G T A E K Y I K L I A N A K T V E G V W T Y K D E I K K F T V T E
BINARY_663 T T Y K L I L N L K Q A K E E A I K E A V D A G T A E K Y I K L I A N A K T V E G V W T Y K D E I K T A T V T E
BINARY_664 T T Y K L I L N L K Q A K E E A I K E A V D A G T A E K Y I K L I A N A K T V E G V W T Y K D E I K T F T V T E
BINARY_665 T T Y K L I L N L K Q A K E E A I K E A V D A G T A E K Y I K L I A N A K T V E G V W T Y K D E T L K A T V T E
BINARY_666 T T Y K L I L N L K Q A K E E A I K E A V D A G T A E K Y I K L I A N A K T V E G V W T Y K D E T L K F T V T E
BINARY_667 T T Y K L I L N L K Q A K E E A I K E A V D A G T A E K Y I K L I A N A K T V E G V W T Y K D E T L T A T V T E
BINARY_668 T T Y K L I L N L K Q A K E E A I K E A V D A G T A E K Y I K L I A N A K T V E G V W T Y K D E T L T F T V T E
BINARY_669 T T Y K L I L N L K Q A K E E A I K E A V D A G T A E K Y I K L I A N A K T V E G V W T Y K D E T K K A T V T E
BINARY_670 T T Y K L I L N L K Q A K E E A I K E A V D A G T A E K Y I K L I A N A K T V E G V W T Y K D E T K K F T V T E
BINARY_671 T T Y K L I L N L K Q A K E E A I K E A V D A G T A E K Y I K L I A N A K T V E G V W T Y K D E T K T A T V T E
BINARY_672 T T Y K L I L N L K Q A K E E A I K E A V D A G T A E K Y I K L I A N A K T V E G V W T Y K D E T K T F T V T E
BINARY_673 T T Y K L I L N L K Q A K E E A I K E A V D A G T A E K Y I K L Y A N A K T V E G V W T L K D E I L K A T V T E
BINARY_674 T T Y K L I L N L K Q A K E E A I K E A V D A G T A E K Y I K L Y A N A K T V E G V W T L K D E I L K F T V T E
BINARY_675 T T Y K L I L N L K Q A K E E A I K E A V D A G T A E K Y I K L Y A N A K T V E G V W T L K D E I L T A T V T E
BINARY_676 T T Y K L I L N L K Q A K E E A I K E A V D A G T A E K Y I K L Y A N A K T V E G V W T L K D E I L T F T V T E
BINARY_677 T T Y K L I L N L K Q A K E E A I K E A V D A G T A E K Y I K L Y A N A K T V E G V W T L K D E I K K A T V T E
BINARY_678 T T Y K L I L N L K Q A K E E A I K E A V D A G T A E K Y I K L Y A N A K T V E G V W T L K D E I K K F T V T E
BINARY_679 T T Y K L I L N L K Q A K E E A I K E A V D A G T A E K Y I K L Y A N A K T V E G V W T L K D E I K T A T V T E
BINARY_680 T T Y K L I L N L K Q A K E E A I K E A V D A G T A E K Y I K L Y A N A K T V E G V W T L K D E I K T F T V T E
BINARY_681 T T Y K L I L N L K Q A K E E A I K E A V D A G T A E K Y I K L Y A N A K T V E G V W T L K D E T L K A T V T E

BINARY_682 T T Y K L I L N L K Q A K E E A I K E A V D A G T A E K Y I K L Y A N A K T V E G V W T L K D E T L K F T V T E
BINARY_683 T T Y K L I L N L K Q A K E E A I K E A V D A G T A E K Y I K L Y A N A K T V E G V W T L K D E T L T A T V T E
BINARY_684 T T Y K L I L N L K Q A K E E A I K E A V D A G T A E K Y I K L Y A N A K T V E G V W T L K D E T L T F T V T E
BINARY_685 T T Y K L I L N L K Q A K E E A I K E A V D A G T A E K Y I K L Y A N A K T V E G V W T L K D E T K K A T V T E
BINARY_686 T T Y K L I L N L K Q A K E E A I K E A V D A G T A E K Y I K L Y A N A K T V E G V W T L K D E T K K F T V T E
BINARY_687 T T Y K L I L N L K Q A K E E A I K E A V D A G T A E K Y I K L Y A N A K T V E G V W T L K D E T K T A T V T E
BINARY_688 T T Y K L I L N L K Q A K E E A I K E A V D A G T A E K Y I K L Y A N A K T V E G V W T L K D E T K T F T V T E
BINARY_689 T T Y K L I L N L K Q A K E E A I K E A V D A G T A E K Y I K L Y A N A K T V E G V W T Y K D E I L K A T V T E
BINARY_690 T T Y K L I L N L K Q A K E E A I K E A V D A G T A E K Y I K L Y A N A K T V E G V W T Y K D E I L K F T V T E
BINARY_691 T T Y K L I L N L K Q A K E E A I K E A V D A G T A E K Y I K L Y A N A K T V E G V W T Y K D E I L T A T V T E
BINARY_692 T T Y K L I L N L K Q A K E E A I K E A V D A G T A E K Y I K L Y A N A K T V E G V W T Y K D E I L T F T V T E
BINARY_693 T T Y K L I L N L K Q A K E E A I K E A V D A G T A E K Y I K L Y A N A K T V E G V W T Y K D E I K K A T V T E
BINARY_694 T T Y K L I L N L K Q A K E E A I K E A V D A G T A E K Y I K L Y A N A K T V E G V W T Y K D E I K K F T V T E
BINARY_695 T T Y K L I L N L K Q A K E E A I K E A V D A G T A E K Y I K L Y A N A K T V E G V W T Y K D E I K T A T V T E
BINARY_696 T T Y K L I L N L K Q A K E E A I K E A V D A G T A E K Y I K L Y A N A K T V E G V W T Y K D E I K T F T V T E
BINARY_697 T T Y K L I L N L K Q A K E E A I K E A V D A G T A E K Y I K L Y A N A K T V E G V W T Y K D E T L K A T V T E
BINARY_698 T T Y K L I L N L K Q A K E E A I K E A V D A G T A E K Y I K L Y A N A K T V E G V W T Y K D E T L K F T V T E
BINARY_699 T T Y K L I L N L K Q A K E E A I K E A V D A G T A E K Y I K L Y A N A K T V E G V W T Y K D E T L T A T V T E
BINARY_700 T T Y K L I L N L K Q A K E E A I K E A V D A G T A E K Y I K L Y A N A K T V E G V W T Y K D E T L T F T V T E
BINARY_701 T T Y K L I L N L K Q A K E E A I K E A V D A G T A E K Y I K L Y A N A K T V E G V W T Y K D E T K K A T V T E
BINARY_702 T T Y K L I L N L K Q A K E E A I K E A V D A G T A E K Y I K L Y A N A K T V E G V W T Y K D E T K K F T V T E
BINARY_703 T T Y K L I L N L K Q A K E E A I K E A V D A G T A E K Y I K L Y A N A K T V E G V W T Y K D E T K T A T V T E
BINARY_704 T T Y K L I L N L K Q A K E E A I K E A V D A G T A E K Y I K L Y A N A K T V E G V W T Y K D E T K T F T V T E
BINARY_705 T T Y K L I L N L K Q A K E E A I K E A V D A G T A E K Y F K L I A N A K T V E G V W T L K D E I L K A T V T E
BINARY_706 T T Y K L I L N L K Q A K E E A I K E A V D A G T A E K Y F K L I A N A K T V E G V W T L K D E I L K F T V T E
BINARY_707 T T Y K L I L N L K Q A K E E A I K E A V D A G T A E K Y F K L I A N A K T V E G V W T L K D E I L T A T V T E
BINARY_708 T T Y K L I L N L K Q A K E E A I K E A V D A G T A E K Y F K L I A N A K T V E G V W T L K D E I L T F T V T E
BINARY_709 T T Y K L I L N L K Q A K E E A I K E A V D A G T A E K Y F K L I A N A K T V E G V W T L K D E I K K A T V T E
BINARY_710 T T Y K L I L N L K Q A K E E A I K E A V D A G T A E K Y F K L I A N A K T V E G V W T L K D E I K K F T V T E
BINARY_711 T T Y K L I L N L K Q A K E E A I K E A V D A G T A E K Y F K L I A N A K T V E G V W T L K D E I K T A T V T E
BINARY_712 T T Y K L I L N L K Q A K E E A I K E A V D A G T A E K Y F K L I A N A K T V E G V W T L K D E I K T F T V T E
BINARY_713 T T Y K L I L N L K Q A K E E A I K E A V D A G T A E K Y F K L I A N A K T V E G V W T L K D E T L K A T V T E
BINARY_714 T T Y K L I L N L K Q A K E E A I K E A V D A G T A E K Y F K L I A N A K T V E G V W T L K D E T L K F T V T E
BINARY_715 T T Y K L I L N L K Q A K E E A I K E A V D A G T A E K Y F K L I A N A K T V E G V W T L K D E T L T A T V T E
BINARY_716 T T Y K L I L N L K Q A K E E A I K E A V D A G T A E K Y F K L I A N A K T V E G V W T L K D E T L T F T V T E
BINARY_717 T T Y K L I L N L K Q A K E E A I K E A V D A G T A E K Y F K L I A N A K T V E G V W T L K D E T K K A T V T E
BINARY_718 T T Y K L I L N L K Q A K E E A I K E A V D A G T A E K Y F K L I A N A K T V E G V W T L K D E T K K F T V T E
BINARY_719 T T Y K L I L N L K Q A K E E A I K E A V D A G T A E K Y F K L I A N A K T V E G V W T L K D E T K T A T V T E
BINARY_720 T T Y K L I L N L K Q A K E E A I K E A V D A G T A E K Y F K L I A N A K T V E G V W T L K D E T K T F T V T E
BINARY_721 T T Y K L I L N L K Q A K E E A I K E A V D A G T A E K Y F K L I A N A K T V E G V W T Y K D E I L K A T V T E
BINARY_722 T T Y K L I L N L K Q A K E E A I K E A V D A G T A E K Y F K L I A N A K T V E G V W T Y K D E I L K F T V T E
BINARY_723 T T Y K L I L N L K Q A K E E A I K E A V D A G T A E K Y F K L I A N A K T V E G V W T Y K D E I L T A T V T E
BINARY_724 T T Y K L I L N L K Q A K E E A I K E A V D A G T A E K Y F K L I A N A K T V E G V W T Y K D E I L T F T V T E
BINARY_725 T T Y K L I L N L K Q A K E E A I K E A V D A G T A E K Y F K L I A N A K T V E G V W T Y K D E I K K A T V T E
BINARY_726 T T Y K L I L N L K Q A K E E A I K E A V D A G T A E K Y F K L I A N A K T V E G V W T Y K D E I K K F T V T E
BINARY_727 T T Y K L I L N L K Q A K E E A I K E A V D A G T A E K Y F K L I A N A K T V E G V W T Y K D E I K T A T V T E
BINARY_728 T T Y K L I L N L K Q A K E E A I K E A V D A G T A E K Y F K L I A N A K T V E G V W T Y K D E I K T F T V T E
BINARY_729 T T Y K L I L N L K Q A K E E A I K E A V D A G T A E K Y F K L I A N A K T V E G V W T Y K D E T L K A T V T E
BINARY_730 T T Y K L I L N L K Q A K E E A I K E A V D A G T A E K Y F K L I A N A K T V E G V W T Y K D E T L K F T V T E
BINARY_731 T T Y K L I L N L K Q A K E E A I K E A V D A G T A E K Y F K L I A N A K T V E G V W T Y K D E T L T A T V T E
BINARY_732 T T Y K L I L N L K Q A K E E A I K E A V D A G T A E K Y F K L I A N A K T V E G V W T Y K D E T L T F T V T E
BINARY_733 T T Y K L I L N L K Q A K E E A I K E A V D A G T A E K Y F K L I A N A K T V E G V W T Y K D E T K K A T V T E
BINARY_734 T T Y K L I L N L K Q A K E E A I K E A V D A G T A E K Y F K L I A N A K T V E G V W T Y K D E T K K F T V T E
BINARY_735 T T Y K L I L N L K Q A K E E A I K E A V D A G T A E K Y F K L I A N A K T V E G V W T Y K D E T K T A T V T E
BINARY_736 T T Y K L I L N L K Q A K E E A I K E A V D A G T A E K Y F K L I A N A K T V E G V W T Y K D E T K T F T V T E
BINARY_737 T T Y K L I L N L K Q A K E E A I K E A V D A G T A E K Y F K L Y A N A K T V E G V W T L K D E I L K A T V T E
BINARY_738 T T Y K L I L N L K Q A K E E A I K E A V D A G T A E K Y F K L Y A N A K T V E G V W T L K D E I L K F T V T E
BINARY_739 T T Y K L I L N L K Q A K E E A I K E A V D A G T A E K Y F K L Y A N A K T V E G V W T L K D E I L T A T V T E
BINARY_740 T T Y K L I L N L K Q A K E E A I K E A V D A G T A E K Y F K L Y A N A K T V E G V W T L K D E I L T F T V T E
BINARY_741 T T Y K L I L N L K Q A K E E A I K E A V D A G T A E K Y F K L Y A N A K T V E G V W T L K D E I K K A T V T E
BINARY_742 T T Y K L I L N L K Q A K E E A I K E A V D A G T A E K Y F K L Y A N A K T V E G V W T L K D E I K K F T V T E
BINARY_743 T T Y K L I L N L K Q A K E E A I K E A V D A G T A E K Y F K L Y A N A K T V E G V W T L K D E I K T A T V T E
BINARY_744 T T Y K L I L N L K Q A K E E A I K E A V D A G T A E K Y F K L Y A N A K T V E G V W T L K D E I K T F T V T E
BINARY_745 T T Y K L I L N L K Q A K E E A I K E A V D A G T A E K Y F K L Y A N A K T V E G V W T L K D E T L K A T V T E
BINARY_746 T T Y K L I L N L K Q A K E E A I K E A V D A G T A E K Y F K L Y A N A K T V E G V W T L K D E T L K F T V T E
BINARY_747 T T Y K L I L N L K Q A K E E A I K E A V D A G T A E K Y F K L Y A N A K T V E G V W T L K D E T L T A T V T E
BINARY_748 T T Y K L I L N L K Q A K E E A I K E A V D A G T A E K Y F K L Y A N A K T V E G V W T L K D E T L T F T V T E
BINARY_749 T T Y K L I L N L K Q A K E E A I K E A V D A G T A E K Y F K L Y A N A K T V E G V W T L K D E T K K A T V T E
BINARY_750 T T Y K L I L N L K Q A K E E A I K E A V D A G T A E K Y F K L Y A N A K T V E G V W T L K D E T K K F T V T E
BINARY_751 T T Y K L I L N L K Q A K E E A I K E A V D A G T A E K Y F K L Y A N A K T V E G V W T L K D E T K T A T V T E
BINARY_752 T T Y K L I L N L K Q A K E E A I K E A V D A G T A E K Y F K L Y A N A K T V E G V W T L K D E T K T F T V T E
BINARY_753 T T Y K L I L N L K Q A K E E A I K E A V D A G T A E K Y F K L Y A N A K T V E G V W T Y K D E I L K A T V T E
BINARY_754 T T Y K L I L N L K Q A K E E A I K E A V D A G T A E K Y F K L Y A N A K T V E G V W T Y K D E I L K F T V T E
BINARY_755 T T Y K L I L N L K Q A K E E A I K E A V D A G T A E K Y F K L Y A N A K T V E G V W T Y K D E I L T A T V T E
BINARY_756 T T Y K L I L N L K Q A K E E A I K E A V D A G T A E K Y F K L Y A N A K T V E G V W T Y K D E I L T F T V T E
BINARY_757 T T Y K L I L N L K Q A K E E A I K E A V D A G T A E K Y F K L Y A N A K T V E G V W T Y K D E I K K A T V T E
BINARY_758 T T Y K L I L N L K Q A K E E A I K E A V D A G T A E K Y F K L Y A N A K T V E G V W T Y K D E I K K F T V T E
BINARY_759 T T Y K L I L N L K Q A K E E A I K E A V D A G T A E K Y F K L Y A N A K T V E G V W T Y K D E I K T A T V T E
BINARY_760 T T Y K L I L N L K Q A K E E A I K E A V D A G T A E K Y F K L Y A N A K T V E G V W T Y K D E I K T F T V T E
BINARY_761 T T Y K L I L N L K Q A K E E A I K E A V D A G T A E K Y F K L Y A N A K T V E G V W T Y K D E T L K A T V T E
BINARY_762 T T Y K L I L N L K Q A K E E A I K E A V D A G T A E K Y F K L Y A N A K T V E G V W T Y K D E T L K F T V T E
BINARY_763 T T Y K L I L N L K Q A K E E A I K E A V D A G T A E K Y F K L Y A N A K T V E G V W T Y K D E T L T A T V T E
BINARY_764 T T Y K L I L N L K Q A K E E A I K E A V D A G T A E K Y F K L Y A N A K T V E G V W T Y K D E T L T F T V T E
BINARY_765 T T Y K L I L N L K Q A K E E A I K E A V D A G T A E K Y F K L Y A N A K T V E G V W T Y K D E T K K A T V T E
BINARY_766 T T Y K L I L N L K Q A K E E A I K E A V D A G T A E K Y F K L Y A N A K T V E G V W T Y K D E T K K F T V T E
BINARY_767 T T Y K L I L N L K Q A K E E A I K E A V D A G T A E K Y F K L Y A N A K T V E G V W T Y K D E T K T A T V T E
BINARY_768 T T Y K L I L N L K Q A K E E A I K E A V D A G T A E K Y F K L Y A N A K T V E G V W T Y K D E T K T F T V T E
BINARY_769 T T Y K L I L N L K Q A K E E A I K E A V D A A I A E K Y I K L I A N A K T V E G V W T L K D E I L K A T V T E
BINARY_770 T T Y K L I L N L K Q A K E E A I K E A V D A A I A E K Y I K L I A N A K T V E G V W T L K D E I L K F T V T E
BINARY_771 T T Y K L I L N L K Q A K E E A I K E A V D A A I A E K Y I K L I A N A K T V E G V W T L K D E I L T A T V T E
BINARY_772 T T Y K L I L N L K Q A K E E A I K E A V D A A I A E K Y I K L I A N A K T V E G V W T L K D E I L T F T V T E
BINARY_773 T T Y K L I L N L K Q A K E E A I K E A V D A A I A E K Y I K L I A N A K T V E G V W T L K D E I K K A T V T E
BINARY_774 T T Y K L I L N L K Q A K E E A I K E A V D A A I A E K Y I K L I A N A K T V E G V W T L K D E I K K F T V T E
BINARY_775 T T Y K L I L N L K Q A K E E A I K E A V D A A I A E K Y I K L I A N A K T V E G V W T L K D E I K T A T V T E
BINARY_776 T T Y K L I L N L K Q A K E E A I K E A V D A A I A E K Y I K L I A N A K T V E G V W T L K D E I K T F T V T E

BINARY_777 T T Y K L I L N L K Q A K E E A I K E A V D A A I A E K Y I K L I A N A K T V E G V W T L K D E T L K A T V T E
BINARY_778 T T Y K L I L N L K Q A K E E A I K E A V D A A I A E K Y I K L I A N A K T V E G V W T L K D E T L K F T V T E
BINARY_779 T T Y K L I L N L K Q A K E E A I K E A V D A A I A E K Y I K L I A N A K T V E G V W T L K D E T L T A T V T E
BINARY_780 T T Y K L I L N L K Q A K E E A I K E A V D A A I A E K Y I K L I A N A K T V E G V W T L K D E T L T F T V T E
BINARY_781 T T Y K L I L N L K Q A K E E A I K E A V D A A I A E K Y I K L I A N A K T V E G V W T L K D E T K K A T V T E
BINARY_782 T T Y K L I L N L K Q A K E E A I K E A V D A A I A E K Y I K L I A N A K T V E G V W T L K D E T K K F T V T E
BINARY_783 T T Y K L I L N L K Q A K E E A I K E A V D A A I A E K Y I K L I A N A K T V E G V W T L K D E T K T A T V T E
BINARY_784 T T Y K L I L N L K Q A K E E A I K E A V D A A I A E K Y I K L I A N A K T V E G V W T L K D E T K T F T V T E
BINARY_785 T T Y K L I L N L K Q A K E E A I K E A V D A A I A E K Y I K L I A N A K T V E G V W T Y K D E I L K A T V T E
BINARY_786 T T Y K L I L N L K Q A K E E A I K E A V D A A I A E K Y I K L I A N A K T V E G V W T Y K D E I L K F T V T E
BINARY_787 T T Y K L I L N L K Q A K E E A I K E A V D A A I A E K Y I K L I A N A K T V E G V W T Y K D E I L T A T V T E
BINARY_788 T T Y K L I L N L K Q A K E E A I K E A V D A A I A E K Y I K L I A N A K T V E G V W T Y K D E I L T F T V T E
BINARY_789 T T Y K L I L N L K Q A K E E A I K E A V D A A I A E K Y I K L I A N A K T V E G V W T Y K D E I K K A T V T E
BINARY_790 T T Y K L I L N L K Q A K E E A I K E A V D A A I A E K Y I K L I A N A K T V E G V W T Y K D E I K K F T V T E
BINARY_791 T T Y K L I L N L K Q A K E E A I K E A V D A A I A E K Y I K L I A N A K T V E G V W T Y K D E I K T A T V T E
BINARY_792 T T Y K L I L N L K Q A K E E A I K E A V D A A I A E K Y I K L I A N A K T V E G V W T Y K D E I K T F T V T E
BINARY_793 T T Y K L I L N L K Q A K E E A I K E A V D A A I A E K Y I K L I A N A K T V E G V W T Y K D E T L K A T V T E
BINARY_794 T T Y K L I L N L K Q A K E E A I K E A V D A A I A E K Y I K L I A N A K T V E G V W T Y K D E T L K F T V T E
BINARY_795 T T Y K L I L N L K Q A K E E A I K E A V D A A I A E K Y I K L I A N A K T V E G V W T Y K D E T L T A T V T E
BINARY_796 T T Y K L I L N L K Q A K E E A I K E A V D A A I A E K Y I K L I A N A K T V E G V W T Y K D E T L T F T V T E
BINARY_797 T T Y K L I L N L K Q A K E E A I K E A V D A A I A E K Y I K L I A N A K T V E G V W T Y K D E T K K A T V T E
BINARY_798 T T Y K L I L N L K Q A K E E A I K E A V D A A I A E K Y I K L I A N A K T V E G V W T Y K D E T K K F T V T E
BINARY_799 T T Y K L I L N L K Q A K E E A I K E A V D A A I A E K Y I K L I A N A K T V E G V W T Y K D E T K T A T V T E
BINARY_800 T T Y K L I L N L K Q A K E E A I K E A V D A A I A E K Y I K L I A N A K T V E G V W T Y K D E T K T F T V T E
BINARY_801 T T Y K L I L N L K Q A K E E A I K E A V D A A I A E K Y I K L Y A N A K T V E G V W T L K D E I L K A T V T E
BINARY_802 T T Y K L I L N L K Q A K E E A I K E A V D A A I A E K Y I K L Y A N A K T V E G V W T L K D E I L K F T V T E
BINARY_803 T T Y K L I L N L K Q A K E E A I K E A V D A A I A E K Y I K L Y A N A K T V E G V W T L K D E I L T A T V T E
BINARY_804 T T Y K L I L N L K Q A K E E A I K E A V D A A I A E K Y I K L Y A N A K T V E G V W T L K D E I L T F T V T E
BINARY_805 T T Y K L I L N L K Q A K E E A I K E A V D A A I A E K Y I K L Y A N A K T V E G V W T L K D E I K K A T V T E
BINARY_806 T T Y K L I L N L K Q A K E E A I K E A V D A A I A E K Y I K L Y A N A K T V E G V W T L K D E I K K F T V T E
BINARY_807 T T Y K L I L N L K Q A K E E A I K E A V D A A I A E K Y I K L Y A N A K T V E G V W T L K D E I K T A T V T E
BINARY_808 T T Y K L I L N L K Q A K E E A I K E A V D A A I A E K Y I K L Y A N A K T V E G V W T L K D E I K T F T V T E
BINARY_809 T T Y K L I L N L K Q A K E E A I K E A V D A A I A E K Y I K L Y A N A K T V E G V W T L K D E T L K A T V T E
BINARY_810 T T Y K L I L N L K Q A K E E A I K E A V D A A I A E K Y I K L Y A N A K T V E G V W T L K D E T L K F T V T E
BINARY_811 T T Y K L I L N L K Q A K E E A I K E A V D A A I A E K Y I K L Y A N A K T V E G V W T L K D E T L T A T V T E
BINARY_812 T T Y K L I L N L K Q A K E E A I K E A V D A A I A E K Y I K L Y A N A K T V E G V W T L K D E T L T F T V T E
BINARY_813 T T Y K L I L N L K Q A K E E A I K E A V D A A I A E K Y I K L Y A N A K T V E G V W T L K D E T K K A T V T E
BINARY_814 T T Y K L I L N L K Q A K E E A I K E A V D A A I A E K Y I K L Y A N A K T V E G V W T L K D E T K K F T V T E
BINARY_815 T T Y K L I L N L K Q A K E E A I K E A V D A A I A E K Y I K L Y A N A K T V E G V W T L K D E T K T A T V T E
BINARY_816 T T Y K L I L N L K Q A K E E A I K E A V D A A I A E K Y I K L Y A N A K T V E G V W T L K D E T K T F T V T E
BINARY_817 T T Y K L I L N L K Q A K E E A I K E A V D A A I A E K Y I K L Y A N A K T V E G V W T Y K D E I L K A T V T E
BINARY_818 T T Y K L I L N L K Q A K E E A I K E A V D A A I A E K Y I K L Y A N A K T V E G V W T Y K D E I L K F T V T E
BINARY_819 T T Y K L I L N L K Q A K E E A I K E A V D A A I A E K Y I K L Y A N A K T V E G V W T Y K D E I L T A T V T E
BINARY_820 T T Y K L I L N L K Q A K E E A I K E A V D A A I A E K Y I K L Y A N A K T V E G V W T Y K D E I L T F T V T E
BINARY_821 T T Y K L I L N L K Q A K E E A I K E A V D A A I A E K Y I K L Y A N A K T V E G V W T Y K D E I K K A T V T E
BINARY_822 T T Y K L I L N L K Q A K E E A I K E A V D A A I A E K Y I K L Y A N A K T V E G V W T Y K D E I K K F T V T E
BINARY_823 T T Y K L I L N L K Q A K E E A I K E A V D A A I A E K Y I K L Y A N A K T V E G V W T Y K D E I K T A T V T E
BINARY_824 T T Y K L I L N L K Q A K E E A I K E A V D A A I A E K Y I K L Y A N A K T V E G V W T Y K D E I K T F T V T E
BINARY_825 T T Y K L I L N L K Q A K E E A I K E A V D A A I A E K Y I K L Y A N A K T V E G V W T Y K D E T L K A T V T E
BINARY_826 T T Y K L I L N L K Q A K E E A I K E A V D A A I A E K Y I K L Y A N A K T V E G V W T Y K D E T L K F T V T E
BINARY_827 T T Y K L I L N L K Q A K E E A I K E A V D A A I A E K Y I K L Y A N A K T V E G V W T Y K D E T L T A T V T E
BINARY_828 T T Y K L I L N L K Q A K E E A I K E A V D A A I A E K Y I K L Y A N A K T V E G V W T Y K D E T L T F T V T E
BINARY_829 T T Y K L I L N L K Q A K E E A I K E A V D A A I A E K Y I K L Y A N A K T V E G V W T Y K D E T K K A T V T E
BINARY_830 T T Y K L I L N L K Q A K E E A I K E A V D A A I A E K Y I K L Y A N A K T V E G V W T Y K D E T K K F T V T E
BINARY_831 T T Y K L I L N L K Q A K E E A I K E A V D A A I A E K Y I K L Y A N A K T V E G V W T Y K D E T K T A T V T E
BINARY_832 T T Y K L I L N L K Q A K E E A I K E A V D A A I A E K Y I K L Y A N A K T V E G V W T Y K D E T K T F T V T E
BINARY_833 T T Y K L I L N L K Q A K E E A I K E A V D A A I A E K Y F K L I A N A K T V E G V W T L K D E I L K A T V T E
BINARY_834 T T Y K L I L N L K Q A K E E A I K E A V D A A I A E K Y F K L I A N A K T V E G V W T L K D E I L K F T V T E
BINARY_835 T T Y K L I L N L K Q A K E E A I K E A V D A A I A E K Y F K L I A N A K T V E G V W T L K D E I L T A T V T E
BINARY_836 T T Y K L I L N L K Q A K E E A I K E A V D A A I A E K Y F K L I A N A K T V E G V W T L K D E I L T F T V T E
BINARY_837 T T Y K L I L N L K Q A K E E A I K E A V D A A I A E K Y F K L I A N A K T V E G V W T L K D E I K K A T V T E
BINARY_838 T T Y K L I L N L K Q A K E E A I K E A V D A A I A E K Y F K L I A N A K T V E G V W T L K D E I K K F T V T E
BINARY_839 T T Y K L I L N L K Q A K E E A I K E A V D A A I A E K Y F K L I A N A K T V E G V W T L K D E I K T A T V T E
BINARY_840 T T Y K L I L N L K Q A K E E A I K E A V D A A I A E K Y F K L I A N A K T V E G V W T L K D E I K T F T V T E
BINARY_841 T T Y K L I L N L K Q A K E E A I K E A V D A A I A E K Y F K L I A N A K T V E G V W T L K D E T L K A T V T E
BINARY_842 T T Y K L I L N L K Q A K E E A I K E A V D A A I A E K Y F K L I A N A K T V E G V W T L K D E T L K F T V T E
BINARY_843 T T Y K L I L N L K Q A K E E A I K E A V D A A I A E K Y F K L I A N A K T V E G V W T L K D E T L T A T V T E
BINARY_844 T T Y K L I L N L K Q A K E E A I K E A V D A A I A E K Y F K L I A N A K T V E G V W T L K D E T L T F T V T E
BINARY_845 T T Y K L I L N L K Q A K E E A I K E A V D A A I A E K Y F K L I A N A K T V E G V W T L K D E T K K A T V T E
BINARY_846 T T Y K L I L N L K Q A K E E A I K E A V D A A I A E K Y F K L I A N A K T V E G V W T L K D E T K K F T V T E
BINARY_847 T T Y K L I L N L K Q A K E E A I K E A V D A A I A E K Y F K L I A N A K T V E G V W T L K D E T K T A T V T E
BINARY_848 T T Y K L I L N L K Q A K E E A I K E A V D A A I A E K Y F K L I A N A K T V E G V W T L K D E T K T F T V T E
BINARY_849 T T Y K L I L N L K Q A K E E A I K E A V D A A I A E K Y F K L I A N A K T V E G V W T Y K D E I L K A T V T E
BINARY_850 T T Y K L I L N L K Q A K E E A I K E A V D A A I A E K Y F K L I A N A K T V E G V W T Y K D E I L K F T V T E
BINARY_851 T T Y K L I L N L K Q A K E E A I K E A V D A A I A E K Y F K L I A N A K T V E G V W T Y K D E I L T A T V T E
BINARY_852 T T Y K L I L N L K Q A K E E A I K E A V D A A I A E K Y F K L I A N A K T V E G V W T Y K D E I L T F T V T E
BINARY_853 T T Y K L I L N L K Q A K E E A I K E A V D A A I A E K Y F K L I A N A K T V E G V W T Y K D E I K K A T V T E
BINARY_854 T T Y K L I L N L K Q A K E E A I K E A V D A A I A E K Y F K L I A N A K T V E G V W T Y K D E I K K F T V T E
BINARY_855 T T Y K L I L N L K Q A K E E A I K E A V D A A I A E K Y F K L I A N A K T V E G V W T Y K D E I K T A T V T E
BINARY_856 T T Y K L I L N L K Q A K E E A I K E A V D A A I A E K Y F K L I A N A K T V E G V W T Y K D E I K T F T V T E
BINARY_857 T T Y K L I L N L K Q A K E E A I K E A V D A A I A E K Y F K L I A N A K T V E G V W T Y K D E T L K A T V T E
BINARY_858 T T Y K L I L N L K Q A K E E A I K E A V D A A I A E K Y F K L I A N A K T V E G V W T Y K D E T L K F T V T E
BINARY_859 T T Y K L I L N L K Q A K E E A I K E A V D A A I A E K Y F K L I A N A K T V E G V W T Y K D E T L T A T V T E
BINARY_860 T T Y K L I L N L K Q A K E E A I K E A V D A A I A E K Y F K L I A N A K T V E G V W T Y K D E T L T F T V T E
BINARY_861 T T Y K L I L N L K Q A K E E A I K E A V D A A I A E K Y F K L I A N A K T V E G V W T Y K D E T K K A T V T E
BINARY_862 T T Y K L I L N L K Q A K E E A I K E A V D A A I A E K Y F K L I A N A K T V E G V W T Y K D E T K K F T V T E
BINARY_863 T T Y K L I L N L K Q A K E E A I K E A V D A A I A E K Y F K L I A N A K T V E G V W T Y K D E T K T A T V T E
BINARY_864 T T Y K L I L N L K Q A K E E A I K E A V D A A I A E K Y F K L I A N A K T V E G V W T Y K D E T K T F T V T E
BINARY_865 T T Y K L I L N L K Q A K E E A I K E A V D A A I A E K Y F K L Y A N A K T V E G V W T L K D E I L K A T V T E
BINARY_866 T T Y K L I L N L K Q A K E E A I K E A V D A A I A E K Y F K L Y A N A K T V E G V W T L K D E I L K F T V T E
BINARY_867 T T Y K L I L N L K Q A K E E A I K E A V D A A I A E K Y F K L Y A N A K T V E G V W T L K D E I L T A T V T E
BINARY_868 T T Y K L I L N L K Q A K E E A I K E A V D A A I A E K Y F K L Y A N A K T V E G V W T L K D E I L T F T V T E
BINARY_869 T T Y K L I L N L K Q A K E E A I K E A V D A A I A E K Y F K L Y A N A K T V E G V W T L K D E I K K A T V T E
BINARY_870 T T Y K L I L N L K Q A K E E A I K E A V D A A I A E K Y F K L Y A N A K T V E G V W T L K D E I K K F T V T E
BINARY_871 T T Y K L I L N L K Q A K E E A I K E A V D A A I A E K Y F K L Y A N A K T V E G V W T L K D E I K T A T V T E

BINARY_872 T T Y K L I L N L K Q A K E E A I K E A V D A A I A E K Y F K L Y A N A K T V E G V W T L K D E I K T F T V T E
BINARY_873 T T Y K L I L N L K Q A K E E A I K E A V D A A I A E K Y F K L Y A N A K T V E G V W T L K D E T L K A T V T E
BINARY_874 T T Y K L I L N L K Q A K E E A I K E A V D A A I A E K Y F K L Y A N A K T V E G V W T L K D E T L K F T V T E
BINARY_875 T T Y K L I L N L K Q A K E E A I K E A V D A A I A E K Y F K L Y A N A K T V E G V W T L K D E T L T A T V T E
BINARY_876 T T Y K L I L N L K Q A K E E A I K E A V D A A I A E K Y F K L Y A N A K T V E G V W T L K D E T L T F T V T E
BINARY_877 T T Y K L I L N L K Q A K E E A I K E A V D A A I A E K Y F K L Y A N A K T V E G V W T L K D E T K K A T V T E
BINARY_878 T T Y K L I L N L K Q A K E E A I K E A V D A A I A E K Y F K L Y A N A K T V E G V W T L K D E T K K F T V T E
BINARY_879 T T Y K L I L N L K Q A K E E A I K E A V D A A I A E K Y F K L Y A N A K T V E G V W T L K D E T K T A T V T E
BINARY_880 T T Y K L I L N L K Q A K E E A I K E A V D A A I A E K Y F K L Y A N A K T V E G V W T L K D E T K T F T V T E
BINARY_881 T T Y K L I L N L K Q A K E E A I K E A V D A A I A E K Y F K L Y A N A K T V E G V W T Y K D E I L K A T V T E
BINARY_882 T T Y K L I L N L K Q A K E E A I K E A V D A A I A E K Y F K L Y A N A K T V E G V W T Y K D E I L K F T V T E
BINARY_883 T T Y K L I L N L K Q A K E E A I K E A V D A A I A E K Y F K L Y A N A K T V E G V W T Y K D E I L T A T V T E
BINARY_884 T T Y K L I L N L K Q A K E E A I K E A V D A A I A E K Y F K L Y A N A K T V E G V W T Y K D E I L T F T V T E
BINARY_885 T T Y K L I L N L K Q A K E E A I K E A V D A A I A E K Y F K L Y A N A K T V E G V W T Y K D E I K K A T V T E
BINARY_886 T T Y K L I L N L K Q A K E E A I K E A V D A A I A E K Y F K L Y A N A K T V E G V W T Y K D E I K K F T V T E
BINARY_887 T T Y K L I L N L K Q A K E E A I K E A V D A A I A E K Y F K L Y A N A K T V E G V W T Y K D E I K T A T V T E
BINARY_888 T T Y K L I L N L K Q A K E E A I K E A V D A A I A E K Y F K L Y A N A K T V E G V W T Y K D E I K T F T V T E
BINARY_889 T T Y K L I L N L K Q A K E E A I K E A V D A A I A E K Y F K L Y A N A K T V E G V W T Y K D E T L K A T V T E
BINARY_890 T T Y K L I L N L K Q A K E E A I K E A V D A A I A E K Y F K L Y A N A K T V E G V W T Y K D E T L K F T V T E
BINARY_891 T T Y K L I L N L K Q A K E E A I K E A V D A A I A E K Y F K L Y A N A K T V E G V W T Y K D E T L T A T V T E
BINARY_892 T T Y K L I L N L K Q A K E E A I K E A V D A A I A E K Y F K L Y A N A K T V E G V W T Y K D E T L T F T V T E
BINARY_893 T T Y K L I L N L K Q A K E E A I K E A V D A A I A E K Y F K L Y A N A K T V E G V W T Y K D E T K K A T V T E
BINARY_894 T T Y K L I L N L K Q A K E E A I K E A V D A A I A E K Y F K L Y A N A K T V E G V W T Y K D E T K K F T V T E
BINARY_895 T T Y K L I L N L K Q A K E E A I K E A V D A A I A E K Y F K L Y A N A K T V E G V W T Y K D E T K T A T V T E
BINARY_896 T T Y K L I L N L K Q A K E E A I K E A V D A A I A E K Y F K L Y A N A K T V E G V W T Y K D E T K T F T V T E
BINARY_897 T T Y K L I L N L K Q A K E E A I K E A V D A A T A E K Y I K L I A N A K T V E G V W T L K D E I L K A T V T E
BINARY_898 T T Y K L I L N L K Q A K E E A I K E A V D A A T A E K Y I K L I A N A K T V E G V W T L K D E I L K F T V T E
BINARY_899 T T Y K L I L N L K Q A K E E A I K E A V D A A T A E K Y I K L I A N A K T V E G V W T L K D E I L T A T V T E
BINARY_900 T T Y K L I L N L K Q A K E E A I K E A V D A A T A E K Y I K L I A N A K T V E G V W T L K D E I L T F T V T E
BINARY_901 T T Y K L I L N L K Q A K E E A I K E A V D A A T A E K Y I K L I A N A K T V E G V W T L K D E I K K A T V T E
BINARY_902 T T Y K L I L N L K Q A K E E A I K E A V D A A T A E K Y I K L I A N A K T V E G V W T L K D E I K K F T V T E
BINARY_903 T T Y K L I L N L K Q A K E E A I K E A V D A A T A E K Y I K L I A N A K T V E G V W T L K D E I K T A T V T E
BINARY_904 T T Y K L I L N L K Q A K E E A I K E A V D A A T A E K Y I K L I A N A K T V E G V W T L K D E I K T F T V T E
BINARY_905 T T Y K L I L N L K Q A K E E A I K E A V D A A T A E K Y I K L I A N A K T V E G V W T L K D E T L K A T V T E
BINARY_906 T T Y K L I L N L K Q A K E E A I K E A V D A A T A E K Y I K L I A N A K T V E G V W T L K D E T L K F T V T E
BINARY_907 T T Y K L I L N L K Q A K E E A I K E A V D A A T A E K Y I K L I A N A K T V E G V W T L K D E T L T A T V T E
BINARY_908 T T Y K L I L N L K Q A K E E A I K E A V D A A T A E K Y I K L I A N A K T V E G V W T L K D E T L T F T V T E
BINARY_909 T T Y K L I L N L K Q A K E E A I K E A V D A A T A E K Y I K L I A N A K T V E G V W T L K D E T K K A T V T E
BINARY_910 T T Y K L I L N L K Q A K E E A I K E A V D A A T A E K Y I K L I A N A K T V E G V W T L K D E T K K F T V T E
BINARY_911 T T Y K L I L N L K Q A K E E A I K E A V D A A T A E K Y I K L I A N A K T V E G V W T L K D E T K T A T V T E
BINARY_912 T T Y K L I L N L K Q A K E E A I K E A V D A A T A E K Y I K L I A N A K T V E G V W T L K D E T K T F T V T E
BINARY_913 T T Y K L I L N L K Q A K E E A I K E A V D A A T A E K Y I K L I A N A K T V E G V W T Y K D E I L K A T V T E
BINARY_914 T T Y K L I L N L K Q A K E E A I K E A V D A A T A E K Y I K L I A N A K T V E G V W T Y K D E I L K F T V T E
BINARY_915 T T Y K L I L N L K Q A K E E A I K E A V D A A T A E K Y I K L I A N A K T V E G V W T Y K D E I L T A T V T E
BINARY_916 T T Y K L I L N L K Q A K E E A I K E A V D A A T A E K Y I K L I A N A K T V E G V W T Y K D E I L T F T V T E
BINARY_917 T T Y K L I L N L K Q A K E E A I K E A V D A A T A E K Y I K L I A N A K T V E G V W T Y K D E I K K A T V T E
BINARY_918 T T Y K L I L N L K Q A K E E A I K E A V D A A T A E K Y I K L I A N A K T V E G V W T Y K D E I K K F T V T E
BINARY_919 T T Y K L I L N L K Q A K E E A I K E A V D A A T A E K Y I K L I A N A K T V E G V W T Y K D E I K T A T V T E
BINARY_920 T T Y K L I L N L K Q A K E E A I K E A V D A A T A E K Y I K L I A N A K T V E G V W T Y K D E I K T F T V T E
BINARY_921 T T Y K L I L N L K Q A K E E A I K E A V D A A T A E K Y I K L I A N A K T V E G V W T Y K D E T L K A T V T E
BINARY_922 T T Y K L I L N L K Q A K E E A I K E A V D A A T A E K Y I K L I A N A K T V E G V W T Y K D E T L K F T V T E
BINARY_923 T T Y K L I L N L K Q A K E E A I K E A V D A A T A E K Y I K L I A N A K T V E G V W T Y K D E T L T A T V T E
BINARY_924 T T Y K L I L N L K Q A K E E A I K E A V D A A T A E K Y I K L I A N A K T V E G V W T Y K D E T L T F T V T E
BINARY_925 T T Y K L I L N L K Q A K E E A I K E A V D A A T A E K Y I K L I A N A K T V E G V W T Y K D E T K K A T V T E
BINARY_926 T T Y K L I L N L K Q A K E E A I K E A V D A A T A E K Y I K L I A N A K T V E G V W T Y K D E T K K F T V T E
BINARY_927 T T Y K L I L N L K Q A K E E A I K E A V D A A T A E K Y I K L I A N A K T V E G V W T Y K D E T K T A T V T E
BINARY_928 T T Y K L I L N L K Q A K E E A I K E A V D A A T A E K Y I K L I A N A K T V E G V W T Y K D E T K T F T V T E
BINARY_929 T T Y K L I L N L K Q A K E E A I K E A V D A A T A E K Y I K L Y A N A K T V E G V W T L K D E I L K A T V T E
BINARY_930 T T Y K L I L N L K Q A K E E A I K E A V D A A T A E K Y I K L Y A N A K T V E G V W T L K D E I L K F T V T E
BINARY_931 T T Y K L I L N L K Q A K E E A I K E A V D A A T A E K Y I K L Y A N A K T V E G V W T L K D E I L T A T V T E
BINARY_932 T T Y K L I L N L K Q A K E E A I K E A V D A A T A E K Y I K L Y A N A K T V E G V W T L K D E I L T F T V T E
BINARY_933 T T Y K L I L N L K Q A K E E A I K E A V D A A T A E K Y I K L Y A N A K T V E G V W T L K D E I K K A T V T E
BINARY_934 T T Y K L I L N L K Q A K E E A I K E A V D A A T A E K Y I K L Y A N A K T V E G V W T L K D E I K K F T V T E
BINARY_935 T T Y K L I L N L K Q A K E E A I K E A V D A A T A E K Y I K L Y A N A K T V E G V W T L K D E I K T A T V T E
BINARY_936 T T Y K L I L N L K Q A K E E A I K E A V D A A T A E K Y I K L Y A N A K T V E G V W T L K D E I K T F T V T E
BINARY_937 T T Y K L I L N L K Q A K E E A I K E A V D A A T A E K Y I K L Y A N A K T V E G V W T L K D E T L K A T V T E
BINARY_938 T T Y K L I L N L K Q A K E E A I K E A V D A A T A E K Y I K L Y A N A K T V E G V W T L K D E T L K F T V T E
BINARY_939 T T Y K L I L N L K Q A K E E A I K E A V D A A T A E K Y I K L Y A N A K T V E G V W T L K D E T L T A T V T E
BINARY_940 T T Y K L I L N L K Q A K E E A I K E A V D A A T A E K Y I K L Y A N A K T V E G V W T L K D E T L T F T V T E
BINARY_941 T T Y K L I L N L K Q A K E E A I K E A V D A A T A E K Y I K L Y A N A K T V E G V W T L K D E T K K A T V T E
BINARY_942 T T Y K L I L N L K Q A K E E A I K E A V D A A T A E K Y I K L Y A N A K T V E G V W T L K D E T K K F T V T E
BINARY_943 T T Y K L I L N L K Q A K E E A I K E A V D A A T A E K Y I K L Y A N A K T V E G V W T L K D E T K T A T V T E
BINARY_944 T T Y K L I L N L K Q A K E E A I K E A V D A A T A E K Y I K L Y A N A K T V E G V W T L K D E T K T F T V T E
BINARY_945 T T Y K L I L N L K Q A K E E A I K E A V D A A T A E K Y I K L Y A N A K T V E G V W T Y K D E I L K A T V T E
BINARY_946 T T Y K L I L N L K Q A K E E A I K E A V D A A T A E K Y I K L Y A N A K T V E G V W T Y K D E I L K F T V T E
BINARY_947 T T Y K L I L N L K Q A K E E A I K E A V D A A T A E K Y I K L Y A N A K T V E G V W T Y K D E I L T A T V T E
BINARY_948 T T Y K L I L N L K Q A K E E A I K E A V D A A T A E K Y I K L Y A N A K T V E G V W T Y K D E I L T F T V T E
BINARY_949 T T Y K L I L N L K Q A K E E A I K E A V D A A T A E K Y I K L Y A N A K T V E G V W T Y K D E I K K A T V T E
BINARY_950 T T Y K L I L N L K Q A K E E A I K E A V D A A T A E K Y I K L Y A N A K T V E G V W T Y K D E I K K F T V T E
BINARY_951 T T Y K L I L N L K Q A K E E A I K E A V D A A T A E K Y I K L Y A N A K T V E G V W T Y K D E I K T A T V T E
BINARY_952 T T Y K L I L N L K Q A K E E A I K E A V D A A T A E K Y I K L Y A N A K T V E G V W T Y K D E I K T F T V T E
BINARY_953 T T Y K L I L N L K Q A K E E A I K E A V D A A T A E K Y I K L Y A N A K T V E G V W T Y K D E T L K A T V T E
BINARY_954 T T Y K L I L N L K Q A K E E A I K E A V D A A T A E K Y I K L Y A N A K T V E G V W T Y K D E T L K F T V T E
BINARY_955 T T Y K L I L N L K Q A K E E A I K E A V D A A T A E K Y I K L Y A N A K T V E G V W T Y K D E T L T A T V T E
BINARY_956 T T Y K L I L N L K Q A K E E A I K E A V D A A T A E K Y I K L Y A N A K T V E G V W T Y K D E T L T F T V T E
BINARY_957 T T Y K L I L N L K Q A K E E A I K E A V D A A T A E K Y I K L Y A N A K T V E G V W T Y K D E T K K A T V T E
BINARY_958 T T Y K L I L N L K Q A K E E A I K E A V D A A T A E K Y I K L Y A N A K T V E G V W T Y K D E T K K F T V T E
BINARY_959 T T Y K L I L N L K Q A K E E A I K E A V D A A T A E K Y I K L Y A N A K T V E G V W T Y K D E T K T A T V T E
BINARY_960 T T Y K L I L N L K Q A K E E A I K E A V D A A T A E K Y I K L Y A N A K T V E G V W T Y K D E T K T F T V T E
BINARY_961 T T Y K L I L N L K Q A K E E A I K E A V D A A T A E K Y F K L I A N A K T V E G V W T L K D E I L K A T V T E
BINARY_962 T T Y K L I L N L K Q A K E E A I K E A V D A A T A E K Y F K L I A N A K T V E G V W T L K D E I L K F T V T E
BINARY_963 T T Y K L I L N L K Q A K E E A I K E A V D A A T A E K Y F K L I A N A K T V E G V W T L K D E I L T A T V T E
BINARY_964 T T Y K L I L N L K Q A K E E A I K E A V D A A T A E K Y F K L I A N A K T V E G V W T L K D E I L T F T V T E
BINARY_965 T T Y K L I L N L K Q A K E E A I K E A V D A A T A E K Y F K L I A N A K T V E G V W T L K D E I K K A T V T E
BINARY_966 T T Y K L I L N L K Q A K E E A I K E A V D A A T A E K Y F K L I A N A K T V E G V W T L K D E I K K F T V T E

BINARY_967 T T Y K L I L N L K Q A K E E A I K E A V D A A T A E K Y F K L I A N A K T V E G V W T L K D E I K T A T V T E
BINARY_968 T T Y K L I L N L K Q A K E E A I K E A V D A A T A E K Y F K L I A N A K T V E G V W T L K D E I K T F T V T E
BINARY_969 T T Y K L I L N L K Q A K E E A I K E A V D A A T A E K Y F K L I A N A K T V E G V W T L K D E T L K A T V T E
BINARY_970 T T Y K L I L N L K Q A K E E A I K E A V D A A T A E K Y F K L I A N A K T V E G V W T L K D E T L K F T V T E
BINARY_971 T T Y K L I L N L K Q A K E E A I K E A V D A A T A E K Y F K L I A N A K T V E G V W T L K D E T L T A T V T E
BINARY_972 T T Y K L I L N L K Q A K E E A I K E A V D A A T A E K Y F K L I A N A K T V E G V W T L K D E T L T F T V T E
BINARY_973 T T Y K L I L N L K Q A K E E A I K E A V D A A T A E K Y F K L I A N A K T V E G V W T L K D E T K K A T V T E
BINARY_974 T T Y K L I L N L K Q A K E E A I K E A V D A A T A E K Y F K L I A N A K T V E G V W T L K D E T K K F T V T E
BINARY_975 T T Y K L I L N L K Q A K E E A I K E A V D A A T A E K Y F K L I A N A K T V E G V W T L K D E T K T A T V T E
BINARY_976 T T Y K L I L N L K Q A K E E A I K E A V D A A T A E K Y F K L I A N A K T V E G V W T L K D E T K T F T V T E
BINARY_977 T T Y K L I L N L K Q A K E E A I K E A V D A A T A E K Y F K L I A N A K T V E G V W T Y K D E I L K A T V T E
BINARY_978 T T Y K L I L N L K Q A K E E A I K E A V D A A T A E K Y F K L I A N A K T V E G V W T Y K D E I L K F T V T E
BINARY_979 T T Y K L I L N L K Q A K E E A I K E A V D A A T A E K Y F K L I A N A K T V E G V W T Y K D E I L T A T V T E
BINARY_980 T T Y K L I L N L K Q A K E E A I K E A V D A A T A E K Y F K L I A N A K T V E G V W T Y K D E I L T F T V T E
BINARY_981 T T Y K L I L N L K Q A K E E A I K E A V D A A T A E K Y F K L I A N A K T V E G V W T Y K D E I K K A T V T E
BINARY_982 T T Y K L I L N L K Q A K E E A I K E A V D A A T A E K Y F K L I A N A K T V E G V W T Y K D E I K K F T V T E
BINARY_983 T T Y K L I L N L K Q A K E E A I K E A V D A A T A E K Y F K L I A N A K T V E G V W T Y K D E I K T A T V T E
BINARY_984 T T Y K L I L N L K Q A K E E A I K E A V D A A T A E K Y F K L I A N A K T V E G V W T Y K D E I K T F T V T E
BINARY_985 T T Y K L I L N L K Q A K E E A I K E A V D A A T A E K Y F K L I A N A K T V E G V W T Y K D E T L K A T V T E
BINARY_986 T T Y K L I L N L K Q A K E E A I K E A V D A A T A E K Y F K L I A N A K T V E G V W T Y K D E T L K F T V T E
BINARY_987 T T Y K L I L N L K Q A K E E A I K E A V D A A T A E K Y F K L I A N A K T V E G V W T Y K D E T L T A T V T E
BINARY_988 T T Y K L I L N L K Q A K E E A I K E A V D A A T A E K Y F K L I A N A K T V E G V W T Y K D E T L T F T V T E
BINARY_989 T T Y K L I L N L K Q A K E E A I K E A V D A A T A E K Y F K L I A N A K T V E G V W T Y K D E T K K A T V T E
BINARY_990 T T Y K L I L N L K Q A K E E A I K E A V D A A T A E K Y F K L I A N A K T V E G V W T Y K D E T K K F T V T E
BINARY_991 T T Y K L I L N L K Q A K E E A I K E A V D A A T A E K Y F K L I A N A K T V E G V W T Y K D E T K T A T V T E
BINARY_992 T T Y K L I L N L K Q A K E E A I K E A V D A A T A E K Y F K L I A N A K T V E G V W T Y K D E T K T F T V T E
BINARY_993 T T Y K L I L N L K Q A K E E A I K E A V D A A T A E K Y F K L Y A N A K T V E G V W T L K D E I L K A T V T E
BINARY_994 T T Y K L I L N L K Q A K E E A I K E A V D A A T A E K Y F K L Y A N A K T V E G V W T L K D E I L K F T V T E
BINARY_995 T T Y K L I L N L K Q A K E E A I K E A V D A A T A E K Y F K L Y A N A K T V E G V W T L K D E I L T A T V T E
BINARY_996 T T Y K L I L N L K Q A K E E A I K E A V D A A T A E K Y F K L Y A N A K T V E G V W T L K D E I L T F T V T E
BINARY_997 T T Y K L I L N L K Q A K E E A I K E A V D A A T A E K Y F K L Y A N A K T V E G V W T L K D E I K K A T V T E
BINARY_998 T T Y K L I L N L K Q A K E E A I K E A V D A A T A E K Y F K L Y A N A K T V E G V W T L K D E I K K F T V T E
BINARY_999 T T Y K L I L N L K Q A K E E A I K E A V D A A T A E K Y F K L Y A N A K T V E G V W T L K D E I K T A T V T E
BINARY_1000 T T Y K L I L N L K Q A K E E A I K E A V D A A T A E K Y F K L Y A N A K T V E G V W T L K D E I K T F T V T E
BINARY_1001 T T Y K L I L N L K Q A K E E A I K E A V D A A T A E K Y F K L Y A N A K T V E G V W T L K D E T L K A T V T E
BINARY_1002 T T Y K L I L N L K Q A K E E A I K E A V D A A T A E K Y F K L Y A N A K T V E G V W T L K D E T L K F T V T E
BINARY_1003 T T Y K L I L N L K Q A K E E A I K E A V D A A T A E K Y F K L Y A N A K T V E G V W T L K D E T L T A T V T E
BINARY_1004 T T Y K L I L N L K Q A K E E A I K E A V D A A T A E K Y F K L Y A N A K T V E G V W T L K D E T L T F T V T E
BINARY_1005 T T Y K L I L N L K Q A K E E A I K E A V D A A T A E K Y F K L Y A N A K T V E G V W T L K D E T K K A T V T E
BINARY_1006 T T Y K L I L N L K Q A K E E A I K E A V D A A T A E K Y F K L Y A N A K T V E G V W T L K D E T K K F T V T E
BINARY_1007 T T Y K L I L N L K Q A K E E A I K E A V D A A T A E K Y F K L Y A N A K T V E G V W T L K D E T K T A T V T E
BINARY_1008 T T Y K L I L N L K Q A K E E A I K E A V D A A T A E K Y F K L Y A N A K T V E G V W T L K D E T K T F T V T E
BINARY_1009 T T Y K L I L N L K Q A K E E A I K E A V D A A T A E K Y F K L Y A N A K T V E G V W T Y K D E I L K A T V T E
BINARY_1010 T T Y K L I L N L K Q A K E E A I K E A V D A A T A E K Y F K L Y A N A K T V E G V W T Y K D E I L K F T V T E
BINARY_1011 T T Y K L I L N L K Q A K E E A I K E A V D A A T A E K Y F K L Y A N A K T V E G V W T Y K D E I L T A T V T E
BINARY_1012 T T Y K L I L N L K Q A K E E A I K E A V D A A T A E K Y F K L Y A N A K T V E G V W T Y K D E I L T F T V T E
BINARY_1013 T T Y K L I L N L K Q A K E E A I K E A V D A A T A E K Y F K L Y A N A K T V E G V W T Y K D E I K K A T V T E
BINARY_1014 T T Y K L I L N L K Q A K E E A I K E A V D A A T A E K Y F K L Y A N A K T V E G V W T Y K D E I K K F T V T E
BINARY_1015 T T Y K L I L N L K Q A K E E A I K E A V D A A T A E K Y F K L Y A N A K T V E G V W T Y K D E I K T A T V T E
BINARY_1016 T T Y K L I L N L K Q A K E E A I K E A V D A A T A E K Y F K L Y A N A K T V E G V W T Y K D E I K T F T V T E
BINARY_1017 T T Y K L I L N L K Q A K E E A I K E A V D A A T A E K Y F K L Y A N A K T V E G V W T Y K D E T L K A T V T E
BINARY_1018 T T Y K L I L N L K Q A K E E A I K E A V D A A T A E K Y F K L Y A N A K T V E G V W T Y K D E T L K F T V T E
BINARY_1019 T T Y K L I L N L K Q A K E E A I K E A V D A A T A E K Y F K L Y A N A K T V E G V W T Y K D E T L T A T V T E
BINARY_1020 T T Y K L I L N L K Q A K E E A I K E A V D A A T A E K Y F K L Y A N A K T V E G V W T Y K D E T L T F T V T E
BINARY_1021 T T Y K L I L N L K Q A K E E A I K E A V D A A T A E K Y F K L Y A N A K T V E G V W T Y K D E T K K A T V T E
BINARY_1022 T T Y K L I L N L K Q A K E E A I K E A V D A A T A E K Y F K L Y A N A K T V E G V W T Y K D E T K K F T V T E
BINARY_1023 T T Y K L I L N L K Q A K E E A I K E A V D A A T A E K Y F K L Y A N A K T V E G V W T Y K D E T K T A T V T E
BINARY_1024 T T Y K L I L N L K Q A K E E A I K E A V D A A T A E K Y F K L Y A N A K T V E G V W T Y K D E T K T F T V T E
BINARY_1025 T T Y K L I L N L K Q A K E E A I T E L V D A G I A E K Y I K L I A N A K T V E G V W T L K D E I L K A T V T E
BINARY_1026 T T Y K L I L N L K Q A K E E A I T E L V D A G I A E K Y I K L I A N A K T V E G V W T L K D E I L K F T V T E
BINARY_1027 T T Y K L I L N L K Q A K E E A I T E L V D A G I A E K Y I K L I A N A K T V E G V W T L K D E I L T A T V T E
BINARY_1028 T T Y K L I L N L K Q A K E E A I T E L V D A G I A E K Y I K L I A N A K T V E G V W T L K D E I L T F T V T E
BINARY_1029 T T Y K L I L N L K Q A K E E A I T E L V D A G I A E K Y I K L I A N A K T V E G V W T L K D E I K K A T V T E
BINARY_1030 T T Y K L I L N L K Q A K E E A I T E L V D A G I A E K Y I K L I A N A K T V E G V W T L K D E I K K F T V T E
BINARY_1031 T T Y K L I L N L K Q A K E E A I T E L V D A G I A E K Y I K L I A N A K T V E G V W T L K D E I K T A T V T E
BINARY_1032 T T Y K L I L N L K Q A K E E A I T E L V D A G I A E K Y I K L I A N A K T V E G V W T L K D E I K T F T V T E
BINARY_1033 T T Y K L I L N L K Q A K E E A I T E L V D A G I A E K Y I K L I A N A K T V E G V W T L K D E T L K A T V T E
BINARY_1034 T T Y K L I L N L K Q A K E E A I T E L V D A G I A E K Y I K L I A N A K T V E G V W T L K D E T L K F T V T E
BINARY_1035 T T Y K L I L N L K Q A K E E A I T E L V D A G I A E K Y I K L I A N A K T V E G V W T L K D E T L T A T V T E
BINARY_1036 T T Y K L I L N L K Q A K E E A I T E L V D A G I A E K Y I K L I A N A K T V E G V W T L K D E T L T F T V T E
BINARY_1037 T T Y K L I L N L K Q A K E E A I T E L V D A G I A E K Y I K L I A N A K T V E G V W T L K D E T K K A T V T E
BINARY_1038 T T Y K L I L N L K Q A K E E A I T E L V D A G I A E K Y I K L I A N A K T V E G V W T L K D E T K K F T V T E
BINARY_1039 T T Y K L I L N L K Q A K E E A I T E L V D A G I A E K Y I K L I A N A K T V E G V W T L K D E T K T A T V T E
BINARY_1040 T T Y K L I L N L K Q A K E E A I T E L V D A G I A E K Y I K L I A N A K T V E G V W T L K D E T K T F T V T E
BINARY_1041 T T Y K L I L N L K Q A K E E A I T E L V D A G I A E K Y I K L I A N A K T V E G V W T Y K D E I L K A T V T E
BINARY_1042 T T Y K L I L N L K Q A K E E A I T E L V D A G I A E K Y I K L I A N A K T V E G V W T Y K D E I L K F T V T E
BINARY_1043 T T Y K L I L N L K Q A K E E A I T E L V D A G I A E K Y I K L I A N A K T V E G V W T Y K D E I L T A T V T E
BINARY_1044 T T Y K L I L N L K Q A K E E A I T E L V D A G I A E K Y I K L I A N A K T V E G V W T Y K D E I L T F T V T E
BINARY_1045 T T Y K L I L N L K Q A K E E A I T E L V D A G I A E K Y I K L I A N A K T V E G V W T Y K D E I K K A T V T E
BINARY_1046 T T Y K L I L N L K Q A K E E A I T E L V D A G I A E K Y I K L I A N A K T V E G V W T Y K D E I K K F T V T E
BINARY_1047 T T Y K L I L N L K Q A K E E A I T E L V D A G I A E K Y I K L I A N A K T V E G V W T Y K D E I K T A T V T E
BINARY_1048 T T Y K L I L N L K Q A K E E A I T E L V D A G I A E K Y I K L I A N A K T V E G V W T Y K D E I K T F T V T E
BINARY_1049 T T Y K L I L N L K Q A K E E A I T E L V D A G I A E K Y I K L I A N A K T V E G V W T Y K D E T L K A T V T E
BINARY_1050 T T Y K L I L N L K Q A K E E A I T E L V D A G I A E K Y I K L I A N A K T V E G V W T Y K D E T L K F T V T E
BINARY_1051 T T Y K L I L N L K Q A K E E A I T E L V D A G I A E K Y I K L I A N A K T V E G V W T Y K D E T L T A T V T E
BINARY_1052 T T Y K L I L N L K Q A K E E A I T E L V D A G I A E K Y I K L I A N A K T V E G V W T Y K D E T L T F T V T E
BINARY_1053 T T Y K L I L N L K Q A K E E A I T E L V D A G I A E K Y I K L I A N A K T V E G V W T Y K D E T K K A T V T E
BINARY_1054 T T Y K L I L N L K Q A K E E A I T E L V D A G I A E K Y I K L I A N A K T V E G V W T Y K D E T K K F T V T E
BINARY_1055 T T Y K L I L N L K Q A K E E A I T E L V D A G I A E K Y I K L I A N A K T V E G V W T Y K D E T K T A T V T E
BINARY_1056 T T Y K L I L N L K Q A K E E A I T E L V D A G I A E K Y I K L I A N A K T V E G V W T Y K D E T K T F T V T E
BINARY_1057 T T Y K L I L N L K Q A K E E A I T E L V D A G I A E K Y I K L Y A N A K T V E G V W T L K D E I L K A T V T E
BINARY_1058 T T Y K L I L N L K Q A K E E A I T E L V D A G I A E K Y I K L Y A N A K T V E G V W T L K D E I L K F T V T E
BINARY_1059 T T Y K L I L N L K Q A K E E A I T E L V D A G I A E K Y I K L Y A N A K T V E G V W T L K D E I L T A T V T E
BINARY_1060 T T Y K L I L N L K Q A K E E A I T E L V D A G I A E K Y I K L Y A N A K T V E G V W T L K D E I L T F T V T E
BINARY_1061 T T Y K L I L N L K Q A K E E A I T E L V D A G I A E K Y I K L Y A N A K T V E G V W T L K D E I K K A T V T E

BINARY_1062 T T Y K L I L N L K Q A K E E A I T E L V D A G I A E K Y I K L Y A N A K T V E G V W T L K D E I K K F T V T E
BINARY_1063 T T Y K L I L N L K Q A K E E A I T E L V D A G I A E K Y I K L Y A N A K T V E G V W T L K D E I K T A T V T E
BINARY_1064 T T Y K L I L N L K Q A K E E A I T E L V D A G I A E K Y I K L Y A N A K T V E G V W T L K D E I K T F T V T E
BINARY_1065 T T Y K L I L N L K Q A K E E A I T E L V D A G I A E K Y I K L Y A N A K T V E G V W T L K D E T L K A T V T E
BINARY_1066 T T Y K L I L N L K Q A K E E A I T E L V D A G I A E K Y I K L Y A N A K T V E G V W T L K D E T L K F T V T E
BINARY_1067 T T Y K L I L N L K Q A K E E A I T E L V D A G I A E K Y I K L Y A N A K T V E G V W T L K D E T L T A T V T E
BINARY_1068 T T Y K L I L N L K Q A K E E A I T E L V D A G I A E K Y I K L Y A N A K T V E G V W T L K D E T L T F T V T E
BINARY_1069 T T Y K L I L N L K Q A K E E A I T E L V D A G I A E K Y I K L Y A N A K T V E G V W T L K D E T K K A T V T E
BINARY_1070 T T Y K L I L N L K Q A K E E A I T E L V D A G I A E K Y I K L Y A N A K T V E G V W T L K D E T K K F T V T E
BINARY_1071 T T Y K L I L N L K Q A K E E A I T E L V D A G I A E K Y I K L Y A N A K T V E G V W T L K D E T K T A T V T E
BINARY_1072 T T Y K L I L N L K Q A K E E A I T E L V D A G I A E K Y I K L Y A N A K T V E G V W T L K D E T K T F T V T E
BINARY_1073 T T Y K L I L N L K Q A K E E A I T E L V D A G I A E K Y I K L Y A N A K T V E G V W T Y K D E I L K A T V T E
BINARY_1074 T T Y K L I L N L K Q A K E E A I T E L V D A G I A E K Y I K L Y A N A K T V E G V W T Y K D E I L K F T V T E
BINARY_1075 T T Y K L I L N L K Q A K E E A I T E L V D A G I A E K Y I K L Y A N A K T V E G V W T Y K D E I L T A T V T E
BINARY_1076 T T Y K L I L N L K Q A K E E A I T E L V D A G I A E K Y I K L Y A N A K T V E G V W T Y K D E I L T F T V T E
BINARY_1077 T T Y K L I L N L K Q A K E E A I T E L V D A G I A E K Y I K L Y A N A K T V E G V W T Y K D E I K K A T V T E
BINARY_1078 T T Y K L I L N L K Q A K E E A I T E L V D A G I A E K Y I K L Y A N A K T V E G V W T Y K D E I K K F T V T E
BINARY_1079 T T Y K L I L N L K Q A K E E A I T E L V D A G I A E K Y I K L Y A N A K T V E G V W T Y K D E I K T A T V T E
BINARY_1080 T T Y K L I L N L K Q A K E E A I T E L V D A G I A E K Y I K L Y A N A K T V E G V W T Y K D E I K T F T V T E
BINARY_1081 T T Y K L I L N L K Q A K E E A I T E L V D A G I A E K Y I K L Y A N A K T V E G V W T Y K D E T L K A T V T E
BINARY_1082 T T Y K L I L N L K Q A K E E A I T E L V D A G I A E K Y I K L Y A N A K T V E G V W T Y K D E T L K F T V T E
BINARY_1083 T T Y K L I L N L K Q A K E E A I T E L V D A G I A E K Y I K L Y A N A K T V E G V W T Y K D E T L T A T V T E
BINARY_1084 T T Y K L I L N L K Q A K E E A I T E L V D A G I A E K Y I K L Y A N A K T V E G V W T Y K D E T L T F T V T E
BINARY_1085 T T Y K L I L N L K Q A K E E A I T E L V D A G I A E K Y I K L Y A N A K T V E G V W T Y K D E T K K A T V T E
BINARY_1086 T T Y K L I L N L K Q A K E E A I T E L V D A G I A E K Y I K L Y A N A K T V E G V W T Y K D E T K K F T V T E
BINARY_1087 T T Y K L I L N L K Q A K E E A I T E L V D A G I A E K Y I K L Y A N A K T V E G V W T Y K D E T K T A T V T E
BINARY_1088 T T Y K L I L N L K Q A K E E A I T E L V D A G I A E K Y I K L Y A N A K T V E G V W T Y K D E T K T F T V T E
BINARY_1089 T T Y K L I L N L K Q A K E E A I T E L V D A G I A E K Y F K L I A N A K T V E G V W T L K D E I L K A T V T E
BINARY_1090 T T Y K L I L N L K Q A K E E A I T E L V D A G I A E K Y F K L I A N A K T V E G V W T L K D E I L K F T V T E
BINARY_1091 T T Y K L I L N L K Q A K E E A I T E L V D A G I A E K Y F K L I A N A K T V E G V W T L K D E I L T A T V T E
BINARY_1092 T T Y K L I L N L K Q A K E E A I T E L V D A G I A E K Y F K L I A N A K T V E G V W T L K D E I L T F T V T E
BINARY_1093 T T Y K L I L N L K Q A K E E A I T E L V D A G I A E K Y F K L I A N A K T V E G V W T L K D E I K K A T V T E
BINARY_1094 T T Y K L I L N L K Q A K E E A I T E L V D A G I A E K Y F K L I A N A K T V E G V W T L K D E I K K F T V T E
BINARY_1095 T T Y K L I L N L K Q A K E E A I T E L V D A G I A E K Y F K L I A N A K T V E G V W T L K D E I K T A T V T E
BINARY_1096 T T Y K L I L N L K Q A K E E A I T E L V D A G I A E K Y F K L I A N A K T V E G V W T L K D E I K T F T V T E
BINARY_1097 T T Y K L I L N L K Q A K E E A I T E L V D A G I A E K Y F K L I A N A K T V E G V W T L K D E T L K A T V T E
BINARY_1098 T T Y K L I L N L K Q A K E E A I T E L V D A G I A E K Y F K L I A N A K T V E G V W T L K D E T L K F T V T E
BINARY_1099 T T Y K L I L N L K Q A K E E A I T E L V D A G I A E K Y F K L I A N A K T V E G V W T L K D E T L T A T V T E
BINARY_1100 T T Y K L I L N L K Q A K E E A I T E L V D A G I A E K Y F K L I A N A K T V E G V W T L K D E T L T F T V T E
BINARY_1101 T T Y K L I L N L K Q A K E E A I T E L V D A G I A E K Y F K L I A N A K T V E G V W T L K D E T K K A T V T E
BINARY_1102 T T Y K L I L N L K Q A K E E A I T E L V D A G I A E K Y F K L I A N A K T V E G V W T L K D E T K K F T V T E
BINARY_1103 T T Y K L I L N L K Q A K E E A I T E L V D A G I A E K Y F K L I A N A K T V E G V W T L K D E T K T A T V T E
BINARY_1104 T T Y K L I L N L K Q A K E E A I T E L V D A G I A E K Y F K L I A N A K T V E G V W T L K D E T K T F T V T E
BINARY_1105 T T Y K L I L N L K Q A K E E A I T E L V D A G I A E K Y F K L I A N A K T V E G V W T Y K D E I L K A T V T E
BINARY_1106 T T Y K L I L N L K Q A K E E A I T E L V D A G I A E K Y F K L I A N A K T V E G V W T Y K D E I L K F T V T E
BINARY_1107 T T Y K L I L N L K Q A K E E A I T E L V D A G I A E K Y F K L I A N A K T V E G V W T Y K D E I L T A T V T E
BINARY_1108 T T Y K L I L N L K Q A K E E A I T E L V D A G I A E K Y F K L I A N A K T V E G V W T Y K D E I L T F T V T E
BINARY_1109 T T Y K L I L N L K Q A K E E A I T E L V D A G I A E K Y F K L I A N A K T V E G V W T Y K D E I K K A T V T E
BINARY_1110 T T Y K L I L N L K Q A K E E A I T E L V D A G I A E K Y F K L I A N A K T V E G V W T Y K D E I K K F T V T E
BINARY_1111 T T Y K L I L N L K Q A K E E A I T E L V D A G I A E K Y F K L I A N A K T V E G V W T Y K D E I K T A T V T E
BINARY_1112 T T Y K L I L N L K Q A K E E A I T E L V D A G I A E K Y F K L I A N A K T V E G V W T Y K D E I K T F T V T E
BINARY_1113 T T Y K L I L N L K Q A K E E A I T E L V D A G I A E K Y F K L I A N A K T V E G V W T Y K D E T L K A T V T E
BINARY_1114 T T Y K L I L N L K Q A K E E A I T E L V D A G I A E K Y F K L I A N A K T V E G V W T Y K D E T L K F T V T E
BINARY_1115 T T Y K L I L N L K Q A K E E A I T E L V D A G I A E K Y F K L I A N A K T V E G V W T Y K D E T L T A T V T E
BINARY_1116 T T Y K L I L N L K Q A K E E A I T E L V D A G I A E K Y F K L I A N A K T V E G V W T Y K D E T L T F T V T E
BINARY_1117 T T Y K L I L N L K Q A K E E A I T E L V D A G I A E K Y F K L I A N A K T V E G V W T Y K D E T K K A T V T E
BINARY_1118 T T Y K L I L N L K Q A K E E A I T E L V D A G I A E K Y F K L I A N A K T V E G V W T Y K D E T K K F T V T E
BINARY_1119 T T Y K L I L N L K Q A K E E A I T E L V D A G I A E K Y F K L I A N A K T V E G V W T Y K D E T K T A T V T E
BINARY_1120 T T Y K L I L N L K Q A K E E A I T E L V D A G I A E K Y F K L I A N A K T V E G V W T Y K D E T K T F T V T E
BINARY_1121 T T Y K L I L N L K Q A K E E A I T E L V D A G I A E K Y F K L Y A N A K T V E G V W T L K D E I L K A T V T E
BINARY_1122 T T Y K L I L N L K Q A K E E A I T E L V D A G I A E K Y F K L Y A N A K T V E G V W T L K D E I L K F T V T E
BINARY_1123 T T Y K L I L N L K Q A K E E A I T E L V D A G I A E K Y F K L Y A N A K T V E G V W T L K D E I L T A T V T E
BINARY_1124 T T Y K L I L N L K Q A K E E A I T E L V D A G I A E K Y F K L Y A N A K T V E G V W T L K D E I L T F T V T E
BINARY_1125 T T Y K L I L N L K Q A K E E A I T E L V D A G I A E K Y F K L Y A N A K T V E G V W T L K D E I K K A T V T E
BINARY_1126 T T Y K L I L N L K Q A K E E A I T E L V D A G I A E K Y F K L Y A N A K T V E G V W T L K D E I K K F T V T E
BINARY_1127 T T Y K L I L N L K Q A K E E A I T E L V D A G I A E K Y F K L Y A N A K T V E G V W T L K D E I K T A T V T E
BINARY_1128 T T Y K L I L N L K Q A K E E A I T E L V D A G I A E K Y F K L Y A N A K T V E G V W T L K D E I K T F T V T E
BINARY_1129 T T Y K L I L N L K Q A K E E A I T E L V D A G I A E K Y F K L Y A N A K T V E G V W T L K D E T L K A T V T E
BINARY_1130 T T Y K L I L N L K Q A K E E A I T E L V D A G I A E K Y F K L Y A N A K T V E G V W T L K D E T L K F T V T E
BINARY_1131 T T Y K L I L N L K Q A K E E A I T E L V D A G I A E K Y F K L Y A N A K T V E G V W T L K D E T L T A T V T E
BINARY_1132 T T Y K L I L N L K Q A K E E A I T E L V D A G I A E K Y F K L Y A N A K T V E G V W T L K D E T L T F T V T E
BINARY_1133 T T Y K L I L N L K Q A K E E A I T E L V D A G I A E K Y F K L Y A N A K T V E G V W T L K D E T K K A T V T E
BINARY_1134 T T Y K L I L N L K Q A K E E A I T E L V D A G I A E K Y F K L Y A N A K T V E G V W T L K D E T K K F T V T E
BINARY_1135 T T Y K L I L N L K Q A K E E A I T E L V D A G I A E K Y F K L Y A N A K T V E G V W T L K D E T K T A T V T E
BINARY_1136 T T Y K L I L N L K Q A K E E A I T E L V D A G I A E K Y F K L Y A N A K T V E G V W T L K D E T K T F T V T E
BINARY_1137 T T Y K L I L N L K Q A K E E A I T E L V D A G I A E K Y F K L Y A N A K T V E G V W T Y K D E I L K A T V T E
BINARY_1138 T T Y K L I L N L K Q A K E E A I T E L V D A G I A E K Y F K L Y A N A K T V E G V W T Y K D E I L K F T V T E
BINARY_1139 T T Y K L I L N L K Q A K E E A I T E L V D A G I A E K Y F K L Y A N A K T V E G V W T Y K D E I L T A T V T E
BINARY_1140 T T Y K L I L N L K Q A K E E A I T E L V D A G I A E K Y F K L Y A N A K T V E G V W T Y K D E I L T F T V T E
BINARY_1141 T T Y K L I L N L K Q A K E E A I T E L V D A G I A E K Y F K L Y A N A K T V E G V W T Y K D E I K K A T V T E
BINARY_1142 T T Y K L I L N L K Q A K E E A I T E L V D A G I A E K Y F K L Y A N A K T V E G V W T Y K D E I K K F T V T E
BINARY_1143 T T Y K L I L N L K Q A K E E A I T E L V D A G I A E K Y F K L Y A N A K T V E G V W T Y K D E I K T A T V T E
BINARY_1144 T T Y K L I L N L K Q A K E E A I T E L V D A G I A E K Y F K L Y A N A K T V E G V W T Y K D E I K T F T V T E
BINARY_1145 T T Y K L I L N L K Q A K E E A I T E L V D A G I A E K Y F K L Y A N A K T V E G V W T Y K D E T L K A T V T E
BINARY_1146 T T Y K L I L N L K Q A K E E A I T E L V D A G I A E K Y F K L Y A N A K T V E G V W T Y K D E T L K F T V T E
BINARY_1147 T T Y K L I L N L K Q A K E E A I T E L V D A G I A E K Y F K L Y A N A K T V E G V W T Y K D E T L T A T V T E
BINARY_1148 T T Y K L I L N L K Q A K E E A I T E L V D A G I A E K Y F K L Y A N A K T V E G V W T Y K D E T L T F T V T E
BINARY_1149 T T Y K L I L N L K Q A K E E A I T E L V D A G I A E K Y F K L Y A N A K T V E G V W T Y K D E T K K A T V T E
BINARY_1150 T T Y K L I L N L K Q A K E E A I T E L V D A G I A E K Y F K L Y A N A K T V E G V W T Y K D E T K K F T V T E
BINARY_1151 T T Y K L I L N L K Q A K E E A I T E L V D A G I A E K Y F K L Y A N A K T V E G V W T Y K D E T K T A T V T E
BINARY_1152 T T Y K L I L N L K Q A K E E A I T E L V D A G I A E K Y F K L Y A N A K T V E G V W T Y K D E T K T F T V T E
BINARY_1153 T T Y K L I L N L K Q A K E E A I T E L V D A G T A E K Y I K L I A N A K T V E G V W T L K D E I L K A T V T E
BINARY_1154 T T Y K L I L N L K Q A K E E A I T E L V D A G T A E K Y I K L I A N A K T V E G V W T L K D E I L K F T V T E
BINARY_1155 T T Y K L I L N L K Q A K E E A I T E L V D A G T A E K Y I K L I A N A K T V E G V W T L K D E I L T A T V T E
BINARY_1156 T T Y K L I L N L K Q A K E E A I T E L V D A G T A E K Y I K L I A N A K T V E G V W T L K D E I L T F T V T E

BINARY_1157 T T Y K L I L N L K Q A K E E A I T E L V D A G T A E K Y I K L I A N A K T V E G V W T L K D E I K K A T V T E
BINARY_1158 T T Y K L I L N L K Q A K E E A I T E L V D A G T A E K Y I K L I A N A K T V E G V W T L K D E I K K F T V T E
BINARY_1159 T T Y K L I L N L K Q A K E E A I T E L V D A G T A E K Y I K L I A N A K T V E G V W T L K D E I K T A T V T E
BINARY_1160 T T Y K L I L N L K Q A K E E A I T E L V D A G T A E K Y I K L I A N A K T V E G V W T L K D E I K T F T V T E
BINARY_1161 T T Y K L I L N L K Q A K E E A I T E L V D A G T A E K Y I K L I A N A K T V E G V W T L K D E T L K A T V T E
BINARY_1162 T T Y K L I L N L K Q A K E E A I T E L V D A G T A E K Y I K L I A N A K T V E G V W T L K D E T L K F T V T E
BINARY_1163 T T Y K L I L N L K Q A K E E A I T E L V D A G T A E K Y I K L I A N A K T V E G V W T L K D E T L T A T V T E
BINARY_1164 T T Y K L I L N L K Q A K E E A I T E L V D A G T A E K Y I K L I A N A K T V E G V W T L K D E T L T F T V T E
BINARY_1165 T T Y K L I L N L K Q A K E E A I T E L V D A G T A E K Y I K L I A N A K T V E G V W T L K D E T K K A T V T E
BINARY_1166 T T Y K L I L N L K Q A K E E A I T E L V D A G T A E K Y I K L I A N A K T V E G V W T L K D E T K K F T V T E
BINARY_1167 T T Y K L I L N L K Q A K E E A I T E L V D A G T A E K Y I K L I A N A K T V E G V W T L K D E T K T A T V T E
BINARY_1168 T T Y K L I L N L K Q A K E E A I T E L V D A G T A E K Y I K L I A N A K T V E G V W T L K D E T K T F T V T E
BINARY_1169 T T Y K L I L N L K Q A K E E A I T E L V D A G T A E K Y I K L I A N A K T V E G V W T Y K D E I L K A T V T E
BINARY_1170 T T Y K L I L N L K Q A K E E A I T E L V D A G T A E K Y I K L I A N A K T V E G V W T Y K D E I L K F T V T E
BINARY_1171 T T Y K L I L N L K Q A K E E A I T E L V D A G T A E K Y I K L I A N A K T V E G V W T Y K D E I L T A T V T E
BINARY_1172 T T Y K L I L N L K Q A K E E A I T E L V D A G T A E K Y I K L I A N A K T V E G V W T Y K D E I L T F T V T E
BINARY_1173 T T Y K L I L N L K Q A K E E A I T E L V D A G T A E K Y I K L I A N A K T V E G V W T Y K D E I K K A T V T E
BINARY_1174 T T Y K L I L N L K Q A K E E A I T E L V D A G T A E K Y I K L I A N A K T V E G V W T Y K D E I K K F T V T E
BINARY_1175 T T Y K L I L N L K Q A K E E A I T E L V D A G T A E K Y I K L I A N A K T V E G V W T Y K D E I K T A T V T E
BINARY_1176 T T Y K L I L N L K Q A K E E A I T E L V D A G T A E K Y I K L I A N A K T V E G V W T Y K D E I K T F T V T E
BINARY_1177 T T Y K L I L N L K Q A K E E A I T E L V D A G T A E K Y I K L I A N A K T V E G V W T Y K D E T L K A T V T E
BINARY_1178 T T Y K L I L N L K Q A K E E A I T E L V D A G T A E K Y I K L I A N A K T V E G V W T Y K D E T L K F T V T E
BINARY_1179 T T Y K L I L N L K Q A K E E A I T E L V D A G T A E K Y I K L I A N A K T V E G V W T Y K D E T L T A T V T E
BINARY_1180 T T Y K L I L N L K Q A K E E A I T E L V D A G T A E K Y I K L I A N A K T V E G V W T Y K D E T L T F T V T E
BINARY_1181 T T Y K L I L N L K Q A K E E A I T E L V D A G T A E K Y I K L I A N A K T V E G V W T Y K D E T K K A T V T E
BINARY_1182 T T Y K L I L N L K Q A K E E A I T E L V D A G T A E K Y I K L I A N A K T V E G V W T Y K D E T K K F T V T E
BINARY_1183 T T Y K L I L N L K Q A K E E A I T E L V D A G T A E K Y I K L I A N A K T V E G V W T Y K D E T K T A T V T E
BINARY_1184 T T Y K L I L N L K Q A K E E A I T E L V D A G T A E K Y I K L I A N A K T V E G V W T Y K D E T K T F T V T E
BINARY_1185 T T Y K L I L N L K Q A K E E A I T E L V D A G T A E K Y I K L Y A N A K T V E G V W T L K D E I L K A T V T E
BINARY_1186 T T Y K L I L N L K Q A K E E A I T E L V D A G T A E K Y I K L Y A N A K T V E G V W T L K D E I L K F T V T E
BINARY_1187 T T Y K L I L N L K Q A K E E A I T E L V D A G T A E K Y I K L Y A N A K T V E G V W T L K D E I L T A T V T E
BINARY_1188 T T Y K L I L N L K Q A K E E A I T E L V D A G T A E K Y I K L Y A N A K T V E G V W T L K D E I L T F T V T E
BINARY_1189 T T Y K L I L N L K Q A K E E A I T E L V D A G T A E K Y I K L Y A N A K T V E G V W T L K D E I K K A T V T E
BINARY_1190 T T Y K L I L N L K Q A K E E A I T E L V D A G T A E K Y I K L Y A N A K T V E G V W T L K D E I K K F T V T E
BINARY_1191 T T Y K L I L N L K Q A K E E A I T E L V D A G T A E K Y I K L Y A N A K T V E G V W T L K D E I K T A T V T E
BINARY_1192 T T Y K L I L N L K Q A K E E A I T E L V D A G T A E K Y I K L Y A N A K T V E G V W T L K D E I K T F T V T E
BINARY_1193 T T Y K L I L N L K Q A K E E A I T E L V D A G T A E K Y I K L Y A N A K T V E G V W T L K D E T L K A T V T E
BINARY_1194 T T Y K L I L N L K Q A K E E A I T E L V D A G T A E K Y I K L Y A N A K T V E G V W T L K D E T L K F T V T E
BINARY_1195 T T Y K L I L N L K Q A K E E A I T E L V D A G T A E K Y I K L Y A N A K T V E G V W T L K D E T L T A T V T E
BINARY_1196 T T Y K L I L N L K Q A K E E A I T E L V D A G T A E K Y I K L Y A N A K T V E G V W T L K D E T L T F T V T E
BINARY_1197 T T Y K L I L N L K Q A K E E A I T E L V D A G T A E K Y I K L Y A N A K T V E G V W T L K D E T K K A T V T E
BINARY_1198 T T Y K L I L N L K Q A K E E A I T E L V D A G T A E K Y I K L Y A N A K T V E G V W T L K D E T K K F T V T E
BINARY_1199 T T Y K L I L N L K Q A K E E A I T E L V D A G T A E K Y I K L Y A N A K T V E G V W T L K D E T K T A T V T E
BINARY_1200 T T Y K L I L N L K Q A K E E A I T E L V D A G T A E K Y I K L Y A N A K T V E G V W T L K D E T K T F T V T E
BINARY_1201 T T Y K L I L N L K Q A K E E A I T E L V D A G T A E K Y I K L Y A N A K T V E G V W T Y K D E I L K A T V T E
BINARY_1202 T T Y K L I L N L K Q A K E E A I T E L V D A G T A E K Y I K L Y A N A K T V E G V W T Y K D E I L K F T V T E
BINARY_1203 T T Y K L I L N L K Q A K E E A I T E L V D A G T A E K Y I K L Y A N A K T V E G V W T Y K D E I L T A T V T E
BINARY_1204 T T Y K L I L N L K Q A K E E A I T E L V D A G T A E K Y I K L Y A N A K T V E G V W T Y K D E I L T F T V T E
BINARY_1205 T T Y K L I L N L K Q A K E E A I T E L V D A G T A E K Y I K L Y A N A K T V E G V W T Y K D E I K K A T V T E
BINARY_1206 T T Y K L I L N L K Q A K E E A I T E L V D A G T A E K Y I K L Y A N A K T V E G V W T Y K D E I K K F T V T E
BINARY_1207 T T Y K L I L N L K Q A K E E A I T E L V D A G T A E K Y I K L Y A N A K T V E G V W T Y K D E I K T A T V T E
BINARY_1208 T T Y K L I L N L K Q A K E E A I T E L V D A G T A E K Y I K L Y A N A K T V E G V W T Y K D E I K T F T V T E
BINARY_1209 T T Y K L I L N L K Q A K E E A I T E L V D A G T A E K Y I K L Y A N A K T V E G V W T Y K D E T L K A T V T E
BINARY_1210 T T Y K L I L N L K Q A K E E A I T E L V D A G T A E K Y I K L Y A N A K T V E G V W T Y K D E T L K F T V T E
BINARY_1211 T T Y K L I L N L K Q A K E E A I T E L V D A G T A E K Y I K L Y A N A K T V E G V W T Y K D E T L T A T V T E
BINARY_1212 T T Y K L I L N L K Q A K E E A I T E L V D A G T A E K Y I K L Y A N A K T V E G V W T Y K D E T L T F T V T E
BINARY_1213 T T Y K L I L N L K Q A K E E A I T E L V D A G T A E K Y I K L Y A N A K T V E G V W T Y K D E T K K A T V T E
BINARY_1214 T T Y K L I L N L K Q A K E E A I T E L V D A G T A E K Y I K L Y A N A K T V E G V W T Y K D E T K K F T V T E
BINARY_1215 T T Y K L I L N L K Q A K E E A I T E L V D A G T A E K Y I K L Y A N A K T V E G V W T Y K D E T K T A T V T E
BINARY_1216 T T Y K L I L N L K Q A K E E A I T E L V D A G T A E K Y I K L Y A N A K T V E G V W T Y K D E T K T F T V T E
BINARY_1217 T T Y K L I L N L K Q A K E E A I T E L V D A G T A E K Y F K L I A N A K T V E G V W T L K D E I L K A T V T E
BINARY_1218 T T Y K L I L N L K Q A K E E A I T E L V D A G T A E K Y F K L I A N A K T V E G V W T L K D E I L K F T V T E
BINARY_1219 T T Y K L I L N L K Q A K E E A I T E L V D A G T A E K Y F K L I A N A K T V E G V W T L K D E I L T A T V T E
BINARY_1220 T T Y K L I L N L K Q A K E E A I T E L V D A G T A E K Y F K L I A N A K T V E G V W T L K D E I L T F T V T E
BINARY_1221 T T Y K L I L N L K Q A K E E A I T E L V D A G T A E K Y F K L I A N A K T V E G V W T L K D E I K K A T V T E
BINARY_1222 T T Y K L I L N L K Q A K E E A I T E L V D A G T A E K Y F K L I A N A K T V E G V W T L K D E I K K F T V T E
BINARY_1223 T T Y K L I L N L K Q A K E E A I T E L V D A G T A E K Y F K L I A N A K T V E G V W T L K D E I K T A T V T E
BINARY_1224 T T Y K L I L N L K Q A K E E A I T E L V D A G T A E K Y F K L I A N A K T V E G V W T L K D E I K T F T V T E
BINARY_1225 T T Y K L I L N L K Q A K E E A I T E L V D A G T A E K Y F K L I A N A K T V E G V W T L K D E T L K A T V T E
BINARY_1226 T T Y K L I L N L K Q A K E E A I T E L V D A G T A E K Y F K L I A N A K T V E G V W T L K D E T L K F T V T E
BINARY_1227 T T Y K L I L N L K Q A K E E A I T E L V D A G T A E K Y F K L I A N A K T V E G V W T L K D E T L T A T V T E
BINARY_1228 T T Y K L I L N L K Q A K E E A I T E L V D A G T A E K Y F K L I A N A K T V E G V W T L K D E T L T F T V T E
BINARY_1229 T T Y K L I L N L K Q A K E E A I T E L V D A G T A E K Y F K L I A N A K T V E G V W T L K D E T K K A T V T E
BINARY_1230 T T Y K L I L N L K Q A K E E A I T E L V D A G T A E K Y F K L I A N A K T V E G V W T L K D E T K K F T V T E
BINARY_1231 T T Y K L I L N L K Q A K E E A I T E L V D A G T A E K Y F K L I A N A K T V E G V W T L K D E T K T A T V T E
BINARY_1232 T T Y K L I L N L K Q A K E E A I T E L V D A G T A E K Y F K L I A N A K T V E G V W T L K D E T K T F T V T E
BINARY_1233 T T Y K L I L N L K Q A K E E A I T E L V D A G T A E K Y F K L I A N A K T V E G V W T Y K D E I L K A T V T E
BINARY_1234 T T Y K L I L N L K Q A K E E A I T E L V D A G T A E K Y F K L I A N A K T V E G V W T Y K D E I L K F T V T E
BINARY_1235 T T Y K L I L N L K Q A K E E A I T E L V D A G T A E K Y F K L I A N A K T V E G V W T Y K D E I L T A T V T E
BINARY_1236 T T Y K L I L N L K Q A K E E A I T E L V D A G T A E K Y F K L I A N A K T V E G V W T Y K D E I L T F T V T E
BINARY_1237 T T Y K L I L N L K Q A K E E A I T E L V D A G T A E K Y F K L I A N A K T V E G V W T Y K D E I K K A T V T E
BINARY_1238 T T Y K L I L N L K Q A K E E A I T E L V D A G T A E K Y F K L I A N A K T V E G V W T Y K D E I K K F T V T E
BINARY_1239 T T Y K L I L N L K Q A K E E A I T E L V D A G T A E K Y F K L I A N A K T V E G V W T Y K D E I K T A T V T E
BINARY_1240 T T Y K L I L N L K Q A K E E A I T E L V D A G T A E K Y F K L I A N A K T V E G V W T Y K D E I K T F T V T E
BINARY_1241 T T Y K L I L N L K Q A K E E A I T E L V D A G T A E K Y F K L I A N A K T V E G V W T Y K D E T L K A T V T E
BINARY_1242 T T Y K L I L N L K Q A K E E A I T E L V D A G T A E K Y F K L I A N A K T V E G V W T Y K D E T L K F T V T E
BINARY_1243 T T Y K L I L N L K Q A K E E A I T E L V D A G T A E K Y F K L I A N A K T V E G V W T Y K D E T L T A T V T E
BINARY_1244 T T Y K L I L N L K Q A K E E A I T E L V D A G T A E K Y F K L I A N A K T V E G V W T Y K D E T L T F T V T E
BINARY_1245 T T Y K L I L N L K Q A K E E A I T E L V D A G T A E K Y F K L I A N A K T V E G V W T Y K D E T K K A T V T E
BINARY_1246 T T Y K L I L N L K Q A K E E A I T E L V D A G T A E K Y F K L I A N A K T V E G V W T Y K D E T K K F T V T E
BINARY_1247 T T Y K L I L N L K Q A K E E A I T E L V D A G T A E K Y F K L I A N A K T V E G V W T Y K D E T K T A T V T E
BINARY_1248 T T Y K L I L N L K Q A K E E A I T E L V D A G T A E K Y F K L I A N A K T V E G V W T Y K D E T K T F T V T E
BINARY_1249 T T Y K L I L N L K Q A K E E A I T E L V D A G T A E K Y F K L Y A N A K T V E G V W T L K D E I L K A T V T E
BINARY_1250 T T Y K L I L N L K Q A K E E A I T E L V D A G T A E K Y F K L Y A N A K T V E G V W T L K D E I L K F T V T E
BINARY_1251 T T Y K L I L N L K Q A K E E A I T E L V D A G T A E K Y F K L Y A N A K T V E G V W T L K D E I L T A T V T E

BINARY_1252 T T Y K L I L N L K Q A K E E A I T E L V D A G T A E K Y F K L Y A N A K T V E G V W T L K D E I L T F T V T E
BINARY_1253 T T Y K L I L N L K Q A K E E A I T E L V D A G T A E K Y F K L Y A N A K T V E G V W T L K D E I K K A T V T E
BINARY_1254 T T Y K L I L N L K Q A K E E A I T E L V D A G T A E K Y F K L Y A N A K T V E G V W T L K D E I K K F T V T E
BINARY_1255 T T Y K L I L N L K Q A K E E A I T E L V D A G T A E K Y F K L Y A N A K T V E G V W T L K D E I K T A T V T E
BINARY_1256 T T Y K L I L N L K Q A K E E A I T E L V D A G T A E K Y F K L Y A N A K T V E G V W T L K D E I K T F T V T E
BINARY_1257 T T Y K L I L N L K Q A K E E A I T E L V D A G T A E K Y F K L Y A N A K T V E G V W T L K D E T L K A T V T E
BINARY_1258 T T Y K L I L N L K Q A K E E A I T E L V D A G T A E K Y F K L Y A N A K T V E G V W T L K D E T L K F T V T E
BINARY_1259 T T Y K L I L N L K Q A K E E A I T E L V D A G T A E K Y F K L Y A N A K T V E G V W T L K D E T L T A T V T E
BINARY_1260 T T Y K L I L N L K Q A K E E A I T E L V D A G T A E K Y F K L Y A N A K T V E G V W T L K D E T L T F T V T E
BINARY_1261 T T Y K L I L N L K Q A K E E A I T E L V D A G T A E K Y F K L Y A N A K T V E G V W T L K D E T K K A T V T E
BINARY_1262 T T Y K L I L N L K Q A K E E A I T E L V D A G T A E K Y F K L Y A N A K T V E G V W T L K D E T K K F T V T E
BINARY_1263 T T Y K L I L N L K Q A K E E A I T E L V D A G T A E K Y F K L Y A N A K T V E G V W T L K D E T K T A T V T E
BINARY_1264 T T Y K L I L N L K Q A K E E A I T E L V D A G T A E K Y F K L Y A N A K T V E G V W T L K D E T K T F T V T E
BINARY_1265 T T Y K L I L N L K Q A K E E A I T E L V D A G T A E K Y F K L Y A N A K T V E G V W T Y K D E I L K A T V T E
BINARY_1266 T T Y K L I L N L K Q A K E E A I T E L V D A G T A E K Y F K L Y A N A K T V E G V W T Y K D E I L K F T V T E
BINARY_1267 T T Y K L I L N L K Q A K E E A I T E L V D A G T A E K Y F K L Y A N A K T V E G V W T Y K D E I L T A T V T E
BINARY_1268 T T Y K L I L N L K Q A K E E A I T E L V D A G T A E K Y F K L Y A N A K T V E G V W T Y K D E I L T F T V T E
BINARY_1269 T T Y K L I L N L K Q A K E E A I T E L V D A G T A E K Y F K L Y A N A K T V E G V W T Y K D E I K K A T V T E
BINARY_1270 T T Y K L I L N L K Q A K E E A I T E L V D A G T A E K Y F K L Y A N A K T V E G V W T Y K D E I K K F T V T E
BINARY_1271 T T Y K L I L N L K Q A K E E A I T E L V D A G T A E K Y F K L Y A N A K T V E G V W T Y K D E I K T A T V T E
BINARY_1272 T T Y K L I L N L K Q A K E E A I T E L V D A G T A E K Y F K L Y A N A K T V E G V W T Y K D E I K T F T V T E
BINARY_1273 T T Y K L I L N L K Q A K E E A I T E L V D A G T A E K Y F K L Y A N A K T V E G V W T Y K D E T L K A T V T E
BINARY_1274 T T Y K L I L N L K Q A K E E A I T E L V D A G T A E K Y F K L Y A N A K T V E G V W T Y K D E T L K F T V T E
BINARY_1275 T T Y K L I L N L K Q A K E E A I T E L V D A G T A E K Y F K L Y A N A K T V E G V W T Y K D E T L T A T V T E
BINARY_1276 T T Y K L I L N L K Q A K E E A I T E L V D A G T A E K Y F K L Y A N A K T V E G V W T Y K D E T L T F T V T E
BINARY_1277 T T Y K L I L N L K Q A K E E A I T E L V D A G T A E K Y F K L Y A N A K T V E G V W T Y K D E T K K A T V T E
BINARY_1278 T T Y K L I L N L K Q A K E E A I T E L V D A G T A E K Y F K L Y A N A K T V E G V W T Y K D E T K K F T V T E
BINARY_1279 T T Y K L I L N L K Q A K E E A I T E L V D A G T A E K Y F K L Y A N A K T V E G V W T Y K D E T K T A T V T E
BINARY_1280 T T Y K L I L N L K Q A K E E A I T E L V D A G T A E K Y F K L Y A N A K T V E G V W T Y K D E T K T F T V T E
BINARY_1281 T T Y K L I L N L K Q A K E E A I T E L V D A A I A E K Y I K L I A N A K T V E G V W T L K D E I L K A T V T E
BINARY_1282 T T Y K L I L N L K Q A K E E A I T E L V D A A I A E K Y I K L I A N A K T V E G V W T L K D E I L K F T V T E
BINARY_1283 T T Y K L I L N L K Q A K E E A I T E L V D A A I A E K Y I K L I A N A K T V E G V W T L K D E I L T A T V T E
BINARY_1284 T T Y K L I L N L K Q A K E E A I T E L V D A A I A E K Y I K L I A N A K T V E G V W T L K D E I L T F T V T E
BINARY_1285 T T Y K L I L N L K Q A K E E A I T E L V D A A I A E K Y I K L I A N A K T V E G V W T L K D E I K K A T V T E
BINARY_1286 T T Y K L I L N L K Q A K E E A I T E L V D A A I A E K Y I K L I A N A K T V E G V W T L K D E I K K F T V T E
BINARY_1287 T T Y K L I L N L K Q A K E E A I T E L V D A A I A E K Y I K L I A N A K T V E G V W T L K D E I K T A T V T E
BINARY_1288 T T Y K L I L N L K Q A K E E A I T E L V D A A I A E K Y I K L I A N A K T V E G V W T L K D E I K T F T V T E
BINARY_1289 T T Y K L I L N L K Q A K E E A I T E L V D A A I A E K Y I K L I A N A K T V E G V W T L K D E T L K A T V T E
BINARY_1290 T T Y K L I L N L K Q A K E E A I T E L V D A A I A E K Y I K L I A N A K T V E G V W T L K D E T L K F T V T E
BINARY_1291 T T Y K L I L N L K Q A K E E A I T E L V D A A I A E K Y I K L I A N A K T V E G V W T L K D E T L T A T V T E
BINARY_1292 T T Y K L I L N L K Q A K E E A I T E L V D A A I A E K Y I K L I A N A K T V E G V W T L K D E T L T F T V T E
BINARY_1293 T T Y K L I L N L K Q A K E E A I T E L V D A A I A E K Y I K L I A N A K T V E G V W T L K D E T K K A T V T E
BINARY_1294 T T Y K L I L N L K Q A K E E A I T E L V D A A I A E K Y I K L I A N A K T V E G V W T L K D E T K K F T V T E
BINARY_1295 T T Y K L I L N L K Q A K E E A I T E L V D A A I A E K Y I K L I A N A K T V E G V W T L K D E T K T A T V T E
BINARY_1296 T T Y K L I L N L K Q A K E E A I T E L V D A A I A E K Y I K L I A N A K T V E G V W T L K D E T K T F T V T E
BINARY_1297 T T Y K L I L N L K Q A K E E A I T E L V D A A I A E K Y I K L I A N A K T V E G V W T Y K D E I L K A T V T E
BINARY_1298 T T Y K L I L N L K Q A K E E A I T E L V D A A I A E K Y I K L I A N A K T V E G V W T Y K D E I L K F T V T E
BINARY_1299 T T Y K L I L N L K Q A K E E A I T E L V D A A I A E K Y I K L I A N A K T V E G V W T Y K D E I L T A T V T E
BINARY_1300 T T Y K L I L N L K Q A K E E A I T E L V D A A I A E K Y I K L I A N A K T V E G V W T Y K D E I L T F T V T E
BINARY_1301 T T Y K L I L N L K Q A K E E A I T E L V D A A I A E K Y I K L I A N A K T V E G V W T Y K D E I K K A T V T E
BINARY_1302 T T Y K L I L N L K Q A K E E A I T E L V D A A I A E K Y I K L I A N A K T V E G V W T Y K D E I K K F T V T E
BINARY_1303 T T Y K L I L N L K Q A K E E A I T E L V D A A I A E K Y I K L I A N A K T V E G V W T Y K D E I K T A T V T E
BINARY_1304 T T Y K L I L N L K Q A K E E A I T E L V D A A I A E K Y I K L I A N A K T V E G V W T Y K D E I K T F T V T E
BINARY_1305 T T Y K L I L N L K Q A K E E A I T E L V D A A I A E K Y I K L I A N A K T V E G V W T Y K D E T L K A T V T E
BINARY_1306 T T Y K L I L N L K Q A K E E A I T E L V D A A I A E K Y I K L I A N A K T V E G V W T Y K D E T L K F T V T E
BINARY_1307 T T Y K L I L N L K Q A K E E A I T E L V D A A I A E K Y I K L I A N A K T V E G V W T Y K D E T L T A T V T E
BINARY_1308 T T Y K L I L N L K Q A K E E A I T E L V D A A I A E K Y I K L I A N A K T V E G V W T Y K D E T L T F T V T E
BINARY_1309 T T Y K L I L N L K Q A K E E A I T E L V D A A I A E K Y I K L I A N A K T V E G V W T Y K D E T K K A T V T E
BINARY_1310 T T Y K L I L N L K Q A K E E A I T E L V D A A I A E K Y I K L I A N A K T V E G V W T Y K D E T K K F T V T E
BINARY_1311 T T Y K L I L N L K Q A K E E A I T E L V D A A I A E K Y I K L I A N A K T V E G V W T Y K D E T K T A T V T E
BINARY_1312 T T Y K L I L N L K Q A K E E A I T E L V D A A I A E K Y I K L I A N A K T V E G V W T Y K D E T K T F T V T E
BINARY_1313 T T Y K L I L N L K Q A K E E A I T E L V D A A I A E K Y I K L Y A N A K T V E G V W T L K D E I L K A T V T E
BINARY_1314 T T Y K L I L N L K Q A K E E A I T E L V D A A I A E K Y I K L Y A N A K T V E G V W T L K D E I L K F T V T E
BINARY_1315 T T Y K L I L N L K Q A K E E A I T E L V D A A I A E K Y I K L Y A N A K T V E G V W T L K D E I L T A T V T E
BINARY_1316 T T Y K L I L N L K Q A K E E A I T E L V D A A I A E K Y I K L Y A N A K T V E G V W T L K D E I L T F T V T E
BINARY_1317 T T Y K L I L N L K Q A K E E A I T E L V D A A I A E K Y I K L Y A N A K T V E G V W T L K D E I K K A T V T E
BINARY_1318 T T Y K L I L N L K Q A K E E A I T E L V D A A I A E K Y I K L Y A N A K T V E G V W T L K D E I K K F T V T E
BINARY_1319 T T Y K L I L N L K Q A K E E A I T E L V D A A I A E K Y I K L Y A N A K T V E G V W T L K D E I K T A T V T E
BINARY_1320 T T Y K L I L N L K Q A K E E A I T E L V D A A I A E K Y I K L Y A N A K T V E G V W T L K D E I K T F T V T E
BINARY_1321 T T Y K L I L N L K Q A K E E A I T E L V D A A I A E K Y I K L Y A N A K T V E G V W T L K D E T L K A T V T E
BINARY_1322 T T Y K L I L N L K Q A K E E A I T E L V D A A I A E K Y I K L Y A N A K T V E G V W T L K D E T L K F T V T E
BINARY_1323 T T Y K L I L N L K Q A K E E A I T E L V D A A I A E K Y I K L Y A N A K T V E G V W T L K D E T L T A T V T E
BINARY_1324 T T Y K L I L N L K Q A K E E A I T E L V D A A I A E K Y I K L Y A N A K T V E G V W T L K D E T L T F T V T E
BINARY_1325 T T Y K L I L N L K Q A K E E A I T E L V D A A I A E K Y I K L Y A N A K T V E G V W T L K D E T K K A T V T E
BINARY_1326 T T Y K L I L N L K Q A K E E A I T E L V D A A I A E K Y I K L Y A N A K T V E G V W T L K D E T K K F T V T E
BINARY_1327 T T Y K L I L N L K Q A K E E A I T E L V D A A I A E K Y I K L Y A N A K T V E G V W T L K D E T K T A T V T E
BINARY_1328 T T Y K L I L N L K Q A K E E A I T E L V D A A I A E K Y I K L Y A N A K T V E G V W T L K D E T K T F T V T E
BINARY_1329 T T Y K L I L N L K Q A K E E A I T E L V D A A I A E K Y I K L Y A N A K T V E G V W T Y K D E I L K A T V T E
BINARY_1330 T T Y K L I L N L K Q A K E E A I T E L V D A A I A E K Y I K L Y A N A K T V E G V W T Y K D E I L K F T V T E
BINARY_1331 T T Y K L I L N L K Q A K E E A I T E L V D A A I A E K Y I K L Y A N A K T V E G V W T Y K D E I L T A T V T E
BINARY_1332 T T Y K L I L N L K Q A K E E A I T E L V D A A I A E K Y I K L Y A N A K T V E G V W T Y K D E I L T F T V T E
BINARY_1333 T T Y K L I L N L K Q A K E E A I T E L V D A A I A E K Y I K L Y A N A K T V E G V W T Y K D E I K K A T V T E
BINARY_1334 T T Y K L I L N L K Q A K E E A I T E L V D A A I A E K Y I K L Y A N A K T V E G V W T Y K D E I K K F T V T E
BINARY_1335 T T Y K L I L N L K Q A K E E A I T E L V D A A I A E K Y I K L Y A N A K T V E G V W T Y K D E I K T A T V T E
BINARY_1336 T T Y K L I L N L K Q A K E E A I T E L V D A A I A E K Y I K L Y A N A K T V E G V W T Y K D E I K T F T V T E
BINARY_1337 T T Y K L I L N L K Q A K E E A I T E L V D A A I A E K Y I K L Y A N A K T V E G V W T Y K D E T L K A T V T E
BINARY_1338 T T Y K L I L N L K Q A K E E A I T E L V D A A I A E K Y I K L Y A N A K T V E G V W T Y K D E T L K F T V T E
BINARY_1339 T T Y K L I L N L K Q A K E E A I T E L V D A A I A E K Y I K L Y A N A K T V E G V W T Y K D E T L T A T V T E
BINARY_1340 T T Y K L I L N L K Q A K E E A I T E L V D A A I A E K Y I K L Y A N A K T V E G V W T Y K D E T L T F T V T E
BINARY_1341 T T Y K L I L N L K Q A K E E A I T E L V D A A I A E K Y I K L Y A N A K T V E G V W T Y K D E T K K A T V T E
BINARY_1342 T T Y K L I L N L K Q A K E E A I T E L V D A A I A E K Y I K L Y A N A K T V E G V W T Y K D E T K K F T V T E
BINARY_1343 T T Y K L I L N L K Q A K E E A I T E L V D A A I A E K Y I K L Y A N A K T V E G V W T Y K D E T K T A T V T E
BINARY_1344 T T Y K L I L N L K Q A K E E A I T E L V D A A I A E K Y I K L Y A N A K T V E G V W T Y K D E T K T F T V T E
BINARY_1345 T T Y K L I L N L K Q A K E E A I T E L V D A A I A E K Y F K L I A N A K T V E G V W T L K D E I L K A T V T E
BINARY_1346 T T Y K L I L N L K Q A K E E A I T E L V D A A I A E K Y F K L I A N A K T V E G V W T L K D E I L K F T V T E

```
BINARY_1347 T T Y K L I L N L K Q A K E E A I T E L V D A A I A E K Y F K L I A N A K T V E G V W T L K D E I L T A T V T E
BINARY_1348 T T Y K L I L N L K Q A K E E A I T E L V D A A I A E K Y F K L I A N A K T V E G V W T L K D E I L T F T V T E
BINARY_1349 T T Y K L I L N L K Q A K E E A I T E L V D A A I A E K Y F K L I A N A K T V E G V W T L K D E I K K A T V T E
BINARY_1350 T T Y K L I L N L K Q A K E E A I T E L V D A A I A E K Y F K L I A N A K T V E G V W T L K D E I K K F T V T E
BINARY_1351 T T Y K L I L N L K Q A K E E A I T E L V D A A I A E K Y F K L I A N A K T V E G V W T L K D E I K T A T V T E
BINARY_1352 T T Y K L I L N L K Q A K E E A I T E L V D A A I A E K Y F K L I A N A K T V E G V W T L K D E I K T F T V T E
BINARY_1353 T T Y K L I L N L K Q A K E E A I T E L V D A A I A E K Y F K L I A N A K T V E G V W T L K D E T L K A T V T E
BINARY_1354 T T Y K L I L N L K Q A K E E A I T E L V D A A I A E K Y F K L I A N A K T V E G V W T L K D E T L K F T V T E
BINARY_1355 T T Y K L I L N L K Q A K E E A I T E L V D A A I A E K Y F K L I A N A K T V E G V W T L K D E T L T A T V T E
BINARY_1356 T T Y K L I L N L K Q A K E E A I T E L V D A A I A E K Y F K L I A N A K T V E G V W T L K D E T L T F T V T E
BINARY_1357 T T Y K L I L N L K Q A K E E A I T E L V D A A I A E K Y F K L I A N A K T V E G V W T L K D E T K K A T V T E
BINARY_1358 T T Y K L I L N L K Q A K E E A I T E L V D A A I A E K Y F K L I A N A K T V E G V W T L K D E T K K F T V T E
BINARY_1359 T T Y K L I L N L K Q A K E E A I T E L V D A A I A E K Y F K L I A N A K T V E G V W T L K D E T K T A T V T E
BINARY_1360 T T Y K L I L N L K Q A K E E A I T E L V D A A I A E K Y F K L I A N A K T V E G V W T L K D E T K T F T V T E
BINARY_1361 T T Y K L I L N L K Q A K E E A I T E L V D A A I A E K Y F K L I A N A K T V E G V W T Y K D E I L K A T V T E
BINARY_1362 T T Y K L I L N L K Q A K E E A I T E L V D A A I A E K Y F K L I A N A K T V E G V W T Y K D E I L K F T V T E
BINARY_1363 T T Y K L I L N L K Q A K E E A I T E L V D A A I A E K Y F K L I A N A K T V E G V W T Y K D E I L T A T V T E
BINARY_1364 T T Y K L I L N L K Q A K E E A I T E L V D A A I A E K Y F K L I A N A K T V E G V W T Y K D E I L T F T V T E
BINARY_1365 T T Y K L I L N L K Q A K E E A I T E L V D A A I A E K Y F K L I A N A K T V E G V W T Y K D E I K K A T V T E
BINARY_1366 T T Y K L I L N L K Q A K E E A I T E L V D A A I A E K Y F K L I A N A K T V E G V W T Y K D E I K K F T V T E
BINARY_1367 T T Y K L I L N L K Q A K E E A I T E L V D A A I A E K Y F K L I A N A K T V E G V W T Y K D E I K T A T V T E
BINARY_1368 T T Y K L I L N L K Q A K E E A I T E L V D A A I A E K Y F K L I A N A K T V E G V W T Y K D E I K T F T V T E
BINARY_1369 T T Y K L I L N L K Q A K E E A I T E L V D A A I A E K Y F K L I A N A K T V E G V W T Y K D E T L K A T V T E
BINARY_1370 T T Y K L I L N L K Q A K E E A I T E L V D A A I A E K Y F K L I A N A K T V E G V W T Y K D E T L K F T V T E
BINARY_1371 T T Y K L I L N L K Q A K E E A I T E L V D A A I A E K Y F K L I A N A K T V E G V W T Y K D E T L T A T V T E
BINARY_1372 T T Y K L I L N L K Q A K E E A I T E L V D A A I A E K Y F K L I A N A K T V E G V W T Y K D E T L T F T V T E
BINARY_1373 T T Y K L I L N L K Q A K E E A I T E L V D A A I A E K Y F K L I A N A K T V E G V W T Y K D E T K K A T V T E
BINARY_1374 T T Y K L I L N L K Q A K E E A I T E L V D A A I A E K Y F K L I A N A K T V E G V W T Y K D E T K K F T V T E
BINARY_1375 T T Y K L I L N L K Q A K E E A I T E L V D A A I A E K Y F K L I A N A K T V E G V W T Y K D E T K T A T V T E
BINARY_1376 T T Y K L I L N L K Q A K E E A I T E L V D A A I A E K Y F K L I A N A K T V E G V W T Y K D E T K T F T V T E
BINARY_1377 T T Y K L I L N L K Q A K E E A I T E L V D A A I A E K Y F K L Y A N A K T V E G V W T L K D E I L K A T V T E
BINARY_1378 T T Y K L I L N L K Q A K E E A I T E L V D A A I A E K Y F K L Y A N A K T V E G V W T L K D E I L K F T V T E
BINARY_1379 T T Y K L I L N L K Q A K E E A I T E L V D A A I A E K Y F K L Y A N A K T V E G V W T L K D E I L T A T V T E
BINARY_1380 T T Y K L I L N L K Q A K E E A I T E L V D A A I A E K Y F K L Y A N A K T V E G V W T L K D E I L T F T V T E
BINARY_1381 T T Y K L I L N L K Q A K E E A I T E L V D A A I A E K Y F K L Y A N A K T V E G V W T L K D E I K K A T V T E
BINARY_1382 T T Y K L I L N L K Q A K E E A I T E L V D A A I A E K Y F K L Y A N A K T V E G V W T L K D E I K K F T V T E
BINARY_1383 T T Y K L I L N L K Q A K E E A I T E L V D A A I A E K Y F K L Y A N A K T V E G V W T L K D E I K T A T V T E
BINARY_1384 T T Y K L I L N L K Q A K E E A I T E L V D A A I A E K Y F K L Y A N A K T V E G V W T L K D E I K T F T V T E
BINARY_1385 T T Y K L I L N L K Q A K E E A I T E L V D A A I A E K Y F K L Y A N A K T V E G V W T L K D E T L K A T V T E
BINARY_1386 T T Y K L I L N L K Q A K E E A I T E L V D A A I A E K Y F K L Y A N A K T V E G V W T L K D E T L K F T V T E
BINARY_1387 T T Y K L I L N L K Q A K E E A I T E L V D A A I A E K Y F K L Y A N A K T V E G V W T L K D E T L T A T V T E
BINARY_1388 T T Y K L I L N L K Q A K E E A I T E L V D A A I A E K Y F K L Y A N A K T V E G V W T L K D E T L T F T V T E
BINARY_1389 T T Y K L I L N L K Q A K E E A I T E L V D A A I A E K Y F K L Y A N A K T V E G V W T L K D E T K K A T V T E
BINARY_1390 T T Y K L I L N L K Q A K E E A I T E L V D A A I A E K Y F K L Y A N A K T V E G V W T L K D E T K K F T V T E
BINARY_1391 T T Y K L I L N L K Q A K E E A I T E L V D A A I A E K Y F K L Y A N A K T V E G V W T L K D E T K T A T V T E
BINARY_1392 T T Y K L I L N L K Q A K E E A I T E L V D A A I A E K Y F K L Y A N A K T V E G V W T L K D E T K T F T V T E
BINARY_1393 T T Y K L I L N L K Q A K E E A I T E L V D A A I A E K Y F K L Y A N A K T V E G V W T Y K D E I L K A T V T E
BINARY_1394 T T Y K L I L N L K Q A K E E A I T E L V D A A I A E K Y F K L Y A N A K T V E G V W T Y K D E I L K F T V T E
BINARY_1395 T T Y K L I L N L K Q A K E E A I T E L V D A A I A E K Y F K L Y A N A K T V E G V W T Y K D E I L T A T V T E
BINARY_1396 T T Y K L I L N L K Q A K E E A I T E L V D A A I A E K Y F K L Y A N A K T V E G V W T Y K D E I L T F T V T E
BINARY_1397 T T Y K L I L N L K Q A K E E A I T E L V D A A I A E K Y F K L Y A N A K T V E G V W T Y K D E I K K A T V T E
BINARY_1398 T T Y K L I L N L K Q A K E E A I T E L V D A A I A E K Y F K L Y A N A K T V E G V W T Y K D E I K K F T V T E
BINARY_1399 T T Y K L I L N L K Q A K E E A I T E L V D A A I A E K Y F K L Y A N A K T V E G V W T Y K D E I K T A T V T E
BINARY_1400 T T Y K L I L N L K Q A K E E A I T E L V D A A I A E K Y F K L Y A N A K T V E G V W T Y K D E I K T F T V T E
BINARY_1401 T T Y K L I L N L K Q A K E E A I T E L V D A A I A E K Y F K L Y A N A K T V E G V W T Y K D E T L K A T V T E
BINARY_1402 T T Y K L I L N L K Q A K E E A I T E L V D A A I A E K Y F K L Y A N A K T V E G V W T Y K D E T L K F T V T E
BINARY_1403 T T Y K L I L N L K Q A K E E A I T E L V D A A I A E K Y F K L Y A N A K T V E G V W T Y K D E T L T A T V T E
BINARY_1404 T T Y K L I L N L K Q A K E E A I T E L V D A A I A E K Y F K L Y A N A K T V E G V W T Y K D E T L T F T V T E
BINARY_1405 T T Y K L I L N L K Q A K E E A I T E L V D A A I A E K Y F K L Y A N A K T V E G V W T Y K D E T K K A T V T E
BINARY_1406 T T Y K L I L N L K Q A K E E A I T E L V D A A I A E K Y F K L Y A N A K T V E G V W T Y K D E T K K F T V T E
BINARY_1407 T T Y K L I L N L K Q A K E E A I T E L V D A A I A E K Y F K L Y A N A K T V E G V W T Y K D E T K T A T V T E
BINARY_1408 T T Y K L I L N L K Q A K E E A I T E L V D A A I A E K Y F K L Y A N A K T V E G V W T Y K D E T K T F T V T E
BINARY_1409 T T Y K L I L N L K Q A K E E A I T E L V D A A T A E K Y I K L I A N A K T V E G V W T L K D E I L K A T V T E
BINARY_1410 T T Y K L I L N L K Q A K E E A I T E L V D A A T A E K Y I K L I A N A K T V E G V W T L K D E I L K F T V T E
BINARY_1411 T T Y K L I L N L K Q A K E E A I T E L V D A A T A E K Y I K L I A N A K T V E G V W T L K D E I L T A T V T E
BINARY_1412 T T Y K L I L N L K Q A K E E A I T E L V D A A T A E K Y I K L I A N A K T V E G V W T L K D E I L T F T V T E
BINARY_1413 T T Y K L I L N L K Q A K E E A I T E L V D A A T A E K Y I K L I A N A K T V E G V W T L K D E I K K A T V T E
BINARY_1414 T T Y K L I L N L K Q A K E E A I T E L V D A A T A E K Y I K L I A N A K T V E G V W T L K D E I K K F T V T E
BINARY_1415 T T Y K L I L N L K Q A K E E A I T E L V D A A T A E K Y I K L I A N A K T V E G V W T L K D E I K T A T V T E
BINARY_1416 T T Y K L I L N L K Q A K E E A I T E L V D A A T A E K Y I K L I A N A K T V E G V W T L K D E I K T F T V T E
BINARY_1417 T T Y K L I L N L K Q A K E E A I T E L V D A A T A E K Y I K L I A N A K T V E G V W T L K D E T L K A T V T E
BINARY_1418 T T Y K L I L N L K Q A K E E A I T E L V D A A T A E K Y I K L I A N A K T V E G V W T L K D E T L K F T V T E
BINARY_1419 T T Y K L I L N L K Q A K E E A I T E L V D A A T A E K Y I K L I A N A K T V E G V W T L K D E T L T A T V T E
BINARY_1420 T T Y K L I L N L K Q A K E E A I T E L V D A A T A E K Y I K L I A N A K T V E G V W T L K D E T L T F T V T E
BINARY_1421 T T Y K L I L N L K Q A K E E A I T E L V D A A T A E K Y I K L I A N A K T V E G V W T L K D E T K K A T V T E
BINARY_1422 T T Y K L I L N L K Q A K E E A I T E L V D A A T A E K Y I K L I A N A K T V E G V W T L K D E T K K F T V T E
BINARY_1423 T T Y K L I L N L K Q A K E E A I T E L V D A A T A E K Y I K L I A N A K T V E G V W T L K D E T K T A T V T E
BINARY_1424 T T Y K L I L N L K Q A K E E A I T E L V D A A T A E K Y I K L I A N A K T V E G V W T L K D E T K T F T V T E
BINARY_1425 T T Y K L I L N L K Q A K E E A I T E L V D A A T A E K Y I K L I A N A K T V E G V W T Y K D E I L K A T V T E
BINARY_1426 T T Y K L I L N L K Q A K E E A I T E L V D A A T A E K Y I K L I A N A K T V E G V W T Y K D E I L K F T V T E
BINARY_1427 T T Y K L I L N L K Q A K E E A I T E L V D A A T A E K Y I K L I A N A K T V E G V W T Y K D E I L T A T V T E
BINARY_1428 T T Y K L I L N L K Q A K E E A I T E L V D A A T A E K Y I K L I A N A K T V E G V W T Y K D E I L T F T V T E
BINARY_1429 T T Y K L I L N L K Q A K E E A I T E L V D A A T A E K Y I K L I A N A K T V E G V W T Y K D E I K K A T V T E
BINARY_1430 T T Y K L I L N L K Q A K E E A I T E L V D A A T A E K Y I K L I A N A K T V E G V W T Y K D E I K K F T V T E
BINARY_1431 T T Y K L I L N L K Q A K E E A I T E L V D A A T A E K Y I K L I A N A K T V E G V W T Y K D E I K T A T V T E
BINARY_1432 T T Y K L I L N L K Q A K E E A I T E L V D A A T A E K Y I K L I A N A K T V E G V W T Y K D E I K T F T V T E
BINARY_1433 T T Y K L I L N L K Q A K E E A I T E L V D A A T A E K Y I K L I A N A K T V E G V W T Y K D E T L K A T V T E
BINARY_1434 T T Y K L I L N L K Q A K E E A I T E L V D A A T A E K Y I K L I A N A K T V E G V W T Y K D E T L K F T V T E
BINARY_1435 T T Y K L I L N L K Q A K E E A I T E L V D A A T A E K Y I K L I A N A K T V E G V W T Y K D E T L T A T V T E
BINARY_1436 T T Y K L I L N L K Q A K E E A I T E L V D A A T A E K Y I K L I A N A K T V E G V W T Y K D E T L T F T V T E
BINARY_1437 T T Y K L I L N L K Q A K E E A I T E L V D A A T A E K Y I K L I A N A K T V E G V W T Y K D E T K K A T V T E
BINARY_1438 T T Y K L I L N L K Q A K E E A I T E L V D A A T A E K Y I K L I A N A K T V E G V W T Y K D E T K K F T V T E
BINARY_1439 T T Y K L I L N L K Q A K E E A I T E L V D A A T A E K Y I K L I A N A K T V E G V W T Y K D E T K T A T V T E
BINARY_1440 T T Y K L I L N L K Q A K E E A I T E L V D A A T A E K Y I K L I A N A K T V E G V W T Y K D E T K T F T V T E
BINARY_1441 T T Y K L I L N L K Q A K E E A I T E L V D A A T A E K Y I K L Y A N A K T V E G V W T L K D E I L K A T V T E
```

BINARY_1442 T T Y K L I L N L K Q A K E E A I T E L V D A A T A E K Y I K L Y A N A K T V E G V W T L K D E I L K F T V T E
BINARY_1443 T T Y K L I L N L K Q A K E E A I T E L V D A A T A E K Y I K L Y A N A K T V E G V W T L K D E I L T A T V T E
BINARY_1444 T T Y K L I L N L K Q A K E E A I T E L V D A A T A E K Y I K L Y A N A K T V E G V W T L K D E I L T F T V T E
BINARY_1445 T T Y K L I L N L K Q A K E E A I T E L V D A A T A E K Y I K L Y A N A K T V E G V W T L K D E I K K A T V T E
BINARY_1446 T T Y K L I L N L K Q A K E E A I T E L V D A A T A E K Y I K L Y A N A K T V E G V W T L K D E I K K F T V T E
BINARY_1447 T T Y K L I L N L K Q A K E E A I T E L V D A A T A E K Y I K L Y A N A K T V E G V W T L K D E I K T A T V T E
BINARY_1448 T T Y K L I L N L K Q A K E E A I T E L V D A A T A E K Y I K L Y A N A K T V E G V W T L K D E I K T F T V T E
BINARY_1449 T T Y K L I L N L K Q A K E E A I T E L V D A A T A E K Y I K L Y A N A K T V E G V W T L K D E T L K A T V T E
BINARY_1450 T T Y K L I L N L K Q A K E E A I T E L V D A A T A E K Y I K L Y A N A K T V E G V W T L K D E T L K F T V T E
BINARY_1451 T T Y K L I L N L K Q A K E E A I T E L V D A A T A E K Y I K L Y A N A K T V E G V W T L K D E T L T A T V T E
BINARY_1452 T T Y K L I L N L K Q A K E E A I T E L V D A A T A E K Y I K L Y A N A K T V E G V W T L K D E T L T F T V T E
BINARY_1453 T T Y K L I L N L K Q A K E E A I T E L V D A A T A E K Y I K L Y A N A K T V E G V W T L K D E T K K A T V T E
BINARY_1454 T T Y K L I L N L K Q A K E E A I T E L V D A A T A E K Y I K L Y A N A K T V E G V W T L K D E T K K F T V T E
BINARY_1455 T T Y K L I L N L K Q A K E E A I T E L V D A A T A E K Y I K L Y A N A K T V E G V W T L K D E T K T A T V T E
BINARY_1456 T T Y K L I L N L K Q A K E E A I T E L V D A A T A E K Y I K L Y A N A K T V E G V W T L K D E T K T F T V T E
BINARY_1457 T T Y K L I L N L K Q A K E E A I T E L V D A A T A E K Y I K L Y A N A K T V E G V W T Y K D E I L K A T V T E
BINARY_1458 T T Y K L I L N L K Q A K E E A I T E L V D A A T A E K Y I K L Y A N A K T V E G V W T Y K D E I L K F T V T E
BINARY_1459 T T Y K L I L N L K Q A K E E A I T E L V D A A T A E K Y I K L Y A N A K T V E G V W T Y K D E I L T A T V T E
BINARY_1460 T T Y K L I L N L K Q A K E E A I T E L V D A A T A E K Y I K L Y A N A K T V E G V W T Y K D E I L T F T V T E
BINARY_1461 T T Y K L I L N L K Q A K E E A I T E L V D A A T A E K Y I K L Y A N A K T V E G V W T Y K D E I K K A T V T E
BINARY_1462 T T Y K L I L N L K Q A K E E A I T E L V D A A T A E K Y I K L Y A N A K T V E G V W T Y K D E I K K F T V T E
BINARY_1463 T T Y K L I L N L K Q A K E E A I T E L V D A A T A E K Y I K L Y A N A K T V E G V W T Y K D E I K T A T V T E
BINARY_1464 T T Y K L I L N L K Q A K E E A I T E L V D A A T A E K Y I K L Y A N A K T V E G V W T Y K D E I K T F T V T E
BINARY_1465 T T Y K L I L N L K Q A K E E A I T E L V D A A T A E K Y I K L Y A N A K T V E G V W T Y K D E T L K A T V T E
BINARY_1466 T T Y K L I L N L K Q A K E E A I T E L V D A A T A E K Y I K L Y A N A K T V E G V W T Y K D E T L K F T V T E
BINARY_1467 T T Y K L I L N L K Q A K E E A I T E L V D A A T A E K Y I K L Y A N A K T V E G V W T Y K D E T L T A T V T E
BINARY_1468 T T Y K L I L N L K Q A K E E A I T E L V D A A T A E K Y I K L Y A N A K T V E G V W T Y K D E T L T F T V T E
BINARY_1469 T T Y K L I L N L K Q A K E E A I T E L V D A A T A E K Y I K L Y A N A K T V E G V W T Y K D E T K K A T V T E
BINARY_1470 T T Y K L I L N L K Q A K E E A I T E L V D A A T A E K Y I K L Y A N A K T V E G V W T Y K D E T K K F T V T E
BINARY_1471 T T Y K L I L N L K Q A K E E A I T E L V D A A T A E K Y I K L Y A N A K T V E G V W T Y K D E T K T A T V T E
BINARY_1472 T T Y K L I L N L K Q A K E E A I T E L V D A A T A E K Y I K L Y A N A K T V E G V W T Y K D E T K T F T V T E
BINARY_1473 T T Y K L I L N L K Q A K E E A I T E L V D A A T A E K Y F K L I A N A K T V E G V W T L K D E I L K A T V T E
BINARY_1474 T T Y K L I L N L K Q A K E E A I T E L V D A A T A E K Y F K L I A N A K T V E G V W T L K D E I L K F T V T E
BINARY_1475 T T Y K L I L N L K Q A K E E A I T E L V D A A T A E K Y F K L I A N A K T V E G V W T L K D E I L T A T V T E
BINARY_1476 T T Y K L I L N L K Q A K E E A I T E L V D A A T A E K Y F K L I A N A K T V E G V W T L K D E I L T F T V T E
BINARY_1477 T T Y K L I L N L K Q A K E E A I T E L V D A A T A E K Y F K L I A N A K T V E G V W T L K D E I K K A T V T E
BINARY_1478 T T Y K L I L N L K Q A K E E A I T E L V D A A T A E K Y F K L I A N A K T V E G V W T L K D E I K K F T V T E
BINARY_1479 T T Y K L I L N L K Q A K E E A I T E L V D A A T A E K Y F K L I A N A K T V E G V W T L K D E I K T A T V T E
BINARY_1480 T T Y K L I L N L K Q A K E E A I T E L V D A A T A E K Y F K L I A N A K T V E G V W T L K D E I K T F T V T E
BINARY_1481 T T Y K L I L N L K Q A K E E A I T E L V D A A T A E K Y F K L I A N A K T V E G V W T L K D E T L K A T V T E
BINARY_1482 T T Y K L I L N L K Q A K E E A I T E L V D A A T A E K Y F K L I A N A K T V E G V W T L K D E T L K F T V T E
BINARY_1483 T T Y K L I L N L K Q A K E E A I T E L V D A A T A E K Y F K L I A N A K T V E G V W T L K D E T L T A T V T E
BINARY_1484 T T Y K L I L N L K Q A K E E A I T E L V D A A T A E K Y F K L I A N A K T V E G V W T L K D E T L T F T V T E
BINARY_1485 T T Y K L I L N L K Q A K E E A I T E L V D A A T A E K Y F K L I A N A K T V E G V W T L K D E T K K A T V T E
BINARY_1486 T T Y K L I L N L K Q A K E E A I T E L V D A A T A E K Y F K L I A N A K T V E G V W T L K D E T K K F T V T E
BINARY_1487 T T Y K L I L N L K Q A K E E A I T E L V D A A T A E K Y F K L I A N A K T V E G V W T L K D E T K T A T V T E
BINARY_1488 T T Y K L I L N L K Q A K E E A I T E L V D A A T A E K Y F K L I A N A K T V E G V W T L K D E T K T F T V T E
BINARY_1489 T T Y K L I L N L K Q A K E E A I T E L V D A A T A E K Y F K L I A N A K T V E G V W T Y K D E I L K A T V T E
BINARY_1490 T T Y K L I L N L K Q A K E E A I T E L V D A A T A E K Y F K L I A N A K T V E G V W T Y K D E I L K F T V T E
BINARY_1491 T T Y K L I L N L K Q A K E E A I T E L V D A A T A E K Y F K L I A N A K T V E G V W T Y K D E I L T A T V T E
BINARY_1492 T T Y K L I L N L K Q A K E E A I T E L V D A A T A E K Y F K L I A N A K T V E G V W T Y K D E I L T F T V T E
BINARY_1493 T T Y K L I L N L K Q A K E E A I T E L V D A A T A E K Y F K L I A N A K T V E G V W T Y K D E I K K A T V T E
BINARY_1494 T T Y K L I L N L K Q A K E E A I T E L V D A A T A E K Y F K L I A N A K T V E G V W T Y K D E I K K F T V T E
BINARY_1495 T T Y K L I L N L K Q A K E E A I T E L V D A A T A E K Y F K L I A N A K T V E G V W T Y K D E I K T A T V T E
BINARY_1496 T T Y K L I L N L K Q A K E E A I T E L V D A A T A E K Y F K L I A N A K T V E G V W T Y K D E I K T F T V T E
BINARY_1497 T T Y K L I L N L K Q A K E E A I T E L V D A A T A E K Y F K L I A N A K T V E G V W T Y K D E T L K A T V T E
BINARY_1498 T T Y K L I L N L K Q A K E E A I T E L V D A A T A E K Y F K L I A N A K T V E G V W T Y K D E T L K F T V T E
BINARY_1499 T T Y K L I L N L K Q A K E E A I T E L V D A A T A E K Y F K L I A N A K T V E G V W T Y K D E T L T A T V T E
BINARY_1500 T T Y K L I L N L K Q A K E E A I T E L V D A A T A E K Y F K L I A N A K T V E G V W T Y K D E T L T F T V T E
BINARY_1501 T T Y K L I L N L K Q A K E E A I T E L V D A A T A E K Y F K L I A N A K T V E G V W T Y K D E T K K A T V T E
BINARY_1502 T T Y K L I L N L K Q A K E E A I T E L V D A A T A E K Y F K L I A N A K T V E G V W T Y K D E T K K F T V T E
BINARY_1503 T T Y K L I L N L K Q A K E E A I T E L V D A A T A E K Y F K L I A N A K T V E G V W T Y K D E T K T A T V T E
BINARY_1504 T T Y K L I L N L K Q A K E E A I T E L V D A A T A E K Y F K L I A N A K T V E G V W T Y K D E T K T F T V T E
BINARY_1505 T T Y K L I L N L K Q A K E E A I T E L V D A A T A E K Y F K L Y A N A K T V E G V W T L K D E I L K A T V T E
BINARY_1506 T T Y K L I L N L K Q A K E E A I T E L V D A A T A E K Y F K L Y A N A K T V E G V W T L K D E I L K F T V T E
BINARY_1507 T T Y K L I L N L K Q A K E E A I T E L V D A A T A E K Y F K L Y A N A K T V E G V W T L K D E I L T A T V T E
BINARY_1508 T T Y K L I L N L K Q A K E E A I T E L V D A A T A E K Y F K L Y A N A K T V E G V W T L K D E I L T F T V T E
BINARY_1509 T T Y K L I L N L K Q A K E E A I T E L V D A A T A E K Y F K L Y A N A K T V E G V W T L K D E I K K A T V T E
BINARY_1510 T T Y K L I L N L K Q A K E E A I T E L V D A A T A E K Y F K L Y A N A K T V E G V W T L K D E I K K F T V T E
BINARY_1511 T T Y K L I L N L K Q A K E E A I T E L V D A A T A E K Y F K L Y A N A K T V E G V W T L K D E I K T A T V T E
BINARY_1512 T T Y K L I L N L K Q A K E E A I T E L V D A A T A E K Y F K L Y A N A K T V E G V W T L K D E I K T F T V T E
BINARY_1513 T T Y K L I L N L K Q A K E E A I T E L V D A A T A E K Y F K L Y A N A K T V E G V W T L K D E T L K A T V T E
BINARY_1514 T T Y K L I L N L K Q A K E E A I T E L V D A A T A E K Y F K L Y A N A K T V E G V W T L K D E T L K F T V T E
BINARY_1515 T T Y K L I L N L K Q A K E E A I T E L V D A A T A E K Y F K L Y A N A K T V E G V W T L K D E T L T A T V T E
BINARY_1516 T T Y K L I L N L K Q A K E E A I T E L V D A A T A E K Y F K L Y A N A K T V E G V W T L K D E T L T F T V T E
BINARY_1517 T T Y K L I L N L K Q A K E E A I T E L V D A A T A E K Y F K L Y A N A K T V E G V W T L K D E T K K A T V T E
BINARY_1518 T T Y K L I L N L K Q A K E E A I T E L V D A A T A E K Y F K L Y A N A K T V E G V W T L K D E T K K F T V T E
BINARY_1519 T T Y K L I L N L K Q A K E E A I T E L V D A A T A E K Y F K L Y A N A K T V E G V W T L K D E T K T A T V T E
BINARY_1520 T T Y K L I L N L K Q A K E E A I T E L V D A A T A E K Y F K L Y A N A K T V E G V W T L K D E T K T F T V T E
BINARY_1521 T T Y K L I L N L K Q A K E E A I T E L V D A A T A E K Y F K L Y A N A K T V E G V W T Y K D E I L K A T V T E
BINARY_1522 T T Y K L I L N L K Q A K E E A I T E L V D A A T A E K Y F K L Y A N A K T V E G V W T Y K D E I L K F T V T E
BINARY_1523 T T Y K L I L N L K Q A K E E A I T E L V D A A T A E K Y F K L Y A N A K T V E G V W T Y K D E I L T A T V T E
BINARY_1524 T T Y K L I L N L K Q A K E E A I T E L V D A A T A E K Y F K L Y A N A K T V E G V W T Y K D E I L T F T V T E
BINARY_1525 T T Y K L I L N L K Q A K E E A I T E L V D A A T A E K Y F K L Y A N A K T V E G V W T Y K D E I K K A T V T E
BINARY_1526 T T Y K L I L N L K Q A K E E A I T E L V D A A T A E K Y F K L Y A N A K T V E G V W T Y K D E I K K F T V T E
BINARY_1527 T T Y K L I L N L K Q A K E E A I T E L V D A A T A E K Y F K L Y A N A K T V E G V W T Y K D E I K T A T V T E
BINARY_1528 T T Y K L I L N L K Q A K E E A I T E L V D A A T A E K Y F K L Y A N A K T V E G V W T Y K D E I K T F T V T E
BINARY_1529 T T Y K L I L N L K Q A K E E A I T E L V D A A T A E K Y F K L Y A N A K T V E G V W T Y K D E T L K A T V T E
BINARY_1530 T T Y K L I L N L K Q A K E E A I T E L V D A A T A E K Y F K L Y A N A K T V E G V W T Y K D E T L K F T V T E
BINARY_1531 T T Y K L I L N L K Q A K E E A I T E L V D A A T A E K Y F K L Y A N A K T V E G V W T Y K D E T L T A T V T E
BINARY_1532 T T Y K L I L N L K Q A K E E A I T E L V D A A T A E K Y F K L Y A N A K T V E G V W T Y K D E T L T F T V T E
BINARY_1533 T T Y K L I L N L K Q A K E E A I T E L V D A A T A E K Y F K L Y A N A K T V E G V W T Y K D E T K K A T V T E
BINARY_1534 T T Y K L I L N L K Q A K E E A I T E L V D A A T A E K Y F K L Y A N A K T V E G V W T Y K D E T K K F T V T E
BINARY_1535 T T Y K L I L N L K Q A K E E A I T E L V D A A T A E K Y F K L Y A N A K T V E G V W T Y K D E T K T A T V T E
BINARY_1536 T T Y K L I L N L K Q A K E E A I T E L V D A A T A E K Y F K L Y A N A K T V E G V W T Y K D E T K T F T V T E

BINARY_1537 T T Y K L I L N L K Q A K E E A I T E A V D A G I A E K Y I K L I A N A K T V E G V W T L K D E I L K A T V T E
BINARY_1538 T T Y K L I L N L K Q A K E E A I T E A V D A G I A E K Y I K L I A N A K T V E G V W T L K D E I L K F T V T E
BINARY_1539 T T Y K L I L N L K Q A K E E A I T E A V D A G I A E K Y I K L I A N A K T V E G V W T L K D E I L T A T V T E
BINARY_1540 T T Y K L I L N L K Q A K E E A I T E A V D A G I A E K Y I K L I A N A K T V E G V W T L K D E I L T F T V T E
BINARY_1541 T T Y K L I L N L K Q A K E E A I T E A V D A G I A E K Y I K L I A N A K T V E G V W T L K D E I K K A T V T E
BINARY_1542 T T Y K L I L N L K Q A K E E A I T E A V D A G I A E K Y I K L I A N A K T V E G V W T L K D E I K K F T V T E
BINARY_1543 T T Y K L I L N L K Q A K E E A I T E A V D A G I A E K Y I K L I A N A K T V E G V W T L K D E I K T A T V T E
BINARY_1544 T T Y K L I L N L K Q A K E E A I T E A V D A G I A E K Y I K L I A N A K T V E G V W T L K D E I K T F T V T E
BINARY_1545 T T Y K L I L N L K Q A K E E A I T E A V D A G I A E K Y I K L I A N A K T V E G V W T L K D E T L K A T V T E
BINARY_1546 T T Y K L I L N L K Q A K E E A I T E A V D A G I A E K Y I K L I A N A K T V E G V W T L K D E T L K F T V T E
BINARY_1547 T T Y K L I L N L K Q A K E E A I T E A V D A G I A E K Y I K L I A N A K T V E G V W T L K D E T L T A T V T E
BINARY_1548 T T Y K L I L N L K Q A K E E A I T E A V D A G I A E K Y I K L I A N A K T V E G V W T L K D E T L T F T V T E
BINARY_1549 T T Y K L I L N L K Q A K E E A I T E A V D A G I A E K Y I K L I A N A K T V E G V W T L K D E T K K A T V T E
BINARY_1550 T T Y K L I L N L K Q A K E E A I T E A V D A G I A E K Y I K L I A N A K T V E G V W T L K D E T K K F T V T E
BINARY_1551 T T Y K L I L N L K Q A K E E A I T E A V D A G I A E K Y I K L I A N A K T V E G V W T L K D E T K T A T V T E
BINARY_1552 T T Y K L I L N L K Q A K E E A I T E A V D A G I A E K Y I K L I A N A K T V E G V W T L K D E T K T F T V T E
BINARY_1553 T T Y K L I L N L K Q A K E E A I T E A V D A G I A E K Y I K L I A N A K T V E G V W T Y K D E I L K A T V T E
BINARY_1554 T T Y K L I L N L K Q A K E E A I T E A V D A G I A E K Y I K L I A N A K T V E G V W T Y K D E I L K F T V T E
BINARY_1555 T T Y K L I L N L K Q A K E E A I T E A V D A G I A E K Y I K L I A N A K T V E G V W T Y K D E I L T A T V T E
BINARY_1556 T T Y K L I L N L K Q A K E E A I T E A V D A G I A E K Y I K L I A N A K T V E G V W T Y K D E I L T F T V T E
BINARY_1557 T T Y K L I L N L K Q A K E E A I T E A V D A G I A E K Y I K L I A N A K T V E G V W T Y K D E I K K A T V T E
BINARY_1558 T T Y K L I L N L K Q A K E E A I T E A V D A G I A E K Y I K L I A N A K T V E G V W T Y K D E I K K F T V T E
BINARY_1559 T T Y K L I L N L K Q A K E E A I T E A V D A G I A E K Y I K L I A N A K T V E G V W T Y K D E I K T A T V T E
BINARY_1560 T T Y K L I L N L K Q A K E E A I T E A V D A G I A E K Y I K L I A N A K T V E G V W T Y K D E I K T F T V T E
BINARY_1561 T T Y K L I L N L K Q A K E E A I T E A V D A G I A E K Y I K L I A N A K T V E G V W T Y K D E T L K A T V T E
BINARY_1562 T T Y K L I L N L K Q A K E E A I T E A V D A G I A E K Y I K L I A N A K T V E G V W T Y K D E T L K F T V T E
BINARY_1563 T T Y K L I L N L K Q A K E E A I T E A V D A G I A E K Y I K L I A N A K T V E G V W T Y K D E T L T A T V T E
BINARY_1564 T T Y K L I L N L K Q A K E E A I T E A V D A G I A E K Y I K L I A N A K T V E G V W T Y K D E T L T F T V T E
BINARY_1565 T T Y K L I L N L K Q A K E E A I T E A V D A G I A E K Y I K L I A N A K T V E G V W T Y K D E T K K A T V T E
BINARY_1566 T T Y K L I L N L K Q A K E E A I T E A V D A G I A E K Y I K L I A N A K T V E G V W T Y K D E T K K F T V T E
BINARY_1567 T T Y K L I L N L K Q A K E E A I T E A V D A G I A E K Y I K L I A N A K T V E G V W T Y K D E T K T A T V T E
BINARY_1568 T T Y K L I L N L K Q A K E E A I T E A V D A G I A E K Y I K L I A N A K T V E G V W T Y K D E T K T F T V T E
BINARY_1569 T T Y K L I L N L K Q A K E E A I T E A V D A G I A E K Y I K L Y A N A K T V E G V W T L K D E I L K A T V T E
BINARY_1570 T T Y K L I L N L K Q A K E E A I T E A V D A G I A E K Y I K L Y A N A K T V E G V W T L K D E I L K F T V T E
BINARY_1571 T T Y K L I L N L K Q A K E E A I T E A V D A G I A E K Y I K L Y A N A K T V E G V W T L K D E I L T A T V T E
BINARY_1572 T T Y K L I L N L K Q A K E E A I T E A V D A G I A E K Y I K L Y A N A K T V E G V W T L K D E I L T F T V T E
BINARY_1573 T T Y K L I L N L K Q A K E E A I T E A V D A G I A E K Y I K L Y A N A K T V E G V W T L K D E I K K A T V T E
BINARY_1574 T T Y K L I L N L K Q A K E E A I T E A V D A G I A E K Y I K L Y A N A K T V E G V W T L K D E I K K F T V T E
BINARY_1575 T T Y K L I L N L K Q A K E E A I T E A V D A G I A E K Y I K L Y A N A K T V E G V W T L K D E I K T A T V T E
BINARY_1576 T T Y K L I L N L K Q A K E E A I T E A V D A G I A E K Y I K L Y A N A K T V E G V W T L K D E I K T F T V T E
BINARY_1577 T T Y K L I L N L K Q A K E E A I T E A V D A G I A E K Y I K L Y A N A K T V E G V W T L K D E T L K A T V T E
BINARY_1578 T T Y K L I L N L K Q A K E E A I T E A V D A G I A E K Y I K L Y A N A K T V E G V W T L K D E T L K F T V T E
BINARY_1579 T T Y K L I L N L K Q A K E E A I T E A V D A G I A E K Y I K L Y A N A K T V E G V W T L K D E T L T A T V T E
BINARY_1580 T T Y K L I L N L K Q A K E E A I T E A V D A G I A E K Y I K L Y A N A K T V E G V W T L K D E T L T F T V T E
BINARY_1581 T T Y K L I L N L K Q A K E E A I T E A V D A G I A E K Y I K L Y A N A K T V E G V W T L K D E T K K A T V T E
BINARY_1582 T T Y K L I L N L K Q A K E E A I T E A V D A G I A E K Y I K L Y A N A K T V E G V W T L K D E T K K F T V T E
BINARY_1583 T T Y K L I L N L K Q A K E E A I T E A V D A G I A E K Y I K L Y A N A K T V E G V W T L K D E T K T A T V T E
BINARY_1584 T T Y K L I L N L K Q A K E E A I T E A V D A G I A E K Y I K L Y A N A K T V E G V W T L K D E T K T F T V T E
BINARY_1585 T T Y K L I L N L K Q A K E E A I T E A V D A G I A E K Y I K L Y A N A K T V E G V W T Y K D E I L K A T V T E
BINARY_1586 T T Y K L I L N L K Q A K E E A I T E A V D A G I A E K Y I K L Y A N A K T V E G V W T Y K D E I L K F T V T E
BINARY_1587 T T Y K L I L N L K Q A K E E A I T E A V D A G I A E K Y I K L Y A N A K T V E G V W T Y K D E I L T A T V T E
BINARY_1588 T T Y K L I L N L K Q A K E E A I T E A V D A G I A E K Y I K L Y A N A K T V E G V W T Y K D E I L T F T V T E
BINARY_1589 T T Y K L I L N L K Q A K E E A I T E A V D A G I A E K Y I K L Y A N A K T V E G V W T Y K D E I K K A T V T E
BINARY_1590 T T Y K L I L N L K Q A K E E A I T E A V D A G I A E K Y I K L Y A N A K T V E G V W T Y K D E I K K F T V T E
BINARY_1591 T T Y K L I L N L K Q A K E E A I T E A V D A G I A E K Y I K L Y A N A K T V E G V W T Y K D E I K T A T V T E
BINARY_1592 T T Y K L I L N L K Q A K E E A I T E A V D A G I A E K Y I K L Y A N A K T V E G V W T Y K D E I K T F T V T E
BINARY_1593 T T Y K L I L N L K Q A K E E A I T E A V D A G I A E K Y I K L Y A N A K T V E G V W T Y K D E T L K A T V T E
BINARY_1594 T T Y K L I L N L K Q A K E E A I T E A V D A G I A E K Y I K L Y A N A K T V E G V W T Y K D E T L K F T V T E
BINARY_1595 T T Y K L I L N L K Q A K E E A I T E A V D A G I A E K Y I K L Y A N A K T V E G V W T Y K D E T L T A T V T E
BINARY_1596 T T Y K L I L N L K Q A K E E A I T E A V D A G I A E K Y I K L Y A N A K T V E G V W T Y K D E T L T F T V T E
BINARY_1597 T T Y K L I L N L K Q A K E E A I T E A V D A G I A E K Y I K L Y A N A K T V E G V W T Y K D E T K K A T V T E
BINARY_1598 T T Y K L I L N L K Q A K E E A I T E A V D A G I A E K Y I K L Y A N A K T V E G V W T Y K D E T K K F T V T E
BINARY_1599 T T Y K L I L N L K Q A K E E A I T E A V D A G I A E K Y I K L Y A N A K T V E G V W T Y K D E T K T A T V T E
BINARY_1600 T T Y K L I L N L K Q A K E E A I T E A V D A G I A E K Y I K L Y A N A K T V E G V W T Y K D E T K T F T V T E
BINARY_1601 T T Y K L I L N L K Q A K E E A I T E A V D A G I A E K Y F K L I A N A K T V E G V W T L K D E I L K A T V T E
BINARY_1602 T T Y K L I L N L K Q A K E E A I T E A V D A G I A E K Y F K L I A N A K T V E G V W T L K D E I L K F T V T E
BINARY_1603 T T Y K L I L N L K Q A K E E A I T E A V D A G I A E K Y F K L I A N A K T V E G V W T L K D E I L T A T V T E
BINARY_1604 T T Y K L I L N L K Q A K E E A I T E A V D A G I A E K Y F K L I A N A K T V E G V W T L K D E I L T F T V T E
BINARY_1605 T T Y K L I L N L K Q A K E E A I T E A V D A G I A E K Y F K L I A N A K T V E G V W T L K D E I K K A T V T E
BINARY_1606 T T Y K L I L N L K Q A K E E A I T E A V D A G I A E K Y F K L I A N A K T V E G V W T L K D E I K K F T V T E
BINARY_1607 T T Y K L I L N L K Q A K E E A I T E A V D A G I A E K Y F K L I A N A K T V E G V W T L K D E I K T A T V T E
BINARY_1608 T T Y K L I L N L K Q A K E E A I T E A V D A G I A E K Y F K L I A N A K T V E G V W T L K D E I K T F T V T E
BINARY_1609 T T Y K L I L N L K Q A K E E A I T E A V D A G I A E K Y F K L I A N A K T V E G V W T L K D E T L K A T V T E
BINARY_1610 T T Y K L I L N L K Q A K E E A I T E A V D A G I A E K Y F K L I A N A K T V E G V W T L K D E T L K F T V T E
BINARY_1611 T T Y K L I L N L K Q A K E E A I T E A V D A G I A E K Y F K L I A N A K T V E G V W T L K D E T L T A T V T E
BINARY_1612 T T Y K L I L N L K Q A K E E A I T E A V D A G I A E K Y F K L I A N A K T V E G V W T L K D E T L T F T V T E
BINARY_1613 T T Y K L I L N L K Q A K E E A I T E A V D A G I A E K Y F K L I A N A K T V E G V W T L K D E T K K A T V T E
BINARY_1614 T T Y K L I L N L K Q A K E E A I T E A V D A G I A E K Y F K L I A N A K T V E G V W T L K D E T K K F T V T E
BINARY_1615 T T Y K L I L N L K Q A K E E A I T E A V D A G I A E K Y F K L I A N A K T V E G V W T L K D E T K T A T V T E
BINARY_1616 T T Y K L I L N L K Q A K E E A I T E A V D A G I A E K Y F K L I A N A K T V E G V W T L K D E T K T F T V T E
BINARY_1617 T T Y K L I L N L K Q A K E E A I T E A V D A G I A E K Y F K L I A N A K T V E G V W T Y K D E I L K A T V T E
BINARY_1618 T T Y K L I L N L K Q A K E E A I T E A V D A G I A E K Y F K L I A N A K T V E G V W T Y K D E I L K F T V T E
BINARY_1619 T T Y K L I L N L K Q A K E E A I T E A V D A G I A E K Y F K L I A N A K T V E G V W T Y K D E I L T A T V T E
BINARY_1620 T T Y K L I L N L K Q A K E E A I T E A V D A G I A E K Y F K L I A N A K T V E G V W T Y K D E I L T F T V T E
BINARY_1621 T T Y K L I L N L K Q A K E E A I T E A V D A G I A E K Y F K L I A N A K T V E G V W T Y K D E I K K A T V T E
BINARY_1622 T T Y K L I L N L K Q A K E E A I T E A V D A G I A E K Y F K L I A N A K T V E G V W T Y K D E I K K F T V T E
BINARY_1623 T T Y K L I L N L K Q A K E E A I T E A V D A G I A E K Y F K L I A N A K T V E G V W T Y K D E I K T A T V T E
BINARY_1624 T T Y K L I L N L K Q A K E E A I T E A V D A G I A E K Y F K L I A N A K T V E G V W T Y K D E I K T F T V T E
BINARY_1625 T T Y K L I L N L K Q A K E E A I T E A V D A G I A E K Y F K L I A N A K T V E G V W T Y K D E T L K A T V T E
BINARY_1626 T T Y K L I L N L K Q A K E E A I T E A V D A G I A E K Y F K L I A N A K T V E G V W T Y K D E T L K F T V T E
BINARY_1627 T T Y K L I L N L K Q A K E E A I T E A V D A G I A E K Y F K L I A N A K T V E G V W T Y K D E T L T A T V T E
BINARY_1628 T T Y K L I L N L K Q A K E E A I T E A V D A G I A E K Y F K L I A N A K T V E G V W T Y K D E T L T F T V T E
BINARY_1629 T T Y K L I L N L K Q A K E E A I T E A V D A G I A E K Y F K L I A N A K T V E G V W T Y K D E T K K A T V T E
BINARY_1630 T T Y K L I L N L K Q A K E E A I T E A V D A G I A E K Y F K L I A N A K T V E G V W T Y K D E T K K F T V T E
BINARY_1631 T T Y K L I L N L K Q A K E E A I T E A V D A G I A E K Y F K L I A N A K T V E G V W T Y K D E T K T A T V T E

BINARY_1632 T T Y K L I L N L K Q A K E E A I T E A V D A G I A E K Y F K L I A N A K T V E G V W T Y K D E T K T F T V T E
BINARY_1633 T T Y K L I L N L K Q A K E E A I T E A V D A G I A E K Y F K L Y A N A K T V E G V W T L K D E I L K A T V T E
BINARY_1634 T T Y K L I L N L K Q A K E E A I T E A V D A G I A E K Y F K L Y A N A K T V E G V W T L K D E I L K F T V T E
BINARY_1635 T T Y K L I L N L K Q A K E E A I T E A V D A G I A E K Y F K L Y A N A K T V E G V W T L K D E I L T A T V T E
BINARY_1636 T T Y K L I L N L K Q A K E E A I T E A V D A G I A E K Y F K L Y A N A K T V E G V W T L K D E I L T F T V T E
BINARY_1637 T T Y K L I L N L K Q A K E E A I T E A V D A G I A E K Y F K L Y A N A K T V E G V W T L K D E I K K A T V T E
BINARY_1638 T T Y K L I L N L K Q A K E E A I T E A V D A G I A E K Y F K L Y A N A K T V E G V W T L K D E I K K F T V T E
BINARY_1639 T T Y K L I L N L K Q A K E E A I T E A V D A G I A E K Y F K L Y A N A K T V E G V W T L K D E I K T A T V T E
BINARY_1640 T T Y K L I L N L K Q A K E E A I T E A V D A G I A E K Y F K L Y A N A K T V E G V W T L K D E I K T F T V T E
BINARY_1641 T T Y K L I L N L K Q A K E E A I T E A V D A G I A E K Y F K L Y A N A K T V E G V W T L K D E T L K A T V T E
BINARY_1642 T T Y K L I L N L K Q A K E E A I T E A V D A G I A E K Y F K L Y A N A K T V E G V W T L K D E T L K F T V T E
BINARY_1643 T T Y K L I L N L K Q A K E E A I T E A V D A G I A E K Y F K L Y A N A K T V E G V W T L K D E T L T A T V T E
BINARY_1644 T T Y K L I L N L K Q A K E E A I T E A V D A G I A E K Y F K L Y A N A K T V E G V W T L K D E T L T F T V T E
BINARY_1645 T T Y K L I L N L K Q A K E E A I T E A V D A G I A E K Y F K L Y A N A K T V E G V W T L K D E T K K A T V T E
BINARY_1646 T T Y K L I L N L K Q A K E E A I T E A V D A G I A E K Y F K L Y A N A K T V E G V W T L K D E T K K F T V T E
BINARY_1647 T T Y K L I L N L K Q A K E E A I T E A V D A G I A E K Y F K L Y A N A K T V E G V W T L K D E T K T A T V T E
BINARY_1648 T T Y K L I L N L K Q A K E E A I T E A V D A G I A E K Y F K L Y A N A K T V E G V W T L K D E T K T F T V T E
BINARY_1649 T T Y K L I L N L K Q A K E E A I T E A V D A G I A E K Y F K L Y A N A K T V E G V W T Y K D E I L K A T V T E
BINARY_1650 T T Y K L I L N L K Q A K E E A I T E A V D A G I A E K Y F K L Y A N A K T V E G V W T Y K D E I L K F T V T E
BINARY_1651 T T Y K L I L N L K Q A K E E A I T E A V D A G I A E K Y F K L Y A N A K T V E G V W T Y K D E I L T A T V T E
BINARY_1652 T T Y K L I L N L K Q A K E E A I T E A V D A G I A E K Y F K L Y A N A K T V E G V W T Y K D E I L T F T V T E
BINARY_1653 T T Y K L I L N L K Q A K E E A I T E A V D A G I A E K Y F K L Y A N A K T V E G V W T Y K D E I K K A T V T E
BINARY_1654 T T Y K L I L N L K Q A K E E A I T E A V D A G I A E K Y F K L Y A N A K T V E G V W T Y K D E I K K F T V T E
BINARY_1655 T T Y K L I L N L K Q A K E E A I T E A V D A G I A E K Y F K L Y A N A K T V E G V W T Y K D E I K T A T V T E
BINARY_1656 T T Y K L I L N L K Q A K E E A I T E A V D A G I A E K Y F K L Y A N A K T V E G V W T Y K D E I K T F T V T E
BINARY_1657 T T Y K L I L N L K Q A K E E A I T E A V D A G I A E K Y F K L Y A N A K T V E G V W T Y K D E T L K A T V T E
BINARY_1658 T T Y K L I L N L K Q A K E E A I T E A V D A G I A E K Y F K L Y A N A K T V E G V W T Y K D E T L K F T V T E
BINARY_1659 T T Y K L I L N L K Q A K E E A I T E A V D A G I A E K Y F K L Y A N A K T V E G V W T Y K D E T L T A T V T E
BINARY_1660 T T Y K L I L N L K Q A K E E A I T E A V D A G I A E K Y F K L Y A N A K T V E G V W T Y K D E T L T F T V T E
BINARY_1661 T T Y K L I L N L K Q A K E E A I T E A V D A G I A E K Y F K L Y A N A K T V E G V W T Y K D E T K K A T V T E
BINARY_1662 T T Y K L I L N L K Q A K E E A I T E A V D A G I A E K Y F K L Y A N A K T V E G V W T Y K D E T K K F T V T E
BINARY_1663 T T Y K L I L N L K Q A K E E A I T E A V D A G I A E K Y F K L Y A N A K T V E G V W T Y K D E T K T A T V T E
BINARY_1664 T T Y K L I L N L K Q A K E E A I T E A V D A G I A E K Y F K L Y A N A K T V E G V W T Y K D E T K T F T V T E
BINARY_1665 T T Y K L I L N L K Q A K E E A I T E A V D A G T A E K Y I K L I A N A K T V E G V W T L K D E I L K A T V T E
BINARY_1666 T T Y K L I L N L K Q A K E E A I T E A V D A G T A E K Y I K L I A N A K T V E G V W T L K D E I L K F T V T E
BINARY_1667 T T Y K L I L N L K Q A K E E A I T E A V D A G T A E K Y I K L I A N A K T V E G V W T L K D E I L T A T V T E
BINARY_1668 T T Y K L I L N L K Q A K E E A I T E A V D A G T A E K Y I K L I A N A K T V E G V W T L K D E I L T F T V T E
BINARY_1669 T T Y K L I L N L K Q A K E E A I T E A V D A G T A E K Y I K L I A N A K T V E G V W T L K D E I K K A T V T E
BINARY_1670 T T Y K L I L N L K Q A K E E A I T E A V D A G T A E K Y I K L I A N A K T V E G V W T L K D E I K K F T V T E
BINARY_1671 T T Y K L I L N L K Q A K E E A I T E A V D A G T A E K Y I K L I A N A K T V E G V W T L K D E I K T A T V T E
BINARY_1672 T T Y K L I L N L K Q A K E E A I T E A V D A G T A E K Y I K L I A N A K T V E G V W T L K D E I K T F T V T E
BINARY_1673 T T Y K L I L N L K Q A K E E A I T E A V D A G T A E K Y I K L I A N A K T V E G V W T L K D E T L K A T V T E
BINARY_1674 T T Y K L I L N L K Q A K E E A I T E A V D A G T A E K Y I K L I A N A K T V E G V W T L K D E T L K F T V T E
BINARY_1675 T T Y K L I L N L K Q A K E E A I T E A V D A G T A E K Y I K L I A N A K T V E G V W T L K D E T L T A T V T E
BINARY_1676 T T Y K L I L N L K Q A K E E A I T E A V D A G T A E K Y I K L I A N A K T V E G V W T L K D E T L T F T V T E
BINARY_1677 T T Y K L I L N L K Q A K E E A I T E A V D A G T A E K Y I K L I A N A K T V E G V W T L K D E T K K A T V T E
BINARY_1678 T T Y K L I L N L K Q A K E E A I T E A V D A G T A E K Y I K L I A N A K T V E G V W T L K D E T K K F T V T E
BINARY_1679 T T Y K L I L N L K Q A K E E A I T E A V D A G T A E K Y I K L I A N A K T V E G V W T L K D E T K T A T V T E
BINARY_1680 T T Y K L I L N L K Q A K E E A I T E A V D A G T A E K Y I K L I A N A K T V E G V W T L K D E T K T F T V T E
BINARY_1681 T T Y K L I L N L K Q A K E E A I T E A V D A G T A E K Y I K L I A N A K T V E G V W T Y K D E I L K A T V T E
BINARY_1682 T T Y K L I L N L K Q A K E E A I T E A V D A G T A E K Y I K L I A N A K T V E G V W T Y K D E I L K F T V T E
BINARY_1683 T T Y K L I L N L K Q A K E E A I T E A V D A G T A E K Y I K L I A N A K T V E G V W T Y K D E I L T A T V T E
BINARY_1684 T T Y K L I L N L K Q A K E E A I T E A V D A G T A E K Y I K L I A N A K T V E G V W T Y K D E I L T F T V T E
BINARY_1685 T T Y K L I L N L K Q A K E E A I T E A V D A G T A E K Y I K L I A N A K T V E G V W T Y K D E I K K A T V T E
BINARY_1686 T T Y K L I L N L K Q A K E E A I T E A V D A G T A E K Y I K L I A N A K T V E G V W T Y K D E I K K F T V T E
BINARY_1687 T T Y K L I L N L K Q A K E E A I T E A V D A G T A E K Y I K L I A N A K T V E G V W T Y K D E I K T A T V T E
BINARY_1688 T T Y K L I L N L K Q A K E E A I T E A V D A G T A E K Y I K L I A N A K T V E G V W T Y K D E I K T F T V T E
BINARY_1689 T T Y K L I L N L K Q A K E E A I T E A V D A G T A E K Y I K L I A N A K T V E G V W T Y K D E T L K A T V T E
BINARY_1690 T T Y K L I L N L K Q A K E E A I T E A V D A G T A E K Y I K L I A N A K T V E G V W T Y K D E T L K F T V T E
BINARY_1691 T T Y K L I L N L K Q A K E E A I T E A V D A G T A E K Y I K L I A N A K T V E G V W T Y K D E T L T A T V T E
BINARY_1692 T T Y K L I L N L K Q A K E E A I T E A V D A G T A E K Y I K L I A N A K T V E G V W T Y K D E T L T F T V T E
BINARY_1693 T T Y K L I L N L K Q A K E E A I T E A V D A G T A E K Y I K L I A N A K T V E G V W T Y K D E T K K A T V T E
BINARY_1694 T T Y K L I L N L K Q A K E E A I T E A V D A G T A E K Y I K L I A N A K T V E G V W T Y K D E T K K F T V T E
BINARY_1695 T T Y K L I L N L K Q A K E E A I T E A V D A G T A E K Y I K L I A N A K T V E G V W T Y K D E T K T A T V T E
BINARY_1696 T T Y K L I L N L K Q A K E E A I T E A V D A G T A E K Y I K L I A N A K T V E G V W T Y K D E T K T F T V T E
BINARY_1697 T T Y K L I L N L K Q A K E E A I T E A V D A G T A E K Y I K L Y A N A K T V E G V W T L K D E I L K A T V T E
BINARY_1698 T T Y K L I L N L K Q A K E E A I T E A V D A G T A E K Y I K L Y A N A K T V E G V W T L K D E I L K F T V T E
BINARY_1699 T T Y K L I L N L K Q A K E E A I T E A V D A G T A E K Y I K L Y A N A K T V E G V W T L K D E I L T A T V T E
BINARY_1700 T T Y K L I L N L K Q A K E E A I T E A V D A G T A E K Y I K L Y A N A K T V E G V W T L K D E I L T F T V T E
BINARY_1701 T T Y K L I L N L K Q A K E E A I T E A V D A G T A E K Y I K L Y A N A K T V E G V W T L K D E I K K A T V T E
BINARY_1702 T T Y K L I L N L K Q A K E E A I T E A V D A G T A E K Y I K L Y A N A K T V E G V W T L K D E I K K F T V T E
BINARY_1703 T T Y K L I L N L K Q A K E E A I T E A V D A G T A E K Y I K L Y A N A K T V E G V W T L K D E I K T A T V T E
BINARY_1704 T T Y K L I L N L K Q A K E E A I T E A V D A G T A E K Y I K L Y A N A K T V E G V W T L K D E I K T F T V T E
BINARY_1705 T T Y K L I L N L K Q A K E E A I T E A V D A G T A E K Y I K L Y A N A K T V E G V W T L K D E T L K A T V T E
BINARY_1706 T T Y K L I L N L K Q A K E E A I T E A V D A G T A E K Y I K L Y A N A K T V E G V W T L K D E T L K F T V T E
BINARY_1707 T T Y K L I L N L K Q A K E E A I T E A V D A G T A E K Y I K L Y A N A K T V E G V W T L K D E T L T A T V T E
BINARY_1708 T T Y K L I L N L K Q A K E E A I T E A V D A G T A E K Y I K L Y A N A K T V E G V W T L K D E T L T F T V T E
BINARY_1709 T T Y K L I L N L K Q A K E E A I T E A V D A G T A E K Y I K L Y A N A K T V E G V W T L K D E T K K A T V T E
BINARY_1710 T T Y K L I L N L K Q A K E E A I T E A V D A G T A E K Y I K L Y A N A K T V E G V W T L K D E T K K F T V T E
BINARY_1711 T T Y K L I L N L K Q A K E E A I T E A V D A G T A E K Y I K L Y A N A K T V E G V W T L K D E T K T A T V T E
BINARY_1712 T T Y K L I L N L K Q A K E E A I T E A V D A G T A E K Y I K L Y A N A K T V E G V W T L K D E T K T F T V T E
BINARY_1713 T T Y K L I L N L K Q A K E E A I T E A V D A G T A E K Y I K L Y A N A K T V E G V W T Y K D E I L K A T V T E
BINARY_1714 T T Y K L I L N L K Q A K E E A I T E A V D A G T A E K Y I K L Y A N A K T V E G V W T Y K D E I L K F T V T E
BINARY_1715 T T Y K L I L N L K Q A K E E A I T E A V D A G T A E K Y I K L Y A N A K T V E G V W T Y K D E I L T A T V T E
BINARY_1716 T T Y K L I L N L K Q A K E E A I T E A V D A G T A E K Y I K L Y A N A K T V E G V W T Y K D E I L T F T V T E
BINARY_1717 T T Y K L I L N L K Q A K E E A I T E A V D A G T A E K Y I K L Y A N A K T V E G V W T Y K D E I K K A T V T E
BINARY_1718 T T Y K L I L N L K Q A K E E A I T E A V D A G T A E K Y I K L Y A N A K T V E G V W T Y K D E I K K F T V T E
BINARY_1719 T T Y K L I L N L K Q A K E E A I T E A V D A G T A E K Y I K L Y A N A K T V E G V W T Y K D E I K T A T V T E
BINARY_1720 T T Y K L I L N L K Q A K E E A I T E A V D A G T A E K Y I K L Y A N A K T V E G V W T Y K D E I K T F T V T E
BINARY_1721 T T Y K L I L N L K Q A K E E A I T E A V D A G T A E K Y I K L Y A N A K T V E G V W T Y K D E T L K A T V T E
BINARY_1722 T T Y K L I L N L K Q A K E E A I T E A V D A G T A E K Y I K L Y A N A K T V E G V W T Y K D E T L K F T V T E
BINARY_1723 T T Y K L I L N L K Q A K E E A I T E A V D A G T A E K Y I K L Y A N A K T V E G V W T Y K D E T L T A T V T E
BINARY_1724 T T Y K L I L N L K Q A K E E A I T E A V D A G T A E K Y I K L Y A N A K T V E G V W T Y K D E T L T F T V T E
BINARY_1725 T T Y K L I L N L K Q A K E E A I T E A V D A G T A E K Y I K L Y A N A K T V E G V W T Y K D E T K K A T V T E
BINARY_1726 T T Y K L I L N L K Q A K E E A I T E A V D A G T A E K Y I K L Y A N A K T V E G V W T Y K D E T K K F T V T E

BINARY_1727 T T Y K L I L N L K Q A K E E A I T E A V D A G T A E K Y I K L Y A N A K T V E G V W T Y K D E T K T A T V T E
BINARY_1728 T T Y K L I L N L K Q A K E E A I T E A V D A G T A E K Y I K L Y A N A K T V E G V W T Y K D E T K T F T V T E
BINARY_1729 T T Y K L I L N L K Q A K E E A I T E A V D A G T A E K Y F K L I A N A K T V E G V W T L K D E I L K A T V T E
BINARY_1730 T T Y K L I L N L K Q A K E E A I T E A V D A G T A E K Y F K L I A N A K T V E G V W T L K D E I L K F T V T E
BINARY_1731 T T Y K L I L N L K Q A K E E A I T E A V D A G T A E K Y F K L I A N A K T V E G V W T L K D E I L T A T V T E
BINARY_1732 T T Y K L I L N L K Q A K E E A I T E A V D A G T A E K Y F K L I A N A K T V E G V W T L K D E I L T F T V T E
BINARY_1733 T T Y K L I L N L K Q A K E E A I T E A V D A G T A E K Y F K L I A N A K T V E G V W T L K D E I K K A T V T E
BINARY_1734 T T Y K L I L N L K Q A K E E A I T E A V D A G T A E K Y F K L I A N A K T V E G V W T L K D E I K K F T V T E
BINARY_1735 T T Y K L I L N L K Q A K E E A I T E A V D A G T A E K Y F K L I A N A K T V E G V W T L K D E I K T A T V T E
BINARY_1736 T T Y K L I L N L K Q A K E E A I T E A V D A G T A E K Y F K L I A N A K T V E G V W T L K D E I K T F T V T E
BINARY_1737 T T Y K L I L N L K Q A K E E A I T E A V D A G T A E K Y F K L I A N A K T V E G V W T L K D E T L K A T V T E
BINARY_1738 T T Y K L I L N L K Q A K E E A I T E A V D A G T A E K Y F K L I A N A K T V E G V W T L K D E T L K F T V T E
BINARY_1739 T T Y K L I L N L K Q A K E E A I T E A V D A G T A E K Y F K L I A N A K T V E G V W T L K D E T L T A T V T E
BINARY_1740 T T Y K L I L N L K Q A K E E A I T E A V D A G T A E K Y F K L I A N A K T V E G V W T L K D E T L T F T V T E
BINARY_1741 T T Y K L I L N L K Q A K E E A I T E A V D A G T A E K Y F K L I A N A K T V E G V W T L K D E T K K A T V T E
BINARY_1742 T T Y K L I L N L K Q A K E E A I T E A V D A G T A E K Y F K L I A N A K T V E G V W T L K D E T K K F T V T E
BINARY_1743 T T Y K L I L N L K Q A K E E A I T E A V D A G T A E K Y F K L I A N A K T V E G V W T L K D E T K T A T V T E
BINARY_1744 T T Y K L I L N L K Q A K E E A I T E A V D A G T A E K Y F K L I A N A K T V E G V W T L K D E T K T F T V T E
BINARY_1745 T T Y K L I L N L K Q A K E E A I T E A V D A G T A E K Y F K L I A N A K T V E G V W T Y K D E I L K A T V T E
BINARY_1746 T T Y K L I L N L K Q A K E E A I T E A V D A G T A E K Y F K L I A N A K T V E G V W T Y K D E I L K F T V T E
BINARY_1747 T T Y K L I L N L K Q A K E E A I T E A V D A G T A E K Y F K L I A N A K T V E G V W T Y K D E I L T A T V T E
BINARY_1748 T T Y K L I L N L K Q A K E E A I T E A V D A G T A E K Y F K L I A N A K T V E G V W T Y K D E I L T F T V T E
BINARY_1749 T T Y K L I L N L K Q A K E E A I T E A V D A G T A E K Y F K L I A N A K T V E G V W T Y K D E I K K A T V T E
BINARY_1750 T T Y K L I L N L K Q A K E E A I T E A V D A G T A E K Y F K L I A N A K T V E G V W T Y K D E I K K F T V T E
BINARY_1751 T T Y K L I L N L K Q A K E E A I T E A V D A G T A E K Y F K L I A N A K T V E G V W T Y K D E I K T A T V T E
BINARY_1752 T T Y K L I L N L K Q A K E E A I T E A V D A G T A E K Y F K L I A N A K T V E G V W T Y K D E I K T F T V T E
BINARY_1753 T T Y K L I L N L K Q A K E E A I T E A V D A G T A E K Y F K L I A N A K T V E G V W T Y K D E T L K A T V T E
BINARY_1754 T T Y K L I L N L K Q A K E E A I T E A V D A G T A E K Y F K L I A N A K T V E G V W T Y K D E T L K F T V T E
BINARY_1755 T T Y K L I L N L K Q A K E E A I T E A V D A G T A E K Y F K L I A N A K T V E G V W T Y K D E T L T A T V T E
BINARY_1756 T T Y K L I L N L K Q A K E E A I T E A V D A G T A E K Y F K L I A N A K T V E G V W T Y K D E T L T F T V T E
BINARY_1757 T T Y K L I L N L K Q A K E E A I T E A V D A G T A E K Y F K L I A N A K T V E G V W T Y K D E T K K A T V T E
BINARY_1758 T T Y K L I L N L K Q A K E E A I T E A V D A G T A E K Y F K L I A N A K T V E G V W T Y K D E T K K F T V T E
BINARY_1759 T T Y K L I L N L K Q A K E E A I T E A V D A G T A E K Y F K L I A N A K T V E G V W T Y K D E T K T A T V T E
BINARY_1760 T T Y K L I L N L K Q A K E E A I T E A V D A G T A E K Y F K L I A N A K T V E G V W T Y K D E T K T F T V T E
BINARY_1761 T T Y K L I L N L K Q A K E E A I T E A V D A G T A E K Y F K L Y A N A K T V E G V W T L K D E I L K A T V T E
BINARY_1762 T T Y K L I L N L K Q A K E E A I T E A V D A G T A E K Y F K L Y A N A K T V E G V W T L K D E I L K F T V T E
BINARY_1763 T T Y K L I L N L K Q A K E E A I T E A V D A G T A E K Y F K L Y A N A K T V E G V W T L K D E I L T A T V T E
BINARY_1764 T T Y K L I L N L K Q A K E E A I T E A V D A G T A E K Y F K L Y A N A K T V E G V W T L K D E I L T F T V T E
BINARY_1765 T T Y K L I L N L K Q A K E E A I T E A V D A G T A E K Y F K L Y A N A K T V E G V W T L K D E I K K A T V T E
BINARY_1766 T T Y K L I L N L K Q A K E E A I T E A V D A G T A E K Y F K L Y A N A K T V E G V W T L K D E I K K F T V T E
BINARY_1767 T T Y K L I L N L K Q A K E E A I T E A V D A G T A E K Y F K L Y A N A K T V E G V W T L K D E I K T A T V T E
BINARY_1768 T T Y K L I L N L K Q A K E E A I T E A V D A G T A E K Y F K L Y A N A K T V E G V W T L K D E I K T F T V T E
BINARY_1769 T T Y K L I L N L K Q A K E E A I T E A V D A G T A E K Y F K L Y A N A K T V E G V W T L K D E T L K A T V T E
BINARY_1770 T T Y K L I L N L K Q A K E E A I T E A V D A G T A E K Y F K L Y A N A K T V E G V W T L K D E T L K F T V T E
BINARY_1771 T T Y K L I L N L K Q A K E E A I T E A V D A G T A E K Y F K L Y A N A K T V E G V W T L K D E T L T A T V T E
BINARY_1772 T T Y K L I L N L K Q A K E E A I T E A V D A G T A E K Y F K L Y A N A K T V E G V W T L K D E T L T F T V T E
BINARY_1773 T T Y K L I L N L K Q A K E E A I T E A V D A G T A E K Y F K L Y A N A K T V E G V W T L K D E T K K A T V T E
BINARY_1774 T T Y K L I L N L K Q A K E E A I T E A V D A G T A E K Y F K L Y A N A K T V E G V W T L K D E T K K F T V T E
BINARY_1775 T T Y K L I L N L K Q A K E E A I T E A V D A G T A E K Y F K L Y A N A K T V E G V W T L K D E T K T A T V T E
BINARY_1776 T T Y K L I L N L K Q A K E E A I T E A V D A G T A E K Y F K L Y A N A K T V E G V W T L K D E T K T F T V T E
BINARY_1777 T T Y K L I L N L K Q A K E E A I T E A V D A G T A E K Y F K L Y A N A K T V E G V W T Y K D E I L K A T V T E
BINARY_1778 T T Y K L I L N L K Q A K E E A I T E A V D A G T A E K Y F K L Y A N A K T V E G V W T Y K D E I L K F T V T E
BINARY_1779 T T Y K L I L N L K Q A K E E A I T E A V D A G T A E K Y F K L Y A N A K T V E G V W T Y K D E I L T A T V T E
BINARY_1780 T T Y K L I L N L K Q A K E E A I T E A V D A G T A E K Y F K L Y A N A K T V E G V W T Y K D E I L T F T V T E
BINARY_1781 T T Y K L I L N L K Q A K E E A I T E A V D A G T A E K Y F K L Y A N A K T V E G V W T Y K D E I K K A T V T E
BINARY_1782 T T Y K L I L N L K Q A K E E A I T E A V D A G T A E K Y F K L Y A N A K T V E G V W T Y K D E I K K F T V T E
BINARY_1783 T T Y K L I L N L K Q A K E E A I T E A V D A G T A E K Y F K L Y A N A K T V E G V W T Y K D E I K T A T V T E
BINARY_1784 T T Y K L I L N L K Q A K E E A I T E A V D A G T A E K Y F K L Y A N A K T V E G V W T Y K D E I K T F T V T E
BINARY_1785 T T Y K L I L N L K Q A K E E A I T E A V D A G T A E K Y F K L Y A N A K T V E G V W T Y K D E T L K A T V T E
BINARY_1786 T T Y K L I L N L K Q A K E E A I T E A V D A G T A E K Y F K L Y A N A K T V E G V W T Y K D E T L K F T V T E
BINARY_1787 T T Y K L I L N L K Q A K E E A I T E A V D A G T A E K Y F K L Y A N A K T V E G V W T Y K D E T L T A T V T E
BINARY_1788 T T Y K L I L N L K Q A K E E A I T E A V D A G T A E K Y F K L Y A N A K T V E G V W T Y K D E T L T F T V T E
BINARY_1789 T T Y K L I L N L K Q A K E E A I T E A V D A G T A E K Y F K L Y A N A K T V E G V W T Y K D E T K K A T V T E
BINARY_1790 T T Y K L I L N L K Q A K E E A I T E A V D A G T A E K Y F K L Y A N A K T V E G V W T Y K D E T K K F T V T E
BINARY_1791 T T Y K L I L N L K Q A K E E A I T E A V D A G T A E K Y F K L Y A N A K T V E G V W T Y K D E T K T A T V T E
BINARY_1792 T T Y K L I L N L K Q A K E E A I T E A V D A G T A E K Y F K L Y A N A K T V E G V W T Y K D E T K T F T V T E
BINARY_1793 T T Y K L I L N L K Q A K E E A I T E A V D A A I A E K Y I K L I A N A K T V E G V W T L K D E I L K A T V T E
BINARY_1794 T T Y K L I L N L K Q A K E E A I T E A V D A A I A E K Y I K L I A N A K T V E G V W T L K D E I L K F T V T E
BINARY_1795 T T Y K L I L N L K Q A K E E A I T E A V D A A I A E K Y I K L I A N A K T V E G V W T L K D E I L T A T V T E
BINARY_1796 T T Y K L I L N L K Q A K E E A I T E A V D A A I A E K Y I K L I A N A K T V E G V W T L K D E I L T F T V T E
BINARY_1797 T T Y K L I L N L K Q A K E E A I T E A V D A A I A E K Y I K L I A N A K T V E G V W T L K D E I K K A T V T E
BINARY_1798 T T Y K L I L N L K Q A K E E A I T E A V D A A I A E K Y I K L I A N A K T V E G V W T L K D E I K K F T V T E
BINARY_1799 T T Y K L I L N L K Q A K E E A I T E A V D A A I A E K Y I K L I A N A K T V E G V W T L K D E I K T A T V T E
BINARY_1800 T T Y K L I L N L K Q A K E E A I T E A V D A A I A E K Y I K L I A N A K T V E G V W T L K D E I K T F T V T E
BINARY_1801 T T Y K L I L N L K Q A K E E A I T E A V D A A I A E K Y I K L I A N A K T V E G V W T L K D E T L K A T V T E
BINARY_1802 T T Y K L I L N L K Q A K E E A I T E A V D A A I A E K Y I K L I A N A K T V E G V W T L K D E T L K F T V T E
BINARY_1803 T T Y K L I L N L K Q A K E E A I T E A V D A A I A E K Y I K L I A N A K T V E G V W T L K D E T L T A T V T E
BINARY_1804 T T Y K L I L N L K Q A K E E A I T E A V D A A I A E K Y I K L I A N A K T V E G V W T L K D E T L T F T V T E
BINARY_1805 T T Y K L I L N L K Q A K E E A I T E A V D A A I A E K Y I K L I A N A K T V E G V W T L K D E T K K A T V T E
BINARY_1806 T T Y K L I L N L K Q A K E E A I T E A V D A A I A E K Y I K L I A N A K T V E G V W T L K D E T K K F T V T E
BINARY_1807 T T Y K L I L N L K Q A K E E A I T E A V D A A I A E K Y I K L I A N A K T V E G V W T L K D E T K T A T V T E
BINARY_1808 T T Y K L I L N L K Q A K E E A I T E A V D A A I A E K Y I K L I A N A K T V E G V W T L K D E T K T F T V T E
BINARY_1809 T T Y K L I L N L K Q A K E E A I T E A V D A A I A E K Y I K L I A N A K T V E G V W T Y K D E I L K A T V T E
BINARY_1810 T T Y K L I L N L K Q A K E E A I T E A V D A A I A E K Y I K L I A N A K T V E G V W T Y K D E I L K F T V T E
BINARY_1811 T T Y K L I L N L K Q A K E E A I T E A V D A A I A E K Y I K L I A N A K T V E G V W T Y K D E I L T A T V T E
BINARY_1812 T T Y K L I L N L K Q A K E E A I T E A V D A A I A E K Y I K L I A N A K T V E G V W T Y K D E I L T F T V T E
BINARY_1813 T T Y K L I L N L K Q A K E E A I T E A V D A A I A E K Y I K L I A N A K T V E G V W T Y K D E I K K A T V T E
BINARY_1814 T T Y K L I L N L K Q A K E E A I T E A V D A A I A E K Y I K L I A N A K T V E G V W T Y K D E I K K F T V T E
BINARY_1815 T T Y K L I L N L K Q A K E E A I T E A V D A A I A E K Y I K L I A N A K T V E G V W T Y K D E I K T A T V T E
BINARY_1816 T T Y K L I L N L K Q A K E E A I T E A V D A A I A E K Y I K L I A N A K T V E G V W T Y K D E I K T F T V T E
BINARY_1817 T T Y K L I L N L K Q A K E E A I T E A V D A A I A E K Y I K L I A N A K T V E G V W T Y K D E T L K A T V T E
BINARY_1818 T T Y K L I L N L K Q A K E E A I T E A V D A A I A E K Y I K L I A N A K T V E G V W T Y K D E T L K F T V T E
BINARY_1819 T T Y K L I L N L K Q A K E E A I T E A V D A A I A E K Y I K L I A N A K T V E G V W T Y K D E T L T A T V T E
BINARY_1820 T T Y K L I L N L K Q A K E E A I T E A V D A A I A E K Y I K L I A N A K T V E G V W T Y K D E T L T F T V T E
BINARY_1821 T T Y K L I L N L K Q A K E E A I T E A V D A A I A E K Y I K L I A N A K T V E G V W T Y K D E T K K A T V T E

BINARY_1822 T T Y K L I L N L K Q A K E E A I T E A V D A A I A E K Y I K L I A N A K T V E G V W T Y K D E T K K F T V T E
BINARY_1823 T T Y K L I L N L K Q A K E E A I T E A V D A A I A E K Y I K L I A N A K T V E G V W T Y K D E T K T A T V T E
BINARY_1824 T T Y K L I L N L K Q A K E E A I T E A V D A A I A E K Y I K L I A N A K T V E G V W T Y K D E T K T F T V T E
BINARY_1825 T T Y K L I L N L K Q A K E E A I T E A V D A A I A E K Y I K L Y A N A K T V E G V W T L K D E I L K A T V T E
BINARY_1826 T T Y K L I L N L K Q A K E E A I T E A V D A A I A E K Y I K L Y A N A K T V E G V W T L K D E I L K F T V T E
BINARY_1827 T T Y K L I L N L K Q A K E E A I T E A V D A A I A E K Y I K L Y A N A K T V E G V W T L K D E I L T A T V T E
BINARY_1828 T T Y K L I L N L K Q A K E E A I T E A V D A A I A E K Y I K L Y A N A K T V E G V W T L K D E I L T F T V T E
BINARY_1829 T T Y K L I L N L K Q A K E E A I T E A V D A A I A E K Y I K L Y A N A K T V E G V W T L K D E I K K A T V T E
BINARY_1830 T T Y K L I L N L K Q A K E E A I T E A V D A A I A E K Y I K L Y A N A K T V E G V W T L K D E I K K F T V T E
BINARY_1831 T T Y K L I L N L K Q A K E E A I T E A V D A A I A E K Y I K L Y A N A K T V E G V W T L K D E I K T A T V T E
BINARY_1832 T T Y K L I L N L K Q A K E E A I T E A V D A A I A E K Y I K L Y A N A K T V E G V W T L K D E I K T F T V T E
BINARY_1833 T T Y K L I L N L K Q A K E E A I T E A V D A A I A E K Y I K L Y A N A K T V E G V W T L K D E T L K A T V T E
BINARY_1834 T T Y K L I L N L K Q A K E E A I T E A V D A A I A E K Y I K L Y A N A K T V E G V W T L K D E T L K F T V T E
BINARY_1835 T T Y K L I L N L K Q A K E E A I T E A V D A A I A E K Y I K L Y A N A K T V E G V W T L K D E T L T A T V T E
BINARY_1836 T T Y K L I L N L K Q A K E E A I T E A V D A A I A E K Y I K L Y A N A K T V E G V W T L K D E T L T F T V T E
BINARY_1837 T T Y K L I L N L K Q A K E E A I T E A V D A A I A E K Y I K L Y A N A K T V E G V W T L K D E T K K A T V T E
BINARY_1838 T T Y K L I L N L K Q A K E E A I T E A V D A A I A E K Y I K L Y A N A K T V E G V W T L K D E T K K F T V T E
BINARY_1839 T T Y K L I L N L K Q A K E E A I T E A V D A A I A E K Y I K L Y A N A K T V E G V W T L K D E T K T A T V T E
BINARY_1840 T T Y K L I L N L K Q A K E E A I T E A V D A A I A E K Y I K L Y A N A K T V E G V W T L K D E T K T F T V T E
BINARY_1841 T T Y K L I L N L K Q A K E E A I T E A V D A A I A E K Y I K L Y A N A K T V E G V W T Y K D E I L K A T V T E
BINARY_1842 T T Y K L I L N L K Q A K E E A I T E A V D A A I A E K Y I K L Y A N A K T V E G V W T Y K D E I L K F T V T E
BINARY_1843 T T Y K L I L N L K Q A K E E A I T E A V D A A I A E K Y I K L Y A N A K T V E G V W T Y K D E I L T A T V T E
BINARY_1844 T T Y K L I L N L K Q A K E E A I T E A V D A A I A E K Y I K L Y A N A K T V E G V W T Y K D E I L T F T V T E
BINARY_1845 T T Y K L I L N L K Q A K E E A I T E A V D A A I A E K Y I K L Y A N A K T V E G V W T Y K D E I K K A T V T E
BINARY_1846 T T Y K L I L N L K Q A K E E A I T E A V D A A I A E K Y I K L Y A N A K T V E G V W T Y K D E I K K F T V T E
BINARY_1847 T T Y K L I L N L K Q A K E E A I T E A V D A A I A E K Y I K L Y A N A K T V E G V W T Y K D E I K T A T V T E
BINARY_1848 T T Y K L I L N L K Q A K E E A I T E A V D A A I A E K Y I K L Y A N A K T V E G V W T Y K D E I K T F T V T E
BINARY_1849 T T Y K L I L N L K Q A K E E A I T E A V D A A I A E K Y I K L Y A N A K T V E G V W T Y K D E T L K A T V T E
BINARY_1850 T T Y K L I L N L K Q A K E E A I T E A V D A A I A E K Y I K L Y A N A K T V E G V W T Y K D E T L K F T V T E
BINARY_1851 T T Y K L I L N L K Q A K E E A I T E A V D A A I A E K Y I K L Y A N A K T V E G V W T Y K D E T L T A T V T E
BINARY_1852 T T Y K L I L N L K Q A K E E A I T E A V D A A I A E K Y I K L Y A N A K T V E G V W T Y K D E T L T F T V T E
BINARY_1853 T T Y K L I L N L K Q A K E E A I T E A V D A A I A E K Y I K L Y A N A K T V E G V W T Y K D E T K K A T V T E
BINARY_1854 T T Y K L I L N L K Q A K E E A I T E A V D A A I A E K Y I K L Y A N A K T V E G V W T Y K D E T K K F T V T E
BINARY_1855 T T Y K L I L N L K Q A K E E A I T E A V D A A I A E K Y I K L Y A N A K T V E G V W T Y K D E T K T A T V T E
BINARY_1856 T T Y K L I L N L K Q A K E E A I T E A V D A A I A E K Y I K L Y A N A K T V E G V W T Y K D E T K T F T V T E
BINARY_1857 T T Y K L I L N L K Q A K E E A I T E A V D A A I A E K Y F K L I A N A K T V E G V W T L K D E I L K A T V T E
BINARY_1858 T T Y K L I L N L K Q A K E E A I T E A V D A A I A E K Y F K L I A N A K T V E G V W T L K D E I L K F T V T E
BINARY_1859 T T Y K L I L N L K Q A K E E A I T E A V D A A I A E K Y F K L I A N A K T V E G V W T L K D E I L T A T V T E
BINARY_1860 T T Y K L I L N L K Q A K E E A I T E A V D A A I A E K Y F K L I A N A K T V E G V W T L K D E I L T F T V T E
BINARY_1861 T T Y K L I L N L K Q A K E E A I T E A V D A A I A E K Y F K L I A N A K T V E G V W T L K D E I K K A T V T E
BINARY_1862 T T Y K L I L N L K Q A K E E A I T E A V D A A I A E K Y F K L I A N A K T V E G V W T L K D E I K K F T V T E
BINARY_1863 T T Y K L I L N L K Q A K E E A I T E A V D A A I A E K Y F K L I A N A K T V E G V W T L K D E I K T A T V T E
BINARY_1864 T T Y K L I L N L K Q A K E E A I T E A V D A A I A E K Y F K L I A N A K T V E G V W T L K D E I K T F T V T E
BINARY_1865 T T Y K L I L N L K Q A K E E A I T E A V D A A I A E K Y F K L I A N A K T V E G V W T L K D E T L K A T V T E
BINARY_1866 T T Y K L I L N L K Q A K E E A I T E A V D A A I A E K Y F K L I A N A K T V E G V W T L K D E T L K F T V T E
BINARY_1867 T T Y K L I L N L K Q A K E E A I T E A V D A A I A E K Y F K L I A N A K T V E G V W T L K D E T L T A T V T E
BINARY_1868 T T Y K L I L N L K Q A K E E A I T E A V D A A I A E K Y F K L I A N A K T V E G V W T L K D E T L T F T V T E
BINARY_1869 T T Y K L I L N L K Q A K E E A I T E A V D A A I A E K Y F K L I A N A K T V E G V W T L K D E T K K A T V T E
BINARY_1870 T T Y K L I L N L K Q A K E E A I T E A V D A A I A E K Y F K L I A N A K T V E G V W T L K D E T K K F T V T E
BINARY_1871 T T Y K L I L N L K Q A K E E A I T E A V D A A I A E K Y F K L I A N A K T V E G V W T L K D E T K T A T V T E
BINARY_1872 T T Y K L I L N L K Q A K E E A I T E A V D A A I A E K Y F K L I A N A K T V E G V W T L K D E T K T F T V T E
BINARY_1873 T T Y K L I L N L K Q A K E E A I T E A V D A A I A E K Y F K L I A N A K T V E G V W T Y K D E I L K A T V T E
BINARY_1874 T T Y K L I L N L K Q A K E E A I T E A V D A A I A E K Y F K L I A N A K T V E G V W T Y K D E I L K F T V T E
BINARY_1875 T T Y K L I L N L K Q A K E E A I T E A V D A A I A E K Y F K L I A N A K T V E G V W T Y K D E I L T A T V T E
BINARY_1876 T T Y K L I L N L K Q A K E E A I T E A V D A A I A E K Y F K L I A N A K T V E G V W T Y K D E I L T F T V T E
BINARY_1877 T T Y K L I L N L K Q A K E E A I T E A V D A A I A E K Y F K L I A N A K T V E G V W T Y K D E I K K A T V T E
BINARY_1878 T T Y K L I L N L K Q A K E E A I T E A V D A A I A E K Y F K L I A N A K T V E G V W T Y K D E I K K F T V T E
BINARY_1879 T T Y K L I L N L K Q A K E E A I T E A V D A A I A E K Y F K L I A N A K T V E G V W T Y K D E I K T A T V T E
BINARY_1880 T T Y K L I L N L K Q A K E E A I T E A V D A A I A E K Y F K L I A N A K T V E G V W T Y K D E I K T F T V T E
BINARY_1881 T T Y K L I L N L K Q A K E E A I T E A V D A A I A E K Y F K L I A N A K T V E G V W T Y K D E T L K A T V T E
BINARY_1882 T T Y K L I L N L K Q A K E E A I T E A V D A A I A E K Y F K L I A N A K T V E G V W T Y K D E T L K F T V T E
BINARY_1883 T T Y K L I L N L K Q A K E E A I T E A V D A A I A E K Y F K L I A N A K T V E G V W T Y K D E T L T A T V T E
BINARY_1884 T T Y K L I L N L K Q A K E E A I T E A V D A A I A E K Y F K L I A N A K T V E G V W T Y K D E T L T F T V T E
BINARY_1885 T T Y K L I L N L K Q A K E E A I T E A V D A A I A E K Y F K L I A N A K T V E G V W T Y K D E T K K A T V T E
BINARY_1886 T T Y K L I L N L K Q A K E E A I T E A V D A A I A E K Y F K L I A N A K T V E G V W T Y K D E T K K F T V T E
BINARY_1887 T T Y K L I L N L K Q A K E E A I T E A V D A A I A E K Y F K L I A N A K T V E G V W T Y K D E T K T A T V T E
BINARY_1888 T T Y K L I L N L K Q A K E E A I T E A V D A A I A E K Y F K L I A N A K T V E G V W T Y K D E T K T F T V T E
BINARY_1889 T T Y K L I L N L K Q A K E E A I T E A V D A A I A E K Y F K L Y A N A K T V E G V W T L K D E I L K A T V T E
BINARY_1890 T T Y K L I L N L K Q A K E E A I T E A V D A A I A E K Y F K L Y A N A K T V E G V W T L K D E I L K F T V T E
BINARY_1891 T T Y K L I L N L K Q A K E E A I T E A V D A A I A E K Y F K L Y A N A K T V E G V W T L K D E I L T A T V T E
BINARY_1892 T T Y K L I L N L K Q A K E E A I T E A V D A A I A E K Y F K L Y A N A K T V E G V W T L K D E I L T F T V T E
BINARY_1893 T T Y K L I L N L K Q A K E E A I T E A V D A A I A E K Y F K L Y A N A K T V E G V W T L K D E I K K A T V T E
BINARY_1894 T T Y K L I L N L K Q A K E E A I T E A V D A A I A E K Y F K L Y A N A K T V E G V W T L K D E I K K F T V T E
BINARY_1895 T T Y K L I L N L K Q A K E E A I T E A V D A A I A E K Y F K L Y A N A K T V E G V W T L K D E I K T A T V T E
BINARY_1896 T T Y K L I L N L K Q A K E E A I T E A V D A A I A E K Y F K L Y A N A K T V E G V W T L K D E I K T F T V T E
BINARY_1897 T T Y K L I L N L K Q A K E E A I T E A V D A A I A E K Y F K L Y A N A K T V E G V W T L K D E T L K A T V T E
BINARY_1898 T T Y K L I L N L K Q A K E E A I T E A V D A A I A E K Y F K L Y A N A K T V E G V W T L K D E T L K F T V T E
BINARY_1899 T T Y K L I L N L K Q A K E E A I T E A V D A A I A E K Y F K L Y A N A K T V E G V W T L K D E T L T A T V T E
BINARY_1900 T T Y K L I L N L K Q A K E E A I T E A V D A A I A E K Y F K L Y A N A K T V E G V W T L K D E T L T F T V T E
BINARY_1901 T T Y K L I L N L K Q A K E E A I T E A V D A A I A E K Y F K L Y A N A K T V E G V W T L K D E T K K A T V T E
BINARY_1902 T T Y K L I L N L K Q A K E E A I T E A V D A A I A E K Y F K L Y A N A K T V E G V W T L K D E T K K F T V T E
BINARY_1903 T T Y K L I L N L K Q A K E E A I T E A V D A A I A E K Y F K L Y A N A K T V E G V W T L K D E T K T A T V T E
BINARY_1904 T T Y K L I L N L K Q A K E E A I T E A V D A A I A E K Y F K L Y A N A K T V E G V W T L K D E T K T F T V T E
BINARY_1905 T T Y K L I L N L K Q A K E E A I T E A V D A A I A E K Y F K L Y A N A K T V E G V W T Y K D E I L K A T V T E
BINARY_1906 T T Y K L I L N L K Q A K E E A I T E A V D A A I A E K Y F K L Y A N A K T V E G V W T Y K D E I L K F T V T E
BINARY_1907 T T Y K L I L N L K Q A K E E A I T E A V D A A I A E K Y F K L Y A N A K T V E G V W T Y K D E I L T A T V T E
BINARY_1908 T T Y K L I L N L K Q A K E E A I T E A V D A A I A E K Y F K L Y A N A K T V E G V W T Y K D E I L T F T V T E
BINARY_1909 T T Y K L I L N L K Q A K E E A I T E A V D A A I A E K Y F K L Y A N A K T V E G V W T Y K D E I K K A T V T E
BINARY_1910 T T Y K L I L N L K Q A K E E A I T E A V D A A I A E K Y F K L Y A N A K T V E G V W T Y K D E I K K F T V T E
BINARY_1911 T T Y K L I L N L K Q A K E E A I T E A V D A A I A E K Y F K L Y A N A K T V E G V W T Y K D E I K T A T V T E
BINARY_1912 T T Y K L I L N L K Q A K E E A I T E A V D A A I A E K Y F K L Y A N A K T V E G V W T Y K D E I K T F T V T E
BINARY_1913 T T Y K L I L N L K Q A K E E A I T E A V D A A I A E K Y F K L Y A N A K T V E G V W T Y K D E T L K A T V T E
BINARY_1914 T T Y K L I L N L K Q A K E E A I T E A V D A A I A E K Y F K L Y A N A K T V E G V W T Y K D E T L K F T V T E
BINARY_1915 T T Y K L I L N L K Q A K E E A I T E A V D A A I A E K Y F K L Y A N A K T V E G V W T Y K D E T L T A T V T E
BINARY_1916 T T Y K L I L N L K Q A K E E A I T E A V D A A I A E K Y F K L Y A N A K T V E G V W T Y K D E T L T F T V T E

BINARY_1917 T T Y K L I L N L K Q A K E E A I T E A V D A A I A E K Y F K L Y A N A K T V E G V W T Y K D E T K K A T V T E
BINARY_1918 T T Y K L I L N L K Q A K E E A I T E A V D A A I A E K Y F K L Y A N A K T V E G V W T Y K D E T K K F T V T E
BINARY_1919 T T Y K L I L N L K Q A K E E A I T E A V D A A I A E K Y F K L Y A N A K T V E G V W T Y K D E T K T A T V T E
BINARY_1920 T T Y K L I L N L K Q A K E E A I T E A V D A A I A E K Y F K L Y A N A K T V E G V W T Y K D E T K T F T V T E
BINARY_1921 T T Y K L I L N L K Q A K E E A I T E A V D A A T A E K Y I K L I A N A K T V E G V W T L K D E I L K A T V T E
BINARY_1922 T T Y K L I L N L K Q A K E E A I T E A V D A A T A E K Y I K L I A N A K T V E G V W T L K D E I L K F T V T E
BINARY_1923 T T Y K L I L N L K Q A K E E A I T E A V D A A T A E K Y I K L I A N A K T V E G V W T L K D E I L T A T V T E
BINARY_1924 T T Y K L I L N L K Q A K E E A I T E A V D A A T A E K Y I K L I A N A K T V E G V W T L K D E I L T F T V T E
BINARY_1925 T T Y K L I L N L K Q A K E E A I T E A V D A A T A E K Y I K L I A N A K T V E G V W T L K D E I K K A T V T E
BINARY_1926 T T Y K L I L N L K Q A K E E A I T E A V D A A T A E K Y I K L I A N A K T V E G V W T L K D E I K K F T V T E
BINARY_1927 T T Y K L I L N L K Q A K E E A I T E A V D A A T A E K Y I K L I A N A K T V E G V W T L K D E I K T A T V T E
BINARY_1928 T T Y K L I L N L K Q A K E E A I T E A V D A A T A E K Y I K L I A N A K T V E G V W T L K D E I K T F T V T E
BINARY_1929 T T Y K L I L N L K Q A K E E A I T E A V D A A T A E K Y I K L I A N A K T V E G V W T L K D E T L K A T V T E
BINARY_1930 T T Y K L I L N L K Q A K E E A I T E A V D A A T A E K Y I K L I A N A K T V E G V W T L K D E T L K F T V T E
BINARY_1931 T T Y K L I L N L K Q A K E E A I T E A V D A A T A E K Y I K L I A N A K T V E G V W T L K D E T L T A T V T E
BINARY_1932 T T Y K L I L N L K Q A K E E A I T E A V D A A T A E K Y I K L I A N A K T V E G V W T L K D E T L T F T V T E
BINARY_1933 T T Y K L I L N L K Q A K E E A I T E A V D A A T A E K Y I K L I A N A K T V E G V W T L K D E T K K A T V T E
BINARY_1934 T T Y K L I L N L K Q A K E E A I T E A V D A A T A E K Y I K L I A N A K T V E G V W T L K D E T K K F T V T E
BINARY_1935 T T Y K L I L N L K Q A K E E A I T E A V D A A T A E K Y I K L I A N A K T V E G V W T L K D E T K T A T V T E
BINARY_1936 T T Y K L I L N L K Q A K E E A I T E A V D A A T A E K Y I K L I A N A K T V E G V W T L K D E T K T F T V T E
BINARY_1937 T T Y K L I L N L K Q A K E E A I T E A V D A A T A E K Y I K L I A N A K T V E G V W T Y K D E I L K A T V T E
BINARY_1938 T T Y K L I L N L K Q A K E E A I T E A V D A A T A E K Y I K L I A N A K T V E G V W T Y K D E I L K F T V T E
BINARY_1939 T T Y K L I L N L K Q A K E E A I T E A V D A A T A E K Y I K L I A N A K T V E G V W T Y K D E I L T A T V T E
BINARY_1940 T T Y K L I L N L K Q A K E E A I T E A V D A A T A E K Y I K L I A N A K T V E G V W T Y K D E I L T F T V T E
BINARY_1941 T T Y K L I L N L K Q A K E E A I T E A V D A A T A E K Y I K L I A N A K T V E G V W T Y K D E I K K A T V T E
BINARY_1942 T T Y K L I L N L K Q A K E E A I T E A V D A A T A E K Y I K L I A N A K T V E G V W T Y K D E I K K F T V T E
BINARY_1943 T T Y K L I L N L K Q A K E E A I T E A V D A A T A E K Y I K L I A N A K T V E G V W T Y K D E I K T A T V T E
BINARY_1944 T T Y K L I L N L K Q A K E E A I T E A V D A A T A E K Y I K L I A N A K T V E G V W T Y K D E I K T F T V T E
BINARY_1945 T T Y K L I L N L K Q A K E E A I T E A V D A A T A E K Y I K L I A N A K T V E G V W T Y K D E T L K A T V T E
BINARY_1946 T T Y K L I L N L K Q A K E E A I T E A V D A A T A E K Y I K L I A N A K T V E G V W T Y K D E T L K F T V T E
BINARY_1947 T T Y K L I L N L K Q A K E E A I T E A V D A A T A E K Y I K L I A N A K T V E G V W T Y K D E T L T A T V T E
BINARY_1948 T T Y K L I L N L K Q A K E E A I T E A V D A A T A E K Y I K L I A N A K T V E G V W T Y K D E T L T F T V T E
BINARY_1949 T T Y K L I L N L K Q A K E E A I T E A V D A A T A E K Y I K L I A N A K T V E G V W T Y K D E T K K A T V T E
BINARY_1950 T T Y K L I L N L K Q A K E E A I T E A V D A A T A E K Y I K L I A N A K T V E G V W T Y K D E T K K F T V T E
BINARY_1951 T T Y K L I L N L K Q A K E E A I T E A V D A A T A E K Y I K L I A N A K T V E G V W T Y K D E T K T A T V T E
BINARY_1952 T T Y K L I L N L K Q A K E E A I T E A V D A A T A E K Y I K L I A N A K T V E G V W T Y K D E T K T F T V T E
BINARY_1953 T T Y K L I L N L K Q A K E E A I T E A V D A A T A E K Y I K L Y A N A K T V E G V W T L K D E I L K A T V T E
BINARY_1954 T T Y K L I L N L K Q A K E E A I T E A V D A A T A E K Y I K L Y A N A K T V E G V W T L K D E I L K F T V T E
BINARY_1955 T T Y K L I L N L K Q A K E E A I T E A V D A A T A E K Y I K L Y A N A K T V E G V W T L K D E I L T A T V T E
BINARY_1956 T T Y K L I L N L K Q A K E E A I T E A V D A A T A E K Y I K L Y A N A K T V E G V W T L K D E I L T F T V T E
BINARY_1957 T T Y K L I L N L K Q A K E E A I T E A V D A A T A E K Y I K L Y A N A K T V E G V W T L K D E I K K A T V T E
BINARY_1958 T T Y K L I L N L K Q A K E E A I T E A V D A A T A E K Y I K L Y A N A K T V E G V W T L K D E I K K F T V T E
BINARY_1959 T T Y K L I L N L K Q A K E E A I T E A V D A A T A E K Y I K L Y A N A K T V E G V W T L K D E I K T A T V T E
BINARY_1960 T T Y K L I L N L K Q A K E E A I T E A V D A A T A E K Y I K L Y A N A K T V E G V W T L K D E I K T F T V T E
BINARY_1961 T T Y K L I L N L K Q A K E E A I T E A V D A A T A E K Y I K L Y A N A K T V E G V W T L K D E T L K A T V T E
BINARY_1962 T T Y K L I L N L K Q A K E E A I T E A V D A A T A E K Y I K L Y A N A K T V E G V W T L K D E T L K F T V T E
BINARY_1963 T T Y K L I L N L K Q A K E E A I T E A V D A A T A E K Y I K L Y A N A K T V E G V W T L K D E T L T A T V T E
BINARY_1964 T T Y K L I L N L K Q A K E E A I T E A V D A A T A E K Y I K L Y A N A K T V E G V W T L K D E T L T F T V T E
BINARY_1965 T T Y K L I L N L K Q A K E E A I T E A V D A A T A E K Y I K L Y A N A K T V E G V W T L K D E T K K A T V T E
BINARY_1966 T T Y K L I L N L K Q A K E E A I T E A V D A A T A E K Y I K L Y A N A K T V E G V W T L K D E T K K F T V T E
BINARY_1967 T T Y K L I L N L K Q A K E E A I T E A V D A A T A E K Y I K L Y A N A K T V E G V W T L K D E T K T A T V T E
BINARY_1968 T T Y K L I L N L K Q A K E E A I T E A V D A A T A E K Y I K L Y A N A K T V E G V W T L K D E T K T F T V T E
BINARY_1969 T T Y K L I L N L K Q A K E E A I T E A V D A A T A E K Y I K L Y A N A K T V E G V W T Y K D E I L K A T V T E
BINARY_1970 T T Y K L I L N L K Q A K E E A I T E A V D A A T A E K Y I K L Y A N A K T V E G V W T Y K D E I L K F T V T E
BINARY_1971 T T Y K L I L N L K Q A K E E A I T E A V D A A T A E K Y I K L Y A N A K T V E G V W T Y K D E I L T A T V T E
BINARY_1972 T T Y K L I L N L K Q A K E E A I T E A V D A A T A E K Y I K L Y A N A K T V E G V W T Y K D E I L T F T V T E
BINARY_1973 T T Y K L I L N L K Q A K E E A I T E A V D A A T A E K Y I K L Y A N A K T V E G V W T Y K D E I K K A T V T E
BINARY_1974 T T Y K L I L N L K Q A K E E A I T E A V D A A T A E K Y I K L Y A N A K T V E G V W T Y K D E I K K F T V T E
BINARY_1975 T T Y K L I L N L K Q A K E E A I T E A V D A A T A E K Y I K L Y A N A K T V E G V W T Y K D E I K T A T V T E
BINARY_1976 T T Y K L I L N L K Q A K E E A I T E A V D A A T A E K Y I K L Y A N A K T V E G V W T Y K D E I K T F T V T E
BINARY_1977 T T Y K L I L N L K Q A K E E A I T E A V D A A T A E K Y I K L Y A N A K T V E G V W T Y K D E T L K A T V T E
BINARY_1978 T T Y K L I L N L K Q A K E E A I T E A V D A A T A E K Y I K L Y A N A K T V E G V W T Y K D E T L K F T V T E
BINARY_1979 T T Y K L I L N L K Q A K E E A I T E A V D A A T A E K Y I K L Y A N A K T V E G V W T Y K D E T L T A T V T E
BINARY_1980 T T Y K L I L N L K Q A K E E A I T E A V D A A T A E K Y I K L Y A N A K T V E G V W T Y K D E T L T F T V T E
BINARY_1981 T T Y K L I L N L K Q A K E E A I T E A V D A A T A E K Y I K L Y A N A K T V E G V W T Y K D E T K K A T V T E
BINARY_1982 T T Y K L I L N L K Q A K E E A I T E A V D A A T A E K Y I K L Y A N A K T V E G V W T Y K D E T K K F T V T E
BINARY_1983 T T Y K L I L N L K Q A K E E A I T E A V D A A T A E K Y I K L Y A N A K T V E G V W T Y K D E T K T A T V T E
BINARY_1984 T T Y K L I L N L K Q A K E E A I T E A V D A A T A E K Y I K L Y A N A K T V E G V W T Y K D E T K T F T V T E
BINARY_1985 T T Y K L I L N L K Q A K E E A I T E A V D A A T A E K Y F K L I A N A K T V E G V W T L K D E I L K A T V T E
BINARY_1986 T T Y K L I L N L K Q A K E E A I T E A V D A A T A E K Y F K L I A N A K T V E G V W T L K D E I L K F T V T E
BINARY_1987 T T Y K L I L N L K Q A K E E A I T E A V D A A T A E K Y F K L I A N A K T V E G V W T L K D E I L T A T V T E
BINARY_1988 T T Y K L I L N L K Q A K E E A I T E A V D A A T A E K Y F K L I A N A K T V E G V W T L K D E I L T F T V T E
BINARY_1989 T T Y K L I L N L K Q A K E E A I T E A V D A A T A E K Y F K L I A N A K T V E G V W T L K D E I K K A T V T E
BINARY_1990 T T Y K L I L N L K Q A K E E A I T E A V D A A T A E K Y F K L I A N A K T V E G V W T L K D E I K K F T V T E
BINARY_1991 T T Y K L I L N L K Q A K E E A I T E A V D A A T A E K Y F K L I A N A K T V E G V W T L K D E I K T A T V T E
BINARY_1992 T T Y K L I L N L K Q A K E E A I T E A V D A A T A E K Y F K L I A N A K T V E G V W T L K D E I K T F T V T E
BINARY_1993 T T Y K L I L N L K Q A K E E A I T E A V D A A T A E K Y F K L I A N A K T V E G V W T L K D E T L K A T V T E
BINARY_1994 T T Y K L I L N L K Q A K E E A I T E A V D A A T A E K Y F K L I A N A K T V E G V W T L K D E T L K F T V T E
BINARY_1995 T T Y K L I L N L K Q A K E E A I T E A V D A A T A E K Y F K L I A N A K T V E G V W T L K D E T L T A T V T E
BINARY_1996 T T Y K L I L N L K Q A K E E A I T E A V D A A T A E K Y F K L I A N A K T V E G V W T L K D E T L T F T V T E
BINARY_1997 T T Y K L I L N L K Q A K E E A I T E A V D A A T A E K Y F K L I A N A K T V E G V W T L K D E T K K A T V T E
BINARY_1998 T T Y K L I L N L K Q A K E E A I T E A V D A A T A E K Y F K L I A N A K T V E G V W T L K D E T K K F T V T E
BINARY_1999 T T Y K L I L N L K Q A K E E A I T E A V D A A T A E K Y F K L I A N A K T V E G V W T L K D E T K T A T V T E
BINARY_2000 T T Y K L I L N L K Q A K E E A I T E A V D A A T A E K Y F K L I A N A K T V E G V W T L K D E T K T F T V T E
BINARY_2001 T T Y K L I L N L K Q A K E E A I T E A V D A A T A E K Y F K L I A N A K T V E G V W T Y K D E I L K A T V T E
BINARY_2002 T T Y K L I L N L K Q A K E E A I T E A V D A A T A E K Y F K L I A N A K T V E G V W T Y K D E I L K F T V T E
BINARY_2003 T T Y K L I L N L K Q A K E E A I T E A V D A A T A E K Y F K L I A N A K T V E G V W T Y K D E I L T A T V T E
BINARY_2004 T T Y K L I L N L K Q A K E E A I T E A V D A A T A E K Y F K L I A N A K T V E G V W T Y K D E I L T F T V T E
BINARY_2005 T T Y K L I L N L K Q A K E E A I T E A V D A A T A E K Y F K L I A N A K T V E G V W T Y K D E I K K A T V T E
BINARY_2006 T T Y K L I L N L K Q A K E E A I T E A V D A A T A E K Y F K L I A N A K T V E G V W T Y K D E I K K F T V T E
BINARY_2007 T T Y K L I L N L K Q A K E E A I T E A V D A A T A E K Y F K L I A N A K T V E G V W T Y K D E I K T A T V T E
BINARY_2008 T T Y K L I L N L K Q A K E E A I T E A V D A A T A E K Y F K L I A N A K T V E G V W T Y K D E I K T F T V T E
BINARY_2009 T T Y K L I L N L K Q A K E E A I T E A V D A A T A E K Y F K L I A N A K T V E G V W T Y K D E T L K A T V T E
BINARY_2010 T T Y K L I L N L K Q A K E E A I T E A V D A A T A E K Y F K L I A N A K T V E G V W T Y K D E T L K F T V T E
BINARY_2011 T T Y K L I L N L K Q A K E E A I T E A V D A A T A E K Y F K L I A N A K T V E G V W T Y K D E T L T A T V T E

BINARY_2012 T T Y K L I L N L K Q A K E E A I T E A V D A A T A E K Y F K L I A N A K T V E G V W T Y K D E T L T F T V T E
BINARY_2013 T T Y K L I L N L K Q A K E E A I T E A V D A A T A E K Y F K L I A N A K T V E G V W T Y K D E T K K A T V T E
BINARY_2014 T T Y K L I L N L K Q A K E E A I T E A V D A A T A E K Y F K L I A N A K T V E G V W T Y K D E T K K F T V T E
BINARY_2015 T T Y K L I L N L K Q A K E E A I T E A V D A A T A E K Y F K L I A N A K T V E G V W T Y K D E T K T A T V T E
BINARY_2016 T T Y K L I L N L K Q A K E E A I T E A V D A A T A E K Y F K L I A N A K T V E G V W T Y K D E T K T F T V T E
BINARY_2017 T T Y K L I L N L K Q A K E E A I T E A V D A A T A E K Y F K L Y A N A K T V E G V W T L K D E I L K A T V T E
BINARY_2018 T T Y K L I L N L K Q A K E E A I T E A V D A A T A E K Y F K L Y A N A K T V E G V W T L K D E I L K F T V T E
BINARY_2019 T T Y K L I L N L K Q A K E E A I T E A V D A A T A E K Y F K L Y A N A K T V E G V W T L K D E I L T A T V T E
BINARY_2020 T T Y K L I L N L K Q A K E E A I T E A V D A A T A E K Y F K L Y A N A K T V E G V W T L K D E I L T F T V T E
BINARY_2021 T T Y K L I L N L K Q A K E E A I T E A V D A A T A E K Y F K L Y A N A K T V E G V W T L K D E I K K A T V T E
BINARY_2022 T T Y K L I L N L K Q A K E E A I T E A V D A A T A E K Y F K L Y A N A K T V E G V W T L K D E I K K F T V T E
BINARY_2023 T T Y K L I L N L K Q A K E E A I T E A V D A A T A E K Y F K L Y A N A K T V E G V W T L K D E I K T A T V T E
BINARY_2024 T T Y K L I L N L K Q A K E E A I T E A V D A A T A E K Y F K L Y A N A K T V E G V W T L K D E I K T F T V T E
BINARY_2025 T T Y K L I L N L K Q A K E E A I T E A V D A A T A E K Y F K L Y A N A K T V E G V W T L K D E T L K A T V T E
BINARY_2026 T T Y K L I L N L K Q A K E E A I T E A V D A A T A E K Y F K L Y A N A K T V E G V W T L K D E T L K F T V T E
BINARY_2027 T T Y K L I L N L K Q A K E E A I T E A V D A A T A E K Y F K L Y A N A K T V E G V W T L K D E T L T A T V T E
BINARY_2028 T T Y K L I L N L K Q A K E E A I T E A V D A A T A E K Y F K L Y A N A K T V E G V W T L K D E T L T F T V T E
BINARY_2029 T T Y K L I L N L K Q A K E E A I T E A V D A A T A E K Y F K L Y A N A K T V E G V W T L K D E T K K A T V T E
BINARY_2030 T T Y K L I L N L K Q A K E E A I T E A V D A A T A E K Y F K L Y A N A K T V E G V W T L K D E T K K F T V T E
BINARY_2031 T T Y K L I L N L K Q A K E E A I T E A V D A A T A E K Y F K L Y A N A K T V E G V W T L K D E T K T A T V T E
BINARY_2032 T T Y K L I L N L K Q A K E E A I T E A V D A A T A E K Y F K L Y A N A K T V E G V W T L K D E T K T F T V T E
BINARY_2033 T T Y K L I L N L K Q A K E E A I T E A V D A A T A E K Y F K L Y A N A K T V E G V W T Y K D E I L K A T V T E
BINARY_2034 T T Y K L I L N L K Q A K E E A I T E A V D A A T A E K Y F K L Y A N A K T V E G V W T Y K D E I L K F T V T E
BINARY_2035 T T Y K L I L N L K Q A K E E A I T E A V D A A T A E K Y F K L Y A N A K T V E G V W T Y K D E I L T A T V T E
BINARY_2036 T T Y K L I L N L K Q A K E E A I T E A V D A A T A E K Y F K L Y A N A K T V E G V W T Y K D E I L T F T V T E
BINARY_2037 T T Y K L I L N L K Q A K E E A I T E A V D A A T A E K Y F K L Y A N A K T V E G V W T Y K D E I K K A T V T E
BINARY_2038 T T Y K L I L N L K Q A K E E A I T E A V D A A T A E K Y F K L Y A N A K T V E G V W T Y K D E I K K F T V T E
BINARY_2039 T T Y K L I L N L K Q A K E E A I T E A V D A A T A E K Y F K L Y A N A K T V E G V W T Y K D E I K T A T V T E
BINARY_2040 T T Y K L I L N L K Q A K E E A I T E A V D A A T A E K Y F K L Y A N A K T V E G V W T Y K D E I K T F T V T E
BINARY_2041 T T Y K L I L N L K Q A K E E A I T E A V D A A T A E K Y F K L Y A N A K T V E G V W T Y K D E T L K A T V T E
BINARY_2042 T T Y K L I L N L K Q A K E E A I T E A V D A A T A E K Y F K L Y A N A K T V E G V W T Y K D E T L K F T V T E
BINARY_2043 T T Y K L I L N L K Q A K E E A I T E A V D A A T A E K Y F K L Y A N A K T V E G V W T Y K D E T L T A T V T E
BINARY_2044 T T Y K L I L N L K Q A K E E A I T E A V D A A T A E K Y F K L Y A N A K T V E G V W T Y K D E T L T F T V T E
BINARY_2045 T T Y K L I L N L K Q A K E E A I T E A V D A A T A E K Y F K L Y A N A K T V E G V W T Y K D E T K K A T V T E
BINARY_2046 T T Y K L I L N L K Q A K E E A I T E A V D A A T A E K Y F K L Y A N A K T V E G V W T Y K D E T K K F T V T E
BINARY_2047 T T Y K L I L N L K Q A K E E A I T E A V D A A T A E K Y F K L Y A N A K T V E G V W T Y K D E T K T A T V T E
BINARY_2048 T T Y K L I L N L K Q A K E E A I T E A V D A A T A E K Y F K L Y A N A K T V E G V W T Y K D E T K T F T V T E
BINARY_2049 T T Y K L I L N L K Q L K E E A I K E L V D A G I A E K Y I K L I A N A K T V E G V W T L K D E I L K A T V T E
BINARY_2050 T T Y K L I L N L K Q L K E E A I K E L V D A G I A E K Y I K L I A N A K T V E G V W T L K D E I L K F T V T E
BINARY_2051 T T Y K L I L N L K Q L K E E A I K E L V D A G I A E K Y I K L I A N A K T V E G V W T L K D E I L T A T V T E
BINARY_2052 T T Y K L I L N L K Q L K E E A I K E L V D A G I A E K Y I K L I A N A K T V E G V W T L K D E I L T F T V T E
BINARY_2053 T T Y K L I L N L K Q L K E E A I K E L V D A G I A E K Y I K L I A N A K T V E G V W T L K D E I K K A T V T E
BINARY_2054 T T Y K L I L N L K Q L K E E A I K E L V D A G I A E K Y I K L I A N A K T V E G V W T L K D E I K K F T V T E
BINARY_2055 T T Y K L I L N L K Q L K E E A I K E L V D A G I A E K Y I K L I A N A K T V E G V W T L K D E I K T A T V T E
BINARY_2056 T T Y K L I L N L K Q L K E E A I K E L V D A G I A E K Y I K L I A N A K T V E G V W T L K D E I K T F T V T E
BINARY_2057 T T Y K L I L N L K Q L K E E A I K E L V D A G I A E K Y I K L I A N A K T V E G V W T L K D E T L K A T V T E
BINARY_2058 T T Y K L I L N L K Q L K E E A I K E L V D A G I A E K Y I K L I A N A K T V E G V W T L K D E T L K F T V T E
BINARY_2059 T T Y K L I L N L K Q L K E E A I K E L V D A G I A E K Y I K L I A N A K T V E G V W T L K D E T L T A T V T E
BINARY_2060 T T Y K L I L N L K Q L K E E A I K E L V D A G I A E K Y I K L I A N A K T V E G V W T L K D E T L T F T V T E
BINARY_2061 T T Y K L I L N L K Q L K E E A I K E L V D A G I A E K Y I K L I A N A K T V E G V W T L K D E T K K A T V T E
BINARY_2062 T T Y K L I L N L K Q L K E E A I K E L V D A G I A E K Y I K L I A N A K T V E G V W T L K D E T K K F T V T E
BINARY_2063 T T Y K L I L N L K Q L K E E A I K E L V D A G I A E K Y I K L I A N A K T V E G V W T L K D E T K T A T V T E
BINARY_2064 T T Y K L I L N L K Q L K E E A I K E L V D A G I A E K Y I K L I A N A K T V E G V W T L K D E T K T F T V T E
BINARY_2065 T T Y K L I L N L K Q L K E E A I K E L V D A G I A E K Y I K L I A N A K T V E G V W T Y K D E I L K A T V T E
BINARY_2066 T T Y K L I L N L K Q L K E E A I K E L V D A G I A E K Y I K L I A N A K T V E G V W T Y K D E I L K F T V T E
BINARY_2067 T T Y K L I L N L K Q L K E E A I K E L V D A G I A E K Y I K L I A N A K T V E G V W T Y K D E I L T A T V T E
BINARY_2068 T T Y K L I L N L K Q L K E E A I K E L V D A G I A E K Y I K L I A N A K T V E G V W T Y K D E I L T F T V T E
BINARY_2069 T T Y K L I L N L K Q L K E E A I K E L V D A G I A E K Y I K L I A N A K T V E G V W T Y K D E I K K A T V T E
BINARY_2070 T T Y K L I L N L K Q L K E E A I K E L V D A G I A E K Y I K L I A N A K T V E G V W T Y K D E I K K F T V T E
BINARY_2071 T T Y K L I L N L K Q L K E E A I K E L V D A G I A E K Y I K L I A N A K T V E G V W T Y K D E I K T A T V T E
BINARY_2072 T T Y K L I L N L K Q L K E E A I K E L V D A G I A E K Y I K L I A N A K T V E G V W T Y K D E I K T F T V T E
BINARY_2073 T T Y K L I L N L K Q L K E E A I K E L V D A G I A E K Y I K L I A N A K T V E G V W T Y K D E T L K A T V T E
BINARY_2074 T T Y K L I L N L K Q L K E E A I K E L V D A G I A E K Y I K L I A N A K T V E G V W T Y K D E T L K F T V T E
BINARY_2075 T T Y K L I L N L K Q L K E E A I K E L V D A G I A E K Y I K L I A N A K T V E G V W T Y K D E T L T A T V T E
BINARY_2076 T T Y K L I L N L K Q L K E E A I K E L V D A G I A E K Y I K L I A N A K T V E G V W T Y K D E T L T F T V T E
BINARY_2077 T T Y K L I L N L K Q L K E E A I K E L V D A G I A E K Y I K L I A N A K T V E G V W T Y K D E T K K A T V T E
BINARY_2078 T T Y K L I L N L K Q L K E E A I K E L V D A G I A E K Y I K L I A N A K T V E G V W T Y K D E T K K F T V T E
BINARY_2079 T T Y K L I L N L K Q L K E E A I K E L V D A G I A E K Y I K L I A N A K T V E G V W T Y K D E T K T A T V T E
BINARY_2080 T T Y K L I L N L K Q L K E E A I K E L V D A G I A E K Y I K L I A N A K T V E G V W T Y K D E T K T F T V T E
BINARY_2081 T T Y K L I L N L K Q L K E E A I K E L V D A G I A E K Y I K L Y A N A K T V E G V W T L K D E I L K A T V T E
BINARY_2082 T T Y K L I L N L K Q L K E E A I K E L V D A G I A E K Y I K L Y A N A K T V E G V W T L K D E I L K F T V T E
BINARY_2083 T T Y K L I L N L K Q L K E E A I K E L V D A G I A E K Y I K L Y A N A K T V E G V W T L K D E I L T A T V T E
BINARY_2084 T T Y K L I L N L K Q L K E E A I K E L V D A G I A E K Y I K L Y A N A K T V E G V W T L K D E I L T F T V T E
BINARY_2085 T T Y K L I L N L K Q L K E E A I K E L V D A G I A E K Y I K L Y A N A K T V E G V W T L K D E I K K A T V T E
BINARY_2086 T T Y K L I L N L K Q L K E E A I K E L V D A G I A E K Y I K L Y A N A K T V E G V W T L K D E I K K F T V T E
BINARY_2087 T T Y K L I L N L K Q L K E E A I K E L V D A G I A E K Y I K L Y A N A K T V E G V W T L K D E I K T A T V T E
BINARY_2088 T T Y K L I L N L K Q L K E E A I K E L V D A G I A E K Y I K L Y A N A K T V E G V W T L K D E I K T F T V T E
BINARY_2089 T T Y K L I L N L K Q L K E E A I K E L V D A G I A E K Y I K L Y A N A K T V E G V W T L K D E T L K A T V T E
BINARY_2090 T T Y K L I L N L K Q L K E E A I K E L V D A G I A E K Y I K L Y A N A K T V E G V W T L K D E T L K F T V T E
BINARY_2091 T T Y K L I L N L K Q L K E E A I K E L V D A G I A E K Y I K L Y A N A K T V E G V W T L K D E T L T A T V T E
BINARY_2092 T T Y K L I L N L K Q L K E E A I K E L V D A G I A E K Y I K L Y A N A K T V E G V W T L K D E T L T F T V T E
BINARY_2093 T T Y K L I L N L K Q L K E E A I K E L V D A G I A E K Y I K L Y A N A K T V E G V W T L K D E T K K A T V T E
BINARY_2094 T T Y K L I L N L K Q L K E E A I K E L V D A G I A E K Y I K L Y A N A K T V E G V W T L K D E T K K F T V T E
BINARY_2095 T T Y K L I L N L K Q L K E E A I K E L V D A G I A E K Y I K L Y A N A K T V E G V W T L K D E T K T A T V T E
BINARY_2096 T T Y K L I L N L K Q L K E E A I K E L V D A G I A E K Y I K L Y A N A K T V E G V W T L K D E T K T F T V T E
BINARY_2097 T T Y K L I L N L K Q L K E E A I K E L V D A G I A E K Y I K L Y A N A K T V E G V W T Y K D E I L K A T V T E
BINARY_2098 T T Y K L I L N L K Q L K E E A I K E L V D A G I A E K Y I K L Y A N A K T V E G V W T Y K D E I L K F T V T E
BINARY_2099 T T Y K L I L N L K Q L K E E A I K E L V D A G I A E K Y I K L Y A N A K T V E G V W T Y K D E I L T A T V T E
BINARY_2100 T T Y K L I L N L K Q L K E E A I K E L V D A G I A E K Y I K L Y A N A K T V E G V W T Y K D E I L T F T V T E
BINARY_2101 T T Y K L I L N L K Q L K E E A I K E L V D A G I A E K Y I K L Y A N A K T V E G V W T Y K D E I K K A T V T E
BINARY_2102 T T Y K L I L N L K Q L K E E A I K E L V D A G I A E K Y I K L Y A N A K T V E G V W T Y K D E I K K F T V T E
BINARY_2103 T T Y K L I L N L K Q L K E E A I K E L V D A G I A E K Y I K L Y A N A K T V E G V W T Y K D E I K T A T V T E
BINARY_2104 T T Y K L I L N L K Q L K E E A I K E L V D A G I A E K Y I K L Y A N A K T V E G V W T Y K D E I K T F T V T E
BINARY_2105 T T Y K L I L N L K Q L K E E A I K E L V D A G I A E K Y I K L Y A N A K T V E G V W T Y K D E T L K A T V T E
BINARY_2106 T T Y K L I L N L K Q L K E E A I K E L V D A G I A E K Y I K L Y A N A K T V E G V W T Y K D E T L K F T V T E

BINARY_2107 T T Y K L I L N L K Q L K E E A I K E L V D A G I A E K Y I K L Y A N A K T V E G V W T Y K D E T L T A T V T E
BINARY_2108 T T Y K L I L N L K Q L K E E A I K E L V D A G I A E K Y I K L Y A N A K T V E G V W T Y K D E T L T F T V T E
BINARY_2109 T T Y K L I L N L K Q L K E E A I K E L V D A G I A E K Y I K L Y A N A K T V E G V W T Y K D E T K K A T V T E
BINARY_2110 T T Y K L I L N L K Q L K E E A I K E L V D A G I A E K Y I K L Y A N A K T V E G V W T Y K D E T K K F T V T E
BINARY_2111 T T Y K L I L N L K Q L K E E A I K E L V D A G I A E K Y I K L Y A N A K T V E G V W T Y K D E T K T A T V T E
BINARY_2112 T T Y K L I L N L K Q L K E E A I K E L V D A G I A E K Y I K L Y A N A K T V E G V W T Y K D E T K T F T V T E
BINARY_2113 T T Y K L I L N L K Q L K E E A I K E L V D A G I A E K Y F K L I A N A K T V E G V W T L K D E I L K A T V T E
BINARY_2114 T T Y K L I L N L K Q L K E E A I K E L V D A G I A E K Y F K L I A N A K T V E G V W T L K D E I L K F T V T E
BINARY_2115 T T Y K L I L N L K Q L K E E A I K E L V D A G I A E K Y F K L I A N A K T V E G V W T L K D E I L T A T V T E
BINARY_2116 T T Y K L I L N L K Q L K E E A I K E L V D A G I A E K Y F K L I A N A K T V E G V W T L K D E I L T F T V T E
BINARY_2117 T T Y K L I L N L K Q L K E E A I K E L V D A G I A E K Y F K L I A N A K T V E G V W T L K D E I K K A T V T E
BINARY_2118 T T Y K L I L N L K Q L K E E A I K E L V D A G I A E K Y F K L I A N A K T V E G V W T L K D E I K K F T V T E
BINARY_2119 T T Y K L I L N L K Q L K E E A I K E L V D A G I A E K Y F K L I A N A K T V E G V W T L K D E I K T A T V T E
BINARY_2120 T T Y K L I L N L K Q L K E E A I K E L V D A G I A E K Y F K L I A N A K T V E G V W T L K D E I K T F T V T E
BINARY_2121 T T Y K L I L N L K Q L K E E A I K E L V D A G I A E K Y F K L I A N A K T V E G V W T L K D E T L K A T V T E
BINARY_2122 T T Y K L I L N L K Q L K E E A I K E L V D A G I A E K Y F K L I A N A K T V E G V W T L K D E T L K F T V T E
BINARY_2123 T T Y K L I L N L K Q L K E E A I K E L V D A G I A E K Y F K L I A N A K T V E G V W T L K D E T L T A T V T E
BINARY_2124 T T Y K L I L N L K Q L K E E A I K E L V D A G I A E K Y F K L I A N A K T V E G V W T L K D E T L T F T V T E
BINARY_2125 T T Y K L I L N L K Q L K E E A I K E L V D A G I A E K Y F K L I A N A K T V E G V W T L K D E T K K A T V T E
BINARY_2126 T T Y K L I L N L K Q L K E E A I K E L V D A G I A E K Y F K L I A N A K T V E G V W T L K D E T K K F T V T E
BINARY_2127 T T Y K L I L N L K Q L K E E A I K E L V D A G I A E K Y F K L I A N A K T V E G V W T L K D E T K T A T V T E
BINARY_2128 T T Y K L I L N L K Q L K E E A I K E L V D A G I A E K Y F K L I A N A K T V E G V W T L K D E T K T F T V T E
BINARY_2129 T T Y K L I L N L K Q L K E E A I K E L V D A G I A E K Y F K L I A N A K T V E G V W T Y K D E I L K A T V T E
BINARY_2130 T T Y K L I L N L K Q L K E E A I K E L V D A G I A E K Y F K L I A N A K T V E G V W T Y K D E I L K F T V T E
BINARY_2131 T T Y K L I L N L K Q L K E E A I K E L V D A G I A E K Y F K L I A N A K T V E G V W T Y K D E I L T A T V T E
BINARY_2132 T T Y K L I L N L K Q L K E E A I K E L V D A G I A E K Y F K L I A N A K T V E G V W T Y K D E I L T F T V T E
BINARY_2133 T T Y K L I L N L K Q L K E E A I K E L V D A G I A E K Y F K L I A N A K T V E G V W T Y K D E I K K A T V T E
BINARY_2134 T T Y K L I L N L K Q L K E E A I K E L V D A G I A E K Y F K L I A N A K T V E G V W T Y K D E I K K F T V T E
BINARY_2135 T T Y K L I L N L K Q L K E E A I K E L V D A G I A E K Y F K L I A N A K T V E G V W T Y K D E I K T A T V T E
BINARY_2136 T T Y K L I L N L K Q L K E E A I K E L V D A G I A E K Y F K L I A N A K T V E G V W T Y K D E I K T F T V T E
BINARY_2137 T T Y K L I L N L K Q L K E E A I K E L V D A G I A E K Y F K L I A N A K T V E G V W T Y K D E T L K A T V T E
BINARY_2138 T T Y K L I L N L K Q L K E E A I K E L V D A G I A E K Y F K L I A N A K T V E G V W T Y K D E T L K F T V T E
BINARY_2139 T T Y K L I L N L K Q L K E E A I K E L V D A G I A E K Y F K L I A N A K T V E G V W T Y K D E T L T A T V T E
BINARY_2140 T T Y K L I L N L K Q L K E E A I K E L V D A G I A E K Y F K L I A N A K T V E G V W T Y K D E T L T F T V T E
BINARY_2141 T T Y K L I L N L K Q L K E E A I K E L V D A G I A E K Y F K L I A N A K T V E G V W T Y K D E T K K A T V T E
BINARY_2142 T T Y K L I L N L K Q L K E E A I K E L V D A G I A E K Y F K L I A N A K T V E G V W T Y K D E T K K F T V T E
BINARY_2143 T T Y K L I L N L K Q L K E E A I K E L V D A G I A E K Y F K L I A N A K T V E G V W T Y K D E T K T A T V T E
BINARY_2144 T T Y K L I L N L K Q L K E E A I K E L V D A G I A E K Y F K L I A N A K T V E G V W T Y K D E T K T F T V T E
BINARY_2145 T T Y K L I L N L K Q L K E E A I K E L V D A G I A E K Y F K L Y A N A K T V E G V W T L K D E I L K A T V T E
BINARY_2146 T T Y K L I L N L K Q L K E E A I K E L V D A G I A E K Y F K L Y A N A K T V E G V W T L K D E I L K F T V T E
BINARY_2147 T T Y K L I L N L K Q L K E E A I K E L V D A G I A E K Y F K L Y A N A K T V E G V W T L K D E I L T A T V T E
BINARY_2148 T T Y K L I L N L K Q L K E E A I K E L V D A G I A E K Y F K L Y A N A K T V E G V W T L K D E I L T F T V T E
BINARY_2149 T T Y K L I L N L K Q L K E E A I K E L V D A G I A E K Y F K L Y A N A K T V E G V W T L K D E I K K A T V T E
BINARY_2150 T T Y K L I L N L K Q L K E E A I K E L V D A G I A E K Y F K L Y A N A K T V E G V W T L K D E I K K F T V T E
BINARY_2151 T T Y K L I L N L K Q L K E E A I K E L V D A G I A E K Y F K L Y A N A K T V E G V W T L K D E I K T A T V T E
BINARY_2152 T T Y K L I L N L K Q L K E E A I K E L V D A G I A E K Y F K L Y A N A K T V E G V W T L K D E I K T F T V T E
BINARY_2153 T T Y K L I L N L K Q L K E E A I K E L V D A G I A E K Y F K L Y A N A K T V E G V W T L K D E T L K A T V T E
BINARY_2154 T T Y K L I L N L K Q L K E E A I K E L V D A G I A E K Y F K L Y A N A K T V E G V W T L K D E T L K F T V T E
BINARY_2155 T T Y K L I L N L K Q L K E E A I K E L V D A G I A E K Y F K L Y A N A K T V E G V W T L K D E T L T A T V T E
BINARY_2156 T T Y K L I L N L K Q L K E E A I K E L V D A G I A E K Y F K L Y A N A K T V E G V W T L K D E T L T F T V T E
BINARY_2157 T T Y K L I L N L K Q L K E E A I K E L V D A G I A E K Y F K L Y A N A K T V E G V W T L K D E T K K A T V T E
BINARY_2158 T T Y K L I L N L K Q L K E E A I K E L V D A G I A E K Y F K L Y A N A K T V E G V W T L K D E T K K F T V T E
BINARY_2159 T T Y K L I L N L K Q L K E E A I K E L V D A G I A E K Y F K L Y A N A K T V E G V W T L K D E T K T A T V T E
BINARY_2160 T T Y K L I L N L K Q L K E E A I K E L V D A G I A E K Y F K L Y A N A K T V E G V W T L K D E T K T F T V T E
BINARY_2161 T T Y K L I L N L K Q L K E E A I K E L V D A G I A E K Y F K L Y A N A K T V E G V W T Y K D E I L K A T V T E
BINARY_2162 T T Y K L I L N L K Q L K E E A I K E L V D A G I A E K Y F K L Y A N A K T V E G V W T Y K D E I L K F T V T E
BINARY_2163 T T Y K L I L N L K Q L K E E A I K E L V D A G I A E K Y F K L Y A N A K T V E G V W T Y K D E I L T A T V T E
BINARY_2164 T T Y K L I L N L K Q L K E E A I K E L V D A G I A E K Y F K L Y A N A K T V E G V W T Y K D E I L T F T V T E
BINARY_2165 T T Y K L I L N L K Q L K E E A I K E L V D A G I A E K Y F K L Y A N A K T V E G V W T Y K D E I K K A T V T E
BINARY_2166 T T Y K L I L N L K Q L K E E A I K E L V D A G I A E K Y F K L Y A N A K T V E G V W T Y K D E I K K F T V T E
BINARY_2167 T T Y K L I L N L K Q L K E E A I K E L V D A G I A E K Y F K L Y A N A K T V E G V W T Y K D E I K T A T V T E
BINARY_2168 T T Y K L I L N L K Q L K E E A I K E L V D A G I A E K Y F K L Y A N A K T V E G V W T Y K D E I K T F T V T E
BINARY_2169 T T Y K L I L N L K Q L K E E A I K E L V D A G I A E K Y F K L Y A N A K T V E G V W T Y K D E T L K A T V T E
BINARY_2170 T T Y K L I L N L K Q L K E E A I K E L V D A G I A E K Y F K L Y A N A K T V E G V W T Y K D E T L K F T V T E
BINARY_2171 T T Y K L I L N L K Q L K E E A I K E L V D A G I A E K Y F K L Y A N A K T V E G V W T Y K D E T L T A T V T E
BINARY_2172 T T Y K L I L N L K Q L K E E A I K E L V D A G I A E K Y F K L Y A N A K T V E G V W T Y K D E T L T F T V T E
BINARY_2173 T T Y K L I L N L K Q L K E E A I K E L V D A G I A E K Y F K L Y A N A K T V E G V W T Y K D E T K K A T V T E
BINARY_2174 T T Y K L I L N L K Q L K E E A I K E L V D A G I A E K Y F K L Y A N A K T V E G V W T Y K D E T K K F T V T E
BINARY_2175 T T Y K L I L N L K Q L K E E A I K E L V D A G I A E K Y F K L Y A N A K T V E G V W T Y K D E T K T A T V T E
BINARY_2176 T T Y K L I L N L K Q L K E E A I K E L V D A G I A E K Y F K L Y A N A K T V E G V W T Y K D E T K T F T V T E
BINARY_2177 T T Y K L I L N L K Q L K E E A I K E L V D A G T A E K Y I K L I A N A K T V E G V W T L K D E I L K A T V T E
BINARY_2178 T T Y K L I L N L K Q L K E E A I K E L V D A G T A E K Y I K L I A N A K T V E G V W T L K D E I L K F T V T E
BINARY_2179 T T Y K L I L N L K Q L K E E A I K E L V D A G T A E K Y I K L I A N A K T V E G V W T L K D E I L T A T V T E
BINARY_2180 T T Y K L I L N L K Q L K E E A I K E L V D A G T A E K Y I K L I A N A K T V E G V W T L K D E I L T F T V T E
BINARY_2181 T T Y K L I L N L K Q L K E E A I K E L V D A G T A E K Y I K L I A N A K T V E G V W T L K D E I K K A T V T E
BINARY_2182 T T Y K L I L N L K Q L K E E A I K E L V D A G T A E K Y I K L I A N A K T V E G V W T L K D E I K K F T V T E
BINARY_2183 T T Y K L I L N L K Q L K E E A I K E L V D A G T A E K Y I K L I A N A K T V E G V W T L K D E I K T A T V T E
BINARY_2184 T T Y K L I L N L K Q L K E E A I K E L V D A G T A E K Y I K L I A N A K T V E G V W T L K D E I K T F T V T E
BINARY_2185 T T Y K L I L N L K Q L K E E A I K E L V D A G T A E K Y I K L I A N A K T V E G V W T L K D E T L K A T V T E
BINARY_2186 T T Y K L I L N L K Q L K E E A I K E L V D A G T A E K Y I K L I A N A K T V E G V W T L K D E T L K F T V T E
BINARY_2187 T T Y K L I L N L K Q L K E E A I K E L V D A G T A E K Y I K L I A N A K T V E G V W T L K D E T L T A T V T E
BINARY_2188 T T Y K L I L N L K Q L K E E A I K E L V D A G T A E K Y I K L I A N A K T V E G V W T L K D E T L T F T V T E
BINARY_2189 T T Y K L I L N L K Q L K E E A I K E L V D A G T A E K Y I K L I A N A K T V E G V W T L K D E T K K A T V T E
BINARY_2190 T T Y K L I L N L K Q L K E E A I K E L V D A G T A E K Y I K L I A N A K T V E G V W T L K D E T K K F T V T E
BINARY_2191 T T Y K L I L N L K Q L K E E A I K E L V D A G T A E K Y I K L I A N A K T V E G V W T L K D E T K T A T V T E
BINARY_2192 T T Y K L I L N L K Q L K E E A I K E L V D A G T A E K Y I K L I A N A K T V E G V W T L K D E T K T F T V T E
BINARY_2193 T T Y K L I L N L K Q L K E E A I K E L V D A G T A E K Y I K L I A N A K T V E G V W T Y K D E I L K A T V T E
BINARY_2194 T T Y K L I L N L K Q L K E E A I K E L V D A G T A E K Y I K L I A N A K T V E G V W T Y K D E I L K F T V T E
BINARY_2195 T T Y K L I L N L K Q L K E E A I K E L V D A G T A E K Y I K L I A N A K T V E G V W T Y K D E I L T A T V T E
BINARY_2196 T T Y K L I L N L K Q L K E E A I K E L V D A G T A E K Y I K L I A N A K T V E G V W T Y K D E I L T F T V T E
BINARY_2197 T T Y K L I L N L K Q L K E E A I K E L V D A G T A E K Y I K L I A N A K T V E G V W T Y K D E I K K A T V T E
BINARY_2198 T T Y K L I L N L K Q L K E E A I K E L V D A G T A E K Y I K L I A N A K T V E G V W T Y K D E I K K F T V T E
BINARY_2199 T T Y K L I L N L K Q L K E E A I K E L V D A G T A E K Y I K L I A N A K T V E G V W T Y K D E I K T A T V T E
BINARY_2200 T T Y K L I L N L K Q L K E E A I K E L V D A G T A E K Y I K L I A N A K T V E G V W T Y K D E I K T F T V T E
BINARY_2201 T T Y K L I L N L K Q L K E E A I K E L V D A G T A E K Y I K L I A N A K T V E G V W T Y K D E T L K A T V T E

BINARY_2202 T T Y K L I L N L K Q L K E E A I K E L V D A G T A E K Y I K L I A N A K T V E G V W T Y K D E T L K F T V T E
BINARY_2203 T T Y K L I L N L K Q L K E E A I K E L V D A G T A E K Y I K L I A N A K T V E G V W T Y K D E T L T A T V T E
BINARY_2204 T T Y K L I L N L K Q L K E E A I K E L V D A G T A E K Y I K L I A N A K T V E G V W T Y K D E T L T F T V T E
BINARY_2205 T T Y K L I L N L K Q L K E E A I K E L V D A G T A E K Y I K L I A N A K T V E G V W T Y K D E T K K A T V T E
BINARY_2206 T T Y K L I L N L K Q L K E E A I K E L V D A G T A E K Y I K L I A N A K T V E G V W T Y K D E T K K F T V T E
BINARY_2207 T T Y K L I L N L K Q L K E E A I K E L V D A G T A E K Y I K L I A N A K T V E G V W T Y K D E T K T A T V T E
BINARY_2208 T T Y K L I L N L K Q L K E E A I K E L V D A G T A E K Y I K L I A N A K T V E G V W T Y K D E T K T F T V T E
BINARY_2209 T T Y K L I L N L K Q L K E E A I K E L V D A G T A E K Y I K L Y A N A K T V E G V W T L K D E I L K A T V T E
BINARY_2210 T T Y K L I L N L K Q L K E E A I K E L V D A G T A E K Y I K L Y A N A K T V E G V W T L K D E I L K F T V T E
BINARY_2211 T T Y K L I L N L K Q L K E E A I K E L V D A G T A E K Y I K L Y A N A K T V E G V W T L K D E I L T A T V T E
BINARY_2212 T T Y K L I L N L K Q L K E E A I K E L V D A G T A E K Y I K L Y A N A K T V E G V W T L K D E I L T F T V T E
BINARY_2213 T T Y K L I L N L K Q L K E E A I K E L V D A G T A E K Y I K L Y A N A K T V E G V W T L K D E I K K A T V T E
BINARY_2214 T T Y K L I L N L K Q L K E E A I K E L V D A G T A E K Y I K L Y A N A K T V E G V W T L K D E I K K F T V T E
BINARY_2215 T T Y K L I L N L K Q L K E E A I K E L V D A G T A E K Y I K L Y A N A K T V E G V W T L K D E I K T A T V T E
BINARY_2216 T T Y K L I L N L K Q L K E E A I K E L V D A G T A E K Y I K L Y A N A K T V E G V W T L K D E I K T F T V T E
BINARY_2217 T T Y K L I L N L K Q L K E E A I K E L V D A G T A E K Y I K L Y A N A K T V E G V W T L K D E T L K A T V T E
BINARY_2218 T T Y K L I L N L K Q L K E E A I K E L V D A G T A E K Y I K L Y A N A K T V E G V W T L K D E T L K F T V T E
BINARY_2219 T T Y K L I L N L K Q L K E E A I K E L V D A G T A E K Y I K L Y A N A K T V E G V W T L K D E T L T A T V T E
BINARY_2220 T T Y K L I L N L K Q L K E E A I K E L V D A G T A E K Y I K L Y A N A K T V E G V W T L K D E T L T F T V T E
BINARY_2221 T T Y K L I L N L K Q L K E E A I K E L V D A G T A E K Y I K L Y A N A K T V E G V W T L K D E T K K A T V T E
BINARY_2222 T T Y K L I L N L K Q L K E E A I K E L V D A G T A E K Y I K L Y A N A K T V E G V W T L K D E T K K F T V T E
BINARY_2223 T T Y K L I L N L K Q L K E E A I K E L V D A G T A E K Y I K L Y A N A K T V E G V W T L K D E T K T A T V T E
BINARY_2224 T T Y K L I L N L K Q L K E E A I K E L V D A G T A E K Y I K L Y A N A K T V E G V W T L K D E T K T F T V T E
BINARY_2225 T T Y K L I L N L K Q L K E E A I K E L V D A G T A E K Y I K L Y A N A K T V E G V W T Y K D E I L K A T V T E
BINARY_2226 T T Y K L I L N L K Q L K E E A I K E L V D A G T A E K Y I K L Y A N A K T V E G V W T Y K D E I L K F T V T E
BINARY_2227 T T Y K L I L N L K Q L K E E A I K E L V D A G T A E K Y I K L Y A N A K T V E G V W T Y K D E I L T A T V T E
BINARY_2228 T T Y K L I L N L K Q L K E E A I K E L V D A G T A E K Y I K L Y A N A K T V E G V W T Y K D E I L T F T V T E
BINARY_2229 T T Y K L I L N L K Q L K E E A I K E L V D A G T A E K Y I K L Y A N A K T V E G V W T Y K D E I K K A T V T E
BINARY_2230 T T Y K L I L N L K Q L K E E A I K E L V D A G T A E K Y I K L Y A N A K T V E G V W T Y K D E I K K F T V T E
BINARY_2231 T T Y K L I L N L K Q L K E E A I K E L V D A G T A E K Y I K L Y A N A K T V E G V W T Y K D E I K T A T V T E
BINARY_2232 T T Y K L I L N L K Q L K E E A I K E L V D A G T A E K Y I K L Y A N A K T V E G V W T Y K D E I K T F T V T E
BINARY_2233 T T Y K L I L N L K Q L K E E A I K E L V D A G T A E K Y I K L Y A N A K T V E G V W T Y K D E T L K A T V T E
BINARY_2234 T T Y K L I L N L K Q L K E E A I K E L V D A G T A E K Y I K L Y A N A K T V E G V W T Y K D E T L K F T V T E
BINARY_2235 T T Y K L I L N L K Q L K E E A I K E L V D A G T A E K Y I K L Y A N A K T V E G V W T Y K D E T L T A T V T E
BINARY_2236 T T Y K L I L N L K Q L K E E A I K E L V D A G T A E K Y I K L Y A N A K T V E G V W T Y K D E T L T F T V T E
BINARY_2237 T T Y K L I L N L K Q L K E E A I K E L V D A G T A E K Y I K L Y A N A K T V E G V W T Y K D E T K K A T V T E
BINARY_2238 T T Y K L I L N L K Q L K E E A I K E L V D A G T A E K Y I K L Y A N A K T V E G V W T Y K D E T K K F T V T E
BINARY_2239 T T Y K L I L N L K Q L K E E A I K E L V D A G T A E K Y I K L Y A N A K T V E G V W T Y K D E T K T A T V T E
BINARY_2240 T T Y K L I L N L K Q L K E E A I K E L V D A G T A E K Y I K L Y A N A K T V E G V W T Y K D E T K T F T V T E
BINARY_2241 T T Y K L I L N L K Q L K E E A I K E L V D A G T A E K Y F K L I A N A K T V E G V W T L K D E I L K A T V T E
BINARY_2242 T T Y K L I L N L K Q L K E E A I K E L V D A G T A E K Y F K L I A N A K T V E G V W T L K D E I L K F T V T E
BINARY_2243 T T Y K L I L N L K Q L K E E A I K E L V D A G T A E K Y F K L I A N A K T V E G V W T L K D E I L T A T V T E
BINARY_2244 T T Y K L I L N L K Q L K E E A I K E L V D A G T A E K Y F K L I A N A K T V E G V W T L K D E I L T F T V T E
BINARY_2245 T T Y K L I L N L K Q L K E E A I K E L V D A G T A E K Y F K L I A N A K T V E G V W T L K D E I K K A T V T E
BINARY_2246 T T Y K L I L N L K Q L K E E A I K E L V D A G T A E K Y F K L I A N A K T V E G V W T L K D E I K K F T V T E
BINARY_2247 T T Y K L I L N L K Q L K E E A I K E L V D A G T A E K Y F K L I A N A K T V E G V W T L K D E I K T A T V T E
BINARY_2248 T T Y K L I L N L K Q L K E E A I K E L V D A G T A E K Y F K L I A N A K T V E G V W T L K D E I K T F T V T E
BINARY_2249 T T Y K L I L N L K Q L K E E A I K E L V D A G T A E K Y F K L I A N A K T V E G V W T L K D E T L K A T V T E
BINARY_2250 T T Y K L I L N L K Q L K E E A I K E L V D A G T A E K Y F K L I A N A K T V E G V W T L K D E T L K F T V T E
BINARY_2251 T T Y K L I L N L K Q L K E E A I K E L V D A G T A E K Y F K L I A N A K T V E G V W T L K D E T L T A T V T E
BINARY_2252 T T Y K L I L N L K Q L K E E A I K E L V D A G T A E K Y F K L I A N A K T V E G V W T L K D E T L T F T V T E
BINARY_2253 T T Y K L I L N L K Q L K E E A I K E L V D A G T A E K Y F K L I A N A K T V E G V W T L K D E T K K A T V T E
BINARY_2254 T T Y K L I L N L K Q L K E E A I K E L V D A G T A E K Y F K L I A N A K T V E G V W T L K D E T K K F T V T E
BINARY_2255 T T Y K L I L N L K Q L K E E A I K E L V D A G T A E K Y F K L I A N A K T V E G V W T L K D E T K T A T V T E
BINARY_2256 T T Y K L I L N L K Q L K E E A I K E L V D A G T A E K Y F K L I A N A K T V E G V W T L K D E T K T F T V T E
BINARY_2257 T T Y K L I L N L K Q L K E E A I K E L V D A G T A E K Y F K L I A N A K T V E G V W T Y K D E I L K A T V T E
BINARY_2258 T T Y K L I L N L K Q L K E E A I K E L V D A G T A E K Y F K L I A N A K T V E G V W T Y K D E I L K F T V T E
BINARY_2259 T T Y K L I L N L K Q L K E E A I K E L V D A G T A E K Y F K L I A N A K T V E G V W T Y K D E I L T A T V T E
BINARY_2260 T T Y K L I L N L K Q L K E E A I K E L V D A G T A E K Y F K L I A N A K T V E G V W T Y K D E I L T F T V T E
BINARY_2261 T T Y K L I L N L K Q L K E E A I K E L V D A G T A E K Y F K L I A N A K T V E G V W T Y K D E I K K A T V T E
BINARY_2262 T T Y K L I L N L K Q L K E E A I K E L V D A G T A E K Y F K L I A N A K T V E G V W T Y K D E I K K F T V T E
BINARY_2263 T T Y K L I L N L K Q L K E E A I K E L V D A G T A E K Y F K L I A N A K T V E G V W T Y K D E I K T A T V T E
BINARY_2264 T T Y K L I L N L K Q L K E E A I K E L V D A G T A E K Y F K L I A N A K T V E G V W T Y K D E I K T F T V T E
BINARY_2265 T T Y K L I L N L K Q L K E E A I K E L V D A G T A E K Y F K L I A N A K T V E G V W T Y K D E T L K A T V T E
BINARY_2266 T T Y K L I L N L K Q L K E E A I K E L V D A G T A E K Y F K L I A N A K T V E G V W T Y K D E T L K F T V T E
BINARY_2267 T T Y K L I L N L K Q L K E E A I K E L V D A G T A E K Y F K L I A N A K T V E G V W T Y K D E T L T A T V T E
BINARY_2268 T T Y K L I L N L K Q L K E E A I K E L V D A G T A E K Y F K L I A N A K T V E G V W T Y K D E T L T F T V T E
BINARY_2269 T T Y K L I L N L K Q L K E E A I K E L V D A G T A E K Y F K L I A N A K T V E G V W T Y K D E T K K A T V T E
BINARY_2270 T T Y K L I L N L K Q L K E E A I K E L V D A G T A E K Y F K L I A N A K T V E G V W T Y K D E T K K F T V T E
BINARY_2271 T T Y K L I L N L K Q L K E E A I K E L V D A G T A E K Y F K L I A N A K T V E G V W T Y K D E T K T A T V T E
BINARY_2272 T T Y K L I L N L K Q L K E E A I K E L V D A G T A E K Y F K L I A N A K T V E G V W T Y K D E T K T F T V T E
BINARY_2273 T T Y K L I L N L K Q L K E E A I K E L V D A G T A E K Y F K L Y A N A K T V E G V W T L K D E I L K A T V T E
BINARY_2274 T T Y K L I L N L K Q L K E E A I K E L V D A G T A E K Y F K L Y A N A K T V E G V W T L K D E I L K F T V T E
BINARY_2275 T T Y K L I L N L K Q L K E E A I K E L V D A G T A E K Y F K L Y A N A K T V E G V W T L K D E I L T A T V T E
BINARY_2276 T T Y K L I L N L K Q L K E E A I K E L V D A G T A E K Y F K L Y A N A K T V E G V W T L K D E I L T F T V T E
BINARY_2277 T T Y K L I L N L K Q L K E E A I K E L V D A G T A E K Y F K L Y A N A K T V E G V W T L K D E I K K A T V T E
BINARY_2278 T T Y K L I L N L K Q L K E E A I K E L V D A G T A E K Y F K L Y A N A K T V E G V W T L K D E I K K F T V T E
BINARY_2279 T T Y K L I L N L K Q L K E E A I K E L V D A G T A E K Y F K L Y A N A K T V E G V W T L K D E I K T A T V T E
BINARY_2280 T T Y K L I L N L K Q L K E E A I K E L V D A G T A E K Y F K L Y A N A K T V E G V W T L K D E I K T F T V T E
BINARY_2281 T T Y K L I L N L K Q L K E E A I K E L V D A G T A E K Y F K L Y A N A K T V E G V W T L K D E T L K A T V T E
BINARY_2282 T T Y K L I L N L K Q L K E E A I K E L V D A G T A E K Y F K L Y A N A K T V E G V W T L K D E T L K F T V T E
BINARY_2283 T T Y K L I L N L K Q L K E E A I K E L V D A G T A E K Y F K L Y A N A K T V E G V W T L K D E T L T A T V T E
BINARY_2284 T T Y K L I L N L K Q L K E E A I K E L V D A G T A E K Y F K L Y A N A K T V E G V W T L K D E T L T F T V T E
BINARY_2285 T T Y K L I L N L K Q L K E E A I K E L V D A G T A E K Y F K L Y A N A K T V E G V W T L K D E T K K A T V T E
BINARY_2286 T T Y K L I L N L K Q L K E E A I K E L V D A G T A E K Y F K L Y A N A K T V E G V W T L K D E T K K F T V T E
BINARY_2287 T T Y K L I L N L K Q L K E E A I K E L V D A G T A E K Y F K L Y A N A K T V E G V W T L K D E T K T A T V T E
BINARY_2288 T T Y K L I L N L K Q L K E E A I K E L V D A G T A E K Y F K L Y A N A K T V E G V W T L K D E T K T F T V T E
BINARY_2289 T T Y K L I L N L K Q L K E E A I K E L V D A G T A E K Y F K L Y A N A K T V E G V W T Y K D E I L K A T V T E
BINARY_2290 T T Y K L I L N L K Q L K E E A I K E L V D A G T A E K Y F K L Y A N A K T V E G V W T Y K D E I L K F T V T E
BINARY_2291 T T Y K L I L N L K Q L K E E A I K E L V D A G T A E K Y F K L Y A N A K T V E G V W T Y K D E I L T A T V T E
BINARY_2292 T T Y K L I L N L K Q L K E E A I K E L V D A G T A E K Y F K L Y A N A K T V E G V W T Y K D E I L T F T V T E
BINARY_2293 T T Y K L I L N L K Q L K E E A I K E L V D A G T A E K Y F K L Y A N A K T V E G V W T Y K D E I K K A T V T E
BINARY_2294 T T Y K L I L N L K Q L K E E A I K E L V D A G T A E K Y F K L Y A N A K T V E G V W T Y K D E I K K F T V T E
BINARY_2295 T T Y K L I L N L K Q L K E E A I K E L V D A G T A E K Y F K L Y A N A K T V E G V W T Y K D E I K T A T V T E
BINARY_2296 T T Y K L I L N L K Q L K E E A I K E L V D A G T A E K Y F K L Y A N A K T V E G V W T Y K D E I K T F T V T E

BINARY_2297 T T Y K L I L N L K Q L K E E A I K E L V D A G T A E K Y F K L Y A N A K T V E G V W T Y K D E T L K A T V T E
BINARY_2298 T T Y K L I L N L K Q L K E E A I K E L V D A G T A E K Y F K L Y A N A K T V E G V W T Y K D E T L K F T V T E
BINARY_2299 T T Y K L I L N L K Q L K E E A I K E L V D A G T A E K Y F K L Y A N A K T V E G V W T Y K D E T L T A T V T E
BINARY_2300 T T Y K L I L N L K Q L K E E A I K E L V D A G T A E K Y F K L Y A N A K T V E G V W T Y K D E T L T F T V T E
BINARY_2301 T T Y K L I L N L K Q L K E E A I K E L V D A G T A E K Y F K L Y A N A K T V E G V W T Y K D E T K K A T V T E
BINARY_2302 T T Y K L I L N L K Q L K E E A I K E L V D A G T A E K Y F K L Y A N A K T V E G V W T Y K D E T K K F T V T E
BINARY_2303 T T Y K L I L N L K Q L K E E A I K E L V D A G T A E K Y F K L Y A N A K T V E G V W T Y K D E T K T A T V T E
BINARY_2304 T T Y K L I L N L K Q L K E E A I K E L V D A G T A E K Y F K L Y A N A K T V E G V W T Y K D E T K T F T V T E
BINARY_2305 T T Y K L I L N L K Q L K E E A I K E L V D A A I A E K Y I K L I A N A K T V E G V W T L K D E I L K A T V T E
BINARY_2306 T T Y K L I L N L K Q L K E E A I K E L V D A A I A E K Y I K L I A N A K T V E G V W T L K D E I L K F T V T E
BINARY_2307 T T Y K L I L N L K Q L K E E A I K E L V D A A I A E K Y I K L I A N A K T V E G V W T L K D E I L T A T V T E
BINARY_2308 T T Y K L I L N L K Q L K E E A I K E L V D A A I A E K Y I K L I A N A K T V E G V W T L K D E I L T F T V T E
BINARY_2309 T T Y K L I L N L K Q L K E E A I K E L V D A A I A E K Y I K L I A N A K T V E G V W T L K D E I K K A T V T E
BINARY_2310 T T Y K L I L N L K Q L K E E A I K E L V D A A I A E K Y I K L I A N A K T V E G V W T L K D E I K K F T V T E
BINARY_2311 T T Y K L I L N L K Q L K E E A I K E L V D A A I A E K Y I K L I A N A K T V E G V W T L K D E I K T A T V T E
BINARY_2312 T T Y K L I L N L K Q L K E E A I K E L V D A A I A E K Y I K L I A N A K T V E G V W T L K D E I K T F T V T E
BINARY_2313 T T Y K L I L N L K Q L K E E A I K E L V D A A I A E K Y I K L I A N A K T V E G V W T L K D E T L K A T V T E
BINARY_2314 T T Y K L I L N L K Q L K E E A I K E L V D A A I A E K Y I K L I A N A K T V E G V W T L K D E T L K F T V T E
BINARY_2315 T T Y K L I L N L K Q L K E E A I K E L V D A A I A E K Y I K L I A N A K T V E G V W T L K D E T L T A T V T E
BINARY_2316 T T Y K L I L N L K Q L K E E A I K E L V D A A I A E K Y I K L I A N A K T V E G V W T L K D E T L T F T V T E
BINARY_2317 T T Y K L I L N L K Q L K E E A I K E L V D A A I A E K Y I K L I A N A K T V E G V W T L K D E T K K A T V T E
BINARY_2318 T T Y K L I L N L K Q L K E E A I K E L V D A A I A E K Y I K L I A N A K T V E G V W T L K D E T K K F T V T E
BINARY_2319 T T Y K L I L N L K Q L K E E A I K E L V D A A I A E K Y I K L I A N A K T V E G V W T L K D E T K T A T V T E
BINARY_2320 T T Y K L I L N L K Q L K E E A I K E L V D A A I A E K Y I K L I A N A K T V E G V W T L K D E T K T F T V T E
BINARY_2321 T T Y K L I L N L K Q L K E E A I K E L V D A A I A E K Y I K L I A N A K T V E G V W T Y K D E I L K A T V T E
BINARY_2322 T T Y K L I L N L K Q L K E E A I K E L V D A A I A E K Y I K L I A N A K T V E G V W T Y K D E I L K F T V T E
BINARY_2323 T T Y K L I L N L K Q L K E E A I K E L V D A A I A E K Y I K L I A N A K T V E G V W T Y K D E I L T A T V T E
BINARY_2324 T T Y K L I L N L K Q L K E E A I K E L V D A A I A E K Y I K L I A N A K T V E G V W T Y K D E I L T F T V T E
BINARY_2325 T T Y K L I L N L K Q L K E E A I K E L V D A A I A E K Y I K L I A N A K T V E G V W T Y K D E I K K A T V T E
BINARY_2326 T T Y K L I L N L K Q L K E E A I K E L V D A A I A E K Y I K L I A N A K T V E G V W T Y K D E I K K F T V T E
BINARY_2327 T T Y K L I L N L K Q L K E E A I K E L V D A A I A E K Y I K L I A N A K T V E G V W T Y K D E I K T A T V T E
BINARY_2328 T T Y K L I L N L K Q L K E E A I K E L V D A A I A E K Y I K L I A N A K T V E G V W T Y K D E I K T F T V T E
BINARY_2329 T T Y K L I L N L K Q L K E E A I K E L V D A A I A E K Y I K L I A N A K T V E G V W T Y K D E T L K A T V T E
BINARY_2330 T T Y K L I L N L K Q L K E E A I K E L V D A A I A E K Y I K L I A N A K T V E G V W T Y K D E T L K F T V T E
BINARY_2331 T T Y K L I L N L K Q L K E E A I K E L V D A A I A E K Y I K L I A N A K T V E G V W T Y K D E T L T A T V T E
BINARY_2332 T T Y K L I L N L K Q L K E E A I K E L V D A A I A E K Y I K L I A N A K T V E G V W T Y K D E T L T F T V T E
BINARY_2333 T T Y K L I L N L K Q L K E E A I K E L V D A A I A E K Y I K L I A N A K T V E G V W T Y K D E T K K A T V T E
BINARY_2334 T T Y K L I L N L K Q L K E E A I K E L V D A A I A E K Y I K L I A N A K T V E G V W T Y K D E T K K F T V T E
BINARY_2335 T T Y K L I L N L K Q L K E E A I K E L V D A A I A E K Y I K L I A N A K T V E G V W T Y K D E T K T A T V T E
BINARY_2336 T T Y K L I L N L K Q L K E E A I K E L V D A A I A E K Y I K L I A N A K T V E G V W T Y K D E T K T F T V T E
BINARY_2337 T T Y K L I L N L K Q L K E E A I K E L V D A A I A E K Y I K L Y A N A K T V E G V W T L K D E I L K A T V T E
BINARY_2338 T T Y K L I L N L K Q L K E E A I K E L V D A A I A E K Y I K L Y A N A K T V E G V W T L K D E I L K F T V T E
BINARY_2339 T T Y K L I L N L K Q L K E E A I K E L V D A A I A E K Y I K L Y A N A K T V E G V W T L K D E I L T A T V T E
BINARY_2340 T T Y K L I L N L K Q L K E E A I K E L V D A A I A E K Y I K L Y A N A K T V E G V W T L K D E I L T F T V T E
BINARY_2341 T T Y K L I L N L K Q L K E E A I K E L V D A A I A E K Y I K L Y A N A K T V E G V W T L K D E I K K A T V T E
BINARY_2342 T T Y K L I L N L K Q L K E E A I K E L V D A A I A E K Y I K L Y A N A K T V E G V W T L K D E I K K F T V T E
BINARY_2343 T T Y K L I L N L K Q L K E E A I K E L V D A A I A E K Y I K L Y A N A K T V E G V W T L K D E I K T A T V T E
BINARY_2344 T T Y K L I L N L K Q L K E E A I K E L V D A A I A E K Y I K L Y A N A K T V E G V W T L K D E I K T F T V T E
BINARY_2345 T T Y K L I L N L K Q L K E E A I K E L V D A A I A E K Y I K L Y A N A K T V E G V W T L K D E T L K A T V T E
BINARY_2346 T T Y K L I L N L K Q L K E E A I K E L V D A A I A E K Y I K L Y A N A K T V E G V W T L K D E T L K F T V T E
BINARY_2347 T T Y K L I L N L K Q L K E E A I K E L V D A A I A E K Y I K L Y A N A K T V E G V W T L K D E T L T A T V T E
BINARY_2348 T T Y K L I L N L K Q L K E E A I K E L V D A A I A E K Y I K L Y A N A K T V E G V W T L K D E T L T F T V T E
BINARY_2349 T T Y K L I L N L K Q L K E E A I K E L V D A A I A E K Y I K L Y A N A K T V E G V W T L K D E T K K A T V T E
BINARY_2350 T T Y K L I L N L K Q L K E E A I K E L V D A A I A E K Y I K L Y A N A K T V E G V W T L K D E T K K F T V T E
BINARY_2351 T T Y K L I L N L K Q L K E E A I K E L V D A A I A E K Y I K L Y A N A K T V E G V W T L K D E T K T A T V T E
BINARY_2352 T T Y K L I L N L K Q L K E E A I K E L V D A A I A E K Y I K L Y A N A K T V E G V W T L K D E T K T F T V T E
BINARY_2353 T T Y K L I L N L K Q L K E E A I K E L V D A A I A E K Y I K L Y A N A K T V E G V W T Y K D E I L K A T V T E
BINARY_2354 T T Y K L I L N L K Q L K E E A I K E L V D A A I A E K Y I K L Y A N A K T V E G V W T Y K D E I L K F T V T E
BINARY_2355 T T Y K L I L N L K Q L K E E A I K E L V D A A I A E K Y I K L Y A N A K T V E G V W T Y K D E I L T A T V T E
BINARY_2356 T T Y K L I L N L K Q L K E E A I K E L V D A A I A E K Y I K L Y A N A K T V E G V W T Y K D E I L T F T V T E
BINARY_2357 T T Y K L I L N L K Q L K E E A I K E L V D A A I A E K Y I K L Y A N A K T V E G V W T Y K D E I K K A T V T E
BINARY_2358 T T Y K L I L N L K Q L K E E A I K E L V D A A I A E K Y I K L Y A N A K T V E G V W T Y K D E I K K F T V T E
BINARY_2359 T T Y K L I L N L K Q L K E E A I K E L V D A A I A E K Y I K L Y A N A K T V E G V W T Y K D E I K T A T V T E
BINARY_2360 T T Y K L I L N L K Q L K E E A I K E L V D A A I A E K Y I K L Y A N A K T V E G V W T Y K D E I K T F T V T E
BINARY_2361 T T Y K L I L N L K Q L K E E A I K E L V D A A I A E K Y I K L Y A N A K T V E G V W T Y K D E T L K A T V T E
BINARY_2362 T T Y K L I L N L K Q L K E E A I K E L V D A A I A E K Y I K L Y A N A K T V E G V W T Y K D E T L K F T V T E
BINARY_2363 T T Y K L I L N L K Q L K E E A I K E L V D A A I A E K Y I K L Y A N A K T V E G V W T Y K D E T L T A T V T E
BINARY_2364 T T Y K L I L N L K Q L K E E A I K E L V D A A I A E K Y I K L Y A N A K T V E G V W T Y K D E T L T F T V T E
BINARY_2365 T T Y K L I L N L K Q L K E E A I K E L V D A A I A E K Y I K L Y A N A K T V E G V W T Y K D E T K K A T V T E
BINARY_2366 T T Y K L I L N L K Q L K E E A I K E L V D A A I A E K Y I K L Y A N A K T V E G V W T Y K D E T K K F T V T E
BINARY_2367 T T Y K L I L N L K Q L K E E A I K E L V D A A I A E K Y I K L Y A N A K T V E G V W T Y K D E T K T A T V T E
BINARY_2368 T T Y K L I L N L K Q L K E E A I K E L V D A A I A E K Y I K L Y A N A K T V E G V W T Y K D E T K T F T V T E
BINARY_2369 T T Y K L I L N L K Q L K E E A I K E L V D A A I A E K Y F K L I A N A K T V E G V W T L K D E I L K A T V T E
BINARY_2370 T T Y K L I L N L K Q L K E E A I K E L V D A A I A E K Y F K L I A N A K T V E G V W T L K D E I L K F T V T E
BINARY_2371 T T Y K L I L N L K Q L K E E A I K E L V D A A I A E K Y F K L I A N A K T V E G V W T L K D E I L T A T V T E
BINARY_2372 T T Y K L I L N L K Q L K E E A I K E L V D A A I A E K Y F K L I A N A K T V E G V W T L K D E I L T F T V T E
BINARY_2373 T T Y K L I L N L K Q L K E E A I K E L V D A A I A E K Y F K L I A N A K T V E G V W T L K D E I K K A T V T E
BINARY_2374 T T Y K L I L N L K Q L K E E A I K E L V D A A I A E K Y F K L I A N A K T V E G V W T L K D E I K K F T V T E
BINARY_2375 T T Y K L I L N L K Q L K E E A I K E L V D A A I A E K Y F K L I A N A K T V E G V W T L K D E I K T A T V T E
BINARY_2376 T T Y K L I L N L K Q L K E E A I K E L V D A A I A E K Y F K L I A N A K T V E G V W T L K D E I K T F T V T E
BINARY_2377 T T Y K L I L N L K Q L K E E A I K E L V D A A I A E K Y F K L I A N A K T V E G V W T L K D E T L K A T V T E
BINARY_2378 T T Y K L I L N L K Q L K E E A I K E L V D A A I A E K Y F K L I A N A K T V E G V W T L K D E T L K F T V T E
BINARY_2379 T T Y K L I L N L K Q L K E E A I K E L V D A A I A E K Y F K L I A N A K T V E G V W T L K D E T L T A T V T E
BINARY_2380 T T Y K L I L N L K Q L K E E A I K E L V D A A I A E K Y F K L I A N A K T V E G V W T L K D E T L T F T V T E
BINARY_2381 T T Y K L I L N L K Q L K E E A I K E L V D A A I A E K Y F K L I A N A K T V E G V W T L K D E T K K A T V T E
BINARY_2382 T T Y K L I L N L K Q L K E E A I K E L V D A A I A E K Y F K L I A N A K T V E G V W T L K D E T K K F T V T E
BINARY_2383 T T Y K L I L N L K Q L K E E A I K E L V D A A I A E K Y F K L I A N A K T V E G V W T L K D E T K T A T V T E
BINARY_2384 T T Y K L I L N L K Q L K E E A I K E L V D A A I A E K Y F K L I A N A K T V E G V W T L K D E T K T F T V T E
BINARY_2385 T T Y K L I L N L K Q L K E E A I K E L V D A A I A E K Y F K L I A N A K T V E G V W T Y K D E I L K A T V T E
BINARY_2386 T T Y K L I L N L K Q L K E E A I K E L V D A A I A E K Y F K L I A N A K T V E G V W T Y K D E I L K F T V T E
BINARY_2387 T T Y K L I L N L K Q L K E E A I K E L V D A A I A E K Y F K L I A N A K T V E G V W T Y K D E I L T A T V T E
BINARY_2388 T T Y K L I L N L K Q L K E E A I K E L V D A A I A E K Y F K L I A N A K T V E G V W T Y K D E I L T F T V T E
BINARY_2389 T T Y K L I L N L K Q L K E E A I K E L V D A A I A E K Y F K L I A N A K T V E G V W T Y K D E I K K A T V T E
BINARY_2390 T T Y K L I L N L K Q L K E E A I K E L V D A A I A E K Y F K L I A N A K T V E G V W T Y K D E I K K F T V T E
BINARY_2391 T T Y K L I L N L K Q L K E E A I K E L V D A A I A E K Y F K L I A N A K T V E G V W T Y K D E I K T A T V T E

BINARY_2392 T T Y K L I L N L K Q L K E E A I K E L V D A A I A E K Y F K L I A N A K T V E G V W T Y K D E I K T F T V T E
BINARY_2393 T T Y K L I L N L K Q L K E E A I K E L V D A A I A E K Y F K L I A N A K T V E G V W T Y K D E T L K A T V T E
BINARY_2394 T T Y K L I L N L K Q L K E E A I K E L V D A A I A E K Y F K L I A N A K T V E G V W T Y K D E T L K F T V T E
BINARY_2395 T T Y K L I L N L K Q L K E E A I K E L V D A A I A E K Y F K L I A N A K T V E G V W T Y K D E T L T A T V T E
BINARY_2396 T T Y K L I L N L K Q L K E E A I K E L V D A A I A E K Y F K L I A N A K T V E G V W T Y K D E T L T F T V T E
BINARY_2397 T T Y K L I L N L K Q L K E E A I K E L V D A A I A E K Y F K L I A N A K T V E G V W T Y K D E T K K A T V T E
BINARY_2398 T T Y K L I L N L K Q L K E E A I K E L V D A A I A E K Y F K L I A N A K T V E G V W T Y K D E T K K F T V T E
BINARY_2399 T T Y K L I L N L K Q L K E E A I K E L V D A A I A E K Y F K L I A N A K T V E G V W T Y K D E T K T A T V T E
BINARY_2400 T T Y K L I L N L K Q L K E E A I K E L V D A A I A E K Y F K L I A N A K T V E G V W T Y K D E T K T F T V T E
BINARY_2401 T T Y K L I L N L K Q L K E E A I K E L V D A A I A E K Y F K L Y A N A K T V E G V W T L K D E I L K A T V T E
BINARY_2402 T T Y K L I L N L K Q L K E E A I K E L V D A A I A E K Y F K L Y A N A K T V E G V W T L K D E I L K F T V T E
BINARY_2403 T T Y K L I L N L K Q L K E E A I K E L V D A A I A E K Y F K L Y A N A K T V E G V W T L K D E I L T A T V T E
BINARY_2404 T T Y K L I L N L K Q L K E E A I K E L V D A A I A E K Y F K L Y A N A K T V E G V W T L K D E I L T F T V T E
BINARY_2405 T T Y K L I L N L K Q L K E E A I K E L V D A A I A E K Y F K L Y A N A K T V E G V W T L K D E I K K A T V T E
BINARY_2406 T T Y K L I L N L K Q L K E E A I K E L V D A A I A E K Y F K L Y A N A K T V E G V W T L K D E I K K F T V T E
BINARY_2407 T T Y K L I L N L K Q L K E E A I K E L V D A A I A E K Y F K L Y A N A K T V E G V W T L K D E I K T A T V T E
BINARY_2408 T T Y K L I L N L K Q L K E E A I K E L V D A A I A E K Y F K L Y A N A K T V E G V W T L K D E I K T F T V T E
BINARY_2409 T T Y K L I L N L K Q L K E E A I K E L V D A A I A E K Y F K L Y A N A K T V E G V W T L K D E T L K A T V T E
BINARY_2410 T T Y K L I L N L K Q L K E E A I K E L V D A A I A E K Y F K L Y A N A K T V E G V W T L K D E T L K F T V T E
BINARY_2411 T T Y K L I L N L K Q L K E E A I K E L V D A A I A E K Y F K L Y A N A K T V E G V W T L K D E T L T A T V T E
BINARY_2412 T T Y K L I L N L K Q L K E E A I K E L V D A A I A E K Y F K L Y A N A K T V E G V W T L K D E T L T F T V T E
BINARY_2413 T T Y K L I L N L K Q L K E E A I K E L V D A A I A E K Y F K L Y A N A K T V E G V W T L K D E T K K A T V T E
BINARY_2414 T T Y K L I L N L K Q L K E E A I K E L V D A A I A E K Y F K L Y A N A K T V E G V W T L K D E T K K F T V T E
BINARY_2415 T T Y K L I L N L K Q L K E E A I K E L V D A A I A E K Y F K L Y A N A K T V E G V W T L K D E T K T A T V T E
BINARY_2416 T T Y K L I L N L K Q L K E E A I K E L V D A A I A E K Y F K L Y A N A K T V E G V W T L K D E T K T F T V T E
BINARY_2417 T T Y K L I L N L K Q L K E E A I K E L V D A A I A E K Y F K L Y A N A K T V E G V W T Y K D E I L K A T V T E
BINARY_2418 T T Y K L I L N L K Q L K E E A I K E L V D A A I A E K Y F K L Y A N A K T V E G V W T Y K D E I L K F T V T E
BINARY_2419 T T Y K L I L N L K Q L K E E A I K E L V D A A I A E K Y F K L Y A N A K T V E G V W T Y K D E I L T A T V T E
BINARY_2420 T T Y K L I L N L K Q L K E E A I K E L V D A A I A E K Y F K L Y A N A K T V E G V W T Y K D E I L T F T V T E
BINARY_2421 T T Y K L I L N L K Q L K E E A I K E L V D A A I A E K Y F K L Y A N A K T V E G V W T Y K D E I K K A T V T E
BINARY_2422 T T Y K L I L N L K Q L K E E A I K E L V D A A I A E K Y F K L Y A N A K T V E G V W T Y K D E I K K F T V T E
BINARY_2423 T T Y K L I L N L K Q L K E E A I K E L V D A A I A E K Y F K L Y A N A K T V E G V W T Y K D E I K T A T V T E
BINARY_2424 T T Y K L I L N L K Q L K E E A I K E L V D A A I A E K Y F K L Y A N A K T V E G V W T Y K D E I K T F T V T E
BINARY_2425 T T Y K L I L N L K Q L K E E A I K E L V D A A I A E K Y F K L Y A N A K T V E G V W T Y K D E T L K A T V T E
BINARY_2426 T T Y K L I L N L K Q L K E E A I K E L V D A A I A E K Y F K L Y A N A K T V E G V W T Y K D E T L K F T V T E
BINARY_2427 T T Y K L I L N L K Q L K E E A I K E L V D A A I A E K Y F K L Y A N A K T V E G V W T Y K D E T L T A T V T E
BINARY_2428 T T Y K L I L N L K Q L K E E A I K E L V D A A I A E K Y F K L Y A N A K T V E G V W T Y K D E T L T F T V T E
BINARY_2429 T T Y K L I L N L K Q L K E E A I K E L V D A A I A E K Y F K L Y A N A K T V E G V W T Y K D E T K K A T V T E
BINARY_2430 T T Y K L I L N L K Q L K E E A I K E L V D A A I A E K Y F K L Y A N A K T V E G V W T Y K D E T K K F T V T E
BINARY_2431 T T Y K L I L N L K Q L K E E A I K E L V D A A I A E K Y F K L Y A N A K T V E G V W T Y K D E T K T A T V T E
BINARY_2432 T T Y K L I L N L K Q L K E E A I K E L V D A A I A E K Y F K L Y A N A K T V E G V W T Y K D E T K T F T V T E
BINARY_2433 T T Y K L I L N L K Q L K E E A I K E L V D A A T A E K Y I K L I A N A K T V E G V W T L K D E I L K A T V T E
BINARY_2434 T T Y K L I L N L K Q L K E E A I K E L V D A A T A E K Y I K L I A N A K T V E G V W T L K D E I L K F T V T E
BINARY_2435 T T Y K L I L N L K Q L K E E A I K E L V D A A T A E K Y I K L I A N A K T V E G V W T L K D E I L T A T V T E
BINARY_2436 T T Y K L I L N L K Q L K E E A I K E L V D A A T A E K Y I K L I A N A K T V E G V W T L K D E I L T F T V T E
BINARY_2437 T T Y K L I L N L K Q L K E E A I K E L V D A A T A E K Y I K L I A N A K T V E G V W T L K D E I K K A T V T E
BINARY_2438 T T Y K L I L N L K Q L K E E A I K E L V D A A T A E K Y I K L I A N A K T V E G V W T L K D E I K K F T V T E
BINARY_2439 T T Y K L I L N L K Q L K E E A I K E L V D A A T A E K Y I K L I A N A K T V E G V W T L K D E I K T A T V T E
BINARY_2440 T T Y K L I L N L K Q L K E E A I K E L V D A A T A E K Y I K L I A N A K T V E G V W T L K D E I K T F T V T E
BINARY_2441 T T Y K L I L N L K Q L K E E A I K E L V D A A T A E K Y I K L I A N A K T V E G V W T L K D E T L K A T V T E
BINARY_2442 T T Y K L I L N L K Q L K E E A I K E L V D A A T A E K Y I K L I A N A K T V E G V W T L K D E T L K F T V T E
BINARY_2443 T T Y K L I L N L K Q L K E E A I K E L V D A A T A E K Y I K L I A N A K T V E G V W T L K D E T L T A T V T E
BINARY_2444 T T Y K L I L N L K Q L K E E A I K E L V D A A T A E K Y I K L I A N A K T V E G V W T L K D E T L T F T V T E
BINARY_2445 T T Y K L I L N L K Q L K E E A I K E L V D A A T A E K Y I K L I A N A K T V E G V W T L K D E T K K A T V T E
BINARY_2446 T T Y K L I L N L K Q L K E E A I K E L V D A A T A E K Y I K L I A N A K T V E G V W T L K D E T K K F T V T E
BINARY_2447 T T Y K L I L N L K Q L K E E A I K E L V D A A T A E K Y I K L I A N A K T V E G V W T L K D E T K T A T V T E
BINARY_2448 T T Y K L I L N L K Q L K E E A I K E L V D A A T A E K Y I K L I A N A K T V E G V W T L K D E T K T F T V T E
BINARY_2449 T T Y K L I L N L K Q L K E E A I K E L V D A A T A E K Y I K L I A N A K T V E G V W T Y K D E I L K A T V T E
BINARY_2450 T T Y K L I L N L K Q L K E E A I K E L V D A A T A E K Y I K L I A N A K T V E G V W T Y K D E I L K F T V T E
BINARY_2451 T T Y K L I L N L K Q L K E E A I K E L V D A A T A E K Y I K L I A N A K T V E G V W T Y K D E I L T A T V T E
BINARY_2452 T T Y K L I L N L K Q L K E E A I K E L V D A A T A E K Y I K L I A N A K T V E G V W T Y K D E I L T F T V T E
BINARY_2453 T T Y K L I L N L K Q L K E E A I K E L V D A A T A E K Y I K L I A N A K T V E G V W T Y K D E I K K A T V T E
BINARY_2454 T T Y K L I L N L K Q L K E E A I K E L V D A A T A E K Y I K L I A N A K T V E G V W T Y K D E I K K F T V T E
BINARY_2455 T T Y K L I L N L K Q L K E E A I K E L V D A A T A E K Y I K L I A N A K T V E G V W T Y K D E I K T A T V T E
BINARY_2456 T T Y K L I L N L K Q L K E E A I K E L V D A A T A E K Y I K L I A N A K T V E G V W T Y K D E I K T F T V T E
BINARY_2457 T T Y K L I L N L K Q L K E E A I K E L V D A A T A E K Y I K L I A N A K T V E G V W T Y K D E T L K A T V T E
BINARY_2458 T T Y K L I L N L K Q L K E E A I K E L V D A A T A E K Y I K L I A N A K T V E G V W T Y K D E T L K F T V T E
BINARY_2459 T T Y K L I L N L K Q L K E E A I K E L V D A A T A E K Y I K L I A N A K T V E G V W T Y K D E T L T A T V T E
BINARY_2460 T T Y K L I L N L K Q L K E E A I K E L V D A A T A E K Y I K L I A N A K T V E G V W T Y K D E T L T F T V T E
BINARY_2461 T T Y K L I L N L K Q L K E E A I K E L V D A A T A E K Y I K L I A N A K T V E G V W T Y K D E T K K A T V T E
BINARY_2462 T T Y K L I L N L K Q L K E E A I K E L V D A A T A E K Y I K L I A N A K T V E G V W T Y K D E T K K F T V T E
BINARY_2463 T T Y K L I L N L K Q L K E E A I K E L V D A A T A E K Y I K L I A N A K T V E G V W T Y K D E T K T A T V T E
BINARY_2464 T T Y K L I L N L K Q L K E E A I K E L V D A A T A E K Y I K L I A N A K T V E G V W T Y K D E T K T F T V T E
BINARY_2465 T T Y K L I L N L K Q L K E E A I K E L V D A A T A E K Y I K L Y A N A K T V E G V W T L K D E I L K A T V T E
BINARY_2466 T T Y K L I L N L K Q L K E E A I K E L V D A A T A E K Y I K L Y A N A K T V E G V W T L K D E I L K F T V T E
BINARY_2467 T T Y K L I L N L K Q L K E E A I K E L V D A A T A E K Y I K L Y A N A K T V E G V W T L K D E I L T A T V T E
BINARY_2468 T T Y K L I L N L K Q L K E E A I K E L V D A A T A E K Y I K L Y A N A K T V E G V W T L K D E I L T F T V T E
BINARY_2469 T T Y K L I L N L K Q L K E E A I K E L V D A A T A E K Y I K L Y A N A K T V E G V W T L K D E I K K A T V T E
BINARY_2470 T T Y K L I L N L K Q L K E E A I K E L V D A A T A E K Y I K L Y A N A K T V E G V W T L K D E I K K F T V T E
BINARY_2471 T T Y K L I L N L K Q L K E E A I K E L V D A A T A E K Y I K L Y A N A K T V E G V W T L K D E I K T A T V T E
BINARY_2472 T T Y K L I L N L K Q L K E E A I K E L V D A A T A E K Y I K L Y A N A K T V E G V W T L K D E I K T F T V T E
BINARY_2473 T T Y K L I L N L K Q L K E E A I K E L V D A A T A E K Y I K L Y A N A K T V E G V W T L K D E T L K A T V T E
BINARY_2474 T T Y K L I L N L K Q L K E E A I K E L V D A A T A E K Y I K L Y A N A K T V E G V W T L K D E T L K F T V T E
BINARY_2475 T T Y K L I L N L K Q L K E E A I K E L V D A A T A E K Y I K L Y A N A K T V E G V W T L K D E T L T A T V T E
BINARY_2476 T T Y K L I L N L K Q L K E E A I K E L V D A A T A E K Y I K L Y A N A K T V E G V W T L K D E T L T F T V T E
BINARY_2477 T T Y K L I L N L K Q L K E E A I K E L V D A A T A E K Y I K L Y A N A K T V E G V W T L K D E T K K A T V T E
BINARY_2478 T T Y K L I L N L K Q L K E E A I K E L V D A A T A E K Y I K L Y A N A K T V E G V W T L K D E T K K F T V T E
BINARY_2479 T T Y K L I L N L K Q L K E E A I K E L V D A A T A E K Y I K L Y A N A K T V E G V W T L K D E T K T A T V T E
BINARY_2480 T T Y K L I L N L K Q L K E E A I K E L V D A A T A E K Y I K L Y A N A K T V E G V W T L K D E T K T F T V T E
BINARY_2481 T T Y K L I L N L K Q L K E E A I K E L V D A A T A E K Y I K L Y A N A K T V E G V W T Y K D E I L K A T V T E
BINARY_2482 T T Y K L I L N L K Q L K E E A I K E L V D A A T A E K Y I K L Y A N A K T V E G V W T Y K D E I L K F T V T E
BINARY_2483 T T Y K L I L N L K Q L K E E A I K E L V D A A T A E K Y I K L Y A N A K T V E G V W T Y K D E I L T A T V T E
BINARY_2484 T T Y K L I L N L K Q L K E E A I K E L V D A A T A E K Y I K L Y A N A K T V E G V W T Y K D E I L T F T V T E
BINARY_2485 T T Y K L I L N L K Q L K E E A I K E L V D A A T A E K Y I K L Y A N A K T V E G V W T Y K D E I K K A T V T E
BINARY_2486 T T Y K L I L N L K Q L K E E A I K E L V D A A T A E K Y I K L Y A N A K T V E G V W T Y K D E I K K F T V T E

BINARY_2487 T T Y K L I L N L K Q L K E E A I K E L V D A A T A E K Y I K L Y A N A K T V E G V W T Y K D E I K T A T V T E
BINARY_2488 T T Y K L I L N L K Q L K E E A I K E L V D A A T A E K Y I K L Y A N A K T V E G V W T Y K D E I K T F T V T E
BINARY_2489 T T Y K L I L N L K Q L K E E A I K E L V D A A T A E K Y I K L Y A N A K T V E G V W T Y K D E T L K A T V T E
BINARY_2490 T T Y K L I L N L K Q L K E E A I K E L V D A A T A E K Y I K L Y A N A K T V E G V W T Y K D E T L K F T V T E
BINARY_2491 T T Y K L I L N L K Q L K E E A I K E L V D A A T A E K Y I K L Y A N A K T V E G V W T Y K D E T L T A T V T E
BINARY_2492 T T Y K L I L N L K Q L K E E A I K E L V D A A T A E K Y I K L Y A N A K T V E G V W T Y K D E T L T F T V T E
BINARY_2493 T T Y K L I L N L K Q L K E E A I K E L V D A A T A E K Y I K L Y A N A K T V E G V W T Y K D E T K K A T V T E
BINARY_2494 T T Y K L I L N L K Q L K E E A I K E L V D A A T A E K Y I K L Y A N A K T V E G V W T Y K D E T K K F T V T E
BINARY_2495 T T Y K L I L N L K Q L K E E A I K E L V D A A T A E K Y I K L Y A N A K T V E G V W T Y K D E T K T A T V T E
BINARY_2496 T T Y K L I L N L K Q L K E E A I K E L V D A A T A E K Y I K L Y A N A K T V E G V W T Y K D E T K T F T V T E
BINARY_2497 T T Y K L I L N L K Q L K E E A I K E L V D A A T A E K Y F K L I A N A K T V E G V W T L K D E I L K A T V T E
BINARY_2498 T T Y K L I L N L K Q L K E E A I K E L V D A A T A E K Y F K L I A N A K T V E G V W T L K D E I L K F T V T E
BINARY_2499 T T Y K L I L N L K Q L K E E A I K E L V D A A T A E K Y F K L I A N A K T V E G V W T L K D E I L T A T V T E
BINARY_2500 T T Y K L I L N L K Q L K E E A I K E L V D A A T A E K Y F K L I A N A K T V E G V W T L K D E I L T F T V T E
BINARY_2501 T T Y K L I L N L K Q L K E E A I K E L V D A A T A E K Y F K L I A N A K T V E G V W T L K D E I K K A T V T E
BINARY_2502 T T Y K L I L N L K Q L K E E A I K E L V D A A T A E K Y F K L I A N A K T V E G V W T L K D E I K K F T V T E
BINARY_2503 T T Y K L I L N L K Q L K E E A I K E L V D A A T A E K Y F K L I A N A K T V E G V W T L K D E I K T A T V T E
BINARY_2504 T T Y K L I L N L K Q L K E E A I K E L V D A A T A E K Y F K L I A N A K T V E G V W T L K D E I K T F T V T E
BINARY_2505 T T Y K L I L N L K Q L K E E A I K E L V D A A T A E K Y F K L I A N A K T V E G V W T L K D E T L K A T V T E
BINARY_2506 T T Y K L I L N L K Q L K E E A I K E L V D A A T A E K Y F K L I A N A K T V E G V W T L K D E T L K F T V T E
BINARY_2507 T T Y K L I L N L K Q L K E E A I K E L V D A A T A E K Y F K L I A N A K T V E G V W T L K D E T L T A T V T E
BINARY_2508 T T Y K L I L N L K Q L K E E A I K E L V D A A T A E K Y F K L I A N A K T V E G V W T L K D E T L T F T V T E
BINARY_2509 T T Y K L I L N L K Q L K E E A I K E L V D A A T A E K Y F K L I A N A K T V E G V W T L K D E T K K A T V T E
BINARY_2510 T T Y K L I L N L K Q L K E E A I K E L V D A A T A E K Y F K L I A N A K T V E G V W T L K D E T K K F T V T E
BINARY_2511 T T Y K L I L N L K Q L K E E A I K E L V D A A T A E K Y F K L I A N A K T V E G V W T L K D E T K T A T V T E
BINARY_2512 T T Y K L I L N L K Q L K E E A I K E L V D A A T A E K Y F K L I A N A K T V E G V W T L K D E T K T F T V T E
BINARY_2513 T T Y K L I L N L K Q L K E E A I K E L V D A A T A E K Y F K L I A N A K T V E G V W T Y K D E I L K A T V T E
BINARY_2514 T T Y K L I L N L K Q L K E E A I K E L V D A A T A E K Y F K L I A N A K T V E G V W T Y K D E I L K F T V T E
BINARY_2515 T T Y K L I L N L K Q L K E E A I K E L V D A A T A E K Y F K L I A N A K T V E G V W T Y K D E I L T A T V T E
BINARY_2516 T T Y K L I L N L K Q L K E E A I K E L V D A A T A E K Y F K L I A N A K T V E G V W T Y K D E I L T F T V T E
BINARY_2517 T T Y K L I L N L K Q L K E E A I K E L V D A A T A E K Y F K L I A N A K T V E G V W T Y K D E I K K A T V T E
BINARY_2518 T T Y K L I L N L K Q L K E E A I K E L V D A A T A E K Y F K L I A N A K T V E G V W T Y K D E I K K F T V T E
BINARY_2519 T T Y K L I L N L K Q L K E E A I K E L V D A A T A E K Y F K L I A N A K T V E G V W T Y K D E I K T A T V T E
BINARY_2520 T T Y K L I L N L K Q L K E E A I K E L V D A A T A E K Y F K L I A N A K T V E G V W T Y K D E I K T F T V T E
BINARY_2521 T T Y K L I L N L K Q L K E E A I K E L V D A A T A E K Y F K L I A N A K T V E G V W T Y K D E T L K A T V T E
BINARY_2522 T T Y K L I L N L K Q L K E E A I K E L V D A A T A E K Y F K L I A N A K T V E G V W T Y K D E T L K F T V T E
BINARY_2523 T T Y K L I L N L K Q L K E E A I K E L V D A A T A E K Y F K L I A N A K T V E G V W T Y K D E T L T A T V T E
BINARY_2524 T T Y K L I L N L K Q L K E E A I K E L V D A A T A E K Y F K L I A N A K T V E G V W T Y K D E T L T F T V T E
BINARY_2525 T T Y K L I L N L K Q L K E E A I K E L V D A A T A E K Y F K L I A N A K T V E G V W T Y K D E T K K A T V T E
BINARY_2526 T T Y K L I L N L K Q L K E E A I K E L V D A A T A E K Y F K L I A N A K T V E G V W T Y K D E T K K F T V T E
BINARY_2527 T T Y K L I L N L K Q L K E E A I K E L V D A A T A E K Y F K L I A N A K T V E G V W T Y K D E T K T A T V T E
BINARY_2528 T T Y K L I L N L K Q L K E E A I K E L V D A A T A E K Y F K L I A N A K T V E G V W T Y K D E T K T F T V T E
BINARY_2529 T T Y K L I L N L K Q L K E E A I K E L V D A A T A E K Y F K L Y A N A K T V E G V W T L K D E I L K A T V T E
BINARY_2530 T T Y K L I L N L K Q L K E E A I K E L V D A A T A E K Y F K L Y A N A K T V E G V W T L K D E I L K F T V T E
BINARY_2531 T T Y K L I L N L K Q L K E E A I K E L V D A A T A E K Y F K L Y A N A K T V E G V W T L K D E I L T A T V T E
BINARY_2532 T T Y K L I L N L K Q L K E E A I K E L V D A A T A E K Y F K L Y A N A K T V E G V W T L K D E I L T F T V T E
BINARY_2533 T T Y K L I L N L K Q L K E E A I K E L V D A A T A E K Y F K L Y A N A K T V E G V W T L K D E I K K A T V T E
BINARY_2534 T T Y K L I L N L K Q L K E E A I K E L V D A A T A E K Y F K L Y A N A K T V E G V W T L K D E I K K F T V T E
BINARY_2535 T T Y K L I L N L K Q L K E E A I K E L V D A A T A E K Y F K L Y A N A K T V E G V W T L K D E I K T A T V T E
BINARY_2536 T T Y K L I L N L K Q L K E E A I K E L V D A A T A E K Y F K L Y A N A K T V E G V W T L K D E I K T F T V T E
BINARY_2537 T T Y K L I L N L K Q L K E E A I K E L V D A A T A E K Y F K L Y A N A K T V E G V W T L K D E T L K A T V T E
BINARY_2538 T T Y K L I L N L K Q L K E E A I K E L V D A A T A E K Y F K L Y A N A K T V E G V W T L K D E T L K F T V T E
BINARY_2539 T T Y K L I L N L K Q L K E E A I K E L V D A A T A E K Y F K L Y A N A K T V E G V W T L K D E T L T A T V T E
BINARY_2540 T T Y K L I L N L K Q L K E E A I K E L V D A A T A E K Y F K L Y A N A K T V E G V W T L K D E T L T F T V T E
BINARY_2541 T T Y K L I L N L K Q L K E E A I K E L V D A A T A E K Y F K L Y A N A K T V E G V W T L K D E T K K A T V T E
BINARY_2542 T T Y K L I L N L K Q L K E E A I K E L V D A A T A E K Y F K L Y A N A K T V E G V W T L K D E T K K F T V T E
BINARY_2543 T T Y K L I L N L K Q L K E E A I K E L V D A A T A E K Y F K L Y A N A K T V E G V W T L K D E T K T A T V T E
BINARY_2544 T T Y K L I L N L K Q L K E E A I K E L V D A A T A E K Y F K L Y A N A K T V E G V W T L K D E T K T F T V T E
BINARY_2545 T T Y K L I L N L K Q L K E E A I K E L V D A A T A E K Y F K L Y A N A K T V E G V W T Y K D E I L K A T V T E
BINARY_2546 T T Y K L I L N L K Q L K E E A I K E L V D A A T A E K Y F K L Y A N A K T V E G V W T Y K D E I L K F T V T E
BINARY_2547 T T Y K L I L N L K Q L K E E A I K E L V D A A T A E K Y F K L Y A N A K T V E G V W T Y K D E I L T A T V T E
BINARY_2548 T T Y K L I L N L K Q L K E E A I K E L V D A A T A E K Y F K L Y A N A K T V E G V W T Y K D E I L T F T V T E
BINARY_2549 T T Y K L I L N L K Q L K E E A I K E L V D A A T A E K Y F K L Y A N A K T V E G V W T Y K D E I K K A T V T E
BINARY_2550 T T Y K L I L N L K Q L K E E A I K E L V D A A T A E K Y F K L Y A N A K T V E G V W T Y K D E I K K F T V T E
BINARY_2551 T T Y K L I L N L K Q L K E E A I K E L V D A A T A E K Y F K L Y A N A K T V E G V W T Y K D E I K T A T V T E
BINARY_2552 T T Y K L I L N L K Q L K E E A I K E L V D A A T A E K Y F K L Y A N A K T V E G V W T Y K D E I K T F T V T E
BINARY_2553 T T Y K L I L N L K Q L K E E A I K E L V D A A T A E K Y F K L Y A N A K T V E G V W T Y K D E T L K A T V T E
BINARY_2554 T T Y K L I L N L K Q L K E E A I K E L V D A A T A E K Y F K L Y A N A K T V E G V W T Y K D E T L K F T V T E
BINARY_2555 T T Y K L I L N L K Q L K E E A I K E L V D A A T A E K Y F K L Y A N A K T V E G V W T Y K D E T L T A T V T E
BINARY_2556 T T Y K L I L N L K Q L K E E A I K E L V D A A T A E K Y F K L Y A N A K T V E G V W T Y K D E T L T F T V T E
BINARY_2557 T T Y K L I L N L K Q L K E E A I K E L V D A A T A E K Y F K L Y A N A K T V E G V W T Y K D E T K K A T V T E
BINARY_2558 T T Y K L I L N L K Q L K E E A I K E L V D A A T A E K Y F K L Y A N A K T V E G V W T Y K D E T K K F T V T E
BINARY_2559 T T Y K L I L N L K Q L K E E A I K E L V D A A T A E K Y F K L Y A N A K T V E G V W T Y K D E T K T A T V T E
BINARY_2560 T T Y K L I L N L K Q L K E E A I K E L V D A A T A E K Y F K L Y A N A K T V E G V W T Y K D E T K T F T V T E
BINARY_2561 T T Y K L I L N L K Q L K E E A I K E A V D A G I A E K Y I K L I A N A K T V E G V W T L K D E I L K A T V T E
BINARY_2562 T T Y K L I L N L K Q L K E E A I K E A V D A G I A E K Y I K L I A N A K T V E G V W T L K D E I L K F T V T E
BINARY_2563 T T Y K L I L N L K Q L K E E A I K E A V D A G I A E K Y I K L I A N A K T V E G V W T L K D E I L T A T V T E
BINARY_2564 T T Y K L I L N L K Q L K E E A I K E A V D A G I A E K Y I K L I A N A K T V E G V W T L K D E I L T F T V T E
BINARY_2565 T T Y K L I L N L K Q L K E E A I K E A V D A G I A E K Y I K L I A N A K T V E G V W T L K D E I K K A T V T E
BINARY_2566 T T Y K L I L N L K Q L K E E A I K E A V D A G I A E K Y I K L I A N A K T V E G V W T L K D E I K K F T V T E
BINARY_2567 T T Y K L I L N L K Q L K E E A I K E A V D A G I A E K Y I K L I A N A K T V E G V W T L K D E I K T A T V T E
BINARY_2568 T T Y K L I L N L K Q L K E E A I K E A V D A G I A E K Y I K L I A N A K T V E G V W T L K D E I K T F T V T E
BINARY_2569 T T Y K L I L N L K Q L K E E A I K E A V D A G I A E K Y I K L I A N A K T V E G V W T L K D E T L K A T V T E
BINARY_2570 T T Y K L I L N L K Q L K E E A I K E A V D A G I A E K Y I K L I A N A K T V E G V W T L K D E T L K F T V T E
BINARY_2571 T T Y K L I L N L K Q L K E E A I K E A V D A G I A E K Y I K L I A N A K T V E G V W T L K D E T L T A T V T E
BINARY_2572 T T Y K L I L N L K Q L K E E A I K E A V D A G I A E K Y I K L I A N A K T V E G V W T L K D E T L T F T V T E
BINARY_2573 T T Y K L I L N L K Q L K E E A I K E A V D A G I A E K Y I K L I A N A K T V E G V W T L K D E T K K A T V T E
BINARY_2574 T T Y K L I L N L K Q L K E E A I K E A V D A G I A E K Y I K L I A N A K T V E G V W T L K D E T K K F T V T E
BINARY_2575 T T Y K L I L N L K Q L K E E A I K E A V D A G I A E K Y I K L I A N A K T V E G V W T L K D E T K T A T V T E
BINARY_2576 T T Y K L I L N L K Q L K E E A I K E A V D A G I A E K Y I K L I A N A K T V E G V W T L K D E T K T F T V T E
BINARY_2577 T T Y K L I L N L K Q L K E E A I K E A V D A G I A E K Y I K L I A N A K T V E G V W T Y K D E I L K A T V T E
BINARY_2578 T T Y K L I L N L K Q L K E E A I K E A V D A G I A E K Y I K L I A N A K T V E G V W T Y K D E I L K F T V T E
BINARY_2579 T T Y K L I L N L K Q L K E E A I K E A V D A G I A E K Y I K L I A N A K T V E G V W T Y K D E I L T A T V T E
BINARY_2580 T T Y K L I L N L K Q L K E E A I K E A V D A G I A E K Y I K L I A N A K T V E G V W T Y K D E I L T F T V T E
BINARY_2581 T T Y K L I L N L K Q L K E E A I K E A V D A G I A E K Y I K L I A N A K T V E G V W T Y K D E I K K A T V T E

BINARY_2582 T T Y K L I L N L K Q L K E E A I K E A V D A G I A E K Y I K L I A N A K T V E G V W T Y K D E I K K F T V T E
BINARY_2583 T T Y K L I L N L K Q L K E E A I K E A V D A G I A E K Y I K L I A N A K T V E G V W T Y K D E I K T A T V T E
BINARY_2584 T T Y K L I L N L K Q L K E E A I K E A V D A G I A E K Y I K L I A N A K T V E G V W T Y K D E I K T F T V T E
BINARY_2585 T T Y K L I L N L K Q L K E E A I K E A V D A G I A E K Y I K L I A N A K T V E G V W T Y K D E T L K A T V T E
BINARY_2586 T T Y K L I L N L K Q L K E E A I K E A V D A G I A E K Y I K L I A N A K T V E G V W T Y K D E T L K F T V T E
BINARY_2587 T T Y K L I L N L K Q L K E E A I K E A V D A G I A E K Y I K L I A N A K T V E G V W T Y K D E T L T A T V T E
BINARY_2588 T T Y K L I L N L K Q L K E E A I K E A V D A G I A E K Y I K L I A N A K T V E G V W T Y K D E T L T F T V T E
BINARY_2589 T T Y K L I L N L K Q L K E E A I K E A V D A G I A E K Y I K L I A N A K T V E G V W T Y K D E T K K A T V T E
BINARY_2590 T T Y K L I L N L K Q L K E E A I K E A V D A G I A E K Y I K L I A N A K T V E G V W T Y K D E T K K F T V T E
BINARY_2591 T T Y K L I L N L K Q L K E E A I K E A V D A G I A E K Y I K L I A N A K T V E G V W T Y K D E T K T A T V T E
BINARY_2592 T T Y K L I L N L K Q L K E E A I K E A V D A G I A E K Y I K L I A N A K T V E G V W T Y K D E T K T F T V T E
BINARY_2593 T T Y K L I L N L K Q L K E E A I K E A V D A G I A E K Y I K L Y A N A K T V E G V W T L K D E I L K A T V T E
BINARY_2594 T T Y K L I L N L K Q L K E E A I K E A V D A G I A E K Y I K L Y A N A K T V E G V W T L K D E I L K F T V T E
BINARY_2595 T T Y K L I L N L K Q L K E E A I K E A V D A G I A E K Y I K L Y A N A K T V E G V W T L K D E I L T A T V T E
BINARY_2596 T T Y K L I L N L K Q L K E E A I K E A V D A G I A E K Y I K L Y A N A K T V E G V W T L K D E I L T F T V T E
BINARY_2597 T T Y K L I L N L K Q L K E E A I K E A V D A G I A E K Y I K L Y A N A K T V E G V W T L K D E I K K A T V T E
BINARY_2598 T T Y K L I L N L K Q L K E E A I K E A V D A G I A E K Y I K L Y A N A K T V E G V W T L K D E I K K F T V T E
BINARY_2599 T T Y K L I L N L K Q L K E E A I K E A V D A G I A E K Y I K L Y A N A K T V E G V W T L K D E I K T A T V T E
BINARY_2600 T T Y K L I L N L K Q L K E E A I K E A V D A G I A E K Y I K L Y A N A K T V E G V W T L K D E I K T F T V T E
BINARY_2601 T T Y K L I L N L K Q L K E E A I K E A V D A G I A E K Y I K L Y A N A K T V E G V W T L K D E T L K A T V T E
BINARY_2602 T T Y K L I L N L K Q L K E E A I K E A V D A G I A E K Y I K L Y A N A K T V E G V W T L K D E T L K F T V T E
BINARY_2603 T T Y K L I L N L K Q L K E E A I K E A V D A G I A E K Y I K L Y A N A K T V E G V W T L K D E T L T A T V T E
BINARY_2604 T T Y K L I L N L K Q L K E E A I K E A V D A G I A E K Y I K L Y A N A K T V E G V W T L K D E T L T F T V T E
BINARY_2605 T T Y K L I L N L K Q L K E E A I K E A V D A G I A E K Y I K L Y A N A K T V E G V W T L K D E T K K A T V T E
BINARY_2606 T T Y K L I L N L K Q L K E E A I K E A V D A G I A E K Y I K L Y A N A K T V E G V W T L K D E T K K F T V T E
BINARY_2607 T T Y K L I L N L K Q L K E E A I K E A V D A G I A E K Y I K L Y A N A K T V E G V W T L K D E T K T A T V T E
BINARY_2608 T T Y K L I L N L K Q L K E E A I K E A V D A G I A E K Y I K L Y A N A K T V E G V W T L K D E T K T F T V T E
BINARY_2609 T T Y K L I L N L K Q L K E E A I K E A V D A G I A E K Y I K L Y A N A K T V E G V W T Y K D E I L K A T V T E
BINARY_2610 T T Y K L I L N L K Q L K E E A I K E A V D A G I A E K Y I K L Y A N A K T V E G V W T Y K D E I L K F T V T E
BINARY_2611 T T Y K L I L N L K Q L K E E A I K E A V D A G I A E K Y I K L Y A N A K T V E G V W T Y K D E I L T A T V T E
BINARY_2612 T T Y K L I L N L K Q L K E E A I K E A V D A G I A E K Y I K L Y A N A K T V E G V W T Y K D E I L T F T V T E
BINARY_2613 T T Y K L I L N L K Q L K E E A I K E A V D A G I A E K Y I K L Y A N A K T V E G V W T Y K D E I K K A T V T E
BINARY_2614 T T Y K L I L N L K Q L K E E A I K E A V D A G I A E K Y I K L Y A N A K T V E G V W T Y K D E I K K F T V T E
BINARY_2615 T T Y K L I L N L K Q L K E E A I K E A V D A G I A E K Y I K L Y A N A K T V E G V W T Y K D E I K T A T V T E
BINARY_2616 T T Y K L I L N L K Q L K E E A I K E A V D A G I A E K Y I K L Y A N A K T V E G V W T Y K D E I K T F T V T E
BINARY_2617 T T Y K L I L N L K Q L K E E A I K E A V D A G I A E K Y I K L Y A N A K T V E G V W T Y K D E T L K A T V T E
BINARY_2618 T T Y K L I L N L K Q L K E E A I K E A V D A G I A E K Y I K L Y A N A K T V E G V W T Y K D E T L K F T V T E
BINARY_2619 T T Y K L I L N L K Q L K E E A I K E A V D A G I A E K Y I K L Y A N A K T V E G V W T Y K D E T L T A T V T E
BINARY_2620 T T Y K L I L N L K Q L K E E A I K E A V D A G I A E K Y I K L Y A N A K T V E G V W T Y K D E T L T F T V T E
BINARY_2621 T T Y K L I L N L K Q L K E E A I K E A V D A G I A E K Y I K L Y A N A K T V E G V W T Y K D E T K K A T V T E
BINARY_2622 T T Y K L I L N L K Q L K E E A I K E A V D A G I A E K Y I K L Y A N A K T V E G V W T Y K D E T K K F T V T E
BINARY_2623 T T Y K L I L N L K Q L K E E A I K E A V D A G I A E K Y I K L Y A N A K T V E G V W T Y K D E T K T A T V T E
BINARY_2624 T T Y K L I L N L K Q L K E E A I K E A V D A G I A E K Y I K L Y A N A K T V E G V W T Y K D E T K T F T V T E
BINARY_2625 T T Y K L I L N L K Q L K E E A I K E A V D A G I A E K Y F K L I A N A K T V E G V W T L K D E I L K A T V T E
BINARY_2626 T T Y K L I L N L K Q L K E E A I K E A V D A G I A E K Y F K L I A N A K T V E G V W T L K D E I L K F T V T E
BINARY_2627 T T Y K L I L N L K Q L K E E A I K E A V D A G I A E K Y F K L I A N A K T V E G V W T L K D E I L T A T V T E
BINARY_2628 T T Y K L I L N L K Q L K E E A I K E A V D A G I A E K Y F K L I A N A K T V E G V W T L K D E I L T F T V T E
BINARY_2629 T T Y K L I L N L K Q L K E E A I K E A V D A G I A E K Y F K L I A N A K T V E G V W T L K D E I K K A T V T E
BINARY_2630 T T Y K L I L N L K Q L K E E A I K E A V D A G I A E K Y F K L I A N A K T V E G V W T L K D E I K K F T V T E
BINARY_2631 T T Y K L I L N L K Q L K E E A I K E A V D A G I A E K Y F K L I A N A K T V E G V W T L K D E I K T A T V T E
BINARY_2632 T T Y K L I L N L K Q L K E E A I K E A V D A G I A E K Y F K L I A N A K T V E G V W T L K D E I K T F T V T E
BINARY_2633 T T Y K L I L N L K Q L K E E A I K E A V D A G I A E K Y F K L I A N A K T V E G V W T L K D E T L K A T V T E
BINARY_2634 T T Y K L I L N L K Q L K E E A I K E A V D A G I A E K Y F K L I A N A K T V E G V W T L K D E T L K F T V T E
BINARY_2635 T T Y K L I L N L K Q L K E E A I K E A V D A G I A E K Y F K L I A N A K T V E G V W T L K D E T L T A T V T E
BINARY_2636 T T Y K L I L N L K Q L K E E A I K E A V D A G I A E K Y F K L I A N A K T V E G V W T L K D E T L T F T V T E
BINARY_2637 T T Y K L I L N L K Q L K E E A I K E A V D A G I A E K Y F K L I A N A K T V E G V W T L K D E T K K A T V T E
BINARY_2638 T T Y K L I L N L K Q L K E E A I K E A V D A G I A E K Y F K L I A N A K T V E G V W T L K D E T K K F T V T E
BINARY_2639 T T Y K L I L N L K Q L K E E A I K E A V D A G I A E K Y F K L I A N A K T V E G V W T L K D E T K T A T V T E
BINARY_2640 T T Y K L I L N L K Q L K E E A I K E A V D A G I A E K Y F K L I A N A K T V E G V W T L K D E T K T F T V T E
BINARY_2641 T T Y K L I L N L K Q L K E E A I K E A V D A G I A E K Y F K L I A N A K T V E G V W T Y K D E I L K A T V T E
BINARY_2642 T T Y K L I L N L K Q L K E E A I K E A V D A G I A E K Y F K L I A N A K T V E G V W T Y K D E I L K F T V T E
BINARY_2643 T T Y K L I L N L K Q L K E E A I K E A V D A G I A E K Y F K L I A N A K T V E G V W T Y K D E I L T A T V T E
BINARY_2644 T T Y K L I L N L K Q L K E E A I K E A V D A G I A E K Y F K L I A N A K T V E G V W T Y K D E I L T F T V T E
BINARY_2645 T T Y K L I L N L K Q L K E E A I K E A V D A G I A E K Y F K L I A N A K T V E G V W T Y K D E I K K A T V T E
BINARY_2646 T T Y K L I L N L K Q L K E E A I K E A V D A G I A E K Y F K L I A N A K T V E G V W T Y K D E I K K F T V T E
BINARY_2647 T T Y K L I L N L K Q L K E E A I K E A V D A G I A E K Y F K L I A N A K T V E G V W T Y K D E I K T A T V T E
BINARY_2648 T T Y K L I L N L K Q L K E E A I K E A V D A G I A E K Y F K L I A N A K T V E G V W T Y K D E I K T F T V T E
BINARY_2649 T T Y K L I L N L K Q L K E E A I K E A V D A G I A E K Y F K L I A N A K T V E G V W T Y K D E T L K A T V T E
BINARY_2650 T T Y K L I L N L K Q L K E E A I K E A V D A G I A E K Y F K L I A N A K T V E G V W T Y K D E T L K F T V T E
BINARY_2651 T T Y K L I L N L K Q L K E E A I K E A V D A G I A E K Y F K L I A N A K T V E G V W T Y K D E T L T A T V T E
BINARY_2652 T T Y K L I L N L K Q L K E E A I K E A V D A G I A E K Y F K L I A N A K T V E G V W T Y K D E T L T F T V T E
BINARY_2653 T T Y K L I L N L K Q L K E E A I K E A V D A G I A E K Y F K L I A N A K T V E G V W T Y K D E T K K A T V T E
BINARY_2654 T T Y K L I L N L K Q L K E E A I K E A V D A G I A E K Y F K L I A N A K T V E G V W T Y K D E T K K F T V T E
BINARY_2655 T T Y K L I L N L K Q L K E E A I K E A V D A G I A E K Y F K L I A N A K T V E G V W T Y K D E T K T A T V T E
BINARY_2656 T T Y K L I L N L K Q L K E E A I K E A V D A G I A E K Y F K L I A N A K T V E G V W T Y K D E T K T F T V T E
BINARY_2657 T T Y K L I L N L K Q L K E E A I K E A V D A G I A E K Y F K L Y A N A K T V E G V W T L K D E I L K A T V T E
BINARY_2658 T T Y K L I L N L K Q L K E E A I K E A V D A G I A E K Y F K L Y A N A K T V E G V W T L K D E I L K F T V T E
BINARY_2659 T T Y K L I L N L K Q L K E E A I K E A V D A G I A E K Y F K L Y A N A K T V E G V W T L K D E I L T A T V T E
BINARY_2660 T T Y K L I L N L K Q L K E E A I K E A V D A G I A E K Y F K L Y A N A K T V E G V W T L K D E I L T F T V T E
BINARY_2661 T T Y K L I L N L K Q L K E E A I K E A V D A G I A E K Y F K L Y A N A K T V E G V W T L K D E I K K A T V T E
BINARY_2662 T T Y K L I L N L K Q L K E E A I K E A V D A G I A E K Y F K L Y A N A K T V E G V W T L K D E I K K F T V T E
BINARY_2663 T T Y K L I L N L K Q L K E E A I K E A V D A G I A E K Y F K L Y A N A K T V E G V W T L K D E I K T A T V T E
BINARY_2664 T T Y K L I L N L K Q L K E E A I K E A V D A G I A E K Y F K L Y A N A K T V E G V W T L K D E I K T F T V T E
BINARY_2665 T T Y K L I L N L K Q L K E E A I K E A V D A G I A E K Y F K L Y A N A K T V E G V W T L K D E T L K A T V T E
BINARY_2666 T T Y K L I L N L K Q L K E E A I K E A V D A G I A E K Y F K L Y A N A K T V E G V W T L K D E T L K F T V T E
BINARY_2667 T T Y K L I L N L K Q L K E E A I K E A V D A G I A E K Y F K L Y A N A K T V E G V W T L K D E T L T A T V T E
BINARY_2668 T T Y K L I L N L K Q L K E E A I K E A V D A G I A E K Y F K L Y A N A K T V E G V W T L K D E T L T F T V T E
BINARY_2669 T T Y K L I L N L K Q L K E E A I K E A V D A G I A E K Y F K L Y A N A K T V E G V W T L K D E T K K A T V T E
BINARY_2670 T T Y K L I L N L K Q L K E E A I K E A V D A G I A E K Y F K L Y A N A K T V E G V W T L K D E T K K F T V T E
BINARY_2671 T T Y K L I L N L K Q L K E E A I K E A V D A G I A E K Y F K L Y A N A K T V E G V W T L K D E T K T A T V T E
BINARY_2672 T T Y K L I L N L K Q L K E E A I K E A V D A G I A E K Y F K L Y A N A K T V E G V W T L K D E T K T F T V T E
BINARY_2673 T T Y K L I L N L K Q L K E E A I K E A V D A G I A E K Y F K L Y A N A K T V E G V W T Y K D E I L K A T V T E
BINARY_2674 T T Y K L I L N L K Q L K E E A I K E A V D A G I A E K Y F K L Y A N A K T V E G V W T Y K D E I L K F T V T E
BINARY_2675 T T Y K L I L N L K Q L K E E A I K E A V D A G I A E K Y F K L Y A N A K T V E G V W T Y K D E I L T A T V T E
BINARY_2676 T T Y K L I L N L K Q L K E E A I K E A V D A G I A E K Y F K L Y A N A K T V E G V W T Y K D E I L T F T V T E

BINARY_2677 T T Y K L I L N L K Q L K E E A I K E A V D A G I A E K Y F K L Y A N A K T V E G V W T Y K D E I K K A T V T E
BINARY_2678 T T Y K L I L N L K Q L K E E A I K E A V D A G I A E K Y F K L Y A N A K T V E G V W T Y K D E I K K F T V T E
BINARY_2679 T T Y K L I L N L K Q L K E E A I K E A V D A G I A E K Y F K L Y A N A K T V E G V W T Y K D E I K T A T V T E
BINARY_2680 T T Y K L I L N L K Q L K E E A I K E A V D A G I A E K Y F K L Y A N A K T V E G V W T Y K D E I K T F T V T E
BINARY_2681 T T Y K L I L N L K Q L K E E A I K E A V D A G I A E K Y F K L Y A N A K T V E G V W T Y K D E T L K A T V T E
BINARY_2682 T T Y K L I L N L K Q L K E E A I K E A V D A G I A E K Y F K L Y A N A K T V E G V W T Y K D E T L K F T V T E
BINARY_2683 T T Y K L I L N L K Q L K E E A I K E A V D A G I A E K Y F K L Y A N A K T V E G V W T Y K D E T L T A T V T E
BINARY_2684 T T Y K L I L N L K Q L K E E A I K E A V D A G I A E K Y F K L Y A N A K T V E G V W T Y K D E T L T F T V T E
BINARY_2685 T T Y K L I L N L K Q L K E E A I K E A V D A G I A E K Y F K L Y A N A K T V E G V W T Y K D E T K K A T V T E
BINARY_2686 T T Y K L I L N L K Q L K E E A I K E A V D A G I A E K Y F K L Y A N A K T V E G V W T Y K D E T K K F T V T E
BINARY_2687 T T Y K L I L N L K Q L K E E A I K E A V D A G I A E K Y F K L Y A N A K T V E G V W T Y K D E T K T A T V T E
BINARY_2688 T T Y K L I L N L K Q L K E E A I K E A V D A G I A E K Y F K L Y A N A K T V E G V W T Y K D E T K T F T V T E
BINARY_2689 T T Y K L I L N L K Q L K E E A I K E A V D A G T A E K Y I K L I A N A K T V E G V W T L K D E I L K A T V T E
BINARY_2690 T T Y K L I L N L K Q L K E E A I K E A V D A G T A E K Y I K L I A N A K T V E G V W T L K D E I L K F T V T E
BINARY_2691 T T Y K L I L N L K Q L K E E A I K E A V D A G T A E K Y I K L I A N A K T V E G V W T L K D E I L T A T V T E
BINARY_2692 T T Y K L I L N L K Q L K E E A I K E A V D A G T A E K Y I K L I A N A K T V E G V W T L K D E I L T F T V T E
BINARY_2693 T T Y K L I L N L K Q L K E E A I K E A V D A G T A E K Y I K L I A N A K T V E G V W T L K D E I K K A T V T E
BINARY_2694 T T Y K L I L N L K Q L K E E A I K E A V D A G T A E K Y I K L I A N A K T V E G V W T L K D E I K K F T V T E
BINARY_2695 T T Y K L I L N L K Q L K E E A I K E A V D A G T A E K Y I K L I A N A K T V E G V W T L K D E I K T A T V T E
BINARY_2696 T T Y K L I L N L K Q L K E E A I K E A V D A G T A E K Y I K L I A N A K T V E G V W T L K D E I K T F T V T E
BINARY_2697 T T Y K L I L N L K Q L K E E A I K E A V D A G T A E K Y I K L I A N A K T V E G V W T L K D E T L K A T V T E
BINARY_2698 T T Y K L I L N L K Q L K E E A I K E A V D A G T A E K Y I K L I A N A K T V E G V W T L K D E T L K F T V T E
BINARY_2699 T T Y K L I L N L K Q L K E E A I K E A V D A G T A E K Y I K L I A N A K T V E G V W T L K D E T L T A T V T E
BINARY_2700 T T Y K L I L N L K Q L K E E A I K E A V D A G T A E K Y I K L I A N A K T V E G V W T L K D E T L T F T V T E
BINARY_2701 T T Y K L I L N L K Q L K E E A I K E A V D A G T A E K Y I K L I A N A K T V E G V W T L K D E T K K A T V T E
BINARY_2702 T T Y K L I L N L K Q L K E E A I K E A V D A G T A E K Y I K L I A N A K T V E G V W T L K D E T K K F T V T E
BINARY_2703 T T Y K L I L N L K Q L K E E A I K E A V D A G T A E K Y I K L I A N A K T V E G V W T L K D E T K T A T V T E
BINARY_2704 T T Y K L I L N L K Q L K E E A I K E A V D A G T A E K Y I K L I A N A K T V E G V W T L K D E T K T F T V T E
BINARY_2705 T T Y K L I L N L K Q L K E E A I K E A V D A G T A E K Y I K L I A N A K T V E G V W T Y K D E I L K A T V T E
BINARY_2706 T T Y K L I L N L K Q L K E E A I K E A V D A G T A E K Y I K L I A N A K T V E G V W T Y K D E I L K F T V T E
BINARY_2707 T T Y K L I L N L K Q L K E E A I K E A V D A G T A E K Y I K L I A N A K T V E G V W T Y K D E I L T A T V T E
BINARY_2708 T T Y K L I L N L K Q L K E E A I K E A V D A G T A E K Y I K L I A N A K T V E G V W T Y K D E I L T F T V T E
BINARY_2709 T T Y K L I L N L K Q L K E E A I K E A V D A G T A E K Y I K L I A N A K T V E G V W T Y K D E I K K A T V T E
BINARY_2710 T T Y K L I L N L K Q L K E E A I K E A V D A G T A E K Y I K L I A N A K T V E G V W T Y K D E I K K F T V T E
BINARY_2711 T T Y K L I L N L K Q L K E E A I K E A V D A G T A E K Y I K L I A N A K T V E G V W T Y K D E I K T A T V T E
BINARY_2712 T T Y K L I L N L K Q L K E E A I K E A V D A G T A E K Y I K L I A N A K T V E G V W T Y K D E I K T F T V T E
BINARY_2713 T T Y K L I L N L K Q L K E E A I K E A V D A G T A E K Y I K L I A N A K T V E G V W T Y K D E T L K A T V T E
BINARY_2714 T T Y K L I L N L K Q L K E E A I K E A V D A G T A E K Y I K L I A N A K T V E G V W T Y K D E T L K F T V T E
BINARY_2715 T T Y K L I L N L K Q L K E E A I K E A V D A G T A E K Y I K L I A N A K T V E G V W T Y K D E T L T A T V T E
BINARY_2716 T T Y K L I L N L K Q L K E E A I K E A V D A G T A E K Y I K L I A N A K T V E G V W T Y K D E T L T F T V T E
BINARY_2717 T T Y K L I L N L K Q L K E E A I K E A V D A G T A E K Y I K L I A N A K T V E G V W T Y K D E T K K A T V T E
BINARY_2718 T T Y K L I L N L K Q L K E E A I K E A V D A G T A E K Y I K L I A N A K T V E G V W T Y K D E T K K F T V T E
BINARY_2719 T T Y K L I L N L K Q L K E E A I K E A V D A G T A E K Y I K L I A N A K T V E G V W T Y K D E T K T A T V T E
BINARY_2720 T T Y K L I L N L K Q L K E E A I K E A V D A G T A E K Y I K L I A N A K T V E G V W T Y K D E T K T F T V T E
BINARY_2721 T T Y K L I L N L K Q L K E E A I K E A V D A G T A E K Y I K L Y A N A K T V E G V W T L K D E I L K A T V T E
BINARY_2722 T T Y K L I L N L K Q L K E E A I K E A V D A G T A E K Y I K L Y A N A K T V E G V W T L K D E I L K F T V T E
BINARY_2723 T T Y K L I L N L K Q L K E E A I K E A V D A G T A E K Y I K L Y A N A K T V E G V W T L K D E I L T A T V T E
BINARY_2724 T T Y K L I L N L K Q L K E E A I K E A V D A G T A E K Y I K L Y A N A K T V E G V W T L K D E I L T F T V T E
BINARY_2725 T T Y K L I L N L K Q L K E E A I K E A V D A G T A E K Y I K L Y A N A K T V E G V W T L K D E I K K A T V T E
BINARY_2726 T T Y K L I L N L K Q L K E E A I K E A V D A G T A E K Y I K L Y A N A K T V E G V W T L K D E I K K F T V T E
BINARY_2727 T T Y K L I L N L K Q L K E E A I K E A V D A G T A E K Y I K L Y A N A K T V E G V W T L K D E I K T A T V T E
BINARY_2728 T T Y K L I L N L K Q L K E E A I K E A V D A G T A E K Y I K L Y A N A K T V E G V W T L K D E I K T F T V T E
BINARY_2729 T T Y K L I L N L K Q L K E E A I K E A V D A G T A E K Y I K L Y A N A K T V E G V W T L K D E T L K A T V T E
BINARY_2730 T T Y K L I L N L K Q L K E E A I K E A V D A G T A E K Y I K L Y A N A K T V E G V W T L K D E T L K F T V T E
BINARY_2731 T T Y K L I L N L K Q L K E E A I K E A V D A G T A E K Y I K L Y A N A K T V E G V W T L K D E T L T A T V T E
BINARY_2732 T T Y K L I L N L K Q L K E E A I K E A V D A G T A E K Y I K L Y A N A K T V E G V W T L K D E T L T F T V T E
BINARY_2733 T T Y K L I L N L K Q L K E E A I K E A V D A G T A E K Y I K L Y A N A K T V E G V W T L K D E T K K A T V T E
BINARY_2734 T T Y K L I L N L K Q L K E E A I K E A V D A G T A E K Y I K L Y A N A K T V E G V W T L K D E T K K F T V T E
BINARY_2735 T T Y K L I L N L K Q L K E E A I K E A V D A G T A E K Y I K L Y A N A K T V E G V W T L K D E T K T A T V T E
BINARY_2736 T T Y K L I L N L K Q L K E E A I K E A V D A G T A E K Y I K L Y A N A K T V E G V W T L K D E T K T F T V T E
BINARY_2737 T T Y K L I L N L K Q L K E E A I K E A V D A G T A E K Y I K L Y A N A K T V E G V W T Y K D E I L K A T V T E
BINARY_2738 T T Y K L I L N L K Q L K E E A I K E A V D A G T A E K Y I K L Y A N A K T V E G V W T Y K D E I L K F T V T E
BINARY_2739 T T Y K L I L N L K Q L K E E A I K E A V D A G T A E K Y I K L Y A N A K T V E G V W T Y K D E I L T A T V T E
BINARY_2740 T T Y K L I L N L K Q L K E E A I K E A V D A G T A E K Y I K L Y A N A K T V E G V W T Y K D E I L T F T V T E
BINARY_2741 T T Y K L I L N L K Q L K E E A I K E A V D A G T A E K Y I K L Y A N A K T V E G V W T Y K D E I K K A T V T E
BINARY_2742 T T Y K L I L N L K Q L K E E A I K E A V D A G T A E K Y I K L Y A N A K T V E G V W T Y K D E I K K F T V T E
BINARY_2743 T T Y K L I L N L K Q L K E E A I K E A V D A G T A E K Y I K L Y A N A K T V E G V W T Y K D E I K T A T V T E
BINARY_2744 T T Y K L I L N L K Q L K E E A I K E A V D A G T A E K Y I K L Y A N A K T V E G V W T Y K D E I K T F T V T E
BINARY_2745 T T Y K L I L N L K Q L K E E A I K E A V D A G T A E K Y I K L Y A N A K T V E G V W T Y K D E T L K A T V T E
BINARY_2746 T T Y K L I L N L K Q L K E E A I K E A V D A G T A E K Y I K L Y A N A K T V E G V W T Y K D E T L K F T V T E
BINARY_2747 T T Y K L I L N L K Q L K E E A I K E A V D A G T A E K Y I K L Y A N A K T V E G V W T Y K D E T L T A T V T E
BINARY_2748 T T Y K L I L N L K Q L K E E A I K E A V D A G T A E K Y I K L Y A N A K T V E G V W T Y K D E T L T F T V T E
BINARY_2749 T T Y K L I L N L K Q L K E E A I K E A V D A G T A E K Y I K L Y A N A K T V E G V W T Y K D E T K K A T V T E
BINARY_2750 T T Y K L I L N L K Q L K E E A I K E A V D A G T A E K Y I K L Y A N A K T V E G V W T Y K D E T K K F T V T E
BINARY_2751 T T Y K L I L N L K Q L K E E A I K E A V D A G T A E K Y I K L Y A N A K T V E G V W T Y K D E T K T A T V T E
BINARY_2752 T T Y K L I L N L K Q L K E E A I K E A V D A G T A E K Y I K L Y A N A K T V E G V W T Y K D E T K T F T V T E
BINARY_2753 T T Y K L I L N L K Q L K E E A I K E A V D A G T A E K Y F K L I A N A K T V E G V W T L K D E I L K A T V T E
BINARY_2754 T T Y K L I L N L K Q L K E E A I K E A V D A G T A E K Y F K L I A N A K T V E G V W T L K D E I L K F T V T E
BINARY_2755 T T Y K L I L N L K Q L K E E A I K E A V D A G T A E K Y F K L I A N A K T V E G V W T L K D E I L T A T V T E
BINARY_2756 T T Y K L I L N L K Q L K E E A I K E A V D A G T A E K Y F K L I A N A K T V E G V W T L K D E I L T F T V T E
BINARY_2757 T T Y K L I L N L K Q L K E E A I K E A V D A G T A E K Y F K L I A N A K T V E G V W T L K D E I K K A T V T E
BINARY_2758 T T Y K L I L N L K Q L K E E A I K E A V D A G T A E K Y F K L I A N A K T V E G V W T L K D E I K K F T V T E
BINARY_2759 T T Y K L I L N L K Q L K E E A I K E A V D A G T A E K Y F K L I A N A K T V E G V W T L K D E I K T A T V T E
BINARY_2760 T T Y K L I L N L K Q L K E E A I K E A V D A G T A E K Y F K L I A N A K T V E G V W T L K D E I K T F T V T E
BINARY_2761 T T Y K L I L N L K Q L K E E A I K E A V D A G T A E K Y F K L I A N A K T V E G V W T L K D E T L K A T V T E
BINARY_2762 T T Y K L I L N L K Q L K E E A I K E A V D A G T A E K Y F K L I A N A K T V E G V W T L K D E T L K F T V T E
BINARY_2763 T T Y K L I L N L K Q L K E E A I K E A V D A G T A E K Y F K L I A N A K T V E G V W T L K D E T L T A T V T E
BINARY_2764 T T Y K L I L N L K Q L K E E A I K E A V D A G T A E K Y F K L I A N A K T V E G V W T L K D E T L T F T V T E
BINARY_2765 T T Y K L I L N L K Q L K E E A I K E A V D A G T A E K Y F K L I A N A K T V E G V W T L K D E T K K A T V T E
BINARY_2766 T T Y K L I L N L K Q L K E E A I K E A V D A G T A E K Y F K L I A N A K T V E G V W T L K D E T K K F T V T E
BINARY_2767 T T Y K L I L N L K Q L K E E A I K E A V D A G T A E K Y F K L I A N A K T V E G V W T L K D E T K T A T V T E
BINARY_2768 T T Y K L I L N L K Q L K E E A I K E A V D A G T A E K Y F K L I A N A K T V E G V W T L K D E T K T F T V T E
BINARY_2769 T T Y K L I L N L K Q L K E E A I K E A V D A G T A E K Y F K L I A N A K T V E G V W T Y K D E I L K A T V T E
BINARY_2770 T T Y K L I L N L K Q L K E E A I K E A V D A G T A E K Y F K L I A N A K T V E G V W T Y K D E I L K F T V T E
BINARY_2771 T T Y K L I L N L K Q L K E E A I K E A V D A G T A E K Y F K L I A N A K T V E G V W T Y K D E I L T A T V T E

BINARY_2772 T T Y K L I L N L K Q L K E E A I K E A V D A G T A E K Y F K L I A N A K T V E G V W T Y K D E I L T F T V T E
BINARY_2773 T T Y K L I L N L K Q L K E E A I K E A V D A G T A E K Y F K L I A N A K T V E G V W T Y K D E I K K A T V T E
BINARY_2774 T T Y K L I L N L K Q L K E E A I K E A V D A G T A E K Y F K L I A N A K T V E G V W T Y K D E I K K F T V T E
BINARY_2775 T T Y K L I L N L K Q L K E E A I K E A V D A G T A E K Y F K L I A N A K T V E G V W T Y K D E I K T A T V T E
BINARY_2776 T T Y K L I L N L K Q L K E E A I K E A V D A G T A E K Y F K L I A N A K T V E G V W T Y K D E I K T F T V T E
BINARY_2777 T T Y K L I L N L K Q L K E E A I K E A V D A G T A E K Y F K L I A N A K T V E G V W T Y K D E T L K A T V T E
BINARY_2778 T T Y K L I L N L K Q L K E E A I K E A V D A G T A E K Y F K L I A N A K T V E G V W T Y K D E T L K F T V T E
BINARY_2779 T T Y K L I L N L K Q L K E E A I K E A V D A G T A E K Y F K L I A N A K T V E G V W T Y K D E T L T A T V T E
BINARY_2780 T T Y K L I L N L K Q L K E E A I K E A V D A G T A E K Y F K L I A N A K T V E G V W T Y K D E T L T F T V T E
BINARY_2781 T T Y K L I L N L K Q L K E E A I K E A V D A G T A E K Y F K L I A N A K T V E G V W T Y K D E T K K A T V T E
BINARY_2782 T T Y K L I L N L K Q L K E E A I K E A V D A G T A E K Y F K L I A N A K T V E G V W T Y K D E T K K F T V T E
BINARY_2783 T T Y K L I L N L K Q L K E E A I K E A V D A G T A E K Y F K L I A N A K T V E G V W T Y K D E T K T A T V T E
BINARY_2784 T T Y K L I L N L K Q L K E E A I K E A V D A G T A E K Y F K L I A N A K T V E G V W T Y K D E T K T F T V T E
BINARY_2785 T T Y K L I L N L K Q L K E E A I K E A V D A G T A E K Y F K L Y A N A K T V E G V W T L K D E I L K A T V T E
BINARY_2786 T T Y K L I L N L K Q L K E E A I K E A V D A G T A E K Y F K L Y A N A K T V E G V W T L K D E I L K F T V T E
BINARY_2787 T T Y K L I L N L K Q L K E E A I K E A V D A G T A E K Y F K L Y A N A K T V E G V W T L K D E I L T A T V T E
BINARY_2788 T T Y K L I L N L K Q L K E E A I K E A V D A G T A E K Y F K L Y A N A K T V E G V W T L K D E I L T F T V T E
BINARY_2789 T T Y K L I L N L K Q L K E E A I K E A V D A G T A E K Y F K L Y A N A K T V E G V W T L K D E I K K A T V T E
BINARY_2790 T T Y K L I L N L K Q L K E E A I K E A V D A G T A E K Y F K L Y A N A K T V E G V W T L K D E I K K F T V T E
BINARY_2791 T T Y K L I L N L K Q L K E E A I K E A V D A G T A E K Y F K L Y A N A K T V E G V W T L K D E I K T A T V T E
BINARY_2792 T T Y K L I L N L K Q L K E E A I K E A V D A G T A E K Y F K L Y A N A K T V E G V W T L K D E I K T F T V T E
BINARY_2793 T T Y K L I L N L K Q L K E E A I K E A V D A G T A E K Y F K L Y A N A K T V E G V W T L K D E T L K A T V T E
BINARY_2794 T T Y K L I L N L K Q L K E E A I K E A V D A G T A E K Y F K L Y A N A K T V E G V W T L K D E T L K F T V T E
BINARY_2795 T T Y K L I L N L K Q L K E E A I K E A V D A G T A E K Y F K L Y A N A K T V E G V W T L K D E T L T A T V T E
BINARY_2796 T T Y K L I L N L K Q L K E E A I K E A V D A G T A E K Y F K L Y A N A K T V E G V W T L K D E T L T F T V T E
BINARY_2797 T T Y K L I L N L K Q L K E E A I K E A V D A G T A E K Y F K L Y A N A K T V E G V W T L K D E T K K A T V T E
BINARY_2798 T T Y K L I L N L K Q L K E E A I K E A V D A G T A E K Y F K L Y A N A K T V E G V W T L K D E T K K F T V T E
BINARY_2799 T T Y K L I L N L K Q L K E E A I K E A V D A G T A E K Y F K L Y A N A K T V E G V W T L K D E T K T A T V T E
BINARY_2800 T T Y K L I L N L K Q L K E E A I K E A V D A G T A E K Y F K L Y A N A K T V E G V W T L K D E T K T F T V T E
BINARY_2801 T T Y K L I L N L K Q L K E E A I K E A V D A G T A E K Y F K L Y A N A K T V E G V W T Y K D E I L K A T V T E
BINARY_2802 T T Y K L I L N L K Q L K E E A I K E A V D A G T A E K Y F K L Y A N A K T V E G V W T Y K D E I L K F T V T E
BINARY_2803 T T Y K L I L N L K Q L K E E A I K E A V D A G T A E K Y F K L Y A N A K T V E G V W T Y K D E I L T A T V T E
BINARY_2804 T T Y K L I L N L K Q L K E E A I K E A V D A G T A E K Y F K L Y A N A K T V E G V W T Y K D E I L T F T V T E
BINARY_2805 T T Y K L I L N L K Q L K E E A I K E A V D A G T A E K Y F K L Y A N A K T V E G V W T Y K D E I K K A T V T E
BINARY_2806 T T Y K L I L N L K Q L K E E A I K E A V D A G T A E K Y F K L Y A N A K T V E G V W T Y K D E I K K F T V T E
BINARY_2807 T T Y K L I L N L K Q L K E E A I K E A V D A G T A E K Y F K L Y A N A K T V E G V W T Y K D E I K T A T V T E
BINARY_2808 T T Y K L I L N L K Q L K E E A I K E A V D A G T A E K Y F K L Y A N A K T V E G V W T Y K D E I K T F T V T E
BINARY_2809 T T Y K L I L N L K Q L K E E A I K E A V D A G T A E K Y F K L Y A N A K T V E G V W T Y K D E T L K A T V T E
BINARY_2810 T T Y K L I L N L K Q L K E E A I K E A V D A G T A E K Y F K L Y A N A K T V E G V W T Y K D E T L K F T V T E
BINARY_2811 T T Y K L I L N L K Q L K E E A I K E A V D A G T A E K Y F K L Y A N A K T V E G V W T Y K D E T L T A T V T E
BINARY_2812 T T Y K L I L N L K Q L K E E A I K E A V D A G T A E K Y F K L Y A N A K T V E G V W T Y K D E T L T F T V T E
BINARY_2813 T T Y K L I L N L K Q L K E E A I K E A V D A G T A E K Y F K L Y A N A K T V E G V W T Y K D E T K K A T V T E
BINARY_2814 T T Y K L I L N L K Q L K E E A I K E A V D A G T A E K Y F K L Y A N A K T V E G V W T Y K D E T K K F T V T E
BINARY_2815 T T Y K L I L N L K Q L K E E A I K E A V D A G T A E K Y F K L Y A N A K T V E G V W T Y K D E T K T A T V T E
BINARY_2816 T T Y K L I L N L K Q L K E E A I K E A V D A G T A E K Y F K L Y A N A K T V E G V W T Y K D E T K T F T V T E
BINARY_2817 T T Y K L I L N L K Q L K E E A I K E A V D A A I A E K Y I K L I A N A K T V E G V W T L K D E I L K A T V T E
BINARY_2818 T T Y K L I L N L K Q L K E E A I K E A V D A A I A E K Y I K L I A N A K T V E G V W T L K D E I L K F T V T E
BINARY_2819 T T Y K L I L N L K Q L K E E A I K E A V D A A I A E K Y I K L I A N A K T V E G V W T L K D E I L T A T V T E
BINARY_2820 T T Y K L I L N L K Q L K E E A I K E A V D A A I A E K Y I K L I A N A K T V E G V W T L K D E I L T F T V T E
BINARY_2821 T T Y K L I L N L K Q L K E E A I K E A V D A A I A E K Y I K L I A N A K T V E G V W T L K D E I K K A T V T E
BINARY_2822 T T Y K L I L N L K Q L K E E A I K E A V D A A I A E K Y I K L I A N A K T V E G V W T L K D E I K K F T V T E
BINARY_2823 T T Y K L I L N L K Q L K E E A I K E A V D A A I A E K Y I K L I A N A K T V E G V W T L K D E I K T A T V T E
BINARY_2824 T T Y K L I L N L K Q L K E E A I K E A V D A A I A E K Y I K L I A N A K T V E G V W T L K D E I K T F T V T E
BINARY_2825 T T Y K L I L N L K Q L K E E A I K E A V D A A I A E K Y I K L I A N A K T V E G V W T L K D E T L K A T V T E
BINARY_2826 T T Y K L I L N L K Q L K E E A I K E A V D A A I A E K Y I K L I A N A K T V E G V W T L K D E T L K F T V T E
BINARY_2827 T T Y K L I L N L K Q L K E E A I K E A V D A A I A E K Y I K L I A N A K T V E G V W T L K D E T L T A T V T E
BINARY_2828 T T Y K L I L N L K Q L K E E A I K E A V D A A I A E K Y I K L I A N A K T V E G V W T L K D E T L T F T V T E
BINARY_2829 T T Y K L I L N L K Q L K E E A I K E A V D A A I A E K Y I K L I A N A K T V E G V W T L K D E T K K A T V T E
BINARY_2830 T T Y K L I L N L K Q L K E E A I K E A V D A A I A E K Y I K L I A N A K T V E G V W T L K D E T K K F T V T E
BINARY_2831 T T Y K L I L N L K Q L K E E A I K E A V D A A I A E K Y I K L I A N A K T V E G V W T L K D E T K T A T V T E
BINARY_2832 T T Y K L I L N L K Q L K E E A I K E A V D A A I A E K Y I K L I A N A K T V E G V W T L K D E T K T F T V T E
BINARY_2833 T T Y K L I L N L K Q L K E E A I K E A V D A A I A E K Y I K L I A N A K T V E G V W T Y K D E I L K A T V T E
BINARY_2834 T T Y K L I L N L K Q L K E E A I K E A V D A A I A E K Y I K L I A N A K T V E G V W T Y K D E I L K F T V T E
BINARY_2835 T T Y K L I L N L K Q L K E E A I K E A V D A A I A E K Y I K L I A N A K T V E G V W T Y K D E I L T A T V T E
BINARY_2836 T T Y K L I L N L K Q L K E E A I K E A V D A A I A E K Y I K L I A N A K T V E G V W T Y K D E I L T F T V T E
BINARY_2837 T T Y K L I L N L K Q L K E E A I K E A V D A A I A E K Y I K L I A N A K T V E G V W T Y K D E I K K A T V T E
BINARY_2838 T T Y K L I L N L K Q L K E E A I K E A V D A A I A E K Y I K L I A N A K T V E G V W T Y K D E I K K F T V T E
BINARY_2839 T T Y K L I L N L K Q L K E E A I K E A V D A A I A E K Y I K L I A N A K T V E G V W T Y K D E I K T A T V T E
BINARY_2840 T T Y K L I L N L K Q L K E E A I K E A V D A A I A E K Y I K L I A N A K T V E G V W T Y K D E I K T F T V T E
BINARY_2841 T T Y K L I L N L K Q L K E E A I K E A V D A A I A E K Y I K L I A N A K T V E G V W T Y K D E T L K A T V T E
BINARY_2842 T T Y K L I L N L K Q L K E E A I K E A V D A A I A E K Y I K L I A N A K T V E G V W T Y K D E T L K F T V T E
BINARY_2843 T T Y K L I L N L K Q L K E E A I K E A V D A A I A E K Y I K L I A N A K T V E G V W T Y K D E T L T A T V T E
BINARY_2844 T T Y K L I L N L K Q L K E E A I K E A V D A A I A E K Y I K L I A N A K T V E G V W T Y K D E T L T F T V T E
BINARY_2845 T T Y K L I L N L K Q L K E E A I K E A V D A A I A E K Y I K L I A N A K T V E G V W T Y K D E T K K A T V T E
BINARY_2846 T T Y K L I L N L K Q L K E E A I K E A V D A A I A E K Y I K L I A N A K T V E G V W T Y K D E T K K F T V T E
BINARY_2847 T T Y K L I L N L K Q L K E E A I K E A V D A A I A E K Y I K L I A N A K T V E G V W T Y K D E T K T A T V T E
BINARY_2848 T T Y K L I L N L K Q L K E E A I K E A V D A A I A E K Y I K L I A N A K T V E G V W T Y K D E T K T F T V T E
BINARY_2849 T T Y K L I L N L K Q L K E E A I K E A V D A A I A E K Y I K L Y A N A K T V E G V W T L K D E I L K A T V T E
BINARY_2850 T T Y K L I L N L K Q L K E E A I K E A V D A A I A E K Y I K L Y A N A K T V E G V W T L K D E I L K F T V T E
BINARY_2851 T T Y K L I L N L K Q L K E E A I K E A V D A A I A E K Y I K L Y A N A K T V E G V W T L K D E I L T A T V T E
BINARY_2852 T T Y K L I L N L K Q L K E E A I K E A V D A A I A E K Y I K L Y A N A K T V E G V W T L K D E I L T F T V T E
BINARY_2853 T T Y K L I L N L K Q L K E E A I K E A V D A A I A E K Y I K L Y A N A K T V E G V W T L K D E I K K A T V T E
BINARY_2854 T T Y K L I L N L K Q L K E E A I K E A V D A A I A E K Y I K L Y A N A K T V E G V W T L K D E I K K F T V T E
BINARY_2855 T T Y K L I L N L K Q L K E E A I K E A V D A A I A E K Y I K L Y A N A K T V E G V W T L K D E I K T A T V T E
BINARY_2856 T T Y K L I L N L K Q L K E E A I K E A V D A A I A E K Y I K L Y A N A K T V E G V W T L K D E I K T F T V T E
BINARY_2857 T T Y K L I L N L K Q L K E E A I K E A V D A A I A E K Y I K L Y A N A K T V E G V W T L K D E T L K A T V T E
BINARY_2858 T T Y K L I L N L K Q L K E E A I K E A V D A A I A E K Y I K L Y A N A K T V E G V W T L K D E T L K F T V T E
BINARY_2859 T T Y K L I L N L K Q L K E E A I K E A V D A A I A E K Y I K L Y A N A K T V E G V W T L K D E T L T A T V T E
BINARY_2860 T T Y K L I L N L K Q L K E E A I K E A V D A A I A E K Y I K L Y A N A K T V E G V W T L K D E T L T F T V T E
BINARY_2861 T T Y K L I L N L K Q L K E E A I K E A V D A A I A E K Y I K L Y A N A K T V E G V W T L K D E T K K A T V T E
BINARY_2862 T T Y K L I L N L K Q L K E E A I K E A V D A A I A E K Y I K L Y A N A K T V E G V W T L K D E T K K F T V T E
BINARY_2863 T T Y K L I L N L K Q L K E E A I K E A V D A A I A E K Y I K L Y A N A K T V E G V W T L K D E T K T A T V T E
BINARY_2864 T T Y K L I L N L K Q L K E E A I K E A V D A A I A E K Y I K L Y A N A K T V E G V W T L K D E T K T F T V T E
BINARY_2865 T T Y K L I L N L K Q L K E E A I K E A V D A A I A E K Y I K L Y A N A K T V E G V W T Y K D E I L K A T V T E
BINARY_2866 T T Y K L I L N L K Q L K E E A I K E A V D A A I A E K Y I K L Y A N A K T V E G V W T Y K D E I L K F T V T E

BINARY_2867 T T Y K L I L N L K Q L K E E A I K E A V D A A I A E K Y I K L Y A N A K T V E G V W T Y K D E I L T A T V T E

BINARY_2868 T T Y K L I L N L K Q L K E E A I K E A V D A A I A E K Y I K L Y A N A K T V E G V W T Y K D E I L T F T V T E

BINARY_2869 T T Y K L I L N L K Q L K E E A I K E A V D A A I A E K Y I K L Y A N A K T V E G V W T Y K D E I K K A T V T E

BINARY_2870 T T Y K L I L N L K Q L K E E A I K E A V D A A I A E K Y I K L Y A N A K T V E G V W T Y K D E I K K F T V T E

BINARY_2871 T T Y K L I L N L K Q L K E E A I K E A V D A A I A E K Y I K L Y A N A K T V E G V W T Y K D E I K T A T V T E

BINARY_2872 T T Y K L I L N L K Q L K E E A I K E A V D A A I A E K Y I K L Y A N A K T V E G V W T Y K D E I K T F T V T E

BINARY_2873 T T Y K L I L N L K Q L K E E A I K E A V D A A I A E K Y I K L Y A N A K T V E G V W T Y K D E T L K A T V T E

BINARY_2874 T T Y K L I L N L K Q L K E E A I K E A V D A A I A E K Y I K L Y A N A K T V E G V W T Y K D E T L K F T V T E

BINARY_2875 T T Y K L I L N L K Q L K E E A I K E A V D A A I A E K Y I K L Y A N A K T V E G V W T Y K D E T L T A T V T E

BINARY_2876 T T Y K L I L N L K Q L K E E A I K E A V D A A I A E K Y I K L Y A N A K T V E G V W T Y K D E T L T F T V T E

BINARY_2877 T T Y K L I L N L K Q L K E E A I K E A V D A A I A E K Y I K L Y A N A K T V E G V W T Y K D E T K K A T V T E

BINARY_2878 T T Y K L I L N L K Q L K E E A I K E A V D A A I A E K Y I K L Y A N A K T V E G V W T Y K D E T K K F T V T E

BINARY_2879 T T Y K L I L N L K Q L K E E A I K E A V D A A I A E K Y I K L Y A N A K T V E G V W T Y K D E T K T A T V T E

BINARY_2880 T T Y K L I L N L K Q L K E E A I K E A V D A A I A E K Y I K L Y A N A K T V E G V W T Y K D E T K T F T V T E

BINARY_2881 T T Y K L I L N L K Q L K E E A I K E A V D A A I A E K Y F K L I A N A K T V E G V W T L K D E I L K A T V T E

BINARY_2882 T T Y K L I L N L K Q L K E E A I K E A V D A A I A E K Y F K L I A N A K T V E G V W T L K D E I L K F T V T E

BINARY_2883 T T Y K L I L N L K Q L K E E A I K E A V D A A I A E K Y F K L I A N A K T V E G V W T L K D E I L T A T V T E

BINARY_2884 T T Y K L I L N L K Q L K E E A I K E A V D A A I A E K Y F K L I A N A K T V E G V W T L K D E I L T F T V T E

BINARY_2885 T T Y K L I L N L K Q L K E E A I K E A V D A A I A E K Y F K L I A N A K T V E G V W T L K D E I K K A T V T E

BINARY_2886 T T Y K L I L N L K Q L K E E A I K E A V D A A I A E K Y F K L I A N A K T V E G V W T L K D E I K K F T V T E

BINARY_2887 T T Y K L I L N L K Q L K E E A I K E A V D A A I A E K Y F K L I A N A K T V E G V W T L K D E I K T A T V T E

BINARY_2888 T T Y K L I L N L K Q L K E E A I K E A V D A A I A E K Y F K L I A N A K T V E G V W T L K D E I K T F T V T E

BINARY_2889 T T Y K L I L N L K Q L K E E A I K E A V D A A I A E K Y F K L I A N A K T V E G V W T L K D E T L K A T V T E

BINARY_2890 T T Y K L I L N L K Q L K E E A I K E A V D A A I A E K Y F K L I A N A K T V E G V W T L K D E T L K F T V T E

BINARY_2891 T T Y K L I L N L K Q L K E E A I K E A V D A A I A E K Y F K L I A N A K T V E G V W T L K D E T L T A T V T E

BINARY_2892 T T Y K L I L N L K Q L K E E A I K E A V D A A I A E K Y F K L I A N A K T V E G V W T L K D E T L T F T V T E

BINARY_2893 T T Y K L I L N L K Q L K E E A I K E A V D A A I A E K Y F K L I A N A K T V E G V W T L K D E T K K A T V T E

BINARY_2894 T T Y K L I L N L K Q L K E E A I K E A V D A A I A E K Y F K L I A N A K T V E G V W T L K D E T K K F T V T E

BINARY_2895 T T Y K L I L N L K Q L K E E A I K E A V D A A I A E K Y F K L I A N A K T V E G V W T L K D E T K T A T V T E

BINARY_2896 T T Y K L I L N L K Q L K E E A I K E A V D A A I A E K Y F K L I A N A K T V E G V W T L K D E T K T F T V T E

BINARY_2897 T T Y K L I L N L K Q L K E E A I K E A V D A A I A E K Y F K L I A N A K T V E G V W T Y K D E I L K A T V T E

BINARY_2898 T T Y K L I L N L K Q L K E E A I K E A V D A A I A E K Y F K L I A N A K T V E G V W T Y K D E I L K F T V T E

BINARY_2899 T T Y K L I L N L K Q L K E E A I K E A V D A A I A E K Y F K L I A N A K T V E G V W T Y K D E I L T A T V T E

BINARY_2900 T T Y K L I L N L K Q L K E E A I K E A V D A A I A E K Y F K L I A N A K T V E G V W T Y K D E I L T F T V T E

BINARY_2901 T T Y K L I L N L K Q L K E E A I K E A V D A A I A E K Y F K L I A N A K T V E G V W T Y K D E I K K A T V T E

BINARY_2902 T T Y K L I L N L K Q L K E E A I K E A V D A A I A E K Y F K L I A N A K T V E G V W T Y K D E I K K F T V T E

BINARY_2903 T T Y K L I L N L K Q L K E E A I K E A V D A A I A E K Y F K L I A N A K T V E G V W T Y K D E I K T A T V T E

BINARY_2904 T T Y K L I L N L K Q L K E E A I K E A V D A A I A E K Y F K L I A N A K T V E G V W T Y K D E I K T F T V T E

BINARY_2905 T T Y K L I L N L K Q L K E E A I K E A V D A A I A E K Y F K L I A N A K T V E G V W T Y K D E T L K A T V T E

BINARY_2906 T T Y K L I L N L K Q L K E E A I K E A V D A A I A E K Y F K L I A N A K T V E G V W T Y K D E T L K F T V T E

BINARY_2907 T T Y K L I L N L K Q L K E E A I K E A V D A A I A E K Y F K L I A N A K T V E G V W T Y K D E T L T A T V T E

BINARY_2908 T T Y K L I L N L K Q L K E E A I K E A V D A A I A E K Y F K L I A N A K T V E G V W T Y K D E T L T F T V T E

BINARY_2909 T T Y K L I L N L K Q L K E E A I K E A V D A A I A E K Y F K L I A N A K T V E G V W T Y K D E T K K A T V T E

BINARY_2910 T T Y K L I L N L K Q L K E E A I K E A V D A A I A E K Y F K L I A N A K T V E G V W T Y K D E T K K F T V T E

BINARY_2911 T T Y K L I L N L K Q L K E E A I K E A V D A A I A E K Y F K L I A N A K T V E G V W T Y K D E T K T A T V T E

BINARY_2912 T T Y K L I L N L K Q L K E E A I K E A V D A A I A E K Y F K L I A N A K T V E G V W T Y K D E T K T F T V T E

BINARY_2913 T T Y K L I L N L K Q L K E E A I K E A V D A A I A E K Y F K L Y A N A K T V E G V W T L K D E I L K A T V T E

BINARY_2914 T T Y K L I L N L K Q L K E E A I K E A V D A A I A E K Y F K L Y A N A K T V E G V W T L K D E I L K F T V T E

BINARY_2915 T T Y K L I L N L K Q L K E E A I K E A V D A A I A E K Y F K L Y A N A K T V E G V W T L K D E I L T A T V T E

BINARY_2916 T T Y K L I L N L K Q L K E E A I K E A V D A A I A E K Y F K L Y A N A K T V E G V W T L K D E I L T F T V T E

BINARY_2917 T T Y K L I L N L K Q L K E E A I K E A V D A A I A E K Y F K L Y A N A K T V E G V W T L K D E I K K A T V T E

BINARY_2918 T T Y K L I L N L K Q L K E E A I K E A V D A A I A E K Y F K L Y A N A K T V E G V W T L K D E I K K F T V T E

BINARY_2919 T T Y K L I L N L K Q L K E E A I K E A V D A A I A E K Y F K L Y A N A K T V E G V W T L K D E I K T A T V T E

BINARY_2920 T T Y K L I L N L K Q L K E E A I K E A V D A A I A E K Y F K L Y A N A K T V E G V W T L K D E I K T F T V T E

BINARY_2921 T T Y K L I L N L K Q L K E E A I K E A V D A A I A E K Y F K L Y A N A K T V E G V W T L K D E T L K A T V T E

BINARY_2922 T T Y K L I L N L K Q L K E E A I K E A V D A A I A E K Y F K L Y A N A K T V E G V W T L K D E T L K F T V T E

BINARY_2923 T T Y K L I L N L K Q L K E E A I K E A V D A A I A E K Y F K L Y A N A K T V E G V W T L K D E T L T A T V T E

BINARY_2924 T T Y K L I L N L K Q L K E E A I K E A V D A A I A E K Y F K L Y A N A K T V E G V W T L K D E T L T F T V T E

BINARY_2925 T T Y K L I L N L K Q L K E E A I K E A V D A A I A E K Y F K L Y A N A K T V E G V W T L K D E T K K A T V T E

BINARY_2926 T T Y K L I L N L K Q L K E E A I K E A V D A A I A E K Y F K L Y A N A K T V E G V W T L K D E T K K F T V T E

BINARY_2927 T T Y K L I L N L K Q L K E E A I K E A V D A A I A E K Y F K L Y A N A K T V E G V W T L K D E T K T A T V T E

BINARY_2928 T T Y K L I L N L K Q L K E E A I K E A V D A A I A E K Y F K L Y A N A K T V E G V W T L K D E T K T F T V T E

BINARY_2929 T T Y K L I L N L K Q L K E E A I K E A V D A A I A E K Y F K L Y A N A K T V E G V W T Y K D E I L K A T V T E

BINARY_2930 T T Y K L I L N L K Q L K E E A I K E A V D A A I A E K Y F K L Y A N A K T V E G V W T Y K D E I L K F T V T E

BINARY_2931 T T Y K L I L N L K Q L K E E A I K E A V D A A I A E K Y F K L Y A N A K T V E G V W T Y K D E I L T A T V T E

BINARY_2932 T T Y K L I L N L K Q L K E E A I K E A V D A A I A E K Y F K L Y A N A K T V E G V W T Y K D E I L T F T V T E

BINARY_2933 T T Y K L I L N L K Q L K E E A I K E A V D A A I A E K Y F K L Y A N A K T V E G V W T Y K D E I K K A T V T E

BINARY_2934 T T Y K L I L N L K Q L K E E A I K E A V D A A I A E K Y F K L Y A N A K T V E G V W T Y K D E I K K F T V T E

BINARY_2935 T T Y K L I L N L K Q L K E E A I K E A V D A A I A E K Y F K L Y A N A K T V E G V W T Y K D E I K T A T V T E

BINARY_2936 T T Y K L I L N L K Q L K E E A I K E A V D A A I A E K Y F K L Y A N A K T V E G V W T Y K D E I K T F T V T E

BINARY_2937 T T Y K L I L N L K Q L K E E A I K E A V D A A I A E K Y F K L Y A N A K T V E G V W T Y K D E T L K A T V T E

BINARY_2938 T T Y K L I L N L K Q L K E E A I K E A V D A A I A E K Y F K L Y A N A K T V E G V W T Y K D E T L K F T V T E

BINARY_2939 T T Y K L I L N L K Q L K E E A I K E A V D A A I A E K Y F K L Y A N A K T V E G V W T Y K D E T L T A T V T E

BINARY_2940 T T Y K L I L N L K Q L K E E A I K E A V D A A I A E K Y F K L Y A N A K T V E G V W T Y K D E T L T F T V T E

BINARY_2941 T T Y K L I L N L K Q L K E E A I K E A V D A A I A E K Y F K L Y A N A K T V E G V W T Y K D E T K K A T V T E

BINARY_2942 T T Y K L I L N L K Q L K E E A I K E A V D A A I A E K Y F K L Y A N A K T V E G V W T Y K D E T K K F T V T E

BINARY_2943 T T Y K L I L N L K Q L K E E A I K E A V D A A I A E K Y F K L Y A N A K T V E G V W T Y K D E T K T A T V T E

BINARY_2944 T T Y K L I L N L K Q L K E E A I K E A V D A A I A E K Y F K L Y A N A K T V E G V W T Y K D E T K T F T V T E

BINARY_2945 T T Y K L I L N L K Q L K E E A I K E A V D A A T A E K Y I K L I A N A K T V E G V W T L K D E I L K A T V T E

BINARY_2946 T T Y K L I L N L K Q L K E E A I K E A V D A A T A E K Y I K L I A N A K T V E G V W T L K D E I L K F T V T E

BINARY_2947 T T Y K L I L N L K Q L K E E A I K E A V D A A T A E K Y I K L I A N A K T V E G V W T L K D E I L T A T V T E

BINARY_2948 T T Y K L I L N L K Q L K E E A I K E A V D A A T A E K Y I K L I A N A K T V E G V W T L K D E I L T F T V T E

BINARY_2949 T T Y K L I L N L K Q L K E E A I K E A V D A A T A E K Y I K L I A N A K T V E G V W T L K D E I K K A T V T E

BINARY_2950 T T Y K L I L N L K Q L K E E A I K E A V D A A T A E K Y I K L I A N A K T V E G V W T L K D E I K K F T V T E

BINARY_2951 T T Y K L I L N L K Q L K E E A I K E A V D A A T A E K Y I K L I A N A K T V E G V W T L K D E I K T A T V T E

BINARY_2952 T T Y K L I L N L K Q L K E E A I K E A V D A A T A E K Y I K L I A N A K T V E G V W T L K D E I K T F T V T E

BINARY_2953 T T Y K L I L N L K Q L K E E A I K E A V D A A T A E K Y I K L I A N A K T V E G V W T L K D E T L K A T V T E

BINARY_2954 T T Y K L I L N L K Q L K E E A I K E A V D A A T A E K Y I K L I A N A K T V E G V W T L K D E T L K F T V T E

BINARY_2955 T T Y K L I L N L K Q L K E E A I K E A V D A A T A E K Y I K L I A N A K T V E G V W T L K D E T L T A T V T E

BINARY_2956 T T Y K L I L N L K Q L K E E A I K E A V D A A T A E K Y I K L I A N A K T V E G V W T L K D E T L T F T V T E

BINARY_2957 T T Y K L I L N L K Q L K E E A I K E A V D A A T A E K Y I K L I A N A K T V E G V W T L K D E T K K A T V T E

BINARY_2958 T T Y K L I L N L K Q L K E E A I K E A V D A A T A E K Y I K L I A N A K T V E G V W T L K D E T K K F T V T E

BINARY_2959 T T Y K L I L N L K Q L K E E A I K E A V D A A T A E K Y I K L I A N A K T V E G V W T L K D E T K T A T V T E

BINARY_2960 T T Y K L I L N L K Q L K E E A I K E A V D A A T A E K Y I K L I A N A K T V E G V W T L K D E T K T F T V T E

BINARY_2961 T T Y K L I L N L K Q L K E E A I K E A V D A A T A E K Y I K L I A N A K T V E G V W T Y K D E I L K A T V T E

BINARY_2962 T T Y K L I L N L K Q L K E E A I K E A V D A A T A E K Y I K L I A N A K T V E G V W T Y K D E I L K F T V T E
BINARY_2963 T T Y K L I L N L K Q L K E E A I K E A V D A A T A E K Y I K L I A N A K T V E G V W T Y K D E I L T A T V T E
BINARY_2964 T T Y K L I L N L K Q L K E E A I K E A V D A A T A E K Y I K L I A N A K T V E G V W T Y K D E I L T F T V T E
BINARY_2965 T T Y K L I L N L K Q L K E E A I K E A V D A A T A E K Y I K L I A N A K T V E G V W T Y K D E I K K A T V T E
BINARY_2966 T T Y K L I L N L K Q L K E E A I K E A V D A A T A E K Y I K L I A N A K T V E G V W T Y K D E I K K F T V T E
BINARY_2967 T T Y K L I L N L K Q L K E E A I K E A V D A A T A E K Y I K L I A N A K T V E G V W T Y K D E I K T A T V T E
BINARY_2968 T T Y K L I L N L K Q L K E E A I K E A V D A A T A E K Y I K L I A N A K T V E G V W T Y K D E I K T F T V T E
BINARY_2969 T T Y K L I L N L K Q L K E E A I K E A V D A A T A E K Y I K L I A N A K T V E G V W T Y K D E T L K A T V T E
BINARY_2970 T T Y K L I L N L K Q L K E E A I K E A V D A A T A E K Y I K L I A N A K T V E G V W T Y K D E T L K F T V T E
BINARY_2971 T T Y K L I L N L K Q L K E E A I K E A V D A A T A E K Y I K L I A N A K T V E G V W T Y K D E T L T A T V T E
BINARY_2972 T T Y K L I L N L K Q L K E E A I K E A V D A A T A E K Y I K L I A N A K T V E G V W T Y K D E T L T F T V T E
BINARY_2973 T T Y K L I L N L K Q L K E E A I K E A V D A A T A E K Y I K L I A N A K T V E G V W T Y K D E T K K A T V T E
BINARY_2974 T T Y K L I L N L K Q L K E E A I K E A V D A A T A E K Y I K L I A N A K T V E G V W T Y K D E T K K F T V T E
BINARY_2975 T T Y K L I L N L K Q L K E E A I K E A V D A A T A E K Y I K L I A N A K T V E G V W T Y K D E T K T A T V T E
BINARY_2976 T T Y K L I L N L K Q L K E E A I K E A V D A A T A E K Y I K L I A N A K T V E G V W T Y K D E T K T F T V T E
BINARY_2977 T T Y K L I L N L K Q L K E E A I K E A V D A A T A E K Y I K L Y A N A K T V E G V W T L K D E I L K A T V T E
BINARY_2978 T T Y K L I L N L K Q L K E E A I K E A V D A A T A E K Y I K L Y A N A K T V E G V W T L K D E I L K F T V T E
BINARY_2979 T T Y K L I L N L K Q L K E E A I K E A V D A A T A E K Y I K L Y A N A K T V E G V W T L K D E I L T A T V T E
BINARY_2980 T T Y K L I L N L K Q L K E E A I K E A V D A A T A E K Y I K L Y A N A K T V E G V W T L K D E I L T F T V T E
BINARY_2981 T T Y K L I L N L K Q L K E E A I K E A V D A A T A E K Y I K L Y A N A K T V E G V W T L K D E I K K A T V T E
BINARY_2982 T T Y K L I L N L K Q L K E E A I K E A V D A A T A E K Y I K L Y A N A K T V E G V W T L K D E I K K F T V T E
BINARY_2983 T T Y K L I L N L K Q L K E E A I K E A V D A A T A E K Y I K L Y A N A K T V E G V W T L K D E I K T A T V T E
BINARY_2984 T T Y K L I L N L K Q L K E E A I K E A V D A A T A E K Y I K L Y A N A K T V E G V W T L K D E I K T F T V T E
BINARY_2985 T T Y K L I L N L K Q L K E E A I K E A V D A A T A E K Y I K L Y A N A K T V E G V W T L K D E T L K A T V T E
BINARY_2986 T T Y K L I L N L K Q L K E E A I K E A V D A A T A E K Y I K L Y A N A K T V E G V W T L K D E T L K F T V T E
BINARY_2987 T T Y K L I L N L K Q L K E E A I K E A V D A A T A E K Y I K L Y A N A K T V E G V W T L K D E T L T A T V T E
BINARY_2988 T T Y K L I L N L K Q L K E E A I K E A V D A A T A E K Y I K L Y A N A K T V E G V W T L K D E T L T F T V T E
BINARY_2989 T T Y K L I L N L K Q L K E E A I K E A V D A A T A E K Y I K L Y A N A K T V E G V W T L K D E T K K A T V T E
BINARY_2990 T T Y K L I L N L K Q L K E E A I K E A V D A A T A E K Y I K L Y A N A K T V E G V W T L K D E T K K F T V T E
BINARY_2991 T T Y K L I L N L K Q L K E E A I K E A V D A A T A E K Y I K L Y A N A K T V E G V W T L K D E T K T A T V T E
BINARY_2992 T T Y K L I L N L K Q L K E E A I K E A V D A A T A E K Y I K L Y A N A K T V E G V W T L K D E T K T F T V T E
BINARY_2993 T T Y K L I L N L K Q L K E E A I K E A V D A A T A E K Y I K L Y A N A K T V E G V W T Y K D E I L K A T V T E
BINARY_2994 T T Y K L I L N L K Q L K E E A I K E A V D A A T A E K Y I K L Y A N A K T V E G V W T Y K D E I L K F T V T E
BINARY_2995 T T Y K L I L N L K Q L K E E A I K E A V D A A T A E K Y I K L Y A N A K T V E G V W T Y K D E I L T A T V T E
BINARY_2996 T T Y K L I L N L K Q L K E E A I K E A V D A A T A E K Y I K L Y A N A K T V E G V W T Y K D E I L T F T V T E
BINARY_2997 T T Y K L I L N L K Q L K E E A I K E A V D A A T A E K Y I K L Y A N A K T V E G V W T Y K D E I K K A T V T E
BINARY_2998 T T Y K L I L N L K Q L K E E A I K E A V D A A T A E K Y I K L Y A N A K T V E G V W T Y K D E I K K F T V T E
BINARY_2999 T T Y K L I L N L K Q L K E E A I K E A V D A A T A E K Y I K L Y A N A K T V E G V W T Y K D E I K T A T V T E
BINARY_3000 T T Y K L I L N L K Q L K E E A I K E A V D A A T A E K Y I K L Y A N A K T V E G V W T Y K D E I K T F T V T E
BINARY_3001 T T Y K L I L N L K Q L K E E A I K E A V D A A T A E K Y I K L Y A N A K T V E G V W T Y K D E T L K A T V T E
BINARY_3002 T T Y K L I L N L K Q L K E E A I K E A V D A A T A E K Y I K L Y A N A K T V E G V W T Y K D E T L K F T V T E
BINARY_3003 T T Y K L I L N L K Q L K E E A I K E A V D A A T A E K Y I K L Y A N A K T V E G V W T Y K D E T L T A T V T E
BINARY_3004 T T Y K L I L N L K Q L K E E A I K E A V D A A T A E K Y I K L Y A N A K T V E G V W T Y K D E T L T F T V T E
BINARY_3005 T T Y K L I L N L K Q L K E E A I K E A V D A A T A E K Y I K L Y A N A K T V E G V W T Y K D E T K K A T V T E
BINARY_3006 T T Y K L I L N L K Q L K E E A I K E A V D A A T A E K Y I K L Y A N A K T V E G V W T Y K D E T K K F T V T E
BINARY_3007 T T Y K L I L N L K Q L K E E A I K E A V D A A T A E K Y I K L Y A N A K T V E G V W T Y K D E T K T A T V T E
BINARY_3008 T T Y K L I L N L K Q L K E E A I K E A V D A A T A E K Y I K L Y A N A K T V E G V W T Y K D E T K T F T V T E
BINARY_3009 T T Y K L I L N L K Q L K E E A I K E A V D A A T A E K Y F K L I A N A K T V E G V W T L K D E I L K A T V T E
BINARY_3010 T T Y K L I L N L K Q L K E E A I K E A V D A A T A E K Y F K L I A N A K T V E G V W T L K D E I L K F T V T E
BINARY_3011 T T Y K L I L N L K Q L K E E A I K E A V D A A T A E K Y F K L I A N A K T V E G V W T L K D E I L T A T V T E
BINARY_3012 T T Y K L I L N L K Q L K E E A I K E A V D A A T A E K Y F K L I A N A K T V E G V W T L K D E I L T F T V T E
BINARY_3013 T T Y K L I L N L K Q L K E E A I K E A V D A A T A E K Y F K L I A N A K T V E G V W T L K D E I K K A T V T E
BINARY_3014 T T Y K L I L N L K Q L K E E A I K E A V D A A T A E K Y F K L I A N A K T V E G V W T L K D E I K K F T V T E
BINARY_3015 T T Y K L I L N L K Q L K E E A I K E A V D A A T A E K Y F K L I A N A K T V E G V W T L K D E I K T A T V T E
BINARY_3016 T T Y K L I L N L K Q L K E E A I K E A V D A A T A E K Y F K L I A N A K T V E G V W T L K D E I K T F T V T E
BINARY_3017 T T Y K L I L N L K Q L K E E A I K E A V D A A T A E K Y F K L I A N A K T V E G V W T L K D E T L K A T V T E
BINARY_3018 T T Y K L I L N L K Q L K E E A I K E A V D A A T A E K Y F K L I A N A K T V E G V W T L K D E T L K F T V T E
BINARY_3019 T T Y K L I L N L K Q L K E E A I K E A V D A A T A E K Y F K L I A N A K T V E G V W T L K D E T L T A T V T E
BINARY_3020 T T Y K L I L N L K Q L K E E A I K E A V D A A T A E K Y F K L I A N A K T V E G V W T L K D E T L T F T V T E
BINARY_3021 T T Y K L I L N L K Q L K E E A I K E A V D A A T A E K Y F K L I A N A K T V E G V W T L K D E T K K A T V T E
BINARY_3022 T T Y K L I L N L K Q L K E E A I K E A V D A A T A E K Y F K L I A N A K T V E G V W T L K D E T K K F T V T E
BINARY_3023 T T Y K L I L N L K Q L K E E A I K E A V D A A T A E K Y F K L I A N A K T V E G V W T L K D E T K T A T V T E
BINARY_3024 T T Y K L I L N L K Q L K E E A I K E A V D A A T A E K Y F K L I A N A K T V E G V W T L K D E T K T F T V T E
BINARY_3025 T T Y K L I L N L K Q L K E E A I K E A V D A A T A E K Y F K L I A N A K T V E G V W T Y K D E I L K A T V T E
BINARY_3026 T T Y K L I L N L K Q L K E E A I K E A V D A A T A E K Y F K L I A N A K T V E G V W T Y K D E I L K F T V T E
BINARY_3027 T T Y K L I L N L K Q L K E E A I K E A V D A A T A E K Y F K L I A N A K T V E G V W T Y K D E I L T A T V T E
BINARY_3028 T T Y K L I L N L K Q L K E E A I K E A V D A A T A E K Y F K L I A N A K T V E G V W T Y K D E I L T F T V T E
BINARY_3029 T T Y K L I L N L K Q L K E E A I K E A V D A A T A E K Y F K L I A N A K T V E G V W T Y K D E I K K A T V T E
BINARY_3030 T T Y K L I L N L K Q L K E E A I K E A V D A A T A E K Y F K L I A N A K T V E G V W T Y K D E I K K F T V T E
BINARY_3031 T T Y K L I L N L K Q L K E E A I K E A V D A A T A E K Y F K L I A N A K T V E G V W T Y K D E I K T A T V T E
BINARY_3032 T T Y K L I L N L K Q L K E E A I K E A V D A A T A E K Y F K L I A N A K T V E G V W T Y K D E I K T F T V T E
BINARY_3033 T T Y K L I L N L K Q L K E E A I K E A V D A A T A E K Y F K L I A N A K T V E G V W T Y K D E T L K A T V T E
BINARY_3034 T T Y K L I L N L K Q L K E E A I K E A V D A A T A E K Y F K L I A N A K T V E G V W T Y K D E T L K F T V T E
BINARY_3035 T T Y K L I L N L K Q L K E E A I K E A V D A A T A E K Y F K L I A N A K T V E G V W T Y K D E T L T A T V T E
BINARY_3036 T T Y K L I L N L K Q L K E E A I K E A V D A A T A E K Y F K L I A N A K T V E G V W T Y K D E T L T F T V T E
BINARY_3037 T T Y K L I L N L K Q L K E E A I K E A V D A A T A E K Y F K L I A N A K T V E G V W T Y K D E T K K A T V T E
BINARY_3038 T T Y K L I L N L K Q L K E E A I K E A V D A A T A E K Y F K L I A N A K T V E G V W T Y K D E T K K F T V T E
BINARY_3039 T T Y K L I L N L K Q L K E E A I K E A V D A A T A E K Y F K L I A N A K T V E G V W T Y K D E T K T A T V T E
BINARY_3040 T T Y K L I L N L K Q L K E E A I K E A V D A A T A E K Y F K L I A N A K T V E G V W T Y K D E T K T F T V T E
BINARY_3041 T T Y K L I L N L K Q L K E E A I K E A V D A A T A E K Y F K L Y A N A K T V E G V W T L K D E I L K A T V T E
BINARY_3042 T T Y K L I L N L K Q L K E E A I K E A V D A A T A E K Y F K L Y A N A K T V E G V W T L K D E I L K F T V T E
BINARY_3043 T T Y K L I L N L K Q L K E E A I K E A V D A A T A E K Y F K L Y A N A K T V E G V W T L K D E I L T A T V T E
BINARY_3044 T T Y K L I L N L K Q L K E E A I K E A V D A A T A E K Y F K L Y A N A K T V E G V W T L K D E I L T F T V T E
BINARY_3045 T T Y K L I L N L K Q L K E E A I K E A V D A A T A E K Y F K L Y A N A K T V E G V W T L K D E I K K A T V T E
BINARY_3046 T T Y K L I L N L K Q L K E E A I K E A V D A A T A E K Y F K L Y A N A K T V E G V W T L K D E I K K F T V T E
BINARY_3047 T T Y K L I L N L K Q L K E E A I K E A V D A A T A E K Y F K L Y A N A K T V E G V W T L K D E I K T A T V T E
BINARY_3048 T T Y K L I L N L K Q L K E E A I K E A V D A A T A E K Y F K L Y A N A K T V E G V W T L K D E I K T F T V T E
BINARY_3049 T T Y K L I L N L K Q L K E E A I K E A V D A A T A E K Y F K L Y A N A K T V E G V W T L K D E T L K A T V T E
BINARY_3050 T T Y K L I L N L K Q L K E E A I K E A V D A A T A E K Y F K L Y A N A K T V E G V W T L K D E T L K F T V T E
BINARY_3051 T T Y K L I L N L K Q L K E E A I K E A V D A A T A E K Y F K L Y A N A K T V E G V W T L K D E T L T A T V T E
BINARY_3052 T T Y K L I L N L K Q L K E E A I K E A V D A A T A E K Y F K L Y A N A K T V E G V W T L K D E T L T F T V T E
BINARY_3053 T T Y K L I L N L K Q L K E E A I K E A V D A A T A E K Y F K L Y A N A K T V E G V W T L K D E T K K A T V T E
BINARY_3054 T T Y K L I L N L K Q L K E E A I K E A V D A A T A E K Y F K L Y A N A K T V E G V W T L K D E T K K F T V T E
BINARY_3055 T T Y K L I L N L K Q L K E E A I K E A V D A A T A E K Y F K L Y A N A K T V E G V W T L K D E T K T A T V T E
BINARY_3056 T T Y K L I L N L K Q L K E E A I K E A V D A A T A E K Y F K L Y A N A K T V E G V W T L K D E T K T F T V T E

BINARY_3057 T T Y K L I L N L K Q L K E E A I K E A V D A A T A E K Y F K L Y A N A K T V E G V W T Y K D E I L K A T V E
BINARY_3058 T T Y K L I L N L K Q L K E E A I K E A V D A A T A E K Y F K L Y A N A K T V E G V W T Y K D E I L K F T V E
BINARY_3059 T T Y K L I L N L K Q L K E E A I K E A V D A A T A E K Y F K L Y A N A K T V E G V W T Y K D E I L T A T V E
BINARY_3060 T T Y K L I L N L K Q L K E E A I K E A V D A A T A E K Y F K L Y A N A K T V E G V W T Y K D E I L T F T V E
BINARY_3061 T T Y K L I L N L K Q L K E E A I K E A V D A A T A E K Y F K L Y A N A K T V E G V W T Y K D E I K K A T V E
BINARY_3062 T T Y K L I L N L K Q L K E E A I K E A V D A A T A E K Y F K L Y A N A K T V E G V W T Y K D E I K K F T V E
BINARY_3063 T T Y K L I L N L K Q L K E E A I K E A V D A A T A E K Y F K L Y A N A K T V E G V W T Y K D E I K T A T V E
BINARY_3064 T T Y K L I L N L K Q L K E E A I K E A V D A A T A E K Y F K L Y A N A K T V E G V W T Y K D E I K T F T V E
BINARY_3065 T T Y K L I L N L K Q L K E E A I K E A V D A A T A E K Y F K L Y A N A K T V E G V W T Y K D E T L K A T V E
BINARY_3066 T T Y K L I L N L K Q L K E E A I K E A V D A A T A E K Y F K L Y A N A K T V E G V W T Y K D E T L K F T V E
BINARY_3067 T T Y K L I L N L K Q L K E E A I K E A V D A A T A E K Y F K L Y A N A K T V E G V W T Y K D E T L T A T V E
BINARY_3068 T T Y K L I L N L K Q L K E E A I K E A V D A A T A E K Y F K L Y A N A K T V E G V W T Y K D E T L T F T V E
BINARY_3069 T T Y K L I L N L K Q L K E E A I K E A V D A A T A E K Y F K L Y A N A K T V E G V W T Y K D E T K K A T V E
BINARY_3070 T T Y K L I L N L K Q L K E E A I K E A V D A A T A E K Y F K L Y A N A K T V E G V W T Y K D E T K K F T V E
BINARY_3071 T T Y K L I L N L K Q L K E E A I K E A V D A A T A E K Y F K L Y A N A K T V E G V W T Y K D E T K T A T V E
BINARY_3072 T T Y K L I L N L K Q L K E E A I K E A V D A A T A E K Y F K L Y A N A K T V E G V W T Y K D E T K T F T V E
BINARY_3073 T T Y K L I L N L K Q L K E E A I T E L V D A G I A E K Y I K L I A N A K T V E G V W T L K D E I L K A T V E
BINARY_3074 T T Y K L I L N L K Q L K E E A I T E L V D A G I A E K Y I K L I A N A K T V E G V W T L K D E I L K F T V E
BINARY_3075 T T Y K L I L N L K Q L K E E A I T E L V D A G I A E K Y I K L I A N A K T V E G V W T L K D E I L T A T V E
BINARY_3076 T T Y K L I L N L K Q L K E E A I T E L V D A G I A E K Y I K L I A N A K T V E G V W T L K D E I L T F T V E
BINARY_3077 T T Y K L I L N L K Q L K E E A I T E L V D A G I A E K Y I K L I A N A K T V E G V W T L K D E I K K A T V E
BINARY_3078 T T Y K L I L N L K Q L K E E A I T E L V D A G I A E K Y I K L I A N A K T V E G V W T L K D E I K K F T V E
BINARY_3079 T T Y K L I L N L K Q L K E E A I T E L V D A G I A E K Y I K L I A N A K T V E G V W T L K D E I K T A T V E
BINARY_3080 T T Y K L I L N L K Q L K E E A I T E L V D A G I A E K Y I K L I A N A K T V E G V W T L K D E I K T F T V E
BINARY_3081 T T Y K L I L N L K Q L K E E A I T E L V D A G I A E K Y I K L I A N A K T V E G V W T L K D E T L K A T V E
BINARY_3082 T T Y K L I L N L K Q L K E E A I T E L V D A G I A E K Y I K L I A N A K T V E G V W T L K D E T L K F T V E
BINARY_3083 T T Y K L I L N L K Q L K E E A I T E L V D A G I A E K Y I K L I A N A K T V E G V W T L K D E T L T A T V E
BINARY_3084 T T Y K L I L N L K Q L K E E A I T E L V D A G I A E K Y I K L I A N A K T V E G V W T L K D E T L T F T V E
BINARY_3085 T T Y K L I L N L K Q L K E E A I T E L V D A G I A E K Y I K L I A N A K T V E G V W T L K D E T K K A T V E
BINARY_3086 T T Y K L I L N L K Q L K E E A I T E L V D A G I A E K Y I K L I A N A K T V E G V W T L K D E T K K F T V E
BINARY_3087 T T Y K L I L N L K Q L K E E A I T E L V D A G I A E K Y I K L I A N A K T V E G V W T L K D E T K T A T V E
BINARY_3088 T T Y K L I L N L K Q L K E E A I T E L V D A G I A E K Y I K L I A N A K T V E G V W T L K D E T K T F T V E
BINARY_3089 T T Y K L I L N L K Q L K E E A I T E L V D A G I A E K Y I K L I A N A K T V E G V W T Y K D E I L K A T V E
BINARY_3090 T T Y K L I L N L K Q L K E E A I T E L V D A G I A E K Y I K L I A N A K T V E G V W T Y K D E I L K F T V E
BINARY_3091 T T Y K L I L N L K Q L K E E A I T E L V D A G I A E K Y I K L I A N A K T V E G V W T Y K D E I L T A T V E
BINARY_3092 T T Y K L I L N L K Q L K E E A I T E L V D A G I A E K Y I K L I A N A K T V E G V W T Y K D E I L T F T V E
BINARY_3093 T T Y K L I L N L K Q L K E E A I T E L V D A G I A E K Y I K L I A N A K T V E G V W T Y K D E I K K A T V E
BINARY_3094 T T Y K L I L N L K Q L K E E A I T E L V D A G I A E K Y I K L I A N A K T V E G V W T Y K D E I K K F T V E
BINARY_3095 T T Y K L I L N L K Q L K E E A I T E L V D A G I A E K Y I K L I A N A K T V E G V W T Y K D E I K T A T V E
BINARY_3096 T T Y K L I L N L K Q L K E E A I T E L V D A G I A E K Y I K L I A N A K T V E G V W T Y K D E I K T F T V E
BINARY_3097 T T Y K L I L N L K Q L K E E A I T E L V D A G I A E K Y I K L I A N A K T V E G V W T Y K D E T L K A T V E
BINARY_3098 T T Y K L I L N L K Q L K E E A I T E L V D A G I A E K Y I K L I A N A K T V E G V W T Y K D E T L K F T V E
BINARY_3099 T T Y K L I L N L K Q L K E E A I T E L V D A G I A E K Y I K L I A N A K T V E G V W T Y K D E T L T A T V E
BINARY_3100 T T Y K L I L N L K Q L K E E A I T E L V D A G I A E K Y I K L I A N A K T V E G V W T Y K D E T L T F T V E
BINARY_3101 T T Y K L I L N L K Q L K E E A I T E L V D A G I A E K Y I K L I A N A K T V E G V W T Y K D E T K K A T V E
BINARY_3102 T T Y K L I L N L K Q L K E E A I T E L V D A G I A E K Y I K L I A N A K T V E G V W T Y K D E T K K F T V E
BINARY_3103 T T Y K L I L N L K Q L K E E A I T E L V D A G I A E K Y I K L I A N A K T V E G V W T Y K D E T K T A T V E
BINARY_3104 T T Y K L I L N L K Q L K E E A I T E L V D A G I A E K Y I K L I A N A K T V E G V W T Y K D E T K T F T V E
BINARY_3105 T T Y K L I L N L K Q L K E E A I T E L V D A G I A E K Y I K L Y A N A K T V E G V W T L K D E I L K A T V E
BINARY_3106 T T Y K L I L N L K Q L K E E A I T E L V D A G I A E K Y I K L Y A N A K T V E G V W T L K D E I L K F T V E
BINARY_3107 T T Y K L I L N L K Q L K E E A I T E L V D A G I A E K Y I K L Y A N A K T V E G V W T L K D E I L T A T V E
BINARY_3108 T T Y K L I L N L K Q L K E E A I T E L V D A G I A E K Y I K L Y A N A K T V E G V W T L K D E I L T F T V E
BINARY_3109 T T Y K L I L N L K Q L K E E A I T E L V D A G I A E K Y I K L Y A N A K T V E G V W T L K D E I K K A T V E
BINARY_3110 T T Y K L I L N L K Q L K E E A I T E L V D A G I A E K Y I K L Y A N A K T V E G V W T L K D E I K K F T V E
BINARY_3111 T T Y K L I L N L K Q L K E E A I T E L V D A G I A E K Y I K L Y A N A K T V E G V W T L K D E I K T A T V E
BINARY_3112 T T Y K L I L N L K Q L K E E A I T E L V D A G I A E K Y I K L Y A N A K T V E G V W T L K D E I K T F T V E
BINARY_3113 T T Y K L I L N L K Q L K E E A I T E L V D A G I A E K Y I K L Y A N A K T V E G V W T L K D E T L K A T V E
BINARY_3114 T T Y K L I L N L K Q L K E E A I T E L V D A G I A E K Y I K L Y A N A K T V E G V W T L K D E T L K F T V E
BINARY_3115 T T Y K L I L N L K Q L K E E A I T E L V D A G I A E K Y I K L Y A N A K T V E G V W T L K D E T L T A T V E
BINARY_3116 T T Y K L I L N L K Q L K E E A I T E L V D A G I A E K Y I K L Y A N A K T V E G V W T L K D E T L T F T V E
BINARY_3117 T T Y K L I L N L K Q L K E E A I T E L V D A G I A E K Y I K L Y A N A K T V E G V W T L K D E T K K A T V E
BINARY_3118 T T Y K L I L N L K Q L K E E A I T E L V D A G I A E K Y I K L Y A N A K T V E G V W T L K D E T K K F T V E
BINARY_3119 T T Y K L I L N L K Q L K E E A I T E L V D A G I A E K Y I K L Y A N A K T V E G V W T L K D E T K T A T V E
BINARY_3120 T T Y K L I L N L K Q L K E E A I T E L V D A G I A E K Y I K L Y A N A K T V E G V W T L K D E T K T F T V E
BINARY_3121 T T Y K L I L N L K Q L K E E A I T E L V D A G I A E K Y I K L Y A N A K T V E G V W T Y K D E I L K A T V E
BINARY_3122 T T Y K L I L N L K Q L K E E A I T E L V D A G I A E K Y I K L Y A N A K T V E G V W T Y K D E I L K F T V E
BINARY_3123 T T Y K L I L N L K Q L K E E A I T E L V D A G I A E K Y I K L Y A N A K T V E G V W T Y K D E I L T A T V E
BINARY_3124 T T Y K L I L N L K Q L K E E A I T E L V D A G I A E K Y I K L Y A N A K T V E G V W T Y K D E I L T F T V E
BINARY_3125 T T Y K L I L N L K Q L K E E A I T E L V D A G I A E K Y I K L Y A N A K T V E G V W T Y K D E I K K A T V E
BINARY_3126 T T Y K L I L N L K Q L K E E A I T E L V D A G I A E K Y I K L Y A N A K T V E G V W T Y K D E I K K F T V E
BINARY_3127 T T Y K L I L N L K Q L K E E A I T E L V D A G I A E K Y I K L Y A N A K T V E G V W T Y K D E I K T A T V E
BINARY_3128 T T Y K L I L N L K Q L K E E A I T E L V D A G I A E K Y I K L Y A N A K T V E G V W T Y K D E I K T F T V E
BINARY_3129 T T Y K L I L N L K Q L K E E A I T E L V D A G I A E K Y I K L Y A N A K T V E G V W T Y K D E T L K A T V E
BINARY_3130 T T Y K L I L N L K Q L K E E A I T E L V D A G I A E K Y I K L Y A N A K T V E G V W T Y K D E T L K F T V E
BINARY_3131 T T Y K L I L N L K Q L K E E A I T E L V D A G I A E K Y I K L Y A N A K T V E G V W T Y K D E T L T A T V E
BINARY_3132 T T Y K L I L N L K Q L K E E A I T E L V D A G I A E K Y I K L Y A N A K T V E G V W T Y K D E T L T F T V E
BINARY_3133 T T Y K L I L N L K Q L K E E A I T E L V D A G I A E K Y I K L Y A N A K T V E G V W T Y K D E T K K A T V E
BINARY_3134 T T Y K L I L N L K Q L K E E A I T E L V D A G I A E K Y I K L Y A N A K T V E G V W T Y K D E T K K F T V E
BINARY_3135 T T Y K L I L N L K Q L K E E A I T E L V D A G I A E K Y I K L Y A N A K T V E G V W T Y K D E T K T A T V E
BINARY_3136 T T Y K L I L N L K Q L K E E A I T E L V D A G I A E K Y I K L Y A N A K T V E G V W T Y K D E T K T F T V E
BINARY_3137 T T Y K L I L N L K Q L K E E A I T E L V D A G I A E K Y F K L I A N A K T V E G V W T L K D E I L K A T V E
BINARY_3138 T T Y K L I L N L K Q L K E E A I T E L V D A G I A E K Y F K L I A N A K T V E G V W T L K D E I L K F T V E
BINARY_3139 T T Y K L I L N L K Q L K E E A I T E L V D A G I A E K Y F K L I A N A K T V E G V W T L K D E I L T A T V E
BINARY_3140 T T Y K L I L N L K Q L K E E A I T E L V D A G I A E K Y F K L I A N A K T V E G V W T L K D E I L T F T V E
BINARY_3141 T T Y K L I L N L K Q L K E E A I T E L V D A G I A E K Y F K L I A N A K T V E G V W T L K D E I K K A T V E
BINARY_3142 T T Y K L I L N L K Q L K E E A I T E L V D A G I A E K Y F K L I A N A K T V E G V W T L K D E I K K F T V E
BINARY_3143 T T Y K L I L N L K Q L K E E A I T E L V D A G I A E K Y F K L I A N A K T V E G V W T L K D E I K T A T V E
BINARY_3144 T T Y K L I L N L K Q L K E E A I T E L V D A G I A E K Y F K L I A N A K T V E G V W T L K D E I K T F T V E
BINARY_3145 T T Y K L I L N L K Q L K E E A I T E L V D A G I A E K Y F K L I A N A K T V E G V W T L K D E T L K A T V E
BINARY_3146 T T Y K L I L N L K Q L K E E A I T E L V D A G I A E K Y F K L I A N A K T V E G V W T L K D E T L K F T V E
BINARY_3147 T T Y K L I L N L K Q L K E E A I T E L V D A G I A E K Y F K L I A N A K T V E G V W T L K D E T L T A T V E
BINARY_3148 T T Y K L I L N L K Q L K E E A I T E L V D A G I A E K Y F K L I A N A K T V E G V W T L K D E T L T F T V E
BINARY_3149 T T Y K L I L N L K Q L K E E A I T E L V D A G I A E K Y F K L I A N A K T V E G V W T L K D E T K K A T V E
BINARY_3150 T T Y K L I L N L K Q L K E E A I T E L V D A G I A E K Y F K L I A N A K T V E G V W T L K D E T K K F T V E
BINARY_3151 T T Y K L I L N L K Q L K E E A I T E L V D A G I A E K Y F K L I A N A K T V E G V W T L K D E T K T A T V E

BINARY_3152 T T Y K L I L N L K Q L K E E A I T E L V D A G I A E K Y F K L I A N A K T V E G V W T L K D E T K T F T V T E
BINARY_3153 T T Y K L I L N L K Q L K E E A I T E L V D A G I A E K Y F K L I A N A K T V E G V W T Y K D E I L K A T V T E
BINARY_3154 T T Y K L I L N L K Q L K E E A I T E L V D A G I A E K Y F K L I A N A K T V E G V W T Y K D E I L K F T V T E
BINARY_3155 T T Y K L I L N L K Q L K E E A I T E L V D A G I A E K Y F K L I A N A K T V E G V W T Y K D E I L T A T V T E
BINARY_3156 T T Y K L I L N L K Q L K E E A I T E L V D A G I A E K Y F K L I A N A K T V E G V W T Y K D E I L T F T V T E
BINARY_3157 T T Y K L I L N L K Q L K E E A I T E L V D A G I A E K Y F K L I A N A K T V E G V W T Y K D E I K K A T V T E
BINARY_3158 T T Y K L I L N L K Q L K E E A I T E L V D A G I A E K Y F K L I A N A K T V E G V W T Y K D E I K K F T V T E
BINARY_3159 T T Y K L I L N L K Q L K E E A I T E L V D A G I A E K Y F K L I A N A K T V E G V W T Y K D E I K T A T V T E
BINARY_3160 T T Y K L I L N L K Q L K E E A I T E L V D A G I A E K Y F K L I A N A K T V E G V W T Y K D E I K T F T V T E
BINARY_3161 T T Y K L I L N L K Q L K E E A I T E L V D A G I A E K Y F K L I A N A K T V E G V W T Y K D E T L K A T V T E
BINARY_3162 T T Y K L I L N L K Q L K E E A I T E L V D A G I A E K Y F K L I A N A K T V E G V W T Y K D E T L K F T V T E
BINARY_3163 T T Y K L I L N L K Q L K E E A I T E L V D A G I A E K Y F K L I A N A K T V E G V W T Y K D E T L T A T V T E
BINARY_3164 T T Y K L I L N L K Q L K E E A I T E L V D A G I A E K Y F K L I A N A K T V E G V W T Y K D E T L T F T V T E
BINARY_3165 T T Y K L I L N L K Q L K E E A I T E L V D A G I A E K Y F K L I A N A K T V E G V W T Y K D E T K K A T V T E
BINARY_3166 T T Y K L I L N L K Q L K E E A I T E L V D A G I A E K Y F K L I A N A K T V E G V W T Y K D E T K K F T V T E
BINARY_3167 T T Y K L I L N L K Q L K E E A I T E L V D A G I A E K Y F K L I A N A K T V E G V W T Y K D E T K T A T V T E
BINARY_3168 T T Y K L I L N L K Q L K E E A I T E L V D A G I A E K Y F K L I A N A K T V E G V W T Y K D E T K T F T V T E
BINARY_3169 T T Y K L I L N L K Q L K E E A I T E L V D A G I A E K Y F K L Y A N A K T V E G V W T L K D E I L K A T V T E
BINARY_3170 T T Y K L I L N L K Q L K E E A I T E L V D A G I A E K Y F K L Y A N A K T V E G V W T L K D E I L K F T V T E
BINARY_3171 T T Y K L I L N L K Q L K E E A I T E L V D A G I A E K Y F K L Y A N A K T V E G V W T L K D E I L T A T V T E
BINARY_3172 T T Y K L I L N L K Q L K E E A I T E L V D A G I A E K Y F K L Y A N A K T V E G V W T L K D E I L T F T V T E
BINARY_3173 T T Y K L I L N L K Q L K E E A I T E L V D A G I A E K Y F K L Y A N A K T V E G V W T L K D E I K K A T V T E
BINARY_3174 T T Y K L I L N L K Q L K E E A I T E L V D A G I A E K Y F K L Y A N A K T V E G V W T L K D E I K K F T V T E
BINARY_3175 T T Y K L I L N L K Q L K E E A I T E L V D A G I A E K Y F K L Y A N A K T V E G V W T L K D E I K T A T V T E
BINARY_3176 T T Y K L I L N L K Q L K E E A I T E L V D A G I A E K Y F K L Y A N A K T V E G V W T L K D E I K T F T V T E
BINARY_3177 T T Y K L I L N L K Q L K E E A I T E L V D A G I A E K Y F K L Y A N A K T V E G V W T L K D E T L K A T V T E
BINARY_3178 T T Y K L I L N L K Q L K E E A I T E L V D A G I A E K Y F K L Y A N A K T V E G V W T L K D E T L K F T V T E
BINARY_3179 T T Y K L I L N L K Q L K E E A I T E L V D A G I A E K Y F K L Y A N A K T V E G V W T L K D E T L T A T V T E
BINARY_3180 T T Y K L I L N L K Q L K E E A I T E L V D A G I A E K Y F K L Y A N A K T V E G V W T L K D E T L T F T V T E
BINARY_3181 T T Y K L I L N L K Q L K E E A I T E L V D A G I A E K Y F K L Y A N A K T V E G V W T L K D E T K K A T V T E
BINARY_3182 T T Y K L I L N L K Q L K E E A I T E L V D A G I A E K Y F K L Y A N A K T V E G V W T L K D E T K K F T V T E
BINARY_3183 T T Y K L I L N L K Q L K E E A I T E L V D A G I A E K Y F K L Y A N A K T V E G V W T L K D E T K T A T V T E
BINARY_3184 T T Y K L I L N L K Q L K E E A I T E L V D A G I A E K Y F K L Y A N A K T V E G V W T L K D E T K T F T V T E
BINARY_3185 T T Y K L I L N L K Q L K E E A I T E L V D A G I A E K Y F K L Y A N A K T V E G V W T Y K D E I L K A T V T E
BINARY_3186 T T Y K L I L N L K Q L K E E A I T E L V D A G I A E K Y F K L Y A N A K T V E G V W T Y K D E I L K F T V T E
BINARY_3187 T T Y K L I L N L K Q L K E E A I T E L V D A G I A E K Y F K L Y A N A K T V E G V W T Y K D E I L T A T V T E
BINARY_3188 T T Y K L I L N L K Q L K E E A I T E L V D A G I A E K Y F K L Y A N A K T V E G V W T Y K D E I L T F T V T E
BINARY_3189 T T Y K L I L N L K Q L K E E A I T E L V D A G I A E K Y F K L Y A N A K T V E G V W T Y K D E I K K A T V T E
BINARY_3190 T T Y K L I L N L K Q L K E E A I T E L V D A G I A E K Y F K L Y A N A K T V E G V W T Y K D E I K K F T V T E
BINARY_3191 T T Y K L I L N L K Q L K E E A I T E L V D A G I A E K Y F K L Y A N A K T V E G V W T Y K D E I K T A T V T E
BINARY_3192 T T Y K L I L N L K Q L K E E A I T E L V D A G I A E K Y F K L Y A N A K T V E G V W T Y K D E I K T F T V T E
BINARY_3193 T T Y K L I L N L K Q L K E E A I T E L V D A G I A E K Y F K L Y A N A K T V E G V W T Y K D E T L K A T V T E
BINARY_3194 T T Y K L I L N L K Q L K E E A I T E L V D A G I A E K Y F K L Y A N A K T V E G V W T Y K D E T L K F T V T E
BINARY_3195 T T Y K L I L N L K Q L K E E A I T E L V D A G I A E K Y F K L Y A N A K T V E G V W T Y K D E T L T A T V T E
BINARY_3196 T T Y K L I L N L K Q L K E E A I T E L V D A G I A E K Y F K L Y A N A K T V E G V W T Y K D E T L T F T V T E
BINARY_3197 T T Y K L I L N L K Q L K E E A I T E L V D A G I A E K Y F K L Y A N A K T V E G V W T Y K D E T K K A T V T E
BINARY_3198 T T Y K L I L N L K Q L K E E A I T E L V D A G I A E K Y F K L Y A N A K T V E G V W T Y K D E T K K F T V T E
BINARY_3199 T T Y K L I L N L K Q L K E E A I T E L V D A G I A E K Y F K L Y A N A K T V E G V W T Y K D E T K T A T V T E
BINARY_3200 T T Y K L I L N L K Q L K E E A I T E L V D A G I A E K Y F K L Y A N A K T V E G V W T Y K D E T K T F T V T E
BINARY_3201 T T Y K L I L N L K Q L K E E A I T E L V D A G T A E K Y I K L I A N A K T V E G V W T L K D E I L K A T V T E
BINARY_3202 T T Y K L I L N L K Q L K E E A I T E L V D A G T A E K Y I K L I A N A K T V E G V W T L K D E I L K F T V T E
BINARY_3203 T T Y K L I L N L K Q L K E E A I T E L V D A G T A E K Y I K L I A N A K T V E G V W T L K D E I L T A T V T E
BINARY_3204 T T Y K L I L N L K Q L K E E A I T E L V D A G T A E K Y I K L I A N A K T V E G V W T L K D E I L T F T V T E
BINARY_3205 T T Y K L I L N L K Q L K E E A I T E L V D A G T A E K Y I K L I A N A K T V E G V W T L K D E I K K A T V T E
BINARY_3206 T T Y K L I L N L K Q L K E E A I T E L V D A G T A E K Y I K L I A N A K T V E G V W T L K D E I K K F T V T E
BINARY_3207 T T Y K L I L N L K Q L K E E A I T E L V D A G T A E K Y I K L I A N A K T V E G V W T L K D E I K T A T V T E
BINARY_3208 T T Y K L I L N L K Q L K E E A I T E L V D A G T A E K Y I K L I A N A K T V E G V W T L K D E I K T F T V T E
BINARY_3209 T T Y K L I L N L K Q L K E E A I T E L V D A G T A E K Y I K L I A N A K T V E G V W T L K D E T L K A T V T E
BINARY_3210 T T Y K L I L N L K Q L K E E A I T E L V D A G T A E K Y I K L I A N A K T V E G V W T L K D E T L K F T V T E
BINARY_3211 T T Y K L I L N L K Q L K E E A I T E L V D A G T A E K Y I K L I A N A K T V E G V W T L K D E T L T A T V T E
BINARY_3212 T T Y K L I L N L K Q L K E E A I T E L V D A G T A E K Y I K L I A N A K T V E G V W T L K D E T L T F T V T E
BINARY_3213 T T Y K L I L N L K Q L K E E A I T E L V D A G T A E K Y I K L I A N A K T V E G V W T L K D E T K K A T V T E
BINARY_3214 T T Y K L I L N L K Q L K E E A I T E L V D A G T A E K Y I K L I A N A K T V E G V W T L K D E T K K F T V T E
BINARY_3215 T T Y K L I L N L K Q L K E E A I T E L V D A G T A E K Y I K L I A N A K T V E G V W T L K D E T K T A T V T E
BINARY_3216 T T Y K L I L N L K Q L K E E A I T E L V D A G T A E K Y I K L I A N A K T V E G V W T L K D E T K T F T V T E
BINARY_3217 T T Y K L I L N L K Q L K E E A I T E L V D A G T A E K Y I K L I A N A K T V E G V W T Y K D E I L K A T V T E
BINARY_3218 T T Y K L I L N L K Q L K E E A I T E L V D A G T A E K Y I K L I A N A K T V E G V W T Y K D E I L K F T V T E
BINARY_3219 T T Y K L I L N L K Q L K E E A I T E L V D A G T A E K Y I K L I A N A K T V E G V W T Y K D E I L T A T V T E
BINARY_3220 T T Y K L I L N L K Q L K E E A I T E L V D A G T A E K Y I K L I A N A K T V E G V W T Y K D E I L T F T V T E
BINARY_3221 T T Y K L I L N L K Q L K E E A I T E L V D A G T A E K Y I K L I A N A K T V E G V W T Y K D E I K K A T V T E
BINARY_3222 T T Y K L I L N L K Q L K E E A I T E L V D A G T A E K Y I K L I A N A K T V E G V W T Y K D E I K K F T V T E
BINARY_3223 T T Y K L I L N L K Q L K E E A I T E L V D A G T A E K Y I K L I A N A K T V E G V W T Y K D E I K T A T V T E
BINARY_3224 T T Y K L I L N L K Q L K E E A I T E L V D A G T A E K Y I K L I A N A K T V E G V W T Y K D E I K T F T V T E
BINARY_3225 T T Y K L I L N L K Q L K E E A I T E L V D A G T A E K Y I K L I A N A K T V E G V W T Y K D E T L K A T V T E
BINARY_3226 T T Y K L I L N L K Q L K E E A I T E L V D A G T A E K Y I K L I A N A K T V E G V W T Y K D E T L K F T V T E
BINARY_3227 T T Y K L I L N L K Q L K E E A I T E L V D A G T A E K Y I K L I A N A K T V E G V W T Y K D E T L T A T V T E
BINARY_3228 T T Y K L I L N L K Q L K E E A I T E L V D A G T A E K Y I K L I A N A K T V E G V W T Y K D E T L T F T V T E
BINARY_3229 T T Y K L I L N L K Q L K E E A I T E L V D A G T A E K Y I K L I A N A K T V E G V W T Y K D E T K K A T V T E
BINARY_3230 T T Y K L I L N L K Q L K E E A I T E L V D A G T A E K Y I K L I A N A K T V E G V W T Y K D E T K K F T V T E
BINARY_3231 T T Y K L I L N L K Q L K E E A I T E L V D A G T A E K Y I K L I A N A K T V E G V W T Y K D E T K T A T V T E
BINARY_3232 T T Y K L I L N L K Q L K E E A I T E L V D A G T A E K Y I K L I A N A K T V E G V W T Y K D E T K T F T V T E
BINARY_3233 T T Y K L I L N L K Q L K E E A I T E L V D A G T A E K Y I K L Y A N A K T V E G V W T L K D E I L K A T V T E
BINARY_3234 T T Y K L I L N L K Q L K E E A I T E L V D A G T A E K Y I K L Y A N A K T V E G V W T L K D E I L K F T V T E
BINARY_3235 T T Y K L I L N L K Q L K E E A I T E L V D A G T A E K Y I K L Y A N A K T V E G V W T L K D E I L T A T V T E
BINARY_3236 T T Y K L I L N L K Q L K E E A I T E L V D A G T A E K Y I K L Y A N A K T V E G V W T L K D E I L T F T V T E
BINARY_3237 T T Y K L I L N L K Q L K E E A I T E L V D A G T A E K Y I K L Y A N A K T V E G V W T L K D E I K K A T V T E
BINARY_3238 T T Y K L I L N L K Q L K E E A I T E L V D A G T A E K Y I K L Y A N A K T V E G V W T L K D E I K K F T V T E
BINARY_3239 T T Y K L I L N L K Q L K E E A I T E L V D A G T A E K Y I K L Y A N A K T V E G V W T L K D E I K T A T V T E
BINARY_3240 T T Y K L I L N L K Q L K E E A I T E L V D A G T A E K Y I K L Y A N A K T V E G V W T L K D E I K T F T V T E
BINARY_3241 T T Y K L I L N L K Q L K E E A I T E L V D A G T A E K Y I K L Y A N A K T V E G V W T L K D E T L K A T V T E
BINARY_3242 T T Y K L I L N L K Q L K E E A I T E L V D A G T A E K Y I K L Y A N A K T V E G V W T L K D E T L K F T V T E
BINARY_3243 T T Y K L I L N L K Q L K E E A I T E L V D A G T A E K Y I K L Y A N A K T V E G V W T L K D E T L T A T V T E
BINARY_3244 T T Y K L I L N L K Q L K E E A I T E L V D A G T A E K Y I K L Y A N A K T V E G V W T L K D E T L T F T V T E
BINARY_3245 T T Y K L I L N L K Q L K E E A I T E L V D A G T A E K Y I K L Y A N A K T V E G V W T L K D E T K K A T V T E
BINARY_3246 T T Y K L I L N L K Q L K E E A I T E L V D A G T A E K Y I K L Y A N A K T V E G V W T L K D E T K K F T V T E

BINARY_3247 T T Y K L I L N L K Q L K E E A I T E L V D A G T A E K Y I K L Y A N A K T V E G V W T L K D E T K T A T V T E
BINARY_3248 T T Y K L I L N L K Q L K E E A I T E L V D A G T A E K Y I K L Y A N A K T V E G V W T L K D E T K T F T V T E
BINARY_3249 T T Y K L I L N L K Q L K E E A I T E L V D A G T A E K Y I K L Y A N A K T V E G V W T Y K D E I L K A T V T E
BINARY_3250 T T Y K L I L N L K Q L K E E A I T E L V D A G T A E K Y I K L Y A N A K T V E G V W T Y K D E I L K F T V T E
BINARY_3251 T T Y K L I L N L K Q L K E E A I T E L V D A G T A E K Y I K L Y A N A K T V E G V W T Y K D E I L T A T V T E
BINARY_3252 T T Y K L I L N L K Q L K E E A I T E L V D A G T A E K Y I K L Y A N A K T V E G V W T Y K D E I L T F T V T E
BINARY_3253 T T Y K L I L N L K Q L K E E A I T E L V D A G T A E K Y I K L Y A N A K T V E G V W T Y K D E I K K A T V T E
BINARY_3254 T T Y K L I L N L K Q L K E E A I T E L V D A G T A E K Y I K L Y A N A K T V E G V W T Y K D E I K K F T V T E
BINARY_3255 T T Y K L I L N L K Q L K E E A I T E L V D A G T A E K Y I K L Y A N A K T V E G V W T Y K D E I K T A T V T E
BINARY_3256 T T Y K L I L N L K Q L K E E A I T E L V D A G T A E K Y I K L Y A N A K T V E G V W T Y K D E I K T F T V T E
BINARY_3257 T T Y K L I L N L K Q L K E E A I T E L V D A G T A E K Y I K L Y A N A K T V E G V W T Y K D E T L K A T V T E
BINARY_3258 T T Y K L I L N L K Q L K E E A I T E L V D A G T A E K Y I K L Y A N A K T V E G V W T Y K D E T L K F T V T E
BINARY_3259 T T Y K L I L N L K Q L K E E A I T E L V D A G T A E K Y I K L Y A N A K T V E G V W T Y K D E T L T A T V T E
BINARY_3260 T T Y K L I L N L K Q L K E E A I T E L V D A G T A E K Y I K L Y A N A K T V E G V W T Y K D E T L T F T V T E
BINARY_3261 T T Y K L I L N L K Q L K E E A I T E L V D A G T A E K Y I K L Y A N A K T V E G V W T Y K D E T K K A T V T E
BINARY_3262 T T Y K L I L N L K Q L K E E A I T E L V D A G T A E K Y I K L Y A N A K T V E G V W T Y K D E T K K F T V T E
BINARY_3263 T T Y K L I L N L K Q L K E E A I T E L V D A G T A E K Y I K L Y A N A K T V E G V W T Y K D E T K T A T V T E
BINARY_3264 T T Y K L I L N L K Q L K E E A I T E L V D A G T A E K Y I K L Y A N A K T V E G V W T Y K D E T K T F T V T E
BINARY_3265 T T Y K L I L N L K Q L K E E A I T E L V D A G T A E K Y F K L I A N A K T V E G V W T L K D E I L K A T V T E
BINARY_3266 T T Y K L I L N L K Q L K E E A I T E L V D A G T A E K Y F K L I A N A K T V E G V W T L K D E I L K F T V T E
BINARY_3267 T T Y K L I L N L K Q L K E E A I T E L V D A G T A E K Y F K L I A N A K T V E G V W T L K D E I L T A T V T E
BINARY_3268 T T Y K L I L N L K Q L K E E A I T E L V D A G T A E K Y F K L I A N A K T V E G V W T L K D E I L T F T V T E
BINARY_3269 T T Y K L I L N L K Q L K E E A I T E L V D A G T A E K Y F K L I A N A K T V E G V W T L K D E I K K A T V T E
BINARY_3270 T T Y K L I L N L K Q L K E E A I T E L V D A G T A E K Y F K L I A N A K T V E G V W T L K D E I K K F T V T E
BINARY_3271 T T Y K L I L N L K Q L K E E A I T E L V D A G T A E K Y F K L I A N A K T V E G V W T L K D E I K T A T V T E
BINARY_3272 T T Y K L I L N L K Q L K E E A I T E L V D A G T A E K Y F K L I A N A K T V E G V W T L K D E I K T F T V T E
BINARY_3273 T T Y K L I L N L K Q L K E E A I T E L V D A G T A E K Y F K L I A N A K T V E G V W T L K D E T L K A T V T E
BINARY_3274 T T Y K L I L N L K Q L K E E A I T E L V D A G T A E K Y F K L I A N A K T V E G V W T L K D E T L K F T V T E
BINARY_3275 T T Y K L I L N L K Q L K E E A I T E L V D A G T A E K Y F K L I A N A K T V E G V W T L K D E T L T A T V T E
BINARY_3276 T T Y K L I L N L K Q L K E E A I T E L V D A G T A E K Y F K L I A N A K T V E G V W T L K D E T L T F T V T E
BINARY_3277 T T Y K L I L N L K Q L K E E A I T E L V D A G T A E K Y F K L I A N A K T V E G V W T L K D E T K K A T V T E
BINARY_3278 T T Y K L I L N L K Q L K E E A I T E L V D A G T A E K Y F K L I A N A K T V E G V W T L K D E T K K F T V T E
BINARY_3279 T T Y K L I L N L K Q L K E E A I T E L V D A G T A E K Y F K L I A N A K T V E G V W T L K D E T K T A T V T E
BINARY_3280 T T Y K L I L N L K Q L K E E A I T E L V D A G T A E K Y F K L I A N A K T V E G V W T L K D E T K T F T V T E
BINARY_3281 T T Y K L I L N L K Q L K E E A I T E L V D A G T A E K Y F K L I A N A K T V E G V W T Y K D E I L K A T V T E
BINARY_3282 T T Y K L I L N L K Q L K E E A I T E L V D A G T A E K Y F K L I A N A K T V E G V W T Y K D E I L K F T V T E
BINARY_3283 T T Y K L I L N L K Q L K E E A I T E L V D A G T A E K Y F K L I A N A K T V E G V W T Y K D E I L T A T V T E
BINARY_3284 T T Y K L I L N L K Q L K E E A I T E L V D A G T A E K Y F K L I A N A K T V E G V W T Y K D E I L T F T V T E
BINARY_3285 T T Y K L I L N L K Q L K E E A I T E L V D A G T A E K Y F K L I A N A K T V E G V W T Y K D E I K K A T V T E
BINARY_3286 T T Y K L I L N L K Q L K E E A I T E L V D A G T A E K Y F K L I A N A K T V E G V W T Y K D E I K K F T V T E
BINARY_3287 T T Y K L I L N L K Q L K E E A I T E L V D A G T A E K Y F K L I A N A K T V E G V W T Y K D E I K T A T V T E
BINARY_3288 T T Y K L I L N L K Q L K E E A I T E L V D A G T A E K Y F K L I A N A K T V E G V W T Y K D E I K T F T V T E
BINARY_3289 T T Y K L I L N L K Q L K E E A I T E L V D A G T A E K Y F K L I A N A K T V E G V W T Y K D E T L K A T V T E
BINARY_3290 T T Y K L I L N L K Q L K E E A I T E L V D A G T A E K Y F K L I A N A K T V E G V W T Y K D E T L K F T V T E
BINARY_3291 T T Y K L I L N L K Q L K E E A I T E L V D A G T A E K Y F K L I A N A K T V E G V W T Y K D E T L T A T V T E
BINARY_3292 T T Y K L I L N L K Q L K E E A I T E L V D A G T A E K Y F K L I A N A K T V E G V W T Y K D E T L T F T V T E
BINARY_3293 T T Y K L I L N L K Q L K E E A I T E L V D A G T A E K Y F K L I A N A K T V E G V W T Y K D E T K K A T V T E
BINARY_3294 T T Y K L I L N L K Q L K E E A I T E L V D A G T A E K Y F K L I A N A K T V E G V W T Y K D E T K K F T V T E
BINARY_3295 T T Y K L I L N L K Q L K E E A I T E L V D A G T A E K Y F K L I A N A K T V E G V W T Y K D E T K T A T V T E
BINARY_3296 T T Y K L I L N L K Q L K E E A I T E L V D A G T A E K Y F K L I A N A K T V E G V W T Y K D E T K T F T V T E
BINARY_3297 T T Y K L I L N L K Q L K E E A I T E L V D A G T A E K Y F K L Y A N A K T V E G V W T L K D E I L K A T V T E
BINARY_3298 T T Y K L I L N L K Q L K E E A I T E L V D A G T A E K Y F K L Y A N A K T V E G V W T L K D E I L K F T V T E
BINARY_3299 T T Y K L I L N L K Q L K E E A I T E L V D A G T A E K Y F K L Y A N A K T V E G V W T L K D E I L T A T V T E
BINARY_3300 T T Y K L I L N L K Q L K E E A I T E L V D A G T A E K Y F K L Y A N A K T V E G V W T L K D E I L T F T V T E
BINARY_3301 T T Y K L I L N L K Q L K E E A I T E L V D A G T A E K Y F K L Y A N A K T V E G V W T L K D E I K K A T V T E
BINARY_3302 T T Y K L I L N L K Q L K E E A I T E L V D A G T A E K Y F K L Y A N A K T V E G V W T L K D E I K K F T V T E
BINARY_3303 T T Y K L I L N L K Q L K E E A I T E L V D A G T A E K Y F K L Y A N A K T V E G V W T L K D E I K T A T V T E
BINARY_3304 T T Y K L I L N L K Q L K E E A I T E L V D A G T A E K Y F K L Y A N A K T V E G V W T L K D E I K T F T V T E
BINARY_3305 T T Y K L I L N L K Q L K E E A I T E L V D A G T A E K Y F K L Y A N A K T V E G V W T L K D E T L K A T V T E
BINARY_3306 T T Y K L I L N L K Q L K E E A I T E L V D A G T A E K Y F K L Y A N A K T V E G V W T L K D E T L K F T V T E
BINARY_3307 T T Y K L I L N L K Q L K E E A I T E L V D A G T A E K Y F K L Y A N A K T V E G V W T L K D E T L T A T V T E
BINARY_3308 T T Y K L I L N L K Q L K E E A I T E L V D A G T A E K Y F K L Y A N A K T V E G V W T L K D E T L T F T V T E
BINARY_3309 T T Y K L I L N L K Q L K E E A I T E L V D A G T A E K Y F K L Y A N A K T V E G V W T L K D E T K K A T V T E
BINARY_3310 T T Y K L I L N L K Q L K E E A I T E L V D A G T A E K Y F K L Y A N A K T V E G V W T L K D E T K K F T V T E
BINARY_3311 T T Y K L I L N L K Q L K E E A I T E L V D A G T A E K Y F K L Y A N A K T V E G V W T L K D E T K T A T V T E
BINARY_3312 T T Y K L I L N L K Q L K E E A I T E L V D A G T A E K Y F K L Y A N A K T V E G V W T L K D E T K T F T V T E
BINARY_3313 T T Y K L I L N L K Q L K E E A I T E L V D A G T A E K Y F K L Y A N A K T V E G V W T Y K D E I L K A T V T E
BINARY_3314 T T Y K L I L N L K Q L K E E A I T E L V D A G T A E K Y F K L Y A N A K T V E G V W T Y K D E I L K F T V T E
BINARY_3315 T T Y K L I L N L K Q L K E E A I T E L V D A G T A E K Y F K L Y A N A K T V E G V W T Y K D E I L T A T V T E
BINARY_3316 T T Y K L I L N L K Q L K E E A I T E L V D A G T A E K Y F K L Y A N A K T V E G V W T Y K D E I L T F T V T E
BINARY_3317 T T Y K L I L N L K Q L K E E A I T E L V D A G T A E K Y F K L Y A N A K T V E G V W T Y K D E I K K A T V T E
BINARY_3318 T T Y K L I L N L K Q L K E E A I T E L V D A G T A E K Y F K L Y A N A K T V E G V W T Y K D E I K K F T V T E
BINARY_3319 T T Y K L I L N L K Q L K E E A I T E L V D A G T A E K Y F K L Y A N A K T V E G V W T Y K D E I K T A T V T E
BINARY_3320 T T Y K L I L N L K Q L K E E A I T E L V D A G T A E K Y F K L Y A N A K T V E G V W T Y K D E I K T F T V T E
BINARY_3321 T T Y K L I L N L K Q L K E E A I T E L V D A G T A E K Y F K L Y A N A K T V E G V W T Y K D E T L K A T V T E
BINARY_3322 T T Y K L I L N L K Q L K E E A I T E L V D A G T A E K Y F K L Y A N A K T V E G V W T Y K D E T L K F T V T E
BINARY_3323 T T Y K L I L N L K Q L K E E A I T E L V D A G T A E K Y F K L Y A N A K T V E G V W T Y K D E T L T A T V T E
BINARY_3324 T T Y K L I L N L K Q L K E E A I T E L V D A G T A E K Y F K L Y A N A K T V E G V W T Y K D E T L T F T V T E
BINARY_3325 T T Y K L I L N L K Q L K E E A I T E L V D A G T A E K Y F K L Y A N A K T V E G V W T Y K D E T K K A T V T E
BINARY_3326 T T Y K L I L N L K Q L K E E A I T E L V D A G T A E K Y F K L Y A N A K T V E G V W T Y K D E T K K F T V T E
BINARY_3327 T T Y K L I L N L K Q L K E E A I T E L V D A G T A E K Y F K L Y A N A K T V E G V W T Y K D E T K T A T V T E
BINARY_3328 T T Y K L I L N L K Q L K E E A I T E L V D A G T A E K Y F K L Y A N A K T V E G V W T Y K D E T K T F T V T E
BINARY_3329 T T Y K L I L N L K Q L K E E A I T E L V D A A I A E K Y I K L I A N A K T V E G V W T L K D E I L K A T V T E
BINARY_3330 T T Y K L I L N L K Q L K E E A I T E L V D A A I A E K Y I K L I A N A K T V E G V W T L K D E I L K F T V T E
BINARY_3331 T T Y K L I L N L K Q L K E E A I T E L V D A A I A E K Y I K L I A N A K T V E G V W T L K D E I L T A T V T E
BINARY_3332 T T Y K L I L N L K Q L K E E A I T E L V D A A I A E K Y I K L I A N A K T V E G V W T L K D E I L T F T V T E
BINARY_3333 T T Y K L I L N L K Q L K E E A I T E L V D A A I A E K Y I K L I A N A K T V E G V W T L K D E I K K A T V T E
BINARY_3334 T T Y K L I L N L K Q L K E E A I T E L V D A A I A E K Y I K L I A N A K T V E G V W T L K D E I K K F T V T E
BINARY_3335 T T Y K L I L N L K Q L K E E A I T E L V D A A I A E K Y I K L I A N A K T V E G V W T L K D E I K T A T V T E
BINARY_3336 T T Y K L I L N L K Q L K E E A I T E L V D A A I A E K Y I K L I A N A K T V E G V W T L K D E I K T F T V T E
BINARY_3337 T T Y K L I L N L K Q L K E E A I T E L V D A A I A E K Y I K L I A N A K T V E G V W T L K D E T L K A T V T E
BINARY_3338 T T Y K L I L N L K Q L K E E A I T E L V D A A I A E K Y I K L I A N A K T V E G V W T L K D E T L K F T V T E
BINARY_3339 T T Y K L I L N L K Q L K E E A I T E L V D A A I A E K Y I K L I A N A K T V E G V W T L K D E T L T A T V T E
BINARY_3340 T T Y K L I L N L K Q L K E E A I T E L V D A A I A E K Y I K L I A N A K T V E G V W T L K D E T L T F T V T E
BINARY_3341 T T Y K L I L N L K Q L K E E A I T E L V D A A I A E K Y I K L I A N A K T V E G V W T L K D E T K K A T V T E

BINARY_3342 T T Y K L I L N L K Q L K E E A I T E L V D A A I A E K Y I K L I A N A K T V E G V W T L K D E T K K F T V T E
BINARY_3343 T T Y K L I L N L K Q L K E E A I T E L V D A A I A E K Y I K L I A N A K T V E G V W T L K D E T K T A T V T E
BINARY_3344 T T Y K L I L N L K Q L K E E A I T E L V D A A I A E K Y I K L I A N A K T V E G V W T L K D E T K T F T V T E
BINARY_3345 T T Y K L I L N L K Q L K E E A I T E L V D A A I A E K Y I K L I A N A K T V E G V W T Y K D E I L K A T V T E
BINARY_3346 T T Y K L I L N L K Q L K E E A I T E L V D A A I A E K Y I K L I A N A K T V E G V W T Y K D E I L K F T V T E
BINARY_3347 T T Y K L I L N L K Q L K E E A I T E L V D A A I A E K Y I K L I A N A K T V E G V W T Y K D E I L T A T V T E
BINARY_3348 T T Y K L I L N L K Q L K E E A I T E L V D A A I A E K Y I K L I A N A K T V E G V W T Y K D E I L T F T V T E
BINARY_3349 T T Y K L I L N L K Q L K E E A I T E L V D A A I A E K Y I K L I A N A K T V E G V W T Y K D E I K K A T V T E
BINARY_3350 T T Y K L I L N L K Q L K E E A I T E L V D A A I A E K Y I K L I A N A K T V E G V W T Y K D E I K K F T V T E
BINARY_3351 T T Y K L I L N L K Q L K E E A I T E L V D A A I A E K Y I K L I A N A K T V E G V W T Y K D E I K T A T V T E
BINARY_3352 T T Y K L I L N L K Q L K E E A I T E L V D A A I A E K Y I K L I A N A K T V E G V W T Y K D E I K T F T V T E
BINARY_3353 T T Y K L I L N L K Q L K E E A I T E L V D A A I A E K Y I K L I A N A K T V E G V W T Y K D E T L K A T V T E
BINARY_3354 T T Y K L I L N L K Q L K E E A I T E L V D A A I A E K Y I K L I A N A K T V E G V W T Y K D E T L K F T V T E
BINARY_3355 T T Y K L I L N L K Q L K E E A I T E L V D A A I A E K Y I K L I A N A K T V E G V W T Y K D E T L T A T V T E
BINARY_3356 T T Y K L I L N L K Q L K E E A I T E L V D A A I A E K Y I K L I A N A K T V E G V W T Y K D E T L T F T V T E
BINARY_3357 T T Y K L I L N L K Q L K E E A I T E L V D A A I A E K Y I K L I A N A K T V E G V W T Y K D E T K K A T V T E
BINARY_3358 T T Y K L I L N L K Q L K E E A I T E L V D A A I A E K Y I K L I A N A K T V E G V W T Y K D E T K K F T V T E
BINARY_3359 T T Y K L I L N L K Q L K E E A I T E L V D A A I A E K Y I K L I A N A K T V E G V W T Y K D E T K T A T V T E
BINARY_3360 T T Y K L I L N L K Q L K E E A I T E L V D A A I A E K Y I K L I A N A K T V E G V W T Y K D E T K T F T V T E
BINARY_3361 T T Y K L I L N L K Q L K E E A I T E L V D A A I A E K Y I K L Y A N A K T V E G V W T L K D E I L K A T V T E
BINARY_3362 T T Y K L I L N L K Q L K E E A I T E L V D A A I A E K Y I K L Y A N A K T V E G V W T L K D E I L K F T V T E
BINARY_3363 T T Y K L I L N L K Q L K E E A I T E L V D A A I A E K Y I K L Y A N A K T V E G V W T L K D E I L T A T V T E
BINARY_3364 T T Y K L I L N L K Q L K E E A I T E L V D A A I A E K Y I K L Y A N A K T V E G V W T L K D E I L T F T V T E
BINARY_3365 T T Y K L I L N L K Q L K E E A I T E L V D A A I A E K Y I K L Y A N A K T V E G V W T L K D E I K K A T V T E
BINARY_3366 T T Y K L I L N L K Q L K E E A I T E L V D A A I A E K Y I K L Y A N A K T V E G V W T L K D E I K K F T V T E
BINARY_3367 T T Y K L I L N L K Q L K E E A I T E L V D A A I A E K Y I K L Y A N A K T V E G V W T L K D E I K T A T V T E
BINARY_3368 T T Y K L I L N L K Q L K E E A I T E L V D A A I A E K Y I K L Y A N A K T V E G V W T L K D E I K T F T V T E
BINARY_3369 T T Y K L I L N L K Q L K E E A I T E L V D A A I A E K Y I K L Y A N A K T V E G V W T L K D E T L K A T V T E
BINARY_3370 T T Y K L I L N L K Q L K E E A I T E L V D A A I A E K Y I K L Y A N A K T V E G V W T L K D E T L K F T V T E
BINARY_3371 T T Y K L I L N L K Q L K E E A I T E L V D A A I A E K Y I K L Y A N A K T V E G V W T L K D E T L T A T V T E
BINARY_3372 T T Y K L I L N L K Q L K E E A I T E L V D A A I A E K Y I K L Y A N A K T V E G V W T L K D E T L T F T V T E
BINARY_3373 T T Y K L I L N L K Q L K E E A I T E L V D A A I A E K Y I K L Y A N A K T V E G V W T L K D E T K K A T V T E
BINARY_3374 T T Y K L I L N L K Q L K E E A I T E L V D A A I A E K Y I K L Y A N A K T V E G V W T L K D E T K K F T V T E
BINARY_3375 T T Y K L I L N L K Q L K E E A I T E L V D A A I A E K Y I K L Y A N A K T V E G V W T L K D E T K T A T V T E
BINARY_3376 T T Y K L I L N L K Q L K E E A I T E L V D A A I A E K Y I K L Y A N A K T V E G V W T L K D E T K T F T V T E
BINARY_3377 T T Y K L I L N L K Q L K E E A I T E L V D A A I A E K Y I K L Y A N A K T V E G V W T Y K D E I L K A T V T E
BINARY_3378 T T Y K L I L N L K Q L K E E A I T E L V D A A I A E K Y I K L Y A N A K T V E G V W T Y K D E I L K F T V T E
BINARY_3379 T T Y K L I L N L K Q L K E E A I T E L V D A A I A E K Y I K L Y A N A K T V E G V W T Y K D E I L T A T V T E
BINARY_3380 T T Y K L I L N L K Q L K E E A I T E L V D A A I A E K Y I K L Y A N A K T V E G V W T Y K D E I L T F T V T E
BINARY_3381 T T Y K L I L N L K Q L K E E A I T E L V D A A I A E K Y I K L Y A N A K T V E G V W T Y K D E I K K A T V T E
BINARY_3382 T T Y K L I L N L K Q L K E E A I T E L V D A A I A E K Y I K L Y A N A K T V E G V W T Y K D E I K K F T V T E
BINARY_3383 T T Y K L I L N L K Q L K E E A I T E L V D A A I A E K Y I K L Y A N A K T V E G V W T Y K D E I K T A T V T E
BINARY_3384 T T Y K L I L N L K Q L K E E A I T E L V D A A I A E K Y I K L Y A N A K T V E G V W T Y K D E I K T F T V T E
BINARY_3385 T T Y K L I L N L K Q L K E E A I T E L V D A A I A E K Y I K L Y A N A K T V E G V W T Y K D E T L K A T V T E
BINARY_3386 T T Y K L I L N L K Q L K E E A I T E L V D A A I A E K Y I K L Y A N A K T V E G V W T Y K D E T L K F T V T E
BINARY_3387 T T Y K L I L N L K Q L K E E A I T E L V D A A I A E K Y I K L Y A N A K T V E G V W T Y K D E T L T A T V T E
BINARY_3388 T T Y K L I L N L K Q L K E E A I T E L V D A A I A E K Y I K L Y A N A K T V E G V W T Y K D E T L T F T V T E
BINARY_3389 T T Y K L I L N L K Q L K E E A I T E L V D A A I A E K Y I K L Y A N A K T V E G V W T Y K D E T K K A T V T E
BINARY_3390 T T Y K L I L N L K Q L K E E A I T E L V D A A I A E K Y I K L Y A N A K T V E G V W T Y K D E T K K F T V T E
BINARY_3391 T T Y K L I L N L K Q L K E E A I T E L V D A A I A E K Y I K L Y A N A K T V E G V W T Y K D E T K T A T V T E
BINARY_3392 T T Y K L I L N L K Q L K E E A I T E L V D A A I A E K Y I K L Y A N A K T V E G V W T Y K D E T K T F T V T E
BINARY_3393 T T Y K L I L N L K Q L K E E A I T E L V D A A I A E K Y F K L I A N A K T V E G V W T L K D E I L K A T V T E
BINARY_3394 T T Y K L I L N L K Q L K E E A I T E L V D A A I A E K Y F K L I A N A K T V E G V W T L K D E I L K F T V T E
BINARY_3395 T T Y K L I L N L K Q L K E E A I T E L V D A A I A E K Y F K L I A N A K T V E G V W T L K D E I L T A T V T E
BINARY_3396 T T Y K L I L N L K Q L K E E A I T E L V D A A I A E K Y F K L I A N A K T V E G V W T L K D E I L T F T V T E
BINARY_3397 T T Y K L I L N L K Q L K E E A I T E L V D A A I A E K Y F K L I A N A K T V E G V W T L K D E I K K A T V T E
BINARY_3398 T T Y K L I L N L K Q L K E E A I T E L V D A A I A E K Y F K L I A N A K T V E G V W T L K D E I K K F T V T E
BINARY_3399 T T Y K L I L N L K Q L K E E A I T E L V D A A I A E K Y F K L I A N A K T V E G V W T L K D E I K T A T V T E
BINARY_3400 T T Y K L I L N L K Q L K E E A I T E L V D A A I A E K Y F K L I A N A K T V E G V W T L K D E I K T F T V T E
BINARY_3401 T T Y K L I L N L K Q L K E E A I T E L V D A A I A E K Y F K L I A N A K T V E G V W T L K D E T L K A T V T E
BINARY_3402 T T Y K L I L N L K Q L K E E A I T E L V D A A I A E K Y F K L I A N A K T V E G V W T L K D E T L K F T V T E
BINARY_3403 T T Y K L I L N L K Q L K E E A I T E L V D A A I A E K Y F K L I A N A K T V E G V W T L K D E T L T A T V T E
BINARY_3404 T T Y K L I L N L K Q L K E E A I T E L V D A A I A E K Y F K L I A N A K T V E G V W T L K D E T L T F T V T E
BINARY_3405 T T Y K L I L N L K Q L K E E A I T E L V D A A I A E K Y F K L I A N A K T V E G V W T L K D E T K K A T V T E
BINARY_3406 T T Y K L I L N L K Q L K E E A I T E L V D A A I A E K Y F K L I A N A K T V E G V W T L K D E T K K F T V T E
BINARY_3407 T T Y K L I L N L K Q L K E E A I T E L V D A A I A E K Y F K L I A N A K T V E G V W T L K D E T K T A T V T E
BINARY_3408 T T Y K L I L N L K Q L K E E A I T E L V D A A I A E K Y F K L I A N A K T V E G V W T L K D E T K T F T V T E
BINARY_3409 T T Y K L I L N L K Q L K E E A I T E L V D A A I A E K Y F K L I A N A K T V E G V W T Y K D E I L K A T V T E
BINARY_3410 T T Y K L I L N L K Q L K E E A I T E L V D A A I A E K Y F K L I A N A K T V E G V W T Y K D E I L K F T V T E
BINARY_3411 T T Y K L I L N L K Q L K E E A I T E L V D A A I A E K Y F K L I A N A K T V E G V W T Y K D E I L T A T V T E
BINARY_3412 T T Y K L I L N L K Q L K E E A I T E L V D A A I A E K Y F K L I A N A K T V E G V W T Y K D E I L T F T V T E
BINARY_3413 T T Y K L I L N L K Q L K E E A I T E L V D A A I A E K Y F K L I A N A K T V E G V W T Y K D E I K K A T V T E
BINARY_3414 T T Y K L I L N L K Q L K E E A I T E L V D A A I A E K Y F K L I A N A K T V E G V W T Y K D E I K K F T V T E
BINARY_3415 T T Y K L I L N L K Q L K E E A I T E L V D A A I A E K Y F K L I A N A K T V E G V W T Y K D E I K T A T V T E
BINARY_3416 T T Y K L I L N L K Q L K E E A I T E L V D A A I A E K Y F K L I A N A K T V E G V W T Y K D E I K T F T V T E
BINARY_3417 T T Y K L I L N L K Q L K E E A I T E L V D A A I A E K Y F K L I A N A K T V E G V W T Y K D E T L K A T V T E
BINARY_3418 T T Y K L I L N L K Q L K E E A I T E L V D A A I A E K Y F K L I A N A K T V E G V W T Y K D E T L K F T V T E
BINARY_3419 T T Y K L I L N L K Q L K E E A I T E L V D A A I A E K Y F K L I A N A K T V E G V W T Y K D E T L T A T V T E
BINARY_3420 T T Y K L I L N L K Q L K E E A I T E L V D A A I A E K Y F K L I A N A K T V E G V W T Y K D E T L T F T V T E
BINARY_3421 T T Y K L I L N L K Q L K E E A I T E L V D A A I A E K Y F K L I A N A K T V E G V W T Y K D E T K K A T V T E
BINARY_3422 T T Y K L I L N L K Q L K E E A I T E L V D A A I A E K Y F K L I A N A K T V E G V W T Y K D E T K K F T V T E
BINARY_3423 T T Y K L I L N L K Q L K E E A I T E L V D A A I A E K Y F K L I A N A K T V E G V W T Y K D E T K T A T V T E
BINARY_3424 T T Y K L I L N L K Q L K E E A I T E L V D A A I A E K Y F K L I A N A K T V E G V W T Y K D E T K T F T V T E
BINARY_3425 T T Y K L I L N L K Q L K E E A I T E L V D A A I A E K Y F K L Y A N A K T V E G V W T L K D E I L K A T V T E
BINARY_3426 T T Y K L I L N L K Q L K E E A I T E L V D A A I A E K Y F K L Y A N A K T V E G V W T L K D E I L K F T V T E
BINARY_3427 T T Y K L I L N L K Q L K E E A I T E L V D A A I A E K Y F K L Y A N A K T V E G V W T L K D E I L T A T V T E
BINARY_3428 T T Y K L I L N L K Q L K E E A I T E L V D A A I A E K Y F K L Y A N A K T V E G V W T L K D E I L T F T V T E
BINARY_3429 T T Y K L I L N L K Q L K E E A I T E L V D A A I A E K Y F K L Y A N A K T V E G V W T L K D E I K K A T V T E
BINARY_3430 T T Y K L I L N L K Q L K E E A I T E L V D A A I A E K Y F K L Y A N A K T V E G V W T L K D E I K K F T V T E
BINARY_3431 T T Y K L I L N L K Q L K E E A I T E L V D A A I A E K Y F K L Y A N A K T V E G V W T L K D E I K T A T V T E
BINARY_3432 T T Y K L I L N L K Q L K E E A I T E L V D A A I A E K Y F K L Y A N A K T V E G V W T L K D E I K T F T V T E
BINARY_3433 T T Y K L I L N L K Q L K E E A I T E L V D A A I A E K Y F K L Y A N A K T V E G V W T L K D E T L K A T V T E
BINARY_3434 T T Y K L I L N L K Q L K E E A I T E L V D A A I A E K Y F K L Y A N A K T V E G V W T L K D E T L K F T V T E
BINARY_3435 T T Y K L I L N L K Q L K E E A I T E L V D A A I A E K Y F K L Y A N A K T V E G V W T L K D E T L T A T V T E
BINARY_3436 T T Y K L I L N L K Q L K E E A I T E L V D A A I A E K Y F K L Y A N A K T V E G V W T L K D E T L T F T V T E

BINARY_3437 T T Y K L I L N L K Q L K E E A I T E L V D A A I A E K Y F K L Y A N A K T V E G V W T L K D E T K K A T V T E
BINARY_3438 T T Y K L I L N L K Q L K E E A I T E L V D A A I A E K Y F K L Y A N A K T V E G V W T L K D E T K K F T V T E
BINARY_3439 T T Y K L I L N L K Q L K E E A I T E L V D A A I A E K Y F K L Y A N A K T V E G V W T L K D E T K T A T V T E
BINARY_3440 T T Y K L I L N L K Q L K E E A I T E L V D A A I A E K Y F K L Y A N A K T V E G V W T L K D E T K T F T V T E
BINARY_3441 T T Y K L I L N L K Q L K E E A I T E L V D A A I A E K Y F K L Y A N A K T V E G V W T Y K D E I L K A T V T E
BINARY_3442 T T Y K L I L N L K Q L K E E A I T E L V D A A I A E K Y F K L Y A N A K T V E G V W T Y K D E I L K F T V T E
BINARY_3443 T T Y K L I L N L K Q L K E E A I T E L V D A A I A E K Y F K L Y A N A K T V E G V W T Y K D E I L T A T V T E
BINARY_3444 T T Y K L I L N L K Q L K E E A I T E L V D A A I A E K Y F K L Y A N A K T V E G V W T Y K D E I L T F T V T E
BINARY_3445 T T Y K L I L N L K Q L K E E A I T E L V D A A I A E K Y F K L Y A N A K T V E G V W T Y K D E I K K A T V T E
BINARY_3446 T T Y K L I L N L K Q L K E E A I T E L V D A A I A E K Y F K L Y A N A K T V E G V W T Y K D E I K K F T V T E
BINARY_3447 T T Y K L I L N L K Q L K E E A I T E L V D A A I A E K Y F K L Y A N A K T V E G V W T Y K D E I K T A T V T E
BINARY_3448 T T Y K L I L N L K Q L K E E A I T E L V D A A I A E K Y F K L Y A N A K T V E G V W T Y K D E I K T F T V T E
BINARY_3449 T T Y K L I L N L K Q L K E E A I T E L V D A A I A E K Y F K L Y A N A K T V E G V W T Y K D E T L K A T V T E
BINARY_3450 T T Y K L I L N L K Q L K E E A I T E L V D A A I A E K Y F K L Y A N A K T V E G V W T Y K D E T L K F T V T E
BINARY_3451 T T Y K L I L N L K Q L K E E A I T E L V D A A I A E K Y F K L Y A N A K T V E G V W T Y K D E T L T A T V T E
BINARY_3452 T T Y K L I L N L K Q L K E E A I T E L V D A A I A E K Y F K L Y A N A K T V E G V W T Y K D E T L T F T V T E
BINARY_3453 T T Y K L I L N L K Q L K E E A I T E L V D A A I A E K Y F K L Y A N A K T V E G V W T Y K D E T K K A T V T E
BINARY_3454 T T Y K L I L N L K Q L K E E A I T E L V D A A I A E K Y F K L Y A N A K T V E G V W T Y K D E T K K F T V T E
BINARY_3455 T T Y K L I L N L K Q L K E E A I T E L V D A A I A E K Y F K L Y A N A K T V E G V W T Y K D E T K T A T V T E
BINARY_3456 T T Y K L I L N L K Q L K E E A I T E L V D A A I A E K Y F K L Y A N A K T V E G V W T Y K D E T K T F T V T E
BINARY_3457 T T Y K L I L N L K Q L K E E A I T E L V D A A T A E K Y I K L I A N A K T V E G V W T L K D E I L K A T V T E
BINARY_3458 T T Y K L I L N L K Q L K E E A I T E L V D A A T A E K Y I K L I A N A K T V E G V W T L K D E I L K F T V T E
BINARY_3459 T T Y K L I L N L K Q L K E E A I T E L V D A A T A E K Y I K L I A N A K T V E G V W T L K D E I L T A T V T E
BINARY_3460 T T Y K L I L N L K Q L K E E A I T E L V D A A T A E K Y I K L I A N A K T V E G V W T L K D E I L T F T V T E
BINARY_3461 T T Y K L I L N L K Q L K E E A I T E L V D A A T A E K Y I K L I A N A K T V E G V W T L K D E I K K A T V T E
BINARY_3462 T T Y K L I L N L K Q L K E E A I T E L V D A A T A E K Y I K L I A N A K T V E G V W T L K D E I K K F T V T E
BINARY_3463 T T Y K L I L N L K Q L K E E A I T E L V D A A T A E K Y I K L I A N A K T V E G V W T L K D E I K T A T V T E
BINARY_3464 T T Y K L I L N L K Q L K E E A I T E L V D A A T A E K Y I K L I A N A K T V E G V W T L K D E I K T F T V T E
BINARY_3465 T T Y K L I L N L K Q L K E E A I T E L V D A A T A E K Y I K L I A N A K T V E G V W T L K D E T L K A T V T E
BINARY_3466 T T Y K L I L N L K Q L K E E A I T E L V D A A T A E K Y I K L I A N A K T V E G V W T L K D E T L K F T V T E
BINARY_3467 T T Y K L I L N L K Q L K E E A I T E L V D A A T A E K Y I K L I A N A K T V E G V W T L K D E T L T A T V T E
BINARY_3468 T T Y K L I L N L K Q L K E E A I T E L V D A A T A E K Y I K L I A N A K T V E G V W T L K D E T L T F T V T E
BINARY_3469 T T Y K L I L N L K Q L K E E A I T E L V D A A T A E K Y I K L I A N A K T V E G V W T L K D E T K K A T V T E
BINARY_3470 T T Y K L I L N L K Q L K E E A I T E L V D A A T A E K Y I K L I A N A K T V E G V W T L K D E T K K F T V T E
BINARY_3471 T T Y K L I L N L K Q L K E E A I T E L V D A A T A E K Y I K L I A N A K T V E G V W T L K D E T K T A T V T E
BINARY_3472 T T Y K L I L N L K Q L K E E A I T E L V D A A T A E K Y I K L I A N A K T V E G V W T L K D E T K T F T V T E
BINARY_3473 T T Y K L I L N L K Q L K E E A I T E L V D A A T A E K Y I K L I A N A K T V E G V W T Y K D E I L K A T V T E
BINARY_3474 T T Y K L I L N L K Q L K E E A I T E L V D A A T A E K Y I K L I A N A K T V E G V W T Y K D E I L K F T V T E
BINARY_3475 T T Y K L I L N L K Q L K E E A I T E L V D A A T A E K Y I K L I A N A K T V E G V W T Y K D E I L T A T V T E
BINARY_3476 T T Y K L I L N L K Q L K E E A I T E L V D A A T A E K Y I K L I A N A K T V E G V W T Y K D E I L T F T V T E
BINARY_3477 T T Y K L I L N L K Q L K E E A I T E L V D A A T A E K Y I K L I A N A K T V E G V W T Y K D E I K K A T V T E
BINARY_3478 T T Y K L I L N L K Q L K E E A I T E L V D A A T A E K Y I K L I A N A K T V E G V W T Y K D E I K K F T V T E
BINARY_3479 T T Y K L I L N L K Q L K E E A I T E L V D A A T A E K Y I K L I A N A K T V E G V W T Y K D E I K T A T V T E
BINARY_3480 T T Y K L I L N L K Q L K E E A I T E L V D A A T A E K Y I K L I A N A K T V E G V W T Y K D E I K T F T V T E
BINARY_3481 T T Y K L I L N L K Q L K E E A I T E L V D A A T A E K Y I K L I A N A K T V E G V W T Y K D E T L K A T V T E
BINARY_3482 T T Y K L I L N L K Q L K E E A I T E L V D A A T A E K Y I K L I A N A K T V E G V W T Y K D E T L K F T V T E
BINARY_3483 T T Y K L I L N L K Q L K E E A I T E L V D A A T A E K Y I K L I A N A K T V E G V W T Y K D E T L T A T V T E
BINARY_3484 T T Y K L I L N L K Q L K E E A I T E L V D A A T A E K Y I K L I A N A K T V E G V W T Y K D E T L T F T V T E
BINARY_3485 T T Y K L I L N L K Q L K E E A I T E L V D A A T A E K Y I K L I A N A K T V E G V W T Y K D E T K K A T V T E
BINARY_3486 T T Y K L I L N L K Q L K E E A I T E L V D A A T A E K Y I K L I A N A K T V E G V W T Y K D E T K K F T V T E
BINARY_3487 T T Y K L I L N L K Q L K E E A I T E L V D A A T A E K Y I K L I A N A K T V E G V W T Y K D E T K T A T V T E
BINARY_3488 T T Y K L I L N L K Q L K E E A I T E L V D A A T A E K Y I K L I A N A K T V E G V W T Y K D E T K T F T V T E
BINARY_3489 T T Y K L I L N L K Q L K E E A I T E L V D A A T A E K Y I K L Y A N A K T V E G V W T L K D E I L K A T V T E
BINARY_3490 T T Y K L I L N L K Q L K E E A I T E L V D A A T A E K Y I K L Y A N A K T V E G V W T L K D E I L K F T V T E
BINARY_3491 T T Y K L I L N L K Q L K E E A I T E L V D A A T A E K Y I K L Y A N A K T V E G V W T L K D E I L T A T V T E
BINARY_3492 T T Y K L I L N L K Q L K E E A I T E L V D A A T A E K Y I K L Y A N A K T V E G V W T L K D E I L T F T V T E
BINARY_3493 T T Y K L I L N L K Q L K E E A I T E L V D A A T A E K Y I K L Y A N A K T V E G V W T L K D E I K K A T V T E
BINARY_3494 T T Y K L I L N L K Q L K E E A I T E L V D A A T A E K Y I K L Y A N A K T V E G V W T L K D E I K K F T V T E
BINARY_3495 T T Y K L I L N L K Q L K E E A I T E L V D A A T A E K Y I K L Y A N A K T V E G V W T L K D E I K T A T V T E
BINARY_3496 T T Y K L I L N L K Q L K E E A I T E L V D A A T A E K Y I K L Y A N A K T V E G V W T L K D E I K T F T V T E
BINARY_3497 T T Y K L I L N L K Q L K E E A I T E L V D A A T A E K Y I K L Y A N A K T V E G V W T L K D E T L K A T V T E
BINARY_3498 T T Y K L I L N L K Q L K E E A I T E L V D A A T A E K Y I K L Y A N A K T V E G V W T L K D E T L K F T V T E
BINARY_3499 T T Y K L I L N L K Q L K E E A I T E L V D A A T A E K Y I K L Y A N A K T V E G V W T L K D E T L T A T V T E
BINARY_3500 T T Y K L I L N L K Q L K E E A I T E L V D A A T A E K Y I K L Y A N A K T V E G V W T L K D E T L T F T V T E
BINARY_3501 T T Y K L I L N L K Q L K E E A I T E L V D A A T A E K Y I K L Y A N A K T V E G V W T L K D E T K K A T V T E
BINARY_3502 T T Y K L I L N L K Q L K E E A I T E L V D A A T A E K Y I K L Y A N A K T V E G V W T L K D E T K K F T V T E
BINARY_3503 T T Y K L I L N L K Q L K E E A I T E L V D A A T A E K Y I K L Y A N A K T V E G V W T L K D E T K T A T V T E
BINARY_3504 T T Y K L I L N L K Q L K E E A I T E L V D A A T A E K Y I K L Y A N A K T V E G V W T L K D E T K T F T V T E
BINARY_3505 T T Y K L I L N L K Q L K E E A I T E L V D A A T A E K Y I K L Y A N A K T V E G V W T Y K D E I L K A T V T E
BINARY_3506 T T Y K L I L N L K Q L K E E A I T E L V D A A T A E K Y I K L Y A N A K T V E G V W T Y K D E I L K F T V T E
BINARY_3507 T T Y K L I L N L K Q L K E E A I T E L V D A A T A E K Y I K L Y A N A K T V E G V W T Y K D E I L T A T V T E
BINARY_3508 T T Y K L I L N L K Q L K E E A I T E L V D A A T A E K Y I K L Y A N A K T V E G V W T Y K D E I L T F T V T E
BINARY_3509 T T Y K L I L N L K Q L K E E A I T E L V D A A T A E K Y I K L Y A N A K T V E G V W T Y K D E I K K A T V T E
BINARY_3510 T T Y K L I L N L K Q L K E E A I T E L V D A A T A E K Y I K L Y A N A K T V E G V W T Y K D E I K K F T V T E
BINARY_3511 T T Y K L I L N L K Q L K E E A I T E L V D A A T A E K Y I K L Y A N A K T V E G V W T Y K D E I K T A T V T E
BINARY_3512 T T Y K L I L N L K Q L K E E A I T E L V D A A T A E K Y I K L Y A N A K T V E G V W T Y K D E I K T F T V T E
BINARY_3513 T T Y K L I L N L K Q L K E E A I T E L V D A A T A E K Y I K L Y A N A K T V E G V W T Y K D E T L K A T V T E
BINARY_3514 T T Y K L I L N L K Q L K E E A I T E L V D A A T A E K Y I K L Y A N A K T V E G V W T Y K D E T L K F T V T E
BINARY_3515 T T Y K L I L N L K Q L K E E A I T E L V D A A T A E K Y I K L Y A N A K T V E G V W T Y K D E T L T A T V T E
BINARY_3516 T T Y K L I L N L K Q L K E E A I T E L V D A A T A E K Y I K L Y A N A K T V E G V W T Y K D E T L T F T V T E
BINARY_3517 T T Y K L I L N L K Q L K E E A I T E L V D A A T A E K Y I K L Y A N A K T V E G V W T Y K D E T K K A T V T E
BINARY_3518 T T Y K L I L N L K Q L K E E A I T E L V D A A T A E K Y I K L Y A N A K T V E G V W T Y K D E T K K F T V T E
BINARY_3519 T T Y K L I L N L K Q L K E E A I T E L V D A A T A E K Y I K L Y A N A K T V E G V W T Y K D E T K T A T V T E
BINARY_3520 T T Y K L I L N L K Q L K E E A I T E L V D A A T A E K Y I K L Y A N A K T V E G V W T Y K D E T K T F T V T E
BINARY_3521 T T Y K L I L N L K Q L K E E A I T E L V D A A T A E K Y F K L I A N A K T V E G V W T L K D E I L K A T V T E
BINARY_3522 T T Y K L I L N L K Q L K E E A I T E L V D A A T A E K Y F K L I A N A K T V E G V W T L K D E I L K F T V T E
BINARY_3523 T T Y K L I L N L K Q L K E E A I T E L V D A A T A E K Y F K L I A N A K T V E G V W T L K D E I L T A T V T E
BINARY_3524 T T Y K L I L N L K Q L K E E A I T E L V D A A T A E K Y F K L I A N A K T V E G V W T L K D E I L T F T V T E
BINARY_3525 T T Y K L I L N L K Q L K E E A I T E L V D A A T A E K Y F K L I A N A K T V E G V W T L K D E I K K A T V T E
BINARY_3526 T T Y K L I L N L K Q L K E E A I T E L V D A A T A E K Y F K L I A N A K T V E G V W T L K D E I K K F T V T E
BINARY_3527 T T Y K L I L N L K Q L K E E A I T E L V D A A T A E K Y F K L I A N A K T V E G V W T L K D E I K T A T V T E
BINARY_3528 T T Y K L I L N L K Q L K E E A I T E L V D A A T A E K Y F K L I A N A K T V E G V W T L K D E I K T F T V T E
BINARY_3529 T T Y K L I L N L K Q L K E E A I T E L V D A A T A E K Y F K L I A N A K T V E G V W T L K D E T L K A T V T E
BINARY_3530 T T Y K L I L N L K Q L K E E A I T E L V D A A T A E K Y F K L I A N A K T V E G V W T L K D E T L K F T V T E
BINARY_3531 T T Y K L I L N L K Q L K E E A I T E L V D A A T A E K Y F K L I A N A K T V E G V W T L K D E T L T A T V T E

BINARY_3532 T T Y K L I L N L K Q L K E E A I T E L V D A A T A E K Y F K L I A N A K T V E G V W T L K D E T L T F T V T E
BINARY_3533 T T Y K L I L N L K Q L K E E A I T E L V D A A T A E K Y F K L I A N A K T V E G V W T L K D E T K K A T V T E
BINARY_3534 T T Y K L I L N L K Q L K E E A I T E L V D A A T A E K Y F K L I A N A K T V E G V W T L K D E T K K F T V T E
BINARY_3535 T T Y K L I L N L K Q L K E E A I T E L V D A A T A E K Y F K L I A N A K T V E G V W T L K D E T K T A T V T E
BINARY_3536 T T Y K L I L N L K Q L K E E A I T E L V D A A T A E K Y F K L I A N A K T V E G V W T L K D E T K T F T V T E
BINARY_3537 T T Y K L I L N L K Q L K E E A I T E L V D A A T A E K Y F K L I A N A K T V E G V W T Y K D E I L K A T V T E
BINARY_3538 T T Y K L I L N L K Q L K E E A I T E L V D A A T A E K Y F K L I A N A K T V E G V W T Y K D E I L K F T V T E
BINARY_3539 T T Y K L I L N L K Q L K E E A I T E L V D A A T A E K Y F K L I A N A K T V E G V W T Y K D E I L T A T V T E
BINARY_3540 T T Y K L I L N L K Q L K E E A I T E L V D A A T A E K Y F K L I A N A K T V E G V W T Y K D E I L T F T V T E
BINARY_3541 T T Y K L I L N L K Q L K E E A I T E L V D A A T A E K Y F K L I A N A K T V E G V W T Y K D E I K K A T V T E
BINARY_3542 T T Y K L I L N L K Q L K E E A I T E L V D A A T A E K Y F K L I A N A K T V E G V W T Y K D E I K K F T V T E
BINARY_3543 T T Y K L I L N L K Q L K E E A I T E L V D A A T A E K Y F K L I A N A K T V E G V W T Y K D E I K T A T V T E
BINARY_3544 T T Y K L I L N L K Q L K E E A I T E L V D A A T A E K Y F K L I A N A K T V E G V W T Y K D E I K T F T V T E
BINARY_3545 T T Y K L I L N L K Q L K E E A I T E L V D A A T A E K Y F K L I A N A K T V E G V W T Y K D E T L K A T V T E
BINARY_3546 T T Y K L I L N L K Q L K E E A I T E L V D A A T A E K Y F K L I A N A K T V E G V W T Y K D E T L K F T V T E
BINARY_3547 T T Y K L I L N L K Q L K E E A I T E L V D A A T A E K Y F K L I A N A K T V E G V W T Y K D E T L T A T V T E
BINARY_3548 T T Y K L I L N L K Q L K E E A I T E L V D A A T A E K Y F K L I A N A K T V E G V W T Y K D E T L T F T V T E
BINARY_3549 T T Y K L I L N L K Q L K E E A I T E L V D A A T A E K Y F K L I A N A K T V E G V W T Y K D E T K K A T V T E
BINARY_3550 T T Y K L I L N L K Q L K E E A I T E L V D A A T A E K Y F K L I A N A K T V E G V W T Y K D E T K K F T V T E
BINARY_3551 T T Y K L I L N L K Q L K E E A I T E L V D A A T A E K Y F K L I A N A K T V E G V W T Y K D E T K T A T V T E
BINARY_3552 T T Y K L I L N L K Q L K E E A I T E L V D A A T A E K Y F K L I A N A K T V E G V W T Y K D E T K T F T V T E
BINARY_3553 T T Y K L I L N L K Q L K E E A I T E L V D A A T A E K Y F K L Y A N A K T V E G V W T L K D E I L K A T V T E
BINARY_3554 T T Y K L I L N L K Q L K E E A I T E L V D A A T A E K Y F K L Y A N A K T V E G V W T L K D E I L K F T V T E
BINARY_3555 T T Y K L I L N L K Q L K E E A I T E L V D A A T A E K Y F K L Y A N A K T V E G V W T L K D E I L T A T V T E
BINARY_3556 T T Y K L I L N L K Q L K E E A I T E L V D A A T A E K Y F K L Y A N A K T V E G V W T L K D E I L T F T V T E
BINARY_3557 T T Y K L I L N L K Q L K E E A I T E L V D A A T A E K Y F K L Y A N A K T V E G V W T L K D E I K K A T V T E
BINARY_3558 T T Y K L I L N L K Q L K E E A I T E L V D A A T A E K Y F K L Y A N A K T V E G V W T L K D E I K K F T V T E
BINARY_3559 T T Y K L I L N L K Q L K E E A I T E L V D A A T A E K Y F K L Y A N A K T V E G V W T L K D E I K T A T V T E
BINARY_3560 T T Y K L I L N L K Q L K E E A I T E L V D A A T A E K Y F K L Y A N A K T V E G V W T L K D E I K T F T V T E
BINARY_3561 T T Y K L I L N L K Q L K E E A I T E L V D A A T A E K Y F K L Y A N A K T V E G V W T L K D E T L K A T V T E
BINARY_3562 T T Y K L I L N L K Q L K E E A I T E L V D A A T A E K Y F K L Y A N A K T V E G V W T L K D E T L K F T V T E
BINARY_3563 T T Y K L I L N L K Q L K E E A I T E L V D A A T A E K Y F K L Y A N A K T V E G V W T L K D E T L T A T V T E
BINARY_3564 T T Y K L I L N L K Q L K E E A I T E L V D A A T A E K Y F K L Y A N A K T V E G V W T L K D E T L T F T V T E
BINARY_3565 T T Y K L I L N L K Q L K E E A I T E L V D A A T A E K Y F K L Y A N A K T V E G V W T L K D E T K K A T V T E
BINARY_3566 T T Y K L I L N L K Q L K E E A I T E L V D A A T A E K Y F K L Y A N A K T V E G V W T L K D E T K K F T V T E
BINARY_3567 T T Y K L I L N L K Q L K E E A I T E L V D A A T A E K Y F K L Y A N A K T V E G V W T L K D E T K T A T V T E
BINARY_3568 T T Y K L I L N L K Q L K E E A I T E L V D A A T A E K Y F K L Y A N A K T V E G V W T L K D E T K T F T V T E
BINARY_3569 T T Y K L I L N L K Q L K E E A I T E L V D A A T A E K Y F K L Y A N A K T V E G V W T Y K D E I L K A T V T E
BINARY_3570 T T Y K L I L N L K Q L K E E A I T E L V D A A T A E K Y F K L Y A N A K T V E G V W T Y K D E I L K F T V T E
BINARY_3571 T T Y K L I L N L K Q L K E E A I T E L V D A A T A E K Y F K L Y A N A K T V E G V W T Y K D E I L T A T V T E
BINARY_3572 T T Y K L I L N L K Q L K E E A I T E L V D A A T A E K Y F K L Y A N A K T V E G V W T Y K D E I L T F T V T E
BINARY_3573 T T Y K L I L N L K Q L K E E A I T E L V D A A T A E K Y F K L Y A N A K T V E G V W T Y K D E I K K A T V T E
BINARY_3574 T T Y K L I L N L K Q L K E E A I T E L V D A A T A E K Y F K L Y A N A K T V E G V W T Y K D E I K K F T V T E
BINARY_3575 T T Y K L I L N L K Q L K E E A I T E L V D A A T A E K Y F K L Y A N A K T V E G V W T Y K D E I K T A T V T E
BINARY_3576 T T Y K L I L N L K Q L K E E A I T E L V D A A T A E K Y F K L Y A N A K T V E G V W T Y K D E I K T F T V T E
BINARY_3577 T T Y K L I L N L K Q L K E E A I T E L V D A A T A E K Y F K L Y A N A K T V E G V W T Y K D E T L K A T V T E
BINARY_3578 T T Y K L I L N L K Q L K E E A I T E L V D A A T A E K Y F K L Y A N A K T V E G V W T Y K D E T L K F T V T E
BINARY_3579 T T Y K L I L N L K Q L K E E A I T E L V D A A T A E K Y F K L Y A N A K T V E G V W T Y K D E T L T A T V T E
BINARY_3580 T T Y K L I L N L K Q L K E E A I T E L V D A A T A E K Y F K L Y A N A K T V E G V W T Y K D E T L T F T V T E
BINARY_3581 T T Y K L I L N L K Q L K E E A I T E L V D A A T A E K Y F K L Y A N A K T V E G V W T Y K D E T K K A T V T E
BINARY_3582 T T Y K L I L N L K Q L K E E A I T E L V D A A T A E K Y F K L Y A N A K T V E G V W T Y K D E T K K F T V T E
BINARY_3583 T T Y K L I L N L K Q L K E E A I T E L V D A A T A E K Y F K L Y A N A K T V E G V W T Y K D E T K T A T V T E
BINARY_3584 T T Y K L I L N L K Q L K E E A I T E L V D A A T A E K Y F K L Y A N A K T V E G V W T Y K D E T K T F T V T E
BINARY_3585 T T Y K L I L N L K Q L K E E A I T E A V D A G I A E K Y I K L I A N A K T V E G V W T L K D E I L K A T V T E
BINARY_3586 T T Y K L I L N L K Q L K E E A I T E A V D A G I A E K Y I K L I A N A K T V E G V W T L K D E I L K F T V T E
BINARY_3587 T T Y K L I L N L K Q L K E E A I T E A V D A G I A E K Y I K L I A N A K T V E G V W T L K D E I L T A T V T E
BINARY_3588 T T Y K L I L N L K Q L K E E A I T E A V D A G I A E K Y I K L I A N A K T V E G V W T L K D E I L T F T V T E
BINARY_3589 T T Y K L I L N L K Q L K E E A I T E A V D A G I A E K Y I K L I A N A K T V E G V W T L K D E I K K A T V T E
BINARY_3590 T T Y K L I L N L K Q L K E E A I T E A V D A G I A E K Y I K L I A N A K T V E G V W T L K D E I K K F T V T E
BINARY_3591 T T Y K L I L N L K Q L K E E A I T E A V D A G I A E K Y I K L I A N A K T V E G V W T L K D E I K T A T V T E
BINARY_3592 T T Y K L I L N L K Q L K E E A I T E A V D A G I A E K Y I K L I A N A K T V E G V W T L K D E I K T F T V T E
BINARY_3593 T T Y K L I L N L K Q L K E E A I T E A V D A G I A E K Y I K L I A N A K T V E G V W T L K D E T L K A T V T E
BINARY_3594 T T Y K L I L N L K Q L K E E A I T E A V D A G I A E K Y I K L I A N A K T V E G V W T L K D E T L K F T V T E
BINARY_3595 T T Y K L I L N L K Q L K E E A I T E A V D A G I A E K Y I K L I A N A K T V E G V W T L K D E T L T A T V T E
BINARY_3596 T T Y K L I L N L K Q L K E E A I T E A V D A G I A E K Y I K L I A N A K T V E G V W T L K D E T L T F T V T E
BINARY_3597 T T Y K L I L N L K Q L K E E A I T E A V D A G I A E K Y I K L I A N A K T V E G V W T L K D E T K K A T V T E
BINARY_3598 T T Y K L I L N L K Q L K E E A I T E A V D A G I A E K Y I K L I A N A K T V E G V W T L K D E T K K F T V T E
BINARY_3599 T T Y K L I L N L K Q L K E E A I T E A V D A G I A E K Y I K L I A N A K T V E G V W T L K D E T K T A T V T E
BINARY_3600 T T Y K L I L N L K Q L K E E A I T E A V D A G I A E K Y I K L I A N A K T V E G V W T L K D E T K T F T V T E
BINARY_3601 T T Y K L I L N L K Q L K E E A I T E A V D A G I A E K Y I K L I A N A K T V E G V W T Y K D E I L K A T V T E
BINARY_3602 T T Y K L I L N L K Q L K E E A I T E A V D A G I A E K Y I K L I A N A K T V E G V W T Y K D E I L K F T V T E
BINARY_3603 T T Y K L I L N L K Q L K E E A I T E A V D A G I A E K Y I K L I A N A K T V E G V W T Y K D E I L T A T V T E
BINARY_3604 T T Y K L I L N L K Q L K E E A I T E A V D A G I A E K Y I K L I A N A K T V E G V W T Y K D E I L T F T V T E
BINARY_3605 T T Y K L I L N L K Q L K E E A I T E A V D A G I A E K Y I K L I A N A K T V E G V W T Y K D E I K K A T V T E
BINARY_3606 T T Y K L I L N L K Q L K E E A I T E A V D A G I A E K Y I K L I A N A K T V E G V W T Y K D E I K K F T V T E
BINARY_3607 T T Y K L I L N L K Q L K E E A I T E A V D A G I A E K Y I K L I A N A K T V E G V W T Y K D E I K T A T V T E
BINARY_3608 T T Y K L I L N L K Q L K E E A I T E A V D A G I A E K Y I K L I A N A K T V E G V W T Y K D E I K T F T V T E
BINARY_3609 T T Y K L I L N L K Q L K E E A I T E A V D A G I A E K Y I K L I A N A K T V E G V W T Y K D E T L K A T V T E
BINARY_3610 T T Y K L I L N L K Q L K E E A I T E A V D A G I A E K Y I K L I A N A K T V E G V W T Y K D E T L K F T V T E
BINARY_3611 T T Y K L I L N L K Q L K E E A I T E A V D A G I A E K Y I K L I A N A K T V E G V W T Y K D E T L T A T V T E
BINARY_3612 T T Y K L I L N L K Q L K E E A I T E A V D A G I A E K Y I K L I A N A K T V E G V W T Y K D E T L T F T V T E
BINARY_3613 T T Y K L I L N L K Q L K E E A I T E A V D A G I A E K Y I K L I A N A K T V E G V W T Y K D E T K K A T V T E
BINARY_3614 T T Y K L I L N L K Q L K E E A I T E A V D A G I A E K Y I K L I A N A K T V E G V W T Y K D E T K K F T V T E
BINARY_3615 T T Y K L I L N L K Q L K E E A I T E A V D A G I A E K Y I K L I A N A K T V E G V W T Y K D E T K T A T V T E
BINARY_3616 T T Y K L I L N L K Q L K E E A I T E A V D A G I A E K Y I K L I A N A K T V E G V W T Y K D E T K T F T V T E
BINARY_3617 T T Y K L I L N L K Q L K E E A I T E A V D A G I A E K Y I K L Y A N A K T V E G V W T L K D E I L K A T V T E
BINARY_3618 T T Y K L I L N L K Q L K E E A I T E A V D A G I A E K Y I K L Y A N A K T V E G V W T L K D E I L K F T V T E
BINARY_3619 T T Y K L I L N L K Q L K E E A I T E A V D A G I A E K Y I K L Y A N A K T V E G V W T L K D E I L T A T V T E
BINARY_3620 T T Y K L I L N L K Q L K E E A I T E A V D A G I A E K Y I K L Y A N A K T V E G V W T L K D E I L T F T V T E
BINARY_3621 T T Y K L I L N L K Q L K E E A I T E A V D A G I A E K Y I K L Y A N A K T V E G V W T L K D E I K K A T V T E
BINARY_3622 T T Y K L I L N L K Q L K E E A I T E A V D A G I A E K Y I K L Y A N A K T V E G V W T L K D E I K K F T V T E
BINARY_3623 T T Y K L I L N L K Q L K E E A I T E A V D A G I A E K Y I K L Y A N A K T V E G V W T L K D E I K T A T V T E
BINARY_3624 T T Y K L I L N L K Q L K E E A I T E A V D A G I A E K Y I K L Y A N A K T V E G V W T L K D E I K T F T V T E
BINARY_3625 T T Y K L I L N L K Q L K E E A I T E A V D A G I A E K Y I K L Y A N A K T V E G V W T L K D E T L K A T V T E
BINARY_3626 T T Y K L I L N L K Q L K E E A I T E A V D A G I A E K Y I K L Y A N A K T V E G V W T L K D E T L K F T V T E

BINARY_3627 T T Y K L I L N L K Q L K E E A I T E A V D A G I A E K Y I K L Y A N A K T V E G V W T L K D E T L T A T V T E
BINARY_3628 T T Y K L I L N L K Q L K E E A I T E A V D A G I A E K Y I K L Y A N A K T V E G V W T L K D E T L T F T V T E
BINARY_3629 T T Y K L I L N L K Q L K E E A I T E A V D A G I A E K Y I K L Y A N A K T V E G V W T L K D E T K K A T V T E
BINARY_3630 T T Y K L I L N L K Q L K E E A I T E A V D A G I A E K Y I K L Y A N A K T V E G V W T L K D E T K K F T V T E
BINARY_3631 T T Y K L I L N L K Q L K E E A I T E A V D A G I A E K Y I K L Y A N A K T V E G V W T L K D E T K T A T V T E
BINARY_3632 T T Y K L I L N L K Q L K E E A I T E A V D A G I A E K Y I K L Y A N A K T V E G V W T L K D E T K T F T V T E
BINARY_3633 T T Y K L I L N L K Q L K E E A I T E A V D A G I A E K Y I K L Y A N A K T V E G V W T Y K D E I L K A T V T E
BINARY_3634 T T Y K L I L N L K Q L K E E A I T E A V D A G I A E K Y I K L Y A N A K T V E G V W T Y K D E I L K F T V T E
BINARY_3635 T T Y K L I L N L K Q L K E E A I T E A V D A G I A E K Y I K L Y A N A K T V E G V W T Y K D E I L T A T V T E
BINARY_3636 T T Y K L I L N L K Q L K E E A I T E A V D A G I A E K Y I K L Y A N A K T V E G V W T Y K D E I L T F T V T E
BINARY_3637 T T Y K L I L N L K Q L K E E A I T E A V D A G I A E K Y I K L Y A N A K T V E G V W T Y K D E I K K A T V T E
BINARY_3638 T T Y K L I L N L K Q L K E E A I T E A V D A G I A E K Y I K L Y A N A K T V E G V W T Y K D E I K K F T V T E
BINARY_3639 T T Y K L I L N L K Q L K E E A I T E A V D A G I A E K Y I K L Y A N A K T V E G V W T Y K D E I K T A T V T E
BINARY_3640 T T Y K L I L N L K Q L K E E A I T E A V D A G I A E K Y I K L Y A N A K T V E G V W T Y K D E I K T F T V T E
BINARY_3641 T T Y K L I L N L K Q L K E E A I T E A V D A G I A E K Y I K L Y A N A K T V E G V W T Y K D E T L K A T V T E
BINARY_3642 T T Y K L I L N L K Q L K E E A I T E A V D A G I A E K Y I K L Y A N A K T V E G V W T Y K D E T L K F T V T E
BINARY_3643 T T Y K L I L N L K Q L K E E A I T E A V D A G I A E K Y I K L Y A N A K T V E G V W T Y K D E T L T A T V T E
BINARY_3644 T T Y K L I L N L K Q L K E E A I T E A V D A G I A E K Y I K L Y A N A K T V E G V W T Y K D E T L T F T V T E
BINARY_3645 T T Y K L I L N L K Q L K E E A I T E A V D A G I A E K Y I K L Y A N A K T V E G V W T Y K D E T K K A T V T E
BINARY_3646 T T Y K L I L N L K Q L K E E A I T E A V D A G I A E K Y I K L Y A N A K T V E G V W T Y K D E T K K F T V T E
BINARY_3647 T T Y K L I L N L K Q L K E E A I T E A V D A G I A E K Y I K L Y A N A K T V E G V W T Y K D E T K T A T V T E
BINARY_3648 T T Y K L I L N L K Q L K E E A I T E A V D A G I A E K Y I K L Y A N A K T V E G V W T Y K D E T K T F T V T E
BINARY_3649 T T Y K L I L N L K Q L K E E A I T E A V D A G I A E K Y F K L I A N A K T V E G V W T L K D E I L K A T V T E
BINARY_3650 T T Y K L I L N L K Q L K E E A I T E A V D A G I A E K Y F K L I A N A K T V E G V W T L K D E I L K F T V T E
BINARY_3651 T T Y K L I L N L K Q L K E E A I T E A V D A G I A E K Y F K L I A N A K T V E G V W T L K D E I L T A T V T E
BINARY_3652 T T Y K L I L N L K Q L K E E A I T E A V D A G I A E K Y F K L I A N A K T V E G V W T L K D E I L T F T V T E
BINARY_3653 T T Y K L I L N L K Q L K E E A I T E A V D A G I A E K Y F K L I A N A K T V E G V W T L K D E I K K A T V T E
BINARY_3654 T T Y K L I L N L K Q L K E E A I T E A V D A G I A E K Y F K L I A N A K T V E G V W T L K D E I K K F T V T E
BINARY_3655 T T Y K L I L N L K Q L K E E A I T E A V D A G I A E K Y F K L I A N A K T V E G V W T L K D E I K T A T V T E
BINARY_3656 T T Y K L I L N L K Q L K E E A I T E A V D A G I A E K Y F K L I A N A K T V E G V W T L K D E I K T F T V T E
BINARY_3657 T T Y K L I L N L K Q L K E E A I T E A V D A G I A E K Y F K L I A N A K T V E G V W T L K D E T L K A T V T E
BINARY_3658 T T Y K L I L N L K Q L K E E A I T E A V D A G I A E K Y F K L I A N A K T V E G V W T L K D E T L K F T V T E
BINARY_3659 T T Y K L I L N L K Q L K E E A I T E A V D A G I A E K Y F K L I A N A K T V E G V W T L K D E T L T A T V T E
BINARY_3660 T T Y K L I L N L K Q L K E E A I T E A V D A G I A E K Y F K L I A N A K T V E G V W T L K D E T L T F T V T E
BINARY_3661 T T Y K L I L N L K Q L K E E A I T E A V D A G I A E K Y F K L I A N A K T V E G V W T L K D E T K K A T V T E
BINARY_3662 T T Y K L I L N L K Q L K E E A I T E A V D A G I A E K Y F K L I A N A K T V E G V W T L K D E T K K F T V T E
BINARY_3663 T T Y K L I L N L K Q L K E E A I T E A V D A G I A E K Y F K L I A N A K T V E G V W T L K D E T K T A T V T E
BINARY_3664 T T Y K L I L N L K Q L K E E A I T E A V D A G I A E K Y F K L I A N A K T V E G V W T L K D E T K T F T V T E
BINARY_3665 T T Y K L I L N L K Q L K E E A I T E A V D A G I A E K Y F K L I A N A K T V E G V W T Y K D E I L K A T V T E
BINARY_3666 T T Y K L I L N L K Q L K E E A I T E A V D A G I A E K Y F K L I A N A K T V E G V W T Y K D E I L K F T V T E
BINARY_3667 T T Y K L I L N L K Q L K E E A I T E A V D A G I A E K Y F K L I A N A K T V E G V W T Y K D E I L T A T V T E
BINARY_3668 T T Y K L I L N L K Q L K E E A I T E A V D A G I A E K Y F K L I A N A K T V E G V W T Y K D E I L T F T V T E
BINARY_3669 T T Y K L I L N L K Q L K E E A I T E A V D A G I A E K Y F K L I A N A K T V E G V W T Y K D E I K K A T V T E
BINARY_3670 T T Y K L I L N L K Q L K E E A I T E A V D A G I A E K Y F K L I A N A K T V E G V W T Y K D E I K K F T V T E
BINARY_3671 T T Y K L I L N L K Q L K E E A I T E A V D A G I A E K Y F K L I A N A K T V E G V W T Y K D E I K T A T V T E
BINARY_3672 T T Y K L I L N L K Q L K E E A I T E A V D A G I A E K Y F K L I A N A K T V E G V W T Y K D E I K T F T V T E
BINARY_3673 T T Y K L I L N L K Q L K E E A I T E A V D A G I A E K Y F K L I A N A K T V E G V W T Y K D E T L K A T V T E
BINARY_3674 T T Y K L I L N L K Q L K E E A I T E A V D A G I A E K Y F K L I A N A K T V E G V W T Y K D E T L K F T V T E
BINARY_3675 T T Y K L I L N L K Q L K E E A I T E A V D A G I A E K Y F K L I A N A K T V E G V W T Y K D E T L T A T V T E
BINARY_3676 T T Y K L I L N L K Q L K E E A I T E A V D A G I A E K Y F K L I A N A K T V E G V W T Y K D E T L T F T V T E
BINARY_3677 T T Y K L I L N L K Q L K E E A I T E A V D A G I A E K Y F K L I A N A K T V E G V W T Y K D E T K K A T V T E
BINARY_3678 T T Y K L I L N L K Q L K E E A I T E A V D A G I A E K Y F K L I A N A K T V E G V W T Y K D E T K K F T V T E
BINARY_3679 T T Y K L I L N L K Q L K E E A I T E A V D A G I A E K Y F K L I A N A K T V E G V W T Y K D E T K T A T V T E
BINARY_3680 T T Y K L I L N L K Q L K E E A I T E A V D A G I A E K Y F K L I A N A K T V E G V W T Y K D E T K T F T V T E
BINARY_3681 T T Y K L I L N L K Q L K E E A I T E A V D A G I A E K Y F K L Y A N A K T V E G V W T L K D E I L K A T V T E
BINARY_3682 T T Y K L I L N L K Q L K E E A I T E A V D A G I A E K Y F K L Y A N A K T V E G V W T L K D E I L K F T V T E
BINARY_3683 T T Y K L I L N L K Q L K E E A I T E A V D A G I A E K Y F K L Y A N A K T V E G V W T L K D E I L T A T V T E
BINARY_3684 T T Y K L I L N L K Q L K E E A I T E A V D A G I A E K Y F K L Y A N A K T V E G V W T L K D E I L T F T V T E
BINARY_3685 T T Y K L I L N L K Q L K E E A I T E A V D A G I A E K Y F K L Y A N A K T V E G V W T L K D E I K K A T V T E
BINARY_3686 T T Y K L I L N L K Q L K E E A I T E A V D A G I A E K Y F K L Y A N A K T V E G V W T L K D E I K K F T V T E
BINARY_3687 T T Y K L I L N L K Q L K E E A I T E A V D A G I A E K Y F K L Y A N A K T V E G V W T L K D E I K T A T V T E
BINARY_3688 T T Y K L I L N L K Q L K E E A I T E A V D A G I A E K Y F K L Y A N A K T V E G V W T L K D E I K T F T V T E
BINARY_3689 T T Y K L I L N L K Q L K E E A I T E A V D A G I A E K Y F K L Y A N A K T V E G V W T L K D E T L K A T V T E
BINARY_3690 T T Y K L I L N L K Q L K E E A I T E A V D A G I A E K Y F K L Y A N A K T V E G V W T L K D E T L K F T V T E
BINARY_3691 T T Y K L I L N L K Q L K E E A I T E A V D A G I A E K Y F K L Y A N A K T V E G V W T L K D E T L T A T V T E
BINARY_3692 T T Y K L I L N L K Q L K E E A I T E A V D A G I A E K Y F K L Y A N A K T V E G V W T L K D E T L T F T V T E
BINARY_3693 T T Y K L I L N L K Q L K E E A I T E A V D A G I A E K Y F K L Y A N A K T V E G V W T L K D E T K K A T V T E
BINARY_3694 T T Y K L I L N L K Q L K E E A I T E A V D A G I A E K Y F K L Y A N A K T V E G V W T L K D E T K K F T V T E
BINARY_3695 T T Y K L I L N L K Q L K E E A I T E A V D A G I A E K Y F K L Y A N A K T V E G V W T L K D E T K T A T V T E
BINARY_3696 T T Y K L I L N L K Q L K E E A I T E A V D A G I A E K Y F K L Y A N A K T V E G V W T L K D E T K T F T V T E
BINARY_3697 T T Y K L I L N L K Q L K E E A I T E A V D A G I A E K Y F K L Y A N A K T V E G V W T Y K D E I L K A T V T E
BINARY_3698 T T Y K L I L N L K Q L K E E A I T E A V D A G I A E K Y F K L Y A N A K T V E G V W T Y K D E I L K F T V T E
BINARY_3699 T T Y K L I L N L K Q L K E E A I T E A V D A G I A E K Y F K L Y A N A K T V E G V W T Y K D E I L T A T V T E
BINARY_3700 T T Y K L I L N L K Q L K E E A I T E A V D A G I A E K Y F K L Y A N A K T V E G V W T Y K D E I L T F T V T E
BINARY_3701 T T Y K L I L N L K Q L K E E A I T E A V D A G I A E K Y F K L Y A N A K T V E G V W T Y K D E I K K A T V T E
BINARY_3702 T T Y K L I L N L K Q L K E E A I T E A V D A G I A E K Y F K L Y A N A K T V E G V W T Y K D E I K K F T V T E
BINARY_3703 T T Y K L I L N L K Q L K E E A I T E A V D A G I A E K Y F K L Y A N A K T V E G V W T Y K D E I K T A T V T E
BINARY_3704 T T Y K L I L N L K Q L K E E A I T E A V D A G I A E K Y F K L Y A N A K T V E G V W T Y K D E I K T F T V T E
BINARY_3705 T T Y K L I L N L K Q L K E E A I T E A V D A G I A E K Y F K L Y A N A K T V E G V W T Y K D E T L K A T V T E
BINARY_3706 T T Y K L I L N L K Q L K E E A I T E A V D A G I A E K Y F K L Y A N A K T V E G V W T Y K D E T L K F T V T E
BINARY_3707 T T Y K L I L N L K Q L K E E A I T E A V D A G I A E K Y F K L Y A N A K T V E G V W T Y K D E T L T A T V T E
BINARY_3708 T T Y K L I L N L K Q L K E E A I T E A V D A G I A E K Y F K L Y A N A K T V E G V W T Y K D E T L T F T V T E
BINARY_3709 T T Y K L I L N L K Q L K E E A I T E A V D A G I A E K Y F K L Y A N A K T V E G V W T Y K D E T K K A T V T E
BINARY_3710 T T Y K L I L N L K Q L K E E A I T E A V D A G I A E K Y F K L Y A N A K T V E G V W T Y K D E T K K F T V T E
BINARY_3711 T T Y K L I L N L K Q L K E E A I T E A V D A G I A E K Y F K L Y A N A K T V E G V W T Y K D E T K T A T V T E
BINARY_3712 T T Y K L I L N L K Q L K E E A I T E A V D A G I A E K Y F K L Y A N A K T V E G V W T Y K D E T K T F T V T E
BINARY_3713 T T Y K L I L N L K Q L K E E A I T E A V D A G T A E K Y I K L I A N A K T V E G V W T L K D E I L K A T V T E
BINARY_3714 T T Y K L I L N L K Q L K E E A I T E A V D A G T A E K Y I K L I A N A K T V E G V W T L K D E I L K F T V T E
BINARY_3715 T T Y K L I L N L K Q L K E E A I T E A V D A G T A E K Y I K L I A N A K T V E G V W T L K D E I L T A T V T E
BINARY_3716 T T Y K L I L N L K Q L K E E A I T E A V D A G T A E K Y I K L I A N A K T V E G V W T L K D E I L T F T V T E
BINARY_3717 T T Y K L I L N L K Q L K E E A I T E A V D A G T A E K Y I K L I A N A K T V E G V W T L K D E I K K A T V T E
BINARY_3718 T T Y K L I L N L K Q L K E E A I T E A V D A G T A E K Y I K L I A N A K T V E G V W T L K D E I K K F T V T E
BINARY_3719 T T Y K L I L N L K Q L K E E A I T E A V D A G T A E K Y I K L I A N A K T V E G V W T L K D E I K T A T V T E
BINARY_3720 T T Y K L I L N L K Q L K E E A I T E A V D A G T A E K Y I K L I A N A K T V E G V W T L K D E I K T F T V T E
BINARY_3721 T T Y K L I L N L K Q L K E E A I T E A V D A G T A E K Y I K L I A N A K T V E G V W T L K D E T L K A T V T E

BINARY_3722 T T Y K L I L N L K Q L K E E A I T E A V D A G T A E K Y I K L I A N A K T V E G V W T L K D E T L K F T V T E
BINARY_3723 T T Y K L I L N L K Q L K E E A I T E A V D A G T A E K Y I K L I A N A K T V E G V W T L K D E T L T A T V T E
BINARY_3724 T T Y K L I L N L K Q L K E E A I T E A V D A G T A E K Y I K L I A N A K T V E G V W T L K D E T L T F T V T E
BINARY_3725 T T Y K L I L N L K Q L K E E A I T E A V D A G T A E K Y I K L I A N A K T V E G V W T L K D E T K K A T V T E
BINARY_3726 T T Y K L I L N L K Q L K E E A I T E A V D A G T A E K Y I K L I A N A K T V E G V W T L K D E T K K F T V T E
BINARY_3727 T T Y K L I L N L K Q L K E E A I T E A V D A G T A E K Y I K L I A N A K T V E G V W T L K D E T K T A T V T E
BINARY_3728 T T Y K L I L N L K Q L K E E A I T E A V D A G T A E K Y I K L I A N A K T V E G V W T L K D E T K T F T V T E
BINARY_3729 T T Y K L I L N L K Q L K E E A I T E A V D A G T A E K Y I K L I A N A K T V E G V W T Y K D E I L K A T V T E
BINARY_3730 T T Y K L I L N L K Q L K E E A I T E A V D A G T A E K Y I K L I A N A K T V E G V W T Y K D E I L K F T V T E
BINARY_3731 T T Y K L I L N L K Q L K E E A I T E A V D A G T A E K Y I K L I A N A K T V E G V W T Y K D E I L T A T V T E
BINARY_3732 T T Y K L I L N L K Q L K E E A I T E A V D A G T A E K Y I K L I A N A K T V E G V W T Y K D E I L T F T V T E
BINARY_3733 T T Y K L I L N L K Q L K E E A I T E A V D A G T A E K Y I K L I A N A K T V E G V W T Y K D E I K K A T V T E
BINARY_3734 T T Y K L I L N L K Q L K E E A I T E A V D A G T A E K Y I K L I A N A K T V E G V W T Y K D E I K K F T V T E
BINARY_3735 T T Y K L I L N L K Q L K E E A I T E A V D A G T A E K Y I K L I A N A K T V E G V W T Y K D E I K T A T V T E
BINARY_3736 T T Y K L I L N L K Q L K E E A I T E A V D A G T A E K Y I K L I A N A K T V E G V W T Y K D E I K T F T V T E
BINARY_3737 T T Y K L I L N L K Q L K E E A I T E A V D A G T A E K Y I K L I A N A K T V E G V W T Y K D E T L K A T V T E
BINARY_3738 T T Y K L I L N L K Q L K E E A I T E A V D A G T A E K Y I K L I A N A K T V E G V W T Y K D E T L K F T V T E
BINARY_3739 T T Y K L I L N L K Q L K E E A I T E A V D A G T A E K Y I K L I A N A K T V E G V W T Y K D E T L T A T V T E
BINARY_3740 T T Y K L I L N L K Q L K E E A I T E A V D A G T A E K Y I K L I A N A K T V E G V W T Y K D E T L T F T V T E
BINARY_3741 T T Y K L I L N L K Q L K E E A I T E A V D A G T A E K Y I K L I A N A K T V E G V W T Y K D E T K K A T V T E
BINARY_3742 T T Y K L I L N L K Q L K E E A I T E A V D A G T A E K Y I K L I A N A K T V E G V W T Y K D E T K K F T V T E
BINARY_3743 T T Y K L I L N L K Q L K E E A I T E A V D A G T A E K Y I K L I A N A K T V E G V W T Y K D E T K T A T V T E
BINARY_3744 T T Y K L I L N L K Q L K E E A I T E A V D A G T A E K Y I K L I A N A K T V E G V W T Y K D E T K T F T V T E
BINARY_3745 T T Y K L I L N L K Q L K E E A I T E A V D A G T A E K Y I K L Y A N A K T V E G V W T L K D E I L K A T V T E
BINARY_3746 T T Y K L I L N L K Q L K E E A I T E A V D A G T A E K Y I K L Y A N A K T V E G V W T L K D E I L K F T V T E
BINARY_3747 T T Y K L I L N L K Q L K E E A I T E A V D A G T A E K Y I K L Y A N A K T V E G V W T L K D E I L T A T V T E
BINARY_3748 T T Y K L I L N L K Q L K E E A I T E A V D A G T A E K Y I K L Y A N A K T V E G V W T L K D E I L T F T V T E
BINARY_3749 T T Y K L I L N L K Q L K E E A I T E A V D A G T A E K Y I K L Y A N A K T V E G V W T L K D E I K K A T V T E
BINARY_3750 T T Y K L I L N L K Q L K E E A I T E A V D A G T A E K Y I K L Y A N A K T V E G V W T L K D E I K K F T V T E
BINARY_3751 T T Y K L I L N L K Q L K E E A I T E A V D A G T A E K Y I K L Y A N A K T V E G V W T L K D E I K T A T V T E
BINARY_3752 T T Y K L I L N L K Q L K E E A I T E A V D A G T A E K Y I K L Y A N A K T V E G V W T L K D E I K T F T V T E
BINARY_3753 T T Y K L I L N L K Q L K E E A I T E A V D A G T A E K Y I K L Y A N A K T V E G V W T L K D E T L K A T V T E
BINARY_3754 T T Y K L I L N L K Q L K E E A I T E A V D A G T A E K Y I K L Y A N A K T V E G V W T L K D E T L K F T V T E
BINARY_3755 T T Y K L I L N L K Q L K E E A I T E A V D A G T A E K Y I K L Y A N A K T V E G V W T L K D E T L T A T V T E
BINARY_3756 T T Y K L I L N L K Q L K E E A I T E A V D A G T A E K Y I K L Y A N A K T V E G V W T L K D E T L T F T V T E
BINARY_3757 T T Y K L I L N L K Q L K E E A I T E A V D A G T A E K Y I K L Y A N A K T V E G V W T L K D E T K K A T V T E
BINARY_3758 T T Y K L I L N L K Q L K E E A I T E A V D A G T A E K Y I K L Y A N A K T V E G V W T L K D E T K K F T V T E
BINARY_3759 T T Y K L I L N L K Q L K E E A I T E A V D A G T A E K Y I K L Y A N A K T V E G V W T L K D E T K T A T V T E
BINARY_3760 T T Y K L I L N L K Q L K E E A I T E A V D A G T A E K Y I K L Y A N A K T V E G V W T L K D E T K T F T V T E
BINARY_3761 T T Y K L I L N L K Q L K E E A I T E A V D A G T A E K Y I K L Y A N A K T V E G V W T Y K D E I L K A T V T E
BINARY_3762 T T Y K L I L N L K Q L K E E A I T E A V D A G T A E K Y I K L Y A N A K T V E G V W T Y K D E I L K F T V T E
BINARY_3763 T T Y K L I L N L K Q L K E E A I T E A V D A G T A E K Y I K L Y A N A K T V E G V W T Y K D E I L T A T V T E
BINARY_3764 T T Y K L I L N L K Q L K E E A I T E A V D A G T A E K Y I K L Y A N A K T V E G V W T Y K D E I L T F T V T E
BINARY_3765 T T Y K L I L N L K Q L K E E A I T E A V D A G T A E K Y I K L Y A N A K T V E G V W T Y K D E I K K A T V T E
BINARY_3766 T T Y K L I L N L K Q L K E E A I T E A V D A G T A E K Y I K L Y A N A K T V E G V W T Y K D E I K K F T V T E
BINARY_3767 T T Y K L I L N L K Q L K E E A I T E A V D A G T A E K Y I K L Y A N A K T V E G V W T Y K D E I K T A T V T E
BINARY_3768 T T Y K L I L N L K Q L K E E A I T E A V D A G T A E K Y I K L Y A N A K T V E G V W T Y K D E I K T F T V T E
BINARY_3769 T T Y K L I L N L K Q L K E E A I T E A V D A G T A E K Y I K L Y A N A K T V E G V W T Y K D E T L K A T V T E
BINARY_3770 T T Y K L I L N L K Q L K E E A I T E A V D A G T A E K Y I K L Y A N A K T V E G V W T Y K D E T L K F T V T E
BINARY_3771 T T Y K L I L N L K Q L K E E A I T E A V D A G T A E K Y I K L Y A N A K T V E G V W T Y K D E T L T A T V T E
BINARY_3772 T T Y K L I L N L K Q L K E E A I T E A V D A G T A E K Y I K L Y A N A K T V E G V W T Y K D E T L T F T V T E
BINARY_3773 T T Y K L I L N L K Q L K E E A I T E A V D A G T A E K Y I K L Y A N A K T V E G V W T Y K D E T K K A T V T E
BINARY_3774 T T Y K L I L N L K Q L K E E A I T E A V D A G T A E K Y I K L Y A N A K T V E G V W T Y K D E T K K F T V T E
BINARY_3775 T T Y K L I L N L K Q L K E E A I T E A V D A G T A E K Y I K L Y A N A K T V E G V W T Y K D E T K T A T V T E
BINARY_3776 T T Y K L I L N L K Q L K E E A I T E A V D A G T A E K Y I K L Y A N A K T V E G V W T Y K D E T K T F T V T E
BINARY_3777 T T Y K L I L N L K Q L K E E A I T E A V D A G T A E K Y F K L I A N A K T V E G V W T L K D E I L K A T V T E
BINARY_3778 T T Y K L I L N L K Q L K E E A I T E A V D A G T A E K Y F K L I A N A K T V E G V W T L K D E I L K F T V T E
BINARY_3779 T T Y K L I L N L K Q L K E E A I T E A V D A G T A E K Y F K L I A N A K T V E G V W T L K D E I L T A T V T E
BINARY_3780 T T Y K L I L N L K Q L K E E A I T E A V D A G T A E K Y F K L I A N A K T V E G V W T L K D E I L T F T V T E
BINARY_3781 T T Y K L I L N L K Q L K E E A I T E A V D A G T A E K Y F K L I A N A K T V E G V W T L K D E I K K A T V T E
BINARY_3782 T T Y K L I L N L K Q L K E E A I T E A V D A G T A E K Y F K L I A N A K T V E G V W T L K D E I K K F T V T E
BINARY_3783 T T Y K L I L N L K Q L K E E A I T E A V D A G T A E K Y F K L I A N A K T V E G V W T L K D E I K T A T V T E
BINARY_3784 T T Y K L I L N L K Q L K E E A I T E A V D A G T A E K Y F K L I A N A K T V E G V W T L K D E I K T F T V T E
BINARY_3785 T T Y K L I L N L K Q L K E E A I T E A V D A G T A E K Y F K L I A N A K T V E G V W T L K D E T L K A T V T E
BINARY_3786 T T Y K L I L N L K Q L K E E A I T E A V D A G T A E K Y F K L I A N A K T V E G V W T L K D E T L K F T V T E
BINARY_3787 T T Y K L I L N L K Q L K E E A I T E A V D A G T A E K Y F K L I A N A K T V E G V W T L K D E T L T A T V T E
BINARY_3788 T T Y K L I L N L K Q L K E E A I T E A V D A G T A E K Y F K L I A N A K T V E G V W T L K D E T L T F T V T E
BINARY_3789 T T Y K L I L N L K Q L K E E A I T E A V D A G T A E K Y F K L I A N A K T V E G V W T L K D E T K K A T V T E
BINARY_3790 T T Y K L I L N L K Q L K E E A I T E A V D A G T A E K Y F K L I A N A K T V E G V W T L K D E T K K F T V T E
BINARY_3791 T T Y K L I L N L K Q L K E E A I T E A V D A G T A E K Y F K L I A N A K T V E G V W T L K D E T K T A T V T E
BINARY_3792 T T Y K L I L N L K Q L K E E A I T E A V D A G T A E K Y F K L I A N A K T V E G V W T L K D E T K T F T V T E
BINARY_3793 T T Y K L I L N L K Q L K E E A I T E A V D A G T A E K Y F K L I A N A K T V E G V W T Y K D E I L K A T V T E
BINARY_3794 T T Y K L I L N L K Q L K E E A I T E A V D A G T A E K Y F K L I A N A K T V E G V W T Y K D E I L K F T V T E
BINARY_3795 T T Y K L I L N L K Q L K E E A I T E A V D A G T A E K Y F K L I A N A K T V E G V W T Y K D E I L T A T V T E
BINARY_3796 T T Y K L I L N L K Q L K E E A I T E A V D A G T A E K Y F K L I A N A K T V E G V W T Y K D E I L T F T V T E
BINARY_3797 T T Y K L I L N L K Q L K E E A I T E A V D A G T A E K Y F K L I A N A K T V E G V W T Y K D E I K K A T V T E
BINARY_3798 T T Y K L I L N L K Q L K E E A I T E A V D A G T A E K Y F K L I A N A K T V E G V W T Y K D E I K K F T V T E
BINARY_3799 T T Y K L I L N L K Q L K E E A I T E A V D A G T A E K Y F K L I A N A K T V E G V W T Y K D E I K T A T V T E
BINARY_3800 T T Y K L I L N L K Q L K E E A I T E A V D A G T A E K Y F K L I A N A K T V E G V W T Y K D E I K T F T V T E
BINARY_3801 T T Y K L I L N L K Q L K E E A I T E A V D A G T A E K Y F K L I A N A K T V E G V W T Y K D E T L K A T V T E
BINARY_3802 T T Y K L I L N L K Q L K E E A I T E A V D A G T A E K Y F K L I A N A K T V E G V W T Y K D E T L K F T V T E
BINARY_3803 T T Y K L I L N L K Q L K E E A I T E A V D A G T A E K Y F K L I A N A K T V E G V W T Y K D E T L T A T V T E
BINARY_3804 T T Y K L I L N L K Q L K E E A I T E A V D A G T A E K Y F K L I A N A K T V E G V W T Y K D E T L T F T V T E
BINARY_3805 T T Y K L I L N L K Q L K E E A I T E A V D A G T A E K Y F K L I A N A K T V E G V W T Y K D E T K K A T V T E
BINARY_3806 T T Y K L I L N L K Q L K E E A I T E A V D A G T A E K Y F K L I A N A K T V E G V W T Y K D E T K K F T V T E
BINARY_3807 T T Y K L I L N L K Q L K E E A I T E A V D A G T A E K Y F K L I A N A K T V E G V W T Y K D E T K T A T V T E
BINARY_3808 T T Y K L I L N L K Q L K E E A I T E A V D A G T A E K Y F K L I A N A K T V E G V W T Y K D E T K T F T V T E
BINARY_3809 T T Y K L I L N L K Q L K E E A I T E A V D A G T A E K Y F K L Y A N A K T V E G V W T L K D E I L K A T V T E
BINARY_3810 T T Y K L I L N L K Q L K E E A I T E A V D A G T A E K Y F K L Y A N A K T V E G V W T L K D E I L K F T V T E
BINARY_3811 T T Y K L I L N L K Q L K E E A I T E A V D A G T A E K Y F K L Y A N A K T V E G V W T L K D E I L T A T V T E
BINARY_3812 T T Y K L I L N L K Q L K E E A I T E A V D A G T A E K Y F K L Y A N A K T V E G V W T L K D E I L T F T V T E
BINARY_3813 T T Y K L I L N L K Q L K E E A I T E A V D A G T A E K Y F K L Y A N A K T V E G V W T L K D E I K K A T V T E
BINARY_3814 T T Y K L I L N L K Q L K E E A I T E A V D A G T A E K Y F K L Y A N A K T V E G V W T L K D E I K K F T V T E
BINARY_3815 T T Y K L I L N L K Q L K E E A I T E A V D A G T A E K Y F K L Y A N A K T V E G V W T L K D E I K T A T V T E
BINARY_3816 T T Y K L I L N L K Q L K E E A I T E A V D A G T A E K Y F K L Y A N A K T V E G V W T L K D E I K T F T V T E

BINARY_3817 T T Y K L I L N L K Q L K E E A I T E A V D A G T A E K Y F K L Y A N A K T V E G V W T L K D E T L K A T V T E
BINARY_3818 T T Y K L I L N L K Q L K E E A I T E A V D A G T A E K Y F K L Y A N A K T V E G V W T L K D E T L K F T V T E
BINARY_3819 T T Y K L I L N L K Q L K E E A I T E A V D A G T A E K Y F K L Y A N A K T V E G V W T L K D E T L T A T V T E
BINARY_3820 T T Y K L I L N L K Q L K E E A I T E A V D A G T A E K Y F K L Y A N A K T V E G V W T L K D E T L T F T V T E
BINARY_3821 T T Y K L I L N L K Q L K E E A I T E A V D A G T A E K Y F K L Y A N A K T V E G V W T L K D E T K K A T V T E
BINARY_3822 T T Y K L I L N L K Q L K E E A I T E A V D A G T A E K Y F K L Y A N A K T V E G V W T L K D E T K K F T V T E
BINARY_3823 T T Y K L I L N L K Q L K E E A I T E A V D A G T A E K Y F K L Y A N A K T V E G V W T L K D E T K T A T V T E
BINARY_3824 T T Y K L I L N L K Q L K E E A I T E A V D A G T A E K Y F K L Y A N A K T V E G V W T L K D E T K T F T V T E
BINARY_3825 T T Y K L I L N L K Q L K E E A I T E A V D A G T A E K Y F K L Y A N A K T V E G V W T Y K D E I L K A T V T E
BINARY_3826 T T Y K L I L N L K Q L K E E A I T E A V D A G T A E K Y F K L Y A N A K T V E G V W T Y K D E I L K F T V T E
BINARY_3827 T T Y K L I L N L K Q L K E E A I T E A V D A G T A E K Y F K L Y A N A K T V E G V W T Y K D E I L T A T V T E
BINARY_3828 T T Y K L I L N L K Q L K E E A I T E A V D A G T A E K Y F K L Y A N A K T V E G V W T Y K D E I L T F T V T E
BINARY_3829 T T Y K L I L N L K Q L K E E A I T E A V D A G T A E K Y F K L Y A N A K T V E G V W T Y K D E I K K A T V T E
BINARY_3830 T T Y K L I L N L K Q L K E E A I T E A V D A G T A E K Y F K L Y A N A K T V E G V W T Y K D E I K K F T V T E
BINARY_3831 T T Y K L I L N L K Q L K E E A I T E A V D A G T A E K Y F K L Y A N A K T V E G V W T Y K D E I K T A T V T E
BINARY_3832 T T Y K L I L N L K Q L K E E A I T E A V D A G T A E K Y F K L Y A N A K T V E G V W T Y K D E I K T F T V T E
BINARY_3833 T T Y K L I L N L K Q L K E E A I T E A V D A G T A E K Y F K L Y A N A K T V E G V W T Y K D E T L K A T V T E
BINARY_3834 T T Y K L I L N L K Q L K E E A I T E A V D A G T A E K Y F K L Y A N A K T V E G V W T Y K D E T L K F T V T E
BINARY_3835 T T Y K L I L N L K Q L K E E A I T E A V D A G T A E K Y F K L Y A N A K T V E G V W T Y K D E T L T A T V T E
BINARY_3836 T T Y K L I L N L K Q L K E E A I T E A V D A G T A E K Y F K L Y A N A K T V E G V W T Y K D E T L T F T V T E
BINARY_3837 T T Y K L I L N L K Q L K E E A I T E A V D A G T A E K Y F K L Y A N A K T V E G V W T Y K D E T K K A T V T E
BINARY_3838 T T Y K L I L N L K Q L K E E A I T E A V D A G T A E K Y F K L Y A N A K T V E G V W T Y K D E T K K F T V T E
BINARY_3839 T T Y K L I L N L K Q L K E E A I T E A V D A G T A E K Y F K L Y A N A K T V E G V W T Y K D E T K T A T V T E
BINARY_3840 T T Y K L I L N L K Q L K E E A I T E A V D A G T A E K Y F K L Y A N A K T V E G V W T Y K D E T K T F T V T E
BINARY_3841 T T Y K L I L N L K Q L K E E A I T E A V D A A I A E K Y I K L I A N A K T V E G V W T L K D E I L K A T V T E
BINARY_3842 T T Y K L I L N L K Q L K E E A I T E A V D A A I A E K Y I K L I A N A K T V E G V W T L K D E I L K F T V T E
BINARY_3843 T T Y K L I L N L K Q L K E E A I T E A V D A A I A E K Y I K L I A N A K T V E G V W T L K D E I L T A T V T E
BINARY_3844 T T Y K L I L N L K Q L K E E A I T E A V D A A I A E K Y I K L I A N A K T V E G V W T L K D E I L T F T V T E
BINARY_3845 T T Y K L I L N L K Q L K E E A I T E A V D A A I A E K Y I K L I A N A K T V E G V W T L K D E I K K A T V T E
BINARY_3846 T T Y K L I L N L K Q L K E E A I T E A V D A A I A E K Y I K L I A N A K T V E G V W T L K D E I K K F T V T E
BINARY_3847 T T Y K L I L N L K Q L K E E A I T E A V D A A I A E K Y I K L I A N A K T V E G V W T L K D E I K T A T V T E
BINARY_3848 T T Y K L I L N L K Q L K E E A I T E A V D A A I A E K Y I K L I A N A K T V E G V W T L K D E I K T F T V T E
BINARY_3849 T T Y K L I L N L K Q L K E E A I T E A V D A A I A E K Y I K L I A N A K T V E G V W T L K D E T L K A T V T E
BINARY_3850 T T Y K L I L N L K Q L K E E A I T E A V D A A I A E K Y I K L I A N A K T V E G V W T L K D E T L K F T V T E
BINARY_3851 T T Y K L I L N L K Q L K E E A I T E A V D A A I A E K Y I K L I A N A K T V E G V W T L K D E T L T A T V T E
BINARY_3852 T T Y K L I L N L K Q L K E E A I T E A V D A A I A E K Y I K L I A N A K T V E G V W T L K D E T L T F T V T E
BINARY_3853 T T Y K L I L N L K Q L K E E A I T E A V D A A I A E K Y I K L I A N A K T V E G V W T L K D E T K K A T V T E
BINARY_3854 T T Y K L I L N L K Q L K E E A I T E A V D A A I A E K Y I K L I A N A K T V E G V W T L K D E T K K F T V T E
BINARY_3855 T T Y K L I L N L K Q L K E E A I T E A V D A A I A E K Y I K L I A N A K T V E G V W T L K D E T K T A T V T E
BINARY_3856 T T Y K L I L N L K Q L K E E A I T E A V D A A I A E K Y I K L I A N A K T V E G V W T L K D E T K T F T V T E
BINARY_3857 T T Y K L I L N L K Q L K E E A I T E A V D A A I A E K Y I K L I A N A K T V E G V W T Y K D E I L K A T V T E
BINARY_3858 T T Y K L I L N L K Q L K E E A I T E A V D A A I A E K Y I K L I A N A K T V E G V W T Y K D E I L K F T V T E
BINARY_3859 T T Y K L I L N L K Q L K E E A I T E A V D A A I A E K Y I K L I A N A K T V E G V W T Y K D E I L T A T V T E
BINARY_3860 T T Y K L I L N L K Q L K E E A I T E A V D A A I A E K Y I K L I A N A K T V E G V W T Y K D E I L T F T V T E
BINARY_3861 T T Y K L I L N L K Q L K E E A I T E A V D A A I A E K Y I K L I A N A K T V E G V W T Y K D E I K K A T V T E
BINARY_3862 T T Y K L I L N L K Q L K E E A I T E A V D A A I A E K Y I K L I A N A K T V E G V W T Y K D E I K K F T V T E
BINARY_3863 T T Y K L I L N L K Q L K E E A I T E A V D A A I A E K Y I K L I A N A K T V E G V W T Y K D E I K T A T V T E
BINARY_3864 T T Y K L I L N L K Q L K E E A I T E A V D A A I A E K Y I K L I A N A K T V E G V W T Y K D E I K T F T V T E
BINARY_3865 T T Y K L I L N L K Q L K E E A I T E A V D A A I A E K Y I K L I A N A K T V E G V W T Y K D E T L K A T V T E
BINARY_3866 T T Y K L I L N L K Q L K E E A I T E A V D A A I A E K Y I K L I A N A K T V E G V W T Y K D E T L K F T V T E
BINARY_3867 T T Y K L I L N L K Q L K E E A I T E A V D A A I A E K Y I K L I A N A K T V E G V W T Y K D E T L T A T V T E
BINARY_3868 T T Y K L I L N L K Q L K E E A I T E A V D A A I A E K Y I K L I A N A K T V E G V W T Y K D E T L T F T V T E
BINARY_3869 T T Y K L I L N L K Q L K E E A I T E A V D A A I A E K Y I K L I A N A K T V E G V W T Y K D E T K K A T V T E
BINARY_3870 T T Y K L I L N L K Q L K E E A I T E A V D A A I A E K Y I K L I A N A K T V E G V W T Y K D E T K K F T V T E
BINARY_3871 T T Y K L I L N L K Q L K E E A I T E A V D A A I A E K Y I K L I A N A K T V E G V W T Y K D E T K T A T V T E
BINARY_3872 T T Y K L I L N L K Q L K E E A I T E A V D A A I A E K Y I K L I A N A K T V E G V W T Y K D E T K T F T V T E
BINARY_3873 T T Y K L I L N L K Q L K E E A I T E A V D A A I A E K Y I K L Y A N A K T V E G V W T L K D E I L K A T V T E
BINARY_3874 T T Y K L I L N L K Q L K E E A I T E A V D A A I A E K Y I K L Y A N A K T V E G V W T L K D E I L K F T V T E
BINARY_3875 T T Y K L I L N L K Q L K E E A I T E A V D A A I A E K Y I K L Y A N A K T V E G V W T L K D E I L T A T V T E
BINARY_3876 T T Y K L I L N L K Q L K E E A I T E A V D A A I A E K Y I K L Y A N A K T V E G V W T L K D E I L T F T V T E
BINARY_3877 T T Y K L I L N L K Q L K E E A I T E A V D A A I A E K Y I K L Y A N A K T V E G V W T L K D E I K K A T V T E
BINARY_3878 T T Y K L I L N L K Q L K E E A I T E A V D A A I A E K Y I K L Y A N A K T V E G V W T L K D E I K K F T V T E
BINARY_3879 T T Y K L I L N L K Q L K E E A I T E A V D A A I A E K Y I K L Y A N A K T V E G V W T L K D E I K T A T V T E
BINARY_3880 T T Y K L I L N L K Q L K E E A I T E A V D A A I A E K Y I K L Y A N A K T V E G V W T L K D E I K T F T V T E
BINARY_3881 T T Y K L I L N L K Q L K E E A I T E A V D A A I A E K Y I K L Y A N A K T V E G V W T L K D E T L K A T V T E
BINARY_3882 T T Y K L I L N L K Q L K E E A I T E A V D A A I A E K Y I K L Y A N A K T V E G V W T L K D E T L K F T V T E
BINARY_3883 T T Y K L I L N L K Q L K E E A I T E A V D A A I A E K Y I K L Y A N A K T V E G V W T L K D E T L T A T V T E
BINARY_3884 T T Y K L I L N L K Q L K E E A I T E A V D A A I A E K Y I K L Y A N A K T V E G V W T L K D E T L T F T V T E
BINARY_3885 T T Y K L I L N L K Q L K E E A I T E A V D A A I A E K Y I K L Y A N A K T V E G V W T L K D E T K K A T V T E
BINARY_3886 T T Y K L I L N L K Q L K E E A I T E A V D A A I A E K Y I K L Y A N A K T V E G V W T L K D E T K K F T V T E
BINARY_3887 T T Y K L I L N L K Q L K E E A I T E A V D A A I A E K Y I K L Y A N A K T V E G V W T L K D E T K T A T V T E
BINARY_3888 T T Y K L I L N L K Q L K E E A I T E A V D A A I A E K Y I K L Y A N A K T V E G V W T L K D E T K T F T V T E
BINARY_3889 T T Y K L I L N L K Q L K E E A I T E A V D A A I A E K Y I K L Y A N A K T V E G V W T Y K D E I L K A T V T E
BINARY_3890 T T Y K L I L N L K Q L K E E A I T E A V D A A I A E K Y I K L Y A N A K T V E G V W T Y K D E I L K F T V T E
BINARY_3891 T T Y K L I L N L K Q L K E E A I T E A V D A A I A E K Y I K L Y A N A K T V E G V W T Y K D E I L T A T V T E
BINARY_3892 T T Y K L I L N L K Q L K E E A I T E A V D A A I A E K Y I K L Y A N A K T V E G V W T Y K D E I L T F T V T E
BINARY_3893 T T Y K L I L N L K Q L K E E A I T E A V D A A I A E K Y I K L Y A N A K T V E G V W T Y K D E I K K A T V T E
BINARY_3894 T T Y K L I L N L K Q L K E E A I T E A V D A A I A E K Y I K L Y A N A K T V E G V W T Y K D E I K K F T V T E
BINARY_3895 T T Y K L I L N L K Q L K E E A I T E A V D A A I A E K Y I K L Y A N A K T V E G V W T Y K D E I K T A T V T E
BINARY_3896 T T Y K L I L N L K Q L K E E A I T E A V D A A I A E K Y I K L Y A N A K T V E G V W T Y K D E I K T F T V T E
BINARY_3897 T T Y K L I L N L K Q L K E E A I T E A V D A A I A E K Y I K L Y A N A K T V E G V W T Y K D E T L K A T V T E
BINARY_3898 T T Y K L I L N L K Q L K E E A I T E A V D A A I A E K Y I K L Y A N A K T V E G V W T Y K D E T L K F T V T E
BINARY_3899 T T Y K L I L N L K Q L K E E A I T E A V D A A I A E K Y I K L Y A N A K T V E G V W T Y K D E T L T A T V T E
BINARY_3900 T T Y K L I L N L K Q L K E E A I T E A V D A A I A E K Y I K L Y A N A K T V E G V W T Y K D E T L T F T V T E
BINARY_3901 T T Y K L I L N L K Q L K E E A I T E A V D A A I A E K Y I K L Y A N A K T V E G V W T Y K D E T K K A T V T E
BINARY_3902 T T Y K L I L N L K Q L K E E A I T E A V D A A I A E K Y I K L Y A N A K T V E G V W T Y K D E T K K F T V T E
BINARY_3903 T T Y K L I L N L K Q L K E E A I T E A V D A A I A E K Y I K L Y A N A K T V E G V W T Y K D E T K T A T V T E
BINARY_3904 T T Y K L I L N L K Q L K E E A I T E A V D A A I A E K Y I K L Y A N A K T V E G V W T Y K D E T K T F T V T E
BINARY_3905 T T Y K L I L N L K Q L K E E A I T E A V D A A I A E K Y F K L I A N A K T V E G V W T L K D E I L K A T V T E
BINARY_3906 T T Y K L I L N L K Q L K E E A I T E A V D A A I A E K Y F K L I A N A K T V E G V W T L K D E I L K F T V T E
BINARY_3907 T T Y K L I L N L K Q L K E E A I T E A V D A A I A E K Y F K L I A N A K T V E G V W T L K D E I L T A T V T E
BINARY_3908 T T Y K L I L N L K Q L K E E A I T E A V D A A I A E K Y F K L I A N A K T V E G V W T L K D E I L T F T V T E
BINARY_3909 T T Y K L I L N L K Q L K E E A I T E A V D A A I A E K Y F K L I A N A K T V E G V W T L K D E I K K A T V T E
BINARY_3910 T T Y K L I L N L K Q L K E E A I T E A V D A A I A E K Y F K L I A N A K T V E G V W T L K D E I K K F T V T E
BINARY_3911 T T Y K L I L N L K Q L K E E A I T E A V D A A I A E K Y F K L I A N A K T V E G V W T L K D E I K T A T V T E

BINARY_3912 T T Y K L I L N L K Q L K E E A I T E A V D A A I A E K Y F K L I A N A K T V E G V W T L K D E I K T F T V T E
BINARY_3913 T T Y K L I L N L K Q L K E E A I T E A V D A A I A E K Y F K L I A N A K T V E G V W T L K D E T L K A T V T E
BINARY_3914 T T Y K L I L N L K Q L K E E A I T E A V D A A I A E K Y F K L I A N A K T V E G V W T L K D E T L K F T V T E
BINARY_3915 T T Y K L I L N L K Q L K E E A I T E A V D A A I A E K Y F K L I A N A K T V E G V W T L K D E T L T A T V T E
BINARY_3916 T T Y K L I L N L K Q L K E E A I T E A V D A A I A E K Y F K L I A N A K T V E G V W T L K D E T L T F T V T E
BINARY_3917 T T Y K L I L N L K Q L K E E A I T E A V D A A I A E K Y F K L I A N A K T V E G V W T L K D E T K K A T V T E
BINARY_3918 T T Y K L I L N L K Q L K E E A I T E A V D A A I A E K Y F K L I A N A K T V E G V W T L K D E T K K F T V T E
BINARY_3919 T T Y K L I L N L K Q L K E E A I T E A V D A A I A E K Y F K L I A N A K T V E G V W T L K D E T K T A T V T E
BINARY_3920 T T Y K L I L N L K Q L K E E A I T E A V D A A I A E K Y F K L I A N A K T V E G V W T L K D E T K T F T V T E
BINARY_3921 T T Y K L I L N L K Q L K E E A I T E A V D A A I A E K Y F K L I A N A K T V E G V W T Y K D E I L K A T V T E
BINARY_3922 T T Y K L I L N L K Q L K E E A I T E A V D A A I A E K Y F K L I A N A K T V E G V W T Y K D E I L K F T V T E
BINARY_3923 T T Y K L I L N L K Q L K E E A I T E A V D A A I A E K Y F K L I A N A K T V E G V W T Y K D E I L T A T V T E
BINARY_3924 T T Y K L I L N L K Q L K E E A I T E A V D A A I A E K Y F K L I A N A K T V E G V W T Y K D E I L T F T V T E
BINARY_3925 T T Y K L I L N L K Q L K E E A I T E A V D A A I A E K Y F K L I A N A K T V E G V W T Y K D E I K K A T V T E
BINARY_3926 T T Y K L I L N L K Q L K E E A I T E A V D A A I A E K Y F K L I A N A K T V E G V W T Y K D E I K K F T V T E
BINARY_3927 T T Y K L I L N L K Q L K E E A I T E A V D A A I A E K Y F K L I A N A K T V E G V W T Y K D E I K T A T V T E
BINARY_3928 T T Y K L I L N L K Q L K E E A I T E A V D A A I A E K Y F K L I A N A K T V E G V W T Y K D E I K T F T V T E
BINARY_3929 T T Y K L I L N L K Q L K E E A I T E A V D A A I A E K Y F K L I A N A K T V E G V W T Y K D E T L K A T V T E
BINARY_3930 T T Y K L I L N L K Q L K E E A I T E A V D A A I A E K Y F K L I A N A K T V E G V W T Y K D E T L K F T V T E
BINARY_3931 T T Y K L I L N L K Q L K E E A I T E A V D A A I A E K Y F K L I A N A K T V E G V W T Y K D E T L T A T V T E
BINARY_3932 T T Y K L I L N L K Q L K E E A I T E A V D A A I A E K Y F K L I A N A K T V E G V W T Y K D E T L T F T V T E
BINARY_3933 T T Y K L I L N L K Q L K E E A I T E A V D A A I A E K Y F K L I A N A K T V E G V W T Y K D E T K K A T V T E
BINARY_3934 T T Y K L I L N L K Q L K E E A I T E A V D A A I A E K Y F K L I A N A K T V E G V W T Y K D E T K K F T V T E
BINARY_3935 T T Y K L I L N L K Q L K E E A I T E A V D A A I A E K Y F K L I A N A K T V E G V W T Y K D E T K T A T V T E
BINARY_3936 T T Y K L I L N L K Q L K E E A I T E A V D A A I A E K Y F K L I A N A K T V E G V W T Y K D E T K T F T V T E
BINARY_3937 T T Y K L I L N L K Q L K E E A I T E A V D A A I A E K Y F K L Y A N A K T V E G V W T L K D E I L K A T V T E
BINARY_3938 T T Y K L I L N L K Q L K E E A I T E A V D A A I A E K Y F K L Y A N A K T V E G V W T L K D E I L K F T V T E
BINARY_3939 T T Y K L I L N L K Q L K E E A I T E A V D A A I A E K Y F K L Y A N A K T V E G V W T L K D E I L T A T V T E
BINARY_3940 T T Y K L I L N L K Q L K E E A I T E A V D A A I A E K Y F K L Y A N A K T V E G V W T L K D E I L T F T V T E
BINARY_3941 T T Y K L I L N L K Q L K E E A I T E A V D A A I A E K Y F K L Y A N A K T V E G V W T L K D E I K K A T V T E
BINARY_3942 T T Y K L I L N L K Q L K E E A I T E A V D A A I A E K Y F K L Y A N A K T V E G V W T L K D E I K K F T V T E
BINARY_3943 T T Y K L I L N L K Q L K E E A I T E A V D A A I A E K Y F K L Y A N A K T V E G V W T L K D E I K T A T V T E
BINARY_3944 T T Y K L I L N L K Q L K E E A I T E A V D A A I A E K Y F K L Y A N A K T V E G V W T L K D E I K T F T V T E
BINARY_3945 T T Y K L I L N L K Q L K E E A I T E A V D A A I A E K Y F K L Y A N A K T V E G V W T L K D E T L K A T V T E
BINARY_3946 T T Y K L I L N L K Q L K E E A I T E A V D A A I A E K Y F K L Y A N A K T V E G V W T L K D E T L K F T V T E
BINARY_3947 T T Y K L I L N L K Q L K E E A I T E A V D A A I A E K Y F K L Y A N A K T V E G V W T L K D E T L T A T V T E
BINARY_3948 T T Y K L I L N L K Q L K E E A I T E A V D A A I A E K Y F K L Y A N A K T V E G V W T L K D E T L T F T V T E
BINARY_3949 T T Y K L I L N L K Q L K E E A I T E A V D A A I A E K Y F K L Y A N A K T V E G V W T L K D E T K K A T V T E
BINARY_3950 T T Y K L I L N L K Q L K E E A I T E A V D A A I A E K Y F K L Y A N A K T V E G V W T L K D E T K K F T V T E
BINARY_3951 T T Y K L I L N L K Q L K E E A I T E A V D A A I A E K Y F K L Y A N A K T V E G V W T L K D E T K T A T V T E
BINARY_3952 T T Y K L I L N L K Q L K E E A I T E A V D A A I A E K Y F K L Y A N A K T V E G V W T L K D E T K T F T V T E
BINARY_3953 T T Y K L I L N L K Q L K E E A I T E A V D A A I A E K Y F K L Y A N A K T V E G V W T Y K D E I L K A T V T E
BINARY_3954 T T Y K L I L N L K Q L K E E A I T E A V D A A I A E K Y F K L Y A N A K T V E G V W T Y K D E I L K F T V T E
BINARY_3955 T T Y K L I L N L K Q L K E E A I T E A V D A A I A E K Y F K L Y A N A K T V E G V W T Y K D E I L T A T V T E
BINARY_3956 T T Y K L I L N L K Q L K E E A I T E A V D A A I A E K Y F K L Y A N A K T V E G V W T Y K D E I L T F T V T E
BINARY_3957 T T Y K L I L N L K Q L K E E A I T E A V D A A I A E K Y F K L Y A N A K T V E G V W T Y K D E I K K A T V T E
BINARY_3958 T T Y K L I L N L K Q L K E E A I T E A V D A A I A E K Y F K L Y A N A K T V E G V W T Y K D E I K K F T V T E
BINARY_3959 T T Y K L I L N L K Q L K E E A I T E A V D A A I A E K Y F K L Y A N A K T V E G V W T Y K D E I K T A T V T E
BINARY_3960 T T Y K L I L N L K Q L K E E A I T E A V D A A I A E K Y F K L Y A N A K T V E G V W T Y K D E I K T F T V T E
BINARY_3961 T T Y K L I L N L K Q L K E E A I T E A V D A A I A E K Y F K L Y A N A K T V E G V W T Y K D E T L K A T V T E
BINARY_3962 T T Y K L I L N L K Q L K E E A I T E A V D A A I A E K Y F K L Y A N A K T V E G V W T Y K D E T L K F T V T E
BINARY_3963 T T Y K L I L N L K Q L K E E A I T E A V D A A I A E K Y F K L Y A N A K T V E G V W T Y K D E T L T A T V T E
BINARY_3964 T T Y K L I L N L K Q L K E E A I T E A V D A A I A E K Y F K L Y A N A K T V E G V W T Y K D E T L T F T V T E
BINARY_3965 T T Y K L I L N L K Q L K E E A I T E A V D A A I A E K Y F K L Y A N A K T V E G V W T Y K D E T K K A T V T E
BINARY_3966 T T Y K L I L N L K Q L K E E A I T E A V D A A I A E K Y F K L Y A N A K T V E G V W T Y K D E T K K F T V T E
BINARY_3967 T T Y K L I L N L K Q L K E E A I T E A V D A A I A E K Y F K L Y A N A K T V E G V W T Y K D E T K T A T V T E
BINARY_3968 T T Y K L I L N L K Q L K E E A I T E A V D A A I A E K Y F K L Y A N A K T V E G V W T Y K D E T K T F T V T E
BINARY_3969 T T Y K L I L N L K Q L K E E A I T E A V D A A T A E K Y I K L I A N A K T V E G V W T L K D E I L K A T V T E
BINARY_3970 T T Y K L I L N L K Q L K E E A I T E A V D A A T A E K Y I K L I A N A K T V E G V W T L K D E I L K F T V T E
BINARY_3971 T T Y K L I L N L K Q L K E E A I T E A V D A A T A E K Y I K L I A N A K T V E G V W T L K D E I L T A T V T E
BINARY_3972 T T Y K L I L N L K Q L K E E A I T E A V D A A T A E K Y I K L I A N A K T V E G V W T L K D E I L T F T V T E
BINARY_3973 T T Y K L I L N L K Q L K E E A I T E A V D A A T A E K Y I K L I A N A K T V E G V W T L K D E I K K A T V T E
BINARY_3974 T T Y K L I L N L K Q L K E E A I T E A V D A A T A E K Y I K L I A N A K T V E G V W T L K D E I K K F T V T E
BINARY_3975 T T Y K L I L N L K Q L K E E A I T E A V D A A T A E K Y I K L I A N A K T V E G V W T L K D E I K T A T V T E
BINARY_3976 T T Y K L I L N L K Q L K E E A I T E A V D A A T A E K Y I K L I A N A K T V E G V W T L K D E I K T F T V T E
BINARY_3977 T T Y K L I L N L K Q L K E E A I T E A V D A A T A E K Y I K L I A N A K T V E G V W T L K D E T L K A T V T E
BINARY_3978 T T Y K L I L N L K Q L K E E A I T E A V D A A T A E K Y I K L I A N A K T V E G V W T L K D E T L K F T V T E
BINARY_3979 T T Y K L I L N L K Q L K E E A I T E A V D A A T A E K Y I K L I A N A K T V E G V W T L K D E T L T A T V T E
BINARY_3980 T T Y K L I L N L K Q L K E E A I T E A V D A A T A E K Y I K L I A N A K T V E G V W T L K D E T L T F T V T E
BINARY_3981 T T Y K L I L N L K Q L K E E A I T E A V D A A T A E K Y I K L I A N A K T V E G V W T L K D E T K K A T V T E
BINARY_3982 T T Y K L I L N L K Q L K E E A I T E A V D A A T A E K Y I K L I A N A K T V E G V W T L K D E T K K F T V T E
BINARY_3983 T T Y K L I L N L K Q L K E E A I T E A V D A A T A E K Y I K L I A N A K T V E G V W T L K D E T K T A T V T E
BINARY_3984 T T Y K L I L N L K Q L K E E A I T E A V D A A T A E K Y I K L I A N A K T V E G V W T L K D E T K T F T V T E
BINARY_3985 T T Y K L I L N L K Q L K E E A I T E A V D A A T A E K Y I K L I A N A K T V E G V W T Y K D E I L K A T V T E
BINARY_3986 T T Y K L I L N L K Q L K E E A I T E A V D A A T A E K Y I K L I A N A K T V E G V W T Y K D E I L K F T V T E
BINARY_3987 T T Y K L I L N L K Q L K E E A I T E A V D A A T A E K Y I K L I A N A K T V E G V W T Y K D E I L T A T V T E
BINARY_3988 T T Y K L I L N L K Q L K E E A I T E A V D A A T A E K Y I K L I A N A K T V E G V W T Y K D E I L T F T V T E
BINARY_3989 T T Y K L I L N L K Q L K E E A I T E A V D A A T A E K Y I K L I A N A K T V E G V W T Y K D E I K K A T V T E
BINARY_3990 T T Y K L I L N L K Q L K E E A I T E A V D A A T A E K Y I K L I A N A K T V E G V W T Y K D E I K K F T V T E
BINARY_3991 T T Y K L I L N L K Q L K E E A I T E A V D A A T A E K Y I K L I A N A K T V E G V W T Y K D E I K T A T V T E
BINARY_3992 T T Y K L I L N L K Q L K E E A I T E A V D A A T A E K Y I K L I A N A K T V E G V W T Y K D E I K T F T V T E
BINARY_3993 T T Y K L I L N L K Q L K E E A I T E A V D A A T A E K Y I K L I A N A K T V E G V W T Y K D E T L K A T V T E
BINARY_3994 T T Y K L I L N L K Q L K E E A I T E A V D A A T A E K Y I K L I A N A K T V E G V W T Y K D E T L K F T V T E
BINARY_3995 T T Y K L I L N L K Q L K E E A I T E A V D A A T A E K Y I K L I A N A K T V E G V W T Y K D E T L T A T V T E
BINARY_3996 T T Y K L I L N L K Q L K E E A I T E A V D A A T A E K Y I K L I A N A K T V E G V W T Y K D E T L T F T V T E
BINARY_3997 T T Y K L I L N L K Q L K E E A I T E A V D A A T A E K Y I K L I A N A K T V E G V W T Y K D E T K K A T V T E
BINARY_3998 T T Y K L I L N L K Q L K E E A I T E A V D A A T A E K Y I K L I A N A K T V E G V W T Y K D E T K K F T V T E
BINARY_3999 T T Y K L I L N L K Q L K E E A I T E A V D A A T A E K Y I K L I A N A K T V E G V W T Y K D E T K T A T V T E
BINARY_4000 T T Y K L I L N L K Q L K E E A I T E A V D A A T A E K Y I K L I A N A K T V E G V W T Y K D E T K T F T V T E
BINARY_4001 T T Y K L I L N L K Q L K E E A I T E A V D A A T A E K Y I K L Y A N A K T V E G V W T L K D E I L K A T V T E
BINARY_4002 T T Y K L I L N L K Q L K E E A I T E A V D A A T A E K Y I K L Y A N A K T V E G V W T L K D E I L K F T V T E
BINARY_4003 T T Y K L I L N L K Q L K E E A I T E A V D A A T A E K Y I K L Y A N A K T V E G V W T L K D E I L T A T V T E
BINARY_4004 T T Y K L I L N L K Q L K E E A I T E A V D A A T A E K Y I K L Y A N A K T V E G V W T L K D E I L T F T V T E
BINARY_4005 T T Y K L I L N L K Q L K E E A I T E A V D A A T A E K Y I K L Y A N A K T V E G V W T L K D E I K K A T V T E
BINARY_4006 T T Y K L I L N L K Q L K E E A I T E A V D A A T A E K Y I K L Y A N A K T V E G V W T L K D E I K K F T V T E

BINARY_4007 T T Y K L I L N L K Q L K E E A I T E A V D A A T A E K Y I K L Y A N A K T V E G V W T L K D E I K T A T V T E
BINARY_4008 T T Y K L I L N L K Q L K E E A I T E A V D A A T A E K Y I K L Y A N A K T V E G V W T L K D E I K T F T V T E
BINARY_4009 T T Y K L I L N L K Q L K E E A I T E A V D A A T A E K Y I K L Y A N A K T V E G V W T L K D E T L K A T V T E
BINARY_4010 T T Y K L I L N L K Q L K E E A I T E A V D A A T A E K Y I K L Y A N A K T V E G V W T L K D E T L K F T V T E
BINARY_4011 T T Y K L I L N L K Q L K E E A I T E A V D A A T A E K Y I K L Y A N A K T V E G V W T L K D E T L T A T V T E
BINARY_4012 T T Y K L I L N L K Q L K E E A I T E A V D A A T A E K Y I K L Y A N A K T V E G V W T L K D E T L T F T V T E
BINARY_4013 T T Y K L I L N L K Q L K E E A I T E A V D A A T A E K Y I K L Y A N A K T V E G V W T L K D E T K K A T V T E
BINARY_4014 T T Y K L I L N L K Q L K E E A I T E A V D A A T A E K Y I K L Y A N A K T V E G V W T L K D E T K K F T V T E
BINARY_4015 T T Y K L I L N L K Q L K E E A I T E A V D A A T A E K Y I K L Y A N A K T V E G V W T L K D E T K T A T V T E
BINARY_4016 T T Y K L I L N L K Q L K E E A I T E A V D A A T A E K Y I K L Y A N A K T V E G V W T L K D E T K T F T V T E
BINARY_4017 T T Y K L I L N L K Q L K E E A I T E A V D A A T A E K Y I K L Y A N A K T V E G V W T Y K D E I L K A T V T E
BINARY_4018 T T Y K L I L N L K Q L K E E A I T E A V D A A T A E K Y I K L Y A N A K T V E G V W T Y K D E I L K F T V T E
BINARY_4019 T T Y K L I L N L K Q L K E E A I T E A V D A A T A E K Y I K L Y A N A K T V E G V W T Y K D E I L T A T V T E
BINARY_4020 T T Y K L I L N L K Q L K E E A I T E A V D A A T A E K Y I K L Y A N A K T V E G V W T Y K D E I L T F T V T E
BINARY_4021 T T Y K L I L N L K Q L K E E A I T E A V D A A T A E K Y I K L Y A N A K T V E G V W T Y K D E I K K A T V T E
BINARY_4022 T T Y K L I L N L K Q L K E E A I T E A V D A A T A E K Y I K L Y A N A K T V E G V W T Y K D E I K K F T V T E
BINARY_4023 T T Y K L I L N L K Q L K E E A I T E A V D A A T A E K Y I K L Y A N A K T V E G V W T Y K D E I K T A T V T E
BINARY_4024 T T Y K L I L N L K Q L K E E A I T E A V D A A T A E K Y I K L Y A N A K T V E G V W T Y K D E I K T F T V T E
BINARY_4025 T T Y K L I L N L K Q L K E E A I T E A V D A A T A E K Y I K L Y A N A K T V E G V W T Y K D E T L K A T V T E
BINARY_4026 T T Y K L I L N L K Q L K E E A I T E A V D A A T A E K Y I K L Y A N A K T V E G V W T Y K D E T L K F T V T E
BINARY_4027 T T Y K L I L N L K Q L K E E A I T E A V D A A T A E K Y I K L Y A N A K T V E G V W T Y K D E T L T A T V T E
BINARY_4028 T T Y K L I L N L K Q L K E E A I T E A V D A A T A E K Y I K L Y A N A K T V E G V W T Y K D E T L T F T V T E
BINARY_4029 T T Y K L I L N L K Q L K E E A I T E A V D A A T A E K Y I K L Y A N A K T V E G V W T Y K D E T K K A T V T E
BINARY_4030 T T Y K L I L N L K Q L K E E A I T E A V D A A T A E K Y I K L Y A N A K T V E G V W T Y K D E T K K F T V T E
BINARY_4031 T T Y K L I L N L K Q L K E E A I T E A V D A A T A E K Y I K L Y A N A K T V E G V W T Y K D E T K T A T V T E
BINARY_4032 T T Y K L I L N L K Q L K E E A I T E A V D A A T A E K Y I K L Y A N A K T V E G V W T Y K D E T K T F T V T E
BINARY_4033 T T Y K L I L N L K Q L K E E A I T E A V D A A T A E K Y F K L I A N A K T V E G V W T L K D E I L K A T V T E
BINARY_4034 T T Y K L I L N L K Q L K E E A I T E A V D A A T A E K Y F K L I A N A K T V E G V W T L K D E I L K F T V T E
BINARY_4035 T T Y K L I L N L K Q L K E E A I T E A V D A A T A E K Y F K L I A N A K T V E G V W T L K D E I L T A T V T E
BINARY_4036 T T Y K L I L N L K Q L K E E A I T E A V D A A T A E K Y F K L I A N A K T V E G V W T L K D E I L T F T V T E
BINARY_4037 T T Y K L I L N L K Q L K E E A I T E A V D A A T A E K Y F K L I A N A K T V E G V W T L K D E I K K A T V T E
BINARY_4038 T T Y K L I L N L K Q L K E E A I T E A V D A A T A E K Y F K L I A N A K T V E G V W T L K D E I K K F T V T E
BINARY_4039 T T Y K L I L N L K Q L K E E A I T E A V D A A T A E K Y F K L I A N A K T V E G V W T L K D E I K T A T V T E
BINARY_4040 T T Y K L I L N L K Q L K E E A I T E A V D A A T A E K Y F K L I A N A K T V E G V W T L K D E I K T F T V T E
BINARY_4041 T T Y K L I L N L K Q L K E E A I T E A V D A A T A E K Y F K L I A N A K T V E G V W T L K D E T L K A T V T E
BINARY_4042 T T Y K L I L N L K Q L K E E A I T E A V D A A T A E K Y F K L I A N A K T V E G V W T L K D E T L K F T V T E
BINARY_4043 T T Y K L I L N L K Q L K E E A I T E A V D A A T A E K Y F K L I A N A K T V E G V W T L K D E T L T A T V T E
BINARY_4044 T T Y K L I L N L K Q L K E E A I T E A V D A A T A E K Y F K L I A N A K T V E G V W T L K D E T L T F T V T E
BINARY_4045 T T Y K L I L N L K Q L K E E A I T E A V D A A T A E K Y F K L I A N A K T V E G V W T L K D E T K K A T V T E
BINARY_4046 T T Y K L I L N L K Q L K E E A I T E A V D A A T A E K Y F K L I A N A K T V E G V W T L K D E T K K F T V T E
BINARY_4047 T T Y K L I L N L K Q L K E E A I T E A V D A A T A E K Y F K L I A N A K T V E G V W T L K D E T K T A T V T E
BINARY_4048 T T Y K L I L N L K Q L K E E A I T E A V D A A T A E K Y F K L I A N A K T V E G V W T L K D E T K T F T V T E
BINARY_4049 T T Y K L I L N L K Q L K E E A I T E A V D A A T A E K Y F K L I A N A K T V E G V W T Y K D E I L K A T V T E
BINARY_4050 T T Y K L I L N L K Q L K E E A I T E A V D A A T A E K Y F K L I A N A K T V E G V W T Y K D E I L K F T V T E
BINARY_4051 T T Y K L I L N L K Q L K E E A I T E A V D A A T A E K Y F K L I A N A K T V E G V W T Y K D E I L T A T V T E
BINARY_4052 T T Y K L I L N L K Q L K E E A I T E A V D A A T A E K Y F K L I A N A K T V E G V W T Y K D E I L T F T V T E
BINARY_4053 T T Y K L I L N L K Q L K E E A I T E A V D A A T A E K Y F K L I A N A K T V E G V W T Y K D E I K K A T V T E
BINARY_4054 T T Y K L I L N L K Q L K E E A I T E A V D A A T A E K Y F K L I A N A K T V E G V W T Y K D E I K K F T V T E
BINARY_4055 T T Y K L I L N L K Q L K E E A I T E A V D A A T A E K Y F K L I A N A K T V E G V W T Y K D E I K T A T V T E
BINARY_4056 T T Y K L I L N L K Q L K E E A I T E A V D A A T A E K Y F K L I A N A K T V E G V W T Y K D E I K T F T V T E
BINARY_4057 T T Y K L I L N L K Q L K E E A I T E A V D A A T A E K Y F K L I A N A K T V E G V W T Y K D E T L K A T V T E
BINARY_4058 T T Y K L I L N L K Q L K E E A I T E A V D A A T A E K Y F K L I A N A K T V E G V W T Y K D E T L K F T V T E
BINARY_4059 T T Y K L I L N L K Q L K E E A I T E A V D A A T A E K Y F K L I A N A K T V E G V W T Y K D E T L T A T V T E
BINARY_4060 T T Y K L I L N L K Q L K E E A I T E A V D A A T A E K Y F K L I A N A K T V E G V W T Y K D E T L T F T V T E
BINARY_4061 T T Y K L I L N L K Q L K E E A I T E A V D A A T A E K Y F K L I A N A K T V E G V W T Y K D E T K K A T V T E
BINARY_4062 T T Y K L I L N L K Q L K E E A I T E A V D A A T A E K Y F K L I A N A K T V E G V W T Y K D E T K K F T V T E
BINARY_4063 T T Y K L I L N L K Q L K E E A I T E A V D A A T A E K Y F K L I A N A K T V E G V W T Y K D E T K T A T V T E
BINARY_4064 T T Y K L I L N L K Q L K E E A I T E A V D A A T A E K Y F K L I A N A K T V E G V W T Y K D E T K T F T V T E
BINARY_4065 T T Y K L I L N L K Q L K E E A I T E A V D A A T A E K Y F K L Y A N A K T V E G V W T L K D E I L K A T V T E
BINARY_4066 T T Y K L I L N L K Q L K E E A I T E A V D A A T A E K Y F K L Y A N A K T V E G V W T L K D E I L K F T V T E
BINARY_4067 T T Y K L I L N L K Q L K E E A I T E A V D A A T A E K Y F K L Y A N A K T V E G V W T L K D E I L T A T V T E
BINARY_4068 T T Y K L I L N L K Q L K E E A I T E A V D A A T A E K Y F K L Y A N A K T V E G V W T L K D E I L T F T V T E
BINARY_4069 T T Y K L I L N L K Q L K E E A I T E A V D A A T A E K Y F K L Y A N A K T V E G V W T L K D E I K K A T V T E
BINARY_4070 T T Y K L I L N L K Q L K E E A I T E A V D A A T A E K Y F K L Y A N A K T V E G V W T L K D E I K K F T V T E
BINARY_4071 T T Y K L I L N L K Q L K E E A I T E A V D A A T A E K Y F K L Y A N A K T V E G V W T L K D E I K T A T V T E
BINARY_4072 T T Y K L I L N L K Q L K E E A I T E A V D A A T A E K Y F K L Y A N A K T V E G V W T L K D E I K T F T V T E
BINARY_4073 T T Y K L I L N L K Q L K E E A I T E A V D A A T A E K Y F K L Y A N A K T V E G V W T L K D E T L K A T V T E
BINARY_4074 T T Y K L I L N L K Q L K E E A I T E A V D A A T A E K Y F K L Y A N A K T V E G V W T L K D E T L K F T V T E
BINARY_4075 T T Y K L I L N L K Q L K E E A I T E A V D A A T A E K Y F K L Y A N A K T V E G V W T L K D E T L T A T V T E
BINARY_4076 T T Y K L I L N L K Q L K E E A I T E A V D A A T A E K Y F K L Y A N A K T V E G V W T L K D E T L T F T V T E
BINARY_4077 T T Y K L I L N L K Q L K E E A I T E A V D A A T A E K Y F K L Y A N A K T V E G V W T L K D E T K K A T V T E
BINARY_4078 T T Y K L I L N L K Q L K E E A I T E A V D A A T A E K Y F K L Y A N A K T V E G V W T L K D E T K K F T V T E
BINARY_4079 T T Y K L I L N L K Q L K E E A I T E A V D A A T A E K Y F K L Y A N A K T V E G V W T L K D E T K T A T V T E
BINARY_4080 T T Y K L I L N L K Q L K E E A I T E A V D A A T A E K Y F K L Y A N A K T V E G V W T L K D E T K T F T V T E
BINARY_4081 T T Y K L I L N L K Q L K E E A I T E A V D A A T A E K Y F K L Y A N A K T V E G V W T Y K D E I L K A T V T E
BINARY_4082 T T Y K L I L N L K Q L K E E A I T E A V D A A T A E K Y F K L Y A N A K T V E G V W T Y K D E I L K F T V T E
BINARY_4083 T T Y K L I L N L K Q L K E E A I T E A V D A A T A E K Y F K L Y A N A K T V E G V W T Y K D E I L T A T V T E
BINARY_4084 T T Y K L I L N L K Q L K E E A I T E A V D A A T A E K Y F K L Y A N A K T V E G V W T Y K D E I L T F T V T E
BINARY_4085 T T Y K L I L N L K Q L K E E A I T E A V D A A T A E K Y F K L Y A N A K T V E G V W T Y K D E I K K A T V T E
BINARY_4086 T T Y K L I L N L K Q L K E E A I T E A V D A A T A E K Y F K L Y A N A K T V E G V W T Y K D E I K K F T V T E
BINARY_4087 T T Y K L I L N L K Q L K E E A I T E A V D A A T A E K Y F K L Y A N A K T V E G V W T Y K D E I K T A T V T E
BINARY_4088 T T Y K L I L N L K Q L K E E A I T E A V D A A T A E K Y F K L Y A N A K T V E G V W T Y K D E I K T F T V T E
BINARY_4089 T T Y K L I L N L K Q L K E E A I T E A V D A A T A E K Y F K L Y A N A K T V E G V W T Y K D E T L K A T V T E
BINARY_4090 T T Y K L I L N L K Q L K E E A I T E A V D A A T A E K Y F K L Y A N A K T V E G V W T Y K D E T L K F T V T E
BINARY_4091 T T Y K L I L N L K Q L K E E A I T E A V D A A T A E K Y F K L Y A N A K T V E G V W T Y K D E T L T A T V T E
BINARY_4092 T T Y K L I L N L K Q L K E E A I T E A V D A A T A E K Y F K L Y A N A K T V E G V W T Y K D E T L T F T V T E
BINARY_4093 T T Y K L I L N L K Q L K E E A I T E A V D A A T A E K Y F K L Y A N A K T V E G V W T Y K D E T K K A T V T E
BINARY_4094 T T Y K L I L N L K Q L K E E A I T E A V D A A T A E K Y F K L Y A N A K T V E G V W T Y K D E T K K F T V T E
BINARY_4095 T T Y K L I L N L K Q L K E E A I T E A V D A A T A E K Y F K L Y A N A K T V E G V W T Y K D E T K T A T V T E
BINARY_4096 T T Y K L I L N L K Q L K E E A I T E A V D A A T A E K Y F K L Y A N A K T V E G V W T Y K D E T K T F T V T E
BINARY_4097 T T Y K L I L N G K Q A K E E A I K E L V D A G I A E K Y I K L I A N A K T V E G V W T L K D E I L K A T V T E
BINARY_4098 T T Y K L I L N G K Q A K E E A I K E L V D A G I A E K Y I K L I A N A K T V E G V W T L K D E I L K F T V T E
BINARY_4099 T T Y K L I L N G K Q A K E E A I K E L V D A G I A E K Y I K L I A N A K T V E G V W T L K D E I L T A T V T E
BINARY_4100 T T Y K L I L N G K Q A K E E A I K E L V D A G I A E K Y I K L I A N A K T V E G V W T L K D E I L T F T V T E
BINARY_4101 T T Y K L I L N G K Q A K E E A I K E L V D A G I A E K Y I K L I A N A K T V E G V W T L K D E I K K A T V T E

BINARY_4102 T T Y K L I L N G K Q A K E E A I K E L V D A G I A E K Y I K L I A N A K T V E G V W T L K D E I K K F T V T E
BINARY_4103 T T Y K L I L N G K Q A K E E A I K E L V D A G I A E K Y I K L I A N A K T V E G V W T L K D E I K T A T V T E
BINARY_4104 T T Y K L I L N G K Q A K E E A I K E L V D A G I A E K Y I K L I A N A K T V E G V W T L K D E I K T F T V T E
BINARY_4105 T T Y K L I L N G K Q A K E E A I K E L V D A G I A E K Y I K L I A N A K T V E G V W T L K D E T L K A T V T E
BINARY_4106 T T Y K L I L N G K Q A K E E A I K E L V D A G I A E K Y I K L I A N A K T V E G V W T L K D E T L K F T V T E
BINARY_4107 T T Y K L I L N G K Q A K E E A I K E L V D A G I A E K Y I K L I A N A K T V E G V W T L K D E T L T A T V T E
BINARY_4108 T T Y K L I L N G K Q A K E E A I K E L V D A G I A E K Y I K L I A N A K T V E G V W T L K D E T L T F T V T E
BINARY_4109 T T Y K L I L N G K Q A K E E A I K E L V D A G I A E K Y I K L I A N A K T V E G V W T L K D E T K K A T V T E
BINARY_4110 T T Y K L I L N G K Q A K E E A I K E L V D A G I A E K Y I K L I A N A K T V E G V W T L K D E T K K F T V T E
BINARY_4111 T T Y K L I L N G K Q A K E E A I K E L V D A G I A E K Y I K L I A N A K T V E G V W T L K D E T K T A T V T E
BINARY_4112 T T Y K L I L N G K Q A K E E A I K E L V D A G I A E K Y I K L I A N A K T V E G V W T L K D E T K T F T V T E
BINARY_4113 T T Y K L I L N G K Q A K E E A I K E L V D A G I A E K Y I K L I A N A K T V E G V W T Y K D E I L K A T V T E
BINARY_4114 T T Y K L I L N G K Q A K E E A I K E L V D A G I A E K Y I K L I A N A K T V E G V W T Y K D E I L K F T V T E
BINARY_4115 T T Y K L I L N G K Q A K E E A I K E L V D A G I A E K Y I K L I A N A K T V E G V W T Y K D E I L T A T V T E
BINARY_4116 T T Y K L I L N G K Q A K E E A I K E L V D A G I A E K Y I K L I A N A K T V E G V W T Y K D E I L T F T V T E
BINARY_4117 T T Y K L I L N G K Q A K E E A I K E L V D A G I A E K Y I K L I A N A K T V E G V W T Y K D E I K K A T V T E
BINARY_4118 T T Y K L I L N G K Q A K E E A I K E L V D A G I A E K Y I K L I A N A K T V E G V W T Y K D E I K K F T V T E
BINARY_4119 T T Y K L I L N G K Q A K E E A I K E L V D A G I A E K Y I K L I A N A K T V E G V W T Y K D E I K T A T V T E
BINARY_4120 T T Y K L I L N G K Q A K E E A I K E L V D A G I A E K Y I K L I A N A K T V E G V W T Y K D E I K T F T V T E
BINARY_4121 T T Y K L I L N G K Q A K E E A I K E L V D A G I A E K Y I K L I A N A K T V E G V W T Y K D E T L K A T V T E
BINARY_4122 T T Y K L I L N G K Q A K E E A I K E L V D A G I A E K Y I K L I A N A K T V E G V W T Y K D E T L K F T V T E
BINARY_4123 T T Y K L I L N G K Q A K E E A I K E L V D A G I A E K Y I K L I A N A K T V E G V W T Y K D E T L T A T V T E
BINARY_4124 T T Y K L I L N G K Q A K E E A I K E L V D A G I A E K Y I K L I A N A K T V E G V W T Y K D E T L T F T V T E
BINARY_4125 T T Y K L I L N G K Q A K E E A I K E L V D A G I A E K Y I K L I A N A K T V E G V W T Y K D E T K K A T V T E
BINARY_4126 T T Y K L I L N G K Q A K E E A I K E L V D A G I A E K Y I K L I A N A K T V E G V W T Y K D E T K K F T V T E
BINARY_4127 T T Y K L I L N G K Q A K E E A I K E L V D A G I A E K Y I K L I A N A K T V E G V W T Y K D E T K T A T V T E
BINARY_4128 T T Y K L I L N G K Q A K E E A I K E L V D A G I A E K Y I K L I A N A K T V E G V W T Y K D E T K T F T V T E
BINARY_4129 T T Y K L I L N G K Q A K E E A I K E L V D A G I A E K Y I K L Y A N A K T V E G V W T L K D E I L K A T V T E
BINARY_4130 T T Y K L I L N G K Q A K E E A I K E L V D A G I A E K Y I K L Y A N A K T V E G V W T L K D E I L K F T V T E
BINARY_4131 T T Y K L I L N G K Q A K E E A I K E L V D A G I A E K Y I K L Y A N A K T V E G V W T L K D E I L T A T V T E
BINARY_4132 T T Y K L I L N G K Q A K E E A I K E L V D A G I A E K Y I K L Y A N A K T V E G V W T L K D E I L T F T V T E
BINARY_4133 T T Y K L I L N G K Q A K E E A I K E L V D A G I A E K Y I K L Y A N A K T V E G V W T L K D E I K K A T V T E
BINARY_4134 T T Y K L I L N G K Q A K E E A I K E L V D A G I A E K Y I K L Y A N A K T V E G V W T L K D E I K K F T V T E
BINARY_4135 T T Y K L I L N G K Q A K E E A I K E L V D A G I A E K Y I K L Y A N A K T V E G V W T L K D E I K T A T V T E
BINARY_4136 T T Y K L I L N G K Q A K E E A I K E L V D A G I A E K Y I K L Y A N A K T V E G V W T L K D E I K T F T V T E
BINARY_4137 T T Y K L I L N G K Q A K E E A I K E L V D A G I A E K Y I K L Y A N A K T V E G V W T L K D E T L K A T V T E
BINARY_4138 T T Y K L I L N G K Q A K E E A I K E L V D A G I A E K Y I K L Y A N A K T V E G V W T L K D E T L K F T V T E
BINARY_4139 T T Y K L I L N G K Q A K E E A I K E L V D A G I A E K Y I K L Y A N A K T V E G V W T L K D E T L T A T V T E
BINARY_4140 T T Y K L I L N G K Q A K E E A I K E L V D A G I A E K Y I K L Y A N A K T V E G V W T L K D E T L T F T V T E
BINARY_4141 T T Y K L I L N G K Q A K E E A I K E L V D A G I A E K Y I K L Y A N A K T V E G V W T L K D E T K K A T V T E
BINARY_4142 T T Y K L I L N G K Q A K E E A I K E L V D A G I A E K Y I K L Y A N A K T V E G V W T L K D E T K K F T V T E
BINARY_4143 T T Y K L I L N G K Q A K E E A I K E L V D A G I A E K Y I K L Y A N A K T V E G V W T L K D E T K T A T V T E
BINARY_4144 T T Y K L I L N G K Q A K E E A I K E L V D A G I A E K Y I K L Y A N A K T V E G V W T L K D E T K T F T V T E
BINARY_4145 T T Y K L I L N G K Q A K E E A I K E L V D A G I A E K Y I K L Y A N A K T V E G V W T Y K D E I L K A T V T E
BINARY_4146 T T Y K L I L N G K Q A K E E A I K E L V D A G I A E K Y I K L Y A N A K T V E G V W T Y K D E I L K F T V T E
BINARY_4147 T T Y K L I L N G K Q A K E E A I K E L V D A G I A E K Y I K L Y A N A K T V E G V W T Y K D E I L T A T V T E
BINARY_4148 T T Y K L I L N G K Q A K E E A I K E L V D A G I A E K Y I K L Y A N A K T V E G V W T Y K D E I L T F T V T E
BINARY_4149 T T Y K L I L N G K Q A K E E A I K E L V D A G I A E K Y I K L Y A N A K T V E G V W T Y K D E I K K A T V T E
BINARY_4150 T T Y K L I L N G K Q A K E E A I K E L V D A G I A E K Y I K L Y A N A K T V E G V W T Y K D E I K K F T V T E
BINARY_4151 T T Y K L I L N G K Q A K E E A I K E L V D A G I A E K Y I K L Y A N A K T V E G V W T Y K D E I K T A T V T E
BINARY_4152 T T Y K L I L N G K Q A K E E A I K E L V D A G I A E K Y I K L Y A N A K T V E G V W T Y K D E I K T F T V T E
BINARY_4153 T T Y K L I L N G K Q A K E E A I K E L V D A G I A E K Y I K L Y A N A K T V E G V W T Y K D E T L K A T V T E
BINARY_4154 T T Y K L I L N G K Q A K E E A I K E L V D A G I A E K Y I K L Y A N A K T V E G V W T Y K D E T L K F T V T E
BINARY_4155 T T Y K L I L N G K Q A K E E A I K E L V D A G I A E K Y I K L Y A N A K T V E G V W T Y K D E T L T A T V T E
BINARY_4156 T T Y K L I L N G K Q A K E E A I K E L V D A G I A E K Y I K L Y A N A K T V E G V W T Y K D E T L T F T V T E
BINARY_4157 T T Y K L I L N G K Q A K E E A I K E L V D A G I A E K Y I K L Y A N A K T V E G V W T Y K D E T K K A T V T E
BINARY_4158 T T Y K L I L N G K Q A K E E A I K E L V D A G I A E K Y I K L Y A N A K T V E G V W T Y K D E T K K F T V T E
BINARY_4159 T T Y K L I L N G K Q A K E E A I K E L V D A G I A E K Y I K L Y A N A K T V E G V W T Y K D E T K T A T V T E
BINARY_4160 T T Y K L I L N G K Q A K E E A I K E L V D A G I A E K Y I K L Y A N A K T V E G V W T Y K D E T K T F T V T E
BINARY_4161 T T Y K L I L N G K Q A K E E A I K E L V D A G I A E K Y F K L I A N A K T V E G V W T L K D E I L K A T V T E
BINARY_4162 T T Y K L I L N G K Q A K E E A I K E L V D A G I A E K Y F K L I A N A K T V E G V W T L K D E I L K F T V T E
BINARY_4163 T T Y K L I L N G K Q A K E E A I K E L V D A G I A E K Y F K L I A N A K T V E G V W T L K D E I L T A T V T E
BINARY_4164 T T Y K L I L N G K Q A K E E A I K E L V D A G I A E K Y F K L I A N A K T V E G V W T L K D E I L T F T V T E
BINARY_4165 T T Y K L I L N G K Q A K E E A I K E L V D A G I A E K Y F K L I A N A K T V E G V W T L K D E I K K A T V T E
BINARY_4166 T T Y K L I L N G K Q A K E E A I K E L V D A G I A E K Y F K L I A N A K T V E G V W T L K D E I K K F T V T E
BINARY_4167 T T Y K L I L N G K Q A K E E A I K E L V D A G I A E K Y F K L I A N A K T V E G V W T L K D E I K T A T V T E
BINARY_4168 T T Y K L I L N G K Q A K E E A I K E L V D A G I A E K Y F K L I A N A K T V E G V W T L K D E I K T F T V T E
BINARY_4169 T T Y K L I L N G K Q A K E E A I K E L V D A G I A E K Y F K L I A N A K T V E G V W T L K D E T L K A T V T E
BINARY_4170 T T Y K L I L N G K Q A K E E A I K E L V D A G I A E K Y F K L I A N A K T V E G V W T L K D E T L K F T V T E
BINARY_4171 T T Y K L I L N G K Q A K E E A I K E L V D A G I A E K Y F K L I A N A K T V E G V W T L K D E T L T A T V T E
BINARY_4172 T T Y K L I L N G K Q A K E E A I K E L V D A G I A E K Y F K L I A N A K T V E G V W T L K D E T L T F T V T E
BINARY_4173 T T Y K L I L N G K Q A K E E A I K E L V D A G I A E K Y F K L I A N A K T V E G V W T L K D E T K K A T V T E
BINARY_4174 T T Y K L I L N G K Q A K E E A I K E L V D A G I A E K Y F K L I A N A K T V E G V W T L K D E T K K F T V T E
BINARY_4175 T T Y K L I L N G K Q A K E E A I K E L V D A G I A E K Y F K L I A N A K T V E G V W T L K D E T K T A T V T E
BINARY_4176 T T Y K L I L N G K Q A K E E A I K E L V D A G I A E K Y F K L I A N A K T V E G V W T L K D E T K T F T V T E
BINARY_4177 T T Y K L I L N G K Q A K E E A I K E L V D A G I A E K Y F K L I A N A K T V E G V W T Y K D E I L K A T V T E
BINARY_4178 T T Y K L I L N G K Q A K E E A I K E L V D A G I A E K Y F K L I A N A K T V E G V W T Y K D E I L K F T V T E
BINARY_4179 T T Y K L I L N G K Q A K E E A I K E L V D A G I A E K Y F K L I A N A K T V E G V W T Y K D E I L T A T V T E
BINARY_4180 T T Y K L I L N G K Q A K E E A I K E L V D A G I A E K Y F K L I A N A K T V E G V W T Y K D E I L T F T V T E
BINARY_4181 T T Y K L I L N G K Q A K E E A I K E L V D A G I A E K Y F K L I A N A K T V E G V W T Y K D E I K K A T V T E
BINARY_4182 T T Y K L I L N G K Q A K E E A I K E L V D A G I A E K Y F K L I A N A K T V E G V W T Y K D E I K K F T V T E
BINARY_4183 T T Y K L I L N G K Q A K E E A I K E L V D A G I A E K Y F K L I A N A K T V E G V W T Y K D E I K T A T V T E
BINARY_4184 T T Y K L I L N G K Q A K E E A I K E L V D A G I A E K Y F K L I A N A K T V E G V W T Y K D E I K T F T V T E
BINARY_4185 T T Y K L I L N G K Q A K E E A I K E L V D A G I A E K Y F K L I A N A K T V E G V W T Y K D E T L K A T V T E
BINARY_4186 T T Y K L I L N G K Q A K E E A I K E L V D A G I A E K Y F K L I A N A K T V E G V W T Y K D E T L K F T V T E
BINARY_4187 T T Y K L I L N G K Q A K E E A I K E L V D A G I A E K Y F K L I A N A K T V E G V W T Y K D E T L T A T V T E
BINARY_4188 T T Y K L I L N G K Q A K E E A I K E L V D A G I A E K Y F K L I A N A K T V E G V W T Y K D E T L T F T V T E
BINARY_4189 T T Y K L I L N G K Q A K E E A I K E L V D A G I A E K Y F K L I A N A K T V E G V W T Y K D E T K K A T V T E
BINARY_4190 T T Y K L I L N G K Q A K E E A I K E L V D A G I A E K Y F K L I A N A K T V E G V W T Y K D E T K K F T V T E
BINARY_4191 T T Y K L I L N G K Q A K E E A I K E L V D A G I A E K Y F K L I A N A K T V E G V W T Y K D E T K T A T V T E
BINARY_4192 T T Y K L I L N G K Q A K E E A I K E L V D A G I A E K Y F K L I A N A K T V E G V W T Y K D E T K T F T V T E
BINARY_4193 T T Y K L I L N G K Q A K E E A I K E L V D A G I A E K Y F K L Y A N A K T V E G V W T L K D E I L K A T V T E
BINARY_4194 T T Y K L I L N G K Q A K E E A I K E L V D A G I A E K Y F K L Y A N A K T V E G V W T L K D E I L K F T V T E
BINARY_4195 T T Y K L I L N G K Q A K E E A I K E L V D A G I A E K Y F K L Y A N A K T V E G V W T L K D E I L T A T V T E
BINARY_4196 T T Y K L I L N G K Q A K E E A I K E L V D A G I A E K Y F K L Y A N A K T V E G V W T L K D E I L T F T V T E

BINARY_4197 T T Y K L I L N G K Q A K E E A I K E L V D A G I A E K Y F K L Y A N A K T V E G V W T L K D E I K K A T V T E
BINARY_4198 T T Y K L I L N G K Q A K E E A I K E L V D A G I A E K Y F K L Y A N A K T V E G V W T L K D E I K K F T V T E
BINARY_4199 T T Y K L I L N G K Q A K E E A I K E L V D A G I A E K Y F K L Y A N A K T V E G V W T L K D E I K T A T V T E
BINARY_4200 T T Y K L I L N G K Q A K E E A I K E L V D A G I A E K Y F K L Y A N A K T V E G V W T L K D E I K T F T V T E
BINARY_4201 T T Y K L I L N G K Q A K E E A I K E L V D A G I A E K Y F K L Y A N A K T V E G V W T L K D E T L K A T V T E
BINARY_4202 T T Y K L I L N G K Q A K E E A I K E L V D A G I A E K Y F K L Y A N A K T V E G V W T L K D E T L K F T V T E
BINARY_4203 T T Y K L I L N G K Q A K E E A I K E L V D A G I A E K Y F K L Y A N A K T V E G V W T L K D E T L T A T V T E
BINARY_4204 T T Y K L I L N G K Q A K E E A I K E L V D A G I A E K Y F K L Y A N A K T V E G V W T L K D E T L T F T V T E
BINARY_4205 T T Y K L I L N G K Q A K E E A I K E L V D A G I A E K Y F K L Y A N A K T V E G V W T L K D E T K K A T V T E
BINARY_4206 T T Y K L I L N G K Q A K E E A I K E L V D A G I A E K Y F K L Y A N A K T V E G V W T L K D E T K K F T V T E
BINARY_4207 T T Y K L I L N G K Q A K E E A I K E L V D A G I A E K Y F K L Y A N A K T V E G V W T L K D E T K T A T V T E
BINARY_4208 T T Y K L I L N G K Q A K E E A I K E L V D A G I A E K Y F K L Y A N A K T V E G V W T L K D E T K T F T V T E
BINARY_4209 T T Y K L I L N G K Q A K E E A I K E L V D A G I A E K Y F K L Y A N A K T V E G V W T Y K D E I L K A T V T E
BINARY_4210 T T Y K L I L N G K Q A K E E A I K E L V D A G I A E K Y F K L Y A N A K T V E G V W T Y K D E I L K F T V T E
BINARY_4211 T T Y K L I L N G K Q A K E E A I K E L V D A G I A E K Y F K L Y A N A K T V E G V W T Y K D E I L T A T V T E
BINARY_4212 T T Y K L I L N G K Q A K E E A I K E L V D A G I A E K Y F K L Y A N A K T V E G V W T Y K D E I L T F T V T E
BINARY_4213 T T Y K L I L N G K Q A K E E A I K E L V D A G I A E K Y F K L Y A N A K T V E G V W T Y K D E I K K A T V T E
BINARY_4214 T T Y K L I L N G K Q A K E E A I K E L V D A G I A E K Y F K L Y A N A K T V E G V W T Y K D E I K K F T V T E
BINARY_4215 T T Y K L I L N G K Q A K E E A I K E L V D A G I A E K Y F K L Y A N A K T V E G V W T Y K D E I K T A T V T E
BINARY_4216 T T Y K L I L N G K Q A K E E A I K E L V D A G I A E K Y F K L Y A N A K T V E G V W T Y K D E I K T F T V T E
BINARY_4217 T T Y K L I L N G K Q A K E E A I K E L V D A G I A E K Y F K L Y A N A K T V E G V W T Y K D E T L K A T V T E
BINARY_4218 T T Y K L I L N G K Q A K E E A I K E L V D A G I A E K Y F K L Y A N A K T V E G V W T Y K D E T L K F T V T E
BINARY_4219 T T Y K L I L N G K Q A K E E A I K E L V D A G I A E K Y F K L Y A N A K T V E G V W T Y K D E T L T A T V T E
BINARY_4220 T T Y K L I L N G K Q A K E E A I K E L V D A G I A E K Y F K L Y A N A K T V E G V W T Y K D E T L T F T V T E
BINARY_4221 T T Y K L I L N G K Q A K E E A I K E L V D A G I A E K Y F K L Y A N A K T V E G V W T Y K D E T K K A T V T E
BINARY_4222 T T Y K L I L N G K Q A K E E A I K E L V D A G I A E K Y F K L Y A N A K T V E G V W T Y K D E T K K F T V T E
BINARY_4223 T T Y K L I L N G K Q A K E E A I K E L V D A G I A E K Y F K L Y A N A K T V E G V W T Y K D E T K T A T V T E
BINARY_4224 T T Y K L I L N G K Q A K E E A I K E L V D A G I A E K Y F K L Y A N A K T V E G V W T Y K D E T K T F T V T E
BINARY_4225 T T Y K L I L N G K Q A K E E A I K E L V D A G T A E K Y I K L I A N A K T V E G V W T L K D E I L K A T V T E
BINARY_4226 T T Y K L I L N G K Q A K E E A I K E L V D A G T A E K Y I K L I A N A K T V E G V W T L K D E I L K F T V T E
BINARY_4227 T T Y K L I L N G K Q A K E E A I K E L V D A G T A E K Y I K L I A N A K T V E G V W T L K D E I L T A T V T E
BINARY_4228 T T Y K L I L N G K Q A K E E A I K E L V D A G T A E K Y I K L I A N A K T V E G V W T L K D E I L T F T V T E
BINARY_4229 T T Y K L I L N G K Q A K E E A I K E L V D A G T A E K Y I K L I A N A K T V E G V W T L K D E I K K A T V T E
BINARY_4230 T T Y K L I L N G K Q A K E E A I K E L V D A G T A E K Y I K L I A N A K T V E G V W T L K D E I K K F T V T E
BINARY_4231 T T Y K L I L N G K Q A K E E A I K E L V D A G T A E K Y I K L I A N A K T V E G V W T L K D E I K T A T V T E
BINARY_4232 T T Y K L I L N G K Q A K E E A I K E L V D A G T A E K Y I K L I A N A K T V E G V W T L K D E I K T F T V T E
BINARY_4233 T T Y K L I L N G K Q A K E E A I K E L V D A G T A E K Y I K L I A N A K T V E G V W T L K D E T L K A T V T E
BINARY_4234 T T Y K L I L N G K Q A K E E A I K E L V D A G T A E K Y I K L I A N A K T V E G V W T L K D E T L K F T V T E
BINARY_4235 T T Y K L I L N G K Q A K E E A I K E L V D A G T A E K Y I K L I A N A K T V E G V W T L K D E T L T A T V T E
BINARY_4236 T T Y K L I L N G K Q A K E E A I K E L V D A G T A E K Y I K L I A N A K T V E G V W T L K D E T L T F T V T E
BINARY_4237 T T Y K L I L N G K Q A K E E A I K E L V D A G T A E K Y I K L I A N A K T V E G V W T L K D E T K K A T V T E
BINARY_4238 T T Y K L I L N G K Q A K E E A I K E L V D A G T A E K Y I K L I A N A K T V E G V W T L K D E T K K F T V T E
BINARY_4239 T T Y K L I L N G K Q A K E E A I K E L V D A G T A E K Y I K L I A N A K T V E G V W T L K D E T K T A T V T E
BINARY_4240 T T Y K L I L N G K Q A K E E A I K E L V D A G T A E K Y I K L I A N A K T V E G V W T L K D E T K T F T V T E
BINARY_4241 T T Y K L I L N G K Q A K E E A I K E L V D A G T A E K Y I K L I A N A K T V E G V W T Y K D E I L K A T V T E
BINARY_4242 T T Y K L I L N G K Q A K E E A I K E L V D A G T A E K Y I K L I A N A K T V E G V W T Y K D E I L K F T V T E
BINARY_4243 T T Y K L I L N G K Q A K E E A I K E L V D A G T A E K Y I K L I A N A K T V E G V W T Y K D E I L T A T V T E
BINARY_4244 T T Y K L I L N G K Q A K E E A I K E L V D A G T A E K Y I K L I A N A K T V E G V W T Y K D E I L T F T V T E
BINARY_4245 T T Y K L I L N G K Q A K E E A I K E L V D A G T A E K Y I K L I A N A K T V E G V W T Y K D E I K K A T V T E
BINARY_4246 T T Y K L I L N G K Q A K E E A I K E L V D A G T A E K Y I K L I A N A K T V E G V W T Y K D E I K K F T V T E
BINARY_4247 T T Y K L I L N G K Q A K E E A I K E L V D A G T A E K Y I K L I A N A K T V E G V W T Y K D E I K T A T V T E
BINARY_4248 T T Y K L I L N G K Q A K E E A I K E L V D A G T A E K Y I K L I A N A K T V E G V W T Y K D E I K T F T V T E
BINARY_4249 T T Y K L I L N G K Q A K E E A I K E L V D A G T A E K Y I K L I A N A K T V E G V W T Y K D E T L K A T V T E
BINARY_4250 T T Y K L I L N G K Q A K E E A I K E L V D A G T A E K Y I K L I A N A K T V E G V W T Y K D E T L K F T V T E
BINARY_4251 T T Y K L I L N G K Q A K E E A I K E L V D A G T A E K Y I K L I A N A K T V E G V W T Y K D E T L T A T V T E
BINARY_4252 T T Y K L I L N G K Q A K E E A I K E L V D A G T A E K Y I K L I A N A K T V E G V W T Y K D E T L T F T V T E
BINARY_4253 T T Y K L I L N G K Q A K E E A I K E L V D A G T A E K Y I K L I A N A K T V E G V W T Y K D E T K K A T V T E
BINARY_4254 T T Y K L I L N G K Q A K E E A I K E L V D A G T A E K Y I K L I A N A K T V E G V W T Y K D E T K K F T V T E
BINARY_4255 T T Y K L I L N G K Q A K E E A I K E L V D A G T A E K Y I K L I A N A K T V E G V W T Y K D E T K T A T V T E
BINARY_4256 T T Y K L I L N G K Q A K E E A I K E L V D A G T A E K Y I K L I A N A K T V E G V W T Y K D E T K T F T V T E
BINARY_4257 T T Y K L I L N G K Q A K E E A I K E L V D A G T A E K Y I K L Y A N A K T V E G V W T L K D E I L K A T V T E
BINARY_4258 T T Y K L I L N G K Q A K E E A I K E L V D A G T A E K Y I K L Y A N A K T V E G V W T L K D E I L K F T V T E
BINARY_4259 T T Y K L I L N G K Q A K E E A I K E L V D A G T A E K Y I K L Y A N A K T V E G V W T L K D E I L T A T V T E
BINARY_4260 T T Y K L I L N G K Q A K E E A I K E L V D A G T A E K Y I K L Y A N A K T V E G V W T L K D E I L T F T V T E
BINARY_4261 T T Y K L I L N G K Q A K E E A I K E L V D A G T A E K Y I K L Y A N A K T V E G V W T L K D E I K K A T V T E
BINARY_4262 T T Y K L I L N G K Q A K E E A I K E L V D A G T A E K Y I K L Y A N A K T V E G V W T L K D E I K K F T V T E
BINARY_4263 T T Y K L I L N G K Q A K E E A I K E L V D A G T A E K Y I K L Y A N A K T V E G V W T L K D E I K T A T V T E
BINARY_4264 T T Y K L I L N G K Q A K E E A I K E L V D A G T A E K Y I K L Y A N A K T V E G V W T L K D E I K T F T V T E
BINARY_4265 T T Y K L I L N G K Q A K E E A I K E L V D A G T A E K Y I K L Y A N A K T V E G V W T L K D E T L K A T V T E
BINARY_4266 T T Y K L I L N G K Q A K E E A I K E L V D A G T A E K Y I K L Y A N A K T V E G V W T L K D E T L K F T V T E
BINARY_4267 T T Y K L I L N G K Q A K E E A I K E L V D A G T A E K Y I K L Y A N A K T V E G V W T L K D E T L T A T V T E
BINARY_4268 T T Y K L I L N G K Q A K E E A I K E L V D A G T A E K Y I K L Y A N A K T V E G V W T L K D E T L T F T V T E
BINARY_4269 T T Y K L I L N G K Q A K E E A I K E L V D A G T A E K Y I K L Y A N A K T V E G V W T L K D E T K K A T V T E
BINARY_4270 T T Y K L I L N G K Q A K E E A I K E L V D A G T A E K Y I K L Y A N A K T V E G V W T L K D E T K K F T V T E
BINARY_4271 T T Y K L I L N G K Q A K E E A I K E L V D A G T A E K Y I K L Y A N A K T V E G V W T L K D E T K T A T V T E
BINARY_4272 T T Y K L I L N G K Q A K E E A I K E L V D A G T A E K Y I K L Y A N A K T V E G V W T L K D E T K T F T V T E
BINARY_4273 T T Y K L I L N G K Q A K E E A I K E L V D A G T A E K Y I K L Y A N A K T V E G V W T Y K D E I L K A T V T E
BINARY_4274 T T Y K L I L N G K Q A K E E A I K E L V D A G T A E K Y I K L Y A N A K T V E G V W T Y K D E I L K F T V T E
BINARY_4275 T T Y K L I L N G K Q A K E E A I K E L V D A G T A E K Y I K L Y A N A K T V E G V W T Y K D E I L T A T V T E
BINARY_4276 T T Y K L I L N G K Q A K E E A I K E L V D A G T A E K Y I K L Y A N A K T V E G V W T Y K D E I L T F T V T E
BINARY_4277 T T Y K L I L N G K Q A K E E A I K E L V D A G T A E K Y I K L Y A N A K T V E G V W T Y K D E I K K A T V T E
BINARY_4278 T T Y K L I L N G K Q A K E E A I K E L V D A G T A E K Y I K L Y A N A K T V E G V W T Y K D E I K K F T V T E
BINARY_4279 T T Y K L I L N G K Q A K E E A I K E L V D A G T A E K Y I K L Y A N A K T V E G V W T Y K D E I K T A T V T E
BINARY_4280 T T Y K L I L N G K Q A K E E A I K E L V D A G T A E K Y I K L Y A N A K T V E G V W T Y K D E I K T F T V T E
BINARY_4281 T T Y K L I L N G K Q A K E E A I K E L V D A G T A E K Y I K L Y A N A K T V E G V W T Y K D E T L K A T V T E
BINARY_4282 T T Y K L I L N G K Q A K E E A I K E L V D A G T A E K Y I K L Y A N A K T V E G V W T Y K D E T L K F T V T E
BINARY_4283 T T Y K L I L N G K Q A K E E A I K E L V D A G T A E K Y I K L Y A N A K T V E G V W T Y K D E T L T A T V T E
BINARY_4284 T T Y K L I L N G K Q A K E E A I K E L V D A G T A E K Y I K L Y A N A K T V E G V W T Y K D E T L T F T V T E
BINARY_4285 T T Y K L I L N G K Q A K E E A I K E L V D A G T A E K Y I K L Y A N A K T V E G V W T Y K D E T K K A T V T E
BINARY_4286 T T Y K L I L N G K Q A K E E A I K E L V D A G T A E K Y I K L Y A N A K T V E G V W T Y K D E T K K F T V T E
BINARY_4287 T T Y K L I L N G K Q A K E E A I K E L V D A G T A E K Y I K L Y A N A K T V E G V W T Y K D E T K T A T V T E
BINARY_4288 T T Y K L I L N G K Q A K E E A I K E L V D A G T A E K Y I K L Y A N A K T V E G V W T Y K D E T K T F T V T E
BINARY_4289 T T Y K L I L N G K Q A K E E A I K E L V D A G T A E K Y F K L I A N A K T V E G V W T L K D E I L K A T V T E
BINARY_4290 T T Y K L I L N G K Q A K E E A I K E L V D A G T A E K Y F K L I A N A K T V E G V W T L K D E I L K F T V T E
BINARY_4291 T T Y K L I L N G K Q A K E E A I K E L V D A G T A E K Y F K L I A N A K T V E G V W T L K D E I L T A T V T E

BINARY_4292 T T Y K L I L N G K Q A K E E A I K E L V D A G T A E K Y F K L I A N A K T V E G V W T L K D E I L T F T V T E
BINARY_4293 T T Y K L I L N G K Q A K E E A I K E L V D A G T A E K Y F K L I A N A K T V E G V W T L K D E I K K A T V T E
BINARY_4294 T T Y K L I L N G K Q A K E E A I K E L V D A G T A E K Y F K L I A N A K T V E G V W T L K D E I K K F T V T E
BINARY_4295 T T Y K L I L N G K Q A K E E A I K E L V D A G T A E K Y F K L I A N A K T V E G V W T L K D E I K T A T V T E
BINARY_4296 T T Y K L I L N G K Q A K E E A I K E L V D A G T A E K Y F K L I A N A K T V E G V W T L K D E I K T F T V T E
BINARY_4297 T T Y K L I L N G K Q A K E E A I K E L V D A G T A E K Y F K L I A N A K T V E G V W T L K D E T L K A T V T E
BINARY_4298 T T Y K L I L N G K Q A K E E A I K E L V D A G T A E K Y F K L I A N A K T V E G V W T L K D E T L K F T V T E
BINARY_4299 T T Y K L I L N G K Q A K E E A I K E L V D A G T A E K Y F K L I A N A K T V E G V W T L K D E T L T A T V T E
BINARY_4300 T T Y K L I L N G K Q A K E E A I K E L V D A G T A E K Y F K L I A N A K T V E G V W T L K D E T L T F T V T E
BINARY_4301 T T Y K L I L N G K Q A K E E A I K E L V D A G T A E K Y F K L I A N A K T V E G V W T L K D E T K K A T V T E
BINARY_4302 T T Y K L I L N G K Q A K E E A I K E L V D A G T A E K Y F K L I A N A K T V E G V W T L K D E T K K F T V T E
BINARY_4303 T T Y K L I L N G K Q A K E E A I K E L V D A G T A E K Y F K L I A N A K T V E G V W T L K D E T K T A T V T E
BINARY_4304 T T Y K L I L N G K Q A K E E A I K E L V D A G T A E K Y F K L I A N A K T V E G V W T L K D E T K T F T V T E
BINARY_4305 T T Y K L I L N G K Q A K E E A I K E L V D A G T A E K Y F K L I A N A K T V E G V W T Y K D E I L K A T V T E
BINARY_4306 T T Y K L I L N G K Q A K E E A I K E L V D A G T A E K Y F K L I A N A K T V E G V W T Y K D E I L K F T V T E
BINARY_4307 T T Y K L I L N G K Q A K E E A I K E L V D A G T A E K Y F K L I A N A K T V E G V W T Y K D E I L T A T V T E
BINARY_4308 T T Y K L I L N G K Q A K E E A I K E L V D A G T A E K Y F K L I A N A K T V E G V W T Y K D E I L T F T V T E
BINARY_4309 T T Y K L I L N G K Q A K E E A I K E L V D A G T A E K Y F K L I A N A K T V E G V W T Y K D E I K K A T V T E
BINARY_4310 T T Y K L I L N G K Q A K E E A I K E L V D A G T A E K Y F K L I A N A K T V E G V W T Y K D E I K K F T V T E
BINARY_4311 T T Y K L I L N G K Q A K E E A I K E L V D A G T A E K Y F K L I A N A K T V E G V W T Y K D E I K T A T V T E
BINARY_4312 T T Y K L I L N G K Q A K E E A I K E L V D A G T A E K Y F K L I A N A K T V E G V W T Y K D E I K T F T V T E
BINARY_4313 T T Y K L I L N G K Q A K E E A I K E L V D A G T A E K Y F K L I A N A K T V E G V W T Y K D E T L K A T V T E
BINARY_4314 T T Y K L I L N G K Q A K E E A I K E L V D A G T A E K Y F K L I A N A K T V E G V W T Y K D E T L K F T V T E
BINARY_4315 T T Y K L I L N G K Q A K E E A I K E L V D A G T A E K Y F K L I A N A K T V E G V W T Y K D E T L T A T V T E
BINARY_4316 T T Y K L I L N G K Q A K E E A I K E L V D A G T A E K Y F K L I A N A K T V E G V W T Y K D E T L T F T V T E
BINARY_4317 T T Y K L I L N G K Q A K E E A I K E L V D A G T A E K Y F K L I A N A K T V E G V W T Y K D E T K K A T V T E
BINARY_4318 T T Y K L I L N G K Q A K E E A I K E L V D A G T A E K Y F K L I A N A K T V E G V W T Y K D E T K K F T V T E
BINARY_4319 T T Y K L I L N G K Q A K E E A I K E L V D A G T A E K Y F K L I A N A K T V E G V W T Y K D E T K T A T V T E
BINARY_4320 T T Y K L I L N G K Q A K E E A I K E L V D A G T A E K Y F K L I A N A K T V E G V W T Y K D E T K T F T V T E
BINARY_4321 T T Y K L I L N G K Q A K E E A I K E L V D A G T A E K Y F K L Y A N A K T V E G V W T L K D E I L K A T V T E
BINARY_4322 T T Y K L I L N G K Q A K E E A I K E L V D A G T A E K Y F K L Y A N A K T V E G V W T L K D E I L K F T V T E
BINARY_4323 T T Y K L I L N G K Q A K E E A I K E L V D A G T A E K Y F K L Y A N A K T V E G V W T L K D E I L T A T V T E
BINARY_4324 T T Y K L I L N G K Q A K E E A I K E L V D A G T A E K Y F K L Y A N A K T V E G V W T L K D E I L T F T V T E
BINARY_4325 T T Y K L I L N G K Q A K E E A I K E L V D A G T A E K Y F K L Y A N A K T V E G V W T L K D E I K K A T V T E
BINARY_4326 T T Y K L I L N G K Q A K E E A I K E L V D A G T A E K Y F K L Y A N A K T V E G V W T L K D E I K K F T V T E
BINARY_4327 T T Y K L I L N G K Q A K E E A I K E L V D A G T A E K Y F K L Y A N A K T V E G V W T L K D E I K T A T V T E
BINARY_4328 T T Y K L I L N G K Q A K E E A I K E L V D A G T A E K Y F K L Y A N A K T V E G V W T L K D E I K T F T V T E
BINARY_4329 T T Y K L I L N G K Q A K E E A I K E L V D A G T A E K Y F K L Y A N A K T V E G V W T L K D E T L K A T V T E
BINARY_4330 T T Y K L I L N G K Q A K E E A I K E L V D A G T A E K Y F K L Y A N A K T V E G V W T L K D E T L K F T V T E
BINARY_4331 T T Y K L I L N G K Q A K E E A I K E L V D A G T A E K Y F K L Y A N A K T V E G V W T L K D E T L T A T V T E
BINARY_4332 T T Y K L I L N G K Q A K E E A I K E L V D A G T A E K Y F K L Y A N A K T V E G V W T L K D E T L T F T V T E
BINARY_4333 T T Y K L I L N G K Q A K E E A I K E L V D A G T A E K Y F K L Y A N A K T V E G V W T L K D E T K K A T V T E
BINARY_4334 T T Y K L I L N G K Q A K E E A I K E L V D A G T A E K Y F K L Y A N A K T V E G V W T L K D E T K K F T V T E
BINARY_4335 T T Y K L I L N G K Q A K E E A I K E L V D A G T A E K Y F K L Y A N A K T V E G V W T L K D E T K T A T V T E
BINARY_4336 T T Y K L I L N G K Q A K E E A I K E L V D A G T A E K Y F K L Y A N A K T V E G V W T L K D E T K T F T V T E
BINARY_4337 T T Y K L I L N G K Q A K E E A I K E L V D A G T A E K Y F K L Y A N A K T V E G V W T Y K D E I L K A T V T E
BINARY_4338 T T Y K L I L N G K Q A K E E A I K E L V D A G T A E K Y F K L Y A N A K T V E G V W T Y K D E I L K F T V T E
BINARY_4339 T T Y K L I L N G K Q A K E E A I K E L V D A G T A E K Y F K L Y A N A K T V E G V W T Y K D E I L T A T V T E
BINARY_4340 T T Y K L I L N G K Q A K E E A I K E L V D A G T A E K Y F K L Y A N A K T V E G V W T Y K D E I L T F T V T E
BINARY_4341 T T Y K L I L N G K Q A K E E A I K E L V D A G T A E K Y F K L Y A N A K T V E G V W T Y K D E I K K A T V T E
BINARY_4342 T T Y K L I L N G K Q A K E E A I K E L V D A G T A E K Y F K L Y A N A K T V E G V W T Y K D E I K K F T V T E
BINARY_4343 T T Y K L I L N G K Q A K E E A I K E L V D A G T A E K Y F K L Y A N A K T V E G V W T Y K D E I K T A T V T E
BINARY_4344 T T Y K L I L N G K Q A K E E A I K E L V D A G T A E K Y F K L Y A N A K T V E G V W T Y K D E I K T F T V T E
BINARY_4345 T T Y K L I L N G K Q A K E E A I K E L V D A G T A E K Y F K L Y A N A K T V E G V W T Y K D E T L K A T V T E
BINARY_4346 T T Y K L I L N G K Q A K E E A I K E L V D A G T A E K Y F K L Y A N A K T V E G V W T Y K D E T L K F T V T E
BINARY_4347 T T Y K L I L N G K Q A K E E A I K E L V D A G T A E K Y F K L Y A N A K T V E G V W T Y K D E T L T A T V T E
BINARY_4348 T T Y K L I L N G K Q A K E E A I K E L V D A G T A E K Y F K L Y A N A K T V E G V W T Y K D E T L T F T V T E
BINARY_4349 T T Y K L I L N G K Q A K E E A I K E L V D A G T A E K Y F K L Y A N A K T V E G V W T Y K D E T K K A T V T E
BINARY_4350 T T Y K L I L N G K Q A K E E A I K E L V D A G T A E K Y F K L Y A N A K T V E G V W T Y K D E T K K F T V T E
BINARY_4351 T T Y K L I L N G K Q A K E E A I K E L V D A G T A E K Y F K L Y A N A K T V E G V W T Y K D E T K T A T V T E
BINARY_4352 T T Y K L I L N G K Q A K E E A I K E L V D A G T A E K Y F K L Y A N A K T V E G V W T Y K D E T K T F T V T E
BINARY_4353 T T Y K L I L N G K Q A K E E A I K E L V D A A I A E K Y I K L I A N A K T V E G V W T L K D E I L K A T V T E
BINARY_4354 T T Y K L I L N G K Q A K E E A I K E L V D A A I A E K Y I K L I A N A K T V E G V W T L K D E I L K F T V T E
BINARY_4355 T T Y K L I L N G K Q A K E E A I K E L V D A A I A E K Y I K L I A N A K T V E G V W T L K D E I L T A T V T E
BINARY_4356 T T Y K L I L N G K Q A K E E A I K E L V D A A I A E K Y I K L I A N A K T V E G V W T L K D E I L T F T V T E
BINARY_4357 T T Y K L I L N G K Q A K E E A I K E L V D A A I A E K Y I K L I A N A K T V E G V W T L K D E I K K A T V T E
BINARY_4358 T T Y K L I L N G K Q A K E E A I K E L V D A A I A E K Y I K L I A N A K T V E G V W T L K D E I K K F T V T E
BINARY_4359 T T Y K L I L N G K Q A K E E A I K E L V D A A I A E K Y I K L I A N A K T V E G V W T L K D E I K T A T V T E
BINARY_4360 T T Y K L I L N G K Q A K E E A I K E L V D A A I A E K Y I K L I A N A K T V E G V W T L K D E I K T F T V T E
BINARY_4361 T T Y K L I L N G K Q A K E E A I K E L V D A A I A E K Y I K L I A N A K T V E G V W T L K D E T L K A T V T E
BINARY_4362 T T Y K L I L N G K Q A K E E A I K E L V D A A I A E K Y I K L I A N A K T V E G V W T L K D E T L K F T V T E
BINARY_4363 T T Y K L I L N G K Q A K E E A I K E L V D A A I A E K Y I K L I A N A K T V E G V W T L K D E T L T A T V T E
BINARY_4364 T T Y K L I L N G K Q A K E E A I K E L V D A A I A E K Y I K L I A N A K T V E G V W T L K D E T L T F T V T E
BINARY_4365 T T Y K L I L N G K Q A K E E A I K E L V D A A I A E K Y I K L I A N A K T V E G V W T L K D E T K K A T V T E
BINARY_4366 T T Y K L I L N G K Q A K E E A I K E L V D A A I A E K Y I K L I A N A K T V E G V W T L K D E T K K F T V T E
BINARY_4367 T T Y K L I L N G K Q A K E E A I K E L V D A A I A E K Y I K L I A N A K T V E G V W T L K D E T K T A T V T E
BINARY_4368 T T Y K L I L N G K Q A K E E A I K E L V D A A I A E K Y I K L I A N A K T V E G V W T L K D E T K T F T V T E
BINARY_4369 T T Y K L I L N G K Q A K E E A I K E L V D A A I A E K Y I K L I A N A K T V E G V W T Y K D E I L K A T V T E
BINARY_4370 T T Y K L I L N G K Q A K E E A I K E L V D A A I A E K Y I K L I A N A K T V E G V W T Y K D E I L K F T V T E
BINARY_4371 T T Y K L I L N G K Q A K E E A I K E L V D A A I A E K Y I K L I A N A K T V E G V W T Y K D E I L T A T V T E
BINARY_4372 T T Y K L I L N G K Q A K E E A I K E L V D A A I A E K Y I K L I A N A K T V E G V W T Y K D E I L T F T V T E
BINARY_4373 T T Y K L I L N G K Q A K E E A I K E L V D A A I A E K Y I K L I A N A K T V E G V W T Y K D E I K K A T V T E
BINARY_4374 T T Y K L I L N G K Q A K E E A I K E L V D A A I A E K Y I K L I A N A K T V E G V W T Y K D E I K K F T V T E
BINARY_4375 T T Y K L I L N G K Q A K E E A I K E L V D A A I A E K Y I K L I A N A K T V E G V W T Y K D E I K T A T V T E
BINARY_4376 T T Y K L I L N G K Q A K E E A I K E L V D A A I A E K Y I K L I A N A K T V E G V W T Y K D E I K T F T V T E
BINARY_4377 T T Y K L I L N G K Q A K E E A I K E L V D A A I A E K Y I K L I A N A K T V E G V W T Y K D E T L K A T V T E
BINARY_4378 T T Y K L I L N G K Q A K E E A I K E L V D A A I A E K Y I K L I A N A K T V E G V W T Y K D E T L K F T V T E
BINARY_4379 T T Y K L I L N G K Q A K E E A I K E L V D A A I A E K Y I K L I A N A K T V E G V W T Y K D E T L T A T V T E
BINARY_4380 T T Y K L I L N G K Q A K E E A I K E L V D A A I A E K Y I K L I A N A K T V E G V W T Y K D E T L T F T V T E
BINARY_4381 T T Y K L I L N G K Q A K E E A I K E L V D A A I A E K Y I K L I A N A K T V E G V W T Y K D E T K K A T V T E
BINARY_4382 T T Y K L I L N G K Q A K E E A I K E L V D A A I A E K Y I K L I A N A K T V E G V W T Y K D E T K K F T V T E
BINARY_4383 T T Y K L I L N G K Q A K E E A I K E L V D A A I A E K Y I K L I A N A K T V E G V W T Y K D E T K T A T V T E
BINARY_4384 T T Y K L I L N G K Q A K E E A I K E L V D A A I A E K Y I K L I A N A K T V E G V W T Y K D E T K T F T V T E
BINARY_4385 T T Y K L I L N G K Q A K E E A I K E L V D A A I A E K Y I K L Y A N A K T V E G V W T L K D E I L K A T V T E
BINARY_4386 T T Y K L I L N G K Q A K E E A I K E L V D A A I A E K Y I K L Y A N A K T V E G V W T L K D E I L K F T V T E

BINARY_4387 T T Y K L I L N G K Q A K E E A I K E L V D A A I A E K Y I K L Y A N A K T V E G V W T L K D E I L T A T V T E
BINARY_4388 T T Y K L I L N G K Q A K E E A I K E L V D A A I A E K Y I K L Y A N A K T V E G V W T L K D E I L T F T V T E
BINARY_4389 T T Y K L I L N G K Q A K E E A I K E L V D A A I A E K Y I K L Y A N A K T V E G V W T L K D E I K K A T V T E
BINARY_4390 T T Y K L I L N G K Q A K E E A I K E L V D A A I A E K Y I K L Y A N A K T V E G V W T L K D E I K K F T V T E
BINARY_4391 T T Y K L I L N G K Q A K E E A I K E L V D A A I A E K Y I K L Y A N A K T V E G V W T L K D E I K T A T V T E
BINARY_4392 T T Y K L I L N G K Q A K E E A I K E L V D A A I A E K Y I K L Y A N A K T V E G V W T L K D E I K T F T V T E
BINARY_4393 T T Y K L I L N G K Q A K E E A I K E L V D A A I A E K Y I K L Y A N A K T V E G V W T L K D E T L K A T V T E
BINARY_4394 T T Y K L I L N G K Q A K E E A I K E L V D A A I A E K Y I K L Y A N A K T V E G V W T L K D E T L K F T V T E
BINARY_4395 T T Y K L I L N G K Q A K E E A I K E L V D A A I A E K Y I K L Y A N A K T V E G V W T L K D E T L T A T V T E
BINARY_4396 T T Y K L I L N G K Q A K E E A I K E L V D A A I A E K Y I K L Y A N A K T V E G V W T L K D E T L T F T V T E
BINARY_4397 T T Y K L I L N G K Q A K E E A I K E L V D A A I A E K Y I K L Y A N A K T V E G V W T L K D E T K K A T V T E
BINARY_4398 T T Y K L I L N G K Q A K E E A I K E L V D A A I A E K Y I K L Y A N A K T V E G V W T L K D E T K K F T V T E
BINARY_4399 T T Y K L I L N G K Q A K E E A I K E L V D A A I A E K Y I K L Y A N A K T V E G V W T L K D E T K T A T V T E
BINARY_4400 T T Y K L I L N G K Q A K E E A I K E L V D A A I A E K Y I K L Y A N A K T V E G V W T L K D E T K T F T V T E
BINARY_4401 T T Y K L I L N G K Q A K E E A I K E L V D A A I A E K Y I K L Y A N A K T V E G V W T Y K D E I L K A T V T E
BINARY_4402 T T Y K L I L N G K Q A K E E A I K E L V D A A I A E K Y I K L Y A N A K T V E G V W T Y K D E I L K F T V T E
BINARY_4403 T T Y K L I L N G K Q A K E E A I K E L V D A A I A E K Y I K L Y A N A K T V E G V W T Y K D E I L T A T V T E
BINARY_4404 T T Y K L I L N G K Q A K E E A I K E L V D A A I A E K Y I K L Y A N A K T V E G V W T Y K D E I L T F T V T E
BINARY_4405 T T Y K L I L N G K Q A K E E A I K E L V D A A I A E K Y I K L Y A N A K T V E G V W T Y K D E I K K A T V T E
BINARY_4406 T T Y K L I L N G K Q A K E E A I K E L V D A A I A E K Y I K L Y A N A K T V E G V W T Y K D E I K K F T V T E
BINARY_4407 T T Y K L I L N G K Q A K E E A I K E L V D A A I A E K Y I K L Y A N A K T V E G V W T Y K D E I K T A T V T E
BINARY_4408 T T Y K L I L N G K Q A K E E A I K E L V D A A I A E K Y I K L Y A N A K T V E G V W T Y K D E I K T F T V T E
BINARY_4409 T T Y K L I L N G K Q A K E E A I K E L V D A A I A E K Y I K L Y A N A K T V E G V W T Y K D E T L K A T V T E
BINARY_4410 T T Y K L I L N G K Q A K E E A I K E L V D A A I A E K Y I K L Y A N A K T V E G V W T Y K D E T L K F T V T E
BINARY_4411 T T Y K L I L N G K Q A K E E A I K E L V D A A I A E K Y I K L Y A N A K T V E G V W T Y K D E T L T A T V T E
BINARY_4412 T T Y K L I L N G K Q A K E E A I K E L V D A A I A E K Y I K L Y A N A K T V E G V W T Y K D E T L T F T V T E
BINARY_4413 T T Y K L I L N G K Q A K E E A I K E L V D A A I A E K Y I K L Y A N A K T V E G V W T Y K D E T K K A T V T E
BINARY_4414 T T Y K L I L N G K Q A K E E A I K E L V D A A I A E K Y I K L Y A N A K T V E G V W T Y K D E T K K F T V T E
BINARY_4415 T T Y K L I L N G K Q A K E E A I K E L V D A A I A E K Y I K L Y A N A K T V E G V W T Y K D E T K T A T V T E
BINARY_4416 T T Y K L I L N G K Q A K E E A I K E L V D A A I A E K Y I K L Y A N A K T V E G V W T Y K D E T K T F T V T E
BINARY_4417 T T Y K L I L N G K Q A K E E A I K E L V D A A I A E K Y F K L I A N A K T V E G V W T L K D E I L K A T V T E
BINARY_4418 T T Y K L I L N G K Q A K E E A I K E L V D A A I A E K Y F K L I A N A K T V E G V W T L K D E I L K F T V T E
BINARY_4419 T T Y K L I L N G K Q A K E E A I K E L V D A A I A E K Y F K L I A N A K T V E G V W T L K D E I L T A T V T E
BINARY_4420 T T Y K L I L N G K Q A K E E A I K E L V D A A I A E K Y F K L I A N A K T V E G V W T L K D E I L T F T V T E
BINARY_4421 T T Y K L I L N G K Q A K E E A I K E L V D A A I A E K Y F K L I A N A K T V E G V W T L K D E I K K A T V T E
BINARY_4422 T T Y K L I L N G K Q A K E E A I K E L V D A A I A E K Y F K L I A N A K T V E G V W T L K D E I K K F T V T E
BINARY_4423 T T Y K L I L N G K Q A K E E A I K E L V D A A I A E K Y F K L I A N A K T V E G V W T L K D E I K T A T V T E
BINARY_4424 T T Y K L I L N G K Q A K E E A I K E L V D A A I A E K Y F K L I A N A K T V E G V W T L K D E I K T F T V T E
BINARY_4425 T T Y K L I L N G K Q A K E E A I K E L V D A A I A E K Y F K L I A N A K T V E G V W T L K D E T L K A T V T E
BINARY_4426 T T Y K L I L N G K Q A K E E A I K E L V D A A I A E K Y F K L I A N A K T V E G V W T L K D E T L K F T V T E
BINARY_4427 T T Y K L I L N G K Q A K E E A I K E L V D A A I A E K Y F K L I A N A K T V E G V W T L K D E T L T A T V T E
BINARY_4428 T T Y K L I L N G K Q A K E E A I K E L V D A A I A E K Y F K L I A N A K T V E G V W T L K D E T L T F T V T E
BINARY_4429 T T Y K L I L N G K Q A K E E A I K E L V D A A I A E K Y F K L I A N A K T V E G V W T L K D E T K K A T V T E
BINARY_4430 T T Y K L I L N G K Q A K E E A I K E L V D A A I A E K Y F K L I A N A K T V E G V W T L K D E T K K F T V T E
BINARY_4431 T T Y K L I L N G K Q A K E E A I K E L V D A A I A E K Y F K L I A N A K T V E G V W T L K D E T K T A T V T E
BINARY_4432 T T Y K L I L N G K Q A K E E A I K E L V D A A I A E K Y F K L I A N A K T V E G V W T L K D E T K T F T V T E
BINARY_4433 T T Y K L I L N G K Q A K E E A I K E L V D A A I A E K Y F K L I A N A K T V E G V W T Y K D E I L K A T V T E
BINARY_4434 T T Y K L I L N G K Q A K E E A I K E L V D A A I A E K Y F K L I A N A K T V E G V W T Y K D E I L K F T V T E
BINARY_4435 T T Y K L I L N G K Q A K E E A I K E L V D A A I A E K Y F K L I A N A K T V E G V W T Y K D E I L T A T V T E
BINARY_4436 T T Y K L I L N G K Q A K E E A I K E L V D A A I A E K Y F K L I A N A K T V E G V W T Y K D E I L T F T V T E
BINARY_4437 T T Y K L I L N G K Q A K E E A I K E L V D A A I A E K Y F K L I A N A K T V E G V W T Y K D E I K K A T V T E
BINARY_4438 T T Y K L I L N G K Q A K E E A I K E L V D A A I A E K Y F K L I A N A K T V E G V W T Y K D E I K K F T V T E
BINARY_4439 T T Y K L I L N G K Q A K E E A I K E L V D A A I A E K Y F K L I A N A K T V E G V W T Y K D E I K T A T V T E
BINARY_4440 T T Y K L I L N G K Q A K E E A I K E L V D A A I A E K Y F K L I A N A K T V E G V W T Y K D E I K T F T V T E
BINARY_4441 T T Y K L I L N G K Q A K E E A I K E L V D A A I A E K Y F K L I A N A K T V E G V W T Y K D E T L K A T V T E
BINARY_4442 T T Y K L I L N G K Q A K E E A I K E L V D A A I A E K Y F K L I A N A K T V E G V W T Y K D E T L K F T V T E
BINARY_4443 T T Y K L I L N G K Q A K E E A I K E L V D A A I A E K Y F K L I A N A K T V E G V W T Y K D E T L T A T V T E
BINARY_4444 T T Y K L I L N G K Q A K E E A I K E L V D A A I A E K Y F K L I A N A K T V E G V W T Y K D E T L T F T V T E
BINARY_4445 T T Y K L I L N G K Q A K E E A I K E L V D A A I A E K Y F K L I A N A K T V E G V W T Y K D E T K K A T V T E
BINARY_4446 T T Y K L I L N G K Q A K E E A I K E L V D A A I A E K Y F K L I A N A K T V E G V W T Y K D E T K K F T V T E
BINARY_4447 T T Y K L I L N G K Q A K E E A I K E L V D A A I A E K Y F K L I A N A K T V E G V W T Y K D E T K T A T V T E
BINARY_4448 T T Y K L I L N G K Q A K E E A I K E L V D A A I A E K Y F K L I A N A K T V E G V W T Y K D E T K T F T V T E
BINARY_4449 T T Y K L I L N G K Q A K E E A I K E L V D A A I A E K Y F K L Y A N A K T V E G V W T L K D E I L K A T V T E
BINARY_4450 T T Y K L I L N G K Q A K E E A I K E L V D A A I A E K Y F K L Y A N A K T V E G V W T L K D E I L K F T V T E
BINARY_4451 T T Y K L I L N G K Q A K E E A I K E L V D A A I A E K Y F K L Y A N A K T V E G V W T L K D E I L T A T V T E
BINARY_4452 T T Y K L I L N G K Q A K E E A I K E L V D A A I A E K Y F K L Y A N A K T V E G V W T L K D E I L T F T V T E
BINARY_4453 T T Y K L I L N G K Q A K E E A I K E L V D A A I A E K Y F K L Y A N A K T V E G V W T L K D E I K K A T V T E
BINARY_4454 T T Y K L I L N G K Q A K E E A I K E L V D A A I A E K Y F K L Y A N A K T V E G V W T L K D E I K K F T V T E
BINARY_4455 T T Y K L I L N G K Q A K E E A I K E L V D A A I A E K Y F K L Y A N A K T V E G V W T L K D E I K T A T V T E
BINARY_4456 T T Y K L I L N G K Q A K E E A I K E L V D A A I A E K Y F K L Y A N A K T V E G V W T L K D E I K T F T V T E
BINARY_4457 T T Y K L I L N G K Q A K E E A I K E L V D A A I A E K Y F K L Y A N A K T V E G V W T L K D E T L K A T V T E
BINARY_4458 T T Y K L I L N G K Q A K E E A I K E L V D A A I A E K Y F K L Y A N A K T V E G V W T L K D E T L K F T V T E
BINARY_4459 T T Y K L I L N G K Q A K E E A I K E L V D A A I A E K Y F K L Y A N A K T V E G V W T L K D E T L T A T V T E
BINARY_4460 T T Y K L I L N G K Q A K E E A I K E L V D A A I A E K Y F K L Y A N A K T V E G V W T L K D E T L T F T V T E
BINARY_4461 T T Y K L I L N G K Q A K E E A I K E L V D A A I A E K Y F K L Y A N A K T V E G V W T L K D E T K K A T V T E
BINARY_4462 T T Y K L I L N G K Q A K E E A I K E L V D A A I A E K Y F K L Y A N A K T V E G V W T L K D E T K K F T V T E
BINARY_4463 T T Y K L I L N G K Q A K E E A I K E L V D A A I A E K Y F K L Y A N A K T V E G V W T L K D E T K T A T V T E
BINARY_4464 T T Y K L I L N G K Q A K E E A I K E L V D A A I A E K Y F K L Y A N A K T V E G V W T L K D E T K T F T V T E
BINARY_4465 T T Y K L I L N G K Q A K E E A I K E L V D A A I A E K Y F K L Y A N A K T V E G V W T Y K D E I L K A T V T E
BINARY_4466 T T Y K L I L N G K Q A K E E A I K E L V D A A I A E K Y F K L Y A N A K T V E G V W T Y K D E I L K F T V T E
BINARY_4467 T T Y K L I L N G K Q A K E E A I K E L V D A A I A E K Y F K L Y A N A K T V E G V W T Y K D E I L T A T V T E
BINARY_4468 T T Y K L I L N G K Q A K E E A I K E L V D A A I A E K Y F K L Y A N A K T V E G V W T Y K D E I L T F T V T E
BINARY_4469 T T Y K L I L N G K Q A K E E A I K E L V D A A I A E K Y F K L Y A N A K T V E G V W T Y K D E I K K A T V T E
BINARY_4470 T T Y K L I L N G K Q A K E E A I K E L V D A A I A E K Y F K L Y A N A K T V E G V W T Y K D E I K K F T V T E
BINARY_4471 T T Y K L I L N G K Q A K E E A I K E L V D A A I A E K Y F K L Y A N A K T V E G V W T Y K D E I K T A T V T E
BINARY_4472 T T Y K L I L N G K Q A K E E A I K E L V D A A I A E K Y F K L Y A N A K T V E G V W T Y K D E I K T F T V T E
BINARY_4473 T T Y K L I L N G K Q A K E E A I K E L V D A A I A E K Y F K L Y A N A K T V E G V W T Y K D E T L K A T V T E
BINARY_4474 T T Y K L I L N G K Q A K E E A I K E L V D A A I A E K Y F K L Y A N A K T V E G V W T Y K D E T L K F T V T E
BINARY_4475 T T Y K L I L N G K Q A K E E A I K E L V D A A I A E K Y F K L Y A N A K T V E G V W T Y K D E T L T A T V T E
BINARY_4476 T T Y K L I L N G K Q A K E E A I K E L V D A A I A E K Y F K L Y A N A K T V E G V W T Y K D E T L T F T V T E
BINARY_4477 T T Y K L I L N G K Q A K E E A I K E L V D A A I A E K Y F K L Y A N A K T V E G V W T Y K D E T K K A T V T E
BINARY_4478 T T Y K L I L N G K Q A K E E A I K E L V D A A I A E K Y F K L Y A N A K T V E G V W T Y K D E T K K F T V T E
BINARY_4479 T T Y K L I L N G K Q A K E E A I K E L V D A A I A E K Y F K L Y A N A K T V E G V W T Y K D E T K T A T V T E
BINARY_4480 T T Y K L I L N G K Q A K E E A I K E L V D A A I A E K Y F K L Y A N A K T V E G V W T Y K D E T K T F T V T E
BINARY_4481 T T Y K L I L N G K Q A K E E A I K E L V D A A T A E K Y I K L I A N A K T V E G V W T L K D E I L K A T V T E

BINARY_4482 T T Y K L I L N G K Q A K E E A I K E L V D A A T A E K Y I K L I A N A K T V E G V W T L K D E I L K F T V T E
BINARY_4483 T T Y K L I L N G K Q A K E E A I K E L V D A A T A E K Y I K L I A N A K T V E G V W T L K D E I L T A T V T E
BINARY_4484 T T Y K L I L N G K Q A K E E A I K E L V D A A T A E K Y I K L I A N A K T V E G V W T L K D E I L T F T V T E
BINARY_4485 T T Y K L I L N G K Q A K E E A I K E L V D A A T A E K Y I K L I A N A K T V E G V W T L K D E I K K A T V T E
BINARY_4486 T T Y K L I L N G K Q A K E E A I K E L V D A A T A E K Y I K L I A N A K T V E G V W T L K D E I K K F T V T E
BINARY_4487 T T Y K L I L N G K Q A K E E A I K E L V D A A T A E K Y I K L I A N A K T V E G V W T L K D E I K T A T V T E
BINARY_4488 T T Y K L I L N G K Q A K E E A I K E L V D A A T A E K Y I K L I A N A K T V E G V W T L K D E I K T F T V T E
BINARY_4489 T T Y K L I L N G K Q A K E E A I K E L V D A A T A E K Y I K L I A N A K T V E G V W T L K D E T L K A T V T E
BINARY_4490 T T Y K L I L N G K Q A K E E A I K E L V D A A T A E K Y I K L I A N A K T V E G V W T L K D E T L K F T V T E
BINARY_4491 T T Y K L I L N G K Q A K E E A I K E L V D A A T A E K Y I K L I A N A K T V E G V W T L K D E T L T A T V T E
BINARY_4492 T T Y K L I L N G K Q A K E E A I K E L V D A A T A E K Y I K L I A N A K T V E G V W T L K D E T L T F T V T E
BINARY_4493 T T Y K L I L N G K Q A K E E A I K E L V D A A T A E K Y I K L I A N A K T V E G V W T L K D E T K K A T V T E
BINARY_4494 T T Y K L I L N G K Q A K E E A I K E L V D A A T A E K Y I K L I A N A K T V E G V W T L K D E T K K F T V T E
BINARY_4495 T T Y K L I L N G K Q A K E E A I K E L V D A A T A E K Y I K L I A N A K T V E G V W T L K D E T K T A T V T E
BINARY_4496 T T Y K L I L N G K Q A K E E A I K E L V D A A T A E K Y I K L I A N A K T V E G V W T L K D E T K T F T V T E
BINARY_4497 T T Y K L I L N G K Q A K E E A I K E L V D A A T A E K Y I K L I A N A K T V E G V W T Y K D E I L K A T V T E
BINARY_4498 T T Y K L I L N G K Q A K E E A I K E L V D A A T A E K Y I K L I A N A K T V E G V W T Y K D E I L K F T V T E
BINARY_4499 T T Y K L I L N G K Q A K E E A I K E L V D A A T A E K Y I K L I A N A K T V E G V W T Y K D E I L T A T V T E
BINARY_4500 T T Y K L I L N G K Q A K E E A I K E L V D A A T A E K Y I K L I A N A K T V E G V W T Y K D E I L T F T V T E
BINARY_4501 T T Y K L I L N G K Q A K E E A I K E L V D A A T A E K Y I K L I A N A K T V E G V W T Y K D E I K K A T V T E
BINARY_4502 T T Y K L I L N G K Q A K E E A I K E L V D A A T A E K Y I K L I A N A K T V E G V W T Y K D E I K K F T V T E
BINARY_4503 T T Y K L I L N G K Q A K E E A I K E L V D A A T A E K Y I K L I A N A K T V E G V W T Y K D E I K T A T V T E
BINARY_4504 T T Y K L I L N G K Q A K E E A I K E L V D A A T A E K Y I K L I A N A K T V E G V W T Y K D E I K T F T V T E
BINARY_4505 T T Y K L I L N G K Q A K E E A I K E L V D A A T A E K Y I K L I A N A K T V E G V W T Y K D E T L K A T V T E
BINARY_4506 T T Y K L I L N G K Q A K E E A I K E L V D A A T A E K Y I K L I A N A K T V E G V W T Y K D E T L K F T V T E
BINARY_4507 T T Y K L I L N G K Q A K E E A I K E L V D A A T A E K Y I K L I A N A K T V E G V W T Y K D E T L T A T V T E
BINARY_4508 T T Y K L I L N G K Q A K E E A I K E L V D A A T A E K Y I K L I A N A K T V E G V W T Y K D E T L T F T V T E
BINARY_4509 T T Y K L I L N G K Q A K E E A I K E L V D A A T A E K Y I K L I A N A K T V E G V W T Y K D E T K K A T V T E
BINARY_4510 T T Y K L I L N G K Q A K E E A I K E L V D A A T A E K Y I K L I A N A K T V E G V W T Y K D E T K K F T V T E
BINARY_4511 T T Y K L I L N G K Q A K E E A I K E L V D A A T A E K Y I K L I A N A K T V E G V W T Y K D E T K T A T V T E
BINARY_4512 T T Y K L I L N G K Q A K E E A I K E L V D A A T A E K Y I K L I A N A K T V E G V W T Y K D E T K T F T V T E
BINARY_4513 T T Y K L I L N G K Q A K E E A I K E L V D A A T A E K Y I K L Y A N A K T V E G V W T L K D E I L K A T V T E
BINARY_4514 T T Y K L I L N G K Q A K E E A I K E L V D A A T A E K Y I K L Y A N A K T V E G V W T L K D E I L K F T V T E
BINARY_4515 T T Y K L I L N G K Q A K E E A I K E L V D A A T A E K Y I K L Y A N A K T V E G V W T L K D E I L T A T V T E
BINARY_4516 T T Y K L I L N G K Q A K E E A I K E L V D A A T A E K Y I K L Y A N A K T V E G V W T L K D E I L T F T V T E
BINARY_4517 T T Y K L I L N G K Q A K E E A I K E L V D A A T A E K Y I K L Y A N A K T V E G V W T L K D E I K K A T V T E
BINARY_4518 T T Y K L I L N G K Q A K E E A I K E L V D A A T A E K Y I K L Y A N A K T V E G V W T L K D E I K K F T V T E
BINARY_4519 T T Y K L I L N G K Q A K E E A I K E L V D A A T A E K Y I K L Y A N A K T V E G V W T L K D E I K T A T V T E
BINARY_4520 T T Y K L I L N G K Q A K E E A I K E L V D A A T A E K Y I K L Y A N A K T V E G V W T L K D E I K T F T V T E
BINARY_4521 T T Y K L I L N G K Q A K E E A I K E L V D A A T A E K Y I K L Y A N A K T V E G V W T L K D E T L K A T V T E
BINARY_4522 T T Y K L I L N G K Q A K E E A I K E L V D A A T A E K Y I K L Y A N A K T V E G V W T L K D E T L K F T V T E
BINARY_4523 T T Y K L I L N G K Q A K E E A I K E L V D A A T A E K Y I K L Y A N A K T V E G V W T L K D E T L T A T V T E
BINARY_4524 T T Y K L I L N G K Q A K E E A I K E L V D A A T A E K Y I K L Y A N A K T V E G V W T L K D E T L T F T V T E
BINARY_4525 T T Y K L I L N G K Q A K E E A I K E L V D A A T A E K Y I K L Y A N A K T V E G V W T L K D E T K K A T V T E
BINARY_4526 T T Y K L I L N G K Q A K E E A I K E L V D A A T A E K Y I K L Y A N A K T V E G V W T L K D E T K K F T V T E
BINARY_4527 T T Y K L I L N G K Q A K E E A I K E L V D A A T A E K Y I K L Y A N A K T V E G V W T L K D E T K T A T V T E
BINARY_4528 T T Y K L I L N G K Q A K E E A I K E L V D A A T A E K Y I K L Y A N A K T V E G V W T L K D E T K T F T V T E
BINARY_4529 T T Y K L I L N G K Q A K E E A I K E L V D A A T A E K Y I K L Y A N A K T V E G V W T Y K D E I L K A T V T E
BINARY_4530 T T Y K L I L N G K Q A K E E A I K E L V D A A T A E K Y I K L Y A N A K T V E G V W T Y K D E I L K F T V T E
BINARY_4531 T T Y K L I L N G K Q A K E E A I K E L V D A A T A E K Y I K L Y A N A K T V E G V W T Y K D E I L T A T V T E
BINARY_4532 T T Y K L I L N G K Q A K E E A I K E L V D A A T A E K Y I K L Y A N A K T V E G V W T Y K D E I L T F T V T E
BINARY_4533 T T Y K L I L N G K Q A K E E A I K E L V D A A T A E K Y I K L Y A N A K T V E G V W T Y K D E I K K A T V T E
BINARY_4534 T T Y K L I L N G K Q A K E E A I K E L V D A A T A E K Y I K L Y A N A K T V E G V W T Y K D E I K K F T V T E
BINARY_4535 T T Y K L I L N G K Q A K E E A I K E L V D A A T A E K Y I K L Y A N A K T V E G V W T Y K D E I K T A T V T E
BINARY_4536 T T Y K L I L N G K Q A K E E A I K E L V D A A T A E K Y I K L Y A N A K T V E G V W T Y K D E I K T F T V T E
BINARY_4537 T T Y K L I L N G K Q A K E E A I K E L V D A A T A E K Y I K L Y A N A K T V E G V W T Y K D E T L K A T V T E
BINARY_4538 T T Y K L I L N G K Q A K E E A I K E L V D A A T A E K Y I K L Y A N A K T V E G V W T Y K D E T L K F T V T E
BINARY_4539 T T Y K L I L N G K Q A K E E A I K E L V D A A T A E K Y I K L Y A N A K T V E G V W T Y K D E T L T A T V T E
BINARY_4540 T T Y K L I L N G K Q A K E E A I K E L V D A A T A E K Y I K L Y A N A K T V E G V W T Y K D E T L T F T V T E
BINARY_4541 T T Y K L I L N G K Q A K E E A I K E L V D A A T A E K Y I K L Y A N A K T V E G V W T Y K D E T K K A T V T E
BINARY_4542 T T Y K L I L N G K Q A K E E A I K E L V D A A T A E K Y I K L Y A N A K T V E G V W T Y K D E T K K F T V T E
BINARY_4543 T T Y K L I L N G K Q A K E E A I K E L V D A A T A E K Y I K L Y A N A K T V E G V W T Y K D E T K T A T V T E
BINARY_4544 T T Y K L I L N G K Q A K E E A I K E L V D A A T A E K Y I K L Y A N A K T V E G V W T Y K D E T K T F T V T E
BINARY_4545 T T Y K L I L N G K Q A K E E A I K E L V D A A T A E K Y F K L I A N A K T V E G V W T L K D E I L K A T V T E
BINARY_4546 T T Y K L I L N G K Q A K E E A I K E L V D A A T A E K Y F K L I A N A K T V E G V W T L K D E I L K F T V T E
BINARY_4547 T T Y K L I L N G K Q A K E E A I K E L V D A A T A E K Y F K L I A N A K T V E G V W T L K D E I L T A T V T E
BINARY_4548 T T Y K L I L N G K Q A K E E A I K E L V D A A T A E K Y F K L I A N A K T V E G V W T L K D E I L T F T V T E
BINARY_4549 T T Y K L I L N G K Q A K E E A I K E L V D A A T A E K Y F K L I A N A K T V E G V W T L K D E I K K A T V T E
BINARY_4550 T T Y K L I L N G K Q A K E E A I K E L V D A A T A E K Y F K L I A N A K T V E G V W T L K D E I K K F T V T E
BINARY_4551 T T Y K L I L N G K Q A K E E A I K E L V D A A T A E K Y F K L I A N A K T V E G V W T L K D E I K T A T V T E
BINARY_4552 T T Y K L I L N G K Q A K E E A I K E L V D A A T A E K Y F K L I A N A K T V E G V W T L K D E I K T F T V T E
BINARY_4553 T T Y K L I L N G K Q A K E E A I K E L V D A A T A E K Y F K L I A N A K T V E G V W T L K D E T L K A T V T E
BINARY_4554 T T Y K L I L N G K Q A K E E A I K E L V D A A T A E K Y F K L I A N A K T V E G V W T L K D E T L K F T V T E
BINARY_4555 T T Y K L I L N G K Q A K E E A I K E L V D A A T A E K Y F K L I A N A K T V E G V W T L K D E T L T A T V T E
BINARY_4556 T T Y K L I L N G K Q A K E E A I K E L V D A A T A E K Y F K L I A N A K T V E G V W T L K D E T L T F T V T E
BINARY_4557 T T Y K L I L N G K Q A K E E A I K E L V D A A T A E K Y F K L I A N A K T V E G V W T L K D E T K K A T V T E
BINARY_4558 T T Y K L I L N G K Q A K E E A I K E L V D A A T A E K Y F K L I A N A K T V E G V W T L K D E T K K F T V T E
BINARY_4559 T T Y K L I L N G K Q A K E E A I K E L V D A A T A E K Y F K L I A N A K T V E G V W T L K D E T K T A T V T E
BINARY_4560 T T Y K L I L N G K Q A K E E A I K E L V D A A T A E K Y F K L I A N A K T V E G V W T L K D E T K T F T V T E
BINARY_4561 T T Y K L I L N G K Q A K E E A I K E L V D A A T A E K Y F K L I A N A K T V E G V W T Y K D E I L K A T V T E
BINARY_4562 T T Y K L I L N G K Q A K E E A I K E L V D A A T A E K Y F K L I A N A K T V E G V W T Y K D E I L K F T V T E
BINARY_4563 T T Y K L I L N G K Q A K E E A I K E L V D A A T A E K Y F K L I A N A K T V E G V W T Y K D E I L T A T V T E
BINARY_4564 T T Y K L I L N G K Q A K E E A I K E L V D A A T A E K Y F K L I A N A K T V E G V W T Y K D E I L T F T V T E
BINARY_4565 T T Y K L I L N G K Q A K E E A I K E L V D A A T A E K Y F K L I A N A K T V E G V W T Y K D E I K K A T V T E
BINARY_4566 T T Y K L I L N G K Q A K E E A I K E L V D A A T A E K Y F K L I A N A K T V E G V W T Y K D E I K K F T V T E
BINARY_4567 T T Y K L I L N G K Q A K E E A I K E L V D A A T A E K Y F K L I A N A K T V E G V W T Y K D E I K T A T V T E
BINARY_4568 T T Y K L I L N G K Q A K E E A I K E L V D A A T A E K Y F K L I A N A K T V E G V W T Y K D E I K T F T V T E
BINARY_4569 T T Y K L I L N G K Q A K E E A I K E L V D A A T A E K Y F K L I A N A K T V E G V W T Y K D E T L K A T V T E
BINARY_4570 T T Y K L I L N G K Q A K E E A I K E L V D A A T A E K Y F K L I A N A K T V E G V W T Y K D E T L K F T V T E
BINARY_4571 T T Y K L I L N G K Q A K E E A I K E L V D A A T A E K Y F K L I A N A K T V E G V W T Y K D E T L T A T V T E
BINARY_4572 T T Y K L I L N G K Q A K E E A I K E L V D A A T A E K Y F K L I A N A K T V E G V W T Y K D E T L T F T V T E
BINARY_4573 T T Y K L I L N G K Q A K E E A I K E L V D A A T A E K Y F K L I A N A K T V E G V W T Y K D E T K K A T V T E
BINARY_4574 T T Y K L I L N G K Q A K E E A I K E L V D A A T A E K Y F K L I A N A K T V E G V W T Y K D E T K K F T V T E
BINARY_4575 T T Y K L I L N G K Q A K E E A I K E L V D A A T A E K Y F K L I A N A K T V E G V W T Y K D E T K T A T V T E
BINARY_4576 T T Y K L I L N G K Q A K E E A I K E L V D A A T A E K Y F K L I A N A K T V E G V W T Y K D E T K T F T V T E

BINARY_4577 T T Y K L I L N G K Q A K E E A I K E L V D A A T A E K Y F K L Y A N A K T V E G V W T L K D E I L K A T V T E
BINARY_4578 T T Y K L I L N G K Q A K E E A I K E L V D A A T A E K Y F K L Y A N A K T V E G V W T L K D E I L K F T V T E
BINARY_4579 T T Y K L I L N G K Q A K E E A I K E L V D A A T A E K Y F K L Y A N A K T V E G V W T L K D E I L T A T V T E
BINARY_4580 T T Y K L I L N G K Q A K E E A I K E L V D A A T A E K Y F K L Y A N A K T V E G V W T L K D E I L T F T V T E
BINARY_4581 T T Y K L I L N G K Q A K E E A I K E L V D A A T A E K Y F K L Y A N A K T V E G V W T L K D E I K K A T V T E
BINARY_4582 T T Y K L I L N G K Q A K E E A I K E L V D A A T A E K Y F K L Y A N A K T V E G V W T L K D E I K K F T V T E
BINARY_4583 T T Y K L I L N G K Q A K E E A I K E L V D A A T A E K Y F K L Y A N A K T V E G V W T L K D E I K T A T V T E
BINARY_4584 T T Y K L I L N G K Q A K E E A I K E L V D A A T A E K Y F K L Y A N A K T V E G V W T L K D E I K T F T V T E
BINARY_4585 T T Y K L I L N G K Q A K E E A I K E L V D A A T A E K Y F K L Y A N A K T V E G V W T L K D E T L K A T V T E
BINARY_4586 T T Y K L I L N G K Q A K E E A I K E L V D A A T A E K Y F K L Y A N A K T V E G V W T L K D E T L K F T V T E
BINARY_4587 T T Y K L I L N G K Q A K E E A I K E L V D A A T A E K Y F K L Y A N A K T V E G V W T L K D E T L T A T V T E
BINARY_4588 T T Y K L I L N G K Q A K E E A I K E L V D A A T A E K Y F K L Y A N A K T V E G V W T L K D E T L T F T V T E
BINARY_4589 T T Y K L I L N G K Q A K E E A I K E L V D A A T A E K Y F K L Y A N A K T V E G V W T L K D E T K K A T V T E
BINARY_4590 T T Y K L I L N G K Q A K E E A I K E L V D A A T A E K Y F K L Y A N A K T V E G V W T L K D E T K K F T V T E
BINARY_4591 T T Y K L I L N G K Q A K E E A I K E L V D A A T A E K Y F K L Y A N A K T V E G V W T L K D E T K T A T V T E
BINARY_4592 T T Y K L I L N G K Q A K E E A I K E L V D A A T A E K Y F K L Y A N A K T V E G V W T L K D E T K T F T V T E
BINARY_4593 T T Y K L I L N G K Q A K E E A I K E L V D A A T A E K Y F K L Y A N A K T V E G V W T Y K D E I L K A T V T E
BINARY_4594 T T Y K L I L N G K Q A K E E A I K E L V D A A T A E K Y F K L Y A N A K T V E G V W T Y K D E I L K F T V T E
BINARY_4595 T T Y K L I L N G K Q A K E E A I K E L V D A A T A E K Y F K L Y A N A K T V E G V W T Y K D E I L T A T V T E
BINARY_4596 T T Y K L I L N G K Q A K E E A I K E L V D A A T A E K Y F K L Y A N A K T V E G V W T Y K D E I L T F T V T E
BINARY_4597 T T Y K L I L N G K Q A K E E A I K E L V D A A T A E K Y F K L Y A N A K T V E G V W T Y K D E I K K A T V T E
BINARY_4598 T T Y K L I L N G K Q A K E E A I K E L V D A A T A E K Y F K L Y A N A K T V E G V W T Y K D E I K K F T V T E
BINARY_4599 T T Y K L I L N G K Q A K E E A I K E L V D A A T A E K Y F K L Y A N A K T V E G V W T Y K D E I K T A T V T E
BINARY_4600 T T Y K L I L N G K Q A K E E A I K E L V D A A T A E K Y F K L Y A N A K T V E G V W T Y K D E I K T F T V T E
BINARY_4601 T T Y K L I L N G K Q A K E E A I K E L V D A A T A E K Y F K L Y A N A K T V E G V W T Y K D E T L K A T V T E
BINARY_4602 T T Y K L I L N G K Q A K E E A I K E L V D A A T A E K Y F K L Y A N A K T V E G V W T Y K D E T L K F T V T E
BINARY_4603 T T Y K L I L N G K Q A K E E A I K E L V D A A T A E K Y F K L Y A N A K T V E G V W T Y K D E T L T A T V T E
BINARY_4604 T T Y K L I L N G K Q A K E E A I K E L V D A A T A E K Y F K L Y A N A K T V E G V W T Y K D E T L T F T V T E
BINARY_4605 T T Y K L I L N G K Q A K E E A I K E L V D A A T A E K Y F K L Y A N A K T V E G V W T Y K D E T K K A T V T E
BINARY_4606 T T Y K L I L N G K Q A K E E A I K E L V D A A T A E K Y F K L Y A N A K T V E G V W T Y K D E T K K F T V T E
BINARY_4607 T T Y K L I L N G K Q A K E E A I K E L V D A A T A E K Y F K L Y A N A K T V E G V W T Y K D E T K T A T V T E
BINARY_4608 T T Y K L I L N G K Q A K E E A I K E L V D A A T A E K Y F K L Y A N A K T V E G V W T Y K D E T K T F T V T E
BINARY_4609 T T Y K L I L N G K Q A K E E A I K E A V D A G I A E K Y I K L I A N A K T V E G V W T L K D E I L K A T V T E
BINARY_4610 T T Y K L I L N G K Q A K E E A I K E A V D A G I A E K Y I K L I A N A K T V E G V W T L K D E I L K F T V T E
BINARY_4611 T T Y K L I L N G K Q A K E E A I K E A V D A G I A E K Y I K L I A N A K T V E G V W T L K D E I L T A T V T E
BINARY_4612 T T Y K L I L N G K Q A K E E A I K E A V D A G I A E K Y I K L I A N A K T V E G V W T L K D E I L T F T V T E
BINARY_4613 T T Y K L I L N G K Q A K E E A I K E A V D A G I A E K Y I K L I A N A K T V E G V W T L K D E I K K A T V T E
BINARY_4614 T T Y K L I L N G K Q A K E E A I K E A V D A G I A E K Y I K L I A N A K T V E G V W T L K D E I K K F T V T E
BINARY_4615 T T Y K L I L N G K Q A K E E A I K E A V D A G I A E K Y I K L I A N A K T V E G V W T L K D E I K T A T V T E
BINARY_4616 T T Y K L I L N G K Q A K E E A I K E A V D A G I A E K Y I K L I A N A K T V E G V W T L K D E I K T F T V T E
BINARY_4617 T T Y K L I L N G K Q A K E E A I K E A V D A G I A E K Y I K L I A N A K T V E G V W T L K D E T L K A T V T E
BINARY_4618 T T Y K L I L N G K Q A K E E A I K E A V D A G I A E K Y I K L I A N A K T V E G V W T L K D E T L K F T V T E
BINARY_4619 T T Y K L I L N G K Q A K E E A I K E A V D A G I A E K Y I K L I A N A K T V E G V W T L K D E T L T A T V T E
BINARY_4620 T T Y K L I L N G K Q A K E E A I K E A V D A G I A E K Y I K L I A N A K T V E G V W T L K D E T L T F T V T E
BINARY_4621 T T Y K L I L N G K Q A K E E A I K E A V D A G I A E K Y I K L I A N A K T V E G V W T L K D E T K K A T V T E
BINARY_4622 T T Y K L I L N G K Q A K E E A I K E A V D A G I A E K Y I K L I A N A K T V E G V W T L K D E T K K F T V T E
BINARY_4623 T T Y K L I L N G K Q A K E E A I K E A V D A G I A E K Y I K L I A N A K T V E G V W T L K D E T K T A T V T E
BINARY_4624 T T Y K L I L N G K Q A K E E A I K E A V D A G I A E K Y I K L I A N A K T V E G V W T L K D E T K T F T V T E
BINARY_4625 T T Y K L I L N G K Q A K E E A I K E A V D A G I A E K Y I K L I A N A K T V E G V W T Y K D E I L K A T V T E
BINARY_4626 T T Y K L I L N G K Q A K E E A I K E A V D A G I A E K Y I K L I A N A K T V E G V W T Y K D E I L K F T V T E
BINARY_4627 T T Y K L I L N G K Q A K E E A I K E A V D A G I A E K Y I K L I A N A K T V E G V W T Y K D E I L T A T V T E
BINARY_4628 T T Y K L I L N G K Q A K E E A I K E A V D A G I A E K Y I K L I A N A K T V E G V W T Y K D E I L T F T V T E
BINARY_4629 T T Y K L I L N G K Q A K E E A I K E A V D A G I A E K Y I K L I A N A K T V E G V W T Y K D E I K K A T V T E
BINARY_4630 T T Y K L I L N G K Q A K E E A I K E A V D A G I A E K Y I K L I A N A K T V E G V W T Y K D E I K K F T V T E
BINARY_4631 T T Y K L I L N G K Q A K E E A I K E A V D A G I A E K Y I K L I A N A K T V E G V W T Y K D E I K T A T V T E
BINARY_4632 T T Y K L I L N G K Q A K E E A I K E A V D A G I A E K Y I K L I A N A K T V E G V W T Y K D E I K T F T V T E
BINARY_4633 T T Y K L I L N G K Q A K E E A I K E A V D A G I A E K Y I K L I A N A K T V E G V W T Y K D E T L K A T V T E
BINARY_4634 T T Y K L I L N G K Q A K E E A I K E A V D A G I A E K Y I K L I A N A K T V E G V W T Y K D E T L K F T V T E
BINARY_4635 T T Y K L I L N G K Q A K E E A I K E A V D A G I A E K Y I K L I A N A K T V E G V W T Y K D E T L T A T V T E
BINARY_4636 T T Y K L I L N G K Q A K E E A I K E A V D A G I A E K Y I K L I A N A K T V E G V W T Y K D E T L T F T V T E
BINARY_4637 T T Y K L I L N G K Q A K E E A I K E A V D A G I A E K Y I K L I A N A K T V E G V W T Y K D E T K K A T V T E
BINARY_4638 T T Y K L I L N G K Q A K E E A I K E A V D A G I A E K Y I K L I A N A K T V E G V W T Y K D E T K K F T V T E
BINARY_4639 T T Y K L I L N G K Q A K E E A I K E A V D A G I A E K Y I K L I A N A K T V E G V W T Y K D E T K T A T V T E
BINARY_4640 T T Y K L I L N G K Q A K E E A I K E A V D A G I A E K Y I K L I A N A K T V E G V W T Y K D E T K T F T V T E
BINARY_4641 T T Y K L I L N G K Q A K E E A I K E A V D A G I A E K Y I K L Y A N A K T V E G V W T L K D E I L K A T V T E
BINARY_4642 T T Y K L I L N G K Q A K E E A I K E A V D A G I A E K Y I K L Y A N A K T V E G V W T L K D E I L K F T V T E
BINARY_4643 T T Y K L I L N G K Q A K E E A I K E A V D A G I A E K Y I K L Y A N A K T V E G V W T L K D E I L T A T V T E
BINARY_4644 T T Y K L I L N G K Q A K E E A I K E A V D A G I A E K Y I K L Y A N A K T V E G V W T L K D E I L T F T V T E
BINARY_4645 T T Y K L I L N G K Q A K E E A I K E A V D A G I A E K Y I K L Y A N A K T V E G V W T L K D E I K K A T V T E
BINARY_4646 T T Y K L I L N G K Q A K E E A I K E A V D A G I A E K Y I K L Y A N A K T V E G V W T L K D E I K K F T V T E
BINARY_4647 T T Y K L I L N G K Q A K E E A I K E A V D A G I A E K Y I K L Y A N A K T V E G V W T L K D E I K T A T V T E
BINARY_4648 T T Y K L I L N G K Q A K E E A I K E A V D A G I A E K Y I K L Y A N A K T V E G V W T L K D E I K T F T V T E
BINARY_4649 T T Y K L I L N G K Q A K E E A I K E A V D A G I A E K Y I K L Y A N A K T V E G V W T L K D E T L K A T V T E
BINARY_4650 T T Y K L I L N G K Q A K E E A I K E A V D A G I A E K Y I K L Y A N A K T V E G V W T L K D E T L K F T V T E
BINARY_4651 T T Y K L I L N G K Q A K E E A I K E A V D A G I A E K Y I K L Y A N A K T V E G V W T L K D E T L T A T V T E
BINARY_4652 T T Y K L I L N G K Q A K E E A I K E A V D A G I A E K Y I K L Y A N A K T V E G V W T L K D E T L T F T V T E
BINARY_4653 T T Y K L I L N G K Q A K E E A I K E A V D A G I A E K Y I K L Y A N A K T V E G V W T L K D E T K K A T V T E
BINARY_4654 T T Y K L I L N G K Q A K E E A I K E A V D A G I A E K Y I K L Y A N A K T V E G V W T L K D E T K K F T V T E
BINARY_4655 T T Y K L I L N G K Q A K E E A I K E A V D A G I A E K Y I K L Y A N A K T V E G V W T L K D E T K T A T V T E
BINARY_4656 T T Y K L I L N G K Q A K E E A I K E A V D A G I A E K Y I K L Y A N A K T V E G V W T L K D E T K T F T V T E
BINARY_4657 T T Y K L I L N G K Q A K E E A I K E A V D A G I A E K Y I K L Y A N A K T V E G V W T Y K D E I L K A T V T E
BINARY_4658 T T Y K L I L N G K Q A K E E A I K E A V D A G I A E K Y I K L Y A N A K T V E G V W T Y K D E I L K F T V T E
BINARY_4659 T T Y K L I L N G K Q A K E E A I K E A V D A G I A E K Y I K L Y A N A K T V E G V W T Y K D E I L T A T V T E
BINARY_4660 T T Y K L I L N G K Q A K E E A I K E A V D A G I A E K Y I K L Y A N A K T V E G V W T Y K D E I L T F T V T E
BINARY_4661 T T Y K L I L N G K Q A K E E A I K E A V D A G I A E K Y I K L Y A N A K T V E G V W T Y K D E I K K A T V T E
BINARY_4662 T T Y K L I L N G K Q A K E E A I K E A V D A G I A E K Y I K L Y A N A K T V E G V W T Y K D E I K K F T V T E
BINARY_4663 T T Y K L I L N G K Q A K E E A I K E A V D A G I A E K Y I K L Y A N A K T V E G V W T Y K D E I K T A T V T E
BINARY_4664 T T Y K L I L N G K Q A K E E A I K E A V D A G I A E K Y I K L Y A N A K T V E G V W T Y K D E I K T F T V T E
BINARY_4665 T T Y K L I L N G K Q A K E E A I K E A V D A G I A E K Y I K L Y A N A K T V E G V W T Y K D E T L K A T V T E
BINARY_4666 T T Y K L I L N G K Q A K E E A I K E A V D A G I A E K Y I K L Y A N A K T V E G V W T Y K D E T L K F T V T E
BINARY_4667 T T Y K L I L N G K Q A K E E A I K E A V D A G I A E K Y I K L Y A N A K T V E G V W T Y K D E T L T A T V T E
BINARY_4668 T T Y K L I L N G K Q A K E E A I K E A V D A G I A E K Y I K L Y A N A K T V E G V W T Y K D E T L T F T V T E
BINARY_4669 T T Y K L I L N G K Q A K E E A I K E A V D A G I A E K Y I K L Y A N A K T V E G V W T Y K D E T K K A T V T E
BINARY_4670 T T Y K L I L N G K Q A K E E A I K E A V D A G I A E K Y I K L Y A N A K T V E G V W T Y K D E T K K F T V T E
BINARY_4671 T T Y K L I L N G K Q A K E E A I K E A V D A G I A E K Y I K L Y A N A K T V E G V W T Y K D E T K T A T V T E

BINARY_4672 T T Y K L I L N G K Q A K E E A I K E A V D A G I A E K Y I K L Y A N A K T V E G V W T Y K D E T K T F T V T E
BINARY_4673 T T Y K L I L N G K Q A K E E A I K E A V D A G I A E K Y F K L I A N A K T V E G V W T L K D E I L K A T V T E
BINARY_4674 T T Y K L I L N G K Q A K E E A I K E A V D A G I A E K Y F K L I A N A K T V E G V W T L K D E I L K F T V T E
BINARY_4675 T T Y K L I L N G K Q A K E E A I K E A V D A G I A E K Y F K L I A N A K T V E G V W T L K D E I L T A T V T E
BINARY_4676 T T Y K L I L N G K Q A K E E A I K E A V D A G I A E K Y F K L I A N A K T V E G V W T L K D E I L T F T V T E
BINARY_4677 T T Y K L I L N G K Q A K E E A I K E A V D A G I A E K Y F K L I A N A K T V E G V W T L K D E I K K A T V T E
BINARY_4678 T T Y K L I L N G K Q A K E E A I K E A V D A G I A E K Y F K L I A N A K T V E G V W T L K D E I K K F T V T E
BINARY_4679 T T Y K L I L N G K Q A K E E A I K E A V D A G I A E K Y F K L I A N A K T V E G V W T L K D E I K T A T V T E
BINARY_4680 T T Y K L I L N G K Q A K E E A I K E A V D A G I A E K Y F K L I A N A K T V E G V W T L K D E I K T F T V T E
BINARY_4681 T T Y K L I L N G K Q A K E E A I K E A V D A G I A E K Y F K L I A N A K T V E G V W T L K D E T L K A T V T E
BINARY_4682 T T Y K L I L N G K Q A K E E A I K E A V D A G I A E K Y F K L I A N A K T V E G V W T L K D E T L K F T V T E
BINARY_4683 T T Y K L I L N G K Q A K E E A I K E A V D A G I A E K Y F K L I A N A K T V E G V W T L K D E T L T A T V T E
BINARY_4684 T T Y K L I L N G K Q A K E E A I K E A V D A G I A E K Y F K L I A N A K T V E G V W T L K D E T L T F T V T E
BINARY_4685 T T Y K L I L N G K Q A K E E A I K E A V D A G I A E K Y F K L I A N A K T V E G V W T L K D E T K K A T V T E
BINARY_4686 T T Y K L I L N G K Q A K E E A I K E A V D A G I A E K Y F K L I A N A K T V E G V W T L K D E T K K F T V T E
BINARY_4687 T T Y K L I L N G K Q A K E E A I K E A V D A G I A E K Y F K L I A N A K T V E G V W T L K D E T K T A T V T E
BINARY_4688 T T Y K L I L N G K Q A K E E A I K E A V D A G I A E K Y F K L I A N A K T V E G V W T L K D E T K T F T V T E
BINARY_4689 T T Y K L I L N G K Q A K E E A I K E A V D A G I A E K Y F K L I A N A K T V E G V W T Y K D E I L K A T V T E
BINARY_4690 T T Y K L I L N G K Q A K E E A I K E A V D A G I A E K Y F K L I A N A K T V E G V W T Y K D E I L K F T V T E
BINARY_4691 T T Y K L I L N G K Q A K E E A I K E A V D A G I A E K Y F K L I A N A K T V E G V W T Y K D E I L T A T V T E
BINARY_4692 T T Y K L I L N G K Q A K E E A I K E A V D A G I A E K Y F K L I A N A K T V E G V W T Y K D E I L T F T V T E
BINARY_4693 T T Y K L I L N G K Q A K E E A I K E A V D A G I A E K Y F K L I A N A K T V E G V W T Y K D E I K K A T V T E
BINARY_4694 T T Y K L I L N G K Q A K E E A I K E A V D A G I A E K Y F K L I A N A K T V E G V W T Y K D E I K K F T V T E
BINARY_4695 T T Y K L I L N G K Q A K E E A I K E A V D A G I A E K Y F K L I A N A K T V E G V W T Y K D E I K T A T V T E
BINARY_4696 T T Y K L I L N G K Q A K E E A I K E A V D A G I A E K Y F K L I A N A K T V E G V W T Y K D E I K T F T V T E
BINARY_4697 T T Y K L I L N G K Q A K E E A I K E A V D A G I A E K Y F K L I A N A K T V E G V W T Y K D E T L K A T V T E
BINARY_4698 T T Y K L I L N G K Q A K E E A I K E A V D A G I A E K Y F K L I A N A K T V E G V W T Y K D E T L K F T V T E
BINARY_4699 T T Y K L I L N G K Q A K E E A I K E A V D A G I A E K Y F K L I A N A K T V E G V W T Y K D E T L T A T V T E
BINARY_4700 T T Y K L I L N G K Q A K E E A I K E A V D A G I A E K Y F K L I A N A K T V E G V W T Y K D E T L T F T V T E
BINARY_4701 T T Y K L I L N G K Q A K E E A I K E A V D A G I A E K Y F K L I A N A K T V E G V W T Y K D E T K K A T V T E
BINARY_4702 T T Y K L I L N G K Q A K E E A I K E A V D A G I A E K Y F K L I A N A K T V E G V W T Y K D E T K K F T V T E
BINARY_4703 T T Y K L I L N G K Q A K E E A I K E A V D A G I A E K Y F K L I A N A K T V E G V W T Y K D E T K T A T V T E
BINARY_4704 T T Y K L I L N G K Q A K E E A I K E A V D A G I A E K Y F K L I A N A K T V E G V W T Y K D E T K T F T V T E
BINARY_4705 T T Y K L I L N G K Q A K E E A I K E A V D A G I A E K Y F K L Y A N A K T V E G V W T L K D E I L K A T V T E
BINARY_4706 T T Y K L I L N G K Q A K E E A I K E A V D A G I A E K Y F K L Y A N A K T V E G V W T L K D E I L K F T V T E
BINARY_4707 T T Y K L I L N G K Q A K E E A I K E A V D A G I A E K Y F K L Y A N A K T V E G V W T L K D E I L T A T V T E
BINARY_4708 T T Y K L I L N G K Q A K E E A I K E A V D A G I A E K Y F K L Y A N A K T V E G V W T L K D E I L T F T V T E
BINARY_4709 T T Y K L I L N G K Q A K E E A I K E A V D A G I A E K Y F K L Y A N A K T V E G V W T L K D E I K K A T V T E
BINARY_4710 T T Y K L I L N G K Q A K E E A I K E A V D A G I A E K Y F K L Y A N A K T V E G V W T L K D E I K K F T V T E
BINARY_4711 T T Y K L I L N G K Q A K E E A I K E A V D A G I A E K Y F K L Y A N A K T V E G V W T L K D E I K T A T V T E
BINARY_4712 T T Y K L I L N G K Q A K E E A I K E A V D A G I A E K Y F K L Y A N A K T V E G V W T L K D E I K T F T V T E
BINARY_4713 T T Y K L I L N G K Q A K E E A I K E A V D A G I A E K Y F K L Y A N A K T V E G V W T L K D E T L K A T V T E
BINARY_4714 T T Y K L I L N G K Q A K E E A I K E A V D A G I A E K Y F K L Y A N A K T V E G V W T L K D E T L K F T V T E
BINARY_4715 T T Y K L I L N G K Q A K E E A I K E A V D A G I A E K Y F K L Y A N A K T V E G V W T L K D E T L T A T V T E
BINARY_4716 T T Y K L I L N G K Q A K E E A I K E A V D A G I A E K Y F K L Y A N A K T V E G V W T L K D E T L T F T V T E
BINARY_4717 T T Y K L I L N G K Q A K E E A I K E A V D A G I A E K Y F K L Y A N A K T V E G V W T L K D E T K K A T V T E
BINARY_4718 T T Y K L I L N G K Q A K E E A I K E A V D A G I A E K Y F K L Y A N A K T V E G V W T L K D E T K K F T V T E
BINARY_4719 T T Y K L I L N G K Q A K E E A I K E A V D A G I A E K Y F K L Y A N A K T V E G V W T L K D E T K T A T V T E
BINARY_4720 T T Y K L I L N G K Q A K E E A I K E A V D A G I A E K Y F K L Y A N A K T V E G V W T L K D E T K T F T V T E
BINARY_4721 T T Y K L I L N G K Q A K E E A I K E A V D A G I A E K Y F K L Y A N A K T V E G V W T Y K D E I L K A T V T E
BINARY_4722 T T Y K L I L N G K Q A K E E A I K E A V D A G I A E K Y F K L Y A N A K T V E G V W T Y K D E I L K F T V T E
BINARY_4723 T T Y K L I L N G K Q A K E E A I K E A V D A G I A E K Y F K L Y A N A K T V E G V W T Y K D E I L T A T V T E
BINARY_4724 T T Y K L I L N G K Q A K E E A I K E A V D A G I A E K Y F K L Y A N A K T V E G V W T Y K D E I L T F T V T E
BINARY_4725 T T Y K L I L N G K Q A K E E A I K E A V D A G I A E K Y F K L Y A N A K T V E G V W T Y K D E I K K A T V T E
BINARY_4726 T T Y K L I L N G K Q A K E E A I K E A V D A G I A E K Y F K L Y A N A K T V E G V W T Y K D E I K K F T V T E
BINARY_4727 T T Y K L I L N G K Q A K E E A I K E A V D A G I A E K Y F K L Y A N A K T V E G V W T Y K D E I K T A T V T E
BINARY_4728 T T Y K L I L N G K Q A K E E A I K E A V D A G I A E K Y F K L Y A N A K T V E G V W T Y K D E I K T F T V T E
BINARY_4729 T T Y K L I L N G K Q A K E E A I K E A V D A G I A E K Y F K L Y A N A K T V E G V W T Y K D E T L K A T V T E
BINARY_4730 T T Y K L I L N G K Q A K E E A I K E A V D A G I A E K Y F K L Y A N A K T V E G V W T Y K D E T L K F T V T E
BINARY_4731 T T Y K L I L N G K Q A K E E A I K E A V D A G I A E K Y F K L Y A N A K T V E G V W T Y K D E T L T A T V T E
BINARY_4732 T T Y K L I L N G K Q A K E E A I K E A V D A G I A E K Y F K L Y A N A K T V E G V W T Y K D E T L T F T V T E
BINARY_4733 T T Y K L I L N G K Q A K E E A I K E A V D A G I A E K Y F K L Y A N A K T V E G V W T Y K D E T K K A T V T E
BINARY_4734 T T Y K L I L N G K Q A K E E A I K E A V D A G I A E K Y F K L Y A N A K T V E G V W T Y K D E T K K F T V T E
BINARY_4735 T T Y K L I L N G K Q A K E E A I K E A V D A G I A E K Y F K L Y A N A K T V E G V W T Y K D E T K T A T V T E
BINARY_4736 T T Y K L I L N G K Q A K E E A I K E A V D A G I A E K Y F K L Y A N A K T V E G V W T Y K D E T K T F T V T E
BINARY_4737 T T Y K L I L N G K Q A K E E A I K E A V D A G T A E K Y I K L I A N A K T V E G V W T L K D E I L K A T V T E
BINARY_4738 T T Y K L I L N G K Q A K E E A I K E A V D A G T A E K Y I K L I A N A K T V E G V W T L K D E I L K F T V T E
BINARY_4739 T T Y K L I L N G K Q A K E E A I K E A V D A G T A E K Y I K L I A N A K T V E G V W T L K D E I L T A T V T E
BINARY_4740 T T Y K L I L N G K Q A K E E A I K E A V D A G T A E K Y I K L I A N A K T V E G V W T L K D E I L T F T V T E
BINARY_4741 T T Y K L I L N G K Q A K E E A I K E A V D A G T A E K Y I K L I A N A K T V E G V W T L K D E I K K A T V T E
BINARY_4742 T T Y K L I L N G K Q A K E E A I K E A V D A G T A E K Y I K L I A N A K T V E G V W T L K D E I K K F T V T E
BINARY_4743 T T Y K L I L N G K Q A K E E A I K E A V D A G T A E K Y I K L I A N A K T V E G V W T L K D E I K T A T V T E
BINARY_4744 T T Y K L I L N G K Q A K E E A I K E A V D A G T A E K Y I K L I A N A K T V E G V W T L K D E I K T F T V T E
BINARY_4745 T T Y K L I L N G K Q A K E E A I K E A V D A G T A E K Y I K L I A N A K T V E G V W T L K D E T L K A T V T E
BINARY_4746 T T Y K L I L N G K Q A K E E A I K E A V D A G T A E K Y I K L I A N A K T V E G V W T L K D E T L K F T V T E
BINARY_4747 T T Y K L I L N G K Q A K E E A I K E A V D A G T A E K Y I K L I A N A K T V E G V W T L K D E T L T A T V T E
BINARY_4748 T T Y K L I L N G K Q A K E E A I K E A V D A G T A E K Y I K L I A N A K T V E G V W T L K D E T L T F T V T E
BINARY_4749 T T Y K L I L N G K Q A K E E A I K E A V D A G T A E K Y I K L I A N A K T V E G V W T L K D E T K K A T V T E
BINARY_4750 T T Y K L I L N G K Q A K E E A I K E A V D A G T A E K Y I K L I A N A K T V E G V W T L K D E T K K F T V T E
BINARY_4751 T T Y K L I L N G K Q A K E E A I K E A V D A G T A E K Y I K L I A N A K T V E G V W T L K D E T K T A T V T E
BINARY_4752 T T Y K L I L N G K Q A K E E A I K E A V D A G T A E K Y I K L I A N A K T V E G V W T L K D E T K T F T V T E
BINARY_4753 T T Y K L I L N G K Q A K E E A I K E A V D A G T A E K Y I K L I A N A K T V E G V W T Y K D E I L K A T V T E
BINARY_4754 T T Y K L I L N G K Q A K E E A I K E A V D A G T A E K Y I K L I A N A K T V E G V W T Y K D E I L K F T V T E
BINARY_4755 T T Y K L I L N G K Q A K E E A I K E A V D A G T A E K Y I K L I A N A K T V E G V W T Y K D E I L T A T V T E
BINARY_4756 T T Y K L I L N G K Q A K E E A I K E A V D A G T A E K Y I K L I A N A K T V E G V W T Y K D E I L T F T V T E
BINARY_4757 T T Y K L I L N G K Q A K E E A I K E A V D A G T A E K Y I K L I A N A K T V E G V W T Y K D E I K K A T V T E
BINARY_4758 T T Y K L I L N G K Q A K E E A I K E A V D A G T A E K Y I K L I A N A K T V E G V W T Y K D E I K K F T V T E
BINARY_4759 T T Y K L I L N G K Q A K E E A I K E A V D A G T A E K Y I K L I A N A K T V E G V W T Y K D E I K T A T V T E
BINARY_4760 T T Y K L I L N G K Q A K E E A I K E A V D A G T A E K Y I K L I A N A K T V E G V W T Y K D E I K T F T V T E
BINARY_4761 T T Y K L I L N G K Q A K E E A I K E A V D A G T A E K Y I K L I A N A K T V E G V W T Y K D E T L K A T V T E
BINARY_4762 T T Y K L I L N G K Q A K E E A I K E A V D A G T A E K Y I K L I A N A K T V E G V W T Y K D E T L K F T V T E
BINARY_4763 T T Y K L I L N G K Q A K E E A I K E A V D A G T A E K Y I K L I A N A K T V E G V W T Y K D E T L T A T V T E
BINARY_4764 T T Y K L I L N G K Q A K E E A I K E A V D A G T A E K Y I K L I A N A K T V E G V W T Y K D E T L T F T V T E
BINARY_4765 T T Y K L I L N G K Q A K E E A I K E A V D A G T A E K Y I K L I A N A K T V E G V W T Y K D E T K K A T V T E
BINARY_4766 T T Y K L I L N G K Q A K E E A I K E A V D A G T A E K Y I K L I A N A K T V E G V W T Y K D E T K K F T V T E

BINARY_4767 T T Y K L I L N G K Q A K E E A I K E A V D A G T A E K Y I K L I A N A K T V E G V W T Y K D E T K T A T V T E
BINARY_4768 T T Y K L I L N G K Q A K E E A I K E A V D A G T A E K Y I K L I A N A K T V E G V W T Y K D E T K T F T V T E
BINARY_4769 T T Y K L I L N G K Q A K E E A I K E A V D A G T A E K Y I K L Y A N A K T V E G V W T L K D E I L K A T V T E
BINARY_4770 T T Y K L I L N G K Q A K E E A I K E A V D A G T A E K Y I K L Y A N A K T V E G V W T L K D E I L K F T V T E
BINARY_4771 T T Y K L I L N G K Q A K E E A I K E A V D A G T A E K Y I K L Y A N A K T V E G V W T L K D E I L T A T V T E
BINARY_4772 T T Y K L I L N G K Q A K E E A I K E A V D A G T A E K Y I K L Y A N A K T V E G V W T L K D E I L T F T V T E
BINARY_4773 T T Y K L I L N G K Q A K E E A I K E A V D A G T A E K Y I K L Y A N A K T V E G V W T L K D E I K K A T V T E
BINARY_4774 T T Y K L I L N G K Q A K E E A I K E A V D A G T A E K Y I K L Y A N A K T V E G V W T L K D E I K K F T V T E
BINARY_4775 T T Y K L I L N G K Q A K E E A I K E A V D A G T A E K Y I K L Y A N A K T V E G V W T L K D E I K T A T V T E
BINARY_4776 T T Y K L I L N G K Q A K E E A I K E A V D A G T A E K Y I K L Y A N A K T V E G V W T L K D E I K T F T V T E
BINARY_4777 T T Y K L I L N G K Q A K E E A I K E A V D A G T A E K Y I K L Y A N A K T V E G V W T L K D E T L K A T V T E
BINARY_4778 T T Y K L I L N G K Q A K E E A I K E A V D A G T A E K Y I K L Y A N A K T V E G V W T L K D E T L K F T V T E
BINARY_4779 T T Y K L I L N G K Q A K E E A I K E A V D A G T A E K Y I K L Y A N A K T V E G V W T L K D E T L T A T V T E
BINARY_4780 T T Y K L I L N G K Q A K E E A I K E A V D A G T A E K Y I K L Y A N A K T V E G V W T L K D E T L T F T V T E
BINARY_4781 T T Y K L I L N G K Q A K E E A I K E A V D A G T A E K Y I K L Y A N A K T V E G V W T L K D E T K K A T V T E
BINARY_4782 T T Y K L I L N G K Q A K E E A I K E A V D A G T A E K Y I K L Y A N A K T V E G V W T L K D E T K K F T V T E
BINARY_4783 T T Y K L I L N G K Q A K E E A I K E A V D A G T A E K Y I K L Y A N A K T V E G V W T L K D E T K T A T V T E
BINARY_4784 T T Y K L I L N G K Q A K E E A I K E A V D A G T A E K Y I K L Y A N A K T V E G V W T L K D E T K T F T V T E
BINARY_4785 T T Y K L I L N G K Q A K E E A I K E A V D A G T A E K Y I K L Y A N A K T V E G V W T Y K D E I L K A T V T E
BINARY_4786 T T Y K L I L N G K Q A K E E A I K E A V D A G T A E K Y I K L Y A N A K T V E G V W T Y K D E I L K F T V T E
BINARY_4787 T T Y K L I L N G K Q A K E E A I K E A V D A G T A E K Y I K L Y A N A K T V E G V W T Y K D E I L T A T V T E
BINARY_4788 T T Y K L I L N G K Q A K E E A I K E A V D A G T A E K Y I K L Y A N A K T V E G V W T Y K D E I L T F T V T E
BINARY_4789 T T Y K L I L N G K Q A K E E A I K E A V D A G T A E K Y I K L Y A N A K T V E G V W T Y K D E I K K A T V T E
BINARY_4790 T T Y K L I L N G K Q A K E E A I K E A V D A G T A E K Y I K L Y A N A K T V E G V W T Y K D E I K K F T V T E
BINARY_4791 T T Y K L I L N G K Q A K E E A I K E A V D A G T A E K Y I K L Y A N A K T V E G V W T Y K D E I K T A T V T E
BINARY_4792 T T Y K L I L N G K Q A K E E A I K E A V D A G T A E K Y I K L Y A N A K T V E G V W T Y K D E I K T F T V T E
BINARY_4793 T T Y K L I L N G K Q A K E E A I K E A V D A G T A E K Y I K L Y A N A K T V E G V W T Y K D E T L K A T V T E
BINARY_4794 T T Y K L I L N G K Q A K E E A I K E A V D A G T A E K Y I K L Y A N A K T V E G V W T Y K D E T L K F T V T E
BINARY_4795 T T Y K L I L N G K Q A K E E A I K E A V D A G T A E K Y I K L Y A N A K T V E G V W T Y K D E T L T A T V T E
BINARY_4796 T T Y K L I L N G K Q A K E E A I K E A V D A G T A E K Y I K L Y A N A K T V E G V W T Y K D E T L T F T V T E
BINARY_4797 T T Y K L I L N G K Q A K E E A I K E A V D A G T A E K Y I K L Y A N A K T V E G V W T Y K D E T K K A T V T E
BINARY_4798 T T Y K L I L N G K Q A K E E A I K E A V D A G T A E K Y I K L Y A N A K T V E G V W T Y K D E T K K F T V T E
BINARY_4799 T T Y K L I L N G K Q A K E E A I K E A V D A G T A E K Y I K L Y A N A K T V E G V W T Y K D E T K T A T V T E
BINARY_4800 T T Y K L I L N G K Q A K E E A I K E A V D A G T A E K Y I K L Y A N A K T V E G V W T Y K D E T K T F T V T E
BINARY_4801 T T Y K L I L N G K Q A K E E A I K E A V D A G T A E K Y F K L I A N A K T V E G V W T L K D E I L K A T V T E
BINARY_4802 T T Y K L I L N G K Q A K E E A I K E A V D A G T A E K Y F K L I A N A K T V E G V W T L K D E I L K F T V T E
BINARY_4803 T T Y K L I L N G K Q A K E E A I K E A V D A G T A E K Y F K L I A N A K T V E G V W T L K D E I L T A T V T E
BINARY_4804 T T Y K L I L N G K Q A K E E A I K E A V D A G T A E K Y F K L I A N A K T V E G V W T L K D E I L T F T V T E
BINARY_4805 T T Y K L I L N G K Q A K E E A I K E A V D A G T A E K Y F K L I A N A K T V E G V W T L K D E I K K A T V T E
BINARY_4806 T T Y K L I L N G K Q A K E E A I K E A V D A G T A E K Y F K L I A N A K T V E G V W T L K D E I K K F T V T E
BINARY_4807 T T Y K L I L N G K Q A K E E A I K E A V D A G T A E K Y F K L I A N A K T V E G V W T L K D E I K T A T V T E
BINARY_4808 T T Y K L I L N G K Q A K E E A I K E A V D A G T A E K Y F K L I A N A K T V E G V W T L K D E I K T F T V T E
BINARY_4809 T T Y K L I L N G K Q A K E E A I K E A V D A G T A E K Y F K L I A N A K T V E G V W T L K D E T L K A T V T E
BINARY_4810 T T Y K L I L N G K Q A K E E A I K E A V D A G T A E K Y F K L I A N A K T V E G V W T L K D E T L K F T V T E
BINARY_4811 T T Y K L I L N G K Q A K E E A I K E A V D A G T A E K Y F K L I A N A K T V E G V W T L K D E T L T A T V T E
BINARY_4812 T T Y K L I L N G K Q A K E E A I K E A V D A G T A E K Y F K L I A N A K T V E G V W T L K D E T L T F T V T E
BINARY_4813 T T Y K L I L N G K Q A K E E A I K E A V D A G T A E K Y F K L I A N A K T V E G V W T L K D E T K K A T V T E
BINARY_4814 T T Y K L I L N G K Q A K E E A I K E A V D A G T A E K Y F K L I A N A K T V E G V W T L K D E T K K F T V T E
BINARY_4815 T T Y K L I L N G K Q A K E E A I K E A V D A G T A E K Y F K L I A N A K T V E G V W T L K D E T K T A T V T E
BINARY_4816 T T Y K L I L N G K Q A K E E A I K E A V D A G T A E K Y F K L I A N A K T V E G V W T L K D E T K T F T V T E
BINARY_4817 T T Y K L I L N G K Q A K E E A I K E A V D A G T A E K Y F K L I A N A K T V E G V W T Y K D E I L K A T V T E
BINARY_4818 T T Y K L I L N G K Q A K E E A I K E A V D A G T A E K Y F K L I A N A K T V E G V W T Y K D E I L K F T V T E
BINARY_4819 T T Y K L I L N G K Q A K E E A I K E A V D A G T A E K Y F K L I A N A K T V E G V W T Y K D E I L T A T V T E
BINARY_4820 T T Y K L I L N G K Q A K E E A I K E A V D A G T A E K Y F K L I A N A K T V E G V W T Y K D E I L T F T V T E
BINARY_4821 T T Y K L I L N G K Q A K E E A I K E A V D A G T A E K Y F K L I A N A K T V E G V W T Y K D E I K K A T V T E
BINARY_4822 T T Y K L I L N G K Q A K E E A I K E A V D A G T A E K Y F K L I A N A K T V E G V W T Y K D E I K K F T V T E
BINARY_4823 T T Y K L I L N G K Q A K E E A I K E A V D A G T A E K Y F K L I A N A K T V E G V W T Y K D E I K T A T V T E
BINARY_4824 T T Y K L I L N G K Q A K E E A I K E A V D A G T A E K Y F K L I A N A K T V E G V W T Y K D E I K T F T V T E
BINARY_4825 T T Y K L I L N G K Q A K E E A I K E A V D A G T A E K Y F K L I A N A K T V E G V W T Y K D E T L K A T V T E
BINARY_4826 T T Y K L I L N G K Q A K E E A I K E A V D A G T A E K Y F K L I A N A K T V E G V W T Y K D E T L K F T V T E
BINARY_4827 T T Y K L I L N G K Q A K E E A I K E A V D A G T A E K Y F K L I A N A K T V E G V W T Y K D E T L T A T V T E
BINARY_4828 T T Y K L I L N G K Q A K E E A I K E A V D A G T A E K Y F K L I A N A K T V E G V W T Y K D E T L T F T V T E
BINARY_4829 T T Y K L I L N G K Q A K E E A I K E A V D A G T A E K Y F K L I A N A K T V E G V W T Y K D E T K K A T V T E
BINARY_4830 T T Y K L I L N G K Q A K E E A I K E A V D A G T A E K Y F K L I A N A K T V E G V W T Y K D E T K K F T V T E
BINARY_4831 T T Y K L I L N G K Q A K E E A I K E A V D A G T A E K Y F K L I A N A K T V E G V W T Y K D E T K T A T V T E
BINARY_4832 T T Y K L I L N G K Q A K E E A I K E A V D A G T A E K Y F K L I A N A K T V E G V W T Y K D E T K T F T V T E
BINARY_4833 T T Y K L I L N G K Q A K E E A I K E A V D A G T A E K Y F K L Y A N A K T V E G V W T L K D E I L K A T V T E
BINARY_4834 T T Y K L I L N G K Q A K E E A I K E A V D A G T A E K Y F K L Y A N A K T V E G V W T L K D E I L K F T V T E
BINARY_4835 T T Y K L I L N G K Q A K E E A I K E A V D A G T A E K Y F K L Y A N A K T V E G V W T L K D E I L T A T V T E
BINARY_4836 T T Y K L I L N G K Q A K E E A I K E A V D A G T A E K Y F K L Y A N A K T V E G V W T L K D E I L T F T V T E
BINARY_4837 T T Y K L I L N G K Q A K E E A I K E A V D A G T A E K Y F K L Y A N A K T V E G V W T L K D E I K K A T V T E
BINARY_4838 T T Y K L I L N G K Q A K E E A I K E A V D A G T A E K Y F K L Y A N A K T V E G V W T L K D E I K K F T V T E
BINARY_4839 T T Y K L I L N G K Q A K E E A I K E A V D A G T A E K Y F K L Y A N A K T V E G V W T L K D E I K T A T V T E
BINARY_4840 T T Y K L I L N G K Q A K E E A I K E A V D A G T A E K Y F K L Y A N A K T V E G V W T L K D E I K T F T V T E
BINARY_4841 T T Y K L I L N G K Q A K E E A I K E A V D A G T A E K Y F K L Y A N A K T V E G V W T L K D E T L K A T V T E
BINARY_4842 T T Y K L I L N G K Q A K E E A I K E A V D A G T A E K Y F K L Y A N A K T V E G V W T L K D E T L K F T V T E
BINARY_4843 T T Y K L I L N G K Q A K E E A I K E A V D A G T A E K Y F K L Y A N A K T V E G V W T L K D E T L T A T V T E
BINARY_4844 T T Y K L I L N G K Q A K E E A I K E A V D A G T A E K Y F K L Y A N A K T V E G V W T L K D E T L T F T V T E
BINARY_4845 T T Y K L I L N G K Q A K E E A I K E A V D A G T A E K Y F K L Y A N A K T V E G V W T L K D E T K K A T V T E
BINARY_4846 T T Y K L I L N G K Q A K E E A I K E A V D A G T A E K Y F K L Y A N A K T V E G V W T L K D E T K K F T V T E
BINARY_4847 T T Y K L I L N G K Q A K E E A I K E A V D A G T A E K Y F K L Y A N A K T V E G V W T L K D E T K T A T V T E
BINARY_4848 T T Y K L I L N G K Q A K E E A I K E A V D A G T A E K Y F K L Y A N A K T V E G V W T L K D E T K T F T V T E
BINARY_4849 T T Y K L I L N G K Q A K E E A I K E A V D A G T A E K Y F K L Y A N A K T V E G V W T Y K D E I L K A T V T E
BINARY_4850 T T Y K L I L N G K Q A K E E A I K E A V D A G T A E K Y F K L Y A N A K T V E G V W T Y K D E I L K F T V T E
BINARY_4851 T T Y K L I L N G K Q A K E E A I K E A V D A G T A E K Y F K L Y A N A K T V E G V W T Y K D E I L T A T V T E
BINARY_4852 T T Y K L I L N G K Q A K E E A I K E A V D A G T A E K Y F K L Y A N A K T V E G V W T Y K D E I L T F T V T E
BINARY_4853 T T Y K L I L N G K Q A K E E A I K E A V D A G T A E K Y F K L Y A N A K T V E G V W T Y K D E I K K A T V T E
BINARY_4854 T T Y K L I L N G K Q A K E E A I K E A V D A G T A E K Y F K L Y A N A K T V E G V W T Y K D E I K K F T V T E
BINARY_4855 T T Y K L I L N G K Q A K E E A I K E A V D A G T A E K Y F K L Y A N A K T V E G V W T Y K D E I K T A T V T E
BINARY_4856 T T Y K L I L N G K Q A K E E A I K E A V D A G T A E K Y F K L Y A N A K T V E G V W T Y K D E I K T F T V T E
BINARY_4857 T T Y K L I L N G K Q A K E E A I K E A V D A G T A E K Y F K L Y A N A K T V E G V W T Y K D E T L K A T V T E
BINARY_4858 T T Y K L I L N G K Q A K E E A I K E A V D A G T A E K Y F K L Y A N A K T V E G V W T Y K D E T L K F T V T E
BINARY_4859 T T Y K L I L N G K Q A K E E A I K E A V D A G T A E K Y F K L Y A N A K T V E G V W T Y K D E T L T A T V T E
BINARY_4860 T T Y K L I L N G K Q A K E E A I K E A V D A G T A E K Y F K L Y A N A K T V E G V W T Y K D E T L T F T V T E
BINARY_4861 T T Y K L I L N G K Q A K E E A I K E A V D A G T A E K Y F K L Y A N A K T V E G V W T Y K D E T K K A T V T E

BINARY_4862 T T Y K L I L N G K Q A K E E A I K E A V D A G T A E K Y F K L Y A N A K T V E G V W T Y K D E T K K F T V T E
BINARY_4863 T T Y K L I L N G K Q A K E E A I K E A V D A G T A E K Y F K L Y A N A K T V E G V W T Y K D E T K T A T V T E
BINARY_4864 T T Y K L I L N G K Q A K E E A I K E A V D A G T A E K Y F K L Y A N A K T V E G V W T Y K D E T K T F T V T E
BINARY_4865 T T Y K L I L N G K Q A K E E A I K E A V D A A I A E K Y I K L I A N A K T V E G V W T L K D E I L K A T V T E
BINARY_4866 T T Y K L I L N G K Q A K E E A I K E A V D A A I A E K Y I K L I A N A K T V E G V W T L K D E I L K F T V T E
BINARY_4867 T T Y K L I L N G K Q A K E E A I K E A V D A A I A E K Y I K L I A N A K T V E G V W T L K D E I L T A T V T E
BINARY_4868 T T Y K L I L N G K Q A K E E A I K E A V D A A I A E K Y I K L I A N A K T V E G V W T L K D E I L T F T V T E
BINARY_4869 T T Y K L I L N G K Q A K E E A I K E A V D A A I A E K Y I K L I A N A K T V E G V W T L K D E I K K A T V T E
BINARY_4870 T T Y K L I L N G K Q A K E E A I K E A V D A A I A E K Y I K L I A N A K T V E G V W T L K D E I K K F T V T E
BINARY_4871 T T Y K L I L N G K Q A K E E A I K E A V D A A I A E K Y I K L I A N A K T V E G V W T L K D E I K T A T V T E
BINARY_4872 T T Y K L I L N G K Q A K E E A I K E A V D A A I A E K Y I K L I A N A K T V E G V W T L K D E I K T F T V T E
BINARY_4873 T T Y K L I L N G K Q A K E E A I K E A V D A A I A E K Y I K L I A N A K T V E G V W T L K D E T L K A T V T E
BINARY_4874 T T Y K L I L N G K Q A K E E A I K E A V D A A I A E K Y I K L I A N A K T V E G V W T L K D E T L K F T V T E
BINARY_4875 T T Y K L I L N G K Q A K E E A I K E A V D A A I A E K Y I K L I A N A K T V E G V W T L K D E T L T A T V T E
BINARY_4876 T T Y K L I L N G K Q A K E E A I K E A V D A A I A E K Y I K L I A N A K T V E G V W T L K D E T L T F T V T E
BINARY_4877 T T Y K L I L N G K Q A K E E A I K E A V D A A I A E K Y I K L I A N A K T V E G V W T L K D E T K K A T V T E
BINARY_4878 T T Y K L I L N G K Q A K E E A I K E A V D A A I A E K Y I K L I A N A K T V E G V W T L K D E T K K F T V T E
BINARY_4879 T T Y K L I L N G K Q A K E E A I K E A V D A A I A E K Y I K L I A N A K T V E G V W T L K D E T K T A T V T E
BINARY_4880 T T Y K L I L N G K Q A K E E A I K E A V D A A I A E K Y I K L I A N A K T V E G V W T L K D E T K T F T V T E
BINARY_4881 T T Y K L I L N G K Q A K E E A I K E A V D A A I A E K Y I K L I A N A K T V E G V W T Y K D E I L K A T V T E
BINARY_4882 T T Y K L I L N G K Q A K E E A I K E A V D A A I A E K Y I K L I A N A K T V E G V W T Y K D E I L K F T V T E
BINARY_4883 T T Y K L I L N G K Q A K E E A I K E A V D A A I A E K Y I K L I A N A K T V E G V W T Y K D E I L T A T V T E
BINARY_4884 T T Y K L I L N G K Q A K E E A I K E A V D A A I A E K Y I K L I A N A K T V E G V W T Y K D E I L T F T V T E
BINARY_4885 T T Y K L I L N G K Q A K E E A I K E A V D A A I A E K Y I K L I A N A K T V E G V W T Y K D E I K K A T V T E
BINARY_4886 T T Y K L I L N G K Q A K E E A I K E A V D A A I A E K Y I K L I A N A K T V E G V W T Y K D E I K K F T V T E
BINARY_4887 T T Y K L I L N G K Q A K E E A I K E A V D A A I A E K Y I K L I A N A K T V E G V W T Y K D E I K T A T V T E
BINARY_4888 T T Y K L I L N G K Q A K E E A I K E A V D A A I A E K Y I K L I A N A K T V E G V W T Y K D E I K T F T V T E
BINARY_4889 T T Y K L I L N G K Q A K E E A I K E A V D A A I A E K Y I K L I A N A K T V E G V W T Y K D E T L K A T V T E
BINARY_4890 T T Y K L I L N G K Q A K E E A I K E A V D A A I A E K Y I K L I A N A K T V E G V W T Y K D E T L K F T V T E
BINARY_4891 T T Y K L I L N G K Q A K E E A I K E A V D A A I A E K Y I K L I A N A K T V E G V W T Y K D E T L T A T V T E
BINARY_4892 T T Y K L I L N G K Q A K E E A I K E A V D A A I A E K Y I K L I A N A K T V E G V W T Y K D E T L T F T V T E
BINARY_4893 T T Y K L I L N G K Q A K E E A I K E A V D A A I A E K Y I K L I A N A K T V E G V W T Y K D E T K K A T V T E
BINARY_4894 T T Y K L I L N G K Q A K E E A I K E A V D A A I A E K Y I K L I A N A K T V E G V W T Y K D E T K K F T V T E
BINARY_4895 T T Y K L I L N G K Q A K E E A I K E A V D A A I A E K Y I K L I A N A K T V E G V W T Y K D E T K T A T V T E
BINARY_4896 T T Y K L I L N G K Q A K E E A I K E A V D A A I A E K Y I K L I A N A K T V E G V W T Y K D E T K T F T V T E
BINARY_4897 T T Y K L I L N G K Q A K E E A I K E A V D A A I A E K Y I K L Y A N A K T V E G V W T L K D E I L K A T V T E
BINARY_4898 T T Y K L I L N G K Q A K E E A I K E A V D A A I A E K Y I K L Y A N A K T V E G V W T L K D E I L K F T V T E
BINARY_4899 T T Y K L I L N G K Q A K E E A I K E A V D A A I A E K Y I K L Y A N A K T V E G V W T L K D E I L T A T V T E
BINARY_4900 T T Y K L I L N G K Q A K E E A I K E A V D A A I A E K Y I K L Y A N A K T V E G V W T L K D E I L T F T V T E
BINARY_4901 T T Y K L I L N G K Q A K E E A I K E A V D A A I A E K Y I K L Y A N A K T V E G V W T L K D E I K K A T V T E
BINARY_4902 T T Y K L I L N G K Q A K E E A I K E A V D A A I A E K Y I K L Y A N A K T V E G V W T L K D E I K K F T V T E
BINARY_4903 T T Y K L I L N G K Q A K E E A I K E A V D A A I A E K Y I K L Y A N A K T V E G V W T L K D E I K T A T V T E
BINARY_4904 T T Y K L I L N G K Q A K E E A I K E A V D A A I A E K Y I K L Y A N A K T V E G V W T L K D E I K T F T V T E
BINARY_4905 T T Y K L I L N G K Q A K E E A I K E A V D A A I A E K Y I K L Y A N A K T V E G V W T L K D E T L K A T V T E
BINARY_4906 T T Y K L I L N G K Q A K E E A I K E A V D A A I A E K Y I K L Y A N A K T V E G V W T L K D E T L K F T V T E
BINARY_4907 T T Y K L I L N G K Q A K E E A I K E A V D A A I A E K Y I K L Y A N A K T V E G V W T L K D E T L T A T V T E
BINARY_4908 T T Y K L I L N G K Q A K E E A I K E A V D A A I A E K Y I K L Y A N A K T V E G V W T L K D E T L T F T V T E
BINARY_4909 T T Y K L I L N G K Q A K E E A I K E A V D A A I A E K Y I K L Y A N A K T V E G V W T L K D E T K K A T V T E
BINARY_4910 T T Y K L I L N G K Q A K E E A I K E A V D A A I A E K Y I K L Y A N A K T V E G V W T L K D E T K K F T V T E
BINARY_4911 T T Y K L I L N G K Q A K E E A I K E A V D A A I A E K Y I K L Y A N A K T V E G V W T L K D E T K T A T V T E
BINARY_4912 T T Y K L I L N G K Q A K E E A I K E A V D A A I A E K Y I K L Y A N A K T V E G V W T L K D E T K T F T V T E
BINARY_4913 T T Y K L I L N G K Q A K E E A I K E A V D A A I A E K Y I K L Y A N A K T V E G V W T Y K D E I L K A T V T E
BINARY_4914 T T Y K L I L N G K Q A K E E A I K E A V D A A I A E K Y I K L Y A N A K T V E G V W T Y K D E I L K F T V T E
BINARY_4915 T T Y K L I L N G K Q A K E E A I K E A V D A A I A E K Y I K L Y A N A K T V E G V W T Y K D E I L T A T V T E
BINARY_4916 T T Y K L I L N G K Q A K E E A I K E A V D A A I A E K Y I K L Y A N A K T V E G V W T Y K D E I L T F T V T E
BINARY_4917 T T Y K L I L N G K Q A K E E A I K E A V D A A I A E K Y I K L Y A N A K T V E G V W T Y K D E I K K A T V T E
BINARY_4918 T T Y K L I L N G K Q A K E E A I K E A V D A A I A E K Y I K L Y A N A K T V E G V W T Y K D E I K K F T V T E
BINARY_4919 T T Y K L I L N G K Q A K E E A I K E A V D A A I A E K Y I K L Y A N A K T V E G V W T Y K D E I K T A T V T E
BINARY_4920 T T Y K L I L N G K Q A K E E A I K E A V D A A I A E K Y I K L Y A N A K T V E G V W T Y K D E I K T F T V T E
BINARY_4921 T T Y K L I L N G K Q A K E E A I K E A V D A A I A E K Y I K L Y A N A K T V E G V W T Y K D E T L K A T V T E
BINARY_4922 T T Y K L I L N G K Q A K E E A I K E A V D A A I A E K Y I K L Y A N A K T V E G V W T Y K D E T L K F T V T E
BINARY_4923 T T Y K L I L N G K Q A K E E A I K E A V D A A I A E K Y I K L Y A N A K T V E G V W T Y K D E T L T A T V T E
BINARY_4924 T T Y K L I L N G K Q A K E E A I K E A V D A A I A E K Y I K L Y A N A K T V E G V W T Y K D E T L T F T V T E
BINARY_4925 T T Y K L I L N G K Q A K E E A I K E A V D A A I A E K Y I K L Y A N A K T V E G V W T Y K D E T K K A T V T E
BINARY_4926 T T Y K L I L N G K Q A K E E A I K E A V D A A I A E K Y I K L Y A N A K T V E G V W T Y K D E T K K F T V T E
BINARY_4927 T T Y K L I L N G K Q A K E E A I K E A V D A A I A E K Y I K L Y A N A K T V E G V W T Y K D E T K T A T V T E
BINARY_4928 T T Y K L I L N G K Q A K E E A I K E A V D A A I A E K Y I K L Y A N A K T V E G V W T Y K D E T K T F T V T E
BINARY_4929 T T Y K L I L N G K Q A K E E A I K E A V D A A I A E K Y F K L I A N A K T V E G V W T L K D E I L K A T V T E
BINARY_4930 T T Y K L I L N G K Q A K E E A I K E A V D A A I A E K Y F K L I A N A K T V E G V W T L K D E I L K F T V T E
BINARY_4931 T T Y K L I L N G K Q A K E E A I K E A V D A A I A E K Y F K L I A N A K T V E G V W T L K D E I L T A T V T E
BINARY_4932 T T Y K L I L N G K Q A K E E A I K E A V D A A I A E K Y F K L I A N A K T V E G V W T L K D E I L T F T V T E
BINARY_4933 T T Y K L I L N G K Q A K E E A I K E A V D A A I A E K Y F K L I A N A K T V E G V W T L K D E I K K A T V T E
BINARY_4934 T T Y K L I L N G K Q A K E E A I K E A V D A A I A E K Y F K L I A N A K T V E G V W T L K D E I K K F T V T E
BINARY_4935 T T Y K L I L N G K Q A K E E A I K E A V D A A I A E K Y F K L I A N A K T V E G V W T L K D E I K T A T V T E
BINARY_4936 T T Y K L I L N G K Q A K E E A I K E A V D A A I A E K Y F K L I A N A K T V E G V W T L K D E I K T F T V T E
BINARY_4937 T T Y K L I L N G K Q A K E E A I K E A V D A A I A E K Y F K L I A N A K T V E G V W T L K D E T L K A T V T E
BINARY_4938 T T Y K L I L N G K Q A K E E A I K E A V D A A I A E K Y F K L I A N A K T V E G V W T L K D E T L K F T V T E
BINARY_4939 T T Y K L I L N G K Q A K E E A I K E A V D A A I A E K Y F K L I A N A K T V E G V W T L K D E T L T A T V T E
BINARY_4940 T T Y K L I L N G K Q A K E E A I K E A V D A A I A E K Y F K L I A N A K T V E G V W T L K D E T L T F T V T E
BINARY_4941 T T Y K L I L N G K Q A K E E A I K E A V D A A I A E K Y F K L I A N A K T V E G V W T L K D E T K K A T V T E
BINARY_4942 T T Y K L I L N G K Q A K E E A I K E A V D A A I A E K Y F K L I A N A K T V E G V W T L K D E T K K F T V T E
BINARY_4943 T T Y K L I L N G K Q A K E E A I K E A V D A A I A E K Y F K L I A N A K T V E G V W T L K D E T K T A T V T E
BINARY_4944 T T Y K L I L N G K Q A K E E A I K E A V D A A I A E K Y F K L I A N A K T V E G V W T L K D E T K T F T V T E
BINARY_4945 T T Y K L I L N G K Q A K E E A I K E A V D A A I A E K Y F K L I A N A K T V E G V W T Y K D E I L K A T V T E
BINARY_4946 T T Y K L I L N G K Q A K E E A I K E A V D A A I A E K Y F K L I A N A K T V E G V W T Y K D E I L K F T V T E
BINARY_4947 T T Y K L I L N G K Q A K E E A I K E A V D A A I A E K Y F K L I A N A K T V E G V W T Y K D E I L T A T V T E
BINARY_4948 T T Y K L I L N G K Q A K E E A I K E A V D A A I A E K Y F K L I A N A K T V E G V W T Y K D E I L T F T V T E
BINARY_4949 T T Y K L I L N G K Q A K E E A I K E A V D A A I A E K Y F K L I A N A K T V E G V W T Y K D E I K K A T V T E
BINARY_4950 T T Y K L I L N G K Q A K E E A I K E A V D A A I A E K Y F K L I A N A K T V E G V W T Y K D E I K K F T V T E
BINARY_4951 T T Y K L I L N G K Q A K E E A I K E A V D A A I A E K Y F K L I A N A K T V E G V W T Y K D E I K T A T V T E
BINARY_4952 T T Y K L I L N G K Q A K E E A I K E A V D A A I A E K Y F K L I A N A K T V E G V W T Y K D E I K T F T V T E
BINARY_4953 T T Y K L I L N G K Q A K E E A I K E A V D A A I A E K Y F K L I A N A K T V E G V W T Y K D E T L K A T V T E
BINARY_4954 T T Y K L I L N G K Q A K E E A I K E A V D A A I A E K Y F K L I A N A K T V E G V W T Y K D E T L K F T V T E
BINARY_4955 T T Y K L I L N G K Q A K E E A I K E A V D A A I A E K Y F K L I A N A K T V E G V W T Y K D E T L T A T V T E
BINARY_4956 T T Y K L I L N G K Q A K E E A I K E A V D A A I A E K Y F K L I A N A K T V E G V W T Y K D E T L T F T V T E

BINARY_4957 T T Y K L I L N G K Q A K E E A I K E A V D A A I A E K Y F K L I A N A K T V E G V W T Y K D E T K K A T V E
BINARY_4958 T T Y K L I L N G K Q A K E E A I K E A V D A A I A E K Y F K L I A N A K T V E G V W T Y K D E T K K F T V E
BINARY_4959 T T Y K L I L N G K Q A K E E A I K E A V D A A I A E K Y F K L I A N A K T V E G V W T Y K D E T K T A T V E
BINARY_4960 T T Y K L I L N G K Q A K E E A I K E A V D A A I A E K Y F K L I A N A K T V E G V W T Y K D E T K T F T V E
BINARY_4961 T T Y K L I L N G K Q A K E E A I K E A V D A A I A E K Y F K L Y A N A K T V E G V W T L K D E I L K A T V E
BINARY_4962 T T Y K L I L N G K Q A K E E A I K E A V D A A I A E K Y F K L Y A N A K T V E G V W T L K D E I L K F T V E
BINARY_4963 T T Y K L I L N G K Q A K E E A I K E A V D A A I A E K Y F K L Y A N A K T V E G V W T L K D E I L T A T V E
BINARY_4964 T T Y K L I L N G K Q A K E E A I K E A V D A A I A E K Y F K L Y A N A K T V E G V W T L K D E I L T F T V E
BINARY_4965 T T Y K L I L N G K Q A K E E A I K E A V D A A I A E K Y F K L Y A N A K T V E G V W T L K D E I K K A T V E
BINARY_4966 T T Y K L I L N G K Q A K E E A I K E A V D A A I A E K Y F K L Y A N A K T V E G V W T L K D E I K K F T V E
BINARY_4967 T T Y K L I L N G K Q A K E E A I K E A V D A A I A E K Y F K L Y A N A K T V E G V W T L K D E I K T A T V E
BINARY_4968 T T Y K L I L N G K Q A K E E A I K E A V D A A I A E K Y F K L Y A N A K T V E G V W T L K D E I K T F T V E
BINARY_4969 T T Y K L I L N G K Q A K E E A I K E A V D A A I A E K Y F K L Y A N A K T V E G V W T L K D E T L K A T V E
BINARY_4970 T T Y K L I L N G K Q A K E E A I K E A V D A A I A E K Y F K L Y A N A K T V E G V W T L K D E T L K F T V E
BINARY_4971 T T Y K L I L N G K Q A K E E A I K E A V D A A I A E K Y F K L Y A N A K T V E G V W T L K D E T L T A T V E
BINARY_4972 T T Y K L I L N G K Q A K E E A I K E A V D A A I A E K Y F K L Y A N A K T V E G V W T L K D E T L T F T V E
BINARY_4973 T T Y K L I L N G K Q A K E E A I K E A V D A A I A E K Y F K L Y A N A K T V E G V W T L K D E T K K A T V E
BINARY_4974 T T Y K L I L N G K Q A K E E A I K E A V D A A I A E K Y F K L Y A N A K T V E G V W T L K D E T K K F T V E
BINARY_4975 T T Y K L I L N G K Q A K E E A I K E A V D A A I A E K Y F K L Y A N A K T V E G V W T L K D E T K T A T V E
BINARY_4976 T T Y K L I L N G K Q A K E E A I K E A V D A A I A E K Y F K L Y A N A K T V E G V W T L K D E T K T F T V E
BINARY_4977 T T Y K L I L N G K Q A K E E A I K E A V D A A I A E K Y F K L Y A N A K T V E G V W T Y K D E I L K A T V E
BINARY_4978 T T Y K L I L N G K Q A K E E A I K E A V D A A I A E K Y F K L Y A N A K T V E G V W T Y K D E I L K F T V E
BINARY_4979 T T Y K L I L N G K Q A K E E A I K E A V D A A I A E K Y F K L Y A N A K T V E G V W T Y K D E I L T A T V E
BINARY_4980 T T Y K L I L N G K Q A K E E A I K E A V D A A I A E K Y F K L Y A N A K T V E G V W T Y K D E I L T F T V E
BINARY_4981 T T Y K L I L N G K Q A K E E A I K E A V D A A I A E K Y F K L Y A N A K T V E G V W T Y K D E I K K A T V E
BINARY_4982 T T Y K L I L N G K Q A K E E A I K E A V D A A I A E K Y F K L Y A N A K T V E G V W T Y K D E I K K F T V E
BINARY_4983 T T Y K L I L N G K Q A K E E A I K E A V D A A I A E K Y F K L Y A N A K T V E G V W T Y K D E I K T A T V E
BINARY_4984 T T Y K L I L N G K Q A K E E A I K E A V D A A I A E K Y F K L Y A N A K T V E G V W T Y K D E I K T F T V E
BINARY_4985 T T Y K L I L N G K Q A K E E A I K E A V D A A I A E K Y F K L Y A N A K T V E G V W T Y K D E T L K A T V E
BINARY_4986 T T Y K L I L N G K Q A K E E A I K E A V D A A I A E K Y F K L Y A N A K T V E G V W T Y K D E T L K F T V E
BINARY_4987 T T Y K L I L N G K Q A K E E A I K E A V D A A I A E K Y F K L Y A N A K T V E G V W T Y K D E T L T A T V E
BINARY_4988 T T Y K L I L N G K Q A K E E A I K E A V D A A I A E K Y F K L Y A N A K T V E G V W T Y K D E T L T F T V E
BINARY_4989 T T Y K L I L N G K Q A K E E A I K E A V D A A I A E K Y F K L Y A N A K T V E G V W T Y K D E T K K A T V E
BINARY_4990 T T Y K L I L N G K Q A K E E A I K E A V D A A I A E K Y F K L Y A N A K T V E G V W T Y K D E T K K F T V E
BINARY_4991 T T Y K L I L N G K Q A K E E A I K E A V D A A I A E K Y F K L Y A N A K T V E G V W T Y K D E T K T A T V E
BINARY_4992 T T Y K L I L N G K Q A K E E A I K E A V D A A I A E K Y F K L Y A N A K T V E G V W T Y K D E T K T F T V E
BINARY_4993 T T Y K L I L N G K Q A K E E A I K E A V D A A T A E K Y I K L I A N A K T V E G V W T L K D E I L K A T V E
BINARY_4994 T T Y K L I L N G K Q A K E E A I K E A V D A A T A E K Y I K L I A N A K T V E G V W T L K D E I L K F T V E
BINARY_4995 T T Y K L I L N G K Q A K E E A I K E A V D A A T A E K Y I K L I A N A K T V E G V W T L K D E I L T A T V E
BINARY_4996 T T Y K L I L N G K Q A K E E A I K E A V D A A T A E K Y I K L I A N A K T V E G V W T L K D E I L T F T V E
BINARY_4997 T T Y K L I L N G K Q A K E E A I K E A V D A A T A E K Y I K L I A N A K T V E G V W T L K D E I K K A T V E
BINARY_4998 T T Y K L I L N G K Q A K E E A I K E A V D A A T A E K Y I K L I A N A K T V E G V W T L K D E I K K F T V E
BINARY_4999 T T Y K L I L N G K Q A K E E A I K E A V D A A T A E K Y I K L I A N A K T V E G V W T L K D E I K T A T V E
BINARY_5000 T T Y K L I L N G K Q A K E E A I K E A V D A A T A E K Y I K L I A N A K T V E G V W T L K D E I K T F T V E
BINARY_5001 T T Y K L I L N G K Q A K E E A I K E A V D A A T A E K Y I K L I A N A K T V E G V W T L K D E T L K A T V E
BINARY_5002 T T Y K L I L N G K Q A K E E A I K E A V D A A T A E K Y I K L I A N A K T V E G V W T L K D E T L K F T V E
BINARY_5003 T T Y K L I L N G K Q A K E E A I K E A V D A A T A E K Y I K L I A N A K T V E G V W T L K D E T L T A T V E
BINARY_5004 T T Y K L I L N G K Q A K E E A I K E A V D A A T A E K Y I K L I A N A K T V E G V W T L K D E T L T F T V E
BINARY_5005 T T Y K L I L N G K Q A K E E A I K E A V D A A T A E K Y I K L I A N A K T V E G V W T L K D E T K K A T V E
BINARY_5006 T T Y K L I L N G K Q A K E E A I K E A V D A A T A E K Y I K L I A N A K T V E G V W T L K D E T K K F T V E
BINARY_5007 T T Y K L I L N G K Q A K E E A I K E A V D A A T A E K Y I K L I A N A K T V E G V W T L K D E T K T A T V E
BINARY_5008 T T Y K L I L N G K Q A K E E A I K E A V D A A T A E K Y I K L I A N A K T V E G V W T L K D E T K T F T V E
BINARY_5009 T T Y K L I L N G K Q A K E E A I K E A V D A A T A E K Y I K L I A N A K T V E G V W T Y K D E I L K A T V E
BINARY_5010 T T Y K L I L N G K Q A K E E A I K E A V D A A T A E K Y I K L I A N A K T V E G V W T Y K D E I L K F T V E
BINARY_5011 T T Y K L I L N G K Q A K E E A I K E A V D A A T A E K Y I K L I A N A K T V E G V W T Y K D E I L T A T V E
BINARY_5012 T T Y K L I L N G K Q A K E E A I K E A V D A A T A E K Y I K L I A N A K T V E G V W T Y K D E I L T F T V E
BINARY_5013 T T Y K L I L N G K Q A K E E A I K E A V D A A T A E K Y I K L I A N A K T V E G V W T Y K D E I K K A T V E
BINARY_5014 T T Y K L I L N G K Q A K E E A I K E A V D A A T A E K Y I K L I A N A K T V E G V W T Y K D E I K K F T V E
BINARY_5015 T T Y K L I L N G K Q A K E E A I K E A V D A A T A E K Y I K L I A N A K T V E G V W T Y K D E I K T A T V E
BINARY_5016 T T Y K L I L N G K Q A K E E A I K E A V D A A T A E K Y I K L I A N A K T V E G V W T Y K D E I K T F T V E
BINARY_5017 T T Y K L I L N G K Q A K E E A I K E A V D A A T A E K Y I K L I A N A K T V E G V W T Y K D E T L K A T V E
BINARY_5018 T T Y K L I L N G K Q A K E E A I K E A V D A A T A E K Y I K L I A N A K T V E G V W T Y K D E T L K F T V E
BINARY_5019 T T Y K L I L N G K Q A K E E A I K E A V D A A T A E K Y I K L I A N A K T V E G V W T Y K D E T L T A T V E
BINARY_5020 T T Y K L I L N G K Q A K E E A I K E A V D A A T A E K Y I K L I A N A K T V E G V W T Y K D E T L T F T V E
BINARY_5021 T T Y K L I L N G K Q A K E E A I K E A V D A A T A E K Y I K L I A N A K T V E G V W T Y K D E T K K A T V E
BINARY_5022 T T Y K L I L N G K Q A K E E A I K E A V D A A T A E K Y I K L I A N A K T V E G V W T Y K D E T K K F T V E
BINARY_5023 T T Y K L I L N G K Q A K E E A I K E A V D A A T A E K Y I K L I A N A K T V E G V W T Y K D E T K T A T V E
BINARY_5024 T T Y K L I L N G K Q A K E E A I K E A V D A A T A E K Y I K L I A N A K T V E G V W T Y K D E T K T F T V E
BINARY_5025 T T Y K L I L N G K Q A K E E A I K E A V D A A T A E K Y I K L Y A N A K T V E G V W T L K D E I L K A T V E
BINARY_5026 T T Y K L I L N G K Q A K E E A I K E A V D A A T A E K Y I K L Y A N A K T V E G V W T L K D E I L K F T V E
BINARY_5027 T T Y K L I L N G K Q A K E E A I K E A V D A A T A E K Y I K L Y A N A K T V E G V W T L K D E I L T A T V E
BINARY_5028 T T Y K L I L N G K Q A K E E A I K E A V D A A T A E K Y I K L Y A N A K T V E G V W T L K D E I L T F T V E
BINARY_5029 T T Y K L I L N G K Q A K E E A I K E A V D A A T A E K Y I K L Y A N A K T V E G V W T L K D E I K K A T V E
BINARY_5030 T T Y K L I L N G K Q A K E E A I K E A V D A A T A E K Y I K L Y A N A K T V E G V W T L K D E I K K F T V E
BINARY_5031 T T Y K L I L N G K Q A K E E A I K E A V D A A T A E K Y I K L Y A N A K T V E G V W T L K D E I K T A T V E
BINARY_5032 T T Y K L I L N G K Q A K E E A I K E A V D A A T A E K Y I K L Y A N A K T V E G V W T L K D E I K T F T V E
BINARY_5033 T T Y K L I L N G K Q A K E E A I K E A V D A A T A E K Y I K L Y A N A K T V E G V W T L K D E T L K A T V E
BINARY_5034 T T Y K L I L N G K Q A K E E A I K E A V D A A T A E K Y I K L Y A N A K T V E G V W T L K D E T L K F T V E
BINARY_5035 T T Y K L I L N G K Q A K E E A I K E A V D A A T A E K Y I K L Y A N A K T V E G V W T L K D E T L T A T V E
BINARY_5036 T T Y K L I L N G K Q A K E E A I K E A V D A A T A E K Y I K L Y A N A K T V E G V W T L K D E T L T F T V E
BINARY_5037 T T Y K L I L N G K Q A K E E A I K E A V D A A T A E K Y I K L Y A N A K T V E G V W T L K D E T K K A T V E
BINARY_5038 T T Y K L I L N G K Q A K E E A I K E A V D A A T A E K Y I K L Y A N A K T V E G V W T L K D E T K K F T V E
BINARY_5039 T T Y K L I L N G K Q A K E E A I K E A V D A A T A E K Y I K L Y A N A K T V E G V W T L K D E T K T A T V E
BINARY_5040 T T Y K L I L N G K Q A K E E A I K E A V D A A T A E K Y I K L Y A N A K T V E G V W T L K D E T K T F T V E
BINARY_5041 T T Y K L I L N G K Q A K E E A I K E A V D A A T A E K Y I K L Y A N A K T V E G V W T Y K D E I L K A T V E
BINARY_5042 T T Y K L I L N G K Q A K E E A I K E A V D A A T A E K Y I K L Y A N A K T V E G V W T Y K D E I L K F T V E
BINARY_5043 T T Y K L I L N G K Q A K E E A I K E A V D A A T A E K Y I K L Y A N A K T V E G V W T Y K D E I L T A T V E
BINARY_5044 T T Y K L I L N G K Q A K E E A I K E A V D A A T A E K Y I K L Y A N A K T V E G V W T Y K D E I L T F T V E
BINARY_5045 T T Y K L I L N G K Q A K E E A I K E A V D A A T A E K Y I K L Y A N A K T V E G V W T Y K D E I K K A T V E
BINARY_5046 T T Y K L I L N G K Q A K E E A I K E A V D A A T A E K Y I K L Y A N A K T V E G V W T Y K D E I K K F T V E
BINARY_5047 T T Y K L I L N G K Q A K E E A I K E A V D A A T A E K Y I K L Y A N A K T V E G V W T Y K D E I K T A T V E
BINARY_5048 T T Y K L I L N G K Q A K E E A I K E A V D A A T A E K Y I K L Y A N A K T V E G V W T Y K D E I K T F T V E
BINARY_5049 T T Y K L I L N G K Q A K E E A I K E A V D A A T A E K Y I K L Y A N A K T V E G V W T Y K D E T L K A T V E
BINARY_5050 T T Y K L I L N G K Q A K E E A I K E A V D A A T A E K Y I K L Y A N A K T V E G V W T Y K D E T L K F T V E
BINARY_5051 T T Y K L I L N G K Q A K E E A I K E A V D A A T A E K Y I K L Y A N A K T V E G V W T Y K D E T L T A T V E

BINARY_5052 T T Y K L I L N G K Q A K E E A I K E A V D A A T A E K Y I K L Y A N A K T V E G V W T Y K D E T L T F T V T E
BINARY_5053 T T Y K L I L N G K Q A K E E A I K E A V D A A T A E K Y I K L Y A N A K T V E G V W T Y K D E T K K A T V T E
BINARY_5054 T T Y K L I L N G K Q A K E E A I K E A V D A A T A E K Y I K L Y A N A K T V E G V W T Y K D E T K K F T V T E
BINARY_5055 T T Y K L I L N G K Q A K E E A I K E A V D A A T A E K Y I K L Y A N A K T V E G V W T Y K D E T K T A T V T E
BINARY_5056 T T Y K L I L N G K Q A K E E A I K E A V D A A T A E K Y I K L Y A N A K T V E G V W T Y K D E T K T F T V T E
BINARY_5057 T T Y K L I L N G K Q A K E E A I K E A V D A A T A E K Y F K L I A N A K T V E G V W T L K D E I L K A T V T E
BINARY_5058 T T Y K L I L N G K Q A K E E A I K E A V D A A T A E K Y F K L I A N A K T V E G V W T L K D E I L K F T V T E
BINARY_5059 T T Y K L I L N G K Q A K E E A I K E A V D A A T A E K Y F K L I A N A K T V E G V W T L K D E I L T A T V T E
BINARY_5060 T T Y K L I L N G K Q A K E E A I K E A V D A A T A E K Y F K L I A N A K T V E G V W T L K D E I L T F T V T E
BINARY_5061 T T Y K L I L N G K Q A K E E A I K E A V D A A T A E K Y F K L I A N A K T V E G V W T L K D E I K K A T V T E
BINARY_5062 T T Y K L I L N G K Q A K E E A I K E A V D A A T A E K Y F K L I A N A K T V E G V W T L K D E I K K F T V T E
BINARY_5063 T T Y K L I L N G K Q A K E E A I K E A V D A A T A E K Y F K L I A N A K T V E G V W T L K D E I K T A T V T E
BINARY_5064 T T Y K L I L N G K Q A K E E A I K E A V D A A T A E K Y F K L I A N A K T V E G V W T L K D E I K T F T V T E
BINARY_5065 T T Y K L I L N G K Q A K E E A I K E A V D A A T A E K Y F K L I A N A K T V E G V W T L K D E T L K A T V T E
BINARY_5066 T T Y K L I L N G K Q A K E E A I K E A V D A A T A E K Y F K L I A N A K T V E G V W T L K D E T L K F T V T E
BINARY_5067 T T Y K L I L N G K Q A K E E A I K E A V D A A T A E K Y F K L I A N A K T V E G V W T L K D E T L T A T V T E
BINARY_5068 T T Y K L I L N G K Q A K E E A I K E A V D A A T A E K Y F K L I A N A K T V E G V W T L K D E T L T F T V T E
BINARY_5069 T T Y K L I L N G K Q A K E E A I K E A V D A A T A E K Y F K L I A N A K T V E G V W T L K D E T K K A T V T E
BINARY_5070 T T Y K L I L N G K Q A K E E A I K E A V D A A T A E K Y F K L I A N A K T V E G V W T L K D E T K K F T V T E
BINARY_5071 T T Y K L I L N G K Q A K E E A I K E A V D A A T A E K Y F K L I A N A K T V E G V W T L K D E T K T A T V T E
BINARY_5072 T T Y K L I L N G K Q A K E E A I K E A V D A A T A E K Y F K L I A N A K T V E G V W T L K D E T K T F T V T E
BINARY_5073 T T Y K L I L N G K Q A K E E A I K E A V D A A T A E K Y F K L I A N A K T V E G V W T Y K D E I L K A T V T E
BINARY_5074 T T Y K L I L N G K Q A K E E A I K E A V D A A T A E K Y F K L I A N A K T V E G V W T Y K D E I L K F T V T E
BINARY_5075 T T Y K L I L N G K Q A K E E A I K E A V D A A T A E K Y F K L I A N A K T V E G V W T Y K D E I L T A T V T E
BINARY_5076 T T Y K L I L N G K Q A K E E A I K E A V D A A T A E K Y F K L I A N A K T V E G V W T Y K D E I L T F T V T E
BINARY_5077 T T Y K L I L N G K Q A K E E A I K E A V D A A T A E K Y F K L I A N A K T V E G V W T Y K D E I K K A T V T E
BINARY_5078 T T Y K L I L N G K Q A K E E A I K E A V D A A T A E K Y F K L I A N A K T V E G V W T Y K D E I K K F T V T E
BINARY_5079 T T Y K L I L N G K Q A K E E A I K E A V D A A T A E K Y F K L I A N A K T V E G V W T Y K D E I K T A T V T E
BINARY_5080 T T Y K L I L N G K Q A K E E A I K E A V D A A T A E K Y F K L I A N A K T V E G V W T Y K D E I K T F T V T E
BINARY_5081 T T Y K L I L N G K Q A K E E A I K E A V D A A T A E K Y F K L I A N A K T V E G V W T Y K D E T L K A T V T E
BINARY_5082 T T Y K L I L N G K Q A K E E A I K E A V D A A T A E K Y F K L I A N A K T V E G V W T Y K D E T L K F T V T E
BINARY_5083 T T Y K L I L N G K Q A K E E A I K E A V D A A T A E K Y F K L I A N A K T V E G V W T Y K D E T L T A T V T E
BINARY_5084 T T Y K L I L N G K Q A K E E A I K E A V D A A T A E K Y F K L I A N A K T V E G V W T Y K D E T L T F T V T E
BINARY_5085 T T Y K L I L N G K Q A K E E A I K E A V D A A T A E K Y F K L I A N A K T V E G V W T Y K D E T K K A T V T E
BINARY_5086 T T Y K L I L N G K Q A K E E A I K E A V D A A T A E K Y F K L I A N A K T V E G V W T Y K D E T K K F T V T E
BINARY_5087 T T Y K L I L N G K Q A K E E A I K E A V D A A T A E K Y F K L I A N A K T V E G V W T Y K D E T K T A T V T E
BINARY_5088 T T Y K L I L N G K Q A K E E A I K E A V D A A T A E K Y F K L I A N A K T V E G V W T Y K D E T K T F T V T E
BINARY_5089 T T Y K L I L N G K Q A K E E A I K E A V D A A T A E K Y F K L Y A N A K T V E G V W T L K D E I L K A T V T E
BINARY_5090 T T Y K L I L N G K Q A K E E A I K E A V D A A T A E K Y F K L Y A N A K T V E G V W T L K D E I L K F T V T E
BINARY_5091 T T Y K L I L N G K Q A K E E A I K E A V D A A T A E K Y F K L Y A N A K T V E G V W T L K D E I L T A T V T E
BINARY_5092 T T Y K L I L N G K Q A K E E A I K E A V D A A T A E K Y F K L Y A N A K T V E G V W T L K D E I L T F T V T E
BINARY_5093 T T Y K L I L N G K Q A K E E A I K E A V D A A T A E K Y F K L Y A N A K T V E G V W T L K D E I K K A T V T E
BINARY_5094 T T Y K L I L N G K Q A K E E A I K E A V D A A T A E K Y F K L Y A N A K T V E G V W T L K D E I K K F T V T E
BINARY_5095 T T Y K L I L N G K Q A K E E A I K E A V D A A T A E K Y F K L Y A N A K T V E G V W T L K D E I K T A T V T E
BINARY_5096 T T Y K L I L N G K Q A K E E A I K E A V D A A T A E K Y F K L Y A N A K T V E G V W T L K D E I K T F T V T E
BINARY_5097 T T Y K L I L N G K Q A K E E A I K E A V D A A T A E K Y F K L Y A N A K T V E G V W T L K D E T L K A T V T E
BINARY_5098 T T Y K L I L N G K Q A K E E A I K E A V D A A T A E K Y F K L Y A N A K T V E G V W T L K D E T L K F T V T E
BINARY_5099 T T Y K L I L N G K Q A K E E A I K E A V D A A T A E K Y F K L Y A N A K T V E G V W T L K D E T L T A T V T E
BINARY_5100 T T Y K L I L N G K Q A K E E A I K E A V D A A T A E K Y F K L Y A N A K T V E G V W T L K D E T L T F T V T E
BINARY_5101 T T Y K L I L N G K Q A K E E A I K E A V D A A T A E K Y F K L Y A N A K T V E G V W T L K D E T K K A T V T E
BINARY_5102 T T Y K L I L N G K Q A K E E A I K E A V D A A T A E K Y F K L Y A N A K T V E G V W T L K D E T K K F T V T E
BINARY_5103 T T Y K L I L N G K Q A K E E A I K E A V D A A T A E K Y F K L Y A N A K T V E G V W T L K D E T K T A T V T E
BINARY_5104 T T Y K L I L N G K Q A K E E A I K E A V D A A T A E K Y F K L Y A N A K T V E G V W T L K D E T K T F T V T E
BINARY_5105 T T Y K L I L N G K Q A K E E A I K E A V D A A T A E K Y F K L Y A N A K T V E G V W T Y K D E I L K A T V T E
BINARY_5106 T T Y K L I L N G K Q A K E E A I K E A V D A A T A E K Y F K L Y A N A K T V E G V W T Y K D E I L K F T V T E
BINARY_5107 T T Y K L I L N G K Q A K E E A I K E A V D A A T A E K Y F K L Y A N A K T V E G V W T Y K D E I L T A T V T E
BINARY_5108 T T Y K L I L N G K Q A K E E A I K E A V D A A T A E K Y F K L Y A N A K T V E G V W T Y K D E I L T F T V T E
BINARY_5109 T T Y K L I L N G K Q A K E E A I K E A V D A A T A E K Y F K L Y A N A K T V E G V W T Y K D E I K K A T V T E
BINARY_5110 T T Y K L I L N G K Q A K E E A I K E A V D A A T A E K Y F K L Y A N A K T V E G V W T Y K D E I K K F T V T E
BINARY_5111 T T Y K L I L N G K Q A K E E A I K E A V D A A T A E K Y F K L Y A N A K T V E G V W T Y K D E I K T A T V T E
BINARY_5112 T T Y K L I L N G K Q A K E E A I K E A V D A A T A E K Y F K L Y A N A K T V E G V W T Y K D E I K T F T V T E
BINARY_5113 T T Y K L I L N G K Q A K E E A I K E A V D A A T A E K Y F K L Y A N A K T V E G V W T Y K D E T L K A T V T E
BINARY_5114 T T Y K L I L N G K Q A K E E A I K E A V D A A T A E K Y F K L Y A N A K T V E G V W T Y K D E T L K F T V T E
BINARY_5115 T T Y K L I L N G K Q A K E E A I K E A V D A A T A E K Y F K L Y A N A K T V E G V W T Y K D E T L T A T V T E
BINARY_5116 T T Y K L I L N G K Q A K E E A I K E A V D A A T A E K Y F K L Y A N A K T V E G V W T Y K D E T L T F T V T E
BINARY_5117 T T Y K L I L N G K Q A K E E A I K E A V D A A T A E K Y F K L Y A N A K T V E G V W T Y K D E T K K A T V T E
BINARY_5118 T T Y K L I L N G K Q A K E E A I K E A V D A A T A E K Y F K L Y A N A K T V E G V W T Y K D E T K K F T V T E
BINARY_5119 T T Y K L I L N G K Q A K E E A I K E A V D A A T A E K Y F K L Y A N A K T V E G V W T Y K D E T K T A T V T E
BINARY_5120 T T Y K L I L N G K Q A K E E A I K E A V D A A T A E K Y F K L Y A N A K T V E G V W T Y K D E T K T F T V T E
BINARY_5121 T T Y K L I L N G K Q A K E E A I T E L V D A G I A E K Y I K L I A N A K T V E G V W T L K D E I L K A T V T E
BINARY_5122 T T Y K L I L N G K Q A K E E A I T E L V D A G I A E K Y I K L I A N A K T V E G V W T L K D E I L K F T V T E
BINARY_5123 T T Y K L I L N G K Q A K E E A I T E L V D A G I A E K Y I K L I A N A K T V E G V W T L K D E I L T A T V T E
BINARY_5124 T T Y K L I L N G K Q A K E E A I T E L V D A G I A E K Y I K L I A N A K T V E G V W T L K D E I L T F T V T E
BINARY_5125 T T Y K L I L N G K Q A K E E A I T E L V D A G I A E K Y I K L I A N A K T V E G V W T L K D E I K K A T V T E
BINARY_5126 T T Y K L I L N G K Q A K E E A I T E L V D A G I A E K Y I K L I A N A K T V E G V W T L K D E I K K F T V T E
BINARY_5127 T T Y K L I L N G K Q A K E E A I T E L V D A G I A E K Y I K L I A N A K T V E G V W T L K D E I K T A T V T E
BINARY_5128 T T Y K L I L N G K Q A K E E A I T E L V D A G I A E K Y I K L I A N A K T V E G V W T L K D E I K T F T V T E
BINARY_5129 T T Y K L I L N G K Q A K E E A I T E L V D A G I A E K Y I K L I A N A K T V E G V W T L K D E T L K A T V T E
BINARY_5130 T T Y K L I L N G K Q A K E E A I T E L V D A G I A E K Y I K L I A N A K T V E G V W T L K D E T L K F T V T E
BINARY_5131 T T Y K L I L N G K Q A K E E A I T E L V D A G I A E K Y I K L I A N A K T V E G V W T L K D E T L T A T V T E
BINARY_5132 T T Y K L I L N G K Q A K E E A I T E L V D A G I A E K Y I K L I A N A K T V E G V W T L K D E T L T F T V T E
BINARY_5133 T T Y K L I L N G K Q A K E E A I T E L V D A G I A E K Y I K L I A N A K T V E G V W T L K D E T K K A T V T E
BINARY_5134 T T Y K L I L N G K Q A K E E A I T E L V D A G I A E K Y I K L I A N A K T V E G V W T L K D E T K K F T V T E
BINARY_5135 T T Y K L I L N G K Q A K E E A I T E L V D A G I A E K Y I K L I A N A K T V E G V W T L K D E T K T A T V T E
BINARY_5136 T T Y K L I L N G K Q A K E E A I T E L V D A G I A E K Y I K L I A N A K T V E G V W T L K D E T K T F T V T E
BINARY_5137 T T Y K L I L N G K Q A K E E A I T E L V D A G I A E K Y I K L I A N A K T V E G V W T Y K D E I L K A T V T E
BINARY_5138 T T Y K L I L N G K Q A K E E A I T E L V D A G I A E K Y I K L I A N A K T V E G V W T Y K D E I L K F T V T E
BINARY_5139 T T Y K L I L N G K Q A K E E A I T E L V D A G I A E K Y I K L I A N A K T V E G V W T Y K D E I L T A T V T E
BINARY_5140 T T Y K L I L N G K Q A K E E A I T E L V D A G I A E K Y I K L I A N A K T V E G V W T Y K D E I L T F T V T E
BINARY_5141 T T Y K L I L N G K Q A K E E A I T E L V D A G I A E K Y I K L I A N A K T V E G V W T Y K D E I K K A T V T E
BINARY_5142 T T Y K L I L N G K Q A K E E A I T E L V D A G I A E K Y I K L I A N A K T V E G V W T Y K D E I K K F T V T E
BINARY_5143 T T Y K L I L N G K Q A K E E A I T E L V D A G I A E K Y I K L I A N A K T V E G V W T Y K D E I K T A T V T E
BINARY_5144 T T Y K L I L N G K Q A K E E A I T E L V D A G I A E K Y I K L I A N A K T V E G V W T Y K D E I K T F T V T E
BINARY_5145 T T Y K L I L N G K Q A K E E A I T E L V D A G I A E K Y I K L I A N A K T V E G V W T Y K D E T L K A T V T E
BINARY_5146 T T Y K L I L N G K Q A K E E A I T E L V D A G I A E K Y I K L I A N A K T V E G V W T Y K D E T L K F T V T E

BINARY_5147 T T Y K L I L N G K Q A K E E A I T E L V D A G I A E K Y I K L I A N A K T V E G V W T Y K D E T L T A T V T E
BINARY_5148 T T Y K L I L N G K Q A K E E A I T E L V D A G I A E K Y I K L I A N A K T V E G V W T Y K D E T L T F T V T E
BINARY_5149 T T Y K L I L N G K Q A K E E A I T E L V D A G I A E K Y I K L I A N A K T V E G V W T Y K D E T K K A T V T E
BINARY_5150 T T Y K L I L N G K Q A K E E A I T E L V D A G I A E K Y I K L I A N A K T V E G V W T Y K D E T K K F T V T E
BINARY_5151 T T Y K L I L N G K Q A K E E A I T E L V D A G I A E K Y I K L I A N A K T V E G V W T Y K D E T K T A T V T E
BINARY_5152 T T Y K L I L N G K Q A K E E A I T E L V D A G I A E K Y I K L I A N A K T V E G V W T Y K D E T K T F T V T E
BINARY_5153 T T Y K L I L N G K Q A K E E A I T E L V D A G I A E K Y I K L Y A N A K T V E G V W T L K D E I L K A T V T E
BINARY_5154 T T Y K L I L N G K Q A K E E A I T E L V D A G I A E K Y I K L Y A N A K T V E G V W T L K D E I L K F T V T E
BINARY_5155 T T Y K L I L N G K Q A K E E A I T E L V D A G I A E K Y I K L Y A N A K T V E G V W T L K D E I L T A T V T E
BINARY_5156 T T Y K L I L N G K Q A K E E A I T E L V D A G I A E K Y I K L Y A N A K T V E G V W T L K D E I L T F T V T E
BINARY_5157 T T Y K L I L N G K Q A K E E A I T E L V D A G I A E K Y I K L Y A N A K T V E G V W T L K D E I K K A T V T E
BINARY_5158 T T Y K L I L N G K Q A K E E A I T E L V D A G I A E K Y I K L Y A N A K T V E G V W T L K D E I K K F T V T E
BINARY_5159 T T Y K L I L N G K Q A K E E A I T E L V D A G I A E K Y I K L Y A N A K T V E G V W T L K D E I K T A T V T E
BINARY_5160 T T Y K L I L N G K Q A K E E A I T E L V D A G I A E K Y I K L Y A N A K T V E G V W T L K D E I K T F T V T E
BINARY_5161 T T Y K L I L N G K Q A K E E A I T E L V D A G I A E K Y I K L Y A N A K T V E G V W T L K D E T L K A T V T E
BINARY_5162 T T Y K L I L N G K Q A K E E A I T E L V D A G I A E K Y I K L Y A N A K T V E G V W T L K D E T L K F T V T E
BINARY_5163 T T Y K L I L N G K Q A K E E A I T E L V D A G I A E K Y I K L Y A N A K T V E G V W T L K D E T L T A T V T E
BINARY_5164 T T Y K L I L N G K Q A K E E A I T E L V D A G I A E K Y I K L Y A N A K T V E G V W T L K D E T L T F T V T E
BINARY_5165 T T Y K L I L N G K Q A K E E A I T E L V D A G I A E K Y I K L Y A N A K T V E G V W T L K D E T K K A T V T E
BINARY_5166 T T Y K L I L N G K Q A K E E A I T E L V D A G I A E K Y I K L Y A N A K T V E G V W T L K D E T K K F T V T E
BINARY_5167 T T Y K L I L N G K Q A K E E A I T E L V D A G I A E K Y I K L Y A N A K T V E G V W T L K D E T K T A T V T E
BINARY_5168 T T Y K L I L N G K Q A K E E A I T E L V D A G I A E K Y I K L Y A N A K T V E G V W T L K D E T K T F T V T E
BINARY_5169 T T Y K L I L N G K Q A K E E A I T E L V D A G I A E K Y I K L Y A N A K T V E G V W T Y K D E I L K A T V T E
BINARY_5170 T T Y K L I L N G K Q A K E E A I T E L V D A G I A E K Y I K L Y A N A K T V E G V W T Y K D E I L K F T V T E
BINARY_5171 T T Y K L I L N G K Q A K E E A I T E L V D A G I A E K Y I K L Y A N A K T V E G V W T Y K D E I L T A T V T E
BINARY_5172 T T Y K L I L N G K Q A K E E A I T E L V D A G I A E K Y I K L Y A N A K T V E G V W T Y K D E I L T F T V T E
BINARY_5173 T T Y K L I L N G K Q A K E E A I T E L V D A G I A E K Y I K L Y A N A K T V E G V W T Y K D E I K K A T V T E
BINARY_5174 T T Y K L I L N G K Q A K E E A I T E L V D A G I A E K Y I K L Y A N A K T V E G V W T Y K D E I K K F T V T E
BINARY_5175 T T Y K L I L N G K Q A K E E A I T E L V D A G I A E K Y I K L Y A N A K T V E G V W T Y K D E I K T A T V T E
BINARY_5176 T T Y K L I L N G K Q A K E E A I T E L V D A G I A E K Y I K L Y A N A K T V E G V W T Y K D E I K T F T V T E
BINARY_5177 T T Y K L I L N G K Q A K E E A I T E L V D A G I A E K Y I K L Y A N A K T V E G V W T Y K D E T L K A T V T E
BINARY_5178 T T Y K L I L N G K Q A K E E A I T E L V D A G I A E K Y I K L Y A N A K T V E G V W T Y K D E T L K F T V T E
BINARY_5179 T T Y K L I L N G K Q A K E E A I T E L V D A G I A E K Y I K L Y A N A K T V E G V W T Y K D E T L T A T V T E
BINARY_5180 T T Y K L I L N G K Q A K E E A I T E L V D A G I A E K Y I K L Y A N A K T V E G V W T Y K D E T L T F T V T E
BINARY_5181 T T Y K L I L N G K Q A K E E A I T E L V D A G I A E K Y I K L Y A N A K T V E G V W T Y K D E T K K A T V T E
BINARY_5182 T T Y K L I L N G K Q A K E E A I T E L V D A G I A E K Y I K L Y A N A K T V E G V W T Y K D E T K K F T V T E
BINARY_5183 T T Y K L I L N G K Q A K E E A I T E L V D A G I A E K Y I K L Y A N A K T V E G V W T Y K D E T K T A T V T E
BINARY_5184 T T Y K L I L N G K Q A K E E A I T E L V D A G I A E K Y I K L Y A N A K T V E G V W T Y K D E T K T F T V T E
BINARY_5185 T T Y K L I L N G K Q A K E E A I T E L V D A G I A E K Y F K L I A N A K T V E G V W T L K D E I L K A T V T E
BINARY_5186 T T Y K L I L N G K Q A K E E A I T E L V D A G I A E K Y F K L I A N A K T V E G V W T L K D E I L K F T V T E
BINARY_5187 T T Y K L I L N G K Q A K E E A I T E L V D A G I A E K Y F K L I A N A K T V E G V W T L K D E I L T A T V T E
BINARY_5188 T T Y K L I L N G K Q A K E E A I T E L V D A G I A E K Y F K L I A N A K T V E G V W T L K D E I L T F T V T E
BINARY_5189 T T Y K L I L N G K Q A K E E A I T E L V D A G I A E K Y F K L I A N A K T V E G V W T L K D E I K K A T V T E
BINARY_5190 T T Y K L I L N G K Q A K E E A I T E L V D A G I A E K Y F K L I A N A K T V E G V W T L K D E I K K F T V T E
BINARY_5191 T T Y K L I L N G K Q A K E E A I T E L V D A G I A E K Y F K L I A N A K T V E G V W T L K D E I K T A T V T E
BINARY_5192 T T Y K L I L N G K Q A K E E A I T E L V D A G I A E K Y F K L I A N A K T V E G V W T L K D E I K T F T V T E
BINARY_5193 T T Y K L I L N G K Q A K E E A I T E L V D A G I A E K Y F K L I A N A K T V E G V W T L K D E T L K A T V T E
BINARY_5194 T T Y K L I L N G K Q A K E E A I T E L V D A G I A E K Y F K L I A N A K T V E G V W T L K D E T L K F T V T E
BINARY_5195 T T Y K L I L N G K Q A K E E A I T E L V D A G I A E K Y F K L I A N A K T V E G V W T L K D E T L T A T V T E
BINARY_5196 T T Y K L I L N G K Q A K E E A I T E L V D A G I A E K Y F K L I A N A K T V E G V W T L K D E T L T F T V T E
BINARY_5197 T T Y K L I L N G K Q A K E E A I T E L V D A G I A E K Y F K L I A N A K T V E G V W T L K D E T K K A T V T E
BINARY_5198 T T Y K L I L N G K Q A K E E A I T E L V D A G I A E K Y F K L I A N A K T V E G V W T L K D E T K K F T V T E
BINARY_5199 T T Y K L I L N G K Q A K E E A I T E L V D A G I A E K Y F K L I A N A K T V E G V W T L K D E T K T A T V T E
BINARY_5200 T T Y K L I L N G K Q A K E E A I T E L V D A G I A E K Y F K L I A N A K T V E G V W T L K D E T K T F T V T E
BINARY_5201 T T Y K L I L N G K Q A K E E A I T E L V D A G I A E K Y F K L I A N A K T V E G V W T Y K D E I L K A T V T E
BINARY_5202 T T Y K L I L N G K Q A K E E A I T E L V D A G I A E K Y F K L I A N A K T V E G V W T Y K D E I L K F T V T E
BINARY_5203 T T Y K L I L N G K Q A K E E A I T E L V D A G I A E K Y F K L I A N A K T V E G V W T Y K D E I L T A T V T E
BINARY_5204 T T Y K L I L N G K Q A K E E A I T E L V D A G I A E K Y F K L I A N A K T V E G V W T Y K D E I L T F T V T E
BINARY_5205 T T Y K L I L N G K Q A K E E A I T E L V D A G I A E K Y F K L I A N A K T V E G V W T Y K D E I K K A T V T E
BINARY_5206 T T Y K L I L N G K Q A K E E A I T E L V D A G I A E K Y F K L I A N A K T V E G V W T Y K D E I K K F T V T E
BINARY_5207 T T Y K L I L N G K Q A K E E A I T E L V D A G I A E K Y F K L I A N A K T V E G V W T Y K D E I K T A T V T E
BINARY_5208 T T Y K L I L N G K Q A K E E A I T E L V D A G I A E K Y F K L I A N A K T V E G V W T Y K D E I K T F T V T E
BINARY_5209 T T Y K L I L N G K Q A K E E A I T E L V D A G I A E K Y F K L I A N A K T V E G V W T Y K D E T L K A T V T E
BINARY_5210 T T Y K L I L N G K Q A K E E A I T E L V D A G I A E K Y F K L I A N A K T V E G V W T Y K D E T L K F T V T E
BINARY_5211 T T Y K L I L N G K Q A K E E A I T E L V D A G I A E K Y F K L I A N A K T V E G V W T Y K D E T L T A T V T E
BINARY_5212 T T Y K L I L N G K Q A K E E A I T E L V D A G I A E K Y F K L I A N A K T V E G V W T Y K D E T L T F T V T E
BINARY_5213 T T Y K L I L N G K Q A K E E A I T E L V D A G I A E K Y F K L I A N A K T V E G V W T Y K D E T K K A T V T E
BINARY_5214 T T Y K L I L N G K Q A K E E A I T E L V D A G I A E K Y F K L I A N A K T V E G V W T Y K D E T K K F T V T E
BINARY_5215 T T Y K L I L N G K Q A K E E A I T E L V D A G I A E K Y F K L I A N A K T V E G V W T Y K D E T K T A T V T E
BINARY_5216 T T Y K L I L N G K Q A K E E A I T E L V D A G I A E K Y F K L I A N A K T V E G V W T Y K D E T K T F T V T E
BINARY_5217 T T Y K L I L N G K Q A K E E A I T E L V D A G I A E K Y F K L Y A N A K T V E G V W T L K D E I L K A T V T E
BINARY_5218 T T Y K L I L N G K Q A K E E A I T E L V D A G I A E K Y F K L Y A N A K T V E G V W T L K D E I L K F T V T E
BINARY_5219 T T Y K L I L N G K Q A K E E A I T E L V D A G I A E K Y F K L Y A N A K T V E G V W T L K D E I L T A T V T E
BINARY_5220 T T Y K L I L N G K Q A K E E A I T E L V D A G I A E K Y F K L Y A N A K T V E G V W T L K D E I L T F T V T E
BINARY_5221 T T Y K L I L N G K Q A K E E A I T E L V D A G I A E K Y F K L Y A N A K T V E G V W T L K D E I K K A T V T E
BINARY_5222 T T Y K L I L N G K Q A K E E A I T E L V D A G I A E K Y F K L Y A N A K T V E G V W T L K D E I K K F T V T E
BINARY_5223 T T Y K L I L N G K Q A K E E A I T E L V D A G I A E K Y F K L Y A N A K T V E G V W T L K D E I K T A T V T E
BINARY_5224 T T Y K L I L N G K Q A K E E A I T E L V D A G I A E K Y F K L Y A N A K T V E G V W T L K D E I K T F T V T E
BINARY_5225 T T Y K L I L N G K Q A K E E A I T E L V D A G I A E K Y F K L Y A N A K T V E G V W T L K D E T L K A T V T E
BINARY_5226 T T Y K L I L N G K Q A K E E A I T E L V D A G I A E K Y F K L Y A N A K T V E G V W T L K D E T L K F T V T E
BINARY_5227 T T Y K L I L N G K Q A K E E A I T E L V D A G I A E K Y F K L Y A N A K T V E G V W T L K D E T L T A T V T E
BINARY_5228 T T Y K L I L N G K Q A K E E A I T E L V D A G I A E K Y F K L Y A N A K T V E G V W T L K D E T L T F T V T E
BINARY_5229 T T Y K L I L N G K Q A K E E A I T E L V D A G I A E K Y F K L Y A N A K T V E G V W T L K D E T K K A T V T E
BINARY_5230 T T Y K L I L N G K Q A K E E A I T E L V D A G I A E K Y F K L Y A N A K T V E G V W T L K D E T K K F T V T E
BINARY_5231 T T Y K L I L N G K Q A K E E A I T E L V D A G I A E K Y F K L Y A N A K T V E G V W T L K D E T K T A T V T E
BINARY_5232 T T Y K L I L N G K Q A K E E A I T E L V D A G I A E K Y F K L Y A N A K T V E G V W T L K D E T K T F T V T E
BINARY_5233 T T Y K L I L N G K Q A K E E A I T E L V D A G I A E K Y F K L Y A N A K T V E G V W T Y K D E I L K A T V T E
BINARY_5234 T T Y K L I L N G K Q A K E E A I T E L V D A G I A E K Y F K L Y A N A K T V E G V W T Y K D E I L K F T V T E
BINARY_5235 T T Y K L I L N G K Q A K E E A I T E L V D A G I A E K Y F K L Y A N A K T V E G V W T Y K D E I L T A T V T E
BINARY_5236 T T Y K L I L N G K Q A K E E A I T E L V D A G I A E K Y F K L Y A N A K T V E G V W T Y K D E I L T F T V T E
BINARY_5237 T T Y K L I L N G K Q A K E E A I T E L V D A G I A E K Y F K L Y A N A K T V E G V W T Y K D E I K K A T V T E
BINARY_5238 T T Y K L I L N G K Q A K E E A I T E L V D A G I A E K Y F K L Y A N A K T V E G V W T Y K D E I K K F T V T E
BINARY_5239 T T Y K L I L N G K Q A K E E A I T E L V D A G I A E K Y F K L Y A N A K T V E G V W T Y K D E I K T A T V T E
BINARY_5240 T T Y K L I L N G K Q A K E E A I T E L V D A G I A E K Y F K L Y A N A K T V E G V W T Y K D E I K T F T V T E
BINARY_5241 T T Y K L I L N G K Q A K E E A I T E L V D A G I A E K Y F K L Y A N A K T V E G V W T Y K D E T L K A T V T E

BINARY_5242 T T Y K L I L N G K Q A K E E A I T E L V D A G I A E K Y F K L Y A N A K T V E G V W T Y K D E T L K F T V T E
BINARY_5243 T T Y K L I L N G K Q A K E E A I T E L V D A G I A E K Y F K L Y A N A K T V E G V W T Y K D E T L T A T V T E
BINARY_5244 T T Y K L I L N G K Q A K E E A I T E L V D A G I A E K Y F K L Y A N A K T V E G V W T Y K D E T L T F T V T E
BINARY_5245 T T Y K L I L N G K Q A K E E A I T E L V D A G I A E K Y F K L Y A N A K T V E G V W T Y K D E T K K A T V T E
BINARY_5246 T T Y K L I L N G K Q A K E E A I T E L V D A G I A E K Y F K L Y A N A K T V E G V W T Y K D E T K K F T V T E
BINARY_5247 T T Y K L I L N G K Q A K E E A I T E L V D A G I A E K Y F K L Y A N A K T V E G V W T Y K D E T K T A T V T E
BINARY_5248 T T Y K L I L N G K Q A K E E A I T E L V D A G I A E K Y F K L Y A N A K T V E G V W T Y K D E T K T F T V T E
BINARY_5249 T T Y K L I L N G K Q A K E E A I T E L V D A G T A E K Y I K L I A N A K T V E G V W T L K D E I L K A T V T E
BINARY_5250 T T Y K L I L N G K Q A K E E A I T E L V D A G T A E K Y I K L I A N A K T V E G V W T L K D E I L K F T V T E
BINARY_5251 T T Y K L I L N G K Q A K E E A I T E L V D A G T A E K Y I K L I A N A K T V E G V W T L K D E I L T A T V T E
BINARY_5252 T T Y K L I L N G K Q A K E E A I T E L V D A G T A E K Y I K L I A N A K T V E G V W T L K D E I L T F T V T E
BINARY_5253 T T Y K L I L N G K Q A K E E A I T E L V D A G T A E K Y I K L I A N A K T V E G V W T L K D E I K K A T V T E
BINARY_5254 T T Y K L I L N G K Q A K E E A I T E L V D A G T A E K Y I K L I A N A K T V E G V W T L K D E I K K F T V T E
BINARY_5255 T T Y K L I L N G K Q A K E E A I T E L V D A G T A E K Y I K L I A N A K T V E G V W T L K D E I K T A T V T E
BINARY_5256 T T Y K L I L N G K Q A K E E A I T E L V D A G T A E K Y I K L I A N A K T V E G V W T L K D E I K T F T V T E
BINARY_5257 T T Y K L I L N G K Q A K E E A I T E L V D A G T A E K Y I K L I A N A K T V E G V W T L K D E T L K A T V T E
BINARY_5258 T T Y K L I L N G K Q A K E E A I T E L V D A G T A E K Y I K L I A N A K T V E G V W T L K D E T L K F T V T E
BINARY_5259 T T Y K L I L N G K Q A K E E A I T E L V D A G T A E K Y I K L I A N A K T V E G V W T L K D E T L T A T V T E
BINARY_5260 T T Y K L I L N G K Q A K E E A I T E L V D A G T A E K Y I K L I A N A K T V E G V W T L K D E T L T F T V T E
BINARY_5261 T T Y K L I L N G K Q A K E E A I T E L V D A G T A E K Y I K L I A N A K T V E G V W T L K D E T K K A T V T E
BINARY_5262 T T Y K L I L N G K Q A K E E A I T E L V D A G T A E K Y I K L I A N A K T V E G V W T L K D E T K K F T V T E
BINARY_5263 T T Y K L I L N G K Q A K E E A I T E L V D A G T A E K Y I K L I A N A K T V E G V W T L K D E T K T A T V T E
BINARY_5264 T T Y K L I L N G K Q A K E E A I T E L V D A G T A E K Y I K L I A N A K T V E G V W T L K D E T K T F T V T E
BINARY_5265 T T Y K L I L N G K Q A K E E A I T E L V D A G T A E K Y I K L I A N A K T V E G V W T Y K D E I L K A T V T E
BINARY_5266 T T Y K L I L N G K Q A K E E A I T E L V D A G T A E K Y I K L I A N A K T V E G V W T Y K D E I L K F T V T E
BINARY_5267 T T Y K L I L N G K Q A K E E A I T E L V D A G T A E K Y I K L I A N A K T V E G V W T Y K D E I L T A T V T E
BINARY_5268 T T Y K L I L N G K Q A K E E A I T E L V D A G T A E K Y I K L I A N A K T V E G V W T Y K D E I L T F T V T E
BINARY_5269 T T Y K L I L N G K Q A K E E A I T E L V D A G T A E K Y I K L I A N A K T V E G V W T Y K D E I K K A T V T E
BINARY_5270 T T Y K L I L N G K Q A K E E A I T E L V D A G T A E K Y I K L I A N A K T V E G V W T Y K D E I K K F T V T E
BINARY_5271 T T Y K L I L N G K Q A K E E A I T E L V D A G T A E K Y I K L I A N A K T V E G V W T Y K D E I K T A T V T E
BINARY_5272 T T Y K L I L N G K Q A K E E A I T E L V D A G T A E K Y I K L I A N A K T V E G V W T Y K D E I K T F T V T E
BINARY_5273 T T Y K L I L N G K Q A K E E A I T E L V D A G T A E K Y I K L I A N A K T V E G V W T Y K D E T L K A T V T E
BINARY_5274 T T Y K L I L N G K Q A K E E A I T E L V D A G T A E K Y I K L I A N A K T V E G V W T Y K D E T L K F T V T E
BINARY_5275 T T Y K L I L N G K Q A K E E A I T E L V D A G T A E K Y I K L I A N A K T V E G V W T Y K D E T L T A T V T E
BINARY_5276 T T Y K L I L N G K Q A K E E A I T E L V D A G T A E K Y I K L I A N A K T V E G V W T Y K D E T L T F T V T E
BINARY_5277 T T Y K L I L N G K Q A K E E A I T E L V D A G T A E K Y I K L I A N A K T V E G V W T Y K D E T K K A T V T E
BINARY_5278 T T Y K L I L N G K Q A K E E A I T E L V D A G T A E K Y I K L I A N A K T V E G V W T Y K D E T K K F T V T E
BINARY_5279 T T Y K L I L N G K Q A K E E A I T E L V D A G T A E K Y I K L I A N A K T V E G V W T Y K D E T K T A T V T E
BINARY_5280 T T Y K L I L N G K Q A K E E A I T E L V D A G T A E K Y I K L I A N A K T V E G V W T Y K D E T K T F T V T E
BINARY_5281 T T Y K L I L N G K Q A K E E A I T E L V D A G T A E K Y I K L Y A N A K T V E G V W T L K D E I L K A T V T E
BINARY_5282 T T Y K L I L N G K Q A K E E A I T E L V D A G T A E K Y I K L Y A N A K T V E G V W T L K D E I L K F T V T E
BINARY_5283 T T Y K L I L N G K Q A K E E A I T E L V D A G T A E K Y I K L Y A N A K T V E G V W T L K D E I L T A T V T E
BINARY_5284 T T Y K L I L N G K Q A K E E A I T E L V D A G T A E K Y I K L Y A N A K T V E G V W T L K D E I L T F T V T E
BINARY_5285 T T Y K L I L N G K Q A K E E A I T E L V D A G T A E K Y I K L Y A N A K T V E G V W T L K D E I K K A T V T E
BINARY_5286 T T Y K L I L N G K Q A K E E A I T E L V D A G T A E K Y I K L Y A N A K T V E G V W T L K D E I K K F T V T E
BINARY_5287 T T Y K L I L N G K Q A K E E A I T E L V D A G T A E K Y I K L Y A N A K T V E G V W T L K D E I K T A T V T E
BINARY_5288 T T Y K L I L N G K Q A K E E A I T E L V D A G T A E K Y I K L Y A N A K T V E G V W T L K D E I K T F T V T E
BINARY_5289 T T Y K L I L N G K Q A K E E A I T E L V D A G T A E K Y I K L Y A N A K T V E G V W T L K D E T L K A T V T E
BINARY_5290 T T Y K L I L N G K Q A K E E A I T E L V D A G T A E K Y I K L Y A N A K T V E G V W T L K D E T L K F T V T E
BINARY_5291 T T Y K L I L N G K Q A K E E A I T E L V D A G T A E K Y I K L Y A N A K T V E G V W T L K D E T L T A T V T E
BINARY_5292 T T Y K L I L N G K Q A K E E A I T E L V D A G T A E K Y I K L Y A N A K T V E G V W T L K D E T L T F T V T E
BINARY_5293 T T Y K L I L N G K Q A K E E A I T E L V D A G T A E K Y I K L Y A N A K T V E G V W T L K D E T K K A T V T E
BINARY_5294 T T Y K L I L N G K Q A K E E A I T E L V D A G T A E K Y I K L Y A N A K T V E G V W T L K D E T K K F T V T E
BINARY_5295 T T Y K L I L N G K Q A K E E A I T E L V D A G T A E K Y I K L Y A N A K T V E G V W T L K D E T K T A T V T E
BINARY_5296 T T Y K L I L N G K Q A K E E A I T E L V D A G T A E K Y I K L Y A N A K T V E G V W T L K D E T K T F T V T E
BINARY_5297 T T Y K L I L N G K Q A K E E A I T E L V D A G T A E K Y I K L Y A N A K T V E G V W T Y K D E I L K A T V T E
BINARY_5298 T T Y K L I L N G K Q A K E E A I T E L V D A G T A E K Y I K L Y A N A K T V E G V W T Y K D E I L K F T V T E
BINARY_5299 T T Y K L I L N G K Q A K E E A I T E L V D A G T A E K Y I K L Y A N A K T V E G V W T Y K D E I L T A T V T E
BINARY_5300 T T Y K L I L N G K Q A K E E A I T E L V D A G T A E K Y I K L Y A N A K T V E G V W T Y K D E I L T F T V T E
BINARY_5301 T T Y K L I L N G K Q A K E E A I T E L V D A G T A E K Y I K L Y A N A K T V E G V W T Y K D E I K K A T V T E
BINARY_5302 T T Y K L I L N G K Q A K E E A I T E L V D A G T A E K Y I K L Y A N A K T V E G V W T Y K D E I K K F T V T E
BINARY_5303 T T Y K L I L N G K Q A K E E A I T E L V D A G T A E K Y I K L Y A N A K T V E G V W T Y K D E I K T A T V T E
BINARY_5304 T T Y K L I L N G K Q A K E E A I T E L V D A G T A E K Y I K L Y A N A K T V E G V W T Y K D E I K T F T V T E
BINARY_5305 T T Y K L I L N G K Q A K E E A I T E L V D A G T A E K Y I K L Y A N A K T V E G V W T Y K D E T L K A T V T E
BINARY_5306 T T Y K L I L N G K Q A K E E A I T E L V D A G T A E K Y I K L Y A N A K T V E G V W T Y K D E T L K F T V T E
BINARY_5307 T T Y K L I L N G K Q A K E E A I T E L V D A G T A E K Y I K L Y A N A K T V E G V W T Y K D E T L T A T V T E
BINARY_5308 T T Y K L I L N G K Q A K E E A I T E L V D A G T A E K Y I K L Y A N A K T V E G V W T Y K D E T L T F T V T E
BINARY_5309 T T Y K L I L N G K Q A K E E A I T E L V D A G T A E K Y I K L Y A N A K T V E G V W T Y K D E T K K A T V T E
BINARY_5310 T T Y K L I L N G K Q A K E E A I T E L V D A G T A E K Y I K L Y A N A K T V E G V W T Y K D E T K K F T V T E
BINARY_5311 T T Y K L I L N G K Q A K E E A I T E L V D A G T A E K Y I K L Y A N A K T V E G V W T Y K D E T K T A T V T E
BINARY_5312 T T Y K L I L N G K Q A K E E A I T E L V D A G T A E K Y I K L Y A N A K T V E G V W T Y K D E T K T F T V T E
BINARY_5313 T T Y K L I L N G K Q A K E E A I T E L V D A G T A E K Y F K L I A N A K T V E G V W T L K D E I L K A T V T E
BINARY_5314 T T Y K L I L N G K Q A K E E A I T E L V D A G T A E K Y F K L I A N A K T V E G V W T L K D E I L K F T V T E
BINARY_5315 T T Y K L I L N G K Q A K E E A I T E L V D A G T A E K Y F K L I A N A K T V E G V W T L K D E I L T A T V T E
BINARY_5316 T T Y K L I L N G K Q A K E E A I T E L V D A G T A E K Y F K L I A N A K T V E G V W T L K D E I L T F T V T E
BINARY_5317 T T Y K L I L N G K Q A K E E A I T E L V D A G T A E K Y F K L I A N A K T V E G V W T L K D E I K K A T V T E
BINARY_5318 T T Y K L I L N G K Q A K E E A I T E L V D A G T A E K Y F K L I A N A K T V E G V W T L K D E I K K F T V T E
BINARY_5319 T T Y K L I L N G K Q A K E E A I T E L V D A G T A E K Y F K L I A N A K T V E G V W T L K D E I K T A T V T E
BINARY_5320 T T Y K L I L N G K Q A K E E A I T E L V D A G T A E K Y F K L I A N A K T V E G V W T L K D E I K T F T V T E
BINARY_5321 T T Y K L I L N G K Q A K E E A I T E L V D A G T A E K Y F K L I A N A K T V E G V W T L K D E T L K A T V T E
BINARY_5322 T T Y K L I L N G K Q A K E E A I T E L V D A G T A E K Y F K L I A N A K T V E G V W T L K D E T L K F T V T E
BINARY_5323 T T Y K L I L N G K Q A K E E A I T E L V D A G T A E K Y F K L I A N A K T V E G V W T L K D E T L T A T V T E
BINARY_5324 T T Y K L I L N G K Q A K E E A I T E L V D A G T A E K Y F K L I A N A K T V E G V W T L K D E T L T F T V T E
BINARY_5325 T T Y K L I L N G K Q A K E E A I T E L V D A G T A E K Y F K L I A N A K T V E G V W T L K D E T K K A T V T E
BINARY_5326 T T Y K L I L N G K Q A K E E A I T E L V D A G T A E K Y F K L I A N A K T V E G V W T L K D E T K K F T V T E
BINARY_5327 T T Y K L I L N G K Q A K E E A I T E L V D A G T A E K Y F K L I A N A K T V E G V W T L K D E T K T A T V T E
BINARY_5328 T T Y K L I L N G K Q A K E E A I T E L V D A G T A E K Y F K L I A N A K T V E G V W T L K D E T K T F T V T E
BINARY_5329 T T Y K L I L N G K Q A K E E A I T E L V D A G T A E K Y F K L I A N A K T V E G V W T Y K D E I L K A T V T E
BINARY_5330 T T Y K L I L N G K Q A K E E A I T E L V D A G T A E K Y F K L I A N A K T V E G V W T Y K D E I L K F T V T E
BINARY_5331 T T Y K L I L N G K Q A K E E A I T E L V D A G T A E K Y F K L I A N A K T V E G V W T Y K D E I L T A T V T E
BINARY_5332 T T Y K L I L N G K Q A K E E A I T E L V D A G T A E K Y F K L I A N A K T V E G V W T Y K D E I L T F T V T E
BINARY_5333 T T Y K L I L N G K Q A K E E A I T E L V D A G T A E K Y F K L I A N A K T V E G V W T Y K D E I K K A T V T E
BINARY_5334 T T Y K L I L N G K Q A K E E A I T E L V D A G T A E K Y F K L I A N A K T V E G V W T Y K D E I K K F T V T E
BINARY_5335 T T Y K L I L N G K Q A K E E A I T E L V D A G T A E K Y F K L I A N A K T V E G V W T Y K D E I K T A T V T E
BINARY_5336 T T Y K L I L N G K Q A K E E A I T E L V D A G T A E K Y F K L I A N A K T V E G V W T Y K D E I K T F T V T E

BINARY_5337 T T Y K L I L N G K Q A K E E A I T E L V D A G T A E K Y F K L I A N A K T V E G V W T Y K D E T L K A T V T E
BINARY_5338 T T Y K L I L N G K Q A K E E A I T E L V D A G T A E K Y F K L I A N A K T V E G V W T Y K D E T L K F T V T E
BINARY_5339 T T Y K L I L N G K Q A K E E A I T E L V D A G T A E K Y F K L I A N A K T V E G V W T Y K D E T L T A T V T E
BINARY_5340 T T Y K L I L N G K Q A K E E A I T E L V D A G T A E K Y F K L I A N A K T V E G V W T Y K D E T L T F T V T E
BINARY_5341 T T Y K L I L N G K Q A K E E A I T E L V D A G T A E K Y F K L I A N A K T V E G V W T Y K D E T K K A T V T E
BINARY_5342 T T Y K L I L N G K Q A K E E A I T E L V D A G T A E K Y F K L I A N A K T V E G V W T Y K D E T K K F T V T E
BINARY_5343 T T Y K L I L N G K Q A K E E A I T E L V D A G T A E K Y F K L I A N A K T V E G V W T Y K D E T K T A T V T E
BINARY_5344 T T Y K L I L N G K Q A K E E A I T E L V D A G T A E K Y F K L I A N A K T V E G V W T Y K D E T K T F T V T E
BINARY_5345 T T Y K L I L N G K Q A K E E A I T E L V D A G T A E K Y F K L Y A N A K T V E G V W T L K D E I L K A T V T E
BINARY_5346 T T Y K L I L N G K Q A K E E A I T E L V D A G T A E K Y F K L Y A N A K T V E G V W T L K D E I L K F T V T E
BINARY_5347 T T Y K L I L N G K Q A K E E A I T E L V D A G T A E K Y F K L Y A N A K T V E G V W T L K D E I L T A T V T E
BINARY_5348 T T Y K L I L N G K Q A K E E A I T E L V D A G T A E K Y F K L Y A N A K T V E G V W T L K D E I L T F T V T E
BINARY_5349 T T Y K L I L N G K Q A K E E A I T E L V D A G T A E K Y F K L Y A N A K T V E G V W T L K D E I K K A T V T E
BINARY_5350 T T Y K L I L N G K Q A K E E A I T E L V D A G T A E K Y F K L Y A N A K T V E G V W T L K D E I K K F T V T E
BINARY_5351 T T Y K L I L N G K Q A K E E A I T E L V D A G T A E K Y F K L Y A N A K T V E G V W T L K D E I K T A T V T E
BINARY_5352 T T Y K L I L N G K Q A K E E A I T E L V D A G T A E K Y F K L Y A N A K T V E G V W T L K D E I K T F T V T E
BINARY_5353 T T Y K L I L N G K Q A K E E A I T E L V D A G T A E K Y F K L Y A N A K T V E G V W T L K D E T L K A T V T E
BINARY_5354 T T Y K L I L N G K Q A K E E A I T E L V D A G T A E K Y F K L Y A N A K T V E G V W T L K D E T L K F T V T E
BINARY_5355 T T Y K L I L N G K Q A K E E A I T E L V D A G T A E K Y F K L Y A N A K T V E G V W T L K D E T L T A T V T E
BINARY_5356 T T Y K L I L N G K Q A K E E A I T E L V D A G T A E K Y F K L Y A N A K T V E G V W T L K D E T L T F T V T E
BINARY_5357 T T Y K L I L N G K Q A K E E A I T E L V D A G T A E K Y F K L Y A N A K T V E G V W T L K D E T K K A T V T E
BINARY_5358 T T Y K L I L N G K Q A K E E A I T E L V D A G T A E K Y F K L Y A N A K T V E G V W T L K D E T K K F T V T E
BINARY_5359 T T Y K L I L N G K Q A K E E A I T E L V D A G T A E K Y F K L Y A N A K T V E G V W T L K D E T K T A T V T E
BINARY_5360 T T Y K L I L N G K Q A K E E A I T E L V D A G T A E K Y F K L Y A N A K T V E G V W T L K D E T K T F T V T E
BINARY_5361 T T Y K L I L N G K Q A K E E A I T E L V D A G T A E K Y F K L Y A N A K T V E G V W T Y K D E I L K A T V T E
BINARY_5362 T T Y K L I L N G K Q A K E E A I T E L V D A G T A E K Y F K L Y A N A K T V E G V W T Y K D E I L K F T V T E
BINARY_5363 T T Y K L I L N G K Q A K E E A I T E L V D A G T A E K Y F K L Y A N A K T V E G V W T Y K D E I L T A T V T E
BINARY_5364 T T Y K L I L N G K Q A K E E A I T E L V D A G T A E K Y F K L Y A N A K T V E G V W T Y K D E I L T F T V T E
BINARY_5365 T T Y K L I L N G K Q A K E E A I T E L V D A G T A E K Y F K L Y A N A K T V E G V W T Y K D E I K K A T V T E
BINARY_5366 T T Y K L I L N G K Q A K E E A I T E L V D A G T A E K Y F K L Y A N A K T V E G V W T Y K D E I K K F T V T E
BINARY_5367 T T Y K L I L N G K Q A K E E A I T E L V D A G T A E K Y F K L Y A N A K T V E G V W T Y K D E I K T A T V T E
BINARY_5368 T T Y K L I L N G K Q A K E E A I T E L V D A G T A E K Y F K L Y A N A K T V E G V W T Y K D E I K T F T V T E
BINARY_5369 T T Y K L I L N G K Q A K E E A I T E L V D A G T A E K Y F K L Y A N A K T V E G V W T Y K D E T L K A T V T E
BINARY_5370 T T Y K L I L N G K Q A K E E A I T E L V D A G T A E K Y F K L Y A N A K T V E G V W T Y K D E T L K F T V T E
BINARY_5371 T T Y K L I L N G K Q A K E E A I T E L V D A G T A E K Y F K L Y A N A K T V E G V W T Y K D E T L T A T V T E
BINARY_5372 T T Y K L I L N G K Q A K E E A I T E L V D A G T A E K Y F K L Y A N A K T V E G V W T Y K D E T L T F T V T E
BINARY_5373 T T Y K L I L N G K Q A K E E A I T E L V D A G T A E K Y F K L Y A N A K T V E G V W T Y K D E T K K A T V T E
BINARY_5374 T T Y K L I L N G K Q A K E E A I T E L V D A G T A E K Y F K L Y A N A K T V E G V W T Y K D E T K K F T V T E
BINARY_5375 T T Y K L I L N G K Q A K E E A I T E L V D A G T A E K Y F K L Y A N A K T V E G V W T Y K D E T K T A T V T E
BINARY_5376 T T Y K L I L N G K Q A K E E A I T E L V D A G T A E K Y F K L Y A N A K T V E G V W T Y K D E T K T F T V T E
BINARY_5377 T T Y K L I L N G K Q A K E E A I T E L V D A A I A E K Y I K L I A N A K T V E G V W T L K D E I L K A T V T E
BINARY_5378 T T Y K L I L N G K Q A K E E A I T E L V D A A I A E K Y I K L I A N A K T V E G V W T L K D E I L K F T V T E
BINARY_5379 T T Y K L I L N G K Q A K E E A I T E L V D A A I A E K Y I K L I A N A K T V E G V W T L K D E I L T A T V T E
BINARY_5380 T T Y K L I L N G K Q A K E E A I T E L V D A A I A E K Y I K L I A N A K T V E G V W T L K D E I L T F T V T E
BINARY_5381 T T Y K L I L N G K Q A K E E A I T E L V D A A I A E K Y I K L I A N A K T V E G V W T L K D E I K K A T V T E
BINARY_5382 T T Y K L I L N G K Q A K E E A I T E L V D A A I A E K Y I K L I A N A K T V E G V W T L K D E I K K F T V T E
BINARY_5383 T T Y K L I L N G K Q A K E E A I T E L V D A A I A E K Y I K L I A N A K T V E G V W T L K D E I K T A T V T E
BINARY_5384 T T Y K L I L N G K Q A K E E A I T E L V D A A I A E K Y I K L I A N A K T V E G V W T L K D E I K T F T V T E
BINARY_5385 T T Y K L I L N G K Q A K E E A I T E L V D A A I A E K Y I K L I A N A K T V E G V W T L K D E T L K A T V T E
BINARY_5386 T T Y K L I L N G K Q A K E E A I T E L V D A A I A E K Y I K L I A N A K T V E G V W T L K D E T L K F T V T E
BINARY_5387 T T Y K L I L N G K Q A K E E A I T E L V D A A I A E K Y I K L I A N A K T V E G V W T L K D E T L T A T V T E
BINARY_5388 T T Y K L I L N G K Q A K E E A I T E L V D A A I A E K Y I K L I A N A K T V E G V W T L K D E T L T F T V T E
BINARY_5389 T T Y K L I L N G K Q A K E E A I T E L V D A A I A E K Y I K L I A N A K T V E G V W T L K D E T K K A T V T E
BINARY_5390 T T Y K L I L N G K Q A K E E A I T E L V D A A I A E K Y I K L I A N A K T V E G V W T L K D E T K K F T V T E
BINARY_5391 T T Y K L I L N G K Q A K E E A I T E L V D A A I A E K Y I K L I A N A K T V E G V W T L K D E T K T A T V T E
BINARY_5392 T T Y K L I L N G K Q A K E E A I T E L V D A A I A E K Y I K L I A N A K T V E G V W T L K D E T K T F T V T E
BINARY_5393 T T Y K L I L N G K Q A K E E A I T E L V D A A I A E K Y I K L I A N A K T V E G V W T Y K D E I L K A T V T E
BINARY_5394 T T Y K L I L N G K Q A K E E A I T E L V D A A I A E K Y I K L I A N A K T V E G V W T Y K D E I L K F T V T E
BINARY_5395 T T Y K L I L N G K Q A K E E A I T E L V D A A I A E K Y I K L I A N A K T V E G V W T Y K D E I L T A T V T E
BINARY_5396 T T Y K L I L N G K Q A K E E A I T E L V D A A I A E K Y I K L I A N A K T V E G V W T Y K D E I L T F T V T E
BINARY_5397 T T Y K L I L N G K Q A K E E A I T E L V D A A I A E K Y I K L I A N A K T V E G V W T Y K D E I K K A T V T E
BINARY_5398 T T Y K L I L N G K Q A K E E A I T E L V D A A I A E K Y I K L I A N A K T V E G V W T Y K D E I K K F T V T E
BINARY_5399 T T Y K L I L N G K Q A K E E A I T E L V D A A I A E K Y I K L I A N A K T V E G V W T Y K D E I K T A T V T E
BINARY_5400 T T Y K L I L N G K Q A K E E A I T E L V D A A I A E K Y I K L I A N A K T V E G V W T Y K D E I K T F T V T E
BINARY_5401 T T Y K L I L N G K Q A K E E A I T E L V D A A I A E K Y I K L I A N A K T V E G V W T Y K D E T L K A T V T E
BINARY_5402 T T Y K L I L N G K Q A K E E A I T E L V D A A I A E K Y I K L I A N A K T V E G V W T Y K D E T L K F T V T E
BINARY_5403 T T Y K L I L N G K Q A K E E A I T E L V D A A I A E K Y I K L I A N A K T V E G V W T Y K D E T L T A T V T E
BINARY_5404 T T Y K L I L N G K Q A K E E A I T E L V D A A I A E K Y I K L I A N A K T V E G V W T Y K D E T L T F T V T E
BINARY_5405 T T Y K L I L N G K Q A K E E A I T E L V D A A I A E K Y I K L I A N A K T V E G V W T Y K D E T K K A T V T E
BINARY_5406 T T Y K L I L N G K Q A K E E A I T E L V D A A I A E K Y I K L I A N A K T V E G V W T Y K D E T K K F T V T E
BINARY_5407 T T Y K L I L N G K Q A K E E A I T E L V D A A I A E K Y I K L I A N A K T V E G V W T Y K D E T K T A T V T E
BINARY_5408 T T Y K L I L N G K Q A K E E A I T E L V D A A I A E K Y I K L I A N A K T V E G V W T Y K D E T K T F T V T E
BINARY_5409 T T Y K L I L N G K Q A K E E A I T E L V D A A I A E K Y I K L Y A N A K T V E G V W T L K D E I L K A T V T E
BINARY_5410 T T Y K L I L N G K Q A K E E A I T E L V D A A I A E K Y I K L Y A N A K T V E G V W T L K D E I L K F T V T E
BINARY_5411 T T Y K L I L N G K Q A K E E A I T E L V D A A I A E K Y I K L Y A N A K T V E G V W T L K D E I L T A T V T E
BINARY_5412 T T Y K L I L N G K Q A K E E A I T E L V D A A I A E K Y I K L Y A N A K T V E G V W T L K D E I L T F T V T E
BINARY_5413 T T Y K L I L N G K Q A K E E A I T E L V D A A I A E K Y I K L Y A N A K T V E G V W T L K D E I K K A T V T E
BINARY_5414 T T Y K L I L N G K Q A K E E A I T E L V D A A I A E K Y I K L Y A N A K T V E G V W T L K D E I K K F T V T E
BINARY_5415 T T Y K L I L N G K Q A K E E A I T E L V D A A I A E K Y I K L Y A N A K T V E G V W T L K D E I K T A T V T E
BINARY_5416 T T Y K L I L N G K Q A K E E A I T E L V D A A I A E K Y I K L Y A N A K T V E G V W T L K D E I K T F T V T E
BINARY_5417 T T Y K L I L N G K Q A K E E A I T E L V D A A I A E K Y I K L Y A N A K T V E G V W T L K D E T L K A T V T E
BINARY_5418 T T Y K L I L N G K Q A K E E A I T E L V D A A I A E K Y I K L Y A N A K T V E G V W T L K D E T L K F T V T E
BINARY_5419 T T Y K L I L N G K Q A K E E A I T E L V D A A I A E K Y I K L Y A N A K T V E G V W T L K D E T L T A T V T E
BINARY_5420 T T Y K L I L N G K Q A K E E A I T E L V D A A I A E K Y I K L Y A N A K T V E G V W T L K D E T L T F T V T E
BINARY_5421 T T Y K L I L N G K Q A K E E A I T E L V D A A I A E K Y I K L Y A N A K T V E G V W T L K D E T K K A T V T E
BINARY_5422 T T Y K L I L N G K Q A K E E A I T E L V D A A I A E K Y I K L Y A N A K T V E G V W T L K D E T K K F T V T E
BINARY_5423 T T Y K L I L N G K Q A K E E A I T E L V D A A I A E K Y I K L Y A N A K T V E G V W T L K D E T K T A T V T E
BINARY_5424 T T Y K L I L N G K Q A K E E A I T E L V D A A I A E K Y I K L Y A N A K T V E G V W T L K D E T K T F T V T E
BINARY_5425 T T Y K L I L N G K Q A K E E A I T E L V D A A I A E K Y I K L Y A N A K T V E G V W T Y K D E I L K A T V T E
BINARY_5426 T T Y K L I L N G K Q A K E E A I T E L V D A A I A E K Y I K L Y A N A K T V E G V W T Y K D E I L K F T V T E
BINARY_5427 T T Y K L I L N G K Q A K E E A I T E L V D A A I A E K Y I K L Y A N A K T V E G V W T Y K D E I L T A T V T E
BINARY_5428 T T Y K L I L N G K Q A K E E A I T E L V D A A I A E K Y I K L Y A N A K T V E G V W T Y K D E I L T F T V T E
BINARY_5429 T T Y K L I L N G K Q A K E E A I T E L V D A A I A E K Y I K L Y A N A K T V E G V W T Y K D E I K K A T V T E
BINARY_5430 T T Y K L I L N G K Q A K E E A I T E L V D A A I A E K Y I K L Y A N A K T V E G V W T Y K D E I K K F T V T E
BINARY_5431 T T Y K L I L N G K Q A K E E A I T E L V D A A I A E K Y I K L Y A N A K T V E G V W T Y K D E I K T A T V T E

BINARY_5432 T T Y K L I L N G K Q A K E E A I T E L V D A A I A E K Y I K L Y A N A K T V E G V W T Y K D E I K T F T V T E
BINARY_5433 T T Y K L I L N G K Q A K E E A I T E L V D A A I A E K Y I K L Y A N A K T V E G V W T Y K D E T L K A T V T E
BINARY_5434 T T Y K L I L N G K Q A K E E A I T E L V D A A I A E K Y I K L Y A N A K T V E G V W T Y K D E T L K F T V T E
BINARY_5435 T T Y K L I L N G K Q A K E E A I T E L V D A A I A E K Y I K L Y A N A K T V E G V W T Y K D E T L T A T V T E
BINARY_5436 T T Y K L I L N G K Q A K E E A I T E L V D A A I A E K Y I K L Y A N A K T V E G V W T Y K D E T L T F T V T E
BINARY_5437 T T Y K L I L N G K Q A K E E A I T E L V D A A I A E K Y I K L Y A N A K T V E G V W T Y K D E T K K A T V T E
BINARY_5438 T T Y K L I L N G K Q A K E E A I T E L V D A A I A E K Y I K L Y A N A K T V E G V W T Y K D E T K K F T V T E
BINARY_5439 T T Y K L I L N G K Q A K E E A I T E L V D A A I A E K Y I K L Y A N A K T V E G V W T Y K D E T K T A T V T E
BINARY_5440 T T Y K L I L N G K Q A K E E A I T E L V D A A I A E K Y I K L Y A N A K T V E G V W T Y K D E T K T F T V T E
BINARY_5441 T T Y K L I L N G K Q A K E E A I T E L V D A A I A E K Y F K L I A N A K T V E G V W T L K D E I L K A T V T E
BINARY_5442 T T Y K L I L N G K Q A K E E A I T E L V D A A I A E K Y F K L I A N A K T V E G V W T L K D E I L K F T V T E
BINARY_5443 T T Y K L I L N G K Q A K E E A I T E L V D A A I A E K Y F K L I A N A K T V E G V W T L K D E I L T A T V T E
BINARY_5444 T T Y K L I L N G K Q A K E E A I T E L V D A A I A E K Y F K L I A N A K T V E G V W T L K D E I L T F T V T E
BINARY_5445 T T Y K L I L N G K Q A K E E A I T E L V D A A I A E K Y F K L I A N A K T V E G V W T L K D E I K K A T V T E
BINARY_5446 T T Y K L I L N G K Q A K E E A I T E L V D A A I A E K Y F K L I A N A K T V E G V W T L K D E I K K F T V T E
BINARY_5447 T T Y K L I L N G K Q A K E E A I T E L V D A A I A E K Y F K L I A N A K T V E G V W T L K D E I K T A T V T E
BINARY_5448 T T Y K L I L N G K Q A K E E A I T E L V D A A I A E K Y F K L I A N A K T V E G V W T L K D E I K T F T V T E
BINARY_5449 T T Y K L I L N G K Q A K E E A I T E L V D A A I A E K Y F K L I A N A K T V E G V W T L K D E T L K A T V T E
BINARY_5450 T T Y K L I L N G K Q A K E E A I T E L V D A A I A E K Y F K L I A N A K T V E G V W T L K D E T L K F T V T E
BINARY_5451 T T Y K L I L N G K Q A K E E A I T E L V D A A I A E K Y F K L I A N A K T V E G V W T L K D E T L T A T V T E
BINARY_5452 T T Y K L I L N G K Q A K E E A I T E L V D A A I A E K Y F K L I A N A K T V E G V W T L K D E T L T F T V T E
BINARY_5453 T T Y K L I L N G K Q A K E E A I T E L V D A A I A E K Y F K L I A N A K T V E G V W T L K D E T K K A T V T E
BINARY_5454 T T Y K L I L N G K Q A K E E A I T E L V D A A I A E K Y F K L I A N A K T V E G V W T L K D E T K K F T V T E
BINARY_5455 T T Y K L I L N G K Q A K E E A I T E L V D A A I A E K Y F K L I A N A K T V E G V W T L K D E T K T A T V T E
BINARY_5456 T T Y K L I L N G K Q A K E E A I T E L V D A A I A E K Y F K L I A N A K T V E G V W T L K D E T K T F T V T E
BINARY_5457 T T Y K L I L N G K Q A K E E A I T E L V D A A I A E K Y F K L I A N A K T V E G V W T Y K D E I L K A T V T E
BINARY_5458 T T Y K L I L N G K Q A K E E A I T E L V D A A I A E K Y F K L I A N A K T V E G V W T Y K D E I L K F T V T E
BINARY_5459 T T Y K L I L N G K Q A K E E A I T E L V D A A I A E K Y F K L I A N A K T V E G V W T Y K D E I L T A T V T E
BINARY_5460 T T Y K L I L N G K Q A K E E A I T E L V D A A I A E K Y F K L I A N A K T V E G V W T Y K D E I L T F T V T E
BINARY_5461 T T Y K L I L N G K Q A K E E A I T E L V D A A I A E K Y F K L I A N A K T V E G V W T Y K D E I K K A T V T E
BINARY_5462 T T Y K L I L N G K Q A K E E A I T E L V D A A I A E K Y F K L I A N A K T V E G V W T Y K D E I K K F T V T E
BINARY_5463 T T Y K L I L N G K Q A K E E A I T E L V D A A I A E K Y F K L I A N A K T V E G V W T Y K D E I K T A T V T E
BINARY_5464 T T Y K L I L N G K Q A K E E A I T E L V D A A I A E K Y F K L I A N A K T V E G V W T Y K D E I K T F T V T E
BINARY_5465 T T Y K L I L N G K Q A K E E A I T E L V D A A I A E K Y F K L I A N A K T V E G V W T Y K D E T L K A T V T E
BINARY_5466 T T Y K L I L N G K Q A K E E A I T E L V D A A I A E K Y F K L I A N A K T V E G V W T Y K D E T L K F T V T E
BINARY_5467 T T Y K L I L N G K Q A K E E A I T E L V D A A I A E K Y F K L I A N A K T V E G V W T Y K D E T L T A T V T E
BINARY_5468 T T Y K L I L N G K Q A K E E A I T E L V D A A I A E K Y F K L I A N A K T V E G V W T Y K D E T L T F T V T E
BINARY_5469 T T Y K L I L N G K Q A K E E A I T E L V D A A I A E K Y F K L I A N A K T V E G V W T Y K D E T K K A T V T E
BINARY_5470 T T Y K L I L N G K Q A K E E A I T E L V D A A I A E K Y F K L I A N A K T V E G V W T Y K D E T K K F T V T E
BINARY_5471 T T Y K L I L N G K Q A K E E A I T E L V D A A I A E K Y F K L I A N A K T V E G V W T Y K D E T K T A T V T E
BINARY_5472 T T Y K L I L N G K Q A K E E A I T E L V D A A I A E K Y F K L I A N A K T V E G V W T Y K D E T K T F T V T E
BINARY_5473 T T Y K L I L N G K Q A K E E A I T E L V D A A I A E K Y F K L Y A N A K T V E G V W T L K D E I L K A T V T E
BINARY_5474 T T Y K L I L N G K Q A K E E A I T E L V D A A I A E K Y F K L Y A N A K T V E G V W T L K D E I L K F T V T E
BINARY_5475 T T Y K L I L N G K Q A K E E A I T E L V D A A I A E K Y F K L Y A N A K T V E G V W T L K D E I L T A T V T E
BINARY_5476 T T Y K L I L N G K Q A K E E A I T E L V D A A I A E K Y F K L Y A N A K T V E G V W T L K D E I L T F T V T E
BINARY_5477 T T Y K L I L N G K Q A K E E A I T E L V D A A I A E K Y F K L Y A N A K T V E G V W T L K D E I K K A T V T E
BINARY_5478 T T Y K L I L N G K Q A K E E A I T E L V D A A I A E K Y F K L Y A N A K T V E G V W T L K D E I K K F T V T E
BINARY_5479 T T Y K L I L N G K Q A K E E A I T E L V D A A I A E K Y F K L Y A N A K T V E G V W T L K D E I K T A T V T E
BINARY_5480 T T Y K L I L N G K Q A K E E A I T E L V D A A I A E K Y F K L Y A N A K T V E G V W T L K D E I K T F T V T E
BINARY_5481 T T Y K L I L N G K Q A K E E A I T E L V D A A I A E K Y F K L Y A N A K T V E G V W T L K D E T L K A T V T E
BINARY_5482 T T Y K L I L N G K Q A K E E A I T E L V D A A I A E K Y F K L Y A N A K T V E G V W T L K D E T L K F T V T E
BINARY_5483 T T Y K L I L N G K Q A K E E A I T E L V D A A I A E K Y F K L Y A N A K T V E G V W T L K D E T L T A T V T E
BINARY_5484 T T Y K L I L N G K Q A K E E A I T E L V D A A I A E K Y F K L Y A N A K T V E G V W T L K D E T L T F T V T E
BINARY_5485 T T Y K L I L N G K Q A K E E A I T E L V D A A I A E K Y F K L Y A N A K T V E G V W T L K D E T K K A T V T E
BINARY_5486 T T Y K L I L N G K Q A K E E A I T E L V D A A I A E K Y F K L Y A N A K T V E G V W T L K D E T K K F T V T E
BINARY_5487 T T Y K L I L N G K Q A K E E A I T E L V D A A I A E K Y F K L Y A N A K T V E G V W T L K D E T K T A T V T E
BINARY_5488 T T Y K L I L N G K Q A K E E A I T E L V D A A I A E K Y F K L Y A N A K T V E G V W T L K D E T K T F T V T E
BINARY_5489 T T Y K L I L N G K Q A K E E A I T E L V D A A I A E K Y F K L Y A N A K T V E G V W T Y K D E I L K A T V T E
BINARY_5490 T T Y K L I L N G K Q A K E E A I T E L V D A A I A E K Y F K L Y A N A K T V E G V W T Y K D E I L K F T V T E
BINARY_5491 T T Y K L I L N G K Q A K E E A I T E L V D A A I A E K Y F K L Y A N A K T V E G V W T Y K D E I L T A T V T E
BINARY_5492 T T Y K L I L N G K Q A K E E A I T E L V D A A I A E K Y F K L Y A N A K T V E G V W T Y K D E I L T F T V T E
BINARY_5493 T T Y K L I L N G K Q A K E E A I T E L V D A A I A E K Y F K L Y A N A K T V E G V W T Y K D E I K K A T V T E
BINARY_5494 T T Y K L I L N G K Q A K E E A I T E L V D A A I A E K Y F K L Y A N A K T V E G V W T Y K D E I K K F T V T E
BINARY_5495 T T Y K L I L N G K Q A K E E A I T E L V D A A I A E K Y F K L Y A N A K T V E G V W T Y K D E I K T A T V T E
BINARY_5496 T T Y K L I L N G K Q A K E E A I T E L V D A A I A E K Y F K L Y A N A K T V E G V W T Y K D E I K T F T V T E
BINARY_5497 T T Y K L I L N G K Q A K E E A I T E L V D A A I A E K Y F K L Y A N A K T V E G V W T Y K D E T L K A T V T E
BINARY_5498 T T Y K L I L N G K Q A K E E A I T E L V D A A I A E K Y F K L Y A N A K T V E G V W T Y K D E T L K F T V T E
BINARY_5499 T T Y K L I L N G K Q A K E E A I T E L V D A A I A E K Y F K L Y A N A K T V E G V W T Y K D E T L T A T V T E
BINARY_5500 T T Y K L I L N G K Q A K E E A I T E L V D A A I A E K Y F K L Y A N A K T V E G V W T Y K D E T L T F T V T E
BINARY_5501 T T Y K L I L N G K Q A K E E A I T E L V D A A I A E K Y F K L Y A N A K T V E G V W T Y K D E T K K A T V T E
BINARY_5502 T T Y K L I L N G K Q A K E E A I T E L V D A A I A E K Y F K L Y A N A K T V E G V W T Y K D E T K K F T V T E
BINARY_5503 T T Y K L I L N G K Q A K E E A I T E L V D A A I A E K Y F K L Y A N A K T V E G V W T Y K D E T K T A T V T E
BINARY_5504 T T Y K L I L N G K Q A K E E A I T E L V D A A I A E K Y F K L Y A N A K T V E G V W T Y K D E T K T F T V T E
BINARY_5505 T T Y K L I L N G K Q A K E E A I T E L V D A A T A E K Y I K L I A N A K T V E G V W T L K D E I L K A T V T E
BINARY_5506 T T Y K L I L N G K Q A K E E A I T E L V D A A T A E K Y I K L I A N A K T V E G V W T L K D E I L K F T V T E
BINARY_5507 T T Y K L I L N G K Q A K E E A I T E L V D A A T A E K Y I K L I A N A K T V E G V W T L K D E I L T A T V T E
BINARY_5508 T T Y K L I L N G K Q A K E E A I T E L V D A A T A E K Y I K L I A N A K T V E G V W T L K D E I L T F T V T E
BINARY_5509 T T Y K L I L N G K Q A K E E A I T E L V D A A T A E K Y I K L I A N A K T V E G V W T L K D E I K K A T V T E
BINARY_5510 T T Y K L I L N G K Q A K E E A I T E L V D A A T A E K Y I K L I A N A K T V E G V W T L K D E I K K F T V T E
BINARY_5511 T T Y K L I L N G K Q A K E E A I T E L V D A A T A E K Y I K L I A N A K T V E G V W T L K D E I K T A T V T E
BINARY_5512 T T Y K L I L N G K Q A K E E A I T E L V D A A T A E K Y I K L I A N A K T V E G V W T L K D E I K T F T V T E
BINARY_5513 T T Y K L I L N G K Q A K E E A I T E L V D A A T A E K Y I K L I A N A K T V E G V W T L K D E T L K A T V T E
BINARY_5514 T T Y K L I L N G K Q A K E E A I T E L V D A A T A E K Y I K L I A N A K T V E G V W T L K D E T L K F T V T E
BINARY_5515 T T Y K L I L N G K Q A K E E A I T E L V D A A T A E K Y I K L I A N A K T V E G V W T L K D E T L T A T V T E
BINARY_5516 T T Y K L I L N G K Q A K E E A I T E L V D A A T A E K Y I K L I A N A K T V E G V W T L K D E T L T F T V T E
BINARY_5517 T T Y K L I L N G K Q A K E E A I T E L V D A A T A E K Y I K L I A N A K T V E G V W T L K D E T K K A T V T E
BINARY_5518 T T Y K L I L N G K Q A K E E A I T E L V D A A T A E K Y I K L I A N A K T V E G V W T L K D E T K K F T V T E
BINARY_5519 T T Y K L I L N G K Q A K E E A I T E L V D A A T A E K Y I K L I A N A K T V E G V W T L K D E T K T A T V T E
BINARY_5520 T T Y K L I L N G K Q A K E E A I T E L V D A A T A E K Y I K L I A N A K T V E G V W T L K D E T K T F T V T E
BINARY_5521 T T Y K L I L N G K Q A K E E A I T E L V D A A T A E K Y I K L I A N A K T V E G V W T Y K D E I L K A T V T E
BINARY_5522 T T Y K L I L N G K Q A K E E A I T E L V D A A T A E K Y I K L I A N A K T V E G V W T Y K D E I L K F T V T E
BINARY_5523 T T Y K L I L N G K Q A K E E A I T E L V D A A T A E K Y I K L I A N A K T V E G V W T Y K D E I L T A T V T E
BINARY_5524 T T Y K L I L N G K Q A K E E A I T E L V D A A T A E K Y I K L I A N A K T V E G V W T Y K D E I L T F T V T E
BINARY_5525 T T Y K L I L N G K Q A K E E A I T E L V D A A T A E K Y I K L I A N A K T V E G V W T Y K D E I K K A T V T E
BINARY_5526 T T Y K L I L N G K Q A K E E A I T E L V D A A T A E K Y I K L I A N A K T V E G V W T Y K D E I K K F T V T E

BINARY_5527 T T Y K L I L N G K Q A K E E A I T E L V D A A T A E K Y I K L I A N A K T V E G V W T Y K D E I K T A T V T E
BINARY_5528 T T Y K L I L N G K Q A K E E A I T E L V D A A T A E K Y I K L I A N A K T V E G V W T Y K D E I K T F T V T E
BINARY_5529 T T Y K L I L N G K Q A K E E A I T E L V D A A T A E K Y I K L I A N A K T V E G V W T Y K D E T L K A T V T E
BINARY_5530 T T Y K L I L N G K Q A K E E A I T E L V D A A T A E K Y I K L I A N A K T V E G V W T Y K D E T L K F T V T E
BINARY_5531 T T Y K L I L N G K Q A K E E A I T E L V D A A T A E K Y I K L I A N A K T V E G V W T Y K D E T L T A T V T E
BINARY_5532 T T Y K L I L N G K Q A K E E A I T E L V D A A T A E K Y I K L I A N A K T V E G V W T Y K D E T L T F T V T E
BINARY_5533 T T Y K L I L N G K Q A K E E A I T E L V D A A T A E K Y I K L I A N A K T V E G V W T Y K D E T K K A T V T E
BINARY_5534 T T Y K L I L N G K Q A K E E A I T E L V D A A T A E K Y I K L I A N A K T V E G V W T Y K D E T K K F T V T E
BINARY_5535 T T Y K L I L N G K Q A K E E A I T E L V D A A T A E K Y I K L I A N A K T V E G V W T Y K D E T K T A T V T E
BINARY_5536 T T Y K L I L N G K Q A K E E A I T E L V D A A T A E K Y I K L I A N A K T V E G V W T Y K D E T K T F T V T E
BINARY_5537 T T Y K L I L N G K Q A K E E A I T E L V D A A T A E K Y I K L Y A N A K T V E G V W T L K D E I L K A T V T E
BINARY_5538 T T Y K L I L N G K Q A K E E A I T E L V D A A T A E K Y I K L Y A N A K T V E G V W T L K D E I L K F T V T E
BINARY_5539 T T Y K L I L N G K Q A K E E A I T E L V D A A T A E K Y I K L Y A N A K T V E G V W T L K D E I L T A T V T E
BINARY_5540 T T Y K L I L N G K Q A K E E A I T E L V D A A T A E K Y I K L Y A N A K T V E G V W T L K D E I L T F T V T E
BINARY_5541 T T Y K L I L N G K Q A K E E A I T E L V D A A T A E K Y I K L Y A N A K T V E G V W T L K D E I K K A T V T E
BINARY_5542 T T Y K L I L N G K Q A K E E A I T E L V D A A T A E K Y I K L Y A N A K T V E G V W T L K D E I K K F T V T E
BINARY_5543 T T Y K L I L N G K Q A K E E A I T E L V D A A T A E K Y I K L Y A N A K T V E G V W T L K D E I K T A T V T E
BINARY_5544 T T Y K L I L N G K Q A K E E A I T E L V D A A T A E K Y I K L Y A N A K T V E G V W T L K D E I K T F T V T E
BINARY_5545 T T Y K L I L N G K Q A K E E A I T E L V D A A T A E K Y I K L Y A N A K T V E G V W T L K D E T L K A T V T E
BINARY_5546 T T Y K L I L N G K Q A K E E A I T E L V D A A T A E K Y I K L Y A N A K T V E G V W T L K D E T L K F T V T E
BINARY_5547 T T Y K L I L N G K Q A K E E A I T E L V D A A T A E K Y I K L Y A N A K T V E G V W T L K D E T L T A T V T E
BINARY_5548 T T Y K L I L N G K Q A K E E A I T E L V D A A T A E K Y I K L Y A N A K T V E G V W T L K D E T L T F T V T E
BINARY_5549 T T Y K L I L N G K Q A K E E A I T E L V D A A T A E K Y I K L Y A N A K T V E G V W T L K D E T K K A T V T E
BINARY_5550 T T Y K L I L N G K Q A K E E A I T E L V D A A T A E K Y I K L Y A N A K T V E G V W T L K D E T K K F T V T E
BINARY_5551 T T Y K L I L N G K Q A K E E A I T E L V D A A T A E K Y I K L Y A N A K T V E G V W T L K D E T K T A T V T E
BINARY_5552 T T Y K L I L N G K Q A K E E A I T E L V D A A T A E K Y I K L Y A N A K T V E G V W T L K D E T K T F T V T E
BINARY_5553 T T Y K L I L N G K Q A K E E A I T E L V D A A T A E K Y I K L Y A N A K T V E G V W T Y K D E I L K A T V T E
BINARY_5554 T T Y K L I L N G K Q A K E E A I T E L V D A A T A E K Y I K L Y A N A K T V E G V W T Y K D E I L K F T V T E
BINARY_5555 T T Y K L I L N G K Q A K E E A I T E L V D A A T A E K Y I K L Y A N A K T V E G V W T Y K D E I L T A T V T E
BINARY_5556 T T Y K L I L N G K Q A K E E A I T E L V D A A T A E K Y I K L Y A N A K T V E G V W T Y K D E I L T F T V T E
BINARY_5557 T T Y K L I L N G K Q A K E E A I T E L V D A A T A E K Y I K L Y A N A K T V E G V W T Y K D E I K K A T V T E
BINARY_5558 T T Y K L I L N G K Q A K E E A I T E L V D A A T A E K Y I K L Y A N A K T V E G V W T Y K D E I K K F T V T E
BINARY_5559 T T Y K L I L N G K Q A K E E A I T E L V D A A T A E K Y I K L Y A N A K T V E G V W T Y K D E I K T A T V T E
BINARY_5560 T T Y K L I L N G K Q A K E E A I T E L V D A A T A E K Y I K L Y A N A K T V E G V W T Y K D E I K T F T V T E
BINARY_5561 T T Y K L I L N G K Q A K E E A I T E L V D A A T A E K Y I K L Y A N A K T V E G V W T Y K D E T L K A T V T E
BINARY_5562 T T Y K L I L N G K Q A K E E A I T E L V D A A T A E K Y I K L Y A N A K T V E G V W T Y K D E T L K F T V T E
BINARY_5563 T T Y K L I L N G K Q A K E E A I T E L V D A A T A E K Y I K L Y A N A K T V E G V W T Y K D E T L T A T V T E
BINARY_5564 T T Y K L I L N G K Q A K E E A I T E L V D A A T A E K Y I K L Y A N A K T V E G V W T Y K D E T L T F T V T E
BINARY_5565 T T Y K L I L N G K Q A K E E A I T E L V D A A T A E K Y I K L Y A N A K T V E G V W T Y K D E T K K A T V T E
BINARY_5566 T T Y K L I L N G K Q A K E E A I T E L V D A A T A E K Y I K L Y A N A K T V E G V W T Y K D E T K K F T V T E
BINARY_5567 T T Y K L I L N G K Q A K E E A I T E L V D A A T A E K Y I K L Y A N A K T V E G V W T Y K D E T K T A T V T E
BINARY_5568 T T Y K L I L N G K Q A K E E A I T E L V D A A T A E K Y I K L Y A N A K T V E G V W T Y K D E T K T F T V T E
BINARY_5569 T T Y K L I L N G K Q A K E E A I T E L V D A A T A E K Y F K L I A N A K T V E G V W T L K D E I L K A T V T E
BINARY_5570 T T Y K L I L N G K Q A K E E A I T E L V D A A T A E K Y F K L I A N A K T V E G V W T L K D E I L K F T V T E
BINARY_5571 T T Y K L I L N G K Q A K E E A I T E L V D A A T A E K Y F K L I A N A K T V E G V W T L K D E I L T A T V T E
BINARY_5572 T T Y K L I L N G K Q A K E E A I T E L V D A A T A E K Y F K L I A N A K T V E G V W T L K D E I L T F T V T E
BINARY_5573 T T Y K L I L N G K Q A K E E A I T E L V D A A T A E K Y F K L I A N A K T V E G V W T L K D E I K K A T V T E
BINARY_5574 T T Y K L I L N G K Q A K E E A I T E L V D A A T A E K Y F K L I A N A K T V E G V W T L K D E I K K F T V T E
BINARY_5575 T T Y K L I L N G K Q A K E E A I T E L V D A A T A E K Y F K L I A N A K T V E G V W T L K D E I K T A T V T E
BINARY_5576 T T Y K L I L N G K Q A K E E A I T E L V D A A T A E K Y F K L I A N A K T V E G V W T L K D E I K T F T V T E
BINARY_5577 T T Y K L I L N G K Q A K E E A I T E L V D A A T A E K Y F K L I A N A K T V E G V W T L K D E T L K A T V T E
BINARY_5578 T T Y K L I L N G K Q A K E E A I T E L V D A A T A E K Y F K L I A N A K T V E G V W T L K D E T L K F T V T E
BINARY_5579 T T Y K L I L N G K Q A K E E A I T E L V D A A T A E K Y F K L I A N A K T V E G V W T L K D E T L T A T V T E
BINARY_5580 T T Y K L I L N G K Q A K E E A I T E L V D A A T A E K Y F K L I A N A K T V E G V W T L K D E T L T F T V T E
BINARY_5581 T T Y K L I L N G K Q A K E E A I T E L V D A A T A E K Y F K L I A N A K T V E G V W T L K D E T K K A T V T E
BINARY_5582 T T Y K L I L N G K Q A K E E A I T E L V D A A T A E K Y F K L I A N A K T V E G V W T L K D E T K K F T V T E
BINARY_5583 T T Y K L I L N G K Q A K E E A I T E L V D A A T A E K Y F K L I A N A K T V E G V W T L K D E T K T A T V T E
BINARY_5584 T T Y K L I L N G K Q A K E E A I T E L V D A A T A E K Y F K L I A N A K T V E G V W T L K D E T K T F T V T E
BINARY_5585 T T Y K L I L N G K Q A K E E A I T E L V D A A T A E K Y F K L I A N A K T V E G V W T Y K D E I L K A T V T E
BINARY_5586 T T Y K L I L N G K Q A K E E A I T E L V D A A T A E K Y F K L I A N A K T V E G V W T Y K D E I L K F T V T E
BINARY_5587 T T Y K L I L N G K Q A K E E A I T E L V D A A T A E K Y F K L I A N A K T V E G V W T Y K D E I L T A T V T E
BINARY_5588 T T Y K L I L N G K Q A K E E A I T E L V D A A T A E K Y F K L I A N A K T V E G V W T Y K D E I L T F T V T E
BINARY_5589 T T Y K L I L N G K Q A K E E A I T E L V D A A T A E K Y F K L I A N A K T V E G V W T Y K D E I K K A T V T E
BINARY_5590 T T Y K L I L N G K Q A K E E A I T E L V D A A T A E K Y F K L I A N A K T V E G V W T Y K D E I K K F T V T E
BINARY_5591 T T Y K L I L N G K Q A K E E A I T E L V D A A T A E K Y F K L I A N A K T V E G V W T Y K D E I K T A T V T E
BINARY_5592 T T Y K L I L N G K Q A K E E A I T E L V D A A T A E K Y F K L I A N A K T V E G V W T Y K D E I K T F T V T E
BINARY_5593 T T Y K L I L N G K Q A K E E A I T E L V D A A T A E K Y F K L I A N A K T V E G V W T Y K D E T L K A T V T E
BINARY_5594 T T Y K L I L N G K Q A K E E A I T E L V D A A T A E K Y F K L I A N A K T V E G V W T Y K D E T L K F T V T E
BINARY_5595 T T Y K L I L N G K Q A K E E A I T E L V D A A T A E K Y F K L I A N A K T V E G V W T Y K D E T L T A T V T E
BINARY_5596 T T Y K L I L N G K Q A K E E A I T E L V D A A T A E K Y F K L I A N A K T V E G V W T Y K D E T L T F T V T E
BINARY_5597 T T Y K L I L N G K Q A K E E A I T E L V D A A T A E K Y F K L I A N A K T V E G V W T Y K D E T K K A T V T E
BINARY_5598 T T Y K L I L N G K Q A K E E A I T E L V D A A T A E K Y F K L I A N A K T V E G V W T Y K D E T K K F T V T E
BINARY_5599 T T Y K L I L N G K Q A K E E A I T E L V D A A T A E K Y F K L I A N A K T V E G V W T Y K D E T K T A T V T E
BINARY_5600 T T Y K L I L N G K Q A K E E A I T E L V D A A T A E K Y F K L I A N A K T V E G V W T Y K D E T K T F T V T E
BINARY_5601 T T Y K L I L N G K Q A K E E A I T E L V D A A T A E K Y F K L Y A N A K T V E G V W T L K D E I L K A T V T E
BINARY_5602 T T Y K L I L N G K Q A K E E A I T E L V D A A T A E K Y F K L Y A N A K T V E G V W T L K D E I L K F T V T E
BINARY_5603 T T Y K L I L N G K Q A K E E A I T E L V D A A T A E K Y F K L Y A N A K T V E G V W T L K D E I L T A T V T E
BINARY_5604 T T Y K L I L N G K Q A K E E A I T E L V D A A T A E K Y F K L Y A N A K T V E G V W T L K D E I L T F T V T E
BINARY_5605 T T Y K L I L N G K Q A K E E A I T E L V D A A T A E K Y F K L Y A N A K T V E G V W T L K D E I K K A T V T E
BINARY_5606 T T Y K L I L N G K Q A K E E A I T E L V D A A T A E K Y F K L Y A N A K T V E G V W T L K D E I K K F T V T E
BINARY_5607 T T Y K L I L N G K Q A K E E A I T E L V D A A T A E K Y F K L Y A N A K T V E G V W T L K D E I K T A T V T E
BINARY_5608 T T Y K L I L N G K Q A K E E A I T E L V D A A T A E K Y F K L Y A N A K T V E G V W T L K D E I K T F T V T E
BINARY_5609 T T Y K L I L N G K Q A K E E A I T E L V D A A T A E K Y F K L Y A N A K T V E G V W T L K D E T L K A T V T E
BINARY_5610 T T Y K L I L N G K Q A K E E A I T E L V D A A T A E K Y F K L Y A N A K T V E G V W T L K D E T L K F T V T E
BINARY_5611 T T Y K L I L N G K Q A K E E A I T E L V D A A T A E K Y F K L Y A N A K T V E G V W T L K D E T L T A T V T E
BINARY_5612 T T Y K L I L N G K Q A K E E A I T E L V D A A T A E K Y F K L Y A N A K T V E G V W T L K D E T L T F T V T E
BINARY_5613 T T Y K L I L N G K Q A K E E A I T E L V D A A T A E K Y F K L Y A N A K T V E G V W T L K D E T K K A T V T E
BINARY_5614 T T Y K L I L N G K Q A K E E A I T E L V D A A T A E K Y F K L Y A N A K T V E G V W T L K D E T K K F T V T E
BINARY_5615 T T Y K L I L N G K Q A K E E A I T E L V D A A T A E K Y F K L Y A N A K T V E G V W T L K D E T K T A T V T E
BINARY_5616 T T Y K L I L N G K Q A K E E A I T E L V D A A T A E K Y F K L Y A N A K T V E G V W T L K D E T K T F T V T E
BINARY_5617 T T Y K L I L N G K Q A K E E A I T E L V D A A T A E K Y F K L Y A N A K T V E G V W T Y K D E I L K A T V T E
BINARY_5618 T T Y K L I L N G K Q A K E E A I T E L V D A A T A E K Y F K L Y A N A K T V E G V W T Y K D E I L K F T V T E
BINARY_5619 T T Y K L I L N G K Q A K E E A I T E L V D A A T A E K Y F K L Y A N A K T V E G V W T Y K D E I L T A T V T E
BINARY_5620 T T Y K L I L N G K Q A K E E A I T E L V D A A T A E K Y F K L Y A N A K T V E G V W T Y K D E I L T F T V T E
BINARY_5621 T T Y K L I L N G K Q A K E E A I T E L V D A A T A E K Y F K L Y A N A K T V E G V W T Y K D E I K K A T V T E

BINARY_5622 T T Y K L I L N G K Q A K E E A I T E L V D A A T A E K Y F K L Y A N A K T V E G V W T Y K D E I K K F T V T E
BINARY_5623 T T Y K L I L N G K Q A K E E A I T E L V D A A T A E K Y F K L Y A N A K T V E G V W T Y K D E I K T A T V T E
BINARY_5624 T T Y K L I L N G K Q A K E E A I T E L V D A A T A E K Y F K L Y A N A K T V E G V W T Y K D E I K T F T V T E
BINARY_5625 T T Y K L I L N G K Q A K E E A I T E L V D A A T A E K Y F K L Y A N A K T V E G V W T Y K D E T L K A T V T E
BINARY_5626 T T Y K L I L N G K Q A K E E A I T E L V D A A T A E K Y F K L Y A N A K T V E G V W T Y K D E T L K F T V T E
BINARY_5627 T T Y K L I L N G K Q A K E E A I T E L V D A A T A E K Y F K L Y A N A K T V E G V W T Y K D E T L T A T V T E
BINARY_5628 T T Y K L I L N G K Q A K E E A I T E L V D A A T A E K Y F K L Y A N A K T V E G V W T Y K D E T L T F T V T E
BINARY_5629 T T Y K L I L N G K Q A K E E A I T E L V D A A T A E K Y F K L Y A N A K T V E G V W T Y K D E T K K A T V T E
BINARY_5630 T T Y K L I L N G K Q A K E E A I T E L V D A A T A E K Y F K L Y A N A K T V E G V W T Y K D E T K K F T V T E
BINARY_5631 T T Y K L I L N G K Q A K E E A I T E L V D A A T A E K Y F K L Y A N A K T V E G V W T Y K D E T K T A T V T E
BINARY_5632 T T Y K L I L N G K Q A K E E A I T E L V D A A T A E K Y F K L Y A N A K T V E G V W T Y K D E T K T F T V T E
BINARY_5633 T T Y K L I L N G K Q A K E E A I T E A V D A G I A E K Y I K L I A N A K T V E G V W T L K D E I L K A T V T E
BINARY_5634 T T Y K L I L N G K Q A K E E A I T E A V D A G I A E K Y I K L I A N A K T V E G V W T L K D E I L K F T V T E
BINARY_5635 T T Y K L I L N G K Q A K E E A I T E A V D A G I A E K Y I K L I A N A K T V E G V W T L K D E I L T A T V T E
BINARY_5636 T T Y K L I L N G K Q A K E E A I T E A V D A G I A E K Y I K L I A N A K T V E G V W T L K D E I L T F T V T E
BINARY_5637 T T Y K L I L N G K Q A K E E A I T E A V D A G I A E K Y I K L I A N A K T V E G V W T L K D E I K K A T V T E
BINARY_5638 T T Y K L I L N G K Q A K E E A I T E A V D A G I A E K Y I K L I A N A K T V E G V W T L K D E I K K F T V T E
BINARY_5639 T T Y K L I L N G K Q A K E E A I T E A V D A G I A E K Y I K L I A N A K T V E G V W T L K D E I K T A T V T E
BINARY_5640 T T Y K L I L N G K Q A K E E A I T E A V D A G I A E K Y I K L I A N A K T V E G V W T L K D E I K T F T V T E
BINARY_5641 T T Y K L I L N G K Q A K E E A I T E A V D A G I A E K Y I K L I A N A K T V E G V W T L K D E T L K A T V T E
BINARY_5642 T T Y K L I L N G K Q A K E E A I T E A V D A G I A E K Y I K L I A N A K T V E G V W T L K D E T L K F T V T E
BINARY_5643 T T Y K L I L N G K Q A K E E A I T E A V D A G I A E K Y I K L I A N A K T V E G V W T L K D E T L T A T V T E
BINARY_5644 T T Y K L I L N G K Q A K E E A I T E A V D A G I A E K Y I K L I A N A K T V E G V W T L K D E T L T F T V T E
BINARY_5645 T T Y K L I L N G K Q A K E E A I T E A V D A G I A E K Y I K L I A N A K T V E G V W T L K D E T K K A T V T E
BINARY_5646 T T Y K L I L N G K Q A K E E A I T E A V D A G I A E K Y I K L I A N A K T V E G V W T L K D E T K K F T V T E
BINARY_5647 T T Y K L I L N G K Q A K E E A I T E A V D A G I A E K Y I K L I A N A K T V E G V W T L K D E T K T A T V T E
BINARY_5648 T T Y K L I L N G K Q A K E E A I T E A V D A G I A E K Y I K L I A N A K T V E G V W T L K D E T K T F T V T E
BINARY_5649 T T Y K L I L N G K Q A K E E A I T E A V D A G I A E K Y I K L I A N A K T V E G V W T Y K D E I L K A T V T E
BINARY_5650 T T Y K L I L N G K Q A K E E A I T E A V D A G I A E K Y I K L I A N A K T V E G V W T Y K D E I L K F T V T E
BINARY_5651 T T Y K L I L N G K Q A K E E A I T E A V D A G I A E K Y I K L I A N A K T V E G V W T Y K D E I L T A T V T E
BINARY_5652 T T Y K L I L N G K Q A K E E A I T E A V D A G I A E K Y I K L I A N A K T V E G V W T Y K D E I L T F T V T E
BINARY_5653 T T Y K L I L N G K Q A K E E A I T E A V D A G I A E K Y I K L I A N A K T V E G V W T Y K D E I K K A T V T E
BINARY_5654 T T Y K L I L N G K Q A K E E A I T E A V D A G I A E K Y I K L I A N A K T V E G V W T Y K D E I K K F T V T E
BINARY_5655 T T Y K L I L N G K Q A K E E A I T E A V D A G I A E K Y I K L I A N A K T V E G V W T Y K D E I K T A T V T E
BINARY_5656 T T Y K L I L N G K Q A K E E A I T E A V D A G I A E K Y I K L I A N A K T V E G V W T Y K D E I K T F T V T E
BINARY_5657 T T Y K L I L N G K Q A K E E A I T E A V D A G I A E K Y I K L I A N A K T V E G V W T Y K D E T L K A T V T E
BINARY_5658 T T Y K L I L N G K Q A K E E A I T E A V D A G I A E K Y I K L I A N A K T V E G V W T Y K D E T L K F T V T E
BINARY_5659 T T Y K L I L N G K Q A K E E A I T E A V D A G I A E K Y I K L I A N A K T V E G V W T Y K D E T L T A T V T E
BINARY_5660 T T Y K L I L N G K Q A K E E A I T E A V D A G I A E K Y I K L I A N A K T V E G V W T Y K D E T L T F T V T E
BINARY_5661 T T Y K L I L N G K Q A K E E A I T E A V D A G I A E K Y I K L I A N A K T V E G V W T Y K D E T K K A T V T E
BINARY_5662 T T Y K L I L N G K Q A K E E A I T E A V D A G I A E K Y I K L I A N A K T V E G V W T Y K D E T K K F T V T E
BINARY_5663 T T Y K L I L N G K Q A K E E A I T E A V D A G I A E K Y I K L I A N A K T V E G V W T Y K D E T K T A T V T E
BINARY_5664 T T Y K L I L N G K Q A K E E A I T E A V D A G I A E K Y I K L I A N A K T V E G V W T Y K D E T K T F T V T E
BINARY_5665 T T Y K L I L N G K Q A K E E A I T E A V D A G I A E K Y I K L Y A N A K T V E G V W T L K D E I L K A T V T E
BINARY_5666 T T Y K L I L N G K Q A K E E A I T E A V D A G I A E K Y I K L Y A N A K T V E G V W T L K D E I L K F T V T E
BINARY_5667 T T Y K L I L N G K Q A K E E A I T E A V D A G I A E K Y I K L Y A N A K T V E G V W T L K D E I L T A T V T E
BINARY_5668 T T Y K L I L N G K Q A K E E A I T E A V D A G I A E K Y I K L Y A N A K T V E G V W T L K D E I L T F T V T E
BINARY_5669 T T Y K L I L N G K Q A K E E A I T E A V D A G I A E K Y I K L Y A N A K T V E G V W T L K D E I K K A T V T E
BINARY_5670 T T Y K L I L N G K Q A K E E A I T E A V D A G I A E K Y I K L Y A N A K T V E G V W T L K D E I K K F T V T E
BINARY_5671 T T Y K L I L N G K Q A K E E A I T E A V D A G I A E K Y I K L Y A N A K T V E G V W T L K D E I K T A T V T E
BINARY_5672 T T Y K L I L N G K Q A K E E A I T E A V D A G I A E K Y I K L Y A N A K T V E G V W T L K D E I K T F T V T E
BINARY_5673 T T Y K L I L N G K Q A K E E A I T E A V D A G I A E K Y I K L Y A N A K T V E G V W T L K D E T L K A T V T E
BINARY_5674 T T Y K L I L N G K Q A K E E A I T E A V D A G I A E K Y I K L Y A N A K T V E G V W T L K D E T L K F T V T E
BINARY_5675 T T Y K L I L N G K Q A K E E A I T E A V D A G I A E K Y I K L Y A N A K T V E G V W T L K D E T L T A T V T E
BINARY_5676 T T Y K L I L N G K Q A K E E A I T E A V D A G I A E K Y I K L Y A N A K T V E G V W T L K D E T L T F T V T E
BINARY_5677 T T Y K L I L N G K Q A K E E A I T E A V D A G I A E K Y I K L Y A N A K T V E G V W T L K D E T K K A T V T E
BINARY_5678 T T Y K L I L N G K Q A K E E A I T E A V D A G I A E K Y I K L Y A N A K T V E G V W T L K D E T K K F T V T E
BINARY_5679 T T Y K L I L N G K Q A K E E A I T E A V D A G I A E K Y I K L Y A N A K T V E G V W T L K D E T K T A T V T E
BINARY_5680 T T Y K L I L N G K Q A K E E A I T E A V D A G I A E K Y I K L Y A N A K T V E G V W T L K D E T K T F T V T E
BINARY_5681 T T Y K L I L N G K Q A K E E A I T E A V D A G I A E K Y I K L Y A N A K T V E G V W T Y K D E I L K A T V T E
BINARY_5682 T T Y K L I L N G K Q A K E E A I T E A V D A G I A E K Y I K L Y A N A K T V E G V W T Y K D E I L K F T V T E
BINARY_5683 T T Y K L I L N G K Q A K E E A I T E A V D A G I A E K Y I K L Y A N A K T V E G V W T Y K D E I L T A T V T E
BINARY_5684 T T Y K L I L N G K Q A K E E A I T E A V D A G I A E K Y I K L Y A N A K T V E G V W T Y K D E I L T F T V T E
BINARY_5685 T T Y K L I L N G K Q A K E E A I T E A V D A G I A E K Y I K L Y A N A K T V E G V W T Y K D E I K K A T V T E
BINARY_5686 T T Y K L I L N G K Q A K E E A I T E A V D A G I A E K Y I K L Y A N A K T V E G V W T Y K D E I K K F T V T E
BINARY_5687 T T Y K L I L N G K Q A K E E A I T E A V D A G I A E K Y I K L Y A N A K T V E G V W T Y K D E I K T A T V T E
BINARY_5688 T T Y K L I L N G K Q A K E E A I T E A V D A G I A E K Y I K L Y A N A K T V E G V W T Y K D E I K T F T V T E
BINARY_5689 T T Y K L I L N G K Q A K E E A I T E A V D A G I A E K Y I K L Y A N A K T V E G V W T Y K D E T L K A T V T E
BINARY_5690 T T Y K L I L N G K Q A K E E A I T E A V D A G I A E K Y I K L Y A N A K T V E G V W T Y K D E T L K F T V T E
BINARY_5691 T T Y K L I L N G K Q A K E E A I T E A V D A G I A E K Y I K L Y A N A K T V E G V W T Y K D E T L T A T V T E
BINARY_5692 T T Y K L I L N G K Q A K E E A I T E A V D A G I A E K Y I K L Y A N A K T V E G V W T Y K D E T L T F T V T E
BINARY_5693 T T Y K L I L N G K Q A K E E A I T E A V D A G I A E K Y I K L Y A N A K T V E G V W T Y K D E T K K A T V T E
BINARY_5694 T T Y K L I L N G K Q A K E E A I T E A V D A G I A E K Y I K L Y A N A K T V E G V W T Y K D E T K K F T V T E
BINARY_5695 T T Y K L I L N G K Q A K E E A I T E A V D A G I A E K Y I K L Y A N A K T V E G V W T Y K D E T K T A T V T E
BINARY_5696 T T Y K L I L N G K Q A K E E A I T E A V D A G I A E K Y I K L Y A N A K T V E G V W T Y K D E T K T F T V T E
BINARY_5697 T T Y K L I L N G K Q A K E E A I T E A V D A G I A E K Y F K L I A N A K T V E G V W T L K D E I L K A T V T E
BINARY_5698 T T Y K L I L N G K Q A K E E A I T E A V D A G I A E K Y F K L I A N A K T V E G V W T L K D E I L K F T V T E
BINARY_5699 T T Y K L I L N G K Q A K E E A I T E A V D A G I A E K Y F K L I A N A K T V E G V W T L K D E I L T A T V T E
BINARY_5700 T T Y K L I L N G K Q A K E E A I T E A V D A G I A E K Y F K L I A N A K T V E G V W T L K D E I L T F T V T E
BINARY_5701 T T Y K L I L N G K Q A K E E A I T E A V D A G I A E K Y F K L I A N A K T V E G V W T L K D E I K K A T V T E
BINARY_5702 T T Y K L I L N G K Q A K E E A I T E A V D A G I A E K Y F K L I A N A K T V E G V W T L K D E I K K F T V T E
BINARY_5703 T T Y K L I L N G K Q A K E E A I T E A V D A G I A E K Y F K L I A N A K T V E G V W T L K D E I K T A T V T E
BINARY_5704 T T Y K L I L N G K Q A K E E A I T E A V D A G I A E K Y F K L I A N A K T V E G V W T L K D E I K T F T V T E
BINARY_5705 T T Y K L I L N G K Q A K E E A I T E A V D A G I A E K Y F K L I A N A K T V E G V W T L K D E T L K A T V T E
BINARY_5706 T T Y K L I L N G K Q A K E E A I T E A V D A G I A E K Y F K L I A N A K T V E G V W T L K D E T L K F T V T E
BINARY_5707 T T Y K L I L N G K Q A K E E A I T E A V D A G I A E K Y F K L I A N A K T V E G V W T L K D E T L T A T V T E
BINARY_5708 T T Y K L I L N G K Q A K E E A I T E A V D A G I A E K Y F K L I A N A K T V E G V W T L K D E T L T F T V T E
BINARY_5709 T T Y K L I L N G K Q A K E E A I T E A V D A G I A E K Y F K L I A N A K T V E G V W T L K D E T K K A T V T E
BINARY_5710 T T Y K L I L N G K Q A K E E A I T E A V D A G I A E K Y F K L I A N A K T V E G V W T L K D E T K K F T V T E
BINARY_5711 T T Y K L I L N G K Q A K E E A I T E A V D A G I A E K Y F K L I A N A K T V E G V W T L K D E T K T A T V T E
BINARY_5712 T T Y K L I L N G K Q A K E E A I T E A V D A G I A E K Y F K L I A N A K T V E G V W T L K D E T K T F T V T E
BINARY_5713 T T Y K L I L N G K Q A K E E A I T E A V D A G I A E K Y F K L I A N A K T V E G V W T Y K D E I L K A T V T E
BINARY_5714 T T Y K L I L N G K Q A K E E A I T E A V D A G I A E K Y F K L I A N A K T V E G V W T Y K D E I L K F T V T E
BINARY_5715 T T Y K L I L N G K Q A K E E A I T E A V D A G I A E K Y F K L I A N A K T V E G V W T Y K D E I L T A T V T E
BINARY_5716 T T Y K L I L N G K Q A K E E A I T E A V D A G I A E K Y F K L I A N A K T V E G V W T Y K D E I L T F T V T E

BINARY_5717 T T Y K L I L N G K Q A K E E A I T E A V D A G I A E K Y F K L I A N A K T V E G V W T Y K D E I K K A T V T E
BINARY_5718 T T Y K L I L N G K Q A K E E A I T E A V D A G I A E K Y F K L I A N A K T V E G V W T Y K D E I K K F T V T E
BINARY_5719 T T Y K L I L N G K Q A K E E A I T E A V D A G I A E K Y F K L I A N A K T V E G V W T Y K D E I K T A T V T E
BINARY_5720 T T Y K L I L N G K Q A K E E A I T E A V D A G I A E K Y F K L I A N A K T V E G V W T Y K D E I K T F T V T E
BINARY_5721 T T Y K L I L N G K Q A K E E A I T E A V D A G I A E K Y F K L I A N A K T V E G V W T Y K D E T L K A T V T E
BINARY_5722 T T Y K L I L N G K Q A K E E A I T E A V D A G I A E K Y F K L I A N A K T V E G V W T Y K D E T L K F T V T E
BINARY_5723 T T Y K L I L N G K Q A K E E A I T E A V D A G I A E K Y F K L I A N A K T V E G V W T Y K D E T L T A T V T E
BINARY_5724 T T Y K L I L N G K Q A K E E A I T E A V D A G I A E K Y F K L I A N A K T V E G V W T Y K D E T L T F T V T E
BINARY_5725 T T Y K L I L N G K Q A K E E A I T E A V D A G I A E K Y F K L I A N A K T V E G V W T Y K D E T K K A T V T E
BINARY_5726 T T Y K L I L N G K Q A K E E A I T E A V D A G I A E K Y F K L I A N A K T V E G V W T Y K D E T K K F T V T E
BINARY_5727 T T Y K L I L N G K Q A K E E A I T E A V D A G I A E K Y F K L I A N A K T V E G V W T Y K D E T K T A T V T E
BINARY_5728 T T Y K L I L N G K Q A K E E A I T E A V D A G I A E K Y F K L I A N A K T V E G V W T Y K D E T K T F T V T E
BINARY_5729 T T Y K L I L N G K Q A K E E A I T E A V D A G I A E K Y F K L Y A N A K T V E G V W T L K D E I L K A T V T E
BINARY_5730 T T Y K L I L N G K Q A K E E A I T E A V D A G I A E K Y F K L Y A N A K T V E G V W T L K D E I L K F T V T E
BINARY_5731 T T Y K L I L N G K Q A K E E A I T E A V D A G I A E K Y F K L Y A N A K T V E G V W T L K D E I L T A T V T E
BINARY_5732 T T Y K L I L N G K Q A K E E A I T E A V D A G I A E K Y F K L Y A N A K T V E G V W T L K D E I L T F T V T E
BINARY_5733 T T Y K L I L N G K Q A K E E A I T E A V D A G I A E K Y F K L Y A N A K T V E G V W T L K D E I K K A T V T E
BINARY_5734 T T Y K L I L N G K Q A K E E A I T E A V D A G I A E K Y F K L Y A N A K T V E G V W T L K D E I K K F T V T E
BINARY_5735 T T Y K L I L N G K Q A K E E A I T E A V D A G I A E K Y F K L Y A N A K T V E G V W T L K D E I K T A T V T E
BINARY_5736 T T Y K L I L N G K Q A K E E A I T E A V D A G I A E K Y F K L Y A N A K T V E G V W T L K D E I K T F T V T E
BINARY_5737 T T Y K L I L N G K Q A K E E A I T E A V D A G I A E K Y F K L Y A N A K T V E G V W T L K D E T L K A T V T E
BINARY_5738 T T Y K L I L N G K Q A K E E A I T E A V D A G I A E K Y F K L Y A N A K T V E G V W T L K D E T L K F T V T E
BINARY_5739 T T Y K L I L N G K Q A K E E A I T E A V D A G I A E K Y F K L Y A N A K T V E G V W T L K D E T L T A T V T E
BINARY_5740 T T Y K L I L N G K Q A K E E A I T E A V D A G I A E K Y F K L Y A N A K T V E G V W T L K D E T L T F T V T E
BINARY_5741 T T Y K L I L N G K Q A K E E A I T E A V D A G I A E K Y F K L Y A N A K T V E G V W T L K D E T K K A T V T E
BINARY_5742 T T Y K L I L N G K Q A K E E A I T E A V D A G I A E K Y F K L Y A N A K T V E G V W T L K D E T K K F T V T E
BINARY_5743 T T Y K L I L N G K Q A K E E A I T E A V D A G I A E K Y F K L Y A N A K T V E G V W T L K D E T K T A T V T E
BINARY_5744 T T Y K L I L N G K Q A K E E A I T E A V D A G I A E K Y F K L Y A N A K T V E G V W T L K D E T K T F T V T E
BINARY_5745 T T Y K L I L N G K Q A K E E A I T E A V D A G I A E K Y F K L Y A N A K T V E G V W T Y K D E I L K A T V T E
BINARY_5746 T T Y K L I L N G K Q A K E E A I T E A V D A G I A E K Y F K L Y A N A K T V E G V W T Y K D E I L K F T V T E
BINARY_5747 T T Y K L I L N G K Q A K E E A I T E A V D A G I A E K Y F K L Y A N A K T V E G V W T Y K D E I L T A T V T E
BINARY_5748 T T Y K L I L N G K Q A K E E A I T E A V D A G I A E K Y F K L Y A N A K T V E G V W T Y K D E I L T F T V T E
BINARY_5749 T T Y K L I L N G K Q A K E E A I T E A V D A G I A E K Y F K L Y A N A K T V E G V W T Y K D E I K K A T V T E
BINARY_5750 T T Y K L I L N G K Q A K E E A I T E A V D A G I A E K Y F K L Y A N A K T V E G V W T Y K D E I K K F T V T E
BINARY_5751 T T Y K L I L N G K Q A K E E A I T E A V D A G I A E K Y F K L Y A N A K T V E G V W T Y K D E I K T A T V T E
BINARY_5752 T T Y K L I L N G K Q A K E E A I T E A V D A G I A E K Y F K L Y A N A K T V E G V W T Y K D E I K T F T V T E
BINARY_5753 T T Y K L I L N G K Q A K E E A I T E A V D A G I A E K Y F K L Y A N A K T V E G V W T Y K D E T L K A T V T E
BINARY_5754 T T Y K L I L N G K Q A K E E A I T E A V D A G I A E K Y F K L Y A N A K T V E G V W T Y K D E T L K F T V T E
BINARY_5755 T T Y K L I L N G K Q A K E E A I T E A V D A G I A E K Y F K L Y A N A K T V E G V W T Y K D E T L T A T V T E
BINARY_5756 T T Y K L I L N G K Q A K E E A I T E A V D A G I A E K Y F K L Y A N A K T V E G V W T Y K D E T L T F T V T E
BINARY_5757 T T Y K L I L N G K Q A K E E A I T E A V D A G I A E K Y F K L Y A N A K T V E G V W T Y K D E T K K A T V T E
BINARY_5758 T T Y K L I L N G K Q A K E E A I T E A V D A G I A E K Y F K L Y A N A K T V E G V W T Y K D E T K K F T V T E
BINARY_5759 T T Y K L I L N G K Q A K E E A I T E A V D A G I A E K Y F K L Y A N A K T V E G V W T Y K D E T K T A T V T E
BINARY_5760 T T Y K L I L N G K Q A K E E A I T E A V D A G I A E K Y F K L Y A N A K T V E G V W T Y K D E T K T F T V T E
BINARY_5761 T T Y K L I L N G K Q A K E E A I T E A V D A G T A E K Y I K L I A N A K T V E G V W T L K D E I L K A T V T E
BINARY_5762 T T Y K L I L N G K Q A K E E A I T E A V D A G T A E K Y I K L I A N A K T V E G V W T L K D E I L K F T V T E
BINARY_5763 T T Y K L I L N G K Q A K E E A I T E A V D A G T A E K Y I K L I A N A K T V E G V W T L K D E I L T A T V T E
BINARY_5764 T T Y K L I L N G K Q A K E E A I T E A V D A G T A E K Y I K L I A N A K T V E G V W T L K D E I L T F T V T E
BINARY_5765 T T Y K L I L N G K Q A K E E A I T E A V D A G T A E K Y I K L I A N A K T V E G V W T L K D E I K K A T V T E
BINARY_5766 T T Y K L I L N G K Q A K E E A I T E A V D A G T A E K Y I K L I A N A K T V E G V W T L K D E I K K F T V T E
BINARY_5767 T T Y K L I L N G K Q A K E E A I T E A V D A G T A E K Y I K L I A N A K T V E G V W T L K D E I K T A T V T E
BINARY_5768 T T Y K L I L N G K Q A K E E A I T E A V D A G T A E K Y I K L I A N A K T V E G V W T L K D E I K T F T V T E
BINARY_5769 T T Y K L I L N G K Q A K E E A I T E A V D A G T A E K Y I K L I A N A K T V E G V W T L K D E T L K A T V T E
BINARY_5770 T T Y K L I L N G K Q A K E E A I T E A V D A G T A E K Y I K L I A N A K T V E G V W T L K D E T L K F T V T E
BINARY_5771 T T Y K L I L N G K Q A K E E A I T E A V D A G T A E K Y I K L I A N A K T V E G V W T L K D E T L T A T V T E
BINARY_5772 T T Y K L I L N G K Q A K E E A I T E A V D A G T A E K Y I K L I A N A K T V E G V W T L K D E T L T F T V T E
BINARY_5773 T T Y K L I L N G K Q A K E E A I T E A V D A G T A E K Y I K L I A N A K T V E G V W T L K D E T K K A T V T E
BINARY_5774 T T Y K L I L N G K Q A K E E A I T E A V D A G T A E K Y I K L I A N A K T V E G V W T L K D E T K K F T V T E
BINARY_5775 T T Y K L I L N G K Q A K E E A I T E A V D A G T A E K Y I K L I A N A K T V E G V W T L K D E T K T A T V T E
BINARY_5776 T T Y K L I L N G K Q A K E E A I T E A V D A G T A E K Y I K L I A N A K T V E G V W T L K D E T K T F T V T E
BINARY_5777 T T Y K L I L N G K Q A K E E A I T E A V D A G T A E K Y I K L I A N A K T V E G V W T Y K D E I L K A T V T E
BINARY_5778 T T Y K L I L N G K Q A K E E A I T E A V D A G T A E K Y I K L I A N A K T V E G V W T Y K D E I L K F T V T E
BINARY_5779 T T Y K L I L N G K Q A K E E A I T E A V D A G T A E K Y I K L I A N A K T V E G V W T Y K D E I L T A T V T E
BINARY_5780 T T Y K L I L N G K Q A K E E A I T E A V D A G T A E K Y I K L I A N A K T V E G V W T Y K D E I L T F T V T E
BINARY_5781 T T Y K L I L N G K Q A K E E A I T E A V D A G T A E K Y I K L I A N A K T V E G V W T Y K D E I K K A T V T E
BINARY_5782 T T Y K L I L N G K Q A K E E A I T E A V D A G T A E K Y I K L I A N A K T V E G V W T Y K D E I K K F T V T E
BINARY_5783 T T Y K L I L N G K Q A K E E A I T E A V D A G T A E K Y I K L I A N A K T V E G V W T Y K D E I K T A T V T E
BINARY_5784 T T Y K L I L N G K Q A K E E A I T E A V D A G T A E K Y I K L I A N A K T V E G V W T Y K D E I K T F T V T E
BINARY_5785 T T Y K L I L N G K Q A K E E A I T E A V D A G T A E K Y I K L I A N A K T V E G V W T Y K D E T L K A T V T E
BINARY_5786 T T Y K L I L N G K Q A K E E A I T E A V D A G T A E K Y I K L I A N A K T V E G V W T Y K D E T L K F T V T E
BINARY_5787 T T Y K L I L N G K Q A K E E A I T E A V D A G T A E K Y I K L I A N A K T V E G V W T Y K D E T L T A T V T E
BINARY_5788 T T Y K L I L N G K Q A K E E A I T E A V D A G T A E K Y I K L I A N A K T V E G V W T Y K D E T L T F T V T E
BINARY_5789 T T Y K L I L N G K Q A K E E A I T E A V D A G T A E K Y I K L I A N A K T V E G V W T Y K D E T K K A T V T E
BINARY_5790 T T Y K L I L N G K Q A K E E A I T E A V D A G T A E K Y I K L I A N A K T V E G V W T Y K D E T K K F T V T E
BINARY_5791 T T Y K L I L N G K Q A K E E A I T E A V D A G T A E K Y I K L I A N A K T V E G V W T Y K D E T K T A T V T E
BINARY_5792 T T Y K L I L N G K Q A K E E A I T E A V D A G T A E K Y I K L I A N A K T V E G V W T Y K D E T K T F T V T E
BINARY_5793 T T Y K L I L N G K Q A K E E A I T E A V D A G T A E K Y I K L Y A N A K T V E G V W T L K D E I L K A T V T E
BINARY_5794 T T Y K L I L N G K Q A K E E A I T E A V D A G T A E K Y I K L Y A N A K T V E G V W T L K D E I L K F T V T E
BINARY_5795 T T Y K L I L N G K Q A K E E A I T E A V D A G T A E K Y I K L Y A N A K T V E G V W T L K D E I L T A T V T E
BINARY_5796 T T Y K L I L N G K Q A K E E A I T E A V D A G T A E K Y I K L Y A N A K T V E G V W T L K D E I L T F T V T E
BINARY_5797 T T Y K L I L N G K Q A K E E A I T E A V D A G T A E K Y I K L Y A N A K T V E G V W T L K D E I K K A T V T E
BINARY_5798 T T Y K L I L N G K Q A K E E A I T E A V D A G T A E K Y I K L Y A N A K T V E G V W T L K D E I K K F T V T E
BINARY_5799 T T Y K L I L N G K Q A K E E A I T E A V D A G T A E K Y I K L Y A N A K T V E G V W T L K D E I K T A T V T E
BINARY_5800 T T Y K L I L N G K Q A K E E A I T E A V D A G T A E K Y I K L Y A N A K T V E G V W T L K D E I K T F T V T E
BINARY_5801 T T Y K L I L N G K Q A K E E A I T E A V D A G T A E K Y I K L Y A N A K T V E G V W T L K D E T L K A T V T E
BINARY_5802 T T Y K L I L N G K Q A K E E A I T E A V D A G T A E K Y I K L Y A N A K T V E G V W T L K D E T L K F T V T E
BINARY_5803 T T Y K L I L N G K Q A K E E A I T E A V D A G T A E K Y I K L Y A N A K T V E G V W T L K D E T L T A T V T E
BINARY_5804 T T Y K L I L N G K Q A K E E A I T E A V D A G T A E K Y I K L Y A N A K T V E G V W T L K D E T L T F T V T E
BINARY_5805 T T Y K L I L N G K Q A K E E A I T E A V D A G T A E K Y I K L Y A N A K T V E G V W T L K D E T K K A T V T E
BINARY_5806 T T Y K L I L N G K Q A K E E A I T E A V D A G T A E K Y I K L Y A N A K T V E G V W T L K D E T K K F T V T E
BINARY_5807 T T Y K L I L N G K Q A K E E A I T E A V D A G T A E K Y I K L Y A N A K T V E G V W T L K D E T K T A T V T E
BINARY_5808 T T Y K L I L N G K Q A K E E A I T E A V D A G T A E K Y I K L Y A N A K T V E G V W T L K D E T K T F T V T E
BINARY_5809 T T Y K L I L N G K Q A K E E A I T E A V D A G T A E K Y I K L Y A N A K T V E G V W T Y K D E I L K A T V T E
BINARY_5810 T T Y K L I L N G K Q A K E E A I T E A V D A G T A E K Y I K L Y A N A K T V E G V W T Y K D E I L K F T V T E
BINARY_5811 T T Y K L I L N G K Q A K E E A I T E A V D A G T A E K Y I K L Y A N A K T V E G V W T Y K D E I L T A T V T E

BINARY_5812 T T Y K L I L N G K Q A K E E A I T E A V D A G T A E K Y I K L Y A N A K T V E G V W T Y K D E I L T F T V T E
BINARY_5813 T T Y K L I L N G K Q A K E E A I T E A V D A G T A E K Y I K L Y A N A K T V E G V W T Y K D E I K K A T V T E
BINARY_5814 T T Y K L I L N G K Q A K E E A I T E A V D A G T A E K Y I K L Y A N A K T V E G V W T Y K D E I K K F T V T E
BINARY_5815 T T Y K L I L N G K Q A K E E A I T E A V D A G T A E K Y I K L Y A N A K T V E G V W T Y K D E I K T A T V T E
BINARY_5816 T T Y K L I L N G K Q A K E E A I T E A V D A G T A E K Y I K L Y A N A K T V E G V W T Y K D E I K T F T V T E
BINARY_5817 T T Y K L I L N G K Q A K E E A I T E A V D A G T A E K Y I K L Y A N A K T V E G V W T Y K D E T L K A T V T E
BINARY_5818 T T Y K L I L N G K Q A K E E A I T E A V D A G T A E K Y I K L Y A N A K T V E G V W T Y K D E T L K F T V T E
BINARY_5819 T T Y K L I L N G K Q A K E E A I T E A V D A G T A E K Y I K L Y A N A K T V E G V W T Y K D E T L T A T V T E
BINARY_5820 T T Y K L I L N G K Q A K E E A I T E A V D A G T A E K Y I K L Y A N A K T V E G V W T Y K D E T L T F T V T E
BINARY_5821 T T Y K L I L N G K Q A K E E A I T E A V D A G T A E K Y I K L Y A N A K T V E G V W T Y K D E T K K A T V T E
BINARY_5822 T T Y K L I L N G K Q A K E E A I T E A V D A G T A E K Y I K L Y A N A K T V E G V W T Y K D E T K K F T V T E
BINARY_5823 T T Y K L I L N G K Q A K E E A I T E A V D A G T A E K Y I K L Y A N A K T V E G V W T Y K D E T K T A T V T E
BINARY_5824 T T Y K L I L N G K Q A K E E A I T E A V D A G T A E K Y I K L Y A N A K T V E G V W T Y K D E T K T F T V T E
BINARY_5825 T T Y K L I L N G K Q A K E E A I T E A V D A G T A E K Y F K L I A N A K T V E G V W T L K D E I L K A T V T E
BINARY_5826 T T Y K L I L N G K Q A K E E A I T E A V D A G T A E K Y F K L I A N A K T V E G V W T L K D E I L K F T V T E
BINARY_5827 T T Y K L I L N G K Q A K E E A I T E A V D A G T A E K Y F K L I A N A K T V E G V W T L K D E I L T A T V T E
BINARY_5828 T T Y K L I L N G K Q A K E E A I T E A V D A G T A E K Y F K L I A N A K T V E G V W T L K D E I L T F T V T E
BINARY_5829 T T Y K L I L N G K Q A K E E A I T E A V D A G T A E K Y F K L I A N A K T V E G V W T L K D E I K K A T V T E
BINARY_5830 T T Y K L I L N G K Q A K E E A I T E A V D A G T A E K Y F K L I A N A K T V E G V W T L K D E I K K F T V T E
BINARY_5831 T T Y K L I L N G K Q A K E E A I T E A V D A G T A E K Y F K L I A N A K T V E G V W T L K D E I K T A T V T E
BINARY_5832 T T Y K L I L N G K Q A K E E A I T E A V D A G T A E K Y F K L I A N A K T V E G V W T L K D E I K T F T V T E
BINARY_5833 T T Y K L I L N G K Q A K E E A I T E A V D A G T A E K Y F K L I A N A K T V E G V W T L K D E T L K A T V T E
BINARY_5834 T T Y K L I L N G K Q A K E E A I T E A V D A G T A E K Y F K L I A N A K T V E G V W T L K D E T L K F T V T E
BINARY_5835 T T Y K L I L N G K Q A K E E A I T E A V D A G T A E K Y F K L I A N A K T V E G V W T L K D E T L T A T V T E
BINARY_5836 T T Y K L I L N G K Q A K E E A I T E A V D A G T A E K Y F K L I A N A K T V E G V W T L K D E T L T F T V T E
BINARY_5837 T T Y K L I L N G K Q A K E E A I T E A V D A G T A E K Y F K L I A N A K T V E G V W T L K D E T K K A T V T E
BINARY_5838 T T Y K L I L N G K Q A K E E A I T E A V D A G T A E K Y F K L I A N A K T V E G V W T L K D E T K K F T V T E
BINARY_5839 T T Y K L I L N G K Q A K E E A I T E A V D A G T A E K Y F K L I A N A K T V E G V W T L K D E T K T A T V T E
BINARY_5840 T T Y K L I L N G K Q A K E E A I T E A V D A G T A E K Y F K L I A N A K T V E G V W T L K D E T K T F T V T E
BINARY_5841 T T Y K L I L N G K Q A K E E A I T E A V D A G T A E K Y F K L I A N A K T V E G V W T Y K D E I L K A T V T E
BINARY_5842 T T Y K L I L N G K Q A K E E A I T E A V D A G T A E K Y F K L I A N A K T V E G V W T Y K D E I L K F T V T E
BINARY_5843 T T Y K L I L N G K Q A K E E A I T E A V D A G T A E K Y F K L I A N A K T V E G V W T Y K D E I L T A T V T E
BINARY_5844 T T Y K L I L N G K Q A K E E A I T E A V D A G T A E K Y F K L I A N A K T V E G V W T Y K D E I L T F T V T E
BINARY_5845 T T Y K L I L N G K Q A K E E A I T E A V D A G T A E K Y F K L I A N A K T V E G V W T Y K D E I K K A T V T E
BINARY_5846 T T Y K L I L N G K Q A K E E A I T E A V D A G T A E K Y F K L I A N A K T V E G V W T Y K D E I K K F T V T E
BINARY_5847 T T Y K L I L N G K Q A K E E A I T E A V D A G T A E K Y F K L I A N A K T V E G V W T Y K D E I K T A T V T E
BINARY_5848 T T Y K L I L N G K Q A K E E A I T E A V D A G T A E K Y F K L I A N A K T V E G V W T Y K D E I K T F T V T E
BINARY_5849 T T Y K L I L N G K Q A K E E A I T E A V D A G T A E K Y F K L I A N A K T V E G V W T Y K D E T L K A T V T E
BINARY_5850 T T Y K L I L N G K Q A K E E A I T E A V D A G T A E K Y F K L I A N A K T V E G V W T Y K D E T L K F T V T E
BINARY_5851 T T Y K L I L N G K Q A K E E A I T E A V D A G T A E K Y F K L I A N A K T V E G V W T Y K D E T L T A T V T E
BINARY_5852 T T Y K L I L N G K Q A K E E A I T E A V D A G T A E K Y F K L I A N A K T V E G V W T Y K D E T L T F T V T E
BINARY_5853 T T Y K L I L N G K Q A K E E A I T E A V D A G T A E K Y F K L I A N A K T V E G V W T Y K D E T K K A T V T E
BINARY_5854 T T Y K L I L N G K Q A K E E A I T E A V D A G T A E K Y F K L I A N A K T V E G V W T Y K D E T K K F T V T E
BINARY_5855 T T Y K L I L N G K Q A K E E A I T E A V D A G T A E K Y F K L I A N A K T V E G V W T Y K D E T K T A T V T E
BINARY_5856 T T Y K L I L N G K Q A K E E A I T E A V D A G T A E K Y F K L I A N A K T V E G V W T Y K D E T K T F T V T E
BINARY_5857 T T Y K L I L N G K Q A K E E A I T E A V D A G T A E K Y F K L Y A N A K T V E G V W T L K D E I L K A T V T E
BINARY_5858 T T Y K L I L N G K Q A K E E A I T E A V D A G T A E K Y F K L Y A N A K T V E G V W T L K D E I L K F T V T E
BINARY_5859 T T Y K L I L N G K Q A K E E A I T E A V D A G T A E K Y F K L Y A N A K T V E G V W T L K D E I L T A T V T E
BINARY_5860 T T Y K L I L N G K Q A K E E A I T E A V D A G T A E K Y F K L Y A N A K T V E G V W T L K D E I L T F T V T E
BINARY_5861 T T Y K L I L N G K Q A K E E A I T E A V D A G T A E K Y F K L Y A N A K T V E G V W T L K D E I K K A T V T E
BINARY_5862 T T Y K L I L N G K Q A K E E A I T E A V D A G T A E K Y F K L Y A N A K T V E G V W T L K D E I K K F T V T E
BINARY_5863 T T Y K L I L N G K Q A K E E A I T E A V D A G T A E K Y F K L Y A N A K T V E G V W T L K D E I K T A T V T E
BINARY_5864 T T Y K L I L N G K Q A K E E A I T E A V D A G T A E K Y F K L Y A N A K T V E G V W T L K D E I K T F T V T E
BINARY_5865 T T Y K L I L N G K Q A K E E A I T E A V D A G T A E K Y F K L Y A N A K T V E G V W T L K D E T L K A T V T E
BINARY_5866 T T Y K L I L N G K Q A K E E A I T E A V D A G T A E K Y F K L Y A N A K T V E G V W T L K D E T L K F T V T E
BINARY_5867 T T Y K L I L N G K Q A K E E A I T E A V D A G T A E K Y F K L Y A N A K T V E G V W T L K D E T L T A T V T E
BINARY_5868 T T Y K L I L N G K Q A K E E A I T E A V D A G T A E K Y F K L Y A N A K T V E G V W T L K D E T L T F T V T E
BINARY_5869 T T Y K L I L N G K Q A K E E A I T E A V D A G T A E K Y F K L Y A N A K T V E G V W T L K D E T K K A T V T E
BINARY_5870 T T Y K L I L N G K Q A K E E A I T E A V D A G T A E K Y F K L Y A N A K T V E G V W T L K D E T K K F T V T E
BINARY_5871 T T Y K L I L N G K Q A K E E A I T E A V D A G T A E K Y F K L Y A N A K T V E G V W T L K D E T K T A T V T E
BINARY_5872 T T Y K L I L N G K Q A K E E A I T E A V D A G T A E K Y F K L Y A N A K T V E G V W T L K D E T K T F T V T E
BINARY_5873 T T Y K L I L N G K Q A K E E A I T E A V D A G T A E K Y F K L Y A N A K T V E G V W T Y K D E I L K A T V T E
BINARY_5874 T T Y K L I L N G K Q A K E E A I T E A V D A G T A E K Y F K L Y A N A K T V E G V W T Y K D E I L K F T V T E
BINARY_5875 T T Y K L I L N G K Q A K E E A I T E A V D A G T A E K Y F K L Y A N A K T V E G V W T Y K D E I L T A T V T E
BINARY_5876 T T Y K L I L N G K Q A K E E A I T E A V D A G T A E K Y F K L Y A N A K T V E G V W T Y K D E I L T F T V T E
BINARY_5877 T T Y K L I L N G K Q A K E E A I T E A V D A G T A E K Y F K L Y A N A K T V E G V W T Y K D E I K K A T V T E
BINARY_5878 T T Y K L I L N G K Q A K E E A I T E A V D A G T A E K Y F K L Y A N A K T V E G V W T Y K D E I K K F T V T E
BINARY_5879 T T Y K L I L N G K Q A K E E A I T E A V D A G T A E K Y F K L Y A N A K T V E G V W T Y K D E I K T A T V T E
BINARY_5880 T T Y K L I L N G K Q A K E E A I T E A V D A G T A E K Y F K L Y A N A K T V E G V W T Y K D E I K T F T V T E
BINARY_5881 T T Y K L I L N G K Q A K E E A I T E A V D A G T A E K Y F K L Y A N A K T V E G V W T Y K D E T L K A T V T E
BINARY_5882 T T Y K L I L N G K Q A K E E A I T E A V D A G T A E K Y F K L Y A N A K T V E G V W T Y K D E T L K F T V T E
BINARY_5883 T T Y K L I L N G K Q A K E E A I T E A V D A G T A E K Y F K L Y A N A K T V E G V W T Y K D E T L T A T V T E
BINARY_5884 T T Y K L I L N G K Q A K E E A I T E A V D A G T A E K Y F K L Y A N A K T V E G V W T Y K D E T L T F T V T E
BINARY_5885 T T Y K L I L N G K Q A K E E A I T E A V D A G T A E K Y F K L Y A N A K T V E G V W T Y K D E T K K A T V T E
BINARY_5886 T T Y K L I L N G K Q A K E E A I T E A V D A G T A E K Y F K L Y A N A K T V E G V W T Y K D E T K K F T V T E
BINARY_5887 T T Y K L I L N G K Q A K E E A I T E A V D A G T A E K Y F K L Y A N A K T V E G V W T Y K D E T K T A T V T E
BINARY_5888 T T Y K L I L N G K Q A K E E A I T E A V D A G T A E K Y F K L Y A N A K T V E G V W T Y K D E T K T F T V T E
BINARY_5889 T T Y K L I L N G K Q A K E E A I T E A V D A A I A E K Y I K L I A N A K T V E G V W T L K D E I L K A T V T E
BINARY_5890 T T Y K L I L N G K Q A K E E A I T E A V D A A I A E K Y I K L I A N A K T V E G V W T L K D E I L K F T V T E
BINARY_5891 T T Y K L I L N G K Q A K E E A I T E A V D A A I A E K Y I K L I A N A K T V E G V W T L K D E I L T A T V T E
BINARY_5892 T T Y K L I L N G K Q A K E E A I T E A V D A A I A E K Y I K L I A N A K T V E G V W T L K D E I L T F T V T E
BINARY_5893 T T Y K L I L N G K Q A K E E A I T E A V D A A I A E K Y I K L I A N A K T V E G V W T L K D E I K K A T V T E
BINARY_5894 T T Y K L I L N G K Q A K E E A I T E A V D A A I A E K Y I K L I A N A K T V E G V W T L K D E I K K F T V T E
BINARY_5895 T T Y K L I L N G K Q A K E E A I T E A V D A A I A E K Y I K L I A N A K T V E G V W T L K D E I K T A T V T E
BINARY_5896 T T Y K L I L N G K Q A K E E A I T E A V D A A I A E K Y I K L I A N A K T V E G V W T L K D E I K T F T V T E
BINARY_5897 T T Y K L I L N G K Q A K E E A I T E A V D A A I A E K Y I K L I A N A K T V E G V W T L K D E T L K A T V T E
BINARY_5898 T T Y K L I L N G K Q A K E E A I T E A V D A A I A E K Y I K L I A N A K T V E G V W T L K D E T L K F T V T E
BINARY_5899 T T Y K L I L N G K Q A K E E A I T E A V D A A I A E K Y I K L I A N A K T V E G V W T L K D E T L T A T V T E
BINARY_5900 T T Y K L I L N G K Q A K E E A I T E A V D A A I A E K Y I K L I A N A K T V E G V W T L K D E T L T F T V T E
BINARY_5901 T T Y K L I L N G K Q A K E E A I T E A V D A A I A E K Y I K L I A N A K T V E G V W T L K D E T K K A T V T E
BINARY_5902 T T Y K L I L N G K Q A K E E A I T E A V D A A I A E K Y I K L I A N A K T V E G V W T L K D E T K K F T V T E
BINARY_5903 T T Y K L I L N G K Q A K E E A I T E A V D A A I A E K Y I K L I A N A K T V E G V W T L K D E T K T A T V T E
BINARY_5904 T T Y K L I L N G K Q A K E E A I T E A V D A A I A E K Y I K L I A N A K T V E G V W T L K D E T K T F T V T E
BINARY_5905 T T Y K L I L N G K Q A K E E A I T E A V D A A I A E K Y I K L I A N A K T V E G V W T Y K D E I L K A T V T E
BINARY_5906 T T Y K L I L N G K Q A K E E A I T E A V D A A I A E K Y I K L I A N A K T V E G V W T Y K D E I L K F T V T E

BINARY_5907 T T Y K L I L N G K Q A K E E A I T E A V D A A I A E K Y I K L I A N A K T V E G V W T Y K D E I L T A T V T E
BINARY_5908 T T Y K L I L N G K Q A K E E A I T E A V D A A I A E K Y I K L I A N A K T V E G V W T Y K D E I L T F T V T E
BINARY_5909 T T Y K L I L N G K Q A K E E A I T E A V D A A I A E K Y I K L I A N A K T V E G V W T Y K D E I K K A T V T E
BINARY_5910 T T Y K L I L N G K Q A K E E A I T E A V D A A I A E K Y I K L I A N A K T V E G V W T Y K D E I K K F T V T E
BINARY_5911 T T Y K L I L N G K Q A K E E A I T E A V D A A I A E K Y I K L I A N A K T V E G V W T Y K D E I K T A T V T E
BINARY_5912 T T Y K L I L N G K Q A K E E A I T E A V D A A I A E K Y I K L I A N A K T V E G V W T Y K D E I K T F T V T E
BINARY_5913 T T Y K L I L N G K Q A K E E A I T E A V D A A I A E K Y I K L I A N A K T V E G V W T Y K D E T L K A T V T E
BINARY_5914 T T Y K L I L N G K Q A K E E A I T E A V D A A I A E K Y I K L I A N A K T V E G V W T Y K D E T L K F T V T E
BINARY_5915 T T Y K L I L N G K Q A K E E A I T E A V D A A I A E K Y I K L I A N A K T V E G V W T Y K D E T L T A T V T E
BINARY_5916 T T Y K L I L N G K Q A K E E A I T E A V D A A I A E K Y I K L I A N A K T V E G V W T Y K D E T L T F T V T E
BINARY_5917 T T Y K L I L N G K Q A K E E A I T E A V D A A I A E K Y I K L I A N A K T V E G V W T Y K D E T K K A T V T E
BINARY_5918 T T Y K L I L N G K Q A K E E A I T E A V D A A I A E K Y I K L I A N A K T V E G V W T Y K D E T K K F T V T E
BINARY_5919 T T Y K L I L N G K Q A K E E A I T E A V D A A I A E K Y I K L I A N A K T V E G V W T Y K D E T K T A T V T E
BINARY_5920 T T Y K L I L N G K Q A K E E A I T E A V D A A I A E K Y I K L I A N A K T V E G V W T Y K D E T K T F T V T E
BINARY_5921 T T Y K L I L N G K Q A K E E A I T E A V D A A I A E K Y I K L Y A N A K T V E G V W T L K D E I L K A T V T E
BINARY_5922 T T Y K L I L N G K Q A K E E A I T E A V D A A I A E K Y I K L Y A N A K T V E G V W T L K D E I L K F T V T E
BINARY_5923 T T Y K L I L N G K Q A K E E A I T E A V D A A I A E K Y I K L Y A N A K T V E G V W T L K D E I L T A T V T E
BINARY_5924 T T Y K L I L N G K Q A K E E A I T E A V D A A I A E K Y I K L Y A N A K T V E G V W T L K D E I L T F T V T E
BINARY_5925 T T Y K L I L N G K Q A K E E A I T E A V D A A I A E K Y I K L Y A N A K T V E G V W T L K D E I K K A T V T E
BINARY_5926 T T Y K L I L N G K Q A K E E A I T E A V D A A I A E K Y I K L Y A N A K T V E G V W T L K D E I K K F T V T E
BINARY_5927 T T Y K L I L N G K Q A K E E A I T E A V D A A I A E K Y I K L Y A N A K T V E G V W T L K D E I K T A T V T E
BINARY_5928 T T Y K L I L N G K Q A K E E A I T E A V D A A I A E K Y I K L Y A N A K T V E G V W T L K D E I K T F T V T E
BINARY_5929 T T Y K L I L N G K Q A K E E A I T E A V D A A I A E K Y I K L Y A N A K T V E G V W T L K D E T L K A T V T E
BINARY_5930 T T Y K L I L N G K Q A K E E A I T E A V D A A I A E K Y I K L Y A N A K T V E G V W T L K D E T L K F T V T E
BINARY_5931 T T Y K L I L N G K Q A K E E A I T E A V D A A I A E K Y I K L Y A N A K T V E G V W T L K D E T L T A T V T E
BINARY_5932 T T Y K L I L N G K Q A K E E A I T E A V D A A I A E K Y I K L Y A N A K T V E G V W T L K D E T L T F T V T E
BINARY_5933 T T Y K L I L N G K Q A K E E A I T E A V D A A I A E K Y I K L Y A N A K T V E G V W T L K D E T K K A T V T E
BINARY_5934 T T Y K L I L N G K Q A K E E A I T E A V D A A I A E K Y I K L Y A N A K T V E G V W T L K D E T K K F T V T E
BINARY_5935 T T Y K L I L N G K Q A K E E A I T E A V D A A I A E K Y I K L Y A N A K T V E G V W T L K D E T K T A T V T E
BINARY_5936 T T Y K L I L N G K Q A K E E A I T E A V D A A I A E K Y I K L Y A N A K T V E G V W T L K D E T K T F T V T E
BINARY_5937 T T Y K L I L N G K Q A K E E A I T E A V D A A I A E K Y I K L Y A N A K T V E G V W T Y K D E I L K A T V T E
BINARY_5938 T T Y K L I L N G K Q A K E E A I T E A V D A A I A E K Y I K L Y A N A K T V E G V W T Y K D E I L K F T V T E
BINARY_5939 T T Y K L I L N G K Q A K E E A I T E A V D A A I A E K Y I K L Y A N A K T V E G V W T Y K D E I L T A T V T E
BINARY_5940 T T Y K L I L N G K Q A K E E A I T E A V D A A I A E K Y I K L Y A N A K T V E G V W T Y K D E I L T F T V T E
BINARY_5941 T T Y K L I L N G K Q A K E E A I T E A V D A A I A E K Y I K L Y A N A K T V E G V W T Y K D E I K K A T V T E
BINARY_5942 T T Y K L I L N G K Q A K E E A I T E A V D A A I A E K Y I K L Y A N A K T V E G V W T Y K D E I K K F T V T E
BINARY_5943 T T Y K L I L N G K Q A K E E A I T E A V D A A I A E K Y I K L Y A N A K T V E G V W T Y K D E I K T A T V T E
BINARY_5944 T T Y K L I L N G K Q A K E E A I T E A V D A A I A E K Y I K L Y A N A K T V E G V W T Y K D E I K T F T V T E
BINARY_5945 T T Y K L I L N G K Q A K E E A I T E A V D A A I A E K Y I K L Y A N A K T V E G V W T Y K D E T L K A T V T E
BINARY_5946 T T Y K L I L N G K Q A K E E A I T E A V D A A I A E K Y I K L Y A N A K T V E G V W T Y K D E T L K F T V T E
BINARY_5947 T T Y K L I L N G K Q A K E E A I T E A V D A A I A E K Y I K L Y A N A K T V E G V W T Y K D E T L T A T V T E
BINARY_5948 T T Y K L I L N G K Q A K E E A I T E A V D A A I A E K Y I K L Y A N A K T V E G V W T Y K D E T L T F T V T E
BINARY_5949 T T Y K L I L N G K Q A K E E A I T E A V D A A I A E K Y I K L Y A N A K T V E G V W T Y K D E T K K A T V T E
BINARY_5950 T T Y K L I L N G K Q A K E E A I T E A V D A A I A E K Y I K L Y A N A K T V E G V W T Y K D E T K K F T V T E
BINARY_5951 T T Y K L I L N G K Q A K E E A I T E A V D A A I A E K Y I K L Y A N A K T V E G V W T Y K D E T K T A T V T E
BINARY_5952 T T Y K L I L N G K Q A K E E A I T E A V D A A I A E K Y I K L Y A N A K T V E G V W T Y K D E T K T F T V T E
BINARY_5953 T T Y K L I L N G K Q A K E E A I T E A V D A A I A E K Y F K L I A N A K T V E G V W T L K D E I L K A T V T E
BINARY_5954 T T Y K L I L N G K Q A K E E A I T E A V D A A I A E K Y F K L I A N A K T V E G V W T L K D E I L K F T V T E
BINARY_5955 T T Y K L I L N G K Q A K E E A I T E A V D A A I A E K Y F K L I A N A K T V E G V W T L K D E I L T A T V T E
BINARY_5956 T T Y K L I L N G K Q A K E E A I T E A V D A A I A E K Y F K L I A N A K T V E G V W T L K D E I L T F T V T E
BINARY_5957 T T Y K L I L N G K Q A K E E A I T E A V D A A I A E K Y F K L I A N A K T V E G V W T L K D E I K K A T V T E
BINARY_5958 T T Y K L I L N G K Q A K E E A I T E A V D A A I A E K Y F K L I A N A K T V E G V W T L K D E I K K F T V T E
BINARY_5959 T T Y K L I L N G K Q A K E E A I T E A V D A A I A E K Y F K L I A N A K T V E G V W T L K D E I K T A T V T E
BINARY_5960 T T Y K L I L N G K Q A K E E A I T E A V D A A I A E K Y F K L I A N A K T V E G V W T L K D E I K T F T V T E
BINARY_5961 T T Y K L I L N G K Q A K E E A I T E A V D A A I A E K Y F K L I A N A K T V E G V W T L K D E T L K A T V T E
BINARY_5962 T T Y K L I L N G K Q A K E E A I T E A V D A A I A E K Y F K L I A N A K T V E G V W T L K D E T L K F T V T E
BINARY_5963 T T Y K L I L N G K Q A K E E A I T E A V D A A I A E K Y F K L I A N A K T V E G V W T L K D E T L T A T V T E
BINARY_5964 T T Y K L I L N G K Q A K E E A I T E A V D A A I A E K Y F K L I A N A K T V E G V W T L K D E T L T F T V T E
BINARY_5965 T T Y K L I L N G K Q A K E E A I T E A V D A A I A E K Y F K L I A N A K T V E G V W T L K D E T K K A T V T E
BINARY_5966 T T Y K L I L N G K Q A K E E A I T E A V D A A I A E K Y F K L I A N A K T V E G V W T L K D E T K K F T V T E
BINARY_5967 T T Y K L I L N G K Q A K E E A I T E A V D A A I A E K Y F K L I A N A K T V E G V W T L K D E T K T A T V T E
BINARY_5968 T T Y K L I L N G K Q A K E E A I T E A V D A A I A E K Y F K L I A N A K T V E G V W T L K D E T K T F T V T E
BINARY_5969 T T Y K L I L N G K Q A K E E A I T E A V D A A I A E K Y F K L I A N A K T V E G V W T Y K D E I L K A T V T E
BINARY_5970 T T Y K L I L N G K Q A K E E A I T E A V D A A I A E K Y F K L I A N A K T V E G V W T Y K D E I L K F T V T E
BINARY_5971 T T Y K L I L N G K Q A K E E A I T E A V D A A I A E K Y F K L I A N A K T V E G V W T Y K D E I L T A T V T E
BINARY_5972 T T Y K L I L N G K Q A K E E A I T E A V D A A I A E K Y F K L I A N A K T V E G V W T Y K D E I L T F T V T E
BINARY_5973 T T Y K L I L N G K Q A K E E A I T E A V D A A I A E K Y F K L I A N A K T V E G V W T Y K D E I K K A T V T E
BINARY_5974 T T Y K L I L N G K Q A K E E A I T E A V D A A I A E K Y F K L I A N A K T V E G V W T Y K D E I K K F T V T E
BINARY_5975 T T Y K L I L N G K Q A K E E A I T E A V D A A I A E K Y F K L I A N A K T V E G V W T Y K D E I K T A T V T E
BINARY_5976 T T Y K L I L N G K Q A K E E A I T E A V D A A I A E K Y F K L I A N A K T V E G V W T Y K D E I K T F T V T E
BINARY_5977 T T Y K L I L N G K Q A K E E A I T E A V D A A I A E K Y F K L I A N A K T V E G V W T Y K D E T L K A T V T E
BINARY_5978 T T Y K L I L N G K Q A K E E A I T E A V D A A I A E K Y F K L I A N A K T V E G V W T Y K D E T L K F T V T E
BINARY_5979 T T Y K L I L N G K Q A K E E A I T E A V D A A I A E K Y F K L I A N A K T V E G V W T Y K D E T L T A T V T E
BINARY_5980 T T Y K L I L N G K Q A K E E A I T E A V D A A I A E K Y F K L I A N A K T V E G V W T Y K D E T L T F T V T E
BINARY_5981 T T Y K L I L N G K Q A K E E A I T E A V D A A I A E K Y F K L I A N A K T V E G V W T Y K D E T K K A T V T E
BINARY_5982 T T Y K L I L N G K Q A K E E A I T E A V D A A I A E K Y F K L I A N A K T V E G V W T Y K D E T K K F T V T E
BINARY_5983 T T Y K L I L N G K Q A K E E A I T E A V D A A I A E K Y F K L I A N A K T V E G V W T Y K D E T K T A T V T E
BINARY_5984 T T Y K L I L N G K Q A K E E A I T E A V D A A I A E K Y F K L I A N A K T V E G V W T Y K D E T K T F T V T E
BINARY_5985 T T Y K L I L N G K Q A K E E A I T E A V D A A I A E K Y F K L Y A N A K T V E G V W T L K D E I L K A T V T E
BINARY_5986 T T Y K L I L N G K Q A K E E A I T E A V D A A I A E K Y F K L Y A N A K T V E G V W T L K D E I L K F T V T E
BINARY_5987 T T Y K L I L N G K Q A K E E A I T E A V D A A I A E K Y F K L Y A N A K T V E G V W T L K D E I L T A T V T E
BINARY_5988 T T Y K L I L N G K Q A K E E A I T E A V D A A I A E K Y F K L Y A N A K T V E G V W T L K D E I L T F T V T E
BINARY_5989 T T Y K L I L N G K Q A K E E A I T E A V D A A I A E K Y F K L Y A N A K T V E G V W T L K D E I K K A T V T E
BINARY_5990 T T Y K L I L N G K Q A K E E A I T E A V D A A I A E K Y F K L Y A N A K T V E G V W T L K D E I K K F T V T E
BINARY_5991 T T Y K L I L N G K Q A K E E A I T E A V D A A I A E K Y F K L Y A N A K T V E G V W T L K D E I K T A T V T E
BINARY_5992 T T Y K L I L N G K Q A K E E A I T E A V D A A I A E K Y F K L Y A N A K T V E G V W T L K D E I K T F T V T E
BINARY_5993 T T Y K L I L N G K Q A K E E A I T E A V D A A I A E K Y F K L Y A N A K T V E G V W T L K D E T L K A T V T E
BINARY_5994 T T Y K L I L N G K Q A K E E A I T E A V D A A I A E K Y F K L Y A N A K T V E G V W T L K D E T L K F T V T E
BINARY_5995 T T Y K L I L N G K Q A K E E A I T E A V D A A I A E K Y F K L Y A N A K T V E G V W T L K D E T L T A T V T E
BINARY_5996 T T Y K L I L N G K Q A K E E A I T E A V D A A I A E K Y F K L Y A N A K T V E G V W T L K D E T L T F T V T E
BINARY_5997 T T Y K L I L N G K Q A K E E A I T E A V D A A I A E K Y F K L Y A N A K T V E G V W T L K D E T K K A T V T E
BINARY_5998 T T Y K L I L N G K Q A K E E A I T E A V D A A I A E K Y F K L Y A N A K T V E G V W T L K D E T K K F T V T E
BINARY_5999 T T Y K L I L N G K Q A K E E A I T E A V D A A I A E K Y F K L Y A N A K T V E G V W T L K D E T K T A T V T E
BINARY_6000 T T Y K L I L N G K Q A K E E A I T E A V D A A I A E K Y F K L Y A N A K T V E G V W T L K D E T K T F T V T E
BINARY_6001 T T Y K L I L N G K Q A K E E A I T E A V D A A I A E K Y F K L Y A N A K T V E G V W T Y K D E I L K A T V T E

BINARY_6002 T T Y K L I L N G K Q A K E E A I T E A V D A A I A E K Y F K L Y A N A K T V E G V W T Y K D E I L K F T V T E
BINARY_6003 T T Y K L I L N G K Q A K E E A I T E A V D A A I A E K Y F K L Y A N A K T V E G V W T Y K D E I L T A T V T E
BINARY_6004 T T Y K L I L N G K Q A K E E A I T E A V D A A I A E K Y F K L Y A N A K T V E G V W T Y K D E I L T F T V T E
BINARY_6005 T T Y K L I L N G K Q A K E E A I T E A V D A A I A E K Y F K L Y A N A K T V E G V W T Y K D E I K K A T V T E
BINARY_6006 T T Y K L I L N G K Q A K E E A I T E A V D A A I A E K Y F K L Y A N A K T V E G V W T Y K D E I K K F T V T E
BINARY_6007 T T Y K L I L N G K Q A K E E A I T E A V D A A I A E K Y F K L Y A N A K T V E G V W T Y K D E I K T A T V T E
BINARY_6008 T T Y K L I L N G K Q A K E E A I T E A V D A A I A E K Y F K L Y A N A K T V E G V W T Y K D E I K T F T V T E
BINARY_6009 T T Y K L I L N G K Q A K E E A I T E A V D A A I A E K Y F K L Y A N A K T V E G V W T Y K D E T L K A T V T E
BINARY_6010 T T Y K L I L N G K Q A K E E A I T E A V D A A I A E K Y F K L Y A N A K T V E G V W T Y K D E T L K F T V T E
BINARY_6011 T T Y K L I L N G K Q A K E E A I T E A V D A A I A E K Y F K L Y A N A K T V E G V W T Y K D E T L T A T V T E
BINARY_6012 T T Y K L I L N G K Q A K E E A I T E A V D A A I A E K Y F K L Y A N A K T V E G V W T Y K D E T L T F T V T E
BINARY_6013 T T Y K L I L N G K Q A K E E A I T E A V D A A I A E K Y F K L Y A N A K T V E G V W T Y K D E T K K A T V T E
BINARY_6014 T T Y K L I L N G K Q A K E E A I T E A V D A A I A E K Y F K L Y A N A K T V E G V W T Y K D E T K K F T V T E
BINARY_6015 T T Y K L I L N G K Q A K E E A I T E A V D A A I A E K Y F K L Y A N A K T V E G V W T Y K D E T K T A T V T E
BINARY_6016 T T Y K L I L N G K Q A K E E A I T E A V D A A I A E K Y F K L Y A N A K T V E G V W T Y K D E T K T F T V T E
BINARY_6017 T T Y K L I L N G K Q A K E E A I T E A V D A A T A E K Y I K L I A N A K T V E G V W T L K D E I L K A T V T E
BINARY_6018 T T Y K L I L N G K Q A K E E A I T E A V D A A T A E K Y I K L I A N A K T V E G V W T L K D E I L K F T V T E
BINARY_6019 T T Y K L I L N G K Q A K E E A I T E A V D A A T A E K Y I K L I A N A K T V E G V W T L K D E I L T A T V T E
BINARY_6020 T T Y K L I L N G K Q A K E E A I T E A V D A A T A E K Y I K L I A N A K T V E G V W T L K D E I L T F T V T E
BINARY_6021 T T Y K L I L N G K Q A K E E A I T E A V D A A T A E K Y I K L I A N A K T V E G V W T L K D E I K K A T V T E
BINARY_6022 T T Y K L I L N G K Q A K E E A I T E A V D A A T A E K Y I K L I A N A K T V E G V W T L K D E I K K F T V T E
BINARY_6023 T T Y K L I L N G K Q A K E E A I T E A V D A A T A E K Y I K L I A N A K T V E G V W T L K D E I K T A T V T E
BINARY_6024 T T Y K L I L N G K Q A K E E A I T E A V D A A T A E K Y I K L I A N A K T V E G V W T L K D E I K T F T V T E
BINARY_6025 T T Y K L I L N G K Q A K E E A I T E A V D A A T A E K Y I K L I A N A K T V E G V W T L K D E T L K A T V T E
BINARY_6026 T T Y K L I L N G K Q A K E E A I T E A V D A A T A E K Y I K L I A N A K T V E G V W T L K D E T L K F T V T E
BINARY_6027 T T Y K L I L N G K Q A K E E A I T E A V D A A T A E K Y I K L I A N A K T V E G V W T L K D E T L T A T V T E
BINARY_6028 T T Y K L I L N G K Q A K E E A I T E A V D A A T A E K Y I K L I A N A K T V E G V W T L K D E T L T F T V T E
BINARY_6029 T T Y K L I L N G K Q A K E E A I T E A V D A A T A E K Y I K L I A N A K T V E G V W T L K D E T K K A T V T E
BINARY_6030 T T Y K L I L N G K Q A K E E A I T E A V D A A T A E K Y I K L I A N A K T V E G V W T L K D E T K K F T V T E
BINARY_6031 T T Y K L I L N G K Q A K E E A I T E A V D A A T A E K Y I K L I A N A K T V E G V W T L K D E T K T A T V T E
BINARY_6032 T T Y K L I L N G K Q A K E E A I T E A V D A A T A E K Y I K L I A N A K T V E G V W T L K D E T K T F T V T E
BINARY_6033 T T Y K L I L N G K Q A K E E A I T E A V D A A T A E K Y I K L I A N A K T V E G V W T Y K D E I L K A T V T E
BINARY_6034 T T Y K L I L N G K Q A K E E A I T E A V D A A T A E K Y I K L I A N A K T V E G V W T Y K D E I L K F T V T E
BINARY_6035 T T Y K L I L N G K Q A K E E A I T E A V D A A T A E K Y I K L I A N A K T V E G V W T Y K D E I L T A T V T E
BINARY_6036 T T Y K L I L N G K Q A K E E A I T E A V D A A T A E K Y I K L I A N A K T V E G V W T Y K D E I L T F T V T E
BINARY_6037 T T Y K L I L N G K Q A K E E A I T E A V D A A T A E K Y I K L I A N A K T V E G V W T Y K D E I K K A T V T E
BINARY_6038 T T Y K L I L N G K Q A K E E A I T E A V D A A T A E K Y I K L I A N A K T V E G V W T Y K D E I K K F T V T E
BINARY_6039 T T Y K L I L N G K Q A K E E A I T E A V D A A T A E K Y I K L I A N A K T V E G V W T Y K D E I K T A T V T E
BINARY_6040 T T Y K L I L N G K Q A K E E A I T E A V D A A T A E K Y I K L I A N A K T V E G V W T Y K D E I K T F T V T E
BINARY_6041 T T Y K L I L N G K Q A K E E A I T E A V D A A T A E K Y I K L I A N A K T V E G V W T Y K D E T L K A T V T E
BINARY_6042 T T Y K L I L N G K Q A K E E A I T E A V D A A T A E K Y I K L I A N A K T V E G V W T Y K D E T L K F T V T E
BINARY_6043 T T Y K L I L N G K Q A K E E A I T E A V D A A T A E K Y I K L I A N A K T V E G V W T Y K D E T L T A T V T E
BINARY_6044 T T Y K L I L N G K Q A K E E A I T E A V D A A T A E K Y I K L I A N A K T V E G V W T Y K D E T L T F T V T E
BINARY_6045 T T Y K L I L N G K Q A K E E A I T E A V D A A T A E K Y I K L I A N A K T V E G V W T Y K D E T K K A T V T E
BINARY_6046 T T Y K L I L N G K Q A K E E A I T E A V D A A T A E K Y I K L I A N A K T V E G V W T Y K D E T K K F T V T E
BINARY_6047 T T Y K L I L N G K Q A K E E A I T E A V D A A T A E K Y I K L I A N A K T V E G V W T Y K D E T K T A T V T E
BINARY_6048 T T Y K L I L N G K Q A K E E A I T E A V D A A T A E K Y I K L I A N A K T V E G V W T Y K D E T K T F T V T E
BINARY_6049 T T Y K L I L N G K Q A K E E A I T E A V D A A T A E K Y I K L Y A N A K T V E G V W T L K D E I L K A T V T E
BINARY_6050 T T Y K L I L N G K Q A K E E A I T E A V D A A T A E K Y I K L Y A N A K T V E G V W T L K D E I L K F T V T E
BINARY_6051 T T Y K L I L N G K Q A K E E A I T E A V D A A T A E K Y I K L Y A N A K T V E G V W T L K D E I L T A T V T E
BINARY_6052 T T Y K L I L N G K Q A K E E A I T E A V D A A T A E K Y I K L Y A N A K T V E G V W T L K D E I L T F T V T E
BINARY_6053 T T Y K L I L N G K Q A K E E A I T E A V D A A T A E K Y I K L Y A N A K T V E G V W T L K D E I K K A T V T E
BINARY_6054 T T Y K L I L N G K Q A K E E A I T E A V D A A T A E K Y I K L Y A N A K T V E G V W T L K D E I K K F T V T E
BINARY_6055 T T Y K L I L N G K Q A K E E A I T E A V D A A T A E K Y I K L Y A N A K T V E G V W T L K D E I K T A T V T E
BINARY_6056 T T Y K L I L N G K Q A K E E A I T E A V D A A T A E K Y I K L Y A N A K T V E G V W T L K D E I K T F T V T E
BINARY_6057 T T Y K L I L N G K Q A K E E A I T E A V D A A T A E K Y I K L Y A N A K T V E G V W T L K D E T L K A T V T E
BINARY_6058 T T Y K L I L N G K Q A K E E A I T E A V D A A T A E K Y I K L Y A N A K T V E G V W T L K D E T L K F T V T E
BINARY_6059 T T Y K L I L N G K Q A K E E A I T E A V D A A T A E K Y I K L Y A N A K T V E G V W T L K D E T L T A T V T E
BINARY_6060 T T Y K L I L N G K Q A K E E A I T E A V D A A T A E K Y I K L Y A N A K T V E G V W T L K D E T L T F T V T E
BINARY_6061 T T Y K L I L N G K Q A K E E A I T E A V D A A T A E K Y I K L Y A N A K T V E G V W T L K D E T K K A T V T E
BINARY_6062 T T Y K L I L N G K Q A K E E A I T E A V D A A T A E K Y I K L Y A N A K T V E G V W T L K D E T K K F T V T E
BINARY_6063 T T Y K L I L N G K Q A K E E A I T E A V D A A T A E K Y I K L Y A N A K T V E G V W T L K D E T K T A T V T E
BINARY_6064 T T Y K L I L N G K Q A K E E A I T E A V D A A T A E K Y I K L Y A N A K T V E G V W T L K D E T K T F T V T E
BINARY_6065 T T Y K L I L N G K Q A K E E A I T E A V D A A T A E K Y I K L Y A N A K T V E G V W T Y K D E I L K A T V T E
BINARY_6066 T T Y K L I L N G K Q A K E E A I T E A V D A A T A E K Y I K L Y A N A K T V E G V W T Y K D E I L K F T V T E
BINARY_6067 T T Y K L I L N G K Q A K E E A I T E A V D A A T A E K Y I K L Y A N A K T V E G V W T Y K D E I L T A T V T E
BINARY_6068 T T Y K L I L N G K Q A K E E A I T E A V D A A T A E K Y I K L Y A N A K T V E G V W T Y K D E I L T F T V T E
BINARY_6069 T T Y K L I L N G K Q A K E E A I T E A V D A A T A E K Y I K L Y A N A K T V E G V W T Y K D E I K K A T V T E
BINARY_6070 T T Y K L I L N G K Q A K E E A I T E A V D A A T A E K Y I K L Y A N A K T V E G V W T Y K D E I K K F T V T E
BINARY_6071 T T Y K L I L N G K Q A K E E A I T E A V D A A T A E K Y I K L Y A N A K T V E G V W T Y K D E I K T A T V T E
BINARY_6072 T T Y K L I L N G K Q A K E E A I T E A V D A A T A E K Y I K L Y A N A K T V E G V W T Y K D E I K T F T V T E
BINARY_6073 T T Y K L I L N G K Q A K E E A I T E A V D A A T A E K Y I K L Y A N A K T V E G V W T Y K D E T L K A T V T E
BINARY_6074 T T Y K L I L N G K Q A K E E A I T E A V D A A T A E K Y I K L Y A N A K T V E G V W T Y K D E T L K F T V T E
BINARY_6075 T T Y K L I L N G K Q A K E E A I T E A V D A A T A E K Y I K L Y A N A K T V E G V W T Y K D E T L T A T V T E
BINARY_6076 T T Y K L I L N G K Q A K E E A I T E A V D A A T A E K Y I K L Y A N A K T V E G V W T Y K D E T L T F T V T E
BINARY_6077 T T Y K L I L N G K Q A K E E A I T E A V D A A T A E K Y I K L Y A N A K T V E G V W T Y K D E T K K A T V T E
BINARY_6078 T T Y K L I L N G K Q A K E E A I T E A V D A A T A E K Y I K L Y A N A K T V E G V W T Y K D E T K K F T V T E
BINARY_6079 T T Y K L I L N G K Q A K E E A I T E A V D A A T A E K Y I K L Y A N A K T V E G V W T Y K D E T K T A T V T E
BINARY_6080 T T Y K L I L N G K Q A K E E A I T E A V D A A T A E K Y I K L Y A N A K T V E G V W T Y K D E T K T F T V T E
BINARY_6081 T T Y K L I L N G K Q A K E E A I T E A V D A A T A E K Y F K L I A N A K T V E G V W T L K D E I L K A T V T E
BINARY_6082 T T Y K L I L N G K Q A K E E A I T E A V D A A T A E K Y F K L I A N A K T V E G V W T L K D E I L K F T V T E
BINARY_6083 T T Y K L I L N G K Q A K E E A I T E A V D A A T A E K Y F K L I A N A K T V E G V W T L K D E I L T A T V T E
BINARY_6084 T T Y K L I L N G K Q A K E E A I T E A V D A A T A E K Y F K L I A N A K T V E G V W T L K D E I L T F T V T E
BINARY_6085 T T Y K L I L N G K Q A K E E A I T E A V D A A T A E K Y F K L I A N A K T V E G V W T L K D E I K K A T V T E
BINARY_6086 T T Y K L I L N G K Q A K E E A I T E A V D A A T A E K Y F K L I A N A K T V E G V W T L K D E I K K F T V T E
BINARY_6087 T T Y K L I L N G K Q A K E E A I T E A V D A A T A E K Y F K L I A N A K T V E G V W T L K D E I K T A T V T E
BINARY_6088 T T Y K L I L N G K Q A K E E A I T E A V D A A T A E K Y F K L I A N A K T V E G V W T L K D E I K T F T V T E
BINARY_6089 T T Y K L I L N G K Q A K E E A I T E A V D A A T A E K Y F K L I A N A K T V E G V W T L K D E T L K A T V T E
BINARY_6090 T T Y K L I L N G K Q A K E E A I T E A V D A A T A E K Y F K L I A N A K T V E G V W T L K D E T L K F T V T E
BINARY_6091 T T Y K L I L N G K Q A K E E A I T E A V D A A T A E K Y F K L I A N A K T V E G V W T L K D E T L T A T V T E
BINARY_6092 T T Y K L I L N G K Q A K E E A I T E A V D A A T A E K Y F K L I A N A K T V E G V W T L K D E T L T F T V T E
BINARY_6093 T T Y K L I L N G K Q A K E E A I T E A V D A A T A E K Y F K L I A N A K T V E G V W T L K D E T K K A T V T E
BINARY_6094 T T Y K L I L N G K Q A K E E A I T E A V D A A T A E K Y F K L I A N A K T V E G V W T L K D E T K K F T V T E
BINARY_6095 T T Y K L I L N G K Q A K E E A I T E A V D A A T A E K Y F K L I A N A K T V E G V W T L K D E T K T A T V T E
BINARY_6096 T T Y K L I L N G K Q A K E E A I T E A V D A A T A E K Y F K L I A N A K T V E G V W T L K D E T K T F T V T E

BINARY_6097 T T Y K L I L N G K Q A K E E A I T E A V D A A T A E K Y F K L I A N A K T V E G V W T Y K D E I L K A T V T E
BINARY_6098 T T Y K L I L N G K Q A K E E A I T E A V D A A T A E K Y F K L I A N A K T V E G V W T Y K D E I L K F T V T E
BINARY_6099 T T Y K L I L N G K Q A K E E A I T E A V D A A T A E K Y F K L I A N A K T V E G V W T Y K D E I L T A T V T E
BINARY_6100 T T Y K L I L N G K Q A K E E A I T E A V D A A T A E K Y F K L I A N A K T V E G V W T Y K D E I L T F T V T E
BINARY_6101 T T Y K L I L N G K Q A K E E A I T E A V D A A T A E K Y F K L I A N A K T V E G V W T Y K D E I K K A T V T E
BINARY_6102 T T Y K L I L N G K Q A K E E A I T E A V D A A T A E K Y F K L I A N A K T V E G V W T Y K D E I K K F T V T E
BINARY_6103 T T Y K L I L N G K Q A K E E A I T E A V D A A T A E K Y F K L I A N A K T V E G V W T Y K D E I K T A T V T E
BINARY_6104 T T Y K L I L N G K Q A K E E A I T E A V D A A T A E K Y F K L I A N A K T V E G V W T Y K D E I K T F T V T E
BINARY_6105 T T Y K L I L N G K Q A K E E A I T E A V D A A T A E K Y F K L I A N A K T V E G V W T Y K D E T L K A T V T E
BINARY_6106 T T Y K L I L N G K Q A K E E A I T E A V D A A T A E K Y F K L I A N A K T V E G V W T Y K D E T L K F T V T E
BINARY_6107 T T Y K L I L N G K Q A K E E A I T E A V D A A T A E K Y F K L I A N A K T V E G V W T Y K D E T L T A T V T E
BINARY_6108 T T Y K L I L N G K Q A K E E A I T E A V D A A T A E K Y F K L I A N A K T V E G V W T Y K D E T L T F T V T E
BINARY_6109 T T Y K L I L N G K Q A K E E A I T E A V D A A T A E K Y F K L I A N A K T V E G V W T Y K D E T K K A T V T E
BINARY_6110 T T Y K L I L N G K Q A K E E A I T E A V D A A T A E K Y F K L I A N A K T V E G V W T Y K D E T K K F T V T E
BINARY_6111 T T Y K L I L N G K Q A K E E A I T E A V D A A T A E K Y F K L I A N A K T V E G V W T Y K D E T K T A T V T E
BINARY_6112 T T Y K L I L N G K Q A K E E A I T E A V D A A T A E K Y F K L I A N A K T V E G V W T Y K D E T K T F T V T E
BINARY_6113 T T Y K L I L N G K Q A K E E A I T E A V D A A T A E K Y F K L Y A N A K T V E G V W T L K D E I L K A T V T E
BINARY_6114 T T Y K L I L N G K Q A K E E A I T E A V D A A T A E K Y F K L Y A N A K T V E G V W T L K D E I L K F T V T E
BINARY_6115 T T Y K L I L N G K Q A K E E A I T E A V D A A T A E K Y F K L Y A N A K T V E G V W T L K D E I L T A T V T E
BINARY_6116 T T Y K L I L N G K Q A K E E A I T E A V D A A T A E K Y F K L Y A N A K T V E G V W T L K D E I L T F T V T E
BINARY_6117 T T Y K L I L N G K Q A K E E A I T E A V D A A T A E K Y F K L Y A N A K T V E G V W T L K D E I K K A T V T E
BINARY_6118 T T Y K L I L N G K Q A K E E A I T E A V D A A T A E K Y F K L Y A N A K T V E G V W T L K D E I K K F T V T E
BINARY_6119 T T Y K L I L N G K Q A K E E A I T E A V D A A T A E K Y F K L Y A N A K T V E G V W T L K D E I K T A T V T E
BINARY_6120 T T Y K L I L N G K Q A K E E A I T E A V D A A T A E K Y F K L Y A N A K T V E G V W T L K D E I K T F T V T E
BINARY_6121 T T Y K L I L N G K Q A K E E A I T E A V D A A T A E K Y F K L Y A N A K T V E G V W T L K D E T L K A T V T E
BINARY_6122 T T Y K L I L N G K Q A K E E A I T E A V D A A T A E K Y F K L Y A N A K T V E G V W T L K D E T L K F T V T E
BINARY_6123 T T Y K L I L N G K Q A K E E A I T E A V D A A T A E K Y F K L Y A N A K T V E G V W T L K D E T L T A T V T E
BINARY_6124 T T Y K L I L N G K Q A K E E A I T E A V D A A T A E K Y F K L Y A N A K T V E G V W T L K D E T L T F T V T E
BINARY_6125 T T Y K L I L N G K Q A K E E A I T E A V D A A T A E K Y F K L Y A N A K T V E G V W T L K D E T K K A T V T E
BINARY_6126 T T Y K L I L N G K Q A K E E A I T E A V D A A T A E K Y F K L Y A N A K T V E G V W T L K D E T K K F T V T E
BINARY_6127 T T Y K L I L N G K Q A K E E A I T E A V D A A T A E K Y F K L Y A N A K T V E G V W T L K D E T K T A T V T E
BINARY_6128 T T Y K L I L N G K Q A K E E A I T E A V D A A T A E K Y F K L Y A N A K T V E G V W T L K D E T K T F T V T E
BINARY_6129 T T Y K L I L N G K Q A K E E A I T E A V D A A T A E K Y F K L Y A N A K T V E G V W T Y K D E I L K A T V T E
BINARY_6130 T T Y K L I L N G K Q A K E E A I T E A V D A A T A E K Y F K L Y A N A K T V E G V W T Y K D E I L K F T V T E
BINARY_6131 T T Y K L I L N G K Q A K E E A I T E A V D A A T A E K Y F K L Y A N A K T V E G V W T Y K D E I L T A T V T E
BINARY_6132 T T Y K L I L N G K Q A K E E A I T E A V D A A T A E K Y F K L Y A N A K T V E G V W T Y K D E I L T F T V T E
BINARY_6133 T T Y K L I L N G K Q A K E E A I T E A V D A A T A E K Y F K L Y A N A K T V E G V W T Y K D E I K K A T V T E
BINARY_6134 T T Y K L I L N G K Q A K E E A I T E A V D A A T A E K Y F K L Y A N A K T V E G V W T Y K D E I K K F T V T E
BINARY_6135 T T Y K L I L N G K Q A K E E A I T E A V D A A T A E K Y F K L Y A N A K T V E G V W T Y K D E I K T A T V T E
BINARY_6136 T T Y K L I L N G K Q A K E E A I T E A V D A A T A E K Y F K L Y A N A K T V E G V W T Y K D E I K T F T V T E
BINARY_6137 T T Y K L I L N G K Q A K E E A I T E A V D A A T A E K Y F K L Y A N A K T V E G V W T Y K D E T L K A T V T E
BINARY_6138 T T Y K L I L N G K Q A K E E A I T E A V D A A T A E K Y F K L Y A N A K T V E G V W T Y K D E T L K F T V T E
BINARY_6139 T T Y K L I L N G K Q A K E E A I T E A V D A A T A E K Y F K L Y A N A K T V E G V W T Y K D E T L T A T V T E
BINARY_6140 T T Y K L I L N G K Q A K E E A I T E A V D A A T A E K Y F K L Y A N A K T V E G V W T Y K D E T L T F T V T E
BINARY_6141 T T Y K L I L N G K Q A K E E A I T E A V D A A T A E K Y F K L Y A N A K T V E G V W T Y K D E T K K A T V T E
BINARY_6142 T T Y K L I L N G K Q A K E E A I T E A V D A A T A E K Y F K L Y A N A K T V E G V W T Y K D E T K K F T V T E
BINARY_6143 T T Y K L I L N G K Q A K E E A I T E A V D A A T A E K Y F K L Y A N A K T V E G V W T Y K D E T K T A T V T E
BINARY_6144 T T Y K L I L N G K Q A K E E A I T E A V D A A T A E K Y F K L Y A N A K T V E G V W T Y K D E T K T F T V T E
BINARY_6145 T T Y K L I L N G K Q L K E E A I K E L V D A G I A E K Y I K L I A N A K T V E G V W T L K D E I L K A T V T E
BINARY_6146 T T Y K L I L N G K Q L K E E A I K E L V D A G I A E K Y I K L I A N A K T V E G V W T L K D E I L K F T V T E
BINARY_6147 T T Y K L I L N G K Q L K E E A I K E L V D A G I A E K Y I K L I A N A K T V E G V W T L K D E I L T A T V T E
BINARY_6148 T T Y K L I L N G K Q L K E E A I K E L V D A G I A E K Y I K L I A N A K T V E G V W T L K D E I L T F T V T E
BINARY_6149 T T Y K L I L N G K Q L K E E A I K E L V D A G I A E K Y I K L I A N A K T V E G V W T L K D E I K K A T V T E
BINARY_6150 T T Y K L I L N G K Q L K E E A I K E L V D A G I A E K Y I K L I A N A K T V E G V W T L K D E I K K F T V T E
BINARY_6151 T T Y K L I L N G K Q L K E E A I K E L V D A G I A E K Y I K L I A N A K T V E G V W T L K D E I K T A T V T E
BINARY_6152 T T Y K L I L N G K Q L K E E A I K E L V D A G I A E K Y I K L I A N A K T V E G V W T L K D E I K T F T V T E
BINARY_6153 T T Y K L I L N G K Q L K E E A I K E L V D A G I A E K Y I K L I A N A K T V E G V W T L K D E T L K A T V T E
BINARY_6154 T T Y K L I L N G K Q L K E E A I K E L V D A G I A E K Y I K L I A N A K T V E G V W T L K D E T L K F T V T E
BINARY_6155 T T Y K L I L N G K Q L K E E A I K E L V D A G I A E K Y I K L I A N A K T V E G V W T L K D E T L T A T V T E
BINARY_6156 T T Y K L I L N G K Q L K E E A I K E L V D A G I A E K Y I K L I A N A K T V E G V W T L K D E T L T F T V T E
BINARY_6157 T T Y K L I L N G K Q L K E E A I K E L V D A G I A E K Y I K L I A N A K T V E G V W T L K D E T K K A T V T E
BINARY_6158 T T Y K L I L N G K Q L K E E A I K E L V D A G I A E K Y I K L I A N A K T V E G V W T L K D E T K K F T V T E
BINARY_6159 T T Y K L I L N G K Q L K E E A I K E L V D A G I A E K Y I K L I A N A K T V E G V W T L K D E T K T A T V T E
BINARY_6160 T T Y K L I L N G K Q L K E E A I K E L V D A G I A E K Y I K L I A N A K T V E G V W T L K D E T K T F T V T E
BINARY_6161 T T Y K L I L N G K Q L K E E A I K E L V D A G I A E K Y I K L I A N A K T V E G V W T Y K D E I L K A T V T E
BINARY_6162 T T Y K L I L N G K Q L K E E A I K E L V D A G I A E K Y I K L I A N A K T V E G V W T Y K D E I L K F T V T E
BINARY_6163 T T Y K L I L N G K Q L K E E A I K E L V D A G I A E K Y I K L I A N A K T V E G V W T Y K D E I L T A T V T E
BINARY_6164 T T Y K L I L N G K Q L K E E A I K E L V D A G I A E K Y I K L I A N A K T V E G V W T Y K D E I L T F T V T E
BINARY_6165 T T Y K L I L N G K Q L K E E A I K E L V D A G I A E K Y I K L I A N A K T V E G V W T Y K D E I K K A T V T E
BINARY_6166 T T Y K L I L N G K Q L K E E A I K E L V D A G I A E K Y I K L I A N A K T V E G V W T Y K D E I K K F T V T E
BINARY_6167 T T Y K L I L N G K Q L K E E A I K E L V D A G I A E K Y I K L I A N A K T V E G V W T Y K D E I K T A T V T E
BINARY_6168 T T Y K L I L N G K Q L K E E A I K E L V D A G I A E K Y I K L I A N A K T V E G V W T Y K D E I K T F T V T E
BINARY_6169 T T Y K L I L N G K Q L K E E A I K E L V D A G I A E K Y I K L I A N A K T V E G V W T Y K D E T L K A T V T E
BINARY_6170 T T Y K L I L N G K Q L K E E A I K E L V D A G I A E K Y I K L I A N A K T V E G V W T Y K D E T L K F T V T E
BINARY_6171 T T Y K L I L N G K Q L K E E A I K E L V D A G I A E K Y I K L I A N A K T V E G V W T Y K D E T L T A T V T E
BINARY_6172 T T Y K L I L N G K Q L K E E A I K E L V D A G I A E K Y I K L I A N A K T V E G V W T Y K D E T L T F T V T E
BINARY_6173 T T Y K L I L N G K Q L K E E A I K E L V D A G I A E K Y I K L I A N A K T V E G V W T Y K D E T K K A T V T E
BINARY_6174 T T Y K L I L N G K Q L K E E A I K E L V D A G I A E K Y I K L I A N A K T V E G V W T Y K D E T K K F T V T E
BINARY_6175 T T Y K L I L N G K Q L K E E A I K E L V D A G I A E K Y I K L I A N A K T V E G V W T Y K D E T K T A T V T E
BINARY_6176 T T Y K L I L N G K Q L K E E A I K E L V D A G I A E K Y I K L I A N A K T V E G V W T Y K D E T K T F T V T E
BINARY_6177 T T Y K L I L N G K Q L K E E A I K E L V D A G I A E K Y I K L Y A N A K T V E G V W T L K D E I L K A T V T E
BINARY_6178 T T Y K L I L N G K Q L K E E A I K E L V D A G I A E K Y I K L Y A N A K T V E G V W T L K D E I L K F T V T E
BINARY_6179 T T Y K L I L N G K Q L K E E A I K E L V D A G I A E K Y I K L Y A N A K T V E G V W T L K D E I L T A T V T E
BINARY_6180 T T Y K L I L N G K Q L K E E A I K E L V D A G I A E K Y I K L Y A N A K T V E G V W T L K D E I L T F T V T E
BINARY_6181 T T Y K L I L N G K Q L K E E A I K E L V D A G I A E K Y I K L Y A N A K T V E G V W T L K D E I K K A T V T E
BINARY_6182 T T Y K L I L N G K Q L K E E A I K E L V D A G I A E K Y I K L Y A N A K T V E G V W T L K D E I K K F T V T E
BINARY_6183 T T Y K L I L N G K Q L K E E A I K E L V D A G I A E K Y I K L Y A N A K T V E G V W T L K D E I K T A T V T E
BINARY_6184 T T Y K L I L N G K Q L K E E A I K E L V D A G I A E K Y I K L Y A N A K T V E G V W T L K D E I K T F T V T E
BINARY_6185 T T Y K L I L N G K Q L K E E A I K E L V D A G I A E K Y I K L Y A N A K T V E G V W T L K D E T L K A T V T E
BINARY_6186 T T Y K L I L N G K Q L K E E A I K E L V D A G I A E K Y I K L Y A N A K T V E G V W T L K D E T L K F T V T E
BINARY_6187 T T Y K L I L N G K Q L K E E A I K E L V D A G I A E K Y I K L Y A N A K T V E G V W T L K D E T L T A T V T E
BINARY_6188 T T Y K L I L N G K Q L K E E A I K E L V D A G I A E K Y I K L Y A N A K T V E G V W T L K D E T L T F T V T E
BINARY_6189 T T Y K L I L N G K Q L K E E A I K E L V D A G I A E K Y I K L Y A N A K T V E G V W T L K D E T K K A T V T E
BINARY_6190 T T Y K L I L N G K Q L K E E A I K E L V D A G I A E K Y I K L Y A N A K T V E G V W T L K D E T K K F T V T E
BINARY_6191 T T Y K L I L N G K Q L K E E A I K E L V D A G I A E K Y I K L Y A N A K T V E G V W T L K D E T K T A T V T E

BINARY_6192 T T Y K L I L N G K Q L K E E A I K E L V D A G I A E K Y I K L Y A N A K T V E G V W T L K D E T K T F T V T E
BINARY_6193 T T Y K L I L N G K Q L K E E A I K E L V D A G I A E K Y I K L Y A N A K T V E G V W T Y K D E I L K A T V T E
BINARY_6194 T T Y K L I L N G K Q L K E E A I K E L V D A G I A E K Y I K L Y A N A K T V E G V W T Y K D E I L K F T V T E
BINARY_6195 T T Y K L I L N G K Q L K E E A I K E L V D A G I A E K Y I K L Y A N A K T V E G V W T Y K D E I L T A T V T E
BINARY_6196 T T Y K L I L N G K Q L K E E A I K E L V D A G I A E K Y I K L Y A N A K T V E G V W T Y K D E I L T F T V T E
BINARY_6197 T T Y K L I L N G K Q L K E E A I K E L V D A G I A E K Y I K L Y A N A K T V E G V W T Y K D E I K K A T V T E
BINARY_6198 T T Y K L I L N G K Q L K E E A I K E L V D A G I A E K Y I K L Y A N A K T V E G V W T Y K D E I K K F T V T E
BINARY_6199 T T Y K L I L N G K Q L K E E A I K E L V D A G I A E K Y I K L Y A N A K T V E G V W T Y K D E I K T A T V T E
BINARY_6200 T T Y K L I L N G K Q L K E E A I K E L V D A G I A E K Y I K L Y A N A K T V E G V W T Y K D E I K T F T V T E
BINARY_6201 T T Y K L I L N G K Q L K E E A I K E L V D A G I A E K Y I K L Y A N A K T V E G V W T Y K D E T L K A T V T E
BINARY_6202 T T Y K L I L N G K Q L K E E A I K E L V D A G I A E K Y I K L Y A N A K T V E G V W T Y K D E T L K F T V T E
BINARY_6203 T T Y K L I L N G K Q L K E E A I K E L V D A G I A E K Y I K L Y A N A K T V E G V W T Y K D E T L T A T V T E
BINARY_6204 T T Y K L I L N G K Q L K E E A I K E L V D A G I A E K Y I K L Y A N A K T V E G V W T Y K D E T L T F T V T E
BINARY_6205 T T Y K L I L N G K Q L K E E A I K E L V D A G I A E K Y I K L Y A N A K T V E G V W T Y K D E T K K A T V T E
BINARY_6206 T T Y K L I L N G K Q L K E E A I K E L V D A G I A E K Y I K L Y A N A K T V E G V W T Y K D E T K K F T V T E
BINARY_6207 T T Y K L I L N G K Q L K E E A I K E L V D A G I A E K Y I K L Y A N A K T V E G V W T Y K D E T K T A T V T E
BINARY_6208 T T Y K L I L N G K Q L K E E A I K E L V D A G I A E K Y I K L Y A N A K T V E G V W T Y K D E T K T F T V T E
BINARY_6209 T T Y K L I L N G K Q L K E E A I K E L V D A G I A E K Y F K L I A N A K T V E G V W T L K D E I L K A T V T E
BINARY_6210 T T Y K L I L N G K Q L K E E A I K E L V D A G I A E K Y F K L I A N A K T V E G V W T L K D E I L K F T V T E
BINARY_6211 T T Y K L I L N G K Q L K E E A I K E L V D A G I A E K Y F K L I A N A K T V E G V W T L K D E I L T A T V T E
BINARY_6212 T T Y K L I L N G K Q L K E E A I K E L V D A G I A E K Y F K L I A N A K T V E G V W T L K D E I L T F T V T E
BINARY_6213 T T Y K L I L N G K Q L K E E A I K E L V D A G I A E K Y F K L I A N A K T V E G V W T L K D E I K K A T V T E
BINARY_6214 T T Y K L I L N G K Q L K E E A I K E L V D A G I A E K Y F K L I A N A K T V E G V W T L K D E I K K F T V T E
BINARY_6215 T T Y K L I L N G K Q L K E E A I K E L V D A G I A E K Y F K L I A N A K T V E G V W T L K D E I K T A T V T E
BINARY_6216 T T Y K L I L N G K Q L K E E A I K E L V D A G I A E K Y F K L I A N A K T V E G V W T L K D E I K T F T V T E
BINARY_6217 T T Y K L I L N G K Q L K E E A I K E L V D A G I A E K Y F K L I A N A K T V E G V W T L K D E T L K A T V T E
BINARY_6218 T T Y K L I L N G K Q L K E E A I K E L V D A G I A E K Y F K L I A N A K T V E G V W T L K D E T L K F T V T E
BINARY_6219 T T Y K L I L N G K Q L K E E A I K E L V D A G I A E K Y F K L I A N A K T V E G V W T L K D E T L T A T V T E
BINARY_6220 T T Y K L I L N G K Q L K E E A I K E L V D A G I A E K Y F K L I A N A K T V E G V W T L K D E T L T F T V T E
BINARY_6221 T T Y K L I L N G K Q L K E E A I K E L V D A G I A E K Y F K L I A N A K T V E G V W T L K D E T K K A T V T E
BINARY_6222 T T Y K L I L N G K Q L K E E A I K E L V D A G I A E K Y F K L I A N A K T V E G V W T L K D E T K K F T V T E
BINARY_6223 T T Y K L I L N G K Q L K E E A I K E L V D A G I A E K Y F K L I A N A K T V E G V W T L K D E T K T A T V T E
BINARY_6224 T T Y K L I L N G K Q L K E E A I K E L V D A G I A E K Y F K L I A N A K T V E G V W T L K D E T K T F T V T E
BINARY_6225 T T Y K L I L N G K Q L K E E A I K E L V D A G I A E K Y F K L I A N A K T V E G V W T Y K D E I L K A T V T E
BINARY_6226 T T Y K L I L N G K Q L K E E A I K E L V D A G I A E K Y F K L I A N A K T V E G V W T Y K D E I L K F T V T E
BINARY_6227 T T Y K L I L N G K Q L K E E A I K E L V D A G I A E K Y F K L I A N A K T V E G V W T Y K D E I L T A T V T E
BINARY_6228 T T Y K L I L N G K Q L K E E A I K E L V D A G I A E K Y F K L I A N A K T V E G V W T Y K D E I L T F T V T E
BINARY_6229 T T Y K L I L N G K Q L K E E A I K E L V D A G I A E K Y F K L I A N A K T V E G V W T Y K D E I K K A T V T E
BINARY_6230 T T Y K L I L N G K Q L K E E A I K E L V D A G I A E K Y F K L I A N A K T V E G V W T Y K D E I K K F T V T E
BINARY_6231 T T Y K L I L N G K Q L K E E A I K E L V D A G I A E K Y F K L I A N A K T V E G V W T Y K D E I K T A T V T E
BINARY_6232 T T Y K L I L N G K Q L K E E A I K E L V D A G I A E K Y F K L I A N A K T V E G V W T Y K D E I K T F T V T E
BINARY_6233 T T Y K L I L N G K Q L K E E A I K E L V D A G I A E K Y F K L I A N A K T V E G V W T Y K D E T L K A T V T E
BINARY_6234 T T Y K L I L N G K Q L K E E A I K E L V D A G I A E K Y F K L I A N A K T V E G V W T Y K D E T L K F T V T E
BINARY_6235 T T Y K L I L N G K Q L K E E A I K E L V D A G I A E K Y F K L I A N A K T V E G V W T Y K D E T L T A T V T E
BINARY_6236 T T Y K L I L N G K Q L K E E A I K E L V D A G I A E K Y F K L I A N A K T V E G V W T Y K D E T L T F T V T E
BINARY_6237 T T Y K L I L N G K Q L K E E A I K E L V D A G I A E K Y F K L I A N A K T V E G V W T Y K D E T K K A T V T E
BINARY_6238 T T Y K L I L N G K Q L K E E A I K E L V D A G I A E K Y F K L I A N A K T V E G V W T Y K D E T K K F T V T E
BINARY_6239 T T Y K L I L N G K Q L K E E A I K E L V D A G I A E K Y F K L I A N A K T V E G V W T Y K D E T K T A T V T E
BINARY_6240 T T Y K L I L N G K Q L K E E A I K E L V D A G I A E K Y F K L I A N A K T V E G V W T Y K D E T K T F T V T E
BINARY_6241 T T Y K L I L N G K Q L K E E A I K E L V D A G I A E K Y F K L Y A N A K T V E G V W T L K D E I L K A T V T E
BINARY_6242 T T Y K L I L N G K Q L K E E A I K E L V D A G I A E K Y F K L Y A N A K T V E G V W T L K D E I L K F T V T E
BINARY_6243 T T Y K L I L N G K Q L K E E A I K E L V D A G I A E K Y F K L Y A N A K T V E G V W T L K D E I L T A T V T E
BINARY_6244 T T Y K L I L N G K Q L K E E A I K E L V D A G I A E K Y F K L Y A N A K T V E G V W T L K D E I L T F T V T E
BINARY_6245 T T Y K L I L N G K Q L K E E A I K E L V D A G I A E K Y F K L Y A N A K T V E G V W T L K D E I K K A T V T E
BINARY_6246 T T Y K L I L N G K Q L K E E A I K E L V D A G I A E K Y F K L Y A N A K T V E G V W T L K D E I K K F T V T E
BINARY_6247 T T Y K L I L N G K Q L K E E A I K E L V D A G I A E K Y F K L Y A N A K T V E G V W T L K D E I K T A T V T E
BINARY_6248 T T Y K L I L N G K Q L K E E A I K E L V D A G I A E K Y F K L Y A N A K T V E G V W T L K D E I K T F T V T E
BINARY_6249 T T Y K L I L N G K Q L K E E A I K E L V D A G I A E K Y F K L Y A N A K T V E G V W T L K D E T L K A T V T E
BINARY_6250 T T Y K L I L N G K Q L K E E A I K E L V D A G I A E K Y F K L Y A N A K T V E G V W T L K D E T L K F T V T E
BINARY_6251 T T Y K L I L N G K Q L K E E A I K E L V D A G I A E K Y F K L Y A N A K T V E G V W T L K D E T L T A T V T E
BINARY_6252 T T Y K L I L N G K Q L K E E A I K E L V D A G I A E K Y F K L Y A N A K T V E G V W T L K D E T L T F T V T E
BINARY_6253 T T Y K L I L N G K Q L K E E A I K E L V D A G I A E K Y F K L Y A N A K T V E G V W T L K D E T K K A T V T E
BINARY_6254 T T Y K L I L N G K Q L K E E A I K E L V D A G I A E K Y F K L Y A N A K T V E G V W T L K D E T K K F T V T E
BINARY_6255 T T Y K L I L N G K Q L K E E A I K E L V D A G I A E K Y F K L Y A N A K T V E G V W T L K D E T K T A T V T E
BINARY_6256 T T Y K L I L N G K Q L K E E A I K E L V D A G I A E K Y F K L Y A N A K T V E G V W T L K D E T K T F T V T E
BINARY_6257 T T Y K L I L N G K Q L K E E A I K E L V D A G I A E K Y F K L Y A N A K T V E G V W T Y K D E I L K A T V T E
BINARY_6258 T T Y K L I L N G K Q L K E E A I K E L V D A G I A E K Y F K L Y A N A K T V E G V W T Y K D E I L K F T V T E
BINARY_6259 T T Y K L I L N G K Q L K E E A I K E L V D A G I A E K Y F K L Y A N A K T V E G V W T Y K D E I L T A T V T E
BINARY_6260 T T Y K L I L N G K Q L K E E A I K E L V D A G I A E K Y F K L Y A N A K T V E G V W T Y K D E I L T F T V T E
BINARY_6261 T T Y K L I L N G K Q L K E E A I K E L V D A G I A E K Y F K L Y A N A K T V E G V W T Y K D E I K K A T V T E
BINARY_6262 T T Y K L I L N G K Q L K E E A I K E L V D A G I A E K Y F K L Y A N A K T V E G V W T Y K D E I K K F T V T E
BINARY_6263 T T Y K L I L N G K Q L K E E A I K E L V D A G I A E K Y F K L Y A N A K T V E G V W T Y K D E I K T A T V T E
BINARY_6264 T T Y K L I L N G K Q L K E E A I K E L V D A G I A E K Y F K L Y A N A K T V E G V W T Y K D E I K T F T V T E
BINARY_6265 T T Y K L I L N G K Q L K E E A I K E L V D A G I A E K Y F K L Y A N A K T V E G V W T Y K D E T L K A T V T E
BINARY_6266 T T Y K L I L N G K Q L K E E A I K E L V D A G I A E K Y F K L Y A N A K T V E G V W T Y K D E T L K F T V T E
BINARY_6267 T T Y K L I L N G K Q L K E E A I K E L V D A G I A E K Y F K L Y A N A K T V E G V W T Y K D E T L T A T V T E
BINARY_6268 T T Y K L I L N G K Q L K E E A I K E L V D A G I A E K Y F K L Y A N A K T V E G V W T Y K D E T L T F T V T E
BINARY_6269 T T Y K L I L N G K Q L K E E A I K E L V D A G I A E K Y F K L Y A N A K T V E G V W T Y K D E T K K A T V T E
BINARY_6270 T T Y K L I L N G K Q L K E E A I K E L V D A G I A E K Y F K L Y A N A K T V E G V W T Y K D E T K K F T V T E
BINARY_6271 T T Y K L I L N G K Q L K E E A I K E L V D A G I A E K Y F K L Y A N A K T V E G V W T Y K D E T K T A T V T E
BINARY_6272 T T Y K L I L N G K Q L K E E A I K E L V D A G I A E K Y F K L Y A N A K T V E G V W T Y K D E T K T F T V T E
BINARY_6273 T T Y K L I L N G K Q L K E E A I K E L V D A G T A E K Y I K L I A N A K T V E G V W T L K D E I L K A T V T E
BINARY_6274 T T Y K L I L N G K Q L K E E A I K E L V D A G T A E K Y I K L I A N A K T V E G V W T L K D E I L K F T V T E
BINARY_6275 T T Y K L I L N G K Q L K E E A I K E L V D A G T A E K Y I K L I A N A K T V E G V W T L K D E I L T A T V T E
BINARY_6276 T T Y K L I L N G K Q L K E E A I K E L V D A G T A E K Y I K L I A N A K T V E G V W T L K D E I L T F T V T E
BINARY_6277 T T Y K L I L N G K Q L K E E A I K E L V D A G T A E K Y I K L I A N A K T V E G V W T L K D E I K K A T V T E
BINARY_6278 T T Y K L I L N G K Q L K E E A I K E L V D A G T A E K Y I K L I A N A K T V E G V W T L K D E I K K F T V T E
BINARY_6279 T T Y K L I L N G K Q L K E E A I K E L V D A G T A E K Y I K L I A N A K T V E G V W T L K D E I K T A T V T E
BINARY_6280 T T Y K L I L N G K Q L K E E A I K E L V D A G T A E K Y I K L I A N A K T V E G V W T L K D E I K T F T V T E
BINARY_6281 T T Y K L I L N G K Q L K E E A I K E L V D A G T A E K Y I K L I A N A K T V E G V W T L K D E T L K A T V T E
BINARY_6282 T T Y K L I L N G K Q L K E E A I K E L V D A G T A E K Y I K L I A N A K T V E G V W T L K D E T L K F T V T E
BINARY_6283 T T Y K L I L N G K Q L K E E A I K E L V D A G T A E K Y I K L I A N A K T V E G V W T L K D E T L T A T V T E
BINARY_6284 T T Y K L I L N G K Q L K E E A I K E L V D A G T A E K Y I K L I A N A K T V E G V W T L K D E T L T F T V T E
BINARY_6285 T T Y K L I L N G K Q L K E E A I K E L V D A G T A E K Y I K L I A N A K T V E G V W T L K D E T K K A T V T E
BINARY_6286 T T Y K L I L N G K Q L K E E A I K E L V D A G T A E K Y I K L I A N A K T V E G V W T L K D E T K K F T V T E

BINARY_6287 T T Y K L I L N G K Q L K E E A I K E L V D A G T A E K Y I K L I A N A K T V E G V W T L K D E T K T A T V T E
BINARY_6288 T T Y K L I L N G K Q L K E E A I K E L V D A G T A E K Y I K L I A N A K T V E G V W T L K D E T K T F T V T E
BINARY_6289 T T Y K L I L N G K Q L K E E A I K E L V D A G T A E K Y I K L I A N A K T V E G V W T Y K D E I L K A T V T E
BINARY_6290 T T Y K L I L N G K Q L K E E A I K E L V D A G T A E K Y I K L I A N A K T V E G V W T Y K D E I L K F T V T E
BINARY_6291 T T Y K L I L N G K Q L K E E A I K E L V D A G T A E K Y I K L I A N A K T V E G V W T Y K D E I L T A T V T E
BINARY_6292 T T Y K L I L N G K Q L K E E A I K E L V D A G T A E K Y I K L I A N A K T V E G V W T Y K D E I L T F T V T E
BINARY_6293 T T Y K L I L N G K Q L K E E A I K E L V D A G T A E K Y I K L I A N A K T V E G V W T Y K D E I K K A T V T E
BINARY_6294 T T Y K L I L N G K Q L K E E A I K E L V D A G T A E K Y I K L I A N A K T V E G V W T Y K D E I K K F T V T E
BINARY_6295 T T Y K L I L N G K Q L K E E A I K E L V D A G T A E K Y I K L I A N A K T V E G V W T Y K D E I K T A T V T E
BINARY_6296 T T Y K L I L N G K Q L K E E A I K E L V D A G T A E K Y I K L I A N A K T V E G V W T Y K D E I K T F T V T E
BINARY_6297 T T Y K L I L N G K Q L K E E A I K E L V D A G T A E K Y I K L I A N A K T V E G V W T Y K D E T L K A T V T E
BINARY_6298 T T Y K L I L N G K Q L K E E A I K E L V D A G T A E K Y I K L I A N A K T V E G V W T Y K D E T L K F T V T E
BINARY_6299 T T Y K L I L N G K Q L K E E A I K E L V D A G T A E K Y I K L I A N A K T V E G V W T Y K D E T L T A T V T E
BINARY_6300 T T Y K L I L N G K Q L K E E A I K E L V D A G T A E K Y I K L I A N A K T V E G V W T Y K D E T L T F T V T E
BINARY_6301 T T Y K L I L N G K Q L K E E A I K E L V D A G T A E K Y I K L I A N A K T V E G V W T Y K D E T K K A T V T E
BINARY_6302 T T Y K L I L N G K Q L K E E A I K E L V D A G T A E K Y I K L I A N A K T V E G V W T Y K D E T K K F T V T E
BINARY_6303 T T Y K L I L N G K Q L K E E A I K E L V D A G T A E K Y I K L I A N A K T V E G V W T Y K D E T K T A T V T E
BINARY_6304 T T Y K L I L N G K Q L K E E A I K E L V D A G T A E K Y I K L I A N A K T V E G V W T Y K D E T K T F T V T E
BINARY_6305 T T Y K L I L N G K Q L K E E A I K E L V D A G T A E K Y I K L Y A N A K T V E G V W T L K D E I L K A T V T E
BINARY_6306 T T Y K L I L N G K Q L K E E A I K E L V D A G T A E K Y I K L Y A N A K T V E G V W T L K D E I L K F T V T E
BINARY_6307 T T Y K L I L N G K Q L K E E A I K E L V D A G T A E K Y I K L Y A N A K T V E G V W T L K D E I L T A T V T E
BINARY_6308 T T Y K L I L N G K Q L K E E A I K E L V D A G T A E K Y I K L Y A N A K T V E G V W T L K D E I L T F T V T E
BINARY_6309 T T Y K L I L N G K Q L K E E A I K E L V D A G T A E K Y I K L Y A N A K T V E G V W T L K D E I K K A T V T E
BINARY_6310 T T Y K L I L N G K Q L K E E A I K E L V D A G T A E K Y I K L Y A N A K T V E G V W T L K D E I K K F T V T E
BINARY_6311 T T Y K L I L N G K Q L K E E A I K E L V D A G T A E K Y I K L Y A N A K T V E G V W T L K D E I K T A T V T E
BINARY_6312 T T Y K L I L N G K Q L K E E A I K E L V D A G T A E K Y I K L Y A N A K T V E G V W T L K D E I K T F T V T E
BINARY_6313 T T Y K L I L N G K Q L K E E A I K E L V D A G T A E K Y I K L Y A N A K T V E G V W T L K D E T L K A T V T E
BINARY_6314 T T Y K L I L N G K Q L K E E A I K E L V D A G T A E K Y I K L Y A N A K T V E G V W T L K D E T L K F T V T E
BINARY_6315 T T Y K L I L N G K Q L K E E A I K E L V D A G T A E K Y I K L Y A N A K T V E G V W T L K D E T L T A T V T E
BINARY_6316 T T Y K L I L N G K Q L K E E A I K E L V D A G T A E K Y I K L Y A N A K T V E G V W T L K D E T L T F T V T E
BINARY_6317 T T Y K L I L N G K Q L K E E A I K E L V D A G T A E K Y I K L Y A N A K T V E G V W T L K D E T K K A T V T E
BINARY_6318 T T Y K L I L N G K Q L K E E A I K E L V D A G T A E K Y I K L Y A N A K T V E G V W T L K D E T K K F T V T E
BINARY_6319 T T Y K L I L N G K Q L K E E A I K E L V D A G T A E K Y I K L Y A N A K T V E G V W T L K D E T K T A T V T E
BINARY_6320 T T Y K L I L N G K Q L K E E A I K E L V D A G T A E K Y I K L Y A N A K T V E G V W T L K D E T K T F T V T E
BINARY_6321 T T Y K L I L N G K Q L K E E A I K E L V D A G T A E K Y I K L Y A N A K T V E G V W T Y K D E I L K A T V T E
BINARY_6322 T T Y K L I L N G K Q L K E E A I K E L V D A G T A E K Y I K L Y A N A K T V E G V W T Y K D E I L K F T V T E
BINARY_6323 T T Y K L I L N G K Q L K E E A I K E L V D A G T A E K Y I K L Y A N A K T V E G V W T Y K D E I L T A T V T E
BINARY_6324 T T Y K L I L N G K Q L K E E A I K E L V D A G T A E K Y I K L Y A N A K T V E G V W T Y K D E I L T F T V T E
BINARY_6325 T T Y K L I L N G K Q L K E E A I K E L V D A G T A E K Y I K L Y A N A K T V E G V W T Y K D E I K K A T V T E
BINARY_6326 T T Y K L I L N G K Q L K E E A I K E L V D A G T A E K Y I K L Y A N A K T V E G V W T Y K D E I K K F T V T E
BINARY_6327 T T Y K L I L N G K Q L K E E A I K E L V D A G T A E K Y I K L Y A N A K T V E G V W T Y K D E I K T A T V T E
BINARY_6328 T T Y K L I L N G K Q L K E E A I K E L V D A G T A E K Y I K L Y A N A K T V E G V W T Y K D E I K T F T V T E
BINARY_6329 T T Y K L I L N G K Q L K E E A I K E L V D A G T A E K Y I K L Y A N A K T V E G V W T Y K D E T L K A T V T E
BINARY_6330 T T Y K L I L N G K Q L K E E A I K E L V D A G T A E K Y I K L Y A N A K T V E G V W T Y K D E T L K F T V T E
BINARY_6331 T T Y K L I L N G K Q L K E E A I K E L V D A G T A E K Y I K L Y A N A K T V E G V W T Y K D E T L T A T V T E
BINARY_6332 T T Y K L I L N G K Q L K E E A I K E L V D A G T A E K Y I K L Y A N A K T V E G V W T Y K D E T L T F T V T E
BINARY_6333 T T Y K L I L N G K Q L K E E A I K E L V D A G T A E K Y I K L Y A N A K T V E G V W T Y K D E T K K A T V T E
BINARY_6334 T T Y K L I L N G K Q L K E E A I K E L V D A G T A E K Y I K L Y A N A K T V E G V W T Y K D E T K K F T V T E
BINARY_6335 T T Y K L I L N G K Q L K E E A I K E L V D A G T A E K Y I K L Y A N A K T V E G V W T Y K D E T K T A T V T E
BINARY_6336 T T Y K L I L N G K Q L K E E A I K E L V D A G T A E K Y I K L Y A N A K T V E G V W T Y K D E T K T F T V T E
BINARY_6337 T T Y K L I L N G K Q L K E E A I K E L V D A G T A E K Y F K L I A N A K T V E G V W T L K D E I L K A T V T E
BINARY_6338 T T Y K L I L N G K Q L K E E A I K E L V D A G T A E K Y F K L I A N A K T V E G V W T L K D E I L K F T V T E
BINARY_6339 T T Y K L I L N G K Q L K E E A I K E L V D A G T A E K Y F K L I A N A K T V E G V W T L K D E I L T A T V T E
BINARY_6340 T T Y K L I L N G K Q L K E E A I K E L V D A G T A E K Y F K L I A N A K T V E G V W T L K D E I L T F T V T E
BINARY_6341 T T Y K L I L N G K Q L K E E A I K E L V D A G T A E K Y F K L I A N A K T V E G V W T L K D E I K K A T V T E
BINARY_6342 T T Y K L I L N G K Q L K E E A I K E L V D A G T A E K Y F K L I A N A K T V E G V W T L K D E I K K F T V T E
BINARY_6343 T T Y K L I L N G K Q L K E E A I K E L V D A G T A E K Y F K L I A N A K T V E G V W T L K D E I K T A T V T E
BINARY_6344 T T Y K L I L N G K Q L K E E A I K E L V D A G T A E K Y F K L I A N A K T V E G V W T L K D E I K T F T V T E
BINARY_6345 T T Y K L I L N G K Q L K E E A I K E L V D A G T A E K Y F K L I A N A K T V E G V W T L K D E T L K A T V T E
BINARY_6346 T T Y K L I L N G K Q L K E E A I K E L V D A G T A E K Y F K L I A N A K T V E G V W T L K D E T L K F T V T E
BINARY_6347 T T Y K L I L N G K Q L K E E A I K E L V D A G T A E K Y F K L I A N A K T V E G V W T L K D E T L T A T V T E
BINARY_6348 T T Y K L I L N G K Q L K E E A I K E L V D A G T A E K Y F K L I A N A K T V E G V W T L K D E T L T F T V T E
BINARY_6349 T T Y K L I L N G K Q L K E E A I K E L V D A G T A E K Y F K L I A N A K T V E G V W T L K D E T K K A T V T E
BINARY_6350 T T Y K L I L N G K Q L K E E A I K E L V D A G T A E K Y F K L I A N A K T V E G V W T L K D E T K K F T V T E
BINARY_6351 T T Y K L I L N G K Q L K E E A I K E L V D A G T A E K Y F K L I A N A K T V E G V W T L K D E T K T A T V T E
BINARY_6352 T T Y K L I L N G K Q L K E E A I K E L V D A G T A E K Y F K L I A N A K T V E G V W T L K D E T K T F T V T E
BINARY_6353 T T Y K L I L N G K Q L K E E A I K E L V D A G T A E K Y F K L I A N A K T V E G V W T Y K D E I L K A T V T E
BINARY_6354 T T Y K L I L N G K Q L K E E A I K E L V D A G T A E K Y F K L I A N A K T V E G V W T Y K D E I L K F T V T E
BINARY_6355 T T Y K L I L N G K Q L K E E A I K E L V D A G T A E K Y F K L I A N A K T V E G V W T Y K D E I L T A T V T E
BINARY_6356 T T Y K L I L N G K Q L K E E A I K E L V D A G T A E K Y F K L I A N A K T V E G V W T Y K D E I L T F T V T E
BINARY_6357 T T Y K L I L N G K Q L K E E A I K E L V D A G T A E K Y F K L I A N A K T V E G V W T Y K D E I K K A T V T E
BINARY_6358 T T Y K L I L N G K Q L K E E A I K E L V D A G T A E K Y F K L I A N A K T V E G V W T Y K D E I K K F T V T E
BINARY_6359 T T Y K L I L N G K Q L K E E A I K E L V D A G T A E K Y F K L I A N A K T V E G V W T Y K D E I K T A T V T E
BINARY_6360 T T Y K L I L N G K Q L K E E A I K E L V D A G T A E K Y F K L I A N A K T V E G V W T Y K D E I K T F T V T E
BINARY_6361 T T Y K L I L N G K Q L K E E A I K E L V D A G T A E K Y F K L I A N A K T V E G V W T Y K D E T L K A T V T E
BINARY_6362 T T Y K L I L N G K Q L K E E A I K E L V D A G T A E K Y F K L I A N A K T V E G V W T Y K D E T L K F T V T E
BINARY_6363 T T Y K L I L N G K Q L K E E A I K E L V D A G T A E K Y F K L I A N A K T V E G V W T Y K D E T L T A T V T E
BINARY_6364 T T Y K L I L N G K Q L K E E A I K E L V D A G T A E K Y F K L I A N A K T V E G V W T Y K D E T L T F T V T E
BINARY_6365 T T Y K L I L N G K Q L K E E A I K E L V D A G T A E K Y F K L I A N A K T V E G V W T Y K D E T K K A T V T E
BINARY_6366 T T Y K L I L N G K Q L K E E A I K E L V D A G T A E K Y F K L I A N A K T V E G V W T Y K D E T K K F T V T E
BINARY_6367 T T Y K L I L N G K Q L K E E A I K E L V D A G T A E K Y F K L I A N A K T V E G V W T Y K D E T K T A T V T E
BINARY_6368 T T Y K L I L N G K Q L K E E A I K E L V D A G T A E K Y F K L I A N A K T V E G V W T Y K D E T K T F T V T E
BINARY_6369 T T Y K L I L N G K Q L K E E A I K E L V D A G T A E K Y F K L Y A N A K T V E G V W T L K D E I L K A T V T E
BINARY_6370 T T Y K L I L N G K Q L K E E A I K E L V D A G T A E K Y F K L Y A N A K T V E G V W T L K D E I L K F T V T E
BINARY_6371 T T Y K L I L N G K Q L K E E A I K E L V D A G T A E K Y F K L Y A N A K T V E G V W T L K D E I L T A T V T E
BINARY_6372 T T Y K L I L N G K Q L K E E A I K E L V D A G T A E K Y F K L Y A N A K T V E G V W T L K D E I L T F T V T E
BINARY_6373 T T Y K L I L N G K Q L K E E A I K E L V D A G T A E K Y F K L Y A N A K T V E G V W T L K D E I K K A T V T E
BINARY_6374 T T Y K L I L N G K Q L K E E A I K E L V D A G T A E K Y F K L Y A N A K T V E G V W T L K D E I K K F T V T E
BINARY_6375 T T Y K L I L N G K Q L K E E A I K E L V D A G T A E K Y F K L Y A N A K T V E G V W T L K D E I K T A T V T E
BINARY_6376 T T Y K L I L N G K Q L K E E A I K E L V D A G T A E K Y F K L Y A N A K T V E G V W T L K D E I K T F T V T E
BINARY_6377 T T Y K L I L N G K Q L K E E A I K E L V D A G T A E K Y F K L Y A N A K T V E G V W T L K D E T L K A T V T E
BINARY_6378 T T Y K L I L N G K Q L K E E A I K E L V D A G T A E K Y F K L Y A N A K T V E G V W T L K D E T L K F T V T E
BINARY_6379 T T Y K L I L N G K Q L K E E A I K E L V D A G T A E K Y F K L Y A N A K T V E G V W T L K D E T L T A T V T E
BINARY_6380 T T Y K L I L N G K Q L K E E A I K E L V D A G T A E K Y F K L Y A N A K T V E G V W T L K D E T L T F T V T E
BINARY_6381 T T Y K L I L N G K Q L K E E A I K E L V D A G T A E K Y F K L Y A N A K T V E G V W T L K D E T K K A T V T E

BINARY_6382 T T Y K L I L N G K Q L K E E A I K E L V D A G T A E K Y F K L Y A N A K T V E G V W T L K D E T K K F T V T E
BINARY_6383 T T Y K L I L N G K Q L K E E A I K E L V D A G T A E K Y F K L Y A N A K T V E G V W T L K D E T K T A T V T E
BINARY_6384 T T Y K L I L N G K Q L K E E A I K E L V D A G T A E K Y F K L Y A N A K T V E G V W T L K D E T K T F T V T E
BINARY_6385 T T Y K L I L N G K Q L K E E A I K E L V D A G T A E K Y F K L Y A N A K T V E G V W T Y K D E I L K A T V T E
BINARY_6386 T T Y K L I L N G K Q L K E E A I K E L V D A G T A E K Y F K L Y A N A K T V E G V W T Y K D E I L K F T V T E
BINARY_6387 T T Y K L I L N G K Q L K E E A I K E L V D A G T A E K Y F K L Y A N A K T V E G V W T Y K D E I L T A T V T E
BINARY_6388 T T Y K L I L N G K Q L K E E A I K E L V D A G T A E K Y F K L Y A N A K T V E G V W T Y K D E I L T F T V T E
BINARY_6389 T T Y K L I L N G K Q L K E E A I K E L V D A G T A E K Y F K L Y A N A K T V E G V W T Y K D E I K K A T V T E
BINARY_6390 T T Y K L I L N G K Q L K E E A I K E L V D A G T A E K Y F K L Y A N A K T V E G V W T Y K D E I K K F T V T E
BINARY_6391 T T Y K L I L N G K Q L K E E A I K E L V D A G T A E K Y F K L Y A N A K T V E G V W T Y K D E I K T A T V T E
BINARY_6392 T T Y K L I L N G K Q L K E E A I K E L V D A G T A E K Y F K L Y A N A K T V E G V W T Y K D E I K T F T V T E
BINARY_6393 T T Y K L I L N G K Q L K E E A I K E L V D A G T A E K Y F K L Y A N A K T V E G V W T Y K D E T L K A T V T E
BINARY_6394 T T Y K L I L N G K Q L K E E A I K E L V D A G T A E K Y F K L Y A N A K T V E G V W T Y K D E T L K F T V T E
BINARY_6395 T T Y K L I L N G K Q L K E E A I K E L V D A G T A E K Y F K L Y A N A K T V E G V W T Y K D E T L T A T V T E
BINARY_6396 T T Y K L I L N G K Q L K E E A I K E L V D A G T A E K Y F K L Y A N A K T V E G V W T Y K D E T L T F T V T E
BINARY_6397 T T Y K L I L N G K Q L K E E A I K E L V D A G T A E K Y F K L Y A N A K T V E G V W T Y K D E T K K A T V T E
BINARY_6398 T T Y K L I L N G K Q L K E E A I K E L V D A G T A E K Y F K L Y A N A K T V E G V W T Y K D E T K K F T V T E
BINARY_6399 T T Y K L I L N G K Q L K E E A I K E L V D A G T A E K Y F K L Y A N A K T V E G V W T Y K D E T K T A T V T E
BINARY_6400 T T Y K L I L N G K Q L K E E A I K E L V D A G T A E K Y F K L Y A N A K T V E G V W T Y K D E T K T F T V T E
BINARY_6401 T T Y K L I L N G K Q L K E E A I K E L V D A A I A E K Y I K L I A N A K T V E G V W T L K D E I L K A T V T E
BINARY_6402 T T Y K L I L N G K Q L K E E A I K E L V D A A I A E K Y I K L I A N A K T V E G V W T L K D E I L K F T V T E
BINARY_6403 T T Y K L I L N G K Q L K E E A I K E L V D A A I A E K Y I K L I A N A K T V E G V W T L K D E I L T A T V T E
BINARY_6404 T T Y K L I L N G K Q L K E E A I K E L V D A A I A E K Y I K L I A N A K T V E G V W T L K D E I L T F T V T E
BINARY_6405 T T Y K L I L N G K Q L K E E A I K E L V D A A I A E K Y I K L I A N A K T V E G V W T L K D E I K K A T V T E
BINARY_6406 T T Y K L I L N G K Q L K E E A I K E L V D A A I A E K Y I K L I A N A K T V E G V W T L K D E I K K F T V T E
BINARY_6407 T T Y K L I L N G K Q L K E E A I K E L V D A A I A E K Y I K L I A N A K T V E G V W T L K D E I K T A T V T E
BINARY_6408 T T Y K L I L N G K Q L K E E A I K E L V D A A I A E K Y I K L I A N A K T V E G V W T L K D E I K T F T V T E
BINARY_6409 T T Y K L I L N G K Q L K E E A I K E L V D A A I A E K Y I K L I A N A K T V E G V W T L K D E T L K A T V T E
BINARY_6410 T T Y K L I L N G K Q L K E E A I K E L V D A A I A E K Y I K L I A N A K T V E G V W T L K D E T L K F T V T E
BINARY_6411 T T Y K L I L N G K Q L K E E A I K E L V D A A I A E K Y I K L I A N A K T V E G V W T L K D E T L T A T V T E
BINARY_6412 T T Y K L I L N G K Q L K E E A I K E L V D A A I A E K Y I K L I A N A K T V E G V W T L K D E T L T F T V T E
BINARY_6413 T T Y K L I L N G K Q L K E E A I K E L V D A A I A E K Y I K L I A N A K T V E G V W T L K D E T K K A T V T E
BINARY_6414 T T Y K L I L N G K Q L K E E A I K E L V D A A I A E K Y I K L I A N A K T V E G V W T L K D E T K K F T V T E
BINARY_6415 T T Y K L I L N G K Q L K E E A I K E L V D A A I A E K Y I K L I A N A K T V E G V W T L K D E T K T A T V T E
BINARY_6416 T T Y K L I L N G K Q L K E E A I K E L V D A A I A E K Y I K L I A N A K T V E G V W T L K D E T K T F T V T E
BINARY_6417 T T Y K L I L N G K Q L K E E A I K E L V D A A I A E K Y I K L I A N A K T V E G V W T Y K D E I L K A T V T E
BINARY_6418 T T Y K L I L N G K Q L K E E A I K E L V D A A I A E K Y I K L I A N A K T V E G V W T Y K D E I L K F T V T E
BINARY_6419 T T Y K L I L N G K Q L K E E A I K E L V D A A I A E K Y I K L I A N A K T V E G V W T Y K D E I L T A T V T E
BINARY_6420 T T Y K L I L N G K Q L K E E A I K E L V D A A I A E K Y I K L I A N A K T V E G V W T Y K D E I L T F T V T E
BINARY_6421 T T Y K L I L N G K Q L K E E A I K E L V D A A I A E K Y I K L I A N A K T V E G V W T Y K D E I K K A T V T E
BINARY_6422 T T Y K L I L N G K Q L K E E A I K E L V D A A I A E K Y I K L I A N A K T V E G V W T Y K D E I K K F T V T E
BINARY_6423 T T Y K L I L N G K Q L K E E A I K E L V D A A I A E K Y I K L I A N A K T V E G V W T Y K D E I K T A T V T E
BINARY_6424 T T Y K L I L N G K Q L K E E A I K E L V D A A I A E K Y I K L I A N A K T V E G V W T Y K D E I K T F T V T E
BINARY_6425 T T Y K L I L N G K Q L K E E A I K E L V D A A I A E K Y I K L I A N A K T V E G V W T Y K D E T L K A T V T E
BINARY_6426 T T Y K L I L N G K Q L K E E A I K E L V D A A I A E K Y I K L I A N A K T V E G V W T Y K D E T L K F T V T E
BINARY_6427 T T Y K L I L N G K Q L K E E A I K E L V D A A I A E K Y I K L I A N A K T V E G V W T Y K D E T L T A T V T E
BINARY_6428 T T Y K L I L N G K Q L K E E A I K E L V D A A I A E K Y I K L I A N A K T V E G V W T Y K D E T L T F T V T E
BINARY_6429 T T Y K L I L N G K Q L K E E A I K E L V D A A I A E K Y I K L I A N A K T V E G V W T Y K D E T K K A T V T E
BINARY_6430 T T Y K L I L N G K Q L K E E A I K E L V D A A I A E K Y I K L I A N A K T V E G V W T Y K D E T K K F T V T E
BINARY_6431 T T Y K L I L N G K Q L K E E A I K E L V D A A I A E K Y I K L I A N A K T V E G V W T Y K D E T K T A T V T E
BINARY_6432 T T Y K L I L N G K Q L K E E A I K E L V D A A I A E K Y I K L I A N A K T V E G V W T Y K D E T K T F T V T E
BINARY_6433 T T Y K L I L N G K Q L K E E A I K E L V D A A I A E K Y I K L Y A N A K T V E G V W T L K D E I L K A T V T E
BINARY_6434 T T Y K L I L N G K Q L K E E A I K E L V D A A I A E K Y I K L Y A N A K T V E G V W T L K D E I L K F T V T E
BINARY_6435 T T Y K L I L N G K Q L K E E A I K E L V D A A I A E K Y I K L Y A N A K T V E G V W T L K D E I L T A T V T E
BINARY_6436 T T Y K L I L N G K Q L K E E A I K E L V D A A I A E K Y I K L Y A N A K T V E G V W T L K D E I L T F T V T E
BINARY_6437 T T Y K L I L N G K Q L K E E A I K E L V D A A I A E K Y I K L Y A N A K T V E G V W T L K D E I K K A T V T E
BINARY_6438 T T Y K L I L N G K Q L K E E A I K E L V D A A I A E K Y I K L Y A N A K T V E G V W T L K D E I K K F T V T E
BINARY_6439 T T Y K L I L N G K Q L K E E A I K E L V D A A I A E K Y I K L Y A N A K T V E G V W T L K D E I K T A T V T E
BINARY_6440 T T Y K L I L N G K Q L K E E A I K E L V D A A I A E K Y I K L Y A N A K T V E G V W T L K D E I K T F T V T E
BINARY_6441 T T Y K L I L N G K Q L K E E A I K E L V D A A I A E K Y I K L Y A N A K T V E G V W T L K D E T L K A T V T E
BINARY_6442 T T Y K L I L N G K Q L K E E A I K E L V D A A I A E K Y I K L Y A N A K T V E G V W T L K D E T L K F T V T E
BINARY_6443 T T Y K L I L N G K Q L K E E A I K E L V D A A I A E K Y I K L Y A N A K T V E G V W T L K D E T L T A T V T E
BINARY_6444 T T Y K L I L N G K Q L K E E A I K E L V D A A I A E K Y I K L Y A N A K T V E G V W T L K D E T L T F T V T E
BINARY_6445 T T Y K L I L N G K Q L K E E A I K E L V D A A I A E K Y I K L Y A N A K T V E G V W T L K D E T K K A T V T E
BINARY_6446 T T Y K L I L N G K Q L K E E A I K E L V D A A I A E K Y I K L Y A N A K T V E G V W T L K D E T K K F T V T E
BINARY_6447 T T Y K L I L N G K Q L K E E A I K E L V D A A I A E K Y I K L Y A N A K T V E G V W T L K D E T K T A T V T E
BINARY_6448 T T Y K L I L N G K Q L K E E A I K E L V D A A I A E K Y I K L Y A N A K T V E G V W T L K D E T K T F T V T E
BINARY_6449 T T Y K L I L N G K Q L K E E A I K E L V D A A I A E K Y I K L Y A N A K T V E G V W T Y K D E I L K A T V T E
BINARY_6450 T T Y K L I L N G K Q L K E E A I K E L V D A A I A E K Y I K L Y A N A K T V E G V W T Y K D E I L K F T V T E
BINARY_6451 T T Y K L I L N G K Q L K E E A I K E L V D A A I A E K Y I K L Y A N A K T V E G V W T Y K D E I L T A T V T E
BINARY_6452 T T Y K L I L N G K Q L K E E A I K E L V D A A I A E K Y I K L Y A N A K T V E G V W T Y K D E I L T F T V T E
BINARY_6453 T T Y K L I L N G K Q L K E E A I K E L V D A A I A E K Y I K L Y A N A K T V E G V W T Y K D E I K K A T V T E
BINARY_6454 T T Y K L I L N G K Q L K E E A I K E L V D A A I A E K Y I K L Y A N A K T V E G V W T Y K D E I K K F T V T E
BINARY_6455 T T Y K L I L N G K Q L K E E A I K E L V D A A I A E K Y I K L Y A N A K T V E G V W T Y K D E I K T A T V T E
BINARY_6456 T T Y K L I L N G K Q L K E E A I K E L V D A A I A E K Y I K L Y A N A K T V E G V W T Y K D E I K T F T V T E
BINARY_6457 T T Y K L I L N G K Q L K E E A I K E L V D A A I A E K Y I K L Y A N A K T V E G V W T Y K D E T L K A T V T E
BINARY_6458 T T Y K L I L N G K Q L K E E A I K E L V D A A I A E K Y I K L Y A N A K T V E G V W T Y K D E T L K F T V T E
BINARY_6459 T T Y K L I L N G K Q L K E E A I K E L V D A A I A E K Y I K L Y A N A K T V E G V W T Y K D E T L T A T V T E
BINARY_6460 T T Y K L I L N G K Q L K E E A I K E L V D A A I A E K Y I K L Y A N A K T V E G V W T Y K D E T L T F T V T E
BINARY_6461 T T Y K L I L N G K Q L K E E A I K E L V D A A I A E K Y I K L Y A N A K T V E G V W T Y K D E T K K A T V T E
BINARY_6462 T T Y K L I L N G K Q L K E E A I K E L V D A A I A E K Y I K L Y A N A K T V E G V W T Y K D E T K K F T V T E
BINARY_6463 T T Y K L I L N G K Q L K E E A I K E L V D A A I A E K Y I K L Y A N A K T V E G V W T Y K D E T K T A T V T E
BINARY_6464 T T Y K L I L N G K Q L K E E A I K E L V D A A I A E K Y I K L Y A N A K T V E G V W T Y K D E T K T F T V T E
BINARY_6465 T T Y K L I L N G K Q L K E E A I K E L V D A A I A E K Y F K L I A N A K T V E G V W T L K D E I L K A T V T E
BINARY_6466 T T Y K L I L N G K Q L K E E A I K E L V D A A I A E K Y F K L I A N A K T V E G V W T L K D E I L K F T V T E
BINARY_6467 T T Y K L I L N G K Q L K E E A I K E L V D A A I A E K Y F K L I A N A K T V E G V W T L K D E I L T A T V T E
BINARY_6468 T T Y K L I L N G K Q L K E E A I K E L V D A A I A E K Y F K L I A N A K T V E G V W T L K D E I L T F T V T E
BINARY_6469 T T Y K L I L N G K Q L K E E A I K E L V D A A I A E K Y F K L I A N A K T V E G V W T L K D E I K K A T V T E
BINARY_6470 T T Y K L I L N G K Q L K E E A I K E L V D A A I A E K Y F K L I A N A K T V E G V W T L K D E I K K F T V T E
BINARY_6471 T T Y K L I L N G K Q L K E E A I K E L V D A A I A E K Y F K L I A N A K T V E G V W T L K D E I K T A T V T E
BINARY_6472 T T Y K L I L N G K Q L K E E A I K E L V D A A I A E K Y F K L I A N A K T V E G V W T L K D E I K T F T V T E
BINARY_6473 T T Y K L I L N G K Q L K E E A I K E L V D A A I A E K Y F K L I A N A K T V E G V W T L K D E T L K A T V T E
BINARY_6474 T T Y K L I L N G K Q L K E E A I K E L V D A A I A E K Y F K L I A N A K T V E G V W T L K D E T L K F T V T E
BINARY_6475 T T Y K L I L N G K Q L K E E A I K E L V D A A I A E K Y F K L I A N A K T V E G V W T L K D E T L T A T V T E
BINARY_6476 T T Y K L I L N G K Q L K E E A I K E L V D A A I A E K Y F K L I A N A K T V E G V W T L K D E T L T F T V T E

BINARY_6477 T T Y K L I L N G K Q L K E E A I K E L V D A A I A E K Y F K L I A N A K T V E G V W T L K D E T K K A T V T E
BINARY_6478 T T Y K L I L N G K Q L K E E A I K E L V D A A I A E K Y F K L I A N A K T V E G V W T L K D E T K K F T V T E
BINARY_6479 T T Y K L I L N G K Q L K E E A I K E L V D A A I A E K Y F K L I A N A K T V E G V W T L K D E T K T A T V T E
BINARY_6480 T T Y K L I L N G K Q L K E E A I K E L V D A A I A E K Y F K L I A N A K T V E G V W T L K D E T K T F T V T E
BINARY_6481 T T Y K L I L N G K Q L K E E A I K E L V D A A I A E K Y F K L I A N A K T V E G V W T Y K D E I L K A T V T E
BINARY_6482 T T Y K L I L N G K Q L K E E A I K E L V D A A I A E K Y F K L I A N A K T V E G V W T Y K D E I L K F T V T E
BINARY_6483 T T Y K L I L N G K Q L K E E A I K E L V D A A I A E K Y F K L I A N A K T V E G V W T Y K D E I L T A T V T E
BINARY_6484 T T Y K L I L N G K Q L K E E A I K E L V D A A I A E K Y F K L I A N A K T V E G V W T Y K D E I L T F T V T E
BINARY_6485 T T Y K L I L N G K Q L K E E A I K E L V D A A I A E K Y F K L I A N A K T V E G V W T Y K D E I K K A T V T E
BINARY_6486 T T Y K L I L N G K Q L K E E A I K E L V D A A I A E K Y F K L I A N A K T V E G V W T Y K D E I K K F T V T E
BINARY_6487 T T Y K L I L N G K Q L K E E A I K E L V D A A I A E K Y F K L I A N A K T V E G V W T Y K D E I K T A T V T E
BINARY_6488 T T Y K L I L N G K Q L K E E A I K E L V D A A I A E K Y F K L I A N A K T V E G V W T Y K D E I K T F T V T E
BINARY_6489 T T Y K L I L N G K Q L K E E A I K E L V D A A I A E K Y F K L I A N A K T V E G V W T Y K D E T L K A T V T E
BINARY_6490 T T Y K L I L N G K Q L K E E A I K E L V D A A I A E K Y F K L I A N A K T V E G V W T Y K D E T L K F T V T E
BINARY_6491 T T Y K L I L N G K Q L K E E A I K E L V D A A I A E K Y F K L I A N A K T V E G V W T Y K D E T L T A T V T E
BINARY_6492 T T Y K L I L N G K Q L K E E A I K E L V D A A I A E K Y F K L I A N A K T V E G V W T Y K D E T L T F T V T E
BINARY_6493 T T Y K L I L N G K Q L K E E A I K E L V D A A I A E K Y F K L I A N A K T V E G V W T Y K D E T K K A T V T E
BINARY_6494 T T Y K L I L N G K Q L K E E A I K E L V D A A I A E K Y F K L I A N A K T V E G V W T Y K D E T K K F T V T E
BINARY_6495 T T Y K L I L N G K Q L K E E A I K E L V D A A I A E K Y F K L I A N A K T V E G V W T Y K D E T K T A T V T E
BINARY_6496 T T Y K L I L N G K Q L K E E A I K E L V D A A I A E K Y F K L I A N A K T V E G V W T Y K D E T K T F T V T E
BINARY_6497 T T Y K L I L N G K Q L K E E A I K E L V D A A I A E K Y F K L Y A N A K T V E G V W T L K D E I L K A T V T E
BINARY_6498 T T Y K L I L N G K Q L K E E A I K E L V D A A I A E K Y F K L Y A N A K T V E G V W T L K D E I L K F T V T E
BINARY_6499 T T Y K L I L N G K Q L K E E A I K E L V D A A I A E K Y F K L Y A N A K T V E G V W T L K D E I L T A T V T E
BINARY_6500 T T Y K L I L N G K Q L K E E A I K E L V D A A I A E K Y F K L Y A N A K T V E G V W T L K D E I L T F T V T E
BINARY_6501 T T Y K L I L N G K Q L K E E A I K E L V D A A I A E K Y F K L Y A N A K T V E G V W T L K D E I K K A T V T E
BINARY_6502 T T Y K L I L N G K Q L K E E A I K E L V D A A I A E K Y F K L Y A N A K T V E G V W T L K D E I K K F T V T E
BINARY_6503 T T Y K L I L N G K Q L K E E A I K E L V D A A I A E K Y F K L Y A N A K T V E G V W T L K D E I K T A T V T E
BINARY_6504 T T Y K L I L N G K Q L K E E A I K E L V D A A I A E K Y F K L Y A N A K T V E G V W T L K D E I K T F T V T E
BINARY_6505 T T Y K L I L N G K Q L K E E A I K E L V D A A I A E K Y F K L Y A N A K T V E G V W T L K D E T L K A T V T E
BINARY_6506 T T Y K L I L N G K Q L K E E A I K E L V D A A I A E K Y F K L Y A N A K T V E G V W T L K D E T L K F T V T E
BINARY_6507 T T Y K L I L N G K Q L K E E A I K E L V D A A I A E K Y F K L Y A N A K T V E G V W T L K D E T L T A T V T E
BINARY_6508 T T Y K L I L N G K Q L K E E A I K E L V D A A I A E K Y F K L Y A N A K T V E G V W T L K D E T L T F T V T E
BINARY_6509 T T Y K L I L N G K Q L K E E A I K E L V D A A I A E K Y F K L Y A N A K T V E G V W T L K D E T K K A T V T E
BINARY_6510 T T Y K L I L N G K Q L K E E A I K E L V D A A I A E K Y F K L Y A N A K T V E G V W T L K D E T K K F T V T E
BINARY_6511 T T Y K L I L N G K Q L K E E A I K E L V D A A I A E K Y F K L Y A N A K T V E G V W T L K D E T K T A T V T E
BINARY_6512 T T Y K L I L N G K Q L K E E A I K E L V D A A I A E K Y F K L Y A N A K T V E G V W T L K D E T K T F T V T E
BINARY_6513 T T Y K L I L N G K Q L K E E A I K E L V D A A I A E K Y F K L Y A N A K T V E G V W T Y K D E I L K A T V T E
BINARY_6514 T T Y K L I L N G K Q L K E E A I K E L V D A A I A E K Y F K L Y A N A K T V E G V W T Y K D E I L K F T V T E
BINARY_6515 T T Y K L I L N G K Q L K E E A I K E L V D A A I A E K Y F K L Y A N A K T V E G V W T Y K D E I L T A T V T E
BINARY_6516 T T Y K L I L N G K Q L K E E A I K E L V D A A I A E K Y F K L Y A N A K T V E G V W T Y K D E I L T F T V T E
BINARY_6517 T T Y K L I L N G K Q L K E E A I K E L V D A A I A E K Y F K L Y A N A K T V E G V W T Y K D E I K K A T V T E
BINARY_6518 T T Y K L I L N G K Q L K E E A I K E L V D A A I A E K Y F K L Y A N A K T V E G V W T Y K D E I K K F T V T E
BINARY_6519 T T Y K L I L N G K Q L K E E A I K E L V D A A I A E K Y F K L Y A N A K T V E G V W T Y K D E I K T A T V T E
BINARY_6520 T T Y K L I L N G K Q L K E E A I K E L V D A A I A E K Y F K L Y A N A K T V E G V W T Y K D E I K T F T V T E
BINARY_6521 T T Y K L I L N G K Q L K E E A I K E L V D A A I A E K Y F K L Y A N A K T V E G V W T Y K D E T L K A T V T E
BINARY_6522 T T Y K L I L N G K Q L K E E A I K E L V D A A I A E K Y F K L Y A N A K T V E G V W T Y K D E T L K F T V T E
BINARY_6523 T T Y K L I L N G K Q L K E E A I K E L V D A A I A E K Y F K L Y A N A K T V E G V W T Y K D E T L T A T V T E
BINARY_6524 T T Y K L I L N G K Q L K E E A I K E L V D A A I A E K Y F K L Y A N A K T V E G V W T Y K D E T L T F T V T E
BINARY_6525 T T Y K L I L N G K Q L K E E A I K E L V D A A I A E K Y F K L Y A N A K T V E G V W T Y K D E T K K A T V T E
BINARY_6526 T T Y K L I L N G K Q L K E E A I K E L V D A A I A E K Y F K L Y A N A K T V E G V W T Y K D E T K K F T V T E
BINARY_6527 T T Y K L I L N G K Q L K E E A I K E L V D A A I A E K Y F K L Y A N A K T V E G V W T Y K D E T K T A T V T E
BINARY_6528 T T Y K L I L N G K Q L K E E A I K E L V D A A I A E K Y F K L Y A N A K T V E G V W T Y K D E T K T F T V T E
BINARY_6529 T T Y K L I L N G K Q L K E E A I K E L V D A A T A E K Y I K L I A N A K T V E G V W T L K D E I L K A T V T E
BINARY_6530 T T Y K L I L N G K Q L K E E A I K E L V D A A T A E K Y I K L I A N A K T V E G V W T L K D E I L K F T V T E
BINARY_6531 T T Y K L I L N G K Q L K E E A I K E L V D A A T A E K Y I K L I A N A K T V E G V W T L K D E I L T A T V T E
BINARY_6532 T T Y K L I L N G K Q L K E E A I K E L V D A A T A E K Y I K L I A N A K T V E G V W T L K D E I L T F T V T E
BINARY_6533 T T Y K L I L N G K Q L K E E A I K E L V D A A T A E K Y I K L I A N A K T V E G V W T L K D E I K K A T V T E
BINARY_6534 T T Y K L I L N G K Q L K E E A I K E L V D A A T A E K Y I K L I A N A K T V E G V W T L K D E I K K F T V T E
BINARY_6535 T T Y K L I L N G K Q L K E E A I K E L V D A A T A E K Y I K L I A N A K T V E G V W T L K D E I K T A T V T E
BINARY_6536 T T Y K L I L N G K Q L K E E A I K E L V D A A T A E K Y I K L I A N A K T V E G V W T L K D E I K T F T V T E
BINARY_6537 T T Y K L I L N G K Q L K E E A I K E L V D A A T A E K Y I K L I A N A K T V E G V W T L K D E T L K A T V T E
BINARY_6538 T T Y K L I L N G K Q L K E E A I K E L V D A A T A E K Y I K L I A N A K T V E G V W T L K D E T L K F T V T E
BINARY_6539 T T Y K L I L N G K Q L K E E A I K E L V D A A T A E K Y I K L I A N A K T V E G V W T L K D E T L T A T V T E
BINARY_6540 T T Y K L I L N G K Q L K E E A I K E L V D A A T A E K Y I K L I A N A K T V E G V W T L K D E T L T F T V T E
BINARY_6541 T T Y K L I L N G K Q L K E E A I K E L V D A A T A E K Y I K L I A N A K T V E G V W T L K D E T K K A T V T E
BINARY_6542 T T Y K L I L N G K Q L K E E A I K E L V D A A T A E K Y I K L I A N A K T V E G V W T L K D E T K K F T V T E
BINARY_6543 T T Y K L I L N G K Q L K E E A I K E L V D A A T A E K Y I K L I A N A K T V E G V W T L K D E T K T A T V T E
BINARY_6544 T T Y K L I L N G K Q L K E E A I K E L V D A A T A E K Y I K L I A N A K T V E G V W T L K D E T K T F T V T E
BINARY_6545 T T Y K L I L N G K Q L K E E A I K E L V D A A T A E K Y I K L I A N A K T V E G V W T Y K D E I L K A T V T E
BINARY_6546 T T Y K L I L N G K Q L K E E A I K E L V D A A T A E K Y I K L I A N A K T V E G V W T Y K D E I L K F T V T E
BINARY_6547 T T Y K L I L N G K Q L K E E A I K E L V D A A T A E K Y I K L I A N A K T V E G V W T Y K D E I L T A T V T E
BINARY_6548 T T Y K L I L N G K Q L K E E A I K E L V D A A T A E K Y I K L I A N A K T V E G V W T Y K D E I L T F T V T E
BINARY_6549 T T Y K L I L N G K Q L K E E A I K E L V D A A T A E K Y I K L I A N A K T V E G V W T Y K D E I K K A T V T E
BINARY_6550 T T Y K L I L N G K Q L K E E A I K E L V D A A T A E K Y I K L I A N A K T V E G V W T Y K D E I K K F T V T E
BINARY_6551 T T Y K L I L N G K Q L K E E A I K E L V D A A T A E K Y I K L I A N A K T V E G V W T Y K D E I K T A T V T E
BINARY_6552 T T Y K L I L N G K Q L K E E A I K E L V D A A T A E K Y I K L I A N A K T V E G V W T Y K D E I K T F T V T E
BINARY_6553 T T Y K L I L N G K Q L K E E A I K E L V D A A T A E K Y I K L I A N A K T V E G V W T Y K D E T L K A T V T E
BINARY_6554 T T Y K L I L N G K Q L K E E A I K E L V D A A T A E K Y I K L I A N A K T V E G V W T Y K D E T L K F T V T E
BINARY_6555 T T Y K L I L N G K Q L K E E A I K E L V D A A T A E K Y I K L I A N A K T V E G V W T Y K D E T L T A T V T E
BINARY_6556 T T Y K L I L N G K Q L K E E A I K E L V D A A T A E K Y I K L I A N A K T V E G V W T Y K D E T L T F T V T E
BINARY_6557 T T Y K L I L N G K Q L K E E A I K E L V D A A T A E K Y I K L I A N A K T V E G V W T Y K D E T K K A T V T E
BINARY_6558 T T Y K L I L N G K Q L K E E A I K E L V D A A T A E K Y I K L I A N A K T V E G V W T Y K D E T K K F T V T E
BINARY_6559 T T Y K L I L N G K Q L K E E A I K E L V D A A T A E K Y I K L I A N A K T V E G V W T Y K D E T K T A T V T E
BINARY_6560 T T Y K L I L N G K Q L K E E A I K E L V D A A T A E K Y I K L I A N A K T V E G V W T Y K D E T K T F T V T E
BINARY_6561 T T Y K L I L N G K Q L K E E A I K E L V D A A T A E K Y I K L Y A N A K T V E G V W T L K D E I L K A T V T E
BINARY_6562 T T Y K L I L N G K Q L K E E A I K E L V D A A T A E K Y I K L Y A N A K T V E G V W T L K D E I L K F T V T E
BINARY_6563 T T Y K L I L N G K Q L K E E A I K E L V D A A T A E K Y I K L Y A N A K T V E G V W T L K D E I L T A T V T E
BINARY_6564 T T Y K L I L N G K Q L K E E A I K E L V D A A T A E K Y I K L Y A N A K T V E G V W T L K D E I L T F T V T E
BINARY_6565 T T Y K L I L N G K Q L K E E A I K E L V D A A T A E K Y I K L Y A N A K T V E G V W T L K D E I K K A T V T E
BINARY_6566 T T Y K L I L N G K Q L K E E A I K E L V D A A T A E K Y I K L Y A N A K T V E G V W T L K D E I K K F T V T E
BINARY_6567 T T Y K L I L N G K Q L K E E A I K E L V D A A T A E K Y I K L Y A N A K T V E G V W T L K D E I K T A T V T E
BINARY_6568 T T Y K L I L N G K Q L K E E A I K E L V D A A T A E K Y I K L Y A N A K T V E G V W T L K D E I K T F T V T E
BINARY_6569 T T Y K L I L N G K Q L K E E A I K E L V D A A T A E K Y I K L Y A N A K T V E G V W T L K D E T L K A T V T E
BINARY_6570 T T Y K L I L N G K Q L K E E A I K E L V D A A T A E K Y I K L Y A N A K T V E G V W T L K D E T L K F T V T E
BINARY_6571 T T Y K L I L N G K Q L K E E A I K E L V D A A T A E K Y I K L Y A N A K T V E G V W T L K D E T L T A T V T E

BINARY_6572 T T Y K L I L N G K Q L K E E A I K E L V D A A T A E K Y I K L Y A N A K T V E G V W T L K D E T L T F T V T E
BINARY_6573 T T Y K L I L N G K Q L K E E A I K E L V D A A T A E K Y I K L Y A N A K T V E G V W T L K D E T K K A T V T E
BINARY_6574 T T Y K L I L N G K Q L K E E A I K E L V D A A T A E K Y I K L Y A N A K T V E G V W T L K D E T K K F T V T E
BINARY_6575 T T Y K L I L N G K Q L K E E A I K E L V D A A T A E K Y I K L Y A N A K T V E G V W T L K D E T K T A T V T E
BINARY_6576 T T Y K L I L N G K Q L K E E A I K E L V D A A T A E K Y I K L Y A N A K T V E G V W T L K D E T K T F T V T E
BINARY_6577 T T Y K L I L N G K Q L K E E A I K E L V D A A T A E K Y I K L Y A N A K T V E G V W T Y K D E I L K A T V T E
BINARY_6578 T T Y K L I L N G K Q L K E E A I K E L V D A A T A E K Y I K L Y A N A K T V E G V W T Y K D E I L K F T V T E
BINARY_6579 T T Y K L I L N G K Q L K E E A I K E L V D A A T A E K Y I K L Y A N A K T V E G V W T Y K D E I L T A T V T E
BINARY_6580 T T Y K L I L N G K Q L K E E A I K E L V D A A T A E K Y I K L Y A N A K T V E G V W T Y K D E I L T F T V T E
BINARY_6581 T T Y K L I L N G K Q L K E E A I K E L V D A A T A E K Y I K L Y A N A K T V E G V W T Y K D E I K K A T V T E
BINARY_6582 T T Y K L I L N G K Q L K E E A I K E L V D A A T A E K Y I K L Y A N A K T V E G V W T Y K D E I K K F T V T E
BINARY_6583 T T Y K L I L N G K Q L K E E A I K E L V D A A T A E K Y I K L Y A N A K T V E G V W T Y K D E I K T A T V T E
BINARY_6584 T T Y K L I L N G K Q L K E E A I K E L V D A A T A E K Y I K L Y A N A K T V E G V W T Y K D E I K T F T V T E
BINARY_6585 T T Y K L I L N G K Q L K E E A I K E L V D A A T A E K Y I K L Y A N A K T V E G V W T Y K D E T L K A T V T E
BINARY_6586 T T Y K L I L N G K Q L K E E A I K E L V D A A T A E K Y I K L Y A N A K T V E G V W T Y K D E T L K F T V T E
BINARY_6587 T T Y K L I L N G K Q L K E E A I K E L V D A A T A E K Y I K L Y A N A K T V E G V W T Y K D E T L T A T V T E
BINARY_6588 T T Y K L I L N G K Q L K E E A I K E L V D A A T A E K Y I K L Y A N A K T V E G V W T Y K D E T L T F T V T E
BINARY_6589 T T Y K L I L N G K Q L K E E A I K E L V D A A T A E K Y I K L Y A N A K T V E G V W T Y K D E T K K A T V T E
BINARY_6590 T T Y K L I L N G K Q L K E E A I K E L V D A A T A E K Y I K L Y A N A K T V E G V W T Y K D E T K K F T V T E
BINARY_6591 T T Y K L I L N G K Q L K E E A I K E L V D A A T A E K Y I K L Y A N A K T V E G V W T Y K D E T K T A T V T E
BINARY_6592 T T Y K L I L N G K Q L K E E A I K E L V D A A T A E K Y I K L Y A N A K T V E G V W T Y K D E T K T F T V T E
BINARY_6593 T T Y K L I L N G K Q L K E E A I K E L V D A A T A E K Y F K L I A N A K T V E G V W T L K D E I L K A T V T E
BINARY_6594 T T Y K L I L N G K Q L K E E A I K E L V D A A T A E K Y F K L I A N A K T V E G V W T L K D E I L K F T V T E
BINARY_6595 T T Y K L I L N G K Q L K E E A I K E L V D A A T A E K Y F K L I A N A K T V E G V W T L K D E I L T A T V T E
BINARY_6596 T T Y K L I L N G K Q L K E E A I K E L V D A A T A E K Y F K L I A N A K T V E G V W T L K D E I L T F T V T E
BINARY_6597 T T Y K L I L N G K Q L K E E A I K E L V D A A T A E K Y F K L I A N A K T V E G V W T L K D E I K K A T V T E
BINARY_6598 T T Y K L I L N G K Q L K E E A I K E L V D A A T A E K Y F K L I A N A K T V E G V W T L K D E I K K F T V T E
BINARY_6599 T T Y K L I L N G K Q L K E E A I K E L V D A A T A E K Y F K L I A N A K T V E G V W T L K D E I K T A T V T E
BINARY_6600 T T Y K L I L N G K Q L K E E A I K E L V D A A T A E K Y F K L I A N A K T V E G V W T L K D E I K T F T V T E
BINARY_6601 T T Y K L I L N G K Q L K E E A I K E L V D A A T A E K Y F K L I A N A K T V E G V W T L K D E T L K A T V T E
BINARY_6602 T T Y K L I L N G K Q L K E E A I K E L V D A A T A E K Y F K L I A N A K T V E G V W T L K D E T L K F T V T E
BINARY_6603 T T Y K L I L N G K Q L K E E A I K E L V D A A T A E K Y F K L I A N A K T V E G V W T L K D E T L T A T V T E
BINARY_6604 T T Y K L I L N G K Q L K E E A I K E L V D A A T A E K Y F K L I A N A K T V E G V W T L K D E T L T F T V T E
BINARY_6605 T T Y K L I L N G K Q L K E E A I K E L V D A A T A E K Y F K L I A N A K T V E G V W T L K D E T K K A T V T E
BINARY_6606 T T Y K L I L N G K Q L K E E A I K E L V D A A T A E K Y F K L I A N A K T V E G V W T L K D E T K K F T V T E
BINARY_6607 T T Y K L I L N G K Q L K E E A I K E L V D A A T A E K Y F K L I A N A K T V E G V W T L K D E T K T A T V T E
BINARY_6608 T T Y K L I L N G K Q L K E E A I K E L V D A A T A E K Y F K L I A N A K T V E G V W T L K D E T K T F T V T E
BINARY_6609 T T Y K L I L N G K Q L K E E A I K E L V D A A T A E K Y F K L I A N A K T V E G V W T Y K D E I L K A T V T E
BINARY_6610 T T Y K L I L N G K Q L K E E A I K E L V D A A T A E K Y F K L I A N A K T V E G V W T Y K D E I L K F T V T E
BINARY_6611 T T Y K L I L N G K Q L K E E A I K E L V D A A T A E K Y F K L I A N A K T V E G V W T Y K D E I L T A T V T E
BINARY_6612 T T Y K L I L N G K Q L K E E A I K E L V D A A T A E K Y F K L I A N A K T V E G V W T Y K D E I L T F T V T E
BINARY_6613 T T Y K L I L N G K Q L K E E A I K E L V D A A T A E K Y F K L I A N A K T V E G V W T Y K D E I K K A T V T E
BINARY_6614 T T Y K L I L N G K Q L K E E A I K E L V D A A T A E K Y F K L I A N A K T V E G V W T Y K D E I K K F T V T E
BINARY_6615 T T Y K L I L N G K Q L K E E A I K E L V D A A T A E K Y F K L I A N A K T V E G V W T Y K D E I K T A T V T E
BINARY_6616 T T Y K L I L N G K Q L K E E A I K E L V D A A T A E K Y F K L I A N A K T V E G V W T Y K D E I K T F T V T E
BINARY_6617 T T Y K L I L N G K Q L K E E A I K E L V D A A T A E K Y F K L I A N A K T V E G V W T Y K D E T L K A T V T E
BINARY_6618 T T Y K L I L N G K Q L K E E A I K E L V D A A T A E K Y F K L I A N A K T V E G V W T Y K D E T L K F T V T E
BINARY_6619 T T Y K L I L N G K Q L K E E A I K E L V D A A T A E K Y F K L I A N A K T V E G V W T Y K D E T L T A T V T E
BINARY_6620 T T Y K L I L N G K Q L K E E A I K E L V D A A T A E K Y F K L I A N A K T V E G V W T Y K D E T L T F T V T E
BINARY_6621 T T Y K L I L N G K Q L K E E A I K E L V D A A T A E K Y F K L I A N A K T V E G V W T Y K D E T K K A T V T E
BINARY_6622 T T Y K L I L N G K Q L K E E A I K E L V D A A T A E K Y F K L I A N A K T V E G V W T Y K D E T K K F T V T E
BINARY_6623 T T Y K L I L N G K Q L K E E A I K E L V D A A T A E K Y F K L I A N A K T V E G V W T Y K D E T K T A T V T E
BINARY_6624 T T Y K L I L N G K Q L K E E A I K E L V D A A T A E K Y F K L I A N A K T V E G V W T Y K D E T K T F T V T E
BINARY_6625 T T Y K L I L N G K Q L K E E A I K E L V D A A T A E K Y F K L Y A N A K T V E G V W T L K D E I L K A T V T E
BINARY_6626 T T Y K L I L N G K Q L K E E A I K E L V D A A T A E K Y F K L Y A N A K T V E G V W T L K D E I L K F T V T E
BINARY_6627 T T Y K L I L N G K Q L K E E A I K E L V D A A T A E K Y F K L Y A N A K T V E G V W T L K D E I L T A T V T E
BINARY_6628 T T Y K L I L N G K Q L K E E A I K E L V D A A T A E K Y F K L Y A N A K T V E G V W T L K D E I L T F T V T E
BINARY_6629 T T Y K L I L N G K Q L K E E A I K E L V D A A T A E K Y F K L Y A N A K T V E G V W T L K D E I K K A T V T E
BINARY_6630 T T Y K L I L N G K Q L K E E A I K E L V D A A T A E K Y F K L Y A N A K T V E G V W T L K D E I K K F T V T E
BINARY_6631 T T Y K L I L N G K Q L K E E A I K E L V D A A T A E K Y F K L Y A N A K T V E G V W T L K D E I K T A T V T E
BINARY_6632 T T Y K L I L N G K Q L K E E A I K E L V D A A T A E K Y F K L Y A N A K T V E G V W T L K D E I K T F T V T E
BINARY_6633 T T Y K L I L N G K Q L K E E A I K E L V D A A T A E K Y F K L Y A N A K T V E G V W T L K D E T L K A T V T E
BINARY_6634 T T Y K L I L N G K Q L K E E A I K E L V D A A T A E K Y F K L Y A N A K T V E G V W T L K D E T L K F T V T E
BINARY_6635 T T Y K L I L N G K Q L K E E A I K E L V D A A T A E K Y F K L Y A N A K T V E G V W T L K D E T L T A T V T E
BINARY_6636 T T Y K L I L N G K Q L K E E A I K E L V D A A T A E K Y F K L Y A N A K T V E G V W T L K D E T L T F T V T E
BINARY_6637 T T Y K L I L N G K Q L K E E A I K E L V D A A T A E K Y F K L Y A N A K T V E G V W T L K D E T K K A T V T E
BINARY_6638 T T Y K L I L N G K Q L K E E A I K E L V D A A T A E K Y F K L Y A N A K T V E G V W T L K D E T K K F T V T E
BINARY_6639 T T Y K L I L N G K Q L K E E A I K E L V D A A T A E K Y F K L Y A N A K T V E G V W T L K D E T K T A T V T E
BINARY_6640 T T Y K L I L N G K Q L K E E A I K E L V D A A T A E K Y F K L Y A N A K T V E G V W T L K D E T K T F T V T E
BINARY_6641 T T Y K L I L N G K Q L K E E A I K E L V D A A T A E K Y F K L Y A N A K T V E G V W T Y K D E I L K A T V T E
BINARY_6642 T T Y K L I L N G K Q L K E E A I K E L V D A A T A E K Y F K L Y A N A K T V E G V W T Y K D E I L K F T V T E
BINARY_6643 T T Y K L I L N G K Q L K E E A I K E L V D A A T A E K Y F K L Y A N A K T V E G V W T Y K D E I L T A T V T E
BINARY_6644 T T Y K L I L N G K Q L K E E A I K E L V D A A T A E K Y F K L Y A N A K T V E G V W T Y K D E I L T F T V T E
BINARY_6645 T T Y K L I L N G K Q L K E E A I K E L V D A A T A E K Y F K L Y A N A K T V E G V W T Y K D E I K K A T V T E
BINARY_6646 T T Y K L I L N G K Q L K E E A I K E L V D A A T A E K Y F K L Y A N A K T V E G V W T Y K D E I K K F T V T E
BINARY_6647 T T Y K L I L N G K Q L K E E A I K E L V D A A T A E K Y F K L Y A N A K T V E G V W T Y K D E I K T A T V T E
BINARY_6648 T T Y K L I L N G K Q L K E E A I K E L V D A A T A E K Y F K L Y A N A K T V E G V W T Y K D E I K T F T V T E
BINARY_6649 T T Y K L I L N G K Q L K E E A I K E L V D A A T A E K Y F K L Y A N A K T V E G V W T Y K D E T L K A T V T E
BINARY_6650 T T Y K L I L N G K Q L K E E A I K E L V D A A T A E K Y F K L Y A N A K T V E G V W T Y K D E T L K F T V T E
BINARY_6651 T T Y K L I L N G K Q L K E E A I K E L V D A A T A E K Y F K L Y A N A K T V E G V W T Y K D E T L T A T V T E
BINARY_6652 T T Y K L I L N G K Q L K E E A I K E L V D A A T A E K Y F K L Y A N A K T V E G V W T Y K D E T L T F T V T E
BINARY_6653 T T Y K L I L N G K Q L K E E A I K E L V D A A T A E K Y F K L Y A N A K T V E G V W T Y K D E T K K A T V T E
BINARY_6654 T T Y K L I L N G K Q L K E E A I K E L V D A A T A E K Y F K L Y A N A K T V E G V W T Y K D E T K K F T V T E
BINARY_6655 T T Y K L I L N G K Q L K E E A I K E L V D A A T A E K Y F K L Y A N A K T V E G V W T Y K D E T K T A T V T E
BINARY_6656 T T Y K L I L N G K Q L K E E A I K E L V D A A T A E K Y F K L Y A N A K T V E G V W T Y K D E T K T F T V T E
BINARY_6657 T T Y K L I L N G K Q L K E E A I K E A V D A G I A E K Y I K L I A N A K T V E G V W T L K D E I L K A T V T E
BINARY_6658 T T Y K L I L N G K Q L K E E A I K E A V D A G I A E K Y I K L I A N A K T V E G V W T L K D E I L K F T V T E
BINARY_6659 T T Y K L I L N G K Q L K E E A I K E A V D A G I A E K Y I K L I A N A K T V E G V W T L K D E I L T A T V T E
BINARY_6660 T T Y K L I L N G K Q L K E E A I K E A V D A G I A E K Y I K L I A N A K T V E G V W T L K D E I L T F T V T E
BINARY_6661 T T Y K L I L N G K Q L K E E A I K E A V D A G I A E K Y I K L I A N A K T V E G V W T L K D E I K K A T V T E
BINARY_6662 T T Y K L I L N G K Q L K E E A I K E A V D A G I A E K Y I K L I A N A K T V E G V W T L K D E I K K F T V T E
BINARY_6663 T T Y K L I L N G K Q L K E E A I K E A V D A G I A E K Y I K L I A N A K T V E G V W T L K D E I K T A T V T E
BINARY_6664 T T Y K L I L N G K Q L K E E A I K E A V D A G I A E K Y I K L I A N A K T V E G V W T L K D E I K T F T V T E
BINARY_6665 T T Y K L I L N G K Q L K E E A I K E A V D A G I A E K Y I K L I A N A K T V E G V W T L K D E T L K A T V T E
BINARY_6666 T T Y K L I L N G K Q L K E E A I K E A V D A G I A E K Y I K L I A N A K T V E G V W T L K D E T L K F T V T E

BINARY_6667 T T Y K L I L N G K Q L K E E A I K E A V D A G I A E K Y I K L I A N A K T V E G V W T L K D E T L T A T V T E
BINARY_6668 T T Y K L I L N G K Q L K E E A I K E A V D A G I A E K Y I K L I A N A K T V E G V W T L K D E T L T F T V T E
BINARY_6669 T T Y K L I L N G K Q L K E E A I K E A V D A G I A E K Y I K L I A N A K T V E G V W T L K D E T K K A T V T E
BINARY_6670 T T Y K L I L N G K Q L K E E A I K E A V D A G I A E K Y I K L I A N A K T V E G V W T L K D E T K K F T V T E
BINARY_6671 T T Y K L I L N G K Q L K E E A I K E A V D A G I A E K Y I K L I A N A K T V E G V W T L K D E T K T A T V T E
BINARY_6672 T T Y K L I L N G K Q L K E E A I K E A V D A G I A E K Y I K L I A N A K T V E G V W T L K D E T K T F T V T E
BINARY_6673 T T Y K L I L N G K Q L K E E A I K E A V D A G I A E K Y I K L I A N A K T V E G V W T Y K D E I L K A T V T E
BINARY_6674 T T Y K L I L N G K Q L K E E A I K E A V D A G I A E K Y I K L I A N A K T V E G V W T Y K D E I L K F T V T E
BINARY_6675 T T Y K L I L N G K Q L K E E A I K E A V D A G I A E K Y I K L I A N A K T V E G V W T Y K D E I L T A T V T E
BINARY_6676 T T Y K L I L N G K Q L K E E A I K E A V D A G I A E K Y I K L I A N A K T V E G V W T Y K D E I L T F T V T E
BINARY_6677 T T Y K L I L N G K Q L K E E A I K E A V D A G I A E K Y I K L I A N A K T V E G V W T Y K D E I K K A T V T E
BINARY_6678 T T Y K L I L N G K Q L K E E A I K E A V D A G I A E K Y I K L I A N A K T V E G V W T Y K D E I K K F T V T E
BINARY_6679 T T Y K L I L N G K Q L K E E A I K E A V D A G I A E K Y I K L I A N A K T V E G V W T Y K D E I K T A T V T E
BINARY_6680 T T Y K L I L N G K Q L K E E A I K E A V D A G I A E K Y I K L I A N A K T V E G V W T Y K D E I K T F T V T E
BINARY_6681 T T Y K L I L N G K Q L K E E A I K E A V D A G I A E K Y I K L I A N A K T V E G V W T Y K D E T L K A T V T E
BINARY_6682 T T Y K L I L N G K Q L K E E A I K E A V D A G I A E K Y I K L I A N A K T V E G V W T Y K D E T L K F T V T E
BINARY_6683 T T Y K L I L N G K Q L K E E A I K E A V D A G I A E K Y I K L I A N A K T V E G V W T Y K D E T L T A T V T E
BINARY_6684 T T Y K L I L N G K Q L K E E A I K E A V D A G I A E K Y I K L I A N A K T V E G V W T Y K D E T L T F T V T E
BINARY_6685 T T Y K L I L N G K Q L K E E A I K E A V D A G I A E K Y I K L I A N A K T V E G V W T Y K D E T K K A T V T E
BINARY_6686 T T Y K L I L N G K Q L K E E A I K E A V D A G I A E K Y I K L I A N A K T V E G V W T Y K D E T K K F T V T E
BINARY_6687 T T Y K L I L N G K Q L K E E A I K E A V D A G I A E K Y I K L I A N A K T V E G V W T Y K D E T K T A T V T E
BINARY_6688 T T Y K L I L N G K Q L K E E A I K E A V D A G I A E K Y I K L I A N A K T V E G V W T Y K D E T K T F T V T E
BINARY_6689 T T Y K L I L N G K Q L K E E A I K E A V D A G I A E K Y I K L Y A N A K T V E G V W T L K D E I L K A T V T E
BINARY_6690 T T Y K L I L N G K Q L K E E A I K E A V D A G I A E K Y I K L Y A N A K T V E G V W T L K D E I L K F T V T E
BINARY_6691 T T Y K L I L N G K Q L K E E A I K E A V D A G I A E K Y I K L Y A N A K T V E G V W T L K D E I L T A T V T E
BINARY_6692 T T Y K L I L N G K Q L K E E A I K E A V D A G I A E K Y I K L Y A N A K T V E G V W T L K D E I L T F T V T E
BINARY_6693 T T Y K L I L N G K Q L K E E A I K E A V D A G I A E K Y I K L Y A N A K T V E G V W T L K D E I K K A T V T E
BINARY_6694 T T Y K L I L N G K Q L K E E A I K E A V D A G I A E K Y I K L Y A N A K T V E G V W T L K D E I K K F T V T E
BINARY_6695 T T Y K L I L N G K Q L K E E A I K E A V D A G I A E K Y I K L Y A N A K T V E G V W T L K D E I K T A T V T E
BINARY_6696 T T Y K L I L N G K Q L K E E A I K E A V D A G I A E K Y I K L Y A N A K T V E G V W T L K D E I K T F T V T E
BINARY_6697 T T Y K L I L N G K Q L K E E A I K E A V D A G I A E K Y I K L Y A N A K T V E G V W T L K D E T L K A T V T E
BINARY_6698 T T Y K L I L N G K Q L K E E A I K E A V D A G I A E K Y I K L Y A N A K T V E G V W T L K D E T L K F T V T E
BINARY_6699 T T Y K L I L N G K Q L K E E A I K E A V D A G I A E K Y I K L Y A N A K T V E G V W T L K D E T L T A T V T E
BINARY_6700 T T Y K L I L N G K Q L K E E A I K E A V D A G I A E K Y I K L Y A N A K T V E G V W T L K D E T L T F T V T E
BINARY_6701 T T Y K L I L N G K Q L K E E A I K E A V D A G I A E K Y I K L Y A N A K T V E G V W T L K D E T K K A T V T E
BINARY_6702 T T Y K L I L N G K Q L K E E A I K E A V D A G I A E K Y I K L Y A N A K T V E G V W T L K D E T K K F T V T E
BINARY_6703 T T Y K L I L N G K Q L K E E A I K E A V D A G I A E K Y I K L Y A N A K T V E G V W T L K D E T K T A T V T E
BINARY_6704 T T Y K L I L N G K Q L K E E A I K E A V D A G I A E K Y I K L Y A N A K T V E G V W T L K D E T K T F T V T E
BINARY_6705 T T Y K L I L N G K Q L K E E A I K E A V D A G I A E K Y I K L Y A N A K T V E G V W T Y K D E I L K A T V T E
BINARY_6706 T T Y K L I L N G K Q L K E E A I K E A V D A G I A E K Y I K L Y A N A K T V E G V W T Y K D E I L K F T V T E
BINARY_6707 T T Y K L I L N G K Q L K E E A I K E A V D A G I A E K Y I K L Y A N A K T V E G V W T Y K D E I L T A T V T E
BINARY_6708 T T Y K L I L N G K Q L K E E A I K E A V D A G I A E K Y I K L Y A N A K T V E G V W T Y K D E I L T F T V T E
BINARY_6709 T T Y K L I L N G K Q L K E E A I K E A V D A G I A E K Y I K L Y A N A K T V E G V W T Y K D E I K K A T V T E
BINARY_6710 T T Y K L I L N G K Q L K E E A I K E A V D A G I A E K Y I K L Y A N A K T V E G V W T Y K D E I K K F T V T E
BINARY_6711 T T Y K L I L N G K Q L K E E A I K E A V D A G I A E K Y I K L Y A N A K T V E G V W T Y K D E I K T A T V T E
BINARY_6712 T T Y K L I L N G K Q L K E E A I K E A V D A G I A E K Y I K L Y A N A K T V E G V W T Y K D E I K T F T V T E
BINARY_6713 T T Y K L I L N G K Q L K E E A I K E A V D A G I A E K Y I K L Y A N A K T V E G V W T Y K D E T L K A T V T E
BINARY_6714 T T Y K L I L N G K Q L K E E A I K E A V D A G I A E K Y I K L Y A N A K T V E G V W T Y K D E T L K F T V T E
BINARY_6715 T T Y K L I L N G K Q L K E E A I K E A V D A G I A E K Y I K L Y A N A K T V E G V W T Y K D E T L T A T V T E
BINARY_6716 T T Y K L I L N G K Q L K E E A I K E A V D A G I A E K Y I K L Y A N A K T V E G V W T Y K D E T L T F T V T E
BINARY_6717 T T Y K L I L N G K Q L K E E A I K E A V D A G I A E K Y I K L Y A N A K T V E G V W T Y K D E T K K A T V T E
BINARY_6718 T T Y K L I L N G K Q L K E E A I K E A V D A G I A E K Y I K L Y A N A K T V E G V W T Y K D E T K K F T V T E
BINARY_6719 T T Y K L I L N G K Q L K E E A I K E A V D A G I A E K Y I K L Y A N A K T V E G V W T Y K D E T K T A T V T E
BINARY_6720 T T Y K L I L N G K Q L K E E A I K E A V D A G I A E K Y I K L Y A N A K T V E G V W T Y K D E T K T F T V T E
BINARY_6721 T T Y K L I L N G K Q L K E E A I K E A V D A G I A E K Y F K L I A N A K T V E G V W T L K D E I L K A T V T E
BINARY_6722 T T Y K L I L N G K Q L K E E A I K E A V D A G I A E K Y F K L I A N A K T V E G V W T L K D E I L K F T V T E
BINARY_6723 T T Y K L I L N G K Q L K E E A I K E A V D A G I A E K Y F K L I A N A K T V E G V W T L K D E I L T A T V T E
BINARY_6724 T T Y K L I L N G K Q L K E E A I K E A V D A G I A E K Y F K L I A N A K T V E G V W T L K D E I L T F T V T E
BINARY_6725 T T Y K L I L N G K Q L K E E A I K E A V D A G I A E K Y F K L I A N A K T V E G V W T L K D E I K K A T V T E
BINARY_6726 T T Y K L I L N G K Q L K E E A I K E A V D A G I A E K Y F K L I A N A K T V E G V W T L K D E I K K F T V T E
BINARY_6727 T T Y K L I L N G K Q L K E E A I K E A V D A G I A E K Y F K L I A N A K T V E G V W T L K D E I K T A T V T E
BINARY_6728 T T Y K L I L N G K Q L K E E A I K E A V D A G I A E K Y F K L I A N A K T V E G V W T L K D E I K T F T V T E
BINARY_6729 T T Y K L I L N G K Q L K E E A I K E A V D A G I A E K Y F K L I A N A K T V E G V W T L K D E T L K A T V T E
BINARY_6730 T T Y K L I L N G K Q L K E E A I K E A V D A G I A E K Y F K L I A N A K T V E G V W T L K D E T L K F T V T E
BINARY_6731 T T Y K L I L N G K Q L K E E A I K E A V D A G I A E K Y F K L I A N A K T V E G V W T L K D E T L T A T V T E
BINARY_6732 T T Y K L I L N G K Q L K E E A I K E A V D A G I A E K Y F K L I A N A K T V E G V W T L K D E T L T F T V T E
BINARY_6733 T T Y K L I L N G K Q L K E E A I K E A V D A G I A E K Y F K L I A N A K T V E G V W T L K D E T K K A T V T E
BINARY_6734 T T Y K L I L N G K Q L K E E A I K E A V D A G I A E K Y F K L I A N A K T V E G V W T L K D E T K K F T V T E
BINARY_6735 T T Y K L I L N G K Q L K E E A I K E A V D A G I A E K Y F K L I A N A K T V E G V W T L K D E T K T A T V T E
BINARY_6736 T T Y K L I L N G K Q L K E E A I K E A V D A G I A E K Y F K L I A N A K T V E G V W T L K D E T K T F T V T E
BINARY_6737 T T Y K L I L N G K Q L K E E A I K E A V D A G I A E K Y F K L I A N A K T V E G V W T Y K D E I L K A T V T E
BINARY_6738 T T Y K L I L N G K Q L K E E A I K E A V D A G I A E K Y F K L I A N A K T V E G V W T Y K D E I L K F T V T E
BINARY_6739 T T Y K L I L N G K Q L K E E A I K E A V D A G I A E K Y F K L I A N A K T V E G V W T Y K D E I L T A T V T E
BINARY_6740 T T Y K L I L N G K Q L K E E A I K E A V D A G I A E K Y F K L I A N A K T V E G V W T Y K D E I L T F T V T E
BINARY_6741 T T Y K L I L N G K Q L K E E A I K E A V D A G I A E K Y F K L I A N A K T V E G V W T Y K D E I K K A T V T E
BINARY_6742 T T Y K L I L N G K Q L K E E A I K E A V D A G I A E K Y F K L I A N A K T V E G V W T Y K D E I K K F T V T E
BINARY_6743 T T Y K L I L N G K Q L K E E A I K E A V D A G I A E K Y F K L I A N A K T V E G V W T Y K D E I K T A T V T E
BINARY_6744 T T Y K L I L N G K Q L K E E A I K E A V D A G I A E K Y F K L I A N A K T V E G V W T Y K D E I K T F T V T E
BINARY_6745 T T Y K L I L N G K Q L K E E A I K E A V D A G I A E K Y F K L I A N A K T V E G V W T Y K D E T L K A T V T E
BINARY_6746 T T Y K L I L N G K Q L K E E A I K E A V D A G I A E K Y F K L I A N A K T V E G V W T Y K D E T L K F T V T E
BINARY_6747 T T Y K L I L N G K Q L K E E A I K E A V D A G I A E K Y F K L I A N A K T V E G V W T Y K D E T L T A T V T E
BINARY_6748 T T Y K L I L N G K Q L K E E A I K E A V D A G I A E K Y F K L I A N A K T V E G V W T Y K D E T L T F T V T E
BINARY_6749 T T Y K L I L N G K Q L K E E A I K E A V D A G I A E K Y F K L I A N A K T V E G V W T Y K D E T K K A T V T E
BINARY_6750 T T Y K L I L N G K Q L K E E A I K E A V D A G I A E K Y F K L I A N A K T V E G V W T Y K D E T K K F T V T E
BINARY_6751 T T Y K L I L N G K Q L K E E A I K E A V D A G I A E K Y F K L I A N A K T V E G V W T Y K D E T K T A T V T E
BINARY_6752 T T Y K L I L N G K Q L K E E A I K E A V D A G I A E K Y F K L I A N A K T V E G V W T Y K D E T K T F T V T E
BINARY_6753 T T Y K L I L N G K Q L K E E A I K E A V D A G I A E K Y F K L Y A N A K T V E G V W T L K D E I L K A T V T E
BINARY_6754 T T Y K L I L N G K Q L K E E A I K E A V D A G I A E K Y F K L Y A N A K T V E G V W T L K D E I L K F T V T E
BINARY_6755 T T Y K L I L N G K Q L K E E A I K E A V D A G I A E K Y F K L Y A N A K T V E G V W T L K D E I L T A T V T E
BINARY_6756 T T Y K L I L N G K Q L K E E A I K E A V D A G I A E K Y F K L Y A N A K T V E G V W T L K D E I L T F T V T E
BINARY_6757 T T Y K L I L N G K Q L K E E A I K E A V D A G I A E K Y F K L Y A N A K T V E G V W T L K D E I K K A T V T E
BINARY_6758 T T Y K L I L N G K Q L K E E A I K E A V D A G I A E K Y F K L Y A N A K T V E G V W T L K D E I K K F T V T E
BINARY_6759 T T Y K L I L N G K Q L K E E A I K E A V D A G I A E K Y F K L Y A N A K T V E G V W T L K D E I K T A T V T E
BINARY_6760 T T Y K L I L N G K Q L K E E A I K E A V D A G I A E K Y F K L Y A N A K T V E G V W T L K D E I K T F T V T E
BINARY_6761 T T Y K L I L N G K Q L K E E A I K E A V D A G I A E K Y F K L Y A N A K T V E G V W T L K D E T L K A T V T E

BINARY_6762 T T Y K L I L N G K Q L K E E A I K E A V D A G I A E K Y F K L Y A N A K T V E G V W T L K D E T L K F T V T E
BINARY_6763 T T Y K L I L N G K Q L K E E A I K E A V D A G I A E K Y F K L Y A N A K T V E G V W T L K D E T L T A T V T E
BINARY_6764 T T Y K L I L N G K Q L K E E A I K E A V D A G I A E K Y F K L Y A N A K T V E G V W T L K D E T L T F T V T E
BINARY_6765 T T Y K L I L N G K Q L K E E A I K E A V D A G I A E K Y F K L Y A N A K T V E G V W T L K D E T K K A T V T E
BINARY_6766 T T Y K L I L N G K Q L K E E A I K E A V D A G I A E K Y F K L Y A N A K T V E G V W T L K D E T K K F T V T E
BINARY_6767 T T Y K L I L N G K Q L K E E A I K E A V D A G I A E K Y F K L Y A N A K T V E G V W T L K D E T K T A T V T E
BINARY_6768 T T Y K L I L N G K Q L K E E A I K E A V D A G I A E K Y F K L Y A N A K T V E G V W T L K D E T K T F T V T E
BINARY_6769 T T Y K L I L N G K Q L K E E A I K E A V D A G I A E K Y F K L Y A N A K T V E G V W T Y K D E I L K A T V T E
BINARY_6770 T T Y K L I L N G K Q L K E E A I K E A V D A G I A E K Y F K L Y A N A K T V E G V W T Y K D E I L K F T V T E
BINARY_6771 T T Y K L I L N G K Q L K E E A I K E A V D A G I A E K Y F K L Y A N A K T V E G V W T Y K D E I L T A T V T E
BINARY_6772 T T Y K L I L N G K Q L K E E A I K E A V D A G I A E K Y F K L Y A N A K T V E G V W T Y K D E I L T F T V T E
BINARY_6773 T T Y K L I L N G K Q L K E E A I K E A V D A G I A E K Y F K L Y A N A K T V E G V W T Y K D E I K K A T V T E
BINARY_6774 T T Y K L I L N G K Q L K E E A I K E A V D A G I A E K Y F K L Y A N A K T V E G V W T Y K D E I K K F T V T E
BINARY_6775 T T Y K L I L N G K Q L K E E A I K E A V D A G I A E K Y F K L Y A N A K T V E G V W T Y K D E I K T A T V T E
BINARY_6776 T T Y K L I L N G K Q L K E E A I K E A V D A G I A E K Y F K L Y A N A K T V E G V W T Y K D E I K T F T V T E
BINARY_6777 T T Y K L I L N G K Q L K E E A I K E A V D A G I A E K Y F K L Y A N A K T V E G V W T Y K D E T L K A T V T E
BINARY_6778 T T Y K L I L N G K Q L K E E A I K E A V D A G I A E K Y F K L Y A N A K T V E G V W T Y K D E T L K F T V T E
BINARY_6779 T T Y K L I L N G K Q L K E E A I K E A V D A G I A E K Y F K L Y A N A K T V E G V W T Y K D E T L T A T V T E
BINARY_6780 T T Y K L I L N G K Q L K E E A I K E A V D A G I A E K Y F K L Y A N A K T V E G V W T Y K D E T L T F T V T E
BINARY_6781 T T Y K L I L N G K Q L K E E A I K E A V D A G I A E K Y F K L Y A N A K T V E G V W T Y K D E T K K A T V T E
BINARY_6782 T T Y K L I L N G K Q L K E E A I K E A V D A G I A E K Y F K L Y A N A K T V E G V W T Y K D E T K K F T V T E
BINARY_6783 T T Y K L I L N G K Q L K E E A I K E A V D A G I A E K Y F K L Y A N A K T V E G V W T Y K D E T K T A T V T E
BINARY_6784 T T Y K L I L N G K Q L K E E A I K E A V D A G I A E K Y F K L Y A N A K T V E G V W T Y K D E T K T F T V T E
BINARY_6785 T T Y K L I L N G K Q L K E E A I K E A V D A G T A E K Y I K L I A N A K T V E G V W T L K D E I L K A T V T E
BINARY_6786 T T Y K L I L N G K Q L K E E A I K E A V D A G T A E K Y I K L I A N A K T V E G V W T L K D E I L K F T V T E
BINARY_6787 T T Y K L I L N G K Q L K E E A I K E A V D A G T A E K Y I K L I A N A K T V E G V W T L K D E I L T A T V T E
BINARY_6788 T T Y K L I L N G K Q L K E E A I K E A V D A G T A E K Y I K L I A N A K T V E G V W T L K D E I L T F T V T E
BINARY_6789 T T Y K L I L N G K Q L K E E A I K E A V D A G T A E K Y I K L I A N A K T V E G V W T L K D E I K K A T V T E
BINARY_6790 T T Y K L I L N G K Q L K E E A I K E A V D A G T A E K Y I K L I A N A K T V E G V W T L K D E I K K F T V T E
BINARY_6791 T T Y K L I L N G K Q L K E E A I K E A V D A G T A E K Y I K L I A N A K T V E G V W T L K D E I K T A T V T E
BINARY_6792 T T Y K L I L N G K Q L K E E A I K E A V D A G T A E K Y I K L I A N A K T V E G V W T L K D E I K T F T V T E
BINARY_6793 T T Y K L I L N G K Q L K E E A I K E A V D A G T A E K Y I K L I A N A K T V E G V W T L K D E T L K A T V T E
BINARY_6794 T T Y K L I L N G K Q L K E E A I K E A V D A G T A E K Y I K L I A N A K T V E G V W T L K D E T L K F T V T E
BINARY_6795 T T Y K L I L N G K Q L K E E A I K E A V D A G T A E K Y I K L I A N A K T V E G V W T L K D E T L T A T V T E
BINARY_6796 T T Y K L I L N G K Q L K E E A I K E A V D A G T A E K Y I K L I A N A K T V E G V W T L K D E T L T F T V T E
BINARY_6797 T T Y K L I L N G K Q L K E E A I K E A V D A G T A E K Y I K L I A N A K T V E G V W T L K D E T K K A T V T E
BINARY_6798 T T Y K L I L N G K Q L K E E A I K E A V D A G T A E K Y I K L I A N A K T V E G V W T L K D E T K K F T V T E
BINARY_6799 T T Y K L I L N G K Q L K E E A I K E A V D A G T A E K Y I K L I A N A K T V E G V W T L K D E T K T A T V T E
BINARY_6800 T T Y K L I L N G K Q L K E E A I K E A V D A G T A E K Y I K L I A N A K T V E G V W T L K D E T K T F T V T E
BINARY_6801 T T Y K L I L N G K Q L K E E A I K E A V D A G T A E K Y I K L I A N A K T V E G V W T Y K D E I L K A T V T E
BINARY_6802 T T Y K L I L N G K Q L K E E A I K E A V D A G T A E K Y I K L I A N A K T V E G V W T Y K D E I L K F T V T E
BINARY_6803 T T Y K L I L N G K Q L K E E A I K E A V D A G T A E K Y I K L I A N A K T V E G V W T Y K D E I L T A T V T E
BINARY_6804 T T Y K L I L N G K Q L K E E A I K E A V D A G T A E K Y I K L I A N A K T V E G V W T Y K D E I L T F T V T E
BINARY_6805 T T Y K L I L N G K Q L K E E A I K E A V D A G T A E K Y I K L I A N A K T V E G V W T Y K D E I K K A T V T E
BINARY_6806 T T Y K L I L N G K Q L K E E A I K E A V D A G T A E K Y I K L I A N A K T V E G V W T Y K D E I K K F T V T E
BINARY_6807 T T Y K L I L N G K Q L K E E A I K E A V D A G T A E K Y I K L I A N A K T V E G V W T Y K D E I K T A T V T E
BINARY_6808 T T Y K L I L N G K Q L K E E A I K E A V D A G T A E K Y I K L I A N A K T V E G V W T Y K D E I K T F T V T E
BINARY_6809 T T Y K L I L N G K Q L K E E A I K E A V D A G T A E K Y I K L I A N A K T V E G V W T Y K D E T L K A T V T E
BINARY_6810 T T Y K L I L N G K Q L K E E A I K E A V D A G T A E K Y I K L I A N A K T V E G V W T Y K D E T L K F T V T E
BINARY_6811 T T Y K L I L N G K Q L K E E A I K E A V D A G T A E K Y I K L I A N A K T V E G V W T Y K D E T L T A T V T E
BINARY_6812 T T Y K L I L N G K Q L K E E A I K E A V D A G T A E K Y I K L I A N A K T V E G V W T Y K D E T L T F T V T E
BINARY_6813 T T Y K L I L N G K Q L K E E A I K E A V D A G T A E K Y I K L I A N A K T V E G V W T Y K D E T K K A T V T E
BINARY_6814 T T Y K L I L N G K Q L K E E A I K E A V D A G T A E K Y I K L I A N A K T V E G V W T Y K D E T K K F T V T E
BINARY_6815 T T Y K L I L N G K Q L K E E A I K E A V D A G T A E K Y I K L I A N A K T V E G V W T Y K D E T K T A T V T E
BINARY_6816 T T Y K L I L N G K Q L K E E A I K E A V D A G T A E K Y I K L I A N A K T V E G V W T Y K D E T K T F T V T E
BINARY_6817 T T Y K L I L N G K Q L K E E A I K E A V D A G T A E K Y I K L Y A N A K T V E G V W T L K D E I L K A T V T E
BINARY_6818 T T Y K L I L N G K Q L K E E A I K E A V D A G T A E K Y I K L Y A N A K T V E G V W T L K D E I L K F T V T E
BINARY_6819 T T Y K L I L N G K Q L K E E A I K E A V D A G T A E K Y I K L Y A N A K T V E G V W T L K D E I L T A T V T E
BINARY_6820 T T Y K L I L N G K Q L K E E A I K E A V D A G T A E K Y I K L Y A N A K T V E G V W T L K D E I L T F T V T E
BINARY_6821 T T Y K L I L N G K Q L K E E A I K E A V D A G T A E K Y I K L Y A N A K T V E G V W T L K D E I K K A T V T E
BINARY_6822 T T Y K L I L N G K Q L K E E A I K E A V D A G T A E K Y I K L Y A N A K T V E G V W T L K D E I K K F T V T E
BINARY_6823 T T Y K L I L N G K Q L K E E A I K E A V D A G T A E K Y I K L Y A N A K T V E G V W T L K D E I K T A T V T E
BINARY_6824 T T Y K L I L N G K Q L K E E A I K E A V D A G T A E K Y I K L Y A N A K T V E G V W T L K D E I K T F T V T E
BINARY_6825 T T Y K L I L N G K Q L K E E A I K E A V D A G T A E K Y I K L Y A N A K T V E G V W T L K D E T L K A T V T E
BINARY_6826 T T Y K L I L N G K Q L K E E A I K E A V D A G T A E K Y I K L Y A N A K T V E G V W T L K D E T L K F T V T E
BINARY_6827 T T Y K L I L N G K Q L K E E A I K E A V D A G T A E K Y I K L Y A N A K T V E G V W T L K D E T L T A T V T E
BINARY_6828 T T Y K L I L N G K Q L K E E A I K E A V D A G T A E K Y I K L Y A N A K T V E G V W T L K D E T L T F T V T E
BINARY_6829 T T Y K L I L N G K Q L K E E A I K E A V D A G T A E K Y I K L Y A N A K T V E G V W T L K D E T K K A T V T E
BINARY_6830 T T Y K L I L N G K Q L K E E A I K E A V D A G T A E K Y I K L Y A N A K T V E G V W T L K D E T K K F T V T E
BINARY_6831 T T Y K L I L N G K Q L K E E A I K E A V D A G T A E K Y I K L Y A N A K T V E G V W T L K D E T K T A T V T E
BINARY_6832 T T Y K L I L N G K Q L K E E A I K E A V D A G T A E K Y I K L Y A N A K T V E G V W T L K D E T K T F T V T E
BINARY_6833 T T Y K L I L N G K Q L K E E A I K E A V D A G T A E K Y I K L Y A N A K T V E G V W T Y K D E I L K A T V T E
BINARY_6834 T T Y K L I L N G K Q L K E E A I K E A V D A G T A E K Y I K L Y A N A K T V E G V W T Y K D E I L K F T V T E
BINARY_6835 T T Y K L I L N G K Q L K E E A I K E A V D A G T A E K Y I K L Y A N A K T V E G V W T Y K D E I L T A T V T E
BINARY_6836 T T Y K L I L N G K Q L K E E A I K E A V D A G T A E K Y I K L Y A N A K T V E G V W T Y K D E I L T F T V T E
BINARY_6837 T T Y K L I L N G K Q L K E E A I K E A V D A G T A E K Y I K L Y A N A K T V E G V W T Y K D E I K K A T V T E
BINARY_6838 T T Y K L I L N G K Q L K E E A I K E A V D A G T A E K Y I K L Y A N A K T V E G V W T Y K D E I K K F T V T E
BINARY_6839 T T Y K L I L N G K Q L K E E A I K E A V D A G T A E K Y I K L Y A N A K T V E G V W T Y K D E I K T A T V T E
BINARY_6840 T T Y K L I L N G K Q L K E E A I K E A V D A G T A E K Y I K L Y A N A K T V E G V W T Y K D E I K T F T V T E
BINARY_6841 T T Y K L I L N G K Q L K E E A I K E A V D A G T A E K Y I K L Y A N A K T V E G V W T Y K D E T L K A T V T E
BINARY_6842 T T Y K L I L N G K Q L K E E A I K E A V D A G T A E K Y I K L Y A N A K T V E G V W T Y K D E T L K F T V T E
BINARY_6843 T T Y K L I L N G K Q L K E E A I K E A V D A G T A E K Y I K L Y A N A K T V E G V W T Y K D E T L T A T V T E
BINARY_6844 T T Y K L I L N G K Q L K E E A I K E A V D A G T A E K Y I K L Y A N A K T V E G V W T Y K D E T L T F T V T E
BINARY_6845 T T Y K L I L N G K Q L K E E A I K E A V D A G T A E K Y I K L Y A N A K T V E G V W T Y K D E T K K A T V T E
BINARY_6846 T T Y K L I L N G K Q L K E E A I K E A V D A G T A E K Y I K L Y A N A K T V E G V W T Y K D E T K K F T V T E
BINARY_6847 T T Y K L I L N G K Q L K E E A I K E A V D A G T A E K Y I K L Y A N A K T V E G V W T Y K D E T K T A T V T E
BINARY_6848 T T Y K L I L N G K Q L K E E A I K E A V D A G T A E K Y I K L Y A N A K T V E G V W T Y K D E T K T F T V T E
BINARY_6849 T T Y K L I L N G K Q L K E E A I K E A V D A G T A E K Y F K L I A N A K T V E G V W T L K D E I L K A T V T E
BINARY_6850 T T Y K L I L N G K Q L K E E A I K E A V D A G T A E K Y F K L I A N A K T V E G V W T L K D E I L K F T V T E
BINARY_6851 T T Y K L I L N G K Q L K E E A I K E A V D A G T A E K Y F K L I A N A K T V E G V W T L K D E I L T A T V T E
BINARY_6852 T T Y K L I L N G K Q L K E E A I K E A V D A G T A E K Y F K L I A N A K T V E G V W T L K D E I L T F T V T E
BINARY_6853 T T Y K L I L N G K Q L K E E A I K E A V D A G T A E K Y F K L I A N A K T V E G V W T L K D E I K K A T V T E
BINARY_6854 T T Y K L I L N G K Q L K E E A I K E A V D A G T A E K Y F K L I A N A K T V E G V W T L K D E I K K F T V T E
BINARY_6855 T T Y K L I L N G K Q L K E E A I K E A V D A G T A E K Y F K L I A N A K T V E G V W T L K D E I K T A T V T E
BINARY_6856 T T Y K L I L N G K Q L K E E A I K E A V D A G T A E K Y F K L I A N A K T V E G V W T L K D E I K T F T V T E

BINARY_6857 T T Y K L I L N G K Q L K E E A I K E A V D A G T A E K Y F K L I A N A K T V E G V W T L K D E T L K A T V T E
BINARY_6858 T T Y K L I L N G K Q L K E E A I K E A V D A G T A E K Y F K L I A N A K T V E G V W T L K D E T L K F T V T E
BINARY_6859 T T Y K L I L N G K Q L K E E A I K E A V D A G T A E K Y F K L I A N A K T V E G V W T L K D E T L T A T V T E
BINARY_6860 T T Y K L I L N G K Q L K E E A I K E A V D A G T A E K Y F K L I A N A K T V E G V W T L K D E T L T F T V T E
BINARY_6861 T T Y K L I L N G K Q L K E E A I K E A V D A G T A E K Y F K L I A N A K T V E G V W T L K D E T K K A T V T E
BINARY_6862 T T Y K L I L N G K Q L K E E A I K E A V D A G T A E K Y F K L I A N A K T V E G V W T L K D E T K K F T V T E
BINARY_6863 T T Y K L I L N G K Q L K E E A I K E A V D A G T A E K Y F K L I A N A K T V E G V W T L K D E T K T A T V T E
BINARY_6864 T T Y K L I L N G K Q L K E E A I K E A V D A G T A E K Y F K L I A N A K T V E G V W T L K D E T K T F T V T E
BINARY_6865 T T Y K L I L N G K Q L K E E A I K E A V D A G T A E K Y F K L I A N A K T V E G V W T Y K D E I L K A T V T E
BINARY_6866 T T Y K L I L N G K Q L K E E A I K E A V D A G T A E K Y F K L I A N A K T V E G V W T Y K D E I L K F T V T E
BINARY_6867 T T Y K L I L N G K Q L K E E A I K E A V D A G T A E K Y F K L I A N A K T V E G V W T Y K D E I L T A T V T E
BINARY_6868 T T Y K L I L N G K Q L K E E A I K E A V D A G T A E K Y F K L I A N A K T V E G V W T Y K D E I L T F T V T E
BINARY_6869 T T Y K L I L N G K Q L K E E A I K E A V D A G T A E K Y F K L I A N A K T V E G V W T Y K D E I K K A T V T E
BINARY_6870 T T Y K L I L N G K Q L K E E A I K E A V D A G T A E K Y F K L I A N A K T V E G V W T Y K D E I K K F T V T E
BINARY_6871 T T Y K L I L N G K Q L K E E A I K E A V D A G T A E K Y F K L I A N A K T V E G V W T Y K D E I K T A T V T E
BINARY_6872 T T Y K L I L N G K Q L K E E A I K E A V D A G T A E K Y F K L I A N A K T V E G V W T Y K D E I K T F T V T E
BINARY_6873 T T Y K L I L N G K Q L K E E A I K E A V D A G T A E K Y F K L I A N A K T V E G V W T Y K D E T L K A T V T E
BINARY_6874 T T Y K L I L N G K Q L K E E A I K E A V D A G T A E K Y F K L I A N A K T V E G V W T Y K D E T L K F T V T E
BINARY_6875 T T Y K L I L N G K Q L K E E A I K E A V D A G T A E K Y F K L I A N A K T V E G V W T Y K D E T L T A T V T E
BINARY_6876 T T Y K L I L N G K Q L K E E A I K E A V D A G T A E K Y F K L I A N A K T V E G V W T Y K D E T L T F T V T E
BINARY_6877 T T Y K L I L N G K Q L K E E A I K E A V D A G T A E K Y F K L I A N A K T V E G V W T Y K D E T K K A T V T E
BINARY_6878 T T Y K L I L N G K Q L K E E A I K E A V D A G T A E K Y F K L I A N A K T V E G V W T Y K D E T K K F T V T E
BINARY_6879 T T Y K L I L N G K Q L K E E A I K E A V D A G T A E K Y F K L I A N A K T V E G V W T Y K D E T K T A T V T E
BINARY_6880 T T Y K L I L N G K Q L K E E A I K E A V D A G T A E K Y F K L I A N A K T V E G V W T Y K D E T K T F T V T E
BINARY_6881 T T Y K L I L N G K Q L K E E A I K E A V D A G T A E K Y F K L Y A N A K T V E G V W T L K D E I L K A T V T E
BINARY_6882 T T Y K L I L N G K Q L K E E A I K E A V D A G T A E K Y F K L Y A N A K T V E G V W T L K D E I L K F T V T E
BINARY_6883 T T Y K L I L N G K Q L K E E A I K E A V D A G T A E K Y F K L Y A N A K T V E G V W T L K D E I L T A T V T E
BINARY_6884 T T Y K L I L N G K Q L K E E A I K E A V D A G T A E K Y F K L Y A N A K T V E G V W T L K D E I L T F T V T E
BINARY_6885 T T Y K L I L N G K Q L K E E A I K E A V D A G T A E K Y F K L Y A N A K T V E G V W T L K D E I K K A T V T E
BINARY_6886 T T Y K L I L N G K Q L K E E A I K E A V D A G T A E K Y F K L Y A N A K T V E G V W T L K D E I K K F T V T E
BINARY_6887 T T Y K L I L N G K Q L K E E A I K E A V D A G T A E K Y F K L Y A N A K T V E G V W T L K D E I K T A T V T E
BINARY_6888 T T Y K L I L N G K Q L K E E A I K E A V D A G T A E K Y F K L Y A N A K T V E G V W T L K D E I K T F T V T E
BINARY_6889 T T Y K L I L N G K Q L K E E A I K E A V D A G T A E K Y F K L Y A N A K T V E G V W T L K D E T L K A T V T E
BINARY_6890 T T Y K L I L N G K Q L K E E A I K E A V D A G T A E K Y F K L Y A N A K T V E G V W T L K D E T L K F T V T E
BINARY_6891 T T Y K L I L N G K Q L K E E A I K E A V D A G T A E K Y F K L Y A N A K T V E G V W T L K D E T L T A T V T E
BINARY_6892 T T Y K L I L N G K Q L K E E A I K E A V D A G T A E K Y F K L Y A N A K T V E G V W T L K D E T L T F T V T E
BINARY_6893 T T Y K L I L N G K Q L K E E A I K E A V D A G T A E K Y F K L Y A N A K T V E G V W T L K D E T K K A T V T E
BINARY_6894 T T Y K L I L N G K Q L K E E A I K E A V D A G T A E K Y F K L Y A N A K T V E G V W T L K D E T K K F T V T E
BINARY_6895 T T Y K L I L N G K Q L K E E A I K E A V D A G T A E K Y F K L Y A N A K T V E G V W T L K D E T K T A T V T E
BINARY_6896 T T Y K L I L N G K Q L K E E A I K E A V D A G T A E K Y F K L Y A N A K T V E G V W T L K D E T K T F T V T E
BINARY_6897 T T Y K L I L N G K Q L K E E A I K E A V D A G T A E K Y F K L Y A N A K T V E G V W T Y K D E I L K A T V T E
BINARY_6898 T T Y K L I L N G K Q L K E E A I K E A V D A G T A E K Y F K L Y A N A K T V E G V W T Y K D E I L K F T V T E
BINARY_6899 T T Y K L I L N G K Q L K E E A I K E A V D A G T A E K Y F K L Y A N A K T V E G V W T Y K D E I L T A T V T E
BINARY_6900 T T Y K L I L N G K Q L K E E A I K E A V D A G T A E K Y F K L Y A N A K T V E G V W T Y K D E I L T F T V T E
BINARY_6901 T T Y K L I L N G K Q L K E E A I K E A V D A G T A E K Y F K L Y A N A K T V E G V W T Y K D E I K K A T V T E
BINARY_6902 T T Y K L I L N G K Q L K E E A I K E A V D A G T A E K Y F K L Y A N A K T V E G V W T Y K D E I K K F T V T E
BINARY_6903 T T Y K L I L N G K Q L K E E A I K E A V D A G T A E K Y F K L Y A N A K T V E G V W T Y K D E I K T A T V T E
BINARY_6904 T T Y K L I L N G K Q L K E E A I K E A V D A G T A E K Y F K L Y A N A K T V E G V W T Y K D E I K T F T V T E
BINARY_6905 T T Y K L I L N G K Q L K E E A I K E A V D A G T A E K Y F K L Y A N A K T V E G V W T Y K D E T L K A T V T E
BINARY_6906 T T Y K L I L N G K Q L K E E A I K E A V D A G T A E K Y F K L Y A N A K T V E G V W T Y K D E T L K F T V T E
BINARY_6907 T T Y K L I L N G K Q L K E E A I K E A V D A G T A E K Y F K L Y A N A K T V E G V W T Y K D E T L T A T V T E
BINARY_6908 T T Y K L I L N G K Q L K E E A I K E A V D A G T A E K Y F K L Y A N A K T V E G V W T Y K D E T L T F T V T E
BINARY_6909 T T Y K L I L N G K Q L K E E A I K E A V D A G T A E K Y F K L Y A N A K T V E G V W T Y K D E T K K A T V T E
BINARY_6910 T T Y K L I L N G K Q L K E E A I K E A V D A G T A E K Y F K L Y A N A K T V E G V W T Y K D E T K K F T V T E
BINARY_6911 T T Y K L I L N G K Q L K E E A I K E A V D A G T A E K Y F K L Y A N A K T V E G V W T Y K D E T K T A T V T E
BINARY_6912 T T Y K L I L N G K Q L K E E A I K E A V D A G T A E K Y F K L Y A N A K T V E G V W T Y K D E T K T F T V T E
BINARY_6913 T T Y K L I L N G K Q L K E E A I K E A V D A A I A E K Y I K L I A N A K T V E G V W T L K D E I L K A T V T E
BINARY_6914 T T Y K L I L N G K Q L K E E A I K E A V D A A I A E K Y I K L I A N A K T V E G V W T L K D E I L K F T V T E
BINARY_6915 T T Y K L I L N G K Q L K E E A I K E A V D A A I A E K Y I K L I A N A K T V E G V W T L K D E I L T A T V T E
BINARY_6916 T T Y K L I L N G K Q L K E E A I K E A V D A A I A E K Y I K L I A N A K T V E G V W T L K D E I L T F T V T E
BINARY_6917 T T Y K L I L N G K Q L K E E A I K E A V D A A I A E K Y I K L I A N A K T V E G V W T L K D E I K K A T V T E
BINARY_6918 T T Y K L I L N G K Q L K E E A I K E A V D A A I A E K Y I K L I A N A K T V E G V W T L K D E I K K F T V T E
BINARY_6919 T T Y K L I L N G K Q L K E E A I K E A V D A A I A E K Y I K L I A N A K T V E G V W T L K D E I K T A T V T E
BINARY_6920 T T Y K L I L N G K Q L K E E A I K E A V D A A I A E K Y I K L I A N A K T V E G V W T L K D E I K T F T V T E
BINARY_6921 T T Y K L I L N G K Q L K E E A I K E A V D A A I A E K Y I K L I A N A K T V E G V W T L K D E T L K A T V T E
BINARY_6922 T T Y K L I L N G K Q L K E E A I K E A V D A A I A E K Y I K L I A N A K T V E G V W T L K D E T L K F T V T E
BINARY_6923 T T Y K L I L N G K Q L K E E A I K E A V D A A I A E K Y I K L I A N A K T V E G V W T L K D E T L T A T V T E
BINARY_6924 T T Y K L I L N G K Q L K E E A I K E A V D A A I A E K Y I K L I A N A K T V E G V W T L K D E T L T F T V T E
BINARY_6925 T T Y K L I L N G K Q L K E E A I K E A V D A A I A E K Y I K L I A N A K T V E G V W T L K D E T K K A T V T E
BINARY_6926 T T Y K L I L N G K Q L K E E A I K E A V D A A I A E K Y I K L I A N A K T V E G V W T L K D E T K K F T V T E
BINARY_6927 T T Y K L I L N G K Q L K E E A I K E A V D A A I A E K Y I K L I A N A K T V E G V W T L K D E T K T A T V T E
BINARY_6928 T T Y K L I L N G K Q L K E E A I K E A V D A A I A E K Y I K L I A N A K T V E G V W T L K D E T K T F T V T E
BINARY_6929 T T Y K L I L N G K Q L K E E A I K E A V D A A I A E K Y I K L I A N A K T V E G V W T Y K D E I L K A T V T E
BINARY_6930 T T Y K L I L N G K Q L K E E A I K E A V D A A I A E K Y I K L I A N A K T V E G V W T Y K D E I L K F T V T E
BINARY_6931 T T Y K L I L N G K Q L K E E A I K E A V D A A I A E K Y I K L I A N A K T V E G V W T Y K D E I L T A T V T E
BINARY_6932 T T Y K L I L N G K Q L K E E A I K E A V D A A I A E K Y I K L I A N A K T V E G V W T Y K D E I L T F T V T E
BINARY_6933 T T Y K L I L N G K Q L K E E A I K E A V D A A I A E K Y I K L I A N A K T V E G V W T Y K D E I K K A T V T E
BINARY_6934 T T Y K L I L N G K Q L K E E A I K E A V D A A I A E K Y I K L I A N A K T V E G V W T Y K D E I K K F T V T E
BINARY_6935 T T Y K L I L N G K Q L K E E A I K E A V D A A I A E K Y I K L I A N A K T V E G V W T Y K D E I K T A T V T E
BINARY_6936 T T Y K L I L N G K Q L K E E A I K E A V D A A I A E K Y I K L I A N A K T V E G V W T Y K D E I K T F T V T E
BINARY_6937 T T Y K L I L N G K Q L K E E A I K E A V D A A I A E K Y I K L I A N A K T V E G V W T Y K D E T L K A T V T E
BINARY_6938 T T Y K L I L N G K Q L K E E A I K E A V D A A I A E K Y I K L I A N A K T V E G V W T Y K D E T L K F T V T E
BINARY_6939 T T Y K L I L N G K Q L K E E A I K E A V D A A I A E K Y I K L I A N A K T V E G V W T Y K D E T L T A T V T E
BINARY_6940 T T Y K L I L N G K Q L K E E A I K E A V D A A I A E K Y I K L I A N A K T V E G V W T Y K D E T L T F T V T E
BINARY_6941 T T Y K L I L N G K Q L K E E A I K E A V D A A I A E K Y I K L I A N A K T V E G V W T Y K D E T K K A T V T E
BINARY_6942 T T Y K L I L N G K Q L K E E A I K E A V D A A I A E K Y I K L I A N A K T V E G V W T Y K D E T K K F T V T E
BINARY_6943 T T Y K L I L N G K Q L K E E A I K E A V D A A I A E K Y I K L I A N A K T V E G V W T Y K D E T K T A T V T E
BINARY_6944 T T Y K L I L N G K Q L K E E A I K E A V D A A I A E K Y I K L I A N A K T V E G V W T Y K D E T K T F T V T E
BINARY_6945 T T Y K L I L N G K Q L K E E A I K E A V D A A I A E K Y I K L Y A N A K T V E G V W T L K D E I L K A T V T E
BINARY_6946 T T Y K L I L N G K Q L K E E A I K E A V D A A I A E K Y I K L Y A N A K T V E G V W T L K D E I L K F T V T E
BINARY_6947 T T Y K L I L N G K Q L K E E A I K E A V D A A I A E K Y I K L Y A N A K T V E G V W T L K D E I L T A T V T E
BINARY_6948 T T Y K L I L N G K Q L K E E A I K E A V D A A I A E K Y I K L Y A N A K T V E G V W T L K D E I L T F T V T E
BINARY_6949 T T Y K L I L N G K Q L K E E A I K E A V D A A I A E K Y I K L Y A N A K T V E G V W T L K D E I K K A T V T E
BINARY_6950 T T Y K L I L N G K Q L K E E A I K E A V D A A I A E K Y I K L Y A N A K T V E G V W T L K D E I K K F T V T E
BINARY_6951 T T Y K L I L N G K Q L K E E A I K E A V D A A I A E K Y I K L Y A N A K T V E G V W T L K D E I K T A T V T E

BINARY_6952 T T Y K L I L N G K Q L K E E A I K E A V D A A I A E K Y I K L Y A N A K T V E G V W T L K D E I K T F T V T E
BINARY_6953 T T Y K L I L N G K Q L K E E A I K E A V D A A I A E K Y I K L Y A N A K T V E G V W T L K D E T L K A T V T E
BINARY_6954 T T Y K L I L N G K Q L K E E A I K E A V D A A I A E K Y I K L Y A N A K T V E G V W T L K D E T L K F T V T E
BINARY_6955 T T Y K L I L N G K Q L K E E A I K E A V D A A I A E K Y I K L Y A N A K T V E G V W T L K D E T L T A T V T E
BINARY_6956 T T Y K L I L N G K Q L K E E A I K E A V D A A I A E K Y I K L Y A N A K T V E G V W T L K D E T L T F T V T E
BINARY_6957 T T Y K L I L N G K Q L K E E A I K E A V D A A I A E K Y I K L Y A N A K T V E G V W T L K D E T K K A T V T E
BINARY_6958 T T Y K L I L N G K Q L K E E A I K E A V D A A I A E K Y I K L Y A N A K T V E G V W T L K D E T K K F T V T E
BINARY_6959 T T Y K L I L N G K Q L K E E A I K E A V D A A I A E K Y I K L Y A N A K T V E G V W T L K D E T K T A T V T E
BINARY_6960 T T Y K L I L N G K Q L K E E A I K E A V D A A I A E K Y I K L Y A N A K T V E G V W T L K D E T K T F T V T E
BINARY_6961 T T Y K L I L N G K Q L K E E A I K E A V D A A I A E K Y I K L Y A N A K T V E G V W T Y K D E I L K A T V T E
BINARY_6962 T T Y K L I L N G K Q L K E E A I K E A V D A A I A E K Y I K L Y A N A K T V E G V W T Y K D E I L K F T V T E
BINARY_6963 T T Y K L I L N G K Q L K E E A I K E A V D A A I A E K Y I K L Y A N A K T V E G V W T Y K D E I L T A T V T E
BINARY_6964 T T Y K L I L N G K Q L K E E A I K E A V D A A I A E K Y I K L Y A N A K T V E G V W T Y K D E I L T F T V T E
BINARY_6965 T T Y K L I L N G K Q L K E E A I K E A V D A A I A E K Y I K L Y A N A K T V E G V W T Y K D E I K K A T V T E
BINARY_6966 T T Y K L I L N G K Q L K E E A I K E A V D A A I A E K Y I K L Y A N A K T V E G V W T Y K D E I K K F T V T E
BINARY_6967 T T Y K L I L N G K Q L K E E A I K E A V D A A I A E K Y I K L Y A N A K T V E G V W T Y K D E I K T A T V T E
BINARY_6968 T T Y K L I L N G K Q L K E E A I K E A V D A A I A E K Y I K L Y A N A K T V E G V W T Y K D E I K T F T V T E
BINARY_6969 T T Y K L I L N G K Q L K E E A I K E A V D A A I A E K Y I K L Y A N A K T V E G V W T Y K D E T L K A T V T E
BINARY_6970 T T Y K L I L N G K Q L K E E A I K E A V D A A I A E K Y I K L Y A N A K T V E G V W T Y K D E T L K F T V T E
BINARY_6971 T T Y K L I L N G K Q L K E E A I K E A V D A A I A E K Y I K L Y A N A K T V E G V W T Y K D E T L T A T V T E
BINARY_6972 T T Y K L I L N G K Q L K E E A I K E A V D A A I A E K Y I K L Y A N A K T V E G V W T Y K D E T L T F T V T E
BINARY_6973 T T Y K L I L N G K Q L K E E A I K E A V D A A I A E K Y I K L Y A N A K T V E G V W T Y K D E T K K A T V T E
BINARY_6974 T T Y K L I L N G K Q L K E E A I K E A V D A A I A E K Y I K L Y A N A K T V E G V W T Y K D E T K K F T V T E
BINARY_6975 T T Y K L I L N G K Q L K E E A I K E A V D A A I A E K Y I K L Y A N A K T V E G V W T Y K D E T K T A T V T E
BINARY_6976 T T Y K L I L N G K Q L K E E A I K E A V D A A I A E K Y I K L Y A N A K T V E G V W T Y K D E T K T F T V T E
BINARY_6977 T T Y K L I L N G K Q L K E E A I K E A V D A A I A E K Y F K L I A N A K T V E G V W T L K D E I L K A T V T E
BINARY_6978 T T Y K L I L N G K Q L K E E A I K E A V D A A I A E K Y F K L I A N A K T V E G V W T L K D E I L K F T V T E
BINARY_6979 T T Y K L I L N G K Q L K E E A I K E A V D A A I A E K Y F K L I A N A K T V E G V W T L K D E I L T A T V T E
BINARY_6980 T T Y K L I L N G K Q L K E E A I K E A V D A A I A E K Y F K L I A N A K T V E G V W T L K D E I L T F T V T E
BINARY_6981 T T Y K L I L N G K Q L K E E A I K E A V D A A I A E K Y F K L I A N A K T V E G V W T L K D E I K K A T V T E
BINARY_6982 T T Y K L I L N G K Q L K E E A I K E A V D A A I A E K Y F K L I A N A K T V E G V W T L K D E I K K F T V T E
BINARY_6983 T T Y K L I L N G K Q L K E E A I K E A V D A A I A E K Y F K L I A N A K T V E G V W T L K D E I K T A T V T E
BINARY_6984 T T Y K L I L N G K Q L K E E A I K E A V D A A I A E K Y F K L I A N A K T V E G V W T L K D E I K T F T V T E
BINARY_6985 T T Y K L I L N G K Q L K E E A I K E A V D A A I A E K Y F K L I A N A K T V E G V W T L K D E T L K A T V T E
BINARY_6986 T T Y K L I L N G K Q L K E E A I K E A V D A A I A E K Y F K L I A N A K T V E G V W T L K D E T L K F T V T E
BINARY_6987 T T Y K L I L N G K Q L K E E A I K E A V D A A I A E K Y F K L I A N A K T V E G V W T L K D E T L T A T V T E
BINARY_6988 T T Y K L I L N G K Q L K E E A I K E A V D A A I A E K Y F K L I A N A K T V E G V W T L K D E T L T F T V T E
BINARY_6989 T T Y K L I L N G K Q L K E E A I K E A V D A A I A E K Y F K L I A N A K T V E G V W T L K D E T K K A T V T E
BINARY_6990 T T Y K L I L N G K Q L K E E A I K E A V D A A I A E K Y F K L I A N A K T V E G V W T L K D E T K K F T V T E
BINARY_6991 T T Y K L I L N G K Q L K E E A I K E A V D A A I A E K Y F K L I A N A K T V E G V W T L K D E T K T A T V T E
BINARY_6992 T T Y K L I L N G K Q L K E E A I K E A V D A A I A E K Y F K L I A N A K T V E G V W T L K D E T K T F T V T E
BINARY_6993 T T Y K L I L N G K Q L K E E A I K E A V D A A I A E K Y F K L I A N A K T V E G V W T Y K D E I L K A T V T E
BINARY_6994 T T Y K L I L N G K Q L K E E A I K E A V D A A I A E K Y F K L I A N A K T V E G V W T Y K D E I L K F T V T E
BINARY_6995 T T Y K L I L N G K Q L K E E A I K E A V D A A I A E K Y F K L I A N A K T V E G V W T Y K D E I L T A T V T E
BINARY_6996 T T Y K L I L N G K Q L K E E A I K E A V D A A I A E K Y F K L I A N A K T V E G V W T Y K D E I L T F T V T E
BINARY_6997 T T Y K L I L N G K Q L K E E A I K E A V D A A I A E K Y F K L I A N A K T V E G V W T Y K D E I K K A T V T E
BINARY_6998 T T Y K L I L N G K Q L K E E A I K E A V D A A I A E K Y F K L I A N A K T V E G V W T Y K D E I K K F T V T E
BINARY_6999 T T Y K L I L N G K Q L K E E A I K E A V D A A I A E K Y F K L I A N A K T V E G V W T Y K D E I K T A T V T E
BINARY_7000 T T Y K L I L N G K Q L K E E A I K E A V D A A I A E K Y F K L I A N A K T V E G V W T Y K D E I K T F T V T E
BINARY_7001 T T Y K L I L N G K Q L K E E A I K E A V D A A I A E K Y F K L I A N A K T V E G V W T Y K D E T L K A T V T E
BINARY_7002 T T Y K L I L N G K Q L K E E A I K E A V D A A I A E K Y F K L I A N A K T V E G V W T Y K D E T L K F T V T E
BINARY_7003 T T Y K L I L N G K Q L K E E A I K E A V D A A I A E K Y F K L I A N A K T V E G V W T Y K D E T L T A T V T E
BINARY_7004 T T Y K L I L N G K Q L K E E A I K E A V D A A I A E K Y F K L I A N A K T V E G V W T Y K D E T L T F T V T E
BINARY_7005 T T Y K L I L N G K Q L K E E A I K E A V D A A I A E K Y F K L I A N A K T V E G V W T Y K D E T K K A T V T E
BINARY_7006 T T Y K L I L N G K Q L K E E A I K E A V D A A I A E K Y F K L I A N A K T V E G V W T Y K D E T K K F T V T E
BINARY_7007 T T Y K L I L N G K Q L K E E A I K E A V D A A I A E K Y F K L I A N A K T V E G V W T Y K D E T K T A T V T E
BINARY_7008 T T Y K L I L N G K Q L K E E A I K E A V D A A I A E K Y F K L I A N A K T V E G V W T Y K D E T K T F T V T E
BINARY_7009 T T Y K L I L N G K Q L K E E A I K E A V D A A I A E K Y F K L Y A N A K T V E G V W T L K D E I L K A T V T E
BINARY_7010 T T Y K L I L N G K Q L K E E A I K E A V D A A I A E K Y F K L Y A N A K T V E G V W T L K D E I L K F T V T E
BINARY_7011 T T Y K L I L N G K Q L K E E A I K E A V D A A I A E K Y F K L Y A N A K T V E G V W T L K D E I L T A T V T E
BINARY_7012 T T Y K L I L N G K Q L K E E A I K E A V D A A I A E K Y F K L Y A N A K T V E G V W T L K D E I L T F T V T E
BINARY_7013 T T Y K L I L N G K Q L K E E A I K E A V D A A I A E K Y F K L Y A N A K T V E G V W T L K D E I K K A T V T E
BINARY_7014 T T Y K L I L N G K Q L K E E A I K E A V D A A I A E K Y F K L Y A N A K T V E G V W T L K D E I K K F T V T E
BINARY_7015 T T Y K L I L N G K Q L K E E A I K E A V D A A I A E K Y F K L Y A N A K T V E G V W T L K D E I K T A T V T E
BINARY_7016 T T Y K L I L N G K Q L K E E A I K E A V D A A I A E K Y F K L Y A N A K T V E G V W T L K D E I K T F T V T E
BINARY_7017 T T Y K L I L N G K Q L K E E A I K E A V D A A I A E K Y F K L Y A N A K T V E G V W T L K D E T L K A T V T E
BINARY_7018 T T Y K L I L N G K Q L K E E A I K E A V D A A I A E K Y F K L Y A N A K T V E G V W T L K D E T L K F T V T E
BINARY_7019 T T Y K L I L N G K Q L K E E A I K E A V D A A I A E K Y F K L Y A N A K T V E G V W T L K D E T L T A T V T E
BINARY_7020 T T Y K L I L N G K Q L K E E A I K E A V D A A I A E K Y F K L Y A N A K T V E G V W T L K D E T L T F T V T E
BINARY_7021 T T Y K L I L N G K Q L K E E A I K E A V D A A I A E K Y F K L Y A N A K T V E G V W T L K D E T K K A T V T E
BINARY_7022 T T Y K L I L N G K Q L K E E A I K E A V D A A I A E K Y F K L Y A N A K T V E G V W T L K D E T K K F T V T E
BINARY_7023 T T Y K L I L N G K Q L K E E A I K E A V D A A I A E K Y F K L Y A N A K T V E G V W T L K D E T K T A T V T E
BINARY_7024 T T Y K L I L N G K Q L K E E A I K E A V D A A I A E K Y F K L Y A N A K T V E G V W T L K D E T K T F T V T E
BINARY_7025 T T Y K L I L N G K Q L K E E A I K E A V D A A I A E K Y F K L Y A N A K T V E G V W T Y K D E I L K A T V T E
BINARY_7026 T T Y K L I L N G K Q L K E E A I K E A V D A A I A E K Y F K L Y A N A K T V E G V W T Y K D E I L K F T V T E
BINARY_7027 T T Y K L I L N G K Q L K E E A I K E A V D A A I A E K Y F K L Y A N A K T V E G V W T Y K D E I L T A T V T E
BINARY_7028 T T Y K L I L N G K Q L K E E A I K E A V D A A I A E K Y F K L Y A N A K T V E G V W T Y K D E I L T F T V T E
BINARY_7029 T T Y K L I L N G K Q L K E E A I K E A V D A A I A E K Y F K L Y A N A K T V E G V W T Y K D E I K K A T V T E
BINARY_7030 T T Y K L I L N G K Q L K E E A I K E A V D A A I A E K Y F K L Y A N A K T V E G V W T Y K D E I K K F T V T E
BINARY_7031 T T Y K L I L N G K Q L K E E A I K E A V D A A I A E K Y F K L Y A N A K T V E G V W T Y K D E I K T A T V T E
BINARY_7032 T T Y K L I L N G K Q L K E E A I K E A V D A A I A E K Y F K L Y A N A K T V E G V W T Y K D E I K T F T V T E
BINARY_7033 T T Y K L I L N G K Q L K E E A I K E A V D A A I A E K Y F K L Y A N A K T V E G V W T Y K D E T L K A T V T E
BINARY_7034 T T Y K L I L N G K Q L K E E A I K E A V D A A I A E K Y F K L Y A N A K T V E G V W T Y K D E T L K F T V T E
BINARY_7035 T T Y K L I L N G K Q L K E E A I K E A V D A A I A E K Y F K L Y A N A K T V E G V W T Y K D E T L T A T V T E
BINARY_7036 T T Y K L I L N G K Q L K E E A I K E A V D A A I A E K Y F K L Y A N A K T V E G V W T Y K D E T L T F T V T E
BINARY_7037 T T Y K L I L N G K Q L K E E A I K E A V D A A I A E K Y F K L Y A N A K T V E G V W T Y K D E T K K A T V T E
BINARY_7038 T T Y K L I L N G K Q L K E E A I K E A V D A A I A E K Y F K L Y A N A K T V E G V W T Y K D E T K K F T V T E
BINARY_7039 T T Y K L I L N G K Q L K E E A I K E A V D A A I A E K Y F K L Y A N A K T V E G V W T Y K D E T K T A T V T E
BINARY_7040 T T Y K L I L N G K Q L K E E A I K E A V D A A I A E K Y F K L Y A N A K T V E G V W T Y K D E T K T F T V T E
BINARY_7041 T T Y K L I L N G K Q L K E E A I K E A V D A A T A E K Y I K L I A N A K T V E G V W T L K D E I L K A T V T E
BINARY_7042 T T Y K L I L N G K Q L K E E A I K E A V D A A T A E K Y I K L I A N A K T V E G V W T L K D E I L K F T V T E
BINARY_7043 T T Y K L I L N G K Q L K E E A I K E A V D A A T A E K Y I K L I A N A K T V E G V W T L K D E I L T A T V T E
BINARY_7044 T T Y K L I L N G K Q L K E E A I K E A V D A A T A E K Y I K L I A N A K T V E G V W T L K D E I L T F T V T E
BINARY_7045 T T Y K L I L N G K Q L K E E A I K E A V D A A T A E K Y I K L I A N A K T V E G V W T L K D E I K K A T V T E
BINARY_7046 T T Y K L I L N G K Q L K E E A I K E A V D A A T A E K Y I K L I A N A K T V E G V W T L K D E I K K F T V T E

BINARY_7047 T T Y K L I L N G K Q L K E E A I K E A V D A A T A E K Y I K L I A N A K T V E G V W T L K D E I K T A T V T E
BINARY_7048 T T Y K L I L N G K Q L K E E A I K E A V D A A T A E K Y I K L I A N A K T V E G V W T L K D E I K T F T V T E
BINARY_7049 T T Y K L I L N G K Q L K E E A I K E A V D A A T A E K Y I K L I A N A K T V E G V W T L K D E T L K A T V T E
BINARY_7050 T T Y K L I L N G K Q L K E E A I K E A V D A A T A E K Y I K L I A N A K T V E G V W T L K D E T L K F T V T E
BINARY_7051 T T Y K L I L N G K Q L K E E A I K E A V D A A T A E K Y I K L I A N A K T V E G V W T L K D E T L T A T V T E
BINARY_7052 T T Y K L I L N G K Q L K E E A I K E A V D A A T A E K Y I K L I A N A K T V E G V W T L K D E T L T F T V T E
BINARY_7053 T T Y K L I L N G K Q L K E E A I K E A V D A A T A E K Y I K L I A N A K T V E G V W T L K D E T K K A T V T E
BINARY_7054 T T Y K L I L N G K Q L K E E A I K E A V D A A T A E K Y I K L I A N A K T V E G V W T L K D E T K K F T V T E
BINARY_7055 T T Y K L I L N G K Q L K E E A I K E A V D A A T A E K Y I K L I A N A K T V E G V W T L K D E T K T A T V T E
BINARY_7056 T T Y K L I L N G K Q L K E E A I K E A V D A A T A E K Y I K L I A N A K T V E G V W T L K D E T K T F T V T E
BINARY_7057 T T Y K L I L N G K Q L K E E A I K E A V D A A T A E K Y I K L I A N A K T V E G V W T Y K D E I L K A T V T E
BINARY_7058 T T Y K L I L N G K Q L K E E A I K E A V D A A T A E K Y I K L I A N A K T V E G V W T Y K D E I L K F T V T E
BINARY_7059 T T Y K L I L N G K Q L K E E A I K E A V D A A T A E K Y I K L I A N A K T V E G V W T Y K D E I L T A T V T E
BINARY_7060 T T Y K L I L N G K Q L K E E A I K E A V D A A T A E K Y I K L I A N A K T V E G V W T Y K D E I L T F T V T E
BINARY_7061 T T Y K L I L N G K Q L K E E A I K E A V D A A T A E K Y I K L I A N A K T V E G V W T Y K D E I K K A T V T E
BINARY_7062 T T Y K L I L N G K Q L K E E A I K E A V D A A T A E K Y I K L I A N A K T V E G V W T Y K D E I K K F T V T E
BINARY_7063 T T Y K L I L N G K Q L K E E A I K E A V D A A T A E K Y I K L I A N A K T V E G V W T Y K D E I K T A T V T E
BINARY_7064 T T Y K L I L N G K Q L K E E A I K E A V D A A T A E K Y I K L I A N A K T V E G V W T Y K D E I K T F T V T E
BINARY_7065 T T Y K L I L N G K Q L K E E A I K E A V D A A T A E K Y I K L I A N A K T V E G V W T Y K D E T L K A T V T E
BINARY_7066 T T Y K L I L N G K Q L K E E A I K E A V D A A T A E K Y I K L I A N A K T V E G V W T Y K D E T L K F T V T E
BINARY_7067 T T Y K L I L N G K Q L K E E A I K E A V D A A T A E K Y I K L I A N A K T V E G V W T Y K D E T L T A T V T E
BINARY_7068 T T Y K L I L N G K Q L K E E A I K E A V D A A T A E K Y I K L I A N A K T V E G V W T Y K D E T L T F T V T E
BINARY_7069 T T Y K L I L N G K Q L K E E A I K E A V D A A T A E K Y I K L I A N A K T V E G V W T Y K D E T K K A T V T E
BINARY_7070 T T Y K L I L N G K Q L K E E A I K E A V D A A T A E K Y I K L I A N A K T V E G V W T Y K D E T K K F T V T E
BINARY_7071 T T Y K L I L N G K Q L K E E A I K E A V D A A T A E K Y I K L I A N A K T V E G V W T Y K D E T K T A T V T E
BINARY_7072 T T Y K L I L N G K Q L K E E A I K E A V D A A T A E K Y I K L I A N A K T V E G V W T Y K D E T K T F T V T E
BINARY_7073 T T Y K L I L N G K Q L K E E A I K E A V D A A T A E K Y I K L Y A N A K T V E G V W T L K D E I L K A T V T E
BINARY_7074 T T Y K L I L N G K Q L K E E A I K E A V D A A T A E K Y I K L Y A N A K T V E G V W T L K D E I L K F T V T E
BINARY_7075 T T Y K L I L N G K Q L K E E A I K E A V D A A T A E K Y I K L Y A N A K T V E G V W T L K D E I L T A T V T E
BINARY_7076 T T Y K L I L N G K Q L K E E A I K E A V D A A T A E K Y I K L Y A N A K T V E G V W T L K D E I L T F T V T E
BINARY_7077 T T Y K L I L N G K Q L K E E A I K E A V D A A T A E K Y I K L Y A N A K T V E G V W T L K D E I K K A T V T E
BINARY_7078 T T Y K L I L N G K Q L K E E A I K E A V D A A T A E K Y I K L Y A N A K T V E G V W T L K D E I K K F T V T E
BINARY_7079 T T Y K L I L N G K Q L K E E A I K E A V D A A T A E K Y I K L Y A N A K T V E G V W T L K D E I K T A T V T E
BINARY_7080 T T Y K L I L N G K Q L K E E A I K E A V D A A T A E K Y I K L Y A N A K T V E G V W T L K D E I K T F T V T E
BINARY_7081 T T Y K L I L N G K Q L K E E A I K E A V D A A T A E K Y I K L Y A N A K T V E G V W T L K D E T L K A T V T E
BINARY_7082 T T Y K L I L N G K Q L K E E A I K E A V D A A T A E K Y I K L Y A N A K T V E G V W T L K D E T L K F T V T E
BINARY_7083 T T Y K L I L N G K Q L K E E A I K E A V D A A T A E K Y I K L Y A N A K T V E G V W T L K D E T L T A T V T E
BINARY_7084 T T Y K L I L N G K Q L K E E A I K E A V D A A T A E K Y I K L Y A N A K T V E G V W T L K D E T L T F T V T E
BINARY_7085 T T Y K L I L N G K Q L K E E A I K E A V D A A T A E K Y I K L Y A N A K T V E G V W T L K D E T K K A T V T E
BINARY_7086 T T Y K L I L N G K Q L K E E A I K E A V D A A T A E K Y I K L Y A N A K T V E G V W T L K D E T K K F T V T E
BINARY_7087 T T Y K L I L N G K Q L K E E A I K E A V D A A T A E K Y I K L Y A N A K T V E G V W T L K D E T K T A T V T E
BINARY_7088 T T Y K L I L N G K Q L K E E A I K E A V D A A T A E K Y I K L Y A N A K T V E G V W T L K D E T K T F T V T E
BINARY_7089 T T Y K L I L N G K Q L K E E A I K E A V D A A T A E K Y I K L Y A N A K T V E G V W T Y K D E I L K A T V T E
BINARY_7090 T T Y K L I L N G K Q L K E E A I K E A V D A A T A E K Y I K L Y A N A K T V E G V W T Y K D E I L K F T V T E
BINARY_7091 T T Y K L I L N G K Q L K E E A I K E A V D A A T A E K Y I K L Y A N A K T V E G V W T Y K D E I L T A T V T E
BINARY_7092 T T Y K L I L N G K Q L K E E A I K E A V D A A T A E K Y I K L Y A N A K T V E G V W T Y K D E I L T F T V T E
BINARY_7093 T T Y K L I L N G K Q L K E E A I K E A V D A A T A E K Y I K L Y A N A K T V E G V W T Y K D E I K K A T V T E
BINARY_7094 T T Y K L I L N G K Q L K E E A I K E A V D A A T A E K Y I K L Y A N A K T V E G V W T Y K D E I K K F T V T E
BINARY_7095 T T Y K L I L N G K Q L K E E A I K E A V D A A T A E K Y I K L Y A N A K T V E G V W T Y K D E I K T A T V T E
BINARY_7096 T T Y K L I L N G K Q L K E E A I K E A V D A A T A E K Y I K L Y A N A K T V E G V W T Y K D E I K T F T V T E
BINARY_7097 T T Y K L I L N G K Q L K E E A I K E A V D A A T A E K Y I K L Y A N A K T V E G V W T Y K D E T L K A T V T E
BINARY_7098 T T Y K L I L N G K Q L K E E A I K E A V D A A T A E K Y I K L Y A N A K T V E G V W T Y K D E T L K F T V T E
BINARY_7099 T T Y K L I L N G K Q L K E E A I K E A V D A A T A E K Y I K L Y A N A K T V E G V W T Y K D E T L T A T V T E
BINARY_7100 T T Y K L I L N G K Q L K E E A I K E A V D A A T A E K Y I K L Y A N A K T V E G V W T Y K D E T L T F T V T E
BINARY_7101 T T Y K L I L N G K Q L K E E A I K E A V D A A T A E K Y I K L Y A N A K T V E G V W T Y K D E T K K A T V T E
BINARY_7102 T T Y K L I L N G K Q L K E E A I K E A V D A A T A E K Y I K L Y A N A K T V E G V W T Y K D E T K K F T V T E
BINARY_7103 T T Y K L I L N G K Q L K E E A I K E A V D A A T A E K Y I K L Y A N A K T V E G V W T Y K D E T K T A T V T E
BINARY_7104 T T Y K L I L N G K Q L K E E A I K E A V D A A T A E K Y I K L Y A N A K T V E G V W T Y K D E T K T F T V T E
BINARY_7105 T T Y K L I L N G K Q L K E E A I K E A V D A A T A E K Y F K L I A N A K T V E G V W T L K D E I L K A T V T E
BINARY_7106 T T Y K L I L N G K Q L K E E A I K E A V D A A T A E K Y F K L I A N A K T V E G V W T L K D E I L K F T V T E
BINARY_7107 T T Y K L I L N G K Q L K E E A I K E A V D A A T A E K Y F K L I A N A K T V E G V W T L K D E I L T A T V T E
BINARY_7108 T T Y K L I L N G K Q L K E E A I K E A V D A A T A E K Y F K L I A N A K T V E G V W T L K D E I L T F T V T E
BINARY_7109 T T Y K L I L N G K Q L K E E A I K E A V D A A T A E K Y F K L I A N A K T V E G V W T L K D E I K K A T V T E
BINARY_7110 T T Y K L I L N G K Q L K E E A I K E A V D A A T A E K Y F K L I A N A K T V E G V W T L K D E I K K F T V T E
BINARY_7111 T T Y K L I L N G K Q L K E E A I K E A V D A A T A E K Y F K L I A N A K T V E G V W T L K D E I K T A T V T E
BINARY_7112 T T Y K L I L N G K Q L K E E A I K E A V D A A T A E K Y F K L I A N A K T V E G V W T L K D E I K T F T V T E
BINARY_7113 T T Y K L I L N G K Q L K E E A I K E A V D A A T A E K Y F K L I A N A K T V E G V W T L K D E T L K A T V T E
BINARY_7114 T T Y K L I L N G K Q L K E E A I K E A V D A A T A E K Y F K L I A N A K T V E G V W T L K D E T L K F T V T E
BINARY_7115 T T Y K L I L N G K Q L K E E A I K E A V D A A T A E K Y F K L I A N A K T V E G V W T L K D E T L T A T V T E
BINARY_7116 T T Y K L I L N G K Q L K E E A I K E A V D A A T A E K Y F K L I A N A K T V E G V W T L K D E T L T F T V T E
BINARY_7117 T T Y K L I L N G K Q L K E E A I K E A V D A A T A E K Y F K L I A N A K T V E G V W T L K D E T K K A T V T E
BINARY_7118 T T Y K L I L N G K Q L K E E A I K E A V D A A T A E K Y F K L I A N A K T V E G V W T L K D E T K K F T V T E
BINARY_7119 T T Y K L I L N G K Q L K E E A I K E A V D A A T A E K Y F K L I A N A K T V E G V W T L K D E T K T A T V T E
BINARY_7120 T T Y K L I L N G K Q L K E E A I K E A V D A A T A E K Y F K L I A N A K T V E G V W T L K D E T K T F T V T E
BINARY_7121 T T Y K L I L N G K Q L K E E A I K E A V D A A T A E K Y F K L I A N A K T V E G V W T Y K D E I L K A T V T E
BINARY_7122 T T Y K L I L N G K Q L K E E A I K E A V D A A T A E K Y F K L I A N A K T V E G V W T Y K D E I L K F T V T E
BINARY_7123 T T Y K L I L N G K Q L K E E A I K E A V D A A T A E K Y F K L I A N A K T V E G V W T Y K D E I L T A T V T E
BINARY_7124 T T Y K L I L N G K Q L K E E A I K E A V D A A T A E K Y F K L I A N A K T V E G V W T Y K D E I L T F T V T E
BINARY_7125 T T Y K L I L N G K Q L K E E A I K E A V D A A T A E K Y F K L I A N A K T V E G V W T Y K D E I K K A T V T E
BINARY_7126 T T Y K L I L N G K Q L K E E A I K E A V D A A T A E K Y F K L I A N A K T V E G V W T Y K D E I K K F T V T E
BINARY_7127 T T Y K L I L N G K Q L K E E A I K E A V D A A T A E K Y F K L I A N A K T V E G V W T Y K D E I K T A T V T E
BINARY_7128 T T Y K L I L N G K Q L K E E A I K E A V D A A T A E K Y F K L I A N A K T V E G V W T Y K D E I K T F T V T E
BINARY_7129 T T Y K L I L N G K Q L K E E A I K E A V D A A T A E K Y F K L I A N A K T V E G V W T Y K D E T L K A T V T E
BINARY_7130 T T Y K L I L N G K Q L K E E A I K E A V D A A T A E K Y F K L I A N A K T V E G V W T Y K D E T L K F T V T E
BINARY_7131 T T Y K L I L N G K Q L K E E A I K E A V D A A T A E K Y F K L I A N A K T V E G V W T Y K D E T L T A T V T E
BINARY_7132 T T Y K L I L N G K Q L K E E A I K E A V D A A T A E K Y F K L I A N A K T V E G V W T Y K D E T L T F T V T E
BINARY_7133 T T Y K L I L N G K Q L K E E A I K E A V D A A T A E K Y F K L I A N A K T V E G V W T Y K D E T K K A T V T E
BINARY_7134 T T Y K L I L N G K Q L K E E A I K E A V D A A T A E K Y F K L I A N A K T V E G V W T Y K D E T K K F T V T E
BINARY_7135 T T Y K L I L N G K Q L K E E A I K E A V D A A T A E K Y F K L I A N A K T V E G V W T Y K D E T K T A T V T E
BINARY_7136 T T Y K L I L N G K Q L K E E A I K E A V D A A T A E K Y F K L I A N A K T V E G V W T Y K D E T K T F T V T E
BINARY_7137 T T Y K L I L N G K Q L K E E A I K E A V D A A T A E K Y F K L Y A N A K T V E G V W T L K D E I L K A T V T E
BINARY_7138 T T Y K L I L N G K Q L K E E A I K E A V D A A T A E K Y F K L Y A N A K T V E G V W T L K D E I L K F T V T E
BINARY_7139 T T Y K L I L N G K Q L K E E A I K E A V D A A T A E K Y F K L Y A N A K T V E G V W T L K D E I L T A T V T E
BINARY_7140 T T Y K L I L N G K Q L K E E A I K E A V D A A T A E K Y F K L Y A N A K T V E G V W T L K D E I L T F T V T E
BINARY_7141 T T Y K L I L N G K Q L K E E A I K E A V D A A T A E K Y F K L Y A N A K T V E G V W T L K D E I K K A T V T E

BINARY_7142 T T Y K L I L N G K Q L K E E A I K E A V D A A T A E K Y F K L Y A N A K T V E G V W T L K D E I K K F T V T E
BINARY_7143 T T Y K L I L N G K Q L K E E A I K E A V D A A T A E K Y F K L Y A N A K T V E G V W T L K D E I K T A T V T E
BINARY_7144 T T Y K L I L N G K Q L K E E A I K E A V D A A T A E K Y F K L Y A N A K T V E G V W T L K D E I K T F T V T E
BINARY_7145 T T Y K L I L N G K Q L K E E A I K E A V D A A T A E K Y F K L Y A N A K T V E G V W T L K D E T L K A T V T E
BINARY_7146 T T Y K L I L N G K Q L K E E A I K E A V D A A T A E K Y F K L Y A N A K T V E G V W T L K D E T L K F T V T E
BINARY_7147 T T Y K L I L N G K Q L K E E A I K E A V D A A T A E K Y F K L Y A N A K T V E G V W T L K D E T L T A T V T E
BINARY_7148 T T Y K L I L N G K Q L K E E A I K E A V D A A T A E K Y F K L Y A N A K T V E G V W T L K D E T L T F T V T E
BINARY_7149 T T Y K L I L N G K Q L K E E A I K E A V D A A T A E K Y F K L Y A N A K T V E G V W T L K D E T K K A T V T E
BINARY_7150 T T Y K L I L N G K Q L K E E A I K E A V D A A T A E K Y F K L Y A N A K T V E G V W T L K D E T K K F T V T E
BINARY_7151 T T Y K L I L N G K Q L K E E A I K E A V D A A T A E K Y F K L Y A N A K T V E G V W T L K D E T K T A T V T E
BINARY_7152 T T Y K L I L N G K Q L K E E A I K E A V D A A T A E K Y F K L Y A N A K T V E G V W T L K D E T K T F T V T E
BINARY_7153 T T Y K L I L N G K Q L K E E A I K E A V D A A T A E K Y F K L Y A N A K T V E G V W T Y K D E I L K A T V T E
BINARY_7154 T T Y K L I L N G K Q L K E E A I K E A V D A A T A E K Y F K L Y A N A K T V E G V W T Y K D E I L K F T V T E
BINARY_7155 T T Y K L I L N G K Q L K E E A I K E A V D A A T A E K Y F K L Y A N A K T V E G V W T Y K D E I L T A T V T E
BINARY_7156 T T Y K L I L N G K Q L K E E A I K E A V D A A T A E K Y F K L Y A N A K T V E G V W T Y K D E I L T F T V T E
BINARY_7157 T T Y K L I L N G K Q L K E E A I K E A V D A A T A E K Y F K L Y A N A K T V E G V W T Y K D E I K K A T V T E
BINARY_7158 T T Y K L I L N G K Q L K E E A I K E A V D A A T A E K Y F K L Y A N A K T V E G V W T Y K D E I K K F T V T E
BINARY_7159 T T Y K L I L N G K Q L K E E A I K E A V D A A T A E K Y F K L Y A N A K T V E G V W T Y K D E I K T A T V T E
BINARY_7160 T T Y K L I L N G K Q L K E E A I K E A V D A A T A E K Y F K L Y A N A K T V E G V W T Y K D E I K T F T V T E
BINARY_7161 T T Y K L I L N G K Q L K E E A I K E A V D A A T A E K Y F K L Y A N A K T V E G V W T Y K D E T L K A T V T E
BINARY_7162 T T Y K L I L N G K Q L K E E A I K E A V D A A T A E K Y F K L Y A N A K T V E G V W T Y K D E T L K F T V T E
BINARY_7163 T T Y K L I L N G K Q L K E E A I K E A V D A A T A E K Y F K L Y A N A K T V E G V W T Y K D E T L T A T V T E
BINARY_7164 T T Y K L I L N G K Q L K E E A I K E A V D A A T A E K Y F K L Y A N A K T V E G V W T Y K D E T L T F T V T E
BINARY_7165 T T Y K L I L N G K Q L K E E A I K E A V D A A T A E K Y F K L Y A N A K T V E G V W T Y K D E T K K A T V T E
BINARY_7166 T T Y K L I L N G K Q L K E E A I K E A V D A A T A E K Y F K L Y A N A K T V E G V W T Y K D E T K K F T V T E
BINARY_7167 T T Y K L I L N G K Q L K E E A I K E A V D A A T A E K Y F K L Y A N A K T V E G V W T Y K D E T K T A T V T E
BINARY_7168 T T Y K L I L N G K Q L K E E A I K E A V D A A T A E K Y F K L Y A N A K T V E G V W T Y K D E T K T F T V T E
BINARY_7169 T T Y K L I L N G K Q L K E E A I T E L V D A G I A E K Y I K L I A N A K T V E G V W T L K D E I L K A T V T E
BINARY_7170 T T Y K L I L N G K Q L K E E A I T E L V D A G I A E K Y I K L I A N A K T V E G V W T L K D E I L K F T V T E
BINARY_7171 T T Y K L I L N G K Q L K E E A I T E L V D A G I A E K Y I K L I A N A K T V E G V W T L K D E I L T A T V T E
BINARY_7172 T T Y K L I L N G K Q L K E E A I T E L V D A G I A E K Y I K L I A N A K T V E G V W T L K D E I L T F T V T E
BINARY_7173 T T Y K L I L N G K Q L K E E A I T E L V D A G I A E K Y I K L I A N A K T V E G V W T L K D E I K K A T V T E
BINARY_7174 T T Y K L I L N G K Q L K E E A I T E L V D A G I A E K Y I K L I A N A K T V E G V W T L K D E I K K F T V T E
BINARY_7175 T T Y K L I L N G K Q L K E E A I T E L V D A G I A E K Y I K L I A N A K T V E G V W T L K D E I K T A T V T E
BINARY_7176 T T Y K L I L N G K Q L K E E A I T E L V D A G I A E K Y I K L I A N A K T V E G V W T L K D E I K T F T V T E
BINARY_7177 T T Y K L I L N G K Q L K E E A I T E L V D A G I A E K Y I K L I A N A K T V E G V W T L K D E T L K A T V T E
BINARY_7178 T T Y K L I L N G K Q L K E E A I T E L V D A G I A E K Y I K L I A N A K T V E G V W T L K D E T L K F T V T E
BINARY_7179 T T Y K L I L N G K Q L K E E A I T E L V D A G I A E K Y I K L I A N A K T V E G V W T L K D E T L T A T V T E
BINARY_7180 T T Y K L I L N G K Q L K E E A I T E L V D A G I A E K Y I K L I A N A K T V E G V W T L K D E T L T F T V T E
BINARY_7181 T T Y K L I L N G K Q L K E E A I T E L V D A G I A E K Y I K L I A N A K T V E G V W T L K D E T K K A T V T E
BINARY_7182 T T Y K L I L N G K Q L K E E A I T E L V D A G I A E K Y I K L I A N A K T V E G V W T L K D E T K K F T V T E
BINARY_7183 T T Y K L I L N G K Q L K E E A I T E L V D A G I A E K Y I K L I A N A K T V E G V W T L K D E T K T A T V T E
BINARY_7184 T T Y K L I L N G K Q L K E E A I T E L V D A G I A E K Y I K L I A N A K T V E G V W T L K D E T K T F T V T E
BINARY_7185 T T Y K L I L N G K Q L K E E A I T E L V D A G I A E K Y I K L I A N A K T V E G V W T Y K D E I L K A T V T E
BINARY_7186 T T Y K L I L N G K Q L K E E A I T E L V D A G I A E K Y I K L I A N A K T V E G V W T Y K D E I L K F T V T E
BINARY_7187 T T Y K L I L N G K Q L K E E A I T E L V D A G I A E K Y I K L I A N A K T V E G V W T Y K D E I L T A T V T E
BINARY_7188 T T Y K L I L N G K Q L K E E A I T E L V D A G I A E K Y I K L I A N A K T V E G V W T Y K D E I L T F T V T E
BINARY_7189 T T Y K L I L N G K Q L K E E A I T E L V D A G I A E K Y I K L I A N A K T V E G V W T Y K D E I K K A T V T E
BINARY_7190 T T Y K L I L N G K Q L K E E A I T E L V D A G I A E K Y I K L I A N A K T V E G V W T Y K D E I K K F T V T E
BINARY_7191 T T Y K L I L N G K Q L K E E A I T E L V D A G I A E K Y I K L I A N A K T V E G V W T Y K D E I K T A T V T E
BINARY_7192 T T Y K L I L N G K Q L K E E A I T E L V D A G I A E K Y I K L I A N A K T V E G V W T Y K D E I K T F T V T E
BINARY_7193 T T Y K L I L N G K Q L K E E A I T E L V D A G I A E K Y I K L I A N A K T V E G V W T Y K D E T L K A T V T E
BINARY_7194 T T Y K L I L N G K Q L K E E A I T E L V D A G I A E K Y I K L I A N A K T V E G V W T Y K D E T L K F T V T E
BINARY_7195 T T Y K L I L N G K Q L K E E A I T E L V D A G I A E K Y I K L I A N A K T V E G V W T Y K D E T L T A T V T E
BINARY_7196 T T Y K L I L N G K Q L K E E A I T E L V D A G I A E K Y I K L I A N A K T V E G V W T Y K D E T L T F T V T E
BINARY_7197 T T Y K L I L N G K Q L K E E A I T E L V D A G I A E K Y I K L I A N A K T V E G V W T Y K D E T K K A T V T E
BINARY_7198 T T Y K L I L N G K Q L K E E A I T E L V D A G I A E K Y I K L I A N A K T V E G V W T Y K D E T K K F T V T E
BINARY_7199 T T Y K L I L N G K Q L K E E A I T E L V D A G I A E K Y I K L I A N A K T V E G V W T Y K D E T K T A T V T E
BINARY_7200 T T Y K L I L N G K Q L K E E A I T E L V D A G I A E K Y I K L I A N A K T V E G V W T Y K D E T K T F T V T E
BINARY_7201 T T Y K L I L N G K Q L K E E A I T E L V D A G I A E K Y I K L Y A N A K T V E G V W T L K D E I L K A T V T E
BINARY_7202 T T Y K L I L N G K Q L K E E A I T E L V D A G I A E K Y I K L Y A N A K T V E G V W T L K D E I L K F T V T E
BINARY_7203 T T Y K L I L N G K Q L K E E A I T E L V D A G I A E K Y I K L Y A N A K T V E G V W T L K D E I L T A T V T E
BINARY_7204 T T Y K L I L N G K Q L K E E A I T E L V D A G I A E K Y I K L Y A N A K T V E G V W T L K D E I L T F T V T E
BINARY_7205 T T Y K L I L N G K Q L K E E A I T E L V D A G I A E K Y I K L Y A N A K T V E G V W T L K D E I K K A T V T E
BINARY_7206 T T Y K L I L N G K Q L K E E A I T E L V D A G I A E K Y I K L Y A N A K T V E G V W T L K D E I K K F T V T E
BINARY_7207 T T Y K L I L N G K Q L K E E A I T E L V D A G I A E K Y I K L Y A N A K T V E G V W T L K D E I K T A T V T E
BINARY_7208 T T Y K L I L N G K Q L K E E A I T E L V D A G I A E K Y I K L Y A N A K T V E G V W T L K D E I K T F T V T E
BINARY_7209 T T Y K L I L N G K Q L K E E A I T E L V D A G I A E K Y I K L Y A N A K T V E G V W T L K D E T L K A T V T E
BINARY_7210 T T Y K L I L N G K Q L K E E A I T E L V D A G I A E K Y I K L Y A N A K T V E G V W T L K D E T L K F T V T E
BINARY_7211 T T Y K L I L N G K Q L K E E A I T E L V D A G I A E K Y I K L Y A N A K T V E G V W T L K D E T L T A T V T E
BINARY_7212 T T Y K L I L N G K Q L K E E A I T E L V D A G I A E K Y I K L Y A N A K T V E G V W T L K D E T L T F T V T E
BINARY_7213 T T Y K L I L N G K Q L K E E A I T E L V D A G I A E K Y I K L Y A N A K T V E G V W T L K D E T K K A T V T E
BINARY_7214 T T Y K L I L N G K Q L K E E A I T E L V D A G I A E K Y I K L Y A N A K T V E G V W T L K D E T K K F T V T E
BINARY_7215 T T Y K L I L N G K Q L K E E A I T E L V D A G I A E K Y I K L Y A N A K T V E G V W T L K D E T K T A T V T E
BINARY_7216 T T Y K L I L N G K Q L K E E A I T E L V D A G I A E K Y I K L Y A N A K T V E G V W T L K D E T K T F T V T E
BINARY_7217 T T Y K L I L N G K Q L K E E A I T E L V D A G I A E K Y I K L Y A N A K T V E G V W T Y K D E I L K A T V T E
BINARY_7218 T T Y K L I L N G K Q L K E E A I T E L V D A G I A E K Y I K L Y A N A K T V E G V W T Y K D E I L K F T V T E
BINARY_7219 T T Y K L I L N G K Q L K E E A I T E L V D A G I A E K Y I K L Y A N A K T V E G V W T Y K D E I L T A T V T E
BINARY_7220 T T Y K L I L N G K Q L K E E A I T E L V D A G I A E K Y I K L Y A N A K T V E G V W T Y K D E I L T F T V T E
BINARY_7221 T T Y K L I L N G K Q L K E E A I T E L V D A G I A E K Y I K L Y A N A K T V E G V W T Y K D E I K K A T V T E
BINARY_7222 T T Y K L I L N G K Q L K E E A I T E L V D A G I A E K Y I K L Y A N A K T V E G V W T Y K D E I K K F T V T E
BINARY_7223 T T Y K L I L N G K Q L K E E A I T E L V D A G I A E K Y I K L Y A N A K T V E G V W T Y K D E I K T A T V T E
BINARY_7224 T T Y K L I L N G K Q L K E E A I T E L V D A G I A E K Y I K L Y A N A K T V E G V W T Y K D E I K T F T V T E
BINARY_7225 T T Y K L I L N G K Q L K E E A I T E L V D A G I A E K Y I K L Y A N A K T V E G V W T Y K D E T L K A T V T E
BINARY_7226 T T Y K L I L N G K Q L K E E A I T E L V D A G I A E K Y I K L Y A N A K T V E G V W T Y K D E T L K F T V T E
BINARY_7227 T T Y K L I L N G K Q L K E E A I T E L V D A G I A E K Y I K L Y A N A K T V E G V W T Y K D E T L T A T V T E
BINARY_7228 T T Y K L I L N G K Q L K E E A I T E L V D A G I A E K Y I K L Y A N A K T V E G V W T Y K D E T L T F T V T E
BINARY_7229 T T Y K L I L N G K Q L K E E A I T E L V D A G I A E K Y I K L Y A N A K T V E G V W T Y K D E T K K A T V T E
BINARY_7230 T T Y K L I L N G K Q L K E E A I T E L V D A G I A E K Y I K L Y A N A K T V E G V W T Y K D E T K K F T V T E
BINARY_7231 T T Y K L I L N G K Q L K E E A I T E L V D A G I A E K Y I K L Y A N A K T V E G V W T Y K D E T K T A T V T E
BINARY_7232 T T Y K L I L N G K Q L K E E A I T E L V D A G I A E K Y I K L Y A N A K T V E G V W T Y K D E T K T F T V T E
BINARY_7233 T T Y K L I L N G K Q L K E E A I T E L V D A G I A E K Y F K L I A N A K T V E G V W T L K D E I L K A T V T E
BINARY_7234 T T Y K L I L N G K Q L K E E A I T E L V D A G I A E K Y F K L I A N A K T V E G V W T L K D E I L K F T V T E
BINARY_7235 T T Y K L I L N G K Q L K E E A I T E L V D A G I A E K Y F K L I A N A K T V E G V W T L K D E I L T A T V T E
BINARY_7236 T T Y K L I L N G K Q L K E E A I T E L V D A G I A E K Y F K L I A N A K T V E G V W T L K D E I L T F T V T E

BINARY_7237 T T Y K L I L N G K Q L K E E A I T E L V D A G I A E K Y F K L I A N A K T V E G V W T L K D E I K K A T V T E
BINARY_7238 T T Y K L I L N G K Q L K E E A I T E L V D A G I A E K Y F K L I A N A K T V E G V W T L K D E I K K F T V T E
BINARY_7239 T T Y K L I L N G K Q L K E E A I T E L V D A G I A E K Y F K L I A N A K T V E G V W T L K D E I K T A T V T E
BINARY_7240 T T Y K L I L N G K Q L K E E A I T E L V D A G I A E K Y F K L I A N A K T V E G V W T L K D E I K T F T V T E
BINARY_7241 T T Y K L I L N G K Q L K E E A I T E L V D A G I A E K Y F K L I A N A K T V E G V W T L K D E T L K A T V T E
BINARY_7242 T T Y K L I L N G K Q L K E E A I T E L V D A G I A E K Y F K L I A N A K T V E G V W T L K D E T L K F T V T E
BINARY_7243 T T Y K L I L N G K Q L K E E A I T E L V D A G I A E K Y F K L I A N A K T V E G V W T L K D E T L T A T V T E
BINARY_7244 T T Y K L I L N G K Q L K E E A I T E L V D A G I A E K Y F K L I A N A K T V E G V W T L K D E T L T F T V T E
BINARY_7245 T T Y K L I L N G K Q L K E E A I T E L V D A G I A E K Y F K L I A N A K T V E G V W T L K D E T K K A T V T E
BINARY_7246 T T Y K L I L N G K Q L K E E A I T E L V D A G I A E K Y F K L I A N A K T V E G V W T L K D E T K K F T V T E
BINARY_7247 T T Y K L I L N G K Q L K E E A I T E L V D A G I A E K Y F K L I A N A K T V E G V W T L K D E T K T A T V T E
BINARY_7248 T T Y K L I L N G K Q L K E E A I T E L V D A G I A E K Y F K L I A N A K T V E G V W T L K D E T K T F T V T E
BINARY_7249 T T Y K L I L N G K Q L K E E A I T E L V D A G I A E K Y F K L I A N A K T V E G V W T Y K D E I L K A T V T E
BINARY_7250 T T Y K L I L N G K Q L K E E A I T E L V D A G I A E K Y F K L I A N A K T V E G V W T Y K D E I L K F T V T E
BINARY_7251 T T Y K L I L N G K Q L K E E A I T E L V D A G I A E K Y F K L I A N A K T V E G V W T Y K D E I L T A T V T E
BINARY_7252 T T Y K L I L N G K Q L K E E A I T E L V D A G I A E K Y F K L I A N A K T V E G V W T Y K D E I L T F T V T E
BINARY_7253 T T Y K L I L N G K Q L K E E A I T E L V D A G I A E K Y F K L I A N A K T V E G V W T Y K D E I K K A T V T E
BINARY_7254 T T Y K L I L N G K Q L K E E A I T E L V D A G I A E K Y F K L I A N A K T V E G V W T Y K D E I K K F T V T E
BINARY_7255 T T Y K L I L N G K Q L K E E A I T E L V D A G I A E K Y F K L I A N A K T V E G V W T Y K D E I K T A T V T E
BINARY_7256 T T Y K L I L N G K Q L K E E A I T E L V D A G I A E K Y F K L I A N A K T V E G V W T Y K D E I K T F T V T E
BINARY_7257 T T Y K L I L N G K Q L K E E A I T E L V D A G I A E K Y F K L I A N A K T V E G V W T Y K D E T L K A T V T E
BINARY_7258 T T Y K L I L N G K Q L K E E A I T E L V D A G I A E K Y F K L I A N A K T V E G V W T Y K D E T L K F T V T E
BINARY_7259 T T Y K L I L N G K Q L K E E A I T E L V D A G I A E K Y F K L I A N A K T V E G V W T Y K D E T L T A T V T E
BINARY_7260 T T Y K L I L N G K Q L K E E A I T E L V D A G I A E K Y F K L I A N A K T V E G V W T Y K D E T L T F T V T E
BINARY_7261 T T Y K L I L N G K Q L K E E A I T E L V D A G I A E K Y F K L I A N A K T V E G V W T Y K D E T K K A T V T E
BINARY_7262 T T Y K L I L N G K Q L K E E A I T E L V D A G I A E K Y F K L I A N A K T V E G V W T Y K D E T K K F T V T E
BINARY_7263 T T Y K L I L N G K Q L K E E A I T E L V D A G I A E K Y F K L I A N A K T V E G V W T Y K D E T K T A T V T E
BINARY_7264 T T Y K L I L N G K Q L K E E A I T E L V D A G I A E K Y F K L I A N A K T V E G V W T Y K D E T K T F T V T E
BINARY_7265 T T Y K L I L N G K Q L K E E A I T E L V D A G I A E K Y F K L Y A N A K T V E G V W T L K D E I L K A T V T E
BINARY_7266 T T Y K L I L N G K Q L K E E A I T E L V D A G I A E K Y F K L Y A N A K T V E G V W T L K D E I L K F T V T E
BINARY_7267 T T Y K L I L N G K Q L K E E A I T E L V D A G I A E K Y F K L Y A N A K T V E G V W T L K D E I L T A T V T E
BINARY_7268 T T Y K L I L N G K Q L K E E A I T E L V D A G I A E K Y F K L Y A N A K T V E G V W T L K D E I L T F T V T E
BINARY_7269 T T Y K L I L N G K Q L K E E A I T E L V D A G I A E K Y F K L Y A N A K T V E G V W T L K D E I K K A T V T E
BINARY_7270 T T Y K L I L N G K Q L K E E A I T E L V D A G I A E K Y F K L Y A N A K T V E G V W T L K D E I K K F T V T E
BINARY_7271 T T Y K L I L N G K Q L K E E A I T E L V D A G I A E K Y F K L Y A N A K T V E G V W T L K D E I K T A T V T E
BINARY_7272 T T Y K L I L N G K Q L K E E A I T E L V D A G I A E K Y F K L Y A N A K T V E G V W T L K D E I K T F T V T E
BINARY_7273 T T Y K L I L N G K Q L K E E A I T E L V D A G I A E K Y F K L Y A N A K T V E G V W T L K D E T L K A T V T E
BINARY_7274 T T Y K L I L N G K Q L K E E A I T E L V D A G I A E K Y F K L Y A N A K T V E G V W T L K D E T L K F T V T E
BINARY_7275 T T Y K L I L N G K Q L K E E A I T E L V D A G I A E K Y F K L Y A N A K T V E G V W T L K D E T L T A T V T E
BINARY_7276 T T Y K L I L N G K Q L K E E A I T E L V D A G I A E K Y F K L Y A N A K T V E G V W T L K D E T L T F T V T E
BINARY_7277 T T Y K L I L N G K Q L K E E A I T E L V D A G I A E K Y F K L Y A N A K T V E G V W T L K D E T K K A T V T E
BINARY_7278 T T Y K L I L N G K Q L K E E A I T E L V D A G I A E K Y F K L Y A N A K T V E G V W T L K D E T K K F T V T E
BINARY_7279 T T Y K L I L N G K Q L K E E A I T E L V D A G I A E K Y F K L Y A N A K T V E G V W T L K D E T K T A T V T E
BINARY_7280 T T Y K L I L N G K Q L K E E A I T E L V D A G I A E K Y F K L Y A N A K T V E G V W T L K D E T K T F T V T E
BINARY_7281 T T Y K L I L N G K Q L K E E A I T E L V D A G I A E K Y F K L Y A N A K T V E G V W T Y K D E I L K A T V T E
BINARY_7282 T T Y K L I L N G K Q L K E E A I T E L V D A G I A E K Y F K L Y A N A K T V E G V W T Y K D E I L K F T V T E
BINARY_7283 T T Y K L I L N G K Q L K E E A I T E L V D A G I A E K Y F K L Y A N A K T V E G V W T Y K D E I L T A T V T E
BINARY_7284 T T Y K L I L N G K Q L K E E A I T E L V D A G I A E K Y F K L Y A N A K T V E G V W T Y K D E I L T F T V T E
BINARY_7285 T T Y K L I L N G K Q L K E E A I T E L V D A G I A E K Y F K L Y A N A K T V E G V W T Y K D E I K K A T V T E
BINARY_7286 T T Y K L I L N G K Q L K E E A I T E L V D A G I A E K Y F K L Y A N A K T V E G V W T Y K D E I K K F T V T E
BINARY_7287 T T Y K L I L N G K Q L K E E A I T E L V D A G I A E K Y F K L Y A N A K T V E G V W T Y K D E I K T A T V T E
BINARY_7288 T T Y K L I L N G K Q L K E E A I T E L V D A G I A E K Y F K L Y A N A K T V E G V W T Y K D E I K T F T V T E
BINARY_7289 T T Y K L I L N G K Q L K E E A I T E L V D A G I A E K Y F K L Y A N A K T V E G V W T Y K D E T L K A T V T E
BINARY_7290 T T Y K L I L N G K Q L K E E A I T E L V D A G I A E K Y F K L Y A N A K T V E G V W T Y K D E T L K F T V T E
BINARY_7291 T T Y K L I L N G K Q L K E E A I T E L V D A G I A E K Y F K L Y A N A K T V E G V W T Y K D E T L T A T V T E
BINARY_7292 T T Y K L I L N G K Q L K E E A I T E L V D A G I A E K Y F K L Y A N A K T V E G V W T Y K D E T L T F T V T E
BINARY_7293 T T Y K L I L N G K Q L K E E A I T E L V D A G I A E K Y F K L Y A N A K T V E G V W T Y K D E T K K A T V T E
BINARY_7294 T T Y K L I L N G K Q L K E E A I T E L V D A G I A E K Y F K L Y A N A K T V E G V W T Y K D E T K K F T V T E
BINARY_7295 T T Y K L I L N G K Q L K E E A I T E L V D A G I A E K Y F K L Y A N A K T V E G V W T Y K D E T K T A T V T E
BINARY_7296 T T Y K L I L N G K Q L K E E A I T E L V D A G I A E K Y F K L Y A N A K T V E G V W T Y K D E T K T F T V T E
BINARY_7297 T T Y K L I L N G K Q L K E E A I T E L V D A G T A E K Y I K L I A N A K T V E G V W T L K D E I L K A T V T E
BINARY_7298 T T Y K L I L N G K Q L K E E A I T E L V D A G T A E K Y I K L I A N A K T V E G V W T L K D E I L K F T V T E
BINARY_7299 T T Y K L I L N G K Q L K E E A I T E L V D A G T A E K Y I K L I A N A K T V E G V W T L K D E I L T A T V T E
BINARY_7300 T T Y K L I L N G K Q L K E E A I T E L V D A G T A E K Y I K L I A N A K T V E G V W T L K D E I L T F T V T E
BINARY_7301 T T Y K L I L N G K Q L K E E A I T E L V D A G T A E K Y I K L I A N A K T V E G V W T L K D E I K K A T V T E
BINARY_7302 T T Y K L I L N G K Q L K E E A I T E L V D A G T A E K Y I K L I A N A K T V E G V W T L K D E I K K F T V T E
BINARY_7303 T T Y K L I L N G K Q L K E E A I T E L V D A G T A E K Y I K L I A N A K T V E G V W T L K D E I K T A T V T E
BINARY_7304 T T Y K L I L N G K Q L K E E A I T E L V D A G T A E K Y I K L I A N A K T V E G V W T L K D E I K T F T V T E
BINARY_7305 T T Y K L I L N G K Q L K E E A I T E L V D A G T A E K Y I K L I A N A K T V E G V W T L K D E T L K A T V T E
BINARY_7306 T T Y K L I L N G K Q L K E E A I T E L V D A G T A E K Y I K L I A N A K T V E G V W T L K D E T L K F T V T E
BINARY_7307 T T Y K L I L N G K Q L K E E A I T E L V D A G T A E K Y I K L I A N A K T V E G V W T L K D E T L T A T V T E
BINARY_7308 T T Y K L I L N G K Q L K E E A I T E L V D A G T A E K Y I K L I A N A K T V E G V W T L K D E T L T F T V T E
BINARY_7309 T T Y K L I L N G K Q L K E E A I T E L V D A G T A E K Y I K L I A N A K T V E G V W T L K D E T K K A T V T E
BINARY_7310 T T Y K L I L N G K Q L K E E A I T E L V D A G T A E K Y I K L I A N A K T V E G V W T L K D E T K K F T V T E
BINARY_7311 T T Y K L I L N G K Q L K E E A I T E L V D A G T A E K Y I K L I A N A K T V E G V W T L K D E T K T A T V T E
BINARY_7312 T T Y K L I L N G K Q L K E E A I T E L V D A G T A E K Y I K L I A N A K T V E G V W T L K D E T K T F T V T E
BINARY_7313 T T Y K L I L N G K Q L K E E A I T E L V D A G T A E K Y I K L I A N A K T V E G V W T Y K D E I L K A T V T E
BINARY_7314 T T Y K L I L N G K Q L K E E A I T E L V D A G T A E K Y I K L I A N A K T V E G V W T Y K D E I L K F T V T E
BINARY_7315 T T Y K L I L N G K Q L K E E A I T E L V D A G T A E K Y I K L I A N A K T V E G V W T Y K D E I L T A T V T E
BINARY_7316 T T Y K L I L N G K Q L K E E A I T E L V D A G T A E K Y I K L I A N A K T V E G V W T Y K D E I L T F T V T E
BINARY_7317 T T Y K L I L N G K Q L K E E A I T E L V D A G T A E K Y I K L I A N A K T V E G V W T Y K D E I K K A T V T E
BINARY_7318 T T Y K L I L N G K Q L K E E A I T E L V D A G T A E K Y I K L I A N A K T V E G V W T Y K D E I K K F T V T E
BINARY_7319 T T Y K L I L N G K Q L K E E A I T E L V D A G T A E K Y I K L I A N A K T V E G V W T Y K D E I K T A T V T E
BINARY_7320 T T Y K L I L N G K Q L K E E A I T E L V D A G T A E K Y I K L I A N A K T V E G V W T Y K D E I K T F T V T E
BINARY_7321 T T Y K L I L N G K Q L K E E A I T E L V D A G T A E K Y I K L I A N A K T V E G V W T Y K D E T L K A T V T E
BINARY_7322 T T Y K L I L N G K Q L K E E A I T E L V D A G T A E K Y I K L I A N A K T V E G V W T Y K D E T L K F T V T E
BINARY_7323 T T Y K L I L N G K Q L K E E A I T E L V D A G T A E K Y I K L I A N A K T V E G V W T Y K D E T L T A T V T E
BINARY_7324 T T Y K L I L N G K Q L K E E A I T E L V D A G T A E K Y I K L I A N A K T V E G V W T Y K D E T L T F T V T E
BINARY_7325 T T Y K L I L N G K Q L K E E A I T E L V D A G T A E K Y I K L I A N A K T V E G V W T Y K D E T K K A T V T E
BINARY_7326 T T Y K L I L N G K Q L K E E A I T E L V D A G T A E K Y I K L I A N A K T V E G V W T Y K D E T K K F T V T E
BINARY_7327 T T Y K L I L N G K Q L K E E A I T E L V D A G T A E K Y I K L I A N A K T V E G V W T Y K D E T K T A T V T E
BINARY_7328 T T Y K L I L N G K Q L K E E A I T E L V D A G T A E K Y I K L I A N A K T V E G V W T Y K D E T K T F T V T E
BINARY_7329 T T Y K L I L N G K Q L K E E A I T E L V D A G T A E K Y I K L Y A N A K T V E G V W T L K D E I L K A T V T E
BINARY_7330 T T Y K L I L N G K Q L K E E A I T E L V D A G T A E K Y I K L Y A N A K T V E G V W T L K D E I L K F T V T E
BINARY_7331 T T Y K L I L N G K Q L K E E A I T E L V D A G T A E K Y I K L Y A N A K T V E G V W T L K D E I L T A T V T E

BINARY_7332 T T Y K L I L N G K Q L K E E A I T E L V D A G T A E K Y I K L Y A N A K T V E G V W T L K D E I L T F T V T E
BINARY_7333 T T Y K L I L N G K Q L K E E A I T E L V D A G T A E K Y I K L Y A N A K T V E G V W T L K D E I K K A T V T E
BINARY_7334 T T Y K L I L N G K Q L K E E A I T E L V D A G T A E K Y I K L Y A N A K T V E G V W T L K D E I K K F T V T E
BINARY_7335 T T Y K L I L N G K Q L K E E A I T E L V D A G T A E K Y I K L Y A N A K T V E G V W T L K D E I K T A T V T E
BINARY_7336 T T Y K L I L N G K Q L K E E A I T E L V D A G T A E K Y I K L Y A N A K T V E G V W T L K D E I K T F T V T E
BINARY_7337 T T Y K L I L N G K Q L K E E A I T E L V D A G T A E K Y I K L Y A N A K T V E G V W T L K D E T L K A T V T E
BINARY_7338 T T Y K L I L N G K Q L K E E A I T E L V D A G T A E K Y I K L Y A N A K T V E G V W T L K D E T L K F T V T E
BINARY_7339 T T Y K L I L N G K Q L K E E A I T E L V D A G T A E K Y I K L Y A N A K T V E G V W T L K D E T L T A T V T E
BINARY_7340 T T Y K L I L N G K Q L K E E A I T E L V D A G T A E K Y I K L Y A N A K T V E G V W T L K D E T L T F T V T E
BINARY_7341 T T Y K L I L N G K Q L K E E A I T E L V D A G T A E K Y I K L Y A N A K T V E G V W T L K D E T K K A T V T E
BINARY_7342 T T Y K L I L N G K Q L K E E A I T E L V D A G T A E K Y I K L Y A N A K T V E G V W T L K D E T K K F T V T E
BINARY_7343 T T Y K L I L N G K Q L K E E A I T E L V D A G T A E K Y I K L Y A N A K T V E G V W T L K D E T K T A T V T E
BINARY_7344 T T Y K L I L N G K Q L K E E A I T E L V D A G T A E K Y I K L Y A N A K T V E G V W T L K D E T K T F T V T E
BINARY_7345 T T Y K L I L N G K Q L K E E A I T E L V D A G T A E K Y I K L Y A N A K T V E G V W T Y K D E I L K A T V T E
BINARY_7346 T T Y K L I L N G K Q L K E E A I T E L V D A G T A E K Y I K L Y A N A K T V E G V W T Y K D E I L K F T V T E
BINARY_7347 T T Y K L I L N G K Q L K E E A I T E L V D A G T A E K Y I K L Y A N A K T V E G V W T Y K D E I L T A T V T E
BINARY_7348 T T Y K L I L N G K Q L K E E A I T E L V D A G T A E K Y I K L Y A N A K T V E G V W T Y K D E I L T F T V T E
BINARY_7349 T T Y K L I L N G K Q L K E E A I T E L V D A G T A E K Y I K L Y A N A K T V E G V W T Y K D E I K K A T V T E
BINARY_7350 T T Y K L I L N G K Q L K E E A I T E L V D A G T A E K Y I K L Y A N A K T V E G V W T Y K D E I K K F T V T E
BINARY_7351 T T Y K L I L N G K Q L K E E A I T E L V D A G T A E K Y I K L Y A N A K T V E G V W T Y K D E I K T A T V T E
BINARY_7352 T T Y K L I L N G K Q L K E E A I T E L V D A G T A E K Y I K L Y A N A K T V E G V W T Y K D E I K T F T V T E
BINARY_7353 T T Y K L I L N G K Q L K E E A I T E L V D A G T A E K Y I K L Y A N A K T V E G V W T Y K D E T L K A T V T E
BINARY_7354 T T Y K L I L N G K Q L K E E A I T E L V D A G T A E K Y I K L Y A N A K T V E G V W T Y K D E T L K F T V T E
BINARY_7355 T T Y K L I L N G K Q L K E E A I T E L V D A G T A E K Y I K L Y A N A K T V E G V W T Y K D E T L T A T V T E
BINARY_7356 T T Y K L I L N G K Q L K E E A I T E L V D A G T A E K Y I K L Y A N A K T V E G V W T Y K D E T L T F T V T E
BINARY_7357 T T Y K L I L N G K Q L K E E A I T E L V D A G T A E K Y I K L Y A N A K T V E G V W T Y K D E T K K A T V T E
BINARY_7358 T T Y K L I L N G K Q L K E E A I T E L V D A G T A E K Y I K L Y A N A K T V E G V W T Y K D E T K K F T V T E
BINARY_7359 T T Y K L I L N G K Q L K E E A I T E L V D A G T A E K Y I K L Y A N A K T V E G V W T Y K D E T K T A T V T E
BINARY_7360 T T Y K L I L N G K Q L K E E A I T E L V D A G T A E K Y I K L Y A N A K T V E G V W T Y K D E T K T F T V T E
BINARY_7361 T T Y K L I L N G K Q L K E E A I T E L V D A G T A E K Y F K L I A N A K T V E G V W T L K D E I L K A T V T E
BINARY_7362 T T Y K L I L N G K Q L K E E A I T E L V D A G T A E K Y F K L I A N A K T V E G V W T L K D E I L K F T V T E
BINARY_7363 T T Y K L I L N G K Q L K E E A I T E L V D A G T A E K Y F K L I A N A K T V E G V W T L K D E I L T A T V T E
BINARY_7364 T T Y K L I L N G K Q L K E E A I T E L V D A G T A E K Y F K L I A N A K T V E G V W T L K D E I L T F T V T E
BINARY_7365 T T Y K L I L N G K Q L K E E A I T E L V D A G T A E K Y F K L I A N A K T V E G V W T L K D E I K K A T V T E
BINARY_7366 T T Y K L I L N G K Q L K E E A I T E L V D A G T A E K Y F K L I A N A K T V E G V W T L K D E I K K F T V T E
BINARY_7367 T T Y K L I L N G K Q L K E E A I T E L V D A G T A E K Y F K L I A N A K T V E G V W T L K D E I K T A T V T E
BINARY_7368 T T Y K L I L N G K Q L K E E A I T E L V D A G T A E K Y F K L I A N A K T V E G V W T L K D E I K T F T V T E
BINARY_7369 T T Y K L I L N G K Q L K E E A I T E L V D A G T A E K Y F K L I A N A K T V E G V W T L K D E T L K A T V T E
BINARY_7370 T T Y K L I L N G K Q L K E E A I T E L V D A G T A E K Y F K L I A N A K T V E G V W T L K D E T L K F T V T E
BINARY_7371 T T Y K L I L N G K Q L K E E A I T E L V D A G T A E K Y F K L I A N A K T V E G V W T L K D E T L T A T V T E
BINARY_7372 T T Y K L I L N G K Q L K E E A I T E L V D A G T A E K Y F K L I A N A K T V E G V W T L K D E T L T F T V T E
BINARY_7373 T T Y K L I L N G K Q L K E E A I T E L V D A G T A E K Y F K L I A N A K T V E G V W T L K D E T K K A T V T E
BINARY_7374 T T Y K L I L N G K Q L K E E A I T E L V D A G T A E K Y F K L I A N A K T V E G V W T L K D E T K K F T V T E
BINARY_7375 T T Y K L I L N G K Q L K E E A I T E L V D A G T A E K Y F K L I A N A K T V E G V W T L K D E T K T A T V T E
BINARY_7376 T T Y K L I L N G K Q L K E E A I T E L V D A G T A E K Y F K L I A N A K T V E G V W T L K D E T K T F T V T E
BINARY_7377 T T Y K L I L N G K Q L K E E A I T E L V D A G T A E K Y F K L I A N A K T V E G V W T Y K D E I L K A T V T E
BINARY_7378 T T Y K L I L N G K Q L K E E A I T E L V D A G T A E K Y F K L I A N A K T V E G V W T Y K D E I L K F T V T E
BINARY_7379 T T Y K L I L N G K Q L K E E A I T E L V D A G T A E K Y F K L I A N A K T V E G V W T Y K D E I L T A T V T E
BINARY_7380 T T Y K L I L N G K Q L K E E A I T E L V D A G T A E K Y F K L I A N A K T V E G V W T Y K D E I L T F T V T E
BINARY_7381 T T Y K L I L N G K Q L K E E A I T E L V D A G T A E K Y F K L I A N A K T V E G V W T Y K D E I K K A T V T E
BINARY_7382 T T Y K L I L N G K Q L K E E A I T E L V D A G T A E K Y F K L I A N A K T V E G V W T Y K D E I K K F T V T E
BINARY_7383 T T Y K L I L N G K Q L K E E A I T E L V D A G T A E K Y F K L I A N A K T V E G V W T Y K D E I K T A T V T E
BINARY_7384 T T Y K L I L N G K Q L K E E A I T E L V D A G T A E K Y F K L I A N A K T V E G V W T Y K D E I K T F T V T E
BINARY_7385 T T Y K L I L N G K Q L K E E A I T E L V D A G T A E K Y F K L I A N A K T V E G V W T Y K D E T L K A T V T E
BINARY_7386 T T Y K L I L N G K Q L K E E A I T E L V D A G T A E K Y F K L I A N A K T V E G V W T Y K D E T L K F T V T E
BINARY_7387 T T Y K L I L N G K Q L K E E A I T E L V D A G T A E K Y F K L I A N A K T V E G V W T Y K D E T L T A T V T E
BINARY_7388 T T Y K L I L N G K Q L K E E A I T E L V D A G T A E K Y F K L I A N A K T V E G V W T Y K D E T L T F T V T E
BINARY_7389 T T Y K L I L N G K Q L K E E A I T E L V D A G T A E K Y F K L I A N A K T V E G V W T Y K D E T K K A T V T E
BINARY_7390 T T Y K L I L N G K Q L K E E A I T E L V D A G T A E K Y F K L I A N A K T V E G V W T Y K D E T K K F T V T E
BINARY_7391 T T Y K L I L N G K Q L K E E A I T E L V D A G T A E K Y F K L I A N A K T V E G V W T Y K D E T K T A T V T E
BINARY_7392 T T Y K L I L N G K Q L K E E A I T E L V D A G T A E K Y F K L I A N A K T V E G V W T Y K D E T K T F T V T E
BINARY_7393 T T Y K L I L N G K Q L K E E A I T E L V D A G T A E K Y F K L Y A N A K T V E G V W T L K D E I L K A T V T E
BINARY_7394 T T Y K L I L N G K Q L K E E A I T E L V D A G T A E K Y F K L Y A N A K T V E G V W T L K D E I L K F T V T E
BINARY_7395 T T Y K L I L N G K Q L K E E A I T E L V D A G T A E K Y F K L Y A N A K T V E G V W T L K D E I L T A T V T E
BINARY_7396 T T Y K L I L N G K Q L K E E A I T E L V D A G T A E K Y F K L Y A N A K T V E G V W T L K D E I L T F T V T E
BINARY_7397 T T Y K L I L N G K Q L K E E A I T E L V D A G T A E K Y F K L Y A N A K T V E G V W T L K D E I K K A T V T E
BINARY_7398 T T Y K L I L N G K Q L K E E A I T E L V D A G T A E K Y F K L Y A N A K T V E G V W T L K D E I K K F T V T E
BINARY_7399 T T Y K L I L N G K Q L K E E A I T E L V D A G T A E K Y F K L Y A N A K T V E G V W T L K D E I K T A T V T E
BINARY_7400 T T Y K L I L N G K Q L K E E A I T E L V D A G T A E K Y F K L Y A N A K T V E G V W T L K D E I K T F T V T E
BINARY_7401 T T Y K L I L N G K Q L K E E A I T E L V D A G T A E K Y F K L Y A N A K T V E G V W T L K D E T L K A T V T E
BINARY_7402 T T Y K L I L N G K Q L K E E A I T E L V D A G T A E K Y F K L Y A N A K T V E G V W T L K D E T L K F T V T E
BINARY_7403 T T Y K L I L N G K Q L K E E A I T E L V D A G T A E K Y F K L Y A N A K T V E G V W T L K D E T L T A T V T E
BINARY_7404 T T Y K L I L N G K Q L K E E A I T E L V D A G T A E K Y F K L Y A N A K T V E G V W T L K D E T L T F T V T E
BINARY_7405 T T Y K L I L N G K Q L K E E A I T E L V D A G T A E K Y F K L Y A N A K T V E G V W T L K D E T K K A T V T E
BINARY_7406 T T Y K L I L N G K Q L K E E A I T E L V D A G T A E K Y F K L Y A N A K T V E G V W T L K D E T K K F T V T E
BINARY_7407 T T Y K L I L N G K Q L K E E A I T E L V D A G T A E K Y F K L Y A N A K T V E G V W T L K D E T K T A T V T E
BINARY_7408 T T Y K L I L N G K Q L K E E A I T E L V D A G T A E K Y F K L Y A N A K T V E G V W T L K D E T K T F T V T E
BINARY_7409 T T Y K L I L N G K Q L K E E A I T E L V D A G T A E K Y F K L Y A N A K T V E G V W T Y K D E I L K A T V T E
BINARY_7410 T T Y K L I L N G K Q L K E E A I T E L V D A G T A E K Y F K L Y A N A K T V E G V W T Y K D E I L K F T V T E
BINARY_7411 T T Y K L I L N G K Q L K E E A I T E L V D A G T A E K Y F K L Y A N A K T V E G V W T Y K D E I L T A T V T E
BINARY_7412 T T Y K L I L N G K Q L K E E A I T E L V D A G T A E K Y F K L Y A N A K T V E G V W T Y K D E I L T F T V T E
BINARY_7413 T T Y K L I L N G K Q L K E E A I T E L V D A G T A E K Y F K L Y A N A K T V E G V W T Y K D E I K K A T V T E
BINARY_7414 T T Y K L I L N G K Q L K E E A I T E L V D A G T A E K Y F K L Y A N A K T V E G V W T Y K D E I K K F T V T E
BINARY_7415 T T Y K L I L N G K Q L K E E A I T E L V D A G T A E K Y F K L Y A N A K T V E G V W T Y K D E I K T A T V T E
BINARY_7416 T T Y K L I L N G K Q L K E E A I T E L V D A G T A E K Y F K L Y A N A K T V E G V W T Y K D E I K T F T V T E
BINARY_7417 T T Y K L I L N G K Q L K E E A I T E L V D A G T A E K Y F K L Y A N A K T V E G V W T Y K D E T L K A T V T E
BINARY_7418 T T Y K L I L N G K Q L K E E A I T E L V D A G T A E K Y F K L Y A N A K T V E G V W T Y K D E T L K F T V T E
BINARY_7419 T T Y K L I L N G K Q L K E E A I T E L V D A G T A E K Y F K L Y A N A K T V E G V W T Y K D E T L T A T V T E
BINARY_7420 T T Y K L I L N G K Q L K E E A I T E L V D A G T A E K Y F K L Y A N A K T V E G V W T Y K D E T L T F T V T E
BINARY_7421 T T Y K L I L N G K Q L K E E A I T E L V D A G T A E K Y F K L Y A N A K T V E G V W T Y K D E T K K A T V T E
BINARY_7422 T T Y K L I L N G K Q L K E E A I T E L V D A G T A E K Y F K L Y A N A K T V E G V W T Y K D E T K K F T V T E
BINARY_7423 T T Y K L I L N G K Q L K E E A I T E L V D A G T A E K Y F K L Y A N A K T V E G V W T Y K D E T K T A T V T E
BINARY_7424 T T Y K L I L N G K Q L K E E A I T E L V D A G T A E K Y F K L Y A N A K T V E G V W T Y K D E T K T F T V T E
BINARY_7425 T T Y K L I L N G K Q L K E E A I T E L V D A A I A E K Y I K L I A N A K T V E G V W T L K D E I L K A T V T E
BINARY_7426 T T Y K L I L N G K Q L K E E A I T E L V D A A I A E K Y I K L I A N A K T V E G V W T L K D E I L K F T V T E

BINARY_7427 T T Y K L I L N G K Q L K E E A I T E L V D A A I A E K Y I K L I A N A K T V E G V W T L K D E I L T A T V T E
BINARY_7428 T T Y K L I L N G K Q L K E E A I T E L V D A A I A E K Y I K L I A N A K T V E G V W T L K D E I L T F T V T E
BINARY_7429 T T Y K L I L N G K Q L K E E A I T E L V D A A I A E K Y I K L I A N A K T V E G V W T L K D E I K K A T V T E
BINARY_7430 T T Y K L I L N G K Q L K E E A I T E L V D A A I A E K Y I K L I A N A K T V E G V W T L K D E I K K F T V T E
BINARY_7431 T T Y K L I L N G K Q L K E E A I T E L V D A A I A E K Y I K L I A N A K T V E G V W T L K D E I K T A T V T E
BINARY_7432 T T Y K L I L N G K Q L K E E A I T E L V D A A I A E K Y I K L I A N A K T V E G V W T L K D E I K T F T V T E
BINARY_7433 T T Y K L I L N G K Q L K E E A I T E L V D A A I A E K Y I K L I A N A K T V E G V W T L K D E T L K A T V T E
BINARY_7434 T T Y K L I L N G K Q L K E E A I T E L V D A A I A E K Y I K L I A N A K T V E G V W T L K D E T L K F T V T E
BINARY_7435 T T Y K L I L N G K Q L K E E A I T E L V D A A I A E K Y I K L I A N A K T V E G V W T L K D E T L T A T V T E
BINARY_7436 T T Y K L I L N G K Q L K E E A I T E L V D A A I A E K Y I K L I A N A K T V E G V W T L K D E T L T F T V T E
BINARY_7437 T T Y K L I L N G K Q L K E E A I T E L V D A A I A E K Y I K L I A N A K T V E G V W T L K D E T K K A T V T E
BINARY_7438 T T Y K L I L N G K Q L K E E A I T E L V D A A I A E K Y I K L I A N A K T V E G V W T L K D E T K K F T V T E
BINARY_7439 T T Y K L I L N G K Q L K E E A I T E L V D A A I A E K Y I K L I A N A K T V E G V W T L K D E T K T A T V T E
BINARY_7440 T T Y K L I L N G K Q L K E E A I T E L V D A A I A E K Y I K L I A N A K T V E G V W T L K D E T K T F T V T E
BINARY_7441 T T Y K L I L N G K Q L K E E A I T E L V D A A I A E K Y I K L I A N A K T V E G V W T Y K D E I L K A T V T E
BINARY_7442 T T Y K L I L N G K Q L K E E A I T E L V D A A I A E K Y I K L I A N A K T V E G V W T Y K D E I L K F T V T E
BINARY_7443 T T Y K L I L N G K Q L K E E A I T E L V D A A I A E K Y I K L I A N A K T V E G V W T Y K D E I L T A T V T E
BINARY_7444 T T Y K L I L N G K Q L K E E A I T E L V D A A I A E K Y I K L I A N A K T V E G V W T Y K D E I L T F T V T E
BINARY_7445 T T Y K L I L N G K Q L K E E A I T E L V D A A I A E K Y I K L I A N A K T V E G V W T Y K D E I K K A T V T E
BINARY_7446 T T Y K L I L N G K Q L K E E A I T E L V D A A I A E K Y I K L I A N A K T V E G V W T Y K D E I K K F T V T E
BINARY_7447 T T Y K L I L N G K Q L K E E A I T E L V D A A I A E K Y I K L I A N A K T V E G V W T Y K D E I K T A T V T E
BINARY_7448 T T Y K L I L N G K Q L K E E A I T E L V D A A I A E K Y I K L I A N A K T V E G V W T Y K D E I K T F T V T E
BINARY_7449 T T Y K L I L N G K Q L K E E A I T E L V D A A I A E K Y I K L I A N A K T V E G V W T Y K D E T L K A T V T E
BINARY_7450 T T Y K L I L N G K Q L K E E A I T E L V D A A I A E K Y I K L I A N A K T V E G V W T Y K D E T L K F T V T E
BINARY_7451 T T Y K L I L N G K Q L K E E A I T E L V D A A I A E K Y I K L I A N A K T V E G V W T Y K D E T L T A T V T E
BINARY_7452 T T Y K L I L N G K Q L K E E A I T E L V D A A I A E K Y I K L I A N A K T V E G V W T Y K D E T L T F T V T E
BINARY_7453 T T Y K L I L N G K Q L K E E A I T E L V D A A I A E K Y I K L I A N A K T V E G V W T Y K D E T K K A T V T E
BINARY_7454 T T Y K L I L N G K Q L K E E A I T E L V D A A I A E K Y I K L I A N A K T V E G V W T Y K D E T K K F T V T E
BINARY_7455 T T Y K L I L N G K Q L K E E A I T E L V D A A I A E K Y I K L I A N A K T V E G V W T Y K D E T K T A T V T E
BINARY_7456 T T Y K L I L N G K Q L K E E A I T E L V D A A I A E K Y I K L I A N A K T V E G V W T Y K D E T K T F T V T E
BINARY_7457 T T Y K L I L N G K Q L K E E A I T E L V D A A I A E K Y I K L Y A N A K T V E G V W T L K D E I L K A T V T E
BINARY_7458 T T Y K L I L N G K Q L K E E A I T E L V D A A I A E K Y I K L Y A N A K T V E G V W T L K D E I L K F T V T E
BINARY_7459 T T Y K L I L N G K Q L K E E A I T E L V D A A I A E K Y I K L Y A N A K T V E G V W T L K D E I L T A T V T E
BINARY_7460 T T Y K L I L N G K Q L K E E A I T E L V D A A I A E K Y I K L Y A N A K T V E G V W T L K D E I L T F T V T E
BINARY_7461 T T Y K L I L N G K Q L K E E A I T E L V D A A I A E K Y I K L Y A N A K T V E G V W T L K D E I K K A T V T E
BINARY_7462 T T Y K L I L N G K Q L K E E A I T E L V D A A I A E K Y I K L Y A N A K T V E G V W T L K D E I K K F T V T E
BINARY_7463 T T Y K L I L N G K Q L K E E A I T E L V D A A I A E K Y I K L Y A N A K T V E G V W T L K D E I K T A T V T E
BINARY_7464 T T Y K L I L N G K Q L K E E A I T E L V D A A I A E K Y I K L Y A N A K T V E G V W T L K D E I K T F T V T E
BINARY_7465 T T Y K L I L N G K Q L K E E A I T E L V D A A I A E K Y I K L Y A N A K T V E G V W T L K D E T L K A T V T E
BINARY_7466 T T Y K L I L N G K Q L K E E A I T E L V D A A I A E K Y I K L Y A N A K T V E G V W T L K D E T L K F T V T E
BINARY_7467 T T Y K L I L N G K Q L K E E A I T E L V D A A I A E K Y I K L Y A N A K T V E G V W T L K D E T L T A T V T E
BINARY_7468 T T Y K L I L N G K Q L K E E A I T E L V D A A I A E K Y I K L Y A N A K T V E G V W T L K D E T L T F T V T E
BINARY_7469 T T Y K L I L N G K Q L K E E A I T E L V D A A I A E K Y I K L Y A N A K T V E G V W T L K D E T K K A T V T E
BINARY_7470 T T Y K L I L N G K Q L K E E A I T E L V D A A I A E K Y I K L Y A N A K T V E G V W T L K D E T K K F T V T E
BINARY_7471 T T Y K L I L N G K Q L K E E A I T E L V D A A I A E K Y I K L Y A N A K T V E G V W T L K D E T K T A T V T E
BINARY_7472 T T Y K L I L N G K Q L K E E A I T E L V D A A I A E K Y I K L Y A N A K T V E G V W T L K D E T K T F T V T E
BINARY_7473 T T Y K L I L N G K Q L K E E A I T E L V D A A I A E K Y I K L Y A N A K T V E G V W T Y K D E I L K A T V T E
BINARY_7474 T T Y K L I L N G K Q L K E E A I T E L V D A A I A E K Y I K L Y A N A K T V E G V W T Y K D E I L K F T V T E
BINARY_7475 T T Y K L I L N G K Q L K E E A I T E L V D A A I A E K Y I K L Y A N A K T V E G V W T Y K D E I L T A T V T E
BINARY_7476 T T Y K L I L N G K Q L K E E A I T E L V D A A I A E K Y I K L Y A N A K T V E G V W T Y K D E I L T F T V T E
BINARY_7477 T T Y K L I L N G K Q L K E E A I T E L V D A A I A E K Y I K L Y A N A K T V E G V W T Y K D E I K K A T V T E
BINARY_7478 T T Y K L I L N G K Q L K E E A I T E L V D A A I A E K Y I K L Y A N A K T V E G V W T Y K D E I K K F T V T E
BINARY_7479 T T Y K L I L N G K Q L K E E A I T E L V D A A I A E K Y I K L Y A N A K T V E G V W T Y K D E I K T A T V T E
BINARY_7480 T T Y K L I L N G K Q L K E E A I T E L V D A A I A E K Y I K L Y A N A K T V E G V W T Y K D E I K T F T V T E
BINARY_7481 T T Y K L I L N G K Q L K E E A I T E L V D A A I A E K Y I K L Y A N A K T V E G V W T Y K D E T L K A T V T E
BINARY_7482 T T Y K L I L N G K Q L K E E A I T E L V D A A I A E K Y I K L Y A N A K T V E G V W T Y K D E T L K F T V T E
BINARY_7483 T T Y K L I L N G K Q L K E E A I T E L V D A A I A E K Y I K L Y A N A K T V E G V W T Y K D E T L T A T V T E
BINARY_7484 T T Y K L I L N G K Q L K E E A I T E L V D A A I A E K Y I K L Y A N A K T V E G V W T Y K D E T L T F T V T E
BINARY_7485 T T Y K L I L N G K Q L K E E A I T E L V D A A I A E K Y I K L Y A N A K T V E G V W T Y K D E T K K A T V T E
BINARY_7486 T T Y K L I L N G K Q L K E E A I T E L V D A A I A E K Y I K L Y A N A K T V E G V W T Y K D E T K K F T V T E
BINARY_7487 T T Y K L I L N G K Q L K E E A I T E L V D A A I A E K Y I K L Y A N A K T V E G V W T Y K D E T K T A T V T E
BINARY_7488 T T Y K L I L N G K Q L K E E A I T E L V D A A I A E K Y I K L Y A N A K T V E G V W T Y K D E T K T F T V T E
BINARY_7489 T T Y K L I L N G K Q L K E E A I T E L V D A A I A E K Y F K L I A N A K T V E G V W T L K D E I L K A T V T E
BINARY_7490 T T Y K L I L N G K Q L K E E A I T E L V D A A I A E K Y F K L I A N A K T V E G V W T L K D E I L K F T V T E
BINARY_7491 T T Y K L I L N G K Q L K E E A I T E L V D A A I A E K Y F K L I A N A K T V E G V W T L K D E I L T A T V T E
BINARY_7492 T T Y K L I L N G K Q L K E E A I T E L V D A A I A E K Y F K L I A N A K T V E G V W T L K D E I L T F T V T E
BINARY_7493 T T Y K L I L N G K Q L K E E A I T E L V D A A I A E K Y F K L I A N A K T V E G V W T L K D E I K K A T V T E
BINARY_7494 T T Y K L I L N G K Q L K E E A I T E L V D A A I A E K Y F K L I A N A K T V E G V W T L K D E I K K F T V T E
BINARY_7495 T T Y K L I L N G K Q L K E E A I T E L V D A A I A E K Y F K L I A N A K T V E G V W T L K D E I K T A T V T E
BINARY_7496 T T Y K L I L N G K Q L K E E A I T E L V D A A I A E K Y F K L I A N A K T V E G V W T L K D E I K T F T V T E
BINARY_7497 T T Y K L I L N G K Q L K E E A I T E L V D A A I A E K Y F K L I A N A K T V E G V W T L K D E T L K A T V T E
BINARY_7498 T T Y K L I L N G K Q L K E E A I T E L V D A A I A E K Y F K L I A N A K T V E G V W T L K D E T L K F T V T E
BINARY_7499 T T Y K L I L N G K Q L K E E A I T E L V D A A I A E K Y F K L I A N A K T V E G V W T L K D E T L T A T V T E
BINARY_7500 T T Y K L I L N G K Q L K E E A I T E L V D A A I A E K Y F K L I A N A K T V E G V W T L K D E T L T F T V T E
BINARY_7501 T T Y K L I L N G K Q L K E E A I T E L V D A A I A E K Y F K L I A N A K T V E G V W T L K D E T K K A T V T E
BINARY_7502 T T Y K L I L N G K Q L K E E A I T E L V D A A I A E K Y F K L I A N A K T V E G V W T L K D E T K K F T V T E
BINARY_7503 T T Y K L I L N G K Q L K E E A I T E L V D A A I A E K Y F K L I A N A K T V E G V W T L K D E T K T A T V T E
BINARY_7504 T T Y K L I L N G K Q L K E E A I T E L V D A A I A E K Y F K L I A N A K T V E G V W T L K D E T K T F T V T E
BINARY_7505 T T Y K L I L N G K Q L K E E A I T E L V D A A I A E K Y F K L I A N A K T V E G V W T Y K D E I L K A T V T E
BINARY_7506 T T Y K L I L N G K Q L K E E A I T E L V D A A I A E K Y F K L I A N A K T V E G V W T Y K D E I L K F T V T E
BINARY_7507 T T Y K L I L N G K Q L K E E A I T E L V D A A I A E K Y F K L I A N A K T V E G V W T Y K D E I L T A T V T E
BINARY_7508 T T Y K L I L N G K Q L K E E A I T E L V D A A I A E K Y F K L I A N A K T V E G V W T Y K D E I L T F T V T E
BINARY_7509 T T Y K L I L N G K Q L K E E A I T E L V D A A I A E K Y F K L I A N A K T V E G V W T Y K D E I K K A T V T E
BINARY_7510 T T Y K L I L N G K Q L K E E A I T E L V D A A I A E K Y F K L I A N A K T V E G V W T Y K D E I K K F T V T E
BINARY_7511 T T Y K L I L N G K Q L K E E A I T E L V D A A I A E K Y F K L I A N A K T V E G V W T Y K D E I K T A T V T E
BINARY_7512 T T Y K L I L N G K Q L K E E A I T E L V D A A I A E K Y F K L I A N A K T V E G V W T Y K D E I K T F T V T E
BINARY_7513 T T Y K L I L N G K Q L K E E A I T E L V D A A I A E K Y F K L I A N A K T V E G V W T Y K D E T L K A T V T E
BINARY_7514 T T Y K L I L N G K Q L K E E A I T E L V D A A I A E K Y F K L I A N A K T V E G V W T Y K D E T L K F T V T E
BINARY_7515 T T Y K L I L N G K Q L K E E A I T E L V D A A I A E K Y F K L I A N A K T V E G V W T Y K D E T L T A T V T E
BINARY_7516 T T Y K L I L N G K Q L K E E A I T E L V D A A I A E K Y F K L I A N A K T V E G V W T Y K D E T L T F T V T E
BINARY_7517 T T Y K L I L N G K Q L K E E A I T E L V D A A I A E K Y F K L I A N A K T V E G V W T Y K D E T K K A T V T E
BINARY_7518 T T Y K L I L N G K Q L K E E A I T E L V D A A I A E K Y F K L I A N A K T V E G V W T Y K D E T K K F T V T E
BINARY_7519 T T Y K L I L N G K Q L K E E A I T E L V D A A I A E K Y F K L I A N A K T V E G V W T Y K D E T K T A T V T E
BINARY_7520 T T Y K L I L N G K Q L K E E A I T E L V D A A I A E K Y F K L I A N A K T V E G V W T Y K D E T K T F T V T E
BINARY_7521 T T Y K L I L N G K Q L K E E A I T E L V D A A I A E K Y F K L Y A N A K T V E G V W T L K D E I L K A T V T E

BINARY_7522 T T Y K L I L N G K Q L K E E A I T E L V D A A I A E K Y F K L Y A N A K T V E G V W T L K D E I L K F T V T E
BINARY_7523 T T Y K L I L N G K Q L K E E A I T E L V D A A I A E K Y F K L Y A N A K T V E G V W T L K D E I L T A T V T E
BINARY_7524 T T Y K L I L N G K Q L K E E A I T E L V D A A I A E K Y F K L Y A N A K T V E G V W T L K D E I L T F T V T E
BINARY_7525 T T Y K L I L N G K Q L K E E A I T E L V D A A I A E K Y F K L Y A N A K T V E G V W T L K D E I K K A T V T E
BINARY_7526 T T Y K L I L N G K Q L K E E A I T E L V D A A I A E K Y F K L Y A N A K T V E G V W T L K D E I K K F T V T E
BINARY_7527 T T Y K L I L N G K Q L K E E A I T E L V D A A I A E K Y F K L Y A N A K T V E G V W T L K D E I K T A T V T E
BINARY_7528 T T Y K L I L N G K Q L K E E A I T E L V D A A I A E K Y F K L Y A N A K T V E G V W T L K D E I K T F T V T E
BINARY_7529 T T Y K L I L N G K Q L K E E A I T E L V D A A I A E K Y F K L Y A N A K T V E G V W T L K D E T L K A T V T E
BINARY_7530 T T Y K L I L N G K Q L K E E A I T E L V D A A I A E K Y F K L Y A N A K T V E G V W T L K D E T L K F T V T E
BINARY_7531 T T Y K L I L N G K Q L K E E A I T E L V D A A I A E K Y F K L Y A N A K T V E G V W T L K D E T L T A T V T E
BINARY_7532 T T Y K L I L N G K Q L K E E A I T E L V D A A I A E K Y F K L Y A N A K T V E G V W T L K D E T L T F T V T E
BINARY_7533 T T Y K L I L N G K Q L K E E A I T E L V D A A I A E K Y F K L Y A N A K T V E G V W T L K D E T K K A T V T E
BINARY_7534 T T Y K L I L N G K Q L K E E A I T E L V D A A I A E K Y F K L Y A N A K T V E G V W T L K D E T K K F T V T E
BINARY_7535 T T Y K L I L N G K Q L K E E A I T E L V D A A I A E K Y F K L Y A N A K T V E G V W T L K D E T K T A T V T E
BINARY_7536 T T Y K L I L N G K Q L K E E A I T E L V D A A I A E K Y F K L Y A N A K T V E G V W T L K D E T K T F T V T E
BINARY_7537 T T Y K L I L N G K Q L K E E A I T E L V D A A I A E K Y F K L Y A N A K T V E G V W T Y K D E I L K A T V T E
BINARY_7538 T T Y K L I L N G K Q L K E E A I T E L V D A A I A E K Y F K L Y A N A K T V E G V W T Y K D E I L K F T V T E
BINARY_7539 T T Y K L I L N G K Q L K E E A I T E L V D A A I A E K Y F K L Y A N A K T V E G V W T Y K D E I L T A T V T E
BINARY_7540 T T Y K L I L N G K Q L K E E A I T E L V D A A I A E K Y F K L Y A N A K T V E G V W T Y K D E I L T F T V T E
BINARY_7541 T T Y K L I L N G K Q L K E E A I T E L V D A A I A E K Y F K L Y A N A K T V E G V W T Y K D E I K K A T V T E
BINARY_7542 T T Y K L I L N G K Q L K E E A I T E L V D A A I A E K Y F K L Y A N A K T V E G V W T Y K D E I K K F T V T E
BINARY_7543 T T Y K L I L N G K Q L K E E A I T E L V D A A I A E K Y F K L Y A N A K T V E G V W T Y K D E I K T A T V T E
BINARY_7544 T T Y K L I L N G K Q L K E E A I T E L V D A A I A E K Y F K L Y A N A K T V E G V W T Y K D E I K T F T V T E
BINARY_7545 T T Y K L I L N G K Q L K E E A I T E L V D A A I A E K Y F K L Y A N A K T V E G V W T Y K D E T L K A T V T E
BINARY_7546 T T Y K L I L N G K Q L K E E A I T E L V D A A I A E K Y F K L Y A N A K T V E G V W T Y K D E T L K F T V T E
BINARY_7547 T T Y K L I L N G K Q L K E E A I T E L V D A A I A E K Y F K L Y A N A K T V E G V W T Y K D E T L T A T V T E
BINARY_7548 T T Y K L I L N G K Q L K E E A I T E L V D A A I A E K Y F K L Y A N A K T V E G V W T Y K D E T L T F T V T E
BINARY_7549 T T Y K L I L N G K Q L K E E A I T E L V D A A I A E K Y F K L Y A N A K T V E G V W T Y K D E T K K A T V T E
BINARY_7550 T T Y K L I L N G K Q L K E E A I T E L V D A A I A E K Y F K L Y A N A K T V E G V W T Y K D E T K K F T V T E
BINARY_7551 T T Y K L I L N G K Q L K E E A I T E L V D A A I A E K Y F K L Y A N A K T V E G V W T Y K D E T K T A T V T E
BINARY_7552 T T Y K L I L N G K Q L K E E A I T E L V D A A I A E K Y F K L Y A N A K T V E G V W T Y K D E T K T F T V T E
BINARY_7553 T T Y K L I L N G K Q L K E E A I T E L V D A A T A E K Y I K L I A N A K T V E G V W T L K D E I L K A T V T E
BINARY_7554 T T Y K L I L N G K Q L K E E A I T E L V D A A T A E K Y I K L I A N A K T V E G V W T L K D E I L K F T V T E
BINARY_7555 T T Y K L I L N G K Q L K E E A I T E L V D A A T A E K Y I K L I A N A K T V E G V W T L K D E I L T A T V T E
BINARY_7556 T T Y K L I L N G K Q L K E E A I T E L V D A A T A E K Y I K L I A N A K T V E G V W T L K D E I L T F T V T E
BINARY_7557 T T Y K L I L N G K Q L K E E A I T E L V D A A T A E K Y I K L I A N A K T V E G V W T L K D E I K K A T V T E
BINARY_7558 T T Y K L I L N G K Q L K E E A I T E L V D A A T A E K Y I K L I A N A K T V E G V W T L K D E I K K F T V T E
BINARY_7559 T T Y K L I L N G K Q L K E E A I T E L V D A A T A E K Y I K L I A N A K T V E G V W T L K D E I K T A T V T E
BINARY_7560 T T Y K L I L N G K Q L K E E A I T E L V D A A T A E K Y I K L I A N A K T V E G V W T L K D E I K T F T V T E
BINARY_7561 T T Y K L I L N G K Q L K E E A I T E L V D A A T A E K Y I K L I A N A K T V E G V W T L K D E T L K A T V T E
BINARY_7562 T T Y K L I L N G K Q L K E E A I T E L V D A A T A E K Y I K L I A N A K T V E G V W T L K D E T L K F T V T E
BINARY_7563 T T Y K L I L N G K Q L K E E A I T E L V D A A T A E K Y I K L I A N A K T V E G V W T L K D E T L T A T V T E
BINARY_7564 T T Y K L I L N G K Q L K E E A I T E L V D A A T A E K Y I K L I A N A K T V E G V W T L K D E T L T F T V T E
BINARY_7565 T T Y K L I L N G K Q L K E E A I T E L V D A A T A E K Y I K L I A N A K T V E G V W T L K D E T K K A T V T E
BINARY_7566 T T Y K L I L N G K Q L K E E A I T E L V D A A T A E K Y I K L I A N A K T V E G V W T L K D E T K K F T V T E
BINARY_7567 T T Y K L I L N G K Q L K E E A I T E L V D A A T A E K Y I K L I A N A K T V E G V W T L K D E T K T A T V T E
BINARY_7568 T T Y K L I L N G K Q L K E E A I T E L V D A A T A E K Y I K L I A N A K T V E G V W T L K D E T K T F T V T E
BINARY_7569 T T Y K L I L N G K Q L K E E A I T E L V D A A T A E K Y I K L I A N A K T V E G V W T Y K D E I L K A T V T E
BINARY_7570 T T Y K L I L N G K Q L K E E A I T E L V D A A T A E K Y I K L I A N A K T V E G V W T Y K D E I L K F T V T E
BINARY_7571 T T Y K L I L N G K Q L K E E A I T E L V D A A T A E K Y I K L I A N A K T V E G V W T Y K D E I L T A T V T E
BINARY_7572 T T Y K L I L N G K Q L K E E A I T E L V D A A T A E K Y I K L I A N A K T V E G V W T Y K D E I L T F T V T E
BINARY_7573 T T Y K L I L N G K Q L K E E A I T E L V D A A T A E K Y I K L I A N A K T V E G V W T Y K D E I K K A T V T E
BINARY_7574 T T Y K L I L N G K Q L K E E A I T E L V D A A T A E K Y I K L I A N A K T V E G V W T Y K D E I K K F T V T E
BINARY_7575 T T Y K L I L N G K Q L K E E A I T E L V D A A T A E K Y I K L I A N A K T V E G V W T Y K D E I K T A T V T E
BINARY_7576 T T Y K L I L N G K Q L K E E A I T E L V D A A T A E K Y I K L I A N A K T V E G V W T Y K D E I K T F T V T E
BINARY_7577 T T Y K L I L N G K Q L K E E A I T E L V D A A T A E K Y I K L I A N A K T V E G V W T Y K D E T L K A T V T E
BINARY_7578 T T Y K L I L N G K Q L K E E A I T E L V D A A T A E K Y I K L I A N A K T V E G V W T Y K D E T L K F T V T E
BINARY_7579 T T Y K L I L N G K Q L K E E A I T E L V D A A T A E K Y I K L I A N A K T V E G V W T Y K D E T L T A T V T E
BINARY_7580 T T Y K L I L N G K Q L K E E A I T E L V D A A T A E K Y I K L I A N A K T V E G V W T Y K D E T L T F T V T E
BINARY_7581 T T Y K L I L N G K Q L K E E A I T E L V D A A T A E K Y I K L I A N A K T V E G V W T Y K D E T K K A T V T E
BINARY_7582 T T Y K L I L N G K Q L K E E A I T E L V D A A T A E K Y I K L I A N A K T V E G V W T Y K D E T K K F T V T E
BINARY_7583 T T Y K L I L N G K Q L K E E A I T E L V D A A T A E K Y I K L I A N A K T V E G V W T Y K D E T K T A T V T E
BINARY_7584 T T Y K L I L N G K Q L K E E A I T E L V D A A T A E K Y I K L I A N A K T V E G V W T Y K D E T K T F T V T E
BINARY_7585 T T Y K L I L N G K Q L K E E A I T E L V D A A T A E K Y I K L Y A N A K T V E G V W T L K D E I L K A T V T E
BINARY_7586 T T Y K L I L N G K Q L K E E A I T E L V D A A T A E K Y I K L Y A N A K T V E G V W T L K D E I L K F T V T E
BINARY_7587 T T Y K L I L N G K Q L K E E A I T E L V D A A T A E K Y I K L Y A N A K T V E G V W T L K D E I L T A T V T E
BINARY_7588 T T Y K L I L N G K Q L K E E A I T E L V D A A T A E K Y I K L Y A N A K T V E G V W T L K D E I L T F T V T E
BINARY_7589 T T Y K L I L N G K Q L K E E A I T E L V D A A T A E K Y I K L Y A N A K T V E G V W T L K D E I K K A T V T E
BINARY_7590 T T Y K L I L N G K Q L K E E A I T E L V D A A T A E K Y I K L Y A N A K T V E G V W T L K D E I K K F T V T E
BINARY_7591 T T Y K L I L N G K Q L K E E A I T E L V D A A T A E K Y I K L Y A N A K T V E G V W T L K D E I K T A T V T E
BINARY_7592 T T Y K L I L N G K Q L K E E A I T E L V D A A T A E K Y I K L Y A N A K T V E G V W T L K D E I K T F T V T E
BINARY_7593 T T Y K L I L N G K Q L K E E A I T E L V D A A T A E K Y I K L Y A N A K T V E G V W T L K D E T L K A T V T E
BINARY_7594 T T Y K L I L N G K Q L K E E A I T E L V D A A T A E K Y I K L Y A N A K T V E G V W T L K D E T L K F T V T E
BINARY_7595 T T Y K L I L N G K Q L K E E A I T E L V D A A T A E K Y I K L Y A N A K T V E G V W T L K D E T L T A T V T E
BINARY_7596 T T Y K L I L N G K Q L K E E A I T E L V D A A T A E K Y I K L Y A N A K T V E G V W T L K D E T L T F T V T E
BINARY_7597 T T Y K L I L N G K Q L K E E A I T E L V D A A T A E K Y I K L Y A N A K T V E G V W T L K D E T K K A T V T E
BINARY_7598 T T Y K L I L N G K Q L K E E A I T E L V D A A T A E K Y I K L Y A N A K T V E G V W T L K D E T K K F T V T E
BINARY_7599 T T Y K L I L N G K Q L K E E A I T E L V D A A T A E K Y I K L Y A N A K T V E G V W T L K D E T K T A T V T E
BINARY_7600 T T Y K L I L N G K Q L K E E A I T E L V D A A T A E K Y I K L Y A N A K T V E G V W T L K D E T K T F T V T E
BINARY_7601 T T Y K L I L N G K Q L K E E A I T E L V D A A T A E K Y I K L Y A N A K T V E G V W T Y K D E I L K A T V T E
BINARY_7602 T T Y K L I L N G K Q L K E E A I T E L V D A A T A E K Y I K L Y A N A K T V E G V W T Y K D E I L K F T V T E
BINARY_7603 T T Y K L I L N G K Q L K E E A I T E L V D A A T A E K Y I K L Y A N A K T V E G V W T Y K D E I L T A T V T E
BINARY_7604 T T Y K L I L N G K Q L K E E A I T E L V D A A T A E K Y I K L Y A N A K T V E G V W T Y K D E I L T F T V T E
BINARY_7605 T T Y K L I L N G K Q L K E E A I T E L V D A A T A E K Y I K L Y A N A K T V E G V W T Y K D E I K K A T V T E
BINARY_7606 T T Y K L I L N G K Q L K E E A I T E L V D A A T A E K Y I K L Y A N A K T V E G V W T Y K D E I K K F T V T E
BINARY_7607 T T Y K L I L N G K Q L K E E A I T E L V D A A T A E K Y I K L Y A N A K T V E G V W T Y K D E I K T A T V T E
BINARY_7608 T T Y K L I L N G K Q L K E E A I T E L V D A A T A E K Y I K L Y A N A K T V E G V W T Y K D E I K T F T V T E
BINARY_7609 T T Y K L I L N G K Q L K E E A I T E L V D A A T A E K Y I K L Y A N A K T V E G V W T Y K D E T L K A T V T E
BINARY_7610 T T Y K L I L N G K Q L K E E A I T E L V D A A T A E K Y I K L Y A N A K T V E G V W T Y K D E T L K F T V T E
BINARY_7611 T T Y K L I L N G K Q L K E E A I T E L V D A A T A E K Y I K L Y A N A K T V E G V W T Y K D E T L T A T V T E
BINARY_7612 T T Y K L I L N G K Q L K E E A I T E L V D A A T A E K Y I K L Y A N A K T V E G V W T Y K D E T L T F T V T E
BINARY_7613 T T Y K L I L N G K Q L K E E A I T E L V D A A T A E K Y I K L Y A N A K T V E G V W T Y K D E T K K A T V T E
BINARY_7614 T T Y K L I L N G K Q L K E E A I T E L V D A A T A E K Y I K L Y A N A K T V E G V W T Y K D E T K K F T V T E
BINARY_7615 T T Y K L I L N G K Q L K E E A I T E L V D A A T A E K Y I K L Y A N A K T V E G V W T Y K D E T K T A T V T E
BINARY_7616 T T Y K L I L N G K Q L K E E A I T E L V D A A T A E K Y I K L Y A N A K T V E G V W T Y K D E T K T F T V T E

BINARY_7617 T T Y K L I L N G K Q L K E E A I T E L V D A A T A E K Y F K L I A N A K T V E G V W T L K D E I L K A T V T E
BINARY_7618 T T Y K L I L N G K Q L K E E A I T E L V D A A T A E K Y F K L I A N A K T V E G V W T L K D E I L K F T V T E
BINARY_7619 T T Y K L I L N G K Q L K E E A I T E L V D A A T A E K Y F K L I A N A K T V E G V W T L K D E I L T A T V T E
BINARY_7620 T T Y K L I L N G K Q L K E E A I T E L V D A A T A E K Y F K L I A N A K T V E G V W T L K D E I L T F T V T E
BINARY_7621 T T Y K L I L N G K Q L K E E A I T E L V D A A T A E K Y F K L I A N A K T V E G V W T L K D E I K K A T V T E
BINARY_7622 T T Y K L I L N G K Q L K E E A I T E L V D A A T A E K Y F K L I A N A K T V E G V W T L K D E I K K F T V T E
BINARY_7623 T T Y K L I L N G K Q L K E E A I T E L V D A A T A E K Y F K L I A N A K T V E G V W T L K D E I K T A T V T E
BINARY_7624 T T Y K L I L N G K Q L K E E A I T E L V D A A T A E K Y F K L I A N A K T V E G V W T L K D E I K T F T V T E
BINARY_7625 T T Y K L I L N G K Q L K E E A I T E L V D A A T A E K Y F K L I A N A K T V E G V W T L K D E T L K A T V T E
BINARY_7626 T T Y K L I L N G K Q L K E E A I T E L V D A A T A E K Y F K L I A N A K T V E G V W T L K D E T L K F T V T E
BINARY_7627 T T Y K L I L N G K Q L K E E A I T E L V D A A T A E K Y F K L I A N A K T V E G V W T L K D E T L T A T V T E
BINARY_7628 T T Y K L I L N G K Q L K E E A I T E L V D A A T A E K Y F K L I A N A K T V E G V W T L K D E T L T F T V T E
BINARY_7629 T T Y K L I L N G K Q L K E E A I T E L V D A A T A E K Y F K L I A N A K T V E G V W T L K D E T K K A T V T E
BINARY_7630 T T Y K L I L N G K Q L K E E A I T E L V D A A T A E K Y F K L I A N A K T V E G V W T L K D E T K K F T V T E
BINARY_7631 T T Y K L I L N G K Q L K E E A I T E L V D A A T A E K Y F K L I A N A K T V E G V W T L K D E T K T A T V T E
BINARY_7632 T T Y K L I L N G K Q L K E E A I T E L V D A A T A E K Y F K L I A N A K T V E G V W T L K D E T K T F T V T E
BINARY_7633 T T Y K L I L N G K Q L K E E A I T E L V D A A T A E K Y F K L I A N A K T V E G V W T Y K D E I L K A T V T E
BINARY_7634 T T Y K L I L N G K Q L K E E A I T E L V D A A T A E K Y F K L I A N A K T V E G V W T Y K D E I L K F T V T E
BINARY_7635 T T Y K L I L N G K Q L K E E A I T E L V D A A T A E K Y F K L I A N A K T V E G V W T Y K D E I L T A T V T E
BINARY_7636 T T Y K L I L N G K Q L K E E A I T E L V D A A T A E K Y F K L I A N A K T V E G V W T Y K D E I L T F T V T E
BINARY_7637 T T Y K L I L N G K Q L K E E A I T E L V D A A T A E K Y F K L I A N A K T V E G V W T Y K D E I K K A T V T E
BINARY_7638 T T Y K L I L N G K Q L K E E A I T E L V D A A T A E K Y F K L I A N A K T V E G V W T Y K D E I K K F T V T E
BINARY_7639 T T Y K L I L N G K Q L K E E A I T E L V D A A T A E K Y F K L I A N A K T V E G V W T Y K D E I K T A T V T E
BINARY_7640 T T Y K L I L N G K Q L K E E A I T E L V D A A T A E K Y F K L I A N A K T V E G V W T Y K D E I K T F T V T E
BINARY_7641 T T Y K L I L N G K Q L K E E A I T E L V D A A T A E K Y F K L I A N A K T V E G V W T Y K D E T L K A T V T E
BINARY_7642 T T Y K L I L N G K Q L K E E A I T E L V D A A T A E K Y F K L I A N A K T V E G V W T Y K D E T L K F T V T E
BINARY_7643 T T Y K L I L N G K Q L K E E A I T E L V D A A T A E K Y F K L I A N A K T V E G V W T Y K D E T L T A T V T E
BINARY_7644 T T Y K L I L N G K Q L K E E A I T E L V D A A T A E K Y F K L I A N A K T V E G V W T Y K D E T L T F T V T E
BINARY_7645 T T Y K L I L N G K Q L K E E A I T E L V D A A T A E K Y F K L I A N A K T V E G V W T Y K D E T K K A T V T E
BINARY_7646 T T Y K L I L N G K Q L K E E A I T E L V D A A T A E K Y F K L I A N A K T V E G V W T Y K D E T K K F T V T E
BINARY_7647 T T Y K L I L N G K Q L K E E A I T E L V D A A T A E K Y F K L I A N A K T V E G V W T Y K D E T K T A T V T E
BINARY_7648 T T Y K L I L N G K Q L K E E A I T E L V D A A T A E K Y F K L I A N A K T V E G V W T Y K D E T K T F T V T E
BINARY_7649 T T Y K L I L N G K Q L K E E A I T E L V D A A T A E K Y F K L Y A N A K T V E G V W T L K D E I L K A T V T E
BINARY_7650 T T Y K L I L N G K Q L K E E A I T E L V D A A T A E K Y F K L Y A N A K T V E G V W T L K D E I L K F T V T E
BINARY_7651 T T Y K L I L N G K Q L K E E A I T E L V D A A T A E K Y F K L Y A N A K T V E G V W T L K D E I L T A T V T E
BINARY_7652 T T Y K L I L N G K Q L K E E A I T E L V D A A T A E K Y F K L Y A N A K T V E G V W T L K D E I L T F T V T E
BINARY_7653 T T Y K L I L N G K Q L K E E A I T E L V D A A T A E K Y F K L Y A N A K T V E G V W T L K D E I K K A T V T E
BINARY_7654 T T Y K L I L N G K Q L K E E A I T E L V D A A T A E K Y F K L Y A N A K T V E G V W T L K D E I K K F T V T E
BINARY_7655 T T Y K L I L N G K Q L K E E A I T E L V D A A T A E K Y F K L Y A N A K T V E G V W T L K D E I K T A T V T E
BINARY_7656 T T Y K L I L N G K Q L K E E A I T E L V D A A T A E K Y F K L Y A N A K T V E G V W T L K D E I K T F T V T E
BINARY_7657 T T Y K L I L N G K Q L K E E A I T E L V D A A T A E K Y F K L Y A N A K T V E G V W T L K D E T L K A T V T E
BINARY_7658 T T Y K L I L N G K Q L K E E A I T E L V D A A T A E K Y F K L Y A N A K T V E G V W T L K D E T L K F T V T E
BINARY_7659 T T Y K L I L N G K Q L K E E A I T E L V D A A T A E K Y F K L Y A N A K T V E G V W T L K D E T L T A T V T E
BINARY_7660 T T Y K L I L N G K Q L K E E A I T E L V D A A T A E K Y F K L Y A N A K T V E G V W T L K D E T L T F T V T E
BINARY_7661 T T Y K L I L N G K Q L K E E A I T E L V D A A T A E K Y F K L Y A N A K T V E G V W T L K D E T K K A T V T E
BINARY_7662 T T Y K L I L N G K Q L K E E A I T E L V D A A T A E K Y F K L Y A N A K T V E G V W T L K D E T K K F T V T E
BINARY_7663 T T Y K L I L N G K Q L K E E A I T E L V D A A T A E K Y F K L Y A N A K T V E G V W T L K D E T K T A T V T E
BINARY_7664 T T Y K L I L N G K Q L K E E A I T E L V D A A T A E K Y F K L Y A N A K T V E G V W T L K D E T K T F T V T E
BINARY_7665 T T Y K L I L N G K Q L K E E A I T E L V D A A T A E K Y F K L Y A N A K T V E G V W T Y K D E I L K A T V T E
BINARY_7666 T T Y K L I L N G K Q L K E E A I T E L V D A A T A E K Y F K L Y A N A K T V E G V W T Y K D E I L K F T V T E
BINARY_7667 T T Y K L I L N G K Q L K E E A I T E L V D A A T A E K Y F K L Y A N A K T V E G V W T Y K D E I L T A T V T E
BINARY_7668 T T Y K L I L N G K Q L K E E A I T E L V D A A T A E K Y F K L Y A N A K T V E G V W T Y K D E I L T F T V T E
BINARY_7669 T T Y K L I L N G K Q L K E E A I T E L V D A A T A E K Y F K L Y A N A K T V E G V W T Y K D E I K K A T V T E
BINARY_7670 T T Y K L I L N G K Q L K E E A I T E L V D A A T A E K Y F K L Y A N A K T V E G V W T Y K D E I K K F T V T E
BINARY_7671 T T Y K L I L N G K Q L K E E A I T E L V D A A T A E K Y F K L Y A N A K T V E G V W T Y K D E I K T A T V T E
BINARY_7672 T T Y K L I L N G K Q L K E E A I T E L V D A A T A E K Y F K L Y A N A K T V E G V W T Y K D E I K T F T V T E
BINARY_7673 T T Y K L I L N G K Q L K E E A I T E L V D A A T A E K Y F K L Y A N A K T V E G V W T Y K D E T L K A T V T E
BINARY_7674 T T Y K L I L N G K Q L K E E A I T E L V D A A T A E K Y F K L Y A N A K T V E G V W T Y K D E T L K F T V T E
BINARY_7675 T T Y K L I L N G K Q L K E E A I T E L V D A A T A E K Y F K L Y A N A K T V E G V W T Y K D E T L T A T V T E
BINARY_7676 T T Y K L I L N G K Q L K E E A I T E L V D A A T A E K Y F K L Y A N A K T V E G V W T Y K D E T L T F T V T E
BINARY_7677 T T Y K L I L N G K Q L K E E A I T E L V D A A T A E K Y F K L Y A N A K T V E G V W T Y K D E T K K A T V T E
BINARY_7678 T T Y K L I L N G K Q L K E E A I T E L V D A A T A E K Y F K L Y A N A K T V E G V W T Y K D E T K K F T V T E
BINARY_7679 T T Y K L I L N G K Q L K E E A I T E L V D A A T A E K Y F K L Y A N A K T V E G V W T Y K D E T K T A T V T E
BINARY_7680 T T Y K L I L N G K Q L K E E A I T E L V D A A T A E K Y F K L Y A N A K T V E G V W T Y K D E T K T F T V T E
BINARY_7681 T T Y K L I L N G K Q L K E E A I T E A V D A G I A E K Y I K L I A N A K T V E G V W T L K D E I L K A T V T E
BINARY_7682 T T Y K L I L N G K Q L K E E A I T E A V D A G I A E K Y I K L I A N A K T V E G V W T L K D E I L K F T V T E
BINARY_7683 T T Y K L I L N G K Q L K E E A I T E A V D A G I A E K Y I K L I A N A K T V E G V W T L K D E I L T A T V T E
BINARY_7684 T T Y K L I L N G K Q L K E E A I T E A V D A G I A E K Y I K L I A N A K T V E G V W T L K D E I L T F T V T E
BINARY_7685 T T Y K L I L N G K Q L K E E A I T E A V D A G I A E K Y I K L I A N A K T V E G V W T L K D E I K K A T V T E
BINARY_7686 T T Y K L I L N G K Q L K E E A I T E A V D A G I A E K Y I K L I A N A K T V E G V W T L K D E I K K F T V T E
BINARY_7687 T T Y K L I L N G K Q L K E E A I T E A V D A G I A E K Y I K L I A N A K T V E G V W T L K D E I K T A T V T E
BINARY_7688 T T Y K L I L N G K Q L K E E A I T E A V D A G I A E K Y I K L I A N A K T V E G V W T L K D E I K T F T V T E
BINARY_7689 T T Y K L I L N G K Q L K E E A I T E A V D A G I A E K Y I K L I A N A K T V E G V W T L K D E T L K A T V T E
BINARY_7690 T T Y K L I L N G K Q L K E E A I T E A V D A G I A E K Y I K L I A N A K T V E G V W T L K D E T L K F T V T E
BINARY_7691 T T Y K L I L N G K Q L K E E A I T E A V D A G I A E K Y I K L I A N A K T V E G V W T L K D E T L T A T V T E
BINARY_7692 T T Y K L I L N G K Q L K E E A I T E A V D A G I A E K Y I K L I A N A K T V E G V W T L K D E T L T F T V T E
BINARY_7693 T T Y K L I L N G K Q L K E E A I T E A V D A G I A E K Y I K L I A N A K T V E G V W T L K D E T K K A T V T E
BINARY_7694 T T Y K L I L N G K Q L K E E A I T E A V D A G I A E K Y I K L I A N A K T V E G V W T L K D E T K K F T V T E
BINARY_7695 T T Y K L I L N G K Q L K E E A I T E A V D A G I A E K Y I K L I A N A K T V E G V W T L K D E T K T A T V T E
BINARY_7696 T T Y K L I L N G K Q L K E E A I T E A V D A G I A E K Y I K L I A N A K T V E G V W T L K D E T K T F T V T E
BINARY_7697 T T Y K L I L N G K Q L K E E A I T E A V D A G I A E K Y I K L I A N A K T V E G V W T Y K D E I L K A T V T E
BINARY_7698 T T Y K L I L N G K Q L K E E A I T E A V D A G I A E K Y I K L I A N A K T V E G V W T Y K D E I L K F T V T E
BINARY_7699 T T Y K L I L N G K Q L K E E A I T E A V D A G I A E K Y I K L I A N A K T V E G V W T Y K D E I L T A T V T E
BINARY_7700 T T Y K L I L N G K Q L K E E A I T E A V D A G I A E K Y I K L I A N A K T V E G V W T Y K D E I L T F T V T E
BINARY_7701 T T Y K L I L N G K Q L K E E A I T E A V D A G I A E K Y I K L I A N A K T V E G V W T Y K D E I K K A T V T E
BINARY_7702 T T Y K L I L N G K Q L K E E A I T E A V D A G I A E K Y I K L I A N A K T V E G V W T Y K D E I K K F T V T E
BINARY_7703 T T Y K L I L N G K Q L K E E A I T E A V D A G I A E K Y I K L I A N A K T V E G V W T Y K D E I K T A T V T E
BINARY_7704 T T Y K L I L N G K Q L K E E A I T E A V D A G I A E K Y I K L I A N A K T V E G V W T Y K D E I K T F T V T E
BINARY_7705 T T Y K L I L N G K Q L K E E A I T E A V D A G I A E K Y I K L I A N A K T V E G V W T Y K D E T L K A T V T E
BINARY_7706 T T Y K L I L N G K Q L K E E A I T E A V D A G I A E K Y I K L I A N A K T V E G V W T Y K D E T L K F T V T E
BINARY_7707 T T Y K L I L N G K Q L K E E A I T E A V D A G I A E K Y I K L I A N A K T V E G V W T Y K D E T L T A T V T E
BINARY_7708 T T Y K L I L N G K Q L K E E A I T E A V D A G I A E K Y I K L I A N A K T V E G V W T Y K D E T L T F T V T E
BINARY_7709 T T Y K L I L N G K Q L K E E A I T E A V D A G I A E K Y I K L I A N A K T V E G V W T Y K D E T K K A T V T E
BINARY_7710 T T Y K L I L N G K Q L K E E A I T E A V D A G I A E K Y I K L I A N A K T V E G V W T Y K D E T K K F T V T E
BINARY_7711 T T Y K L I L N G K Q L K E E A I T E A V D A G I A E K Y I K L I A N A K T V E G V W T Y K D E T K T A T V T E

BINARY_7712 T T Y K L I L N G K Q L K E E A I T E A V D A G I A E K Y I K L I A N A K T V E G V W T Y K D E T K T F T V T E
BINARY_7713 T T Y K L I L N G K Q L K E E A I T E A V D A G I A E K Y I K L Y A N A K T V E G V W T L K D E I L K A T V T E
BINARY_7714 T T Y K L I L N G K Q L K E E A I T E A V D A G I A E K Y I K L Y A N A K T V E G V W T L K D E I L K F T V T E
BINARY_7715 T T Y K L I L N G K Q L K E E A I T E A V D A G I A E K Y I K L Y A N A K T V E G V W T L K D E I L T A T V T E
BINARY_7716 T T Y K L I L N G K Q L K E E A I T E A V D A G I A E K Y I K L Y A N A K T V E G V W T L K D E I L T F T V T E
BINARY_7717 T T Y K L I L N G K Q L K E E A I T E A V D A G I A E K Y I K L Y A N A K T V E G V W T L K D E I K K A T V T E
BINARY_7718 T T Y K L I L N G K Q L K E E A I T E A V D A G I A E K Y I K L Y A N A K T V E G V W T L K D E I K K F T V T E
BINARY_7719 T T Y K L I L N G K Q L K E E A I T E A V D A G I A E K Y I K L Y A N A K T V E G V W T L K D E I K T A T V T E
BINARY_7720 T T Y K L I L N G K Q L K E E A I T E A V D A G I A E K Y I K L Y A N A K T V E G V W T L K D E I K T F T V T E
BINARY_7721 T T Y K L I L N G K Q L K E E A I T E A V D A G I A E K Y I K L Y A N A K T V E G V W T L K D E T L K A T V T E
BINARY_7722 T T Y K L I L N G K Q L K E E A I T E A V D A G I A E K Y I K L Y A N A K T V E G V W T L K D E T L K F T V T E
BINARY_7723 T T Y K L I L N G K Q L K E E A I T E A V D A G I A E K Y I K L Y A N A K T V E G V W T L K D E T L T A T V T E
BINARY_7724 T T Y K L I L N G K Q L K E E A I T E A V D A G I A E K Y I K L Y A N A K T V E G V W T L K D E T L T F T V T E
BINARY_7725 T T Y K L I L N G K Q L K E E A I T E A V D A G I A E K Y I K L Y A N A K T V E G V W T L K D E T K K A T V T E
BINARY_7726 T T Y K L I L N G K Q L K E E A I T E A V D A G I A E K Y I K L Y A N A K T V E G V W T L K D E T K K F T V T E
BINARY_7727 T T Y K L I L N G K Q L K E E A I T E A V D A G I A E K Y I K L Y A N A K T V E G V W T L K D E T K T A T V T E
BINARY_7728 T T Y K L I L N G K Q L K E E A I T E A V D A G I A E K Y I K L Y A N A K T V E G V W T L K D E T K T F T V T E
BINARY_7729 T T Y K L I L N G K Q L K E E A I T E A V D A G I A E K Y I K L Y A N A K T V E G V W T Y K D E I L K A T V T E
BINARY_7730 T T Y K L I L N G K Q L K E E A I T E A V D A G I A E K Y I K L Y A N A K T V E G V W T Y K D E I L K F T V T E
BINARY_7731 T T Y K L I L N G K Q L K E E A I T E A V D A G I A E K Y I K L Y A N A K T V E G V W T Y K D E I L T A T V T E
BINARY_7732 T T Y K L I L N G K Q L K E E A I T E A V D A G I A E K Y I K L Y A N A K T V E G V W T Y K D E I L T F T V T E
BINARY_7733 T T Y K L I L N G K Q L K E E A I T E A V D A G I A E K Y I K L Y A N A K T V E G V W T Y K D E I K K A T V T E
BINARY_7734 T T Y K L I L N G K Q L K E E A I T E A V D A G I A E K Y I K L Y A N A K T V E G V W T Y K D E I K K F T V T E
BINARY_7735 T T Y K L I L N G K Q L K E E A I T E A V D A G I A E K Y I K L Y A N A K T V E G V W T Y K D E I K T A T V T E
BINARY_7736 T T Y K L I L N G K Q L K E E A I T E A V D A G I A E K Y I K L Y A N A K T V E G V W T Y K D E I K T F T V T E
BINARY_7737 T T Y K L I L N G K Q L K E E A I T E A V D A G I A E K Y I K L Y A N A K T V E G V W T Y K D E T L K A T V T E
BINARY_7738 T T Y K L I L N G K Q L K E E A I T E A V D A G I A E K Y I K L Y A N A K T V E G V W T Y K D E T L K F T V T E
BINARY_7739 T T Y K L I L N G K Q L K E E A I T E A V D A G I A E K Y I K L Y A N A K T V E G V W T Y K D E T L T A T V T E
BINARY_7740 T T Y K L I L N G K Q L K E E A I T E A V D A G I A E K Y I K L Y A N A K T V E G V W T Y K D E T L T F T V T E
BINARY_7741 T T Y K L I L N G K Q L K E E A I T E A V D A G I A E K Y I K L Y A N A K T V E G V W T Y K D E T K K A T V T E
BINARY_7742 T T Y K L I L N G K Q L K E E A I T E A V D A G I A E K Y I K L Y A N A K T V E G V W T Y K D E T K K F T V T E
BINARY_7743 T T Y K L I L N G K Q L K E E A I T E A V D A G I A E K Y I K L Y A N A K T V E G V W T Y K D E T K T A T V T E
BINARY_7744 T T Y K L I L N G K Q L K E E A I T E A V D A G I A E K Y I K L Y A N A K T V E G V W T Y K D E T K T F T V T E
BINARY_7745 T T Y K L I L N G K Q L K E E A I T E A V D A G I A E K Y F K L I A N A K T V E G V W T L K D E I L K A T V T E
BINARY_7746 T T Y K L I L N G K Q L K E E A I T E A V D A G I A E K Y F K L I A N A K T V E G V W T L K D E I L K F T V T E
BINARY_7747 T T Y K L I L N G K Q L K E E A I T E A V D A G I A E K Y F K L I A N A K T V E G V W T L K D E I L T A T V T E
BINARY_7748 T T Y K L I L N G K Q L K E E A I T E A V D A G I A E K Y F K L I A N A K T V E G V W T L K D E I L T F T V T E
BINARY_7749 T T Y K L I L N G K Q L K E E A I T E A V D A G I A E K Y F K L I A N A K T V E G V W T L K D E I K K A T V T E
BINARY_7750 T T Y K L I L N G K Q L K E E A I T E A V D A G I A E K Y F K L I A N A K T V E G V W T L K D E I K K F T V T E
BINARY_7751 T T Y K L I L N G K Q L K E E A I T E A V D A G I A E K Y F K L I A N A K T V E G V W T L K D E I K T A T V T E
BINARY_7752 T T Y K L I L N G K Q L K E E A I T E A V D A G I A E K Y F K L I A N A K T V E G V W T L K D E I K T F T V T E
BINARY_7753 T T Y K L I L N G K Q L K E E A I T E A V D A G I A E K Y F K L I A N A K T V E G V W T L K D E T L K A T V T E
BINARY_7754 T T Y K L I L N G K Q L K E E A I T E A V D A G I A E K Y F K L I A N A K T V E G V W T L K D E T L K F T V T E
BINARY_7755 T T Y K L I L N G K Q L K E E A I T E A V D A G I A E K Y F K L I A N A K T V E G V W T L K D E T L T A T V T E
BINARY_7756 T T Y K L I L N G K Q L K E E A I T E A V D A G I A E K Y F K L I A N A K T V E G V W T L K D E T L T F T V T E
BINARY_7757 T T Y K L I L N G K Q L K E E A I T E A V D A G I A E K Y F K L I A N A K T V E G V W T L K D E T K K A T V T E
BINARY_7758 T T Y K L I L N G K Q L K E E A I T E A V D A G I A E K Y F K L I A N A K T V E G V W T L K D E T K K F T V T E
BINARY_7759 T T Y K L I L N G K Q L K E E A I T E A V D A G I A E K Y F K L I A N A K T V E G V W T L K D E T K T A T V T E
BINARY_7760 T T Y K L I L N G K Q L K E E A I T E A V D A G I A E K Y F K L I A N A K T V E G V W T L K D E T K T F T V T E
BINARY_7761 T T Y K L I L N G K Q L K E E A I T E A V D A G I A E K Y F K L I A N A K T V E G V W T Y K D E I L K A T V T E
BINARY_7762 T T Y K L I L N G K Q L K E E A I T E A V D A G I A E K Y F K L I A N A K T V E G V W T Y K D E I L K F T V T E
BINARY_7763 T T Y K L I L N G K Q L K E E A I T E A V D A G I A E K Y F K L I A N A K T V E G V W T Y K D E I L T A T V T E
BINARY_7764 T T Y K L I L N G K Q L K E E A I T E A V D A G I A E K Y F K L I A N A K T V E G V W T Y K D E I L T F T V T E
BINARY_7765 T T Y K L I L N G K Q L K E E A I T E A V D A G I A E K Y F K L I A N A K T V E G V W T Y K D E I K K A T V T E
BINARY_7766 T T Y K L I L N G K Q L K E E A I T E A V D A G I A E K Y F K L I A N A K T V E G V W T Y K D E I K K F T V T E
BINARY_7767 T T Y K L I L N G K Q L K E E A I T E A V D A G I A E K Y F K L I A N A K T V E G V W T Y K D E I K T A T V T E
BINARY_7768 T T Y K L I L N G K Q L K E E A I T E A V D A G I A E K Y F K L I A N A K T V E G V W T Y K D E I K T F T V T E
BINARY_7769 T T Y K L I L N G K Q L K E E A I T E A V D A G I A E K Y F K L I A N A K T V E G V W T Y K D E T L K A T V T E
BINARY_7770 T T Y K L I L N G K Q L K E E A I T E A V D A G I A E K Y F K L I A N A K T V E G V W T Y K D E T L K F T V T E
BINARY_7771 T T Y K L I L N G K Q L K E E A I T E A V D A G I A E K Y F K L I A N A K T V E G V W T Y K D E T L T A T V T E
BINARY_7772 T T Y K L I L N G K Q L K E E A I T E A V D A G I A E K Y F K L I A N A K T V E G V W T Y K D E T L T F T V T E
BINARY_7773 T T Y K L I L N G K Q L K E E A I T E A V D A G I A E K Y F K L I A N A K T V E G V W T Y K D E T K K A T V T E
BINARY_7774 T T Y K L I L N G K Q L K E E A I T E A V D A G I A E K Y F K L I A N A K T V E G V W T Y K D E T K K F T V T E
BINARY_7775 T T Y K L I L N G K Q L K E E A I T E A V D A G I A E K Y F K L I A N A K T V E G V W T Y K D E T K T A T V T E
BINARY_7776 T T Y K L I L N G K Q L K E E A I T E A V D A G I A E K Y F K L I A N A K T V E G V W T Y K D E T K T F T V T E
BINARY_7777 T T Y K L I L N G K Q L K E E A I T E A V D A G I A E K Y F K L Y A N A K T V E G V W T L K D E I L K A T V T E
BINARY_7778 T T Y K L I L N G K Q L K E E A I T E A V D A G I A E K Y F K L Y A N A K T V E G V W T L K D E I L K F T V T E
BINARY_7779 T T Y K L I L N G K Q L K E E A I T E A V D A G I A E K Y F K L Y A N A K T V E G V W T L K D E I L T A T V T E
BINARY_7780 T T Y K L I L N G K Q L K E E A I T E A V D A G I A E K Y F K L Y A N A K T V E G V W T L K D E I L T F T V T E
BINARY_7781 T T Y K L I L N G K Q L K E E A I T E A V D A G I A E K Y F K L Y A N A K T V E G V W T L K D E I K K A T V T E
BINARY_7782 T T Y K L I L N G K Q L K E E A I T E A V D A G I A E K Y F K L Y A N A K T V E G V W T L K D E I K K F T V T E
BINARY_7783 T T Y K L I L N G K Q L K E E A I T E A V D A G I A E K Y F K L Y A N A K T V E G V W T L K D E I K T A T V T E
BINARY_7784 T T Y K L I L N G K Q L K E E A I T E A V D A G I A E K Y F K L Y A N A K T V E G V W T L K D E I K T F T V T E
BINARY_7785 T T Y K L I L N G K Q L K E E A I T E A V D A G I A E K Y F K L Y A N A K T V E G V W T L K D E T L K A T V T E
BINARY_7786 T T Y K L I L N G K Q L K E E A I T E A V D A G I A E K Y F K L Y A N A K T V E G V W T L K D E T L K F T V T E
BINARY_7787 T T Y K L I L N G K Q L K E E A I T E A V D A G I A E K Y F K L Y A N A K T V E G V W T L K D E T L T A T V T E
BINARY_7788 T T Y K L I L N G K Q L K E E A I T E A V D A G I A E K Y F K L Y A N A K T V E G V W T L K D E T L T F T V T E
BINARY_7789 T T Y K L I L N G K Q L K E E A I T E A V D A G I A E K Y F K L Y A N A K T V E G V W T L K D E T K K A T V T E
BINARY_7790 T T Y K L I L N G K Q L K E E A I T E A V D A G I A E K Y F K L Y A N A K T V E G V W T L K D E T K K F T V T E
BINARY_7791 T T Y K L I L N G K Q L K E E A I T E A V D A G I A E K Y F K L Y A N A K T V E G V W T L K D E T K T A T V T E
BINARY_7792 T T Y K L I L N G K Q L K E E A I T E A V D A G I A E K Y F K L Y A N A K T V E G V W T L K D E T K T F T V T E
BINARY_7793 T T Y K L I L N G K Q L K E E A I T E A V D A G I A E K Y F K L Y A N A K T V E G V W T Y K D E I L K A T V T E
BINARY_7794 T T Y K L I L N G K Q L K E E A I T E A V D A G I A E K Y F K L Y A N A K T V E G V W T Y K D E I L K F T V T E
BINARY_7795 T T Y K L I L N G K Q L K E E A I T E A V D A G I A E K Y F K L Y A N A K T V E G V W T Y K D E I L T A T V T E
BINARY_7796 T T Y K L I L N G K Q L K E E A I T E A V D A G I A E K Y F K L Y A N A K T V E G V W T Y K D E I L T F T V T E
BINARY_7797 T T Y K L I L N G K Q L K E E A I T E A V D A G I A E K Y F K L Y A N A K T V E G V W T Y K D E I K K A T V T E
BINARY_7798 T T Y K L I L N G K Q L K E E A I T E A V D A G I A E K Y F K L Y A N A K T V E G V W T Y K D E I K K F T V T E
BINARY_7799 T T Y K L I L N G K Q L K E E A I T E A V D A G I A E K Y F K L Y A N A K T V E G V W T Y K D E I K T A T V T E
BINARY_7800 T T Y K L I L N G K Q L K E E A I T E A V D A G I A E K Y F K L Y A N A K T V E G V W T Y K D E I K T F T V T E
BINARY_7801 T T Y K L I L N G K Q L K E E A I T E A V D A G I A E K Y F K L Y A N A K T V E G V W T Y K D E T L K A T V T E
BINARY_7802 T T Y K L I L N G K Q L K E E A I T E A V D A G I A E K Y F K L Y A N A K T V E G V W T Y K D E T L K F T V T E
BINARY_7803 T T Y K L I L N G K Q L K E E A I T E A V D A G I A E K Y F K L Y A N A K T V E G V W T Y K D E T L T A T V T E
BINARY_7804 T T Y K L I L N G K Q L K E E A I T E A V D A G I A E K Y F K L Y A N A K T V E G V W T Y K D E T L T F T V T E
BINARY_7805 T T Y K L I L N G K Q L K E E A I T E A V D A G I A E K Y F K L Y A N A K T V E G V W T Y K D E T K K A T V T E
BINARY_7806 T T Y K L I L N G K Q L K E E A I T E A V D A G I A E K Y F K L Y A N A K T V E G V W T Y K D E T K K F T V T E

BINARY_7807 T T Y K L I L N G K Q L K E E A I T E A V D A G I A E K Y F K L Y A N A K T V E G V W T Y K D E T K T A T V T E
BINARY_7808 T T Y K L I L N G K Q L K E E A I T E A V D A G I A E K Y F K L Y A N A K T V E G V W T Y K D E T K T F T V T E
BINARY_7809 T T Y K L I L N G K Q L K E E A I T E A V D A G T A E K Y I K L I A N A K T V E G V W T L K D E I L K A T V T E
BINARY_7810 T T Y K L I L N G K Q L K E E A I T E A V D A G T A E K Y I K L I A N A K T V E G V W T L K D E I L K F T V T E
BINARY_7811 T T Y K L I L N G K Q L K E E A I T E A V D A G T A E K Y I K L I A N A K T V E G V W T L K D E I L T A T V T E
BINARY_7812 T T Y K L I L N G K Q L K E E A I T E A V D A G T A E K Y I K L I A N A K T V E G V W T L K D E I L T F T V T E
BINARY_7813 T T Y K L I L N G K Q L K E E A I T E A V D A G T A E K Y I K L I A N A K T V E G V W T L K D E I K K A T V T E
BINARY_7814 T T Y K L I L N G K Q L K E E A I T E A V D A G T A E K Y I K L I A N A K T V E G V W T L K D E I K K F T V T E
BINARY_7815 T T Y K L I L N G K Q L K E E A I T E A V D A G T A E K Y I K L I A N A K T V E G V W T L K D E I K T A T V T E
BINARY_7816 T T Y K L I L N G K Q L K E E A I T E A V D A G T A E K Y I K L I A N A K T V E G V W T L K D E I K T F T V T E
BINARY_7817 T T Y K L I L N G K Q L K E E A I T E A V D A G T A E K Y I K L I A N A K T V E G V W T L K D E T L K A T V T E
BINARY_7818 T T Y K L I L N G K Q L K E E A I T E A V D A G T A E K Y I K L I A N A K T V E G V W T L K D E T L K F T V T E
BINARY_7819 T T Y K L I L N G K Q L K E E A I T E A V D A G T A E K Y I K L I A N A K T V E G V W T L K D E T L T A T V T E
BINARY_7820 T T Y K L I L N G K Q L K E E A I T E A V D A G T A E K Y I K L I A N A K T V E G V W T L K D E T L T F T V T E
BINARY_7821 T T Y K L I L N G K Q L K E E A I T E A V D A G T A E K Y I K L I A N A K T V E G V W T L K D E T K K A T V T E
BINARY_7822 T T Y K L I L N G K Q L K E E A I T E A V D A G T A E K Y I K L I A N A K T V E G V W T L K D E T K K F T V T E
BINARY_7823 T T Y K L I L N G K Q L K E E A I T E A V D A G T A E K Y I K L I A N A K T V E G V W T L K D E T K T A T V T E
BINARY_7824 T T Y K L I L N G K Q L K E E A I T E A V D A G T A E K Y I K L I A N A K T V E G V W T L K D E T K T F T V T E
BINARY_7825 T T Y K L I L N G K Q L K E E A I T E A V D A G T A E K Y I K L I A N A K T V E G V W T Y K D E I L K A T V T E
BINARY_7826 T T Y K L I L N G K Q L K E E A I T E A V D A G T A E K Y I K L I A N A K T V E G V W T Y K D E I L K F T V T E
BINARY_7827 T T Y K L I L N G K Q L K E E A I T E A V D A G T A E K Y I K L I A N A K T V E G V W T Y K D E I L T A T V T E
BINARY_7828 T T Y K L I L N G K Q L K E E A I T E A V D A G T A E K Y I K L I A N A K T V E G V W T Y K D E I L T F T V T E
BINARY_7829 T T Y K L I L N G K Q L K E E A I T E A V D A G T A E K Y I K L I A N A K T V E G V W T Y K D E I K K A T V T E
BINARY_7830 T T Y K L I L N G K Q L K E E A I T E A V D A G T A E K Y I K L I A N A K T V E G V W T Y K D E I K K F T V T E
BINARY_7831 T T Y K L I L N G K Q L K E E A I T E A V D A G T A E K Y I K L I A N A K T V E G V W T Y K D E I K T A T V T E
BINARY_7832 T T Y K L I L N G K Q L K E E A I T E A V D A G T A E K Y I K L I A N A K T V E G V W T Y K D E I K T F T V T E
BINARY_7833 T T Y K L I L N G K Q L K E E A I T E A V D A G T A E K Y I K L I A N A K T V E G V W T Y K D E T L K A T V T E
BINARY_7834 T T Y K L I L N G K Q L K E E A I T E A V D A G T A E K Y I K L I A N A K T V E G V W T Y K D E T L K F T V T E
BINARY_7835 T T Y K L I L N G K Q L K E E A I T E A V D A G T A E K Y I K L I A N A K T V E G V W T Y K D E T L T A T V T E
BINARY_7836 T T Y K L I L N G K Q L K E E A I T E A V D A G T A E K Y I K L I A N A K T V E G V W T Y K D E T L T F T V T E
BINARY_7837 T T Y K L I L N G K Q L K E E A I T E A V D A G T A E K Y I K L I A N A K T V E G V W T Y K D E T K K A T V T E
BINARY_7838 T T Y K L I L N G K Q L K E E A I T E A V D A G T A E K Y I K L I A N A K T V E G V W T Y K D E T K K F T V T E
BINARY_7839 T T Y K L I L N G K Q L K E E A I T E A V D A G T A E K Y I K L I A N A K T V E G V W T Y K D E T K T A T V T E
BINARY_7840 T T Y K L I L N G K Q L K E E A I T E A V D A G T A E K Y I K L I A N A K T V E G V W T Y K D E T K T F T V T E
BINARY_7841 T T Y K L I L N G K Q L K E E A I T E A V D A G T A E K Y I K L Y A N A K T V E G V W T L K D E I L K A T V T E
BINARY_7842 T T Y K L I L N G K Q L K E E A I T E A V D A G T A E K Y I K L Y A N A K T V E G V W T L K D E I L K F T V T E
BINARY_7843 T T Y K L I L N G K Q L K E E A I T E A V D A G T A E K Y I K L Y A N A K T V E G V W T L K D E I L T A T V T E
BINARY_7844 T T Y K L I L N G K Q L K E E A I T E A V D A G T A E K Y I K L Y A N A K T V E G V W T L K D E I L T F T V T E
BINARY_7845 T T Y K L I L N G K Q L K E E A I T E A V D A G T A E K Y I K L Y A N A K T V E G V W T L K D E I K K A T V T E
BINARY_7846 T T Y K L I L N G K Q L K E E A I T E A V D A G T A E K Y I K L Y A N A K T V E G V W T L K D E I K K F T V T E
BINARY_7847 T T Y K L I L N G K Q L K E E A I T E A V D A G T A E K Y I K L Y A N A K T V E G V W T L K D E I K T A T V T E
BINARY_7848 T T Y K L I L N G K Q L K E E A I T E A V D A G T A E K Y I K L Y A N A K T V E G V W T L K D E I K T F T V T E
BINARY_7849 T T Y K L I L N G K Q L K E E A I T E A V D A G T A E K Y I K L Y A N A K T V E G V W T L K D E T L K A T V T E
BINARY_7850 T T Y K L I L N G K Q L K E E A I T E A V D A G T A E K Y I K L Y A N A K T V E G V W T L K D E T L K F T V T E
BINARY_7851 T T Y K L I L N G K Q L K E E A I T E A V D A G T A E K Y I K L Y A N A K T V E G V W T L K D E T L T A T V T E
BINARY_7852 T T Y K L I L N G K Q L K E E A I T E A V D A G T A E K Y I K L Y A N A K T V E G V W T L K D E T L T F T V T E
BINARY_7853 T T Y K L I L N G K Q L K E E A I T E A V D A G T A E K Y I K L Y A N A K T V E G V W T L K D E T K K A T V T E
BINARY_7854 T T Y K L I L N G K Q L K E E A I T E A V D A G T A E K Y I K L Y A N A K T V E G V W T L K D E T K K F T V T E
BINARY_7855 T T Y K L I L N G K Q L K E E A I T E A V D A G T A E K Y I K L Y A N A K T V E G V W T L K D E T K T A T V T E
BINARY_7856 T T Y K L I L N G K Q L K E E A I T E A V D A G T A E K Y I K L Y A N A K T V E G V W T L K D E T K T F T V T E
BINARY_7857 T T Y K L I L N G K Q L K E E A I T E A V D A G T A E K Y I K L Y A N A K T V E G V W T Y K D E I L K A T V T E
BINARY_7858 T T Y K L I L N G K Q L K E E A I T E A V D A G T A E K Y I K L Y A N A K T V E G V W T Y K D E I L K F T V T E
BINARY_7859 T T Y K L I L N G K Q L K E E A I T E A V D A G T A E K Y I K L Y A N A K T V E G V W T Y K D E I L T A T V T E
BINARY_7860 T T Y K L I L N G K Q L K E E A I T E A V D A G T A E K Y I K L Y A N A K T V E G V W T Y K D E I L T F T V T E
BINARY_7861 T T Y K L I L N G K Q L K E E A I T E A V D A G T A E K Y I K L Y A N A K T V E G V W T Y K D E I K K A T V T E
BINARY_7862 T T Y K L I L N G K Q L K E E A I T E A V D A G T A E K Y I K L Y A N A K T V E G V W T Y K D E I K K F T V T E
BINARY_7863 T T Y K L I L N G K Q L K E E A I T E A V D A G T A E K Y I K L Y A N A K T V E G V W T Y K D E I K T A T V T E
BINARY_7864 T T Y K L I L N G K Q L K E E A I T E A V D A G T A E K Y I K L Y A N A K T V E G V W T Y K D E I K T F T V T E
BINARY_7865 T T Y K L I L N G K Q L K E E A I T E A V D A G T A E K Y I K L Y A N A K T V E G V W T Y K D E T L K A T V T E
BINARY_7866 T T Y K L I L N G K Q L K E E A I T E A V D A G T A E K Y I K L Y A N A K T V E G V W T Y K D E T L K F T V T E
BINARY_7867 T T Y K L I L N G K Q L K E E A I T E A V D A G T A E K Y I K L Y A N A K T V E G V W T Y K D E T L T A T V T E
BINARY_7868 T T Y K L I L N G K Q L K E E A I T E A V D A G T A E K Y I K L Y A N A K T V E G V W T Y K D E T L T F T V T E
BINARY_7869 T T Y K L I L N G K Q L K E E A I T E A V D A G T A E K Y I K L Y A N A K T V E G V W T Y K D E T K K A T V T E
BINARY_7870 T T Y K L I L N G K Q L K E E A I T E A V D A G T A E K Y I K L Y A N A K T V E G V W T Y K D E T K K F T V T E
BINARY_7871 T T Y K L I L N G K Q L K E E A I T E A V D A G T A E K Y I K L Y A N A K T V E G V W T Y K D E T K T A T V T E
BINARY_7872 T T Y K L I L N G K Q L K E E A I T E A V D A G T A E K Y I K L Y A N A K T V E G V W T Y K D E T K T F T V T E
BINARY_7873 T T Y K L I L N G K Q L K E E A I T E A V D A G T A E K Y F K L I A N A K T V E G V W T L K D E I L K A T V T E
BINARY_7874 T T Y K L I L N G K Q L K E E A I T E A V D A G T A E K Y F K L I A N A K T V E G V W T L K D E I L K F T V T E
BINARY_7875 T T Y K L I L N G K Q L K E E A I T E A V D A G T A E K Y F K L I A N A K T V E G V W T L K D E I L T A T V T E
BINARY_7876 T T Y K L I L N G K Q L K E E A I T E A V D A G T A E K Y F K L I A N A K T V E G V W T L K D E I L T F T V T E
BINARY_7877 T T Y K L I L N G K Q L K E E A I T E A V D A G T A E K Y F K L I A N A K T V E G V W T L K D E I K K A T V T E
BINARY_7878 T T Y K L I L N G K Q L K E E A I T E A V D A G T A E K Y F K L I A N A K T V E G V W T L K D E I K K F T V T E
BINARY_7879 T T Y K L I L N G K Q L K E E A I T E A V D A G T A E K Y F K L I A N A K T V E G V W T L K D E I K T A T V T E
BINARY_7880 T T Y K L I L N G K Q L K E E A I T E A V D A G T A E K Y F K L I A N A K T V E G V W T L K D E I K T F T V T E
BINARY_7881 T T Y K L I L N G K Q L K E E A I T E A V D A G T A E K Y F K L I A N A K T V E G V W T L K D E T L K A T V T E
BINARY_7882 T T Y K L I L N G K Q L K E E A I T E A V D A G T A E K Y F K L I A N A K T V E G V W T L K D E T L K F T V T E
BINARY_7883 T T Y K L I L N G K Q L K E E A I T E A V D A G T A E K Y F K L I A N A K T V E G V W T L K D E T L T A T V T E
BINARY_7884 T T Y K L I L N G K Q L K E E A I T E A V D A G T A E K Y F K L I A N A K T V E G V W T L K D E T L T F T V T E
BINARY_7885 T T Y K L I L N G K Q L K E E A I T E A V D A G T A E K Y F K L I A N A K T V E G V W T L K D E T K K A T V T E
BINARY_7886 T T Y K L I L N G K Q L K E E A I T E A V D A G T A E K Y F K L I A N A K T V E G V W T L K D E T K K F T V T E
BINARY_7887 T T Y K L I L N G K Q L K E E A I T E A V D A G T A E K Y F K L I A N A K T V E G V W T L K D E T K T A T V T E
BINARY_7888 T T Y K L I L N G K Q L K E E A I T E A V D A G T A E K Y F K L I A N A K T V E G V W T L K D E T K T F T V T E
BINARY_7889 T T Y K L I L N G K Q L K E E A I T E A V D A G T A E K Y F K L I A N A K T V E G V W T Y K D E I L K A T V T E
BINARY_7890 T T Y K L I L N G K Q L K E E A I T E A V D A G T A E K Y F K L I A N A K T V E G V W T Y K D E I L K F T V T E
BINARY_7891 T T Y K L I L N G K Q L K E E A I T E A V D A G T A E K Y F K L I A N A K T V E G V W T Y K D E I L T A T V T E
BINARY_7892 T T Y K L I L N G K Q L K E E A I T E A V D A G T A E K Y F K L I A N A K T V E G V W T Y K D E I L T F T V T E
BINARY_7893 T T Y K L I L N G K Q L K E E A I T E A V D A G T A E K Y F K L I A N A K T V E G V W T Y K D E I K K A T V T E
BINARY_7894 T T Y K L I L N G K Q L K E E A I T E A V D A G T A E K Y F K L I A N A K T V E G V W T Y K D E I K K F T V T E
BINARY_7895 T T Y K L I L N G K Q L K E E A I T E A V D A G T A E K Y F K L I A N A K T V E G V W T Y K D E I K T A T V T E
BINARY_7896 T T Y K L I L N G K Q L K E E A I T E A V D A G T A E K Y F K L I A N A K T V E G V W T Y K D E I K T F T V T E
BINARY_7897 T T Y K L I L N G K Q L K E E A I T E A V D A G T A E K Y F K L I A N A K T V E G V W T Y K D E T L K A T V T E
BINARY_7898 T T Y K L I L N G K Q L K E E A I T E A V D A G T A E K Y F K L I A N A K T V E G V W T Y K D E T L K F T V T E
BINARY_7899 T T Y K L I L N G K Q L K E E A I T E A V D A G T A E K Y F K L I A N A K T V E G V W T Y K D E T L T A T V T E
BINARY_7900 T T Y K L I L N G K Q L K E E A I T E A V D A G T A E K Y F K L I A N A K T V E G V W T Y K D E T L T F T V T E
BINARY_7901 T T Y K L I L N G K Q L K E E A I T E A V D A G T A E K Y F K L I A N A K T V E G V W T Y K D E T K K A T V T E

BINARY_7902 T T Y K L I L N G K Q L K E E A I T E A V D A G T A E K Y F K L I A N A K T V E G V W T Y K D E T K K F T V T E
BINARY_7903 T T Y K L I L N G K Q L K E E A I T E A V D A G T A E K Y F K L I A N A K T V E G V W T Y K D E T K T A T V T E
BINARY_7904 T T Y K L I L N G K Q L K E E A I T E A V D A G T A E K Y F K L I A N A K T V E G V W T Y K D E T K T F T V T E
BINARY_7905 T T Y K L I L N G K Q L K E E A I T E A V D A G T A E K Y F K L Y A N A K T V E G V W T L K D E I L K A T V T E
BINARY_7906 T T Y K L I L N G K Q L K E E A I T E A V D A G T A E K Y F K L Y A N A K T V E G V W T L K D E I L K F T V T E
BINARY_7907 T T Y K L I L N G K Q L K E E A I T E A V D A G T A E K Y F K L Y A N A K T V E G V W T L K D E I L T A T V T E
BINARY_7908 T T Y K L I L N G K Q L K E E A I T E A V D A G T A E K Y F K L Y A N A K T V E G V W T L K D E I L T F T V T E
BINARY_7909 T T Y K L I L N G K Q L K E E A I T E A V D A G T A E K Y F K L Y A N A K T V E G V W T L K D E I K K A T V T E
BINARY_7910 T T Y K L I L N G K Q L K E E A I T E A V D A G T A E K Y F K L Y A N A K T V E G V W T L K D E I K K F T V T E
BINARY_7911 T T Y K L I L N G K Q L K E E A I T E A V D A G T A E K Y F K L Y A N A K T V E G V W T L K D E I K T A T V T E
BINARY_7912 T T Y K L I L N G K Q L K E E A I T E A V D A G T A E K Y F K L Y A N A K T V E G V W T L K D E I K T F T V T E
BINARY_7913 T T Y K L I L N G K Q L K E E A I T E A V D A G T A E K Y F K L Y A N A K T V E G V W T L K D E T L K A T V T E
BINARY_7914 T T Y K L I L N G K Q L K E E A I T E A V D A G T A E K Y F K L Y A N A K T V E G V W T L K D E T L K F T V T E
BINARY_7915 T T Y K L I L N G K Q L K E E A I T E A V D A G T A E K Y F K L Y A N A K T V E G V W T L K D E T L T A T V T E
BINARY_7916 T T Y K L I L N G K Q L K E E A I T E A V D A G T A E K Y F K L Y A N A K T V E G V W T L K D E T L T F T V T E
BINARY_7917 T T Y K L I L N G K Q L K E E A I T E A V D A G T A E K Y F K L Y A N A K T V E G V W T L K D E T K K A T V T E
BINARY_7918 T T Y K L I L N G K Q L K E E A I T E A V D A G T A E K Y F K L Y A N A K T V E G V W T L K D E T K K F T V T E
BINARY_7919 T T Y K L I L N G K Q L K E E A I T E A V D A G T A E K Y F K L Y A N A K T V E G V W T L K D E T K T A T V T E
BINARY_7920 T T Y K L I L N G K Q L K E E A I T E A V D A G T A E K Y F K L Y A N A K T V E G V W T L K D E T K T F T V T E
BINARY_7921 T T Y K L I L N G K Q L K E E A I T E A V D A G T A E K Y F K L Y A N A K T V E G V W T Y K D E I L K A T V T E
BINARY_7922 T T Y K L I L N G K Q L K E E A I T E A V D A G T A E K Y F K L Y A N A K T V E G V W T Y K D E I L K F T V T E
BINARY_7923 T T Y K L I L N G K Q L K E E A I T E A V D A G T A E K Y F K L Y A N A K T V E G V W T Y K D E I L T A T V T E
BINARY_7924 T T Y K L I L N G K Q L K E E A I T E A V D A G T A E K Y F K L Y A N A K T V E G V W T Y K D E I L T F T V T E
BINARY_7925 T T Y K L I L N G K Q L K E E A I T E A V D A G T A E K Y F K L Y A N A K T V E G V W T Y K D E I K K A T V T E
BINARY_7926 T T Y K L I L N G K Q L K E E A I T E A V D A G T A E K Y F K L Y A N A K T V E G V W T Y K D E I K K F T V T E
BINARY_7927 T T Y K L I L N G K Q L K E E A I T E A V D A G T A E K Y F K L Y A N A K T V E G V W T Y K D E I K T A T V T E
BINARY_7928 T T Y K L I L N G K Q L K E E A I T E A V D A G T A E K Y F K L Y A N A K T V E G V W T Y K D E I K T F T V T E
BINARY_7929 T T Y K L I L N G K Q L K E E A I T E A V D A G T A E K Y F K L Y A N A K T V E G V W T Y K D E T L K A T V T E
BINARY_7930 T T Y K L I L N G K Q L K E E A I T E A V D A G T A E K Y F K L Y A N A K T V E G V W T Y K D E T L K F T V T E
BINARY_7931 T T Y K L I L N G K Q L K E E A I T E A V D A G T A E K Y F K L Y A N A K T V E G V W T Y K D E T L T A T V T E
BINARY_7932 T T Y K L I L N G K Q L K E E A I T E A V D A G T A E K Y F K L Y A N A K T V E G V W T Y K D E T L T F T V T E
BINARY_7933 T T Y K L I L N G K Q L K E E A I T E A V D A G T A E K Y F K L Y A N A K T V E G V W T Y K D E T K K A T V T E
BINARY_7934 T T Y K L I L N G K Q L K E E A I T E A V D A G T A E K Y F K L Y A N A K T V E G V W T Y K D E T K K F T V T E
BINARY_7935 T T Y K L I L N G K Q L K E E A I T E A V D A G T A E K Y F K L Y A N A K T V E G V W T Y K D E T K T A T V T E
BINARY_7936 T T Y K L I L N G K Q L K E E A I T E A V D A G T A E K Y F K L Y A N A K T V E G V W T Y K D E T K T F T V T E
BINARY_7937 T T Y K L I L N G K Q L K E E A I T E A V D A A I A E K Y I K L I A N A K T V E G V W T L K D E I L K A T V T E
BINARY_7938 T T Y K L I L N G K Q L K E E A I T E A V D A A I A E K Y I K L I A N A K T V E G V W T L K D E I L K F T V T E
BINARY_7939 T T Y K L I L N G K Q L K E E A I T E A V D A A I A E K Y I K L I A N A K T V E G V W T L K D E I L T A T V T E
BINARY_7940 T T Y K L I L N G K Q L K E E A I T E A V D A A I A E K Y I K L I A N A K T V E G V W T L K D E I L T F T V T E
BINARY_7941 T T Y K L I L N G K Q L K E E A I T E A V D A A I A E K Y I K L I A N A K T V E G V W T L K D E I K K A T V T E
BINARY_7942 T T Y K L I L N G K Q L K E E A I T E A V D A A I A E K Y I K L I A N A K T V E G V W T L K D E I K K F T V T E
BINARY_7943 T T Y K L I L N G K Q L K E E A I T E A V D A A I A E K Y I K L I A N A K T V E G V W T L K D E I K T A T V T E
BINARY_7944 T T Y K L I L N G K Q L K E E A I T E A V D A A I A E K Y I K L I A N A K T V E G V W T L K D E I K T F T V T E
BINARY_7945 T T Y K L I L N G K Q L K E E A I T E A V D A A I A E K Y I K L I A N A K T V E G V W T L K D E T L K A T V T E
BINARY_7946 T T Y K L I L N G K Q L K E E A I T E A V D A A I A E K Y I K L I A N A K T V E G V W T L K D E T L K F T V T E
BINARY_7947 T T Y K L I L N G K Q L K E E A I T E A V D A A I A E K Y I K L I A N A K T V E G V W T L K D E T L T A T V T E
BINARY_7948 T T Y K L I L N G K Q L K E E A I T E A V D A A I A E K Y I K L I A N A K T V E G V W T L K D E T L T F T V T E
BINARY_7949 T T Y K L I L N G K Q L K E E A I T E A V D A A I A E K Y I K L I A N A K T V E G V W T L K D E T K K A T V T E
BINARY_7950 T T Y K L I L N G K Q L K E E A I T E A V D A A I A E K Y I K L I A N A K T V E G V W T L K D E T K K F T V T E
BINARY_7951 T T Y K L I L N G K Q L K E E A I T E A V D A A I A E K Y I K L I A N A K T V E G V W T L K D E T K T A T V T E
BINARY_7952 T T Y K L I L N G K Q L K E E A I T E A V D A A I A E K Y I K L I A N A K T V E G V W T L K D E T K T F T V T E
BINARY_7953 T T Y K L I L N G K Q L K E E A I T E A V D A A I A E K Y I K L I A N A K T V E G V W T Y K D E I L K A T V T E
BINARY_7954 T T Y K L I L N G K Q L K E E A I T E A V D A A I A E K Y I K L I A N A K T V E G V W T Y K D E I L K F T V T E
BINARY_7955 T T Y K L I L N G K Q L K E E A I T E A V D A A I A E K Y I K L I A N A K T V E G V W T Y K D E I L T A T V T E
BINARY_7956 T T Y K L I L N G K Q L K E E A I T E A V D A A I A E K Y I K L I A N A K T V E G V W T Y K D E I L T F T V T E
BINARY_7957 T T Y K L I L N G K Q L K E E A I T E A V D A A I A E K Y I K L I A N A K T V E G V W T Y K D E I K K A T V T E
BINARY_7958 T T Y K L I L N G K Q L K E E A I T E A V D A A I A E K Y I K L I A N A K T V E G V W T Y K D E I K K F T V T E
BINARY_7959 T T Y K L I L N G K Q L K E E A I T E A V D A A I A E K Y I K L I A N A K T V E G V W T Y K D E I K T A T V T E
BINARY_7960 T T Y K L I L N G K Q L K E E A I T E A V D A A I A E K Y I K L I A N A K T V E G V W T Y K D E I K T F T V T E
BINARY_7961 T T Y K L I L N G K Q L K E E A I T E A V D A A I A E K Y I K L I A N A K T V E G V W T Y K D E T L K A T V T E
BINARY_7962 T T Y K L I L N G K Q L K E E A I T E A V D A A I A E K Y I K L I A N A K T V E G V W T Y K D E T L K F T V T E
BINARY_7963 T T Y K L I L N G K Q L K E E A I T E A V D A A I A E K Y I K L I A N A K T V E G V W T Y K D E T L T A T V T E
BINARY_7964 T T Y K L I L N G K Q L K E E A I T E A V D A A I A E K Y I K L I A N A K T V E G V W T Y K D E T L T F T V T E
BINARY_7965 T T Y K L I L N G K Q L K E E A I T E A V D A A I A E K Y I K L I A N A K T V E G V W T Y K D E T K K A T V T E
BINARY_7966 T T Y K L I L N G K Q L K E E A I T E A V D A A I A E K Y I K L I A N A K T V E G V W T Y K D E T K K F T V T E
BINARY_7967 T T Y K L I L N G K Q L K E E A I T E A V D A A I A E K Y I K L I A N A K T V E G V W T Y K D E T K T A T V T E
BINARY_7968 T T Y K L I L N G K Q L K E E A I T E A V D A A I A E K Y I K L I A N A K T V E G V W T Y K D E T K T F T V T E
BINARY_7969 T T Y K L I L N G K Q L K E E A I T E A V D A A I A E K Y I K L Y A N A K T V E G V W T L K D E I L K A T V T E
BINARY_7970 T T Y K L I L N G K Q L K E E A I T E A V D A A I A E K Y I K L Y A N A K T V E G V W T L K D E I L K F T V T E
BINARY_7971 T T Y K L I L N G K Q L K E E A I T E A V D A A I A E K Y I K L Y A N A K T V E G V W T L K D E I L T A T V T E
BINARY_7972 T T Y K L I L N G K Q L K E E A I T E A V D A A I A E K Y I K L Y A N A K T V E G V W T L K D E I L T F T V T E
BINARY_7973 T T Y K L I L N G K Q L K E E A I T E A V D A A I A E K Y I K L Y A N A K T V E G V W T L K D E I K K A T V T E
BINARY_7974 T T Y K L I L N G K Q L K E E A I T E A V D A A I A E K Y I K L Y A N A K T V E G V W T L K D E I K K F T V T E
BINARY_7975 T T Y K L I L N G K Q L K E E A I T E A V D A A I A E K Y I K L Y A N A K T V E G V W T L K D E I K T A T V T E
BINARY_7976 T T Y K L I L N G K Q L K E E A I T E A V D A A I A E K Y I K L Y A N A K T V E G V W T L K D E I K T F T V T E
BINARY_7977 T T Y K L I L N G K Q L K E E A I T E A V D A A I A E K Y I K L Y A N A K T V E G V W T L K D E T L K A T V T E
BINARY_7978 T T Y K L I L N G K Q L K E E A I T E A V D A A I A E K Y I K L Y A N A K T V E G V W T L K D E T L K F T V T E
BINARY_7979 T T Y K L I L N G K Q L K E E A I T E A V D A A I A E K Y I K L Y A N A K T V E G V W T L K D E T L T A T V T E
BINARY_7980 T T Y K L I L N G K Q L K E E A I T E A V D A A I A E K Y I K L Y A N A K T V E G V W T L K D E T L T F T V T E
BINARY_7981 T T Y K L I L N G K Q L K E E A I T E A V D A A I A E K Y I K L Y A N A K T V E G V W T L K D E T K K A T V T E
BINARY_7982 T T Y K L I L N G K Q L K E E A I T E A V D A A I A E K Y I K L Y A N A K T V E G V W T L K D E T K K F T V T E
BINARY_7983 T T Y K L I L N G K Q L K E E A I T E A V D A A I A E K Y I K L Y A N A K T V E G V W T L K D E T K T A T V T E
BINARY_7984 T T Y K L I L N G K Q L K E E A I T E A V D A A I A E K Y I K L Y A N A K T V E G V W T L K D E T K T F T V T E
BINARY_7985 T T Y K L I L N G K Q L K E E A I T E A V D A A I A E K Y I K L Y A N A K T V E G V W T Y K D E I L K A T V T E
BINARY_7986 T T Y K L I L N G K Q L K E E A I T E A V D A A I A E K Y I K L Y A N A K T V E G V W T Y K D E I L K F T V T E
BINARY_7987 T T Y K L I L N G K Q L K E E A I T E A V D A A I A E K Y I K L Y A N A K T V E G V W T Y K D E I L T A T V T E
BINARY_7988 T T Y K L I L N G K Q L K E E A I T E A V D A A I A E K Y I K L Y A N A K T V E G V W T Y K D E I L T F T V T E
BINARY_7989 T T Y K L I L N G K Q L K E E A I T E A V D A A I A E K Y I K L Y A N A K T V E G V W T Y K D E I K K A T V T E
BINARY_7990 T T Y K L I L N G K Q L K E E A I T E A V D A A I A E K Y I K L Y A N A K T V E G V W T Y K D E I K K F T V T E
BINARY_7991 T T Y K L I L N G K Q L K E E A I T E A V D A A I A E K Y I K L Y A N A K T V E G V W T Y K D E I K T A T V T E
BINARY_7992 T T Y K L I L N G K Q L K E E A I T E A V D A A I A E K Y I K L Y A N A K T V E G V W T Y K D E I K T F T V T E
BINARY_7993 T T Y K L I L N G K Q L K E E A I T E A V D A A I A E K Y I K L Y A N A K T V E G V W T Y K D E T L K A T V T E
BINARY_7994 T T Y K L I L N G K Q L K E E A I T E A V D A A I A E K Y I K L Y A N A K T V E G V W T Y K D E T L K F T V T E
BINARY_7995 T T Y K L I L N G K Q L K E E A I T E A V D A A I A E K Y I K L Y A N A K T V E G V W T Y K D E T L T A T V T E
BINARY_7996 T T Y K L I L N G K Q L K E E A I T E A V D A A I A E K Y I K L Y A N A K T V E G V W T Y K D E T L T F T V T E

BINARY_7997 T T Y K L I L N G K Q L K E E A I T E A V D A A I A E K Y I K L Y A N A K T V E G V W T Y K D E T K K A T V T E
BINARY_7998 T T Y K L I L N G K Q L K E E A I T E A V D A A I A E K Y I K L Y A N A K T V E G V W T Y K D E T K K F T V T E
BINARY_7999 T T Y K L I L N G K Q L K E E A I T E A V D A A I A E K Y I K L Y A N A K T V E G V W T Y K D E T K T A T V T E
BINARY_8000 T T Y K L I L N G K Q L K E E A I T E A V D A A I A E K Y I K L Y A N A K T V E G V W T Y K D E T K T F T V T E
BINARY_8001 T T Y K L I L N G K Q L K E E A I T E A V D A A I A E K Y F K L I A N A K T V E G V W T L K D E I L K A T V T E
BINARY_8002 T T Y K L I L N G K Q L K E E A I T E A V D A A I A E K Y F K L I A N A K T V E G V W T L K D E I L K F T V T E
BINARY_8003 T T Y K L I L N G K Q L K E E A I T E A V D A A I A E K Y F K L I A N A K T V E G V W T L K D E I L T A T V T E
BINARY_8004 T T Y K L I L N G K Q L K E E A I T E A V D A A I A E K Y F K L I A N A K T V E G V W T L K D E I L T F T V T E
BINARY_8005 T T Y K L I L N G K Q L K E E A I T E A V D A A I A E K Y F K L I A N A K T V E G V W T L K D E I K K A T V T E
BINARY_8006 T T Y K L I L N G K Q L K E E A I T E A V D A A I A E K Y F K L I A N A K T V E G V W T L K D E I K K F T V T E
BINARY_8007 T T Y K L I L N G K Q L K E E A I T E A V D A A I A E K Y F K L I A N A K T V E G V W T L K D E I K T A T V T E
BINARY_8008 T T Y K L I L N G K Q L K E E A I T E A V D A A I A E K Y F K L I A N A K T V E G V W T L K D E I K T F T V T E
BINARY_8009 T T Y K L I L N G K Q L K E E A I T E A V D A A I A E K Y F K L I A N A K T V E G V W T L K D E T L K A T V T E
BINARY_8010 T T Y K L I L N G K Q L K E E A I T E A V D A A I A E K Y F K L I A N A K T V E G V W T L K D E T L K F T V T E
BINARY_8011 T T Y K L I L N G K Q L K E E A I T E A V D A A I A E K Y F K L I A N A K T V E G V W T L K D E T L T A T V T E
BINARY_8012 T T Y K L I L N G K Q L K E E A I T E A V D A A I A E K Y F K L I A N A K T V E G V W T L K D E T L T F T V T E
BINARY_8013 T T Y K L I L N G K Q L K E E A I T E A V D A A I A E K Y F K L I A N A K T V E G V W T L K D E T K K A T V T E
BINARY_8014 T T Y K L I L N G K Q L K E E A I T E A V D A A I A E K Y F K L I A N A K T V E G V W T L K D E T K K F T V T E
BINARY_8015 T T Y K L I L N G K Q L K E E A I T E A V D A A I A E K Y F K L I A N A K T V E G V W T L K D E T K T A T V T E
BINARY_8016 T T Y K L I L N G K Q L K E E A I T E A V D A A I A E K Y F K L I A N A K T V E G V W T L K D E T K T F T V T E
BINARY_8017 T T Y K L I L N G K Q L K E E A I T E A V D A A I A E K Y F K L I A N A K T V E G V W T Y K D E I L K A T V T E
BINARY_8018 T T Y K L I L N G K Q L K E E A I T E A V D A A I A E K Y F K L I A N A K T V E G V W T Y K D E I L K F T V T E
BINARY_8019 T T Y K L I L N G K Q L K E E A I T E A V D A A I A E K Y F K L I A N A K T V E G V W T Y K D E I L T A T V T E
BINARY_8020 T T Y K L I L N G K Q L K E E A I T E A V D A A I A E K Y F K L I A N A K T V E G V W T Y K D E I L T F T V T E
BINARY_8021 T T Y K L I L N G K Q L K E E A I T E A V D A A I A E K Y F K L I A N A K T V E G V W T Y K D E I K K A T V T E
BINARY_8022 T T Y K L I L N G K Q L K E E A I T E A V D A A I A E K Y F K L I A N A K T V E G V W T Y K D E I K K F T V T E
BINARY_8023 T T Y K L I L N G K Q L K E E A I T E A V D A A I A E K Y F K L I A N A K T V E G V W T Y K D E I K T A T V T E
BINARY_8024 T T Y K L I L N G K Q L K E E A I T E A V D A A I A E K Y F K L I A N A K T V E G V W T Y K D E I K T F T V T E
BINARY_8025 T T Y K L I L N G K Q L K E E A I T E A V D A A I A E K Y F K L I A N A K T V E G V W T Y K D E T L K A T V T E
BINARY_8026 T T Y K L I L N G K Q L K E E A I T E A V D A A I A E K Y F K L I A N A K T V E G V W T Y K D E T L K F T V T E
BINARY_8027 T T Y K L I L N G K Q L K E E A I T E A V D A A I A E K Y F K L I A N A K T V E G V W T Y K D E T L T A T V T E
BINARY_8028 T T Y K L I L N G K Q L K E E A I T E A V D A A I A E K Y F K L I A N A K T V E G V W T Y K D E T L T F T V T E
BINARY_8029 T T Y K L I L N G K Q L K E E A I T E A V D A A I A E K Y F K L I A N A K T V E G V W T Y K D E T K K A T V T E
BINARY_8030 T T Y K L I L N G K Q L K E E A I T E A V D A A I A E K Y F K L I A N A K T V E G V W T Y K D E T K K F T V T E
BINARY_8031 T T Y K L I L N G K Q L K E E A I T E A V D A A I A E K Y F K L I A N A K T V E G V W T Y K D E T K T A T V T E
BINARY_8032 T T Y K L I L N G K Q L K E E A I T E A V D A A I A E K Y F K L I A N A K T V E G V W T Y K D E T K T F T V T E
BINARY_8033 T T Y K L I L N G K Q L K E E A I T E A V D A A I A E K Y F K L Y A N A K T V E G V W T L K D E I L K A T V T E
BINARY_8034 T T Y K L I L N G K Q L K E E A I T E A V D A A I A E K Y F K L Y A N A K T V E G V W T L K D E I L K F T V T E
BINARY_8035 T T Y K L I L N G K Q L K E E A I T E A V D A A I A E K Y F K L Y A N A K T V E G V W T L K D E I L T A T V T E
BINARY_8036 T T Y K L I L N G K Q L K E E A I T E A V D A A I A E K Y F K L Y A N A K T V E G V W T L K D E I L T F T V T E
BINARY_8037 T T Y K L I L N G K Q L K E E A I T E A V D A A I A E K Y F K L Y A N A K T V E G V W T L K D E I K K A T V T E
BINARY_8038 T T Y K L I L N G K Q L K E E A I T E A V D A A I A E K Y F K L Y A N A K T V E G V W T L K D E I K K F T V T E
BINARY_8039 T T Y K L I L N G K Q L K E E A I T E A V D A A I A E K Y F K L Y A N A K T V E G V W T L K D E I K T A T V T E
BINARY_8040 T T Y K L I L N G K Q L K E E A I T E A V D A A I A E K Y F K L Y A N A K T V E G V W T L K D E I K T F T V T E
BINARY_8041 T T Y K L I L N G K Q L K E E A I T E A V D A A I A E K Y F K L Y A N A K T V E G V W T L K D E T L K A T V T E
BINARY_8042 T T Y K L I L N G K Q L K E E A I T E A V D A A I A E K Y F K L Y A N A K T V E G V W T L K D E T L K F T V T E
BINARY_8043 T T Y K L I L N G K Q L K E E A I T E A V D A A I A E K Y F K L Y A N A K T V E G V W T L K D E T L T A T V T E
BINARY_8044 T T Y K L I L N G K Q L K E E A I T E A V D A A I A E K Y F K L Y A N A K T V E G V W T L K D E T L T F T V T E
BINARY_8045 T T Y K L I L N G K Q L K E E A I T E A V D A A I A E K Y F K L Y A N A K T V E G V W T L K D E T K K A T V T E
BINARY_8046 T T Y K L I L N G K Q L K E E A I T E A V D A A I A E K Y F K L Y A N A K T V E G V W T L K D E T K K F T V T E
BINARY_8047 T T Y K L I L N G K Q L K E E A I T E A V D A A I A E K Y F K L Y A N A K T V E G V W T L K D E T K T A T V T E
BINARY_8048 T T Y K L I L N G K Q L K E E A I T E A V D A A I A E K Y F K L Y A N A K T V E G V W T L K D E T K T F T V T E
BINARY_8049 T T Y K L I L N G K Q L K E E A I T E A V D A A I A E K Y F K L Y A N A K T V E G V W T Y K D E I L K A T V T E
BINARY_8050 T T Y K L I L N G K Q L K E E A I T E A V D A A I A E K Y F K L Y A N A K T V E G V W T Y K D E I L K F T V T E
BINARY_8051 T T Y K L I L N G K Q L K E E A I T E A V D A A I A E K Y F K L Y A N A K T V E G V W T Y K D E I L T A T V T E
BINARY_8052 T T Y K L I L N G K Q L K E E A I T E A V D A A I A E K Y F K L Y A N A K T V E G V W T Y K D E I L T F T V T E
BINARY_8053 T T Y K L I L N G K Q L K E E A I T E A V D A A I A E K Y F K L Y A N A K T V E G V W T Y K D E I K K A T V T E
BINARY_8054 T T Y K L I L N G K Q L K E E A I T E A V D A A I A E K Y F K L Y A N A K T V E G V W T Y K D E I K K F T V T E
BINARY_8055 T T Y K L I L N G K Q L K E E A I T E A V D A A I A E K Y F K L Y A N A K T V E G V W T Y K D E I K T A T V T E
BINARY_8056 T T Y K L I L N G K Q L K E E A I T E A V D A A I A E K Y F K L Y A N A K T V E G V W T Y K D E I K T F T V T E
BINARY_8057 T T Y K L I L N G K Q L K E E A I T E A V D A A I A E K Y F K L Y A N A K T V E G V W T Y K D E T L K A T V T E
BINARY_8058 T T Y K L I L N G K Q L K E E A I T E A V D A A I A E K Y F K L Y A N A K T V E G V W T Y K D E T L K F T V T E
BINARY_8059 T T Y K L I L N G K Q L K E E A I T E A V D A A I A E K Y F K L Y A N A K T V E G V W T Y K D E T L T A T V T E
BINARY_8060 T T Y K L I L N G K Q L K E E A I T E A V D A A I A E K Y F K L Y A N A K T V E G V W T Y K D E T L T F T V T E
BINARY_8061 T T Y K L I L N G K Q L K E E A I T E A V D A A I A E K Y F K L Y A N A K T V E G V W T Y K D E T K K A T V T E
BINARY_8062 T T Y K L I L N G K Q L K E E A I T E A V D A A I A E K Y F K L Y A N A K T V E G V W T Y K D E T K K F T V T E
BINARY_8063 T T Y K L I L N G K Q L K E E A I T E A V D A A I A E K Y F K L Y A N A K T V E G V W T Y K D E T K T A T V T E
BINARY_8064 T T Y K L I L N G K Q L K E E A I T E A V D A A I A E K Y F K L Y A N A K T V E G V W T Y K D E T K T F T V T E
BINARY_8065 T T Y K L I L N G K Q L K E E A I T E A V D A A T A E K Y I K L I A N A K T V E G V W T L K D E I L K A T V T E
BINARY_8066 T T Y K L I L N G K Q L K E E A I T E A V D A A T A E K Y I K L I A N A K T V E G V W T L K D E I L K F T V T E
BINARY_8067 T T Y K L I L N G K Q L K E E A I T E A V D A A T A E K Y I K L I A N A K T V E G V W T L K D E I L T A T V T E
BINARY_8068 T T Y K L I L N G K Q L K E E A I T E A V D A A T A E K Y I K L I A N A K T V E G V W T L K D E I L T F T V T E
BINARY_8069 T T Y K L I L N G K Q L K E E A I T E A V D A A T A E K Y I K L I A N A K T V E G V W T L K D E I K K A T V T E
BINARY_8070 T T Y K L I L N G K Q L K E E A I T E A V D A A T A E K Y I K L I A N A K T V E G V W T L K D E I K K F T V T E
BINARY_8071 T T Y K L I L N G K Q L K E E A I T E A V D A A T A E K Y I K L I A N A K T V E G V W T L K D E I K T A T V T E
BINARY_8072 T T Y K L I L N G K Q L K E E A I T E A V D A A T A E K Y I K L I A N A K T V E G V W T L K D E I K T F T V T E
BINARY_8073 T T Y K L I L N G K Q L K E E A I T E A V D A A T A E K Y I K L I A N A K T V E G V W T L K D E T L K A T V T E
BINARY_8074 T T Y K L I L N G K Q L K E E A I T E A V D A A T A E K Y I K L I A N A K T V E G V W T L K D E T L K F T V T E
BINARY_8075 T T Y K L I L N G K Q L K E E A I T E A V D A A T A E K Y I K L I A N A K T V E G V W T L K D E T L T A T V T E
BINARY_8076 T T Y K L I L N G K Q L K E E A I T E A V D A A T A E K Y I K L I A N A K T V E G V W T L K D E T L T F T V T E
BINARY_8077 T T Y K L I L N G K Q L K E E A I T E A V D A A T A E K Y I K L I A N A K T V E G V W T L K D E T K K A T V T E
BINARY_8078 T T Y K L I L N G K Q L K E E A I T E A V D A A T A E K Y I K L I A N A K T V E G V W T L K D E T K K F T V T E
BINARY_8079 T T Y K L I L N G K Q L K E E A I T E A V D A A T A E K Y I K L I A N A K T V E G V W T L K D E T K T A T V T E
BINARY_8080 T T Y K L I L N G K Q L K E E A I T E A V D A A T A E K Y I K L I A N A K T V E G V W T L K D E T K T F T V T E
BINARY_8081 T T Y K L I L N G K Q L K E E A I T E A V D A A T A E K Y I K L I A N A K T V E G V W T Y K D E I L K A T V T E
BINARY_8082 T T Y K L I L N G K Q L K E E A I T E A V D A A T A E K Y I K L I A N A K T V E G V W T Y K D E I L K F T V T E
BINARY_8083 T T Y K L I L N G K Q L K E E A I T E A V D A A T A E K Y I K L I A N A K T V E G V W T Y K D E I L T A T V T E
BINARY_8084 T T Y K L I L N G K Q L K E E A I T E A V D A A T A E K Y I K L I A N A K T V E G V W T Y K D E I L T F T V T E
BINARY_8085 T T Y K L I L N G K Q L K E E A I T E A V D A A T A E K Y I K L I A N A K T V E G V W T Y K D E I K K A T V T E
BINARY_8086 T T Y K L I L N G K Q L K E E A I T E A V D A A T A E K Y I K L I A N A K T V E G V W T Y K D E I K K F T V T E
BINARY_8087 T T Y K L I L N G K Q L K E E A I T E A V D A A T A E K Y I K L I A N A K T V E G V W T Y K D E I K T A T V T E
BINARY_8088 T T Y K L I L N G K Q L K E E A I T E A V D A A T A E K Y I K L I A N A K T V E G V W T Y K D E I K T F T V T E
BINARY_8089 T T Y K L I L N G K Q L K E E A I T E A V D A A T A E K Y I K L I A N A K T V E G V W T Y K D E T L K A T V T E
BINARY_8090 T T Y K L I L N G K Q L K E E A I T E A V D A A T A E K Y I K L I A N A K T V E G V W T Y K D E T L K F T V T E
BINARY_8091 T T Y K L I L N G K Q L K E E A I T E A V D A A T A E K Y I K L I A N A K T V E G V W T Y K D E T L T A T V T E

BINARY_8092 T T Y K L I L N G K Q L K E E A I T E A V D A A T A E K Y I K L I A N A K T V E G V W T Y K D E T L T F T V T E
BINARY_8093 T T Y K L I L N G K Q L K E E A I T E A V D A A T A E K Y I K L I A N A K T V E G V W T Y K D E T K K A T V T E
BINARY_8094 T T Y K L I L N G K Q L K E E A I T E A V D A A T A E K Y I K L I A N A K T V E G V W T Y K D E T K K F T V T E
BINARY_8095 T T Y K L I L N G K Q L K E E A I T E A V D A A T A E K Y I K L I A N A K T V E G V W T Y K D E T K T A T V T E
BINARY_8096 T T Y K L I L N G K Q L K E E A I T E A V D A A T A E K Y I K L I A N A K T V E G V W T Y K D E T K T F T V T E
BINARY_8097 T T Y K L I L N G K Q L K E E A I T E A V D A A T A E K Y I K L Y A N A K T V E G V W T L K D E I L K A T V T E
BINARY_8098 T T Y K L I L N G K Q L K E E A I T E A V D A A T A E K Y I K L Y A N A K T V E G V W T L K D E I L K F T V T E
BINARY_8099 T T Y K L I L N G K Q L K E E A I T E A V D A A T A E K Y I K L Y A N A K T V E G V W T L K D E I L T A T V T E
BINARY_8100 T T Y K L I L N G K Q L K E E A I T E A V D A A T A E K Y I K L Y A N A K T V E G V W T L K D E I L T F T V T E
BINARY_8101 T T Y K L I L N G K Q L K E E A I T E A V D A A T A E K Y I K L Y A N A K T V E G V W T L K D E I K K A T V T E
BINARY_8102 T T Y K L I L N G K Q L K E E A I T E A V D A A T A E K Y I K L Y A N A K T V E G V W T L K D E I K K F T V T E
BINARY_8103 T T Y K L I L N G K Q L K E E A I T E A V D A A T A E K Y I K L Y A N A K T V E G V W T L K D E I K T A T V T E
BINARY_8104 T T Y K L I L N G K Q L K E E A I T E A V D A A T A E K Y I K L Y A N A K T V E G V W T L K D E I K T F T V T E
BINARY_8105 T T Y K L I L N G K Q L K E E A I T E A V D A A T A E K Y I K L Y A N A K T V E G V W T L K D E T L K A T V T E
BINARY_8106 T T Y K L I L N G K Q L K E E A I T E A V D A A T A E K Y I K L Y A N A K T V E G V W T L K D E T L K F T V T E
BINARY_8107 T T Y K L I L N G K Q L K E E A I T E A V D A A T A E K Y I K L Y A N A K T V E G V W T L K D E T L T A T V T E
BINARY_8108 T T Y K L I L N G K Q L K E E A I T E A V D A A T A E K Y I K L Y A N A K T V E G V W T L K D E T L T F T V T E
BINARY_8109 T T Y K L I L N G K Q L K E E A I T E A V D A A T A E K Y I K L Y A N A K T V E G V W T L K D E T K K A T V T E
BINARY_8110 T T Y K L I L N G K Q L K E E A I T E A V D A A T A E K Y I K L Y A N A K T V E G V W T L K D E T K K F T V T E
BINARY_8111 T T Y K L I L N G K Q L K E E A I T E A V D A A T A E K Y I K L Y A N A K T V E G V W T L K D E T K T A T V T E
BINARY_8112 T T Y K L I L N G K Q L K E E A I T E A V D A A T A E K Y I K L Y A N A K T V E G V W T L K D E T K T F T V T E
BINARY_8113 T T Y K L I L N G K Q L K E E A I T E A V D A A T A E K Y I K L Y A N A K T V E G V W T Y K D E I L K A T V T E
BINARY_8114 T T Y K L I L N G K Q L K E E A I T E A V D A A T A E K Y I K L Y A N A K T V E G V W T Y K D E I L K F T V T E
BINARY_8115 T T Y K L I L N G K Q L K E E A I T E A V D A A T A E K Y I K L Y A N A K T V E G V W T Y K D E I L T A T V T E
BINARY_8116 T T Y K L I L N G K Q L K E E A I T E A V D A A T A E K Y I K L Y A N A K T V E G V W T Y K D E I L T F T V T E
BINARY_8117 T T Y K L I L N G K Q L K E E A I T E A V D A A T A E K Y I K L Y A N A K T V E G V W T Y K D E I K K A T V T E
BINARY_8118 T T Y K L I L N G K Q L K E E A I T E A V D A A T A E K Y I K L Y A N A K T V E G V W T Y K D E I K K F T V T E
BINARY_8119 T T Y K L I L N G K Q L K E E A I T E A V D A A T A E K Y I K L Y A N A K T V E G V W T Y K D E I K T A T V T E
BINARY_8120 T T Y K L I L N G K Q L K E E A I T E A V D A A T A E K Y I K L Y A N A K T V E G V W T Y K D E I K T F T V T E
BINARY_8121 T T Y K L I L N G K Q L K E E A I T E A V D A A T A E K Y I K L Y A N A K T V E G V W T Y K D E T L K A T V T E
BINARY_8122 T T Y K L I L N G K Q L K E E A I T E A V D A A T A E K Y I K L Y A N A K T V E G V W T Y K D E T L K F T V T E
BINARY_8123 T T Y K L I L N G K Q L K E E A I T E A V D A A T A E K Y I K L Y A N A K T V E G V W T Y K D E T L T A T V T E
BINARY_8124 T T Y K L I L N G K Q L K E E A I T E A V D A A T A E K Y I K L Y A N A K T V E G V W T Y K D E T L T F T V T E
BINARY_8125 T T Y K L I L N G K Q L K E E A I T E A V D A A T A E K Y I K L Y A N A K T V E G V W T Y K D E T K K A T V T E
BINARY_8126 T T Y K L I L N G K Q L K E E A I T E A V D A A T A E K Y I K L Y A N A K T V E G V W T Y K D E T K K F T V T E
BINARY_8127 T T Y K L I L N G K Q L K E E A I T E A V D A A T A E K Y I K L Y A N A K T V E G V W T Y K D E T K T A T V T E
BINARY_8128 T T Y K L I L N G K Q L K E E A I T E A V D A A T A E K Y I K L Y A N A K T V E G V W T Y K D E T K T F T V T E
BINARY_8129 T T Y K L I L N G K Q L K E E A I T E A V D A A T A E K Y F K L I A N A K T V E G V W T L K D E I L K A T V T E
BINARY_8130 T T Y K L I L N G K Q L K E E A I T E A V D A A T A E K Y F K L I A N A K T V E G V W T L K D E I L K F T V T E
BINARY_8131 T T Y K L I L N G K Q L K E E A I T E A V D A A T A E K Y F K L I A N A K T V E G V W T L K D E I L T A T V T E
BINARY_8132 T T Y K L I L N G K Q L K E E A I T E A V D A A T A E K Y F K L I A N A K T V E G V W T L K D E I L T F T V T E
BINARY_8133 T T Y K L I L N G K Q L K E E A I T E A V D A A T A E K Y F K L I A N A K T V E G V W T L K D E I K K A T V T E
BINARY_8134 T T Y K L I L N G K Q L K E E A I T E A V D A A T A E K Y F K L I A N A K T V E G V W T L K D E I K K F T V T E
BINARY_8135 T T Y K L I L N G K Q L K E E A I T E A V D A A T A E K Y F K L I A N A K T V E G V W T L K D E I K T A T V T E
BINARY_8136 T T Y K L I L N G K Q L K E E A I T E A V D A A T A E K Y F K L I A N A K T V E G V W T L K D E I K T F T V T E
BINARY_8137 T T Y K L I L N G K Q L K E E A I T E A V D A A T A E K Y F K L I A N A K T V E G V W T L K D E T L K A T V T E
BINARY_8138 T T Y K L I L N G K Q L K E E A I T E A V D A A T A E K Y F K L I A N A K T V E G V W T L K D E T L K F T V T E
BINARY_8139 T T Y K L I L N G K Q L K E E A I T E A V D A A T A E K Y F K L I A N A K T V E G V W T L K D E T L T A T V T E
BINARY_8140 T T Y K L I L N G K Q L K E E A I T E A V D A A T A E K Y F K L I A N A K T V E G V W T L K D E T L T F T V T E
BINARY_8141 T T Y K L I L N G K Q L K E E A I T E A V D A A T A E K Y F K L I A N A K T V E G V W T L K D E T K K A T V T E
BINARY_8142 T T Y K L I L N G K Q L K E E A I T E A V D A A T A E K Y F K L I A N A K T V E G V W T L K D E T K K F T V T E
BINARY_8143 T T Y K L I L N G K Q L K E E A I T E A V D A A T A E K Y F K L I A N A K T V E G V W T L K D E T K T A T V T E
BINARY_8144 T T Y K L I L N G K Q L K E E A I T E A V D A A T A E K Y F K L I A N A K T V E G V W T L K D E T K T F T V T E
BINARY_8145 T T Y K L I L N G K Q L K E E A I T E A V D A A T A E K Y F K L I A N A K T V E G V W T Y K D E I L K A T V T E
BINARY_8146 T T Y K L I L N G K Q L K E E A I T E A V D A A T A E K Y F K L I A N A K T V E G V W T Y K D E I L K F T V T E
BINARY_8147 T T Y K L I L N G K Q L K E E A I T E A V D A A T A E K Y F K L I A N A K T V E G V W T Y K D E I L T A T V T E
BINARY_8148 T T Y K L I L N G K Q L K E E A I T E A V D A A T A E K Y F K L I A N A K T V E G V W T Y K D E I L T F T V T E
BINARY_8149 T T Y K L I L N G K Q L K E E A I T E A V D A A T A E K Y F K L I A N A K T V E G V W T Y K D E I K K A T V T E
BINARY_8150 T T Y K L I L N G K Q L K E E A I T E A V D A A T A E K Y F K L I A N A K T V E G V W T Y K D E I K K F T V T E
BINARY_8151 T T Y K L I L N G K Q L K E E A I T E A V D A A T A E K Y F K L I A N A K T V E G V W T Y K D E I K T A T V T E
BINARY_8152 T T Y K L I L N G K Q L K E E A I T E A V D A A T A E K Y F K L I A N A K T V E G V W T Y K D E I K T F T V T E
BINARY_8153 T T Y K L I L N G K Q L K E E A I T E A V D A A T A E K Y F K L I A N A K T V E G V W T Y K D E T L K A T V T E
BINARY_8154 T T Y K L I L N G K Q L K E E A I T E A V D A A T A E K Y F K L I A N A K T V E G V W T Y K D E T L K F T V T E
BINARY_8155 T T Y K L I L N G K Q L K E E A I T E A V D A A T A E K Y F K L I A N A K T V E G V W T Y K D E T L T A T V T E
BINARY_8156 T T Y K L I L N G K Q L K E E A I T E A V D A A T A E K Y F K L I A N A K T V E G V W T Y K D E T L T F T V T E
BINARY_8157 T T Y K L I L N G K Q L K E E A I T E A V D A A T A E K Y F K L I A N A K T V E G V W T Y K D E T K K A T V T E
BINARY_8158 T T Y K L I L N G K Q L K E E A I T E A V D A A T A E K Y F K L I A N A K T V E G V W T Y K D E T K K F T V T E
BINARY_8159 T T Y K L I L N G K Q L K E E A I T E A V D A A T A E K Y F K L I A N A K T V E G V W T Y K D E T K T A T V T E
BINARY_8160 T T Y K L I L N G K Q L K E E A I T E A V D A A T A E K Y F K L I A N A K T V E G V W T Y K D E T K T F T V T E
BINARY_8161 T T Y K L I L N G K Q L K E E A I T E A V D A A T A E K Y F K L Y A N A K T V E G V W T L K D E I L K A T V T E
BINARY_8162 T T Y K L I L N G K Q L K E E A I T E A V D A A T A E K Y F K L Y A N A K T V E G V W T L K D E I L K F T V T E
BINARY_8163 T T Y K L I L N G K Q L K E E A I T E A V D A A T A E K Y F K L Y A N A K T V E G V W T L K D E I L T A T V T E
BINARY_8164 T T Y K L I L N G K Q L K E E A I T E A V D A A T A E K Y F K L Y A N A K T V E G V W T L K D E I L T F T V T E
BINARY_8165 T T Y K L I L N G K Q L K E E A I T E A V D A A T A E K Y F K L Y A N A K T V E G V W T L K D E I K K A T V T E
BINARY_8166 T T Y K L I L N G K Q L K E E A I T E A V D A A T A E K Y F K L Y A N A K T V E G V W T L K D E I K K F T V T E
BINARY_8167 T T Y K L I L N G K Q L K E E A I T E A V D A A T A E K Y F K L Y A N A K T V E G V W T L K D E I K T A T V T E
BINARY_8168 T T Y K L I L N G K Q L K E E A I T E A V D A A T A E K Y F K L Y A N A K T V E G V W T L K D E I K T F T V T E
BINARY_8169 T T Y K L I L N G K Q L K E E A I T E A V D A A T A E K Y F K L Y A N A K T V E G V W T L K D E T L K A T V T E
BINARY_8170 T T Y K L I L N G K Q L K E E A I T E A V D A A T A E K Y F K L Y A N A K T V E G V W T L K D E T L K F T V T E
BINARY_8171 T T Y K L I L N G K Q L K E E A I T E A V D A A T A E K Y F K L Y A N A K T V E G V W T L K D E T L T A T V T E
BINARY_8172 T T Y K L I L N G K Q L K E E A I T E A V D A A T A E K Y F K L Y A N A K T V E G V W T L K D E T L T F T V T E
BINARY_8173 T T Y K L I L N G K Q L K E E A I T E A V D A A T A E K Y F K L Y A N A K T V E G V W T L K D E T K K A T V T E
BINARY_8174 T T Y K L I L N G K Q L K E E A I T E A V D A A T A E K Y F K L Y A N A K T V E G V W T L K D E T K K F T V T E
BINARY_8175 T T Y K L I L N G K Q L K E E A I T E A V D A A T A E K Y F K L Y A N A K T V E G V W T L K D E T K T A T V T E
BINARY_8176 T T Y K L I L N G K Q L K E E A I T E A V D A A T A E K Y F K L Y A N A K T V E G V W T L K D E T K T F T V T E
BINARY_8177 T T Y K L I L N G K Q L K E E A I T E A V D A A T A E K Y F K L Y A N A K T V E G V W T Y K D E I L K A T V T E
BINARY_8178 T T Y K L I L N G K Q L K E E A I T E A V D A A T A E K Y F K L Y A N A K T V E G V W T Y K D E I L K F T V T E
BINARY_8179 T T Y K L I L N G K Q L K E E A I T E A V D A A T A E K Y F K L Y A N A K T V E G V W T Y K D E I L T A T V T E
BINARY_8180 T T Y K L I L N G K Q L K E E A I T E A V D A A T A E K Y F K L Y A N A K T V E G V W T Y K D E I L T F T V T E
BINARY_8181 T T Y K L I L N G K Q L K E E A I T E A V D A A T A E K Y F K L Y A N A K T V E G V W T Y K D E I K K A T V T E
BINARY_8182 T T Y K L I L N G K Q L K E E A I T E A V D A A T A E K Y F K L Y A N A K T V E G V W T Y K D E I K K F T V T E
BINARY_8183 T T Y K L I L N G K Q L K E E A I T E A V D A A T A E K Y F K L Y A N A K T V E G V W T Y K D E I K T A T V T E
BINARY_8184 T T Y K L I L N G K Q L K E E A I T E A V D A A T A E K Y F K L Y A N A K T V E G V W T Y K D E I K T F T V T E
BINARY_8185 T T Y K L I L N G K Q L K E E A I T E A V D A A T A E K Y F K L Y A N A K T V E G V W T Y K D E T L K A T V T E
BINARY_8186 T T Y K L I L N G K Q L K E E A I T E A V D A A T A E K Y F K L Y A N A K T V E G V W T Y K D E T L K F T V T E

BINARY_8187 T T Y K L I L N G K Q L K E E A I T E A V D A A T A E K Y F K L Y A N A K T V E G V W T Y K D E T L T A T V T E
BINARY_8188 T T Y K L I L N G K Q L K E E A I T E A V D A A T A E K Y F K L Y A N A K T V E G V W T Y K D E T L T F T V T E
BINARY_8189 T T Y K L I L N G K Q L K E E A I T E A V D A A T A E K Y F K L Y A N A K T V E G V W T Y K D E T K K A T V T E
BINARY_8190 T T Y K L I L N G K Q L K E E A I T E A V D A A T A E K Y F K L Y A N A K T V E G V W T Y K D E T K K F T V T E
BINARY_8191 T T Y K L I L N G K Q L K E E A I T E A V D A A T A E K Y F K L Y A N A K T V E G V W T Y K D E T K T A T V T E
BINARY_8192 T T Y K L I L N G K Q L K E E A I T E A V D A A T A E K Y F K L Y A N A K T V E G V W T Y K D E T K T F T V T E

### Table S9. 56-Residue Chains Sampling Random Sequence Space.

RANDOM_1 A G G G H C S V K R S I N K V V S N E E I W V V K F R V F H N K S E V W R D G S A I S T G N T R E E H L E L F L
RANDOM_2 N E E Y T P E A V E F S S G A A K A T Q K T G S Y D L R F E S L H L R S D I E A Q F S S H P S Q V K N D S Y Q R
RANDOM_3 R S K Y R E D Q L S A K E T Y I A Q A S V F L K N F M R N Y T V I L L A A G S T N D R D W P Q S T L A L T S L K
RANDOM_4 V V K K V N G D L L K L H T T Q N I P S R T A Q I I L I S A M F G A V P E L P C L Y P P R L P R G F T F Q N Y S
RANDOM_5 K A L V V R A Q F V S P G G V E G P T I D V P V N R P I I T F L N V V S L N D E V G K D E H R T Q G L Q E T S A
RANDOM_6 Y N M K R S K R S L N S V P Q V E N R Y A L G R V A A D K F Q N P L N I E P E R V S V F I F D A Q G Y T D P V P
RANDOM_7 D I R S Q G G V T E G Q R F A S M A N A S D V L L S V S G K S L V S A G Y Y V D A H P D C F R S D L H L L L F T
RANDOM_8 T V T T Q P P A P G Q F G D E L D S K I G A V D R Q V Q L P V D G E S A A Y G S P G P H L E V L C D R I E Q C C
RANDOM_9 L C A I R E L K Y L L Y S N N D L Y R G F H V D L N I M A L P S M S K V S F P R S P R R P S L T T S I N A G L G
RANDOM_10 V T G R N N L T P A E N L L Y P S V H H M N C Y F D L E Q K I R D L Q V I E L P L Q V Q L H L D L L Y K G T G I
RANDOM_11 I N M D D I G Q K P I V A E E I D T N K K V K Q V L L A E E S S G M D I P F N G E Q Q S Q K G Q S G L N E S Q G
RANDOM_12 E R Y K E I Y F I L Q N L N A S G Y G I F F F D F G G D V P G F V E N D F R Q I P S T L T S N R Y V V K T D I N
RANDOM_13 E A A A K W S K S E V E P N L S G L I D A S K F I F M C V C N N G Q R V N V D G G V S K M N D P G F S F A V D L
RANDOM_14 Q L V L I I V C F T W A R Y L T F R M H R L K G F D S D H A T T S E D P T C F T T G G Q E N E L L T S V S F F T
RANDOM_15 H C V P S F L L N A C T T T D V I I E M R R E L L T T G S I H T L T P T S D V V G F E D V Q P S G D T V R I P I
RANDOM_16 K Q Y S S E R N Q E D R D F N I T S P E A L S R V F F I A G F H S G R S A L K Q P G A L Q Q Q R V K R K D I R L
RANDOM_17 G T S L D D P L T I A D L L H Q R P S H S Q S N N H Y Q E P Q P W I R A Y C A P A V P L Q T N P E A V Y L G G D
RANDOM_18 L R P L Y E R A M D W W G N F I C A A V N K R G S G D M I F I M G P R V R P T K D E T Q G E L V E L T E W L E E
RANDOM_19 N D L Q Q H N L S G R I L A K C F L G C V L M E P G E H T I E K E V G D K D G T R W G C T L L Y V F Q M Q G P P
RANDOM_20 A W L R P L T H T Y D G P V N S H N V G Y E Y D S P P S G L E K C G L L E F V R R P T N F T S S D V S A F Y K F
RANDOM_21 C R Y G W N E G N I F R P Y K R L A I I V Y E L V A A I V T F F G F T L K L V A G T G G K L Q A G N D V K E T R
RANDOM_22 Y Q F K M Q N F T L P A E P G L V N I S A H Q D R A L I F L I R L S T L N I L V M S S A Q N R L P K L E P Q Q C
RANDOM_23 Q L N W S R V D M A K T N N G I I Q L A Q R L S K K T L S G K L H E R I T F Q D G P T G N A E G T E H E T L A S
RANDOM_24 G A H Q K L S T E K P M A Q I R G A K A C L N K S S E E G P Q P S E G R I L T G D D D A R N Q T A I R I Q Y G E
RANDOM_25 A F L T V L L Q V F L L V T F F S E V G Y G L Y I I M T F N Q C N I D D T V L Q K H K N N A A K I T N L E V E V
RANDOM_26 P L R Q E Q E I L G N P K L K E S N D I Y W V A P S A T G P D K R A L H A G G H A T S L E T Q G K G A A T L K E
RANDOM_27 A G P K Q A M Q E Q S W Q G S F D W A S S P F K K V I G G G A L T W G G V K G V R S Q E Y I S L L A G G F T E V
RANDOM_28 P M C C R I R E N A T C L V L Y V E N P D L E A V H E E K A N F T S G R F A H E Y L D Q K E I A E V L A I L F C
RANDOM_29 D A G K G H P R S N T I G T A F F H G R E Q W V G I P G T S A I A Q L V A E G T G Q E I L T A L A M V E N F N S
RANDOM_30 D W E I N C C R L L A P Q L G Q W G E L K A H Q K G K I A D D K T V P N G K S I E K D L A G E T H R A Y A K D T
RANDOM_31 A D P I L V H E K C D L K A R P D L N D D V G D F N L K K I K R A E G V D E Q C D Y L K S V G V S I A A V R I G
RANDOM_32 G T F S M F G T Y F L P A V D N P Q L Y V S S D L A P S E E D Y K L F D V I M T H F I K D L N R C D A E Y S N D
RANDOM_33 H V Y S N R R S A E R T Q F R G F I G G M G L F G P E V E Y N F V T G M F G R W V A P H S D D T S F I Q N G T L
RANDOM_34 R F L S A F F R V Y P R F P N F K K S L Q K T K D V T T D G Y Q V L Y G V F E K V I E R L S D I A P K Y G G F D
RANDOM_35 K I A D G N V G S S D K N F N V E R Y F A I A R T I Y T V G Q Q K G Y T P G W V G D T V T N T L R R L A P K D G
RANDOM_36 L L Q L K K F F A E A A Q A D F A D A Y S A A P P L C A G A T M T E A Q D E K A A G D T E L H V Y A H R L C K I
RANDOM_37 S V P F Y G G D W E T Q S T L K H D S F N Q V I L I K E C Q A S G F G M L L L L L Q M R M K N V T S L S E I Q S
RANDOM_38 V A N V K F I R K P L I F K V I L R T Y S D K P S A S E P T W V R D N S L L G V C T M C H D G Y K S H F T N D S
RANDOM_39 G Y P L P G L P C W K D I E I T E M K I G Y G A M F I T K R E I E E V Y G F K S T V W E R V Y L A W M I S F D V
RANDOM_40 G I R R S D R I L D P A T P T E G G T L M S A D G N G I I L G C M P D K L N G I A E K T V Q F S Q I S L K R E L
RANDOM_41 A V K R D D T S V F L Y P G T I G P A C L K M Y I E K K E E Q T F I F I V E P A T N E G Y M E E D E Y A S L I L
RANDOM_42 Y G F K D G A W T E P S N M A T C F A G L T R G Q P S G Y A C E L D A T K I A V S P A L F V M C V S L I Y I P T
RANDOM_43 L S D V D V G D K P L P S L L F N H M A R A V L S H P T P C L S V N E E Y G R A E H E I R T K L R N L G Y I A F
RANDOM_44 V T H V V L K D A Y L A S T F Y S V V Y F P H P N V D S S N Q S S M R R R Q I L N R Y V T P C R T S L Q Q E H I
RANDOM_45 T P F E L L F T G H N A A R I L E W L S Y E L K K I L R H V E T T T L G E Q F G A E D N L F V R G Q N I T G G V
RANDOM_46 K E L E I V E K T F N M V L I S H G N L L K R L G N C K G K F H T T A S A Q N M A A D V D S H L L T P G R H K C
RANDOM_47 K I S A Q G S D G P L S L L N N V S C A V M M N A M E F F D I E M R D Q G A C G R H L S D G T A R N Q C C H S L
RANDOM_48 P A G G G Q K L D F E E D P S G A F Y M E A E H L F Q D T F Q T S M P A R P S E P A P K Q Q G R D N A F Q V L G
RANDOM_49 K L C T F A Q G P N L E R R E G R D T L L Q N T C D R L K N L Y L H C K D A S S K C P F V C A T S D A P S P K H
RANDOM_50 K Q L S C G V K P S T D R G M I I K D G Y R T M P N S P D A G Y E V Q R T R M S I A T M A V L E P Q D L Q K A R
RANDOM_51 V Y Y T A K P D V K R A P E T V P H S G K G Q L P G L T V T L S W T C E P A T D A D C K G A W K H M L M D L L T
RANDOM_52 V K V A I T G D G L G T Y K L I K G E Q K K T R S N A D R A P G L E L F Q L I N W S M A L V G V G T L L G D E V
RANDOM_53 A G S L V P A K Q Y L A V V D V T K H L P R L K T Q S S I S D E P V N Y G S P V L G T Q K V L S F I P H V M M T
RANDOM_54 H A Y T T A E S E E K G E F L F A D D L R L N R E P F A T N L R G M D T P L C T W N G K L N I L H E K L T H P G
RANDOM_55 E L R L I P A S E E S T E K F Q F A E F A E P F L Y D S Y I I L A F W A N I S A K D K N N E E L F Q V A R L P I
RANDOM_56 G N Y V L R V Y E N M S K P T C Q S L A S Q Y C N W E M L L P E H A G L Y S S L F A A K S F L S P S Q N W S E S
RANDOM_57 K E P T A T K Q A D F Q N M L V S Q D V Q I E E Q S Q S V E I G D G T G S Y G N D P V I Q E E S D K R M A L P Q
RANDOM_58 Y T G V G F S L G Q S C E V Q Y N A V F D Q R F L F I S P S T P M F E T S T L I N D L A L S I L G T K R I A D T
RANDOM_59 V T I T S V P S E R V T Y Y P G F P H C A E L E F E P A P L T N E R V P K F Y L A V C R G P K S D L K L G L S A
RANDOM_60 R Y L T S T S L N E D E L Y Q V A T H L G Y T G G R D G L I V L H P E G T H F N D L I A N N T P L Q A T C V N V
RANDOM_61 K C A R D Q A T L G V Q G Q R M S G F H T E Q G G P E N M F T H K F Q E H N E V E Q I P L N Q G D M A C A V C D
RANDOM_62 Q I A K E F N Y K Q F S H V S H E E N P V T E L D Y A L G C R I F G T H M Q A S K D N E D S L T R F G H N G R S
RANDOM_63 L E T T T M P D A D Y G G E Q S A Q P H E I L M R K R F A T L N F A M D A A E A T L Q C H N L S D G V E G E Y G
RANDOM_64 K L E G S I S A N L V L N I Y D V F Y I I N M T F I N L I I E M Y A A I G L E V T F G L S L G V V P A I D D G F
RANDOM_65 R C I F S E E E P V C N I K D Q G F A M R Y T I R P D S F L Q G L V K D S T L V R Q K N H A G L K T N R F L G E
RANDOM_66 E F T L H D H S A V D L L D D D Q D Q P E L L I I I E G D I E L K L R L E V Y A F L I V V E F I D W N L K E P C
RANDOM_67 I R E P S I G L Q N N E A A N F V I S Q G L E I R E V K G S A V T L L F S A N P L S D V N A H N V R A P L V E H
RANDOM_68 N R A K G T I A T L I R G Q R A R F L P N I R K Q A E I R I A V S E T D V Y V F E E P K S A T Q K E A G D C V R
RANDOM_69 G L W I E R I T I A N P D Q I E V G S Y A G G I N V L T P A A K K E A E K L G G W G A K M C S D D S G S K I A E
RANDOM_70 K E F D N E M D E S P P P L E S L R E S T E W M T S G G G T G P A R R K C K A F S S G L R M I V I I L A K A C F
RANDOM_71 I V T K F A R P H G L S D E S M L E M V Y D L K L G E S T R L E K D I A Q Y R R K S L L L A I K T K T P G E P L
RANDOM_72 L T E S L T G G R P A C I K I G I D A T V H C K A I L D G Q F V P L P H R A L G K S P L G S P E D K Q I Y Q D A
RANDOM_73 S E A R L L Q S Q E N A T K C A N G G P E M I V V E G P L S V Y A E Y D E S K I I Q A P L P V A N L T P S F K I
RANDOM_74 S M G D D I D G T H K L M L K G K G Q I K R A T K S A I K L Q Y M T T I V D D Q P A A D D I L N D R G P L L T I
RANDOM_75 Y H L N E E P A S D A F M K I D H A E L G K P C P D K E R I S D S R G G V Q Q K E Q Y W C T E P I L A L V A W E
RANDOM_76 G W V Q D I S C L D A Y F L K C I S W C Y A G T K G L T I R A I G Q A Q S R E V G M K I A S Q A G Q T T C Y S Q
RANDOM_77 L R V P P Y L R I Y V N Y L L N P K K E L F P E L R C G T L N G V R I P A A V P G R P Y A S A R N A P G Q Y T T
RANDOM_78 A N A I D V T G F I C R L T I G I K T I T V P T I P F I V P V A I D G G K H T E A E S E K P E I D M L G T W P K
RANDOM_79 S I V K H I V Q S K S P S E N E Y V L N T L L R R R A A F K P W R L G G Q P E Q K R S N E A T K L T H A L D N L
RANDOM_80 L S P I I G Q K Q T Q Y D R A L V D N P A S P P L T T S F F F A S L C D Y Y P S R Q L Q D T R V P I W P A R E S

RANDOM_81 N V L G T R G R L A Y N R L A D L L E V A Q L T Y K G I T Y G L R Q V S M L L L Q H A F G T K Q F E I T M Y Q V
RANDOM_82 S H L L S I D F F V G K P V P L Q Q G D L F M L G L Y E S L S E W G G A E M G T T N S K L S K H S T V T A I S M
RANDOM_83 L L A L L S Y K S N Q A G R W Q V R D V S F Y T E Y N E D N P I G R Q G S V G K N L G L L R K K V A A V Q T A F
RANDOM_84 V A V N L K S G P D K A I T Y C E G I P H I F Y K Q R S E Y Q R N L D L S Q E C T A M Y E V K L P L L N G D S A
RANDOM_85 A D E F A A L A S Q M S E H E S D A S S R K R Q C D S F R R S D D P K N E H R G R I V T Q F W S R R L I E N S L
RANDOM_86 A A T Y T H A F G E T V I A F A E P R L S V P Y E T R V W H K K E V H E S A P S L L Q G T Q W T Y D A W V S K S
RANDOM_87 P S G L V G L A A G Q T R E L H F P E E D T I Y Y Y G G L Y G D A R G F T A L P L N K F E R Q E E I E G L M V P
RANDOM_88 A I F S V S A Q S Q L E K T Q L M D E K L L S A I I N M Q H L Y L R A R R V K N L T H H D P G G W V E A R E E F
RANDOM_89 R L I D Q G C S R I T A I P R P T E N G F E F P Q G S P G K N F Q D L E L A Q A V M S G I K V V V R E K L F D Q
RANDOM_90 L R L F A K I F C G L R L F Y D E T I F L G Y L G I W K K D M L E N P A G Y I I M D G E L P L T T P H L C A G K
RANDOM_91 P N V Y P S L T Y H T E E R A Q G A A S L V R N N G A V N V H E P T D V I F A C L N H V V V C K L T M Q V N K Q
RANDOM_92 P R H N F Y H S C K Q H D L N T L I M V R T E W L T R H T L S L P S A Y D L N L S H A K P N V F G I E R D K E G
RANDOM_93 R N G G D T A A G A P D S K K G L F A F G C Q G N L Q V Y D F L W A V A L S V N Q R G L L L E L D A F L K L Y G
RANDOM_94 V Y G I L E V K C S L P A K P T H L S Y A S V E L N H K E W V N P L A L R Y N A N D E D H F I P V A I V Q C W S
RANDOM_95 L K L S L V T M A L P H T R N M K D G Q A R Q T N A Q N G K L T L Q T V A G F S T S Y V A Y F D N V L I N M S T
RANDOM_96 G K N D L M M I L Y A G G V V L V G L W P R K G D G R G G F F S L T A L I K E S G Q L V S P M Q F V V D S N E V
RANDOM_97 P N L S I I E D A V A T K L Y V L L L A D S A G H S H R E D N R S G G V N F Q T V K F A E D P N Q R A N E E L V
RANDOM_98 V N N D E K V L I H S Q R A E E M E E L F R Q S E R I H G T Q T V G W Q A V G L H N C I E T D T Q L L S L A N I
RANDOM_99 G P E Q A M N E E I F P A K L L Q G A M V H D K L M N L P K S D V F H V L L G G D K K L T G Q K S V K N V A N I
RANDOM_100 V L Q V D N L R A I S P Y S S Q A E I L A Q A W Y K N G P E D F D G F N Y N G V L I L I Q N T R P A V F K C A R
RANDOM_101 C G F K G G V T G P T N T K P H P R G V K A F K A A R A Y F Q R Y M G C A G V K N G I S T V D A I P D M I R W T
RANDOM_102 G S S V F V P K N M K V L N P Q S K A D W I N S H N S G R K T L Q L R G M N R L L G F M L T A V V H E I G R A E
RANDOM_103 D L S A C W E F K H D Y R D A F Y P W I V D L N D K D Q V G M L L L F T P L V E F D G N C E S S N D C T W E I M
RANDOM_104 D R H K Y G D V H F A I A S N G C S N C A V F R G A A P D L P F V S Q L A A T R Y S A S G G D Y V L E M D Q C Q
RANDOM_105 R R L N Q H G R H Q A K N R A A V C V G A R G K N N D T V I E Y P T A K E G T G I L K R W D Y S Y G P G L R G L
RANDOM_106 S T P N P G D I I L Q V V N D G G P D Q L T N T L S G E M V L E N N V V Q G G E G V E D A E K I D K Y G K S F F
RANDOM_107 Q V S S V G L D L T Y I I G E E L S V G L I Q E P V G T V M D R A I L K E Y A G A T C D R L T E L D K Q G I R S
RANDOM_108 D H V P G V T S N K S T F R K I N G S R L G L N T G L Y A S M S N S V D A I D A E P A D V G T R P Q E A L E N E
RANDOM_109 L A L A R R S V S P I F L E P I T S L N L I A K Y A Q P T S A M R I L D V M K Y P N E A E L C R G K N T C Q Q T
RANDOM_110 C R S Q R D P S A S W G D I S V G R Y S W Q E V G P I E P S P G S T R S A K E D H Y S M G S E L A D A S A K P P
RANDOM_111 S I Q E T M T K Y E L P V V R L T P V N I H S S S R D K A K G R L L Q A F C N I W W V A R L D L C A A P S G K G
RANDOM_112 C S V E Q A D Q E Y M I E F V F S N G R L S K F E P R D M K S E V N K T F L R P P V L K G E G K E L G E P I G N
RANDOM_113 G G I G F I F K K P A L H N K Y E K I P E G S S Q Y E G C L G V A V S A E R K V A F A T T T E G Q N T F E I A P
RANDOM_114 I Q S N P D T L R Q L D E K Y L Y F H G I F K V P T V I A L Y F P Y A T S P L V L R P E D E L M C I I P V P V V
RANDOM_115 S S N W E L S M L G L M L T F V T E Q Y A D T L K E S K N E S R P N I L S K K E S R L T T L L E N V Y D T Y I H
RANDOM_116 P N G L L F L S L L L G S S P E L L L A A K M G N P A K Q A E H G K R R L P V G K I I V E D K G Q A D N N F N E
RANDOM_117 M E V A Q L V I S Q T R A G P L L F D K K I L E I T A Q F K C E G F C Q P K D V M T G I T T F I V N I K P P F F
RANDOM_118 E T P R K K P D L G I H R Q I R G V R L D R G F D G S I I D N N L D L S C G T E R S V F Y S N E S V V Q M S E E
RANDOM_119 A R R G P V P S C T L W D Q G H R N T R Q E N K N G S L K A R M L T E F M N S G F F G E N V F T L V G L P N G H
RANDOM_120 D C M N E I V R L S E L A R Y D L V I V G A N L T Y T G T N S A G Q F Y L V N H G H R P V V G P Q C D H M A N P
RANDOM_121 S W A L Q V D F P P T F Y Y I R L D N I L R S E N G L K T G L R I Y A P R V A A E N F K T S I T S K L S T F A F
RANDOM_122 Y N T V H V S F T R K H Q S Y I E A L D E P E V P K Y R R L M A R G K A E Q N P L S C S N T D Y W Y G N L D R I
RANDOM_123 A G S L F A V A Y V L M E T L L A D K I V D I D R Q Y T T G L R M E V S Q A K N R I D E A S R V S N F K M D K L
RANDOM_124 Y Y P L P W A G V A R A P C G K L P I Q E I E Q L L L S P G D Q V L T S I E V S F L P I A P V L M Y K T P Q D T
RANDOM_125 L V G N A K F T Q P N R S Y L D E V N D G I E A I I S M L M A N P H Q A V T I C G Y I E A S A I I S L Y G N E H
RANDOM_126 G F S E D P M P R E K K L H H A D E K D N R N V D N S L K L K P V A K Q R Y D N R K P K V L D C G M H M T P T P
RANDOM_127 T R A L V K N G M R E I P P V P E M P V R K Q A M H F R V M D I T C F D R A Y V R V Y S A M M E D Q T A N S V T
RANDOM_128 V Q P F G L N E N A L T G E D P A V S E E P D I R G L E S S P E Q V R L N N V W Y L K M T R V A S D L N Q S H R
RANDOM_129 A E K T M V V G D K G D G W I T K T T K C T K Q N W F P L F Y E C A F I S L N T G D D L P A S D W D L A V D P Y
RANDOM_130 G S Q V D A V T C V K G A S A R V S D L K P G F N V T N V K I E E R L D P V P A V D E N R H I D Y H V D P D Y G
RANDOM_131 R G K N N L E K V S V K N F A K G T F L L F E R S T V L L G P D V P L D K G K L E P E T G L I S K S K D S K R V
RANDOM_132 D W G A S L A R E R D G S V Y L S I S T G S P S K G K S A L R N Q A I M I M T E V S I P R A Y G T G N T L C F D
RANDOM_133 P I V T Q A E A F T D T S L N N L S A Q W L Q A G L L N P E Y P V D K G K K A K A E S V R S G A L R A R Y C G G
RANDOM_134 D T Q G G I K M G P M P D V L P R T D G A R K S P T S F A D D P S S T W R S T I L A P A V L L K V N D C V R G T
RANDOM_135 P N G D P V P A E D D E E I D E D K A V T V I W F Y L Q D T L H A I S C R G G G Y G H I A A H Q I V K P F K V N
RANDOM_136 V I E A C P V E A Q G D K R T W D R I L N L R A V A L A H I D L L C L F F P E L I S F D E P I P H Q C F G S P I
RANDOM_137 K T S D F E A T R T F T L Q T I F K R E D N P E S E G P D G T E R K L Q F G V L T I F R H N G G D T I V E L G E
RANDOM_138 E P V S T W S F Q L L A K G E G I H S V M A L K F V G K D N E S Y E V N G S Y S T F R S W N L K V L A Q I A L P
RANDOM_139 E W A A Q P P F L D Y L Q P R Y V G L N N S P V Q R S Y S M Y A I S D E L E S S L C E A R I G N L T D P A G I Q
RANDOM_140 R L A Y K S V K T K G G N L K K I E D A D S N G V Y K A L W L A E Y I E S K N Y I A Q A T N G L D R G I A S V L
RANDOM_141 H H G L Y S P K G E L H L R C Q K I V G A N S N I E H H N A S M M V S S G N I S M I L L I F N V M D E T G S P F
RANDOM_142 V R M I R S F A G A N A Q I A N V T G S V K S A Y Y M V V G T R L A D T G S A L L E I R F V V F V V G K G I K T
RANDOM_143 V C S E E T L Q E S D E E T F I L I I A Y A T R W R L Y H E G E T V G T F F L C R Q T E D L T Y M E N E P E M I
RANDOM_144 I P G L D K M S F P E G L T S P N A S W F S G C N H R K S R R G S L F P T A P N A L L E Q V S R F F D Q C V F I
RANDOM_145 T I I R A S S F L Q F A N L D T I W V E V R R S E P A E Y T H I P A V E N N F D S Q L G R A E L K A V P Q V L W
RANDOM_146 K E T Q C H V D H A D C K C T A V W H I F V N L D S G E R A F A S Q V E L E D G V S F K G S G G A R S L R V G T
RANDOM_147 S G L S H N V Q S Q V A R L A D A Q K A F K Q A G A S D V L P G Y L R D P P A P F G L T Y N G Q P L V Y G C P P
RANDOM_148 M S I Q T G N T R H N D A Q P Q F Q F G L K S P S K K M R N C G V K I E N G V R G A G E D R Q Y M L P K A A R E
RANDOM_149 Q T P L T A T K D K S G D F G S P I G E S T Y C L M C I G L L R Y A I S K P R E G C R L D T D I A A K S L N D L
RANDOM_150 I E I S T C Q G S I K K G A R V F A N Y A I F N A E V V E H R N A I K Y S L V S Y L A T K I T E F G W L Y I S F
RANDOM_151 A V M C E K N M V S A D G M Y L T K S V L G N L V T I E G A L A D A L K A K V T D L P L E F L R Q G E C F D E F
RANDOM_152 P G M L Y T F V N M A S E G A K P A Y Q T Q F G C M R K C R A V L I S V V K R E A R S Q I P M G W H N D A T N M
RANDOM_153 T G M G S Y Q F D L S D G I A I E E E V A Y I L Y A K D I G G P A A R G V Q Q R R S F G S L L F I V A G P P R I
RANDOM_154 K V E V Q T I W E G D D R H A A V W T K S E V S S H P G S K L A G A T R E G G N I P K A A Y P Y D S Q N G R S Q
RANDOM_155 E V T I G L R Q G K P Y G R G K N T G F T F R G T K G L G H L L I P Y L P K Q D V V V K N L R P V Y M Q G F G C
RANDOM_156 D Q Q F Q G L K S S A L D P I R L Q C K L H S Q L P V V Y R H K L V P T L T T E D R C K Q Q S G S L L A V S T E
RANDOM_157 G F V R D R T L L H T A G R T L K F E R H W Q T S M A N F K L K G D I Q A F I N S L L K G K K L E I N N C E P K
RANDOM_158 V V T G R Q Q A S I A A R I T C I D T E L F S S S I D E P A G N G D L E N S I S D N L R Q L V W S K Q A P D R L
RANDOM_159 T R Y D P Q P G M G I S A N L G K N T L A N Q P M L R F G V K V G G R L L N L L F L T N N S R K K M S E S V P A
RANDOM_160 Q E P L F Q V S D C S P E E L S L M G T K D M Q L C A K I S K K D Q G K K A H Y E G S D A E A A I L L E L A V S
RANDOM_161 H I S K A A T V R V K S Q D E I V E G E M Y K G S A R T I S D Q G F L A L A H I E W P F L T G G K A T S H L E V
RANDOM_162 L F A G S G Y N T N S S G N R A E D F N V I R Q E P K L G E I V A I W L D I H F A C N I P I S F D T K L L F I D
RANDOM_163 V S S M A V E R L K E V T D L S E Q E G A Y V D K C Y V F V D N D D N P E D E P L K G A S S P F Y S V P H T V E
RANDOM_164 L A S P N A S C N L A M Q I E Q E A T N V N E E G H Y G E K S N L I N K C T S S G F K L V I A L E T F C M S I F
RANDOM_165 K L S L H F V N S E D K E L K M D V A N S R R P Q S Y H Y L E F E N K E L R A I V A M M I T V S P P V R S T D Q
RANDOM_166 V L G S K F P V A L H G Q I I F I V K A I K D F G G E E A S G L K I V T Q N Y L E H Q S V I P S Q D A T D V E C
RANDOM_167 S I A V L Y L A D C F D I P V P G V S I K A R Y A S C S M T R S S F Y V S M R G Y I A N Y I E C K G W F T E Q P
RANDOM_168 F R G A A I F L G P Y T V D C A T S E A Q N D S A Q A E A P S Y A G R V G A S E P F A V K G R I C I Q S W I N S
RANDOM_169 Q T S T H E D P E F E A E T I I D Q I I M M T Q Q E S A V G Q S I L I K I R T G D I E I L E L D P N G G G Q T V
RANDOM_170 P D E V T W S Q V R A D S C Y V S V F N H H Y N K D N S K E G A L A F K K A N H V V M Y T G Q C K P S Q E H K G
RANDOM_171 G P P V R G K A R K G E P S P R K L M R E K N V T P K P K P A L N T S H P A M K L K P G N T T P I C G T V G R V
RANDOM_172 C F V N S K C T P T L T P G V N R E Q T N S I L L L E V P K V T K R T H L E Y R Q V P K S G R G D P I P D E Q L
RANDOM_173 A Q L S T E E M D A A L I H L I I F A L N L P I Y R G Q G I R I R A E V F K K R D I K S C A P P A E G I F A S Q
RANDOM_174 D C S F G Q D D V K E A G T E V L L Q R I G L H E N G K Q D M S A Y V Y L D L P K S D C I V A L L L P V H V L Y
RANDOM_175 T Y D K P E I Q S A P V P A L L I V Y D A R Q G V R C Q K E R S E I A G N V A V L V G V A G G T P T A T I A H H

RANDOM_176 K S C E L K L G K P A G G V Y C W Y A S S E D L G S G A I P G L A A S P F S H K R I F K M D V G F A E E S L H N
RANDOM_177 A G L D R V N L G L A P P A T A G R F E V L A L V E R A A Q S I L R L Q I A S T E P A N S N F Y T H P G P F N L
RANDOM_178 F D K M I G A P L L D R T A G E V G G A E E S P D F T R P V K T V M D L D Y A E G R A D C F L L N D P L D E K V
RANDOM_179 L P G D E D Q N I V A P E A P A A P K Q A R I N M E N E P R I N G L E G N E I A T G N V T H K I T L T Q V T N Q
RANDOM_180 Q E I L M L H W T D E S V V Y S L L G V I E T G T R V T R R A L G I Y L D N E I K G V V F E V G N S S V G L Y L
RANDOM_181 L K P S N H R P L Q Q G G R I S R N W T L I F A L L L L T S P G A L F A L K N L A V T Q K I G V C A A G H T A T
RANDOM_182 N F G L L E G G G W S F T L S F E A Q L V E L D E C Y L V K S P A G P T E P R A E V L M A V S I T V K T T C K A
RANDOM_183 N K P P T Y Q S F A T L A E L Y H F K L L Y G L H A W Y N I T K G S L N A S E N Y D G Q P N V T F F P G P T C R
RANDOM_184 G D A A R S S E Y L V V Q Q P F Q G I D K M F I S S E R P L A R S D V R V D G L A E I D R L L N D I K C G T I L
RANDOM_185 L G D G S L I A K T L P P T K E D S N D T E R G S P G D E H F Q E T L A A F Y K E S V S K I E E E P A G V R P T
RANDOM_186 E R K S F G Q S L S H D L P F K L C L G S E E G K I K I G E S P K T D S A M P M P L K N N L R Q Y K I T W T F T
RANDOM_187 S G T T N Q C C E H I E S E S G N Q I E A R T R A L G A H S P K E G L S R V T G V D M T P P R K T H K D I L P T
RANDOM_188 V F M Q A C L W S D S V K S F F Y P H F A E I A A G F Q I A L L L M D Y V K P E I N N A D T K H P H T H V V G Q
RANDOM_189 K G P E D T N I T N R A A T G Y R N N T L I Y K L K L T Q H G Q E L H D T E A I A G P L C A I L F E V K A Q G L
RANDOM_190 L N K E V N P D V R H P K N I S K I P V G R E Q L E A S Q P A E W L I D L P R R T A T E A D Q K P L K L F R Y I
RANDOM_191 G L E S G V C S V V L C I A Q I A K T R L V G R R I F S N G I P K E H L S V L F I A A L K S N A N L F A L Q A D
RANDOM_192 K C L P S N Q G F T A L E P H A D Q R T L A E R A R M C D G A T Q K G L A S R S Y K N A E A V L A K G T N E N V
RANDOM_193 L E E A L Y I E F V N L N E N C R T R K D F G V K R F D R E V S I C K D G T Q K R P T E G Y A L K W L G I L E G
RANDOM_194 Q Q G R I V R S L A Q E L Q S K E R R Q P S M N R G L R K S G R N H T T A D S P S T D H M Q M Y R H E G S C E H
RANDOM_195 P D P I S S F D V P P D S A T R P V V G Q Y Q W S F F M M I D Q F N E V N M A L R K L P G F S N G T N V S A D G
RANDOM_196 Q S M S I I Y G A P V G D P Y G F G K E Y R S S R T L L L K T P P L A Q T W T S F A Q G A T N R S N V G R N L A
RANDOM_197 K V L L L E N P G V Q A E T L Y H G N G L T D D V I K N K E G Y S L V P K S F S E H P E A L D A S L R A S Y S D
RANDOM_198 K A D E W P N Y W L E N S H H K G S E K P P S L K Y A N Q I Y G A P E E P T F S L A S Q F Q G L P P H I V W L M
RANDOM_199 M S H G W I S T A Q T T D D E E G A Q I A A D T E Q A A Y F L G D L V I N T S V R S I A R K R S R S S S K E Y D
RANDOM_200 L H V R A S Q R P L A K C V T A G Q E E R V V L K G D A E G V L K G K E Q K D I Q A A T R A V G S K I Q F T R D
RANDOM_201 D D D V D P Y E N G F T R C A T D L W L R G T G P N W V N Q E R Y N A Q G M V D R L C I L T A Q H H P A S A K N
RANDOM_202 F E S S H M N G G I G L G P S G I P M S I W Y H H I R Y Y Y Y V K H E P P L R G I V A L I I S V V T T S Y W G L
RANDOM_203 T N D I E A Q N I L E L G H E I E G E A A Q A D D F S K A E H D H K P A I K D C P R P Q L Q E L N A G V I H H K
RANDOM_204 Q I I R I I S A R G L Q P G I I T A P P L K V K T R T A I M R P E V N K N M F V S R P P L R V K T E A D R R I M
RANDOM_205 N S L Y R L E N K D R R G K C G C L Q S R N R R L L Y S R V A M K F Q D K N F G E Y Y C E Y M L L V Q A D L H T
RANDOM_206 P I P M L S S W F N F L E G E R Q S H Q K A L L T K P S K L P Q I K N D L E E P K S C S G V G L N E D H G P S I
RANDOM_207 D E L G E H I K A T K G L Q Y G E G A T S A S A A Y S E L V F P Y Y H Q L E A V D P V N T N Y R D L F A P R G G
RANDOM_208 V S V S G A E G L K C L P R E A V R K G V L N G T D L T L K P S V A A A V V Q K L K Q N G I C R S C I L Y S Q V
RANDOM_209 A M A S I F G A I G V M T E K I R L I R D D C N A Q G C R C S Q W G L A Y G I C I V E T G D E S L R N L P S I P
RANDOM_210 N A S R N P E R V V G I Q P K K E E H A L S M V L G G C E Y G S L L D D C A F V I P E D I Y F A R E T Q A I Y R
RANDOM_211 F E Q Y A N G W I I D A E G F G D D K K E L N I N H P V F L L K P G A A A S P M E A R G P T G I R L H D L E R D
RANDOM_212 R L L R T R L F E M D S K S Y N D E T D P Q F R A M D A M A E L T E T L N Q P P I P P A R D F S E L S A E R S L
RANDOM_213 S Y L D S A M P W T D A G N T V H H N I T P Q V F T D A M S G D L A D S V V K R K L L E R L K N S E R I D D Y R
RANDOM_214 H R F G N K Y M R K M G S R E S L T L K V L R Y L S N Q G I G Q Y A E K L P G G R E F A D T Q E G A K T G K I R
RANDOM_215 E I N L G T R L C D A L F L A G E K T S E N G Q Q R T F E E K N L E D L K D R N Q R T E F V S G I H R Q T V F R
RANDOM_216 P G N A K I V E Q Y Y A K Y L C V E L G E K G R S E Q T T C T Q H W A A P L K H I Y G M H L G E P T S W P M D Q
RANDOM_217 N D D T I G G L T S L K A W A S F A Y A T R P V I L L S S G A A F D L A E L S M M E L L S D H K S K N V N E G Y
RANDOM_218 A D P R V V V I N V E T I N H S L D A V F V R P E K E Q D G E D T V G A A T E F Y A A D A T D A V D L F A L L K
RANDOM_219 K L G A S G N Y K L N Q I G G D L R L I T F P S E Y V E R G E D I G Q S Q I G N D Q P K I D D F H E G F A E S D
RANDOM_220 N E L G Y T N V G S N S P Q P F A D L T W E G L R L H H S Q Q K A Q V K V P I K V E V S L M I Q R L F P I N A L
RANDOM_221 A F L I G T R G T L T Q H Y D P A D S Y Y V T R A Q N Y I L M A S K K L L A H A L L A S R F H W M N E V Q A N T
RANDOM_222 S F P D P T N P F H H Q I T P P S S V A Q R R I F R T H A G P C T K A L G G V M E S F S P S T S G A K K D R N I
RANDOM_223 T G E N H T N L K N V L V N S L S T V D H D E G F R N T I F L V Q P K M E P I D K L M L G L V K N W L V S N N E
RANDOM_224 K I K H S D Q E P I K P G D E N I V K D Q V I H V V E N R Q M I P D I D L L I S W K L Y Q Y K T R L T W I S Q L
RANDOM_225 F A A L I I N L A H H E H G E P A T L H R I D V E V D R T A T R A M E E L K G N E A C L L G G D G S F I I T F S
RANDOM_226 M L T L S S V F Q A A L T N Q F G L G L L T G A G G F V E E M A K L T E V P E R A A N I R Y S G A E Q L A T S H
RANDOM_227 Y L L F W A D V G M K E H S I G C Q M Y I D T D Q P L V P T T K A K P T T P R Q R I S P F T E Q P Q L V V T G D
RANDOM_228 K Q I D P C I F R N Q G D G A L A F R S G G T R V A H V K H L Q A L A I L M I E E T S T H L S S W E Y Y D G C N
RANDOM_229 E A I E Y A V H A Q E N Q D S D D M G I E T V E I E A P I I L A V V Q S G P V T G L G K N L R K A L E T V I D P
RANDOM_230 D Q G F T L T V I G K K S M H C L G P P P E L R G G W R T Y K R K T P G I P G S A K Y L G H A K D P V K K A G R
RANDOM_231 S V L I G N A Y I S C W E R V F L G S L Y A I Y E E L L W F P G D N V D S V S L I L Q V Q Q E L L M V V E K C P
RANDOM_232 V G L R R L R D E L F K Q D I R A V Q D S V I L L S A I D V H Q S T E R Y F T E A N A F G P L I C S F Q D H K P
RANDOM_233 F Q K L N S M N V N V A Y D M T P Q K M M A D M Y T D A I W D I I R S F E A S N G N D R G V L Q V G V P Y A Q T
RANDOM_234 I A L S E V E S E S V L T M P A S F P V T A R W L I R T V G R K A D E A S F E N S V V L P P L R A S D A G T V A
RANDOM_235 G Q Q A Q A F I G A L D Q L A A T T V N W S E V P S S S T D A V H V C K I R A S C G Y M I K E L D T S S H V Y I
RANDOM_236 P S N I M Q S E G T H Y I W P L D I E C N A E S K T M N T S E K E K W C A C P V V S G T T N E R P L V R V A L V
RANDOM_237 T G I S N E W Q S S C D S R L A A P E D P E L K Y V V G E D P P I G S V T G E K P H T A G N T E D I M Q S S L T
RANDOM_238 S N N D V D Q A H I Y C N L P T G Q Q F P R F Q Q S L D M N R S R S N E F S I V A S F K E A K W S W I R P A K T
RANDOM_239 E P R S S G K Q T A D N W G H R V K R C L S G R K G L F G S D Y V I V C R C A A A D D R G L I D I I G I R N I S
RANDOM_240 I I K I V V T S D G L I R I R C A Q E I I I S F M G V K A F A V A P K W E Y D D G C F H I D A L G G F P L S D K
RANDOM_241 P A P N Y A P L P S Y N G G L H E N V L N G L N E S A T G L S H P N Y H T P G K G E A D V G W Y D I T D E N A V
RANDOM_242 I G C D K V V T G G T I V T V H S G S A I F G A K G M G D A L S G L E N E A V K I P K C T E L P L D G N G C Q L
RANDOM_243 F S D C D V T K G L Y T E R I P R S Y W Q G D L L N L W I M R D Q V D K Y A P R G F Q Q Q F V F L Q S G T R F A
RANDOM_244 K A R G V D E V E G G N R R T P Q K F Y L T V K D M W E F E A Y P V T L D I A P N L V G P E L A K A K H W S R E
RANDOM_245 V K K L G K E I V A R G I C A P C Q D I M E Y T A K H P L V L K T R D E S I A S E A T C T A F D A Q E S Y L L R
RANDOM_246 K G V F K Q F T V P L A I L G A N H D V K Y E I Q E F G V V V H T K L A G G G A P A V K I S P P N W K V F C K H
RANDOM_247 P V R T C K N D D L K M A A C R G S E R N I E D D E G G Y Y Q A N S E I L Y E N R Y S G P F V H I S L S T G N P
RANDOM_248 E D C E S L P F T I P A M T R K P Y L T S G N A Y S S Q P E G G Q G I G G N F T H A R S E F E L L D R N N P L V
RANDOM_249 T V T T S Y R L A V P K K S L Y V T I L Y D A E K Y R Y P N C G R L Q L L S G T W P K R I L L E M N V Y R F L G
RANDOM_250 F K S E A G G Y K L S S L P M N Q R Q N I P P D T H L Y L K K W S S Q I R D P S Q G R N S Y Y E Y Y P L R S V R
RANDOM_251 E G I A N K F A S F V S A K T C A Y D D N F R F P F A D M D Q M E A T E K V P T V G G Y N V N I Q Y S A K R D G
RANDOM_252 N K L E N S S R K I L P N I Y Y S R D K G R Y K A T L D E A W L Y I N Q G E K R A V V A L F T N L G A V V T D P
RANDOM_253 I R S A G D A R A S T V F T V K Y A M V G G H V G D L G Q N A T S L E F V V A E S F T D D S L E K G Q T Q Q D E
RANDOM_254 I D M F H V Y I E V G R V D R A E E E A S Q D I P A S A T L F G N D V G V L K A A V F D K K E V G T R V N V L P
RANDOM_255 F V H S S E G I K I Q D S S P V G G T D E K K K L T H F A I E V Y L C G E F Y I Q F G R D T I Y A A D M Q Q V G
RANDOM_256 T E R P T G F N G D R G S F E S F Y L G A N N H R L A G T L L Q S K K C D C I G E W P I R V D P K H V D A P E N
RANDOM_257 K L T F I D T A V K V L G R P V A R G F S F N K A N I I F T L A E S Q L A F D S A G S A N P H N F G V T K C H V
RANDOM_258 Q P S G L K E L Q E D R E N R P P S A K S Y E R N I W E E R K D G F L T L N L K A E N A D P L V Y N D M K P G Q
RANDOM_259 Q G L F N N D R Y N P E D T K Q P D G R M S T L Q I C S T M R V W K A T S P G T I T Q E D P I G T R G S V F T I
RANDOM_260 S L K A S Y E Y K F Y E S E N R Y S V L Q S L Q K P P E A V D E H N F D L L P A H T Q N L V S G S T S L A H S E
RANDOM_261 K N A G L K K Y G R L P V L V T D Y K S D F R P V S K G D E Y K L E R G V A H G Q W Q S G D E L L R N M S I G F
RANDOM_262 D L H N M Q Q L I Q N P I K Q D D A L I Q V G G A F P Y Y T T Q G L R R L P A Q A R I T G P D Y I T A L R N A D
RANDOM_263 E W N Y H A G S V P S D T S L L L P E I V R T T S E A R Q E E W M G M Y S K I E P K N K S N N G L G N D Y H A T
RANDOM_264 N A S T R Q Q L N N S V N E T Q E V A G A W I Q E E R A G F R R L G T I L D G F A V F E R E L N N K G I Y G V N
RANDOM_265 R V R L L Q N L I S A R G F A S D Y Q P E L P S L D E D I V S V P P V R E F G D A N D S G Y D V I Q I P C G T Q
RANDOM_266 L G M S N Y R K P L P N D T R K T T G N L N S R N G S E V Q D A K E K G G T L A D V S S G R S K L V F Q F G F V
RANDOM_267 G I Q L F V L E P E K A T H A A Q E C L S R E D V D S G G I F G C G R F A T L E A D G L I G K E M L K V T T L F
RANDOM_268 L T T L V L A V Y N R C T W F D K L K A I M L D K P H L F G G N D F K Q A H R E R A G A Q G T I G H D R G N Q D
RANDOM_269 S G I A L Y R L G E K A S M Q N T R T W T A R N G V P T T K E M V Y T P N H E I R P D R T P M I Q S S M E R T E
RANDOM_270 F I T R F L P V Y E W K L D G A K L K L R K D A K G T V L R Y R L L R W I V N F Y L T R C S A E R K L L L I Y K

RANDOM_271 L L A V T F N D K I I F D D A D L V V A V Y K L S I A E F A R G D P T R T C A K K A V N G R H T E L G N L G K G
RANDOM_272 I G I L I L M A K H L T M R D A A I A N A T Q C N E R L V M V Y V S Q P S R K G A I R K A N S K R T I L P E G C
RANDOM_273 Q G T E S G L V E Y D L E P Y H G G I V K W N A R V A S N Q P D K T Q L L D K D V A L V P L S T A L F V A K V K
RANDOM_274 F I K T F L K F I R P N A F I K W L T D P A K N P E M K D L D S V S F L A D K R L E P Y L A K A R V D A T I A R
RANDOM_275 D M E N S Y L F H M M I G D A T G F V E Y N I K S T L S L D I M K W E K V F E V Q D R P N L I G H G E I T Q G G
RANDOM_276 D T A H D K H Q S H T L R L A M A V Q H A D N I G T Q K V T Q L N L Q Y A K D G S A A V E I P Q Q I V N I I E K
RANDOM_277 T R V P K A N S L A Q L R C V K D E N Q S S M V G E R Q L G G D F F S E L I S Q V Q T P F T E T E N K R M K P K
RANDOM_278 A P Q A E R E R G G N P S V V N A N K I S L E D R L V S I F S G R N G V K S I P L A S N A Y A L L V L I K R G S
RANDOM_279 D V S L E A P P V I L D G I A P K I I I V N K A A E K S Q G E M E Q G W E D N L L L A K R D L V S T G E S Q G T
RANDOM_280 N H A L D V E V F A P F L L K Y D A G D C E S Q E E K A H M Y P Q Q G I F L K L V K F P D T F A C L N C V E L Q
RANDOM_281 F Q N P G D P Q S E D L V D V V M A R N T L G E M M V E L H Y Y R E A D I G S I R G D D K M G E V I Q S D V C L
RANDOM_282 A C A A T V Y W T P A Y V T P A S K E L H A A D C G T L L N F W Y Q L G K Q G F L E G A H L E V A G S L E N W T
RANDOM_283 Q L A P F L A R P R V Q K R L P R T V R T P D P S I A D A Q Q P G E S E Y A K I R T V I H S F L N G S N A V V K
RANDOM_284 G E C L M F T M L Q V Y E V E A R D N F F N Q K V G Y D M T I Y Q T K S N C A S G K D V Q K S N A S K Y Y F T A
RANDOM_285 R L E S P A E M L P Y T K C T I H D F V P K S G A V K E P F C E Q G K E V E T M K E T E S R H S P A N M C R K V
RANDOM_286 R Y P V P G A R M V I G P D H H A L C P G A V D L S N G F E I E N R T P N K Y E Y V G P S S L E Y C Y R E K S I
RANDOM_287 I Y G E P P S R T V H I G G H E L E I Q G C L M I R L S Y S S E R A Q G S R R L K G K G P A T D A I P Y R P S D
RANDOM_288 G P I K N D V R A P S Q H L E A P D T T R M K V A D S A N T S M E Y N L S N K V G N E L M K M L K R V L T H V T
RANDOM_289 A L D K P E P K G A F Y G A Y Q P W G I S N V K G R A Q S L Y L L L Y S G T L C L K F D T M G N T R T T D V R G
RANDOM_290 S P H L R V L Q P N I Y F P W H K D I E V V S K N Y R D T A P W T V L T P A F Y K V D E I D P A W L S R R T L Y
RANDOM_291 Q L L I I I K K G W L G L R L D L I E I K E L E F M Q V D K A V V D N L F N Y F G Q E C K F R N N L A R R P T V
RANDOM_292 S Y L Y A K F G R E F I H D S S A F K H E K G L Q S K L K N A P L V W F R A I A G A R R P D P P H A P K A P K M
RANDOM_293 D W V A L G H G G A G R S Q Q L F G V M I S L D A I N L A H I V A L L P K Y D L A F R S T E E L F F I D Y H I R
RANDOM_294 Q V P I S H L A T T H S V R T R G A Y K L V E S A P A Y E K L T Y E I P S L E K Y A V V F N Q F D N P I I H D M
RANDOM_295 H L V E G Y T P G E G S N G Q E Q D P A N V I E V Q K W Q I Y T T E H E Y V R F F A S E A K K E G G W A E W M T
RANDOM_296 A H P V V D F L A N I C D V P A D Q W A G K Y R H N L A V P F E G S R P E Y A F P W A H Q G F E G D T V A D R I
RANDOM_297 A N E C L C G A Y S A F P I F V T S D D D P L C I I G G G H Q K E D R G E N N F I P L P R H G M E I I M L A H K
RANDOM_298 L N G N G I S A A T N D S T Y G K S S G S P A N K R V Q A G Q D S A F L S K W T P F R G G P D P A F I C R Q M P
RANDOM_299 L L H G G L I N E D L G I I T T E T L I F V G K K F M A R G G A K D R L G V K H K S R Q G R A R K I E I V V K L
RANDOM_300 I T M A I S N N A L S L K P K F C Q Y E L S K D T C S L G V D F V L H T Y L S Q E D P E G V G N T I R I Y T V T
RANDOM_301 G M R Q T E A N S F G D D G Q L M F R D W T P L F L I T L P I S W E L G Q P Y H K G L R H P Q L D N P L F Q I S
RANDOM_302 V S I H F F S A K A F T G E Q N S A I Y G G R R S C Q R L A S G T K G Q F P D V H G S F D T F L P L T K M C F T
RANDOM_303 N M D I T E E I L A G A D G Q T V I A D R C S V G E N T G F L L K K S K R I A W A E G T D T Y R H A R S Y A N S
RANDOM_304 F G A R Q S N N G D A Q V D T S G N R S V P F S P F N T V K A N E D A H G L L T F N A Y Y V R S L E A L A E S M
RANDOM_305 I R R N V L K E T E T A S E A P L F D H E Q R S L I V Q A D L G T V R E K A D G E P V V P E F I A L G G S E P E
RANDOM_306 P E L T V N H H P L V I E P P G G S S P K S Q A I M L F C F L E I A M F G Q R T D N P A Y W T P A N R S T V Y V
RANDOM_307 D S N H G I D K E V D L C A P Q E P V D N Y E T V D Q G G I F L Q I L I D N S K I P F Q I G P G A N E E G T C S
RANDOM_308 L S N I K P V E T S S N K C G F G I I W S T Q E L R N D G L R I Q L G Y T F S I L S S L S L A N F D L G S A A D
RANDOM_309 N K K G Q D I A N F I F V A G H N P D G A S I M T S K F M L C E K L A A S A P V L S M C Q Q P H G L H L V G L P
RANDOM_310 M K V I L V E L V D A S F P T N L F N V G G K C G D R S P H P R V D T D E Q P A Y I G R V G G T N D F A P G L G
RANDOM_311 K P E Y T L K N S T L S S Q T L T T G N I L D S V A G A L T R Y V R A N H R L S V A Q N L S F G D T S I D W V A
RANDOM_312 A E D G L A L E Y A Y I D I H I V V K A Q C T S A K T S P F V G F S N V G L R Y S A K A K L H S V K L F S E N N
RANDOM_313 A S E Y K D I S K T A L I W G E D G L K G S C D F S L N F F R Q L K V F H A M S I T E N S T D R P S W V G T M G
RANDOM_314 V R A V G E P D G G D T A I A G P S V L Q R D H N R G G T L E A L E C Q T G E L W A S V A L A S A V S N L P L E
RANDOM_315 K L G A T T K G S A H G N E R G F V A W E K E T G P V R K V V S L D S Y D K M R T R K V F Q A N G V Y K R R R L
RANDOM_316 T K R Y I Y P V E G T S A T Q G A G E A E F R G N F S L G Y Q M D N S P N N L P H L I A A Y G L S I A G R H L N
RANDOM_317 R P L Q L Q H E T V A R A H W N E Q A P T D G V A E V L F A Q W D D D G P N Q W D P D W Y E L Q I F E N S R Y E
RANDOM_318 S F Q V L T K A E L E I D G D Q Q H T I F L A P K D K V E N S L N S V L F P L V L D P T N I T F P D N S A P H V
RANDOM_319 P D S A N G L Y V C C K R V A T R A K L D T E P A T S L I S L C R P D W S T C V C L L Q S K M V K F G A K G F Q
RANDOM_320 V Y I G T T L C M E G A Y P E L S A Q C S V R S S Y K K T S C G Q C A E L L M H R N D G K S E D S C D M A G V D
RANDOM_321 K V L H V E M H G E K V A R L I N C S C L L F G L I K A L S Q Q H D N P L N H L G L A G T L E R H F G G L D T P
RANDOM_322 A T L V P G I R T E L D C V I F V L I S G D N N I N R S G T Q G R P Q S T G E I L D N R G A T I M L V G Y P Q L
RANDOM_323 N G E A N G T C N D S G M P P P Q M L R S I L P A G L A R D K S T V G F L K P I S C G L H P L G Q S S D C T F L
RANDOM_324 I L E R A T I K N V N K Q Q A L T W S G G K L M E H S M L N N A A D L L L G M Y K D L G K R K L D A L P E W S S
RANDOM_325 R S K W K V S N P A T P H T F M P A K V A D K K T T P D D Q T E T G E I D N H E L R R G T V A L A C G N Q E Y W
RANDOM_326 G Y P S T P T R L I R R L N I L Y I Q G K A M G L K S L P G M G S P R P L S A R T A P D A M G E D Y W C K Q I H
RANDOM_327 H T L H M Q K N Y T L V K P L I R P Q R D L V C Y Y I S W L I K G D N C K L P L D A D P S T T I D R S L S G A W
RANDOM_328 Q Q G G L I A F L E T C N L D L P D Y G A P P S E A H R A G K P H E I Y P A T L L D A Y N R I E Q L G V Q P C K
RANDOM_329 N Q T S A M Q N I D N K C L Y K L D S A Q N L S G Q S D E A N V L T Y S P D K G R P P Y G T N L G T S K F Y R T
RANDOM_330 R R V G M W S K K G I S S R L V N S L R S T G H R E Y L C F Y E H L T K I D Y E E Y K Q V N F K V T G Q S T N L
RANDOM_331 I D M C E P G S G H K I V R D R A E D E L D E R Q L L L T Q S W K I Q T K V G I M Q R G K L E R L F G K Q T F K
RANDOM_332 I S W G Y A S S L K Q V A N N Q L G E I L L A D E W G Q D A V V T V I Q I A S S C T L S T G V K T T F A S G I P
RANDOM_333 L V F G L R L L V V V G T G L C S A A V G K S N G L S T Y R A H E L T Q Q D G D P R S S S G G A F A T P L L A L
RANDOM_334 S E L P M D Y I F F S Y R S D W L S E V V K S P F P W N F D D I C R V Y Q I D R L I D Y P N T D T E Q T R I M K
RANDOM_335 H D R P P E A C T A G I E N N A F Q G L C P N R G T Q V P P K A C L R S Y L P E Q N V Y K G W L A F E I V K Q A
RANDOM_336 E Q T R V P Y I D L I T S T F W W A L Q A F I A A V L E L I P G E A N I V S G G W N I Q A F I A V L N K F L Q V
RANDOM_337 G G V A Q K G R C V G G E D A K V Q A M G M T S T M T L I F T S V T V A V L W A G D L G G V W L E P A R K H L V
RANDOM_338 L S A G R T T Q R L C G K C P E L L F E K P F P F R Q P Q L C S T A I Q E S P V F K L Q A K I F R G L F A Y P P
RANDOM_339 G D S L T G H N H P V N L T Q C C A D Y V E C Y V P G A N P G C R Q Q E L K M V L S C V H Q K T F C P G M K S D
RANDOM_340 G D F I Y D N S K G H L C A D V S D V F C D L T P N D W L S A G Q C D L P D A L M L S S S D S Q D N I R E G D I
RANDOM_341 V G E F E Y R S R C S T P A Q A V T G N L S K S S N R N L L P G L Q I N T P Y L K N R S L A Q P T V Q L L V Q S
RANDOM_342 P Q L Q H E T F P D K F S E G P G P S I C Q D P R P T A L G V D H D T K N G R A N P S F Q L E D E M A F I N I Q
RANDOM_343 T L D T P G Q S L G K G S L T L P W G S P D T Y E G R K Q S F D P L D P L K D K V A K A R L S C N V G N Q V C L
RANDOM_344 V S P G I W T A M P G L A A G A V L P A Q S G R L V D M S E S P Q Y D M V D H Y L G S K Q I M N P D H V A D R Y
RANDOM_345 N S N L Q T T V A L V T N W T K A R T P R N A P A K I A V Y C T I E C L G A N L V V E S P R L N P E E K S K G M
RANDOM_346 R I D D L V A Q G T Q G L T N D P K P S D K R G Q L N I I Y L P S L S D Q R T Q T F V E L L L S V I V D S E L S
RANDOM_347 A E G G R D S G Q F V D S G G G I C V V G H S Q K A I P H K D I V E L M V K A F Q S Y P N N S S D G T L A Q K D
RANDOM_348 L P S K F K L F S Q I T A P A F S T G M R V M W G C D T E T K W E G Q R D L L L Q A M Q L G A D W I S S G V S M
RANDOM_349 F S C N F E T G I G Q G E P R F L A N D V D K R V W C V E S E C G Q K L L E V F D V L A S N P L G N D S V Q P R
RANDOM_350 S A N D P R L Y S K V D Q G R Y S K C R E D F N A S E A N L Y S T Y V I L M F I R F Q P D Y E G S I Y S E Q A T
RANDOM_351 I I V M I P N A I Y V G L I T A E D T C L Q T S M P L S G K P A V K S F V D N A P K P G R V I N E S L Y A F T P
RANDOM_352 P I T G I P Y I N F G R A R W G K E R E D A R P K C D L G I P W L L N T Y D N K R E A G V C Q L K Q R Y F I F T
RANDOM_353 P Q N R G L F M Y L A L P H T P C H K D I Y N K I A T T W N V M L C K L R D S C A V N L F D S L Y Y V W K C A G
RANDOM_354 M D N E P Q W G M M K Q L G Y T L T G R A L I R E A N A E F A M A R F A A T T Y T Q E L I W K F I L Y P K V I F
RANDOM_355 T E R S S Y R H S P I P I E F S F S G S V V L T V L S A E D D P L L I G A T G Y F G T T D I L T P F K P E T D I
RANDOM_356 R S M L I N I G K V V G L T L G K S L L N P S S Y Q T N T S P M M F S D L K D E G N L L Q T K D F T A K L V I D
RANDOM_357 K T T D E A V P G M L T L P E P I L K R L C V S G L D A G A E G V A K G D P L R L V I A S E Y L M L T R A P V T
RANDOM_358 T L E D F I K I V D N C S V L I L M S I S L M S E R G R F K N F A K F K I K L N S N I F A Q C M E M E K Y G P D
RANDOM_359 S G F F D V T N P S H G N R I V L K Y N T E E V V T H K L K V F V F S Q E G K L Q R K S H V E C V K G N I E V W
RANDOM_360 K Y F K R A Y T G P K L Y K L A D G A Q T L A P L L D I E V S H E N P A P D G P A I S Y G L T T T V Y F K C L D
RANDOM_361 S T P N R V Q V W K E A G V V A I L G P Y A L N S S H T P T C I S A E D D K V L K H A G Q E N N K G T F G G S H
RANDOM_362 C P E L G Y A V S R A T H N G L R L S F A V S L F S C G R G N T A P H R K I G Y I E L D I Y S A Y S D K L Y E Q
RANDOM_363 G D Y G A R F D R T I F A C E E E Q G I W R T A V V E N L N V N C L M Q A R S E N G E L P G L K L R Q N L K H T
RANDOM_364 S R V I G L I T N S S V L H D A R N S L A E E T L V Q Q T D Y P G R A A T R C H K E S V I G S T S A R Y D F D G
RANDOM_365 K L V T K E S F Y A T K K V V Y S A A N I S S A S G L E Q M A T S Q S K Y L D D L R M D P E I D D S D D T K V K

RANDOM_366 R L S L A L K E D R G V I S G K R L L R E F T G A K V P E H S T L L E S C R E R R L V F K Q T D A G T N Q F D R
RANDOM_367 D Y C A P E G M T A K R L A P P K C A G C N E V L R V E T G S V Q R H A V A E V H K A P N G F I I G A L S A S A
RANDOM_368 M E Q P V E C L V V G G F K G L N M N N E K D F T R A A T D E A W L L D K I D L M D T I G T D R F V L F P V A E
RANDOM_369 Y H S N K V V P E I S R D Y L L L T H H L Q V K S T Q G R A T L K I S W L E R E N L H P A S T A S L T L V L E P
RANDOM_370 G L V I Q T L L V A Y Q S L T G L D A S P A R A R E A A M M G C C K I F G F I N E L E D C P Q R S G T G Q K E L
RANDOM_371 I Y P T G L T C F R R R K G R I K C V V I P D E I R A K R T R E Y N P L G I L T H F I Q G P K N S P L V S E S L
RANDOM_372 R L Q P P I Q A G A A W S S M G Q Q V A C H S A W L K T W S S Q Y F S L A S N V R V I F A V R R I C V Q S D S E
RANDOM_373 Q S V Q F S Y T E T Q C G S V R M A L V L T L L C L M W K L S Q T T E N P L E W E L E D F G K R G E V L A A E G
RANDOM_374 G E D I D G P A S I L K P M C I R V K Y A V Y S A L A D D R K G H A L T D T T R F I N G R S V D L V F L I K K Y
RANDOM_375 G V A H L N S D L L K T I F V Q A P G V Y N K L K G G T K R K R M G T E A I D I S I G V R A S A M F G V F D R Q
RANDOM_376 L A R P K F I Q L Y R Y E G H D D N T T H S T F A F G S F K L F G E S R A H N P G G E S Q K A P A K F N F N Y G
RANDOM_377 I L L K I L L P L A A E G D F C S I H I Y M L N E Y P P R T T A G N G K K A I S G V Q T P R N T G L L V P Q E E
RANDOM_378 R K D M R P E L Y V P H K V D E A P Y G I D S C K R K T K S P D T R V N G V D P W A M I G L Q Y K E E T K L A E
RANDOM_379 G H G R P Y S S H N S R T S D R S S K H F T P T Y K S T A L N Y A E E A K E A K S Y S R S K Y Y T C F I A E K R
RANDOM_380 D T P N S S S I L Y W G I Y F Y P F R K L L P T L L M S D F A H S D Q L K L P P G I F D D D K T C F D D R K A V
RANDOM_381 C E R N C D D Q A Q D L D D A P L F G L G H Q V Q N V E S V G R P K I Q D D R L A S P K R E L P L R C T R P V S
RANDOM_382 I S P G E G E W D I S K S R G D S S I F E A G G K P T H T N P A L V L R I R F K F T Y A S M L P D T G I D L A T
RANDOM_383 Q E D G A F D A R D E G V P L R H A S S G H L K W C L S L V R R E E N I T E G L L R A A S I W V M G K L K S M V
RANDOM_384 A I L E L E S C G A R I I E S N D D E L L S A D S N S A K A P A C G C V K G P T I M L W N L I F A D M V Q D G A
RANDOM_385 G S L A H N L F I L E N R A A G S P A Q V K P I T S P D Y R L T I S Q C V S K F N Q Y T A Y Y T A S Y I L G S N
RANDOM_386 G Q Y S T V S T G L A R G L V P E M T A V V R F R Q L S K I S T E N V G G M R E L A C D L A A T V T S M S A D G
RANDOM_387 D N L K Y R S E Y N W N G P I T H M A G A V M L H A Y D N G V L G T V G I E P G T S V F W S Y T T V F I I S D G
RANDOM_388 V G A G L K M T K N A S R T L A Y K F C G P E E T M G Q T R I A V G N E S L F Y E F P I A K A D Y S G V Q S T Y
RANDOM_389 S M P F E G V D V S Q D S L C T R Y H Q H M T A E S R G R R E I Q V S Y R K Q M L S N M E P K K E Q N K L F G I
RANDOM_390 E G I L G K F M A V G I D L P I F F T S F G A L G L G V S G V V V S K E S N F E L C R P L L P K Y A S M C V K E
RANDOM_391 D S W S I T D T V S Y H W P P A P G K I D L F I K G R P I G A Y I N K I P P P R G R D Y D F A R F R P A S E E S
RANDOM_392 E G M T L K F P Q R S V Q L W F E E K G P P V A F E P A Q V I D M I A L K E W A S K E A P G G Y M A A S G P K S
RANDOM_393 T S N I C I C G P N Q I S D G Y P W T P A E K L V A L L F N K L G K I H T R R S A T R G N Y I H G N P I K R T T
RANDOM_394 R Y T D L G V K A N L W G K A L E P A G N V P Q A K R M F P S V P G I D K K R P A A L A E S A H S F P L P S A I
RANDOM_395 D L I P A R M R A D F E L E D A S A P L G D T C G G N N I L I V L A L E R P I A R E I G S L D Q K L A W S E W S
RANDOM_396 H L V G S K T G S V F G I G W K P Q S I L Q L E H A I G A L S P I Y Q P L K G D R S N D P L K E L D G G M K A R
RANDOM_397 H Y V V Y M H W N P K G L Q Q N L I Y E A E F A A N R V R F I A Q Q N Q E K P Q P K K G T Q Y G S S R S E I G G
RANDOM_398 D I I S M V D L Q A G F W C Y A G D A T G K K V N K R S T E R F G V P S F G L S M V G R L N S G I L G H G K S T
RANDOM_399 D R L C S V E G G K S G G R K Q Y N G D Y T T H Y C I F K E I R E P S D L N V Q L A N M Q V N I P Q N P K W G K
RANDOM_400 N T P K R H V V F L V V A E E I V S Y I T K W P L V A R S I L M L L G S D A A A S K G A L T R F F H G R K P P N
RANDOM_401 E I A I I K R L K F L L R R G R L V M E A S P A V V Q A I G M S I G T Q T R R Y A F A Q R S L R Y L L T G M S E
RANDOM_402 K R A V M D T E K T K P I L N Q G I A K Q A G M V P V C I R T Q A E G V T W E G V Y L M S L A T L D I T N L E D
RANDOM_403 I I Q P A C Q P G K V G I D E L L V K Y N D R T G L F T K G K T L A Q Q A N N H G I G A G E V K A Q P K Q A P G
RANDOM_404 T F V G I L Q R E D R D A K A A G A R A A F I M V P V R Q F Q A T G T W F G R M Y Y T R L D G E N L L N L V R R
RANDOM_405 R E A T D G F S S L T L C M V L W F V Q L A E S P I D G P D F P V G A F V C M C G D N N V A V G T P Y P S Q F E
RANDOM_406 F L F V Q F L N K R S G S R Y Q A L S L E W K L N A T T D S R A S V L T M L V T P H D V T I G S K L A R K A S K
RANDOM_407 Q L P Y A K S Q I I L E T I Q A G T Q W L I L Q H F D P G A F V V K R K G A L Q R A N T S T G R C Q L A R K W L
RANDOM_408 G K Q E I L S L Q D Y T T G Q L E K T S V L K S L L E L Q A Q M D F V H D T R L R K F E H P P E G S M Q T I P E
RANDOM_409 K T L G M M Y E R L L A L T D T V T P P D A L G P N R E T R C T S K Q T M N A S R T F S L T P S D T H V N L R W
RANDOM_410 R N K G T G V F S P Q G I E N K G V T D G R G Q Q S S F R M A V T A F Q N T P L G K S A Y E R T L W V S T A D A
RANDOM_411 G P W K A H N A Y C E T K G T L Q Y R E N G S N T A V A W R V F I G L V N T S A L R C N E A F L S Y S I R A K S
RANDOM_412 S K Y Q L W R R E F L N Y E E Q L L E C V R A V K S I I Q N A Y K C Q F F L S R F D E Y T N A W N D T S Y A A T
RANDOM_413 T A P A E R R N W D M E S G I D A S D L L S V P T S L P L P G R S K A L Q P Q E R F G D I F T R C R L D W L K D
RANDOM_414 W L L K I W Q E R L G G A G P V G I D L P L D G F V G N M F I M E L F G Q L D S I Y N K A K V I F L F Q L P S E
RANDOM_415 A R V N A G E T V M S L P G D D N C E Q N R S R F G N Q D P L C V V S M V V Q A E F K Q Q D M L V F D T A P V L
RANDOM_416 L I L Y L P V L F I Q W T F L G L D D L P Q W V F G P T S F A K G Y Y A P E R R G P P Q E I G P A H K A G F G R
RANDOM_417 N N A S K I F A E V N N V A S T H T R R A K R P E D F R S L P D E K I L V V F A K H Q M G I A S Y Q Q G R I R L
RANDOM_418 S E A V P Q C F E D H D S Q K G D H L G A A R R T Y E V Q T A Y G T S Q P V P G D K I T A C V V V G L L Q A T N
RANDOM_419 L H V N G R S A G S Q F Y R K G A L S V T F T L E E G G N K Q P E S G E A G K M T E I G E K L L A T G D T D Y Y
RANDOM_420 A W Q Q I T I K S V N S G V D S V G R M L F G Q F A P R F E W F C L F D S W A L T S R T N R C E R L G N P T S I
RANDOM_421 Q A G V G A G T E N S G N L T T G N E N P M L V E I P K P L F P Q G G F Q Q T C T G E A D K L L P S A P F I T R
RANDOM_422 D Q V L D G E S V V K H L D L S E L V D A F G T M K A T Q S F E Q A S K V F T G K I Y E K I D L H D K E H R P R
RANDOM_423 S G P A R G Q M L R L I E E G D S V T A R P D Q A F D T W S Q L T N E Y G G T K G L P Q G G I Y P Y A Y L G V S
RANDOM_424 N K I G S G K V I H L F N S D I I F A V S K I N Q V N Y K T R K S Y F S H H A S A A F A H G S A P K L S Y E T A
RANDOM_425 M F Q E M V P P N I A Y L S V K I R Y Q P E A T T Q L A F L H A N K S L K A N A C A T L M D S A Q I A S L G L C
RANDOM_426 F Q I Q R N N F D A P S K V Y T N Y K S V L L R V E A Q G D C D Q I P Y E A S E A D A G V S N A T S F W V S R F
RANDOM_427 A A D Q A D D L C I L N T R D E L P V D P N K R N L V D A L R G D V C A N G T L L R T F T V D S L L S Y S E D P
RANDOM_428 G L D K A V K M K D P G D F P S I Q T A G R Q K H T F Y L R I R K L S N L F Q G S T R K T A Q W I G M E L K V G
RANDOM_429 S D N V A H G P L S E Y N F C D N Y P G S V M N T G F H L E G L H E Q A S P K T K P N V A M Q A G A S P L S P A
RANDOM_430 P Q V V K G V V N A S V L P A V L A P G T K Q Q K K F M W P L V G A W Y G K T T L T L K R S A T N I V V T A N A
RANDOM_431 W P V I V A A H S A D Q L L A I T D G G R W L S V S G V T S N S N G M M A L D K A L S H R L P L I K Y D L D L F
RANDOM_432 H K V P T L D E K V V A A E Q K S A F W V G E G T N T K V E V W F Q Q I T E G P I M E T N W G H N T D V Y L Q L
RANDOM_433 E S V F M N V V G G I K R M S E D P V A T F T I N K D Y H I M P T S Q V C Q S A A S K T E E A S V I P I E G K T
RANDOM_434 A E R C F A I V R K T F S T Q I E K F T G V T L D G E G L I F V E Q Q A N N C G G V F V T V S V A L Y C V T R A
RANDOM_435 P M L I Q Q R S S S R T K G L A A L Y V N P N V Q L S L A K A L D A Q L I V V A Y L R I E I A Y T P H C A L I R
RANDOM_436 K Q L V A A R P E P L G P S A S E Q C I I W R E D Y R E Y V R A V A K G T F S Q S K A A I P K E N W T G P E G L
RANDOM_437 V G K I V K K L I K P S G N K K A V G Y I V I N A E N P I T A A D S R E Y L P T I P A C W M P V V S I Y S E G P
RANDOM_438 W S W T S N W P N L K M R Q I K E C F I C C H A L S S A A P F H G P L A L G A D L V S S R L N A T V N Y G N V L
RANDOM_439 D E K V F Q N M K Q L T Y E A T C L K Y L N S S G F D K Q R F H A W Q T M R S F V T A E G T G E D Y Q V T G K Q
RANDOM_440 L V V L G E C V H G P G D R K G C R N Q T V F H I V N L V M E K H N E D L R L F G F P G V D Y Q V V Y Q E D D D
RANDOM_441 L V N N H L P L E H M G T Q M V N C L V E L A S C A G L Q A G A L K C R I L D C L P G Y G V R T P N E N I Q T N
RANDOM_442 I Q Q G A K T K E L A G D V H V E Q G R K T G L G C G L L W W M N N E P T R E F S E S K K E S Q K A I R D F S E
RANDOM_443 F K E T A L L A A L K N K L V T T L L L F N E D W M F T Y K D Y Y A V V S T K E D L P N T E V I F K S L L P P A
RANDOM_444 L G W Y Q Q I I F A A L L L T G M E R A Y K Q T K R P A K E S L R L L V C G W R F I P C A G K E N K S D F L F G
RANDOM_445 C I C P V M I R R G G K L L I K V I N G P S I G G M D S P I S V G I L K G C F L S G L S F F N F N V N R E E K P
RANDOM_446 N K K Y S L A R G I F M T A I R P N I D F Y A L L S D Y V R N V F P E W S R A W G A I N G M P T M R M E R I E T
RANDOM_447 S G P Q N P M L T T S Q W A Y C D D E A Q T R Q V A C L V A Y K N F V F E K I T T N S V E V E G P S K F G K N A
RANDOM_448 Q E K H D A G N V E H R T W H A S H S A I T R L S N C A G I A D S N F S P G N V F L F G A N A R K T I E I A E V
RANDOM_449 L A S A L H N M Y I S T K K K A I T K L I Y R I Q L L E S R E E Y T L D P L T L G V K Y A N G C N Q K G Q Q T M
RANDOM_450 G F M G C G N M P G A L K Y P P H A C S S S R L S F G D F T Y C S I P D N L T P S L E S N D L A R L K S G G E K
RANDOM_451 D R F R Q L E A K T E D D G P L E G P N H I S G G N D V A E L G N I A F G C M G S E Y E S E T N Y G R D N E K R
RANDOM_452 A V F L P I G A T S T F E Q F L M A A P K A Q A P V P K V S R E K R P P D I V N I A D L Y D I S A I E M P R R R
RANDOM_453 I Q M K V Y L I M F M G S Q Q M A T G N M V W P E T C R A V R K E S L S Q N E M P V S I Q Y D V S N L A T Q M E
RANDOM_454 H K G N T Q Y P A C G Y E A T P C Q N T F D G E V D A A E I V T D E L W L E N I D E A V K A N E F I K I I L V I
RANDOM_455 V T C G A H M S F S P I Q A L L N K T P K H R E N L I S D R N F L A C E T S R T K L L Q A L E H A M Q G G P G T
RANDOM_456 P L M Q L C I A G F H C H G Q N F I I E C I E E F D K M A M I M S P A M R Q R P P S P L C I P A F S V E T D L R
RANDOM_457 G L Q R N Q S R G G W E R P P D S I F V R T V N I D K P P T Q G E A S D A K E F L H I C K F D D T F E V A Q T P
RANDOM_458 S W T L L S S N A F L L L M L Y F L P P A I D P K V P I A T E K C D S L D Y A Y T D G M R Q D R L G R V I T L F
RANDOM_459 G F E S I Q L M V L G A S Y L V S Y C T V T L L S Y N S Y R V V Q S E I N K S R E A T S C G L S R V N P S C G A
RANDOM_460 E V K S K E N H I G Q T D V P K A Q K G G L L L Y A T L L L G L A K T G K K Y S P Y P A E T M L L S A N T Q K T

RANDOM_461 E I L L N L E E N V Y P A L T D Q E V L S L P T L A S V S E L N A Y A I P K L V D T Y L E P M S A S P K S V Y A
RANDOM_462 W E T H D V P C A T V H D S A N N G P I D Y V S I A K V L A P A G Q I A G L F S I T A P K R P F I Q A D A A R L
RANDOM_463 K C H D P L A V L V S I Y G C G F N T V K I V H D A L K T R E I G Q K S H G G V Y V F D V M V L L L E D Y F G H
RANDOM_464 I V I V G C G E N G G A A Q V N R S A D C P M Y K S G A K L R Y V Q G Q E N V N K C R Q E D L T L E L K D A E S
RANDOM_465 A G V K G S H V S S E M S Q R C A G G C P A N K N V A D S P P I A N N I P P H S L P N E G L V G E M A L T A R R
RANDOM_466 I Q G S L T V I C A S G C S A G W L S G E E A R I I Y V A Y L N E D N P P S V N P R P H L G S D D L E N S L R H
RANDOM_467 I T L L E G P T G L H L V E G I S Y T D K Q L I D Y S C V L K G E G F T H T Q S N L I Y K N R F L M K Q K R L F
RANDOM_468 S Q L R V A A R P S A G I K H W T A A F K V R I K R L G D I N M D P K L H N L R P K S R S N H D K S S Q T K D L
RANDOM_469 R R N D L A F Y N I G G W H T V D V F R L S R E K E G K E E S Q P L A M A W A T Y F P M T K I A W L P A P A G F
RANDOM_470 C A M I G A S I E G D P L E E S S R A L I P N S F V T G A Q N C N L H S M Q R W L D P G R N L F R R D S K P F I
RANDOM_471 L G T T D Y K S C N K S A V D L H D R D G G A E E F W G P L G F C R A Y Y R P E E Y P V N T H L Y V A S E F K F
RANDOM_472 F V A K Q R A K L L F S M I E M A S K K D D N I T K I I T S N G N R N G L V T R R R A D L N E Q S A V R R I P V
RANDOM_473 N P E L I D Y L I H L A V R E Y R D E E E K G T E A I K S D V V M I I V V I W E C D L K K H L R K D F D D Y E D
RANDOM_474 K G M G Q S K P S A S E S G G T F D V A S I N M T L I K E I T F A V M Q Y T R T G V T P R P F K D R H G S G G M
RANDOM_475 Q I K T R F N W R V H N L Y R A G R G T A P K G K T M G L E P L I I N S V Q I L D R D I G Q N D R I T Y I G A T
RANDOM_476 T T Q V Q C S S M H R A S D Q R Y W M P F C T K Q S I L C Y T L C N Y Y L G A G T R L D Q K T L L T A G S E P R
RANDOM_477 W L V L L V G C T T N F T E R T Q C R V L I M L A F V S S R V P T Y D F G V G E N E N P T K R L C M I K S P K E
RANDOM_478 I R Y G P A H L V E G P G C G G T L K N M L A S I T Y F S I S L E L L L I L M N F A P S P G N V I F R G V F A E
RANDOM_479 H L E V R I Q V N Q Y E R C N L N G I K A V G P L T S Y N L F Q R R R I G V R T F I P R L W L T L E S I D I P I
RANDOM_480 L S I T Q N P V G G L P E T T V S V E A S C A G E R S T V Y A N D E S S E S C A G A R L Q V S D G K R S D A I Q
RANDOM_481 L T G I P K R G L S H A W V E E L L E G L K S A Q T T M G I V I I T Q D D E L S I T V P S D L A R A L L V K K G
RANDOM_482 L R I T V T Y G T D S L R P P G A L A A M Q E Q E D V Y S N D F Q Y I A F A V Y S L S H T S A T I V W S T F K K
RANDOM_483 A H A L A E Q P R T A L N V G P E E T D S M R T S S W C V Y V G P E L I A D F K R G Y G N S S T E T R N A G R K
RANDOM_484 P E S A S V Q L R I I T K A V N M V Q V S E T D S F R V I T Q S I C I V Y L A I D Y P E Y V L K E F Q D Y Q K L
RANDOM_485 K G T S K P F F D L V V R I H R L G H E S Y R G P P T P T N R G G L I F Y Q S D G P A N D P P G G F Q P D K N D
RANDOM_486 T R E S H T H T L H N H L Y E L I L E H H Q G V P T S G E Q V Y K C C P I Q A I A R P G A A R Q N F S I V H K S
RANDOM_487 Q V K T N Q I W Y N Q P L L G F T R N D S H A A F H P V A Y P L V E K R I S C S L T I I V S L L V Q D D G T R E
RANDOM_488 E H A K W L R D E D R M G F H F N G T A V D F I E E E Q Q T N D P A Q A D D S E S F D D S I D F R Y P T M Q G L
RANDOM_489 V D T R G N Y Y T L A R F Q Q G P A A G I Q Q A R E A C S F G K V E Y K L D K P F S W S V R L L S A K G Y Y S V
RANDOM_490 E N S E I P V A I T N T L I F P V F Q E R F Q T R D T Q Q R L G D G L K G G V S E E T R S D P L S P Q A P Y I T
RANDOM_491 A C Y K R E A K D T S G G A A L P E A G E S W H V K R S D K S V L D N W D G E L D T E K E C S G K F V S P C V Q
RANDOM_492 G K V G G N Y K L V A D A K S A W D R T Q A A L S A S L E T P A S R E G L V H L M C V L S L M L E E V E R F E C
RANDOM_493 Y E L A Q R K G D C I S G S E L L C Y A P H A W L G N I Y P I N D R S G Y T L D M E L P A A R P D S Y P I K H V
RANDOM_494 R D G D N T T K S W M D F P Q Y V T I V T L T Y Y E H P D A K G Y I L S S L V E C G D L D F D F N E S E Q L S K
RANDOM_495 Q V A V L A T D L H I L G E G C G I P L G D P C Y A T I A E V E A S Q L R L Y A L A L S Y T K I L Y P Q Q T K G
RANDOM_496 Q E I C T P L S S Y I S H A L C C A R I Y L A V Y T R T F S S K D L G S T V A Q A H D G D K I R T K C R G M Y F
RANDOM_497 H V E C A K N K V A M N F G Y E V M L E Q I H Q G K V F I V A M S S F E S L T W L M I T L L I I I I S L N C W F
RANDOM_498 A Q I T H R A V L N D Q G S K W P T T W G I T H E A M N T M S H M C I S L T K I H P D S I I E Q G L L S R S G N
RANDOM_499 I Q E R W G T P K I L S K R I T S T A D L Q V R A D I S S E F P Y S V K E L M T A C Q G D R G L L S W A G E G D
RANDOM_500 S G Y V E K D L Y N E V E V D F E T D S H K S T A F L P N L R T K R C E E D R H A P E S V G L L G T I L E Y A F
RANDOM_501 G Q Q E K L V Y K G T S I A T G A K S P K N S L T Y V T P M A Q A T T V R L G V T A M L A R T T I M A N L L E R
RANDOM_502 D I Y G K Y P M I S S R F A K P A V A A G M G M H P C L V E Q V H A E S D T Q D P L V Q E R E H M K S L G F R F
RANDOM_503 R L E K C G S L G T G V V I R R S S S S R K Q S K G R I T D I L P L N I M M Y M A P M D G I K L E A W F G N P K
RANDOM_504 L N I G K D S A T C S A E G K Q P K L L G H A T P S M L K N F S R K L I K F V H A A V A L K G R K N S I K V P R
RANDOM_505 S C H E Q S S Q K D L A E K L K K V L K V T T V C E A L I Y R Q K L N T Y R P P R A S S R A Q Q E N G P A E I N
RANDOM_506 L K E V S L F R Y A V A I D R S A E S E A Y N K P T F A A Q W D V A G P V N E W I R F I S Y A D L P F N F I L Q
RANDOM_507 S V N P A V T E E L G V K N M K A N K H I M G T D H E S I D T A G E G S G S A I L N G L I T T V A C A M S S H H
RANDOM_508 V S T A C A T D L P Q A N V S H L V P Y T P Q P N L C R H Y R I N N K G Y V S Q M T Y G N L L G S I H D W I V A
RANDOM_509 R E A Q A K L P L M D L V A N S S F K M E D A G F Y A P E N T C P E L S I C D H D C E F I L L I D K A S A I P I
RANDOM_510 S S D Q A S A F W P G D R W Y C L K K G R N V D S K N Y L K K N L P I A L N K K V A H G G D I E I I S G N V F P
RANDOM_511 S G V S T D N R Q L G M G Y V I G D G D V G N G V F I I N S N E G P Y Q Y H G E I I D F P T T T A P G D P A D F
RANDOM_512 L N D Q I E Q C D V V G I L E Y F G A P A S H P G A D F A D S D V E K G G S F H H P K N N I P D V P I A H A L Q
RANDOM_513 M L R L H K S R G H L R I I L A E P I F Y L N L A G F R E S C Q K D N W G A K V K G V V F Y I G P S G F G D G P
RANDOM_514 D E Y R A S I G M M N S D D K G D M N K D C V A T K K C F C I R T L L L V T G F I S P P L Q I A K D G A A E A A
RANDOM_515 D P R N L A D K F D L S I S D R F G S A A S S Q W S F L P L A L N N D A F W Q H K S S E V E T H T L Y S I R P K
RANDOM_516 S R P L V N P K T I A S T D P F R W P A G L G M A V N C A S Y A G E G E R W F G A G A P S D T D A I Q T Q A S R
RANDOM_517 M F L D P D V S F V L D Y T T K V Y E I I S A E K G L Q L A I F T A K H E I V A E P L V S Y K R V R F C C F C S
RANDOM_518 L T S P L W R L G E Q T E P E V T N K L K W G D G S Y A E P L V A H G K R H D Y A F V I A A G I A T N Q S G Q S
RANDOM_519 S V K E L R F E P E A E V I M A N S F S A T S N T N M K I I S D G D E F L Y I F D D L T P C I G P K L E V H P S
RANDOM_520 R N D D G A Q S D L E T S R L E Y K Q R N A S K S C K L N P I T K K A A D F S S R P E M H L D I T L G A S I F M
RANDOM_521 S I T F V V K G P P L F I G T R Q N E V G S K C D G S A R D I D G C H W Y T M E R R N N L A R T G G A G N V E L
RANDOM_522 I M S L A Y T E L G S H I S L L I K D F S K T P V R S P M N K D W V G A K A L A L G G N D Y V F N K V G T D G A
RANDOM_523 D E L W N W G V V W P S I P W T R T R D I Q G T L E T V R K N H K P E P P I C K Q D L N P A N V N R H W H A A M
RANDOM_524 N I S G V V T L L K L S C F T R R T A R C L D E E G A R Y S I T S N D Q T D A T Q C S C P F S E C P K A E S I I
RANDOM_525 G L A E L N D A I S T S I G L L S G R A I A A Y S Q L F R A K K A L A V T C G I H E I I R S R I C L N G E Q E F
RANDOM_526 G A K E F T L G G A N K V H A E T K V A N R I A V Y I V E A I K A R F T P V K S I L K G T N K L D S A N A D I A
RANDOM_527 H W I R A G K K Q H A N W E A V K Y C V V P F C T G K K D D A V E E S D H R N S I N Q L G R P S L S S A F T R I
RANDOM_528 Q A E E L S A V R K A D F V R S T I P F I L D T L K D C V K F F M A H A V P K L V Y P K L R K V N A D L R G F I
RANDOM_529 W E Q G L V V A M S L K G C N L L L E L I N F P G Q D S G A G I G D V V A G E A L L D L H L E T A Y G R Q G I L
RANDOM_530 T K A K F Y K Y A T L R A T D P Q D N V P G L T K G E A H A E D W I K R Q S A A E I G I R I L L E G P Y V T T L
RANDOM_531 I N T Y P V D D N K F L E E Q I E E K G P K I V S I R H W E R R G L V D V K K T E V N S E K E A L R T F K G F S
RANDOM_532 H T D E I Y W W D G C F K G H P G V L R L E G P L G P R T N N A F E E K L E F G T T E H K R A K L L A P N V V G
RANDOM_533 F G L T E S S A L P A E Q A A D N E H N V L V S D L K V D G V L L A K T N T L A E H E Y I I T G C H E L S V N G
RANDOM_534 K A K H L K I I T E K A L F R L D G A I R A T F L P T G G E P D G P T Q S P K S G G P V T K T M E L G N L I R A
RANDOM_535 L F V G W K L Q G F N V M S E H G C A E I Q T A F A E R D Y Y E T A M D E Q L Y S E P E E G N C C K G A Q S E Q
RANDOM_536 Q A Q A Q L A K D S G K E G A G T I K Q D A W M N A P L L A A L K L Q G F P A A E A M F K I G Q V E V N A L N D
RANDOM_537 E Y F L I K A T M C N T Q K D N G D K R G P T A S S R I I I N C Q R A M F E E S S Y S E A F I F R V D Y V P R L D
RANDOM_538 G C A S V F E L G S I I M P P E S S K L V T E K D H E E L R I K R V D Y L G S G G T T L S G M G R A L P D E D A
RANDOM_539 P E S R S H A A A V G P V D Y E S I S Q K G S Q N N K F D S L R S E N S G Y P F A L T Q R I A T S N D A H A D I
RANDOM_540 G E S Y I D L A L V M H A N V T N T P H K L R V H T K Y K D V R F E E F Q A R G D L S E K C A P D W A A K D A G
RANDOM_541 K S L E F Q G K N Y R V F D Q P F A L N A G P G S A R P A A E G D Q G P C G C F F T L R M I D T Q R W R E K S D
RANDOM_542 I G A N L K T H N D S G S L K K E L G R A S S F Y I F V V V R P D I K A T G Y F E S R S S P F S N S S F Q S N A
RANDOM_543 V G S E A Q E K A P S S D S N Q V G L D Y F L T P I Q T E A K I A V G G A E E F W A A C A N G Q C I G L I P C M
RANDOM_544 A E D T W T C C M S H V W P G Q S I D N G F V H W N G T R S Q C K R L S I T V E S F G I S P I W A K A V Y D F M
RANDOM_545 Q T L L W G D A A T N N F E H G D N D G A Q M L L S E A V I V T C E Y P I F D L V T S M S L K K C Y H N V H C M
RANDOM_546 S T P S F Y W E V E G G F L L I L I K S S S I I K D G S N L N Q R G K E K F T D P F V A A A K K Y H D K D K P M
RANDOM_547 G L Q V S W D K M W S D I P K T C P R R K G T L N I E K K W S S D E S R G D K V Q V G Q D K S D K S D S R T S S
RANDOM_548 T V A S D K G L F R K P E I M H L A K L C K T A V Y G T S K A E T M H A I T N L G G T F L L M K E P F Q A T K G
RANDOM_549 Q Q I R H K G D G T T S S Q E A F G G L N S G G G M T V R P W L E D G Q D G K R V T V A Q A Y G G H L G N E N D
RANDOM_550 R N G T P H C K P L V N L N M L T K V E E D L T E P A P L E I S S R R S P L L R G S S F V I S M R P T T H I L V
RANDOM_551 C D G H S A F V D G L E K V G E K N S Y L S G R C V R C I Q S N Q G V N N S N M T M E Y S Q D Y I E G A G V E E
RANDOM_552 T A R R R I L N E R Y H N K G G T R G L M S G L K T D A D V N E W A E P L L A Q V R N E L A V E V K N V T Y F S
RANDOM_553 F L S P G T A G D S C G T A G V G T V R R K L R Q P S L M A T Q K E V M D A Q L I E R M S M S V G S L S N K S H
RANDOM_554 P E R K S A I M S E L A I C Y G P F I L H P L K S P F M G G F V I N E E F M L A R Q L A K Y A R K K G S L T N T
RANDOM_555 V I A V K L G I P N G V G K D E L Q Q L N K H L R K Y K Q E Q D E L V I G T Y R E F A G S T T Y F P Q K Y I D P

RANDOM_556 F I L S P A N Y S S D Y F T V C E R Q Q Q L A T V T L I H T R T D T H A Y K V L K Q Q N M M T D T Y Y C K C T Y
RANDOM_557 V L M S T W V E E L D F D P S Y A G L I G N P D Q A R V L V T A D V E T S G S T F S E S D Y Q Y L I N V K A K N
RANDOM_558 Q I P V G Q E P Q N Y A F S T T D N I K E W A Y A E L Y F S S T P P V S A P T T F L V V A D K L G L P I T G G R
RANDOM_559 P E Y A Y N K S D P S G L G V D A T H I P T S G A T E H H F C T G V N E G F L S A M H D R F L I E S D N W D S L
RANDOM_560 E S P P I L K A T T K F K G C S F H L E T F S F K G G G L S T I F Y G G I A Q F R L A K L R W F M K V E V I N N
RANDOM_561 S H W R Q W N H M F F A V L F A A L I G Q K R V E A I P K E K S S E R A V R F G L Y A G T E I A E D G K F D K C
RANDOM_562 S E Y L L D L I D K A E L G E T R M R P C V R N G H I R D P N G S L M I E M P A H D L V P I D F R Y A V L V E Y
RANDOM_563 F Q A V P W D E A R R G T I V G G K F V I K Y G P V D E N K L G S T W S Q L I I Y H L N K S A S Q N E S V G V V
RANDOM_564 Y N G L Q T G A Y V E W G P E E S W G P A V Y V E W N Y C L C G T N S L S E K F G G E Q I C D K V G I A L G G Q
RANDOM_565 T T T S A T K R L S E L D L T F T W A E I T Y D I N I M P L T D V A V D K E V L Q T S S H N D G K D D A V L T E
RANDOM_566 G D L I E L V N A P C D L Q I L E I A E A V E T D M N A K M E V F L G E N G D K S I L N T E D H E E S S Q E K V
RANDOM_567 A L V R L F E A Q M P Y L Q Y M L L P K P C F S D K Y H R V Y K G L T S K D G L R G L T N N K D S G I S W L A D
RANDOM_568 R R L A V G W Q E N D M S E A S I I D K V T L T F Y W D C Y R M Y H L A Y A C C D V M Y S R S V L M K Y P R L A
RANDOM_569 V P N N A G N E I E F F G I E A R R N G G A K K G L A K M K G E D A A N S Y N L Y I K T T I S F Q L K L I V G Y
RANDOM_570 T Q D P P N D R V R D A I G E P V T E C Q N F M D V I S S V K K E A A R D L T E V G T Y S T E L M C V K A I F P
RANDOM_571 N T S Y G L V K Q V P K S Y D L L D E S G Q S H Q P A I D Y V K R D V E D S D S N A G D E E D A L D I R D Y L P
RANDOM_572 S V M S S E L G I A H Y Q H A L A L P G G A E D L G N I K A G K R S L L V H S A I S G T E L E T M G P R Q G V C
RANDOM_573 A S R A E N I M E S A A R I M L L K T L F I F D W R P C D P S P E R F N P T N P E F R E A D G L R T S V E R W I
RANDOM_574 A G V K R L F P K I K Q Q I D F M N R E Q K R N Q V T G H P E K L S L E E K F S P Q P F P D S T Q K K T L A N H
RANDOM_575 A I L D L D E W S A E N M S G S F A S N V L N N R V A A T I I R G T M F N N T D G G A Y R L T V I N S N Y T K E
RANDOM_576 I Q S P L T Q E V S G P E L A A D S L L K S S Y A F N F P H Q K L F W A V V L A S T L E L S P S E T R I D S V E
RANDOM_577 Q E F G A C A K G F L Q T K I Y L I G V I D Y I V G A L N T S G E L T Q I P L S V D L C H D D D N C I V L H L G
RANDOM_578 N N Q D D E A R Y L L Y L M P L E Q G T C S K M F D V Y E G F N E T Y W V R Y C F F C Q P G N L A V Y N V H K I
RANDOM_579 M N S I S P D L S Y F L E R E C F A F P T R E D G G C T R M Q F L K K A S W G M S I N I T S L V N S S A R G M N
RANDOM_580 C L V I V V N I C I V A E L E F T V G V L L E N L E A Y T S S M S R V G A G N F N E M L L T V Q T K C M F I T K
RANDOM_581 Y I T S A I C V L M V E L G P P N S T R F P I C P S A A T K I P Q Y D A G M R R T F D D L F R R G Q L V T W A R
RANDOM_582 M G Y D A C K L H T A L Q D Q G R G V K M A F G D V E R L R T K K S D T T Y R A L I P T G R K Q S S G C S K V F
RANDOM_583 V P E Q K H V F T Q S L E E N D W N L S N V G D N E V I Y G Y G G N S E N V P F W D K A P H Q M E N Y A N A K V
RANDOM_584 I V G A I A G T G Q P K L V V F E P T E F V V Q W Q N R T M I H Q F S G K A S R V R D G M I C D T S E S L L L N
RANDOM_585 P T G A D T E Q D A P E T M W T I A A G A F S F I F Q L M S L K P V D T A I D H E A A E F S P R K A L P A V R P
RANDOM_586 E K M F Y R M G F S A A E R C L L G V K Q R C F F Q A I T G Q V P T V N E S I V H T A V A N G S Q R I D A L D R
RANDOM_587 D Q E T P E G V K P V S N S P W I W D E A K H K E D S W T Q C G V A N E R E L V V F L K A D V A G G Q A K S G S
RANDOM_588 E S N M D L F E D L F H F D F G M G V G G E L G D H R V N Y C E I I G T R S V L V D G G Y N G G G K G T H L V H
RANDOM_589 C K R A C L E R K D A N K R K I V R C H S S G L P S D R G L N S K E F A T E K K I I N A S V F P V Q V V A E A A
RANDOM_590 V N L R D D E K N R G K F S A L T N G Y E D N D D M I T I L G M L E T Q T M T G R Q K S E A S I E A C F V I L H
RANDOM_591 V I D V D F A M V P G Y Q L V F G K F N D A D T V N E T F F C V K G Q I A D A S L L L K W G M G R Q Q L Q V N Q
RANDOM_592 L S C A N Q V L V S E K P T I Y S T E V P F T G L S I S R T T I I A K A F A I G K E G Q G G L F I K D V K I G Q
RANDOM_593 H T Y Y K L N F Y N R G L D K L A L Y N C G L Q R S G K T S E V E D G D E M P L V R V V H T S L W T V T Q Y I K
RANDOM_594 G E G S G F G P P L D V G V K H V G P T L L S E L I R W W T H F N C V E I S E R T C E S C K P Q L K H I R L S I
RANDOM_595 I I E A V G Q G E E G E S L R Q V P E D L S F D V E G H S W P L D T T I S V F T Q N R A Q K M N L A V P A K P K
RANDOM_596 P T L G L T P E T S S H V V G E A K F R H G I T N P V D F R D P S M R N R N D S R L S G S W L G V I R G L A P W
RANDOM_597 A D G V M V L A W V L T L S Q D V L S R A E G W V L L L F I R K Y G T G G V C W N E L Q P T P E P E G Q A G I K
RANDOM_598 R P V G K Q A A H V V S A R C H R N R I A I A Q P L D P P V K F L K R L N A S G A S K T E T R P S K R Q C I W S
RANDOM_599 I T I A S D V I W R A V Y M V E H E Q G E T E M G Y S R K T F T E P T N L D L F S I S N H E N A A D R F V H S E
RANDOM_600 L L L G G L V Q G S R S A R S F W I D K L A N R R G N S L G P D M T I N L L M A S F G F A G G K T I E K F M Y E
RANDOM_601 D S A N H L C L L I I Q A V H N P T R S A Q Q K E F E F L W P A D D L R F A R S I M Q S P G L I L G L D K F L V
RANDOM_602 D E E R I C S D V R S I D I V N C V N D V K M Y L S A N P N Q F V R E E L L Q I D K S T E F G L F Y L A Y R S Q
RANDOM_603 V K E L V W Y M A F A G A S D G W H S F K Q A D R G I F P G K D H A L C A I G L L P L T T R E D A R K W L V T K
RANDOM_604 D E E Q L C Q V L R P K R L A A L D V E L R R N K A L L F K Y P Q S F D T F L Q T P V G I V R L K D F P F D L N
RANDOM_605 P E L G H Y R G Y R V D A R T K A E Y Y M Y A P A L K F S L T F L G L N V A P Y K S M A L H P N R R H S R F R E
RANDOM_606 N Q F L L T F A I P T L L A L Y T Y G Y Y I D T L Q E E G K A V Q K R D E K E P A A F G R V K N F V K S K L L G
RANDOM_607 G A K E R R V S Y A G D T A C R A D P L I G V D E E E L K F R D L P A L L L N S P P N P D Y P G Y I A N D T F Q
RANDOM_608 I T R P V L E V N S P T S H S M G E M Q P E V D A I S V Q R M L T F A V N P S E F E A R Q D F Y Y R S S A V K A
RANDOM_609 A Q L A M D P L G Q E K Q N S P S D I G V R A D L E P H A E G I K F L V K N L T F L E P P Q N M T E F M N I K C
RANDOM_610 P G G I L T A G G G R A P E K K G L W H K R G C P L I G T Q D S G Y L C V L Y S F I K P E A G A T H T E R C G I
RANDOM_611 R Q S P A P K L E N L N D Q P L A S E K S F G V N D S Q E A I V P T T T R E N L P G S P E A A L Y Y W C S G L T
RANDOM_612 T G V Q K V P G P L D L A C I L K K K K W Q T L T P L K L C Y L V S E F M R V Y M V Y N D H R V E K L R M T H K
RANDOM_613 G E Y T L E S A N T R F P V D H V N A E L L V F I P V N Q H E R A V Q N V E S L P Y Y I S D D N G G P K P R G L
RANDOM_614 D G R K E C E M G P S K T A S L L E C N R L G T I I D R N Q G E L R T K S D S S L W F L V Y G Y W W A K G T S L
RANDOM_615 I S Y H D G T G T Q Q T A I Q Y K V Y G G A K N T N D I Q P T V P F I L V F G N A H F K N F P C K R G E Y G G G
RANDOM_616 R I F V V D A N L H D V A L V N M R G K K F L S R Q I A A R L F P L T Y Y G T W W L E A I D K H H G V L L G R V
RANDOM_617 T L Y T D L K E T M W I I E T K D P P R D G W R T G R Y L S R T D S R N D I G P M I A P S G A L I A A S K F T I
RANDOM_618 T K R G P S N V I A V C P Q Q A W V T R K T T W I E L E N D S S E G N E K C G L L D F Y G M L G Y M Q P T Q G P
RANDOM_619 L H T M I D G G S K L R G A V S C D K I G D A Y A R L S T K E Q P P K D L A N K G V S H C E T V W A S A I E V R
RANDOM_620 P L Y M V K I G R K Q V P R V V F N H T D N E G V D S L L Q M H Q Q R R G Y T G L S F N V A Q I P E I T I D E G
RANDOM_621 S M V A Y I R G R E A N G Q F N R Y P M H M I L V Y G A M F N T T L A C L V P S L F E S N A A K V T P D Y F L A
RANDOM_622 G L T F L N A S G I L R Y E Q T L N V L G G P C K H T G L P C Q K V T G I N V P N E L T D Q D S K L E A A Y G P
RANDOM_623 A N V N T V M H R L I Q F Y T A P V A E L D Q I H F S V F G V G L F A K E I I A H K S F A A R F M G A S V T L M
RANDOM_624 L D V M P Y K A P T E S G A S A G V S L A V I A T L E T A A G A A P R D K V L E G T Y V W I P K A I S H Y I L A
RANDOM_625 F F N K L T A K A F D F D Q L Y E I T P V H L G N V P T L E Y K P E I E A L R W L I H F A G D T D V F R S A A K
RANDOM_626 R P P G D H L P L K I E G W Q I Q P K T F L R R A K T V D L A D S C E T R L K T G H N H L V P S G E I V R L S S
RANDOM_627 I S K L K K N N N K G V G Y N D K N S S F K I M V T H M G Q R P G L V S Q Q K H G D N Y F D A V H V M L E A G A
RANDOM_628 A F V S P L V V K P K S Q A P E R K S T T F D S R N R V C I R N C R P M N D Q D N M I F C R R I E L S A V P E E
RANDOM_629 P Q K V Y N E K S V L F I K T A T N A P T D Y A I Q H E F S L D M E Y S K H G E S G V G A T E D P D F T I Y K D
RANDOM_630 S S A E P R K K A V M G I H V N G N V F L V P T S I M T S G D A P S I I R A E T V R L P E K L V N C V V Q P E T
RANDOM_631 D V A S D S I T H F Y A E P A P T T W S T H K S E A T G F D I E A L M K K K T T N L R P F K G G E S E E G T L F
RANDOM_632 E S I S R D R G D Q T P S E A D P M L A E Y I T R L L N G S T Y N E Q I Q S E L F S L E P M P D I D G K S K D N
RANDOM_633 L A T N N S T E L Y L T Q D W L V I E I F V P Q K I D F I N D F E A E N C H G I N S M Y L G A P H F D A Q A T D
RANDOM_634 I C M K R G S F A S D I V K G L S K L Q T A A S V G F D G L K A D A W E G Q V I D W F K D V D A V A Y G I F E G
RANDOM_635 R C S L L G Q L T P G L V S V L Y N S V A G G C Y G K E H R V E I I M P A V R K V I V E R Y R E T K N D L S F R
RANDOM_636 G F S D A V Q L T G R Q R P G D A P T D E N T R P L Q S L I F R V A E S I K A M P A K S T S F Y E P C V L Q G L
RANDOM_637 V L R N T P Q F T V A L M L R S F A I L I Q K A S L D F S M T F Y A V A V D R F T V G E P N G L R T F P G A L T
RANDOM_638 F D Y F A T N T L A W K G L N V Q L A T E L K A I H A A Q V C L T P A I Q K R W N A F M K T N T R Y R H G L D V
RANDOM_639 K L N A T C G I P L I S G A I I H L Y G A G K T E I S E L R L D P V E F T A N M A S M S A P A G E P P A T C N L
RANDOM_640 G V T Y A A R G N N A R N L V V G T R S S L D R K T T L A I N S S T Q Q Q R R E T T M N E D K K E M A S F P A F
RANDOM_641 N S I C A D L P L K G D K Q N A A V V L H S R G R I I Q V Q P A E G A L G E S L I P Q G P H N S A E S H T V L M
RANDOM_642 I T D E F S V Q G S L G E Q S L Y K G S F K R F R G P H E G F R V F D S D V G D A A E L M Y V N S K K Q I S T G
RANDOM_643 P G T N S E D L K A G L R D P G E F T S E S T Y C K G T V W V L I R R A S D Y Y T E W L Y E S V A A V K A L F E
RANDOM_644 S D E I P A N M Q V V S K V F E L N E T D R R E I C A A L V A K I A D S S W D L K S V I A G E E A L L K L Y K M
RANDOM_645 V A D V F L R M L N K G A N T V D R Q P W L L R N N L H E Q S T T A K D A D V N F H K W Q L I A L V N P S Q S E
RANDOM_646 F K L S G L D Q G A F S Y A I S T E C S S N Q P P L V F N L R K G I L S S L K V E A A L M N M T M H E H N Y M A
RANDOM_647 W D G A S R V G R L L T Q P A A T L G M A L P D F S G L W A E I K S P E V R S D N C K E D L R T S I A L R G P S
RANDOM_648 A A A K Y L P C T A L I F V Q A K A S N A N E G S S Q F T P H C F G A S M E S N A Y D D N V L R D D G R H A I A
RANDOM_649 Y Q V K G I L A I F N L F S T E F L V V S T Q R N A N I E R L H S L N E C T T Q R F A I N E P D A L T T K T F P
RANDOM_650 A S A M D H T P E L S L S P D G V P I L R D K S A G A L Q V Q P F V S Y Q A D L L R Q T L D S K N K I A G I K E

RANDOM_651 S S L G L C G A Y I R L S G L Q N G A V V E S F A L I K R L S I T V G N V G L G R L S E A L R S G R S E Y G L V
RANDOM_652 G I N D D Q L G R Y N L L G N W N K L D I F V A K D R V L D K Q P Q T F L Y Y K L L Q V S E H V D F R G S I N T
RANDOM_653 S A P Y E M L A G V A P Q V K N G F T P K G G N I V N I F L T Y G S Y C S E M Q L G T R T V V Y H N G M T W Q R
RANDOM_654 V I E C K E K N V G D L I A G R L A R I Q I Q W M A E P T R I Q A S A F A C E M N W N A G V P I I G I L M Y T S
RANDOM_655 V I Q E L R D K P I E G E E F N L I K H R E V I T S M A R E D T L L E P L G T E I G S M D Q L G F A P G D S V E
RANDOM_656 P A R P P V A V T Y G S A T V G G K S K R N K S K D F L F F A T L V R A K A V T E V G F P E V V G S H L N N E P
RANDOM_657 T Q K A D S P N F K D Y V M F G I K T L S I V R A L Q F V G S A K S S E I F L S A L E V A R P I N E M F L W C G
RANDOM_658 F Q L H P S A R D S L L D G S A L S K I F V P S T T E I S G V Y T H P S L H Q Y V T K R S L V V V V C T N I A T
RANDOM_659 G T G S E T V D A M R G A E G S D T G D A L F A K V Q I S E T A R G P M S D F R H Q E L A V L E S A N A P A T G
RANDOM_660 G K L S V N Q Y L L W S A G S S C P G Q L G E L N L P M P C K S S I K T G P D P A Y F N N A L E T P Q D D M S T
RANDOM_661 G L K D F I Q S K E E Q E T D N G P A D P P D S K E R G P I E V I S S L Q A H G A D E L E R P L T L L T P Y L R
RANDOM_662 V I T R V K N V E Q L V W E E R I S V K K M F Y K K L R G L G A D L T Y A Q I D G F S E S H A S I S E Y D T I I
RANDOM_663 V G S T K N A D P L L R T Y T T G V A L L I H A M G S A L N P L T A N S F L A R L K R W N H R S V A L G W Y A S
RANDOM_664 S R K G K Q M D C P G F Q G D C Y I T Q S S N A M V V V S E A I I I Q T G S H K Q S E Q E R A L A R Q G L G E M
RANDOM_665 I G W T T D G T D F V G R Q S T K D K V L S T V I S S S I N A P P D T G Q T A H E G G G S K H P T S R D E S L H
RANDOM_666 K L S K G F L S Q V P Y P Q Q N A V T D S I K L E F E V V Y K H I F M L S I A T W T I G S T Y E T K R I Q C Y F
RANDOM_667 H F Q R M Y K G D Q G N D V Y G G T M S K N C S D F I A M V Y G F K D I P D S G A I H K R A G E T V V G K S H E
RANDOM_668 S T I S W K S Q V A T N D S T T G D T T G L M Q D Q E E K D M H L L S Y P E L A L K T C W D L S H E E R D R A L
RANDOM_669 R K N N D F P A I K L P L A T A K F T S V K V V L R T R Q E C R R P H K A A V I L A T E A S T C K C S N N R A K
RANDOM_670 A M W L T I Q R A A T E D R D G L F A A L K V V V G N A I F F A S L L V L V R I E Q G K M D R N G A R I E R L V
RANDOM_671 P R S L I Q D F V A A V E I S V K G D T E N L Q D L N T E D W N I S C F G Y P N L W G P N R E S N N E D L E Y S
RANDOM_672 M S N H D H R K L K L R N D T A F T A N G T F C I R I Q G I H G L L D G T R K L Y Q S T E F G A I C M K H G V E
RANDOM_673 G D P L R R T L G W Y G S K V G R H Y P E L L L Q P L S Q A H S D V V C F P L F A K W I A V P S Q A W Q R V F V
RANDOM_674 A T V L I K P T E R M L T H R S L L V D V E L P P Q E L P L P N E P D G V R E E V Y S G Q I K F V L A E G Y I H
RANDOM_675 D T G E E G F F E S V S N S R A V P E T L K V P L C D I I W E Y R E L L F G E A T A D E T S I L N Y H L A I I E
RANDOM_676 G M K Y S A L G I A D G D A T H H E F K K M S R P I Q F P F Y Q R E D L T C A G D T S T P Y K R N V E S K T R W
RANDOM_677 R L L G T G P K I H D E P R I T L N R P A R D E G G L K A P I R R T Y R N R G Q V S F Y S S R R Q Q V A P A S D
RANDOM_678 E N K R I T K V A L M D L D D Q F V W P R G D G K Y K A A Y L E N N D A V S D S D A Q Q A L P L L N P N Q E M R
RANDOM_679 L L Q T E I I S S P E R D G K D M L Y N L Q V D A Q D D Y D D E P M E S E V A S R I T D K A W R E C L T H F D R
RANDOM_680 D G M Y K T L Q M R S S E P E Q G L F S Q F L E R N S E A P Y M L L Y S T G T T N H R N C A S N E A Q A G L E H
RANDOM_681 G Q A A L T G F N K T P Q M V A A D K P Q M T P Y D L A E F C A K N F C N G I A N A P Q Q E H Y G Q M L T K S W
RANDOM_682 I M G G G A Y L V R L G Y L I Y I I E C E D I D F V E G L G C P K S S R Q K G L T V G H D G Q V I Y F P L D E R
RANDOM_683 P L W A Q K C E V W G L L E G E L A K V Y L M R F S A L Q W E F I E R R Y G E N P T I G P K Q K K K K L G E N P
RANDOM_684 Y V A P T T G L K I Q R F H L V H N V E T Y H T V P P K K F G A T I E Q D H P V R K I N L L W S T L V A N C V D
RANDOM_685 E R F A R T V G P S Q R L D A V Q E M P T G L D N R F S E W H K I A Q K R L C P S A A V C S L A L K R G N G G P
RANDOM_686 C E D A H Q T V S Y V E T I P P N Y V K S K Q R A N V N L I L Q S R K V E Q A E I P A A K S D E R Y C F S L K I
RANDOM_687 C E A Y D L L S D F I K P S M I V T V P K F H E A E L P K K G T W I G S E A I D T R L L I W G T A E I M G H T L
RANDOM_688 G F T I P T W D G P P R Q K Q K I M F G F H K K G I Y F M V G S L Q Y D A H A T R N E L K V M F V E G N T V I S
RANDOM_689 I I V T D G C K T K K Q E V P G L D S D N S Q C S F H V R T N L C K M N P S F G K P P Y K I V P R E F D S E S P
RANDOM_690 H A A S N N G G E D D P S A N L A R T P A L R F S P L G A R S A V E Q H P I A G H A A P T C E P T S V P E I S G
RANDOM_691 L N V E H K Y M H L W G G D W P N Q M L G P S R D N Y S L Y D S E W E K A V P D P S Q P S A H Y S S Y L Q K L Q
RANDOM_692 A A G M F A Q M L V P P Q E P G V N L T T D P G V S D V G G S S G I S V Q N L S L G K I T P G C T P G C E I E F
RANDOM_693 V E F S C R A K F G P D T V A C A L I P F L D A V G S T E D T F G G V A L Y E P K G T L N V R S L M L G N R K Y
RANDOM_694 D G S C I Y I H Y A R M E N I Q I P G L E K F L V N S E E L N A D T L K S R A Q T I D E E L V E C Q F L F K K D
RANDOM_695 I K V R V G P A D A S V N L A D Q C L G V A T C K N T L G K G I Q F A D E E M A A T R Q A P P D A K Q G A T R A
RANDOM_696 L S Q M S G K D T Q A N T V E S L G H V F G N G N L S H T D S E E P A R A M V Q S E L V L H S K S A I S P L D S
RANDOM_697 N T A D G W D N M R K M M A Q G M Q V A V G Q L L F E P P G H E E E L V S D S L D G T V C L N P V E Q A V V S C
RANDOM_698 S T A V R L A G S L T C P F K P G A S D G G D K V G K L E K R G L A T I E F L E K Y P E S K P G R W A K G V I E
RANDOM_699 I K L Y T Q E G G K Q K Y L S K E T D L E L P G Q P R E E K I I P L D N Q L A F G T Q N E K I M K D Q I R L K A
RANDOM_700 D S L Q D F V D N V G L G N A R G G A I N M N T A A G R W G I V A P R L A D T T I G V L L R P C A G C R I A G A
RANDOM_701 E E G T E E Q V N P A K D Y E R E M L H V I S H W Q T M E C I S Y Q S V Y R A F S V N S T A A S E F Q S D D G Y
RANDOM_702 Q R G L Q A E N Q G C K M V R I V L P A Q L P A Y K C F S K T W S F N I K A Y Y P D N Q R E I V Y T V D Q S G A
RANDOM_703 N A Q E M S V L L Q L K Y A P P E S R P H V I E S I N E D K L I S C I L L Y S R F L G E A N G I C L C R R N K Q
RANDOM_704 Y R T V S A K D W Q L K S L E D A K D L F L L A I N A E F T A S V F E R T E A Q L A K L I S L L S S I E E Q Q D
RANDOM_705 Y E A C G N E V A L P E P A V F T K K L N C Q A S I Q G Q T V L P L A S G T R A P D W G S K Q P I T L K H A E L
RANDOM_706 I P E D N L T A S L A V A T T D C Q S V C P G R V S I V L I P G E I R A Y I N A E V N H H R A D A V L S V T I L
RANDOM_707 M G A D S C T S L H G V Q F W R N L P G K V H N A R S M T S A P L L K L D N Y G P K I P S T P V K F L E D E A E
RANDOM_708 L Y P R H D V S N D S M E Q N E I K T A D D F P I L R K K G T P Q G S E L T L H A E N V A F G K R T L P D Y I T
RANDOM_709 F L S Q S L K T E G P T R L D R L G E G F M L E C P P S C S V S S V A G R Q S K A A Y L L H Q S E G L R P A S Y
RANDOM_710 Y R I R N R L C L D D V M V F E C K L S D L G D I C P W F F I L A L V D Q A W I V M T L P G A P K L G D K G G N
RANDOM_711 W C R G T I M L L M Q L V E E V P H T R L G S M G A G L D W L G F E Q E S P L V N L V Q Q A C T H C V E E P P A
RANDOM_712 F N E L G G V S C Y D V N E D P K G E G T R I A A T L S D A T L Y I S Y E K T V I E S F K I P F Q D A V P V K E
RANDOM_713 G R L A K R N G Q T F D Q S Y F I T E P Y I I L Y D K Q L W A S E Q A Q L I K G I V E I L G F L V R N L F E L S
RANDOM_714 G Y T P K M G C S I I I E N H L L V S L P H M E P P N A F Q V G C G K T L A I L P Y S G L A R E E W Q M H D K L
RANDOM_715 Y L G L T D V P R D G N K V T K D Q I K I Y T V M T S L K S H D F K P F A N G C Y C T G W A K I G I A L K E T K
RANDOM_716 N I F S S K Y S N S Q T D W E S V K A S L K C A Q E S L L L S P G D F F A I I R K V R L E N L L S E G E Y A Y P
RANDOM_717 T L M K G L A A L W G E E E F G V V D C R L Q V Y K M K S Y K F S D N S C Y H I N G K L S R P S A I E K P P E Q
RANDOM_718 L V S P D V L S D R T F T N G E G F A S L P S P D A T R T E A G T D E S G T Q S M I N V G Q S E S E N I N A A S
RANDOM_719 A F T K A K G T I C M Q R F S G I T V T T W D N W K H Q K N T Q F L Y C T S K A S T P Y D G E D T A T D A I H S
RANDOM_720 P N A E V G L G K S L K H A R M L M L L M V F D D A K V A A D Q L L D H L E A S F R S L R G T L S G N V Q V M L
RANDOM_721 R V T P I R E V I D G V D I A L M E K M H S P Y Q D R S N G L S D W R L G N D K Y T H P I L T R G R Q H K Y K V
RANDOM_722 P A K D M S D E A C S C E P Y A E L R F S K E K P L R V L D Q L L G G P D F L S M N T G L L P I Q I T A F D F W
RANDOM_723 T E I R V A L A G S M M E P F H G G A E I N G A P C A T N F L H A C F K T R K D D W A K G V P K L S N R I S K L
RANDOM_724 G I D G E S H S I E L A E K E Y S V R F K N M I Q G E I L S Q C I A S P K L T A Y M V R D P Y T E K V E V Q L D
RANDOM_725 H T A L I A L A N W Y G G V F H I Q Q K N Y C D H Q W Q K Y P T W Q G G R M R V C E I I S D L D R S L A T I A P
RANDOM_726 D A A V A P L L D R S T E V Q I S E D M G N E S Q S Q A T N E P I M M N R P G K P L S N G G T A L S S F G K D T
RANDOM_727 P R L G E K D E L E L V R G P D Q E H F D L I L S I K K I F F E V T D Q K W M T L V V R V A S Y D L T Q D A G K
RANDOM_728 A I V P S I K K F G G M S Q T Q K L P K I T L A A S L A G A K L A G R P D K H P D P V G S R E S A P Q Q H V A C
RANDOM_729 I A T K L L C F A D V V G R N H F K W G D L T S P D K V G I K E P A F T G G Y V D L T V N S N T S Q A L G W R F
RANDOM_730 R L Q G W D G K N N K V N I S K C S Q T G G I L A P L C V S N N A S I E F S K A F L I R Q C N F S V G V K N A L
RANDOM_731 P M L Y K N K A I T S A E D L K V E Y R G N H E I D I E I A T S V V T R V A S A L T F I C T A I P E A C S D P A
RANDOM_732 K G Q F P Q A V V R M T E T I I G E G V L D A L P L S L T V V G D Q S S G E K V R L G N A L N V G P W P V Q E N
RANDOM_733 K D S G C P T F G A E A D E K A S H Y E T W G L S Q G D Y S I S N E I V A E F A L N D P L N E S M C S K S K I N
RANDOM_734 P L L D G S I R R G I L R S A S Q V A N S G S A N G M K A M T I L E K G S A R I T E V T G I E G Y S L D R D T L
RANDOM_735 F L N D P L R G C R A Y V C G G P V D H P L E G E V R G A P T M N I Q A T D W V V P N G S P L G M W F S S S K Y
RANDOM_736 L V R R Y E K S N Y V C E V P L L L F Q R G P M L E K P C E V T E G L F M N K Q G S G K V I S H T K L V E V A I
RANDOM_737 N A D M V K I T Y P K A I P F N M I Q E G L E S S V V S S S E A Y P G T R L Q P Q M L S A L D G N L S A R A Q C
RANDOM_738 G I F K N T Y H T A P A E S T S F K P R H A V E L L Y H V I K T I N Y F L V L S P Q F G A E V M S Q M T Y V T T
RANDOM_739 I L A N I K T I G A T V I C P R I V R Q E V G G S N R D P S L S P L R V G E D G F Q K P Q I E L E G V P N L L F
RANDOM_740 W R D Q G E Q D G E P E N N S A M E A V R T L Q T N F D D L F P R T G G Q N V Q Q I L K I L A N R G D K H K S Q
RANDOM_741 I D Y E I H G P S G A A P A E N P E E Y R V E Y T K Q L L Q I S D I K R S S L V T I T P R A E F D G V Y T L A L
RANDOM_742 T A P A K V S Y F R P P P T T T Y K E L S T P G R K L D G T V V G R E G L I D A E N V L M Q Q T T K S S S S P L
RANDOM_743 T T T V R E K N R R E Y T F K I I A L K G C T E G V S I P P F L K D S T E Y E P P E M K K K F G I A S G R K G G
RANDOM_744 N T I L M D T H M Y D T A Q A L P S P L A L D V A V N R W V V A F K G K R P K G T E R G Q L P S L K P H V T N I
RANDOM_745 P P A S G K N G K Y P N Q G L V P H L I L D S T T E V L F Y M A L I L I T I D L E W V T T G Y Q E Q N G S T P P

RANDOM_746 N P L F E D R H N V I S C P P H P T L N G I S P T I V S Y Y C T W A Q F D F P N A I L K A Q A P S G R K A G G P
RANDOM_747 K N P A A H Y I I S E G A M F S T S V G F A I A G G K F Q I N Q V H E V N P V E Q V Q A S Y T I A T V L R H A S
RANDOM_748 S I A G A Q F T Y L N D G T Y V A R E M S D K T I E Q I G V F N V V T L R V I L H V H S Y E A N V G E N G K S I
RANDOM_749 H N F G A L G D D W H N Q D V L L M P V F F V L Q A V L I F N L M Q E D K L D I G M C G V M N G Y A R R S R T L
RANDOM_750 N F A V D T L V A G K H T H L P G A E H T G R T A E W V P R S W V F G I H V R T R D V N G G I Q R P K D D P T P
RANDOM_751 A V K G A L L S G S H F V A K P G P A I A E D K M C P G E K G H N W A V G R N K K M A S T E L K I L Q E D R A T
RANDOM_752 P C L R G S I L D N F N R G G T K Y L E A M D R C R T P V K A R D V K G E G I K N D M T V P D V D R E E T Y H F
RANDOM_753 D Q E H T P I F E D I V Q A Y A D K P P Q K P R L V L T P Y I V H I A N K S N G L I S I R K G E N G A D K L Q S
RANDOM_754 F F S D F I M Q N F R F T T G R L Q P S I R S S T R A V T P R L A A I A W V E R H P S S S T I G S K K Q K V N W
RANDOM_755 Q K C Y M N C T N A Q H A L F T G D Q V N C H G E K D S D C G Q A T K R S L P Q Y M N K N H P W L E G V Y H P E
RANDOM_756 D D K K W M S A S V L G R K V Q M S L F T S M D L F N Q D E G V Q A Q V S L A P Y A Q G L S D S Q S S S D I F F
RANDOM_757 G L Y L I N T T R T S K L Y R L V Y S L V S L E M G Q Q K S L F A G D Y L L A Y L C N M E Y Y F P G S A K E V S
RANDOM_758 Y L F Q G S F G P D T I K M D A K N Y G P D L A L H Q P R C P K F L C L F I L W E R Q T I G A D G K P E V S L H
RANDOM_759 L V F K V Q L R F V D L I E K T D F A T T N S L E Y Y V C N I E R A V K R K C A I C E P A N V T D S V I F Y R S E
RANDOM_760 L L R V V G E K K Q E L K K G L D A N T D P Q F L A I D E R P G P E Q V C G G K A L A G P Q A K G P T N G N P F
RANDOM_761 P G G V D T T F V R R S K L A G D R K Y R T Y P L Y S T K K H H Q I T P M L T L F K H L I L E A A F L N F G V L
RANDOM_762 L M K F T Q I P S N F S S V S F A I G A K R T S P S R G T L I L Y G I R A C R T C D N W F P L V L L Y Y D Y W I
RANDOM_763 Y V L V V S A S P L P P G K I K A I W E N E A E T S G G E V H G M Y S R E H A Y N R T A V R L D K G S V D G L D
RANDOM_764 S M C L Y I R A A W P G W E V V K T G H I Q L C A R L L N T V H T L V E L L K A F S P R G I N S I N A N L L L S
RANDOM_765 A R M Y S K R A P M P V L A E I A L K T T T Y N Y N V L Q E D W Y G G P K E T N G G S S D N G K M K I L F S K Y
RANDOM_766 I T H T L A L V Y L T T C D A S H T N P E L S T N T C G F A R D G G Q C D G A F M P V G K L T R L E E N A G Y L
RANDOM_767 I T N L R G T E T N F G N R C V R F Q Y Q V E R Q K I G K Y A K L T S V G E K L I D E D A L E S A T F N N V P I
RANDOM_768 L G S R K A H P N M S G W A L L D N S Q E Y D K V H A Q F N L T L V R I S T T G A D L N G M M F Y F G E S S R N
RANDOM_769 L A N A S N W D D V L C L V R I S G Y G R D H L S L E A A I A V Q P K L F R Y P S I F L R L K E Y R K W Q V W L
RANDOM_770 G K T L A Y D G A I I L S A I D T V N G I E R T S E V F A T G E Y L L L C E Y S G A A T G L I Q R F G V S N I I
RANDOM_771 I I A T A G G Y Y M E K S L T R I V S A S G P C T P P Y T I A A N Y F K N L N P W T K I M G P F F E S S W M Q S
RANDOM_772 S G D E G L P K S E V E E K L K P V I N E A S Q L K E L S H C R S H G I G S D L A A T V K D K D L G N Q H H F S
RANDOM_773 L L A G P Y A E T K I D A I P V D P E I Q T L Y G V P P S Y A L H R F K Y P Q H C M C V L P G F I R Q Q F R T R
RANDOM_774 L V A S T K L S P Y L E G E P Y V L V L L T A F E R L T A L A F A Y L T N S V S D F A K S E F F A A Q T K G R A
RANDOM_775 N A T K D F T S S H L L R I Y D A A R P S I I N A S E R L L A S Y G F I K S A K S E R T R K L M S S P C Q L F E
RANDOM_776 E L K I E F A R D H T T R D P K I R K R T Q D K V R G L R A A N Q T S E E R T G V R N P Q F N I D R A D D E T F
RANDOM_777 V L R T Q A H D R G H G K A S S A R D C Q K E A L S R E K D H C V N M V R E M R C G N F T P S V N A R L G K W K
RANDOM_778 D E F V V V G A H K V H V W R E H F T Q R K R S C M K A P A I G A G D K N Y L N E C P A K P K S I L V E S A K I
RANDOM_779 R K K E I I S G E V F E P I T K A V S I K V A D G E K S E I S N Y L E L T I P K E P K R R D R I T R F V Y R T V
RANDOM_780 A R Y P D L A A P A G N S T T Y A G I G K P Y M Q N Q T R T D I H L Q A D D A F A S D Q F G I S L Q L Q C G A M
RANDOM_781 D T L I D D E I L R V L S C G A L E N L S F V L L C M A Y E T I G G T N L L Y D D G K G N F K E L L Q A E D V S
RANDOM_782 S D I A V L L E R E K P A L R V G K P W R A E E D S A H S L G T M F P S S P T I F G I P T L T A T D P V L T N K
RANDOM_783 L C A G E I L A E S R E S V L S F A C E V E G L H S A F R E F V P G E L S D L E G H R G K D T S Q A F I F Q G Q
RANDOM_784 N G S E N R A S M G L P E F P D T R T F Q A N K I K V E M T F E T Y Q G S Y Y E K D S F S Q R T F Q Q V R H V I
RANDOM_785 M Q P R G C G A E N G G G G Q S D E N I H P P E W V V N L D R I G A S S K S I L E K F K T N Y S E N C H K Y D Q
RANDOM_786 N N L G C A K E D T S T D L P D G R Y N S N H F C V S E P A N M N I W T G V G P V G Q H V S Y P D D E E A L A G
RANDOM_787 T L T S A G P N A G S E R L G T Q A N H A T V P N Y D G P Q T V R N S E A Y G Q N G A Q R S R Y N T N M H Q Y S
RANDOM_788 L L E N I A I G D I R P I R G M E E H V S P Y R T E G K G L A L L K T T R K F P N F A K L K L P Q E K T S Y Q T
RANDOM_789 E S D F D S R A T Q Q S D P K K L T D G V A K A M P L P G R T A L P G S L Q N D V A N D G I I P G R Y L N K A G
RANDOM_790 M R L V T Q S R P A K V I S Y L H E S L G L Y F L G V R F I R R L L Y K P Y F G P Y G A V D K G W K H E G S H L
RANDOM_791 P F N M T E E A L A A P N T F I G L G P D E K L S L D T V N L D L N S D L A S C P G G G S P A S E A V G Y K V C
RANDOM_792 G E A Q N P V Y L I H K A D A D Q A T T H I C E V H N P E K L D P Q A R L S T I N D L E G K D A I K G A T F V S
RANDOM_793 L I M V I R E P R D G N E E K A V G V L E I R L L H P K R T S F A V S T I R E I D K G T L R V A D Y L L P P L D
RANDOM_794 P Q Y E W H D L G K P E A H G G L P L G E V N M P L A P K E I V F L K Q R R V T M I F C A R N E A G S T Y T K L
RANDOM_795 N L M R A T T G L I S G R N A G L E L E V I T S R A A L Q P R A L K M S G S A F I A D A N L L S G V V Y I K R E
RANDOM_796 D V I L A G Q P L P V G D T V H D Q M H V L D N N K T C I N I C H K D R G S I R D T I E K N E F T I E L Q G G M
RANDOM_797 T R V L K A V A H G T L Y G A T V F G S W Y E A K E S L L I V L V L I E Q L T Y A T N I I N L T R Q T D N S S I
RANDOM_798 P N C L K H G F W P T M G I K S V C Q Y T E S M R P W L K K P M D V G Q T S S D A G W H R S T C E F P Y L L A P
RANDOM_799 E N D T Q Y H R S Q G K T S Y V D R R V G L L T I G L A T D L G L E R S V A P Y L E G A L S S Y A T G D F I D H
RANDOM_800 P R N D P P P G K R R C I A A N G P S R Q L E G K H E T P K Y A R G F T N V A G I L T L F I G G G N K S N P E L
RANDOM_801 M S E V G F G M H S L Q V A T E Y D F L V T S V G E A T A D R I V V D E D T L T N T A Y R I I N G A A V I Q E K
RANDOM_802 R R E I L V Q K R P L F S A L I H A S S K Q E H L Q L I C Q L S S Q F V V P V S C Y Y A C E H L R M N C G Q V K
RANDOM_803 Q R S D L A H W L V A K E S Q L A Q T K G K L F L P C D V L G N I Q F S A G P P H P C A L R K K E R G A S G L T
RANDOM_804 Q Y Y D R K K V D S L M A D S V T I Y E T Y Q S I A S P Q M G R N M A Q Q W S T S R L S S Q Q K R N S Y N Q S T
RANDOM_805 Q N E A R A S G V N K D T P P T A A A R I Q L P N V S I Q I K S C S G G A M R I A W K Y E D M R G I E L M P N P
RANDOM_806 A G M F P Y R K G P Y F P G Q D K E S C L Y A I F D I N K F G S L A L T R N K V C F P L D K D W M G L D L K K D
RANDOM_807 F E V E R S A R L G R L I A N S P R G Q F Y T L A S K A A I V V P K G F V K R E L S L F L H T K K Q A P R A Y S
RANDOM_808 I K S G Q L Q I L L N L G D Q L K S I I D P I P L L N G C C Y D K P D P A T G Q V S G S E F N T P D F E P Y Q V
RANDOM_809 K R S P G N T V R G L E F E Q N K K E R F P R P I L L T D H V Q A I L G I S Q L I A N T E L H E G F D L D S E N
RANDOM_810 K E R I G K M K G E C G S E M D K G G E D A G P D Q A S V L G K F L E R N L G A F I G H Y S I R L F Q I H R Y M
RANDOM_811 D D A Q G S R Y L Y E A K A K P K S D I P P Q D Q N L D L A S F V Y E E I I S L D I T S V R P V Q I T E L P P L
RANDOM_812 F Y G V P N L Y D I K Y V W F Q A T A E N R S A P D Q H F P S L S V K A P D K P R R R D L S G S A I G V S Q E V
RANDOM_813 M Y T V C F M M K T G L L L T H Q K D E V A N I K V E F V G E Y K S F S R P N G G I M D N I N S D G E A A E R K
RANDOM_814 V S V K K T F R L P I L S E G I G T D R T G L P K P E F G K M S Q G H L L G G L G I D D L Y K R A V I L P A K I
RANDOM_815 L P G V G G S V E S V D L L G S L E Q I K Q R Y C I E Q E G I S A V Q R G S F P T K E Y G A G T S V D L A D H G
RANDOM_816 S L M F T T H K L L Y V T H Y N F V N A F E H I Q A I K P K N D L R S N L Q K E V K S Y E G A I G L E D K E A V
RANDOM_817 P P P T K T R I I N L P R K I S Q Q S L K P Y G S D L Y V N V S E C F W A R W L Y E E R N E L P S Q T A Y F A G
RANDOM_818 Q E F R E K G K K A F K Q N P R L S T F E D R G H S H V V D T R N L P R N Q F K C G A L D N D D R G N P L G L N
RANDOM_819 W A R R L E H L T N D S L C N I V G G I Q C I K M L G Y N S Q H R M V M I N T V E V H I F A I A L E V S T G A S
RANDOM_820 L Q T G A F E T A R L T G Q K A T T G G Q M Q E F F C P A C Y V R A C A L V D A V G T L L E L N P E K L K P L E
RANDOM_821 S H K T K P G E S I V R I D G D R L K Y T K E S K T G R C G R A V K R L G R V L G I F G K Q D A P I A L D K V F
RANDOM_822 S E G S S L T Q F G Y D A F A G A T A Q G T H G R Y V V N G Q G G Q K D C Y Y R R T G F C H L F A S L K I G A K
RANDOM_823 I M D G K Q L Y I V W D E L I E V S C S Q E D R E L Q I C C Q D G A K E L D A A Y P F V M Q Y P L M L I A A F G
RANDOM_824 T T R D N T G P E P M H R D P H H M F Y L A D P G T R L E G A G T Y A W A R P E A S C S L L S S N A P T D G E R
RANDOM_825 S Y T R T A R Y N W L K K W V C M E P L G D G S D K V L N P M K D S K E L A T E L G T P P G M S S D D I T L L E
RANDOM_826 R L P N W E S N V P G M R G L H Y G G V R I I L R K V T D V I V A I L D Q S L G K F T Y T E A V W E L I F R A L
RANDOM_827 F L W R S R T L S N L N E F P N K A S T L A K I V L Q E V G A F D P V F A L D Y A Y T R H S A S Q V V I W C A D
RANDOM_828 Q Q V N G V K L E P P V K P K V K N A N S V F C H G G D H V L L M G Q S Y V N G K F A K R D Q P G E R V L S G L
RANDOM_829 G A K D A L D T M F L V V W K E E K E R R S L S T A V S E G G F T S G P S G H K V A V V I L A R G Y G Y F I N S
RANDOM_830 I Q T L E Y S Y Y T I W T W C E A S Y T Q I Q Q Q V T S R L S G F Y V L G R T V H S Y L L L E I L P A D S Q V G
RANDOM_831 S A G P S D G L L D L E Y L V G S V T G A V R D V A E D L P E L D L H N I E R A T S R A F W T Y S V S P Y F D D
RANDOM_832 N H P P A N Q T T N W G H Q K S E D P P H K E I N G I K S C A S A G G N E C Y P V V Q L A L G L L A N P S M S T
RANDOM_833 Q D H G P A K K H T Q F I P Q Y T P N A C G A Q E E T F P L A D Q T V F S V D G K S D A V K P A L Y Q H T C C K
RANDOM_834 A S P A S G R D D R V G C H I S L F G Y G Y F L Q A I A T I S M A L E M L S L W S H G L R L S T G I E S P S K L
RANDOM_835 S F I G V L S S G Q T E Y Y D Q G G M H Q S R S L I P L A S K L E D A I V S D L G S E K A T P L T E R S A Q N E
RANDOM_836 G N H G G V N S V V P G F S S E C E T P Q H A S A N V A W A S E W G H K C L E A S T G P N V L A W I D N A Q Y E
RANDOM_837 D S D A R P T Q N V A E A Q I A Q L L F L Y S A A V M P L L L E G F D V V T A L F E I A S T P I V Y R S A T I C
RANDOM_838 K I E G G F E L M A M L P L D N D L C T K L S A S G L E G A N D T K A G V I V Y R C C S F Q F I K E L T I E I S
RANDOM_839 R P A P H S V L T Y I P L A Y V K K L G N N P L C L I N M M W A E L P S L R Y L Y T F D E L V I A V L L T N K Y
RANDOM_840 I F R A W G G K D G A H L H S Q E Y D E G G F R R E S S I L K A D F V F K A K D V Q N F G N S V L E G A N C L K

RANDOM_841 V V S D Q A E Q V C T E F V P L L V L P P N Y R S S R P G T L P N A T L D L E P Y A R N L S Q G Q L L E Q D E A
RANDOM_842 N N R A A S T F I M G R G E R H P P W V S G D D K I L F E D V T G N G Y R C V A A F V F R G V P Q K I A D E V A
RANDOM_843 G L Y N A E E G E K P M V Q I V A P V Q W I V L G Y P I T G E Q T L N D H D E G L V E I D T G R Y G G D V Q S G
RANDOM_844 L K L V D I T T L T V A E L S T L H W N G E K S W S V Q C G S G P R T E Q H V A A G G R K P T E V T Y E D F S D
RANDOM_845 A D T P F Q D D P C D W S I S T H D S E Y T H L S P E L G L V T A C E S Q D L P L K T S S K K E I Y T W D L A M
RANDOM_846 E N F R D K K G E S S R G A R A Y N Y K D K L N G V A K T I T E L G N D Q T R A E S E T I E S D V S G A T Q K K
RANDOM_847 R H G A T Q V S N N H C D Q K T G N L V I G F L C H S P T R S M R H M I Q S L H C K L Q P A M A V D P I L N S T
RANDOM_848 C A N E T E E K Q P G V N Q G K T T F P N L V C L Y G Y M N K D L P N A N P M T P R G N A L H R D V G T I F S S
RANDOM_849 N P R G V G Q V D P C N V S F G A D I R E G S G E S K H H L P T G D Q N R A H T K G S T G P T D T S D V I D A E
RANDOM_850 C W K Q S G V D Y Y V A A D N V Y A I E G C L L L V Y K G K V T G G E N P G L D P V L V D W G E P A S M L I K F
RANDOM_851 D K I G G T T S E R L T S V R W G M Y Y K L L P L T A A E T V I S G I L A N E L N I S L S S A P G G T L T N K A
RANDOM_852 F E L K D Y A Q Y Q D K A I G I S L Q K C Q R A G E E G S T D Q W Q A T S D Y G F A A W G V G A H T I K N G L R
RANDOM_853 Q R D I G M I F H I T L S P T N W T L S C V F E G I L L Y G R A Q V E I F T K N L F P P H N Q I V V L G C P G L
RANDOM_854 D L V Q T G P E F I F T L C S K P Q K N S I A L I P K F G H Y N F Q W E P G F K K D L A D V N M K S F P I L L D
RANDOM_855 L G H Q I G T F C C N S A A A E H Q S G E N Y T T G I S P I E G V K T N A V D S K F A G H W S P E R R S M L Q Y
RANDOM_856 Q D V Y N H T L R T L A N A G F I L E L Q R K Y T G Q N N T V L A K T R S N P D R A C F S T I M V P G D N E G K
RANDOM_857 T A L G E G F S G G L K V K G R S L S R T D P T G D E Q S I E F K L K F N Q V E C W N Q T Q P K K N I A P G M D
RANDOM_858 I V T T S I T Q P P I Y R R V S L A E S L W G I G E L V G A S V T L R S K M K L D Y G D C P I W G V A K P G T V
RANDOM_859 V D V L V C K E D G L R G V A V R N I S W Q R H P N E D P V Q G S I V R A A N V A G A V G L S E Y I N K V N A L
RANDOM_860 T R G F C A S I A M H S T L T D K A G V V L T A L L M I F L I P S N F G G L P S R A E Q F S P G D D S G V T V G
RANDOM_861 M S P C I A V R T E S S S K A K C R L G A Y I P P Q D A P H K V K L V V P D S P K A C E C R E V A K N N G V I C
RANDOM_862 C E G L G N V K L V L L K S T L S V S V K S I W E V H E K A E Q K M E P A L L A A L T K E L S S S R P D A D S R
RANDOM_863 L L K V V S S K K S V S P L H N R N E A V R G T L L F D A S T G G S Y E L K M A E I S L E T V R F C R A D L A T
RANDOM_864 K V Q P N K I R R G P N N V E R C R G I N Y I L K S A Y I R D A L E R N Q T L P Q N L N T L C A Y Q E R Q M T G
RANDOM_865 F E I I A I H L A F L G G A P Q W V V S D M C N T V R L V D D M A K L N V G T V G R C N V E T S G Y E R W D E E
RANDOM_866 N G D V G P T N N F G T E G S I M G I V K K K I A S N W D P C E R F S L W G R G K D A I G Y W S L H Y S A E P R
RANDOM_867 I F A L E S H V R A R I K T L A G G D F I L Y Y R E N C R V H V L S W G T E H I R K Y L R I S G V H L L E A E A
RANDOM_868 Y H Y M Q P N R Y P L G P V G L L C C S S M M H S V V F Q M A Y V F L C D M I E G F T G H I N D K T K E A A K F
RANDOM_869 V T H Y M L K K A M H F E L G D L M K D I T M S E H L L T V K S Q E D S T V D S I P P I D A S D K V S E L T I P
RANDOM_870 G G G S G R L E K G P P E L E L I V F K A F I I D Y A R A L F D S Y D L P T R L S R E E M L K R P M T K V P D Q
RANDOM_871 P P W A L E K T Y E V K V S A S P T L R I A R T P L A Y Y D Q T K G Q A D P G S R D A G V A E Q L M L A Y F N H
RANDOM_872 Q N C P L R W L A E C R A G P T G S C N L S I L A K A C Y K S R P R Y D L R R E N S C D R K K F S T S F Y L V L
RANDOM_873 V M K P V R T L K G Q W G N K D N M T R K I S T F S S G V L P K Q A S K H H I H V D Q G V L T A D L G V G Y Q Y
RANDOM_874 D H V R K A T Y R N F R G Y A G P K D E Q A N A E N L Y V I L D F S E A L P F N T R R S G S F H D R E A V R N N
RANDOM_875 I T I T D C L P G N E I Y A C K V H D G Y E V V R L Y S D S Y V V S T K P I K S K C L I V S G C V S L S C A F N
RANDOM_876 H F W I V W K V M Q C G G I N D P L P G P R Q K E K V L K P N Q L E H N G L M Y V L H R F Q E P K V T K T G L G
RANDOM_877 L S I T A C Q P W A I I V L V M A A V Q R Q E N Q N V W F L V S E V L D Q A W T A E G Q V L M G I E C V H P N L
RANDOM_878 F N S A E P V A D R A L K L K F A D N L F R N R N D A I S P S I T A L Y P E G K S K I V N V D S G G R T G G V L
RANDOM_879 R Y G C F E K S S Y G E H A Y F E I F V A A A L A V V L A N D I A D Q E I F L P S I V N K A M D K K Q P D P M G
RANDOM_880 V E L C L A P R T Q S V D A S D T W I W K T S Y W V P E R D S C I D L V K M D M Q R N P F T A L N I G K S A K A
RANDOM_881 S A I Q F G A F N I V Y R E G A V Q R D G F S L K D C T S G S Q A L E I L Q D A K L T N R T T T N L G G K A F D
RANDOM_882 Y W A E G K K Y V R E A L E K Y V G G Q A G D G S S Y L R G P L D Q E S K T T D D F K V L R P G G F D G G A I L
RANDOM_883 P A E E I F C H E S L T E G I Q S K G A M A S V Y R T D L L Q A E P K S A G Q T V G S R S A K P S L V T T V L D
RANDOM_884 E H W D Y V T P E P L A F L P D D T K L C C N D D A I F T V D R S T I R G S K L D W F L L S C Y E Q C Q G S I P
RANDOM_885 R E L K N E C T V I L R M P P T V E W G S I Y K L H E C L L N N E E A V F L K D L I E G M T N H G D S R N G L G
RANDOM_886 I M R D V H L L A Y Y M V P M A E Q P F R E G E P V Q G T T S Q R Y R S L S I I R P P K V G M V L C A L T D S V
RANDOM_887 P F A A I S G A S V Q I L V A S F L S S D Q L I G T L S I E G N G A Q L M Y E D P A D A Q E E F K D N V P N V S
RANDOM_888 A S Q R E Q F K S P D L W T F L N A G G I N T F F G A I Q D H L P S D D E A S C L K A N L P Y P L L V P K S T G
RANDOM_889 G A K Q L I P Q I V L K N I L D Y P E Q E R S L D S R V D E L L T F N I T E G Q S S K I V E R K I K E G M S L K
RANDOM_890 K Y I T G P W S D A P V L L K E R M Q I K A G P E V R F A F E L N C P A S R R S K T D L G E N G L P G R L K G G
RANDOM_891 L L P A V I Q H G S D A Q Q L S V V V S K N A D V P R R N N D T S S L I A N A S C L V I V S T G T V R E P R L T
RANDOM_892 S L T H A I F T T T P P V A M K A E S G I R D S V G N E R A P I F R L P D F K S A H D R T K P G V D R G G T S R
RANDOM_893 Y E T S V R C D A S G Y P G S V Q K I E H V A A M A R R E S Q D P L E L A N V A A L Q D V D E A K D E D L K F V
RANDOM_894 I S E E L A G H D N N E G A A K G E Q N L S G F V S S A R L E S L Q D E C M R V S L Q W A T R S T G A I K L E T
RANDOM_895 L L I N G Q N T G Y R H F A N K D L R G E L V L A S L S E L R D P I V F T M A P T R R P R D K K L R R N Q E F L
RANDOM_896 G S L S W L D E S V K A D G E D V S L V K A D K A V A K P P Q S N E K A L F P N T Y K Q Y E T S G K D E A G E A
RANDOM_897 Q S L A V A N Q K L R Y R P F R N E V N T Y Q S L S E I T T Y T K K Q R L E S S N L L N P S E E R E A E L L A G
RANDOM_898 C C L Q A K K L I V D N N Y R T T D S E E R S N S L R K N M A M Y N S M Q E T V R E V Y F Y I R H Y L A L V I L
RANDOM_899 V R Y I A S R K R E A Y S I A S T E T F K T V A I Q R W Y L F V K V Q E V P F W W T A G V N N S E D A V Q P A R
RANDOM_900 Y Q L I Y N L K N H K N F L Q N L V L S V L L T T D E N E N Q H K C G K S S N S G R S Q A K L P Y P T I V V G D
RANDOM_901 V Q K P A I R Q V G Y M A M S L N I N V F R E E N I I P G G V C E M Q D N T E F D D D A A A G E C M N S A P Y Y
RANDOM_902 V A Q Y K A Q E W G M Q L T N E G S I R F E E K F V M V G A P I I L D N A R S A S I D G N E L E T L M F I K V I
RANDOM_903 D C N H F I G Q Y T P M N T I A T D C R F Y G D Q L I Q P G E G A N Q L T K F N V P N N V S L L R V E H K D Q Q
RANDOM_904 A E D N S Q T S N V A A G F T G E N M R A F N K F F K I T A K W D G S C Y C V L A S D N L L E G E P A H L R I H
RANDOM_905 N Q K Q G S G C Y A A A A N L C D T S D N C F N E H V S D I Q G R E K A L S D V T R G V L E V M F S S K W F E M
RANDOM_906 A G A T F E Q K D I D G V Q G I K T R L L L L G N S S N I G E S M L I G S D S D N E G S Q V K Y Y K E I S E G L
RANDOM_907 V Q Q P S E E P R G R Q I E D T G G L G I P V R W T A T G I G E S L K K R R G R W R F V V W A S N R Q G Y P K A
RANDOM_908 S I G Q V S S G S T A R A Q S L E E K N G E K Y E Q H M K A F P E A V K S I F M G Q S I Y Y G T D L F N T I L L
RANDOM_909 S D Q H K P K P M F P N I L L I T E I Q P L R Q F P V I R F F L F V K R T T A Y G D E R S T L S S P P I K Y F L
RANDOM_910 L V N F T E N H K T N K C E G N R K N K D L A E I L T L F L Q R S I D Y S S A G W V S A I A Q T F L Y I L V E L
RANDOM_911 K D D V V A L L R D T M V K Q H P S F Q F R F L P L Q V Q Y K Y E K V D M D E G V N V M T G L A Y N F P S D D F
RANDOM_912 R Q V W D H D D F G I P V P G E S P Q P E I V V N R Q T Q E L W H L N D P V C A N R Q N I V E I S R A G P N S V
RANDOM_913 P D V A F G R D E A D L A L G L I L E D S I L E D L I E G D D F S A P Y E G R C S P I V K I N K D Y W S S L M I
RANDOM_914 L Q Q P R I G T A P Y R R I L V G K F H E S L Q R F Q L F G V N V E D E G S L N N G E Y K E R P K T L Y Y P R E
RANDOM_915 Q K P N G V K R Q P I L Y F S P N G G F E D A P V G P S P E R P K G L P S I Y K N Q R Q T L F Q L V S D D P T I
RANDOM_916 V G P F L N Y Q D G T R D Y T S G A H A T D L R E S W L G C Q E A S P L N K R H V I L S L K D I T R V K T N T K
RANDOM_917 A N P L T N K S N W I Q K M N K K E S A S D N T G K T F E D S A L A K V E E I T M D D F T V R E D K R L K R G T
RANDOM_918 Y E L I D F L A N S I E L F K A R S R K R K G K R D A S F K G T V Q A L V L S V T F I T I S S I I F C Y K E V L
RANDOM_919 S Q S W V I P L T D L C G V N Q P K C R I L W A T P V S L Q L G T C I K D K L R L F Q E V L G E A T A S G V T D
RANDOM_920 E G Y A Y V T G D D K L F V T T L V V A H Q S V G A T V S F S E A A K T V T T N F H I P M L T Y D P C L S L D P
RANDOM_921 S L T E T L F P S S S A R G P G V E M G N E R V T T G A H V M E S V W Y F K T D K T S R A P T Q V A K R P L Q A
RANDOM_922 T V L R I L P G E S G N N S K P S R N L Q A H R N A G H V S D S F P S L E D V P K E N A V T L H N D A G S L F Q
RANDOM_923 A G N R V E I G A G R G L F E V K F D K D S L L V H N V S A H A H I S Q R M L G L Y R R G I N E G Y P M F I N E
RANDOM_924 R S V S G V I A A L F D K E A D I E P A A V L A K G L D P R V I K M M T D K S H N G P T C A P S S L T Y A A Q H
RANDOM_925 L F G G S C I H V G I V A P K T T D D R K H E L L L E Q D V E L I K C S I L T T K S D K G F M V T G S H L A G N
RANDOM_926 M H W Q L V F S Y K R Y T N N P A W M M L F Q N T F S V D Y T T H M L L A G N G E L A A A F L K H D K G P L R F
RANDOM_927 A T Q G L Q Q K A H Q S K S P E V D E Y P H W C T I Y K K T D T D N N V K N T L V F Y T G I P P H A K Q R S L V
RANDOM_928 L A R L V E K S Q V H P S Y L A A R R G Q R A R S T P D S M Q N P N I E E F N V N V D G F E Q T S S K R P Q T L
RANDOM_929 I Q E N G S E H S N C C P N D L G V V S A T S V I W A M E R C K G D Q F D P E G Y E A S R R D S V Y G A H D K P
RANDOM_930 T K D A I L S K K I V N R I D F W S F Y A G V S Y D E S K G M Q D I R G H T E N G I N D G V T G V K F T S V G G
RANDOM_931 R A N G W M V T E D S V V R T E A E D L I E E R N W Y S L K R G R T E G T V V K D T F P R K Y G M P P G P L R L
RANDOM_932 I K L T N T R N G K G G S N N K T Y G V Y N V H Y A P P S P N Q T Y A K L I Y A A D S E N V V Q L K G M W N S E
RANDOM_933 H E L A N T K Q K P I L C G H R G D H E A V D T R S D S V D E P K D H M D I A F N G A I L R A E R D K F C G G N
RANDOM_934 D R I E V A R P A E F S S A R F V N S A A W L V R L A T Q D Y C G P M T E K A D M E L N G S V A A Y L D Y I S M
RANDOM_935 A L V A S S N L M R L G C V V Q F Q A D W L D E F N V K L N Q I S T I A T G K P G Q L A F V A R A K V K I L L M

RANDOM_936 V E T R G D K F C A A N R T K R C P V K S N P L G E N A L V E W V S I K D T P Y W E A S E K D I E A T V R L E L
RANDOM_937 C K N G E E E G A S P L K E K H R R K D L R E P R C C Q N H C T R L A R A L L E A E V G S G G V T V V I P N I K
RANDOM_938 E Q E D L K T D E R K A L L V L F E K A G D A S A S P H K K T G E D T E K E G P L P N P P T A G W A S T G N G A
RANDOM_939 L R V I W G S V S L A A N G Q N P K L A F T I P E R D N V D S K T N K T E I V I T M R K V M C L V A K N P F G C
RANDOM_940 S N L T N R K H I S V R T I V I A A L L F K I P M K N E R V G K A W Y Y S W A S T Q S M H T P H G E S L P G K P
RANDOM_941 W N L T A V T R S T H S V I S S G K K Y L S M R V R F H Q C L K Y D T H V E I T S A V D G P L P N D K S S P N P
RANDOM_942 G N A L N N L L A V I D V F R K E G L G P A Y V A G G G A R F V L L L S R N D T D L C V V Y R D V A S A A F S C
RANDOM_943 A Y K F G P V V P L P D A E G C L H A G A L R D E T E V N S G Y S E Y S S D H K I R A E C G Y L E E I F L A S E
RANDOM_944 G I I N N G E R A R V N G W F S T E A T F Y P E Y V L R M W L T Q L C R L G A D V A M M E V E F E L P T T D V L
RANDOM_945 S D I Q G M K S D Y Y G I K T N R G Q S S I L S V N S S W N L M Y D E A V K P P E Y Y Q A G R L L R A C I R D K
RANDOM_946 Y V A A L D G W G E N T A L N F A T E Q R V P T Y E N F A P P S Y E G L C A I T N E L N C I R L K F S K Y I T M
RANDOM_947 K A S D K W P F D E K A P T R S V G T C E L H R L T A N T W Q G D T S R H E K A D I L T P R M R E L C F E E R P
RANDOM_948 K L L R L A L K K K L E W A A E T E T G R L P Y P S E K T G T M I N S F G A E M R I K R A T L Q P V N V I D V V
RANDOM_949 D N I E R I D I K Q F L V D T F N Y H D K L E K S Q L G G V S F L S G D V S L K L F I L P A T V K F G A S C T D
RANDOM_950 Q T L N F A A Y T K T E G G E V Q K T M I T L M I I E G P F L A I F Y P S S K Q I R V P C L C S Y A T D A N A I
RANDOM_951 A P R M I Q G E G L A V D K E K I M H K I A Q C S G C S K A D R L L V F D G I Y S I K A R I H D A P T S S T P P
RANDOM_952 K V P F E N K L D Y H S G L N E A R G L V D H Y Q P G T A C S A R A A L T F L H T D D Q L C L T G T D R S L E R
RANDOM_953 S E A I A V S T P L K V D T K L Y P G A T Y Y V Q V G S P L S A K L G E F W E H T G P E Q P K L A I K R C A A Q
RANDOM_954 L T T R I A L F Q K C Y E V G A L T L S Y S L G E L Y S Q E S T P F V V Q I V G R D I S A A V P L E Q A V H D F
RANDOM_955 E S K A N I P S G I T I D R Y T I L I F S E R A K S F S P T S E K G S L N M G K R G G Y A S R C L W I V S E R K
RANDOM_956 D I F M M N E W G P D P A D H L E V Q K A A R I G K G P S A N E V V V L K Q V L E R A D K F T G G Q W N V A I L
RANDOM_957 S R H K R P S G K G S N L T I M R I I H T G R M F N A L V A A A Y E V S Y G G R T H E V V F E M Y A V F A A A P
RANDOM_958 T D R G G S E N N V S L G Y I S N R D F H L T T T E N G L F L S A V H S Y D D S A N A D D R A L F R T F K D Q Q
RANDOM_959 P Y D Q I V Q M D H T Y P Q V F A L N V D D G A D L T Q E L I L D S T V A D F P P L V H R S P R T T K N K S F H
RANDOM_960 R A A N K L T T A C S L R S R W K R P R L I I Y T V A S F D K F E I K F S S Q E R L D W S M S V G P V G A S N M
RANDOM_961 T L L L N I R V I S L E T A A V S E R N C G T E I K G G Y H T S V C V D V R N L Q K M S N Q T I A R A T L D R R
RANDOM_962 Q G K T I E I S G M V D A D A V F T Q A T P D S E Y N D K K D I I T E R Q Y V I A Y P K Y I L Y M I L L Y A D K
RANDOM_963 S S G I V L D P Q N Y R E T L L V F L V T S G I G F M L D N N P D L A L D I S L D F R I L K V G P A E S N I P S
RANDOM_964 P E Q V S C N Q V F W R F M W G E G S D V C K F S P S G I Q R F A Y S I R Y F S L A P M S G F D E V F L N E V L
RANDOM_965 A V N S I F I R A Y T F V I Q H G L S D N T N T N F N L V K A E F C A I R K L T M I I K G A D N A K P I A H C I
RANDOM_966 A G I R D S G T S R F G E V Q V G R I I D R S L Y D E E C K N E I Y F P K L I A L V L L I K T A T L K L N N I Q
RANDOM_967 Q C I V K Y F D R A Q N L S R S P S A A E T T G G H V L G G E E F A E L I G T F G A D P K G L V C P N Q F R T L
RANDOM_968 A A T V S A D R V P L G A L T G W S T S P N S Y E V P N V Q G I L E S D L Q V N E G Q F A G D R S K R W L D G F
RANDOM_969 K K E G R E L G G G G A V V V L Q A L T V Y S A F T L K L I G R V Y E I A L L K I G L S R Q I D P R V T R N N V
RANDOM_970 H G G T W L C K H S E R V M K G R A L T L A N Q I D A K G Y E R G D P S P L G R N G D Q D I R D D V T R G A A V
RANDOM_971 A P A P V C G Y Q A M R T Y Q G L K Q K L R L W K F P G D L N W D R R I E G L W R E G E A L E F G V G G T A T T
RANDOM_972 L R V G K L P I P L I L G G N A V L A S Q D I T E R T T N T G I L W S A H W R Q A V G C A P R L L H K E F G A Y
RANDOM_973 I L P N E E R Q S Q I P L S R E T T T S W I S F L H P A S L L M P P W A T V G Q K H S M Q L L C R P R P E R K P
RANDOM_974 S L P P M V D S M G Q D L A L S T R R A E L T V I K Y N K V G H A R F M V L E V P E K K F R N K N A W N V A W R
RANDOM_975 F L Q F L V T P G I G G F Y L V S D Q G R A S Y Y A N V A K F I I M I L A D H L T L C E G Q E G G F L A E S R E
RANDOM_976 V R I P I V G G P D L K V T I L V P N Q L E A M T G T I Y F R N A R N R G F L D K V A F Y P E V A E E M R A F A
RANDOM_977 N G T L S Q E G S Q M E S R V G L G Y D T P N T L A L M I C H C V E D K K G L G Q D Y Q P D G K K L S Y P H H F
RANDOM_978 A N S A D T V L I A Y R A R W P A S L I N P R G V A T L K Q V A F L A Q D Q V T V W G L D S A P E V E N N E A I
RANDOM_979 G I F E Q G L D S L V L I W F S E Q V D A N K A V P P P P L C Y Q A V I N Y R E G N K M R A I V N E N R D I A E
RANDOM_980 F N T V Q P L R D L S K P G I Y T E T T T P V M P I G L V I T Q T H S F N G K T K Q R E I T P L Y R P V A L L F
RANDOM_981 S K A P F G D E R E A R Q P S Q G T D D F Q D V Q V M V K S K V K S R Y C K Q T F D E E V T I G D E G T L G I A
RANDOM_982 L T L L K C A E I A L Y C Y V N C K T V L L G T D E P E P E H K W I T P G K A I P I T V V V V A N D G A L C I A
RANDOM_983 P L C M W V M Q V Q T S E V N L L E A T D N M M Q F F W P E I G D E D A A P Y N I G V L F D Q L V D K L G G L F
RANDOM_984 C A E A G M V L P L Q E G P C Y W D E Q T F A K V A E M T A R F K L Y K G G E D F D P Y V I L L E D D E F L P Q
RANDOM_985 N P H K D D T F M A M R K D T V D I G M T V G L G G E N R L A T R N V I M G S P S I S I L I N K I S V L K K D P
RANDOM_986 D P T F Y P G Y S V L D V L Y K E L S F E G G N K L E E S G L V A G N S C K V R H A G Q W T R P T L S S L K G L
RANDOM_987 G G E N S S K S Q P P E G I K M E G D K V G Y S S G Y K M V A I P Q R A D S E N D N P R T P G Q K H L Q L F T Y
RANDOM_988 L E V D T P R W A L G R V L H A D S I Q K P I R Q G L P E L L T Q P Q G P I N C Y T K K G M I P A D N P L G A Q
RANDOM_989 H F R N M C I M N T G S W H V T E S N I N G E L Q L G G S D N K R D R A K S A S V I E A I A T Y I A G Q G L C I
RANDOM_990 V P E T S A L Y R E A Y R V S D R K V F V S G R S R S L V H F S P A G Q Y T Q D Q S I E K A K S L G A S N V D E
RANDOM_991 A M L N R G P Q A R S T R G E K S K T E K A T N L E D L T N E E N S N N G N G K Y K N V N H W P F K G S W L P E
RANDOM_992 P A W G K L K L F Q C A E R G L A I V A D F E D R D N Q A Q E L C V G A T Q A Y Y E S Q Y A G D K Y P V S P F G
RANDOM_993 T H R R D G C E E Q G V A Y S V L Y L L S S P R P T R D S W E L M S L L Y Q G S G A R D C H L I F E D H L L L I
RANDOM_994 Y Q D S G S G K L I W T H Y K Q Q P N Y S M A R H E L Q M V L S N V N K G S L A S S P E K I L L F S A L V C N S
RANDOM_995 V S S P L R G Y S R Q V A H Y V V F P R E I S G I N R P N E V C A I H P V A R Q R Y L D V S N L T D I T T L C F
RANDOM_996 I I A E S W R E S A T Y N S D T I V L S Q V A L L I F C K S K V G V C E L M V I K L L D G A G L A K G T Q A F D
RANDOM_997 K D Y L I Q F I R K L K T W C H G H S T Q I K K T S E A N K A V S L Y E E K F E L E Q A G K P T Y G G Q R V A Y
RANDOM_998 V E R K N A D F S Q I C N D K T I R W L A M P V A F I A E S V D R V L R R P A L D R D Q D S G M M E A H E E M L
RANDOM_999 L F L E E A G I P F S A V G I N I N C Y D K I F K M A N I F D E F N I L C A T E V L Y F L L T F V G S N E S L R
RANDOM_1000 R Q I L I L A T Y E A Q G L L V K V K N P I L A V D C M V G L K S V V A Q E S M E V F A R F V F G A L L I Q I S
RANDOM_1001 G V S S G E A V F E N L T E K A V K G Q V T L S P E I Y A E R V E R N E L P I M P G L Y K I I Q D G I S A V E L
RANDOM_1002 P Q L T R I P Y A M C G T I N Q Y D K I L D D Y I Y G G Y H N L K D Q E R A W N A W A V W Y V V A F V S W K Q G
RANDOM_1003 T G L S L H K Q Q A S R F A G P L Y F S I R T E I Y C L G I L N D R S Q I R I A S E L P V G G L C Q A A E V L L
RANDOM_1004 P E I V F F D Q Q D A E E L K G V V D Y L T V A S K Q K V E E M A R V P Y L G I T C G E P A C F S M K C H G E V
RANDOM_1005 F A H R H R G E C Q W L V G K G A L L V D G G D S T S A T L R C A D R R N L S L P A N I W C L S R C V N E M G N
RANDOM_1006 Y A G I K W L D A K L A R P A N I I Q D G E K L K M D N G A N S Q P P P G S G D G G P V T A P K D T K S W R G K
RANDOM_1007 A G D D S A V S D G A L S W G P I K V L S R I A S T G G A D S E L Q Q G Y P P A V I S S F G P V L Y S M I G W T
RANDOM_1008 L C W G T E D R I L E K D K T K E N S N A R G Y L Q G V G I E Q I N A L T F V L P G W P D L D A F G K D R K I K
RANDOM_1009 G M L G V A L K L V K L T L P K N F H G D E C A A L G S K G L G D A G S I E G E S Y E V D V D P Q K I E K T T L
RANDOM_1010 T G Q W I G E R D R L A P N I W R V A Q H Y P K K D E P I N E I D G P V T Y L E F V V G Q S A Y A R G A L Q K P
RANDOM_1011 P G I R S P T E V C I D V N G G G S R S T S V Y A G M A C S I N S Q V R L V E Y P A N N E T F V V M L I K N N Q
RANDOM_1012 T E T E Q G L K Q L N V L G Q V S R K G Q G K V N I P T R W L A K P I D A L V S C K H R A I S G Y T S P T P E P
RANDOM_1013 R D S L R M T E G L C N A T P N Q H V R G E R V G D P I T S I C T S D L G L G W L A C C P A G V W E H G L F L G
RANDOM_1014 T L K E T Q R Y K A F L A R M R V G T S R Y S E N P H V L M Y G R E A F P V D G T T A L D L K Y E S K L A N Y A
RANDOM_1015 G S M C F A C A Q I V S T A V I A Y V G I N L V G F T R P A T S Q T F K H K S A V T D K E E A I K L K V I R G H
RANDOM_1016 K N I V Q T Q D C I K G K P K P D K A Q G A D Y N H D L F D E N F L T G V N H S D A N C P V R S L G M E I V R Q
RANDOM_1017 L W E A R G G N F V E A L C D L D A F H D L E N K A E F L A N K A A G C P A F K I L T Y K L V V D M A R E S L E
RANDOM_1018 L T D E Q T L E K N Q E R P V T E G T R L V G L K E G S D G K V L A G W K D T A Q K A A T C F E G Q D Q V Q M H
RANDOM_1019 S T F Y E N G S L K T R S F G A F W Y R F N H C C P L E G V P Q T T W R S L W F T T P L K F S T P M G W P L P P
RANDOM_1020 G F V S V S V L S N V M A P V W R N W D S Q L T P W R P G F E A G S E V A L L L A E Y D A V L M D L Q G I W D I
RANDOM_1021 D D V G T C M D P L N M T R T F T S I D S L E N N A E S F L I Y Q K E T Y Q D P N Y P H T E P V G E L K M D S Y
RANDOM_1022 A R K S L K A T S E V V G S G M G R A N S A F N A R V A T N R I L M M D W I S L S D S T G W V T R V P L L C L A
RANDOM_1023 N R A Y V G D G L D A E P S Y F D P A P A R S V V S S E E P P D I M K R E M L D T I A M Q Y E N T V C T N V A F
RANDOM_1024 T D D R S G D K P V G Y K A Y A A T I Y A I K T D R G I I L V K A G D S P F Q K N C G K G R A T P H E R L K G K
RANDOM_1025 E A E I L Q K Q S E E G S G G N G S K E P A E V P P A L P A F A R Y A I Y R R R K R A N H G G N T N T Y G N E A
RANDOM_1026 A V G H I Q I A G D E L T E I I T S P V S P N A W S A L C Q S T Q E H I Q C G E L L I W H K G K Q G S P N D D L
RANDOM_1027 V E P K Y H G I L Q A T G T G E C N A V V S Y S G V A K D P I S V E E E S I Q P T E L S R F E E G D D H A T E N
RANDOM_1028 D P R K G E Y S S R L D D L S N V G G S E G L T K H L I K A L K E L R G V K T K G Q V M S A S A S D I L E Y F R
RANDOM_1029 Y D K Q I S I K S D K R E G S W V G H D K G E M G R G M G V V L L K D E I G W A V E D E E R C E F A I L A E L D
RANDOM_1030 S G V P T I D G S L I A S P L A H A Y G Y L R G H R D D N K D Q G G N Y Y D L Y F M G D R K P S A E Y E V A G K

RANDOM_1031 T T A G K G T T R R G K L K K F D L L Q H G L N P F R K W K L P S L L T S R D V T R V V D V Y F V P L I A T C L
RANDOM_1032 Y T V K Q A T Y T I K V Y Y A S K Y E A I A K I N H G Y G K G V L T L Q L P H C P F T R V P A L A N E A K F D K
RANDOM_1033 Y K E P R E I L C V F L T I E T D K L N D N A F A L E L H P I T A S A G K V K R K G H T K A D A A K L L D P R S
RANDOM_1034 M L L F V A W W A P S A Q G T E R G T N V K A M N K S K H I I P G K T M P R M N S V T V F C S S V F P E T E K S
RANDOM_1035 D K D L I E F G E K H D G P N R E N L Y K I I F R V Y I H I W G D I I G V K C K A S D P Q S M D Q L S T G A Y L
RANDOM_1036 K G E K T G P A G K D W Y A S C I K I T G T F T L Q D C M S L I N D V V F R A Q Y N T S A Y S E R L E Q N E K C
RANDOM_1037 M G L D S V E L P T G T F A H E Q V V N M L D H G V A K R S Y E L L G L A P Y G Y T D R L Q M S E L E K A L S T
RANDOM_1038 K L F K D P E V C L Q E G A L L V E V G L N M L T I S A L R L P Y A Q G E Y G T S P K I Q L E H L C E A I G P L
RANDOM_1039 I D L F E A L D I Q Y D W Q V T T Q T I E D N T N C G G G E D D V V V F Q A G G S N S K P C A I Y G N V E K Q L
RANDOM_1040 V S I C K D G P K S P D Q K M A N G R R S T A A P A F G S G L S K L P D W D T N L I N F I A S A P P V S P M N K
RANDOM_1041 E L I Y G Q L P S L Q S G V F A V S K G S L R D I W N W K I Y C E S D R L R I L Y L T S A Q E S E N D E E K G V
RANDOM_1042 I R E K P N D R S D S V F T V Q V A L M N P A K K T K V N T G R C T A F L K E R D S D Q E R G A D T H S E E N S
RANDOM_1043 I H F R P N V L K R H I G A E Y D I P N A A N P V N S L E D E K K S H D E Q H K S Y P T D L T E A T R F V E P A
RANDOM_1044 L A W M V P K D G T N L I G G Y L V L D E A I T N V K S K T A L S W F N V T E A K S I K M Q E P A E F P V S P P
RANDOM_1045 L P G A I A R M A E S V K T E L V A G G P Y R P P W I F D L G A V H R G F I A P S P N V R T I G G G V A A V R R
RANDOM_1046 H S P M E K E F N E L T G K A L K S R S A F G I S L H A I Q E D V P K P C T Q F Y N S V A D R N L E A M A F L A
RANDOM_1047 S D W G W W C V W S E R L Q M C K L C G R E V K R R A R R Q S I A D H Y T L G K T E N L P Y H E Y R Q A K M P E
RANDOM_1048 N A P T G I E D V T L I G K L R L K C S L K V N R Y E V L S D T A K I H G Q G F I V G F V V D F V V W Q P A L Y
RANDOM_1049 V I R F L S F Q V R Q N D Y A T F T N A M V K K H L G V F R N S D V S I L I L W G D F S E A G L T H T F L V S Q
RANDOM_1050 E Y S F S N G S A A S E K Q E V D C G L L L E Y P A V F V I E S L A T T P A P R M I K R L H D P C F Q I T K L W
RANDOM_1051 G S N A K G P N Q A P T I P R D G A G K Q S A Y G W K M Y K A S C A P T K S Q S T M I F N T I Q V T S S K P I N
RANDOM_1052 G V I V N E L A T F Y G V E H D P T C R S V S C I G K A Q F L A P S S I Y G R K M V P F M I N T L M L L V W D G
RANDOM_1053 G V K L L V L K I T G F S V S L P L T Q R Q W C K K A T R K T L V A S I P S D H S I A G N T Y V L T Q L N I R D
RANDOM_1054 S S T V I M V G L N A D V L V G S E L S H S K T R E S P Y T S N Q V N A M F R L K A T N K A D D S H V S Y G R I
RANDOM_1055 A G P V Q D Y E T G P Q G I E G Y M L P L S D Q S V S T C T V L R L T I Q M R L E R I R P A G Q R S T A I L L T
RANDOM_1056 P E Q L S G G P D F E I K H A L G K E S K L N H F L Q G T V N V A L R G S L Q T V M L I R Q L I E M P T N L P P
RANDOM_1057 P Q A P Y D V G R S S S K E I I C S T P Y F M I A H F T L G G D Y L P T T E N D G S F W V T F Y Y N Y I Q L K N
RANDOM_1058 Q A Q V K K T E H R A E A A L L V M Q I G A W L Q V N A R R I A F G E I V L F C A K Q Q L D M M R I N C I M D D
RANDOM_1059 V V A N M C A S K C C V R I R V R R I V V C V G Y N G H V Y E T P D M L A G S V S P G A V L L D V G A R V S V K
RANDOM_1060 K D W S P W L Q V Q V Q I S Q I P L N Y N F E A E V L N Q E P Q G W D M A Y A N A Q G A F L Q K V E P R D L S R
RANDOM_1061 I V C G D K Y P K E D I N W N T P S L K K R L A A Y N H P S P K T A L I A N K K D R Q S V G T G N K I G T A D A
RANDOM_1062 P A Y T T P D S L E M L A A G G L F S T T H D I S Q N L E G L E V A L L E V F A E M C L T T E C K M G Q Q A L V
RANDOM_1063 D F L Y R A N S C N D L D S H G T G F E N S D L M N H S H I Q K Y Q G V S R V G H L G G T D G S F T G I D S Q T
RANDOM_1064 G L V P Q N L C T I E S F R N P I A Q M M A I K H T P K E D H I T R K Y S D Q T F C H N M L C R T I T A F F G L
RANDOM_1065 E G K N L Y T I R I G S A S P H E G T Y N L L L V S E Y T I M E D N G N I A S I E D M G L G S M R P Q E L E R I
RANDOM_1066 L T E D L L K R Q E L I L M H E M E T K F R G N T Y S P S S L D R L L D R Q L V M V K L L D E D F Q F P K L L V
RANDOM_1067 R S K K D F G K D S M E V C A A H I V A Y Y L L S I R R A S N I H H A F Y F M L F C M G L G L M Q D K T A Q F V
RANDOM_1068 N G V G S S V D T Q D Y K Y G S G I L A K E C E L K S K E M A W C Y L D E L S E S T V R I P F E P E K F L E P N
RANDOM_1069 E V C T I A R A D T I V P T F T I L S V L S T R S P P E E D K V D D C A R P F G V H A A S G D D E G G L I D V D
RANDOM_1070 L S Q K L Q R R T W E S D Q D G E L K Q S D V Y R K M G G V E R G V Y V F E S G T A I Y F Y V I T Y G T A G I V
RANDOM_1071 L W E N L T A P S N F L S Q I I P V N G V L G E I K L W E Y K V G G V Q S E V D T N T L V K Q L V Q G G V R I I
RANDOM_1072 I S I L G K G F L C E R D A G K L T P K R R V A E D L T Q Y F F M I E A M E S S C G Y D M S N V I P N S H L S W
RANDOM_1073 M A V L G L S G Y L A P S A A Q D T S C S S S V S F A F M I H T S R D K K A G K R S Y A M A E L I L I K F V D Y
RANDOM_1074 L Q F C I P V F Q N P E P S H R G C P P N L P M V R I G P R S L H D W S Y V S L L P F F R V Q L K G G D F M A G
RANDOM_1075 S T K R M I P R V P Q I G M S R A I L N T T K P E A L G V W H D S V G D Y V S E G W A S D A G A K D T Q H A K Y
RANDOM_1076 G V K A K L V S N G E C F C N A G Q K L A F R L L L D I E L M R K L A L V Y P I I E S V T I K T F F E R N R A S
RANDOM_1077 C C R I A Y A L A I R G E G A V A H V L Q L N L V H E N N E K R N K P V L S Y L K A G Y G G P T E Q D G G T F S
RANDOM_1078 I V L T Q Q E R V M G G R R L T A G T L C G Q K R A S V P M N H G K N N V A A T Y Y H D M A G F L R K G D N T A
RANDOM_1079 H G K G Y D I I L L E D I L S E I L T V G Q S A S K T I Q D D V N R Q E S R H S F R G S K V T R D I G S G C G L
RANDOM_1080 T A E N D Y L F N E W L K P K A Q E P G I N T N V L T L A N A G V R V Q L D N T L L Q P S F N L F G E L V G H G G
RANDOM_1081 A K K I D R S K L S K D P T D D A L Y S I I E K D S F K M N A I V R V K N V F V G L A I I F H V G D G R A E E D
RANDOM_1082 A I L G G Q K L Q N S I K S I M R S Q P I S N N G T A S L A K G E R L D I P C E T Y L S K H P C A V A M P I I I
RANDOM_1083 T R F L V A E E S T R F F K P R V C E K Q A I Y T T V L F P I V P T D H F T H V K G F L M D L I F Q I F Q N S A
RANDOM_1084 T K F Q A Q T D P F I A D F F S F E E P G H G Y K V T K P L S H L I P G R Q T F I R V Q G M V D Q S Q K N R L Q
RANDOM_1085 G G S L L K K T A Y D Y V P T R L Q I N V V S S L K P L H L C A G R F R G G G T A P T P D D V L V E L M P T P N
RANDOM_1086 T F Y F D E D N S N L F M G P A R L V T G L Y E S S P D A H V Y Q T E L E I V Q W T P S Q G Q M V L Q A V L T M
RANDOM_1087 F G T T S I Q Y G S A G F L L A D C Q L A G N S A L D G S G N M E K Y Q F L R A S L V N V I K Y A Q I N V Q A A
RANDOM_1088 S I I P L A C G M K T D I E E C D Q L V D C L T H P Y G G G D I T Q D D K G Q I V V E W T A A S D L W H S C Q V
RANDOM_1089 T R E C T G Y N W R G R S E V K G I V I W P A V V A I S T G R M V L N K V Y H V F Q Q M L G G A Y C P E E I S V
RANDOM_1090 Y S E D V E S Q N S F G A I F A F G G R L L H G V L M K G V P D G R M S G T C L Y C F Q K T E L V E A R P V G T
RANDOM_1091 C N S G T L H D V R R R L V G T V K L C T F A P Q Y K D G P Y G E T A Y L Y R T L I I W E L M M R I K E A G G I
RANDOM_1092 N S D L M M A S R T S A S D T T A N M T R P V R L Q T A G V K L K N D V S G A A S N V S R L I E T G F Q H I A L
RANDOM_1093 R N G I D R I A E T A R H E T K N L T H V W S T E A L E N G V S Q S P L G G R N A P G P E Q V C R E L Q K T E I
RANDOM_1094 K E H Y L K N E Y V G M Y P D L Q Q L A S W D C F V D E A L L T G G G A E L S Q I F L H T T Q G F N N L E R P R
RANDOM_1095 A E N V F D P D G E V P D A V G G E I V P R V Q S G P L G L I D L Y E S R L A I G V M Y P A E L V S I V A C D G
RANDOM_1096 V E E S K V C A R L A F A D T Q V T P P D G K S G W S E T R A H E L F H V C I S A E H S S G A S A A E G S Y L I
RANDOM_1097 M S K N T G N S N T I K N I D S M S Q V V V P N N T G E G A C A Y F H V A E E M E N A S D G A S L A F T C G S G
RANDOM_1098 N A S C D N V L M S T R I C R I V A F S G G V K A V A L P G A F L C P F G L L I K T T P K F G I Q Q R A G A C L
RANDOM_1099 L G A A R V E G Q P Y A L S P W Q K Q K A Y S V V P K G V F F T D G Y E T A A E S V R R P V Q Q L F D S K E D K
RANDOM_1100 W T L N R V F C G T G Y E T L V V E Q T T P E P E S G V G Y Y S G Q W L Q I W E Y I R P V L K L V D A N F R K I
RANDOM_1101 T D G A Q P N L I R Q K S E V R Y Y A G R P K S M D E Q F M G Q S L L E M Y G H Q G I F I T Y Q T W S R E L Y I
RANDOM_1102 K V T K L F T S V F L F V T N R V A I C D M G S H Y A L Q C N A L T Y N A D P V E M K L H A A D R E M D G S T S
RANDOM_1103 K A I R D E V N Q I Q P A Y V D A S I F L G D E R S K P Y G P Y S A M N L I T V T K R P Y I L S E A L A L A D K
RANDOM_1104 Q E F P M T A D S F R G A R F A K S A P T Q A E E I P F P L T I M V M V T I V K V N Q G S G P T A N R K T A Q T
RANDOM_1105 N C L A Q E F N Y P W V K Y P E K N K T G E K V R I F R I R D R L W I H N G V T V E H G E L V K P A E E L I L C
RANDOM_1106 H S G D S G L Y S D S V M D T W P T T N E D E I I L D L W R D T D P L V I L M A I R G G Y A T V L E G T V V G D
RANDOM_1107 S S Y L V E I T K I A H R T W T G R P T P L F Y L G M T S T G G M V S L N N Y V P F Y P A H E A S E F R P C F D
RANDOM_1108 P P P D T F Q L A L K K L L F S L N Q R F L K Y D V K G Y L E E N D L G I V A A G A L Q R K T T V V L V S S L F
RANDOM_1109 G H F V V P G A G L P L E T R S T H K A V D A V K Y P I S L S Q P S K P D R Q Q G S G S A H D L M C A T G T Q L
RANDOM_1110 L A G I I N Q N C T D K N V E V P G F L T V E V T E G I I K T Q E L E S R R T D Y G I T G D P F I V G K F P A T
RANDOM_1111 I G D H S N L W H V S E A N E F G T R D S F A E E A L T K F F E E T T N A K K P Y D T Y L H A S L H K N A G G F
RANDOM_1112 H N F N D M S R P Q C G L G E L E V D V R R D L G H L A Q S G K P E Q M P S K P V V S F S R I D D V T R D R I L
RANDOM_1113 P R K G L A T Q G E T T Q H D Q I R G I D N V K P G Q S N I V V G S T A V L P R V G Y Y F E G Y T L T S N G G A
RANDOM_1114 D V Y H A T Q L Y A E H N Q V S I Y K L F S K R S F V D S Q D S L E N V L W S L P D D C K W Q V A I C S S P P L
RANDOM_1115 C D S L Q M R I R D G L I F W A L S N E W Q I R S S V D D D A L Q P L G V R G N P V K N E S S L G N L E A L E L
RANDOM_1116 N Y N N K N E Q S D P V G C R Q V P K G K T A C T T D P P E T D D A G H D D D S V D S L R F E R E W Y L I Q M S
RANDOM_1117 P M I Y W E N Q L L L Q L G P E V K P G N N D M V L L F P A S I E A E P Y D V F G F I P K W P Q S S P V D K G Q
RANDOM_1118 P A E G T V V K T V L L G G G R S P N I V Q L S P P R D V K L M V P E F M K F S F R L T Y I D K G P L Q E S C G
RANDOM_1119 N I E Q G E Y C I G L C T A D D K G M L Y I R G C L R F E V T S Q G L C P M H P R L A E E L K M C P L A V K R T
RANDOM_1120 I P D R K K K P M L P P M L A E A G Q N G Q K C G P W N Y W Q F T L K V D I Q G K G A K G K I E D F W I G N T D
RANDOM_1121 S A G T G D W T P L A F G D V A S E S I T S K G C Y R D L Y D G T R A G A S G D P V A M V P Y K Y Q E T K A D G
RANDOM_1122 Q N G L E P Q K V K S A T P V P A P Q G T M A T E L E M N V N A K F L E I K R Y R R S V E D G Y C P E I S N N I
RANDOM_1123 M G S S G Q R D V I D I E L G N L L S Q L P T L T L E G D I S C Y G D F E L R D K L Q S Y F I D L H G G S E L T
RANDOM_1124 L L L V V A V E Q K Y M A A I K P C E L K A I A T D T S S T L E W E H R L R R F S L L L R L G Q G V Y S G E L I
RANDOM_1125 D L G W L S Q I S E L P L L N F I R R V A Q P N T S G S S I S W W T I R G G R K T I A Y C E L I I Q E I S E L K

RANDOM_1126 F H L Q T E R S T Q I N G S N F R A F E A G S V C D D G T I D Y T D F C D R Q T R Y I S R S N V K E S S L G Y Y
RANDOM_1127 T K L E T D T Y Q T S P S F A Y K D P P V K H T M K A V R C P A R G L N I D S F G D S S L N R G M K C N N L K K
RANDOM_1128 G R S A P D P G G L T G L R M K M L P G D E V I D Q T T S G R F L N I F F Q S G H Q N T R A S R Y Q F L S E T A
RANDOM_1129 T Q C D A H E T Q L E V P S V M Q P S N L D Q S T S Q I E S V A A I D E L D T S M I K S V A D D S E D V G L T I
RANDOM_1130 H L I R K V D G Q V V V R R F S A C T Y G G L G F S R I G E I D K T G E R L S P T E K P S C S D V S L H A H P G
RANDOM_1131 R S L E L L K S P A K I D C C G H L R E P A A L I L I H H A G P L R N G T A R C P R Y C E C R K T A L G T F E L
RANDOM_1132 M R G E D A T V R Q L N A M M A E S D E G G L S T E K D Y Y A L V I V N L R K D V P V D F A S E S E R V R C F R
RANDOM_1133 G I L K Y E V V F L W T I S S I L K N V L N H T K E F H H L L L S L Q R A L L P E F I L L E N Y E P M F K F F N
RANDOM_1134 N D M Y R E C G Y V T T C R Q L N G A R S A L K Y V E P A R V P F D G V K Y N P A V N H G L N P I R T L A G H S
RANDOM_1135 I L S Y I F D S L R A Y T C W N N G L G K G T N Y S N I H K G V L S Q A K T R L N F Q I T K V N D N K V I K Y F
RANDOM_1136 L S P L R E L T G S M D E C S P R R P W R S A C R V G Q G L D L P H S F E I G T L L C V G A H S M V A K S Y E D
RANDOM_1137 L A L V I G L L D L S N I C G P D D Q K S G F Y Q S E K L K F P L K L C V T L A A T F F I P K D K R M L N E D K
RANDOM_1138 V T R K S L F E I T V T T Y T V I R E S Q I A V A A E I K Y S I A G Y S Q K M I Q C T L G L D R I Q E A W Y W K
RANDOM_1139 F V D H D R L Q A S P L A Q V K K G A G S K D S A F T G N N T P A V Y L I A C A A G I T S K V F L H R T I A M N
RANDOM_1140 A I R I L N T V T V K V T G E R G V L G P F T T Y V D R L E A S K W A H N R K P Q A G P H L D F T A F M E N T G
RANDOM_1141 S C V Q S S P T V E Y V G P A N R P P M E F V S S I L D A N S K S S L S T R W K R A G H E F V V Y K A L V L L Y
RANDOM_1142 E V C L V P V C T K Y Q Q G K V D N G L A L A C L R L L C A Y L K L D R N Y D A Y R C S T K F S S K D D N I N Y
RANDOM_1143 K T L H G V T P P Q L V S S R G W G D R V P V E R Y I L N K E A Q C Q K H G T I V K R Q D T L S W V Y V K R N V
RANDOM_1144 H L V V L V I F V G A P A F T L E V V S K P S Q G V R E E S K K T C L A Y E V M T D L N L L A E T V E L L V F H
RANDOM_1145 R S H S N T S E Y A A D R A T K W K V I L V T L L F R R E R K Q S E L R D V G D F I L S E E L P T L A Q P A A P
RANDOM_1146 L M E P H K L F T K S C L A G A L V P P C D A T K M A T R A A F P T M R P N P D F E E F Y Q F E E F M G L P V C
RANDOM_1147 R V G T Y R P E E L T R E V D P W E Y I R Q S L V A A I L G R Q L S K E G L G E S I P A D V L H A K T T G G T L
RANDOM_1148 L S E C L K F D F A A E K K C P C H T G G K W G T L G V V P T F D S H M G V P A L A R Q S D T G N L P S I W C I
RANDOM_1149 R Q V N R S L L Q A K V R E A D T K P V A D S F L Y P Q N D I E V P K L N S L L R L R R G R Q L D Y T C A A A P
RANDOM_1150 F G A F K L F T L E A T P E C W L F S G L E L S D P E H G G K V V P T D I I N G S M P W Q F V D I S Q A K N G T
RANDOM_1151 L K P G Q D L H T S D I V E E S P T L G L A V S V E A F L E K E K C K K G L K R T A A F S D H L V G R E H H R I
RANDOM_1152 I K V T M H P T T A G V T V S G P H E F T V R G M I A A E G M L T G I F V S V H T L F E S C D L V H A S A A V S
RANDOM_1153 M A L E A F R G K L L V F L K F A I V M S Q S K Q Q E L K G H A V A S A V E E I G S K F D W K Q D K R S G E C Y
RANDOM_1154 S P T Y P E A M E Q D K L Q E G L W L N G I K N L Q N A L C T I E V D P T N S T K R R M V A T D K D H S V K A L
RANDOM_1155 I G D Y M Y S G R I C K R L Q I G K L M A T I G G M W D F Y R I Y A G I V L Q A F P S T A K K N T P S P K S C L
RANDOM_1156 E M G R F S D Q G S H M G N I E P E F Y N G R K I T L L P P N P F D G A Y K T E K T P K R T K N S L C A I Y Y R
RANDOM_1157 P L W E R T I F V S S Y C E L P P W G L S Q Y R S F T N K L S P S R A Y W N F G I G A G T K Q S S S I G I A R Q
RANDOM_1158 G K F E A Y G Y V I K N G S A I Q H K S V V T Q A T L W F G E D Q I D L P L I S R I I A T N S Y S L K D V N Y S
RANDOM_1159 T E G I S I E K F W S N Q L R F Y Q T T S V S A C A G P F R T E K Q L K E E E I Q H F D V V T V S K V V D H M M
RANDOM_1160 G Q I E G I L L F E V P P M C E L Q L L N E I S I T I D E L S T Y P Y S S N S E D I Q K Q V L I Y K T T P R R T
RANDOM_1161 P P P V E A M A K A M E G P P N Y I L D G R Y E D Q L C V R V A S A G L G G L L C V V P W F T S E P Y Y L C E S
RANDOM_1162 I W S F S R L S S S T G L A S L I I I V P S Q K F G S V E T A D M H T Q P V P Q S V F V N A I E I L K P P C M E
RANDOM_1163 P K N G V Y T S P S G C W T A R N R A A P A I D I L L A G S S Q V D Y Y S V K F K K G G D Q A E E L E I R K D G
RANDOM_1164 L M A K F I E Y Y F L L D L T G R V E P I P G Y Q D K A A A E A L V A F N S Q D F K W K E F F S V D I D S Q A N
RANDOM_1165 A L P W L V F F E P S P E R P E W Q V E L S P A V T Q R G D L P F L T T P S S L E V N F L S G G S S L L T P M L
RANDOM_1166 L W G K R K Y T P A F S S Q V Y P A V I Y I A L M P E N T F R C K K L L Y V P T Y A D F Q Y K D T G I A P P T V
RANDOM_1167 R M Q D H N R Q P E R Q I E P W G G R A E C V H K T Y I G H Q T I K W S K K R L G R V A Q P R D F K A E Y S M Q
RANDOM_1168 M Q I N E C I R K Q A E G T D W W T K F K W Q V W V K Y P L T G E F L L D P F D E T Q V Q L D P I A F F A A C S
RANDOM_1169 G A N R W P P G F L V P T E S L W L L A W C E S L A Y R Q H D M V Y I L S N C G D V V S T T I I P G D I S T L S
RANDOM_1170 L P G L Q V K L R F C Q L V L V V R K K L E D I T R C D G T E Y P T G H H P N N E M I V V P N L A E N Y R E T A
RANDOM_1171 T K I C A T T V Q S L G F R T L G P M P E A L E N I M S K S Q H R V N T N G V V A L V W Y A K D S N L E P F G P
RANDOM_1172 S T S G A K F N G R I L L Q T S I I P R Q G E P T P A S S S S Q C L G S A K G E H C A V V R V L P W E R Q K F V
RANDOM_1173 Q G F L D R F A N P T E S A N P D C L H A H Y P F E G Q K E T D V Y V E L G L A E L L F P S R Y T Q F S F P A S
RANDOM_1174 S A L N R K A D Y P L A R P L G Q E Q D R L V A F R S E H E Y Y A L T V N T K R P L S N Y D S H A S K G H L P I
RANDOM_1175 Q S P Q Q S E E F T E D A S L N C A D G G E T L K H C R M H V G I V V I A L Y T S V P G P P Q G N T A L T E G R
RANDOM_1176 F T A K A S I A K L M F C F L A K P D R G K A V C E R A K T R V V E K V I S R A L T A L N S E A R I E I S C T E
RANDOM_1177 N D E L D P K D T F D V A A R G E L N L N S Q M D P R V A F S V P M D A E A N P Q G R V Q A D G T G P T K K R L
RANDOM_1178 V N T G R S E K E T E Y E C R T D F R R V E W L V E P D P T S L N R G E A P G G V L Y K L A A N G N A A M G V G
RANDOM_1179 E G S Q D A W D Q Y E H A T R F G I N E P I N V W P F Y T Q W T Y V I P H L I D A G D A Y V G S N N N R D A K T
RANDOM_1180 Q A I S S D D S S W Q S E G L N Q Q S N S R E T I S A S L E M R D E D A L E R A A L T L M R M V A T A H T E A K
RANDOM_1181 L Q Q L V L F F Q V Q R D N Q L Y I L S N V A I F S A N A P A D G Y A N T G Y P M V L K L G L V L K V D L E A L
RANDOM_1182 F L L A A L Y I I K S P K E P Y E L E M M F K A F I I E S K C E T K Q P D R I V S E M G S L F T P H T F I A G S
RANDOM_1183 V S A K I S K A S L L D P S L S L V T G G C H G N A L T A G G S S T E Y M H R S T L R G N K S N P G G S P I K T
RANDOM_1184 S P D V M V V G S S F R E F S V S L L K A P F Q L I I D V A R L L A K E H T C P L Y N P Y L Q L I C F S S K Y E
RANDOM_1185 K Q D A Q S N R L A R E Q L G Y S V K G N M P P V P Q T G V T N L Y D I K L A E M F S S L T Y R F D I G H F P G
RANDOM_1186 I E N M L V K K S D G G V R D R N L L G S R N A D S S Q V L T K P R P L W S A N K K R P V W E S S G G C S Y P P
RANDOM_1187 L L P C A L P P H S K W H P G T V L R K N S D A R P A R L G W S D F D G G E L V R T E T F L R S P G G G I L A R
RANDOM_1188 A E E V Y A L D G Q S A Q A N G L S S L D G V Y D S P F E F N H C T F A I R N E G G P L I M P Q A L G M L V P Y
RANDOM_1189 K E R D G K P Q L Q L V P S T P Q T H V S I W S A L G K Y T L Q G D K L Q V A L V T N S P L F N T D A L F R N E
RANDOM_1190 L L T S Q K S G K S S L T T S V M G S I L P K E N T F R G S Y D S Y H A K S Y I R A R A G I Q C M A L R T A M L
RANDOM_1191 L A T I F M F D A E L N R L Q N G I S W D N T G G D M Q L F V W L G H N T N K D F N K V F Q I V V I S S Q V T S
RANDOM_1192 G V I S L I D Q T R E S E M A Q V Q A K F Q R P D S C R M Q H L V L Q L N Q I S H G N A P D V R Q K S K A A F M
RANDOM_1193 S L Y V L S I V F V K P V R L H N H L E A G H G T Y S K S A F E A L G R L Y S I R P G D P K D A Y R S K G I C F
RANDOM_1194 T P E L G Q F R G G P P G Q W F K Y S R E H Q S G A K L D L K V I L V L F P V N V F V A K N H F R Y N C L T N T
RANDOM_1195 S G D V L G V F S V G Q H N D K P A N F L K G K A H V Q L V E P V Y G V V T G G S Q L S S L S E T K G S T R K V
RANDOM_1196 G E Q I Y P E A C H N A E A L K I C L A E G V C L K H S Y Q D D D V C E L F Y G A Y I K G I L I P A F A D G S I
RANDOM_1197 M P L E E T P K V Q R A E V Q E D L F R V Y P S L R P D V D A Q S R R E F A V D K V W S G L K S C K A H F V C D
RANDOM_1198 D I V E R S N A V C R Q D L L G Q S L E V M D Q G D E K N S S I V G A K P G V L C V P A G S K I L T D M Y I A Q
RANDOM_1199 T M L D P K V M V T L M S K A F E R G D Y S I V A E F I L G Y K C D P S I V F Q K A G D G R K K T G S L P D M E
RANDOM_1200 P L T N N K C R N G R Y L V T M G A K Y S W R I S E K Y T W Q G T Y S V R Y L Q G I T R L E A R A E Q N P L L E
RANDOM_1201 P D D K T L Y K V L F E I A A Y T L V M P K V K E G A S H V V G L K G G C N D K V F K M L W C K T P K A E I G K
RANDOM_1202 V S R K L E Q E P T P P S V F V I F E S T D F L H W A T E F A L E L N S V D I E N S G M L G R T V R Q L T R I T
RANDOM_1203 E Y R L S A N V W T A H V K D T I P L V M N L T P H S T G S C D D G T L Q R Q E R A G N Q G P G G R L L I C W G
RANDOM_1204 N D F F Q N M V G T F G A T T T D H A V D L L A E A F R L A R K E D S E E D V I T S I A Q P P P H P E A P K S T
RANDOM_1205 V P A Q S G L K G L A R K A R R K R K E G P P L E I P A K D V R Q L D R S D K K A W A G M E A C T N L D R F L L
RANDOM_1206 G F A T L D S T L G H L K P L N L Y I F C L E I F R L K L I W A S G G K P A W L A I R G F F D Q P D P Y I A N A
RANDOM_1207 A G P T R I N K L F T P E P A S L T T A G K D P S G C I G R M N F F D F F N E G A R C S E V E L V G A A M V F L
RANDOM_1208 D H A H A F Y F L A R E N Y L A G L R V L I N L L P K E T L E I Q I L L L P I V L L F A E I D Q C I F V L A S D
RANDOM_1209 E D F S R T D G Q W M S V N E E T T R E G C V F R E V V H G L L S R T Y S E E L P A L T V R P V R L T D C N G D
RANDOM_1210 Q E F R N F L E A L N D V T L A K D T F H I Y L G M P H E R P P R C V G R V E R W E V T E K T A R S S T G V P E
RANDOM_1211 C F P S K I L R D K P E R E F A A P D M V R A K L N V G S D Q A L E K K K P K S S I A V M C G Y G T Y C A G G P
RANDOM_1212 E K V S V Y V V V M W C N F P F K I A K A D L A G W M V Y D E M G S T T K L V E W F V V N Q A S E M T A T D D S
RANDOM_1213 R L F E R E G D N N L C M R I E R P K G V N W D S S H L P M I I E S F M F I G F Q I L H I D T S E N L L D G I A
RANDOM_1214 G D E F P P I T F L G P A V E V F P A L R G Q E A E L I A V T N S K S A R K K A K M S E F C A V T H R Q L A N S
RANDOM_1215 F D D A Q L A T K F Q L P D Q P A T V A S P D L C V Q R V Y D H A C V L I V E E R A D G D E K M L Q G L K Q Q N
RANDOM_1216 I D S Y R V G F S H R M G E A G T N F N F Y N T K Q E A M L S D G C G E R H H A Q F Q Q D L T T Q S A P A Q W T
RANDOM_1217 Y F Y I Q V M R K I A A T V E E F L N V D Q R F E L R N P R Q D M C E R E G E G P K Q S Y Q I V E L L Q M K Y T
RANDOM_1218 P M E T W F G Y H E P L F A T E P V R N D D R R L I I E V M D E G R L I K P E V I S A A P D D D F E H H S C L M
RANDOM_1219 P E A S P Y L I D W A C T A M H H P L A Y E G S H G N Y L L N F W H M I P M L S T P S G S E S I I V T P F R V R
RANDOM_1220 S A R K R T E I H T K D D G Y F I Q S A A T K E G D E S D R A Q V R A L N S S A V G A E G L D K S Q L G V R L S

RANDOM_1221 K R L E L G K S V S H A G V M A F D T R N L K T G K G F V G H C W I E W E Q L I N K A F V A M E L S T A R V G L
RANDOM_1222 D H T K I G G S R T P G T Y L F P M H E W A D A L R P D R P C G P C L G A L P I N G Y A D L G G R V L P P T L G
RANDOM_1223 K R L M N I A V Q S K R Y V I W A D S D Q D T W I F Y Q M G R V E T Y S N S V G E A H M Y A S P N G V R Y S V F
RANDOM_1224 F F I I V Q N L R T V D P F E R Q S V P G R C S G V R I F T L G S Q F V T F Q T T I P R G F E G S N T M K Y P K
RANDOM_1225 L Y G R W G E N S E N S P E P I G P A I T S N T L Y E L K P P S N S T K T K Y C V P G T V D I G T N D V L V K R
RANDOM_1226 Y P L V K A D Q T K K R A K Q D Q L H C A K I R N Y A Q K F V Q E T E I Q C Y N R W S T H T Q G P V Q P P L S L
RANDOM_1227 K A L M W G E F V L S E P R A R Q P G L A Q G K K E T Y F N D S W I A I H P D H I I L P T F K L K G I F N E Q N
RANDOM_1228 L A T H E I N I R A L S S A D Y V A N M N K N F D F I H V R R E T F E F S A T P W E D A A L S L R G A I L K P L
RANDOM_1229 A A K N N I V A A I S L P K V P V R F L I M I D H L H D S A L E S S G S E K V V L L R A T A E P D I D E M T L N
RANDOM_1230 D H R V D Y P I L L S V Q A G T D E D E A A S V G I L D K L I C T Y L Y V Y L P T Q E E F D V N V V T G R L L C
RANDOM_1231 L I D M S K D R E Y E T E M T T P F I Y P E L L K Y S F Q W Y R V T S D L A E P F Q T I W K P E V V T T V A S Q
RANDOM_1232 Q E G T A T Y F Q V G V D L Q T T F S Q V S R Q R T D L R E L L L P G M V V D V T I P L Q I L G K N H S I G I E
RANDOM_1233 T F S E T L A R G V H E Q G P G C P E A L S M R S K Q L A D H Y H T G P A I L F E A K P F L C T G V F P H R I Q
RANDOM_1234 N V F A P K D Q V F Q Q A Q F K P V Q K Q H E L Q E T T K T F Q S D S L R K A E A V P Q S W S T A G Q T Q G E V
RANDOM_1235 A I K Q G S G Y L Y G L I W D E A T T I F P E L A R D I L E A D P V D E S P L L V Y N L I K L I G R E A S E I E
RANDOM_1236 E L E I G D S T F R P D S V V V F A N G Q L L W V K V S V P D Q G T Y G T N A Y L N Q I T I V S T L A E F F I Y
RANDOM_1237 A I E L H N L A V N I T T V L C N K L Q L V Q G P D A V V H I Y T V S A V L N W F A D T T G H K K G I C I A V S
RANDOM_1238 S T E S F D F A N T R V P I Y N L I K I S P T F D F V F A D E Q G P A S G P L E E P S P I G Q A E T F W N N I K
RANDOM_1239 F D G S D L N I A Q F M F G C L W N V T T E F F D L D K K F G L D Q L F E W E Y R P W G L E K L Q G V F V S D P
RANDOM_1240 R D L D W A C E D R A F A P G H G D L T I N E K G H T P A V H G G P Y W L R I Y M T Y Q L K T L L K L A K K I K
RANDOM_1241 G K D I T S Q A S T A P M L W A D R E D G L G E K S C C N W A I R E S Y F Q P Q A P E K D Q C I G L L E E L S Q
RANDOM_1242 R I Q A R N G G L L P L G Y I P W V K D T H I D G A G M G A S A R D V V V F Y Q K A I E R G R A A R Y S H A A K
RANDOM_1243 T E R L T R I V C S Y Y G V G L V Q I P I S G E F L Y E Q V L D V S I S S L M T H I P V P Q P D D D G Q A T Y S
RANDOM_1244 I D G A R S V Q D L L F K S S N I D L D V V A Q K I K E K A A C N S L C N D I P T M E I I V N M I L K I M F K A
RANDOM_1245 P Y F G I L V A E V W T N Q L R V F G K Y P S A A G S A C E V N L S I E G I K A D H R E P L V L N H R Q E A G S
RANDOM_1246 C S F G I G F Q P K F Q Q C G C Y I P G E V S K L I S K G V S P K Q V S F Y N C D M L D P C L P P E H T N T A A
RANDOM_1247 V D E V Y A R Y N L Y E T A P E T P K I G N C E T F G H G I F G C A V T C S A S Q S P R L A K D N Y Y T D I G H
RANDOM_1248 S L V T K T I G V F T T G T K I E Y V D L R G A L P N A L N L W F F W W E L K E L R Y T Q I G A A F T F V M Y V
RANDOM_1249 E G T D A S L S A A T A L E S F T I T S P A Q R V R N R D F Q V P T L S S Y D L G R T I N I Y K V C N N M S W T
RANDOM_1250 M V V E L E P C Q P F R D W E K S E G K T Q G Y P V A T I S N E G F P C T R A S T C E K A A D K Q A W P P S P A
RANDOM_1251 A L D P A W C S E K Y V M T S F E L Q L N W E D M A K Q A C G R D L V H T L A V V S P A S L T K E F E N L K D R
RANDOM_1252 H K Y E W E D A A V L Y Q A A R C K R V A E M T S G T R L D G E N A P V I R Q S D H Y E T P S Q L T K S L W E Q
RANDOM_1253 P N F L S A G F R Q E F I I T G A Y A F I H L T E D E Y D W T N N Q S G P G T V T K N K P D V E R F I L V E A S
RANDOM_1254 Y L Q G W H C D N T S L N R W E V L Q S A L H A E I C Q P S T Q L H S D Q Y E R K L S S P D T W W L Y Q G S D V
RANDOM_1255 C G I N L S N D N N I F S V T S S F G T Y T T S T L C V C I P R S R Q Y H V V S P S D I V G H L S P N R T Y D Q
RANDOM_1256 S D E G C V S A Y D A R R A G G V M A S G Q E C R V P T V R L F G K V L G V N V R A T E P G V I L V G F S A G A
RANDOM_1257 N A T F G T A K L T D D K R L I T L V L L L Y T V C K T S Y K S M Q S L R G V T N T L S F N L S P N I I F E K F
RANDOM_1258 Y I W H N K E D L I E L R P V A E N P D N P A T K Q S I T V Y F T F I D A K A A L K A W L C S N E D Q T G K W T
RANDOM_1259 G S I Q L I E L T G D L M L E A S I T Y Y A L T R D R L V S T G L I E A T G Y V N A S P T R A D D L M L L L G I
RANDOM_1260 G L I I K D T P R Y E K H R G T A T H S G T R F S D K L C L G S G N R N I G P A F F I T G P L Q P E P L G H S F
RANDOM_1261 G I N G A N V P A R A R E Q S R K A D L S V V P S P V N G T D E D R T N P V P T A G C D T Q N R S R N T E Y D L
RANDOM_1262 Y Q M G E W Y Q F D F R S N E P S A R R G Y Q T C L A P Q Q L S D K V G K E L D T M S P R G H G P S G V S N R L
RANDOM_1263 A H Q K A C T R T Y A L L R H S A D W S A L Q A L K R F A F L G S V P Y V D E Q Q K V G T R N A W G S K L N P L
RANDOM_1264 G V E E S G K T K I F E I L T E R E I G A P S E P K Q A N C G Y Y G G V P E I H M M P D M T D I A S T P K L L R
RANDOM_1265 L E D T W L A I A I E R S K E V G K F F R Y H F K L C K L L Q T Q M K L Y D P T D A K A F Q A S R W T Q K V P T
RANDOM_1266 V H P N A E M Q F S D K L A T D A A K E H N R G D T N P K Q W L H M E K Q Q K W I N D G S V V R L Q I R L A N T
RANDOM_1267 P L F S A A G I T A K S E N C R S A C W F S V F E V G I A A S N L G Q F V S L G S S H R T G G D Q W T V T K A Q
RANDOM_1268 E L D I K V V V E I A D N Y K P G I F L R T A D A S T K D E I K K C G P L C L A E S Y F L M L V D N Q R Y A P F
RANDOM_1269 P I Y N E R F F K S I K L Y N L G L Y E R T K W L T N M M L F G S F R V Q I V L V L L L R A T V N S V D V G A K
RANDOM_1270 K P Q R G D G T S Q M G I I Y M F Q S K A Y G N P Y D A T F W L A K V L P A G S R H R V H Q G D F D C I N S H G
RANDOM_1271 L A S R T R S E P G I N H W P V T F K F G V N E F G R N L F V F F P F F F H Q K V F E T N T S Q T S V G N A F R
RANDOM_1272 M K K I Q G Y P P D F E Y T L L H S N G G P S K E P T E P A K M K E K S K T R T A L R D N P V W Q D P A A T G L
RANDOM_1273 G Q C P I R D G R V L G E K G D K A N H T S R K D P H V R L N L R S T Q S P L S A E G L W Y Q L C M W N S L A K
RANDOM_1274 F G E A T F Q L A D I V T K E Q E E T L H M H M L K C A L E S F D N P A S S T V S A N S R T D R I M G V R Q G I
RANDOM_1275 R A Q L I F K E W S N Y V L L K K C A A F K P S N E E L A Q L S S R D H L T L K I I K Y V S L P Q Y N T R P L C
RANDOM_1276 I L E V T Y F C D S Q G V D L W Q G T I K S P Q L Q A V T Q V Y V A F L S V A I L S L V I V D S V E V V G I D D
RANDOM_1277 A M Y R D K M F M R V E P A D V S A E N Y K A L F K N L A N Y A K S L E I I T G E A L K E A H A C E R L N P K E
RANDOM_1278 I D G I W R D G V L A D R L V K I T L L E C Q V C D H N L L L N S I C S E E P K S L I S R A R D R A S K Q Q V A
RANDOM_1279 K Q A V V G V A L T L A K G P K K Q G E A D L S S V L A L F S L E S K A K Q T Y L F S Y E F S L M T L L E V I A
RANDOM_1280 V T Q D L T Q N G K G F W N D K Q L V G G V I S V D G I C P T E F P S S I K P V V E M L E N V D E G A S A W M E
RANDOM_1281 P K T P V D D S A F I E R W G S H L D E L C Y D A G M V L F G I L S S N P G P A A P N D A E K S Q A E Q L R I A
RANDOM_1282 Y W R K K L I C V G M R S Y N G C T V Y T Q N I S I A A K V N A C E G L T I F R C F A I D M V S A R R D A Q Q T
RANDOM_1283 I T V A V A A T A H A K M I C Q D L T T K T A W E A M T V F G A K N E A K A L R E M G S V G A A I K R N K I P N
RANDOM_1284 G R I L G G I A I D G Q N D K I S T I N I N T T C G K F V A S F S R D R D Y L L S V S I K P L A I T Y L P S E N
RANDOM_1285 S E R Q S A K P K V K S F D A G N H T K S S Q V N N A A K N K D E C S D S T I E M I H T A G V Y N R D R E V Q F H
RANDOM_1286 P T G A N A A A Y D I L N V L C C E V G E L F A E C I Y D A A Y R E F N S V G L G V P Q D E M I T L C E S F R Q
RANDOM_1287 S V F L K M P V S M D G H F Q D T N A A K Q V E A A S A Y N A N Q Q E K A I N G L P G D Y V P E I T G I V V L D
RANDOM_1288 L H S V T R I F R A V G Q T T G W T V G N S C V H S R Q A E N V F I S F G Q L P S T V K V D A T K F A L A T S A
RANDOM_1289 I G W S L T K P R N K L R P G N G E P E G C Q Q A M N A L L R N Y T K L D Y V N E K A L A E A T R D V I E R G G
RANDOM_1290 A A F I K H G F K D S E V D V R N Q S D Y A P N Q P T W T S P A Q E A P I R M S I A C K E R I S A P A L K Y D V
RANDOM_1291 Y V V G I Q R P K V A L M E Y P A A D D V Y S Y A S K D Q L D R M G A Q A D F I L D S I L D T L A N D V A E I A
RANDOM_1292 G E S T P D N E M V I F N L T K P V M F I Q Q A L L A E M G I L L R W L C V E I A A N P S K I A E A I R A N E L
RANDOM_1293 E E K E C Y G T S V L G S K G L K R K I E A H I Q C F P I R R N L Q L D F E D S I G S I H S F H L D L T N R T S
RANDOM_1294 L E I M P C I N Q L H L T V V E Q V L L K V G S G E D Q Y M N Q W D L S V Y A T R G D Q L R P R V S S I G I K L
RANDOM_1295 L S M Q I E I K D S E R T S A E N R I A V K M R W V V G V Y G H A L Y T A Y G M K E N E S E R Y P K T S Y G H N
RANDOM_1296 E K V D M H V I L E L G R H F K L F L E D V T A N E Q L S L L T D W N N D D S D G F V K V L I V T V T V S R R W
RANDOM_1297 R L I D L V E Q L E K N E A T A T H E S F A M E D K R S F K L I N Y E W Y N K G S H Y I R A D T P N G N L I E L
RANDOM_1298 M Y F T S H G S K Y R E P D K V F D V G N G R V K A C V A N C S L I L A I E L E N I F P F C D P Y H S A D T S T
RANDOM_1299 S P V S K K K G R D S I F F I K A V T P V K S Y S N L K G G A Q I E R L Y G E S L R Q K A S D W Q L L A F E A G
RANDOM_1300 G L N K K A P D S H A T G K W S F E V L D D S N A G P G V T V V A G A K R D T T G M D A T G G M E K T L F E K G
RANDOM_1301 K L L Y G H E V V D R A Y L S W A G E M D F N L K Q Q T L S R I E Y I C H G Y L F N V L T A Y S N A I L S A N K
RANDOM_1302 P L P R T T T T S V F G T K G P I P T L K G Q L F Q N W A Y A A I S V H L V T I D N A I N G E L N W Q K L G P G
RANDOM_1303 A R I T D S P A G V H K S E M A Q C L N Q G T V A V L G V S R T L S P E T S W N G R T V N I L D F R A F S L I G
RANDOM_1304 I L G V Q L L G A I Q Q W K E S E C T L R L I G D S I S N R N Q G G V S D M K K D S H S G P I A P T D W Q I G N
RANDOM_1305 D V C C T V L C C D L E N C S A I T R G A A L P L S A V A I A Y E E D F P A A L G T E N V S Q P A R N H D G D I
RANDOM_1306 Y K I C M V A M Y E K L M W I A L L E V S N L W S E E I S D E C K R C N F I N F G K A T A E S G R C L S R M E E
RANDOM_1307 Q K I K V S N G F M T D S R N V E G A R D T H F Y A L R D A Q D T L G M Y N E V P A F Q G Q Y V N E Y T S V A N
RANDOM_1308 S Y N A I K T D D V Q T Y T E P M G A K D V Q E W Q I T L P G P A A L Q K I N A V I A R N G S C E G H L P S G E
RANDOM_1309 A E W G N N L S I T K S Y K K R P I Y G T A I E L R G P E I V D N K I V Q P P L T W I V L Y P G A I D G T K G S
RANDOM_1310 C H R E Y T T I V V W R A F I L E A G V I D P K L G R N G R R G S A V R S K R A K N P C A I S A M T C A S D R M
RANDOM_1311 Y P S G G K Q N I L A T F V Q V S R V E H G E P F T V N S I C N I A D T H S L V A S N G G I P N D E L R S P W P
RANDOM_1312 Q E H L N G F R I T Q F D F W Y L L M A Y D K Y M A F C M C N T R H A S S S R P N D D P K G D V V W G G I D E Y
RANDOM_1313 R F A K L H F Q I Q G N K M L R K D E T N L G Q Q L K D M N C F R I G H Y G E L T V N I E P V A T K L S Q S S Y
RANDOM_1314 M S R F K Y N M T L K E K I A F P G K P A Y V I I Q R L G P P G G S E H D G D S E I K T P Q C E P S L L T L L S
RANDOM_1315 R K C H H P G T L Y F D P M L A Q Y L D T D S P K N R A P V S V T L W A F P F Y T V P E I K G Q L T T L F P V R

RANDOM_1316 K N H I A M T G T E G L I R K V D Q G D F S D Y V E E E K L G V E P S G H V I V D F R R I F V V H K K A V L P E
RANDOM_1317 G Q G E Y Y V Q D I M K F M A I D D L F L L A A N R N M R G R T R E N Q S K A H W E V E V D L D Q L A M G H T S
RANDOM_1318 N L R R G V R V A A L S T P L R Q T Q F T W Q A T Q A G V W T C S I V V G A F I Q G E P E T C K V R T F V P K P
RANDOM_1319 S A C L G S E P V Q D N E K G A L A S D G G G T G A T E Q P N V S K V E D A A K S S I W M E G L S E N R P N D M
RANDOM_1320 I Y V D R K D N P Q L T S S N T P S V A N Y I T N T D I Q V D N L V T N I A A Q L V G L I A A H Q V G A E L T L
RANDOM_1321 A F R V V F S D I L V A W Q Q V Q T P T I Y G S G A V L G W L W L M R K E V S R D N N F L A G A Q F D D G Y V A
RANDOM_1322 R R Y L W Y N S T F H E Y G L R Q C E D P F A N L K V Q A D F C D H E N L A K K A G W L T I N S T S A D P P D V
RANDOM_1323 Y D A R F D N S S L Y L K M L M L Q L T Y S I G Q G W N G G G S Y K N D L E K N K D G L R T I I T E S V G C L W
RANDOM_1324 A D R A L C G A S H N R S F T S S P V A P E I P R V L A V A S Y Q E A N Q N L L K R K K H T M R Q E G S Q F Q E
RANDOM_1325 P V T V I G K S K G N G S S Q M K D S G H D N G P G G D L K E G A S N G F V T T A A M V G L A G Q V L V C E L S
RANDOM_1326 L A L I P G F H A P G G Y S V L F Q H A L H D S Q Q L F I S D H E A D D Q G G N F C Q D N S M K A L R A V L H E
RANDOM_1327 L V I L P G D P N A P V G L T K W K T A A H L G A D L G A Q Y M T F F D C N L D F G C E S G S L S Q V L E D L L
RANDOM_1328 L T K N P N Q A T L N G A I S S V P S A G L I G A D A S F K R P D R D M H G G R G E Q P Q I T V C I G R G C D H
RANDOM_1329 Q E L E S G E W N S I H G P E V E A K P E T F N L N I K R L P A P V S S Q F K G K L L A Y F C E V I E W L Y I L
RANDOM_1330 I T L L F N Y A Q C K N V P V M G S T G A V G F G A V S E Y G R E P A D K G A R M Q L L F N T G N H V G P N M S
RANDOM_1331 D A C P K S M E L V V P T G D A E G N D H Q L A S W K G I F E G R T K Q L H S D D A S I T Y N A V L H L S N S D
RANDOM_1332 K E S I P V S E G D E T I K Q P A L Q E E Y E M S G D L S A P T H F Y T I K D H Q M D I D G N I M A A F L L D H
RANDOM_1333 T N F T A E T Y H M Y K Y V G S N D D R L V D Y N I T P A I V V A A C L H F F I D R E T K G H V R P A L E S N L
RANDOM_1334 W Y K L I L G S L E G S R N D G S L T M K E R L E S G Y L L I F D C Y P N R S G L S A S F L G Q I P G K G M E P
RANDOM_1335 T V A S A D L L S S I K E K K P A L C S V G F E A N P R F V V A T C L G G V E I W K L P L F D A L S I A L K H S
RANDOM_1336 V K D R D V E Y K S G P Y G A I I V N K C Q G G A V C E R R L A Q T N T E A R F W K Y P E P P S I K M K N Q N I
RANDOM_1337 A D A I L D V V A L P L N Y P Y T S S I G W L F V R P F T A G D A I P G S D A M D I I A T P D N V A W T T L I K
RANDOM_1338 E I A A V Q K A R Y I L R P Y Q W G L T D R H I Q E Y L D Q T G P M T S D N K L T G D F P T M Y A S L M K S E I
RANDOM_1339 E M F L Q M M L F K E S Y L V E Q V Q I L R D E Y Y R D A M D L H G Q T V F Q F I F S R T R G L E E R T K D K P
RANDOM_1340 N T K L K S V T V R N T Q E E W V P V V L I E S A M S E Y I T P W L E E T M A G L S Q K V R T L Q L Y V S E K E
RANDOM_1341 T T L T P S I V S I I W F Y N T T F H S L S I C D P E L C N N L D K Q K P Q C P L A G D I A D L E V T A P H W Y
RANDOM_1342 S P I E G R I F A I G Q L W R S I S K K L Q L F I F H N T K D G D E Y L R D E S N G L T L S Q I E H V G N L G E
RANDOM_1343 T S Q N Q D S I V M L S Y F I K G I I T A L L A Q E H G E Y A D E T P K R D L V T T F F M T N D T E W E I G G S
RANDOM_1344 Y P L P G L M G I N D V D Y I L D P L P V L I A A N T V T V S S H Y K T E V L A W L N F L Y V P Y G D L P L R A
RANDOM_1345 N R W K I A R R I Q K A K V S S P V I G G T Q L K Q R R N G G T V M T L V L V R K M C G L L K S A E I S T V Y P
RANDOM_1346 C K K E Q G I K S F H S L Q P R Y T S E V I T N Q M L A L E S H E P D I G T V M G N V E H Q G G D L P V V S I Y
RANDOM_1347 N K D S G I Y D F N D L S K E F D V L G S G Y Y Y A V L F D P T S K T I V P L L F P V H E Y S L L N A V N C L V
RANDOM_1348 A S V R G V Q N R L Y P D V N L D E L Y T I K G L A G M S D W A R Q E R P A K T C N S Q G A V L E I R N T A T Q
RANDOM_1349 V I G P F I A T L T E T G I Y E A C P T Y E P F T E H T A R V N S K G M V E A K A M D S T A G D P T C N L R V L
RANDOM_1350 F G A R R Q W E F F S F D G L K Q E G E R L A D A E E F K D E Y E S E T H D L F C V D A D S M T F L Q A N F N K
RANDOM_1351 F H P E N L A P T L A V V A L R Y R Y W K P T T D M A H P E K E M L L E V G P N N S V V P I R C A E S Q K G K C
RANDOM_1352 K S V V Q A I N D M S V A S K T Q K S H Y G A A S Q Q A D H S P S S P F T P K P G R V F A S L G F N T E V R L R
RANDOM_1353 L P F E Q P I V F S R V G V I R A G P G T C V D G N S L T A P E F S Y G D S V T I P S T L K V V Q P F C G G L L
RANDOM_1354 L A Y K K Q S L I K E F R A Q F N N L K N V P A E E R L R F G K K G L G A K K L G G K A A N H T A V Q G Q A S R
RANDOM_1355 Q W V F N L Q G L A L M A E L V I K T L A I T K Y S L G A D F T F F L G N K D P G T A D E K T K H A D P A Y V E
RANDOM_1356 A V T F Y K L K K A S G D E V T G G Y E L V E F S P V G I A S W L Y A L K N R M D H P G I Q F P T V T N L T L I
RANDOM_1357 G E M L L P I N A L G Y A V K D C K K K P W E S A A E L R A N F P P A I T F L G A E G K N W F S L Y P I L K L S
RANDOM_1358 L K P T R S H A R E W P S L T I H C P D G G L K L P Q L V V L C K I L E S H I I K G R V Y Y T T G Y S Q E D I K
RANDOM_1359 P R S K L A G L A G L Q D N N A V A G K K R P K D G R E T D R A A D R S S E P P D H G S I Y P F F T T T C G L L
RANDOM_1360 A F L Q G S A Q T T R G Q R T R I R V P R V S A R S Y G W A L S C F P G L S F E Y F Q R T G I S T E V S M I A F
RANDOM_1361 S L L Q I V M A T S Q G S S V G V I T T S A R E R G G M N V F W E H K L G Q K I K I S R H G C D F A V L G G L Y
RANDOM_1362 Q V C D R R W D G L T H K L G R T H K Y A Q Y Y N E N I N K S P E G G P R R V R F T L E E A A R V T C T V V I C
RANDOM_1363 L T S L H I N T S M R S Y N P D H N A E P I R V Q H A G V V S S E F N E F Q A V S R H V R N G P G N T A V I P E
RANDOM_1364 P Q S T H V E S I L R G V R G A G Q V S E I L K G P L A G A N G V N T Y A A T K L I A M S A G A A L S F A D E P
RANDOM_1365 S Q V R D S F H T M K L T G T I V G L G H L M G R H R F I D L L S K Q N I D N S N K T S E L L M G E D G S E G L
RANDOM_1366 A T I G T A E E P L K I S G Q V L N V I A K K S L I S A F L T F R V T F D F D H A H T T Q V N N R I I G V I T A
RANDOM_1367 M F K Y A G E K F F W G I Q F V A L S Q L A N I E G L L H A F Y A V I E D A L S T A S R R P T N G K S I V E G Y
RANDOM_1368 G R D S P T S A G N I K Q A F L D R N S L E T W D G V D S P A V T S D T R V V E M N V Y G H V Q L V K N R C S L
RANDOM_1369 K A H N T L M N A A I Q E Q G C Q V I T L I G E A I F K K A Q Q E S T K P L G K I Q L E A Y V T S D N G E A V C
RANDOM_1370 R A P F V A N S S C D P W T S D V V F S G P P L T A R F T T R D Q F Y D K I E S R G Q L D R Y R P Y L G L L F G
RANDOM_1371 D Q H E G V A N T P S L K Q D Y E S D C V T M D A Y L K I T L E P V A S M T D E T R L Q I G G I N T E L E F T H
RANDOM_1372 I A P N K V V S T G N T D K I P G Q N L Y H Q F Q A I T V Q G M T Y Y E E A N P A E N A V E C H T V H S S V T P
RANDOM_1373 P G L W L E R T P K A R A L P I Q A E S T E A G A S I D D K A L N Q A E F R F V L Y S I S G N S D N N D G V T K
RANDOM_1374 D F V S D I A T T S Q T K N L Y S N T P F L I W P N K Y V S K G T A G K S D T S E I S G Y H G V S A G Y T R D T
RANDOM_1375 P A A P C Q K K K V T D A H R L M K E M L L V K D N Q P S T K F T F Y L T Y M M Y A F F Q R Y R D K V E Y T M Y
RANDOM_1376 G S F I G L R P Y L S I I N R D Q R S S S P T P E S A A A K H T C Q L C G F E E N L Q K F D K A E N T F T V E M
RANDOM_1377 L T A P N S T S D V H K G E L Q S R Q G P A L H T P P F G I K F E T Y N T Y L A A Q S K T H M G D N L Y T C G K
RANDOM_1378 D Q I L W S L W P C V Q G M K K G V L P H K L T G L T N T Q L R K V L T N L K V G L T T H K P Q N L V Q L I I L
RANDOM_1379 V D K N N Y L V P R H E I G L E I S Q R A L P C I S L F E Q G D F V L L V P N G V F G E P L A Q S G H V I V L L
RANDOM_1380 S P L A A K H L T V T H T D L L Q G K P I A D F A K D T I S G G P D P T M E S S F T P P R A C T C F S A H I W F
RANDOM_1381 I Q L F D P S F T R Q A S Y T T F S T L D S P S S L Y F N A M V N A L V T L V L E Q A Q D Q Y D E S L T I V D D
RANDOM_1382 R S P K I I L E K T P K V L L T S L E L G V F T H Q S Y H N K K P V T A S A D K F M S M E F L R I D N D E N P R
RANDOM_1383 A D H R D S G L G D N D D A L P A F V Q E L I I L K N G L Y S A P G P T G T T K H W G L Q A Q S F Y M I G V A L
RANDOM_1384 L L V V P P I L Q C W L D R G R L M L R G C L D E F G L N L T R Q V I A R C E A I N F K G N R S S R N Y S T V D
RANDOM_1385 K G S N N G I T A G A G E E V A N A D T E I Y E D H F E E K G S G L V A N Q T L V I S S V G P N Q T K Q S D A T
RANDOM_1386 T T V E D S I L D I T S I P T A V S L F K S L N H M E C F G K G P G S T S R D G G I P E P K A G P D T A S K N K
RANDOM_1387 T L T F C G S T H E A M L G G H S N A P A V D Q P N V N M R P P S S I Y P P K R G Y Q T W G N S G F M E G G T V
RANDOM_1388 T V S M I I V G M Q T K S E Q T G E D H P M T F V A F Q G A T V Q T I L L E L N R K G M R H W E L P Y N G E N A
RANDOM_1389 P V L A C F N S E P A E W L Q L A C T S V L K I D K A K K R M S L T Q P V T E A G A N E A N G Q A N L N A L A G
RANDOM_1390 T A L L R E A I N T L R K A T R I W V A V S Q V G D A K G N V E S R C M A R V P G P T K E L E M P I T E A P R Q
RANDOM_1391 E E K A T V K Y R T P I K L G S D L G E W G A D T V V A L M S A I H R L T D D A Q S L G V S I R A L T A N G Q S
RANDOM_1392 T P G M A D E R L L T S V D G D A D G E G P D G G V E T V A L I L N V T A A E H K P L L L C E G T T K T T D H D
RANDOM_1393 K T I F L Y E K E L C G D T G L D F S P R V S I L G L A I V F V L V P Q A T S G N C S S G E I N I N C I Y G N T
RANDOM_1394 E L I A L T E Q Q I S N T F K C F K K D N G V R G D F D A F A Y R K L M L F Q D L D C Q K E C V P P K G L Q P T
RANDOM_1395 Q G S A V L I L E R E G Y F D I F A H Y E K V P K Y V V K L S C L E K G L A M W P F T G G F Q S S L A I A V E V
RANDOM_1396 K S K A E D F R A S S G V P A Y F E F Q G E T N L T T H N L E K N L S I S A I Y K G A R D V T E A V F S V F H V
RANDOM_1397 I F L P D S F K K P I V S M A D I G G A G T R E L G F L P M N L I L N T S L F C P Q L E F P S R I S V L S H K L
RANDOM_1398 L I N R S I N I K P I L H K M V F R D G I L W I S E I G R F S F T G A K K G A A H K P I M A D A H E E F R V V H
RANDOM_1399 A K Y E T L Q P P C S G R S S A H H E L S P N V A L H K P L L Q S R A L G L T V V T A T E Q T A P E M G T F T H
RANDOM_1400 H S S T L D L C P C A V F Q P K V N M D E Q G H D Q S F N V E A V Y G D K E I K L P I A Y R N M I A G T V A R I
RANDOM_1401 P K D K T V S R G E F Q Y A V A L P G L D N I E E Y D S S R Q F L E G I A I E F E V G F S Y Q D I W G E R A D G
RANDOM_1402 D S T G N V W A Y K R F D K H V G D R I K D A A Y S R D L A R L G G L K D E D S P H D K A R T K N Q V S I L G Q
RANDOM_1403 N K R I E V I F I Y K G A P N S E N I A K E S E G L I I Q M P Q S D N G L Q R P A K N E A Q E R Q R T I F L L L
RANDOM_1404 G F Y D C K I L R H C R S W Q I L T N L Q K Y I V S Q M S F R S A T A D K P L N K L D L P K I H K F V F K S V K
RANDOM_1405 M G V E K V P P D K E L G Q R A R L I G D Q S Y F K A E E S I R E P T G E N T D G L S T L D N D R S M A L L T D
RANDOM_1406 E G F E S P V L K G N L A I P P A I P M M R I V G V G E Q T R Q P S R A Y L D K T M Q K Y K A M T G L A F G T V
RANDOM_1407 E E D A E I E Q G E S Q V M R N L R D A A P I Y P V M L S L R F K G D S W S Y Q I C K I D L F T V F L T C V I A
RANDOM_1408 H M A R P E G P D H V G R I E E L V L E W I E T P S F M F G C M P P T D V S M A M V P Q T G S F S L W K E N K L
RANDOM_1409 V T L P G S L P P N D A T D L G R V L H P W L R F V K I V G K W S E Q W D K V N L F G V E E V L D N K D R A S F
RANDOM_1410 P L L S S N L Q Y T S R A A K P P L L H G L M S A F Y G F Q S G T T I T P Q K N S S H S L Y T R E K M V G A E W

RANDOM_1411 S I S S Q K N N K S I P V M E A Q K V L A L A K I A I P I D A C R H N H Q F G F E I V M D P V G E A D K E A H F
RANDOM_1412 I N K R S Q P E P T V L C S S V S V P G K K N F H L G E L Y T A H N E V C F L L A M M L G A L A N G G R K Y F P
RANDOM_1413 E V D T Q E R A D H I E K V T D V V D I P I I F I M S E P R G D H K T R S V S A I H H D F R V W R V S F Y M I P
RANDOM_1414 T L M A Y R A I G A R G C F S W P L S I A I D F E D G V C N S A L L E G S N H H M C S D A A D A T K L A R K Y D
RANDOM_1415 E C P V T H R R L Q W T T D A F Q S T Y R A R T G S P N K H L T T T A A G H L W P L I L S F H S P A Y A V N F Q
RANDOM_1416 I H D P F I L M G A I S R L W M T N Q T A A S Q L S G P N G Y M Q E V V T S V S N S G A L Q G S A D T F A L I K
RANDOM_1417 Q R N A K I F V E D A T V G K S M A G F E N Y Q T K P G R Y P D L F S D N A V V T V R D G V V L V G K T N S H P
RANDOM_1418 I S F G V E L D K I L E W I G L A Y K G G N A Y K R G S R K Y C P V V Q A F A I E G A K E L A Y S G R M T T I N
RANDOM_1419 L N S S N Y R R P Y N K Y L P F G G T V L R W L Q A R R S N I S P L Y P Y E Q R T K R L W V C V R Q Y N R E E L
RANDOM_1420 Y Q V M H V G W V Q W A V P M F H S G K Y P S A F A Y A E I G R N R V D A V T G A S T D L R K T T C R A A S I L
RANDOM_1421 I P V D I A Y A R G K P D K Y R A H I Y V P L S I V M N A Q V L L T L T A A A P E M K N S C K K S S T L E A T E
RANDOM_1422 S Q R S F F V N E N D A K Q R A P F V G A Q L A T A A I D M L F L D W S E G M L R G E T I Q F I T G R F E T F G
RANDOM_1423 V T F G D Q V P V E K N W K R E Y G E E D K P Q H V M A R P D A T F L Y D G P H D R T L Q H S V A V V A D I A L
RANDOM_1424 I A P D E T L R R I N S R T Q S R I D Q I G I L G G T N L S L M C G T S L Y G Y F W F Q G E L S S P Y Y I P Q F
RANDOM_1425 L G T W N D R D L W V V S S G A P L N E A G I H V E L A C K E Y E L E K N Y D N S E Q P D K N H E Q V P K P L L
RANDOM_1426 D G F R V T D L S F E I N T G A P V L P H C T M Q A P H D A I P A L D S F N K G M Q G V Y T I M T V K G V L I M
RANDOM_1427 A M E A N A W L N D G V K L S E T S P H L N T Y S L V L I S S V G R F E L K A L V S T D N M R A N Q K M T M L L
RANDOM_1428 R A K Q A A K S Y M S D L A E D A V N V Q Y S I E G K R L N V R G A D N L D A F I C H M D P N N L T A C T E G T
RANDOM_1429 D G E L E K I V E A Q E S K P K L A H D S S S P S G Y I A Q S S K A V V L P S I N V C N S K R C A L L V H K L A
RANDOM_1430 L P F K A S H S C V N I E M R D E M R R I V S S E T A K V A I Q T A V F A A S N R E I P Y Q D F L N G N K G N E
RANDOM_1431 L N A H Q K L T G R K T L S I V F A A F E L T F K L P S T T P G P R N T G A G Q G S S V S Y N I V S H M T W D V
RANDOM_1432 R P K S S S I D I R V D S T S S G V L S D K A F I T S G V M D D Y A S K V L Q A A T I N N N H S E D R D N L G Q
RANDOM_1433 Q P D P E D V S R L G G V V A S Y R P L V K G C V Q T P G G Q G E Q G L R A S M T G S D M S Q Q L S L S S A Q Q
RANDOM_1434 T Y L S G A D E C L L L R L F R G G I L H V V E A S A D T C A E E I T A G D I H R K A Y A A D I G Y G W D T C L
RANDOM_1435 G S P L V N D R Q M S K N P L L F Y P D P L R E E N N S Q E Q L E I M I A T L E I V N V L S V S T G V P Q V I E
RANDOM_1436 L K I P S P I G G A R H T T E S L R E A F N G T K R T M T A H D R K G V R H S T D Y Q Q G R G E G H E L G A A Q
RANDOM_1437 G I I C S P E T L P L L R S V H I Q I V K F Y D G S C T T R N L G A L S P K G D V E E G V M D Y D D K R V Q K C
RANDOM_1438 F T G E I Q I D E G E N A A T Q S A G F F S S D R V G P D G V I S R T L W L P K C I S S G S P L A K A P M T F S
RANDOM_1439 N D L Y T Q R I V E W I Y K V T K R P Y I I G V C V K H I K V V V G L D L L T L G A Y T C G N L I A G D D K H E
RANDOM_1440 L G L D T V T E S S S G D N N L G H R Q G W V Q G N A F P D P L E D P E V R S I P L Y H T C E E D S N T G A S P
RANDOM_1441 E M K A L L D P K M H L L A A A S I S F D D E Q Q A G A E M R G W E Q R Y R I P P F R S K A I Q V A G K R G G R
RANDOM_1442 T R E N S C Q L I Q P G V F K C A E L V A R F W L T F G D V G R A D Y P G P A P C H F S F Y S Q D K R T E N T L
RANDOM_1443 V N G S L R R G T Q G F C C L L R A P E T A T K A Q T Y T W K S V A L F A D A R S G Y L M N Y L H L S F S T N V
RANDOM_1444 N L D V P D G S V V L W E T Y N S F V G C V N H R H P G G T K I A F A I Q K T D G N H I M L H V C N L S D A M Y
RANDOM_1445 P A E V A F T H Y D S L E A L I G P I S C E A T L C F D A P G D D A P D V K E E I P N L L S S I G S A G D V K T
RANDOM_1446 V V S V T C I P R T T E D Q E L W P C D D Q G R L Q Q E F T G L E S G G E Y T L Y P C Y P T P K L F G E E A S N
RANDOM_1447 T G E T D H I N M R S I S L N E P S R D E P R A P R Y E E G V Q V H E M D N Y P S A N M P F E K R N R R E H E Y
RANDOM_1448 L A G D L E M V R V R N E I P I I A Q L P K L Y G M V K H E E Y V T D S T T E Q H N A L S P W N F V F K T E R S
RANDOM_1449 H T S L K L E R V V E F M V P E C V S H N N G I S I T T N L G Y L G A T Y L S E M Y T F E R A S H S F S I W T L
RANDOM_1450 V A R W A D I I M R A F V R M Y E K Y A E K T I A E P A A F N E F H W I V P G T V T S T T G E A W Q I C L Q G D
RANDOM_1451 V S S N A G L K A V Q G I S E P N Q S V R T E A N S G P P M F Q E C Y M T V N K K S G E V G A S T A K W P Y H A
RANDOM_1452 Y I V N F P I Q I N G L I G H L H C N N R G N D V P G M A A R E S K E L K P N E L L V L A E S L T R V R S N P G
RANDOM_1453 Y Q F K S Q I P I G T L T Y A K R Y L E S H K V V G Y I N R V S F N T T T T G F T R D K A Y E Q G T P H D G E Q
RANDOM_1454 A A D L W S E I D S Q C N R N H P D G K M W D L F L S K A A R K V L T L V Q Y K T S S L I F L R Q C T L Y L E K
RANDOM_1455 S H Q E G K D A G F H S V K E G M E A L K P S F G N L A D L N R M C G M V E K V P T E S T F H A P G R E T E P H
RANDOM_1456 A L S H I C S A P E D T P Y A G N A E Q K T G I L Y S R D A P K T V P T G P G F S Q L R E S V E D H Q S H N D N
RANDOM_1457 K S R S N K S G F P G A E P L L G S F S G M V P C H P H L I E L K P C G P S P A P L A Q S G E P S E I E E G S A
RANDOM_1458 E G M L G L N R S A G T A Y L G G S G Q E T A V D S V L L T R Q M A L E V L P F N P Y E E K R L L F I G R Q L L
RANDOM_1459 D R H I T T F S A L C A Y W V L P V A E T V L I H Q A Q N Q R F R I F S N P K L D V P L Q G S A Q G S T L V D Q
RANDOM_1460 N N L N N T A C K H G G H E Y A Y S A L Q N N Y Q D A Y Q G Q A C P V R F S K I T F G P K V K G K G V Q A L N S
RANDOM_1461 P R R I A M I V E Y R I E L S I K C R S H P F G A S D G V V K V R M S D M S Q N L V K L K P V R E L T P V G T D
RANDOM_1462 N R V F E Q A R Q V R R P I C E V M F K F R Q P E G P I G K N D K V D C P D P P F W A A L S C F N H C H S C V L
RANDOM_1463 P R T Y I G G L R W G K G H S Y F Y S A P T I T K H S R E S V P D Y W A Q S H E N K L G E A S V N P L A P C Q K
RANDOM_1464 F F Y G G Q R A V S G K Q G E G R R K C T E L E P V R I M M M L N L A H T A D S I C P N S V R Q Y S V P V V K P
RANDOM_1465 V R R A M P Q F G W S K A R A A T N L I N Q A N G V A F K I D L R V S G D A E N G M I F R L R A L D T A I K P S
RANDOM_1466 A T G R D G I K R V I K V I L W F A A N R V H T P M R I I A H V S K F L F Y S I N V L K D Q Y G P L L I V F P E
RANDOM_1467 P G V V N C D I Q V N G V Y K A K E S T S E V N D K G N T P C I H R E L F P S D V G V Q F Y K K D L M E L N S G
RANDOM_1468 P G P Q R S L R L Y R K K E N C N M D S E S A D A N K A T K V A E T K E I R H Y R V C D M K R E P E L A P L N T
RANDOM_1469 K V L I A A P G K E R C D G N I F E T Q S V T T H G F A K S W H H L R S S I G T I V A S S V I G S V Y K A R G G
RANDOM_1470 D Y S L S A K W I S G F N A N I E V G Q L K V I M E V V P K R D F N L T G E V D K K Q S S I D S V V A K R D I L
RANDOM_1471 Y N R G L K R F P V G A L D F C E A S S S R G I A L R G Q N G V V V L S T S T V Y L G T V F Q L A P P K A E K L
RANDOM_1472 K Y D G I I K Y L V S T V G A T L I N A I D V V D G I N Q Y F E V N G A A A D V F G Y K Q E K A D A V K P T Y G
RANDOM_1473 I L I V I G N R K S Q A N S C P A Q V P Y S G L R S D R F N D N S A Q P L A I R G D V F I D L C T S L F I D S V
RANDOM_1474 V N S N L L E R E K T R S K Q Y P A V R Q S R P G R L E D K G P A K R F F P N T S H P P P T P T Y F L P D G L L
RANDOM_1475 K I F P I K A D L F K A A Q D V E M T I F A N P N G R G T W N S L L W D M S K R F F T E R Y V P T D R E P H V M
RANDOM_1476 E A I G Y H D P Y T K F V P E G D D S S E G G E V A Q D S V L Y R V D Y E A D G S Q I S I M A G K A I R T Y K E
RANDOM_1477 Q K F T P L V L V D K K H E A S E V G K Q A A N G L Y T P L D L S K G R Y L A I E N N Q T D E V T K D G K T D P
RANDOM_1478 I S G G C E Q P A A P P S A S A S F F Q C D N P E S N K S W L P V R G D C G M S C N A E Q Y L L A T Q L S E D M
RANDOM_1479 I V T D F T S L R Y W L R Y S K S H Y L Q S T R M V L Y L L S K L C Y I P G G M V Q Q Q R K G K G L A V K Q Y L
RANDOM_1480 S T H L R L E G I G V L Q V T G E Q D L G E Q G L L P K I V D D V L P S R K G K A Q K P K L V L G P D T L H F F
RANDOM_1481 L L S G T T S I L G L G A E W K S I Y L E G S V T V V I I R A I V E H Q V F L S T N S K A P Y Y K N T R E T K D
RANDOM_1482 V S S G P G K K I W H E S L A W E L T S S F A I T G Q L H Y F L L S P M N I M N E K K P A T V N T L E S G H L A
RANDOM_1483 E I C P I E E S V A C L E A L T A R E H E Y L D Q T R H P G L I N D Y T I I I L Q A E L F E E E L E D I N G G K
RANDOM_1484 S G S P I Q A V Y P P P V G D K A E K Q C E D N N Q G G G Q F E D I L N T G S R A P S G K K S F Y F D T W N V E
RANDOM_1485 A T K A T V L E N A R L L G V K R C S V D C T P T D V H A H T I N P A N L P Q I S G E A T A A A N E G L Q T L M
RANDOM_1486 A E P V E I C V H S A D V N G E N C S P V P V D C L S V V R K R K M D R G A P Y V Y D R Q G V A I E N R G S E V
RANDOM_1487 R D I I E F L S I V A G D Q A L M K K A N V R A D N P I F Y T M P G A G M N S Y F L E S K L Y S G I D A L T L G
RANDOM_1488 Y T D I R Y C G I G Q K S F H R A E A A V Y S V C L A W Q I I L F A A P H E S I G T Q P V H T I R T R R N G V I
RANDOM_1489 A K L A L L E S V C D A T A Y L S V G N L S P N E D N W V R C L P N D T Q A W Q V S A R P L L D K M Q W R A G Q
RANDOM_1490 V T Q E S S L K S A A T L D K T L E F V E T K Y I Q G I F D V Q K V P N Q R F E D Q L E I N R I A K V R I V P L
RANDOM_1491 Q A I L G Y K S I D G M P L S E T N L D F E V G L A R G Q C D E A L C P N Q L H T R F M V V L L L M H N Q S T K
RANDOM_1492 A S L Y P L T I Q S A A D G V E S F P A S V G L R V R V A I M L Q E L P G L Q S N E M G S N I L L A N L P P S Y
RANDOM_1493 F G R S L L T A S C G M R L A P Y V K V R P R A L L P A V I I H I I A Q M M H L P E S W A G P E E A Y L K M K V
RANDOM_1494 T R D Y L G K S T P G G R S H S V A L P Y L G A V G G A P L D E E R G S H I V N I I W T I C V E L L A P A E P E
RANDOM_1495 Q V A V S T L L D A I L T Q E S L L G V I V M V G A E V V R A G V A S R L K Q T K I E L D E L E L A F P G Y L N
RANDOM_1496 Q S S V L D T V K L Q N V Q D K L P G E T M L Y I R S S I D A T E L E K D D K P I E M C A P H I A E Q N I S S E
RANDOM_1497 K R L N L E K F V A T A K E P A G V L P N S Y V G T Y N S V D A F A N L M E G K A G A D E A F P Y I S F C E P V
RANDOM_1498 R R P S G V W S I L G V L Y P Q G E E K F A P G P A I Y G F L G E S R R D W T V I T T K S T R P E Q P I R D N N
RANDOM_1499 K K I A N N V N F A E M F T P S Q G L V Q M E A G Q T A K N L I E F L G R P S A G Y I V Q G V S L Q T M Q S S D
RANDOM_1500 V G I D D K A V Y A S S I R V P Y N A A D Q L E D L L D D T L N H S N A L K L A M N E K L M Q V G T P T K T E E
RANDOM_1501 T Y N I S E Q G C G K K V I Y I N P F S V K Y K F R V L G G Q R A K N I R V P M A L V N S G R P T D A G P H A Y
RANDOM_1502 T V Q I T R I F Q F A T D Y V L F E R T L M D M S S I D S L W R G L L D K V E I M E K E S E V T V I V V R E E V
RANDOM_1503 Y E T R L Y G T T V L P Q R V R N E S V T L R A V S L Y I S L G G G D V N V S L Q T I H E V T L Q N I N E L L N
RANDOM_1504 H E Q E A L T Q L P E A F A K V L F D G K Q T D T R R S E A L A G A G G P G C V Q K T N H C K A T E N R W L V T
RANDOM_1505 S A L E Y D I V G G T K G T C L G M C I R A T G E P N A G I F P S Y E F Y R I A S A K A I P G R D N C E N A F L

RANDOM_1506 S T A E A Y A R Q A T A F P A P K E A K E K I P T E P A H L L E D M F T V T P A H S P E I E F T L A P R L F V H
RANDOM_1507 Y P I S G G N G N L Q T A A D R S P E V V T D A Q S I G F E T L Q L V L H D E N Q A E V N L S I D S L S T H P N
RANDOM_1508 E T T D I G F I D A I S S K D Q G T W C L R V A L V A L Q G I L S G I E D T D R A T G K N Y L H G L S K Q L N K
RANDOM_1509 V E A T K Y L D N K G G T E A L N K C L S V A L P R T L I P A L A S Q C A D H A L V L A P Y D S K F G H R V D I
RANDOM_1510 A I T G Q R G P E G G A C L S Y S T P G I G K G F H A G Y T G A S R T D S C R D G Y L P C S D K K V D H L A N I
RANDOM_1511 G S R V N E S P V T P E E S G L P G G E P A W V Y V Q Q R D R K F N L Q L T S K L C R Y H A K N N K P P N Q P P
RANDOM_1512 H M T D Q A E H P S A P E S S Q S T E P D T D P R E T S S S R M D R E Q A V V T A A E Y D W G L N Q G H C D E V
RANDOM_1513 R I A I L T L R D P S M A S G V L F D Q L S W L G R I L E A P I D C K V K E W K F L H S A I V S R A T A L N V F
RANDOM_1514 Y E L A V L D E L E L E N Q L L S G F Q H A T V K S M Y P L Q R Q G A L G G R C K N P L M G R K M I V Y S N Y M
RANDOM_1515 R M Q D T V G V D N L P L G A K G I E Q L E L S S S L A I V G F R L A I K F V A V L T S I R W E L I I A S F F L
RANDOM_1516 D T Q A G F T L I Q Q Y Y L Q Y R A A K V S A V K I Q S E I A V L C T H R N V Q L A A N S H C A T N I K I G G K
RANDOM_1517 L D P R F E F H C D T L F T E S K V A N V T G N Q Q N V V F A R G K T E H G A A D R R L E P T V A F N I T A N Y
RANDOM_1518 Q Y G Y E D S S N L K V L N L E H E P L S L L D L P L V P K E A T E S E I T G H N T E Y V G S Y G A L S F M N E
RANDOM_1519 K G N G E Q T E A A A L R M V I S L A L G E I S V T V F V P A G E R V N S S E V W N S Q V Q G V S K L K Q V G M
RANDOM_1520 M T H Q L T N D L E H E E V G A H V W K T R N G G Q T M G T A P I A W N T G Q N S K T Q L S P W Q T I T L K E K
RANDOM_1521 V T T T I I K G G H A L R K G E V P S S I E V K A E G Y D W E N V I C A V A T I F R V H H D V Q E S N S V K T D
RANDOM_1522 T G Y A L S V L L S R L A R H T R V V A S N N A D Q R T A S I S V E R E S E G I A G K N C I Y C E K E H T T A P
RANDOM_1523 N K S G D D C T I K P R A A S R Y Y E G A N T S Y P F P G P E F R V Q G E F P S A M A K V M A I D V E T I A F Y
RANDOM_1524 D V S A D R E G L A G F G I N K D V M P D H T K G P E K K A S A R F L D V S S E E D L D N F E Q I L G L R V L A
RANDOM_1525 Q R Y Q E D D I R A G K W Q S S I T G E Y L G S Q T P P S R L Q C N G G F I A S F P M T S D A L A K Y V F N N N
RANDOM_1526 E L I F V K D A G G D K R G R K F R R G T D V D S I I C F G D T V M Q S C I I R W E Y Q H L D Q L Y L K L K H K
RANDOM_1527 A G L I R W Y R G E D A N K L L R V Q K D D R N Q G G N S L F W N Q V Q T M S I T S T F R M E S E S V M L F Q Q
RANDOM_1528 D T E K P C G K K G R E S T P V T I A G P K L R A A A G L R D G H K G P L E E P Y M S L P I E C E D L M R K N N
RANDOM_1529 V I S A R T V K S A K P H F G R V D L T R V L T V L D K T V V T G Y K R N D A E A W G F I P R P A A L A A N T P
RANDOM_1530 E K R L G P L V W T A C W L V Y P E K P V T L M T S C H Q P G P F F D T K D T P Q K L Y D F F F Q F I I E S I N
RANDOM_1531 P Q P N Q V H L G I E A A P E E R D L C S F P K E N A F L S S S K Q K W W L T E V L Q G L C D T T I I M F G K E
RANDOM_1532 Q W S P N R Q P E D Q T P L A Y M D S P P P L E V V Y G L R G I E A Y Q N A Q F T S L A H I P R S L K S V G T L
RANDOM_1533 C T E T P C A V A E I E Y D I L D L K A R C P R L L Q F I C P G L P P A K C K D W I L I D K L A A A A T Y S C Y
RANDOM_1534 L A R T T T C Q T T A Y S H S A S R T F S G W G G R S G V W E Q G A E L V A G R H T A N C L K N R F V I S W A Y
RANDOM_1535 E L K T P I D F V F P L I N Y T G M R W R V Q R T V P M T S G M K H P P T R N P E Q G Q M E L T L A L V T Y K I
RANDOM_1536 A G D P C I W T N L W Y L R G G F A L R T G W N I D E C L K P M V S N M E A R A S S V E N G Q R A F H E D A L T
RANDOM_1537 E H K Q N T N A V T H T A A T N P E P M A A V R N D Q T M A N F I V T L V Q Q T C W G L E L K G G I A P V K G A
RANDOM_1538 V A V I D D G Q N E A L K P L L H N L S E S R R G G Q F E G T V E E E P I I Y H V M K N G S E C N A F N G I E I
RANDOM_1539 L Q N Y E S R L T S G E Y G T S Q V I T Y H A M Q R I A K R G L Y V E D D D A Q F H V D K T N S R V N A G W G L
RANDOM_1540 H A D E Q A F S R G A K D N L P E S S G D A L S I R G Q G V V G A V E L V E C I G P N K P A V D T V K A Y L P D
RANDOM_1541 N F K N E K N Q Y L E V S G I H K E D F V D I D K P S G S A N M N L S P A L R A L V K V E V T D K T I E P T P Y
RANDOM_1542 D R K T V E P K F T G G M Q F G H F E C G E T E L A P C N S E D Q K E N R L P G P D P Q R S E F V L A D L L E D
RANDOM_1543 P E P Y K N D P L S T D D D L N V C V L Q Q G I I K A L Q G F I I R A V N E L L Q A G L D Q F F V D D V A I S N
RANDOM_1544 D E L L K V A R E P S R P R F D K G L I L E A S E S R K R A A E L E Q E A N F Q D L N T T K L E N C P I G L L S
RANDOM_1545 E P L S D A E N G A M G K D D K G S G S D S L R I A N V V E S M N D P L R T T L F V L H R H G D E A T F E V P N
RANDOM_1546 E D P D F P M Q C T T F S S V K G T G D G C M I I N N F H L Q V S E E F E L L Q A N E Y A V N G Y S V N V H E K
RANDOM_1547 N V Y C N D T H Q G Q E T N G K D K L H L P L T A V D Y M V K N F E P S T K A D Y Q G Q P L D V P T S E F I V P
RANDOM_1548 Q K A K G R N A N G Q L A Q H E L A P C D A I G I A C V A A M G V H A G H F L Y T I E S R A A L F E Q E L E S A
RANDOM_1549 G T L N D M V L D R T I Q S V S A R V K L V T L G D M G R Y P D Y T R R Y P T N S A C E L D S M D S S V G L E E
RANDOM_1550 S I C L G P P K D P A T E E I N G K S P W S G T P G N I F I P W S L V A I N L K F P I M V D A E H S L I S Q I V
RANDOM_1551 F C S N D P C D P T G T N N Q I N S T G M M I P E D Q A G Q H R M M A N G I E P T F E A R L T R P Y S L A I K P
RANDOM_1552 C A A L V W A E W G R S T S H G A E S L E H A P A D F R H I R N G Y P V A L S V S Y E S L I R E S S E G I L P K
RANDOM_1553 I S E K T K F L E S I Q A G I N G N T S N Q R T A I D S T G T L R L Q N R V D R I P A E K T G G E A I L D M A R
RANDOM_1554 M Q V V Y S E T Y V L K P T G A Q R H S T K E R S S T N C V N N Y L E A V Y L Q A G H E T A S R D R I E C I Y Y
RANDOM_1555 L P T P K V G F A A E N P R H T L L H F H I G K L N L S G R D G A A Q P G S H K S E N D G L P Q N G Y N L K G Q
RANDOM_1556 G S C Y N Q V M L P S I P A P F F E T L S A K P G K T R T S D W R D D L P R L R Y A A N I T E G H E T L F N S G
RANDOM_1557 G T I T Q I L S A G C T K K L F P L R D Y K K P P H G N K S Y M F T L H L K E I P V F F P D L Q T L P D D P F P
RANDOM_1558 I Y P V A V S P Q S I A I S L R F A S V P Y L I R G L L S H G T C S Y S N G R Y V G E P L W F A L Q K R G E D N
RANDOM_1559 V Y G D A Q E P L D V Y F T I E S T T T V P E A D R N S F Q G M S A V V E C Q V Y T T A A L K A E T S Q V G V P
RANDOM_1560 R V A I A L A A P P Y N E D C S N W L N T A S H L L L T E S A S E S P T I K E A T D Q S F Y E M L T T P D S A I
RANDOM_1561 K G T A L K I N G A C C D T L K C Q E C H A K V A G S V T R S F F G Q H F D I D I S V R R K A P Q F L S H C S E
RANDOM_1562 T L F Y A S L I V G D G S N K Q D Q A K K P S R N D I K S L L L L K A M S N M G L R L M R L V S N E S D E R D N
RANDOM_1563 N R S T K L Q A R F A I A S S G G L T A E E F G E Y A P Q I A Q K V G K S A E A F F T G P D A Q E C L Q Q P Q Q
RANDOM_1564 P D Q A L N E S A E P C I D L G Q D D E C N G V T T P S V H V Q S D K A G A I E F A D L Y E Y D I L T L V V G P
RANDOM_1565 K D L T C D F K A G S D W A D Q Y T S A T Q D E V K I D H G L M A S A P S A P T E G G S A C T Q S T K V K V L E
RANDOM_1566 T L G T H I N T G C D R M D K I N R T N G D E Y W S E L Y Q F V G P K F S E D V T W T C G Q F T A A G D D L Q S
RANDOM_1567 G G Q A K K V T R S V Y K S T D R C R S A G H A S L A A G I S T S N R R Y S I G L H K S S A L E P P G R I Q T I
RANDOM_1568 S V F V I G K L L F S Y Y K M V I P I G Y S R E K V S N I L L S V I K S C L A S G A A V E R N A S A L D T S S S
RANDOM_1569 G Q G Q E S G I N K R V K R A I R N V G R E E S P R I P P L V Q L L K D S Y W R W F V F G V D Q I L D T A G P Q
RANDOM_1570 L G I D D N A G G A R Y F L D L R C K E N Q V I L G N T L A A N L S S A G E A R P G G N L V S T F E V E Q L Q S
RANDOM_1571 L D V V R K L I L E G S G L K K N L Q V A Y T I G A R V T A Q R V Y K F F I S G G K C D L N I V P Q F D Y T V C
RANDOM_1572 T L A T Y V H G S T A L L D E E V A D K W P S L S E A N D E F A D C I E A F G A L R Y L I E P G E K I L T D K D
RANDOM_1573 H A V L V K E K Q K K K Q N G A Q A S Q D S S D H L T D Q I G N K N L T R Q L D A G G T M M P N V N F T L G N S
RANDOM_1574 G D A K L G G F Q L K L Y L D K L P A S N I N V T I Q L E T K S E I D T D P S L R E I Q N S F G N V S G P Q I Y
RANDOM_1575 T P V G L K E N N A E C Q A S P G D C F T T E F A G D N Q N H E A K R L P R H G A A I V A G S T L L E H S W V E
RANDOM_1576 Q V L Q A W M Q A G A A V F F V K L V T L A H V F L K T E R P N K G E K G Q G S L P T G G V S G G F S G H E T V
RANDOM_1577 I L N T H F L A V G G P A Y Y A G R N S I I Q R V D A A C K G F V I S T F L N P N D F E G A N F E S N L V E A E
RANDOM_1578 R Q Y T M D L F Y Q V A T Y Q V D I P Q K T L R S R D G V S A S S Q M K Q S A L Y T H K R N L H Q L S A C V E G
RANDOM_1579 P Y S N V I P Q Q A P A S G P A V I A H Y G F Q A A M V N G G V C T S Y Q P Q A E T F E A Q Q A E H I D A L S A
RANDOM_1580 G G Q L C L N D S P L S S L S G C L R W A D S F T T D R C V K G L C D I S K T T D L L D Q G P C Q G D I G S E I
RANDOM_1581 Y A L V T T A K C P S S A C I G W A T L V E T L P N V D P Q P K R Y S D S N M N D A T S M E N G T M G T K P H D
RANDOM_1582 T S D S T E M I E A A A K R A I I A T Q V S P W D A V G H W E D L M K N K Q V V A W A S E A Q E M L G G N N S Q
RANDOM_1583 E A P V E N G T A T A L T P G Y G L L S M L F D I N V E E L E H P P T S V R W N T S S E G A K N S E S Q L Y A K
RANDOM_1584 V V S V G E Q G P I R V R D T Q L N Y I K T F E G G Y N Q Q N K Y E V T A L I A P P S N Q F Q A F A E G L F N W
RANDOM_1585 N P K I Y E P T L N V V A A Y D A D I D T G T D A V L G P V K K R P L V A R A K P A A V E A K R V Y D E E M N S
RANDOM_1586 A A C S L S V K S E Q Q K E L D T S N A I L T S Y T L W D V R I Y Q V E G R G F K R P D M D G L D V D F L S D Q
RANDOM_1587 P N R E A N W E Y E D R D P L A N P N S K L C E I V V A A G N L K C V M I G S D E N E A H M A L A Q T E L I N W
RANDOM_1588 S K P A R T D A E E A Y F D E P A N S E Q L Q I D F A V I I P A R F V R Y Q Y P A G E H R A D V Y K R T F N G M
RANDOM_1589 N A K P Q V R S P I Y S V C F R P D E A L S L H N V R S A P P E W L K S D F Q P G S I W Y A D G S D L Y T R M I
RANDOM_1590 S V H A A N S G P Y V L S R G L K E L S H H L F Q T Q G A E R C G T D V R N E H P L G V S F M M P F Y T Y L P L
RANDOM_1591 P N P S Q A V A S T C A L Q D L K D A A D I S I K G D L C H L E P A P I E E L P V L I R L T F L N W S L I E S Q
RANDOM_1592 M E G A Y T M C A D A P S M I E Q L V G T P G R V R P I P A G C T K N F Y D K V Q E S H Y G K T G K N G G P I G
RANDOM_1593 P L L A N N N S H K K H I D A K R L P P E G I W L P N L A G R N M S Q D A F E R F S K T G T G C C R G S W R H P
RANDOM_1594 A L E L K L N G L Y K L L S N V A V I L T I L I L G L G D T A I P V K E M W Q I A L M T D E N D A T D D N S I T
RANDOM_1595 C D G M I E A K A H K H D E I V R A T P I T R H I E G M D I D M S H V C T T L V S I Y N V D I I Y G V K E C G N
RANDOM_1596 E V S A E G N H F I N K T T R A A T I G H P F R L I I S E N D V C L R V E I V Y E N L D I V S G G N K G A F A A
RANDOM_1597 F A Q K F K K K R Y A C A R L Y V E K N S M N N S Q I F K W P L Y E P T P V K K K H G V N P G E D S P Q L R S G
RANDOM_1598 Q G T D P N K N N S D Y G V C H T P G Y L V I G M V L A F S A I S A L L S G D G Q V A D G S G S V G K L A G E S
RANDOM_1599 M M L Q T M R L I L T C A F D D P L F S S S T H A D E D P G G L D N F L N H M S L A D A N E D S G G D E L M F G
RANDOM_1600 H G N P F A Y P L D L S P K I L D F R L E A N Y A L L Y T I G D P S I R G R V N K V T R Y G L Y L G K V R F G K

RANDOM_1601 Q T G E V D E G E S I L N V V E A N G S K K Y N V E V S D V D P A S S F G S K L V E V L T Q G D N D P E V A Y K
RANDOM_1602 K R V A M D A N S S I C E W V F W D K T G G L P M W Q R A V K T V N E S A T L E R Q A R F N H M D W M A T A H V
RANDOM_1603 N M N G E L R N S V E I H G L L V K H G C G L E A G F N A T S K P Q E F E L H Y L L V I D V H V D L P A K S A G
RANDOM_1604 N S G I N A K A Q V I L R D L N L A S P T V R I K R L K T L L K C R F V G D S G Q L T Y S E K I A E L K G E E P
RANDOM_1605 N K E G Q I Y N G W G I N V L L E M N V I M D T G T D I G Q F H I C V V R M A V R N G F T I M S L S K S E T S R
RANDOM_1606 Y T A V M R A W L V F K M N C T D K I L H R L Q R R A G V P G K D A V A R L Y Q K L I Y L R A E L L R G H E A K
RANDOM_1607 P R S A W S A K M V R L T E P V C N G S H F L D I T L R S V G S G L R F S E Q L I I E S N Q L S A D D A C D K Y
RANDOM_1608 R L L C Q V Q M V G G G N L S Q K G F L P R P N G G Y F W P R I F D A T S Q E G Q T Q K A G P L Y A L S Y S I G
RANDOM_1609 S D T R P T V N I I K Q V Y M R V E L Q A T P P K K T V V P L A R G S C P D N E G Y K A V L P V H I C K Y S A D
RANDOM_1610 L L V W T G K I M E A R A A P Y L Y L M P I I Q K R D Q G R V S A N Y W A E Q T P E I F Y T P V K S L S L D I T
RANDOM_1611 R I P V I V E N E D L L M E E A Q L A S A L Q R P E L K C R E P I D L L K R F N G K L G F T V D V A Q N D A G Q
RANDOM_1612 M L V R Y R T R E L Q R S N S S N D Q F E G M A T N T T A D V Y S L L A G A G N R M K L E P Y D A S L P H D D D
RANDOM_1613 P R Y L R S F D L D M R K L D S G T E D L D C Y D V K F R F P G S D G F R F Q S T A I K G V N K S H I M W S C V
RANDOM_1614 R Y P M Y P K T S K K T G H Q S P G P S A T S F Y S G V K A A E A L V T E K R R V E V V F V K C D G A T G F Y G
RANDOM_1615 N I L N T V H D P Q V C P L Q L T S Y L L N S N W K A D K A N I P Y S F V K S V L G A C S E A I I G E R G K E V
RANDOM_1616 P R T S Q Q G F W G G I Y S C V S S I D P V F T W H D I A W G V G V A R S E P L G S K F T T L S E G D N Q A L N
RANDOM_1617 A E I C E E G A D N T L E E A I K G M S L T V P P A L T P P G I T E K G V S F G D R N W I G T M L A K D N S E A
RANDOM_1618 E A G Q Q T N A S Y D R C S L K L R E W Q D L T Q G V K A D F N N V K A F F V R A S S Y S V P T K S S Y V F Y N
RANDOM_1619 R G L R C E D F K R E N E L E R L L T T D V Y R L S E D L F S P G N A E N I V A G Q N V L L L L P I E F N V A G
RANDOM_1620 P P K L V T L L P A L K L C R S L S N D E G K L P F H S K Q E V T K S I D L E K I S I S A D K D E E S R K K S P
RANDOM_1621 C P Q T A C G I L E N E Q E L F L S D A Y K E A A Q K V A Q G S A E K F D N E Y R G V L P Q L N N V I A R T R F
RANDOM_1622 P G K F A V T V E K I N S D R P N L R G V D D S G H R S V Y K S Q Y M P K W K L G P V P G I S T G T M T G V G N
RANDOM_1623 T R T G R E A R G Y G E K L K P P I T G S G I I V V S E R Q V G D V A V V K N Q A P Y A V Y L A N G S G L S S P
RANDOM_1624 V S Q I N I N N H E F F L E Y D K P Q S Q A R V D E A R R D P P T K T S D R A K S P G L W R M S P A G A N V E S
RANDOM_1625 C E E L S E D Q G R Q Q D L R F N P Q R W V G D V V I K I Y M K S D S D A V H I S A I I D G Y S R I Y Q G G D S
RANDOM_1626 L V M L C G F Q F N P E H G W M R E L L E I F Y F F V R K T R I R H R V P Y K L F T V R K Q G R N I I A H C D Y
RANDOM_1627 C V E L L W R Q S L A K W M T R L P I Y V M G P P Q S A D K A V L S A I D D F L S G I C L E D A V L F Q V S L I
RANDOM_1628 A K Y K G L F E P N N L N K P G T A P A L S E N I I S F Y T F A V P N V E I E M Q P R E A T E S G P Q P G V D A
RANDOM_1629 E H K D S A F I K A Q F D E S E T P Q G P G I E V G G G A A Q V G R L E P E T F D W R S W A A S R Y P D V T P A
RANDOM_1630 F P Q S A G K E V A L R E P R L A N K H R G I D R M Y Y D L E R K E P W A K E V R P H L A N S K I E R G T S I R
RANDOM_1631 R V L F A S N S A E N A S G K Q S I T A V S L T S P S D W K E K P S Y A S P D F N Q S A I I W A R Y P L L I V T
RANDOM_1632 L Q A E S A E V P I E Y S E I N V Q K T C Y L A E Q C I F S A V V I F P V V K L E D Y D K L K E Y T E S G R L V
RANDOM_1633 S S S L V M A S Y G R N C V V L L K K G A V R N F H H I A H D N V T G A A V L I H D I S Y A G L H E E G H V D M
RANDOM_1634 R G K S G D W G H G G N Y V D E T D Y Q L V E G D D S N T D L Y I K L S L A K T V L Q K A V Y D A A S V V T A K
RANDOM_1635 G D H A R R D A F K A V M I N V P R G G H M I W P L P V D G L G K A L T T S R N T N L F A E A E Q L P A D Y L L
RANDOM_1636 I A E A N V L H S W C S M S N F D L I E D Q E V A N S P H L W N N N A A T A S T E G Y A E I D S D D K E L Q I Q
RANDOM_1637 Y Q G E N L Y I G T D P A E F L K D Y H N A R N I Q R V G L N E V H S I D F L E G Q I K I V Y I P K A T V V D I
RANDOM_1638 T G L L V L E S R L L K A E L G W Q K Q I A H N F G T V D T E S W E N E M I D S D T G N K F F G H V R L K A Y E
RANDOM_1639 H E G K G S P Y E F G T H S L L E A S K I K V I S M L V S G D L K T D A G T K T I T T Q G D C A V E Y I E E N L
RANDOM_1640 K E D Y L L T E S G A G G T E G D K N S D L P H E D T T K M V Y G P V C Y G S F I T E A E G R K Q L P V H Q R G
RANDOM_1641 I T E G V K K S L S F W H Y Q K V L L N A P A V L E L Q A N L T R D L E P P A E G P A E E D L G R I E D Q G T R
RANDOM_1642 T F E L P G S L Y H T Y P L S D L Q H Y Y D E V T T V N K L P P T N P R N A K L G D E S G L N L T H K K L A Y T
RANDOM_1643 G E P L G S H F N D E L Q T E H R S K C A E S A K D E V S F E P V L R E S S N R K F S K G A N F S I D S R L K I
RANDOM_1644 S G K L E N A K V G Y M D S S Y R A N D N D N Q N G E K L G D F M E S Y R T G Q I Q T D I F E R I A E L A L A V
RANDOM_1645 A I M S P V L Q L Q R F A E L N R C Q P L L P M Y S D A T Y L G G A H A R P E M L P L D S V N G M G F Y H A K E
RANDOM_1646 T G L P A N Q F K L S R I D M M I P Q G I N P E A E L V L R C T S L R A I L T V A C W L T E P L Q K S N N H Y L
RANDOM_1647 A I A D V F I V V D P L F D A L H T D F A A T F A L T I N E Q I P F K E T T F A A I K Y I K P N Q Y W C C T N D
RANDOM_1648 L F Y R I G S N S F A A E E M Q T L F L R L A A S V R E Q S F L K L D N G L S S I V Y A K G D Q Y I F V E E L C
RANDOM_1649 I A A A V D C W E S Y F S G F Q S V V Q P D R N A N Q A T L C F Y W T G M S R I L C G Y K K A E K N Q F A E Q C
RANDOM_1650 I H V G Q P D Q G A L K L Y T D L K Q T S K P Q T C S R S G G T F Q K G L G T A E R C L K D A K R L D L G S I I
RANDOM_1651 S E L P A L L R L A N E V P G A E W S C W S F I D Y H E D A K I V Q L G H A R A Q P Q Q G L R L V S E S I T Q R
RANDOM_1652 N S Y C K D E P P T L D V E D H S Y A I E R K E G V S R H I I L R F S I G S V F M N P G Y K L L G S H T T M K N
RANDOM_1653 V I G V H G N K D S H M R A D K E T I F R W F V A Y S Q S I A K A G A R L K P L T M W A A N S K R K S S A A A L
RANDOM_1654 E Y S A G V T L P K G V E N S P G S I K I E H A K S E I E S D E G P V E V T K D T P S G L P A A K N R K V P D S
RANDOM_1655 W A E K C E I N N I V S G T L Q K Q L L A F Y P F E I R D I T T S C Q V K L D L Q G H S S K T N K L E G Y N L L
RANDOM_1656 S Q G T Y T G R E G S K A L G K T G T N E V K Q E K V G L K A F V Q W D R L D S S P Y V G S V G S E A T A V K S
RANDOM_1657 L E Y L D K F T K I V S H T V Q G E S K A G S V S G P I G N D L V Y S T T D E Y K D D M P L G G V N W K I I K H
RANDOM_1658 D E L K A D D N P E V P P T L I R F G K S G F G T S G K M I E G T S L I L L P I D T Y V K A K C K A L K Q C M K
RANDOM_1659 F A I T R H E V R A H G Y S V K P L K L Y G D A V L A P D N C W E N A A D T A P A D G F Y N N G S K R R A N Y R
RANDOM_1660 V Y D G Q P F Y L L V H H R P H A D P R V F Q D T M F G F L V G S E Q R I T M E W E E I M T T P L S I L W Y N H
RANDOM_1661 E S E I E T E F P F M H K A I L W C P C M P L L D R I A E L K Q G V S R G K D A L A S H T Y A G R R I T H A M E
RANDOM_1662 E N A G D E I G N H A A H L T T A D T E A L K K F Q K M V D E P V T F T K E Q P L L Q D G G P S Y N P G S R K D
RANDOM_1663 L K V V P D G S D E A P L G I Q T S H K D W I G S M K F L Y I V T F P E R D L G G T V Y N D A T A G P N S K E A
RANDOM_1664 V S D T L T R T G F F T E R T V L D A N Q A L E W D Q E T D L A N I F G V K Y H T Y N K L R R T D I N D C L I V
RANDOM_1665 E R A K L E P N H A N L H R A R Y Y K N P G R S P Y D A G S V E S P R G T G D Q L G L V L K V E Q A M H M S A S
RANDOM_1666 N P G L E W K V R C Q W A P R A V T H L T E K V A H G E W A T H R I W K E R M D F F A L K L S K A T T H G F V A
RANDOM_1667 S T S I L Y S N Y I P G I V S Y S V S S S D N G T H L D N C Q C E T V L N V S L S I S T V L C K V C A K K S G L
RANDOM_1668 P I R Q M G G N G K S L V S H H D S S I E L K N I K F L T D D D R A A I F D I Y A L C G G V I V Q R A A T K G A
RANDOM_1669 M K L D E R L M G E G E N E V V E L P D I H N W L I E V A K N K N P R A E S G E T V Q S N K V A P T G G N D Y V
RANDOM_1670 P I L R A G F M D P I I G P K A N K L E L K A Y Q R I I S T S S G S E Q I T E D S R W H S W N K T C V K A P D E
RANDOM_1671 L G E M L R F L S R L R R D P K L F F V D E Y S V C N C N T W S E E Q L E A L T Y R C L A Q T S G G P H P L F S
RANDOM_1672 I L P E R A P G D R A G I K V N Q T S I S F I K E N Q L V T G S G E K G P A D T L A D T I N A M S A K I R A L P
RANDOM_1673 T E L Q I L L K Q A G T Q M D I L H L I E A I D V G S D I R F V D L E F T P S P L L G L S P G Q Q E Q Y S D K F
RANDOM_1674 P D T S D L M E A S S I S F I K S D E T L T D P L F R V D L G N N V K V K W V P H G Y A L E A G K K I N V L V A
RANDOM_1675 K V L S A N L S G F F L G K K L T Q L H C H V R L S S Y T G V I A R I Q R K S N N T A R V T E V L S S V A V P L
RANDOM_1676 I F Y H S A I E P K R D E P E K D E A S G L Q L R F K P E T E N G T S D S A K D D G F M K R T A D S A F P Q G T
RANDOM_1677 D N S P S T N K S Q S I V R K K I G G Q L A H G V D C L G M K K Y H R L F F E G E P N G G E T N A I N G L Q V Y
RANDOM_1678 V R D P R S C M D A Q V G N G F T D V F M I A N S Q F K G E R S R G N W I V K M T E A Y R T S I K L P P K V A R
RANDOM_1679 I Q L G D P V V L Y H V F W G I Q V I I V V M E V K Q L V V Y S V F R L S N C L G A E D I T P N C Q R L V Q T G
RANDOM_1680 T T S T W Y Y K A V F H F V D V Q D S E V Y N P S E F V P D K F R D A A K L Q I H Y F R N G L A V L S S F L L L
RANDOM_1681 P F V P M L T D H R D T R L R W T A S Y L L L F Q S V L P A V G V F V V A Y V L K G I R G P N K K P C V G F S I
RANDOM_1682 M L H D D D D F S G L V F H Y C L S K G H Q R I N A S I Y A P P D C F L A V N A D S K S R I Y L D K L L A C K E
RANDOM_1683 T E T V S N I K Q A V S T V I K S P D N G P K L A E E W S L A G M V Y K G L D D K L A E V C R T L T K M R E S Q
RANDOM_1684 Q D M S S S A V G A A P L D S G A R N L S F I I T R A S L I A L I N Q F Q P S L L M F P A L E L A I D Y A A L Y
RANDOM_1685 P D R N L E E Y F A P V R S H K L M E C G Q Y N V L I L G R R K P S E H V S Q H S G Y N M G A A P E M P T T Y T
RANDOM_1686 D D W T S H S K L E L Q S Q M E S I N R Y Q S A K E E V P L K P M V G H M S L P G G L V L D C L M A S I Q S N C
RANDOM_1687 D P I N D A K A V A V L H L K G I S Y D P Q L G L M L Y L I V E A A Y L N Y S V I Y T S E K Y K V K C T Q K I V
RANDOM_1688 C V V N S Y Q L P S E G T R M L T E T R L V L V L L F D E A T T S C R R G V H E Y E Y L L D T D N R A I G C P E
RANDOM_1689 L G C F V L L L D K L L F R G I P S P T D S G E I G E A E K V R L D D P M C L Q H W Y T E P S N D D V F K S G Q
RANDOM_1690 K K K D Q V E T C V T D R D E T I Y S L F A S K K W F Q L K N A L P S N M E D A E Q Q G L H Q P T L G G G A F E
RANDOM_1691 V N L R S F L A I F W Q T Q T I R A W N L P Q D E L F Y L Y A R G L E N V E Y K N T E Q V G R M C N S S G C L P
RANDOM_1692 V T A V P I T T V Q E I I M I R E D F A V L A D D L Q Q T F E T Q L K V A A A A T L R K Q E L S G H S E E R S A
RANDOM_1693 D N D D S S S L K L R R V Y R C R D S A A Y V F M A I L G W P V L G A K L V G W L Q R I G L K V G N Y Q G V V N
RANDOM_1694 E R H D I K E G F G Q K K A L L T G D S Q P T S G T V S Y V E H L L V S C G N P G E Q V I F S I P A E T M R L T
RANDOM_1695 R Q Q Y V L E S P C S S A G S S A N V V T L G T K H H G E S I F V G L S V V P S I N P T V H F D S A V C T I I T

RANDOM_1696 Y A W L I Q S V C K D I W L W N G G E L H L L E S D S S V V T L M S T A Q G R E S S Q D I I G P F V A M M R L L
RANDOM_1697 D F F N D G E F D Y T S G Y S G P S Q G A P E N T H T E C H R F V Q H V P S E S V C M E G N F V G D I W F P P P
RANDOM_1698 S K F A L G F A L L C N P S G Q I T E S S L D S Q R E Q S T I V N K M R F Q T Q V P Q W T R Q E Y R E V T V R T
RANDOM_1699 R Q H K F S S F V E V P D C I F Y T R N E R D D Q P E V M V Q S A R I F E Q S G S I T V L Q Q T I L L K H E L L
RANDOM_1700 K K Q T Q D A V K A H L D A A L S D E A V D G E T W I P E E G T V I T T S E R L T P V D I R S E V S D P T V F V
RANDOM_1701 E S I N F F D K N Q S N S T D E F L L A D Y D S A D G Y E V Y F S V P T I K K R K K I T V D R I Y V K V R K E L
RANDOM_1702 N M Y G S D S H K K N I A K A P T W G R Q G L L L Y F E K G G Q S A K S C D P N P R Y G N V I Q L T A V S T L C
RANDOM_1703 T C G K S A V K I R D D D L T V V K G S C S L Y K Y K E M T A F L I M R Y A H D A N R G E P L Y T E M I M P F F
RANDOM_1704 S I R K A E V A E I L V K I N G G R N Y N V K D D Y R Q L P G I N N P F S P C E T E S C T D Y T K L D D A R R D
RANDOM_1705 N S N C Y H L D P D Y L E E W L E Q A M N D V Q N A R R E R Q L M E E Y L L I W T L V Q V T S T K G L V A S L Y
RANDOM_1706 T L G S L I Y S H G L Y D F A T G Q P L I H F S L T S T T P E R V G D A F G G D I E H M Q P S T V E L E S G G L
RANDOM_1707 A G H A G R R N R R E E A L Y R A L R L Y A S R L F H I S W N R E K N H K V P G R Y V S Y T A L D K I S Y W L T
RANDOM_1708 D L L L K F G Y C E M L L R T G H F D I I D W V I T N I D F A V A R G L S F Y D R V R G I L R V L S A F I G H S
RANDOM_1709 R E E D E V R S V H G W L S L E H V L S K I K G G F I A C T R L G K L H A A S I G K M S K N L S L T L A E P L Q
RANDOM_1710 A F F L T R I L L C Y G T N L M A M Y E F F Y V S Y L L T L Q E V N N I H F T I E R F A G Y Q N G G D C S L Q R
RANDOM_1711 M G N R P K V A P G I A M K R E S E D L F F L D P L N V L T N N T M P S I V W D G Q D V N I P G P A P S G W G T
RANDOM_1712 S Q T Y K R N Y D F L S V Q A H L G F I G Q S K E K A K Y F D Q S N R C S G F V K Y Y S E T V E Y G L M M D V E
RANDOM_1713 G R A F A I S Q L P N Q L G G P E N T P T D S L L T V K A V S I M A A T A Q C F R V A L K I V V F K G I G L A L
RANDOM_1714 L Q K V Y H E I G P D K G R A E G T T T K S I V T V T L D I R V G D P L I L L Y S V R L T T R I M K T A Q M N A
RANDOM_1715 Q E A Y G Y E R G G N Y K H A I L H D V A I T L K I S G K I G G A V P S S P N H V A S V R S R G H G M A I L Q N
RANDOM_1716 R G P V F A D G H E A Q P S T D T E A T G H V E T Y D G K S L T I D I S A S P V G Q T A V T F M L V S Q P E A K
RANDOM_1717 R G S A E D D A T K K V C G E F A V V K S A G G N P L D G S A Q P C V L N F K M L N L A S D L K Q Q D A V G N Y
RANDOM_1718 M V S V L G Q T A D Q E A K D E D A Q Y R Q A R S C Q R A A Q L K Y S D F Q A I P L V F F E Y G T V T I G V A P
RANDOM_1719 M F N V L K H N Q K V P K A I Q Y R P T W A S P K R L S G A A L E E V Q Y V I T R E I S T F N A F C L L S S D E
RANDOM_1720 R A A S G L V H E D G T R Q V A M L I C G D L G N Y L V R P C K G I A Y A V L S T R R I L A D D F F D C F V G I
RANDOM_1721 R K P K Y N L F P S T W T G T A L D V F R C I V V G S S F G R Q R I E A D T L F V S S K P Q W Y I P K K D L L G
RANDOM_1722 W D G S R Y T P S D R K G E A R C T Y L V R F V A E Y Y Y A M H Q A E K N A G C P K G V E T G A L I S E R N E G
RANDOM_1723 N R S S M A G P N F Y T D R A A S P K Q S K F K A T T K Q D C D H V A L A Y L A V G K K V V S K R Q V D Y C P F
RANDOM_1724 P V I H S D R V Y C V V R E D L S E A A K R V A M V N E G I H F I V D I G E L Y N F I A F S V E V A R E G Y G D
RANDOM_1725 K K T I T E A Q D P R W M G G S N L L S G N G V R D P K A W R Y P N E A G S T Q A Y M D N V V N Y P R G L P T V
RANDOM_1726 D E E K N N R T K A I N I K P N A S T Y H Y C R P S T G L G P T K F P A S V L G I I M L S C Y A N K K H Q D I A
RANDOM_1727 T Y G R F P A Y Y S N D T M G V Q Y R R R C S N V E E A K F E A N F L G L K E C E F E L E K E S I Q H F C D S Y
RANDOM_1728 M E V D H G G T F S V V I F I N E G S K I T T S A T E A G N N W G A K A I F G A I F F A Q L S K Q Y L Q S E L L
RANDOM_1729 P V Q E W S R L P G I S V T N L A S T Y Q S Q T R V T Y G V N Q Q P Q L L N H V F C A Q S S N V V E Y F L L E I
RANDOM_1730 T P N T Q E L V V L H S V M P N E V K D Y V E Y S K Q A L E R V E G M W L Y L V S K Y E C N N Y Y D D N G L G S
RANDOM_1731 V L P L C N W V F H T A N L A G G I F L K N A A K C A E L G L Q N T Q L S R C K A L I F P G F G Y Q C K E F L N
RANDOM_1732 G Q H V V E S M G L A H E H C Q R S N L G V R L D P T I T I Q P D V Q P N F V A E D I E G G G Y V S Y R P E R N
RANDOM_1733 T L M K V R L Q K L G I V L K L R N R M G D A D A S Y L L A E K R H S L Q G S E P G P G F A I A T G T P G T S L
RANDOM_1734 T H E S N G Q R Y K A S P N S G H V S L K G A E A D G A P E E R S A Q E D K L V S L R T D E Q P K E I I I L S D
RANDOM_1735 F E I V L R L L S R V P A P K L R P I A T N I V G F D A R S L R N P I D Y Y I T A L P P P Y L M T S A D Q L C C
RANDOM_1736 D E T F L L I Q M A C A S P G N Y H F Q S E K M F M A D F A T L E A D E R G Y E A N R P N H P G L E L G F K F N
RANDOM_1737 L L V E K K G K K F L K S Q F G T T R F L M N R V L T I A N G Y P H I L L S Y S V M E N A E Q G I G E M L N T I
RANDOM_1738 Q L D Y A K I Y G G T V F V L E S F K L M K E A G N A K S S Q C D Q I L V Q P I I P S L F A G L G D I L D A L D
RANDOM_1739 D G E V G G S F V V N N N E E S K D C D D T R I Y S S S H N R T H G G Q T K P Y A K A T K G D Y F P A P E V H M
RANDOM_1740 A F A M E V I K L H A L A P A L G S R L A R S N K P F N N G D E R H I R E A G G E L S H V Y V K D V E P V C T M
RANDOM_1741 D Q F E I E L S E D V L I T W S W I S V F D S L T E E L N Y S L V P D D A A L Y H F L N F L C I G K K N I R T N
RANDOM_1742 Q P L K A P S D F D S G R W V K K V R E V F F I Y N L G G L I V I R L L L V N E Q M G A V I A I A F S G T S V R
RANDOM_1743 A L A T P G E A T L L G L G Y E T V V P R I I Q P Q G L V G A K V T T L L L M T D Y R A H I N S V R D Q V V T V
RANDOM_1744 T R D D F G F Q G T E R K K M R K I A D K S C C N H E Q A Q F E A E H A S E E Q V Y C R L Q V A S I L I L I N Y
RANDOM_1745 Q A A S I N E S S E D S D A D P E H I T I V E G G E D S F F Q L L T R K V A E P Q E D K N R L P S L S A A A I T
RANDOM_1746 L M V E P A C V P Y S I V K D V C M P Y D F P L R L H K L Y L Q Y S I A D Q E K E N H L L R Q A K R S Q V G A T
RANDOM_1747 K L F D F S S D L S L D Q K R K H R A M L P S L W A D L N I I K R H L D V L S G D L N S R I G L T G S T K D I R
RANDOM_1748 L L R S A P V S D S E L P Q T K R I A L A I D D A Q D T T H F T K V F R E K L R P K A Y K C A F T L S F L A E M
RANDOM_1749 E T D V M D G G R R R L T R R N E E M E V I R F D Q I L P Y L E L R M G A V I G P S R D R D T T D Y N L C E T G
RANDOM_1750 K L A L R V R Q G I V W V L A K R L A N Q Q F L H L G F L P Q I T T R W P P I N V E R Q L G V L W H V A T D R G
RANDOM_1751 G A L V W N A P I R R V G D Y T P Y R E Q K A V A L Q C K F L T L S R I E R V W M T L S I D L I G N E N L F K R
RANDOM_1752 Q K G L T A E D L A A K K F P L F I K E F D S W Q M A T N G D S I P C L A V E C D G Q L D A H L S Y V W G I K Y
RANDOM_1753 S P L A V P A G L Q G T Q P S G A Q L I G G D L L A A N G A T S K D L A Y V A G Y A T Q Q V E E Q P L N M A N E
RANDOM_1754 I I K E S L Q V G G V F G Y P N N S L Q D C I D L A V S L I A Y Q I G T L V L M N G T F A I F Y N W S P K A F I
RANDOM_1755 S T Q K N L S G P V A K T R M K C P V F Y N L I H R K S E F E P A Y L G P V S P P L R E Q E V D P M R P R P E G
RANDOM_1756 T Y D R L G L T T G K A Y F H L T N S A L G W T S L Q K K N E V N L Y I V F I H V A K H P Y R A P D E V Q Y E G
RANDOM_1757 D I S S A Q G V R I E N A F V L K G R D E E G I C E N D E R D R A C Q R G A S N Q T K S A A T E H E L V L Y F S
RANDOM_1758 S M R P R N L M Q K T W I V A Q D Q A M L N Q R F A A M R L E R R S D E V T I V H V C V N L G A P R L V G F T L
RANDOM_1759 I F R H I E G L I R I L L K E A T D Y A M N R S T G Y V L C C I C F S M T I G P V S R S S E I S E L F I M A M Q
RANDOM_1760 E S L D S E D T A D V E Y A W L Q T T N C I L E L Q I S T S Y T V R A L I E Y E N K R I I A P S V V V C F R G Y
RANDOM_1761 E N G L A Y C K E L Q T K I K H H L L Y Q P L I L Q K G Y I I N T D A P E K L G F L I A R R F V E S L C K T F D
RANDOM_1762 H G Q E Q G I L S E S R A Q P T R T I V S P I A P N G Y V N R A K N D S G K T K V D A A G L S Y Q K M G P D V I
RANDOM_1763 L C P E S P G L T Y E L L L L A T I V I Q E D I D V T R V N T V L P L I D Y V A F G M Q D K L V I R L A F A D A
RANDOM_1764 D I S S Y Q V T F L S G I S S T V G Y R A V R K E S I D Y T V M V V E T H P E K R I E R S N V K I E N S H Y L E
RANDOM_1765 P P G L D L Q A A D K Y Q Q Q E E W K G Y G G F H C W P S Y E Y V N G V M Q A R L A R I Q I N Q E P L T I K A L
RANDOM_1766 N E K E D G K G R Q T T Q T Q F E V D G C E L V E G N M W V L E M T I N N K V K K V R N S R Q Q D D D M T Y Q S
RANDOM_1767 I P N L W P V T F C G P L V L A Y K S A G V D T C T M S S A Q S A V T T L F V Y I T L V A L K E N G M E Q L A D
RANDOM_1768 D K W S I G G S D F L D I L F Y L I K R Y P T S S V L S F P Y S R E K D T P T V D G V G F G Q Q C Q L V A I D A
RANDOM_1769 D I A A T L F S A K L T L K T E Q I K N E M Q A S R Q I M P S F Q D P C A G P Q L Q K T T I L I R P K L T A I R
RANDOM_1770 V G K A R E L K P V R Q L W R A V L D G P L P T A A C Y S T S S C V I A T C A N K K Y E A D G A L H H G P C D V
RANDOM_1771 A D E Q V G T D G E G L N A V F F L E T M G S A T P Q R P E A Q T F R L T C A L S A S L Y K A K K S A K Y G Q K
RANDOM_1772 K T W R E F K S G P G W Q L H N S V V I D T I S G A G A P Y F L Y K V M N P S V D S F Y P I W A T D Y R G T A Q
RANDOM_1773 L G C L M E K R A S G E N F L T G P S S P T A E N N P L I A G M P R S Y L T Y G E S N I N V K P C Q H G K N P D
RANDOM_1774 L A E K P T P D N P G Y V H K T M R T P L H V T T N I D R L N L A P P S R A S G G P F P V Y K T F V E G G T A D
RANDOM_1775 E M S K A D R N G I K D N S S P D A M I Q F N F A G Y I Q K S T S L N R S K E V S T K A L A P C R L E S Y E T K
RANDOM_1776 D N L K F G M V T Q N A G D L P T S R E V K Y K L F L K K N F N P V K R S T T V I G V E D M S D L G T N R E I L
RANDOM_1777 Y N V D G L K L S N Y G T I D L D A V P E A K L N D H P G E L E E D V N G G A I T T R P T K S V F F Y E L I H D
RANDOM_1778 E F G T P F E I E I E L K Q P R F L I D S F T L Q K S H E T L D G F T V L I Q A L G D A M A D H R A R S K E G N
RANDOM_1779 K L Q G Q A E Q F I I W H N R I D L E I N W R V R V I S A D R K L S N L D G F C T L V L C L I G S S A F P T F K
RANDOM_1780 V Q K S F S S E W T F K G A D P N E F N E G Q T T R A S Y D P N W A R D E S I V R S E Y V R Y P T L V A W S G K
RANDOM_1781 V P V I D M T T N E G R L N P V S K F D V E P Q S A M N F H V E S T E G A I L G K A L T L M D E K M A D H T Y L
RANDOM_1782 K L V I L S V W G E M T P K D T F V L I V I S L I S L V A A K L G S I G P N R E F I P I L V N K K I Q A T L A R
RANDOM_1783 G L V M K L R E S A L P P C K L S F K R E G A N S L L D I N E G L S K Q S S L S G S R V L K L I G D M H A S Q L
RANDOM_1784 G L S G A E S K A Y S L W K N I V T F Q S H G I R T K P W T V A V S G R M A L L T P T F P G E K F N I D K M G A
RANDOM_1785 N L M V W K P R K Y G A R E Q A D P E A S P S P A K T E A N E A L E K A Y L Q L Y P V G G L T W E F H G K L G S
RANDOM_1786 T L S E K A P P V G R T E P N L E N A D V Q D S G T L S S N E Q L R P I S F G K A Q L Y E S G S V V V L V R P Q
RANDOM_1787 K E H L N L L S R Q Y G P G K H G P N V N V A E G C T H L H E W G S N P T K E G V K P E N W N T D A F D H S L I
RANDOM_1788 T A L N L P D E D I R N Q L M Y D T T L A P E S R R S V G D F L R N L P V M P D Q M S I L K P D Q A F D M L D F
RANDOM_1789 M H D G A V A E V G G A M S E R G Y N E I E K T V Q N T I V D L S Q Q F Y A I S P N I Q G C D G R S A V G F G A
RANDOM_1790 T E D D T A F R L Q F F C A A A F A H T G S F V L L Y P V T Y K Y V S A L G Y N I F P P H G H V A Y G Y V E L T

RANDOM_1791 I P A E G G I W I A L I P N R G A F R Q L E G L R E A M E A P S Q G G W P D Y S P D L F D L V V E G K T L T G G
RANDOM_1792 S M C M C R R P S F E R A S A L V L K A V D Y Y E V K G L R Q T A K H W G R R E M V V D I I S C I M A G L S K S
RANDOM_1793 V H Y T L H R P Q K N C L E G D Y N M L I T K W R A E E A G S L E V R I A K T I S P L N H W D R T D Q W C T N T
RANDOM_1794 P S R V S L S T S T L I Q D I M S C V S K P G L S M Q N T S I P E T K L E A Y K R A M E I T S R T S E K H E E S
RANDOM_1795 I T D S S G N L D Y V R E Q W S G G Y T L K F G I G V K K V T G K G A G I S T K I D K N P Y N S K Q T V S E A G
RANDOM_1796 A Q R S G I N K A M G D G C V P A S Q G C Q G D K E Q S V T A Q T T Y S A V F T E F P G Q W D G I A S L G T L L
RANDOM_1797 L I L D H I A Y N F Y G L K Q D M V L L N A D M F L E K G S H N F D E T D G Y P R L A E N E M G A I S I K A P M
RANDOM_1798 C D K P N A S I Q V L L V N L Q A T D T E P C I K S D E G I R T L T K A P A P E T S G E E A R I S D P R Y A N V
RANDOM_1799 N G T I I L E H D D A I T S L A P H P V R R A K T V A P A P P L N P G P R R G W R K T S D T I I I L C V Q V P R
RANDOM_1800 A A D S G F Q D L E V S K E T V T E A Q Q V L E G V P T V K T D I D L L D C Y Y F P N G E I A G I A E T C S L I
RANDOM_1801 L K H A V M E N R F D K T C L V L G G S Q Y E K F D F K E R H I S T V L H Y A M D P C P Q A K E P I V I E G F D
RANDOM_1802 Q R P S V R A G L P G T S D E D S P E N H P Q G D G K S L L A P P N S T L S L V D P T Y P T E R E P E L K K G A
RANDOM_1803 G A Q G L R L Q D D A P H S P P Y F T E T L T I S E N E S E S G P Y K T T H G E E L I Q F E W T V E C Q E S S V
RANDOM_1804 S Q E E V C I T A E A D W E V L V P M A T E S Y T E S Y P K Q D L Y H E T I Q D Y Y D G A G V G D I Q Y T F I K
RANDOM_1805 P S Y S P V M I L I F P G W G E P A G A A T R S K A S V Q I A M S K K D A E D L E A K V P Y S S T V G G F R M M
RANDOM_1806 F E C Q Q P D K A L Y V A N F F F G G K R F G G T L W F Q D W L Q G A L E F T N K K K P L H M K D A S D G E C E
RANDOM_1807 F T L K G A Q R V H K K D A F G D G K G N D A T L G E P K L K F D R V P F V G C T L K E K I P P P H E R V G L D
RANDOM_1808 V E I H Q E V T L D T L R F F V T L K H T E G P K E S S V V L L I D V Q K I S L C Y K F A K D Q A L I I V E W A
RANDOM_1809 P T G P N E V D C A G E G F R A R V V V L F I T S E L Y S R G E T I P E H V D S Q A F P A G G I K A L K A S S K
RANDOM_1810 L A L Y I D K W F A L P P E S H A P A E I G A D I S K L R H S G L V P M E E V K L N Q G H F S A W D Q L Q Q F F
RANDOM_1811 A G M L S S S L S W G P G P L I S N T E E D V Y L V G M V S H E K M I P I Q S V P S R E Y Q D A E S P P G A Q H
RANDOM_1812 N D P I K A P P D K K S E P L G D C V S A T K I K C S P H P P L F G A Y Q L S K R L L A E V F Q A I R L T Y Q N
RANDOM_1813 Q E Q I G G K T G Q R S T S L R K Q D K V N G T F T W F Q T F K I D T I V V N V I S G P Q H D V T Q K N D E I N
RANDOM_1814 Y R E D E K D R T Q I P L D F Q T R A T I V N R R A T L A D L G L A A I N V A G V I V V T R R N H L H G G L Q S
RANDOM_1815 I Q N V V G L S T G N Q G P T H T G T V T E I N N P S D N G E I T E K P L R F N F N Y S S W L V P L V T I I A V
RANDOM_1816 S D S D L F N Q D F T M E A W D T M A A T L M Q L G E I W A L A T L H E D N R V K S L L F V P K I F L G V P L G
RANDOM_1817 V R A R C D C D Q N M T L V S T W G V I E M E F Q E Q G P A V I F D M Q T L T K L P T P L G S V G R A L E V C L
RANDOM_1818 R V I L G T A P D L G E S D Q G V E K G T I Y E N A T L E Y T P A T A R G L R T Y T G A I V F F A S R D L G P G
RANDOM_1819 A F E K R A Y Q V S A E Q P Q Q G A Q L L A N L Q R R K E W P V E F Y K S L F I I L K Y A R Q A I T L Q K T W S
RANDOM_1820 I P G L P R R Q T L S N K G N K N K N G L L M Y P S R I H H R E D T Y R D F L V K N Q V S E N W S R H R P M D N
RANDOM_1821 F V I Y I Q R D V V R A E L A L W S E W G E R N E G K M L Y H A L E S A V D A E I R V F E D V Q K P Q K Q P F N
RANDOM_1822 K I F L E F R S Q Y E A Q G K A L A V P A C D A P T A V D D S G L K C A P K P L L E N Y E A Y Y G Q R S S N Q V
RANDOM_1823 R M T A A R R P D I S D K K K C A T Y V S S E D N L K L L L V S L A T T Q L P P Q Y E K L D I E W T P D L V M
RANDOM_1824 N I W Y T A L N I F T P V D A N L C E P K A T L T Q G G K V F E L D A D G L T A G Q T T I N V V G R L E E D D W
RANDOM_1825 M L L K T I L A M G I D E I Y A A D V L S L A S S P K Y H C P S E P A A Q G N V E G D N L I A P A K Y F G W G G
RANDOM_1826 W S G D R Q S S E T V T L Y S Q T R C A N F G A K D D P A I A S A S P K Y Q Q W F L D Y F H T A R L T T M L S R
RANDOM_1827 E G G E S W E V G G Q K D A K D G L V Q G I S S Y P C D K K Y G G Q L M H A V R K S W A I S S R A C H N S L G A
RANDOM_1828 P V L D P S S Q N S R S D V R A E S C D H T I Y Y Y A Y T D I K V A K L N H V D D G P S K V D Q L E T G I I T Y
RANDOM_1829 D G A I F H A L M I G F H N F R D H E D L L E K I L V E S T P N L D N G K S E P Y A K Q M N V W R P V P H S N T
RANDOM_1830 Y K R P P F I V Y A A Y E P I A R E R I A A E D G P T N F W S S G T K E R L S A A N Y A I A G G R E H D W I L S
RANDOM_1831 N A L T Q S T A V L W W Q N F Y P S G W C R I L S Y A E D G F I G K A T E P G K L C L H S R M R Q D Q T S F I A
RANDOM_1832 K K P M L K P K P K L A L C Y D D G Y A I M F I A V S V D S Q S V L R G C T Y E N L G Q L T V S W G I Y R L H G
RANDOM_1833 K F P D Q A T A T G R T E S A P D M I T L Q T H A A Q K G Q I E Y E A P W R H G L Q Q V N D H G L A L L A N T S
RANDOM_1834 K R V P N P F R L L R R R I V V K A E L L V I S L V S N I F M I V F K W F Y D T C L H Q P E S N T K D G S Q Y K
RANDOM_1835 S A K V T Q N W V K V R E P N E S E K A Q P D A N F E R L G L G A L K H K E D E K R G D A C P K L P A Y E G V T
RANDOM_1836 A K F M E F A V A A D W A Q Q P I N V P V P I E K R D T L D E L V S V G L S S Y K G M L A E L D L N D R C K F V
RANDOM_1837 G M L A G V A A D K D H V R L S C S G W Y K D S T N G K A R H V P R E K M T T Q A T V G G K Y Q N R K S D A S D
RANDOM_1838 A G D I Q Y Q A A E K Y L Q S P P L L D Y R T L P V N V S G H K Q D T E C G G P K R I M A F A Q L T D T D G F Q
RANDOM_1839 E L S S G T K W L L M S E R W D T F L N G I R G G D F F D I S A L V T D R P F N Q A T F R W A M L N F N G A P V
RANDOM_1840 V A L Q R D P Y H G D K A C Y P T R Q V S K L T L E A L L A T H H F V G G S T T R S Q Q R Q Q P F L P Q P G P G
RANDOM_1841 I D T T V Q S S T T K V P L P E T T I S F P T P A G G F A P R P C G E I T K L I D H E E T D S K R E A S I K L P
RANDOM_1842 R L E A G Q C N Y V R V G E E P L S N N D D S C R Y E E Q S S K L K T V P A K P S R Y Y V V A P E C G R P V E F
RANDOM_1843 N Q I W F E S N S L N E I G A N K P D C L K P I P V A T G I S L E I D K I H S Q P V L G F K R G A L H R F S S W
RANDOM_1844 S L P K L C P P V G T R P V E M T F G A S G D A L Y T S V E L G V M T G F K K V T A S I I D Q G I F Y D I E L K
RANDOM_1845 A H G K A M L W N F V T Q W V I T T Y A V T L I Q N R F S S L I G I S S E S S F N G S T I N L M E Q G F I F M I
RANDOM_1846 Q M N A M N T E G I E N L H A N L K R D E T R P Q N A L F I I G V H V K V V H G L G R T L K I Y K V L I S I R A
RANDOM_1847 F Q K V I K N L V F S H D N D L G G Q L G A K A D A A D K G G T F A F H R S D D W Q P S D P N S K L N N A S E V
RANDOM_1848 I Q T D E V S E P A S E K R L C Q A D N S Y T G H S I L A D V L E G I A E Q F P L S R I V G D L E Y L G R N E V
RANDOM_1849 Q D E Y S M A N T N P N Q H G L V P F K L R M Y Y M T W Q K M L S I L A K W H C L I P T N E S K K S D G S E P P
RANDOM_1850 H V V A R P L A N L D Q V C M C F P G R Q S F V M L V H I F T R F F T K D I A I G F T S T S F K T R C N E E P D
RANDOM_1851 E M T A L L N V D V A A K I H V P F P L D P M Q G L E S T E K G T A G T V I N K N G L K I W D A T A I P A C L V
RANDOM_1852 T D T Q S P I L Q H F E E N S A M R K A K E E Q G E R G Y Q R S L P F F P R G Y G T K I L Y S A E N S R A F H H
RANDOM_1853 L T D T Y T S H I L I P C V H S N Y T R S F H I E A A K V L M K N D A D M V I L K L I A A Y Q S P V Q V Q T D C
RANDOM_1854 I R T M R S E A C G D K R E D L C I K K K R G L K I E T L L L R D R L V W L S E H F G F T G V C V D D P R T N N
RANDOM_1855 P L F T Q R G G F R V H K P L V W N I Y V G F I R V A G V P T L N R S F F T V H A F K E G Q L F L P I A C T K D
RANDOM_1856 Y Q V G K T S G M G I S G L L K E L S L R V L M A A L P K T D G P T T E F L L I T A S V S C N S V N G L S S N S
RANDOM_1857 L S V T G T K M L I H L K F Y A T A E G F L R F D G R S Q K F L A R D V N S D S P H D Y F F M S F T K K G G E E
RANDOM_1858 A N K Y C A Q Y G Y S A S N K V T T D E A F S N A Q L I D R G W S L A T P S G T P S C A D G C D V A K D V K I D
RANDOM_1859 P A G A P D Y T I N N N A L V Y W G L Q A G P C N S Q L V R K I K T N E E L A V D T S T G P T P I S L V K R D A
RANDOM_1860 K V S N M E L K F Y C T V R E K S L N A S K P I A D K I K Q D C D A I R H Q R I A N L H K E L R S E P M P C V A
RANDOM_1861 Y D A L R D C L I P N I L A A G K I R L A V E R L A V M F W A W P L P D N A Q E E G N M N I G V Y T A S D G R A
RANDOM_1862 K A R S Y M L Y D N E F G E E F K Q G Q P T Y P F G S F V D D F C G S K T V T G V D G W P S T A T D H S S L W Y
RANDOM_1863 I A K S I F H H C A G A C L A I F D E L M W E T A I V P L S H V E H N I G Q Y V S N R L L N F A F T K K N G V G
RANDOM_1864 S V S L S S K T I G R A F W P E T F T I V K S F E T T S G L G F Q H G T L Y A V L P L V G K F L V D I E I K L A
RANDOM_1865 S A C D A G Q D I I E V Y K S P K R M R S G G A A C T V L L R A F D H E M S N N S G A F A E Y G F R A K C L Y S
RANDOM_1866 L Y L C A G T E V K L L M V T I E Y N H V N Y R P G K C G P V A D K Q S P R A P R E L L F E V L I E S P V V T N
RANDOM_1867 P P Q N W T A V A W A A A L I G L M P F V S Q D A S L D L S T R A G G A M T Y T A L A N A R A I G I N S G E P I
RANDOM_1868 M K F M D D Y E I A D E V R D P R Y C E K G S P R S I W A E M G L G D G V G K I E T R K L N G M E C C N E G D K
RANDOM_1869 I R E D E N H R R Q R F S R H K D L A Y M C Q R N S L D K P I R S A D P L G G K Q V A T F A H M E F H E K V G F
RANDOM_1870 E L A F S A R Y L L F A E P L P E M D I Y N P L S G M P A L Q P R V W V L L A I E T N I F L A S N K P S F I I V
RANDOM_1871 S S L L R G S A F V I E R Y K E A L G Q F V T A S S I T V A N H V Y A W F R E C E G T F Q T E R E I E A K L F P
RANDOM_1872 A P F T L T F E G K L E P D M M P R N G L F G I S G I C A T C M N V E K G L F L I I K Q L K P I N W T V A N S M
RANDOM_1873 S A Q I P A P Q G P P N L A G I M Q E D G V K R V F H C D H V L A A V G A T N V T V Q D N V C K R R D Q D R Q S
RANDOM_1874 M G M N G Y I V G A F L Q V N Y I P G G K V I S N L L Y L R I T A R R D L S D D P A C R Y G G L A V M R G P M P
RANDOM_1875 V D E A F S G I V L R K K D L V R Y A C R R N D N I G P P V K H S I T Q K S G H G I L A K C L N S T N P Q E T I
RANDOM_1876 K E F A L S R I K K A F A G D P R S I L I V V K D T T D S H K K V L T T L M I K T G L N E T L F S Y I S E F E L
RANDOM_1877 D V T S V K E Y Y A T R L A T N T T N C Q I A L L D P P A A Q A F D T F F Y L P V A I E A T V P L G V D P R P R
RANDOM_1878 D P K P Y L N F E K F S G V M G P N I V P V G D L R S G L A H P T P T S I L N N L Q S Q G K D Y V N Y D H S E R
RANDOM_1879 A T S A D K C S T E E T P S E S Q S D L K L R M E N A K N G A G L G D I T A F N K I I Q T N F T R T G A V Y T S
RANDOM_1880 L R L H M E Y I E S I Q A K F E A I K P L R V E S D N R R M V D T A M P Y E F L V S A C A A G I W V D L N T L I
RANDOM_1881 L P D E R T Q W E T N D N K D D S F H I G S Q G T H D E A E I N T S C S G W S S K Q N A N I L V L V V C E K L T
RANDOM_1882 R S K A S Q A F M L V D A A D S Q T V N R A G S F A S V A A S A G C L D R E V L E T G P D E M V N R A P P V A F
RANDOM_1883 G F A D S P K Q K I M S N E C V I L N S A I L R L A V K A S V T R S E R V I A F Q N A S Y K I H P H L L A L Y F
RANDOM_1884 V L C G T F K L K K Q N A V S H K E G E R G L T T R R P T M S I V A F F L S K L Y S I V T S N A E A I D S E N G
RANDOM_1885 L T T W A S R C L I S E E I I P N E E T Y V S G D E M Q A E T D K P D K N Y G E F V Y P D K P G R T A N R I I R

RANDOM_1886 M T I R P R P K D P C T V R M E E P E K R G E T R F L V N L P T A K T D S R E Q N V Q F A M I Q A A F P E R Y R
RANDOM_1887 A S K S A T P G P S K N E K D N L L E V L S A Q L P E A L F C F L V S A R L R A L A L I L V I V C I K G V V D G
RANDOM_1888 Q M S A H N T V I E V V N N K A C C A E Q L E E A N K P V Q I S R V L M F G T D A C M T R P S K L S A T L D A Y
RANDOM_1889 L E E Q P A Y E L D E P W F H E V P D I F G P Q S T L D P T D Y V S R S L Y E S I K F V D M Y G S Y M N R G G T
RANDOM_1890 I S G D E F G K D L M V F V R A R K I C P Q K A V P E L K I V P K V P I G M E N E G D T W I K V R A A L V E V S
RANDOM_1891 V V H G P P R T C L G T G G N N Q F L V N V A K K G F I W R N G D D F L I V T W P I S I N P G T A I A T P S I G
RANDOM_1892 Q E R V K A N K F D T F S L G L T R I F K V V S T D E V K K Q T P L E I G E V N V R G V S G E A Q P I L E P C E
RANDOM_1893 S G N L A D Q N V H C H G A F Q P H Q I L D S I N R R I P A Q R G R E G D K D K N Q E D F H D G D D G E L V S T
RANDOM_1894 A I D V T S K A F S R I P E S G P M D V K V G G S L I N H S S V D Q Q S V L G N L L I H P I R T N H A G I D H S
RANDOM_1895 L K L P A I Q E R L E P I N N F G E N S D P A L A V S P V M N A P L K T M S G R G S S K Q A K N L E L L A C L A
RANDOM_1896 R V Q I P R K K T R V S L Y S S I A G L G K I P A H C F L L D R S M S S N C M K G T P W V G S F S A S V I P E V
RANDOM_1897 R G A A K E S Y N H H G F G P G V S Q V V H I Q I H K A V T G I T Y M G N R E F I S Q T S D A C V E L S V Q T M
RANDOM_1898 S P V I A E R I S G D N L R F V D E K P P D A P I T D I F D E E Y T L I G R D C Q L L R R G Y L G P G L K G T G
RANDOM_1899 K S M V C L T Q F W L Q L N S L L L E P R F W L Q G T H P G R N T Q S N M P G K P W A E R C E D T P D E E Q R L
RANDOM_1900 E E D I L K W R I D L F L E Y C S H Q S R V K A N V D A L V T T R R T S F S E Y R F E T T P R Q P G L R S S A L
RANDOM_1901 H P L G T Y E G N Y G V F S P K K S L Q G R L S N V H R F C A Y E A K L S D L S F L Y F Y M S F V R A D S V K I
RANDOM_1902 R H I L N S L A V A V I Q T E T N L N W P C G A L F C Y S T S Y V H R R A I R L T S Y W A M E S I N S G N K W E
RANDOM_1903 P L S K D Q Y T Q A K Q T V N G N D H S H A I I K V K N R K F T E G G D P Q H A L L G H A A R L N W Q D A R P V
RANDOM_1904 T S S G I K S P W L D A I H P G G R V Q T C G I S A Q K L R D Q R L L W K I S G I Q F C R I A T T G L R K T Q Q
RANDOM_1905 A A A S L F D E K E L F Q F G E D L G R G L M G P F R G Q G R E I Y H K L Q D L F A Q E E S P S D F P D E T K P
RANDOM_1906 Q S I E V I V D K V F N Y P S R Y M K T R R D L P H Q R L C T T I I L L K H E E N N G N A D E A M I D H S A T V
RANDOM_1907 T G Y G D T V H H H A N I I M I G T I T C L N M T L N A E I E A H C F A A G R I R I E R W K A I A K S D G F I N
RANDOM_1908 P Q I N S W E G G R D A S E R Y C S E L I I R Q G S E L A I N G H T F F V P Q P T A Q F K K L T I S F L A A Q I
RANDOM_1909 G S D R D P A N Y Q G R D T Q A G P G L N N H G P E F P S K I D R G T T I L K V T C W N I M I A R L S T W V L N
RANDOM_1910 T D F A E R I S L R E D E R E R D I C D L V S N V K E L K C M Y S N K T K Y T F I G C L V V C T H A G S T V A V
RANDOM_1911 L Q R K K H I G L L G D S T L K N V I T F C K A W G V R S L K E I A V L S G M A E F P K E R A R K F G R I N N K
RANDOM_1912 G M S I L K D V R G I V S N K G W V V I E V V T A I T T V H G Q K C A V P P A K H L C K Q I G M P D K R G S E A
RANDOM_1913 Q R V Q T V P P G R V Y K Y N S R H M L S R S D A R F A T I G L Y T R G Q R Q P V T R G A D K K K P Y N V H Q R
RANDOM_1914 I E L E V E G F I M T I D T A S S A Q C T R N A E Q T T R I P Y L N V L M P G T A E V V T E H S R Q K L R E V P
RANDOM_1915 P P Q H P G A N I W G G R H F G S Q P R F R A L I S S P D L A R G R Y D H N S D K L S G R E R C H H A R P K A Q
RANDOM_1916 A A T D N F K H Q V D K G L L A N A A I V V H K A D F G E F P S V T G R R V G Y A E T L V G N Y K A S L Q D V N
RANDOM_1917 V K V E K I S K H M S D P F L G I S G L A G E Y P T K Q V A T K N K Y S M S E K L T G L E P L I L L D F N N G V
RANDOM_1918 G P E Y V P T K F E K V E F V V P L L Q A S T K L S R S A V R V L K W S E Y L S L A E V D V V A C I R T V Q Q Y
RANDOM_1919 H P M Y I L R Y I N E N Y L K I E L P F P V G D S N Q N A S S P D D L C Y S N P F A H S Q A E S S N P R W D S I
RANDOM_1920 G A R T K P M N R H V V A E L L S V S P V D S S R Y R A Y N K Q L Q Q Y V T M A G K E V G R T Y K A I H D C G V
RANDOM_1921 A I H E V G A E G N T T I E L V A P T Y H A P C T W H G I L H F W T K V T G A L L E G D I M V P G A K A P A E F
RANDOM_1922 Q R P G L A S K I H S Q A E A Q I R G A G S S L L M L E D N I Q D R Y G H I V Y I L G F W D R T A P I I V D P Q
RANDOM_1923 G D E V A L L S L Y R V L L L G P G I E S R I P P K I I G K G D V Y D A A L T D T L F T P T F L A G N E F G E E
RANDOM_1924 K S P K N A E E R K V A I I Y N G W G K L F L E E V L L T R S I S F K N A L S K D V P L V S F G A K K Y K G G A
RANDOM_1925 L L L Q A S E S R L E G I R H A I P V R G I R N V V T P Q A F R M D K P E R L G L S L V I W H R G R Y E K V T D
RANDOM_1926 I G L D Q V K G G A L K K D D S C E G R E D L L A H T Q N P Q C H P T A K H A F E R V T M N G F D T K R E Y N H
RANDOM_1927 Y A L S T L N V H D G K S S L P E P N I W A R T V L A E P F I I G G E V A G K T K E D N G I L R L L R D G A T K
RANDOM_1928 M A A H S L L V K E D R W L W I A C Q L T Y V Q I L L S Q R K F L Q E L V E Q T L L N P N V G L N V A I S I T T
RANDOM_1929 Y L S L W R I P P D D E D E V I P K N F G A D A K L D V S N G L E R P T T A E N A Y R A S N F F Q I P V V E R P
RANDOM_1930 A D Y T N G I L V S L V N G L C G H S C F N T W V N K D E T Y R L P I I V V V R T S Y E S T W E T N A L M F P S
RANDOM_1931 E L R F E E P R F D D D V S P A K V T S V E S S A T P T F V E M D Y H V K K D T Y G D T E P W E P F S A T V Y Y
RANDOM_1932 G K G K V L R M W K R R V K T A S L V V D D Q A Q V T V R D K V P S K Q N R A V Q F F I G I I F Y Y W V S D D K
RANDOM_1933 G S Y T V R S G V Y R V T A I P M T I M K R D E E K N Q N L S R C P M S A Y G G A D S N R K K K S E S V L N E D
RANDOM_1934 T F E Y T G S N I L H S I A E C A I D G K K A S L Q G L Y E P Y F K S S A R R I Y Q R N M D G W P D I L D P S R
RANDOM_1935 E K G D D K K Q M E Q R Q G S E T T I S E Y P S V F K Q C G G N T H A R P K A E P F K G Q T Q T S C I L A I R Q
RANDOM_1936 V A A V P S T L D I L P K T G Q S A F T R I K F L L M H C Y S A E G Q T T R I G L G V D R A K Y E E E R Q F A P
RANDOM_1937 L L Y V L D K T N N D P L R E K Y K L E L Y P N V H A P V G A Q S L N R N Y F E T A P E L L R G E G K C E A M A
RANDOM_1938 E L S D S Q F I L M F P P Y Q C L H H E L V K P Q F I G V I Y C K A G A G T G D I I I L A D Q I L M V Y I G N G
RANDOM_1939 S E N D A S M K L Q C I T F T D L R K S G P T Q L R G Y G S G I T A Q P E T I P Q P I Y E V K E Q G D G P G D L
RANDOM_1940 T R A K Q P C A V F F D Q H V N K V D A T H V S S N R I E E A F C T G C D P A E R V L F S M E P H Q L S V A P E
RANDOM_1941 Y P T L Y S I A I G D E A K T L Y E Y L P K K L K E T L Q Q T R I C E Y Y I A E E W S T E L S T A S G N L G S A
RANDOM_1942 N D P L P G A N E I V L F C A Y V K I A S M V K K R P F E E A G A Y L A D D E G E V S L L A H T T R G S G D V A
RANDOM_1943 A N L G A E L M L I P K S E P M D L E R P G V Q I V F D P I L F S N I Y N Q M K D A Q W A V D T S E T D I Y T L
RANDOM_1944 V F P P L E R N Y Q C L Y R S S V P F L Y G L E A V A V I K Q E F G D G S A I Y T V K S K V T A T G F D A L F W
RANDOM_1945 P N S S D N N T G G N S S R S A N G A V V C V Y F E I R V F V C K D A A L L F P F P T Q V L D G M S G N A K M D
RANDOM_1946 H L A H D I T D S W T E R A E S S N I E V S R N E L D P S R S M A R N K T F G A Y E Y F N G A P T L L Q D K Q P
RANDOM_1947 V I T E S P S T Y G C I V R M T V Y E V E K K L E S G V Q K M D M I Y S S T V N E I P T R R R P S S I K P K L T
RANDOM_1948 A V S H C R N H T S Q A Y L A K R S L T W F E G L F F I R D L L P H Y L E S E A V C Y G A Y E E N C A S S Q L L
RANDOM_1949 F A M E V N V S D L T K V R F P E S F C A V L I P Y L L L A T G T K A L T K P R G G S F I R H P L T D Q I R V M
RANDOM_1950 V D G R P D A A R F G E I D T H T G T G L V G G Q A D S C V S E Y N T S C V D R L I V E T Q A A F P G E V G I R
RANDOM_1951 C I R A G L Y T P G V T K R Y H T A V A I V G S Q R F H G R P A I N L R R D E M V L V R V P L S P R F D E A A I
RANDOM_1952 H G L R G S V L D M T S H P A R K A S N D P E E A V L E Q T M A S S S D D S S N G G C V N Y R K N Y G T D H A G
RANDOM_1953 H D T T F Q Q D P A V Q T L S P G F S E C V G L Q C L A S L Y I V P P A N E S T G N L P G D C D N E Q S V P P K
RANDOM_1954 T Q F G A P L I H I L L N K A F P A N H N C G P K F R T K D V D Q N F P G N K F V G S S S L K Y R N P S L K N Q
RANDOM_1955 D F C G I P Q D T T S P W C V E P S G A T C D H T Q Y S I E L P E D M A G G E V T K I M P S L A P E M Y I A E G
RANDOM_1956 W D A I P G S N T P D T A R L A I C E D Q D A A H H V H A E E K V E L G E K M H F F I P T M R Q L H T E K L W F
RANDOM_1957 A L T T K V N K V E A E R M A I R R K A R K M M A T S S V A A E G I M L I K K L D G K R I K G P L Q M G H G E D
RANDOM_1958 Q G G Y D M H E D R Q H M G V Y I R L C P G I A C K T T E L G D Q S R S E V S Y L I G K L K G A E C R R T L E K
RANDOM_1959 I K S H L D F E T P G Y S M D L R A A H E F S V E M Q P E Y A P F Q N T A A M N F R G K T T V Q P L K R K T L W
RANDOM_1960 K A W H R V D G D A A Q V V L G L F S A V G L S G T D P N S G M N V I R L L Y G P W F A P V G G F D K E T E G G
RANDOM_1961 D N A L A L P A I F P P R S A L D Q G Q E Q F H I D P G Y C Y R M L E T P E P L R A M I G A L A V A Q L P T P N
RANDOM_1962 I N R P Y I V R E A T N V V C E Q A V T D G I L L K Q R V S G I E F D H D E N L G G C Y R S S K Q F P K G K S Y
RANDOM_1963 V Q Q A S L C K G E S M E C N K K D P N L G H E H L V L T Q G L N Y L N S A G T E L Y V N R S M E G R T L V G I
RANDOM_1964 K F D D I V S G T Y Y C K H G V S R S L S A D A D L K E A E T M P R L R S I L A P S G E N E D M Y C F Q S L K V
RANDOM_1965 S V S F N P Q V L G G K A R S T Q G F Y L L K P T M T Y S G I Q Y T P G S S S A T L V Q A G I A D I N F V L S V
RANDOM_1966 N I A D F N G T N R A N R N L R G Q M A I M S F S L T Y G K K L N G N R Y F K V D R N T E C K A S I L S M T S G
RANDOM_1967 Q N K P F G G V H T E R W W S G I M T V I A S V I A Y R D C A W T V V D T P A T V L P G K F P G P E R L A K T S
RANDOM_1968 L G P K T D P T P D A D L S G T P G G V L G Q S N T V S D L E V S V S P V C S L N P L A W V K K V N D N T D N A
RANDOM_1969 L L Y S R I L V E T G T L V S S V R S L R V S K A M L G V A V N W A Q L P K M S S L A W G K R N I L F E R M N A
RANDOM_1970 T G P W N A P S N C N K Q I F K D M W P P E G S K P V S I W K N A R T W V G A H I N K L C F K I H A G V S G G A
RANDOM_1971 S K K H E G S F L G T P V N D S V S L N E D A S S G D I L K L A I F C L A H W R A E H P S G K P L N E T Q S V P
RANDOM_1972 A T W Y Q A V Q G A Y G A L D N A V L T A S F C D A I Y T L A N G S Y Y D T G P S Y A K R V L L S L I P E A S V
RANDOM_1973 I S I R K F N V W V E I K L E G Q A P E L A K Q A P E S A N R Q V F L E G F L V R Y D I S A K T K S L L G Q V N
RANDOM_1974 A I A V I V M M F P A D S V G S T L W V N N G G P G N R G G Q R T A S L A K T Q E T P L L P N K Y D A L S P A V
RANDOM_1975 A E G K D A V L S H V I W F D K S G T N G Q P M S D L V I K V P N H E E L S D F I D G L D K V D K R A G L D P P
RANDOM_1976 R G F V S Q N C G D D V P C M G P P I L S F E I A K V S K T S A G G P P F L Q F K Y Q T R G S F K S E Q G W E L
RANDOM_1977 P G I K K Q S L P A L G T G T G A S N Q R Q E Y Y R L H V K K H Q R I I K T I D R D I K E C P K A T Q L K D L L
RANDOM_1978 N Q Q S T I I G E L S L L R R S L E I F R L R E P K T Q G L G A G L M T T K F G E E Q S D A G P I S L R S D T L
RANDOM_1979 T S R E P D L E W A Q S L S S A Y K M E F S K I L E G N T P P P H P G F C A L A A N F A Q N P K N K G G L A L N
RANDOM_1980 G V Y G M H M D E L D S I G L K T V G S K D Q E E F D R E S A A C E A L T D I L A H L L C D S A I E R A E F S R

RANDOM_1981 L E T H W R V E L L M P Y M A I L I D I H V A T E G I V L L L P L V E M L P K S A P R T P T I P P N T T P M K T
RANDOM_1982 D A K D A E R K D D T A D T L D M Y R V T D A K V H T D T R A T K S K R G L A R G K F L K H D G C E Q T L E S I
RANDOM_1983 A Y P S Y A G L V V I A R W A T Y W F W Q C G G L N L N D L T L S I W C N P L S L T P G S G L H F A A W T G T P
RANDOM_1984 T H P F E V L P S M G K A L W V S V L A T S L R Y P L K R A M L L C L E P D D P E V E S H F S E H L Q S A L S F
RANDOM_1985 P L N S D G Y G K I P L T P A G P E T P W L A G I E R R S D P Y K K N C Q Q N A E R I T Q F L D R M G I V R K D
RANDOM_1986 K D A K D P K P K F K S K G P A Y T L M G E N I N V A E L Y K A S W N E N F D T A R L E P L S E K K S M G M D V
RANDOM_1987 P E D F R R D E V N H L K K H A R G T S A V V Q E N C K K T C K R R K Q L I D H D A R G L Q S K L L P M S V S I
RANDOM_1988 T L Y K Q K G G T G P S D L Q E K A D V L A D P N N T L G L G P V V L K Q Q R G D I C T T I D D I K A F I Y L F
RANDOM_1989 I V A A S N Q H K P R S I E C L M F D P G E V S K M H L P W H S K I L Q L K T G G G Y G P F A C A G G N L A K E
RANDOM_1990 E F R E G E K T A M I R S H S T I A Q A N N P S M A R W N G F A D P E H F Q H D E F D G P P A D F I A M G V I S
RANDOM_1991 E K K S V Y T E A A I I L Y V I D L E V Y P A I N N A D D L E V M D L A D D N P C V A S Q T C T E R T V A E E K
RANDOM_1992 F S V L T G D L A G D L D G G F F L D I L L V Y K Q A G N I S S Y S P F I S L L T N V E A G L D G V I Q K Y C C
RANDOM_1993 I E K E T N Y E L V I M C K R G C S G S C I Y E A S L K E F T P S M W E S M V M G K L Q I S L G N P D P V S S S
RANDOM_1994 N P T K V P R V Q K M H L M V L G A R Q L Y L I K R V N G L K E Y D T L G H A P E D V R K L V T Q V A R K C L Y
RANDOM_1995 T A K R I L P G R L D L Y H M V P Q P A V E V K T T K R D S Y V T R D L N Y P N D S S A N E V Y D P F A G I G S
RANDOM_1996 A Y S V T S F K C Q S K S A F Q T R I F F G A F E D R R G P L G W C C A G Y P T D G G L G S R R V N N T P R K K
RANDOM_1997 M D Q S A R K I Y A E Q M E P F L G G G I E G N V L G F A M R S M F M L V S K L A L E E P N E N D I G N Y E L V
RANDOM_1998 T A G D L T C T G G C G S L H I I L A L R A A R A K S A V A V I Y A D W T E T S R M P V F A V S P K T C L K Q T
RANDOM_1999 S R D L G L R R D F I K C R L V I N G P S G S S L N V I G C V Q R R A R C I R N A V S P A V T G V P T Q M S M V
RANDOM_2000 A K L A E P P L R M S L M E E I S V I I F Q M N A I K V P K V A T A I H H C N K E P M A Q W P V N L D S Q S L R
RANDOM_2001 T F Q T K Q E M L P K S D E S N E T I H K V E S M L R I E P L T Q Q H T I T H C K A F Q V G L Q P L S V F W R Q
RANDOM_2002 K A L V C E G L K L A G K L I F M N G G T Q D I L L A A V R R M W D K W T S G T R G F C K K N E A G E E Y N V G
RANDOM_2003 T M S A V G V H L K S L S L K P R R F E Y L M R L G F L V S G F E G E V S L M K A G V Q Y K E S A V E P V P L G
RANDOM_2004 D E S N G S L S R V S E A Y A K T Y G S R T K F G K A W W L V Q K V D G P G N S S R N P G S S V S G L M L S I A
RANDOM_2005 H Q S S F D K C K L C A G L T G L D T D F L G S V T L V G D K F S R T W G V D S L K L P L G C D L G E E N E G F
RANDOM_2006 L R K G F K S S N P K I I E L A A C Q V E Q P D Y V P A P S S E Y A H V L P A E N Q T V D K Y R K H Q F E T I L
RANDOM_2007 E T R E C T R L I F E K N I V D E L T G I T D C L I F Q N N T K G N D M S A D I W G P A S D D K E D A Q N Q K P
RANDOM_2008 Q K E N L Q T N G E T R V G N Y M V Q R R A K T I L R A L L P K T I S E E N D K F V M Y T D A T L P S S M N G A
RANDOM_2009 E D T V G E L S G S S P S H H N K S S R R T Y Y A V D S K S P Q I S N S D N R L E D E Y E N F A L E L S V R I H
RANDOM_2010 V D A V A Y I R Q R A A V L Q C A P F L G N V D L G H P L S V N V G D V C L K N V G I D V E G F D K I P E Y K T
RANDOM_2011 Q L I E C G E H N E L S G F L T G Q V P S V L I I C N T L D A P I E R P G F V G L P E R F Y V S L L K N G T L T
RANDOM_2012 L D L S V F Q A D E P E Q W I K R L K T F Y L I Y D E K V A S K P H Y E A G F D N K L T S S E S E T S I I S M A
RANDOM_2013 H N D V V K G P R T K P V F L P E V H R V K K V M G L R S R F R L T E T V L R N S T L A A Y I E T R P E P V P P
RANDOM_2014 C D S Y P S L Y P R N P P L K N L H S N T Q I F Q E R I F L D S G G Q G P V R S G K A T V D T G V V R V V A K G
RANDOM_2015 Y F S S G E L K Q P T V G A W Y S A D Q N L K A S I C Y P L D A E A I V A L N N R W S D V I A Y S E D G A L T A
RANDOM_2016 R E P D L P S A Q G W V Y S E H V I V M A F P G S D T Q V V H M M A A G T H P D W K E A G T C L I A E D P G A S
RANDOM_2017 S D K K A E A Y T S A G T T S S F K L V Q D T L L R Q G K G H P T A F Y A E K N D H L T H E E V N L R A K Q D Y
RANDOM_2018 E T V L Q E A H N L L T A Q T I N K S L M L R E R A R Q Q F G R A F I D E S G I V G E G D D G T E N L A F M N T
RANDOM_2019 L S Y R P P D A K A N I A Q L Y V I A E R R A T D I F L S S G L E G R H E N T K D R F N K D A D V G H P D E N H
RANDOM_2020 T I R G I E W I M A I C L P G E S I S E A Q R H H I Q S P R G N T S D A A V L S E K E A P R Y C T A V T L E K N
RANDOM_2021 N F N S N H D N L C S R D E N G A D V D N G V S I T Y F E V P N K S R D P F L Q K Q D T G T N K L A L D P G N T
RANDOM_2022 A Q N T Y S K K T P N V N C V P P A K V A M N S K V F L E Q R P Y L K V K A T N S A A D I L K H F Y Y M P L F L
RANDOM_2023 Q E Q D R H S Y E E L N H I L P A N L S I C L Y G E A D A T Q G E K E K S G D R S V L G L V R N R A L V L Q L L
RANDOM_2024 A E S Q R T E M I Q G L V L F T W Q F G Y A F D T P N R L P A I L T F E G A T Y D S I L P V V K S I N D V K D S
RANDOM_2025 F N S D T D C G G Y V N Q E A G M V K I S S K F V N I V A Y A R E S L Q M L E L Q M K D V M L N P L L D D A L S
RANDOM_2026 G K R A E V D L Y D Q R G S A I S K W V G S C S V I R A E A W R P K K S Y V L H A C A L C I D L L A G F L C W A
RANDOM_2027 K P Q N G L W A D S G G A N D V D I T N P V Q D K G V M A K D S T I G I N G S A L L L L E G V D K K L K A W A E
RANDOM_2028 G P G F T H L Y V V T Q A L M R E F K P Q C M S R Y G L G S F A N L K P Y P T M E F G T Q V D A M L E L L P Q N
RANDOM_2029 V N A H Q F I T K D T M T L L N G E S D L C P G G K V I D L L D L T G L S G G H G V M Y L P V N F L E R F G A N
RANDOM_2030 A G A M C G V S A M H F A P D T C Q L E K V Y E L I K T L I T G A A E N Q H K Q A Q G P E L L E G K I S L A N E
RANDOM_2031 S E P F G K L Q A A R R G S F N G A W G V K W V M S N G M I A T E A C I H Q L R V E T Y A A G E D N W A V R A T
RANDOM_2032 I L K L D C L S A E D K D K T S E S K D E N L P Q K P G T H N I E V M Q T E G L L N T G W N N S P G C G Q Y G N
RANDOM_2033 D N D N M C I K K K G P G D I D L I E L E H I Q F S L R E G P F R S M G A K G L S K L E V H T Q M A F Q E S A P
RANDOM_2034 G Q N D I A I Q N D D L H I E L L V T N L D C V A E H N E T G A Y L C E A G K A G P Y Q D A G Q Y T K N K P G G
RANDOM_2035 V T M E Q G I P Q K P V Q E L S D Q R V G E Y L L Q L R F L L E K R M V E H D P I L S H V Y T M P S V E Q I E N
RANDOM_2036 G M E V E N T T D S I Q D Q R E A N T F N I F L K R Y E D M Y L G A A D K M A E S S Y R F P A I V K N D K H A S
RANDOM_2037 N T A Y A P V L S A G E F L S I A L I P K E A K V P N N A R E A L N A I N L M N Q T A G F K Q P A L A M G Q V R
RANDOM_2038 K S D S L L L F S A Y K I V E D E M K V V G L N V E G N V K Y C E L V T L G S Q L R Y P L R L C S F C G W S S I
RANDOM_2039 S A P I D W A Y M H S L E P T D L N L S M T P D R H L K H P T L R D A Q G T I C L T E L I R W H D E D G M K A W
RANDOM_2040 V I M P S W L I V N F A T I I I A C I F N V P L V A L S R G V Q N E G S G D Y V D E I R V R P K R D L I N A A C
RANDOM_2041 G Y L E N D M F P E Q S P S G S P A K E C T I F D Q L T T A D N M E S T M S D L G A R K S S G I F P M L I L N A
RANDOM_2042 L L S L G K D D P R E L G P R S K E M P Y Q L V A P V H R P M N V K L E A L E Q K P G R Q V D A L Q R T G L A R
RANDOM_2043 R H L S V I R L Q Y G D N L R Y V K M V F F A S E L G L Q P G V Q L R S A L N D Q K F S E L W G I V R E T E D E
RANDOM_2044 G E Y V P A Q N G K L K P A R D T S D G E E G R L H S D D P D V G P Q D A G Q E D S R R R P E F T E G Y V F E L
RANDOM_2045 S Q V T N A G R E V W D S L G E E R L L K S S H C G C F K V Y Y R A R W W I A L G N K E L A A D T L S V K P P C
RANDOM_2046 N P P L Q E S N S L K S F Y M L E I C R N L Y H A L N N S Y F R W G E P P F K D C S R K A L I N K A M S A E D R
RANDOM_2047 Y R N R E E G E S P H N E D C K G R G E G R P A A V G C S I A S I F N Q T G I E F Q G Y I H F D V Q C R L S P K
RANDOM_2048 L A K P W F Q P A S G I T G Q T T A Y D Y S W L T G V F D S D L V N A V L N G D L V A S S K E P N K Q V I G E I
RANDOM_2049 G N K S G G N Y W H A V S D D S M N G P I S D V G E Y H Q V A D N Q T G T I E L P Y I L E V Q N H A N N F Q A I
RANDOM_2050 G F Y K S D G M Y I K N K T S E G L Q H Q T V G F Y N D F L S V V N R S E P V I G E K L A F E Q K N I I P V G G
RANDOM_2051 D G T C S S A P F A S Q D A D E A V F Y S A G T G I D M V L V K F D I E V I I N S A R K T R L A E L L S Y T H I
RANDOM_2052 M D D L R P I E K R K G E S V L G T T M H F R K G A L K N G A V N K A R V A H S C G L R L P A A Q V E R S V F T
RANDOM_2053 Q I A L E W M H G S V A V D V G K A P A G E D T S L V K A Q M P P Q L H L A W E V S E P R D D K L L T L P R I D
RANDOM_2054 I G T G V A I E G P S G D N F P D K W A K A A Q Q L I G Y T G L L G V R F K T R M W I M I I G R E L S G T C M G
RANDOM_2055 S S F S R I V W Q S S R L Y S G H N Q P A H E L E D W Q I K F S E V G P T E S P L L K A M N N F E N N Y V T L T
RANDOM_2056 S V S W T E A S N S A V Q W K G G N S M T V K A D Y Q A S W G I M I N F A R H P A L I A K N A E P A H I S S Q T
RANDOM_2057 I D F G R S D G V G G N D E D P Q A L Q V L Y I E P F V V C A G I I N N V E V G I E L M H L H S A E N A F S R W
RANDOM_2058 A M W T V S G T S G V F K R V Y P L T I P K M I F Q A G D L L V E A V R L G V P L K G T I Y D S I L A A T I R L
RANDOM_2059 L H K Y I F M V D P P K L L R V M I S I N T A S W Q E Q K K H R S F L F M E Y P V N F K K T Q R D T I G F L P I
RANDOM_2060 L F R D E K K Y C T K V P A C K D D T P W P F L G L W S R N M Q R E Y K G R F H V S Y I R P G Y L F T I V L V L
RANDOM_2061 G K C Y L T T C M F S H L T Y V F G R P S D H S S I N T P Q H P C A T S N L H A V E I W V P E K I A P V K A L I
RANDOM_2062 H F R L V Y L S A E R T Y N E V L P L R I K W P S R P V D V T Y K G K A V L P G E Y P Y L P P T Q M A L E G S L
RANDOM_2063 V P L N K A A L Y R G S L V Q A L L N G T D M V F E G L Q S T P D P K W L S K P F E E T A L E G E G L R T I P Q
RANDOM_2064 S A G M T L T P K L R L I R K I F F T A D T E K V T Y V T Q N L S G A I M K R N A E F Y I S F P Y S R D S Q W D
RANDOM_2065 T F A V P R K Q A H Q L S Q T G Q R L G Y F D V V K H Y A S F S P Q V I L R M T E S E V S R G R R A R F N W L E
RANDOM_2066 Y P L K A K D P F Y S L E G V E D V W E P G E D K A L A D M V A C P A P V G V K A L K F S D L D K Q V I R K P V
RANDOM_2067 S E L L L L A L E I A S S T A V A S D T R A A L K F N I I E I S Q F A Q D I N D I I L K F G S Y W R I G F V I S
RANDOM_2068 D N T L I K N Q A P L A F A A Y R H E V L K A K D C M Q A Q F V G R I A V S L G K G S D Y F K F M D Q R K R Q V
RANDOM_2069 E K E T E T N E C S A I Q E L L H G A W V K N G T H Y A H E E K N I A E E E T K I F M I V Q G N R N L L Q T S G
RANDOM_2070 A I E Y N G Q D P T T A K G D P L F W T V N P D T S A V E G G R E R V I F F L A C P R H E P E T G S C F Q A L N
RANDOM_2071 F T Y V H Q M N A R R A D P A A V R Y R Q I N T D Q T A H V G V F L A Q F S V N L L P S H E T P F R E S S S L Y
RANDOM_2072 L D G V R S V K A L Q L E S Q E G K I S I F M Q L L S S S D P I S G V R A P L F A Y L L P S R Q E Q A L D R W R
RANDOM_2073 L Q Q M F E E E A S E S M V P V T F F R G C V C P K I F K M V L G A T P R R I E S T M T I A N E I C R C W G I Q
RANDOM_2074 T E K E L K T T A R A Y Y V E S I G D N L G G E G D Y P E V K E I Q L V S R G F V I A L A N K S E W S S P E S L
RANDOM_2075 D R S R H I V L E A V N M L R T K D N K V K Q V S N V I F L P G C A K Q L N T V M T P T G A E S L S S V Q N N I

RANDOM_2076 R T E Q P P E K F E A W G S A A I A F N E A F P V Q G K S I T T Y I Y A I Q Y G N Y Y R S R Y A T S C V F M C N
RANDOM_2077 A S W T L N T D A L I F L L M M L N A H N T R P G P R V D N L D Y V P V D A S T C W N A A G S L A D V M G R Y I
RANDOM_2078 A S D V V H Q G V A V P S I G L L N A P R I K G R S T G G H A K Q R I G Q D I R K T S D W T Q C G K A G I G G D
RANDOM_2079 D L D Q D K M G K N D A S I E E T E E T V H E C N M N V R G R A R F P E S T H T E I Y S V S K N G N D K L L G T
RANDOM_2080 A K D R I I I E V I S T K G G I P A E R E D Y R S P A H C S G P F A K T A H K H S K L S N V F K N N Q Y V A Y F
RANDOM_2081 R T L D C K Y V A V S T A D P S A T G K S R M G F V V E P K Q P A M D N E I I G M W L P W T K E A P Y F V Q L H
RANDOM_2082 D R G V F I E A L K K V K I R E D K H A T S A N I F F R I F E F T N L E G S S G R E S N E L F A T F I T F A L P
RANDOM_2083 S E G G G H G A T G M K A C Q V K F I R F I I H G W G K I S S F F A Q A I S D F D V L N P L R I T K M S E N W E
RANDOM_2084 T P Y R V M P I N T F L R N Q A C K D M W G T V E P N S Q D R Y I M V G Q I L P R Q E V G E L S K G M Y Q D F L
RANDOM_2085 P A M N E L V L R L S L T W F S A H F D A E K A K C N L I L G H I D A R G G T E T L S L T K I P V T K P M Q L D
RANDOM_2086 L S V C G L G V Y A F M E N P Q S S R T L L N T Y L G N G V R Q L T L E R E S K K D E R T A L S D H A N P L S H
RANDOM_2087 R E V E K K L N W R I I F R E L G F F L I S F T V F L K N A R L M V T T T E Q Q F R S P T S R A E G V A F I P I
RANDOM_2088 L S D G I R C K A L L E I R L K L Y W I G M G V L E A Q V I Y F A V F S G A V D V A S A G E I V E P V D D G W T
RANDOM_2089 A L T L Q A L V P V T W E A A Y G S L G A N S G I T L T N Y S A V E L L Q V V K E L A Q G Y P K S V F R V M L T
RANDOM_2090 Q A M A K V V Y K T A S A L S L I K L G W L S Y R S L S L K C K N M S L P V D F T K V G E I T V L S D G I L G F
RANDOM_2091 N S C Q E Q R I V E M K I L E V S N E E N K T A H A E R L M I V P R P L M S S Q S F F I T K G H L I N Y S E K N
RANDOM_2092 P S L V Q L S R G L F F L K M T S D G R G H T S R I P L T E D A Q A K P D V G N G F R R V P Q S S S A N E V S N
RANDOM_2093 N E N I Y K L Q E L Q A L V Y S I L K R Y V D I G K T V R L E K C F L H E E E S L E T T P F D V A H I A K Y S R
RANDOM_2094 T D G R S G P S A I D L L G S R L A S G S N R D A F N I S R E D N H G Y Q L V Q T R C R E G V G Q L N S R E R S
RANDOM_2095 Q T T E Q N K Y G D G A L A L C F Q G I C L D I G F R E S T D I V Q T D N A P K L W G M F F G P Y R K T N L M D
RANDOM_2096 D V E Q T T L R Q S S R I A V L A D N I E C H N N R G A N K H K M V P G R T V L D P Y F V L V K G S L P E K A V
RANDOM_2097 N V D V N Q I D K S E P E R T K K G T A V K G K V D P A I T I R A F V D R L L T Y T D T T R G G S N F E S A D D
RANDOM_2098 I E T H S L E T Q I E T P K S S D Q T K G L P N E G P V V L V Y T T R G Y P M L K P D L P M A K G E H G V Y E R
RANDOM_2099 V G L I T S A V N W G N L A C I S D P L P G V H V W G A A L E Q N T R A R L D V D P S S G A F L S Y L P S G E F
RANDOM_2100 E Y N G A F Q G Y P P I S E P K L K K Q W Q M F Y S V I G V S L L I S A V P F V E K F V R P R A E Q V H A L E L
RANDOM_2101 L E Y R K N L V D R K F R I A Q T H K D E V D S V W E K L A D F P S T H K D V H R L F A K M R A G T A K A D P D
RANDOM_2102 F G V L Q G S A E E I I Q M T K R N S T S E R T E D L M E G Q T L R L N L T Q P V A P V L D Q T T R Y R K R V K
RANDOM_2103 A T C L E A S F T R L P L K S K I G W D E I T L R S G R D H F P T E K F E T P P P L R S D T Y Y S N T G V I E M
RANDOM_2104 L L K L P Q L L K V I Y C V Q I S S R T G P D V E S A F A G G A L L Q A Q K G T W Q P L R C I S R P R Q L D L H
RANDOM_2105 K L K K F N E I F Q I V K S E A N T G W W Q Q A R C L V R D A I K K H N K D N L Y R T F L F Y M I E K P A Q D F
RANDOM_2106 G S A D S Y T A D H P L N A V T L G C A L V E K K N L E I A N P I R D S P L T L A Y N G P R E I V P I A Q N D A
RANDOM_2107 D V E M S E M F E I Y S P T T E L S E V E I S D V L R R I K L S A G A I K M L K W M G L G A M G L A S A P T L S
RANDOM_2108 S N G D E V A N A K G I V N A F L G F G P A L H L V A V K S E Q I V P P C F W D Y Q S A H V T S D A Y K R C K V
RANDOM_2109 W E P A Y Y Q V S R N N F T N P Y T D C I E P N P I W V D Y T C F Q I S E E V L A F P I C N V P T S L K G T K N
RANDOM_2110 A I N R V L N S F A M A V L E P L L R E H D L A T T P C I E A L Q Y A Q R Q K A K Q D A S Y Q S G S Q D E Y H R
RANDOM_2111 S R T Y D K W T A G A L L N T F R L D D T W D A L S I S K G V H N R F G D K V G R L I A N K S G E E Y I G S M T
RANDOM_2112 T L I A L V K V C Q T K P L L R Y K P P L Y A D T G M A A S L T L Y F L S H Q N V H V E R F K T D G L S A P R P
RANDOM_2113 T D S E G A S N T F Q V L K T R E V A A R S R T D A G D G T L D E L N G F G D L I A G K K E C E L S I T N G S V
RANDOM_2114 V T A N D N F A Y K P H T A I P G G Y A N L L D G Y Q N I M F A D P V N L P G G A G S N V M A Y F Y K V V L K L
RANDOM_2115 G F K S P D L T S L M I T G G D V A E S F K I V Y A E M K F E E V L K K G D K H F P Q T A T P E W R S G V A L A
RANDOM_2116 T P R E R I G L A S Y K M V D Y D I A I S T V V K A K M E K I F T C V A I L D P W H G A R V A L V N S P D N W I
RANDOM_2117 V G K R N S G V L D I Y P R L H N D R C L R W K A I Q S C E Y R P G E D E R L Y K C H I Q I S K A D F Q R T V V
RANDOM_2118 D I V L V E R Q F K A V I N W N I T F D D C K A K A S K L K T K E T S D A L E R Q A A L V E E A T R V I L T L P
RANDOM_2119 D G V D T L G E D S G R G D P A T N Y A A W G K T G K E G M P E K M H V L A D S V S S L N K I L I L P G E G G N
RANDOM_2120 D R T S E K Q I T V I E R K A A V D F A G M R K D T A Y A V A R P I A K K T I M R V A D G A L K S T I G Y Q R W
RANDOM_2121 A L L L T R P G W L E K Q G I Q E N F N A G V N S E R Y N G G M Q K V P L P D F E E R G A K H Q S L L K S K I L
RANDOM_2122 L P T F F V N V V L G V Y Q M L V A S I I F S S K A V D L A L A S W E I S I R C A N T A L N Y H S R S K P D F Y
RANDOM_2123 G I G N S K S T I A K C L V K Q G G P E C M L V V P A S A Y L T P F A P E L E I F D E I K Y M T P E H R E T H R
RANDOM_2124 V Q Q I Y V A I A I D N N V G I L C Q R F T P T M H F L T Y F S T V R M G R H L E Y M V A G G G L L Q D S V E D
RANDOM_2125 W D Y V T N G T K Q M K Q L A V R N E L S T E L P A P W D R G A I S T R W T N D A E D L D L A G G Y P S P R F D
RANDOM_2126 G T E R T E K E P I Y M R E K V G M R L V A L D E R P L L L S V P P N G N S A R S E Q I A F P P F H A V P L R D
RANDOM_2127 P S F C K R D T R T Y G A K D V E T D Q L P Q Y Q F P S T M D T Y N V T K L D P E S E E I L L D A K S R P K E L
RANDOM_2128 L D L S K K Q A A T R L A I Y F M F I Q S M S E G K L E R S Q A G S A N K T Q E F N I H T R A R T I D T K L C E
RANDOM_2129 I D W L I R E P K K S K F D N P G L H Q I I D T F T T N S D D L Y Q H Y K K L Q F T G V H E G V C D S S G G C G
RANDOM_2130 A Y R N A I T K H K F V A D E I D A A G Y E Q I Q K H V E H A P E V P N N R G D E Y D K S F D P T T K F E T I Q
RANDOM_2131 Q F K R R N P T Q F I C Y W R V N L D R E R F Q D V E P S I C T R T V L V G A Y K V Q A I C V I P H R A V F F R
RANDOM_2132 T D I K Y A T N P P M L A Y V A A R V T F I N P I N N A S G N V F S G R F I N N V T A E D M F V G P A K V R K T
RANDOM_2133 K M H S E A S W V P C F S N C D S T K Y E T L E V L F P Q K K V R L A K G G S P D Y E V P H K K A L S F G Y P D
RANDOM_2134 A V N P G R R L R Y R G Q E K T Q M V V V R K T Q G L M R N A S D S K R C T A V R E E C N E A S R S K T F V P G
RANDOM_2135 A N L K A E S L F E Q W R K Q G H L F K A L R V Y I N Y D R N V V D L E V F P A A A T T H G H M L G N V D A Y D
RANDOM_2136 K E C K P V E V T P A L Y S K L E G P S S L L A T E T D L L L G P V A T E V L S V R K Q T S L I Q N E G G I T Q
RANDOM_2137 Q G N D L G Y N E V D L D E S H F M A P D L E T P L L Q R M F P M S S T Q K P G P I E C P G Q Q H Q Q A G L G H
RANDOM_2138 Q F A D K Y L M T H P L A V P P I A C P Y C S E A A W K A D R H R N P G A E R V I E A N A S L F I G V R Q E A V
RANDOM_2139 E S V T N E L V G I G M V D L Q V P K Q D D Y K I S G Q R E L T M G I P S G R E R S D V T M C V Q R Q I P F R L
RANDOM_2140 F R S K A K M A V E Y E E R G L E G C T L F R L V L A Q S F C S R F R K F F S T K I T V I T T C L D K L R K V F
RANDOM_2141 I S S L I R P E S D S Y F I Q R D N D A D A E P A F G G Q L G D L M D Q S V V P S A A T T T I Q E K A K L T N R
RANDOM_2142 D E S A S T V M L I Q F N F K Q E K K P D Q E K D D V K W P S G D K A H M F D F A K Y A N T A Y F L A E P E E I
RANDOM_2143 I I G T G H E P N I E A E L F R L T L I A M G F S A T D V V N S K V F A D P T R Q L Q I G A L Q R E F L A L K I
RANDOM_2144 S R D V S Y D R R S W F R A E Q Q A S K T A E E I A F G Y L G R R T L P L K L K L F T P P G A N R S P Y K K V A
RANDOM_2145 N G P V S S P S H A Q D Y H Y L L H N E A S K N G E A R M D G K R G N P G K S E A E Y C N P Q P G V F H N I I L
RANDOM_2146 H R P R S A Q V N Q I P K M G T N E I N M L M G G T F K K R R R S G E T A D A I R P G I Q V R A N S S C T I I A
RANDOM_2147 G K G K R L D T F F V R I R N T N K L I D N A C L G Q W L Q F E A R Y E H F A K D S D P Q A R G T I P N T C T I
RANDOM_2148 K C M V K I Y G D A E R R A K K E A G I L D M P I L L T H F P Y K Y A L Q R E K N W I L V S Q M I M N L A D I S
RANDOM_2149 I L V S C N Y V A Y T G W T S M F G E E G K T S I R W D N P T R T L E T A T A V R G E R G E R L V K A R L D K L
RANDOM_2150 D Q K Y F D T I L I T A N V G V S A Q S V K M T T G L D S H Q I A L K G L S A A W C C V P I G K A F V A I F K H
RANDOM_2151 I K C F I N Y S A S K A L P L A A P S V E D N S I A R Q E D L P I T L P V P Q F S V V G M A T Q E P I G P C T G
RANDOM_2152 L E S C T D R I R P T F K S L G S E H L L T R N E V P R K V D F G R C S F E A N P D A L M L F Q D S L T P I T I
RANDOM_2153 D T I T S A A C C C Q A N E V F G I T T Y V G L S T L L D V G A V Q L A V Q L Y I A I A L R C K S P S A F Y E K
RANDOM_2154 P E L G S F E L P L E N V L P L Y F N D Y P R Q R G Q F Y V I Y K A N E A V S K S Y R R A S R T L I H I K G K E
RANDOM_2155 A T P A V A C C Y V L W R E Y G D V E E E Y Y A L N L Q F D A L K D M N L R Q Q A L V V L L F P K G E A R S L I
RANDOM_2156 A K A P A I G C T A A Q Y N M L I K Y S L S F A W N L E A E A E D P Y R Y Q V S F Y G G T F M D N R R R P T R T
RANDOM_2157 L T V D D E K D V E S P Q T A H N S S F M T A A P P L D N S G K A T N Q D V S H K I D P P T N G L L D S P T N K
RANDOM_2158 P S S F G D C T K C P P F H V I E L I Y I N P E C E G F R S R S D G A Q P E T P D F H V W S L L I Y L F S M A G
RANDOM_2159 E S E T G E L L S I P N G D T H K K T L S Q I L R D R T A V K L R P W T H V N S D S E N E A L T I T I Y E L R D
RANDOM_2160 N D L V R T E F I G L L L V N I D S L V T V A V T S P G D V S G R S F I Q V T E L M L N H T A K L I Y K A G S H
RANDOM_2161 P W Y L I A K K I V L W V T V M T G D P D T T T E K Y I G E V S M G R T V Q E S S K Q P G L Q S I H F E E C G E
RANDOM_2162 Q P F W S K R D F A T M I E T L I T N I Y L H T K D A F G R T K Y H A S G F T Y T R G G A T Q Q N G T Y L M Y L
RANDOM_2163 P G M A Y E K R R A A G F C S S R V I S V E W A I G R V W P G Y L V Y V G P L V F P F V T K G Q M H F Q V A Q R
RANDOM_2164 P P A P D Q K E A S T V I D P A G C A R L Q S F G S V C N L A E E D G K A N F N R D H S S M E Q H G F G P L Y S
RANDOM_2165 V N Q G V C E I V M I E P E T Y D L P T L E P F V A T Q G I H A T L T G E H E E I G L E V A V V L G G Q L A I V
RANDOM_2166 K T E L W S R V K N G T Y Y E G Q E M E S T E I S T M F L G M L K S M L A D Y P F S S K N P K R P G H H E L A S
RANDOM_2167 K A R S I H R V E Y A H L E I P V E C W V H H S G F I R L E V V Q A L I T L G Y R S L A L I V A F I W P Q Q G Y
RANDOM_2168 Q V E A E V R C A A K N V D S T M Y R I S E A S W S A K K L L I K L E W L Q I S Q K E R T S P V A P G A Q R I T
RANDOM_2169 L D E I R D S G P C M R Y L G Q T G I I V G V A H L C N L G I V S G G A L D R W Q T N E K T F F P Q S A F F S S
RANDOM_2170 F Q A T H E M P P I T P L G G L N L L M N I D T L N K E N P I K C L F C A N V S Q T K V R C F M D S K P R K M R

RANDOM_2171 N N E G K G E L S Y Q K S T A M H N L K D T E N N Q Y Q L C Y K A Y H H K L V L H G K P H R S S A G R A P N W Y
RANDOM_2172 L N F I K K P R T L S I H A K T W L D A K H A S E E I I I K I S A E S Y R P A N L P V R E S Q G P G G P P A G L
RANDOM_2173 Q L E E D E L P K S A I Y K G P R D V D S K T G V D A M N I V A R S R S G Q E K E E Q N L H N A V Q P D E K M E
RANDOM_2174 E W K R E F V C T E I E F P T E V Q N Y A G V I P D D V G A V S P T Y S N D S H G T A G Y T V G M R A K S P N L
RANDOM_2175 K D P S S K F S G G H V L S G N E E N C D Q L F Y V Y L K E N E Q A G F L G V G H A V H I S A L A P F W E T G R
RANDOM_2176 V F I E E L A Q A P S G V S E R S D V I L E Q A T D R I F P A L M I Q G G V L N L T L V S G P R A M S I N K E T
RANDOM_2177 V I D D S M R D T V I A R H L A T T R N Y A G P S E R L N R D R T V S K D Y R Q S V R T P W T E Q L L H Q S V P
RANDOM_2178 I S D V T L E R R S N T G K D V F P R M D Q L L S P G Y Y W S L R A A N E P K S A F C D S E E V E Q Q H N L L D
RANDOM_2179 L Y L A L H R P A A L Q P L S G N K Q L D L K S V Y K G S N F F S L R R S Q T H D P H Q M N K L G P P E D S N D
RANDOM_2180 E T R L T R S D C S T W S L L N E V W I Q K Q P P G R A L R S L P K S L D M M V I L G G P A I Q M L K T I L L G
RANDOM_2181 D G P S V N Y V H G F Y M K D G A G H E E Q V R I V V E D V A P L F C T H G G V Q L A V P E I K H D R V Q N D D
RANDOM_2182 S L A A P E S K N S F Q G G P K S A T S V K R C Y P S R S E S V P T E P C T P A V G A E A P I Q P A E A C A G L
RANDOM_2183 K A L E Y P G F L G G R G Q I V P T L D A C V S E I E L E P M G Y Y L S G G D P S T G F D C A M A P N K L I I Q
RANDOM_2184 A G A E Q V K S H I D E E P Y L L A R G A E R T V A S N I W E G C E S P T M D S E S W R A S L V C F L K Q T P S
RANDOM_2185 P N R G S G L Q L E E V V L N N C Q N V S G R K S F L R Q G G D Q D F M I L A N V L G G G S F F S I T E S E S A
RANDOM_2186 K K P F D G H H V K N S K V T V I K L H A A C G E E S D A Q T A I K N G L R V S S F Y I L T A E L N S V T E V L
RANDOM_2187 T D P A S C V Y E T L T T S Q W T D A W K A Q K E P N M N V K K E A A P K L K S P A L D K M A M L R V G H E E G
RANDOM_2188 Q A A F P A I G H F F I T S D A V N R A G E Q L D P S R P C E L F G N I G F L G P L A G S Y I S S I L Q A S K A
RANDOM_2189 Q A L L Q A P L Y V R M R S W K S C K G M D E N I K F F D L S F E Q E I I E T K E F S K T Q R L L I G E L V K E
RANDOM_2190 L P D Q K L W A K R L S I S V E G F G Y S G S G I V G D G T C L L I V D C C E A G A L G K S A K L A A G K M L D
RANDOM_2191 V E G S T F P Q K G S E G R K G H Y L F E R L L E L N P A V N N E G T G G N I T L Q D G E H P Q C G Y C G E T A
RANDOM_2192 S S I Y G F I N N A K P K V Y T K L G Y R A K C S R C A T M W I T D L A V G L D P F A L P E N G S I K F V L E N
RANDOM_2193 W Q C I K P S L I L A I N N I G R L P M H K Y A G S L G D G A G Q K Y N R P I D L E V G D F E V S L K T E T F W
RANDOM_2194 R E D R L L C L I E L Q G F A V P S I T P T L R Y P K A V G K S E R L I Q N E L E N T P W T E L Q H G L K L G E
RANDOM_2195 K G G T K L P T I A D F T K K N S P L V Y G P Q P F W L W V N A P L F M N T V S F G H V L H M A L G A S V L T W
RANDOM_2196 K E V P F E P S T T A L C S F D D N V S G L T D R A L I L M K A Q T Y T R H T D V E T M P Q T F P Q C D S D K Y
RANDOM_2197 T S R R E A S D I S V V S N I K Y D P T R L K R S E D E S T P P V L L I F D P S A A S F K I L G A L R M Q E F E
RANDOM_2198 Y Y H V P I Q R K V G H N K V K M L D Q I R D R A R Q P P Y C G D S P W I K I V S E K P K D T D Y S H V S E I M
RANDOM_2199 P D I V H P A N S Y A T A V K Y S T T S E L Y V D C S L I R G A G E K T L R I Q E C T Q A L T E A P K D C C P I
RANDOM_2200 G E K P F D K N K Q V K K S A R A N D P E W K P L N K Y D S N R V L I D E Q T I M N M L V T D L P F N S Y G I E
RANDOM_2201 F A G A K T K L L S S M A F T I F E R A Y L G Q R Y L G S F P P Y V H S K M L T C C G T E D A L D K L G I Q R A
RANDOM_2202 E M V L G F S Y D A C I D Q P L V I K G I S L K R L E R E K R N M G G A R S G A P N G P I G Q V L P Y R T A L Y
RANDOM_2203 S K G Q F V N K L Q C G L D S N I K D V H G P N R E A A V K A Q L A V S C V R V Q L K D S V K I T S G L G I R Q
RANDOM_2204 G A I L G G N V D N H E L Q W Y N R L E K H V K N V S L V L I A S E G E S L N P S Q M N S M Y E S V T G N Q L A
RANDOM_2205 D V K L G E Q N Y Q G F C E K R K E F K G G L P V Y P G N A T E P T S Q R S A L R P L T A P G T R I P A A R E E
RANDOM_2206 M K Y R C L Y N S I L N A D L S P V N L K S K K Y D K H N L G I A Q L E P K H A S D L T R A N F M N D E D V S A
RANDOM_2207 P K A V M E W M G S P S P L V A V F Y T D G L F E L T S I G L Q Q C P R V A T G K S P L L T G P G A W L G D N L
RANDOM_2208 F K G P T R A K G N K N R E C Q Q R L D Y E G Q P K Q E R C L V F A V T F H A M K S E N T G N Y T W G C D A L A
RANDOM_2209 L T L L Y I Y W K F V D L C V E T D D V S Y D R M I G L D S K Q Y E H L F V M A F G K Y F R L P G P Y D S E L M
RANDOM_2210 G K S L D P I G D W D E A K E T A L G L N A Q N A V F H M E Y K E I G P P D G Y W L A G M A M T N D V E Y R I E
RANDOM_2211 T A P H I R G V H P S V W V R L K S T D Q Q L V C V A I P V V G Q L K E H S E W L E R S L D S M N N A L E T W A
RANDOM_2212 P D A V T Q Q D I G S F I C E Q L A R D L F T E F A P A G A L V I S T N I R A E L A V L L L K C R C N T P L S Q
RANDOM_2213 G F A N E K E S R L D G E R D V P P E F K P V C A V P G V T N G D R A L L D L P F S Q R V G H R V R S G R M S E
RANDOM_2214 E Y A K K G K S L S T C F A A P T A M F T G M I E E K G L S T G N S P V L F N P R K K L K I E S R Y A P V V N Y
RANDOM_2215 P W D N I T T G E A N E G H S S G E E Y G E T T K R Y P H S E Q F S T G T G A V Q I D A F S A K K I G A K T L T
RANDOM_2216 L Y V F E I R F Y G L C R T C L G V R P E I G N D C G N Y L A I S M M E I R R F A A E I D P I A P F S D Y D F S
RANDOM_2217 E L H N Q V I K E V Y G Q T V C R K K K P N G T D Y K Q G I T A W I V A L K P N E G S F R D V G Q G T A C A F E
RANDOM_2218 F K N W V E D R K D W L K K M I G K N K V P A L C S A G V I N S N Y M L E V E A L L G T P A A D G Q L L T S H S
RANDOM_2219 V G V D D P A P W E K T L V R V R S G D E Y N I L E V P P W Q T R L V C D A H R R L A F R E K Y A A T N G G S A
RANDOM_2220 L A A D L C Y M P L R N L N T A Q S G S L K T Q D I N D L Q F A D E F P P S D T D A H F A T M L R G L L Y K M K
RANDOM_2221 L G A T A K R G S A M L E K A Y T V T K R K T K G I S G E V L D F G I L E L A A E L D L V R I N P P D S R D C S
RANDOM_2222 G E R W R L I D H P F N A I M A E E S C Q Y G V G D I G Y H T V T Q G L P A Q G E A N G L D L D T C P E Q Q A I
RANDOM_2223 Y T P I C D E P K I I V R L W T T D P L G S L A K I D F Y P M S F A G Y R G A N Q Y C H G K S P G G P Q V A T D
RANDOM_2224 S N N V Q L L S Q K Q G K F N S S R L P T T I A E R T A L Q F V T K S P A L R L W A G T V S Y G C N L F E L L V
RANDOM_2225 C Q E D F T L S F K Y A E L L L F Y R D T Q S Y L F P K P L S T L P M T D I V S F S A Q I L Q V E G A H T E S R
RANDOM_2226 C N E Q L D I L K G A E L I T I E A Q L H V E Q D R C H R Y R R G L M R G E I V G S F M A L I P M K A S L Y D S
RANDOM_2227 C I T D G I N D L R P M M T A Y E N H L I M F K Q P Y V R N L W S F L I D S W W K Y A M V G I I N Q L V I T D P
RANDOM_2228 M T T A D N S C F V T T A F T Q P S T T L F I G D Q N R K M A K V K G T S M I M N A T L W L D V Q V N I F V I L
RANDOM_2229 K F V K T C N I T A R I E N H P R G L Y E E Q R Q A W K K V V K F F T R V S E V Y A K R T Q S N S K N E G R I F
RANDOM_2230 L E V L L T S W N D D P T P T I V N T Q L E P F S I V R N T L T S T V G E L W G V W S T D G Y G I F V S L T G G
RANDOM_2231 P C A E H V E Q D N S K Y L K I L K I G P H P V I P T D A G Y G Q N R I I D G T K A E H L Q D E R Q R H K P L E
RANDOM_2232 V H E V V N R N N T G N E R E H P R S A Y N L G I V V D L A E Y L E G K F N E S V V Q I C G D Q G Q E I S H D P
RANDOM_2233 L Y T E R V H K L T T L L I A V F Y Q Q A Y I L N G E C F E V D G V M T S A L S V P Q E E D M N Q I E V N D L W
RANDOM_2234 A P K T L V V R N E M R R H Q S G P V K K I W D I V L T A E A E R E L G V Q L G L D G P L W S G R Y F M N E R G
RANDOM_2235 S D P V L V K F M L N L V G L A R T L M E I G K E K W A G Q P D N L A F S A P T E N A G G Y E E F N S K P G P D
RANDOM_2236 K F S E E A M N P A I L G I V E V A P S L G T L T L K N F T A T I V D F I P A T I L T V K V K L L Q A Y K N S D
RANDOM_2237 D S A V L V D Y M V A L S D I L Q M S T R A L L D I E T K M K E A K D K A F Q T V S T V A Q N F T L N V S P M K
RANDOM_2238 M Q P Q A A E D A P Y R L D A P G R E T P V A T L G Q Y L S S G P I N A Q A V E T L R S S N A F F N N L Q N A E
RANDOM_2239 V R V A D V W R E I R A C E E H D S Y Q V A L G P F S G E Y I P I G Q V Y A K G Q V F I L I S A R L P S L S S A
RANDOM_2240 A H E V R A A Q I Q S A S C F N T Q C S Q I D T I T T A S A H I G L K D C P D P V S E A A S N L G R F S C Y Q V
RANDOM_2241 S T L V V S V T N T A S R D M V P I L C V Q E L V L G A T I V F I K V E E S T E V A P V M E E G L E W L I F A K
RANDOM_2242 I E E D G S L S K L I T R T C Q W N R T S S P S S S I N G F S K G A S V E G E P H E F E M K S T S G G C S F H S
RANDOM_2243 R S E T S G A T R S P A N T G L S V R R N S P I A S N C Y A K R I R A E D D G S G A K E A D E L V E D S S T A L
RANDOM_2244 S T F L N D G M W T D S V M H N V H Y N N R P K S V K D V V Q G K C E R G G G I L T E Y W S Y P V E D S S A N D
RANDOM_2245 S I S Q T L W E F A R T A L L L A A E A A L T L W P H F Q A Q K Q I G H T H K A P S S I D S A S Y G Q Y H D R A
RANDOM_2246 E G E G L P K I A G P G G P Y F Q E L D P N F Y A I I K L M S G D Y P L D Q G H G A E S T V M M Q S Q R P R L R
RANDOM_2247 I R T D S E L K V P A G L L V D H G T F M T R E V N I P Y D D L L R V N F R M G W D P L S A S R R K A T D D E E
RANDOM_2248 K D K G T T V Q K F S S Q E A Y G S K Q I M R H R A S D A I L L D A P I S G G K Y G E L E R S N I I T G A R L S
RANDOM_2249 S M G I Q G G E P K S E G Y K F P G L Q T A K G D T F P V M I R F E S S A N A K V C N K L E A R D A Q E A A Y A
RANDOM_2250 G L R Y H D A K S G S V E A F D T N L V Y N L T E F V H N L C S P T M R K E S D A K E T Y M T D E Q D T T Q G W
RANDOM_2251 W P P K Y R D L D V K G M K S Q D A Q W P G R S Y D K S E G S F N I C I I D V L H L E V A K S K G H K M E G C L
RANDOM_2252 G P L Q A C T D K A A C L N S N N E P K D E R E T L H T A G L M L L V T G D L R P Y R P G R E R Q H E A P Y A C
RANDOM_2253 D S T R T A A L N R R G R E K T K A P F S I L L K K A L V V E T E A G L E K P Y T R D I A K P L G I K A N I S R
RANDOM_2254 C Q I V G Y D P M A S K T H A Y L N A R E R S V I K N V V Y S Q A A M K H I R I E V G H L K Q E P E H L K P I P
RANDOM_2255 E P D M T L G F D I E I R T L A I D E G A T G V N L F T T I I N N A R D E I R L R T E G K V K D E L Y A V Q D Q
RANDOM_2256 N S G V S H A D L V H S L A S G Q I A R V E T E R D A S Y P H E F V Q L W N A R E N V N G F V N T Y R A I P Y A
RANDOM_2257 S L P R I A A K A I A T V T G S P T S G K A G E P R R Q D S E R A P N M Q P V A E G E V Y E S G S D T S N G L T
RANDOM_2258 D R W V N H V K I G M K S I I D V A A V L L Y V M E G V A R D V E S F N T E T M V C K V I I A T Q E K V S F A N
RANDOM_2259 R Q A G A G E V G V Q M H A F D I P G T S G R Q F A P V D F W E N E P P L A L E Q N A M S P G E V N V L S R A V
RANDOM_2260 N S S F V L S D W K V E Q A R K L H I G G D T L L S L Y E S A A G A S D M E A A L S T V D Q F F L Q L Y D L L K
RANDOM_2261 I K P Y V F T Q D K D P V P K T I G G T L M I V S Q F G G N H P F S S Q W G P L T A D D M S G K C Y L G E V R L
RANDOM_2262 L N E T L G H G D A K A P P F L K H T R T Q K D P K M T W L T S S A V K T P L L M I L C F V Y C C G T W P F I A
RANDOM_2263 Q G Q I I V T W V F N T V D S L F A I V N K P K P A A N F A R H K G F S M E S F P S I S L Y E K I E F Q V G R D
RANDOM_2264 T S L A T R Q V Y I E S L R G S I W H G N C T G R N S M F A Y Q R D T A V I T C A I V L Q I W V H E T E E F I V
RANDOM_2265 Y E G G A E A E A Y D A N N E A D I A P A Q A Q A G E D R N S L G T K R Y T F A F W P R G S E I R R I H G D A F

RANDOM_2266 N L Q F N W K Y E T W Y N L T K M A G A P Q T T T K L N F R S R L P S L Q E V S K R T M T R A V Q H H V L K L D
RANDOM_2267 V G S N T S C T G S H I R N T K F Y K P E Q R A G R M L E A F L T C R V S A E P C Y L P L A E G P V L V R Y I K
RANDOM_2268 L N G K L T K T M Y G E A H I A G T I F I I E L S H S G G I Q T T V P F S C V D F T G P R I M V D D G V A A T E
RANDOM_2269 T N D N A D S H R T E L S K F R L A G S L L K A Q F R A R E V F T W P T A I K H S S W N L E K Y Q R E P A D N H
RANDOM_2270 S F E T S Q T M V E F P E Q G S M L T W A H H Y E L Y S K E G I F S Q V W E F A Y A Q M R I N G S S L D T R G E
RANDOM_2271 K T A G D V Y K A N N T I D P M A C A Q W P E E E E A W P A G V I P H L V S G S H S A D S D P G E L S S P E H F
RANDOM_2272 T S N C G V K G K F F R G V I I V V D Y H E N G H R E K L L S I S A R E L R M T T W F I M T Y T L E N T Q E K Y
RANDOM_2273 K R V A A L R H V E N F Y C A S K E K R E L E H N A K E S T G Q R P L E K V L C D Y Q A F P T L D Y K E D I Y D
RANDOM_2274 Q Q D L K A E D G V Y V S S S D P F L N C G H M I K D I A F V V V V D V M R R N G V D L Y G V V H R K D K M Q R
RANDOM_2275 K W P E V S H T G N A R R Y P P N P A F R Q Y V A V N V N I W K S F R W Q V C K Q E T V A F T L E T G W K D P V
RANDOM_2276 R E L A P Q R I Y S R I V S Q M T A I I G K I T G F Q A G A P D G I G I L V W K V R K K L C E S P Q N N F S N F
RANDOM_2277 Y N T T I S E T I K C H E M L E P D N S S L E L Q E S S C E S A K L I G A C P N R M G I K G A Y T S P M L M V D
RANDOM_2278 E E I A G P G V T R S L L E L I Y Q S H N K F I A D F L L Q L G L W W A T F S L I K H V C F L L P Q E S P N S T
RANDOM_2279 F G G S L V P I A S G L V G D T C T F W M D T G L F V D K L T M S Y M A Q G R S P N S I S Y R I N L N I C W K Y
RANDOM_2280 Q K R S H R A R I G E K K E K L G S A N A R K K I P D Q D G C I Q R L C Y G A V E I M T P V L E D V M E Y L V W
RANDOM_2281 I K G N R D S E G Y E R T V S G K D Q T K I C M F E G Q T E K D R E H P V T K N C E G D M V R C S A E G G L T H
RANDOM_2282 I A G S I G G F W P I P K E I L P E F T A N A E D D L F K S E S K D S A G Q N E G N T F S A I T R L K M G K T S
RANDOM_2283 I V G S W Q P D D E Y T N S K A L G R F Y P T I L G P L H G V K P F S A P R Q S Y Q S H A D S H S S A V L I Q E
RANDOM_2284 P E V K H D I S A N P M I A P C M L G K L R P E Y G W R Q H D E N R Q L S V R I I D Q K R N T D E R G N P T D G
RANDOM_2285 S R G K Y R W H F P D H V K T R F D L K K L T H D F G S L V I P S Q N E E K C G S T G N Y T E S P G Y A Q A G L
RANDOM_2286 G A C G K N T P L L S E F R Y S K G A Y A L G G V N T A C V C G R Q S K A D D L K K L N T V A S L L L F G H V D
RANDOM_2287 R T V V I E K I C T N G K M G M G T R G R D L F R L T G R W D L V E K I E D K G T E D S E I L E I L L D G I G G
RANDOM_2288 L A V L M P K P P R G Y P L N D G K L T C K A V V P K W T L N D L I I D L E Y W T W C F S L C N K S V P F L S G
RANDOM_2289 Q Y R T D G H C P Y W E E Q I I H E H Q V P G S S E P E T N S I K Q G K T R S A T Y L V V K L T S Q K P S K Y R
RANDOM_2290 N T D P L E H Q A N K G K S V D Q R A P Q D K L C R Q L E N R A T N V R A I I M V A D K G L L V K I S A I W E I
RANDOM_2291 S V P D G Q R D W Y E L Y F L K L L Q R N A Q N S R I D L G S R P N A K E V H Q V V N G A P E F S M K Y A K R P
RANDOM_2292 V H I S Y A P K Q G L A G H R T D T P Q H I N S L N R N S G A E G F L K R Q Q N G Y T V I S L P P S L S N G E K
RANDOM_2293 T T R I P T Q L L G E M K I L G I R T Y P G L E I D A S G E G E Q K G S W M D A H R Q I E K M M A G V Y A H L G
RANDOM_2294 T S S A P D N R E P D Y I N V S L S I N V P G G R S K P S G S Q I L T L T V L L G D A A M R F R F L S L A R K N
RANDOM_2295 I V G A V M S F H G P I L P T V Q P N Q L P Y D D S G M E G V Y A Q D A F E E R R D L T Q G N R G P G C Y N D Y
RANDOM_2296 D G F K P A Q S C I L R T E Q T V S S L P I A K L K Q V E E D V S S H T T K K S G A V A V N L S L V Y L F V L Y
RANDOM_2297 P T A T Y S N R F I R L G T T P A M R A R E R V V Y T R G L G S A V Y R F V N P C V Y A G N L E L F D M L S R D
RANDOM_2298 N L S D A Q L I H P E T G N M H K T E A T S S T T L T F N V T P S N Y L S F S N L I G H L K S P S Y V L S D K N
RANDOM_2299 D K N T E I L D I A L E N H I A M P S N T S E R P I E G A G V R S E D T E Q R D I N Y A S P R Y E Q H P L W D L
RANDOM_2300 R L K D P D F G H F Y P T L L L M R Q D W P N T Y G Q L A G G Q L K H R P V S G E S P T W D T A F R G E V P I C
RANDOM_2301 A Q P L R L L D L K D A N S R M A T P A V V L P N Y N S K P P K C T K I G Y P T M L W D R P G P Q L M G K E Y E
RANDOM_2302 D G S D H V K N S A R E V S E Q G F G V S Y V A S E K F S D E V Q T D Q I Y S E I I K R D V G R L N V R A T M L
RANDOM_2303 L T L S L L L W L S N S G L Q Q E V P L Q R M G D E G Y D V Q L C I Y G G M N T I Y E E D S A P D Y L K Q M M D
RANDOM_2304 D E D N L Y Q C V W A E D H N F E Y S M G L G V Q Q L H N A L S V K L D Y T T K A V Q K R D I S F L R A S G L Q
RANDOM_2305 E F E T V R V D P R N K M T I L V D P K N R V Y N I F T L W R I T R I A R D S K S S L V V D I T L G D L Q V N T
RANDOM_2306 Q E Y M E L S T E G V Y R C E L K P D Y D K P S N L R Q E A D S E R T E A N I E R S Q R I F D P N G R I L I A L
RANDOM_2307 E P K D N R S G V N L K C N K V T M H A Y G P M G S I V F N K L N Y M L L N S H G L P L E D D E F T E H P I Q R
RANDOM_2308 G A T H V D E Q G W A I A R I G P G L F F G K M L Y E K T A S G L Q T A T K Q D K A N T Y K S A S G V L N I L K
RANDOM_2309 G D G Y D Y R P W S Y T T L G I T K T K L D T K C G A Q T R I V M A Y N S Y G S M D E S I K E S P E G N R V N D
RANDOM_2310 L V E Q T G E P A K L V H G K F V D G R R N I T N H H I S K L V R V G Q F R H C P K R G Q V P A M L V N D F T C
RANDOM_2311 E S N L I Q C T A G E L L P P P W A T E E L Q F V D S R D W D D G K R L K D H A R A D G S R K K R A F T Y K L N
RANDOM_2312 Y G T V E I L P G K Y H A L L D V I R A E M Y N E A T L L K V L P P R T D L Y K R E Y L S P E T A K L L G W D I
RANDOM_2313 I D A D L R P E P A G S A S F S G P L T V T K I S V E V L T D L D M L S N L E Y P G P G P L L K R D W V S N W Q
RANDOM_2314 E Q H T E A Q A T L V M C E N Q C L A A Q L E V T P N F Y P P K Y P G H S N G Q V A V L G E V C D T A Y P A S L
RANDOM_2315 T A A R A W T D V R G W P Q A T D L Q P N G L S M C A T D R S N C T V G H M A W L L Q L S G V S L E I V W E I K
RANDOM_2316 L D A S K L Q G R F Q L D E R A N A T P T D G R L N V I L F I D S P P G T I G K P P G A D N L L G R S E K K T Q
RANDOM_2317 N P S I T E V M E L N E K E V G C V Y E A N T F E A A D L D P F L F E D W R L D C K Y P K Y Y G T A Y R S G T L
RANDOM_2318 Q A N R M R E D T C D P L S S Q G Q L E T E T L S D K L E A S T L Y P G D A D L T Y T R P H V R S E S E P V L G
RANDOM_2319 R E K R V P A A D R Y N V R A S L C K T F Q Q K G P T S D C G T G D T I R S V P Y K I Q S H E G R W T L N S L G
RANDOM_2320 Q R S Y K P N R M H E G N M L E M Q R Q I P G S D G L D T I A N K S L R Y S Q E V L S S L C D N A T F V A N Y E
RANDOM_2321 L K H L F M S K A Y F N G L M K E D G K K A P I K T E N E K S L E L F S E D K Q A N P I N L F G D R P F T A C H
RANDOM_2322 Q S E Y H Y I P V A I N D I A M F K A D G G T E I E E M K G W D K H D P V R Q F A I F V V R R K D H E E W R G N
RANDOM_2323 G L K A N N N A V V P R N T E Y D S P E R D F F K T N E P F P D I L Y D G T T D V V H K I C F L E G F M N D Q E
RANDOM_2324 L A K D K G L P Q S Q K S N I L Y P A H W R N F D T P M L R K L H Q S E Q Q S F I K F T N M N T E L K L G L F G
RANDOM_2325 R L S K V S S Y L W G K I I L P P P N A F P C L L W A L E E T L Q G A L A Y P D T H C D I G S S L D D S Q R S E
RANDOM_2326 L S R S P G V M Y M Q F A C L Q Y G Q D R E Q I G G E Q V G G R E V E L P K K S V T R T H L L G V A G I N G D I
RANDOM_2327 A M S F N D G P R P S G L L T F Q D K N D G A A N L K E I T D S E G Y Y L L E L Y D V N I D K D P Y R M E K G I
RANDOM_2328 A G Y F C E A V G A R E S T S P A K E N K S L N I E E E I T R P L M Q K F Q A R R K H S G G E W I G R D H E Y G
RANDOM_2329 W F N H N V T L S I A D S I A A S E R A K M N F L M T L S P G S E S D I H S K T F L I C P N E A Q I L I E M V V
RANDOM_2330 K S P I S Q A R E D Q Y R L R I I A Q H F N N K E V K K V A L E P S S L I V K V Q A D V T N C T E A G G S K Q E
RANDOM_2331 Y A E L S R E R S E P D E G M A L C F S C P K S T K D V V L T L R A S K K T N F Q V A K G L V R R K R M Q I A G
RANDOM_2332 Q K M K L G S I Y A K Y Q A K F T A S G F D P R G T V G A N P E E A D G Y D D F E T D N P P D F A L D K E A A N
RANDOM_2333 P E V T F E F D I R F N V L V L A V Q P E D L T E G G Y D Q Q P F S A H V D K P Y L R A T T N Q N V K Y R E L N
RANDOM_2334 T I T N G D P V L I V R Y K Q I P Q N I E K I K G Y G L Y Q H K I G L Y D S K D V E A K G E K L P G L E T T L P
RANDOM_2335 G I F K V G L T S G L V P L E A R S L A L L Y T S M A N W D L L K A S R T D T P T V W S A V Y V R K V T A F P L
RANDOM_2336 L D L C P K L Y F L R P P L L D E D D L S P T R L A V L T K P F S G T C E R Y Q R M E S R L I I R P G H Y G V P
RANDOM_2337 R M K A N W E A L T G K V G Q C M G W E E L L F S P V M E A C L D P S E A F L D D V I E E T C K G T K S N F P D
RANDOM_2338 S F H T S L L V R N K I L N E D S T F F G A P P G T I R R T A G H L H I A E L R E I N G A S D S A E S F R E G S
RANDOM_2339 L S D K Q E V F K P Q Q I V E Y R H E D K L K K L Q F K R W I T H L R N T N K V P F D Q I G A Q I R Q T H S L E
RANDOM_2340 P L S K V D F K Q L S Q V Y Q I Q R G W N K T L K K F V T D E T S A T Q G S L C D E S S S L K G V E L F S P K A
RANDOM_2341 G A R L A L I D G K S V R T G G R H C A T Q R L V L Q S A T G T P G I S L H P T L A L Q Q I H K Y R L L M V S P
RANDOM_2342 R G K G E D K L G L H D R E G R Q H E G C C P G I A F S P L K F T G N A H Y A V S S K E R V H E E W I K S D D G
RANDOM_2343 I N K E P V A E T V G N R K T E A M I K G D G R L I F Q V L I M P L T P N S A Q E K A E G Y L R T G A T V L V R
RANDOM_2344 L G G A Q R I A L S L V S Q S W L T G E P L E T E Q T I K N I G G T G L N S V R G A G T Q D T G V I D Y V R A N
RANDOM_2345 K D L K N G A P S I V A E V K T V L S S A L D P G D A S W S D A E R D T A K R T A E G N V S Y K V V K T D S Q C
RANDOM_2346 K R R T L V N A P L V Q L M A V A L S E L R T T R C H A Q G A N E I L S T D E V E Q V I A N G R G T T S L R Q R
RANDOM_2347 K Q M Y L T Y W K Q E H I P L C S R F P P K G K I V D G S L L R V A L E A N D Q I L A A E R G S A S H Q S H G H
RANDOM_2348 I R P E G D D D T D A T T L D M F Y N G S D I K M T V P S T I K K P E P Y C Q P F L E W I F L G K V V Q D K E E
RANDOM_2349 Q F N E S Y E L G C V W D A S A T S I N Y W M G F Y L Q P V G A N V V I L A Q A L D E Y L P L K Q F E V A L F S
RANDOM_2350 I P K N N F L C Y E L G G A R V L L L S L E G E E I P K K P T P A Q T A Q P F I T Q F Q Y V N Q A A R T A L E I
RANDOM_2351 L S L G F A R F Q G H A A H K L S G Q D E M T R E P V K K M N I Q V V I I V D D P D S E R R K M C M R D L I D L
RANDOM_2352 T P D M L T T S Y Q E S V Y F N V Y E Q Q Q N G K E G E P T K E S T E L N T M N A A S K N M F N V K G A S K L T
RANDOM_2353 V A P N E V V T G L C S G N S N R Y K V A K A L G Q N K Q V S V A I K V I I R T S I G K K V T D W D L T A R P A
RANDOM_2354 Q T T N S L K P V K S E L P K S H I L S T L H K G A L S Q K N A K Y F K V F R D D G P V K R T H G V N I Y G I Q
RANDOM_2355 L D A S S P S A E A C E N G W E E P G L G R A M G S D I D K P I S E L G I V L A G S A C G S Y E A L T T A T D V
RANDOM_2356 A L R Y V Q I G P F A K T D S H I I K L T L K Y K E T Y A Q K P E A Y K F K D Q D E E R I A H S V A M Y V Y C I
RANDOM_2357 Q E G A G N S S L V G L M V S G L T K S I F I D K R D S E T F E R G V A V P Y Y E S P A K L K L V T Y S S E L A
RANDOM_2358 I T P H A V K P N L G G T R C Y F C W L K E N I L G K R S S R V E I H G D L V T Y E R F E A P T I N V S F S E R
RANDOM_2359 N L V L P R A A P L I L E D E K P W S H T F W G G A K A K L K G P A T A D E A R K H G D D V G E Y P V L D V E R
RANDOM_2360 F D C I V V I Y C P L D V Q P S E E V R R R L N E W S T G G F D K F S D I V S Y I S F L S L S L G T I L L A H P

RANDOM_2361 L E H P F Y R E F T V A P V P Y L L L L I L L K P Y G N V R Y L A K L L S A S Q T N F S S D P A V Q F I F V V I
RANDOM_2362 W A N K P I E A E N W H K R L C D G P I Y V F D L Q A S E D K F V T K G H N V R T I Y G A P N K Q Y G G P D L Q
RANDOM_2363 N A T Q Y A T L D N M L L L E D S T Q I L F R P L L S A L P I L E E R S M V V L L Q E N K C A A N V G S P N P L
RANDOM_2364 T S D R L M T E D P S A G L G M C Y E M Q L Q I A N L G D S L R V G T Y V G L P E T P G V E T L I W G K G N I A
RANDOM_2365 R D I A G I T A L Y Q I V K G K Q I D L G E A E Q Y Y L Y D L Y T K Y Y A Y V I G T R F K M P D G E P N P D N E
RANDOM_2366 G G G P G N G L L E G L G E A L L D N E I K G Y T F T N Q K W G K G E S A W C T Q A D P G P A V I R E K Q T A G
RANDOM_2367 G S R Y A R P F E D G V R K V P L M D P S N K A R Y K A V L I I E A I L Q I T E D Y M K Q A A A N A F L S I V N
RANDOM_2368 C G K L G E T R V Q E A N P G C P V Q M R R D P P D A L V Q I F G V D R V Y R K P V N L Y V E I D L W S N C P K
RANDOM_2369 C D P V L R W W A D T A P G G G Q G K L R H S G D P C R Q L V S S G I Y A I D G E V N N E E N I D G R L P L E T
RANDOM_2370 I D E Y I V K N C W A T E S S V A H M R Y L S G P E D E A L D N A T T D S A S P T L L A F S K S Y L H T G N E M
RANDOM_2371 S G V Q N K D Q K D S G K N S G G K D S H T V F N R R L G H N G R L I E S V W E L G E N V V G A T Y T L S N L Y
RANDOM_2372 S K S L A L H A V L P D N S L R E N Q P A I T C C K P E F I R A L L L K E Y K S G A P Q K V C D E Q A S V A I E
RANDOM_2373 M E S E F S I G K G T L V V F R K D G L I G Q T H K S I Q G W A V G E A T K G I L N L Q G Y D F F Q A L W E E A
RANDOM_2374 V G P I Y S V E L D R D R D V Q N Y F S Q N S G L G P L N P N G L T L R H D Y E D K M T I N A Y A T D A T E G A
RANDOM_2375 T S G N V E P P T V C L T L H V S K Q V T T D L V D T F E Q S P T T F L Y F R D Q D I T A P I N Q G E D E S H D
RANDOM_2376 R R M H F K F W L L S E R N K D D D A T L P D F V D A F D V E H L L E R N V K E F S V L Y N G Y H L D P K S T H
RANDOM_2377 K E T Q T R G S P K G Q I K F A K A G Q G A V R D A D S A S G Y G V I Y L S P Q A A N P P P K H L P Y D E L N S
RANDOM_2378 T E P E V S D E N L V M R P A E N C D K S T S D Y S Q G Y E P E V N K G T E C G S T I F G V H I S K T E V R K G
RANDOM_2379 C E G R T V K Q I F A Y S Q R A P E D P Q V T Y K R D L P T N M N I I I E L A E V T N Q V R R D P G G E L P D P
RANDOM_2380 P V L R D K I E G R N E Q T A K G R A E G V D S A R H A S H Y L L W K E W R S F G I L V G Y G H P G P C H L N L
RANDOM_2381 C C L K Q T V A G T E V A V V K P G Q V K E D Q P C M L I S G D W R F P G E E T K L G I L N E P A S L N M S E G
RANDOM_2382 A E S Y R T L N A R K K N Q L V Y Y P S T E K L S L G V W R G K E Q Q A D G D E N K K G L Y F G S L I R S D V V
RANDOM_2383 G S V T T T G V Y Q R G V A D L A P G L K R S I V Q I L N E D T L R G Q L F V V C Q R F S D S P Y D M T D S C I
RANDOM_2384 Y S T I I P M N I G T V Q E D N Q L V H T E S L K R I P Q Q V F I S I V P V R A T Q I G A D E A W D S S T G K C
RANDOM_2385 R W G W Y P E D F R K V L N R L H F P A Y F S K A F L G I G V K Q P H F L I P D K F P S S I A L N P A A T C V I
RANDOM_2386 A N G S E D T E I N P M S F G V D Y E K E D R I L F K V P A N W V A Y L V F H C S F L F R P M P E D A G T L G L
RANDOM_2387 S Q E C L R K V S Y T E H T G L R L R D L Y A T Q C E T E A T A F N G S H Y Q L P Q L Y D I S M R C V L A K N A
RANDOM_2388 N G W E P G A V H V F H E F D V F E Y D V E K K Q K Q D L K S R T E P K I E V D V Q L T E S G S Y E F P G S V H
RANDOM_2389 K F H H A T F S S V W E L G G K P K S Q V D R S H R A D R K V T A T I V P V A N P R S Y E S T Q G K W C R A V S
RANDOM_2390 C G C S L A Y T N L G A L A I A D L C L E A S G E N R D R N V E D G G H A N G R T I L E K I E S E D Q A A L V Q
RANDOM_2391 L D V G M S E V A G W S I Q D H D N A L Q A L Y N T C P E T Q C I G L L K G S T L E P L G G T D G G F L F Q R L
RANDOM_2392 P T S E D A I L M K Q P I E F C A P P A N S L S E D V S Y N P S P A L L K P T S K V T V S M F V D C A A T I G E
RANDOM_2393 G Q P R H S R K T K S Y L P H G A Y M N L N L N G V R H E V C Y R P E G K S A D Q V W S P A Y K R A W T Q V H Y
RANDOM_2394 Y M A S V A F P I D G R R I S D F W S S R M P E K G N E P A T I L D R A R A V H I E H V R T Q L L Y S S A M A E
RANDOM_2395 D Q S I L L F N K S H G I S D S C A D V F S F N S A G N A L G L R F I S N K F K V I D G L A K H V S M T A G V P
RANDOM_2396 E P D T F E G G K E K V A W H N P S I G P R G S R D K I E L N A P V F R E N S K L V R G S N F F R P T T V Q T A
RANDOM_2397 Q A F R I V N A D R Y S R T A E D T L P L P L I N V D V N T I T F A P S G M Q S V Q D T M A P L S I F L N E E G
RANDOM_2398 E W F S E A D V T G A N A D E G A Y L Y G R H P S P G N T K D S I R P F Y S S V V L N S D S R T C S A K S P H V
RANDOM_2399 E N A V E Q R Q Y L P H M E P P W P Y G N H I R A H F S S V D I S S D S C V Q D R V A H E Q A S L L D A V M V I
RANDOM_2400 C P N K I G A S A I Y S E K S D D K G A W G I L A A P M P T I P H Q P I A L L L I N I D G G T E K T Q S M S Q R
RANDOM_2401 V S A K E L A V Y V F K C Q K A K S F P L R A G V F Q Y S L A I G S F M S D Y Y I K V L D D A T S A Q Y V F R G
RANDOM_2402 K L H I G D W L H I Q A P E G N L A Q R L L D R D G K L Q S T R A T T D G D P E K R A E V M D I Y S E V P E C L
RANDOM_2403 D N Y E P L G R G I Y P A T V P L S D L L A S V Q F S D N P Y T G E A G L S I M Y Y G K T D K K I T K T L D G A
RANDOM_2404 S I Y A C R H V W I L L E P N K P E T A T K L A P S I I K L L Y K W V I V K T V V E R P N V S A A P S R I N I T
RANDOM_2405 I M Q K A V N E S V G F L A N L W F P C S K N I W V M E S A P L I A Q I L S M T G A L V H P L L Q P L C N I G L
RANDOM_2406 L E I A T S N P K F K E H G S D W G A N S I I D S T L Y T L A T A G N L R Y A E V K G V L T I V T V Q N N G V V
RANDOM_2407 K N G W N A P N K A D E E F Y V S G T V S E S V L A D S I L E V G R I N R K A M S N I K K V L T N V L A D A Y R
RANDOM_2408 Q R K L G E N K K W D Y M G D R L F V F G E G G D R G P F A E E A N H R A I P T K G E L T V I F P K K D P F G I
RANDOM_2409 G L D G L E N F Q G G M V V K V T P E D L H D A E I A N D P C H R S V Q A D G A L I T S L P H G K I F I S K N A
RANDOM_2410 K R N L G Y V D D I A L L L Q N D G V I S N S I E D T V R N T M I G V F Q K Y N I L I P L A S Q D G I A L S L R
RANDOM_2411 K A E G Y S I Q H A T V N K A G L L F G N K R F V C L F A D S L Y N I N F Q W H N Q K S F R Y R A P P L K P T G
RANDOM_2412 D L M Y E T R T F Y L F F C R K Y V E E V T G G G Q V L E Y N Y V S Y F T A T T A F P L D V I P G Q H V L A V F
RANDOM_2413 Y D T G V K N E Q A I E L Q G G K L L V Q I S N C D I K E S T S I Q N D D D F D H I N I H L W S F L D A P D I T
RANDOM_2414 P Q T Y I A K F A S E G Q L Y K H S G P P G N F C E G E V L F S C I L S V S R T G V P P I M S S L S L V R S E H
RANDOM_2415 K S E Y P D W A P M Q K S I K S A A S I G T G G Q T E S G P D S S S V D V W S C E R G I N C E L I A K Q D R G K
RANDOM_2416 Y K E A F P V K V T Q L A V T F A K S G V N T T G Y A S K D F K D G A K S K L P L N V S N L I L N E G I C D F E
RANDOM_2417 Y D A P M I F Y D L N S L F E R A D K Q S I S F P L T A A A V P I K I T Y T E V I L T S R L L I R A N C P H L A
RANDOM_2418 T P D P S W K Q I R R G G V D F L S Q F G N T G D S N G K L K Q L L P A T S H K I A A N E K I T A G Q M N G I M
RANDOM_2419 G N F V R P N E R T L G R M E C I G I D D E Q G P F E R Y L E T I D S I R G E A K S G Y G Q E A N F A S A Y Q H
RANDOM_2420 S G A L A L R K N S S P I L A G E Y Q S V G V P H G D D P V A S R S T I Q R D Q S V N V C I E A D V V E N M E P
RANDOM_2421 A F M D I N R N Q I P L G I G L P L K L N Q L T F V V T Y E H L I T N A R K Q R W M P K K K N S G K V K P V A A
RANDOM_2422 E A E D Y T E T E D V C N S E I L H W A P S L K Y K T Q Y E V T I D L S D G C Q L R K Q E M E A H D P E R P M Q
RANDOM_2423 E L G S L G T P R K I R G T F S I Q L I S D C V G D F R G M S E N R S V R V N Q L D Q D V G D V Q A T Q A T T V
RANDOM_2424 Q R R E A R C F E V H H T K L N V S K P I R I V T G A V M R I I Q K Y D G S D A K L K Q G N S L H A I Q I S E G
RANDOM_2425 A L I Y Y F Y K R I R E D K N D C A A T M A G V R D T K S V G S Y S C T L A N L L Q K H V I G V F S Q C A G V M
RANDOM_2426 I S A H N L E I T S L E L D C A L S V S M G D Q F L A L P V V L G T N T S G R P A D N T S R F A R E I D Y A N S
RANDOM_2427 L K R T P Q V S A T Q G M R N Q R G K P L Q G V I Q G S I P G G L Q E K G L I S S L L Q F P K Y D T P G C N L I
RANDOM_2428 G T Q V Y Q L T T H S H R K P I K L L D P R E P H S V K P I I N P N T D F M M Y P G T N S F Q V S V H P Q C Q Y
RANDOM_2429 G R E L D L F W D V A G S T F P V Q K A R P Y Y A I I D N D V G P C L D L A R A V M L S N E G K N F A N G Y G F
RANDOM_2430 G G L I V E V A E A Q R A Y L A M R G K E E A K E R M K K V F Y K R C E A E N E Q E D W C A E E D S E Q L E D I
RANDOM_2431 V R S Q V Y S D I L C A P M R S T L A R Q E G I E I N P E D K L G A A P F I G S E K F E G A H Q S D N Q T L I A
RANDOM_2432 D A W G Q K K N Y K T N Q E N R V L P E R T L V V L I I F P E P N E L S K Y V N L V R S R P S G I Q E N E T D A
RANDOM_2433 V S L A R T A V P E G G V G L R L L R Y C D A V I G G G T F K Q R Q N L F T A L F D H P P P P S G L A H V I W I
RANDOM_2434 H R T C A R L A K I P Y Q G K N G R G L I V S Q K F N K S S D V L V R G K D P S G D V D Y P S E S Y Y S T E K L
RANDOM_2435 L D N S G L T P G V D G Q G Y A A V L Y S E H G E G K W I T D Q L T A L E A K G P R P T S A K H Y P V A E L N A
RANDOM_2436 G E A D W N K K P L R P D Y I D G T A G C I L S D R T Q Y Y K K K A L F D S G A S I A T S P T M P Q F V A H Q G
RANDOM_2437 R N V S Q R K P R P T I K A P D L F M S M K L P D R G S N D G G D P N S I G I E I K Q E V A P E K N G R D E D Y
RANDOM_2438 S Q D K Y E T G Q K M N V I T Q A Y A G C D S L L N G D T R L F G G A P M T T C I S A D K T L I R D S S V F P E
RANDOM_2439 A E R S E Q S Y D L N R Y S T R Q D F H Q P R D A Y G G K L E A G T L M P S Q F S Q Q A V L S N K P L V P Q G K
RANDOM_2440 S L T Q E A R T C C P Y A N V F D G F I L S T F K E A L F I L E M A F G M L T D V C S T K R K R S M Y S S L E L
RANDOM_2441 K V S I P S R T Q V R K I P A G Y I N S V Y H R C F W Q K T S S E K E V I L E G L P L A A I A P R V L G R D M D
RANDOM_2442 N E D F Q D C G K A G E I I S G L Q L E V K K T P P Q E Q K P A G A R S G W A R Q E R H T R A A F F N E M C F E
RANDOM_2443 K C T M V I D I G S G V G G A L A E R P T L L N G F V G L D N T T S K K P H V Y E K P V R I I I I V S K T Y L W
RANDOM_2444 H I S Q V T E P A G L W S I I R H P K N F S E T V V Y E I E G K W A C E K E Q E T T T V N S R I S N I K C M E N
RANDOM_2445 Y S G N N N F Q A L K A P G G H V Y M Y D T L K E Q P T T V P L F K G L G N T G N T D D E Y A E Q D I A S N C L
RANDOM_2446 G D A K I T D K H G A A T I Q Q T E E A S Y S P S G D L L P K D M K W F S K G V D K S S E A R A T K Y S A R K E
RANDOM_2447 S N T F F N K F N I Q G T R K E E G C T N S C Q K Q K Q K M N H C S D N G V N F C L K M L P F E L W P H V S S A
RANDOM_2448 D C R V L R P Q S F I E L L L G L A A S W K T L S S S S E G R D G A E K T F N A S P T Y R F P L V R L L N I L V
RANDOM_2449 D D T I G A W S Y V G Q I R H V K I D S R S N G Y H Q F L N G L N R G G L E F R T V V Q R Y L P A I L S A E F S
RANDOM_2450 L D H I N D E V M T A Q A S E R L S A N R M Q A K E A S L S P T T F P N D E T D R C G A A E R H D G N A Q P M Q
RANDOM_2451 Y A L I S I A F L P A K C D E R N M P A R W L R N N I I T P N N G S H V V S G A A E E V I L V A Y E A V L F E L
RANDOM_2452 G P A M L C Y E K C E L Q E L R C A P Q P I N D A G S Y G S H N R F N I R R A S C T P L T V K E Y R T V Q S R E
RANDOM_2453 W V Y T Q L G K K N K V E A S L A A I K G Y D L A L T K S D K E Y G T N S E I D A E S T P S V T E I L L G R N K
RANDOM_2454 L I P G A R S S H K V L V A N L A P P N W Q I D E F H W E P L P N Y P A V P Q Y T A I T C G H I Y T Q D N Q A F
RANDOM_2455 L C P P E G V K V V S V Y V P I S Q L Y S T T G G L T T H E N Y T R K D N L A A L M G I F N V T T I L Q G V H E

RANDOM_2456 S E H K Y Q F G G F R F L W V Q R S E G R T L K S R F I K K R K G K L A L T D T Y Y T Q L F R E D P L I L I I K
RANDOM_2457 V T D I P E I R K C S D A K N W L L F Q A D V G V A I I T G R R R E K I A D I W C M L L E G T D P P K A R M K L
RANDOM_2458 A P L Q E S T R Q L G D S F A L D G R F A Y L R L G A H L H G S Y L R E A Y F K I D K D H A L D I G F Y P P E P
RANDOM_2459 C G D A T P E Y A G T C T L I K K W P P Q D L P R G Y L H S A S Q D W T T N V I A C A V K N Q S P F S K E T H K
RANDOM_2460 D H L M N D G T A A P A Q D L T A K R L E A K S R N P L P M L E G T R L R Y S T V M K A R A G E N A K Y R F C V
RANDOM_2461 E D V I Y V R Y A E Y K L S F A K N V S I S A L L S T Q L T L Q R D N I A K S P I A G S P V D F N P P Y T S K K
RANDOM_2462 L L D T L F G P L L F W S A I K I D K S I L H N T I D H E S E Y F G S D M E F E V Y M K E I S H N V V V P G D V
RANDOM_2463 T S S L A T K I G G P A N K F W R P V V K P A V N A E A C N I S V N S R S N N Q Q V L Q N V G G L W G Q L D T E
RANDOM_2464 W L R G L S P K L V L D T L R F V Q F K E E L A K N V D H S R V F T L Y L P P G M I Q F T K L N A R N W L A V S
RANDOM_2465 S N A G L L S Y K I D L L Q R E F C S H K Q K Q S N P N A N F A W V V E F N S G Y L T Q L C I S K N A T L R A P
RANDOM_2466 G G S E K G P Q I S L D L R K G L F N E Q N K G P R F Q I T L T V S N M R P E G W I S A Q W K G E P G Y P A N K
RANDOM_2467 K T N I A F E I Q A G E Q R S S A P P E V R C Y E R Q L P I H G D A F I S F G L K D G K I A G I D Y L Y H H E N
RANDOM_2468 A V P E S D A S K E A A V R F D C L T A N I R D I V S Q G V V G L A Y G D Y Y R Y G L C K E F V N M G T N G P D
RANDOM_2469 A Q D Q Y P Q K D K Y P G R L E A A F I L Q L E S I F A S C V V L L A I S F Q S Y M V P N T R L F E S R G G D T
RANDOM_2470 P E P R W R K P Q V H A F D N S M I S T G A L G L Y E M A Q L K L N K A P T V L N E L G V S F M L F N H T I N A
RANDOM_2471 G A L K P F D R R I G D P P P H K G K T P F V T V G R P Y E S A G V Q F T C V N G S D V Q K V E A G V K S K L A
RANDOM_2472 A Q A L L E I Q Y V I S K V S Q T L R I F G S R V S G P F F K F Q K V A I P Y M P K P A N V C V D K D Y S L F L
RANDOM_2473 P A P A Y S N L D N S P Q N S I F K F M V N I Q K S M L W L C I S E L P L G A R K Y G P A L S L V N A F S P L A
RANDOM_2474 N D I P R I V G A Y L A L I L K G N H D T K T D K E D A G V S E S E D G S V V V T S A S W A I A A T L N W Q K L
RANDOM_2475 D C N T F G S F D Q T G F D R T S K S G Q I I A Y I T P S P L G Y G E Y N F V I P L S F T H A N K L K R E Y T L
RANDOM_2476 T G I I Y P K K D K L K M S L V S R G D E S F H R E L F K N V L Q S L L E R K G N S T K N L N H A D S V S V G L
RANDOM_2477 G D M I Y Y S M P K L T G R Y A C A S F Y S V V K R R P I R S E G A L T A E S Y L N R K D G G N S T W E D L D V
RANDOM_2478 D R K D E G L I K P G L G S S M Q S N K Q H P H S S V K S H G T V K R K R N L M N K L T Y K C N S T T Q D K C M
RANDOM_2479 D M I N R G M V I V L V N A K P A Y A I T Y G S A E I G P S I I N N T F D H L P A Q Q A V G T R L E R W G D R S
RANDOM_2480 N K L E Y V N H G A P M V V K I A N F Q E M G I A L A P P E S A L S T L T V S E M R W S T V L I C S W D V L S N
RANDOM_2481 G I P E M S P G R Y I H A L A P P E L E S A R A N R S Y N T P R F D F F D G S G A G Q R S A K P G M E N N M L D
RANDOM_2482 M A F Y A C G D K K Y E G G A F Y N S W G R L K N L F L A G G D V K D A N R E I L E M L T S S G G G K E F P V P
RANDOM_2483 K N E W H E P F I E R L N S R P R C N R D G G T L P D K I A N Q N N V M S T D R W A M A V I I A E A V P Y G D R
RANDOM_2484 L M I A I D D Q R T Y I S G S S V Q C D A F Y S K A I K K T A P K E M V W K P I L F Y K E L G L D E G L H F L A
RANDOM_2485 D I G G A C S I N S P P G F L G T L S S L K H S G T E M I A F K T A H M V Q G L L V Q G K T F G C M S I H Q A Q
RANDOM_2486 L L V F I T N D V Y Y K V Y A T D A L G Y I L N R A A D Q N P P K I Y E D L N K K G P R A T L R R D S T D E R H
RANDOM_2487 E E Y S T I D F L A N Q N K G E E Q E T S L Y R D A P M R S S D G D E S A S L G V D I N G F G K P N E F S T E S
RANDOM_2488 A V T P Q C T T I L A A A N L F T L Y K T E L A F M E A G L D H I P P A E N E F A K G N L L G M Y T K S Q G K P
RANDOM_2489 T A K L F R G V L E E S E K R S V K Q T W N A S R N Q R T R V P I G P D Y K A L I D M M A T T E T Q A N E E S W
RANDOM_2490 Y I E L L T D K S C V N A I D S I R E T G M S S H G T R K Q L L A F V V G G T V F E L L D P E S P G C H V A R D
RANDOM_2491 P G D L G G H L V P F E K S P V E S C D E M H P D L L P E A I Y K H K G P V E R G E P S P F S P Q Q T T L V D Q
RANDOM_2492 D L C T G S G G R K A R V N G S G R Y K P H T K V R Q V V I D M T D P L V W S S V G F Q S L G M Y V V A V P K L
RANDOM_2493 Y W V G G S A S S P N A D N F E D T E N S G G D E D V L G E G Q A K D K S V W S G A S F V T L P S V I T N P G M
RANDOM_2494 F I H W T T S S L M A G I Y K L Q T N L Y D T N N Y T V G R P S M A E E T Q S Q I Q F G Q Q A M L A D Q F E V I
RANDOM_2495 A E K F E R R D G P S N Y R V S H G Y R S S H E V L R D E E C P G P L E F S E I T S H S T K V T W W L Q P V S T
RANDOM_2496 G T S G L L A I K S T L K A G A Q N N T V C K K I F G L L Y T F T S S Y K L G V R Q I V V E D G Q H K S G V H A
RANDOM_2497 S L V A I K R V R Y S K S D T C W E V A R D H R P F E L F Q R E Y S C F A P G G K D P N L E V L R G V V G A L L
RANDOM_2498 E E T N D Q A K L I I R V L Q T A F G H L E C Y P E Q G D W E L V V K K E M N A V N D G P V K D K I L V R T P V
RANDOM_2499 N N S L G K V G K Q I T V A F E D L R G G V Y C P S S Q G F I F K I N D S N Q V P T D N E C G K R E D E P D K K
RANDOM_2500 D S K A G H S G V D P F Q I H V I V G E H I T P I P K C D L R D L R E E L F N W N G T T V T F M K Q I L Q A D E
RANDOM_2501 S N H G M Y F M Q P F G I P N A N T D Q K L S Y I D P L K C I E V L L N L C T E P V P Y S R P A E K K D T E T P
RANDOM_2502 T A S E L L V Q A S C L A V V F A V R N P D E A G V P I R A C R Q E S F P K L V E V L F T L M N A F K K P E E Q
RANDOM_2503 S Y E N G L P M Q S I T L L R P D R Y G A P P V P C N Q L Q V T G N L D I S T R N L L A Y A T N Q D W T Y F E M
RANDOM_2504 L I D I E L E P D V L C L A K V R G C A R F I F R P L S L E A D D Q A G E F A L K P T V E G S T C A N T A V L D
RANDOM_2505 P D G Q H L A D L D D L E G G F Y L N K S M T L N F S D L M M D M A P M E N R I K T G A V A A Q F P R Q P H E S
RANDOM_2506 M V N V R A H G N T I L E E S G A N A S L L C I T W G I D K P L T R G A E T A A G A F P I A A G A M H F K L C V
RANDOM_2507 K L F K P P N V P H S I E S Y L E C P A P M A R L Q E G D C N P A F T T R P D N G L E F A Q I E M N C V A R Y A
RANDOM_2508 P L R M K I G F Y K P E S V N I E P G S L G G L T K D K Q D Q G G A G I E T E W G Q T A C L G F V C T G Q N A G
RANDOM_2509 I V Y L S L R E R I S Q K Y G E K F L A T S T A L L A K I D E H M K F R H P L L R S E C Y K M R L G T H T V D A
RANDOM_2510 D L L D L D E P Q A I I N F D E A T S S S Y G T T Q L I L D T T S Q P G V P S G T H R I K G T I I S D K E M K F
RANDOM_2511 I A R P S I T P T F K T F M T Q D V T G A I I K L N S S P I K K K Q V D G P R Q P Q A G A L K A A R I T P N N D
RANDOM_2512 Q K A Q G A M E S P C V G P R I P A L Y C A D G Y F Q S E D T A L L D C A R M D E V G H S F D Y N S D C S E L K
RANDOM_2513 N T L Q M G E C R E E G N Y V G Q Y Q L D M G S C Y N V F D L Q E C A G W P I G Y C R S Q F A S L T V L L H A N
RANDOM_2514 A I A D S L Q V C E N P S K N G E H P N A D N I Y V Q A A N R S S L S A V G Q H C Q I C G K L A L L Y E Y V P M
RANDOM_2515 P S G N A N L I I H A G I L P L A A T R G A A V R T K G G N S Y L R L L I Y G V A Q T S G R V K Y H D K R I N I
RANDOM_2516 G S T P A W K R E I G N V P T L A R E G K A C V G V T V I R W L F R S N V I L Q K D L D T F G P H D N S V R F F
RANDOM_2517 K V S Y L Q G M L A E I L S K G I Y S P S D Q K R G G V K L E A P T L W G A A T Q Y C I H V L N E D V C F A A E
RANDOM_2518 V L G F D P A F V N R E D L T E K C K M V L G N N P S F E T S T S P V T N D L G L S R G E D S M E Q Q D K V Q P
RANDOM_2519 E S S S F R K R F C V T V L I S R A E P V K S A F Y V N S A D D D K A M K E V A V Y Q R Y N L K N L S E I A T K
RANDOM_2520 Y L S P D S T D Y R N V V A E T A N P Q N G W P F G E E D D E L V Q M K S V F K F I P A I Y N K F H E E I V P S
RANDOM_2521 S R Q E S N Y I I W P A I S Y S V R V F R D L D P Y R V W E Q N V W T I N G G F L P D P R V P G E T E V R V L R
RANDOM_2522 R F C I P E S A P G S Y S L T A G F G G T L F F Y L V K A D I V M G F A V D G A D V T Q G E K K G Y A R N Y L N
RANDOM_2523 A F D A K K E R S Y K S A S A P V A Y Q K R L T T L T G R V L A Q E Q P Y D T L A N L P P E T N E N D W E V T P
RANDOM_2524 D K D D H K H F K N L E Y S E T V L C T C P P D R L M M I L G E I F Y N G A N E R Y S G S V I N H V K V R I I V
RANDOM_2525 S C P C P I E L G L K T H S V L D F V A H G N Q H E A H E I L T T G L P K L P R S L S A T R H A V T C D I R S E
RANDOM_2526 I T G Q T R R V F E N G S K Y I G T K D A D P S P N Q D S K P S T G G R V S S S F R A L Y F D E D A A P I Q G Q
RANDOM_2527 A A D S C V G I V Y E P L I L G I K I Q D Q L N A K L S A P T P V S K K N C D A T G L L P Q W E K G H R G R P I
RANDOM_2528 V V P M E W P V E L C K S A E L V A N F K C E E D E R S I R A C P S Q N S A C G A F A K R R R E E N A F D I F I
RANDOM_2529 E I Y A T G Q Q H K K S L S P L R A K L E S M V D G R E A L P L D R F D V H L D M V K R A I I L A A C D A G S S
RANDOM_2530 T N K R F K R P I G G L E T N A S N P V Q V N V K Y E S A G E F K S F V Y A K F P Q D V L P H V I F K R Q R S G
RANDOM_2531 L V I P T I H G P S K A E K D A G M N N N I A S Y A P G R M I N T S V G K S N E P V T S L K F K R Q Q S Q P G S
RANDOM_2532 T G F A T G T S F T T Q C V F N D A C M K Q E G N R E D A E P C E Q V L L K N D T L H T S S D K Y K L K Y N K L
RANDOM_2533 F E A N K T G E S S G R F A Y D R T V D F Q K L S T Q L M A G G Q K G K C T Q P V K N F T I G R Q L L L G G N L
RANDOM_2534 T E L Y E L G S S I L N F K A R Y P Y K Q I P Q E A V S A I K K F V Q T P G K P S N Y N I M R V H L V G A K P A
RANDOM_2535 M V M Y V K K L T G G T T D G D A D S K C P L V C F D V A S W L W D V L L K I V N H I V G R K T V A C K F G E I
RANDOM_2536 K Q G K L P P T I H L T P A L K T G C I P F Y V F I K G P A R I R T P F D V L G T I I R V F L A L R A K P L Q A
RANDOM_2537 F P F A T G L A D I D D F I L V L A G K A F D G C N L Y E G P G E L T E F N Y S D T K R S L D Q L Y E I V A Q T
RANDOM_2538 G V R S V S Y T V D R S E E V G D F Y N L I S G L G P L D L V G N K A E K A S I C T Y S C E V E T D C R P A C L
RANDOM_2539 V H L N F S D N S L F E V Y L R G K R S A L I Q L G K V V K G H E D A P I Q Y T N A T T V C A K N V P D G S S M
RANDOM_2540 S W R D I E G A A S E Y E L V N R A F D Y K F R N G T T I E D Q T F P V H C S S Q F L I T I I I T L D S S F D G
RANDOM_2541 L P N V A R G E V M S V S G H A V F F R N F K N N A N A Q G D N R A L K K W D I V I V W A F D V Y T L G F K W N
RANDOM_2542 A Y M R D G F H V L Y L E A P Q K L A L V P C I P T V E Q A Y P L Y S V P S G K D K I N I V K D C V G T N G D A
RANDOM_2543 A V E I L D E G L D W L V M D K G K P N N S K M A Y I Y S L A K D L L A F K S K F D R E E G Y P F I E A Q V T A
RANDOM_2544 I R A S S L T S K E Y L A S T M L A G D V Q R A E E T K V S I R V K I N V A I C N N H K S A G K A L K T V A Q N
RANDOM_2545 I A T H R Q E W V A V L T I L M V H L Q A P R R S Y G E I A V P P K E P E L T P S T L G D E G A Y A V G E D Y F
RANDOM_2546 K G R A L Q V L P V T A L G L I I I A N Q Y W L S D I T V F V T T F E K K V N G A D E I V C R W V F P L I Q Y V
RANDOM_2547 H A S A S N V A S G S R Y V D Y R T E L E R Q M I R S P T C P P E A R V Q T C H T W G L K K S F I S Q N H S N S
RANDOM_2548 Q S K K Q K I L I K K Q P L T G L L Q M F G H P P A E L A F T F L V K S E I D K R V L L Y L H Y S G L Y E E S P
RANDOM_2549 T L N S R Q S N N G R Q A F G I D R T V A K R R K Q G D A T Q D L P V P A R N T K A V K M N E N K Y V T I Q P V
RANDOM_2550 A V T A A H M L M S I S N N A I E I G P K C L G L S T P M E M S T M P Y I K T T L N L P G I L G M A D T G G N A

RANDOM_2551 C K S L G K K M F R R N Y P T L D D R K S G G T V Q V L Y T H K R K A A S G Q T K S R G G V G S K S L A Y P V L
RANDOM_2552 L A L Q C L F R A N E T G L E T P W D F Y Q K R V P G M N E Q N S W I A W Q V R G S A R M P Y M A G C A L C G P
RANDOM_2553 D A L K V F H A I T L N A H L S G S F G R S S A D N Q T V F V C K M T E I I E E D A S I K E S A G E W T S L Q Y
RANDOM_2554 Y A A I L K P R Q N R L V P L S R I W D R P N K G I N A Y L S M A Q S P L C P K A H Q A E T D N K M T I F G A P
RANDOM_2555 I I R L N S E C E P W L A R I S P P T N L G I K K D S M P N L I E A A T G D L I V T A C L Q G V C S D E S E V P
RANDOM_2556 N E A G N I L P K G P T P I V H L A E S L S G C T D G G G D Q R W Y K V N A G H F S D V V D E N Y E L A A I T G
RANDOM_2557 A S E Y G T E N I R N V L A Y R I K E G F F A V V A V Q H H A T G G E S Y F E Y S L F D W S S E F F G E T R S V
RANDOM_2558 L R T K Q I F P S R G T F T I A V L G D S S K E S H G A Q K S P Q A T L A A N A G R F A I E K Y N K Y L E T K V
RANDOM_2559 G Q K E Y K A Y P Q A L C V L I G P L P A I C G E R M L K T L I G Q P R R E R A K A E V D I M K C A R P A E K D
RANDOM_2560 S A A Q D P V L S V I C K P P R V L E V R C F R S A K S G P G F G K G P A G I I F N Q P C S D L P E E G L V Y G
RANDOM_2561 K N D K A M R P P S G G E G C R I R I K R H Y S L V V N P S I N L K S G A D Q T G A A R Q V P L V A N E K L R Q
RANDOM_2562 R A L C A D E M H E G E T R T S R H G K D R L A S V N L D P S D G R F A S D I K C E T D L T D H Q E D T Q A V T
RANDOM_2563 Q L M L D M Y F T L G V L L S F P P P R K E V T A Y L L L K L P V K S D E I L S H Q F D E S V Y W E V V S S T K
RANDOM_2564 S M G A E P P M V T K G L W L A K V P N N A F L R L Y H E S G D G F T S S L A G A V L N L C C V G A N G S R N V
RANDOM_2565 K D A S D I R A K S K P N S S G D L L F S N S I L K S Q V V F R E D L F T E L I I N T P G V C D T D Q R L G F W
RANDOM_2566 I S K N E L K Y S A Y E M S P P Q G L N T E R W I T R I L E H S T R A K Y V Q K L N G R K S I D N D Y D H K S V
RANDOM_2567 H Q C D G S D I K G G F A G T A G P Q I I I E T G D T V A Q F Q Y G A G T Y A K Q V I K C L S T V L Q S Y T R A
RANDOM_2568 V S Q S E D R G L I S I C W E E D A F N S L D A G S M Q W V L G I R S D E A V P D L L S N D I K Q S V G I G L T
RANDOM_2569 G D G K I S R H T A L E A I L P S H E E E D D I S I D E R T N N N G L P S C S E Q G P S N A E D T T G G N Q V V
RANDOM_2570 T L Y P S S T M A I T A N I G T N Y K H D G N G M D F W A D L E S Y Q A P A N I E Q E V L L C K K D V L V Y K M
RANDOM_2571 S F P G L N R G H I P A K E T G F P A L S V T Q S A A M D R Y G D P N A S Y L G C M G R I V M D A S T D W S L L
RANDOM_2572 P R Y H T D L V L G T D Q I G C K V E F Q K K V F F A A S G Y P G G S E F V A T L T N M F I N Q Q C V T I V H E
RANDOM_2573 D L V G V K Y E N I V K L A L Q V F V S D E R K R Q S G A S Q T G G Y A L G W R S W A I R N Y T V R Y S T A L E
RANDOM_2574 L I G P I N K A H S T G T V P E T E V M S Y N Q G D M Q T A P G Y L A N T T L P I L S L G I L A A S N L V V L W
RANDOM_2575 Y H L Q M S Q L V V R S P R A Y R E L E S V D E P R L L Y D T G K S L I I Q L L E L K S A G S V I T Y S C H A V
RANDOM_2576 Q G P F L E V R K A L F P G L H Y P T I N M F K L L G L M V K K N S H K I C A E E Q L T S S I K S S D E A N D L
RANDOM_2577 F A G K V C E S R Q G Q T H G K Y Y A N P N I K A D R F A V T R G R D R V L P L M D N A I W D E S D Q D E D K S
RANDOM_2578 F A D A F N L G E K N E E Y N S Y V N T E E D E L I A Y G S R M E V H L V P A S T L V E P N G H T C T F L C H M
RANDOM_2579 L K D L A A A L I S D I C I K E P G K V T D P P A P V D G G G K M C V V S I K T K Q D Q T S F R Y G N K G Y V L
RANDOM_2580 A E I E Q P L P N Q A K Y P L R D T V I F Y D P R T A M A Q K C K V T L E T L A N T S L A D Y G G L T S I E K L
RANDOM_2581 Y V G R F D L E S D M T K C Q G K V Q A A L I R Q Y G L L E I S S R C C I P P V A T K D S L N S G S S V H L P R
RANDOM_2582 N S G L W E L R Q L A K M N N H P K D E C I G D Y E G E Y P L P P G F P F A M C N E T P P L V E D G E G N G P N
RANDOM_2583 K R C Q I S R Y K N S G G L G R N D I E E V I G L S P T A N A S Y T W N V K I R S P H I D W I R K E L F F G F Y
RANDOM_2584 Y D E P G G K F F Y S V Y Y H N W G A F L T D V Q A S R L G D G K N L V N T A F Y V P D H L M T A R Q D L I T Q
RANDOM_2585 I R L V K L D Q S G I V Y E S H S L Q S S F T T D A I Y A W D L E A G N V R S V Q D I Y G Q I R S I D S D P Q V
RANDOM_2586 N L R P S S V I K N F N I A A A E A H L K S Q I D M A A Q L K E E V Q N P T Q E G G E T K D A N M R I L A K G A
RANDOM_2587 V H G T R S V N G P N L L L V P I F T E W R A M G S E L G P Y V N L S P L A T D L I H L N N N T R D A E G G T L
RANDOM_2588 E R I D T S L A L V N T Y T T I R A P N P L R N S R G A Q K K A G S K M N G H K Y Y E L T L G G T I T K N K Q F
RANDOM_2589 A L D N R A I P A D E L E K A M G D Q P A A Q Y P S E I V Q V D H G Q S H S K A K D L S M G P F L D L G E S A L
RANDOM_2590 G S G F T T S L G E F W A E L I L G L G M V I T I L K D G L E P S S H D M R E N A F T R S N E D Q G W C Q L C F
RANDOM_2591 V L M A R C V Q F L K L A S D S R D F H C V Y D R P I E S S R V C A V I N Y L A R F R I Y I T E K L A P L I E M
RANDOM_2592 C E L R R T T D I A P T C R L V N M P S A I L L E L L T T M L D S E T N G T A L A L G S T E A R R I S S G L V V
RANDOM_2593 L Y I Q S S L K P G K A G L F R Q H P R W D P Y A P L I L V W V P P L E N S E G R E Y R D T P L K Y R E D V L H
RANDOM_2594 T D N H A S N Y G E I C I T P S P S T N P D T R F S I A Q S V S Q M V A G R K Y A N T V T E Q A N Y M L Y H E V
RANDOM_2595 M S R A E L A E L I E G V H D D M S G L A E A K D P L R T S E I H V R R S V Q I H K N S P R V H I S D L K G K A
RANDOM_2596 M A N K Y S H R R R P F N T P P D K R P V E I V Q A Y T T T W C P H L R D G D G S E A G G N A N M E R V G Q K F
RANDOM_2597 R D L C K M Q K N E G V Y I I I R C I V R E P E G H A L S G K M N G G L E N F T L N G V K E K I E G S Y I D E F
RANDOM_2598 K C S T T Y H A R K S S Y L L S D N L S N E S W D S R Y D S L E I G D L E V G A I H K S V S L T E S S A L A S M
RANDOM_2599 Q D S W L R A R E W Q I N E T A V D L E I A E D L T D E T P N Q R E G D F Q K Y N D K G E G H S A G N S R I A L
RANDOM_2600 V K L G N M A P G L Q R T L Q E L P D D Y F K K F K R K T R R A E D V P G H L Q D L V Q V K V F Q S E G V H A Q
RANDOM_2601 P C G G T R G P A G R W I V E A R V T V N I S K P P L P P F D E T K I F D R H T G V S P R H W D K P E A G Q T W
RANDOM_2602 G G I G A T F T D L L I T V G K L K S E T G L G A F L H R C K Y N N T E G I T F G Y A M G D S H P A L A A L E E
RANDOM_2603 R T I A S Q H L A V I E T G E E L T G V E T S G E E I D K V P T L S T K K R T P K G K V E S G D G V S F S L P I
RANDOM_2604 K I V F G K W K G R V T H T F Q K L L I R L D A A H P T A E S L Q E H T T N P T S F F I N V A D D D P F G V R I
RANDOM_2605 T L I W I K S T A R M R E A A R G Q R Y E I V Q I G E C P Y T H T S R Q G K L E L L G R I D C N L G V I R L L A
RANDOM_2606 G T S G P Y I L H K G T N D M G N N D V S V I L A A A L E I F S R D E T F L M A T L L M E V R G L R I L A S G A
RANDOM_2607 D A P L S L P L A E L V A S A V F A H E Q V I I M V S V L P N M A V V T K V F A A R G F S L N M W H A R E L I L
RANDOM_2608 V P Y P I R R E E G S S V D A A I G I T S K R K V S T Y F L L F A E L A S V N E D T C F F I T E G V F R A E L G
RANDOM_2609 A S S I L D D T L S I L C N L T E L L E G E E M H K A G G T E T Q K P Y L V V S Q T Y T L C L P N I M I S I A I
RANDOM_2610 N V H A I K A G C H P C L H V H H A V I S E T T T V D M N L G D E E N L F T S G K S L S S S R S V C G P G D G D
RANDOM_2611 C E N G R F E R K F F N L D F V T K V C L P T L Q V S G S S D K E P K A R L E S T K G F P P S C N L P V S G H Y
RANDOM_2612 L K L L L D R N R D I I L T R P H T V K D Q T L V E V L T I A D A G P G A L N E E L D L A L F R D R R P D K N H
RANDOM_2613 S A A Y P S P M A S L L N I I S K A S N R M S L L M K W S I G L P T K K N D T S F Q Y R S G L T I E E E T H K P
RANDOM_2614 R A V S Y P K E K D T V Q E K R V A Q G S T E C N V D P L K A M V I L I S R T F N N R N S P A R K A L A D E F M
RANDOM_2615 A Q L R L A W G V N F D V V L P D P R L I M E L S E L K N F I L A A F T R A L A Q L R K G V L V R I K G S T L S
RANDOM_2616 A M S K G I T P G K I T K N L E E E G P R V Q R S F H V S W R L E L G P N C E E T N L Q H T L R H P V A K D Q C
RANDOM_2617 L S L L D E M I V N E D Y R P M Q F G E G T L L S E G K W I S F D V W I I E K E S C Q F L D S V G S W K S V Q Y
RANDOM_2618 T T W T W A M G K L Y I S N H G W E A K A T K V R K D S I P D P G G G K T T M T G A I E S A A Q V G I A G S D G
RANDOM_2619 L S I P N C L Q Y N P P E L L L G L A I T V A V P A R P E G E V N F A Y M H D G S Q F Y D R D F F H T T V T I M
RANDOM_2620 M A L T I C C K K P A A L P A G G T Q N H A A T L A E K D K D T V V G Y K R L P K S L A L N Q N Y F R L G G L D
RANDOM_2621 M I L S N S S G Q W A C L D K L K K Q L A T L A S S V R D V L N S P H L V Y A T V Q Y Y V F M P N C S P M Y A S
RANDOM_2622 D R A Y L N K T A G C G M C V V E C P I A D N C S P T V M Q A T F Q H I V R P W W R S G I Q T P A G L I Y Q K I
RANDOM_2623 N T I A K P D C L F T G P S P I E E V M S R D M R R G T K N F L I R Y F S P N G E G M N H S L I E E E E A L G L
RANDOM_2624 G V R H D E T V K P P G R R A E L L S R E F L N A N V D V Y G S S A E K I S D K Q L Y F V I L R T L K F C H D C
RANDOM_2625 V G M G L K A H R K L I A D G S K Q S I G V S L L T P E T D I W P F M Q S Y D A K N R K L R V G E V N A A G S R
RANDOM_2626 E S T R R T V A A Q T R P M E V W A R I L L L G G K S M H A P I V R I P F H V F I R M N I L S K I A T V P L S L
RANDOM_2627 V D C M K L N L L N M F R V K W Q R E Y G T S V P S G R N S R G I G I N N Q E I I R G S D P L R G R P N N K G R
RANDOM_2628 P P E V N F T Y A E I L V G D P V G Q E Q I R T A F D N A Q L D L K N V L C I K P T G I S K K D E P A K F L K V
RANDOM_2629 C S I Y F P D Y I I A K L K C R D A I G V P G T G G A I I D K V R T F S L Q A T V G R K S T L Q A R E R K L N G
RANDOM_2630 G A N L E L S L A D D T R R Y P V D A T I N K F D S G D L D D L P M K D Q L G R E N G I R I L T F N W A K Q V R
RANDOM_2631 I M L A P R T T W P D A P N V S D I G R G I N L R P G A N D A V V Y N Q S K N V G S E N M L S C V G I Q N L L Q
RANDOM_2632 S F L M N I Q W E A V L S R V N N M C H T T A S V N Q G F S Y R A E P G Y L V P P L D V L I G H R Y C W I G R G
RANDOM_2633 S P A A Q V A V Q I Y T C F G G D R K E K T T G R D G E D D G S G Y L L D A P N I I D L E V V E K G A R I T G A
RANDOM_2634 S G E S D M F E W T N L G A S W G R E K R T D Y A V V Y G C E I V E D V V Q E H A S R R Q D S R E R R E I T Q T
RANDOM_2635 V N I E G C T R E A D L R K S S V E H S G Q Q E L A S L C V V L E R A E E N P T H L E R G A L L S K G H S E T P
RANDOM_2636 Q T M H D S A L S D S S V G W D N Q F A S K S D E I T F K D I N R R P V F P S G H E E P T S Q G Y E F M V R L L
RANDOM_2637 E D Q I F I I S D K A G E K D S Y T G E P K P T L H R D L D D F N S A P A T V D R S L A P R I S L I K V L T N R
RANDOM_2638 L A G Y Q A N T N L N L I H E E V N L A T Q S N E L S L T R S A S D T P P D C E L L F R N P D S A W I H L E G K
RANDOM_2639 P L C D S A Q I C V H I K E G I M Q S L E V T K V I I P N R T W H P F T P R L P T P N R T V V H G E I A A I D K
RANDOM_2640 A L R G L F S Y N F R F R G N D Q G A Y E Q V Q R R F A S A D A V N D K P P D E P L P I E Q L L G L A Q F N K G
RANDOM_2641 T M V N K L L T I L G S W N W L A P H T P G I N R G A E S V K K A L V I V E Y A L W N D S G N G E E E S D E F G
RANDOM_2642 L D L N L P V G T A L A K K G I L P V R I S Y T I A Y E L D G A S Y D K A L N K A T Y Q P F R M V K V K Q L Q Q
RANDOM_2643 P G S C W G Q E I T R L L W K A I D R T N Y K Y E L I T C H L F T T R H L K F P P A F A S W L T T V S G H R G I
RANDOM_2644 R D E S H Q D N R D E D L V G S S T R P L T D S L L L S L L A G S A D G A L Y K S E R V E P G V C D L Y S W V I
RANDOM_2645 K M N K L N C S K T I Y G A A D G K N E R S A T E A F F G D N V P G R A Q M L T S V I E T F P V F N M V E D L T

RANDOM_2646 L P P C Q I R A P I L N G E P A D T Q V C K A R W Q V E A F E V R K K W E N A S Q V D V I G T S C V L Q L V R L
RANDOM_2647 P G K Y P A N K L I A R A S L E A R P D I T E R V A K Q A D W L E A Q G G L N L Y P G K P R R C E E K V I G E I
RANDOM_2648 S L L E S K A L S P E N T P V G E N Y W Q E S A S V F G F V D N G Y S G G R L A N S L K S L Q I A L M V S N R L
RANDOM_2649 D T V A D L A F A S E Q L S L S I A E D D K I P S P G P E D L K S Q D A R S E N W A T M L L A I R K T R F S S D
RANDOM_2650 G G I K T G T E E E P R W M S G D C H L K K W S S N H Q G K C R S D I T Q D P T E F L L T Y L T L L L N W Y F N
RANDOM_2651 V F L S N N W T V W F Q Q L I G E P D R G T K R A L D D P F V I N T V Y G G E R D A T Y V H T L P V S E G D Q R
RANDOM_2652 E Y K M S P R V G G N S Y G E N M E R A S Q M A K G E S L M S L A N I Q D K T V L Q L V I T Q T G G S E S I S S
RANDOM_2653 G E P D I H G S I D G V H K S H P Q T G N S H E A R C L V D G A A E G L V Y G K Q E S V F P G A G F S S L G N S
RANDOM_2654 M D C A P L V D W I V F P V Y T V N V G Q N Q R Q S T D V V S A S W N E F L G P S V G L M D K Q P L S S L F S N
RANDOM_2655 T Q V M P V L I E Y G R A R K G I R M V E R V E T E C C P P E A S T L T L L G E K S L N L T M R Q N T A K T F R
RANDOM_2656 G K I F I D M Q A M P L D P K V L Q S T F L L E P N I D S R P E T G E T L G M S P A E T V Y S L L V G S N D T S
RANDOM_2657 F S R R C I A L T S R A D A E E E I V V A V D N Y G V P C V R P L H I G L L G S K E T R F I N D G V A V S V V L
RANDOM_2658 S F C Y R V P L H E S D V N L W G G A I D G T H E V T N T V S L T R L L N S L M S V L G T L A H P Q A R G R Y I
RANDOM_2659 D L V E N S F I L A R G E I F M W M G K E I D Q I K N S V E N D M H M Q T E R R F T G S A Y R N G N N R G V R E
RANDOM_2660 P N P T F G D G Q L A C M N L T P K K G T P G S P L E H S L S P G D F V W P D T A G L L R W H C L Q L V F G L K
RANDOM_2661 K N A E N Q L A G N L N R T V N R L L P E E E A A I L V K G E R F V T K F E V S T L A Q P L F I Q A T R Y Y P R
RANDOM_2662 M V V R K I V G L A E A K G Y S F G Y M P Q L E S K S L I V D P P D L G S M Y A H P H C G K E V F K R L V I W G
RANDOM_2663 T W T E T P K Y S K I S T I S E A S D Y P Q T D L Y L R V D G I I S V G R Q F L T K L F D A K I F D S A N N T A
RANDOM_2664 N Q V I L I D R I Y Q Y T D E Y K C K T K P T D S N D L Y A Q R G I I F D M N L A S V L K P L R L A I V Y V F I
RANDOM_2665 S N I L G Q R C L I F L V Q V T V I G L K N E A G C G V A K A N A T V E K E S V N F L D L F I L I K M V S F P M
RANDOM_2666 N T G D F M K R A K R L L K P V D L M L H I N I M R D Y G R F S N L K L F P S V E L G R E V N H I H G K G V F K
RANDOM_2667 L F I S E G Q G T R I E W I Q E D H A V F K N G G S I M S V V D V S D D H Q Q Y R R E W I V R T C A T L R E R F
RANDOM_2668 G E V L Y L N G V S M R C V N G N G I P A K D L Q G T V S K F L K K N M K L Q D D L A S E K L P P F K Y T L D K
RANDOM_2669 S S F L A H L G D E H A L M L S K E G L V Y P Y I S P G H K T C S Q E M K V V A D K I L L V K G L L R S S C H A
RANDOM_2670 R F V I G S P L E A T E A T Q S E Q A I G S K E N G Q Y H L K G A S L T G I I N F L F C E V T H A W A W N E Y E
RANDOM_2671 S L S L A M V G N E V V K D E R S L D A A W R K R E A Q S P K D E L R K L E M E H V S I F Y D F S F C V G Q Q T
RANDOM_2672 P Q D L I F K K I G R K P V L L R D Q R H T G I G D P F Y G L H G S K K G R Y F G D T S A E D Y I V E I S R R D
RANDOM_2673 V L Y H L L R T S H L G P T P R P K N Y L A S H N M Q G K T Q G F N V V V I A T Y N P Q P I E K A S R E V N L Y
RANDOM_2674 S S G E F T W T V C A R Q I T R L G R P G G N L N L N C I L S N R G I E P V W G S D F P K E V I N K H G E S G T
RANDOM_2675 I V S T F Y D P S P K L L G P A G I H M V G S P A K D S D K V V L P V C S V K I N K Y Q L E R E A Y H T R K T D
RANDOM_2676 K L A K G N N F N N I L T A I L N A N Q M G M D I L G V G T Q F A F K Y N E P E L V F E C L G K I A A G D P K G
RANDOM_2677 A L G D A R C I R G Q R F F S E A A A K N N R N A L A F G S D N T Q A I G P W P T I F I R S G G Q F V L P Q G T
RANDOM_2678 E V K V L M A K D T A L E K K M D S D I G T M R S H F R T D I E V N S F G D T L D L L I F E R D R F D A D E E C
RANDOM_2679 H I K Q D M A P G D I D Y K L E D I Q R T G M L I R E Q N R E V G P R F E Q D N E N L M K V W L P A V A R R G G
RANDOM_2680 E T S T K I K R K F E G E S K M R D L R G S E S N D G F S G Q G D R A A N R G E C V G R E V L W S G G Q K V A V
RANDOM_2681 I P K E M Q G H A D R S D E C A G P Q Q Q H V S V P T H L N G A G A R S M S V T G K P V L A E I D G A L I H G G
RANDOM_2682 D Q H G L L N L D K L P I R I D L V S Q E D W A E K A R G D L N C P N L G S K L N A H P G A S E G L Y G T G L L
RANDOM_2683 P T A G G E N M E G P T L I P G C V V D K E Q G L H S Q G A I R Q P K L D T K Y N D R G A A Y T L R K G V D D S
RANDOM_2684 E D I V N N A L N G G M I T F V A L E R M P A P S A D N R R V L S A T A K H Q G T Y D T T T D G A L W G E G S L
RANDOM_2685 T H T F A Q S S V G P D M R Q Q D A Q D I A L S Q N V F E K E L E G N I E N I P Q A G L S M Y V R V K D E E F S
RANDOM_2686 P K F L V A A P T Y L P P V R E V L N S F H K I D E A S V A C V R N L V E L R M E L V R N G T Q K I C K G V R I
RANDOM_2687 Q L N V E Y T L L K I Y A R S T F S N A T K P A N Q V E T C L A Y P D N C I N S E Q N S Q I A N S G V D G G D G
RANDOM_2688 H F Q A T G L G V A N Q T Y F S P G T A F I T R P T I K N F M S F Q F C D D G L E Q C F A L E R E S N L E E T Y
RANDOM_2689 Q I M V E S K M S H R Q P D N N E A T E L L H R S S Y L E S G N H L R G A M S M R A H A N A P T G S Y R N Y V G
RANDOM_2690 V K Y T C I C T V V G Q A R G N L S V V G P K S M C L R Y R T I V A Q C T K V L S G V P N D N N S G K F D G D G
RANDOM_2691 P M A N I S N S S E S F A S G F E R I E N P K G V P N T I Q L C A R H A E L L T R H V Q E V D A Q A T N R K P F
RANDOM_2692 W V Y G V V K H D Q S F R P V K T D T L W F D Y C F G Q V D L L G A C R Q Q P D T L L Q K I Q A L V K S V I V V
RANDOM_2693 G D A H D L T T Q R F I G G P D P I S P E G R V N I A G N W I N L G T E Y Y M V G F C A C A R E E F L D F N N C
RANDOM_2694 L G T H H Q A Q E Y S E K Q G F S G E G V T M P R A I A C I W A E K Y S S K Q R L S L V N F I D I E R R S H G F
RANDOM_2695 H L F L F N D K K L S V A S M F Y N Y P E I N V V Y T Y V F S F V S A S K A K Q A R S Q K A T V D L N T S E N E
RANDOM_2696 P A R Q A E H F A P L G L A G G E N R T W N A T V K I R V K I M W Q Q L S L T R P L C I A I P P V W T F T S T S
RANDOM_2697 S R T A K I I W Y N G H P G P G D T V E G E E S V M T T R F S K C D A H F S D N F C K K A P M V F R T N A S L I
RANDOM_2698 N N E A I K L I E T G T I W T S S N G L I D K L K S V S V E P M L P S D L T G Y S V A G K R D K E N H N I T F I
RANDOM_2699 E P Q E V V V S G V Y N K P M L M Y D N L Q P G E T P V Y H F G E N F L A G S F R K D D K E F A Q V W A K K I V
RANDOM_2700 H C K D C G T E S P C Q I T P V A E Y E R S F N T Y Q P G S D V T S T L A A T K I K C R S L S D V Y P T A K Y F
RANDOM_2701 K T A G E R R T V G I P S V T N N Q K V V K T N K I Y D R R N A S S L Q K K E V G S D T Y K H I M I K Y L E H S
RANDOM_2702 Y A V V V L K G E T G L A R V G A A P Q T L R I G V L V D I T N T I V S I S S E Y E R V H L Y G N E D Y N D A V
RANDOM_2703 M Y L M K E A C V G L F P F L Q S A M V H N D N N C S N G L A P P C F N T L T K G M C S V G G T A L P T P D L R
RANDOM_2704 R N L Y T F K G Y S D P H L V F V L A A K G S F Q R A Q M S L Y E G L T N Q L I V S S A P A T S L T T G D V R E
RANDOM_2705 V V E M N A S W Q Q L V V V E N L V I I E S L N Q D D A T T K V G A E K D G N V P E M N G A G Y Y G G V F V T N
RANDOM_2706 G T P S T A A L R L V I G S F A Y S D L A I R D Q M K K D C R S L N N N M L I L E S I A F K P I A R L T T A Q L
RANDOM_2707 A K K G P K S D L S Y R N K G E D C Y A D W Q L K T E Y G N L K L S I N Y I G W F T E K E V T L V Q S K K V S V
RANDOM_2708 T Y H P G R A N C E V I I G H A G T Q P D Q G E R N N A S Y S D S T P L A E Q L I V C D P I F L A Y T E D D N A
RANDOM_2709 T G I G L H R G C S T G S Q T V F N L Q Y C F R F R G F Y N R K E Q R H G E P K D Q L R P T T T L G K M A K H Q
RANDOM_2710 K D Q A S G N D L I V A T P L K A E S Q V Y N R E A A E L E P T G V L E Q S E Q V I Q P C S L A K L R T K G A S
RANDOM_2711 K L Y I D Q S K T V F N E G V P T P P G K T E T K L A P L V G N F E K E K R F N A A E H S M D S V L D Q S K A I
RANDOM_2712 Q Y P V F P V I P M D G A R P H Q S A I S N M K L D R D L D D A P G T A C V D A Q V G A G F Q I A S I S I T V I
RANDOM_2713 T G S L R S G L I D V R Q P T E G S P A A K V K A R E A R S A L L K A M W D L K G K E N T A N P F Y V L R C P D
RANDOM_2714 H F R D D H H W I S L G A L E N N V Y N L G R P V Y P L A V F G R I E G K P P S M G T T E P D F G I Q I A S G R
RANDOM_2715 S G G P Y V Q V G L A L Q D N K S K E S N E R G R V Y Q A V A G L P R M Y V P H H D F L A A Q L E M I F V I N L
RANDOM_2716 G M G T V Q K N S L R S R C S D I I A V A Y L D A R D E I L I S I D V H L E V V D G S F T A P L V A M A L N F L
RANDOM_2717 E W N Q L L D P L P E T M L S P D S A E I T L A T R T Q N G K I Q V K L R G E P L G D E A V E T T I L A S A N D
RANDOM_2718 V R S T K V H W Q Q L I P T Y T Q H I V N S R I G K I K Q S P P Y R N I N A A P G P F A N R L G K T T N S L L A
RANDOM_2719 F R T V C C I K Q G D A M I M H E E V H P G S E G A R T I I K P G N D V V V I E E M A I D W T G V V G E K D C D
RANDOM_2720 I L A I P S N Q D P M Y A S P N Y I Y S Y I W S S V N H S M V S I E Q D S F T F D V R P A W Q S E N N D V L S L
RANDOM_2721 C V D Q T Q V V A F Y L L K S N R A N Q I V M A V Y G I S F D R V S F G Y S T W T G R E E P A G S I Q K D L D K
RANDOM_2722 N R R N T A L D N V V K S D R E H E L F M L E E D A N K T Q E L A L A P G A Y Q H V Y A A N A D Q V V P K V V I
RANDOM_2723 N A G A M L I Y S S V G L L F N A A N V V R K V P P L S Q I K G R T P P A V P V D R N I R D D V N G V S T Q G T
RANDOM_2724 L T C E A P F L E I E V I S V H Q C A P E L A S S T V Q N G K A R S L S A R G M L T L F E P V N T R M D F V V P
RANDOM_2725 G G G A V L K Q F C V K H K V Q E R E T G S N D V E F I F P P S L Q V K G T R C L G A P D N V M L G R C D R E I
RANDOM_2726 Y S C D E T E D V Y P V Y F L L A N I R P E Q E A F E K Y A G T N W D S R A L V C C A C V E T P L Q L N R L R G
RANDOM_2727 W E L L S G N A T Q N P F E R R T I F R A Q L R N S F L V C G R A R V N I F T W L A K V L F V G H V A Y A A R E
RANDOM_2728 S S T Q K L M M I G V D G S V M S A R W T K G A G Y V A I L R M V N Y R A A S M N Y A F G M K M L S P I F F R L
RANDOM_2729 C N P V L S V A S T Y L A H V Y E V S V P E Y V V E S L N D S P R D I V S Y K V K E L Y V E T H E L H C R P V F
RANDOM_2730 R Y R A V P N A F G N N Y A A G P S L A G T Y G Y I T S S D V S Q S F I T G T T R P V A D G S D T W A E Q P E D
RANDOM_2731 K T Y E A A R I E L I F K K E T T D Y D G F T G R I N G L K I R S A S S T V L Y K G A P A C S S L P D I L S W S
RANDOM_2732 N E D G C E T V I L L L L E V T D V S Q G L S C W H Q G W Q E G R K A R L T S L K E S G C L A R L G V V G G A C
RANDOM_2733 T Q R D R L Y V T P G V R N V A A A L L A G S K F S F V L W T V T I R T P P G P S T L L Q S A L G C R G E N E S
RANDOM_2734 S V D F L D D V A L K E Q L A L V L A R V G Q S T M W I A W G V F S K Y F P G P Q P V S G Q Y R M A M I S N P E
RANDOM_2735 G L M G K K Q H F P L N I E S G V L K I S E S H H I R H S T L T A Q A L S D L L E C D S G F K E S S G C K D A E
RANDOM_2736 Y D Y V A A L K S H L I H D A G G A L S L K D C A V E T I P R L P L A Y E P T W L A D K N M E P V N F S G V A Q
RANDOM_2737 S Y R I S F A I P I V S Q C D V L F R Q I G R K L A R A G A T S M K W R V G R S E A S L E W S E T L E P P G A R
RANDOM_2738 K T F G Y P S A A A K I S Y N E K I T G T S I E G Q V T K I L I A E A Q W A A T Y L Y T W G V A P A D A E Y A A
RANDOM_2739 A A S S H R G T F F L L V K A S P K L K S M A N T T L D Q D R D T P M D H E G G G L A N F A G D Q L T A Q F S R
RANDOM_2740 P T E P R E P K G F S A L K E L S A G P P S Q Q Q V S V L S C T Q L T T M E D Y C A S G T F P V F F K G L K V C

RANDOM_2741 I Q G I I S A G N T R I Y C H V A G T G A Y T A K D G M Y D N V S F C A A C P K D Q Y R A G M E V L I M V R G V
RANDOM_2742 K L I R G S G K D E I E F G K N D A G G Y D K G P T E V C W A I P R S V P S T T Y N D Y V Q Q I L D E F T E Y T
RANDOM_2743 S V Y K Q E I A D D T G A T D L L Y R S A E E H P G M L Y P F G V N Q D Q V D F G P T G V C T A I G A T F Q R Q
RANDOM_2744 G I K Q I I V D N P A E A L P L Y L M Q Y G D I P S D E Q W P P Y C L A R K L M K P P P D T K A P G S E N L L V
RANDOM_2745 E F K A G I R Q V S L E G R A F L L R Q P K Y D T A D M Y L V V L G S V S T L K S G Y P V A T K F A V E A Q C V
RANDOM_2746 D K W K Y S E Q H V K Q D G M Y S G C G A R V K R L A L A F F P K A K S D R F F E D A E V Q S S V G G E S R S N
RANDOM_2747 D D L E P F R M R G E K F A F Y P T Y L K T S A G S Q T L A G D S I G G P Y H A N T A A E V L E S E A G L E K E
RANDOM_2748 G A F K T M T A L L H L G A I W E H G R G A S V Y T Q D E D I L L C N H P F V N N L R Q I I Y R E E D S L M I E
RANDOM_2749 L S S G I T V Q E V Y A T I P A S S T S K E I L L K V I F A E W T A N M N V E W Y N D L P D L Y A A R A R F T S
RANDOM_2750 E K S H Q K L I R M L H Q F D N C P E M S A V P N T S A Q K L R I S T G D L H L V Q D G L M I Q A D Y G T E C G
RANDOM_2751 A F D G Q Y K A D V A T M P R G V Q F A M S E P I V Q S T G N L P R M H V G V T P L S N V G K R P E Y K L I T W
RANDOM_2752 N N V P A K A E L D S Y A T T S L L I I N G G P A I V K Y N N G S M L S V S Q N A K I F D Q D R H A C G V G A G
RANDOM_2753 E Q Y P G L L P Y A R M K D G S S T E K I N K F C V T M K V P D A I D T A S V K G L E T C N S P W C K V T N Q G
RANDOM_2754 G Y Q G E A L Q C D A L Y N L G Q L E E Q N A Y V D I T R A S L D A N S K F D Y Q S P K F F W I R I T G Y V A H
RANDOM_2755 K A V L A A V N A I T G I T E S N T G S L T R D T K H S L S T H S T P R E V K H L Q Q S I C L A S Y F Y R E V I
RANDOM_2756 G L Y F L Q M M F L L I A L Q T L G L I S P A F I R D V R K V R G F H I V T G V L P N A R I C G R W V L S E P Y
RANDOM_2757 K T W T A V R R N T E N G A P I K E A E E A D A A Q I K P A L K N I K V V D L L A A R A H D H E D G T H K P F F
RANDOM_2758 K M G V D L A Q S A I V E E D E L L K V T V N D Q G H E W G M T Q S E N E G Y Q G E S N Y A P N K G D G N N K I
RANDOM_2759 T D N A R N M K K K V T N T L F M L K G D E M Q E V Y S C L T D Q E L P Q V L S E L A I F A S P K A R I L T G E
RANDOM_2760 I D S G D N I P S C P G S N L V I T K L A V G W R T Y D A G G P Q M E Q S A S D A S A F L I P R E K L I G K A A
RANDOM_2761 F E Q Y A N D S R G L W A T S A A E T M P W L A L R L D R I E A M K F K N G G T T N S E N L K D E T S E R V D R
RANDOM_2762 F G E G Y R D Y C Y W Y E I A Y L R A Q S H T H E D T I K L R Q T S V V H I P P R R S G C L P A E G L Y Q G A S
RANDOM_2763 V D I G L R P M N I K I A A I Q P A V D G A A T S K Q M V N V A P V N S G T H G V V S A E M P R R K G V V Q D H
RANDOM_2764 T F T M V D G A H R L N L K H A K W Q S S Q Y L K E P P L R K F R T V H Q T F A Y L L K G C M V L E V A K P R L
RANDOM_2765 W T D G L L A E M F I V C Y W I Q S N S I S R A L N Q F Y R A T T L L Q V Q V D E T R E K N G G D T V T L Y L T
RANDOM_2766 I L Y E A V E G L I F G L G L E G A N T R I V L A V Q G T K T T E G G E R R G L L L G G S K G Y G P P A G T L G
RANDOM_2767 D K S P T P V L K D D A L K H I L E Y G L L V E N Q F S A S P R K F S K A T S Y I C E E K Y H H V K M C A S Y S
RANDOM_2768 Q R T F P T V P H F F L L F M V V N V E I Q T I C S N V S G L F V V N D V L L S S L L D Q V I L T A T F E A K A
RANDOM_2769 T V E D P K E A A I T P T R R I A V N D K A S G A Q F F Q L V P I D K C L F R S C G R E F Y Q A K V D N M C L G
RANDOM_2770 T D N E S P H P Q L D R A V C R Q K A P L Y R R V P Q V A Q T S A A K R A N S A T E Y T I G E L E T V T P G N P
RANDOM_2771 D V M E P L I L E L G I V L N E Q I S T A A S G L D L P N K F S E D T E Y D G T T C G L G V A V G W Y S L S L K
RANDOM_2772 P D Q R L R P D V V Q K T A T H N G Q A S E M N M T L L V T D S G P K R N K I T L V R T S E K E N A Q T K R I N
RANDOM_2773 L E I Y W D R Y W I C L N I V E C K I M L K Q E F E M A P S K N G I F I A G G P E G P P P S K G H L K F V V F F
RANDOM_2774 S N L P N R R P T G V R N S T K S Q V A F S L E L H A K V S N S P E E T A T S N T G L T W E L G N K F E D I G F
RANDOM_2775 K Q Q V I P R E K T V G S W L Y A W A G K V L K C L G I Y L T R D V T T R H L R Y S G S T Q T R V E S L N N N G
RANDOM_2776 F T T G P E V P S C G E T N R S G Q Q Y V W D G E L D S E L V S S S I F E Y L E A L E E L N L A N A V K D A P L
RANDOM_2777 Y F G P E G P K L G G D A G Y T Y F Y G R P E E Y G T A Y T N R V T I A P R Y G G N L K N H I L S P G Q L T E L
RANDOM_2778 A Y P F E A Q C T P G I L L I D R A W P Y S T A S Y T L E W W I N V D E I V S A S Y A P F K G K N E A G I G S G
RANDOM_2779 W S S S A G Q A M Q F K F S G Q C G I K P N N D G K T A I W Y M R V K E N R T N K C W R D Q L Q T N A T Q P N F
RANDOM_2780 L H L F S L G T R L N N Y A H K N G A F I T G V G A A G C E F N H N T N L T E S R G N R L V H M P D D K K A L D
RANDOM_2781 N K N W N T L M K P D G G Q A W P D W S M S G S I Y D I A T G T W A A E L S W G L D I K I G R L V E T M W Y L T
RANDOM_2782 R Y P V T M G D K K D D M W L Y L E Q S C M E I W N R K R A S Y L E T F D R Q A S I K I I S D L N D F K A K V D
RANDOM_2783 S L S S A L S Q S L A D E A S L N H F K S S S A S P R P V L M G G R K F E F N E A T M P D V A W H F N N F S Q A
RANDOM_2784 G R K A Q N M A L G I S I H C P E Q D P S A L G L G Q H G G I R A Q V G P S E E A L G Q P L A Q K Q L S K D A R
RANDOM_2785 G F E V V E S R I L P S Q L L T K H S K T Y G Y V A S D T N P P C I R F S A A F D K E S D V P P Q E V I D Q I V
RANDOM_2786 T I Q E P S E D P N L L T T K L L K V L I A D T N E H E F I S H D V A F P I C R K A P E D S L S V G A R N Q S V
RANDOM_2787 G L D E T G R T P H E D F A G Y P K N S V G Q G P L T D R D K C C A I M I A V I E V H N G R F Q G A R I A T T D
RANDOM_2788 A V I P L Y S V F E Y N V P E L L Q Q G V D S R D Q E S C L L C N R T L A R P F R S F D E I Y E A R H G D L G G
RANDOM_2789 R N S Q C L L G T P Q H G T P L R S G D L M S Q I L L P S G C I R H S R A V W Y P F A E T I S I L F R T Q H S H
RANDOM_2790 K Q I T Y P R I G T F P A M D V F D H G F N G R M G G L L F Q I V V A E R L G K H L A G E K E E R D G V T I D A
RANDOM_2791 R F F N I G R F M W Y S Y I T V A V K R G T A V F Q N F A E D L Y G C N R Q G K V F S V R L I D R P I F T T Q Y
RANDOM_2792 I P N A D P N S P Q H V H W H K S D G T V P L S P T Q A K L S Q K G A A E A R Y P H F G L L T A I Y S D G I T D
RANDOM_2793 L T S K A L I I R N R Q S S Q A G P Y S V L Q Y R D Y D L S L R G A M K V L L C T Y L S G L V K L A R C R A L T
RANDOM_2794 E E E D I D A T K D G A G E V A I T G Y S W V N A N C E I Y A L S K V G M T G E G T M F I A G L M E N E L H V V
RANDOM_2795 S V W D T L Q E G G L G N F P D L K P G K G E S I A A E N V E P Y E G Q F N Y R A L P G C D A T S G V P R Q E Q
RANDOM_2796 L A N Q H I K A F Q K T T Q T C Q L D F A P V A S W C R W G T Y W C C H C G N P V Y R M A P T L E E S M Q V W I
RANDOM_2797 D R L S Q A E G P E K A C Q Q E A I I G L L V V R P P A D G N Q A V T L T A G W C F E Y K V T P K N Q F R D V S
RANDOM_2798 I G Q P V R V Y L I N Q A Y T S G L Y Y K I K F I A G T A A P Y A N H T H L V P A C D M S H Y Y T F Q W M G R C
RANDOM_2799 T K M S T V A N R D G E L N I T I E N E G D P I R A Y Y D G C R L S A L W L G E A H L C A G M L L G G T V D L K
RANDOM_2800 L I V A L A R N E G R Q W Q Y D E C E V Q K C R P R P H F V Q V I P H L E R S R Q W R S R K C S D L M L H F R A
RANDOM_2801 G K T K F E L A P L E K D A G H E Y I G F M F S V E I T P L L E K A D G E S E F Y V A H Y K T G L V V R Q D V D
RANDOM_2802 N G K E R V D H H K E F I R D R L G P D T S A N E F G L S N L W T S H G A I P I P S L Q I K E L S S M G N L S W
RANDOM_2803 W N A E E Q Q Y C K I L K Q D D L A R L F S Q Q Q H N T Y K K E L G L V I S C N L T F V P Y G D P H E E N N P A
RANDOM_2804 R R M T H Q N G T Y D Y R E T V T A K G S H E T G K R I G E Q R R L G R H V M R G S N P D R Y P K T L A V A Q W
RANDOM_2805 L D V V P S H D S D F K G E T P D F N Y R S R R K K R I R K L R L V A E N N V A S S F R A Q E A K A S G T W I K
RANDOM_2806 L P P H Q N N A G I G P H N L G S L G Q M L D T M A L C P A M P S P N G A L N V P T I Q S S C W D A I T V Q W F
RANDOM_2807 G P N G L V T N S A M P K D I D N E A F K E A F S L T K D S G Q Y Y G I C P R S T G K S K F Q A R E Y K E V F L
RANDOM_2808 D S A S T D D H E K Y R P Q F G G C A G I R A I A L Y S G S F N L Y H R I D A E T K L M I A P S L T G S F L Y K
RANDOM_2809 I W R G C S V H E G S R R E I S N D T T L V G G W V Q I V R S Y L G L H K G P Q G G A V H H G C K V Q F T N E V
RANDOM_2810 A N A I A L L R A K E I G G F A R I L T Y G N L G A V A L V T T S T M Q S L I P F A D S G K D A A S K T R N S H
RANDOM_2811 E H L Q F V F K E S I A G P G R I A Y T Q R R A S F D N P L D S L V S E N G L Y P F T I V T E F D S T L L L A V
RANDOM_2812 A N H H S Y T I S L I I F R T R T N P A P G T I D I A R I N G L Q K I I S S I G P K V L P C N S S K P F F Q I F
RANDOM_2813 E Y P T E Y R E F D D F L P E T E F A P C A K D I K F P I G E P P L T A G I A E V S S F T P V R H Y T C F T T G
RANDOM_2814 I Y R R L L S E A A V D F G I Y N Y S S L F A G I M T S A R T V P V V R Q E Q K H S A F S E G L W Y G A V T V A
RANDOM_2815 Q T A C L F D T A R D V H A R K I W A Q P V T G G N Y N K M L Q A L E K H E T S L K F P G C V T A A G A C N L R
RANDOM_2816 P S Y P Q D T S T T T S F K G Q E S F V F N G C N S I Y S I I L F E I A F V A G S Y P R L P A V V T C P G T Y N
RANDOM_2817 G C D P T N E K A S A G I E F R T Y T L A K N L L D P Y N I A P L F N G E A L S T L V R T G Y A W P K A A I A K
RANDOM_2818 S D N S I L D R M W Y G L D G W F Q R V A Q I L R L A A P K L E D C L H Y D S N Y G T F A D L G K D Q W F I Y L
RANDOM_2819 S P E K N R A V E H V A G Y E I C R D F D R N P L E W A E Y M S L I P L P Y Y H Y Q H T A L T Y I R W M Y D F K
RANDOM_2820 R R Q L E V A G M S L C S P Y D L L Q S N H S F V M Y K I S V S F L V F E H P L L E S P Q R A L Q G E Y C Y E C
RANDOM_2821 A R F L E I M D F A S P L P V T T Y H Q A Y V M S T A A S R C L E V L L L R R E G G A P A G S N G G E R H V V L
RANDOM_2822 I D W N P L L I T A S Y E R Y I V N R H R F A I T I G G T S A H L M N K N K V A G V Q P P G Q V Q K N F F R E A
RANDOM_2823 I A R T Q S E T R N K E T L D G E S E T V N R K N L L R V A K P G A V I Q K L M F S G I V V I P S L L L V I K S
RANDOM_2824 R E M Y Y L Q P E P L Y P I K L S V R F L S V R Q V V W G V A L K Q P Q V A Y Y R D A P E N Y S A R D Y I Q R K
RANDOM_2825 H N M R N E H G G A Y L S Y G L G S K A H P N Q N L W K L S Q G V T K L N D L S M K I G P S E E D Q A N G Q T K
RANDOM_2826 S Q P V A L I L S P M G T E N A N G V Q K L I G L G V K W A S Y E E Q Y N T K L T V E N L Y K M A R T C A V I S
RANDOM_2827 F G M I F F A G V N K E T F V I D D F Q N Q V D A F F W T I M H D Q Q L Y Y P K N V W F F L P G N Y G A A N P G
RANDOM_2828 M A E A G G F V V L G P V I S I S D R Q R T S G V S Y A K G S G V K K D E L L L N G A R Q V G E I F E K Q R K G
RANDOM_2829 L D K F F W P L E I N E K S E N L R E Y L E W I I A T M F P K Y F H P Q F M D M P L Y Q T A S N L S T S V Y N M
RANDOM_2830 S F K A N L T L L V I C G G P K L G R L H K Q S L R S V S R S S A Q R P D K T P H S V K L E K V T Q E A K L S L
RANDOM_2831 E L L A A E N P T V M N I D V G S L G V D M S M L Q L L S Y Q A Q K H Y L I I Y G V D T L D P T M A T A V K F I
RANDOM_2832 E K N V L N E N L R T K H Q I R V G R Y H D T Q K L T I K G A Y I R R S R A H N N A V H K P G E P P A T T P Q L
RANDOM_2833 Y V A S L D W D S D R V C P G P A F R G I D Q W I S R L K Q K A A G T R P T F N Q I M N H S V Y G P T M L P A T
RANDOM_2834 E W G Q L N S Q K R Q A A D L A E R G I Q L D D W S G K I L V Q L S K V D G Q M V V I P D G V A P G P Q G A Q A
RANDOM_2835 A D R G V L A R E S M P G L R W L R L T T K K A S G L K V G T Y K Q R D T S A G D A G F R L S T G M K S I A D A

RANDOM_2836 R I Q M K K M T F L N H K S V T Q I H F V D V Q M R W V L Q S D P L V L V A E L A L E R E P Y E S A P R S Q R N
RANDOM_2837 A P V E C R A D Q E L N E M S S G F T G I S D D V L L F L V K G T L A Y W S T K N A I E P E W I G M V S V V S N
RANDOM_2838 L E Q M D G L L K D H P D L I P E A A I K A N R Y L T H G I T P L P K D C G F G L N W F S H P F P I P L L L Y P
RANDOM_2839 S N P S E A L L R R T E E L L E F Q S T H T A I S S D Y E Y S N T S E Q K T Q D P I N S R S L D A T Y S G A T E
RANDOM_2840 Y R R F D A A G A E V S F L G F P S E H I V G G D V C K E F P V I N T V A A F Q L R L G H L G L G E P V L L Q A
RANDOM_2841 L T I V G S T R S I L A G R W F L A V S T W C N D W N K D Y F A K E L K R V K W I D G E C T L S N Y L C L K S A
RANDOM_2842 L Y C E V N R K T S S E R H D Y N G I P A D G N L A T G D A I T E Q I K P F K V V R P H E A L W G K P R V E A E
RANDOM_2843 P L A R T P L D L E A D P E A S V T F A G R S N V V H Q G S I N R E T K N S G R P A I Y D Y Q F G A P L G T E V
RANDOM_2844 A S V L E R G L E M D Y T A K A T N P I K L T F I P L V P D I F I T G L V L F K W T L K L L S Q G N C F T K C K
RANDOM_2845 P D V G S A K N I D K L I A S N L P E A R L Y I L L A T V L C D T A Y F D G A Q D G G F N I M D S L Y T R T T D
RANDOM_2846 Q P G F R V G V V V S I D R K D R N G E G Q Q Q P L L Q S E E S E S W V Y K R E T S L F M S V R Y C A E Q V P K
RANDOM_2847 S P S L D L W F V A N S D A A S K I L A E V P T D Y T I F G A M Q L S R L K S K A A E W I D R C I S I Y G Y Q W
RANDOM_2848 S A P F E K C A S G A H L E Y T S D D D Y S M C S N R L G D T L S L S R Q V V T D L S H A H Q E P D N Q T Q H I
RANDOM_2849 S N P L P F R A L K A L S S G G C A G V F Q A V L S H P F L Y S S N N Q V T Q N P I S M N E L S V H K N T E W H
RANDOM_2850 E R T P Q P Q P I G Q E E K F S P L Y P K Q R L S Q D I H M L M A D E A S V Y P W R K L M A T S K M K K E T P S
RANDOM_2851 I S A P V L K A S V P I P W V H A I G K L A Q R R I H H D L A R H Y R R H M E A T L Q G G E R I T T A A V G N A
RANDOM_2852 L P R Y G D T F V S T E R V T S A N E S M E G F I P P G K F I P N A P Y T L I G Q D S T K L P N P P N D N P Q N
RANDOM_2853 V H V M A V C P R E V L M R A D Q K E L T M N S Q R P S D G I D L S Y I V S V T K N Y Q T M I F L M G E E S I S
RANDOM_2854 N I R G P Y L F I K L V I E V N A Y L K G A G Y P C N T E S Q A D E D W R F Q P Q L V E H M N T T V Q P S K I C
RANDOM_2855 A R P D P I R Y T Q W G T G L Q C L Y I K G P Q L F I K I K F S G N M H T E V E P G L K L V L N G E L R G K R V
RANDOM_2856 G C A G M T V L T A S L K V K P L P A T N S S S Y C H G E R E I E Q Q I K S R Q R Y G F A M G Y G Q P P S T P T
RANDOM_2857 N V S H L I R N L I A T A G E E L P M T D K R D K L M Q L Y D S V D C N I E L V Q V P N E I E D S K V A K W F I
RANDOM_2858 R K G G R A I T E R S A D R C T P P A D E S T R A K T C P Q G K F R L L Q F A M T S T L T I T C A S G D H I S K
RANDOM_2859 N F E S L P L V S A M L N M L A T E L L A H L H K K R M I V Q A Q L N G M L T S L Y I A A S L L K Q T E G L W E
RANDOM_2860 M K R K K K P S E A A H T G R F K D K C L Q E C P I G I N S S H P L L D H N N H L E Q G V D F L N L E Q W T L G
RANDOM_2861 L P A G L L N F V Y H L F I R L K Y K E M M H T H S F V I L L Y I D Q L Y P K W A N H L D C H G H V I R Q S Y G
RANDOM_2862 L K S L P Q C L L W L I A K N A K M S V V R I S V E T C C W Y A S V I T A S Q G F D K N R L Y A S A G A V E G Q
RANDOM_2863 V S C G K P A D N E Q M L S G L K S T M K R S R D A P S D S Y T E L I A T L S R A N Y C V I I N D E N G R C F A
RANDOM_2864 Q Q R N P E I I L N A V R E I V R T E G D K K A T A K C P R S V L R L N P C E L V A C T I A S P K D S R G K V E
RANDOM_2865 L G E V V H D V L A Q D N H G L K P N V T S L E R K E A L M S D Y L A G A M E A N P D W I E L Y L L I F E K A A
RANDOM_2866 T R S A I R C A N A D G G G D H V A K L W L G W K E S E T E A I D G S T G H A R A E S A S K S T Q E Q D W G K G
RANDOM_2867 H S E N F Q A N N K D S S P V A E A P Y D P S A Y Y A G T K D L G F Q L E Q R E F H F E K P L T R Q I T W H T S
RANDOM_2868 A F G R Q G S G S D V K T S L T V D Q C V A Y R A Q G G D H R L L P S F G Q C Q P I Q F P D H R S E V P P V T Q
RANDOM_2869 P K A L M K G E T I A L K V C L E M G K V D L G V M T A P M P S W W L T I S V Q W V N K S S T Q I S D T D S S D
RANDOM_2870 E K A G Y G D Q N Y T G T I G T P D P S I F F L T K A T L T P C A G A F E T G C V E L K K L L S N S F Y Y F Y G
RANDOM_2871 T V S L E V E D E G E A W Q L W V V A I V Q S M G L R E W T Q S P P N I A M D L R P A M S L S N S N D H P D E W
RANDOM_2872 A E A I V K E I D N P K E S V L K G K I A Q G V K E R Q D V K I A S T V Q L P T H V A G I S I I R T T L E S C N
RANDOM_2873 H Y L P A T F L G T F I A R R Y A L S P G V E F W A F D A S G K G K Q V F L W F L K G R C T I T D E E L W P Q Q
RANDOM_2874 L G K I S D G E S S T S Y F V L R F G A L G L Y D L K L G S C K D V S Q L T E L T G N R L G G W L A H N Q D N R
RANDOM_2875 E L L N D S R S K I C L V K E E A Q Y T F V I P E R T E S E E S Q S Q Q G R V Q K T Q Q R T A V K W R C W K A R
RANDOM_2876 E P L L S S I G R F C A H A P R L R L S M Y K Q G P W F I E A K F D P H I C W L P A H C A E L K V N P W L V L A
RANDOM_2877 Q N Q A I F A N G A G P T V L V L Q N P K V D K A S P E C S T W N P S D V R P V I G T S P I P R Q S A A P I G S
RANDOM_2878 N D D D L S I G V N N K L C I E T P Q Y S I W G P N K I N G T R A G T P T A E A T A E K E Q H A V L Q S G F Q A
RANDOM_2879 A E D H V C S I M V G T W F T N D E V A D L C S K G V A Q L I N H L G Q I K L E E Q D T G T K G G I I M R V A E
RANDOM_2880 N A G P L N V L D L L M L H R T L G C R A T L N S G V E K F A A E P A I P G V G C K Q F N H E K G R R Q L T W Y
RANDOM_2881 Q G G W I V W M P V L D K A L V L L L V E A G Q A P A A E P L L M Y L K D L S G L S S S G E P I L S L P E A V T
RANDOM_2882 E A K Y E M S G H D Q C Q C C V M V Q K N A Q G V L T R P L V L V A I S Q L H Y G Q N I L Q T E C Q E F G L K R
RANDOM_2883 T A V L L K P W V F Y S R P G S C N E V L T V G Q K K F E A K T P T S K P G E K G T E R P F A N Y G K G L M N G
RANDOM_2884 G K E D Y G S Q L N G P I G E Q A S P Q P E Q N P E K R Q K A Q E K D E Q L C W H V I F V A D F R C G R Y Q G V
RANDOM_2885 F Q S R D Y L G T Q L E N A D I Y A Y S H T G D D S A M M L E C A R Y E E P T P F N A A L T H A G D R L T S E A
RANDOM_2886 V V R A K H P A R S I Q Q S E L Y L H K A N E A I R T Q C L Q R K K R P G V I E A N T L K T C L R V T G N K A I
RANDOM_2887 L V I Q P L K S G G Y T K E Q L S W R M E T H I D L R E G A K A D A V R I C I E K R N S E Q K R N V W V A R N N
RANDOM_2888 G I E A A I D V T K A S Q V M S D E W G E T K C Q Q G Y V S V W K S I T H A N Y P G D E C F S H N E S K I R W L
RANDOM_2889 T H V N A R I A T Q W P I M D D G G M N D D G S L M F D A L M L Y S A P P R V N S T V C T L D A S P V L R A A Q
RANDOM_2890 G Q T A I A H S D V E Y A R M G E P P M I P N A T L E L T T Q Q K V D P N L I W V F A P F A L Q N R L E N G N L
RANDOM_2891 R L G F S V N G Q P S A K T E S T L I S N L L V T V F A S G E G K R K E S V A T S A V G D C V Q R P I P Q G G S
RANDOM_2892 D P G E R D G R M L G S I E L D K K C Y L R K Y W V W T L V P Q R G F Y P E V R S T D I E G Q Y K R Q G G V R D
RANDOM_2893 D D I V L T H R L E E L A N K S L L I K G P L Q D D Y C A Q H L R L D I P L A D Q V E R V D V V N P R T L Q L L
RANDOM_2894 I T Y V F D N V V S L R T L K C D S A P D V P D P K D D P Q T C S T N L L Q L Q R T P R V S G G W G D P H L N I
RANDOM_2895 E N Q L R A N I H A M L R F A A I H N D V L L T D F P K T P K L T G I M L I L R G D R D P Q G P K L F V R E L S
RANDOM_2896 L V E T R L R S L S D G L E V R C L H P L I C V D G V V P L L G M I A H I A T A S S P I G M G E G L K K A T K P
RANDOM_2897 E F K Y R N L W Y S L L L A R S N D L L L L G I G R Y R L T L S D R Y A A L R T A D N D S T Y I R V W A R K Y S
RANDOM_2898 P R Q E N L T L L K N D N A S K E A L C L R G L L M N S S E D F F C E I G T T R V W D Q R K D F K C D T P G Y I
RANDOM_2899 Q S N S T L T P S K I V P F E F L E D K L S K F P I N A S I Q G M M D A G S K A A K K F T Q K E M N S D Q W I V
RANDOM_2900 Q E I V Q K M N G A F A R Q T P N G M P T R D S T N G A K P T S R S G L W A L E A Q K P A L R A L Y E V L Y V D
RANDOM_2901 W A P I T Q Y T V I H A G Y R H S E Y D N E M N L G E R S D V G D V A T S A L N L T K L S L I I C K T K G N P T
RANDOM_2902 S F A G E A L T R L T D I Y P F L I G V I G V G D Q T P S V K D T I T E D D T I F C H R V E R R L G V M K P I M
RANDOM_2903 A V G Y M M H L R G G V G A K I A N T I R G K L T V K L Y F K E L T P R V N L R R I E A I R D E C Q L S T Q P I
RANDOM_2904 I N V A H K G P I P L K F T D W I A V R Y C V G I F L G P N G D P K S D A N N K D H K R P I M Q G D R G L G I P
RANDOM_2905 M K R K G L P F I V L G Q F K S F L G Q R R G Q A K A D S M V E D E Y Y G C R V Q D D F S E E P S V R V D G Q S
RANDOM_2906 A D M W E L A N V L F W N K Y T K C I A E S Y I A C N G D I V P I Q V L K A R E K D I Y S P P A W R T L P D H D
RANDOM_2907 A L N G L P S V A E C R L F G N N K D G G T N T G D Q K C K F D S L Y P R G T K T M C F T R D A D V D E A V Q V
RANDOM_2908 L E V P V H T T L I G T M A I T R V C L A L L S F R Y T L N S R H D I A L I M V S T V P V T I L I A D Q D P G I
RANDOM_2909 Y V E V V K Q A G A V E G V Q H S N Q D K S I P K L S S P L E I L L L H V E K L S V K A F V M F L A A R G N L N
RANDOM_2910 A R N A E K V S S N E G S S E D G N L I P N L N I A D N M L R P F A S R G R A D E H L A I G P L G L S A L G P K
RANDOM_2911 H L K A T D E K M P K Y L M L N A S D F A L A A T L R N Y T R P S I P A I Q Q L A I F L V L R P A Q G G L R S D
RANDOM_2912 L S V H K F Q D S H G F V Y N V V L H R S K D S S Y L S P H A L P A D L V A G D R L L R D Q K S I S W G K Q T V
RANDOM_2913 R D D V T K V P L S N V K S E Q E M R V S T V I D G L T H N N S V R S S A S A S T Y R R G L N P I E Y P E V M L
RANDOM_2914 K C E S P H A L F T R D E Q T S D L N H L E K T Y S T T S S S I P N L E D D L N P Y G K V K G P V A P T D T G S
RANDOM_2915 V I T K R K D F N M Y E P D T H R V C T E V V D V I K S A V E G M L P S Q A A K A L D N I P K Y Y K E N S L I N
RANDOM_2916 A K L K P E A H F D A P G W R I S I A P P V K Q G G N A I K D V P Y W G E R D Q F R A R H A T Q Q N S V I I T V
RANDOM_2917 E L P K M C F T E G S I G L A S L V R R V P R A G V K Y T C K P G S G T N W I A N A A L N V A L E G N Y S D H N
RANDOM_2918 H K Q Q V N A A P V K A R G N T V C A Q A A L A C F A P L V I A V G D Y V L L L Q T L I S W K S P Q R T S P Y L
RANDOM_2919 H N L M G P G K Y M V G S K A V S L V L E G S N R P Q V R A E K E E I L L S G V V V S A W L K D M G N D N K M V
RANDOM_2920 L D M M N Y S P K C F A G D S R V T G L A G F S E T R L Q M L N F V S S N F R R V P F S S G K A L E K Q F L Q I
RANDOM_2921 R G D F V C G K L K V L F L F V G D S R F E C K F G M G A V P N A V Y D I Q E I C A D L Q E K T A E T D D Q D Y
RANDOM_2922 G N E D T I D G G T M K A K S L H V A F N K I M E C Q G P R Q P A Q H T T T D G S N H Y Q L G V S R P V K D C V
RANDOM_2923 L C D R T Y D I V E K K G G A A K S D A S F P D D V Y M K E K I L L T E K E A K V E G S P T G L N P T A G L K F
RANDOM_2924 Q G I W V L R L F E S I S S P Y T L G N V E T W D N A L S T H V P P E I K S S G I S F S V E S V G R W G Y G T K
RANDOM_2925 E P N Q L F S Q L S A F A Q V G D V G P A I T F E G N A Y P A D S P N C Q P V A T I Q A L T E H S L Q L Q Y E D
RANDOM_2926 D P L G S S S I N E L V G G E Y K S L I L Y V L S I P G D Q S A R M A G Q I H I L C S P G K I S Y D G L A L Q L
RANDOM_2927 V W L G G L G N N V N T A G V L T G E L L V G G W P N S P F Q C A S P T L N V L S W T T R E T G V I W V R V C L
RANDOM_2928 S G I N L V F Q K K K R A S A Q A S A P N R A G T Q F F V A A E E K S H L D M K I S R M N A S R D E Y S P V A N
RANDOM_2929 C N F Q K P S M R E L L A I E S G S S D V L R L T A K R Y G V G K G D K P P M R C V Q G P E K V L D Y A A G G H
RANDOM_2930 A E I P N R G I E L W H E M G I D A V L P P K L A E G M S K I D F I V P L D F A T F C I D H Y F C T T T V V I H

RANDOM_2931 R L T V F F Q D E Y G K D T P V S K Q D W R G K P G K E K D E C L D A E G R N N Y E L P R D V W A Q E P P S D E
RANDOM_2932 P L L I H T F V S D V L A Q R K L L M A I S I T E K L R L I N L G M A N A V D L M F Q G E N V A P E Q G G E V L
RANDOM_2933 D Q G P T S M K L V L G T A V Y K T L D S E K R N Q Y C P S L N G T R R T G V D G S D S S R G R R D N L E G V A
RANDOM_2934 P C F D G P S K D P N T E P G L A V I T I C A K G R G P A G C V M E L A E T G L Y V R L N S Y L P K Y T A S E S
RANDOM_2935 G Q L T R G P Q A P N P G L V G V N Q Q Q Y P L R T N A R I K T S N A W G S T L L E A L E A A M K D L R F P I A
RANDOM_2936 A H N A K P V R A S E R G I Q F G A W I I Y V K L D A D T D T Q N Y V T P C G H H N E K V V V F V L T L Y V P L
RANDOM_2937 S T G K L G V D P T L W D R S I N S Q L K G P P D L V L K S Q P K T L K I Q Y H P L I N A N K T K P G L G M N Q
RANDOM_2938 Q L T D S V D P A T G R H G K K S G D H K L R K E N Q C S L W T P S V D G M A S V I L C F T P M E C S F S E T W
RANDOM_2939 V P P P F A L T G Q L N P S V F K S V Y L L R S L A S G A K I Q D K G E P S A A N N Y V E L F W I A I C L Q F T
RANDOM_2940 K K L N A V S K R R R F A T K P R L A Y K E T A M V A S N A Y I S D R I T Q G Y P D A S A V K N G C K N L Y Q I
RANDOM_2941 V Q V A H A A S S E D G P G G A S T N L S F T D T K G L Q E W F G D T R Y D E G C T M N L C W G P R V W L F V V
RANDOM_2942 A D K R N R P G I G I T A N T T V D E L A E A L M V F I E Q P D H P G L L G E I T P A L K K H Y A P Y D K G K L
RANDOM_2943 L N H F E N N L E E I G D A E F L P S P P T T C C L T L R E Q S T G T V Y P T E S L D M G M D P T A G F K A R L
RANDOM_2944 T D Q T L K A K V D S A E R Y A G P N T W K T S G V V I G A T C A M G D L A T Y N G E L I A Y D A I L P P V T K
RANDOM_2945 V Q N L L K F L V F G W F V T T I K E S D R T Q A L G W A Q I L L S L V N D N G E L A R T C R A C F Y T G F G A
RANDOM_2946 K L S T H P I S S P Y W C W E R E F G I Y N V T K G L P I L W I M I Q I H K N A W F S G E N V T Q H A E L Q L L
RANDOM_2947 G A I R A L F Y Q I A L Q R K N G V E R G S V I Q P S D Q V T S T K M H R V C V L F T D V V L S E R E L N K E C
RANDOM_2948 E N I Q P L D K D G L R V L A L S K S R A S L E K L S T S C G M G A A V Y A F D K P Y L L T G P T L T L L R K E
RANDOM_2949 Y P T S C L R R R A F S L Q I E G F V S Q S Y L N F T P W M A K V L G V Q S E K L T Q V S Q P L T V A W V I S T
RANDOM_2950 I S H P V L A P I S K I R Q P I G M K L H P L E Q G Y Q I Y K K Q K S Y T Q L T G T N G K K N P S G V G N T E R
RANDOM_2951 R N G L K L A G L T N L K V N V L H R T T M A L R N G M L A Q S I Y A L L Q S L C D G A D E Q G G A T D G T G I
RANDOM_2952 P K P R K G D L N Q R D Q V P D M V P P L G S D I V V N L S M A E L G S E K P L S P I Q V Q V T I V L A R H T G
RANDOM_2953 N A C T G L Y R L R D K W V P S E D Y T Y A Y P A L F G M L L V E T H P I K D G L K L I T I L A G N F V K P S F
RANDOM_2954 I R S F F Q A P A A D G N A Q Q K G S Q G L W Y E L L S A D M R D A A S F N N A M T L P L S Q P L D S T C W N M
RANDOM_2955 W T S F L A K T T N V T L G S F D Y A F R Q G R Q T G G L E F A D G E G V I K S G Y R K A E L N E R G K A S G R
RANDOM_2956 T E A D D Q F F R G Q E A A A A S D H G P L P T R A G L Q V E T D Q Q H S D F G H K T Q S Q D A N H G M E L G A
RANDOM_2957 H K T Q E K E I V F E L G N P H G T R V E T V E Y P E I S L T G A K L L S F Q K T A S H Q C G R K L D L M K Y L
RANDOM_2958 S T V F L E E S S K A L T T W V G V Y T M V R H P D A G L I S E V D P M G T L R A I S P C V G S A R S L H P A P
RANDOM_2959 Q A E R V P H S P T H R S E W K T P S V E R Q L W N N Q T N G D T T P T E N S L A V E V N Q F S A F F T N A V A
RANDOM_2960 V G G Q F L I Y T G A G V K D K E D D T T I Y S S V A P Y L T V V G P V F Q S T F I H R Q R F T D D A K S M S R
RANDOM_2961 V V N D T G K C R P C E V V R S V S F E I Q G G L A L T L V L C K Q V L D D V E E K P G K H A R E G Q G Q F L S
RANDOM_2962 N T I A F N D F H G N K L F Q C L C D K N P M K P C C L L V A I S Q R L L K G T R Q A V T L L F N T R N S S V D
RANDOM_2963 I E L R D A L Y P L E D L V L D G L K F A L G L Q L S T R M N W L I R R V D N H S V A L R V K K P A H V E L E I
RANDOM_2964 Q P G T G I E E S A L Y C V T V G T I S A M G I G S E P K W E A S T S L R E N E D E P C L D Q K L D P Y M L Q N
RANDOM_2965 A I T F A Q A K A A R G K C S R G L A V N K A A Q A V D P D P K H P M L E I V S D P T S F Y V H D P R A P L K V
RANDOM_2966 E A R T F T D G V R W D L V C K V W A A L G V H K Y G I S S N T G A S F G P K I G E Q N E E N P A G A L W K L I
RANDOM_2967 Q T K D F G I G L T V L T P C D V S P F D E R M R M N A P N Y G D D E K N D P T A F L M V G E D G S A R L T N Q
RANDOM_2968 S N L E Q K E A V C A V P C I L E Y A I A R A A D R H E R T Q L P L C L N W A Q G I R L A F Y R G M N V D A T V
RANDOM_2969 A Y F V R F E F M D V Q Q A D H P S L I V N W E C R T M G L S W G V T I E S S T R G R C G N R P K G T R P E S Q
RANDOM_2970 R V R A V S I E M S S W F L G E T P C T V A V E V V P S R E G S A S M G R A Y H L I A E A A Q K T L E A E T S L
RANDOM_2971 A G I R V Q Q L G V G S Y A Y F C D A G L T P G E K Q M S G D K Q S G K E S H S V A L L V G N L K L N F I P E L
RANDOM_2972 T L A T A V V C L T V A D T G M P F G L L N V Q P E S H V L N A V R V K G E A P D E F A E P R Y E I T L A D F I
RANDOM_2973 S M Y L P N V E F D V N T L L L L W K L K V E V V I R V I T E P E Q K R E D Q K V V V M T E Q R G I N Q L A H V
RANDOM_2974 I M P T L R N V A L E D L G K K E P A K I T D I R I A T V D Y S P V F N N E R A V N E Y M H I V K K F Q Y G L F
RANDOM_2975 I S E L I S L A P I T L C Q I K N E D T Y I S V G K F D K P A R E T P L G R I D N W Y L I R V A C S A N N F G V
RANDOM_2976 L I S A L K T T R R R D S V W S Q Q K I D S I A V G K A A P G D I E S Y L D F V K Q L V G L D Q N V D Y V D K N
RANDOM_2977 G Q Y W G V T R K P G I L R T D A W V R L N I S R Q V M Y L D F Y T A K S D G V T D K S K P P M N F V Y Q R L L
RANDOM_2978 I L L K L I G I T K L C T Q D K H S A F H T H M D H H N V I D G A Y R R R T F N V D A A E R L D Q S V A Q K V N
RANDOM_2979 C G I V D T T G F Y A I L R N W Q F A L S N A K K K N S Q H G S F K P G T Y H E V A P G A A K Q G V K T Y P L R
RANDOM_2980 E R E P T F G A K A K N R Q F S F G D A R S D E V F I L S N L M P I M I H R Q V S Q M Q H H K L T E D V W E S A
RANDOM_2981 L P N V W I L E D T G N V F A E A E G S I V D P Q G P K L P N D A V L K G Y F S Q I D D H I T G I N G T V K G Y
RANDOM_2982 F L Q G A G T S P L V D Y L T E D D A D A Q L Y Y L S N E A E P R E F D G R K F K V L P W G L T D N R S L A C L
RANDOM_2983 I A K N G L I W T T P H A H M W G V Y G A R G I F A V L A K K Y F D Y E T M D V E P M F N L C R E V P M Q Y S T
RANDOM_2984 F L T G V V H S E S K L T E S V R V A G K S S K Q Q G G C G R H T Y T I D R S P G D N T A N W E L R E D L Q L C
RANDOM_2985 I P K Q W P C H Q Y A K R L E A I S H D D I Q F A V V I T E G Y T E G E T P G F C G A L A A S K C D R T E H D D
RANDOM_2986 P Q I Y T E D C K M L M R T A H T E H W S I K A D L L C S N E E H E R S K A I Y A G S Q A L F C F S N R D I I D
RANDOM_2987 I D P T H T F D A F Q S P E K K F V N Q T S D T R D A P I L T Y M L T M T R R L P I Y N M H S P S R T T L D D R
RANDOM_2988 R E V G G G Y Y C T S L E Q L D N G A S F A W H H G N E L E M C R E E H Y A T S S F I V F A K Q E T F I Q N F L
RANDOM_2989 A N E M L N A K T V G A E A S Y G R F F S C Y C P R P D K A F T P K V R Q M P S L G E S Y E F M L L W L Y Y K M
RANDOM_2990 M R T Q P A H A A E L A Q Y Q A T N E I P S L G S D N R A I P E F R S G P T K Y T Q W L H Q A V A A R I G L A V
RANDOM_2991 E E F I H Q A T E D D A A S E N L L F T T F G R Y G I I E Y L A R E A W F Y S H Q A E T F I T G N G N G R K G I
RANDOM_2992 G T T G M N P S V V D V P Q S T Q S P E R L S F K V L Y T D I V A P D R Y G A L A K D F N D R V K H E L S D V A
RANDOM_2993 L D R V L L N T H E R V K R N F I I K S A F E E Q R Q A A A T E E L T T E A N Q D L L R L Q E D D E N S A P L V
RANDOM_2994 C I K G L A L E Q K A V P T K L K I G Y E G D N R I L G I G R L V I T T G R K L L C D G D V I S K P C T M I P P
RANDOM_2995 C V S H F R R V C C Q C F D C G Q T E P I I G S R D G F V Q W G A H L A V I K C L T Y A E A G L S E V R S K D D
RANDOM_2996 L S G F R L N A R N G K R E H K D L G K V D A G P S Y E F C N V D K P T F P T D Y S L G G S G P E L D C T T G K
RANDOM_2997 R L H A K T P F K P G P L R V L K F G G R A K D N Q Y M G A E T F K G P S S G D P S K T V F Q K L V S A I L G L
RANDOM_2998 A L R R K F L F L P A F K K S S M Y L D K S S L G G T L G N I I Y M G Q A A P D Q S I N E D D V H D V W A F V A
RANDOM_2999 M Q R Q A S P L D R T L T L S S G V D G A V V W T K C D E R I L S R R L G N L N N K D L A P T A S Q G M H N S E
RANDOM_3000 R A E R Y P D L Q T T E L E S G G T A T G N M V A K E R F L R N V S R E D I W G R V S V Q G S N A P F P N P K L
RANDOM_3001 D S R R I E R K L G Q G P I D I T G S S N I G S G I R N E T Q L I V N T T F D Q C C P D A E H D A G T L Y V F H
RANDOM_3002 G K L S V S P L Q F K T E F S T E A G I E D A V N L A P S R P E H I P K E K V Q L N S K V S N E T K G G K F V S
RANDOM_3003 K K K S S S L R G H Y I A K I N T Q A P F W K R V S S W Q D R V V Q H V Y E A Q Y V S Y P P C Q S R Q D T E M P
RANDOM_3004 S A Y P E L K Q F D S F G N E K L Y Y R W R G E K V P A K P P L K V S S Q G G K S Q N V P S F A G Q W M F F R G
RANDOM_3005 C L N A F E L N S V C G H T S K P F E G F K L K D P S Y A Q A V T I P A F V G R A T S R I Y A R L D S N P E D I
RANDOM_3006 R L L P T F A L L F L Y P I S A E L P S P I Q L E S A M G L S D A D F G C T S D P L P T R N G W I Q T A H A T P
RANDOM_3007 L Q E P M C F F G P Q K S K K Q F Y F R D R L G Y D E L Q K Q A V A P I Q F R E A A R A T P N E T G H I V V I N
RANDOM_3008 L K I G T G E T L G P E Q S P R R F V T G P D S W G C S R T E A Q N T A P Q S T E W M I M A S R L N A F G M G W
RANDOM_3009 E F N G N W L K G F L E I L A N K G R H P K D Q V D A L V K F L N D L R S F K L L S E Q V V A A L K S S Y Y A L
RANDOM_3010 S E S E S K C W S G L D G R A L R T A L Y A A A K V R P P A R K D V Q F T L P R I S P E F A K G V I G P K K M K
RANDOM_3011 Y R Q F L A F A Y L V S R N E G N F A Q V L M K G Y E S I I C F P S S S A S P C T V L L L S Q P D C Y S S N R A
RANDOM_3012 Q R T H F E K A D E L P G N P P P P Y G S Y A P I Q S N P L A V I W K T V K G G A I I T V S E P N L Q K V G F A
RANDOM_3013 S T R C A S S Q T L K D N Y M Y L L A R N V P Q M V A Q P A A V R N G D L V C Q D Y L G V L N G W A K G K N A N
RANDOM_3014 R F G Y P I N M V V V D A R K N I W V K N A P T G T P H G E P Y K F M A A L G E S V V T D K C V E Q D N L L F S
RANDOM_3015 T V E N V A V L C S K E Q D Q R G S R Y L F C L S N Q A L L E Q N I R E F L L M F A V K P V G R P S G A L V Y F
RANDOM_3016 D V S V Y E A D G S H T Q T D P K L T V Q L D A H K V F L T T I P N N I S A E D I C K L T A H I A Q T A R S D Q
RANDOM_3017 A F K M T E P A E N F G K R E K N T A E V S H G Y L Q A L S L L G K S K G L T K V E I V G K A E S K F D Q P E G
RANDOM_3018 A K V G F F A L A N F S S I G A V K K A E L A C V A Q T S L V P F P L G F D L G P K G E T T W V Q E R P V G T E
RANDOM_3019 N D S K A K P E K P A A D S N Q N E P R G Q G I L K Q A K P A L L D K D S I S G L A I E M W D A R D L S H R D V
RANDOM_3020 A W M A I Q A C S G I A T G H L M T Y S P K D Q L I E S L A D A N K A I L Q F T S Y A E N W F S G Q V E Q C S P
RANDOM_3021 T P V A E A C S G L R T P A E I E I T I K H Y L I S G E S K H S K V K H K K I G E H R K P F A V K L I T R S S G
RANDOM_3022 A W S E K A E I A D F Q R D S P I E L A V D T P S T I M L I W T S A V G Y G I L S A Q N G N L N K T G S Q D R L
RANDOM_3023 Q Q I L L E N N K E G K D E S T S I T S I K L V L R I R K D T S L L L A P Q D R V G I D G E E F K A T E P N R A
RANDOM_3024 R G S A K V P K C V T L A Q N I G P L P T C D N G Y H G I K G F H L D P L S N P R Y E A L F N A Q K A A K M F W
RANDOM_3025 K G Y R P F D K V I P E T D V E A G S I I N H D L F H K K M T D P L P D P P P E S K P K S F P G V G E R C L Y A

RANDOM_3026 V L S P F S S F A I I E L K V L K V A E P G Y A F Q Y H L Q R T L F A A L D H H Y R K C L R E Q R V S A Y C R E
RANDOM_3027 L L T P K K R C G P R I H Y E A F H S L M A K R F T E A N I D V G V Q Y L K H F E G D G T S T L M V K T L L T V
RANDOM_3028 G I S K L S N Y L N D I K E L I H N M K G Q A T L Y V L F A V S P T D I Y L P C S F M M S T G L A K I T L F M L
RANDOM_3029 F S A G Y A V A V G A K E R T V H G L Q E G E W W K N G P G A P L L W Q M L S G E H S L F Q I E R F H S Q I A E
RANDOM_3030 A Y V G P P V E L P T H F H T M L N T I I N L E S V D Q S K H E P G F T V Y L R R R K M G D R E G P C N I E V A
RANDOM_3031 E I F C V L Y S N K G Q G F T A L L S L E Q Y L V L V S K L D Q R C A I T P T S S I E F E G E T W V S E K N R G
RANDOM_3032 A W L S P V W E E M S T R K E S A A N E R Q A Y P A N P S L Q I M K E E R R S A Q K A T P P M M F F Q G L A C G
RANDOM_3033 N A P C T I N A A V V E I E A C T R S N T E R T A S V L E A C G W E S A V G T T G K E G Y P I P E T S I P K P L
RANDOM_3034 T E L S P P E N D Q K G I L F L L I L Q S S M P I Y F L R I D A S M N D S T Y V T S K V P E K A L G T H C F Y E
RANDOM_3035 A R K V T A T G Q M S I A G D F T L L K D E T R L R A H N K K I A Q L Y P C Y T G M F N A I L K K S L S V M I G
RANDOM_3036 G P S S G K S L V M G R G K V R K D F C G F F K N V T I N F N E D R Q E H G A K D N T P D N T D M I G V V E V Q
RANDOM_3037 T T L Y D R A G F S N S R L G L M F V Y V S G L A F G L S A N C H P G F W L L L Y E E G Q Q L A N C T K A P V I
RANDOM_3038 P I F H E V N D M V P R T E W D F Y S V A G P S P G S S N D C T S Y D V K I P Q L N P S R P D M G V N H F S P G
RANDOM_3039 F T S Q E L R P S T Q R K W D I V L V E A I H A Q L R L P K I S C G L S K L L R S R I I I D F W L A V L L C K D
RANDOM_3040 S E H N S A P K T G P L T F S V P K E R K A S L R T I H W T L I E G I T G L K M G A L Q T N N L V H S D G T Q K
RANDOM_3041 S T L Y D T V Q M G T A T N G Y I R A T R G R Y L P L F S E K T K L M S C A L G D D L D A E A C I G E I F T N G
RANDOM_3042 Q Y E L R A Q A N I I V G H T A C E L G Y V E L E Q D E P L K G K G A Q L L L R G Q S Q D A D T G S P R L Q K P
RANDOM_3043 T T N D D Q L G Y F Y I P M L A P Y Q E S V N K I N E V I R I L F P R Q K V W T A K A N G L Q H G R K S R I M E
RANDOM_3044 F S P H W T T V L L V R S P N R N W Q V Q R T P H A G N M L G Q Y G D L T G D S T S Y C D D K N G T L K S V Q A
RANDOM_3045 Y S S A L P T V L H G D L S G E K P G C K A S C N R R S S N Q F K S K K R V S L H E L P I D P D S L E E A R T V
RANDOM_3046 P L R S Y D H G Q L M L R V A L G G G K Y L I T F A L I H L Q G T I I E H S S D E K L L A V T S I L T H R S G L
RANDOM_3047 P S T R T E G W H K A Y K D G L P L I A T F G F K L D S D V S E F A G G R G L D I V G D P V N K K F R N G E S I
RANDOM_3048 G T L A D R V S T Q H G V R T I D L E C V L R F Y G S D K N L L I I K D A V H S V V H T C V V E I D E Q S P I Y
RANDOM_3049 V S G E I A I W A D I V S P M M P R Q D L T Y Q L G H V G W Y R P V C F I H I A A R P A I S E T G V D F F E L Q
RANDOM_3050 L E T G L L L E L D G K H L H I N P A K P R T K S K V L Y L L T Q K P G L F T P E P K V T S K R T D D L P E Y V
RANDOM_3051 G A M K T Q F K A G I E P T L T Q G N P S A S K S V V H V F H Y Q P S R I T I V V V H S P A A G M Q C A G L L R
RANDOM_3052 P A R K V L L E F R P G E A G L Y V K R R A E D N I H H F D S R P E A M Q L R Q E Y L G H V G L C Y E L H S N A
RANDOM_3053 P S D Q G D I S P W G L I D D G L Y T A M T K L A K G D N F R M F A S V E K V D R S S E E H A A G E V Q V G G A
RANDOM_3054 L P P L Y V F G L Q E I L G E A N P N S G L Y Q E T L V A A S A E F V Q L E T M I M H T G E F V K F E V F L V Q
RANDOM_3055 L T Y R D D L V L A T G T K S L G A L G L V S L L L S V D S V S L T G I V A T L L G K L C E E R T G G R I V F K
RANDOM_3056 N Q I F V G L V Y T V I I E L K V R V S K T Y M G E R L L L G P D S F R S V K I E I S G V D K N I Q A F A S I T
RANDOM_3057 E P Q L S L Q K L E M K T M T L D L K I G G L Q Q D I P Y Y L I E Q E F V I V C L K K L V H N A V Q A M T P E L
RANDOM_3058 L S L A G Q K R F L K E M P T V R E N V Q A D I Q I P F G N A P L R L H P E D G S D P S G Q R I S S F S I P F Q
RANDOM_3059 K G V V D P N D K G L M K T K R D T G K T E P H W H S I G K I A T A S L V T F K E V T A A V Q F D S P T L R R E
RANDOM_3060 E I T D G K N K F N N V D G E S L E W R C T Q T A N W H S Y F S E S L L I N L S S A K V T Q L P D I N S S T W F
RANDOM_3061 M Y A F K E I S A R P W N N E P R K G A T N V V Q I M F L R G N T L K A K V F D G S W L I S K Y V G A P F A L F
RANDOM_3062 G T A H H S Q H D S G G R L H V V N K G C E F H D S T F S L M R A K N I A N T H P L L Q R F S G M E G L Q L V Q
RANDOM_3063 S K M E P S T L N T A S P G I G N W V T E E L K R V P T F T H I V G V Q G K V G H Q V M M V E C D K I R Y S A N
RANDOM_3064 F E V T V T T A A L D G Y K K P P S R L D E T I L M G P K C S L D D A F Q K C L S D L G I F Q G F R G N E L S T
RANDOM_3065 N L V S T L P A P V G S D T H I L C E L V L L Q D T G T D E H D K I Y A T K K E G G I I L A A S F T K E D E H G
RANDOM_3066 G G G G V P K A W E G V G K D T W E E V L S M T E A Y L R K Q R S E R I S W T C S T L K N I C Y I P Q T G R R V
RANDOM_3067 L W D T L S D P N S Q E E E A L E A A T M V I C R V M Q E R Y S A L Q S K Q S V G P P A L Q L L D N Y S N N F D
RANDOM_3068 S V L C K Q D H H A H N D E T Y K R I A I F E H Q R L T F P V Q L D R L Y D L P A I A L T S I M T S D L S T E K
RANDOM_3069 L Q E T G G E P T K K D T P K V P S Y P A S H E Q E Q V T L Y P K Q Y I L D A W V T P E D N L R E S V Y T T G K
RANDOM_3070 L L M E Q A Q V A A K A L D P F L L V K G A S I R I R N H H E I T Y V L C V V V V A G R W L V V D D A D V Y Y G
RANDOM_3071 K A G L K Y A W G V E L F D A P I I V S V P E V I P Y Q G R P N I L W A W I F E T T K T V H R Q Q F P L I T A L
RANDOM_3072 G A I Y W G R E G R R A K D S D S A C F V I I L I E I E N R F D Y Q T V V E K G S R A E A G H G N S P C L R G E
RANDOM_3073 N H K E P G A W C T I K G F D E R I C L L Q D Y L T I S T E A H A V Y E I Q E R G L A F V I L D I A V I A A I T
RANDOM_3074 G G R T T N L V G S T P Q P Y K K T A D I T D E G S V H D M I F Q Y Y L E T R L K T K L L S I E I L G L K P A G
RANDOM_3075 S K V S G L L A Y K I Q Q E S H T R P L Y I E P F V C G K Y V Y D P G P I C D Q L N G K H L I H L D P N T T H A
RANDOM_3076 T S P E G S S L G P C A K F F D A K R A E A L N K I F T N N G P A I S L N M Y L I S I L V S V Q L Y P T Y G S L
RANDOM_3077 V M A Q Y R T L D G S Y N V K E R T S V F G K G S F L K R K L R Y L G W N P C R S I V M D G Q T A L I H Y D S Y
RANDOM_3078 I Y Y W K H H D C T F L T K W C K H E E K T V G L N S K Y N A D R P G W T F S I E E E L K E L F P D D S G T Q N
RANDOM_3079 A E L C A T W A C P K D T V W P Q V P H E L E A A A N M K K E I A P L G E N G V A K S I V L A A E L S K N F H P
RANDOM_3080 P E N E L H G Q A F R A V P A I G F A S G R H E T D V Q T P W C S V V G D V F S P A T T K A R R I V L F Q T K F
RANDOM_3081 E C F V D I V P W F G Q L S I A I E F D A I K L K M D S A H P R R E M I C S F V N V E E L H L F K D K T W Q D I
RANDOM_3082 A R Y D S P Y R E Q Y Y D G D K Q F R N T S L L G T A G A Q P W V S Y K Y C A D L V L E T E F P A R Q T V N E T
RANDOM_3083 N R Q D S H I W K S L L G D M Y R S L N R C R A H K G Y R K E S A W S A H G A D R G P I A E R H M L K P C K S A
RANDOM_3084 E W G S P N M R P A G D V T I R T R S L T E S A V W I T T T K I A G M R Y H A A A E I L F F Q C L G N T L Q S T
RANDOM_3085 I C M A P D V P N E M K S L A Q S K V P E G E R L A L V W Q N P T D V M G L H D G T F G A F S I K Y Q L M D D R
RANDOM_3086 E V K R P H R I I L S N D K Q G H V Q G F Y V L T V G H S G P R Q T N C N Q D E T N I G W E A E V L I K A K K V
RANDOM_3087 C L V N R K V Y T V Q I E L N G T F I Y E H Q Y G A P E A G D T M N E N D T G M W F T D S R T Y F A Y V R F A T
RANDOM_3088 V G D R A H E A F E F L Y E K S K E P G T W E P D F V K W V I L E E T W P N S P T E A G S C S K A D R N K Y G G
RANDOM_3089 W S K F D V S G L K G G T S M S T E G L N V R R L I Q L D D L W G M G C E C T P M V G A P M L Y G E D Q G S F L
RANDOM_3090 K I T A T D T H D F L K N V S V V S F Q K D R T Y L S T V T A L E R I R A E G E H H L E S C R I Q L D D Y N K C
RANDOM_3091 V M P D S L T R G N G S G W L E Y N V I Q L Q G K K Y I G A N D P M K H N I G S L R G D L Y E G N P N S R D L L
RANDOM_3092 A G R A L T A A S A V V E R F A P H G L H N K D I V F Y T P P R L S E G N V S K V G R G E G Q R V G E D H H A L
RANDOM_3093 R E A L Q I D P V L V K S R S N V K M L P Q Q Q E G F A A T E S K P K T L K T P S S Q T R N F R V D G K F Q A L
RANDOM_3094 Q H L L G W W K V E H H L D A F V G T P E E S L K A T L S I N F V A N T A G I H G G P A F D V T E R A R T N V F
RANDOM_3095 L R K R M Y K D G S V Y V R Q N S G I F F M H Y R H V N V L Y F V T A C E P D S G E L I N N T L E A V W G E H R
RANDOM_3096 V W N R V A S E G Y A K A Q P A N R P S T A Q Q S V A F Q K T I K N D I K G I L A N K E G Q N A T Y Y T G S N L
RANDOM_3097 A K N R E D Y E R S N V Y L K W S D G V F E S K P L G M Q L L R K E T R S V H K D S V D S A Q S P S E P A D I Y
RANDOM_3098 H Y L V A S E D A N Q A Y Q P G V A E L E E Q A D A N V K P E C A Y K M G Y K K V L E K I E R T S E M D I S Y M
RANDOM_3099 Q R A R A S K Q T C K N L G I F P H Y R Y G D A K D I Y L K A C H G R L P R T K Q T S Q V G W S K L S D V F D T
RANDOM_3100 C V G K L I P P C A P V G S L N Q D S K Y V Q A E L A R P R G T P E H P I V R E V F T S P K H V T A R W A A G Q
RANDOM_3101 Q E N G L L A Y G E Y G R L C S A R D T K L A V V A A E R S R Q F R E T K R S A I T R V R L T S I V F K E F D Q
RANDOM_3102 V S G G D D A Q Q P V K Q L P G D G T K E M G L G V R A A S S P R S V I I I L I L L I Y Q V F V M Q S S S S K R D
RANDOM_3103 W L V D Y H T P T S S I A V G A S D H L L F M G G S T C G H P L S R V A A I L G I A E G S L T H G E W P M H S I
RANDOM_3104 A T L I Q I A A F F S Q L K Q S L V V L D G C S V D A V I R N G A M A G C G A I L L V P E D D E A F L T S I R F
RANDOM_3105 G I G S T P D E K I W S S D V A L A L F N F D T H L S V N S L W C D G N S S I D S R E S S S I T N R A V L L Q G
RANDOM_3106 G L A D A E G I T Q G P N W P K E L L Q H R L V S Y Q G T D G T L G W G T A A H P A T K H N L T T A T D S Y N D
RANDOM_3107 I A N A A Y L V H I K V E T Q F A Q L Y A N Q H Y S I W W L Q E E S F P K E L I I S D A L T G I I L G E F L T D
RANDOM_3108 A E K I L L D A H E N K N G G F A T N C G S L E N V L P I Y A D I T K L E V D P A H Y T F M Q E S N N P E M E S
RANDOM_3109 R K Q R M F Q A I I M Q L T D R E M A S N T A R A N Y K P L S K S Q I G I A P G E K R E Q G I L E H G W I V G T
RANDOM_3110 P N G F T D E G M C F V A W F Q E Y P V I T A A I W F S L H R N E R K N L E E R F M G K D V W T N G S I Q F S Y
RANDOM_3111 L K R N G G M G T L V M E N D I D A Y P K F D P Y P I D L A K G V R T L L D R L A L S L E E L F M M N D V E D K
RANDOM_3112 E L P E Y A V N E Y T A K A P F G L P E Y Q T P P W K E A P S T N L L R G P Q G A L A D L D M G K P D N M L I I
RANDOM_3113 M E K P G S F P G D S G D S Q P D P V E F G A Q G I M Q K A K N H L I Q N T R I D G S T Q W K V A S Y D K R S R
RANDOM_3114 Y I R R R Q K L N T A L M V L I I R Y F R I K A Y N C K C G E L H L E E S H D I G L A G L N D N K D G V S G R S
RANDOM_3115 Q F L E E V T I H N V M V A L T I D K L T F L G E N G N T E G G P A A A W L I E L V N N I I S I P P A L H P Q R
RANDOM_3116 F T N R G E T E V R F G K L S S G G A M A P R N V G T V S K T V H P S V G F L P G Q Y G S A M Y C T L R P M E R
RANDOM_3117 G S F Y T R V G K Y A D L K R L I V A D N Q A G E L A A K G L A Q Q P A G E F D G L G S S E H F L P L F S V M V
RANDOM_3118 S S V E V I V E V L T Y L V V D M T N S L S A F L A A G D R S F D D T V A S A Q S P G M W N P A Y I V R G R T L
RANDOM_3119 L A E F V E R L A V A S G R L E I G N F P Y N N A D I N V V V A M A N L G D Q S Y A R D H S V L N N G R P Q E G
RANDOM_3120 T N C G N T P G R V V L R T T S I L S K G Q I D W K Q A V Q E L F Q Q D V T V I W K S K I A E Q H P H R A Q Q W

RANDOM_3121 F G M I S R L V P S P Y A G M K E D C Y E S T A T G P V V Q E R D Q F S P N V L N Q D D P F E K F E H A L E T N
RANDOM_3122 S S D V P L P I E N T I H G K F P G Y F L W L S C N T L E K M S V L P S G S L L E E E N K Q S M A V L P A R A L
RANDOM_3123 P L T N S L S P E F Y F G N T W N F D V D M I Y I K L P G E N L P R R E R K P R Q N I I V I I Y V P K S E P V L
RANDOM_3124 E R S Q L F I K K D T A D I L E G S F Q N Y N L N S Y F T E N D I P M F T G C T V P P G R F E I I L S L A P K C
RANDOM_3125 R V L G R C A F V S A V A S A A A G R A N V R L K S D D G L R D V D W F R R N V A L C S A L K T L I L S H D L N
RANDOM_3126 N G S G G G Q E D G T E L T F F A F Y N T V D C H I N K P S K N R G L H R E K M F Y C S L F E T A V A D E S I R
RANDOM_3127 T R V T G L L E V G E H T V P E C T V H A I K L I G P D F T T S G V D Y I L I D D L T Y E S K C E T N L I N A E
RANDOM_3128 C G P D I H D K K T E N L R A E Q G S E L V A R P S G P R G L Q H A P R D C R F K E G R L L W L L F I A Q C T T
RANDOM_3129 I Q P K H K A L V S K C R A D S Y A F Q R N L Q T C V E S L H L V Y A H G T M W K A Y K A R K Q T S S I M D R V
RANDOM_3130 N M I S D K R G K N A G E G G V S V A R P E K T Y R S P T P V G P G R A A S V Q A A K L S N S V T S T W N Q W S
RANDOM_3131 I W N T S N E E H A D S A N C E K D K G G P K Q T L P E K P K G N I K D S P T R D S Q L A Y A W G G A V S N F F
RANDOM_3132 G Q C F R L S S G Y P S C M I V L K S V S G V T G A V Q A P T A A Y R E R N K S T L I T D L R D Q A C Q K N E D
RANDOM_3133 M K Y T G Y I G A T W H P D E N S A E I V G R C K E G I S N Q S L Y V M K D F S A D L E D A V L L D P L H L G P
RANDOM_3134 G L A R I H K T P L R D A A L Y E G Q S G P S C E A A V R A N L A A L N V L K Y L Q S Y V G E D S E G A A L P H
RANDOM_3135 A Q Q C E V H K I T L D T A N G F T D M V D K W D R I E V G P D H Q L L A Q P C D G Q T E H E G Q A H N D P K Q
RANDOM_3136 M P R Y A E F S H D A R G K P Y P L V Y W N V R T C H K H H S R L E V N I C A A R V R E G I E D P Q D S Q M T A
RANDOM_3137 V P S S I P M P P I G R N C I I E N S H G G E S D F G A C C F V L D L G N Q C S D L V T L A E V G E K A G N L S
RANDOM_3138 Q L V I L A S I E Q V T S L K H A L S W V Y G K F C H E K Q F T V P T Q K L D D V N Y C I A L E P L T S L G S W
RANDOM_3139 N G W I A V E V L R I H G H I C D Q S Q S Y L V A L K Y H L V L V V L K V T A I M R A L Y S D S H T G S Q R T Y
RANDOM_3140 N Y E M D R L D V K S K D A V S P M L S P A G Q P D V D F V E N D G A L D I V L A R F G G T T A P R V W A D D E
RANDOM_3141 H H D P A L P F N S G I P D S L N R N E D Y V Y N V Y L P T A T N D T K E P D V A T N I S R M V A D W R N M A Q
RANDOM_3142 S T A L E C G V F M G A Q N F K M R S F W K Y S E L K M T R V A Q R N P N L T A S F V V K H I E L A L E D L F P
RANDOM_3143 E T S D P P A A L A D M D F K C D E D E T Y I A A K A D A S N D E V A S V S N R N I E P P L R E D E T L E R W N
RANDOM_3144 F V L E S S Y E I V M L N T I F S K F A Y E E D N A G N S E T L E E T A E D D P F A F Q N I T T R K F G L Q Q A
RANDOM_3145 D H F P R T D D F L N P S S I C L L G K R T Y K C V R A S R M K N L A E R N A N Q S I Y S L P G G L W F L T I Y
RANDOM_3146 C Y P R A K T P P K D P G F F F V D N L K S F A I A V A E N E S D E N R T R T Y L F G A I L T G Q K N S L C L E
RANDOM_3147 M Q Q K E D S G T D Y G K R I S C V V K P K P F E L L T K V P T L M P D E L D K A D F L T R T N G L D L Q G R G
RANDOM_3148 T K Y C F L V S M L V M T A C K L E L T A T F T Y S K V S E E E V E Y E T K F K M Y N Q V S Q R P I R L F L D Y
RANDOM_3149 D A V P H T R E D R G R L K R P G A T M F L S M N Y T P N S L Y A D T H H H P W E P A P R A D E Q F M E T I F R
RANDOM_3150 I K I I D S V D F I L A D V Q V L T Q S R R F N P T Q G G D S E S W G E E G N A M T K G L S E L G S M I S N Y V
RANDOM_3151 S S D S V A A W G C N P P G E L W S H F S A S V D Q F H V K R A P S F A Y A K L L V A H L L I L R R G V K T T A
RANDOM_3152 Q R W H S S G P Y T R T T E F E W G A G A N R A I L S V Y S M G F N V I Y K G K P L D A P M H F G A D G G F G L
RANDOM_3153 S E D W K L C M L L L K Y V T P D M T R S E I M S P I D N T F T L C A G S S V A V T L D A S M G K S N L E E R L
RANDOM_3154 V S S W Q G G A F V H L I L R G H A E A S Y C R V V A C K L K G P I T E F C L S D S D E Q S V V L L F I R K G T
RANDOM_3155 E E L S R I P S V S A M N I C S L F N S A G N A W L F E D Q Y V V V E D L P K W G G P R P G K V V E L D R C F V
RANDOM_3156 Q N P P P R N R V A A L S I G L E S K N D G Q L N A F P P Q E L M L Q Q K S G M C I T S V N H K I V K R L N S S
RANDOM_3157 L R P L H D V F G I E C F L E E I R E K G I D L D C A F F V L L C S A V S P V S I K T Q M K L K R T L E D V S N
RANDOM_3158 I S V V L T P Y A E L K I F K R K R E N T L T P Q H I L R L D A F L A R R V P D L H Q L Y H I F P A R W L R K Y
RANDOM_3159 I H S S V R L T L S M V G S A E N D V P E K P L L Y Y L T N Y E Y L V T M P C Q C R Y N E T K E M G L C A E D C
RANDOM_3160 K V I Q A L T T M P Y A G F S H G D I A M H G P D S K T K G H I S S L T S L L E G R F L A S H G I K S M D N P P
RANDOM_3161 E L C E G D T I P E L C L A R S A E S T D V V R F K F K Y S S G W A C T G T A D A K W S G N N R S Y T T A F G M
RANDOM_3162 V N L H K G K S I T A Y H Q A I L E N L F S T N R Q C H F E M P P N W I E G D L S P M E L A K T P S T Y T V S Y
RANDOM_3163 L N V T A Q P A F S T K H L A N L G A T A C M S Q K D A R A R N K D Y Q Y P G T N V G A A N V K G D R L S V R P
RANDOM_3164 R E D Q M R V V S S P Q S N R S L P P D I E F R S R D Q S S Q G A I E T A E H N I N I S L I Y I Q F S M N S K A
RANDOM_3165 A E E Y I S C G F N R Y L V L T L D I K G A P G L L D N T L L P A F F I T Y R E N A K T D E G H D K N P Y R V Q
RANDOM_3166 G W R V L N F K T R Y C E Q S S D Y T M S S E P V R F D E P L A G K K K N V G I F A Q P T L P G A D E E N R R N
RANDOM_3167 Y P L C L G A Y Y W G E I I R A Q S N Y M T F K S G Q Y T R L L L Q V V K S S T A M L D F E V C S S D E E G C L
RANDOM_3168 M S D F E V L L K L R V E D A T E T R K E D I G G V L E I A H K G I E L I M N K C K R M T W K W E T R P L G T N
RANDOM_3169 W V T L S Q R S D I T L E G F G L N V P K D F E A A G S G R I V K A G N E F F P Y M V P K D L R V W R E H P L F
RANDOM_3170 F R Y V R T E Q E S P G R F H N G A G L P W P Y W S D N L P F R P F Y M G T G V I R K V V L E I R R T V D R D G
RANDOM_3171 R T G L P G E K G Y S E G L A G Q V F D Y G T P A P G K G P G Q A R D K I D A Q N K L L V G G S L H A S R K T H
RANDOM_3172 F G Q T N L L S K N F I D S P G N M L S Q R S L P P P G L A A G K S T S Q Q E Q Q G V A T K K N Y A V E A I L E
RANDOM_3173 L E Q F M T E C R N F S R L E D L F L L M F E P S Q S K W E L A F I P P R D Y G L S E A N L Q E D D G G Q Q M D
RANDOM_3174 K N L P I F K G Q K L G N S G A V A T G Q D D L Q I H N R K Y V A Q D H Y S G V Q E S L E V L K S R R P G T T R
RANDOM_3175 K S C L L Q R T L P K R I I T N S L A E E L V S P T G A R S T L G H L A S A L T R Q P P Y E I A L Q F W A T S V
RANDOM_3176 I V R G L A S V S Q A V D S P T V L R T V P T L R G G D Q S N N G Q C M L T A A A I P N D Q T I D A I T I R K M
RANDOM_3177 K E N V S L T M L K M G K G W E E R V V E A L L M R F S F L R Q D S T D Y I M P K R H A P A T R V E V E Q T H F
RANDOM_3178 R I V G V S T L R V P S L A N T R H V L D D P M L S R P H S Q G M T L T E S F K R G D L A P G L R I A S S V M K
RANDOM_3179 A G N A A K Y E L D S E T N A S L G I S R A G I P C V A S V Y S I A G R P D G L L D I Q S V P I Q P L S C Y T S
RANDOM_3180 K C V N R Q E E K C V R F S T P I R T T F R G S I T H G K E Q G C Y S T W T L T P Q E F Q G T C F A R C N A R T
RANDOM_3181 S E T Y D E V N N W I K P S A E G L M L E D V K L R I P D F W V G H M G G A R R V S T N D A H E R V E N R A V I
RANDOM_3182 L Y L D A S M D A G K G T L R L A P E A N Q S N D D F N I Q H S V R N L Q V L P G Q V P R E D D N S V T M E P N
RANDOM_3183 Q V A G R G V A E I S S Q A I C D G S L K T L P P S G L P E T L H V S I S G G T S V Q H G I R S L G V L V M Y G
RANDOM_3184 V S I P T T G H S R I T S T L E P I T E K T S A Y M L L F P R G D G I A T V R F N T A G S G A D V P Q L G V P Q
RANDOM_3185 D E L G K R M I N V Y A D L M T P G P I V E E H V T S G I E D G S I Y F T C D R P D G L M G S Q E Y G V F E F D
RANDOM_3186 R L W Y G V N N T P L P N L T G P P T A Y Q V E N I K D N N G R F R A D P N Y T H K A R A K T A L R A D P T M C
RANDOM_3187 A P N T T K E G I F R I L T S Y Q D C H L A H L E T N E G A D V I L V S L A K G Y T N E I L A S H N T T T E Q A
RANDOM_3188 V R Q I Q F E K K E Y S P N L E C F Q E L F R S L R E N I V A I V N A C H E M D R H T R K N D L K K E Y G P N L
RANDOM_3189 V L D E T A D W R Y F D A V N Y Y G D P E S G H P T P D L P S N S Y N V L D R S V L Q A Q G P K E G N E Y L Q A
RANDOM_3190 N E H A K L R I Q F C T N N P V Q I R G P L G I T D T L V R A G G N I S V P Y L L L K I I A L A Q V D A G A G V
RANDOM_3191 D N G K N Y V S D W D K I A Y E L A G C S W L L S E Q S V E K P G K L K W R K S A T I K S I I T S P A Y T Y L E
RANDOM_3192 S G R L N L K Q N W D R D L W D A L L A C I G S Q Q E L D I N Q K C I I A R L L I D R P C N C C G Q D E S V R V
RANDOM_3193 S R I F V E V G L M E N G G G K L N S A G A V I V V D P G K T V S V V K E Q A N V I E E E D S E N Y F N R R S Y
RANDOM_3194 W K L N Q D L I S F E G S H P G E G L D P E R M P L L K L S V H K G R Y A S S R P V K T Y P V S L V L L G M F T
RANDOM_3195 W M D K T K N R T G S G L G A T A H L S S L G L G C V C V V I V Y S L F F G M C L A I P T P F V R N I S L A R P
RANDOM_3196 P N Y K D R C P L V Q D M P D D A E Y N V R A W V S P E M E A V V L D D S G G V A L Q K R Y A D R A E T L D G K
RANDOM_3197 K L V A H S S V N E Y H E G K N L I K T L D L A K D A A R G M E S S N E G D H G P N E A A G E T K I Y A N T S T
RANDOM_3198 I L P Q L L P E G L T K N E K T G V L E L D D D C F S T K D G I A K T P L D L D S E I C I E V K A A E V A R R L
RANDOM_3199 G Q A K E S E C L K I T R D E P W A V V N F E A N N I G N W G D M Q T L E I T N I D R A H S F S P V W L G S S A
RANDOM_3200 Q A G K T F Q Y A V E I P H K T L A L S A W D Q S S H S W Q E N S Y P S Y N N L L H G T T Y L A L H L N P S M G
RANDOM_3201 C H Y P N A D L E H E L F V A V R L P K P G K K D R I R Y A A V G P L A H E G E V L Q F T A Y E V E R A Y F S L
RANDOM_3202 S T S L E V V D L C D G F V D Y I N L N E K S L N L K L D N F A C S K N L Q L C N S A R Q V S L H L K Q T V K F
RANDOM_3203 L E V S N P I E E T P G A D N A G W H S A Q Q A V K A L E V G R L G N L F M M T C R Q I S V E Q K I N K E F A R
RANDOM_3204 L E K L Y E G D Y G R G L Q G G R L S I V V S R L S C G K V V V G C D T I G D R G R Y M P S C A V R G V R S R F
RANDOM_3205 Q T H Y Q Q F R L G Y Q T T I L R P D M G M R K P I G V D L E L L N L L G V I I V S R E N V T I A R P N A C T A
RANDOM_3206 S T D L S S L P F I A R P D L G R P S Q L K N A E E L R Y R R S A Y R L G E L Y T P F L T P T S V S P G P H L Q
RANDOM_3207 F N H E D Y D L T S C N Q R I V T R L Y N N Q T N V G N K A G V K S P P T L E K L K V V N T Q D E I I Q K R V A
RANDOM_3208 A D A G M Q P F V A K K E L L N L L T L S L S F D Y S V H Q W P G E R H T Q R V N L K E P S E P I A G P K T E F
RANDOM_3209 V P E P V V I A K S A I N S T R C K P T L L N K S L N A K L N K P S R H A G C G T A S F L G P G P L D H K A T P
RANDOM_3210 G H K E D S A T L G S L A M V P P N S A A P D F Q W A T V L R Y E T N K F G K K D D M S L I M Q A V P V L T A I
RANDOM_3211 V E R A P G V M A L V D I S N L A G L K K Q S Q E P I S L K D R A Q N A R P V L I K F T C I K I S V L G L M Y T
RANDOM_3212 E C G P I K Y D A P L N K G V C D G M L A P I A S S F A P I S G Q K G T D L T D K C D L Q Q T K L Q V L P V D D
RANDOM_3213 A D T R G Q G L P S Y E V L G R Y S A G E F V V G S H P T A N G D G T D A V S E C M L L I A D I L F Q R K W Q N
RANDOM_3214 T R L Q H L A G L T K P V R R Q G L Y C D N Y H D K L E D F T R Y T L Q A K K V N V G P S K E I V A K A S K I Q
RANDOM_3215 Q M H S D S T G G D L G A H F I E T I T K K T C K Y L L G Y R R E M V V V I S L A T L A F V R N P Q F D G G L N

RANDOM_3216 L S G G S E Q S P E E N T P L H K G L C S Q D P E V I K S S R G F R M A K L G G N Y R N R L K T N I N K Y S K L
RANDOM_3217 T Q G L L S P E M C V S R S M I V G A V Q S I A W N E S K L I K I T E N C L M S I D V Y A N A L D D H D I V A R
RANDOM_3218 N A P I P G L P M V V N R S R Y G T D L E G P K N E L E V E K Q Y I P L T V R G T R I D P Q G I E G K S F I N S
RANDOM_3219 L P F V T C D V G A P C G V I Y A L I R P L T D V I G T V L K S M A F T A D R I C K F S A L G L G E K N S N A S
RANDOM_3220 T E G Y S M Q L P A L L D V G R L S A H I E A F H H T M P Y L K T P I D V M A G F L L G L V F T A D I D R E A K
RANDOM_3221 G S L Q S N F Y Q N H K Q F N Q S R Q M L P A G A T S E A A Q R G L K R G T T C C G T V M V K M F I D V Y S A G
RANDOM_3222 R G G T L M T T S Y K N N Q L S P Q R I F C L L E A S G S P T T K I E P L G T P Q T A V P V K I P D E K L V G A
RANDOM_3223 L I V A L S N C G T L I C L W K S R L R V V H A K S F S N P L Q D S L I G L V E D K D I I E V H I E S G A G S Q
RANDOM_3224 Y D M R K K R L F C K S G A C V L V L Q T D P E R W L G S L I D G T T K R L C F T E Y M H A S S L Q A A G N V Q
RANDOM_3225 K S R G F I G D D V V P G L E A D P Q V A S E D E T D S E A D K V A A P S H S E I K D G H P P D L A G G I D P S
RANDOM_3226 K R I Q V E L F Y L G G S H Q N F F K D V A I R K A T L G A S D F G S Q A R W I A P V I P A V S G P Y S Q G E I
RANDOM_3227 S V L G F F P M Y A A K E S V N G V E Q T N R G T A C I S S D V N V V E K A T K S K R G A L I K L G A Q A N R D
RANDOM_3228 A M Q K K A A S C I R F S V A E A Q F Q F Q S D K F F R C E D R Y N S V V V P F R L M D L V I E E Q F I G V G Q
RANDOM_3229 D I L L L K V Q G S E W L A E W R V N L R T S I E E E L K C G H M S L M L T E G C A L F E L P S G A G S R L T Q
RANDOM_3230 D V G V W D V G I G F M R F I E G S M A N N V V L A M L G E I T I S A R S S D P L F V T P S R K D E V F R A L H
RANDOM_3231 T A Y K R A L G H A V G I I L A E G I Q V N L K K R G T E L P V F G Y R P L Q G S Y A F E V R L V I A T I W I E
RANDOM_3232 V S Q L C T F K G M S M K L F Q D E G I K Q R P Q M R V K Q A L Q A P C A I A T T N L P N R D E S G G A V A L F
RANDOM_3233 D L D G I K A V S E L N H G K V P R R P D H G A P K I S Y T P T S D T T M E N G W N S P S N G R P F L H H T G D
RANDOM_3234 L H I K I F G H V H N Y G R N F A V F F V G T C K E A D S D V Y S P E T F D T H Q A L V E I F P V H T Q L T R P
RANDOM_3235 E D S D R P L F G I D V Q F L L A S K I C V F V D L A A M D L K K Q A E K L A C Y L D N N V I L F R L D T N K Q
RANDOM_3236 Y S G Q W A M V A E T V F I E A L K G D L V V H P T I S T R N G K R C I A T A A Q W C V K P V R Q E E R L V L Q
RANDOM_3237 Q I H I E K D L L N Q S N K I Y G T I L P S V G Y C A T S M L Y L T M K I V V Y F M V V V C G M E P I R I F I G
RANDOM_3238 D V F N G T L G L N E C K G F G K G P K G D E E I E N Q M S G R P G L N L D G S I L K V S M C R T A Y T E W L R
RANDOM_3239 M S K F S P V K D P D P S T P L S S V S Q N P Y I T G L I I F G S L I E A V N S P L E I E D N I P K T C I N I L
RANDOM_3240 S Q D H L V Y E A N F I V L F C P S T L K L K D S P T I L E S H E E Y N E R V D I V F L Q S P C K R E I R T I R
RANDOM_3241 S L I Y A D A V T G I V K P V P D G E E G E E R N V D D S Y K D T R D G F S R S W T P I I Y V Q H N L D Y L Y A
RANDOM_3242 L T P R R V G G V V F D F M K S C G I L F Q G H V F H D A K V L K Q T L V P D P I I N H F V E G D K I I F L M A
RANDOM_3243 E F L E P T Q P A F K G V V P G P G P V E L L R I R L K K Q S L K I A A R Q F L G N L Q P D V K K K W E S K S I
RANDOM_3244 S E L T S S Q P M R R D C P K S E F N S T L N E V G G M G R S F A F T L F S E R T P T G I G G H V I Q L P A Q F
RANDOM_3245 R Y E K W L Q L I S Y Q M K S P E Q M G P Q I M L I E K F N E L G A R A A A D V N L A G N T E A W K R S G E I W
RANDOM_3246 I E T E A A K D A D I G Y D A T E A M D P A D I A I T Q Q S P P T A Q K F G L F Q A P E V Q P T D M Q S N Q S I
RANDOM_3247 C K K P K L T N L E D L M Q D P Q A A K I K F C C T N V N K S A L G V L P T D H N E V E L P F K A R Y N G Y D A
RANDOM_3248 S E E K M A K G C P H D E T K N K L F L A G A I V A C A E N H V S T Q G Y D E E K L K L A L R Q T G L W C L G D
RANDOM_3249 C H C A L P C K L L P C L D S V L I K Y I G L F S L C S L A A G N M V I I R N L G Q N D S Y N D Q I D A P K Q E
RANDOM_3250 D G H T A M P V E L G L K V S V T R L V P A N K Q V Q F S V A P T G L I F I D K I A I Y K K T Q G G G R G A I A
RANDOM_3251 L G P H W V V K F E F A L L R S P P A S S L P N Y I F F N L M Y I V D W N D S P Y F P Q L E Q I N M S I L K S D
RANDOM_3252 V Q K I L M L W S A T I Y T T D D R I G L T V A N S A N A F V E I I S T F N C T S I C W Q D E C P I A L K I T S
RANDOM_3253 G K G S A E M Y W R P S K E N L M N S G I L L G I K E V A Y L V T V P G K V R N K Q P A V D E E D D F E A L L R
RANDOM_3254 F R V L K M L P G G C A V L G V G N K T G E N R E N C I Q E S R P V L V S Y P F M L G T G P F L Y G A G L H Q L
RANDOM_3255 P A I P G I D N L Q R Q E R S S F N R T L N Y I Y M P F A D L R K F N T D R A D V G D T G W N A V A I I R L I E
RANDOM_3256 L I E H T T Q G K V E R Y L Y A I V G V T T Y H A G E W L E R I A L L T H V E H Y N A P P D S A L L P G D Y L L
RANDOM_3257 V D G C P A L Q S N S L D L Y E T S V Q L D A G A E V G L K K A S L D Y L P S F P S V P T N I D S L V Y K L I R
RANDOM_3258 M A R H I Y E G G Y I G K G G S D Y G G D Q V L S L L P H T E I V A A F V A M A A C P D T L D D P Q S A T I E F
RANDOM_3259 N L S N I E P P L H E Y S S T F N G G W R E G S Q A E A M P G K Q F V L N Q H K I T K V G G S Y T N N D G Y V L
RANDOM_3260 Y N L S D I L F P R V N L V E V I G C A P N K C K C Q L R N A A Q P T N D F V Q L S Q T G L A I D V Y L I E G R
RANDOM_3261 R A N V V F V Q M R S R Q F L R D E E Y E I K R H L L K F E I V S Q V N A V R P G Y E G T L M L T D L P Q Q S T
RANDOM_3262 G Y Y L E L K Y P G E K A L V L G E L K M P L S D M I P F T I E K A G R A N P Y N K P S F T G K V Y A S P K S P
RANDOM_3263 N M G K P T K G L A D M D T Q F F D Y A R D Y N S E E A L C L Y W T V T F G L T F N L Q E D A E V D A M A C P G
RANDOM_3264 D L E A T R K V W T N L L L G N Y V F P L D N T P R V Q I R T S Y T Y I Q P L Q K V L T A N I E H V S I E R P V
RANDOM_3265 V E I S S L A G C N W F N V W G N C V S L F E A S V Q F D E G L E S G D L K G Q S L K F A G E C A S R E G V I G
RANDOM_3266 L A S N C F N Y D H F V S A D A S S K T E L T S T L I D V A S R L D L E D G L K K Q E S Q V V Y Y V N I T I T I
RANDOM_3267 F T R I E S L P L D M E E M L A F Y P V A C T P A I M A Q K I W K I Q V G F Q H K E A R V E K V H D C D I V P H
RANDOM_3268 K Q T A A E G H L K N Y I V L M T G V R E Y R E E G Q E G L A A K F R A T I S R K I V G E R L L D T A L N E G T
RANDOM_3269 A T P Y I R A R Y K R Y G Y R I I P T K Y P I M I A G K L P S T M D M A R Y M E N G P N N N L G N M T N C C N V
RANDOM_3270 C G S S N E E P S L E Q I L V D G G D W R G L R K T G A V K A L P K L C N L P A E S F N V W R N V W T D L Y T M
RANDOM_3271 L Y M Q Q F S A V E D G K A S R A R S L L Q V E V L R F E K A S W T R S P A S K Q G V D C K S P Y N R D K I I E
RANDOM_3272 V S N E L L I A Y M G T W N S P E L K Q L E P S D L R V S G E F D E P E N I P L N G A G H K V E E L A R F Y V G
RANDOM_3273 T V R G D P G R E V E G T K A D I G S R D S K D N Q N T A P P G G N S D S I D A V V V S N E C D H A G V L S E N
RANDOM_3274 G I K V P A F R F L G T R S Y L F P R G L I C T T S D D L A Y E A A Q D L C L G H Q I V T D Q R S V E Y L D T P
RANDOM_3275 F P R F E S F S K M K I L K I V A T Q I N K I T G L K H T L M D V A D T Y E K W R Y S A G A G L V L E N P T C I
RANDOM_3276 H W I S P N D N G E D V D G V E L I M V A T S M G S S T L S L M H W I D Y V A L P L H L L L S L C S K A D L A P
RANDOM_3277 A K W K C T N N V A E L P P K A E Y L G F Y L T E F G R N Y E R E D N S Q T A S C E N A L K S N P D G M A M E R
RANDOM_3278 S V T A Y S F Q L E Y A L K L I L S S I A L D E L N V V T W V G Q V P D R V D A Y N A F N K A A Q H Q M G P G I
RANDOM_3279 L L V S V V K R P G K S L L A P V F V F K A L C N D M Q G R F G Q R E R K C V K Q T I T A S F P E V S R G R G V
RANDOM_3280 G G V R Y I V A N P D V P P L N L Y L I Y D A P R R K I G K V G V Y A Q V P W S A D Y P A T A A S M G P D D R T
RANDOM_3281 D E L L M P L V V N N G M L A P T Q D A L R Y V C K F I K K L K S E F C M V L L T I P F C R I K L E A Q D I T E
RANDOM_3282 A D S Q Y V H A A K T L G K I M P S D S I G A G T K V D D L I K Y V K S L E L E A A C S Q D Q F G G V F H W L R
RANDOM_3283 F H L A E K Y V A V N A S G C E N Y N I G F S N K P E Q E D S D G G S P Q L I R S Q W A T F Y N D E V G R L R E
RANDOM_3284 E K Q S E F D P L Q I R L D N P G R V F G G R Y A G G D N N L D D T I A F L M K G Q K D A A H S Q P I I R L Y G
RANDOM_3285 S N L Q N G K G E K G A P A S G H N P D P K Q P E F E Y C W F R D D K R V L K F R N I D G S A G V Y Q S G G T V
RANDOM_3286 D D L T D Y L Y T E A Y A E A S K W Y A K E T L P R I P I I I F Y I R M A T L F K G S V K K A Q T G Q D Q F K E
RANDOM_3287 V D S I N K L V N L P Q S K S A I G T H F L T V I D G C R D R T M A P I P L K V Q Q V S F E E N R S V F A S L C
RANDOM_3288 E T Q A S H R K E G S C M L N A M R G A K L I M V D K T T G D V S A R S H F G S C G V T A R G K Y N R S K V C V
RANDOM_3289 R R R P T L E A Y T T D F P T A A E Q D K P E S N H T K M P D R M A V L E S E S G Q I I G A D T A T E T C D V Q
RANDOM_3290 F F T Y T L T E K M C T E W P S D G R L D E T T T V K Q V L V M S W S I S F L G P G A E L D Q F N L Q A W K L F
RANDOM_3291 C G Q Y A Q E S G N V G T Q A E E G S G L A P R P E G F I N I S C A E R G Q A V N Q G D E P V D G D Q R R V N S
RANDOM_3292 D F A P P G G P S R S I G P K Q V G A P N P G V V T E R E G S M A I C Y Y A Y S W N V E R K K F F A T K Q S N A
RANDOM_3293 R G T L D P G E L G L E N S K I G Y S L F A Y S E W S G N Y Q L F G K S Q C S D K E N A E R S P P I Q H K T F K
RANDOM_3294 D N K V S V H V T E F A G P K W F Y P A L T A V I L E I R K T I L I N V I E R D R E E Q T L Q L I I R D I H G A
RANDOM_3295 M A A V E V Q S G E I P L Y V N A S G S P A N L Y E C L A V Y T S G E F A P I N Q T S R S A G D A Q R T S T S F
RANDOM_3296 L V L T R E P S R N C D L E L A H G T Q E L P H E R V E R Y V S S V L G Y V L T T S L L Q Y G K S W M E A Q Y I
RANDOM_3297 S Y D K L H P P A H C L C L S F E K T K N Q S W A L Q K W F G Y D A H S L P L I I I F S F A S M L L Y E L V D R
RANDOM_3298 V K M L R I E E L L S P M P S L P I Q P G E V L R K V T M K G A E V P T D G L N P G Q K K L S L L K E L T W E I
RANDOM_3299 L M G L S L D D S Q E E L F G S V G Y Y Q E Q L E Q A I A L H H A L T I R K N C N G H S I M P F R D K T N F V Q
RANDOM_3300 A D T P E K L P C L M L V A L G L S K S T E S S L T C N D N L A E L V V D L Q G E L P L G E N R A R L T L I P I
RANDOM_3301 H R E V D F L M S E T G A A D N T Q G I D A H G V A R A R Q D I G K T D R A D T Q K D K E I D G N G E N Q R L S
RANDOM_3302 T Y K Y Q K C P V Y Y L D Q K V R N I G V V L V G T K C I A I H C A S A G A A M A D W V K S N I S D D K M G P A
RANDOM_3303 H E K I Y S A I S K L T S V T S E P T Q W K Q M L F H E P N I Q L S G E K H Y E P Q G P N P P L N G C A I Q T I
RANDOM_3304 E C G N I R Q Y N S D G N V G Q L V F I Y D L T V E I W L R L Y R Q C K N P M P G S L G I M V H T M L A W A L V
RANDOM_3305 W K N V K V S L P N P K S D L R M L G S G G G G N V A W H V A G F D A V E S G D E E N T P F F E W E L V I L L D
RANDOM_3306 Y L P K G E A L Q W G G P K R A F Y L E T D T P Y D S V L P E G H G L V S V T R D F K G D V G V R V S L P G S V
RANDOM_3307 M F T A L E P Q G E Y L A T C K G S G D E L T K P M G D V F I N V L E Y R V K V R F L G C T G L N S G V E G P N
RANDOM_3308 Y A R S N N I D Y E R S Y A L G G C N T D G L Q T K A A A Q G A Q L G Y A H V G L V W L N I Q D R K A D I A M I
RANDOM_3309 A K S G S T E P F R D D R T P L E L K A A T T E R V K P D E G V M G F K V S L P D E K V E I P T E L A V L T D A
RANDOM_3310 F M N E T Q E G T K K V D D E T D G I E H I N L E Q D S K C I D N G D S Y A S M R Q S Q P D F V K E P V Q R S Y

RANDOM_3311 E F S D L N E A A A K N V K M K E Q N K N A V N T G I T L P A D Q T H F G C A I F D P D L Y F K L E A R T A N F
RANDOM_3312 T R A G S S V L S K P L N I H E V T D S A P A K Q S F L E A N L E C P T T F K R I S E R Q T D V T Y A D L T N A
RANDOM_3313 T A R R I I S G A Q S Q E N H G T Q R G Q Q G A M A E V P E E N S M T S E R G L L G L E D L H S L H R A S N Q Y
RANDOM_3314 D H A T A V L V Q Y V R S G K N K T E S T V L L L F A P L L P F R C V S G D S R S R W L R A V G K D E I A A Q H
RANDOM_3315 S S V R Y G T V L R Q I V E I V R T L H S N D E H F N R G L C L L P Q R N V V G S P L P K V F S L G F K H K V G
RANDOM_3316 P L V E A K Y Y K S E P Y A K A F A N R D G D Y S S L I G D T A L T D S W L S V K R D G S C T Q C S I L D E V V
RANDOM_3317 H T S R I D P A L I Q K G D G V R P C Y N G L Q D F L G F V W Q L V G Y V E G R K I I N L M V R Q P E N L P L K
RANDOM_3318 L A K P R E G A L A A A K G Q D T M S Q S A G V V W Q L V A S V T R F G G V L D G G D A Q I D Q K S Q C Q V D A
RANDOM_3319 L P P M C G V K V T H K Y Q L S V T L V Q D T A A Y L E K A L V R D M P K E Q A I D Q S P N K L L N D N T A P Q
RANDOM_3320 L S N A E E F F T F R H P A V F M G L L D P P L A T V I V V G N F V M G K K R S G G L P T T F Q D Q N A S Q H T
RANDOM_3321 M A T K K F I V K S D G S A M Y E V L D W P R A R I K A K S G N P S G E V R T D G A R N P K L S D K N G V G G N
RANDOM_3322 A S L G T P L L K D S N D T G K A G T P L Y A E L R D A S A G L T M G N E E A T S G A Q L L V V N G F R R D A L
RANDOM_3323 N D V K C G Y K S S N I A I A I M R N K L K P S A N P S R E Q S L T N L E Q V S T G E L Q E S T A T I A A A A G
RANDOM_3324 M A S G A S I H S H A M A P T L L H V R G P L L G L A S Y A K R L T A A R D G V D D G T A G P D E L Y M T A D F
RANDOM_3325 G A F I N V M K K L L R A P A I P S L T H V D Y E T L V P F V V V A A C R D D E D R V A D F T K A W A E A D S G
RANDOM_3326 V V T H A A L P V V V A L L K A R Q H W E N K V P Y R P Q M T E A A T W V N R Q P M G E A E R P P K K C A N L P
RANDOM_3327 K V C K M E D R A E A E R E N A A T H R R A A Y K N E A L W N Y L G E R M M R P L T L A G G E R Q V Q N I I P A
RANDOM_3328 A I V G P E S S G G R M D T V P Q V L T L M F P T A G T V E G H Q E T L L L Y L C E T R L P I R H G L F Y R G L
RANDOM_3329 A H P G F L S G K K D D D G V P R V Q P H P R T S G L L W P Y M G K S I G F N L T C L F C V L L S I V K T G E S
RANDOM_3330 Y P L W G W S V Q I A P G Q W I P H H L Y N R K V S N I V L R G M S S R I Y A T I L E S K V V S G E H N A K E I
RANDOM_3331 F D G T H G Y A V S F L P H L M I K P T V V T M K Q E Y Q Y L A G S L I Y G T L A L W N L P L G A V A C D K D S
RANDOM_3332 R H M H K F V H H I K P P I F I Y Q F A W K K L S E L E L I V E E S P G P D D K A V R R A T Q Y M Q V A S N I G
RANDOM_3333 S Y L T P Y A E K P G S D V M S Q D F Q A A M L D A I W V S K K T P D E A H A N S L E P G W D E V T A D P E F G
RANDOM_3334 Y D V F T K S G V D N F N P E A N S E E Q D T K L R S D T C T Y E W I V L G L V D G S W Y D T V L M W H V N H P
RANDOM_3335 D G D V G V D L D Y E F M L M L D F P F Q F G E H Q P R Q E E F G Q G K A T C Y I R K L V S S T S A P F A R F Y
RANDOM_3336 S G P R T T K L G F Y R V S E T K S V C N A E S W P K Q S A V F H D T K M M D L T G T S D R I A L I K C L D V M
RANDOM_3337 A Q K E R Q R R V F D G N A Q L I T T E L T D D P R C Q F T Q A K P R A A V D S Q P V D S K D V T I G D T Q E V
RANDOM_3338 W Y Q E V A A S S S P Q Y N E L L V N S K L L A S K L L N A G R V I A D G E E P A D G V L S G K N R L T G L M T
RANDOM_3339 A E W N A L N A Q I A Y S S D L A Q V T T R K L K S H W C G V G S I D A D G S R D V R L A C Q L Q W A T G S E S
RANDOM_3340 S D E G T A F E K A P F N Y Q T R H P Y L L H I V S N G K I K A Q V G E N S V N S G R F R Q R P T L L W F G P L
RANDOM_3341 T R V N G V L A A E N E A S R C V C H I K I G R M T S G Y P V D S G N N A L S G H T L R S N Q S G E P A T G L L
RANDOM_3342 I C L S S N F V R L N C N S L N A R V A S N Q Q Q D L Q A S C L F P F S G K T G V T K Q L P L G V K V I L A L P
RANDOM_3343 E I L K T W T S I S R S D P F R K L N L A V P K A C I T A A T V A Q I I Y L G H S L F T V R L G A D T G K L T P
RANDOM_3344 A E G V Q M K L I G V S V P D N G E E P T L H K V V I A K V P V M R I V A T E G L K K L K P K K N F S V P I P N
RANDOM_3345 S C S Y M H I N R V L A T D V A T S L P S K T L E T L Y K N G G E A G I A Y C G K V P C Q A K H S A A P A I S E
RANDOM_3346 V G P V S N R V G L T P L C K L A I D M P V G F S Q G T G G I S K F H K C R I I S V T S D R W G C D G L F V T K
RANDOM_3347 T G R E S V R G V L N P I A R S R P V W D L K T E N E G L T A R V G A I V P Q G P A G N I G S N P R S T G P A A
RANDOM_3348 I S A P Q L V S V N D S L A S C D N I V E L G R L A E F Q Q A P S G G R N N S T C C L L F M R E L V P K V I T T
RANDOM_3349 E A D A A S S S I T P L V P A L P G A N L L D K A Y D D I F K A L S I L V K S C V D F T E D L V P C H S L T K G
RANDOM_3350 T I Y D S G P V M V H G E G E K M F N V I L A E G N D K K L T K L F L A E N M P L G A D N M R E I N V L V K R D
RANDOM_3351 I I E D G D K D W V G T Q M A A V D E C A S D A S M S H A V L L I T V E S V E V V L S P A G I K C P G V L L S S
RANDOM_3352 G M A S N A F T T I L G E D L P P T A T G G Y F T Y E L V F A T L L H A T K D S L P Y E S A P Y D V V K S T N D
RANDOM_3353 L C S I S Q A S T T P I W I F K S Q K N V G S L I Y F S W S K G V E C T I G Y L H N K L T V A K E S V R H M S Y
RANDOM_3354 A Q A Y K G S L I Q D L S R L T V R S I R A P A T V R P K P T K R A R L A R R T C T R N G N D A A D S D R A I I
RANDOM_3355 L N A C K L V F S N I R G S K I L E G V V C S K P N A L S M E V E L A N S R M L R E A M T W L D Q A I G P Y H K
RANDOM_3356 L R N L T P V V A G L H D P E S N K Q E T L D K F G V D G M K E T F L F D W L C G C L E L I E A V L Q N H A R T
RANDOM_3357 E S S I G Q I C G M G W G K L S R G L N F I R C S G F Q V G G D L E T D Q S G R K N E M T G H L E R G L C A H L
RANDOM_3358 E S T K C F Q N K T L Q G S Y R S T L S L K M G I H A L S I D I E N V L K K E H A I F A D P H R Q G P P P G Y M
RANDOM_3359 N Q Q S D Q A I Y T I T D Y N E A G E P K G G R A K C R Y A P K P Q C Q T A S S H W W L L P H I L F W M K Y E V
RANDOM_3360 W Q L R K I I W L N A A E G E V E F N L G V I N D L P K K L S M A V V N W G Y S P Y F S N F K Q L H A T R Q L S
RANDOM_3361 S R T A F I G N Q A M L I G T E I A Y V R S E T T G T R S L S L Y G Q N L G E R I G N E P V P V P M Y T C T L A
RANDOM_3362 P A H E D S F K K D A G V D Y E N S T S P F R P S W R L R P T Q V L V R Y P S D A N R Q S C V A R I I D E T T D
RANDOM_3363 Q S A D T L L L A D A S I G E D F L K A K E D K R N L A E L E H P L T P P L P A S G Q V H A I G I Q F G A H N D
RANDOM_3364 T Q T T K R Y D E I Q I R L S I S F F E N M A L L T S S N G V I Y I K Y G A Q A D L L V S Y L S A C M A C S R V
RANDOM_3365 M G E T N S V A D M Y K I F T S Q V M A G V G W P P K K P I G K P G K A A N F R L M D T E M S G A Q N I E L Y L
RANDOM_3366 D Q Q L S A R H I A I L V H L T H N T A G G S S Q E T Q V G Y I Y R S K N L G D Y V E P L D D I E P T V I S E Q
RANDOM_3367 L Q L L P K E V S W T E Q P E R P A A I T I R S D H T R A L V D N E E F S V G L L H F P V T Q E S D Y S F A M E
RANDOM_3368 P T I I T S R Q S R I D Q S E G L H S L E L T S Q N Q P D S R E N K P L L R N I N T G V L N L I P K Q T P K K S
RANDOM_3369 G G S T L P I A N A G L E E A Q P E I L W D S E P Y E K K S A D I D P A I E L S A G P S I Q K S E P K Q M K Q R
RANDOM_3370 N L L V S E L L G G H Q Q P V C L V Y H T L T F S N L Q A G L A L Q H V N V V L A A R K T E T F G G F I N K T F
RANDOM_3371 A G I L N T K S I W P R E A L I S T F V L L D G N K G E I S I Y R C I T R D K Q L I E P M R N N Q L S R Q Q G P
RANDOM_3372 L G K K L Q G Q S A E I T E V M R E G G G A S D P Q M Q V E F C G T A L R V R G A F V L V N A Y S K A S L I L I
RANDOM_3373 G R A S T E P R V V I D K N R D I M D N C V A E K G C Q L T T M A V K R R D I Y T L P G D P S G L A N A N T G A
RANDOM_3374 Q G N C A Q T S P Q I E G R A N I I L L V C S V A V R H I W T Y P T H S E E L N Q K Q W E T G L D I T D F A C P
RANDOM_3375 M S D G K T H P T K T S V T T D G W K I A K G A A T V A M L S V K L A P L M Q Q L L I E T R T P T R F D D N L I
RANDOM_3376 P E Q G G V E V L F E S K V V P E R S K R L F A N S Y A S Q W V Q V Q A Q G N N I N Y N E D E S L S G T T G G R
RANDOM_3377 I K I A S G K I V E M N P G C T D S R G V K P D L E D H S A L F C S L M V K V T M F A G L T L N R V P A R A K I
RANDOM_3378 A V G P E F I K T A L F A G L D F D C I P G S R T A D V W Q A E K T L D L G L K I G Q E D D F T H A L M L P W L
RANDOM_3379 V G S G G N K V L Y V A S P Q K L M N L I T L T F I I F Y W Q I I P R S H E N P S N L G Q L A N R L F G P F G R
RANDOM_3380 N L N R V E T C I E V I E E Q R T E F I G R S P S N M E T L H H H G P T D K I P Q V R A E D G T E Y P D K F D T
RANDOM_3381 P D A K E A F Q R L T E I A G I V V P Q T R L R N I N G A L L I F E K I N Q V P D V I L P D V I E Q L D G F I Q
RANDOM_3382 Y H F N A N Y T V A G I N L R I Q A Y N A A D L S S T G H L L G D S M E F G W T Y V A G T L I D W R P G Y L D A
RANDOM_3383 S Q S R R T K E S S V V G M E A P S L D K R A I W H R D E F T M T L P N Q P V C S S T Q R V N Q F S R E P K E K
RANDOM_3384 E Q N T C I R T S T V V V L K L T S E V L K G I Y P A L E S G G F P N I A D P H S Q K P L K E F V L T N D T R V
RANDOM_3385 L V P M P P S F G C D Y T T Y P E A R H T F K A K S L H Y Q A S T S L A N R P V W A C E Q G S G R L I T D D K E
RANDOM_3386 L S G A R V S G A E V K G P A I L Y G V G R D C L Y G L S V S K E N T G Q C L L A Y T N S R Q Y E Y G E G F E L
RANDOM_3387 E G I C G I R V I K N G E S V M W T A L T I R A L V S K I P L L A A V P M Q A K S G N D L Y E L V F E C A S I Q
RANDOM_3388 T S G N Q A K R S S G I A A W G L T G F T S L H F I A L G I I L W I W A Q S L V A P S R L N M H G A S Q A V S R
RANDOM_3389 A N P V D A T E G V K D F T H P S D I A V A C A S Y K V R S D A A V E F R Q R E K S C F V L R G C I E K P T D R
RANDOM_3390 C A G T L R I G S G E D P R D F R D D K L E H V A V L P L N K E N F Q R V L G D S W G P T R R K Y A G G A N I E
RANDOM_3391 G C I D V R G I V K V V A Y L R T C E L A I D L I N K F A Q C G V A L K E H V S G Y G D A A S D D F T A I N L N
RANDOM_3392 S A D P G D Y T S G N N N E I Q I L P A A S V Q P D A K T R Q V D T D L G G W T Q S E L N V K S L W P S D M Q G
RANDOM_3393 G A D H D P R E I G E G Q A A V Q V K N T L I F R K P T L T E S C V E T A H R T P I V G N I P G S V N A D A A C
RANDOM_3394 L G D L E R L E L V S E V R I D G T D K T G A E Q G K A F P Q T R Y I S W Q A V R D A S N Y N P A T T S A T A F
RANDOM_3395 E V P D V D A F K P V C R S A A L T M N I G G L P T Y N H F L H E A G Y L V T K V T A E L L V Q C I P S Y G D A
RANDOM_3396 P T K E Y F V D I D L K L S L L Y K T L S R T S K V K T L K K F G S I T H S T E T L E T T N K D I G T S R K M Q
RANDOM_3397 D R S Y D P E L I T K T I Q P L N F E L T S I V D A E R Q L A S R C F I G R E I Y L V K Y G K S V G P A L P G G
RANDOM_3398 G G L D R V K S R D H G A Q Y L A C S M I H P L E L D R S S L I E D L R R Q S Y M R T S S I Q K K E A Q G L K R
RANDOM_3399 H L W T N N K G G G A L S Q K F E V N F C A T D K T K V D Q Q Q L P C Q I K I L I L S P Y A K Q V L V G L M S D
RANDOM_3400 R Y W P K Y P Q T G S G L G P P K N A S G S V V F R R I V S L A T D V H N T I D L V T S N F D A S K L G P N Y S
RANDOM_3401 T A A S Q I F G F V T P G Y M L V K T P V G N D E L V G L I F S N M T D T H A T S G Q L V L C D K W H V K L M Q
RANDOM_3402 N E S L R T P E D C S F A C N P W I M L A A G H K Q N V S A Q L L V F A E V H A N D H F V C S N T M W K I L G G
RANDOM_3403 S Q Q V G D Q L E G P A R S T E S R P V M K G E D T L G K K V V V V I M I S Q R E L F S F H P L G E A L E K M S
RANDOM_3404 F K N V F A T R E L Y P M E G T A A E T N I D V A N A S E K A T Y T L L L K V I T L V G F G F R Q G K S S V P W
RANDOM_3405 I G A I D F Y N Y K P I C A D D A A D G S P Y N A D G N R D L Q S R E Q V L F K N P S T E G P Q K A S I R L W V

RANDOM_3406 Q K S A K P T K Q N N S K L L R Y L H L T L D D E K L V F N I T V R I R H K K P Q A M N T N S M G S G A L L D P
RANDOM_3407 W G R P S C K A R N N S Y S S G S N P Y A G P C A E D G E S L S Y K S V A S P K H Q R G K P R C E I G G V P L V
RANDOM_3408 V L W T T I D R Y I I V G A A F H T A K L S D S E T N I L V P A T P E M A T Q E H V E G T K T V R A H V V K L F
RANDOM_3409 N C L G Y S A H L P V I A F E R P S V D T V V E L V S G T S R I A K L M A S A N E D E Q T S A F L G Q H L N L G
RANDOM_3410 C V G V S L E S G I A V E Y L S L I S Q C P L D D F Q H L L I H M T L D Q E K G L Q G T P V K S R P F D M S E T
RANDOM_3411 L S A I A P G T V R V P G L K A H M N Y K A A N N P Y R L N E A E L M S P K D N V P A R A A L F R R R K V V Q R
RANDOM_3412 A A V L N C R I I V A S H N A D D G D S N P V A T S G T F A Q S R A S K E V F A D L M V L S V A V V T S Y E D K
RANDOM_3413 S D D L A R G Q A G E E K A H V G C K D L T F K D A A Y Y N G P S R D E V Y S R E K A L F Q S I K I G V R N E R
RANDOM_3414 K E Q P A N E W A A P P L N P G N A Y Y T D A S G V T R T L L K Q E A N L A S L D L P F A E R A K A K R L G K G
RANDOM_3415 F E S K G K I V T V F I A L A S L Y N K K D S V N V D V A M E P T F D N S I E G V G S C G K V A F I S Q E R A L
RANDOM_3416 E Y I A D A K T V R F K S E R E A A Y Q I E N G T E D H D A T I D S Y K I P I L L S P Q L E R L E P F P F V F E
RANDOM_3417 G L N K P Y T Q G A G E S A G D S E A L T K I D E T K L E E K I H K C H T N R I L G T M R E I S N L K N L S R E
RANDOM_3418 N E P Q R L R L T L D K S G M P M T V Q Y S G M V H L H E P S G L D I D A A A L D Q D A G F D F G N D P R S Q S
RANDOM_3419 D A N D K T L L L E F F L L K K A K C M P K Y Y I P V E K D A E K S E A E I N L L H F K A V N A D L P T E W D A
RANDOM_3420 A A P Q L G L V L V L E E G E V L E V L F V R A V I F Q A S A G F S Q V S Y S A P M H V L R H K V A D E I S I I
RANDOM_3421 K D I S W L G G G E R D A N L T D C N S D Q W K S A M K G G M V Q K T Q V V K D S N N N R G T P A N E E E G R R
RANDOM_3422 L E F E S V F W H E L A T Q T E I N A D K M T K L N V G P K L H D L G S A N D R S A K L N C R L V P H T E G V A
RANDOM_3423 T S C T A G K G K L V E L L R A K E V S S R Y L F T L Y I L S N Q I E N S P L L A I T L L N S P I E D D E T Q S
RANDOM_3424 N V T W A A K L F T K V R Q D Y A G G L N I T I R I I T E F L S K T N I N I I P R K R I V G Y L Q H T L G L C Q
RANDOM_3425 A A E S D N R H I T R V L T S I H P K L L E F V S G V K G P K A G K K N R K F A G P S K T C R I K S L I E V I P
RANDOM_3426 F G D L S V L T T L G D Y Q L P T L V V K S D S L N Y Y V D V D E I V M P G E G A A A A C V A L D E H A E L V D
RANDOM_3427 G F C V K I V T R H E S T L F C I F Q P T E L K L Q I Q I F A P Q A F T Q R N D Q F P F D L T R N E I P G L M G
RANDOM_3428 I A R G L G E S S T I C D D G N N L R D P V A A D L K E Q V A D D L M E T G V G T K C V R A V D V T V R K S P Y
RANDOM_3429 V Y A Q I T L Q K S G H C I D F G E K K G C G I K K K W F G N N V Q E P A K L E L P S L D Y E K L A N S S G Q G
RANDOM_3430 D I L L D A L K P Q L V L L I N T A V R W E L F G E R I K I P P C E G T K I G F D E R D Y A L W I F Y H A S P T
RANDOM_3431 H S D S C G A R T H V H N R S L A S Y S E E Q I N M L D L L G C V F K A Y Q E L A G V I G K D K T L S G Y Y L R
RANDOM_3432 K N V E T V L E T D P L E T D K L S E I C G K I A I K G Q D L C K V S L T L L E W K M Q S K Y T P K F A G E L E
RANDOM_3433 P I M L V A D V S A A T T S K D L L T G D A E E I L F V I V D D P C D Y N L G P F A Y S L W G S K R E S L F L R
RANDOM_3434 K F L V A A Y E R A D I S C T K D M L E F G T P I L D C G Y Y V H R H G L I R S E L C L F M F D L I K A G Y K V
RANDOM_3435 Y K K R I T C R I T E E K K Y A E P Q I N F S R H R P K Q V I I R D L R Q G V A D R N P G D W G I R L R S Y P H
RANDOM_3436 N M D G V F Q T D L T E M S K A L G H V P L K T K W S I P Q M N V G F L S A V L S T E Q D N E A R P G F T C L L
RANDOM_3437 W M Q R P S I I L V A I A R K E K Y I D P K L G R G Q Y V L A V V N N F G Q K E E W I I M M G L L T E K F R G F
RANDOM_3438 L N D Y Y E C M L G G L A R P E L P T T V G F G G T T A V V S Q Q P F P I F A D I H L K K S P L T Y T Y A V Y M
RANDOM_3439 T R Y E V A L D S G Q V R P N M M C R R T W D Y K A Y Y L Q N Q A T F C V K K V P G V F N N T N M G S G Q E A I
RANDOM_3440 W Q K C F K V M L Y K A K G G N Y V A N K P D Y R E F Q T K L A I R T L G K A Q F D K Y Y E S S D P T V R I A H
RANDOM_3441 N Y L A Q D C G F N T S D A D A L V W D L T N Y M I F N Y G K S A H D D F E L K N R H S P Q E N W T T R T G K Q
RANDOM_3442 H S K K M R I E M E Y M L E P W D L Q N T L A S V F G S E G K W C K V L N D V K G V E E Q E D S G D H T L Y P G
RANDOM_3443 T I Q T V V T V P A D V G S Q P R F G T K L Q V L V L K K N I L S K S H N P F A S C H K G S D A L T V D P I E P
RANDOM_3444 A F Y I W A W V R Q S T L S Q A G H A A S S A T T L S T V R D G L E P A G A F K L I A V F K V K E H L T S K M I
RANDOM_3445 P L L N I N A G T I E L H Y Y H A H H L Q L P L A V H V G A D L R V K V L P P V S D K P S Q N S A V C K D G S R
RANDOM_3446 K T G F W A T Y N T G E Q K I S S N N H M E F T K M G Q Y A S E I D G H K G A Q L T M I R D W G P N V D M G K V
RANDOM_3447 L I T T R T D G D V D L D D Y C Q T A S R T N R P K K T E Q E R P E P P T S H C S V Q V F P D H Y L K G I A T S
RANDOM_3448 W D F S G G N V P L T L L F C S F W V Y H I P H F Q N T L H R V M K S L S L F G I R L K S V Q A S A G G S S A K
RANDOM_3449 K L N P N G M G R D Y C C S E V D A Q D H K G D P S F S Q N A E P S G L A S A S S Y V V C A G N I C L I S F F T
RANDOM_3450 S C I N Q L H G R E L P R S K E E T V K P E G V I K I W K G A T R T R T Q K Q K R P K V I M N E E N D V A N N A
RANDOM_3451 V A P Y V C V L T S Y V Q K A K L A I S I L S S S V G K T V K K A I A L E K E L S I I Q P I L V F Q E I V G G D
RANDOM_3452 M Q P G G K L Y A P N S K K E R G I G L V K T L V S R G A G R N A S T V R R E C A P G Q L E M P Q I Q A M I A I
RANDOM_3453 Q E D K Q I R E S Y G V L L L E Q V N W Q L T K F R K K W S V I R Y M K N D P S I Q V A G W Y H D P L E S L T L
RANDOM_3454 G L Y D S A A V E T D N A A K K I Y Y E E H G V G R N T L Q P L E Q S S E K V A R V V S G Y A G Q S N T F V S L
RANDOM_3455 G P A E P T E A S V L T R P D P G S T A A E A I G Q K E P F D R S Y S F P T I Q A Y T G A S L P S I S E V I Q W
RANDOM_3456 G P L Q S L S Q S F R L G S R N T R Y A N P R N V R A R E P P A I E V K N N Q A N T P K T G K H A K S P P G G Q
RANDOM_3457 S C V L L P C K T F D M E L Q S S S N P Q A N K G N R N D S Q A P E N L R G E Y Q A P G L K V S D A I Q L K V F
RANDOM_3458 E T I L C W A I P S Y T A A R E F W T G A S W F L L F V G H S D W I C G N I K C D V Q P Q S L H K V S A R D E S
RANDOM_3459 D D G A L A E W H F E S S T T L L P M A K V A A K Y A G G E L D T Y C G A N T G H L M Q G K S T P L S S Y L D Y
RANDOM_3460 V G S V Y E M D V K L R R D T S E V S S S I Q I N D K S I E F F E E Q V T K S P E R V D S P A E H R A C T F I N
RANDOM_3461 V H I A D S K H L R E F G A R V E N E L T A R G R S C L V A A F D P Y P A A W A S A A G G E I V I G L R L L N T
RANDOM_3462 A S A M A N I T T S Y Y V S K Y R I R P L I P L S A P R K M P K S M N I R A Q L H S M T Y Q Y V P A L I A P K T
RANDOM_3463 N P I K D L T Q P G V I G Y S Q H V P H K T S S F V R Y F Y E F V L C S T D E N L L A D T V L E D Q R A I E S L
RANDOM_3464 G S G G A V W T K T P E R S A S E P K S N L S K E E V S T S L S T I D W G G G G D Q T A E R S D V E R T L G A R
RANDOM_3465 T V N M E C S Y G S P N G R Y L D I V T T H W C L M D L Y Y K A D I R N K A L N G N S V L R L G D N L L T V D F
RANDOM_3466 D F R N I V T Q Q S F I K E G D S R L H F L R H V I E R A T P E S F V D M E C V G L K A R K Q A Y I H S H Y V A
RANDOM_3467 T D L L E A A T G K G R L S R T S H N L G G T N R R W P T A I D T V K G Y S P T L A K I V R E I L A I W G A G E
RANDOM_3468 S Y V T S E A S L K S N R T F G D K A N E L A F K G A V C R L G P Q L H R T P P D R S C F I G S N L P A W V Q L
RANDOM_3469 Q K E I A L G E V F L L V L A A L I N F P F T F P P I W P Q G V S G Q Q K K E A P K A R K Q S T A P R P L L P Y
RANDOM_3470 P H T E G E E D I L I G S D L L Y Q Y T I E R T V E I G A L D R T G T K I E Q Q Y A E A D G R G P A K E D Y M V
RANDOM_3471 L N D V V Y G N W P F K F Q D A S L L V N G E Q S L E I K P T I S E P E K L A H A T F L H L A P A A P I G L E I
RANDOM_3472 S L R T S K F G T M F L K S E G L A W Y D G T G K P S F P T F D T L A K E V E R E R I S P L S L A G D G L S G Y
RANDOM_3473 K S A F G E D C R V D Q N E V A K R G A R F G H K H S N V S P T L L P G E L V T G Q F T L E I T R L D L V M N F
RANDOM_3474 A G K V V G G K D S Q I W I C V E R L S N A G N G G H D L Q L T P F S T Q G L I Y S V P C K K S F D G G E Y W G
RANDOM_3475 S P D I F L G D L S Y Q A S W F E Y F N R N V E F I R K E Y Q P S K C I T N S V R A P I K Y M E E G S E L L L V
RANDOM_3476 T M A E G V P V E L E R K A D G M F A Y P P F P L M A T C E L G I R G R L H S N W R T C V M H F N D E S G T K C
RANDOM_3477 K D V F A E A C G S V Q N V A V R A C P A D D V K S I E Y D N H R A V P N A G D D E T T Q V K Q A I L I M S V G
RANDOM_3478 V N S F S E R V Q T S P K S K P V E G D F G V L K V V H A L I C G K R N D Q Y P D Q E T A A A H W A A T E V M V
RANDOM_3479 F D F L T C N Q L Q V V V F I H T I I A A S D N D C F S V Y L R E G E E R N S V A L I Q N C R Q T Y F G A E R G
RANDOM_3480 R G T W P A S D D T L S R Q L Y P P S L T R K L N N L S S Y N C A V Y I L R Q G A R M I K G I L P V K L E P T C
RANDOM_3481 K L L D K I G G P A H G L S V A E S L P E D V C E F F L R D T D V S C R L Y D Y Q K S R L M S G M T C S A A I L
RANDOM_3482 P P F K N I L M K I H T L G L V N N D G S L S D A C D S T G C A S D L G A G S L N N G N T R T F E T L Y S I D L
RANDOM_3483 S G R E F A G T V K T L G G V V H A F L L L K C L F P F V V N I I G K P V K Q I K N S E Y S K F E A N D L L P R
RANDOM_3484 R I E N I D L L Q T F G P T C V L D L E T R G V V A L P A R D Y E N L N H N A T E A Q M H F K Q S N Y P L C L K
RANDOM_3485 L S N L F I Q R K V R G V T L E H E L V N A D A E F N L R G A G P Q K I Y A D Q I Y A G D G L P G P D G S E S I
RANDOM_3486 D V A G Q R Y Q A S Y S L G L D P Y A K R N F K V T P W Y N T V A I V P E L K F V Y I L G S L Y P P E L V M P G
RANDOM_3487 C R K L S R P I Y G V N L T N R Q S L S R D E T I Y V G Y N S M T P R E S N Q H M V A A G V Y D P T N N G F W R
RANDOM_3488 F T G A Q G D V Y S A V D G A R E E V L S V L K R K G W S P Y K A E R L Q N F L T L S C S E F D P L F R T M R Y
RANDOM_3489 P R P I K F A Q L Y G R Y G K S G T F N E E H L K L A I K R G V I E K I K P Q R W D E S D V F F A L T G A T I S
RANDOM_3490 Q H H K G D A A I M R W K A Q I G R D A G N G H E S K V H A H N G S T S S A S K R P Y Y A I E Y D G S E S I C D
RANDOM_3491 S K L E D V F V A E Q D H C F S K Y D G T N T L T E Y R V T M P N L M N F I E P H V A M F F E Y A H R M V S L Y
RANDOM_3492 N K K S A P S Q M V T E G A V S F A E R R R N L A F R F R W G G L G C S T A G V Q G N V L K C P L D G Q F D P K
RANDOM_3493 S K G D A Q G T A H L N I Q R L P S V D G Q L E S L P D V D L E I A L P S H E T G S S Q I S T D N R L L K S S T
RANDOM_3494 Q H K A K Y N A K R M N Q K A Q G D Y A L E Y K T G E S G L N F F L G F I C A V I T V L I A H V A I D L Q A A S
RANDOM_3495 S S P Q E E K G K I P L I D P S I V H K E Y T L L M C E S D V T A A S L S G E L L T E G W L I P L Q G A A N N L
RANDOM_3496 S T E N S A W A L I L L E S E C S K K Q L A V H S A L R V I Q S A D Q K N N S P Q S S L D S G D Y S P M K N P N
RANDOM_3497 V W G G I N T K S R Q Q H E V R A A P S F E I R D A R A E Y A P N N Y L L V G G A E A S G A K T I H G Y V E P N
RANDOM_3498 S P V A T W C C K V F H T L H F E S P Y A W Q V S G V H T P D C R T F M E P N N S G S H A A P L R T H M T S I E
RANDOM_3499 L L L N A S E W F H N E D G T N G T P P Q S S Q M T D F T C R A D E E P H T K S L K I H A G G S Q L D K P D G S
RANDOM_3500 V C N E V A D S K F T L A C D S D S E R K G V K F R A L R F C Q G T R S I I Y P G H S P L G E K C L I W D Q G

RANDOM_3501 L A K I T E A E C Q H T A P F S P G L V R T T P K L N K T Q Y G G V S T D V M G L V L P T V L R V E L N H T R E
RANDOM_3502 Y E F A I R D K N A D D V G D S L A I T A T P N L E R I S L A Q E V I T A D I Q Q L E A V G Q S S V A L P D K G
RANDOM_3503 H S E H T E R C I T S E I I A L S E Q E S N L L G T D D S K R V E E S Y S I R A V Y A I R A Y T G S F T Q T Y R
RANDOM_3504 V F W S K Q C M F V N L T P L K K D N I N K A L A G E E A V L H I V W I Q L S Q Y F C E S R P R K C N H L S F R
RANDOM_3505 S E L K L S T K G S N S V L R T G S N V H E Y S R T A H G L T T L E N V L L S G L D A A Q P A A H P P K I G K T
RANDOM_3506 C M D A T M L P V A A L E T K N L P S T Y V S Y Q G L G S Q L M Q M E L D F R E M D V P S V V R S E G D T E R L
RANDOM_3507 G H L D G N K G S I P A D I N W K F V T L K V L D F F L Q E A R V V P Q K G A P L H Q L A A Q E I S K A E S N L
RANDOM_3508 N G A G Q E E V K K L Y P G S A D G S V D K T V Q I T Q N S A P C E G D V F E P T G L Y T V K S A T L H F G P L
RANDOM_3509 G D D M I R E N L L A N R L L A G L D C D N C K K L K Q Q Q F Y W E D Y G P I S S K N T H A C G L H S C I T A L
RANDOM_3510 S D L N T L S R S L G P M A S D N T S Q V A D N D H V V K S F E D R L R Q T T D A Y V G G R G S S P G A D R Q K
RANDOM_3511 M L T L H L V T F L E L Q Y C T I T S E V C N A L K G Q D F D P D T I S D R Y T G G P N I G V T S Y E S E K H T
RANDOM_3512 C G R P F D D P F S M N G T A K N L K S A L Q L L L P D L L G D Q L A C Q K K L L L A N Q G V P S T A A K V C T
RANDOM_3513 A L H V P A L S I E K Q R R A S L N F T S K H V F W N K V A G K P K Y L H F K P Q K L G N T A S L R A R R I T P
RANDOM_3514 N L F V G A H S A R T D N I G E N K Y E A K A Y K H G A L L A S A V N V D W V R K H T S D L P W G T Y Y S K G G
RANDOM_3515 G G G A G A M E C G L K E Q G L P L S K T V V R V M L L S P R C Q N Q V E D V W S D I G D N H N V F L T R A P F
RANDOM_3516 V L D Q L N S T S S S V K S P D Q D E T Y D R H L M R A R P T W V L E G A Q R A G V P T E H L V D Q Y L L L G R
RANDOM_3517 F D R V A H I L V D E Y L H R G E E N S C W P D T S T I S D T S L K E L A G V F S Y K M Q E Y R F D V E D S T I
RANDOM_3518 E E D R V N K T T K F S Y A T A S F K I Q A Q L V Q D G E S K S T F K K S T F S S L L G P S R S L N A A Y V S G
RANDOM_3519 E A N Q S A T L N E A R N N G E A A L G S S D I T H G V C D G A N F I E F R T L A Q K D C F K A R S L G D A V A
RANDOM_3520 G H P Y G R F R L V E E G N Q A S E D Y I G N E I Y V K S L G M D R S M Y D I Y L A T N N N G W I S V W K P E S
RANDOM_3521 T D D E W C H H I R V S S G F N G A W E T C Q L R N A S G K R W S E V S R S Q G G E Q I F D V A I D P G V V N P
RANDOM_3522 G E W N M L D M D N G I S S L N S R R D S L R P M V A D K Q D L T T D Q K E I G K Y S S P E L N Y I M L S Q G L
RANDOM_3523 L N K F V M I I S L T S G L E T K M Q T R E V R V D M P Q H A A G K G L F S R A G K D P V E S N G L T R G G D V
RANDOM_3524 G G C Y C L P F T Q V V Y Q D V N I R V F D S F E V A K S L D T L A N M P A T L D L H L L L R A Q D E M R D K S
RANDOM_3525 Y A Q D K E K D D Q A Y G D C Y L R S V N P V V P Q E F N H E S L Q T M R G K N R T L G E C V Y R L H T L T Y S
RANDOM_3526 V N L L N S V D E S L V K L L G R R C N L Q A L S A H V T L P P G L Q F V A G K G F R M L N L M A L E Y D M Y D
RANDOM_3527 D A A G G P P L E E V N I T N T G S N N D G G D V R Y K N N V S A N S E P V D Q G K A D S A N E G G N P T G G I
RANDOM_3528 G T R I S V Q A G M P C M Q S E T D L S Q D R V S P K V F N T R G E Y D P R A E D G G Q E I L A L T I Y L W R H
RANDOM_3529 R G I K P T G L E E F W L I G N V P K F F T S R R P V A Y M S Q P C S Y H R S C S Y K E P S L P F Y Y K L S D E
RANDOM_3530 G T D Q I G F E W I V A G L T I E M K Q Q S C I V T N P D I K R K M T M M Y Q D D V D A G Y N W K E L S D C T P
RANDOM_3531 L A D L K Q V D V L V Q G H P S C F C L L S A V L Y N P N D G Q K E G G T D E L A W Q G Y L K P M I S H P S K V
RANDOM_3532 K R E T L A T E T A D E S E E A I L G V S R D V F P E E D I L Q T K A I S V A S L L C A K R V H L I Q G S S D L
RANDOM_3533 E R T D P E E T D G V P S F P Y V I H G Q D L Q V E K D M T S A R K T L P R Y Q R S I G I P F L L F F G R G P E
RANDOM_3534 Q A S N N D G D V L Y G D Q H V K G S N Q D C T F S D V P E E K H L P L S T R G G G E N P R H G P T N L V N R M
RANDOM_3535 I N L R D N M S P W P D Q T P H G K T N E Y D P V D S C D A D S F I E E Q K D K Q P N P V V N F A D A S K G Q T
RANDOM_3536 T Y Y F H L M E V L T T N L K A Q G K L T A L A A N N C L R G G K T D S L E V F S F V L S L A E E A R S A C I T
RANDOM_3537 D S A I I G P A R G L T D N Y L E A A R L S K G D S C E D F C Q S A T Y L R L T R G T L K Q G E T F L E Y E I C
RANDOM_3538 N L A S L L S K K V S G T L F N V N M H A P L F L V R D I S N L L R Y N Y L K S Q P N H T V Y R Q L I S K I G P
RANDOM_3539 K R F Y A A S Y Q R Q G I R S Q A G R G F M H C I I E S A W L K S L I Q I S L G A M E E G T D I I R Q E C Q L P
RANDOM_3540 M Y G P Q D L E I E N S T L A E M G P A E T G C E A L S A V Y E Y Y G R V F W R N A R L C Q I G T T N A N A P E
RANDOM_3541 K M R A L Y L M P S S Y P I Y E S W I Q F V Y G P L V R S S L K Q S E I A N G D L I H F H P D S A K I A T L L A
RANDOM_3542 R A L C I G C I I C L E P C P Y L P F L T T G I G R Y N I F N V P V P T A A K K V G C V T D E T G W R S E E I R
RANDOM_3543 R K T R E K H T P A I F V G P R A E L D G A T D K D T R P C G N V L D D G P D D Y W V E C Y C I M L L E F Q K P
RANDOM_3544 V A P R S G L L E K K P K F P P P D L R E Q K V G L D R L S N W I I G Q S D A K N T S R N V E N K W N S S F N A
RANDOM_3545 L R S R A A K A G V Y D R I L G D I L S G C L N A G E K T T S K L S W N K Y N Q R G T A Y S Y G N L S Q V I G A
RANDOM_3546 V V K T H L G G E S L P F D M I A R V K M D E T S R T E G P P M A Y P I G R W T C R S K Y V E Y W A T D D I P R
RANDOM_3547 V H Q E P D L V K K G T K A K N G L A I W K G Q S A I E D S Y A I L S F T G T Y E V G L N D L T A P G Y V G S G
RANDOM_3548 V T G T E A D P V V T V L W E D R K V V Q S L P S F L V T D L A E V V I R P P G V G S P R P S Y E Y E H V E A E
RANDOM_3549 G N H E S G C I R L Q K I K V L A A E P D S P I V I S R S I I Y G E A L A K K R N A R T L V A V E Y G V S S A R
RANDOM_3550 T I V L A L P T P Q G G H L V A M D I P G Y M E V I S L G N N L S G F N Y L M T L L Q K D Y S D F R L V F D T K
RANDOM_3551 L I T N G H A Y I P L L K V K P T A L T E K N A E L P K R Q Q S N K T P A S K Y K G V M R A F V V P D L A G K P
RANDOM_3552 S G Q G K I R N K E R N P A E T G S S C A A H G D V D K L N S T V Y P P K P K T N F V Y S E K R T W F L W G L G
RANDOM_3553 V D L M E W W T H N V P M G D N K C N A Q E Q N D D C D K R A E F P P L L S T N F D A P E K L E R N L L I V V C
RANDOM_3554 V Q V Y F L A F A G V H L T I Q N L Q P P Q Q E P K V M R S T A Q F I D L T L R E K T C I N P G Q E P E L Q P P
RANDOM_3555 K S G K P L H L N S V E K Y R T N F L V S Y V L K L M T E R E D V N V G T L L V S S V T R K T S Q G E R D A I G
RANDOM_3556 M F F G G A H K R A A L Y I S V A I K T M V Q D N A A C E S L L F Q G A K G N P R V R N Q K G R Y D I L R L A G
RANDOM_3557 S H Q L D F T H W T V R L E T P L D N K N A A L V E M S L V R Y R S G R Y M Q M A Y L K D R L K K R E S M G K L
RANDOM_3558 P I E D R L G S D S D L E S S A E G N N V K N H I G T I S S G N G F R L N R L L A I A V N S E A Q M L V Q E M D
RANDOM_3559 F T T L G M N D I E S R I S I Q G V I G L A G L A G E I G N A G P N A V M E D I L P I L E V N G A V P A V Q G S
RANDOM_3560 G I F E L T G T V W V S V A W T W T V N A L S I P T F C L D C K V D K G K S S V T F L G F I Q A N K P L A H D V
RANDOM_3561 F F S G P A N K H S D L R G P I F Q A C N E T F F T V G A I Y G Q T K R L L Q L N G V S R D V M E W H C I R T N
RANDOM_3562 A S L E L R N A H Q M K P P Y M M I K K N P P S D T H L G K C S F T I L H R K E P E V T A F V G V Q L M Y F E P
RANDOM_3563 R L P R V Q R H E G G S N L A Q H S N A V D D H L E G Q D K A T V L E L G S E T G T R K D D A C K N L I H Y T I
RANDOM_3564 I E W K T F V A E A N Y T H E R V Y L L E P K E M A L L R A G G L A N E R C S Q L E D W A L L G Y S R L K Q P G
RANDOM_3565 I G E N G Y R W E E S L L V S E P L I A S L S N G R D S V D L N Q G T M I L H L K V V A R D V I V Q T G P S A H
RANDOM_3566 Y V P P A G A G D V G N E A L A E F E T A A E L S G T V V G A N V V V L A G E D K L L I P T V H E F T E P L T V
RANDOM_3567 L V E D Y P R A H T A G A D S V I F L L M L C G N V G T V H R I T E A I R L L L T F L L S S A T D V T L T S D E
RANDOM_3568 A A Q V H L V R S S R R L I M G S K V K Y N N D G A R G P H D R C S L E G H G T F G Q A L M P P R I E Y S L G H
RANDOM_3569 K M F S I P E A L L K E D L E V S S E C R I G A S I G T G I L I Q A G T I T Y N V P Y C N V I N V D A R I R K T
RANDOM_3570 H V Q E N V R D P L E T N I I N D H D A P A L R N R L E M L C C A G M I M Q I P V R G T Q N K V I Q L S Q Y I Q
RANDOM_3571 Y L F R A M Q P T L P H F N A Q S T G R P G A K L G L R R T S S L G E N V S A K G T K L N P E V H Y D Y L T W V
RANDOM_3572 R L I L N K N R Q R C N E R R L V Q Q S G T L H D R V L S M C V Y L K G G A R Y R S K F L D E E L E I L G S T V
RANDOM_3573 L E N D F M P M S M R Y W V N R G P K Y Q M N D P Y L T G K P S D T L D S I M D D I N Q N S T K V E L Q T I R A
RANDOM_3574 N Y L D R E Y V G D M F V W H K I N L S P R S E Q L G S I F V Y L E V I F N A A G V D V G L S D A A P I I P E Q
RANDOM_3575 A M A K D G R P G A R P I E S Q S D A R I W Y F W D P N N E A D P G L I L L M D E F V I E E T E P D Q F F L N Y
RANDOM_3576 S P G E P K R E L S C Q L V G V N Q P Y K Y A I Q S A A V P C L G P P F G N L H E G S D E N N N R I P A P G G L
RANDOM_3577 C V A K S K W P K E Q I L G R D Q V E E L S P C L R I I L I S R E Q G S T V S L T R V A K G G K C D K V K V V L
RANDOM_3578 L P V H G P N T S V L Q K D R K R G D H L H D A S Q W Y I S L A T R F S E G I Y P W R V N S L G L I M F P Q W K
RANDOM_3579 L A H L K A K V L P L R E R E L S C V K Q A L A C P R K L R T K A L S T N T E L P A C L W T V L I K F S I E G S
RANDOM_3580 G T R P V I C T D E I S P C W F D G I S P R I K K I T L S S A Y S G Q I R G V F H M F G S Q N P C E C L L T Q R
RANDOM_3581 W G G T F A F G L E A N T R E N I K D S Y S G I S K L D T M Q P E L A E V V E G F N T R D I I V F T S Q N K N T
RANDOM_3582 K N R V Q S E L S E E T R M G L P V S R E A D D S C L K V E R Q C D G S V E V F V E H S N E K E E P L A L K P Q
RANDOM_3583 Q G G S L R K E D Q L G D D L R S P L T S L V D T G F T E G A H L T L A K S Q F D T L K A D F T T E H D Q S G G
RANDOM_3584 V F S P L T F R R L F Q G H S A M C T D I T T H M G V Y K A K V L V A H E Y T V P I A E S N V L L C L A E I G W
RANDOM_3585 E L G L E G R L G P E D S A L Q I V G H Q D S H W V D A R A N G L V Y M T G S Y P E W D C G G V I P T L C A P G
RANDOM_3586 Y V Q N K T K I K P K E G Y R G V L S L P I T V G R R E Q P L I S D R E P V G K R A G H F S S P T E E V Y L V F
RANDOM_3587 K L I L R D E V G R A L I P G R Q P S N R G F H L S D E Y L L K S L V S M A D V R Q F L D F K E L A K P G D A V
RANDOM_3588 L T S R T M G K K W V V P D Y L T P L I K F N V R G Q W A T V K F G Q G D R C K I V E F A E T K I R F K L G D L
RANDOM_3589 A S G H I G T I M V Q T L K I C R N K A K V Y R E C L E R R H L G Q Q D G I S P V L L I V F Y N G F S E M P S L
RANDOM_3590 N V D F Q R K F T S D H N L R L P G D I L E L K Y Q G K Y L R G L V R L L V R I T V L K A P L L K K K K S K D R
RANDOM_3591 F P Q I A S S S A T Y E H L K W T G C K N K G P S Q K S R K V F A Q D N N E L L F L T I V S T E H S L L L H K G
RANDOM_3592 R T T P E E L E L N E S M L L D H I G T F L A D K K P E D L A M P T L G K R K N L E A P F P S A F A D L E F Q S
RANDOM_3593 S T S A P I D Q I L G A A D V A S H L I G F A M A S F V N E F I Q E H V Y K R N A E D L L H N V Y T L K I Q N T
RANDOM_3594 D T C G K V P S C K S L I A L N V H G P A I H D I N Q S C T R L V P L G E W G G R Y S S K T M I N E V F A A A G
RANDOM_3595 H S S G H Q L Q F D E G V K W Q F S D E S R I E A G A T Q E P E T L P D T G V N L C I P G Q V A Q S N Y Q T G D

RANDOM_3596 C T Q G P R T Q R G N S A G H V T D A Y L W T F T K A A S R Q G G D S L E G G Q R W L W S T V I D N N E Y K P A
RANDOM_3597 S V W A C A N V V Y Q L G G P L N A F G K D R L A Y E L Y F V R R V R E S V G P V A V T R V D A A G F T L V T S
RANDOM_3598 A D V W L L M V M F A E I I E F V M L A A V S S P P T W V D D M K P A Q V D D K M T L R R L F C A V L T V V G I
RANDOM_3599 A R W V L R S K A P D L A G D T K I I N N F A W K C I T G R K N D R L S M A L L A P Y L N N V A P N K K F T S L
RANDOM_3600 Y M A S T F S I L L L A Q N I Q Y S A T S D G Q K T W V Q Y H T V G L N D M N T M G T K G Y P V K K S T L E P G
RANDOM_3601 Q S S T M P V T S G K L R S R D I D L P G T T A K E I F N Q F Y A N T L V K N S Y G H E A C G G K Q Y L Q T T F
RANDOM_3602 Q I H A N L T G L M Q N I K Y L Y V L V I Y S L H S G N H G L S Q W I G T H N N K D D T S V Y D S A T E G M T W
RANDOM_3603 T G A V A T I K P G E W G E T E I I A G G T F T P L V P S A A A K D K A N E D L V C M N K A F S S G P F E Q I S
RANDOM_3604 W Q A V I G I F Q T H P P D A K Y R T I V E C V R S K S E R N I S E V L F N E T Q D R G L N S N M G P L Y S K I
RANDOM_3605 L M D E G V I E G F T K T T S N A N G T K Q P T D L L K L F V S I K H A D M Q I N L S P G L E E G Q I R E F D G
RANDOM_3606 E P E N Y G A M R F G P T L I E T E K F A E F T R V S T N N V V Y T T T S A M A R L S P G V D I L E L G D K Q M
RANDOM_3607 S L V P V S K L I V E D I A L D H P C R E W L A G H G Y S H A V I P A I K G I T S Y T E M H Y R Y A Y P E T S L
RANDOM_3608 G A L G E M N D N D V W A I A L R K T W G K W M Q A M P R D V P D D D K L V S V Q A P S G G D H A A E Q V F H R
RANDOM_3609 R H F M T S S P R W V G G E Q Y T T N D G P F R L R E Q A K G N S A V Q G V G H R A E Y P P L D C D Y G F V W H
RANDOM_3610 L N L V K K E E E E L Y S A I C I M Q D L E D S C V G Y A R V N I Q E M P K L G S S N E S V K A P M R R F F A L
RANDOM_3611 N L L L V I T Y I L T P N L G S I I H V G T M P E E T F Q A D V F R A A R M V F L L S Y V K E E L I S D Q K D A
RANDOM_3612 D E L N I S D V L L E P Q K L G S Q D N L T F L V I R T F W W S E Q R P I E P G I S E L V I Y L F Q C K Y E Q E
RANDOM_3613 C L W A A C P G E T T P V D Y G T A T V S I F E D D D L A V Q L E Q S L F L G M D L V I I V L I P K P S I E R V
RANDOM_3614 I I S I F K W V K S A R I A K D A F I P M W G I Y I P P A G K E S K D E D S R K P L K A F G F R L V A Y A N A A
RANDOM_3615 F R D K K K T L K N D L Q S R F S L T D F G A G T E A A L P G R Y E P G W L L P N V A P S V G G M S G Y K K Q G
RANDOM_3616 K F L N F R E E N W T K E V Y G A G E V L R Q G E Q I G D G G G V A K L L Q E F N W N P S L N G Q L D L P G P E
RANDOM_3617 L K T G L G Q V V G R L I G P V N Q N L G A S L F E Q Q L G L E E A D D L S H G A G M F A K H R F A A S P G F V
RANDOM_3618 V L R M S H N A D N A T A G E A A C E C W D S F Y R L L V Y F I M V L R R V E R I A E V V F D W A M V Y N L G E
RANDOM_3619 G L D S I S W P T T R P S T L S Y C L N K N C L F I Q L E D T L R K S E Y R D A V Q G G N S P A G E L L G L I I
RANDOM_3620 P A G Y T K Y A L G S A I P N S F S G R M R R P R L Q P Q L E N L K E R V N A F L S P A D S Y C R T L A R C S K
RANDOM_3621 I G F D Q A S C S S L E P V S S E L A Y R V D V P E M S G V Y M A P G N G A W R I N A D I P L Q A S F M H D N S
RANDOM_3622 W Y G P T D A S A L G P F S N R R F Q K S S S P R A R T Q Y C F Y Y Y S T G I V N A K R F L S L E F T A N K H Y
RANDOM_3623 I Y T N P L L S S K E G T E S R N V K Q W F I D L Y M Y G Y N L D S K H Y Y S D E H Y D F E L M V A L D L R Q A
RANDOM_3624 P K K A Q G T E G T I E V I L D G C Y Y T S V R A A Q D R V P S Y E I A L P R S P G K E C C D V P L C T Q V L Q
RANDOM_3625 A C K A V V D N A Y D L K F G R L F R R L K P R S L K R I T I R Y S V D R P H R K R E A E Q V V I Q D G F N N F
RANDOM_3626 G S P R I Q S G Y S D A K K N A M Q S D P L T I Y A A N V D S A D K A D L D G Q C D T R A A G S N L L E R L C A
RANDOM_3627 D L L P Q S L I Q M S H F R A W L V L T D A L R S D S S A A T A G Q V Y K E V K T L L F E L I L L I L K D V F D
RANDOM_3628 H W F A T H I A G A I Y Q D Y W H T G S A Y P I K R G A P L P V E I V P D F D E E R G A E G S N F G V K F V C E
RANDOM_3629 L E A K G L T A K R I I M A T L Y Y V A K R I L L M S A R V V E L A E Q E V C E S T T S F H P S V D A T D L V F
RANDOM_3630 T G I D L G D E I M G L R V P R P V K R R F Y I A V T C Q I D D L K F G C Q T P F K R W I T M S Y K K Q G A S P
RANDOM_3631 Q A L I A L E L R Q F K G N S L G P L I V E Q S V Q L G D W G R L Y H R V W E G I G A G A P E L A I K T D R D V
RANDOM_3632 V N K G A V I I K E I I E A G L N P D Y G I E L Q G L Q R K G I I C W F D N D L F E Q L K P I V N V F T P S S S
RANDOM_3633 K S Y L R K I P T A K I L K D G L H R E L K V W E Q K S D S V L A Q L F G E P M L V T P G W A Q V Q G A C I S R
RANDOM_3634 F K S N L Q H P G T D Y S S Q P V N R V G A T I A G A A G L R E A F A G G Y S A A G L R W E H V R D P R K Q G H
RANDOM_3635 L T D V K K H D V K E K T Y S F A H E E G L L L A E K C I C K S N C I T S V P C S L R H Q E R I L G Q L P Y P L
RANDOM_3636 K I S Q L M P F N Q T D D E T T N L T L V E M N S K G R C F T T A V P K V F G D A E Q E G Q F T N E A P L V E E
RANDOM_3637 E Y N S S G V F H I L K R S Q E V V D W A R R A H T T P L L R P M L C W S Y P A Q A I P P K P S L C E R Y L Q G
RANDOM_3638 Q A A I K S F P Y L C G A T V L V M V S A V T C K D R D L G Q Y E S N R S L P I T A Q L T I L Y P A R M S K M D
RANDOM_3639 Y D V R A L S A K E P G L A D K R R A T N W V L K N I Q S D E D A S L E W I R G S L V F P Q F K Q S G S Y G P I
RANDOM_3640 W M Q G L A F D G Q S A T S E G E E A A I R N A A D V L S F D F A F T K E L T R N C T R R L R F D L K K K N I K
RANDOM_3641 T A D C A R P G N A N K H N R A Q M N P L S A L L S N T K I L S V Q V I L A T S L F V I P D R A L E Q T K V T G
RANDOM_3642 S V A M P Y T E M T L Q C S F T S S A F K Q Y M G V L A H Y E P K G W P K E V S K P L R K L V E V G D E A L A E
RANDOM_3643 A S L P S E V K V I L L S M K P Y G A R A E G Q F G E Q R D F V C N S Y V S V N F A A Y I V R P A P I K Y G F G
RANDOM_3644 E K E T N N E P C D R A V S L V F N M A V D G P V A S P S L M T R C W D P W E G K H G S G A D M I R G V N S L G
RANDOM_3645 D T A T Q N E I Q R V T F E L F P H F H L T K P C F E V R V K E E A I T Q R A K V F K G L V Y E K A I N P A H P
RANDOM_3646 D G S F S L Q Q V L K K P P R R I Q I G L V H I N V R N R D P S Q L P A P V E M Q L T V N Y T K M P Q Y F L D I
RANDOM_3647 T M P R I K F G H P R K G C I Q A D P A M R R Y K F I R D W T S I S V W A S R K N L P L V S V D D V K E N S N Q
RANDOM_3648 V N A E I F Q E I V Q L E R A R C G V S E T Q T E K I G T G V A E E K N C G I E A P E P S F L R G L F S K L E E
RANDOM_3649 Q E K G Q R N K W T K I G T I L G H F I P D V V L I L K K K S S T D F F N E L K D K L L T V R A G G K K V L A Y
RANDOM_3650 Q V E N Y L Q T Q K V E H A Q N L E A N V N P S A M F D V D A G L D L K L E W N A L D A E R G P K E S Q R V D S
RANDOM_3651 V I E V H A N G T H R E D V R T D N F S P V R Q V S S G H I L V T G L S F E W I L L V N A A P F M A N Q E T W D
RANDOM_3652 I A P S L I Y I T A C Q Q M P I C L E N S E P A S S V G H G I I I S A N A A L C G S D D L D E L T C L V D F P R
RANDOM_3653 P N R Q Y K L L D I L V C T D G R Q V N K N Q I R D I S H L N G L W L Q K F P S K S A R L F Q H A Y K G D H L V
RANDOM_3654 N A V S G Y P I D G Q L M G A F D V N G K S F F S R M I S S I I G L Y M E F S Q I A G V G Q W N S N E G E S Q S
RANDOM_3655 Q G V R A S G G S H A E C A T S G H A S I R S T M L L S L E A I V H R L G T K L K G M D P L Q L K L G E A G F W
RANDOM_3656 N G P S K Y F K S D L G D S E S G K D C F V G C E R E A P A V S F S L L L N T R F F L G V P D N P A T W T T K A
RANDOM_3657 F L H Q L N Q V T D R T Q R Q R Y M K R R K P G G E A M G A P A C N A Y F E Q L P V E G S K G A S P M T T Y D W
RANDOM_3658 P A H Y P E R H L V P A W Q A N G T G E Q K S D P V E G V E W S T P P Q G V L F S G G L L I C V T A T N V T K Q
RANDOM_3659 A T F G F L I D V D L S K M G G L N G K P N G A Y E Y Q A D I L P V G I P F I T R Y V T G I G I E Q M L P R A D
RANDOM_3660 F V I L Q L W A G G D A D M E T T D V S N A V G Y K K K A M E G A R Q S C K G M P G L N I M F M E K V P R P E S
RANDOM_3661 S R F N T W K L L E L S G A S E P I V I P I G G F R D A K G L F G R E F S A V W Q G T T P L I A I L G F T P D W
RANDOM_3662 I K F F N L N N V E S Y L P L T I F A G A I L P G V E G K E D G A I P Q A P Q K E L A Q N L T T R T M I A P F L
RANDOM_3663 F T N Q Q G G L A T V E K L F F G N E E H M V M G R S Y L S S A P V L M R T G F S H V D S D S Y F L F N A H L S
RANDOM_3664 K E P H E S P K S R G T R Y Q V A S V G V G D Q L L L L S N L A K F G D P I S Y G E M P K Y N V A Q P Q D I G S
RANDOM_3665 D E A L I F P G S S Q K T V S E D N P T G Y P K T V L Y H F F L A F G I S S I S N T G D K T P K A G Y A E S A L
RANDOM_3666 N T Q Q S R N C T S K E F L V A S N F T P D Y S K G A D L P N H E V E K K F D D E S L D D L A W G G R L L A L Y
RANDOM_3667 V P P K R L L S E I H A A T R N D D H F D Y V K H K P A M P S T V T G S H T R N I Y F D R A N G S F Q R E D V F
RANDOM_3668 L E E F E L Y V L G N G L K S S F L E V P D L S F I L F S Y K E Y N G S A P W C T A D D E I M T V T S L F G L H
RANDOM_3669 G K P N G M Q I R E E G I V D D G S G N Q S T V L C C N L A M T E S G S L P F L L R K R G G L A K T I S K L P A
RANDOM_3670 F L K D W P R A V G L D S R D L P A A K T D S V P M K V D L I T Y I T D V A A C P G P W L M V A R T V A C Y F S
RANDOM_3671 G S T A I V D L T D K N I P V P Q V S N P Q A H E S N E L D N I G V Q G G I E R V N E H A A T Y L V L D V V R Q
RANDOM_3672 S L S T F G E E F L P H A H D N K V H A H D T E E V D P G F H K M A G A L L K Y L H T S K I R T A M H W T I R V
RANDOM_3673 I I D A I L S Q L Q D F I N L I G G V P P D Q R L Y Q Q Q L A K F Y Q H N V G Q S P E P A G S Q P I N S S P I P
RANDOM_3674 M D A N S L E R K H N N K Y N G I G K I P I T G V Y M V E K N H H Y Q W N A H I G E A A T K F E S A F P V E R V
RANDOM_3675 L Q D D E M K V S Y C F A D I K G A P D A L G D E F S C P T G V R Q F L I L G M K E A D R K N E V Q R S P E K E
RANDOM_3676 P T S A T P F A V S S P E Q E E F S T Y G I I G P T K L Y P S E C P D V I V K F I Q I Y S L V N A Y E N V S V L
RANDOM_3677 N L E T T T D V L E S E S A L Q A K Q I C S K I E R R V I G V E A S I R G M Q S I S C C M A G S H A L A P T Y C
RANDOM_3678 V W Y D T N T E N S V Q L N M K S G G R A D K S K E T L R A A I L W S P D G A E E L D T Y S D N I Q I V S V K T
RANDOM_3679 Q I P V L R L L R T D N S S R N G I L M N V L C F A V I V N G N K H T E G F A D Q N V Q T Y A E E S T S Y S K P
RANDOM_3680 A D T I E V L A S N N L R R S Q A V V R L R F E E T L M A D M C Q L D N L Y S E A Y E G T G S H Q P G D K H H L
RANDOM_3681 H I I E V Y Y P V E R L S E H A L K V V F D G C S F D R V M G P K A P Q Y M D G N H R S V A H N T S Q E Q Y F A
RANDOM_3682 Y S E N V L C G D L S N R K T L A G E G N P V A W I I N A I D P H A A Y S K I E G R V L C D E A I T D R A G L D
RANDOM_3683 G A R I S S A P S I H A D N L V L P D S A A S L I D I W L P T K L N T A Q G V S L G N T Q W T G T V S V G K W C
RANDOM_3684 Q F G N D Q F E F Y D D I G I G A E V K L F F D G V E V A P S V W C F Y V G A D R S V G R L D V A A R P P S V A
RANDOM_3685 P L D Q F S P G W G K D K G L I A V Q F G S Q E H G R K V D K E E R P I R V T V I N G S A C A A P K G I E A A K
RANDOM_3686 D G A G H S D V S S D V V I K A S G E F Q A E T G A W S E K E I R Y L K Y A F S D V G P P R G G E S G A Y K K S
RANDOM_3687 T P P E E K S V I H L K V F K R R T S F L P A L G M S A P T V I T T Q P P G I Y S T D M T D E Q T V I G G R N D
RANDOM_3688 G A N K W R S S L I F C P D H L K S P E V F R R T R R C G T M V D D P I M C V F R E L K S E G A T S G P T L E A
RANDOM_3689 L W V R L C A K K A F K L D D A L K T S A R L E R N D H K C L T G C G D F I W S I V R T V L L Y A L L A T N Y P
RANDOM_3690 I K R T W T L A G K G K L V K S I F A P I H S S L A Q L F M R N S L L N E W K I S G Y L A T N A A P I S V P R P

RANDOM_3691 T R E N L L T V N K T L L V T V A R I F G L I F Y E C V G P R L E S S E D S G I P A V L S T E T D V K L R Q S K
RANDOM_3692 I R F C Q I E G L H K K I D K L D C P K N A N G D Y E V A V G Y Q T Y I G N R A V G A P K L I N Q C S G H D T A
RANDOM_3693 R S L L P K N A I N A E S P Q A P D S I M E E D M P Q F K D C L V E V E L P C Y L W F R R A D E N L I N M L S N
RANDOM_3694 Q A L A V A V S A L F R I Y I H G D V E R I G I I Y A L L A D K P I A G E F E G L S A S A Y N G F Q M T V E F V
RANDOM_3695 A E A L Q G E L P E T A D K Q C L G G F K D A D H T K L D P T R P K V S P L F T R S T D N Q I E M D S S A L R Y
RANDOM_3696 S V T L V E P F T A S D I N A G H P N N K S M N V Y D V L D A L Y A G E E S W N G T S A R C L Q I R L E V G W V
RANDOM_3697 A R K H N T P A Y A L T A Y V C N Q D Q R T E D L R T M V Y S E Q C G G N N Q N I T I I K L E N I G N P I K V N
RANDOM_3698 P I F E P Y P N M T R R Y P P V F P N S A S K K F Q E K P I S R Q R N Y S Q E P L E N L K A R L M Q N C R M R G
RANDOM_3699 G A C T E S V V T K P S D E S C L H P K D G N N A E Q Q A T A A P E N Q D A H L S N E I K T R M G M A I Q A T S
RANDOM_3700 S Y L S S H A P D A L W A R G V G G E A M P Q L P D A T R L M R L G S E V G Y R I V H P I A C G G D I Y Q N A R
RANDOM_3701 K Y Y F A L P G G L E D A E S S G T E P D D V A N P A Y V E Y D Y L L N Q T P N Y I G I K C E F G G Y C T L E Q
RANDOM_3702 F Q G R S G D L S P H V M N E M D F L R N G Q W H P E K T E Q R V N A M V W L V A G R K I E T I D R K Q L G S T
RANDOM_3703 Q G R L N G V Q G I R Y S T K T V S D F F F T A D Y P M N E P N I R D A A D I D A A D M D R G A Q S M S Q L T S
RANDOM_3704 D S E H G D L V T W I F R L P H H Q R I T M A R E K Q D D R A F S H K Y R K A G N T V K T K P L S V D Q V R P V
RANDOM_3705 N R A H G H V P T L F R A S H E S S A E F T W A V E F L L N T V E S D K R G P A G S A E R F L N D D D L P I G I
RANDOM_3706 T P P P I N F V G C A G V L V S S V Q L K I R L A R D E A D G Q P G N E I F A R R E T R I V G A M A E F P L S R
RANDOM_3707 N P M L T L K V Q V G N I A P L W G S G D G H A L L F S D K A V A I F I A A A A N Q Q A S R S R C V K G S D D Q
RANDOM_3708 A Q L S Y H C K G Q P S Q G T Q E I E E L L K Q M P L S Q T P N E I G L V K E N T G D R Q S I S D P D G G H R Q
RANDOM_3709 E C L V R H S P A Y Q W V D K L S S K P P A S A R M F I T T Q K T I N G A R D I D N Y K T Q V P C L A A L Q T D
RANDOM_3710 A Y I A N K D G Y A T N Y S V H K T E I N A S D E S N I N H N L E G D T L E R P L S E D S S G S K L N D N L M V
RANDOM_3711 Y V Y P A I P S R K E G H D Y T T S S Q S A G K V T T G I R A R A L Q Q D S F A R T V A V E R S L K K K E R Y M
RANDOM_3712 T R G N V E A H P F P N Q S I F R M D D A Q P E D A L Q P K Y V M T E N L V Q P N M A Y A C G D I H P K I C K L
RANDOM_3713 S Y G P V L L G D M L S D I A C R P I E D H A W S V I L C T P Q W E T V D Q D H E E Y K R L K P E Y S V L N T V
RANDOM_3714 V G D I S I L L E E G E P R E E K P Q I C R P K G G T G R F Q N Q D W L Q A S P A F E P L Q L P E L V A K E T P
RANDOM_3715 T G E C N T S Q R N Y I M S D Y D I C P D Q Y D R V L L Q F L E M G H M R I L K Q L Y H P I V V K S L V F I Q A
RANDOM_3716 P G A S E D E A R F A T Q R D D D G S S L N H N L M T A L V F R L H K R T R V W R M N A H I K R R S M Y L E Y N
RANDOM_3717 T I G Q S D H V G G E R C T D P M T V H D E G T S I H P S D D G V N R P R Y H S K Q L T Q V S T G I N Q G G Y R
RANDOM_3718 A L Q Q E K L A Y T A S T K I V S E S D Q P G S G K A D R F T D R D D G T I I G E K E T Y Q V N G S E R A L E N
RANDOM_3719 G G I S Y I L D V A Q E D M N L Q S S K A I R G A L L A Q N V F Q A P P T K N R K E E R G F G L K D Y G L A F D
RANDOM_3720 P G D R A N L H S P Q V S G R L S A N K A S D F G W I G P Q N S K P D F V M S T R H D K W E L R L L I T S V F P
RANDOM_3721 N G T K P G M L S P Q V L T I A V S H A A Y L I F E P R H I S D R N L S N R S K F L L F S N F V S K I N E L L H
RANDOM_3722 V N V T F C I C S P Y A V C P A S D F I R A L A M N G A F A P G N T D Q F F R G E H S I V R N L D N I G L I K L
RANDOM_3723 T D K L S F G L C A L R K K Y P S R A R A A K A T V K H I V P N C Y R F A H S L R H E Q A E L Q T V H I D T E V
RANDOM_3724 E Q P S D Y S L T I L V M G I T S P G T C P E Q V V V G D R A L I V C A Y L C L D I K R T G Q G T D G I K K L S
RANDOM_3725 T G R F L L P E E F L T N A C T I A V V L L L M T L E A V T I K S I E A V S D F A I K S C T E G A I V L V A L I
RANDOM_3726 K E A E R P A G Q P W L S K S D R N Q P Y L K V E G C S T F S S S V I L A A A L T T G L L W P N N E F L Y T L P
RANDOM_3727 I S F K S H N M Y K S E Y R Q K V Y Q A G F K A K G E I L V I E V N A A E R S R S L A H D A P D I M A R V N T Y
RANDOM_3728 K L G I Y L Q T M T D D Q F V A T T D E H D G R S G I S T T N C V S T I K Y K I V P V P R A L R I I E Q D M S V
RANDOM_3729 I K E A A L I M K K R V M P A V A V G K A S R T N K A E L H L N D F A V V L R Q A K G R V A C V A L V V S S S G
RANDOM_3730 P V L N S G P R W I D I E D Y R A L A G I H L K W A M Q T G K L W L P A I P R I D W E I E V I L A Q P R L Y L M
RANDOM_3731 F A N A A P L H K G G S P R E K T S G K V S R L I E R S R A T S G F F P V H P C P V C E D K E A A V Q I T S L H
RANDOM_3732 I A S T G T G K G I R F A A H L R K S K A P F E M H S G D R G A V V A V L G Q L Q M L D L T I T R G I A F A W Y
RANDOM_3733 K N K A D I P T G L K C D E C I G Q V W A T I M T Q Q G L G G R P P K A Y K D C F L M F W L N A K V E P N A L V
RANDOM_3734 I S E P L S P G L V S G T W S F E F I S C L R G R F N Q L I N L V T R L L T K L I D P E I G L R V A D A G F E T
RANDOM_3735 L D A A Q E F V Q P P D L E L G E V Q G V Q S P R L V E E E G A Y V T K Q D L E R A E K D V S I P P V Y P E V L
RANDOM_3736 F Q E A L K V N S A V A R P S A R L F D D E V M A N W T P Y P V I E E M V T L L Y G F T Q S S I N G A R G E Q R
RANDOM_3737 S E H I A L K A G K Q E R T N Y L V L R E D N A E T E K C N R F Y D V A V Q K V E D R M S P S N S K H G G R W N
RANDOM_3738 Y D T I P E K N T E I V S E I S Y A R S E A G P L N G D E R T R E S N Q L V Y L M I E W N C Q L G N Q N G F F E
RANDOM_3739 V N T R L Q I F R P S L L C K A R H S S G D P I G E G S P E E Q S A K S D L N E A A T A H L P Q V Y M E L Y M G
RANDOM_3740 A N N N V P T H S L R L T G C H F T T G T E L P R P M A D L E S R P S F P I T W L A R D V A E F Q H V Q T M V I
RANDOM_3741 C G T D Y T T I L T K N G S A A R Y R I D P S H K F N R I L C G R T R N S E L G G K I K Y K D L T C K E F R L K
RANDOM_3742 F N A D S L G I E D F E M T L G H D H S E I E R S M S S Y K Q K D F P L E E S K F V T V A G K M I A I Q L S D A
RANDOM_3743 L K V W G G T Q I W K T K T T D R A F N T G V C P L E Y E T N E F R A L L M V F A S F N R V M I E F P A L G L H
RANDOM_3744 E V D V T P N K T L M T P L N Y A A I V E H I T S R P S A N Q N D L A E F S P G R Y A F K A K G F I M I A F I S
RANDOM_3745 S L R G F N A P G A S V T S Q K G A Y G L G T R I P E M I D L R W P E M S G K D T P S G A N L Q S L P D Y I A A
RANDOM_3746 R R K V V P P G D Q D L Q I F S G G V L P R M Q K G G L L S F R L N S A A S W T A E D N F G G S T D D E G Y K T
RANDOM_3747 P E I V E M G R V G A E P R E Y A C I S L E G N H R K G K D L A K A W Q G K Y S A K Y Q T D F R G P V V A E R H
RANDOM_3748 C V L V A P Q E V G D L R V E E G M S Q T A K L F A Y F K Q V R Y G P C P S G R H V P T V D T A E S E S L T V P
RANDOM_3749 M R H Q Y D S R P A D E Y L A Y K T K L V T L Y S L R T Q V L C F C P V A N F K V G T Y F V S V A D L E R L A Q
RANDOM_3750 A N L I E K L N S K D I I R T A Q E P S L S I M W G P R G N N M L Y L S S V V L Q L S Q K E I V Q Q L L G T E E
RANDOM_3751 D Y H I E G L P R A D C A G P P V A R D R M K W D R K R P V G L G S G L L A K K Q Q D G N E F D L L R A T P F Y
RANDOM_3752 K L L G N G L S P W W G L S A A G E D N L M L I L L N E A T R P A D D T T R D F G Y K E E V K W D Y R R V G G S
RANDOM_3753 E G E L A S C A A D N Q Y L S K Q W W A F V P E V H A V A T S I D F V V R I K E L K A R T N I D A A G N L P T T
RANDOM_3754 N G H D S L T K M A Q P E Y R A Q A R I A E R D W D Q D L V G A N D G Q L R P V R D E D D L G Q A K T D V R P A
RANDOM_3755 T W S Y C V F Y G A S K Q I C G L N T V K I A L Q D E K I R M Q F G E C S V I V H V N A A A F L P F T W P Q K V
RANDOM_3756 L E L T Q R G T N N Y K F Q E A R K K T L M A N H H Q E G L G T Q E D I A T Q L G W A D G L K I Q I I C A I V R
RANDOM_3757 W K A S Y I G Y D K S R V Y A G P S I I S F S M D L P N K V T W E S F A A S A F P S T R P G L N V E A K C E I R
RANDOM_3758 N F L L S V R A N D Y Y R P T G N Q A G V L A D G V R L A E K D L L W L M L P P L D L V S S A I I C P D V S K N
RANDOM_3759 H S S W Y Y F Y S D F A N I S E T E K E S C H E Q P E T L R A H E T L A R E A S L A V P V A G P K L M T D L E S
RANDOM_3760 Y S L L G R K V L P P E G F V G S M T D L T F K A S T Y G S A S M T G W E L P G R K Y G G E C E L K F C A N F K
RANDOM_3761 F M F A T H L V D P H S C L K G V V N S C G F T R E E T H S Y L Q A I K I H M M Y L V L P T E K H H V D D S F G
RANDOM_3762 N V D V T T G T D D G T F Q A H I I N P K I P G S A A K N R M H C P A K S R P S N T E Y Q W P T L G K N W Y L A
RANDOM_3763 V S V D Y P G N W C P H I Y S G M A I A M F P A K V D E I S R V N L D N R L V V V C D R E G F N G Y S A M Q A M
RANDOM_3764 P V P K P N G A I A A T A H K A E P G S N R G H F N S Y S V R A G R D G H V G E C Q V S L K G D S A S E T S L Y
RANDOM_3765 G L P S A A P W Q Q C E A L Y K Q L V V H V D K G E T S P L L P L Q S I W L K E I L G L N A V K L Q T H G I H V
RANDOM_3766 D S C T K P E Q T K P G N W G K P K D G L L G C I Q F M R L Y A S S L S R P P P S V G L V Q R N T Q K Q I P S N
RANDOM_3767 L I G I N Y I F A I T A N D S Q G V G H V Q N V W Q A H L P M D C K E K S D A G A I R C A T Q G V T A R G D R Q
RANDOM_3768 F S N S Q G S L P P V N Q T L N A K S P N P M G P V N D R I P L Y P F Y D F E E L M L A E A H A A K H L G S A S
RANDOM_3769 E V D M V T S G D G R H S Q Q L V A V T R L H A A A V W G N P L E E Q A R S Y D Y G R L S Y K K Y E D L D L D N
RANDOM_3770 V S L P V N D D N L Q K P D N D P A N R S G L S V W K P T K A L I T F A R S G G G F I H S T T K S I F L A R V A
RANDOM_3771 E K E W S Q R H T T Y T N A V R D L Y S D E E V G F P E D D K E Y A V E K Y I N H A Y K G F A T T S K G R G R T
RANDOM_3772 K V T V A K D S I K G K Q L V A G Y E F R C I Y T S Q R L L E P P A S A A A T M S V P G G N P V A P L A I G A I
RANDOM_3773 E I N A D F V F L G E V W G E D R K P L D N T L K Q E Y N P S L S L V D G H D S F T E F H Q Q P L Y E P E V K R
RANDOM_3774 A S D L Q I V A R L A E P G V R Q L C S E K Q D K V T E A S Y D N C D D L K D F M D E Y V H L L P A S S F H G A
RANDOM_3775 G D D N K A N T T K I T P T A Q E L G L L K A F D M G K A L V L A T V S P G S L V K R T P M G D T R F W E F A V
RANDOM_3776 I K S N R K A A Q N A P N E P L Q S Q L V N G E P L D M S T K N F E C K S P E G D N S V I G L Q C R L I S G E A
RANDOM_3777 V N G N S G K K A A Q A G F L L D A K M K G M A G Q E Q L G V Y S S P Y D L S F Y S T M E I I R N D R K Y K W T
RANDOM_3778 I S F N E N A V G G K M T A L E G E A S I N T S T G Q Y V Q V N K A D G A L Y C A S L G S F E W V F P L W Y G P
RANDOM_3779 S P D K A G K S A A G N I L W M E S E K A R V E A L G T I L I S V T G S V Q C H I V D G S G G E H G L A K I A L
RANDOM_3780 P Q I P A G V V D V I A L T A K A R K R F P H F E Q S G R N R A L L L I S L I G T G L Y T A R H L L R V R H S G
RANDOM_3781 S D G Q S V N D A F D L V T M R Q G A R I E Y L N A K V V N D V V D G L G T D N I T G L P Y A N E F V V R L T C
RANDOM_3782 T F L L R Y T S L A S A F N K E N F L A G P L V N D R Q V D S E K V N T E E A A K T E Q L F V S A I I Q V R L E
RANDOM_3783 K T A D D T G A G Q E N L R D T C V E S Q D E L E P G I L P R N P V S V G I N M S S Q F V I V G G K Y A F D L V
RANDOM_3784 D Y V T Y E D N A D L E T C G F Q W E W S V I D A P G E T T E E L K Y K D G H E S L K D I I L I L N T D S I T P
RANDOM_3785 T C N A F H A L H I K Y T Y G T Y R H R F R L Y R T S S D P L V K G E P N P D L T G A E Y I G E P Q I P L P G A

RANDOM_3786 S G N Q H K N L E S N D Y P M G K A H I K T K Y Y L C V K V P L V G V I D E D N R D E S F P N A A T P K S K K P
RANDOM_3787 R D Q K G L P P G F G T R E I K N R R I M V G W L L K R F T G Q L Y V K D A I L V R S K V V S S G K S Q L V D Y
RANDOM_3788 S N Q D H A A P I I V G L A M H V P R L N G I A I A L D A T D D D V A L D E P T R Q S N I N I E K K R D W T Q G
RANDOM_3789 P I G F R K S N V V C A L G S E E R F S A S L T F F R K N N Q F R L E D L A Q L R G V Q I T A F S P K Q M R A L
RANDOM_3790 M N T A Q T A A S S K I W E D L A E D L F K R M L P A L S V K Y E E Q W G G T V D E D Q V H G L L D T I L L P F
RANDOM_3791 W E A L G Y I T E D I N H S S A A Q Q N I S P E R T E V H A S V G F C W R D L T A Y G E D R W V E K P K T A E V
RANDOM_3792 G L N H S S T T S G Q H M G T E T K R S D T Y R V F S W Y G Q F I K A A R H D E G S F M G F E I Q A K R S Q P S
RANDOM_3793 N S Y Y R D A I D G E I L K V R Y T P H F N G Q K F E K T S P G G L L S K N T W G R A A M T R L N N I L F L G P
RANDOM_3794 L H V M G L D I A K L S R V S H I L N T N Y Q L K Q S T G D G P A K E P V E L T T P I R L N G V D N T M T H S S
RANDOM_3795 V D P I T K T I H D R T G L D V C R A P Q A K K L A N E P I L T T P C A E R L W C A E T F A D Y L N L V P W V K
RANDOM_3796 E P V P I Q I D L G E K D E G G A C L N S T T D A V R M L Q T A F G T E G Y V R I S I S T R G N S E T N S C E A
RANDOM_3797 I R N N E A L E A G S N N L D L I N V R A I G K W C S S Q Q S E V L A R D A Y L T E M P A A K L S G K M L G V Y
RANDOM_3798 F C Q P G G N I S Y T G W P E R L K S V I G H P F A N G K W A P V P R S I T I N S V G G K T Q D S P H N E I K Y
RANDOM_3799 T C R D A V K P D I A W A L V F G L L M A Y I R L N L S A G D T G R R R M R A R A R L R L V A R P Y H A V I I L
RANDOM_3800 S G E E V D E E E L K F A Q C K T L E H E L Y G M M P S Q P F N K I G G I A V T R I S S P K M K R E E H H N G A
RANDOM_3801 R G S T T H V G D A A L M S S W T A E V L V I A S C S A Y K K V D I A H P I H K I P V L G A A V S L G S G T K S
RANDOM_3802 Q L G V G G F F V A S L Q P A K G G A I Y M E P V I K I Y K L E P V I R L T K K V G D T P Q T G R I N A G E N G
RANDOM_3803 K R R V E T R K V G E N F K D K S L Y K N S L V A A F I I I E L L A M I N V K G R N A N G I G Q K F S D R D N Q
RANDOM_3804 L A H Q S N Q L N T A Q N P D A L S N S I Y C K A G S R N D G V S Q L L E A A I V K S F F D L V E M E H D T V P
RANDOM_3805 W P D Q L K M L D N D E M R Q P T A A P L G F L G R E V P A D A R I L L L G D A D N S D C K G C T E K T K K L I
RANDOM_3806 N P F V L E H Q R R F N V S V A F M T P A S C K R E R D I N G G F R G L R G L L I L Y S K N G S K N R Y P G V E
RANDOM_3807 P R K I D L M E L I F A L G P N I M E M F F P V A F G V P E V G I V S A L L A K E S A I H R V A P L Y I I P G S
RANDOM_3808 G T G H V G T S A P Y Y Y Q V E D K H A T Q R D S T N C K I I A T A W I Q L E T Q S I S K T N H K Q E I P D Q T
RANDOM_3809 D R R T K I P L G H G S A V L A N D V D R D I R A P V K F Y V T R A V V F I R F V F M Q N F D E Q L V F L R R T
RANDOM_3810 E F L L E L G P S C I A I I G I L T Q N G N N I F F P V V E S K R H K T Y K L L K K Q F I A V S S Q K M C M I G
RANDOM_3811 S K A Y D E S Y N V S A E Y L D C H V V A R K H L D L P A R K D S L A L T N V N Q Q G S S M L L R V Q G G S L M
RANDOM_3812 F N Q N P Q A Q D P K D N V D Q V L A Y Q W E A D P M T S H A L P E Q G R D R S Y F R D S D A F Q I P S A V R A
RANDOM_3813 Q E S Q E D V G P I S F P D Q V Y I P I H L R K E S A V R E G R R V N W L A D T L Q H P I I I L L P D I E R S I
RANDOM_3814 H G V S T A K M R H F T Y L I L H E V G A W Y M M G T L S E N L A G V F Q N E Y E E S S N H N N A T E P G M L W
RANDOM_3815 K S Q W G T G R G I Q N Y R E S P S I H V V K D S V P T G K D T C D V L L Y K K Y D P L T W V K I K Y L V V A F
RANDOM_3816 A G A T T R E S D N W V L V G S G N W V D L F K A N T I Q L P V S L D D P L N G L S A E K V P L E G F S C G S Y
RANDOM_3817 K M A S N R E Y H D E G W G L G F V G A V E I G D H A Y K G G S P G F Y A V V E G G E R M I F D D E S G T S D I
RANDOM_3818 D I R G P T W A T L K Q L F L N N L I D S E T S K D G M T T K D M D K P I G V W I P A Q Q G E N S Q C A I H G V
RANDOM_3819 I Q Y N K H Y L S K L S I L V P A P G L T C P E N G R K W P D E L Q G L H A L E L F T P G P F Q S G L Y A S L S
RANDOM_3820 G L P M I A S S F I V S E V S L Y R G D L T Q T Y A N I Q P I L P M Y S R I K P Q W T R P T N D P F T G L L L P
RANDOM_3821 L P V L L D T M N Y N G S D Q S L N A K Q S Q L Q A A I S G W D Y I E A Y T G P G D R L K Y K L L S P R A V T G
RANDOM_3822 V H L M K S L E M K N Y D C C G K T P D K G G S S L N E P T Q A D K S V T T G P K R T S I E L E R V D Q N L Q M
RANDOM_3823 D A G T D Q Y G L E A P T G L G L T I A I K H V F S S I E G A S T E P L A P Y V G T L K R F G R A C S K A D R L
RANDOM_3824 K G T V Q L A I R V D Y A I E Y L R H A C Y A T N M D G G D W C V V D A L G F Q T F I L G L D G L G P T V N P L
RANDOM_3825 V L E T V G A L S T S W A T D L V T G G L F A K G K V D T Y D R S A F L F L E S K R Y G V D E F V N F M T E S P
RANDOM_3826 E E A E L K T G L E K F I M D S E F G H F T S R V T R L H D N T K T A Y T A N L C P Q H R T K Y L L T I K F R S
RANDOM_3827 L L V A A K T I Y K D P P V L Y E E L I P A R P N L A N N E L L S T P T E N V V L Q F S E S Q D I L K R G V S N
RANDOM_3828 L D C R A A C D T A G R L P A S Y S Y A H R F S L N A Q N L Q S D L A R S F S L F Q A R P N C A I I I D A A T E
RANDOM_3829 G C I S I V Y R Y K P G D A N T L S Q V Y K F K P R V T G D P H L F G C R G A G G T G A G E D I E S P T H M G Q
RANDOM_3830 R N E E L D R L N A Q I K V P Q Y D I L Q G I G L N Q L L C G Q R V R G G S Y L S G E V C S Y P I D I S L A N H
RANDOM_3831 G R A L N L S L I Q N I P R A Y V K C Y R D I E M S D Y V K A T G F S T Y K A R R D E Q R A R H C K V T V D L G
RANDOM_3832 S L I K N A R G D K I S V I L T N A D H A P K D R T T G R L G L G T L P S D S Q S F I E P G K S Y N I G G R M T
RANDOM_3833 G P F M D S P S I P P S R W G M E V V E V W F M A A I R S S E S M G L V T Y G S T I I K V V F V L V Q Q N Y W E
RANDOM_3834 D L C G T R N T L D R H G F I I R L Y K L L E E N P P D H F F G E E F S V N K R D Y G E E N S L Q S L P P A K V
RANDOM_3835 R E F G I K P A S A F P F L A E F L V R Q Q S L G K L L R L C M S G P L W V K N E T E Q F S V D A E V S A K V Q
RANDOM_3836 V P N K G D E V V V G A D F V T N P G Q P P R F A V G V H A F P I P A L E T Q L Q N P D A P E K L K R S R E Y G
RANDOM_3837 K R A N R V L L L P T C L R A H R P F E K M S F A E G M G A P H K L F P R P S Y I E F Q R R S Y V N R S N M F F
RANDOM_3838 G S P V N P S D F T A F L L S P V V K E N A M D F K F F K G K P L R K I C S E N C S W V F I E A C V K E Y S F Y
RANDOM_3839 W V E F D Y D V G K L R N K F D F G I A P G D A N M C G I V F Y V I K G A F K R V R T G V Q G I V N S L L S L G
RANDOM_3840 L G S A E K A S F I H K E V R S E S H S K D C N L C L R V S I R T V K P A T P E V Q K V Y K K A Q E T L L G D N
RANDOM_3841 E G R G N M G D F D L D I L I V G Y T D M K S A S R E V I G L M L V I Y A L T G E S G Y D P A E N N D V P Y G L
RANDOM_3842 Y Y R A L E E V Y W R L D L G I Y D D S I G H G H A A L R P E T L L E S H Y G T L I H A S D P Y G F Y S K D G G
RANDOM_3843 S G N D G D W V T W E P V L S G A M E K P V L A S L D A F P T T S H P L G A D W H K D Y C G T Y L K F A L F A D
RANDOM_3844 S P V P P V I K V R N R I V I T V V A G T Q S Y G V R G A Y N C V K I S H G D T S V F S D P C A Y F G E S G G I
RANDOM_3845 L S Y E V K L S R G Y V V Q Q A P A M V P D I D S F V E Y K T V S W H R I E E S A G G P K A S G K F P F S N P E
RANDOM_3846 F E S C H A A V A Q S Q R P Q V P K D I L L D N Y M V P N A L S E L A D Y G L D Q G A F G L W E M A A L D L K L
RANDOM_3847 S G D L L G Y G G L L A Q E P Q S Y R N S G S P Y D A A R S V A D A S S L Q R K A V N G V F Q N V F E I I R L T
RANDOM_3848 H W R N T V F E V V A I A K A V D D E M S D D H G G R D I C C T A K D D L Q L L A L D R T V P G S F L A F L A M
RANDOM_3849 R D N T E P S D E F D G K V L R N R C Q K V R P P L R L E H K H T I A N L F Y L S A V R Q G E Y H A V D L D L A
RANDOM_3850 H D K P A K D V D C A E R I Y G G A E M F K W V R N R G E K T E Q V L Q Y R M R T V G G H E K W K E N H T L K E
RANDOM_3851 Y Q M P S G D V T T K K A C G R E K I W I R L L D R F G T A T S M N Y K E E A L E T A F D Q W Q L M I A S K D G
RANDOM_3852 K V F V L L P M T H R Y V T V Y T N D D K S N R G T K T L M L K S V L G P P L R G I N G L D V T R Q T S H Y V L
RANDOM_3853 R A E S L K S Q T V A A F L V G A K M T F H L Q A R R K E S F E L S I Q Y R S C L L C R S E S N G S A K A I N C
RANDOM_3854 E K Y G Y F V L S T G M C P R H I G K E Q L A I R P C E L T K N I G L K A A V G F E A L G H D K C V L A P T F N
RANDOM_3855 K N G N G S T H F A C M R T T L R E K S A Q D G V R G L Q P E Y Q L S W E P L S G D C E G V I S K I A S D I Q D
RANDOM_3856 D I M L R G G N G A G L T L S C M C T D P I P C P A V D M L V I E K N A I G G W L L N Q A C P N E I L T D E K I
RANDOM_3857 M V G G D L V L H A R R V D R E G G N K H S N R L F S M Y V I G M G H F E T T F I M A Q L A Y D N S E G A N G V
RANDOM_3858 M Y I Q W F T D K E A I L R Q W A S P I M S Q H M E I S C F C S F Q G V I R C T I N V I D D S L S L V S G K R V
RANDOM_3859 D P Y D F F G Q G R N F Q Y C S P G S I P L F L A Y F I V L I D I Y D H V D V K E R R T V E Y P D C L T R T D D
RANDOM_3860 I R P P M L P C R G L G C L C V F H Q Q V M S L A E D G W D L D H G A F A T V Q A G P K K T H S E T E Y P Y K I
RANDOM_3861 M L P Y P L E S E P V R C D L H Y K I I K L T T I E N S N W W P T F E C L E E P K K F K N K P L G C L I L A I I
RANDOM_3862 D V H C V K A G L F F C F N V L W N T A P G S D W I G V M G Q R T K R H P S Q A P H S E L S V V L P E I G N T G
RANDOM_3863 Y A I A L S T S T L L L E G A Q V A E Q T K F T H R I N S A L E A G A S G C A A V P G T L I E E E K D K D A S K
RANDOM_3864 M D G N G P D R M L L H Q Y V L E A L A D S Y S A D C K M K H A R R G G S S G A G Y P M H A V P I H T L K K P V
RANDOM_3865 M P D S V G R Q F K S V T Q N G F S C G E K L D Q R G N G S G L T K E G L C N L I L V D E L V L D K S F V T P L
RANDOM_3866 T S N M A M Y I L R M F F L L Y N N Y R Y K A V L S P D N T D I K L N L K G K V K D I V K Y Q T E Y T V A Q L R
RANDOM_3867 P Y D K M K A A M A T G L E S D K F M T H E K E K K V T A A S I Q R T R G Y V K G C N S E P N V G L A P P F K T
RANDOM_3868 G M H C L G K G I Q R E G F E S D A N G G K D A L L G E F G D N T T K R E K T N I Q P L V G L W V K T S E K R S
RANDOM_3869 I D T G K V L H M I K D R V S D L I N H A F R D V P H Q A P G Y K L T L L I L T A S A C T R A A V L S Q L E E L
RANDOM_3870 V M Q W K T P S K S L K G G T S E Q P L S E L K L T V D A F E A I S C E L S G T R G S K P A D S L G D E V E I I
RANDOM_3871 P P K I E E Q V G T S S D D H F H E T A K F A P P F P V K D G T E V H D K L N L F V E L D T T A R K G N R K Q A
RANDOM_3872 E Q I R I A K V P T A Q P T L Y Y Y G K E K Y E G I A F Q G L R E W Y L G N I E S K K I V Q V Q E D D T Q G Q G
RANDOM_3873 M S K W I A A H G S D K G S A K S W G N I A V Y V T I F N F L E K L K L V K H P Y A K P L Y I Q S G V P C I L K
RANDOM_3874 N A V Q G T S I G S T F P F K D K T N V P R E W P Y I T E A I S K E Y S D F R R N R E N P L G Q A G Y E S F P R
RANDOM_3875 L E T K E E K K F L D Y R N L V V P L R C A Y T K Y R C P N R D E L G V I S Q T K P N E N P G T R R K T T A G L
RANDOM_3876 K L V E S G A K R L L L P Q M D G K G N Y D R I Y Y L H Q R R L A S F S A E F C E G N P G C S T V L I A R P S L
RANDOM_3877 S G Q L D T A V M Y S V L L K E P M V A A P A M T K G E M E Q N Q N N K I K Q L G C A I Y R G N T P G I D P H A
RANDOM_3878 S T I L A L V E I D T H T A T T T Y T R I R D E P H S P M D I A C S S A D V R S E R V D S M L L A G E L E A E D
RANDOM_3879 P E G Q L V R Q V N W S T S S G G Y I T S P F T I F S R P E E V E N R L I K Y L P V A S P R G K A I D Q Y M D M
RANDOM_3880 K Q V N P R S P T E A E D A R Y T I L I F Y S P N F G K A V K V M T C L R Q P L V R G Q A S R S V Y E A T I I G

RANDOM_3881 G L F Y S V L K K V T K T I D P C Q P I E L S R A W A F N E T H A G C L D A I G A T L W I G A G C I L K G S R L
RANDOM_3882 L P P I K S L D S T N K Y E W A R A W W S I K I S I V F K V I K N I A S P W D S R L S V K E D D E A R G G A A E
RANDOM_3883 A L L K F H E N T N C V A L H S L F G R Y S A L K Y P P M E R G Q S I E P L T I N L L G L G L A G A C C R E I D
RANDOM_3884 T S V L R C V H P D Q R E V L K E A P D Y S L N G M R M G F G K T A T P T H N S D P R R H A E G V L D Y V P E V
RANDOM_3885 G Q F R A P S E E A K I A K G L E E S T I Q S H V T N I D A V F A Q D G S L G W T Y K F G G Q G S R Q A K T D N
RANDOM_3886 L T W W V P N R E P N E L T A A L E A N Y P S N T D P N T S I P H P K N D W E T K I M Q E G Q T A K A Q H A H K
RANDOM_3887 Q G E Y D L V G D A V P H V A V E Y M S A Q T I D G A S Q Q L M S G D R A A A R E A P A A K N L F L D Q N K L A
RANDOM_3888 P N A L I Q N Y I G D C I Y F S P N Y F I E K G I V Y M A G P N G F A T I K A I N S S Q S A Q E T T V P S G V V
RANDOM_3889 N R G K R R H E G T S V A L N I A L L P A D V L N D G I A P Y L L F N E S I N I Y L D T T E G P D V M Q E Y T N
RANDOM_3890 F N A V L P K D I H T S N C F Q D A P N A D E K A P I S N A G G R F N L A L S F Y S Y I D N I Q Y I N G G R M S
RANDOM_3891 K P D I F G N R F N L I V P W Y I T D G S K F M F K T K N E N Q F K V E F I M G S P S E Q K T A L W A Q A A E L
RANDOM_3892 S Q A P E L E L Q C G K T S P T F A A D N E V A F T L I N P H L D F Q G V G S G P S E Q R E I E L S N P Q D M D
RANDOM_3893 P N D A L L E L P A S D Q S P A F A I L A T C K F A P M E E S I V E T V S E R E E V A E K C P R F G T R A N T C
RANDOM_3894 V H G I A P E L L A A V G L V R E P A S A Y D M H G T Q K T L Y W H G M K E G T Q L V G T I T G H V R N G N G K
RANDOM_3895 Y L D V T P A N A L T G V A R L K T A K L H G D I K P T C Q V S I A K M I I F I Y V S E A S A P P V G Y N Q S A
RANDOM_3896 I V G V E N Q G F R H G A H G R W R Q D V L L S K T L V L H F A A K S W E A P D K L E T E G T C A V N R G I M R
RANDOM_3897 A P N A T D S F N K D E T W V D Q A T F V H I V R V D N P T D V T C D H F R T P A I P D T S P K A R L S T M V V
RANDOM_3898 Y Y T G W N R Q C A R W L V A R L C T E L G I L A I G P V L V L K A T I N S A K N S Y L I V Y Y Y T N C T D P I
RANDOM_3899 A S R I A I E R G W A I P V R M H R A F N N K A Q R F S A A G I S D L Q Q V L F P R T G Q A D F C Q G L R W P G
RANDOM_3900 E C L Y A T S R S G N L W A L T G M K T E R R I A E D R V S R R R S I G M D Y V P Q L D G F Y L M G D D A V E L
RANDOM_3901 S I R G N S P D L L K G T K G G T P C H D V G S G P H P L D G D K V G D L A G I C R G I S T L S G G G A L A V R
RANDOM_3902 K D A P G Y P L K S Y I Y Q G D I E G Y S Y R A S L M D G H G M F I S M W G D P I L A L N K V Q V L V A R I Y L
RANDOM_3903 T W D S L L S I D A D Y F N M A A L D G Q V T D E I N S L V P V N V G M N M Y R G L G D K S D K E D D E P I T N
RANDOM_3904 R I M K N K V G H V K K R R F Y M F R K K N V D R I D M G N Q K T D E L M A K F L G Y N A T C R D P V T F K I Q
RANDOM_3905 D S A R D P N Q I L N L L A G N S G E M Q H T S R T H V Q D A A C G P A S Q W N L A A A A V C Q Q S G E G M A E
RANDOM_3906 L Y N T E S L T P P Q R P F M V L T G D L Y E M R I P Q T H D I S V G P T G M I Q Q G K F W D P I E T S E L D V
RANDOM_3907 Y G R F L S T S K I Q R P L R L K T V H K F N T N S A F R D T E R V L R L G R K H Y G L A L A P G D P V D L R E
RANDOM_3908 L L S R I L S P K N N P Q V A E H D L E V C N E L V A N L F E S T E T K Q A E A S D E L T R Q A L A N C R S I V
RANDOM_3909 F R Q Q A A P G F V N D N N K G C Y Q Q T E L Q D A P D I P I Q A E P T P Y C E R W I C S P G I N V N G P L P F
RANDOM_3910 I P R F K A S E H S V G A G A D T W E G V P G T T V L Y T V I F S K V V H L E F G T H L I P I L V R V L L V D E
RANDOM_3911 R E I R T F I L Y E L P D A N R L L A S D A I E Y A V E D L E A Y L G H V E M P G S G A D A Q N N R A V A F N G
RANDOM_3912 K I N V I V T N L G R F Y Q K D D R I Q F L Q N K D L F G T K K K D Q V R L G E S L S H L E Y L T P D M T Y A F
RANDOM_3913 A I G S S D E V S G I S A S C N Q T V K Q E Q P V I A V R C I S L V L A E H L V G V N Y S S T R D V K T L C V V
RANDOM_3914 M L L G E A L F F G G W F E L A E A S Y A D V D G R E P F D I A C V S D I R G P S L W I D E Q F A Q D S E A L A
RANDOM_3915 F S V K L H A T G K G I M R N T K L G S K Q V V A P L L E R A R N K G S A D I L A A D D Y H N L E E L G R G Y G
RANDOM_3916 G K G S K Y A S L V P E P D A L S L L P Q V N K T F Q S N D P Y V I C H V F R I I Q M D E Q Y T M P L G T S T R
RANDOM_3917 P D D V R A A S G I T Q G T P I G I G E H K Q V L G S L T E Q W G I H F Q E N S G T Y F I A D L I K D G Y G L M
RANDOM_3918 A P L E H I E E P P R D G L K Q L E D K D N Q L I C T F H H S I S E D N T L E N G I G D E L G R S T V I H P R G
RANDOM_3919 R P C P G A K K S A T P Q T N S P A L T N F V S M G T K V A M D L T Q N R G S L S P V L N K T E K R L L H N P T
RANDOM_3920 M Y L Y S P M R L R G C A R S I S E S K T S D A H T N R I C D S N Q A A E L V Q K N L G M S A P V I D E K S M P
RANDOM_3921 Q F Q M G A A E F A G A I P R A G A A L R K Q Y N N S N Q L N E L F V S N M I M S A S E I P L A E L T V P D H T
RANDOM_3922 K E W V L N R Y T A A T L Y F N A G C E E V A A F T N E E D S D E G E C V I L W L S C Y L N Y V D A Q C I K P K
RANDOM_3923 G W R P E M S L A R L S I K G E G H C Y T I T K K M L F E P L T A P R I N Q K A Q R I A I T C Q L G G L P L T V
RANDOM_3924 S V D R A W T D D V K L R K G R I F G V V S F K Y N T C A D Y A V Q L G L G A E N N F S K D V E R K M L A P W P
RANDOM_3925 N S E L V C P L K V P Q T S L K G E A F F M R I D N I G D Q I G V A I M S A T R I Y R D G I F Y V V L L Y R K R
RANDOM_3926 N A G S D H C Y N L S M A N T T N D E K N N P E G P H A H T K L A A L S F G I A Y A R I I W D C R A E N V Q R F
RANDOM_3927 V F C K W H T L Q S S Y W G I A M V W H G N Q C I D A V P F L A E R R E V L E K L P I P Q N T R S R I T H L V G
RANDOM_3928 L T S N E I V N A D R I S L V L I K D T C A P S Q N V P V L V E P G L K A V Y H Q S A H H M S A R T K D H T A G
RANDOM_3929 N V S F L L K G R W N L I L S L V R A S E P I Y V T N D P K T I F A F P S R D G L S L L L S L V Q L I L L R T T
RANDOM_3930 K I K H Y A V P A D G K E T N Y L V E Q H R R D D T P R P H L V A T P R W D S R P L E Q D P W F Q D K L C H D L
RANDOM_3931 G T P A Q G T L V D L W E E K M S A G A V Q Y S S A I Y C V S D M L L R L K F D A L T E N K E K L E P I I W K N
RANDOM_3932 T P L T P E K C A V M G D G R T L V A P I V I A R S S S P R D A N T N T G R P T H T L T W A D S G M G P N S M G
RANDOM_3933 L I G G R L V S D N T R G F G Q Q R K A E K N A P Y P C K S G A K S G V V L D F Y D L P F Q I E A E K S Q N Q A
RANDOM_3934 T V K N I E L T Y I T K Q T G S E A P L S A E M W N H A W G S G A T I I G F V L F G R L L V Y D C Q A P F A P D
RANDOM_3935 R R L Q D I L A M Q V K I G Q G N N D Q R P K L H P A Y K L Q S G E E F E D L P E A A Y F I R V L A G G A K P I
RANDOM_3936 S A N I D F A L T W L L A G T C G P L Q V S N L F I R T H A H D A T E K A Y A I D A G K M N F Y P A E E F E E E
RANDOM_3937 D L V F A M L R D W E K L L E L L T N K G G V R I P W E A L I P D V A H Q L S Y R N Y Q F E R P G R H S T I G V
RANDOM_3938 I V F W G F E F D D F V L D I D Y V N K H V T G Y G W P T Q Y Q E N Q I Q G P L I L A S L A F F S G S L S R N T
RANDOM_3939 N A T R H K M A Y S E L Y S A S A Q T Y G F A I L P P R D G R F G L G F E Y H D N I T Q A V D N D S P S Y W G N
RANDOM_3940 V N R V I K V A A E N L K H A Y N S D I Y R Q E N D H T A K K R T Y Q V Q I A N K T G G P R P S A S A G P G L Y
RANDOM_3941 V V G Q L M P A V I S A Q N H L G E S R A D T S P G G E T G I I L P S L L C N P T Q Y G T P E D T D T D P K S S
RANDOM_3942 A L A I V Y W P Q E D L F K S L K S E T C M F S G M E P L Y A S S N Y D P G D I N D N K A P N H V Y V T V N K R
RANDOM_3943 F L H V I S D N R I S R L G K F G M L G L I A K N G N S E T N V A D G C E A H G D V T D R Q A N G G N A R S L A
RANDOM_3944 E P S A P A P A E E N A F P Y L F D A H S L V I Y G R K P E V R I K A H R D R V T K F D H W N T G C T N A Y G S
RANDOM_3945 S P G D Y S S P D V K I V N L Y T L V Y Y Y D L D H F H A L V L V G S L V A D I I G F G G V L T C S E Y D Y K N
RANDOM_3946 G R K D G G Q P P P L K W V S D L D K P M E R D Y S I K P Q V T I S L G G G S G T K Y V S S D L T S L R V E D H
RANDOM_3947 I I C Q L L A S S S P L E S L L M D N L R E A E V L S P K E D T A V T S K L A A S G T H R M E Y R L P A Y S Q K
RANDOM_3948 K A C Q S G V S E A K I Q P S P S A S D T G L Q A N K T C A V N V L D D W L K F P G T I K T T H I M L D Q D V S
RANDOM_3949 A W K T K I E L E T K E M P V A Y E A V G T N T P N Q C M P Y T P L G Q R G F T V A G D L P L Q S Y T T Y E E T
RANDOM_3950 W K G P T P V N C R M T H H S I P P I K Q R E E G E S S F K I N D P T R A R K E D N R K E R P P C I S R I E L K
RANDOM_3951 N F L F Q L E D E L D N V E A S Q N F Y L Y Q C D A S V W H G G V D K P R R R T C I E E T E T A F F S I I P V N
RANDOM_3952 E D L A K A V A E T A V K P N L D T P T I K K G I E T V I F R G I S D L G D N T Y I A D L E T F T R I A L G P K
RANDOM_3953 L E G A A E C T E T I D S V Y A P P T P E Y S K T V S A G S G L L N T G L K V G A G S S Q L T I Y E I A A K K S
RANDOM_3954 L K A L G K K L V K H L A G K D D D A N H I K S K I R V L K T S Y C K A I D F Q P R M L I H V N G A S V M D N
RANDOM_3955 R K G T R G A I K S P S C A Y S G S L I Q P P Q I A S A G D V L H L K I A E S S Y E T K L Q A P R K K W Q Y A R
RANDOM_3956 I P H S S E N D R T D K F K F V P C K R L T I F A R Y Q E I G H S S I L G E E N N P F S N Q P V L R T E G M C R
RANDOM_3957 A S V E D V L L K A V Q V Q N G L A I T V Q K F K R Y V S K L L H N S G D K G R L A L T V Y H L E L R F W R M V
RANDOM_3958 T A R L R S E H G A Q Q K V P N F G F S W L L T V S S Q G S L Y T E G V T F T L E S L L G C S N G Q I V S Q Q F
RANDOM_3959 E A L A R A K E N E G S G K A S V E I A S N A G G A G M W K F N W N L Y G G E Y S L V V T F T V V P E A H Y P S
RANDOM_3960 A G Y S F G W P A T N V A M R D A L N S A R W W H S D L G P D D F M N E S L G E P F E W N N M K H E L T T S F P
RANDOM_3961 S D Q N N D G A D Q E K R D S A D V E Y S P R L A Q Q K K K G D T A P E K V F P S E H L I P S G G G K K A V S I
RANDOM_3962 E G N Q T D N N P V L S I K A G V E K E F L Q L I N G D A D K A W D P P P T I S Y L A D N R M S L F C R L F R S
RANDOM_3963 E S R P T K E L P A R V G K G M S P T V I P T D C I K K V V Y S H I L S Y S V L E V T E F V P F R L T G V L R Q
RANDOM_3964 K A G R I F T S G R K R W C V L C S N I Y G C P Q T Q P G A D S A F S Q P G T C M V V Q S A E R K A M L S A S I
RANDOM_3965 G S S G E I I T R A W V V N L D L L K M P R L V N M V C L M S N L L G L M I R C C P E P K A K N C T G H G Y M K
RANDOM_3966 I F C T L P E K S M S E D V D G L L D I L I G A G K L A M G L R D N V P M V S D L Y A N E P L P N E H K D L L T
RANDOM_3967 I L S E M A N S E W L A F K E L I E G Y N K V N Y A S L P N E D V T N K G T S M E Q Q L H K G A L C E I G T P P
RANDOM_3968 Y V T E P G L G L G F Q V T F Y M H G D W D I S G C N E T G L V P A V S P A S G S R Y G L K T G F K S R C M R L
RANDOM_3969 D S N A D W A V H N S S L E K G G D M P F S P V E T N L E V S T Q N P I A R I F K S K G V P G L L V D S N Y Q Y
RANDOM_3970 G C S I P G I E R T V D S G L E D N H N P R L P T D R L R G V I A K D N T M C E T K E P V Q E S A S S P S L E A
RANDOM_3971 N L A I L E L Y H Q D K L A A I N D S G V T K I L E I R N K H S L G E P E F L L L N K I A E P D E T E A G L K V
RANDOM_3972 F P V D P L Q L T E E Q K R L E M S Q N D K T V I E C K D L N V D N G K S I T E A F C M D E K C R G A N N D S K
RANDOM_3973 G A G A A E Q A N S S H K P A M N L E G E L H M A E A D M T K I N T I V L S S C I W S G V G K A Y D T A L V D I
RANDOM_3974 P A F C G K G D N S S D E C G V R E E E K M G R G P N T K Y E Q L R K M R G D S D N V I L N V K V A V E S P I L
RANDOM_3975 Q K Q D K I R F L G A D T P C A Q G L I P I V L K L T D A G K A L Y A R R R S V R D E E I P A G I G K K D G K V

RANDOM_3976 F F A M G T K W I P E M P A S E A Q G V E M P D P K P T D G E T A F S F N D L S L I T T E P Y A Q E E P E D E I
RANDOM_3977 E R V K S I P G A L V F E M C A F A H P G V G E V A L G K Q A Q A A L G F P A L A W I Y E T L N P G A I K Y S G
RANDOM_3978 V V Y I R T S E C K E M Y G Q K D E R V V T Y A E G I P T K P V G K Q D D K P D S L D E I A D E F N K T A E S A
RANDOM_3979 L V F D P A K W H T P G E L D V I I S H K L G G T W L P N K T V R E R N S S S K G A T Q D K A K K Y V P A S S S
RANDOM_3980 V V Y G S V T W N A E E G R T E V L H P T E E P Q P S L I N V L H M Q N V R T A V F A F T S N V K M A C Y L Q L
RANDOM_3981 A L A S T T V T G G G S Q I N Y R I G T K Y Y A F S H K Q F S A V I L E V G E I A M K Q R D I L R N F M N E G Y
RANDOM_3982 T A L L V K R G P G I D V V P L S S T S H F T A P E A C Y V S E E K T D Q N S A I V E L K D A I Q Y A G A V P T
RANDOM_3983 K A K E N K A G T P Y V E F C W E R V F D G V G W P F T F D D F M T E E K R L K V K R E A H S L T K N R T Y A G
RANDOM_3984 A P T S I I V L N S E T F N Y I P G F D N L L N G N S Q S Y N A P D F V D E L A N N T A P R S P E M R E R E Y V
RANDOM_3985 L N I F F G C F D A N A Q I I E L N P R S L G R E F H L D V S D M T C K K P A Y N E A A A V S L Y P V K L M D S
RANDOM_3986 Y G S V K M A Q W W I P D Y Q K E K Q S A A Y E N S A A N S N E S G C G P D D E L G A T A E D S F G L Y L Y L P
RANDOM_3987 E P F A S T S R Y L P K S D F I F N P C G S N G I N L A Q I I T F P E K E H F E N T V I T F N R D T K P S G V I
RANDOM_3988 L S N L L D R K L R K K S I F L L Q A V Q C M V N E R V Q R D G G K I T K R T S I G V W T R Y V R T G T R I D L
RANDOM_3989 T A G Y I E R I P W Q S E V T Y A V G R E S A E Y W D I R D M L P F E Q M K S D L Q N A K R T P C F G G R F I E
RANDOM_3990 R E P T I V M I L I G E V Q D Q G I G V A F M P A D V A I A K N Q Y E C F L S G E F G V F R N E D Y W R G G G T
RANDOM_3991 L D I V L K C K D V T I I A F T V R P P R V T L L G V N K L S G A M F D T F F A L P Y A E S L L L C H A F E T V
RANDOM_3992 I E L G Q S E L E I H R L S H D V G G H S Y K G F N M I S E L S L Q E K A P N G L A M I L A T G E Y C K E N H W
RANDOM_3993 G Q K E Q I M S E F P E R S T N L D T C S S N A F V T L L N D A V V Q E E G F T K E H L N R S T A L C E Q Q K F
RANDOM_3994 Y V K S N G F V N K H G K R C H D D V T T A T M A G N P H A R K A L F T D D Q E T T T K F V F K M L T T Q E I A
RANDOM_3995 D S E C F C I K D P L K V A A A N N A Q N C G E T G S R I D G N L K K K P A T L D R E K A E T G R F R S F E V V
RANDOM_3996 R T A E L G N S P G D K F K V R V A A A N L F H G V R Y P L A G S N D L P A E M P I T G D S A F G L A E G G E Q
RANDOM_3997 V E G S L V Q V A S T S L I E I D A Q A Q R G F D M L V H Y I Y V Q G L K T Q A D P D G A E A L Q R A M E S K F
RANDOM_3998 V I A K R T Q F D R Y L L S A G G N T W I E G N V V A G A T G Q P A N G Y K D G S M Y A D C A S K N E C T L V F
RANDOM_3999 L I S R A M S F T S S T L F P H H P P Y K S V Y L R V H P A R F K Q D I G Y A E E L G P F G G A E R M Q A T L I
RANDOM_4000 V F P P V D P G S L K L N V N L R S R V T V M L D N F A V F A N S E G E D G A E P T L S A Y F T E S K S Q A E G
RANDOM_4001 L M G L S I F W V N R L I W A Q G N G K T N S G L G N S L S L I I G C L G V E A S P R L G Y S V S L L R L K S S
RANDOM_4002 H N V Y G L F N C K V S A T N T P D L Q Q E F L E D D V D T L G W E T E M I S V N E K Y D A S V M V V R Y V E F
RANDOM_4003 L I A L T I G C S N R S T A V F A T T S L Q Y K A I F K T F I G D S I D Q L K A I L E N A K Y V V V V L K S C L
RANDOM_4004 F D T S Q A H T R E L M L Y F G A D I I G L M V N V C G I F M H N N L T L K T I G C A E C S F N E A V T R D L S
RANDOM_4005 P G P Q T G F Y D I Q K I I T S V R P K M L C P V S G E Q L A Y L G I E L G L F E L R L K S V G G L L P A F R I
RANDOM_4006 W S L L R A G N R F S S G P P L V N A N Q H S W V L P L P I E V A S S V P K R G D T A G R L Y S P G A I N R D I
RANDOM_4007 S E A K V S D A G S G M A S H R K R S S C E K A R K K W S G A N K V P A R Q S Y V H A Q V F I L Q Q H D A S I P
RANDOM_4008 D N V E G Q R V K N V A P V R I Y D P S S L T D Q E D A M Q R L I P A W N F E R K N A G I L K K P I L I S K P S
RANDOM_4009 W E N G S K L E L L K D R G L L T R A V E V G S N A E L W I M N Q Q V K A L S L Y T M C A A P E V M R D I T S E
RANDOM_4010 W V D T P D N K S N Q Q F K Q R R D D R K Y L E T P Q T R I I E I S W C G A N L I R P S S A Y G I N V T V V D L
RANDOM_4011 I T N M N D L C D V N I T N L C Y H D D L F T G V C K W M F S T L V I Y G Q I D R E F A A F K L K I F D G G F I
RANDOM_4012 D P R E A G H C F S K V R Q L G D T V Q L K G N S V R Y L I S T E L V I E V V I V P M Q D L D P L D E E T Y S K
RANDOM_4013 T A H F N D Y D I S Y E D S L H W K L L A V K A G D E V T M V L N V R G G V C W D H K S T T L P L L Y V K V A Q
RANDOM_4014 M V L I F S K I Q G T K T L V R T D G A I K A A D K R L R N I K S L F P F V A N P M P I L F T I K L W A V S C D
RANDOM_4015 F Y I G D A P R D V T A A N S H A P D T V V F V Q C G G Q S A E R K C G E G V Q K K P G G R Q R Q I N V D A D L
RANDOM_4016 N L P V M L S N E K S G I A V G T D T T L L Q Q A K D C A D E I M V E G A T I P N L K A S D L Q I V H L P N T K
RANDOM_4017 F L R R K E L V V I S R S D A P L E D P H G V R Q T A A Y S T L E G T S L L H T T P D S V D V M I C C F G M E C
RANDOM_4018 W F E R V Q K A K K E S G I L I I A V P T M Y G I P G E I H E D M I G K Q N S I D G D H G K V L S E S K A T A V
RANDOM_4019 T Q S K G V K C E V T I S H G E K D E Y S P H N A Y T A R C F G T P K K G E P P G R H A R N L T T L D V V E F R
RANDOM_4020 D E P F P K K G N P I Q S A L P L E G E G P L E K F P S R M I T T V L D F I D N S V S S E S S T S N A V A I G Y
RANDOM_4021 L S A V S D R L V K H A S K T K S T Q E K Y G R L D R L S A L L D R P E N V S I V M R V I A C V S T Y L S N S R
RANDOM_4022 L S H R L P L P A G S K I T K K Q I L Q E C A Y A E T G A S I K F D S P P Q C S L N T V V H I A E E V T A L N L
RANDOM_4023 R N G T E P D T V Q A A V V R S K F N A Q K S E Q T A A S I L S A D N A P G W Q A I H T P H I I D T E Q T V P I
RANDOM_4024 C G K M K K E Y M K A T L P K Q S P H D G N L F L K L T C A K M E A T C T I S E Q E P L G V N I K N A G L N E G
RANDOM_4025 S E T A R G Y D R K N C G E L L M G A I N T A D L S H V K E P F E E R M T A V G A E P G L R D V T A S L T D G T
RANDOM_4026 D D H I V S T K D M N G E S L R G T H A P A F A E I G I I I V N L D N E A L T K I E M Q Q A N L A G D L Y P A L
RANDOM_4027 G K L E W Q K S Y V N V Y E S V F E F V G E S F T C Q E R A G E G P F L T N G K P Q V A P Y L G A G M G S L G L
RANDOM_4028 D E M S L P L N V E G R F P T D W L D L V S S E I I N M K K E M I L V S G N G I E R L A V T P M G E A L E V N T
RANDOM_4029 N V V N H L R D Q A S Q C R N L A L G S R S H R L V S D D K T G G I P T G N G G K T R P L T N W A F S S L T G V
RANDOM_4030 V N T R R M N W L G C S G K V Y I S Q V S P A S P T F S E S S R Q L Y P E D V H H Q E I G A D P D F L S K A W A
RANDOM_4031 A P K S L A L V T K A L A T D L T Q D L Q I L L F D T P S T V P L N I R I A K R T R P F T S Q T G G D P W V N Y
RANDOM_4032 W M S L Y G I P Y E K R E M R Q A S Y N N Q N E Q C V Q E A D I R T G Q G T K Q A E T K L W V A N N L Q T P Q D
RANDOM_4033 K V V N V K R T K P L L A A K Q C I Y W D Q G A D V D I Q A S T N I R S D D L V F S R G L P H E T Y H F A L G R
RANDOM_4034 F K F A N I Y H D E K A S L A P K R C Q L K G D K P S Y K D L Q Y D Y N L G F W V T P A G K D T G T D N S K A Q
RANDOM_4035 S I G G K A Y Q G T G S T S S K R I N R I L S N S S T V A Q A A R D P M A A T G G V N C L T L S Q S S A G G F D
RANDOM_4036 V G I V A Y R I I Q K N E P H I D G G G S G M Q P L E A L K L Y G D L V V D A A M E Y V S S R Q T I A L Q E T E
RANDOM_4037 D T T Y V K P I P P D N G H G I Y F G L V R F T G L A L L K V G A I C S S D I L P S S Y T T T T T L I E Q F A G
RANDOM_4038 L E D E D D L E E K A L D A I I N V K I N F I D G E N N P V S E L L E I E P T V Q G D S D I A T G L A Y Q A V G
RANDOM_4039 V R R N D F H G G D F D S E H P Q A S E E L L M T A G K P P G K M L V K E A T T D F S D L P E F E I G L A V H M
RANDOM_4040 H R D K K A I P D G P Q Y G M T M R K F F S Q Q R T G D G S E D Q G L P V R D L I K K V E L S E Q R W W Q G L E
RANDOM_4041 S K F G D V F S H Y K S D F S G H N K A N A S D S N M C K W Q T L Y Q E S V S G L T G P C T C S Q R M T M A H P
RANDOM_4042 V L T I L T A I K L Q C G G K S Q D G N M C E R P C E E K R V N N A P R L E V G Y P V S A A D S I G V R I A L I
RANDOM_4043 Y L G R S E M R V H E T P A Y P L H E W T S L P K D R E I E I G R R T G G E Y Y G P I P C N H T A G K T N G L V
RANDOM_4044 T I D F S C R V P D L P M Q D G I G F S Y A Y G A N G D P L R H S I T L N K L G I S I S W P R W K G E R P E R G
RANDOM_4045 H H I F L K Q T L P G L V V A L R V F V Y A F K A N G R V S S E A E N N G L Q V D S E Q T L A E D Q A V V G M D
RANDOM_4046 V Q L H A S D V L K S H E I K L E P S M E V E N A I H F P D L P A V T N S K A D L L H V L F E H G P S T E N V A
RANDOM_4047 N K V G L A G T P M L N F E L D L K W M T E N S S T T T D L G S I I T P G A Y T N K L E P G L K D F V K R E W Q
RANDOM_4048 A Q E Q G T L Q V D W I F V A E R Q T D P D K N C G G K Y W K G T D G M M D Q L P A D A I P F L E Q D G G Q A A
RANDOM_4049 A F N T L T M R G A E S Q S H Q L T G F V S A Q N A L S P M Q Q K P D E T L S P W F S S P C M E T G A I C I K T
RANDOM_4050 P C F I P E A S N K D A E V E T A H R S I S A T A P E E A L E P E A S F V Y L L L E G A L A A R D L A V S K E F
RANDOM_4051 P K E S N K S W K L L G H N V R P Y L K I G Y L L A Y I N D L G E A M R M G H D I I S G I V F C V V V V I I F K
RANDOM_4052 G C N G E S V Y D H L A V N V S R L T D E K L A E K L Q L S V M C I Q Q I P W S L G I D T I T P D F R E L A A L
RANDOM_4053 P L S I K I V A L V V H H E A H K K Q D A G P Y A W S R F N S G S E A G L N E H A R A E E A V G A L Q V I S G G
RANDOM_4054 V P L G L E E L L D F V S L K Q L I M V I K F P Q G G L Q E I N S W L S N T V L T V K F L S A E P Y G P L L R S
RANDOM_4055 K S D K P K L R F L P L E D Y P A Y S K S L T L G I E K L V P P P P S K F R Y T G C A G G T G T I S W Q V V S T
RANDOM_4056 T K E A G D V I F A L W L M G G I P Q S G W F V P A A S D S L T E E N M T M Y S G V A N R G F G H H M T L I K R
RANDOM_4057 G S E G R R E E I D F L N T S L F P P S A Y T A E F C A A L T G T D Y N V N I P D V N S D D E L E G D M C L L L
RANDOM_4058 R V L V V D K I F L F A K M N W S E F W V A P P A G A E Q V L E N E P A E A V V Y P E A T P Y L L Y E F N E A S
RANDOM_4059 S L L V W V Q Y V Q K D K A T M G I G P S L N G S C V L T R D Y A E Q I E N D E W E V G P K N T Q Q D E C S V A
RANDOM_4060 P T D R R V L Y A A A V V K N E L H P C R A P K F G G G R V E E F Y E N K V A Q T L V L N V F D P V L D L H P S
RANDOM_4061 I L F V G L G P V M Q L N V Q Y K A K E S T K S D A G A G K I L G V Q K M P Q N Y G K N I V R G D V S A D R A Y
RANDOM_4062 R I L V V L G E F P E P K I A D E K C K P E F V A V D P L D T L A N S A V V F Q A A Q L S T I A M K D N N K K T
RANDOM_4063 S M I V K I A Y A P L D A G D V R R Y G D C R T E I G S A M D A D G G R M E G G T L N R A G T L D R Y K L K G E
RANDOM_4064 W E L A I G L D K L F I V K V L G K K L C R Q D I T E A V T I A W H L K W T V Q N A D L A S R Q T L W A L H N V
RANDOM_4065 F Q S G E N E S H S I F L T D A L G S A F P I C K Q G Q G W Q F N G P K L Y N P K T L T S E C V K I T S L S G M
RANDOM_4066 Y Q T D K L V Y G F A K K P P H L P A A C G N T F P E I A P V R Q W A L I L F L P H H D E E S E I K T Y T I L V
RANDOM_4067 H D T A R T L L I C F G D R S Q V K D Y D A L P R P G L I Q V K A H R A P Y R I D D V R S D P I G Q S Y E I L T
RANDOM_4068 K K V A D L V S F V E D R S M G N F T L D D E D G K V R P I T L A V G T S S Q E T S K L L E D Q A L I M F D R G
RANDOM_4069 P F R S A T R N V R N Q N I V N E T M Q G E G L A I T V Y Y T Y R D G Q E E Y D A P E A T P S D D I S L A N S F
RANDOM_4070 Y T D A T Q M L G M P L A G Q A G E G G L L N S R E S E D D V A Q D L Q T V L M N V C K Y K D K Y S F G F A E N

RANDOM_4071 Q E Q G N A P Q F Q A N G K D P Y S E T S E S S D A N G A L M N K L Y E K A V I A S K C L F G N N F R Y S K I R
RANDOM_4072 S G Y A S L A D G A Q E A A I M S L T N K S I E L F A E P W L T I S L G G T V G E S F E C R D G P T Q A L A F S
RANDOM_4073 T S V F A A G Q F G T V I D D W E C D S N G F T G S V A S T T W I A G K A D F L E G R K S E E L E C I S F Q I I
RANDOM_4074 F D L M D R L R E T S D N Q E G S K L S L N E Q N H N F T I N D K V G S L P K I L E A K A A S A K E V L K V P D
RANDOM_4075 V P A Q A S N L R V Y A G D V Y I A S V A L D K C H P D M G L N T V G F T C T D G N T I M L A E L T Y E L N L F
RANDOM_4076 F Q E I K D P Q A I V I V E N E A S R T P G M S V E I A E A I N I A S P E K I H D K S N Q A Y V R A K T K H V H
RANDOM_4077 M P L N E L K E C H Q N I I A C R G Y L G K V E D L I L L Q K K V S S G L T T S I K A S R P R V T E N G P F G G
RANDOM_4078 S R R S E G K K R A L L V V Y Q G M G A D G T G A G G F D Q G L G R H E T C E S A R E S A G D K S L E K T I E A
RANDOM_4079 A I T N S N D K N K C D D F E T P C Q A Y D N H M N A V L M Y N P F R T L A P Q I P E K L D L L E D K K R A Q P
RANDOM_4080 R S K I R R A Q S Q G F R G F D T F G G S T D P T H D G R M N I A I V E S C G L R T V D P C I L G Q R E S N K V
RANDOM_4081 E Y L I Q S L T K V F Q S Y L D H L A W G G S Y Q F E S G I G N R L E S W T A C A G I T D V D L S S V L W M S R
RANDOM_4082 Y T L F T P N I I A E S K G F F T V D D G E A T T P Q D R Y L G G L N S E A L D R S D G T N G I V S S G S N A F
RANDOM_4083 D S E P S T T V A G Y G E N P M N T N R D S I E A C G P G R M E Q A F N A A L L Q A P I R L L E C G R S I E G R
RANDOM_4084 L N N G A V M E E E L L K T F T G R A A N P S M M K C R A K S P T S T V E H V N D S Q L Q R N L F R N I A P A C
RANDOM_4085 P E E A T K K K H V D A L L Q V Q N L S L A V F V V E T E N T L L S S D I P E F S F G N R A E Q L M Y K S R G T
RANDOM_4086 G A D Q S F I C H T A E V S E L H S T A Y F D M P V V T W S D Q E D T L K R A S L E D M K G K R P S L T A V S N
RANDOM_4087 S W W A K M S A Q D G M A L H F G G S P V C S S Q S R R E S L P L Y H L H I A I Q P V F V K A T C F G E K K L N
RANDOM_4088 T L E T T I T Q K K I L A N E L P K G A Q K G W Q P T M L P P T N G A D Y L P R S Q S I L T L S E C M G S P G R
RANDOM_4089 I N L R G G V A A F L P Q V S S L W S L I E A T A V R L S D T R A R R L N G V D C Y P M V V N L M Q W L A S I H
RANDOM_4090 I Q V Q V Y I I E R D G Q E G V A K R G V K K D D K K E N Q P T T G E A E V G Y R S G D A L G E E T V K L L A T
RANDOM_4091 N G L L A H S A V L P G E M V I F T F K F G L A Y V I Q N G K G E Y G L V R C L W S K G D D D A G H R G D T R Y
RANDOM_4092 A H G L G E R G L I S T R K E P T K A N S A F I T T P L L K H G K E T P P Y F D P K C D I N K Y F K D R M N A A
RANDOM_4093 S V W S D N E A F D S A Y E S F R L F M P M L I K R E G S Q H L I T A G C G A E T A V K A R V C N S Y T S H M G
RANDOM_4094 F I E K R T A M L R A S D E A T S Y L W Q K V A C T L P I Y K D L G A E T K I V N G L S V E F M N F P H Q M T T
RANDOM_4095 Q D S F G F Y Q K S I P E N T G G E F K E T L H M R S M F G G S N Y F F F A I T T V E Q S W Y L L N D G E K A V
RANDOM_4096 E V Q R G S K Q F N F G K C A Q F K V Y E R I I Y T G H D F Q S F K P E E G E S I D S E P A I K E I I G E V Q P
RANDOM_4097 E G K F A E T V L V G Q H L F K L Q R R V S A V F Y H S D D G C I I R Q E A G F K Q E S K I V T E M P R A G W Y
RANDOM_4098 S F K H F A H T H C T E N S I A P V G L R S E M S D L Q G G F T T P G W N G H G F Q V G L E A K R V P G H G F F
RANDOM_4099 V R E R A V V G S F I D L Y N D A L E N P V E L P H A V Q L R S H P T M I Q H G P C N K S Q N M Q F K V N S G Q
RANDOM_4100 P A A L K L Q A G I T N K A P V P M S P S A A G K D K V E M R T L C R G V A N V A E T Y F S K L L P V A K G A G
RANDOM_4101 K H K S N V V P F T F V E K L K C N I S S V F G D M M G R K R L I T D A E E D A Q I L D A E Q G K K G T R A E G
RANDOM_4102 K E R G A E A K E E T S H C N D N T S I E H A A P H V P G A K V L I V P G K M S S F I V F N I S Q L N S E H K C
RANDOM_4103 F T Q W L T G V V F S E C T S K N G S F V L R K P I Y V Y K L K P V A T V P D I S K G L M S Q N S G F L G K L S
RANDOM_4104 L S T S V G G L Q K D N T S Y Q G P I L T C A A A P L P L P S P Y L F L A T R P I K S S L G P Y R E N F M D F K
RANDOM_4105 V V R V A P A F K P I T G G S E Q D E F E E N A L G V G K R I Q H K N I L Q D Y Q I K K T S I A G L F R P E L K
RANDOM_4106 T M G E L K P E V D K T K K N I K G N V A R L S A R Q V V K W A Y A K G F N R I R T S V E N F F F R N R P I A T
RANDOM_4107 D P L T L V K V R D K G F A H K N C K S T S P D E P M G S V G G V S D P G E D G T G A G S A V A F G W W C D Y A
RANDOM_4108 A R T N G N G T F R I N V Y S Y H E G E F P A E L R P S R V V P A L G G D D R R D E M T T P M P L D L F L P S G
RANDOM_4109 E L A R G K N L S V E K A V A E K C G E G F V P A F R A L T Y G R R D S T P S P E L A G Y I K L G E P D S G I V
RANDOM_4110 L A V G A K S Q A T K C A V C T H S I V C E P R D D G A S G P D S L E D K N E T L Q L S M F M P I K T R T K F I
RANDOM_4111 H W D V G L P L M G R R K A D P K D A I S F L S A C E Q L G F S P C R A A L A K R N E R I Q I E F P T S D A K G
RANDOM_4112 S V A Y M F R D P L K I L N T A E A R A R R R S G I D N E F A A M D I G W E V E F Q E G V Q F S S A C T C Y T S
RANDOM_4113 P C V T L H Q G I H Y E Y D L N L Q R S G K E G I I V A I T L L K L C N L V V S L T T L Q I G E D S T V R Q D W
RANDOM_4114 V G L C R K Q Q L L I R G F R Q M R V K K G L Y T G V L F R L R K L E G F L F V N V G G E N V G G A T T L E T D
RANDOM_4115 T L A K Q Q G L V R I G H N Q A A K Y K N Q A L S R K G C I Q F S I D G A Q D V N N T L Y K D S N E C Q N V K P
RANDOM_4116 A M L R E V P G T T L K A L L T V D Y S E P Q N A P E R A R G E M S K L A G F T L L I Y L F T R V G G A R P H S
RANDOM_4117 S P M P N V D Q A D T E S N S A D T Q W W V N L T I L V L V L M P I E D K N R E I D Q P S A P E H L A R Y N T E
RANDOM_4118 G M P P K Q I A G L I G H Y Q A S A S D R G A K A D N A P Y Y H E V L Q A R A A P K T K Q T D A T F V R S W I L
RANDOM_4119 A A D C P K T V N V M V P K R I V V K F N C P V Q R S L Y L Y L G G F A E D S G K A S L N T A S E N L S L S V I
RANDOM_4120 V P G E A C G L E E Y M E I Y V L L A E R R Q G Q K Y D I I P V T I D W F C V L N T L A F P N Q S Q V A V G Q P
RANDOM_4121 L P R D V G C A V A I E T N A A P I K D S S T P L L V N L F E G N V D S L A Q F V G S C K C L V D K P S F I I G
RANDOM_4122 V Y L S F S H K Q F F H E M G S A P Q D I A I A D T I P T F K K H D I A E D P G A A S Q S G L K F K V R S G L G
RANDOM_4123 N R W S M S I V I E D N V V G K T Y D R F M L K D Q K E A G S L L L G W Y G A V H T D Q E E A A L G N V A T F P
RANDOM_4124 L R K V T K D T F H S G L Q E A C I A K V F A Q I L E A I Y P K L Y L K K R F F M S L C P A Q I C D E T F E T A
RANDOM_4125 Y P F W Y T M P Q I F T E I L A G A M K S T R G D R S A I Q V Q P D E A F V A W A G L A V S P I L P M F Q S T K
RANDOM_4126 Q K E K P Q N R K S P A A K F L K T E A S I K G Q L Q G S Q W F S E E E A V G S E N K N Q D F S D A V T I K F M
RANDOM_4127 C M L S G C Y N Q D L S T V V D S C A G P W I S A A L L N Q T A A V K R D K A A V H N T T N D F V P A L Q C L I
RANDOM_4128 R F L L M G A N V K E K V A D F S I I S S L V N A G K D D F R E S R M L P Q S A A F S K I I L G L S E A N E S V
RANDOM_4129 P I A I G V W E L K E K S E L M V S D K F C D L I K I L S A L R S S P N D F I A A D A T R E A K T C K I L V V L
RANDOM_4130 K N N F D Q W L C S E Q V N P I D L Q T I L L T A E R I Q L N Q L L T V P V S A V W S E I A M S V N H V K A K V
RANDOM_4131 G A I T N K L L L I Y A L Q T I G P I W I Q A I V Y E I C G Q S I F L Q T M V S G H G T Y G G G T V C T D H P P
RANDOM_4132 N I E E W V T Q I K A A G H G Y Q C T L E T L E D F T P Y N G S E R R V P I R C M N I P P R F P G F A G H L T L
RANDOM_4133 N S A G A G W K L G A M L S T L V N L E E W R Q R A R G M N D F L Q E K P G G G F D Q H Q D L L T A E L H R V G
RANDOM_4134 K F S N Q T T R T P I H N V S I I R G F Q L A V C E P L E A D P Q R P F P L I K G T A T F L K E Y I P P V K D P
RANDOM_4135 R E L D L N L S G R L F F H I P H E I F Q D S T Q S Y I V F T N K L S V M Q G P L L S T L A S G I F D Q A A L S
RANDOM_4136 K I S R F M G V A C L I I V G V T K A L Q V V A V T P G E T M G Y K V L T L T V N I L I L K S L I Q L V D L S D
RANDOM_4137 P F L K Y E N T R L S L F A I R F P K A S G D Q A K N D A M S S R K D N R L I S L F F L K I N S V F R S K I E F
RANDOM_4138 T N G S R T T K I A I H I A F D C L G K S L L V G N G E A K F K D D R I L G Q T F V A L Q F M N E N M V M Q L D
RANDOM_4139 G V V T T Q H D E T S A A Y K E M K D G T I T I H E E P S N K P L P R R S S C L D T I F D Y V E I W L Y E E E N
RANDOM_4140 S D K S S G Q A P C H S A S T T V N E F T D L S N I H D R A A S V E Y V E G T R K L E N F L N E Y Y N R M V N L
RANDOM_4141 G L G V G L R V P D I D G N L Q P L Q F E R W G L G W G R T G Y L A T R S V I P C A Y Q E Q S D S E A P L A V Q
RANDOM_4142 S N A K L L E I S V V G S L A D L L V G E S I T G A F A R I D P K G T L Q V A L M P A E G D T D H E C T Q N H N
RANDOM_4143 N R E P S R R P P N K F G G D A G R T K G R D R I T L H P F W T L N D F D M T P S L Y A N D H D G G E E T P F Y
RANDOM_4144 K A K G S E V G F Q H N D D R L L L P P S R E I I A Y D A V P V L Y T T K H L P C A D E S F K V D S R F G G G R
RANDOM_4145 T V I L L D K N Y N S G M F T A Q L V H L T Q I K G P L F Q I I I G L D P Y G G L T L I D S T T R Y E N T T Q T
RANDOM_4146 Q G D D N S K D G P E K C L G E K K Y R Q S L S A L T A I A S T A W K A F P V F I R A E L I L I F V V L L P G E
RANDOM_4147 D V A P L N E G L K Q P L R Q R E G L L M G G D R Q V Y A L K D G Q G C E S P A N F A S M K Q R L P M A A G K L
RANDOM_4148 Q I E P L L P E N Q L E V T V T S I I A V K A V A N L E F N R S C T E V R D I D R E F Y P H A Y P S K M C E A L
RANDOM_4149 M L V M M L S M V D V G Y P F R I M V K M L D G N L Q A C L R K A D R K D Y K L S K S A G S I M A M G S E P G Y
RANDOM_4150 G S I N N S E K T G C S K V Y C L H K N R D W I G L C L V S R R M R K G C Y D F S S N L R W V D F G R I A Q F K
RANDOM_4151 R W Q D S L V V D A E L V L K M M F G Q D D A F K R E L A E E G S N I L T S P L L V G F Y P R S E A K E L S P L
RANDOM_4152 D D P R Q P N A F L W I F A C H P N T P L E E V D W S R V V L M G G V M D K T A V P G F G E M V S H F V I S V I
RANDOM_4153 G A Q G L S S P L N I G V N I G D K H R D Y T E R L G E F D P S P Q R D D S D S G E R L S S I K Q S G I G K E V
RANDOM_4154 C L D L E P S L S S V L I K G E Q N S V F L S I L S K R K D V K L F S L P N L A K A K D L L N A T L P L V V P E
RANDOM_4155 V S Q Q K K N K M A L L A C P Q F I A Q C L E S S K M L F Y I A E L H Q E E S R N T R G T F T S A P G L K M N I
RANDOM_4156 A P F S C A R G V D T Q Y L C L L D T L P L G R M M G G S D A T E M E N I N F V L T V G K D F H R L E P N D L E
RANDOM_4157 W A C L Y H W D R G H E T L S F L N G A I F P G E I H I I Y G R Y V G T S V V H S V T N D N E E V E V R Q V G E
RANDOM_4158 I G M L G F A I G S D P Y L Y Q V P Q Y M A L N D P I N V I K D F K A A S G E N K Y L Y P P S G V V M V T G F N
RANDOM_4159 C A L Q Q L K F T P D S I I M I P K D A G N R V T P F I V T K P D E G K N C L R P S E S N R Y E V T D V Y A T S
RANDOM_4160 T S A K Q M F I R S F H E C P T A Q T I A L T Q Y R P S D Y Q R K T T G S W I Y T W D A Y E P F A P Y L N M I P
RANDOM_4161 F P K R S K M E L H Q M F Y D S E R V R G H N V S D G G A K M L T G F Y M D G C M G I F G F A A A A D G K F S K
RANDOM_4162 E F S P V E L F A A C Q V T R I M L I M N D P E G N S S T L Y L V V K R T G F R L S P F T A R Y R S V W E K K D
RANDOM_4163 P V E P E G P T L S P L T P P L K N Q E L L D L H R T K F Q R M L T K A S I S A N K L K D V G L R T P T T I I F
RANDOM_4164 A D T V T L M L R W G S G E S K H D C M T D T W L G P S E L V D F A E E V T A Y R V L F C L G V T T R G G G V N
RANDOM_4165 L G R V N A F A S G L P N P S T F K K A F P K S M F I Q R P N A T L S K C I A Y Q G Y E G N A K C G I G Y L S A

RANDOM_4166 W V I V Q S E L G L D G P G P N S L Y A F D N F E L R G P K H G P R T Q Q R A N M T E A C S G T L P V L P E S N
RANDOM_4167 R T S P N D Q M R P A V P M I G P D G E H M I N A G N I W Q V A L M A Q I R Q D L G H T D R T R L A P N L D L L
RANDOM_4168 V A T G L I K H L S F K S T E T T S P N T T V N P H F K I T T V L P D V D M P T F G W Q I L L Y S L A D H K L F
RANDOM_4169 N A N F G G L A Y K E Q L Q A A P G Y G Q P T I C L T E I S E D V Y V G T V L E Q T A E T R E G K Q N E D L A R
RANDOM_4170 D P C V N Q T T M P S N L A A G T L V Q T V F Q A N G M A P K S Y C Y A L V N R T L V E F A T T P A L P A I L M
RANDOM_4171 T A L G A D G N W A P D G P E P H K L S P Q G V G Y P P K T Q V K L T E T L N N K N P H L D Q R P I S P G T G A
RANDOM_4172 I E V P N E A V Y R L T I Q T V N T K D V A W Y L V A K V G L W E H S P R V L Y L N D L F S G I T S K R L P G H
RANDOM_4173 I K L I T W E T M W K Y N C S A E G S P Y A F I Y G F P S L V T V M L K T R T I A Y K C P Y K P N A T N E Y N C
RANDOM_4174 T K D L S G Y V I P G K D R T N S A N L T H E F L G F V G V V D E T A I N M L A T D R R I K K A P S A D S F N E
RANDOM_4175 N Q T E S L S D S V N R T G M F M A N L I R W V A Y S P L M P K A S P L V N D K R G Q E S L K A F T N S G Y G A
RANDOM_4176 D Q S A F S M E V S W D E Q P G M G R D Q P D W T A K P D K V L L K R Y Q P F K F P G F A E D T I G R E Q L T A
RANDOM_4177 V F S L Q T A L M P G P E G S T G R L A T N D S L G F C V V M D N G Q L N A A L G A L M L P E F S A Y T F V A L
RANDOM_4178 L Q E F P F P Y P V L I G E C G L I M P V P C K V G G D V T I K L E C K L T N P G Y P L L M S V I L S V A G D Y
RANDOM_4179 V S D T I A E P H C R L N M K I A Y V E K N T P L G D H Q K F V T K A L L E T E I E P G Y Y C L V V Q A I P S G
RANDOM_4180 A S P S N N D P F I E L E S M D F V E G F N S S T Q L I G Y D K I E F L E G L V L K Y A V Q Q A F L W A V E E W
RANDOM_4181 P Q K L V R L L G N E P H F H P N V S V K E H E Q A P Y C M R L I G K T I F V Q L E S V K L E E V I R V I M S N
RANDOM_4182 Y D G S V T M I F A P Q F E G L T V E T V W I R R T S P L L G S R L F D A R T Y M F R K G R N K A E G T I G C L
RANDOM_4183 V P V M I R V K S L E E D N I K I H D Q E A D L G G A M Y V Q I T K L I Q G L G T E L N D I I S A V P A R E L N
RANDOM_4184 V G T L F S G C G L S L L S G M K Q R S L Y L P P N T H R G I W L G I I H I G D I D E T E R R V S S L L S V S L
RANDOM_4185 I S V R K I T F E L E P Q C A N T N V P Q K I D G A C P R D T N P I F N L M S T Y L N Y Q T R G M S F A L L M N
RANDOM_4186 E D G G I S D C G E A Q I K P G G F E A D A C L Q P P S P K R V I A A S G R D Q F V I I V L E P T C A G W E E I
RANDOM_4187 D G S G G E P L F G K K E F V E M W K P T S R S P M G E S C I V N M Q V T L T S L N F Y Y A E K E S D H T E A E
RANDOM_4188 Q G I V K G K I K S R I D S R H S L G A F K V G K I L E G T L H V A V E S L Y N I T S C P F K M E L L L G Q N V
RANDOM_4189 I L E H E Y K L D G L R I P C A N T F G N S C N L L I Y E G A Y S S D V R Y D Q K P L N G D H R D K T F V Y E H
RANDOM_4190 R A D G R S F A L E L P D P K G H L P V N C R F A P I A N K G S T Y R P D N N K A T V I T K D R E E D R L E I S
RANDOM_4191 T P R G V G I E Q V I L T T M S L L D R Y T L F E E T I T L T R N V W P R G Q I Y A E N C A E D D P T N T R Q V
RANDOM_4192 H L E G T T P E T L A G N G G F Y T N W M F N K G R G L G T T F T T V P E S R M F D T K P T Q P E I V Q D M Y T
RANDOM_4193 K G S I V Y E V S V K L P C D R R Q S A V R Q S C L L F F A Q L F V D L A N L G Y P Y S E I D K E C Q T A L L R
RANDOM_4194 T L L G S F N W F L N G V A I N R T T F N S L L H Q V P F Y S M Y A R N G A S R L N V S A A D V G H Q L Y Q Y L
RANDOM_4195 E Q Q V L G Q S D E K L A E L L S N A S D D R L G N L S S T A E P R K T Q G G D K P K S N I G L A V R G P S N A
RANDOM_4196 Q Q L V I V L K P P N I N L G L G L R S T D A W S C Q G I G A S A R T P R V T M W E T L N I E V A G R E G L S S
RANDOM_4197 R E D G Y V S A E L T G D E K D F Y K C L E D F M G G G P L H F D V A F W L D C T I A P E Q L F K S M W M S Y S
RANDOM_4198 K L E V F S D W S T K V V T L D L R T A G D L E V P K A V G V F Y R T E V Y G F C E Y T S D P M V R S G T Y T K
RANDOM_4199 V C V P V E Q G A S S A D L K N F I E V F Y L V I F I A K E C L K H L V K I P A N S A R R L G R P V H S K S N I
RANDOM_4200 Y S S S Y S I A K I L V G R N F G P Q W Y T D I A S G G N G A Q P S V Q A H I M S T L A D Y H Y G V A A S R A S
RANDOM_4201 P M E M K A M C A V F Q V C K A I L E F A K G T C I T Q A L V V Y A L A A I E S S P K D I Q E G F V N K C S V I
RANDOM_4202 G D D V D W G L I Y G A A T V D A C C I P V P V L S N E N H A S S T R I I I G L E G S G A K E A D L N C V A G T
RANDOM_4203 V D D F A Y P T L G P K P F K P P L L G S I Y G D D D R G E I S L Y E A G E T G A I K R L D K N V L V G R I T F
RANDOM_4204 V Q L N A H A I G V Y G L R Y F I K G E K A K L S E V H Q M Y T T V L K D A A A M K T D A T N P W N K K V D S P
RANDOM_4205 V P M K F D E T T L E I G H L H D R L D V A A I Y G A I F I V Q V S M A L G N V L E Q E E M K E A G G A R C P F
RANDOM_4206 W G K T G A Y N S R F Q E S L N G H A F H A L V T G Q S W I E L H N T Q P C K I S E K D L H I L W L R K P S K K
RANDOM_4207 A G V F S D Q K M A R M I L P W T G G T H F R N L E P I V N D W A C D P Y M K P S G G Y A S I G E H L D P N L D
RANDOM_4208 V T R K Q N T D W T P V F V G L P L E Y K C E A E L P F K Q Y H P Q A N L L M E L K V Q P R P N R T G T E P K F
RANDOM_4209 A A G I V F M P G D C K N M G N F A F C H I G Q T D P R V E L S Y Q Q F F P P E A A A A F G V D M R P Q L L M P
RANDOM_4210 V E K A E K F I I P I Y E L V R A I P V S R L E D V Q Q P A E E P K L A R R V G L K A D A A E P E S G D R G N T
RANDOM_4211 K Y N E L I V R T V L S P I R R E G K G L T K S S D F I R L V E K A R G I R A S E F V P G P S G E D P L R R S S
RANDOM_4212 L K I T R T R G S V T T V L V M D S G G G R F K T N N K T L K P Q I W A I A D P Y I T P D E M M L G P M S S Y D
RANDOM_4213 L A P K V T K S S L S V R R A A L R P I N V S E D L I Y C T S N L E S G A N D Y T M V N F V N Y T S F N Q G A F
RANDOM_4214 I T S D Y V K E I G Y R P V G G M V Y L D E D L I L Q Q S D G D D Q V T L I C K D F S T K V L D S D P C A E T S
RANDOM_4215 D L E W S D L L N F P D T L K L K G F A I E Q K E K E P E Y A Q V S E A F T N N D L I N F G A K L S I L A R T A
RANDOM_4216 K K P D P E R D S Q D P E I L T R F I L F Q P G H R P G L E L V R N V V A A N G C D P S P V N F M V E N S I N E
RANDOM_4217 N G L A T D V T V H S V E L Y A Y M P R A Y L A S H L S G G P G R H Q L Q L V S Y S N D I C C S P Q S K Y K M T
RANDOM_4218 G C A F I E R E G K P G V T A E K S T Q N S V E V K E I K Q I G P M E L I G L V D C R L A K G Y A T Q P P H S F
RANDOM_4219 L K A A S G D E I E G T M R P Q N H A A A T F A R E Y C G L K A S E K P P R G I N G A S N Y R P G Y P G I G H I
RANDOM_4220 T L M Q G D I A L M E D G C G R D A P S V Y N V V K C L D W T Q K I T C A L I H L S A L N K Y I V I D E K I Y G
RANDOM_4221 R Y T G K R L I A G L P L L T K Q N D E S L R C G S T D V A I L R Y C Y P E R A V L V M D S Q P V R V G E F T G
RANDOM_4222 C V M S E A V P L V T L I R I E F I I Y G D V F Q Q P K Q N I E L L L S P I E E S G F L K Q W L G G I A R E Q Y
RANDOM_4223 D A V N I T I K R P K A G A G D G Q F V D I R V T T R L Y A D P C K A S A P A A R M V D P D S L R A K A T L K L
RANDOM_4224 K G K V M G A V R I L R Y K S L V L C G M E Q A N V A E S L I A T P S E R D A F R S W H L I L M W N F I N L N K
RANDOM_4225 N D Y A G E L A L A K K L A L G E V I L D F D L G A R T K D M L A T A N P L M I Q E T T N K Y A S G V L I Y L S
RANDOM_4226 Y Q Y H I L G A N T L E D S G T M R M N G K R G P K A Q L G V P S G Q T C A Q T G L K E P T S R T S T A D A G S
RANDOM_4227 A L L V P R M M D V N G I L C N L T S T R A F I E I L A C D Q S L S R F L I R Y V I I A N S L M T K I L W E Y A
RANDOM_4228 L G Q V A G R G E S K N E V S D L D K A V V L K T G N N R N A T I T M H I R V G K A P I D P Y I P K R Q L G R G
RANDOM_4229 N M I G F S S T T L H V Q S F P S T I L I D S E R A M V E E S L K A S W G I K K L I Y G R L A Y I P L D Y G R I
RANDOM_4230 D S A C S V P G N G F G S S F C R Q C Y E V A V L K D V L S L T G V Y G P F S T L R V P Y N I G V T F I G A D L
RANDOM_4231 L V T C S E P M T R P V T S T E G V D N A L S E A R A P A A Y L Y K G H M R N A S T S N H Q A L A N K D L W H S
RANDOM_4232 N I V F A G V K P N K L I K A L T Q L T V Q R N Y I L D S V Q G S F K W T P S T N G D K M L E P R G A A M D S T
RANDOM_4233 L T P D S S D H S L N H A K E Y G A E E D L K T K S E H E D S T A G V N T A R L T V K A D G P T H N E A L F L W
RANDOM_4234 F L L S T A I A T N G L Y D A T P Q N E A Y C T L A F H A R F T I L Q S L E V T T F L D N A W L A V S N M S D N
RANDOM_4235 L S K S H L Y G P E K V E T E R V G K P A A P Q S S E Y N S T L Y G I R G I G V G V L K E L E K G D T A G Q F G
RANDOM_4236 K S Q S F F K D E E T E D N S A Q Y G L N T G G Q D I D I Q R R S G S G T K P P Q Y G M K K Q A Y Q Q G E S V D
RANDOM_4237 P W Q C V G P S G W L K Q V A A D W S M A C A H V K A G N G R G V E D G D H A G R E V E K A P V D P L T E T K G
RANDOM_4238 D N N I P E E T A Q D T Q P E P R I L G A R A I L K K K G A W G Y G R L Q H G Y Y S N A P Y L A P F L A A L A H
RANDOM_4239 S A L K V A T T G P G Q D S V L S A W I K A V C D E C C T L F F G A L V C G D E A R G V L N D A Q I A L R G A P
RANDOM_4240 W L M L L D A P E S F T L A T S A M L I R F N A K E T I P Q H E N R L G T S P G N L S M Q M V G S Q V K P D A Q
RANDOM_4241 P H H E A D Q V K I L F L S H L C G R M A V R P L L L S V M K G F I S V K S S S L N A Q L T A A V H N F T T S F
RANDOM_4242 E G E I T I H R R E N D I K A A A I N F K A A M Q A L G A E P M G R T H L A A G A V V V T R L K M Y C D E F K G
RANDOM_4243 G T K H Y A R I N M V Q G E H Y E R C T Y G D S A V G M S M I V L A Q I T L E E E F E I V I N E E G L L A S G M
RANDOM_4244 Q W Y C E S V R D S S V I D G Q T T E E D N R K E S S P C P S G A T F P I T I S G E L T I T A A G N G M A K Q Q
RANDOM_4245 E R R Q W D D R T I G Y S A T P R I H N E Q E N P T Y R K L N S L T D L V D K G V E R Y S D F Q K I N P H E R V
RANDOM_4246 G R R E S T G P G V N Y Q Q L Y A M V A I T R Q N S G R K L M Y I A Q K V H S V L E K L T M G R K N R N V G D N
RANDOM_4247 R G Q Y V K G Q E T L F V L H L D K T S G I T F K K N S A T M G T K M C L R E T Q P A L R P C I P F P P L K A Y
RANDOM_4248 A D E P W L S A C S A A T F K I P D R H G V P R M A S S A K M N Y T F M A H W T E C A L V K R T F E D P N F V A
RANDOM_4249 E F A T D G K G R T E Q G T C C G T I W G F S K D G E K D L R C G D A I P Q Q L P S H L N P E L K Y G T Q F V S
RANDOM_4250 Y S A A L R K T F T L K I D I F G K R L V K H S D D Q N I P A M E A K L G I D S M L T A E K S G T L A Q A Q R S
RANDOM_4251 L E Q L S G Y E E P Q Q I F C N A S K V A K S I A T A R A S E F L G G V A I S T G A Q T W E N T K G S E V N S V
RANDOM_4252 N L R L Y P G A E F R G K M H N L N C P S G A S C A Q V A E G A C R G R L C H S P E S Q S A Y K A A D P S T E Q
RANDOM_4253 G W I V V G L R V P K Y E I K I V L L A F P G F R G P Y K N P T P R Y K V L K F R I V S G A R V F K I D N A M S
RANDOM_4254 G T V N K K A V A L A K P T I A L N I N W V G K P C Q D E L V T T L P F K K P G G M S T I Y V I R H A Q S Q P F
RANDOM_4255 D A N I S L Y L V H N A G D A V F K F L E L G F A V Q G L S V Y K L S V T I F V R R N A S M S V L S S L D E L A
RANDOM_4256 N K S G A L K L Y V F A D Y L K L V I P E L K D L M E G A A R A T S E T T T K S A S N V A G T L D L Y S F L T P
RANDOM_4257 L F E S K E L G P V S E N V L M N L T A K T L G E I M I I L E K L A K P F D D E A V S L H D A E T E S K T I E V
RANDOM_4258 D F T S A E M K P E A V G T R L E K W V V V E L E H S N E K E K A V D G T P G E T L V L C S L L Q A P V D K I I
RANDOM_4259 V D M V I K V C K I T V K Q Q M N S L G T A W F G S I S S R V R R L W E P A A I L K P K S D V S E S R T A P D R
RANDOM_4260 A E M E I L D E L I P A F R A W P H V G D V K K P Q S F K Q Q I R R S M A E L L S W R D G F K F I R K P R S D E

RANDOM_4261 V R Q G C K F R V N F F H E P S H I L E Y T S G E W Y P R L M I F L A E E D V T T A G E C Y I K N K L E N D K A
RANDOM_4262 L A S Y M I Q S G V T A V E L E N V K V P P Q G I K A R V S G H C R Y N E R Y G L C F R V E W Y A N G I M T I G
RANDOM_4263 L D A G W E E L C N T R A L K K C K L R P V K N P M V E K E T E V P R K E A T S Q M I F V I R D L H L K V S V W
RANDOM_4264 I N D A K A Q E D F S I E D H M D G G L V G L S D A E D I N V K L D D H A T R S R S I A M Y T S T M V P D T L D
RANDOM_4265 A P L T K M V R T V T A R N N H Q Q Q S K E N L T I L A R D D R A S K G N F G E N V I T V R V K H I S G V S F A
RANDOM_4266 A E A L L Y S A N P S D G L L W I F C P R P N T A M N A D A A T A G Q G L I D K D T F T C P E S G V K E C F R P
RANDOM_4267 S N G D V S F N F V S Q R D L I S V W E R L V N Y P H A A A Y R D Q N G G S G D S E Q W L D F I N I D V L C G S
RANDOM_4268 G K A E P Y G V T S Q R A N F G H Y R T E R I P M Y V N F V L P Q V L K T F Q I E Y A A R H N L K L D S K T L L
RANDOM_4269 N E N V T R P R G K E V C I L R K M N N H C M S D G A S S R L G L L F N G Y L V R E E S T M P C I D D I P A G S
RANDOM_4270 A P A T F M T Y L N N I D Q G E W S Q I C K A I G P Y R G D W N S E I T H N M T A D K E T A S A L L S N D L H P
RANDOM_4271 K E S E Y I R V F R E E Y D V P N Y Y P A T T L K A R L E D I R K G F K R D K K I L I E G T Q W N A A L E E G I
RANDOM_4272 K K L D N Q G R F T P V P D F Y D S N I D Q H S G T T A P D E L L A M I F K G S S K M K L A P T L G R R W K N W
RANDOM_4273 E S T Y E W T T K L G L R L V I C K T I G M V N P L L E S E H Q W L L N S D S C E E K P V Q E I T V V W P P C G
RANDOM_4274 N V Y Q C E Y E G I D T R A E L S Y V Y A R M T L T P V D N L S A S E N K K T T I V L L D K S R Q Q F L V S Y V
RANDOM_4275 A L R T V S Y G R D F E L Q F H T G S K Y N I V E P T L S K D N F D T S V P L W T Q Y E L S T I D G Q V R S L E
RANDOM_4276 P S I F Q R I E N V W T A L L P E L G L N N R H K G L P F L V G W W L Y R G A V D T H Y N I M S N R E V L N L Q
RANDOM_4277 M Q R E E S K I Y V L V L Q R P E L T L G G P P S T Y A Y M G T N A S D P H Q R A N Y M K K S T P T A K A Q G P
RANDOM_4278 P E G D G A S D E R E D E C C N L T P S Y Y F S L E N L L M K L D F M T E R L T F Y G D E I V C A P E E L C L C
RANDOM_4279 S S L E V F L T Q N G Y K S E T I I D W I A L W P S A L V K E E Y T L V T F K D S E A F G F I I S E G S D V K E
RANDOM_4280 D A P E F N L M A R V R L H G C K N G I T Q K L E G N N P T Q A P S C L P A E E D E F G W I K V W K L L A R C A
RANDOM_4281 Q G Q G T N L A G D V K L C S I G D P F S S V N L T S P Y E H N T K D D L V H L P L Q R I N T V R T E G M E V V
RANDOM_4282 I E K W D G V V R A I G V V E K P K L Q L E A E K A G R I K G G A P W V D I I G H L N S E A P Y R L G T R A E A
RANDOM_4283 E T E L R R A F I T K S D G E L W T Q M C S H N L G E N P F I N S D K C Y S S S Y C P W R P G C G L G K Y A V I
RANDOM_4284 C G K I Y S L W D A A V T L D R V H M I V T R R L L G K S Y G L V D F Q P S T Y I D D V K G L G F L D R V V K Q
RANDOM_4285 Q N W I L K T T M M L D V N G Q G I T L K V I N V H D S P V S V W A P G R R P N T W K Q D L G T L K P L K I A R
RANDOM_4286 K I S P S D L L R G R H N A V R G E V G T F E E N V N S E L R T V E K V N R I Q D Y P E K W I D S P F C A I L A
RANDOM_4287 T D A A D F E H A E G S G K R S G A A A G Q L P W V A R F K Y V F M Y A R S W K L F V I H G D I S L K Q T K M D
RANDOM_4288 D F E G Q E R A C H L D D L V V P D L N M Q K A L I M S Q E A S S I E V D E T Q I A N G S V D I A V V P A L H I
RANDOM_4289 L A G S T E R G P M V R S D L M R Y A L V T G A V L E R L S K D G D A R Q T I R H R V L S C N E S W K Y N D V S
RANDOM_4290 T L V T Q V S F S C K T K T V A E S I R V L S P A A V V N M A P N T S L N S P A D D E E V N V F S Q N T I F D T
RANDOM_4291 V I E D F Q L D E K L L R E R V P V L N F M L L V T F L K E W M S V I E I R N Y C T E R C R S I C T P I S F A E
RANDOM_4292 M C S E E N G I T I E G C M G K C E P A Q H Q L G D L Y V G E I S V N N R L N I M H Q M D L P Q T A E I A G F Q
RANDOM_4293 P P F Y F D P P D R W I N M R V K D D W T V T V P H I Y P R L D G I R S D I L V T Q K S D S K K V W L G V I A T
RANDOM_4294 E G F T S A T I M V Q N Q R A R F I E N K S T Q A A A V I W L A F T K V R G I A L N Q D Q A P K E R R V A K Y D
RANDOM_4295 L E H L D D V L Y L A F A E D Y Y T E A R I V N N E V P L K L S E M L Q R K P D A E L L V Y M F C L L E G K L N
RANDOM_4296 A K G T E G Y D T L R K M P A F A S G N K L N T A Y K R Q I D D K N L G N V T E N D L L G L V Q Q D Y E E G N L
RANDOM_4297 V M R H F S G N E L T Q N T I L K K E K F G A A L K E T L Q K K A S D Q V T E Y T T A E S S K D I Y K E I Y H S
RANDOM_4298 P I C F Q R A V Y G T E V N G P P L A Y G P S S A I R S L S H R N G E L F M V A H S D K V S E S L P R I F V G A
RANDOM_4299 G E T P L S I A L F Q C A R D C D G S K N L K G C S N F F Q Q T L L S G A S A Y N G T E T A M G F D H K P I A P
RANDOM_4300 H C D R I Y W Q N V T G G Q G I M K T A L I A P S K S L N G S T L L N E V H D K A T E P C Q T G V I R C D I L L
RANDOM_4301 A D C M Y D D D G K T A L F H T I G I T V P K R N D G V N Q D S F V A I G E A N F A E H S V H R V C A V F L G P
RANDOM_4302 R K K S G S E G T F F S V T P N L L Q H Q K S Q Q E W L E R E L L V P S I V V S T P S P C S Q R I K I S M L P V
RANDOM_4303 C S A R D G E A C V A L P E Q E L A N K A A D I G C E S C Y L Y N N G G G R C S R F D Q R K A D T Y G T S R V Y
RANDOM_4304 C D F D P T G P Q G V S S P R W Y R A E T G D G V G D L R I K Q V A A F W S A L P K F E V E K V N L P C E I V L
RANDOM_4305 P N P S K N V T P I L D Y L E Q R P A I G V T R T S I R Q N C W V W T S I T G M V K Y Y R V S D L W G Y W I L L
RANDOM_4306 Q I Y F P Y I S Y Y V G N K T N S G L L E N A V P S L A F I W P N Y T L N M N Q R N E K M T G Q R N D L A A R T
RANDOM_4307 Y Y V D E V E D A S G M V V G E L N G G V L P N P K E N L Y E A L N V Y I D R M V S D V R S T P A H R P A P P V
RANDOM_4308 S Q M M Y A N G E T G A N T L T A K R G S I Q G V K V Y T S D Q H E Y A V T L Q G S L K M F G L D E S T A M P A
RANDOM_4309 E R K C H E R I D W G A W E T D K I P D R E K R G F R A Y E V K L V Q I R L N C G Q I N G P L F R R L L K H E S
RANDOM_4310 D I E R H P T Y G P L T Y M T R A F S S N Q W D I A D L E M A D F Q I V S S P V E D I V K G L Y T F S P A S R Y
RANDOM_4311 K I L D R L Y Q D G T R E F Y V R G C S A A S D A Y L A A A F E G F Q K R M C P K V A S P M T M V L L L A I R L
RANDOM_4312 F K K H H V P T T Y E T L L D Q A F H C W D A T K T N L Q S P E H S V K N V T G T Q K I R K A R Y R A R L S V E
RANDOM_4313 A T V S R P E N F T E R R S S K D L K D I P T V V P S L M G A M L Q Q F M K T C K I Y Y T F E M F S L L P M F T
RANDOM_4314 V A A D V L Q P H L E R D T S A G Y G P I L Q T E N K G K M A S N Q A S D G L K D P E I P L P L T V D R V V K P
RANDOM_4315 A V I T R M L L N I V R L A A T F A I T L A D V T V A N S V I K L H L S F V L P L D T A T V T T K E V E I V Y V
RANDOM_4316 G V V L E L L V Q L E P Y G G T L F S P P L E D L D N G W E P D A K K Q D P M Q E K A A L G K F V P G G L D W P
RANDOM_4317 D N G L A V G K K G L D T E P S C D L K N R T E K E D E G P I Q K T S V A N G N I Y Y Y V G D L G V Q F Q D Y T
RANDOM_4318 Y Y T H Y L P L E T Y A I D S L V V P S R G N V Q D L E Y N T A N R V H P R R I D L P K A L R V T F L D T G F S
RANDOM_4319 R P F L F A G N L R D R L Y G V F Q C K Q Y T S S G Q G Q M G R E S A C G L I G S R P T Y F G C N I R N R A V T
RANDOM_4320 E P E V L V P N P P D A T I G V K K N Y R S G V A K L M L G T K A V S L A A D S G V S S G N L V Y W E F Y E R P
RANDOM_4321 L G K K N N S V F R D K F L K K T K V M K R E K S D F G Q V I K T T G A E E V A M E T A T S E P H W F W Y E D D
RANDOM_4322 L P A L R A L A I P Q D R T P E H H A S T N A F K Q F D A G E H G S Q S G G S C F A N S R S G L H W P K M V P K
RANDOM_4323 A R S D I T N F N T S A N G F Y Y V F C F C G F K L Y S L K S A V F L Q K L E A G E P G K A R I V I T P L S T E
RANDOM_4324 S V A P K T M D N V D K T G Y Q P V V D P A D T W P N I R S D I A Q F S Y P G R N F N T V D E G F G S C G D E W
RANDOM_4325 V D M T F S W G G R P K N Y I Q R A Q Y S P H F E R D P I S D H T D D S R I R T G M D A Y F L S T K V Y E A S R
RANDOM_4326 G G Y D Q I D L L G M A T N W D L P N S L M R I G L V N S P G S L G N Y A M S G Q L Y Q K T Q D N E H H F R M T
RANDOM_4327 V W H D K Q E C E L P T P D G G I I L R N W S A N S I S K R L P G M E E T G A F R I V A L N I Q R R R P D L Y R
RANDOM_4328 C T T E I V A S D N L T K S T R E T V G T V K R P T K T I I M V V K I M L G P Q L L N D T D G W R I S D I N P E
RANDOM_4329 C H G K G F E G Q E R M Y C L A V S P P G E D V T L D S G G S G Q R V D V P L T G L M L Y K M S K S A L I E V P
RANDOM_4330 H L A V Y F K E Q L N I A R H D Q Q R E M E P A Y A L L L E T A N P D D R T T V T G S G T V F P V T Q R V G N I
RANDOM_4331 G G T Y A G T D R K H L R R S F K A V S L L Q T G L E A V G E V E L S D S P A K P N G A T I I I A I P T G K D A T
RANDOM_4332 G F I P V P L D G Y L D I Q L C S D T N F F T G P R L T L N K K K R V T A V L R L K L L S L L G F P L S T I I L
RANDOM_4333 W R A L E P A L N T G H P T L L E W A S T C H M A D K T L V K D T P Q V E Y V G L S H Q Q S A T H S N S F T A V
RANDOM_4334 V A Y K G L G I S S L T D V N L A E T V W L L A V D L G I K I G P G R N K S A A D H I E N K C L I V T V A D T E
RANDOM_4335 G Q F D F K Q L D D V Y K T P K K D P P P R D F G A W P Q P A I I D V E R S N D P V G Y V A F T L K Y S G K E Q
RANDOM_4336 F G L E K N A I D R I V S A N D D Y Y K K L A D I Q Y M V R M K S A A K T A P V A G K S I Q P D T V E T Y G R V
RANDOM_4337 K V K H L G L F L T E Q S Q G T E L I A F N P Q Q L G S E G F T R S G A E N G P Q K E I C L I P K E P G D A M Q
RANDOM_4338 H C V R E L S Y G I Q P R Q I I G Q T G V I Y V A A S E Q L A E I Q R G H C D L S R T A D R A F A E R D C A F I
RANDOM_4339 K T G G R G F T I V D L A M L N I Y N L S P F I R L D D Y S S L T A K L T T F A L D P A V S T I S P E I R R K L
RANDOM_4340 R V M L Q Y T A H V T T C G E N T I P K P V I R L W A D L Q V L T S A Q I S C E N S R Q L A L N E Y M D L P S Q
RANDOM_4341 D N V R R S K A G G F E C D Y I K P N Y V C N G L D R N Q P V R C G Y E P A R P E G D E S Q N T L C M E G V H A
RANDOM_4342 K L K T P R I S R Q E S C P K I G T R V V S K E F S L T G Q Q T N D D G K I F M V C I P S S V R I S E Y D N K Y
RANDOM_4343 V G E A Q K I G K A G F A D R Y G T A H K L D A F M Y L P N N C L Y L S D K F K T P S F D L L L F S A T K G E S
RANDOM_4344 Q F M P G K S G G T G R S Y V I S T I C W G Q I E N I H D S P Y T I N T L P I L S R P G A Y P K R S G L F D V G
RANDOM_4345 T G M R D I W A L K H G L G S L Q A A G N P P K R A V S A T Y K D I N Q I A G I K G F G K K Y V T A E Y R S S A
RANDOM_4346 G V G G F A C S W F P H A S G S A L E R G G T S L D F P N K H N R A Q V M M V S A T A A E Y I K N S E E T S V I
RANDOM_4347 F V P P Q D G F S H T A L L M A V F S C P K L G S N R L H E L G S S T D K S H A S R G S V T H Q T K H G P G G W
RANDOM_4348 L A E N E Q L A V S P A T N K D L Q A E L T G A Q G A H S T T A T S P S N Q A C A G Y G L I V P K F Y A L Q F C
RANDOM_4349 L V Q G K A P V D M T H A R S A T R R P E K E G N R I L N V F R K P A S C V K K V L F K V R D P S Q E P D L A G
RANDOM_4350 Y N P Q I M K Q P T T C G D G P R Y G T C V G N C R R C A F F L E W E S D N K G E L Q P T E Y L V Q N S S M L V
RANDOM_4351 D V L G H G S E Y D L I V A F G K Y N A N W G V S V C G F L M T A N V R E N R T G Y S R N A D W V S F S L S W E
RANDOM_4352 A L V L W K Y Y V S Y W H D Y L N V Y M P I T K I L P S F E F I Q P W E G S T P D R N V I R E V Q E Y W G V S E
RANDOM_4353 V R N G G G A A I D T G H Q I T L L P D N L P R T S P I P G T T M A H T K L V T G L D A P A G I S L G T G V A A
RANDOM_4354 G G V R D S E C A V G G L K D P A L Q P V H G R Q F E K M S S G P T N S L V F Q N F E S E D A Q K T R A Q H D G
RANDOM_4355 F D S R T R A E S M M S A D N R Y V P V G R I Q D W K L L M T N N Q L D P T G T K S M G L P F K G G C P N G V P

RANDOM_4356 I Y K A V I L T R R A E F N R A N R Y K K L I C L A Y E Q K V E F W N A A T H A F T P G S A Y V F E I K V K S E
RANDOM_4357 S L F P T V R L M A L R P L K P P E F T W A H D M T K K S F N S I S M A S S L F I C G A L V P R S G E G H C T S
RANDOM_4358 T P E G P R G E A S D S K C I F F R G G L A Q I Q K P K V S L L Y G P S K H R G S L Q S S A R T L G D S K K G Q
RANDOM_4359 D I A E P Q V Q I C R G S D L A V L V Y G G F K A T A T Q Y M L L A S P A T H G A D K V D C M G Q W K E R Y C T
RANDOM_4360 V A P A P F A D C R G D G H I P G A P D W A I E D T V D N H I E P T N T I G H D H K R T P K L N Q A E L A D Q S
RANDOM_4361 E E F L G F E G S V L R Q S Y R K G K E K A E L L G G K N Y G L K V C S S A T L H Q H V N V E M I G D F G K T E
RANDOM_4362 L R S A Y K K N G D G R M A V D I S D R T A Y S L V Y S L W D R N P L G S A E T E L K N E L L L G Y E E S P D V
RANDOM_4363 Q V A F F L E R N Q I T A C R L G P T L D T D A A Q N V G Q V H L E K Q K E G Q V N I G V S G A S Q V A G M N K
RANDOM_4364 V V L V L L I E T D S E I D T C P A F R E K S F K G N D G K L S N D F S S G Y I L P L E K K L L S A R T G L V V
RANDOM_4365 E T K G E G P L S H A S D A E N K D P G T K V G H P E E R F E D R G E E Y C Q A R N K N D P S I L N S I A K R I
RANDOM_4366 R A S G F N A R Y H D K F A A G R A T F K Q P F L V G D A H N L E T E T L F A L G G L A Y D D A S P S S V G S M
RANDOM_4367 I V T I R A P P P L G E E C K A A N P A L N R S M G G G D E A S G P L N N P T K R F I T G K C N I N S H N K P P
RANDOM_4368 R M D N L P S S E M T A T L G I T Q K G N A P L G Q M W Q V Y K P A K L G C P I A F P H G A V Q E M G T K C D I
RANDOM_4369 D R T E L T L T L H A R R V S A L K K P D V E G Q P F R V W I S K L L K A P K A S F I F H K V S V G Y V A R F N
RANDOM_4370 L R L A E Q L D H K W P C I G V A K D T E R L S P T A K Q T S K S I S E G E V L D H N E V A T T A K T P N S P I
RANDOM_4371 C K A P S S I G I H C I T A K G L P R I N R M G N G D L I K G R E I E T A L D V I T S A L D M I Y K K Y M W S S
RANDOM_4372 F A E C K A T E T P S N K K E F P H E T L P G Y R V S P R E F M K K L C S N L R V H A I G G N T L D A D G E L S
RANDOM_4373 K V N P C E C Q D R L V Y F E N Q D W A K E G V H G L S M I Q G V L E D S L F Q P E V D V V V V L D P A E V I A
RANDOM_4374 L T F T D P D D Y C T R E L L W N S P E Q G V S A K S F L L V L C G G S K A P P T E N I W A S H R T A M R K G D
RANDOM_4375 V Q Y K E S N H F M G G V N Y F Q K E F L W T H T G L E P N P N T P L G N F P Q G P T R R Y P Y F D H D A N G A
RANDOM_4376 D Q A I G S E R Y V T D G S D V D S I R I C P A F Q A R Q I T T D L D W A D E E N N T G H I E I N S D D D G D A
RANDOM_4377 G H A T V N L S Q V S A T A V R P N L N F V N N E W E R V S P A T D D I V T V Q G T L L L C E L F G L M N G A V
RANDOM_4378 S D E M W P E S T L I P Y A S M S G N P L P V E F I E M T Y S T T S A D A L L P D K D Y E E A T R L E E T I R Q
RANDOM_4379 I G Q E L L N G G F Q C A A R D L L N L I L P E V H R D D M F R S E N G E Q S G G P L A D L C T G C L E E E T W
RANDOM_4380 I G P I P W I P W F V L K Y Y W D D N I V G Y N K D A S V S G L R R L N R R P L G I S T A G A K E E E T T C S I
RANDOM_4381 T S V S S A V M V P I D G G K W C Y G L M N I Y K A V Q M C K V A L E I R F T M T Y W E Y P D P M N I K D S L H
RANDOM_4382 G P R R E A E V M D Q P V E E H N G W Q H N N L R A E E W A L H D T Q G Y R L C T A Q E I L E G Y L C Q A I I S
RANDOM_4383 L P E N E K K G R R D S Q K K A V D T C A A S A G S T T S R W D I K L I S A F V Q L D I N H P A E P A E V S G N
RANDOM_4384 Q L E K K L R P E D K D Q E N I T G R Q C L S K A V D R L W N T A Q S N K I T K D L P H I L N P A S A E S D L L
RANDOM_4385 C H A L F G P A L T P G S R L F N F M L Q T V F D G E P A M Q D M G Q S I P A A L I I P S P L R L Y T E D Y I G
RANDOM_4386 I R F V V A Y E W R L N A T S R K K S G A L F K I S M L V V E F H V L L E C R R Q L H S L F P D P L M T L V L H
RANDOM_4387 Y G L G V A S S T C F S V S E Y C A Q D G E T I S Y C D M S K R H Q K F N W I L I R G Q K D S S G G P Y V L P A
RANDOM_4388 P F Q I A T I P L T R E R L L K D P C C E R N D I K D L M V L R A P E N N C G R M G N Y Y E T S T I R P A P K A
RANDOM_4389 Q A V A M G M V M F T A N T A A S P A V T E P T Q C C A L A T P L A K L K A A N A G Y T G C K K R M L P G A A A
RANDOM_4390 T A V M N K A Y F T V F G S E S G P R M S F D T A N W Y P Q A T S L L A D T E T G R P Q A W N R Q H N M D K N K
RANDOM_4391 T R R V V N E T E A I C L G Q A Y N Y N N Q A G S D L P S A P F G G I S T M A E N D G A A A D S F L W I T L E P
RANDOM_4392 T D T N V L T N V K P I Q L K T Q L R G A W K G N E A A F L G T I H K E V I L G V G G T L F H K Q Y V F A V R S
RANDOM_4393 L C P A P V D K N R R F E A P T E K G G K E S I F T N T V Y G F T L V S A D H E K K F L N T D Y E L D G I F E S
RANDOM_4394 A M E G G N Q R D V H M N Q P R G A V Y A S Y D T V V L L D I M F R V E A Q H G Y S K S L S Q S D N Q D R S D A
RANDOM_4395 C A Q A G S K L N G L G N S E I C L L M K I E C P F I F L Q I Q A S L L S K L A L G R R T T P I W D Y R G E D S
RANDOM_4396 P E G K F I H R M K L L T K S A S V T T N P W N T I N A Y L D L G P R E L V V Q A K V T H G L G Y A R S T E V D
RANDOM_4397 K A S E K T T N R T N R F G V K V S G S Q T K G N M I R D S E G S Y R E I P T V F M V Q L M P S C N E Q V L R T
RANDOM_4398 F D G S A T E L K N D Q R V Y L Q V R L G G R N P G G F D F V V S I E E E I A N K P E H E P H Y F V R L I Q V G
RANDOM_4399 I Q F L L R I Q S T Q G Y I R S R L V H T P A G L V I H S Q G K G I G E I Y S V V I V M A G M L G S S A S M R L
RANDOM_4400 C Y K H V H M W A T D P V E Y E A G Y I E G E Q E N I V I G S T G G P D E L D T A V L H R V T D V S T E V R V T
RANDOM_4401 H L V L F Y G S G L G A P Q Q P N S R V G P F I E G S W G L T V M H L L D I T I Q L D R A E D A V V K P E V I G
RANDOM_4402 P L E D S A A G H R D A S I A I V E G G K R V S I K A D G V L L L G R G A S Q L Q K R Y I L S L G S E A S H Y Q
RANDOM_4403 G H V I Y I N I A E R V I P S C I Q G F F S L G Q L G Y E L W M K Y V G W P E R T F V S L P K S C S A K R L V A
RANDOM_4404 V L W F I T V G E S D G K Y N V S G W L T I G L R A P N M N E A G K P T A E E R R K M E G N R H L M S G K W T T
RANDOM_4405 V Y E D G E A Y G Y A I V G G P T C P G R R L V V T A A Q Y A I F K N E K Q L I E I L W G V T L R T S H L I A F
RANDOM_4406 R T T Q L A V P I N T L I C R S W P R R K C E D D S A D Q E T M K I D R I N A Y P G N F K A L Y K T A N G P I S
RANDOM_4407 E A S F G N Q P A T T T S R P A T P L G I F T D V N A T P K S Y W V L S M A I G L T D T G L L N A T L T M R A G
RANDOM_4408 M V Q I Y L T V F V R V E T V T Q K P A H E Q S G A S N F C Y F A G K F M K I L P D K E A M Q K L L V R G P G L
RANDOM_4409 Q T C V G L F S H F M G N K E L K E K N A T V R L K N E D Q E R I Q D A S P T T C D A L S A Q H P I S N T S H S
RANDOM_4410 N Q S Q G K R A K K L N L L V E P E F K N E D Q V V G R F M L H T R T D L D A N A P C K A E E Y K M A N D V G M
RANDOM_4411 Q D M A Q Q D L T A Q T R Q L G V Q Q V A T A F V D R D A W L Y D P E V M V G I Y S I G E K A M Y S M L V P S Y
RANDOM_4412 K Y F S C G Y G V D I A L G I E R T V A P D D A S V I I E R V D G E I K N S K T F D L N R P Q W N T D D L N D I
RANDOM_4413 I N L T A L Q G R V F R W D S K S L I G T L F T Q K S Q Q L M E S G L F R N K F L Y R F E A T D K E S A A L D R
RANDOM_4414 E S A I V T L S L S Q D V I S K D M C S E I R G H V D A D M I A I F D L G L P L N E A R I Q Y F N T I L G G S Q
RANDOM_4415 R A E M P N R G K F E S L V G R E S G Q G N I P K V K Q R R G L T F V G Y N E G D Y L Q E C Y N M C N Q Q E T A
RANDOM_4416 L Q S E V R V M S A P S S H F E I T T G G I I E S I D D L E D I I A L P L A K D Y S A Y E R A T C P G P S I L P
RANDOM_4417 Y L R M M L I L L V H H P F E H E A V V M I I S Q N A L Q N E V E F G D F L A H I A A R C V Y K M H T V I E A S
RANDOM_4418 S A K A G A V L I A D N A T R I Q F E A N Y P P H V Q K K H L V E P D H G D L W D K L R Q R I E F K S P E V L T
RANDOM_4419 Q W P Y K T V A I V D V A E N T L T L E P I Y L R E H K V V T S Q K V I G P P V Q P H T V P S M R S I R E S T E
RANDOM_4420 A K P I A K P S K G Y A L G G P Q L N F L A T E I N R G D F E M K G Y R T F G K F Q Q T F T S R I N S H I L E F
RANDOM_4421 S Y A P L E Q A V V W R F G M L D T D D F V T E S S N R L L R L E L I R H D M Q V K I T V G S A S Y R A A F S L
RANDOM_4422 K F V E I T S R S S T V F L I K K V V G V Q T T E P N C P L W L I Q D Q R G G C S T V D Q V E Y C G G S L F K V
RANDOM_4423 V F A N E Q C K S L A A A L W R I I L A A E G V I H S R I D K I L P A N S L Y T T A T A G N E A V P M R E S E L
RANDOM_4424 V T Y G A T L T G P Q L Y E L P S M T A D Q G E A N G Y L E I N A F Y L G G I Y A C M D M T N K T S E W E W A H
RANDOM_4425 I C A N F G T A K F L E L S K E N E E R H S I F R N I E Q L E I L G V E D V I D A A E S G E L S T G P L N C K L
RANDOM_4426 N D P A N S G E E V M F I F A S W S I Q S S P A K P D D P P E S F Q G A K G K T W S W P H F E N A L L L A L P S
RANDOM_4427 Q L I A Y N N G P Y D L Y G Q C H Q E S V I S F R K V P T K A D I S L V Q G T G T Q G S F E R Q R K D E G V E W
RANDOM_4428 I E G A S M N R P S D E Q Q L L S V T L E F A Q K V E P N T R D L I K I V E P V S F E Y R R P I P D L T V H L T
RANDOM_4429 A A D K L H L T G R S L A G F D L F L T A K A I L R W K E D V S N K D L H W A V E I L I Q V T R C H R K C C L E
RANDOM_4430 G D S P H G T R A S L D A T K E V M N A V I M T I C W A I P Y F N V N N E A A R A Y A V L E R A V K G L A R G G
RANDOM_4431 T N S R M E P I A D S G F G N K G V D D E G E V V I S K D Y S A G G T P L N Q L T A G L L I K A S V D A Q L Y F
RANDOM_4432 V H P W A N M K G A K S S V E N F T S Y T S S G D W T E L R G I D I P F P G Y R A M P A L F M L S K Y D Y S G S
RANDOM_4433 V E G L F S L G A K D G F T S A P T K F K R P L S N Y V N S T T K S E T D H K Q R E S P P T L Y N N P E L A A P
RANDOM_4434 L S N R F E C R S V V S G I I S F D T S E K G A R V F V G G Q E Q P D Q L V L Q K I R Q A N T R E I T E Q Q K V
RANDOM_4435 L E A D I K P Y D Y L H P P E Y L M N K H Y P A K Q Y N Y A K A S R Q K Q G L P L Q P W A R N Y N Q G F E W V A
RANDOM_4436 E R G R C F K I S T R W A Q R T K L G P H V S E A E G R K T S A S S L A L S G G F K V G G V P L C E A P Y G R P
RANDOM_4437 K G F H N S E D I K T G F L T H A A I D G P F L F T K L G H V Q K P P G P F M L W S Q F L A E L A K I V T R N N
RANDOM_4438 A L A A L C D E D Y I F A A L A V D G A P A Q K A T T F E P N Y L D A Q Y F H C I R Q G D I G P S V F A S M V V
RANDOM_4439 L S I H C G S Q H R L G P H G V H E D P D P S P V L A S A W E L S P F N G S A L A T Y E N F V L G Q S L S C V S
RANDOM_4440 D H S A G G P I G E I T D I G I L H A K S P E L G W R D S G L Q E R T V S I K Y W L A N E M P K T Y L A V C R K
RANDOM_4441 V I T M A A S N V F R V R I E M H F K M G M L E W S A L D K L K F A V G L A E Y E S K P A G V V V P S K K G R G
RANDOM_4442 L F R S K D K V K K E R L V Q N D G Y V N L R S R L V A G R Q V L T M I L L Q L K T S S L A N A R G D C E D D A
RANDOM_4443 Y K L S N A V H S S I K Q L F N R Y Q L S N Q Q C Y N S F A A D K P A W G V A Y T G R C S A V V N S L A Y S S L
RANDOM_4444 Y P H C L V K D G V A T L C N M Q N H G T Y N L I M T A S V K L P P I L F G L V D Q S L T T P A Y L S P V K M R
RANDOM_4445 T L D A G G Q N Q L R S T T C I F A T L C P G C P A V Y E T T Y I L S G A G A A T C G N E D G G I S V Y E F A L
RANDOM_4446 Y R G S E F N A K A V D T P R F P R I E A S E W V T S T E Q T H N S A K V H S I V L N V N Y K S T G Y M M D P T
RANDOM_4447 F N S S Q D H E R Q K K C S Q I A G S R D V G G V I L L I F Y L V V G T Y N F R Q N D Y P D G A C A V G S R K F
RANDOM_4448 S P E G S E A K A D C C A R H G H R Q V T Q L A E P S W L V L T R D E N I L N D F G H P G S L M C Y L D G E G L
RANDOM_4449 T I E T A L H F T N S L P V Q R S K T T V F D I F I V H Q L P L F L F N V P S E I H E T N D G I L R A E L L K F
RANDOM_4450 A W W V G M L Y K P P V W Q N F F V W R V G N G R G H S Y Y E V E N C D T L G E K K N I K P H I M A G I K Y P N

RANDOM_4451 S A F I R A A K L G C P S I F I S A S I C S Q M E V F L V V E A V I V D H A N L S A E A K V Q A V N H G K I D V
RANDOM_4452 D A E E P S A E N Q D R I P K F T E V Y R V E Q P M L S I H T V A N W Q D Y V H V G M L C S K T G T D T R Y G A
RANDOM_4453 N L L T A C T A E G E F V D G R N P K F L E A V R D D Q D P I N V K S G M L K D P D R N A G D K V P H G R E V C
RANDOM_4454 F V V T H A M W W F L G F I G K N G S L A V T K Q F L N H R K T H F K E P Q Q I E L M E G A P Q G S Q T Q V Q P
RANDOM_4455 A F W R Q E P T P D I L I F C K D Y E F I Y W Q F D K L D K L P T L Q N D Q N E C Q D L F G L P F L S T L G L K
RANDOM_4456 A F V P D R L A E I L S D L M D G G R D I L L E E P W I S P K V K S P V T I I K S D R K T N F R R V K K A H F V
RANDOM_4457 G L S F Q F F Y V S L F T R R T E T I S S S R Y I L Y H K I L K P E H S G Q I V L F A R V H D M L L P V Y R G F
RANDOM_4458 E C Q G A G V F T K R I G A N R S F H F L R S V G S T R G P M S E A T L S D K K S S L R K L L V F P D G A V S E
RANDOM_4459 L C Y S Q L E K A F A K Q G L I K V G F V A P Y N Q P N F V R T T A I E G S A L L K K G M F F F G N R V I L W V
RANDOM_4460 S T Q A V N V V D G E T D A P A F E Y A K D Q M V T L L V A A D L T I H A E R A K F T K N K G M D V R I N Y R P
RANDOM_4461 N N S L R Y R T A D D L C Q K R S S H C A I V L S A M G M T Y V S R D G L G A L S L A K Q T A F L A G D S G N T
RANDOM_4462 D D K G A L V Y A T V M S N T I A F D E G S A I F G D Q V K T D K L A E E F I R I D S T K N M V L F D T G T I M
RANDOM_4463 E M D D S K L K N Y S G T K V E V D G L V E E A S R F D M N H A S V V I A S A S L P Q L L G W L A R Q N A L L W
RANDOM_4464 Q V K T V L G R Y L P A M A S G R F Q E T E K N V Y H M D H M M H P R V E G K V R L V T F C S D L Y G W E Y G P
RANDOM_4465 L R Q E E S T H F V L S G S K F T K E D L V P Y V Y S G A T N I F A A T L V M V G L S E Y A W E H E F K K L S V
RANDOM_4466 L R H W C L L E S P L K G D D Y A Y L R R N L V A V A S N E R N Y W T S V P P T L L E G T G Q Y I R T L G S K A
RANDOM_4467 T S E G L P C L M D K M A S Y I G S S D I E E D A A A D S N A D E H R T I T L D D N Y I L S L P G V L Q N S D K
RANDOM_4468 A H L Q T V E S V G N Q L E K G V D L L G G A S P I L C E F N W S V S E N T P P H S A I T T S T E T Q K K S V I
RANDOM_4469 F N A G N N R K Q K L T S T H T F L I A G Q P V P D K T T T K L S N S P A N F L D I S S H Q V D A M Q I V G A T
RANDOM_4470 D C G E V N G P E V E Q K K E V D E R A D S A Q E N W Q D G L T K K G E C A N Y A G W R I A R A L R L Y I I G V
RANDOM_4471 N E P D S G R L G D G G T I A A Q E S T A D M D R Y D V A Y G Q T V L K V L N R F T G I K E V E H N G N K G Y Q
RANDOM_4472 Y N R I L V T R R P T I D K R T R V M S S S H G T K Q L D L P V Q S L L C P G K V L R A S S S P H D L E G V L C
RANDOM_4473 E G V P L G A K D L S T D L V N S K S R K S Q F R F V E D F M C H I S A S L P A N D T T Q A A D L E E T S Y T G
RANDOM_4474 A G V Y S I D E G R L V E G L A D W K G K E F D R P D G G E T L I K K V K G H Q P P N A R C L V K N Y A E L W T
RANDOM_4475 A T D F Y I R V I E F K L T L W E K L W L M S T M L A G Q T D K G S G L Y V G S D P K L T H W V D K Q S S A D P
RANDOM_4476 D K S P G S Q P S V D V S K G R D V T G R L N K D W S D K E S G A E A P T K K C E E G C Q G R I T A K E H F N I
RANDOM_4477 A P L A N S Q D N H L R E G Y W T V P Q Y T N G C T Q S S I Q K C W G V I R K I S S K F L V Y L P S N K G C P R
RANDOM_4478 L F D D P K Y S V G W T L K G L L V T H F L L W T N A E K K E N T G Q K G G V L G E A L Y I C N E H D K T D Q D
RANDOM_4479 T L L R V S K L R T Y P I V M F I N G M F G S Q T A L G S K E M R W S G I S R S W L S V P F A T P D K L G A V F
RANDOM_4480 L K W R Y C L A R S G K F G M D G K E T L L S G D V P G I G R V G F Y L G P K L Y S G R A L Y L A A T D D I S R
RANDOM_4481 E A S K F A E G K Q F P L F T V D L T G A P N E W G T L N L R Q S S E L V T A I S A A V N I I E Y K Q P V T Y L
RANDOM_4482 Q S A K D E A F Y V G S L L C T S E P R E V R V I K I Q C G Q Q G R D S T F F D M T V K W A Q H I L R N V L W L
RANDOM_4483 Q D F H S S L R S C G A E A K T G Y I V I G I A F N G Y E P M A G E E K L F E N A A L D Y G E V S Q D R A V D K
RANDOM_4484 L F E N N L P R L A I N S Q I P I K A I G H L S H I N N A V A I R Q Q I S R L I P S F K A V F A L G G S S T L H
RANDOM_4485 I V T M N F T V Q S M T A A A F N Q L K N I L Q K N N P L Q Q L I V Q Q Y L S S N G D K A V F R M K R L W L T D
RANDOM_4486 M A E Y N P K L V E H E L A C E S L A S V G K R Q I Q K G L I K F P H E W K R D G D K G A D I L S L D I R A K H
RANDOM_4487 M D P G V W I S N W D T P Y L I P N M V S G Q Q E Q K N F Q H G V R V I G I M N T E V S A P F E L P G Q L L V K
RANDOM_4488 K R P E T W K E G V A I A F R T Y G Q F E H C I Y L V K L S T I G E S Y P A T G G L V E T Q R R T G V I N K G K
RANDOM_4489 E V R L E P D G G V V H S S D P S D A M G L F R L T R F K S L S A V R L E H E E H I P E S A M E L L R E G M P K
RANDOM_4490 Q Q P Q N G I A S C F L F L P E L Y F D P I D D T G E L S S Q T P E G V L T L F A E N T S L V R L P S S D P S K
RANDOM_4491 A V C L A G M S E P D I K E E D D Y R V S N R D G T M A E G G M H N L Y S K F L N T P L R C E K M G E M G D G A
RANDOM_4492 N D A E D L T M Y D G S L P Y I W A Q M V Y N T P G K D I G I M T S S L F P P V G Y F G K S L P F V H E Y I P G
RANDOM_4493 T R T T F S E C Y T G L D F G K A E L L G E F T S Y V A K F N L G A F P N V N K S S D V R V P Y Q A Q Q V N P A
RANDOM_4494 I G D P S D L D I I T N W G S V A L M V R L N L D R E T C E V V K T E Q P L T V L R T R K L S A D R D K K K R S
RANDOM_4495 I F R G V R A L T L R P I V R M R M I M T G G A V F R L N Q R Y T W P D L K L L F S E P N E V M I E V S K E N P
RANDOM_4496 L L T R L S A N W E I S L E L A A Q Q A A E E T E A N Y K F Y R L P L D D N A T W Q W T K I L C L N M I P Q L L
RANDOM_4497 L G L Q A M D E L P V P D V L A H N D A L N L D V A R P Q N T M T C L L P Y W A S N L I A A A N R E V V N I Q L
RANDOM_4498 Q Q G G D T M R R T R E P C Q F E P T Q F I V Q A S I N T T S V G T I T S G A P H D K F K K K K D D G K G H T E
RANDOM_4499 C F E S D R E L M N S Q S N Q R C W L L K Q P P W V G S R V A P A G N R E Y P S A P T Q D N E C V L F E S A S L
RANDOM_4500 T G L Y I K R N M E T G S R G L N D G G T E I M P D V S R I S R E P Y R N S D H A Y L K E E F R A S V S T S I K
RANDOM_4501 K G G Q G M G A S N E G L N L L V K P Q V S V N R P V L S E A G H G T R Y K D K G H L E M E P T L S I H N K T T
RANDOM_4502 Y C G E K A C A A G D A T K A T R P R A L A G S S D T E T I M L L T S A Q R V S R S I D D A Q D A G C L V Y S A
RANDOM_4503 L V W W S D W G G E F S P R R K I I N Y H I Q G L G Y K T V A Q Q K G F K R E A P I F K A M L G T S N S L T P E
RANDOM_4504 L Q L K D P Q S T W F E R Q K G T G N Y S R C A N A K K I I N Q Y D A D L F Y S P S R A L R A A G T A E P R F S
RANDOM_4505 L D L G L R S K E P V D P L E A P E R N P H K H A I Y S L G V D I N D P V L H W H P V A K G T T V A T Q E M Y L
RANDOM_4506 P T W P E L V F G N G N G A L L T S R Q T S A L A F Q L T L Q D R A P G V H G L S V R M I R S N T W E V W S Q N
RANDOM_4507 C M F E T M I L R F D A I I K E L S M Y N I R V D R M T D D A N N E F I V A F T V S T L A G T G R Q S L G R S V
RANDOM_4508 E G G P F T E D F C T I S E L T E E T D Y A T L A G S T R E A Y G D V I N N V A D I Y L L H S L Q A I Q L G L T
RANDOM_4509 Q L S V A I D H A V E Q F A M L V Q D S V L R M K G H K S P K F G N L Q T I P I T G Q L E Q T I C V H A A C P H
RANDOM_4510 L G I N R A S I T G Y D S G K E D N N G K I Q V G N D L A G R S R S N T F L R L S Q L P L K D G Y V L C V G D K
RANDOM_4511 P L I V L D I K P A G G G S D K E D I A D Y P E A R P C M E K G S C L S K I V V R D Q Q P G Y D S P A E A T Q N
RANDOM_4512 T A K S L Q K G S L A L L G N G G N D A S L F L T K K K N C S W I L K G Y A E L A V Q I N K F E T Q E P A P I G
RANDOM_4513 N I S E C E G I K L K D H C L G A L R S L K L Q F I L M N D E R K R H V N G I T S V D T Q F L S R L L E A A N K
RANDOM_4514 S N S A Q G R E G P K V N V Q P Q K P F I E K L S M E C S T A F A V E A G E I E G I G R S N R A R A D F R A C V
RANDOM_4515 T A R T V T F P D N C K Q E R E L G G H A N Q R H K N S E W L Q A Y G T A A F N Q G R D G S E G F W I A S E V L
RANDOM_4516 Y S V R T K R S K I D Q V H G R E R V G R A S Q S S G P S N M R D E C K K N A P D L Q L G T S E Y I I C N V N G
RANDOM_4517 N P K S L N A A L H E R M D L K I R L G A D E I T Q N C Y N Q M A A W R Y P D C N S E K T R Y Q S L Y A L D Y A
RANDOM_4518 V E L N V Q K D Y R I Y Q L G L G N E V H L D Y N A N S I Q S F E A E S E S A I R I Y G L N N I K Y Q E A D L I
RANDOM_4519 C N Y M S V A K P A Y P N K I H S S N Y T T I Q G V S Y V I R I G S S S D R A N M T F K L Q L S Q L T A E R I L
RANDOM_4520 V A L N V V K E A Q E G A A T K Q V K E S S S Q D N T C C A E G C G L Q D L M S D L H D E T D W A N V V S T H H
RANDOM_4521 S R G D I V L F F G I K G D R T E K Y S R Y F C G R F A M R E N P G V E D T G L S E I D K S W T M N A R K L I R
RANDOM_4522 P F L H W L L D V M K Q V D E K G I K G R L S A T W Y V P S E A L T D V D I V P E G S T T G I G F V T E N C D S
RANDOM_4523 R V Y L L L A S G V I H A P G S V Y V T E D S L L Y L V L Q K Y A L L T K R A R I G A S A L L M I S G G A L F S
RANDOM_4524 T P Q D P K I V M G H E D E T R N L Y V L E L Q F A M T K V A G R R V S L A F A F R A C T K S E D V K E T Y R R
RANDOM_4525 C K D L K K R S L V N A K H E S T Q E D F S P T N D G D T G I V G K F Q N H A E D L P E G G G I D E L S K D H W
RANDOM_4526 S N K Y Y K G I I N E P Y P N L M C I F H F G V K A N F V F V V A T P A V F F T L T Q D S A D G M E F N M G Y Q
RANDOM_4527 V H N N I K E G H G G T D P S F V S P I R V G V F V E S K I A E E A N G D E E S S M D G A D M W D I G N L I P P
RANDOM_4528 G Q V C D D G Q N Y V R L Q T A R L R G A L E D S A L I L S Q I Y D K K D W K A E S G D R G M G D T E L V K G L
RANDOM_4529 D R K L N E A D F T R R C G V L G G D M L P V R P A S P V M N D N A G V A Q K F I Q L I N R P D C S A W V P E G
RANDOM_4530 L Y P V E I I K K H P S A G S L K S T Q S S A I A F S G D P N C D L A P C G A P K D S M D D T V Y Q Q P A S N N
RANDOM_4531 G K R Q L R Y Q K Q A L L D L G K S L M S D I S V Q L P L L A A L A R E G F I T P I G N Q E Q G A D E A C E L L
RANDOM_4532 N L S L L D F Q P T Q I P N N M R V L E R G E N G K T S N S V F K F T P N D S Q V E K R F G I S M L G L S C V Q
RANDOM_4533 H M H A Q P A A T L S R I S V Q W S A M N S I V I D V P Q V S A S Y C R G H Q K K E R N T D W D T K P V I F L N
RANDOM_4534 P V K V E P Q K L E I E D K R L G V G L T L L L D G A R K D R C Q D L D V R S D G A Q I V T K H L P K P G S G I
RANDOM_4535 T L L E W I S A H F F G A L L K P M L P V V S T G L S Y I N L A K E A P R G L K G G I D R N R E A L I A R P G A
RANDOM_4536 T F L S I E V Y F A R V R L I K Q I R N E D G F I R F D L L D S S P Y V L L D P M P V R M Y W S L F G R R D E A
RANDOM_4537 R S D F A R T E K P K A G N A L G A L C G T T V I Q A C F P K A Y A G H Y G I P C I C V N M K T I E P V V Y R E
RANDOM_4538 T R M T A S M Q A C L P V I Q A D R F S S E S T L R Q E S T L V E V A V D Q V R E K T D S S N I T F I F L H D P
RANDOM_4539 T E V L T R F F G G Y I S A A S A Q A A T P W N N A S R R K K H G E V D K M P D L D T G Q Y V L Q T E F V N G K
RANDOM_4540 C E N P D E V K T P K I H P A S R L I N I V F L A S S Q V D A E G I M K A F V R E I K G T I Q Y K S S E Y F N P
RANDOM_4541 I L M Y F M Q N I I A D T K H E P L Y Q E K A L P G M E G C K V Q D I G P D R T F R T F R Y T G Y I I K G L I T
RANDOM_4542 S A G M T Q P P D D G S F V L P E L A I W D C C L V F E S I R A D L N I S I Y L P D I E E D K W S S I Y H L R P
RANDOM_4543 E D L D C L T D T T T I D Y Y Q P S M G P W N L P E S I V P L T L G T D G P A D E H E I Q V P L A E G A K K G V
RANDOM_4544 G G S S H N Q D K E P E S G D A A F F L K E R S V R N G R A A Q V W S S T I N A K D L A E C P Q T W G A E M S F
RANDOM_4545 G T R K C N I C V S H G T S G L K D S L R P N T A L P S I P A R S G G P S Q D S L G F H S N R N Q W E N P Q E F

RANDOM_4546 G V M K W V D R K E A L T V S L Q A H E Y T L T S N L L L Y E M S C G S I S R N K G V G A I P W N P Y C D K S P
RANDOM_4547 E Q M G G V D N P S I D Q L E L E A Y D V M K D V A A E A P N A R E A I F G L E L N N P W A G G G L C A I K L D
RANDOM_4548 I A S E E D F E V K V K Y Y Y L T L M L M F V L G L R D F C C V F F T K R L F P V L V N P E T H S W L R H I Y K
RANDOM_4549 T P Q E R G Q K E H K M P S F A P Q W Q N T W L N P L A P S V V N M N A L L L P V W Q L R V E I Y W T Y P N E P
RANDOM_4550 D M L S E V A G M E W W V K Q L N G S K I T R G T L K C G A Y R E I H F S S P A S G S I T P R L D G G T V T F W
RANDOM_4551 Y S Y I R S L L S Y L Q Y V G E W A F M D G K F N A A E T K L Q A E D T I A Q V K S P I T D P Y R Q G V A E C L
RANDOM_4552 Q M A G A N L S T V G A A A W R Y I F L N Y A H T P F L Q D H E S H S E D S R N M Y T M E V P V W D A V V K Y M
RANDOM_4553 N S Y G V D A I S L F A I L S S T D K T K D P V S I L L K E P E K G V H I N K K K S I S R L Y L A Q Y H N D A L
RANDOM_4554 N H Y M A P K D T A L Q G W S K S R P L V V K R Y T S V E E R F N N H T V S E S L V I L D N D Y S H E N T T I D
RANDOM_4555 E V Q A G G I R R Q C R T F K H A V S T S T N G L W G V F G F N K S D G Y K F A D P S S L L S P P R P V K N D A
RANDOM_4556 A L T L T Y V G K W G N L L E L H A L K P V I Q G D T A G F R A T G L R N T D P D I V T P G S I L R G E P T M T
RANDOM_4557 G H P T I I P N T P K N L V I G I P K R I D R G A A L P Q L P G T H F T E N Y Q Y D F Q E E N Q K W E K D Y F A
RANDOM_4558 P D K L L V D E E T K S G S A Y K G Q K A P N S L A P S V L A R K D H I P P L D L V S P V E H T G I P N Q G A L
RANDOM_4559 I I W T L A G N G S L A Q M D A P S F A R Q V N C T S D D A G N L A L D E R E G M K W R S A D C S L L E A Q D G
RANDOM_4560 H V K A F C F F G E A K L H S K A E D K T M A Q P S A S G T P Y E M C A K S E D L V V C L I H L N V A R E L K D
RANDOM_4561 K S R Y A I G L T Q Q E Q V Y R A S A A I V E H A E K R A Q N E V F G N L P A Q L M A Q M V L L Q K G A S V T S
RANDOM_4562 S H K T E T A N N L A S R V S L R N R M P S K V Q L I D A Q L R E K K K R F A N L G N G G T L S V P E S E R R L
RANDOM_4563 A S F G I K F R M A S A G C T G E T I S N L P T P D S L L A K L M G E C T V E I R G K K F V A N F Q R G W F S S
RANDOM_4564 D P K V A P D T I K V P G G I G V W T G Y E Y L G R H T G V E K L E D G V G E R L K G Q H N M N G W D T P S Q L
RANDOM_4565 G I E S R L V M L S K E H R I A P T Q I F Y G T A S L P V T S G L E A P T L N A F Q L T I F K P D A K L G C R P
RANDOM_4566 E M Y A A I S E E L A N K A V F L L A R R I H N F L K P G G N Q D Q I K D E P Y F L G A I A G W V P L V S P N D
RANDOM_4567 I D I A K E P E S D I Q A T A L R L C V Q G P I N E N L N Y E K L P R L G R Q S N P T W T K P I Q D E N Q V E L
RANDOM_4568 E S A D C M D I P D S Q L E G S E L G S R W S L N A N I F A T D M L M A T L T V T A T I P M R L S N A V G R D Q
RANDOM_4569 E G R Y I D V I A S A D H T I N V T F P F G D D V F R K L C I H F N F E Q S S D L Q D S L V G A H T L S I D E A
RANDOM_4570 A K I E A A K T S V V E Y E Q K S R G D P M G N P I F R G L P Q Y W P D F A K T C R E K G R M P D K L F W L A K
RANDOM_4571 F R L R V Y S T P G C F E T F E S V L R T L A T K D S D C N G D Q G R V I T A W A A C A F A Q Y K N P G F F T V
RANDOM_4572 I Q A I T M N T L T K M L R F E G P M E S L N E L A A S N A Q E F K G L C G V F S C K V H N P L T P V S H A E V
RANDOM_4573 P A E H D G K V V E A P L A T D S F N C R P L Y S T L T N G C I A Q V G Y S G R R F L S V T Y I S N T M P K L K
RANDOM_4574 R Q C D Q Q T S G L H N A G T L A K D F A W L S V N Y L R I L L H D F C L Q E E H Y V A P R P L A V Q K V K K T
RANDOM_4575 N E F V L D N Q S Y A P L Y V Q Y C A F T H L K K S G A D V A S M W N L G R Q E V W E D K Y V L R G K S G E V I
RANDOM_4576 A E C L S D G S Q F G S P G T V R G Q Q Y T I I R N S Q S S C T L L R I G Q Q G W G T V K L D A L V H T G E M W
RANDOM_4577 I A P L Q G S G D K P E K E D G R D N L P K M A I W L G I T L I Q A A P S L F E Q C I A T N R S E V H L F E T V
RANDOM_4578 M L I P V T I L A R I S E T G V P N I E M L Y D G F K K G P E G A E N A G A D S I H E G I I T N F V V K V G T S
RANDOM_4579 E L W I I L V V T G N L T S R A T D A I F F I T D S W F N L K F V D A I P P H S V K L C Q L N D G L T S P V G K
RANDOM_4580 Q D D L D L T M R L N M G D V L T P R Y V P T L P Y F I L E Q I S K R N F E V E S G N N E Y Q G D L F T L T G P
RANDOM_4581 G G P V A C M S L V L S Q L G L Q F S R G Y M F Q I D E M L A N V F E G L V T Q S F G G W L A T L Q Q G D V I Y
RANDOM_4582 T E S E K K K I S F T D D F F L C H G D N Q Y S S N V A S D G I V T V Q Y N Y A K L M G T I I R R S G G L R K I
RANDOM_4583 S F G S K L P Y N L V M S I I S M P A C T E A P A M K Q L H S P R E T G D M E P D N Y D K G N L V I R E A G R T
RANDOM_4584 N M G S P G V S L G E E M G E L L P T M L K R M D V R N P D R K L K L I Q A T D N V T K D P T H T P H P S E T S
RANDOM_4585 L V L Q E V A N G L D G R V L E K S S I R Y M P D Y N P A S N D E Y V N I I S A K L W K L F A P L S D F D G A W
RANDOM_4586 Y V R V S Y I K V L D V G Q Y E R T E D I I E S V T A G I G Q V F N F C F I T E S F A L H R G P N G N E L N R E
RANDOM_4587 Y P D Q Y S G T S R G L V T T L Y S V I Q P G D Y V S H S V T D T N R A A T L F D V N Y S F P K C E A D Q G G V
RANDOM_4588 F G W L A S N E Y T G I L V L P V Y N A T L Q Q P T D S Q S R L A N I H H S E S D I V V D T E D G N E D F P L Q
RANDOM_4589 H L K V L A R M A G G Y V R N R K M Y L E F F G V D M G E R K S Y D D G P R V E I P K G V D T R A K F A H A G A
RANDOM_4590 I M Q M G E L F G I L Y K N H G D R Y G N S Y A Q N K E Y A L P I K T D G D L L N P N E K V A M Y L Q S G L T V
RANDOM_4591 D R D A V R G V K L P L K L A K I H I T T G F G S N F D L T S E P G L T I L H T K W G H P A E I L Q G G M V E G
RANDOM_4592 K A L E S T T I V S G V L L N L T N S S D Q P C N D D Q E I T V E N R A K Y R E K D V Y P E A A S V R A F A G A
RANDOM_4593 E A K K Y G T L A E G P C S F F T V H E E E A K T D G C I S L S I N V N L T N G G L F F G M I T E P M L I I K K
RANDOM_4594 S G Q F N V E H P N D K H L E N R Q Q G R T N P A N D S N F L D R S D E D I S V G A H E T Y E K K T W S E T A T
RANDOM_4595 S K E V E F A E A G V K Y T R S A P D Y V I Y K I T S R A H R I V E N R A A F S I V M I V N S D L L Y E R L S N
RANDOM_4596 S L T L C F Q T K S E F C T G Q T C V A E V R E S E L V N T N D K K L P E Y T D I L Q T R E A N K R R S G Q G G
RANDOM_4597 R P L V Y D K I L E S I R G R G G D R G K P A E E W N L R M I E E A L G Q F L W Q Q G R I T L N D D N K R F P Q
RANDOM_4598 L F L G K Q V A E A G C G A M H G D K F W A V T N A Y F V K E S A D Y I Q A L L F R R V T H Q A V P T L A M K R
RANDOM_4599 T A G E E I P E F P R K F R F P D T L R P A V D I L V S A M F L D P T N R L V T T V Q S Y S S Y D S L A R R S M
RANDOM_4600 P G N S Q R D G Q D D P P E K I C R S D L K R S W I V A R D G G N R P N A S L L P R A F P D A R E T T L E L V C
RANDOM_4601 G C C T R G V V V N E V K Q A L P I Q K A V G G D A A Y L R D H A A Q G P E A S M L T Q F L K L S P L D I K E C
RANDOM_4602 G Y D I V L I H K N Y P D K V H K Y C K E S Y F A N G L F R T H Y F L Y Q R V P P H E W M I Q M T D K D L C E D
RANDOM_4603 G G T E P A R G R S V Q S G I F G E A P Y T K E T G L R H V S P E S E L P C M A V R F H S E D E A E F S Q A K K
RANDOM_4604 G S C E V H M A S T K E V N L I L T R S V A K I E K S S T C E A R E P E I G E L C P K K R Y Q F H T R L P G Q A
RANDOM_4605 D S D I Y D N I G Y A V K P L L L P D L E L A P S S A G I F L K T W W L P L N N T V E R H S T P D C F A I E V N
RANDOM_4606 E I R S A S C V D K N L L G D C S N A S W G K E Q H R T E S V M D I T E R N Q S D R Y E D Y G S E L T L C L S V
RANDOM_4607 Y K P L T M V K C P R M I L N V P T L A A L R T K A G S R I S D G E S E P V N V D E G W Q P T L H G Y V P G K W
RANDOM_4608 Q E N L A C K T R L P V G K A V E P G S A E L V R S H E S C Q S D M T G L L T V A Q V L I S E S S L F D N F E K
RANDOM_4609 E N L G R S I R C Q V D P L T E E V V F T K S P N C A K I T T R G N R I S D H D H F T K D N A G N I L Q A T E L
RANDOM_4610 M N M V G L G R A V Y P C R D V A R E P H G E T L R T C T G S V E R T V P T A S L M Q A F S R F G Q L F V N A A
RANDOM_4611 L A L V F S D S P A L N I T D K Q E Q L D S I Q S R V S C T W A P V H T A L D G G V A Y S Q A Q N I D C E G T N
RANDOM_4612 H V A D S Y G Y A V A Q S R P D M Q N A G C N E P I G A K L V D C K C K P Y L A T K H M F N V T A C Y V S E A L
RANDOM_4613 F L I T K D L F D T M N D P S C E D A A V T L S A I A R G V D T N P C E D V D K N L S N I P L L K D L E V E E L
RANDOM_4614 T V G E E L M S L D S H S I D L T P F I F R Q G W V L R A G T W E G E V M L Q K I K D L K L V E P L L P A Y D C
RANDOM_4615 A N S T P E F P T P P A I R R G R Q G S W K D N M L T C V L A R C A E V H K A V Y A T Y V L G P C S Y Y T L A T
RANDOM_4616 R K D F E P S H L G C A G G N V T G T F L F D N S K A A M K A T G A G F S C N K A E C A P L E Q R R L N L E C S
RANDOM_4617 H A Q K A K K R D W K A Q A N L N L G R A A T L Q Y P Q A S V G L P E S M V F A L R Y N I V S Y S D N D L L A I
RANDOM_4618 D T T R E T T M E L E P F F L W P P E H W P A Y R E E L D P P R W S V S N K R I K I V I N T A K S A P T G S S Q
RANDOM_4619 E T A P Q D E G T S R I A W V G L H G Y H G N T R F H N E K K S N L W I I L G L S M R N W A R E E F L E K K R F
RANDOM_4620 D A T D L V T A A H L Y L G Q L C V V N H A S G I N G G V S N K N G T S L S Y A Y K G T P C E C S L Y E C H F D
RANDOM_4621 P P V T Y L D L R E F H A K E T K V S D T V L S L V E K R V N N D T S P A H C V E E V Y L T G L I D S G M R P F
RANDOM_4622 S G R R L E P T Q V E K S S E K R L P T Y L D R K L L V K G R S S T P Q V G P M F L W T T A P N G K S E G E R S
RANDOM_4623 L F L P N I S E L N M V K V S G A I E L L A T E S S M Q G L Q V H A A Y H F G A P T A L Q K W K E E P A M I G K
RANDOM_4624 Q Q K L A S Y G K G K S Y C D N S N L A T R V G M G R S N P L V P Y C S K I V L E E L Y A T A S F W T E C Q L S
RANDOM_4625 N E S Y H V R T G L H V A D V D A A L V G V P G A R D E C E S C F E I S D N E V P R M S A Q Y F A Q V A V V E D
RANDOM_4626 P K Q S R Y S A I M G D C A L P F G G H P S R T L V D D P S K L K E K P R L N N E E G Y D F Y V A A C G L I Q P
RANDOM_4627 D G A M K D T Y S L L L G S V L S E L Q A T S E K M V Y F N L L G W R R G G K V E A Q R P A Y P S K S I F H D E
RANDOM_4628 N D K Q R A M A N T L F S V N H L G A G L N G P E D P E H I S I L R F C R A T V S A I K T L L E V I T F G Y I C
RANDOM_4629 L P G T R N K I Y N R K S G P G Y E F N E T G L Y S G A R V R N I I N H N N L D E G L G D R I M Q E A D G V G Q
RANDOM_4630 K I H L V I D D S S T L T K V P T A V L V R P L D Y Q N V G A S P G E M L V T S G E L D E G I A S A K I T P F S
RANDOM_4631 P L Y V P S T G I Y K L L I K L T I I R Y S A D W I P V S V F S T S G Y N N V K G D N A Y D F K P G G C P D D T
RANDOM_4632 G G T L A K L A H W L F P V G R Q Y H E L S K K P S T G Y S T G T A G Q G F M N E G H A G K P Q G A R S F R E T
RANDOM_4633 A D L K E I A Q S T S L D G Y V P C E L V Q R K R T S T G D V G E Y A L A D I A Y F I L E V N R V V M A T S M A
RANDOM_4634 I V Q T L S G D V V L E Q D L T I A K L C L A E E E T C A C N G Q D R P T K F K L E E M D S N A V E E L E Y L Q
RANDOM_4635 I A E P K H I I T G F P S D E S E V P R H R K A A L T G G E I G A K Q E N T P H V V R G M T P V E V F E T L F K
RANDOM_4636 Q H S L Y T A L T D K P K F C K Y A N Y L D V T K E K A Y K R M L S E E R V Q K E S K I F M P N Y M L G V E Y P
RANDOM_4637 T A F H N K T A V L F L V S P A Q R A R I V I E A R I S Y F L K G K L V K Y A T S Y R S T H Y K L T V F F F G F
RANDOM_4638 I G K G I G G D G M K H T V L P E V R K V K K D R A S G G S K K D L D A S F P E A E E E G R I N L G S K T S I A
RANDOM_4639 L A E T M N N G L T D V R N S N G K T T M W I Y F S D L R E D A K P Q K R E H N E L V N Y A I V I A S R C K R F
RANDOM_4640 D A V N A P A L E T R P V K T T P L V D V S L P G G E S Q L P G S L A Y C S G F E A I L E K A S A T D G N A R R

RANDOM_4641 E R S K V W T H V P R H L Q G Y W Q E A L Q H S E V Y V I D V V T L H R K L Q T N V C T L G C N D A L V V E N S
RANDOM_4642 P H Y N G C K M N P T H A N G F D L K L R N M V T S T A S T L V T T I V T E N I T E I F I E P V G A K E A R L R
RANDOM_4643 N R K C K I S K S V E F L L T A E K F Y W S C T P K N L L D P T Q P L K L Y A G F N N G P Q A P P E V D A N Q N
RANDOM_4644 L Q N F G V F A N S D V S S H K C E G A V S G P T D G S L T Y K K I V W E D D R L D S R K S A A S S M P D D T I
RANDOM_4645 Y R K G G Q I F K R I E E E A A E K K I S R W R A C E G F P A C A A L V C T D T E K I L S Y P R A Q A R R A E V
RANDOM_4646 D I E L Y T P P G Y P R D G L P E K A R A G A P V Q E D S E V E Q T N L Q K G T S S V E Q L K F K E T N K N S Q
RANDOM_4647 A R L Q L S H N C S I G V V T Y M S K K Q T G E D G D T G C G E A D S S G E G L L M S T N G F S L L A S Y F R V
RANDOM_4648 E Q S P A F V A N L K V G V R E I A I N S L G E I R P I S H E L R F E Q C S G L T N D M N P Q R L L C W K H C F
RANDOM_4649 K S A G E M Q S Q D V I G V G E P N R E A G E A F I L E G G V R I S H F L R A A P Y D G F E F L K N T S R S A S
RANDOM_4650 V T F L S L G A A S A D Q L L W E F N S S Q F K L A V M I S D L A Y P E S A N R I M D N S V L E S E D M R A F D
RANDOM_4651 G G T A T E R S L L D Q R D S T G D T R E N V E L N V I D T L V S A D S S L A T Y M Q V L A D I G I Q H T G Q N
RANDOM_4652 P V Y V P D R N K V F G K Q A K G D R G P S A G D I N T E A L N E L L A Y K E L D I T V K W K D R K V A H T E D
RANDOM_4653 L K S G Y L D Y Q K Y Q K V G E L E Q L S E K L N L E D S V V T K Q P R D N G V G L W K A M S R I P N N A T R S
RANDOM_4654 A D T K A R G L I L G A T V L A M G V D E D S H L N E G G F Q W L I S S R G V G L K S T N I R I L Q Q A N Q N K
RANDOM_4655 W D G N D S H A T Q L G S N Q G F L L E T N V T L T H G G S D Y G A P E M L F T A N P G G A F T Y A N Q R A T I
RANDOM_4656 G A G V E W L G L Q D V G P P E R G D R L V A M I V D K L F A I Y Y W D N A M M S N Q V A H I C L N R P S V D A
RANDOM_4657 K L G V L V A V S A S N T V S L S V E F P R T L G L Y C I P E A L V V A L N S L F E K V G M V C G Y K P R K Y A
RANDOM_4658 Q W S Q D H G D L G S K L K L G L M S D P T L V S T D F E L H Y E N A R G A L A A N S W K V E E R K W P S G A D
RANDOM_4659 K C A S C G N L C L L R E V P V C N G L Q R E G G C V A Y Q K R D P L F V Q I R V E G L C Y D I Y F A N S F L F
RANDOM_4660 A L H G K T Y L G F G E S P G G M H G T L P K T L D F D G N N A M L V Q D A K W M L H F K Y G K I L V N L S T S
RANDOM_4661 T K L A E E E L T T I V D T L A D W C P I M N L A I S A D V P S L G V R A E F I R R D F P H C V N K A P R T F D
RANDOM_4662 A D I G D K A V L S E E A K K S L S E V Y W R Q R P I G L V S F P V G S E G V L P S W D L I L I T A P Y P P A R
RANDOM_4663 L V F H F N D D Y T E S E K I V D A D E Y R S E C G L T W Y P A E S F V F G L P A P G I L W I R G S Y R A V A L
RANDOM_4664 G A R F L K Y K N G T L I E Q A L P L L I C E E Q N D L A G Q E C K L S A M D L A C M T F I Q S T A D T L V V R
RANDOM_4665 L Q L I A K T R G G S S G Y M G V Q L E S S C D I G P V W G M I D G T F H S V K F F N V G S D A E W N P E R W G
RANDOM_4666 H E R R A A Q L E E C E A A Q P T L V T L N F V L E G R T V I K G N N D L Q H F I L E L S L I D S L G G A V G P
RANDOM_4667 A V A G H L V G V R Y R Y T G L L H N H P V R K M S A K Q L R E L A L T K Q A E L S I R G E A C A E V A R Y G C
RANDOM_4668 V S N T K V S I S A P L R E Y P C A G N K D E G Y S N G I V A R S K T T A G R P I R I N F L A A V D S K Y S F M
RANDOM_4669 P H C E R F L F L A A Y R F I T I K K F D F Q E I G K V V F S A M E N L S Q N R L V L P F V R T F H I S I N D D
RANDOM_4670 L T S R D F D F D L T K P T W P L F D K V H C N S L V S V D A V L F Y D V P G C I G L P Q S F N K K Q K L D Q G
RANDOM_4671 K L M T L G A K N Y N V K V G K A P D T H T T P E N V L Q A T V Y G Q V N L G R L K F Y D Q D Y P L L F R T G E
RANDOM_4672 S L L P V E V F L P E R I K D I D G F T N F H G F T F L L I P S T A N V E A I K L R E T K D R E L E V G P H V K
RANDOM_4673 C A V Y R S V T P R H A E T I L L T E E K C Q H N L I V Q R P W K A P M G R N E K W L D S E R V D N A W D A K Y
RANDOM_4674 K G Y E A P A L W H A A E N S R E D A L D M E D V H M I G G K E L S E L G A R T S Q P F E H W G G E V H S Q W A
RANDOM_4675 K E F L G Q P I F V A R A P T S Q Q P A P T S F R E E T Q C H F W V H R D A H E I L R S G D T I F E L S V V S D
RANDOM_4676 I T S G R W F G D I H K Y N E T E R Y M K L I V L I D C R L K K R A Y S H S F G D S N K K A L F E F P L G P L I
RANDOM_4677 G T F S N S N N P P K C V R R G T E E L I S V L L Y G S I N D F I L D N I R A M A D R K R S A D P T A P I S S L
RANDOM_4678 F K L E E S P F D Q R R C I R R S K C F F Q R P R L G W P M A E R L S P F D I E G S L G Q P P H V S P E I P L R
RANDOM_4679 G T I A V N Q G C E N K L E K R D F V F F F T Q L Y G F D P L Y E M D T R Q L G L L S F E S M S Q A T D V N V T
RANDOM_4680 P D V I G H L D R A L G K L E L P G I E D F G S G S G Y I D F L A Q I Y S I A A T Q Q E N V Y A L P D H E R R K
RANDOM_4681 Y V I C D Q K A L G S R L D G F T E I V L D R A D Q P I M L F A L D G A Y V F H T Q V G A V P P Y K Q L F Y F G
RANDOM_4682 L G G S T S L L D P R S H S L D G A I H E V G P D A S M E V G N N Y A P S G V D Q Q F R V K G L S V L C V Q P D
RANDOM_4683 A A Y V R S G R T P P F Y L K P E F L K G R G V T Q S H L C A I T A G G N K D P R L S N E Y S D I Y N D L N P L
RANDOM_4684 A E D R D P M L S N A L F P D D T A T L D T Q F P G H P A S D K A T R P G Q G V Y G W R P E T P M S R K N S T R
RANDOM_4685 A H Y Q A R R Y N D S T N V E K F L P V W P N V N D S F L R L I I L F D S G A L A V S R A R V Q L I R T L P A V
RANDOM_4686 S G C V N R G S N S R G L E A P S E A L P M R L V V V V K M A N R K I P A L G Y N A A K E W W F E S K D E F A S
RANDOM_4687 N E Y D Q V S L N L S G A K Y I E K Q P F L V E D P S S L T A F S R F D C E L T E K L I D I E P I Y V L V K G H
RANDOM_4688 E G P I S G T P L G M W R G F P Q G L Y D A L V L L I E M E Y E H E A P P Y T T S I E L R P C Q R P A H E R V K
RANDOM_4689 G G P W G T F N A T G G V P D R N P Y K Q A D L E G N G F N V T K Y T Q V Y L R E L R I K N D C L N N F N K A F
RANDOM_4690 Q L I F K D T W Y I V P G L V L K E L L A Y Q C V L L L L R A H Q G I T Q P W W I A Y G V Q P F D T N R C S F G
RANDOM_4691 T V E D W K N M V V D S A E V Q I Q S M R I S A T G R C M H S Q L I W I N K P S R A D Q K L I S L K A N G E P T
RANDOM_4692 P I L S R D T T P E E A K V M A D K P V H Y D E G D R Y F T I Y I G N F Q I V K V S K V V E T G E S F A K A D L
RANDOM_4693 L S L K R S K Q I L T L E T F D P L I F A K K A T A L G A G V A D K N F H R F N E N N N F R T S E A Q S L S T K
RANDOM_4694 N K P G F L I E G L A V W L A Q I W R E T L I C A P L L V V N A P L Y Y K K S A L G S K K I C G A R P S K Q A R
RANDOM_4695 N F V T K P A I R A P A V Q Y I K R K V V Y V H V N E A L T K T K G G A L I S R I P C V N I T L K A L L I L G Q
RANDOM_4696 T F L T D P N P G V Y D G K V V D Q V E Y I L G N V S I H K C I E V L T S T G R A Q E A W K I V R S E Q G R A S
RANDOM_4697 L V P P D D V S V D T D K I S G Q V L A L S P Q V G W K D F I N I V T S C A V L E A I N A A G H K G T F M F I G
RANDOM_4698 A N S G A D G S E V L V T A E E S C P G I H A I V S Y E H G G E R I S D I N S D I C K E Q C D G T I K L T S L S
RANDOM_4699 P H R E A M A N Y E T F E R I F A G G H E Q G D V K A P S T G L I T S D E R P G T A P D H L E V K K R A V N G T
RANDOM_4700 D E I R M L R D V D P M E S I T D I T Q G M D S I R F E D D G W S T C L K L T F I R Y A P A K N P H G V V C V N
RANDOM_4701 E A Y G P G P K M H V D E K D L T Q I W T G F N L R Y L F A H S S E G V E E Y N T E V V F E K L T P A V Y K I A
RANDOM_4702 E S R K A F G Q S R Y I C S N F Q P P T T Q G S K L Y G G G T V Q V V L C R M M R T K S F M P P D R Q S E A H L
RANDOM_4703 N L H N I P A I R N D R V P G I S N E F Q N G R A A A G I G A S D F T P P Y F T V A I Q L N A E S V S M A V S K
RANDOM_4704 A T T N A E T H S G A L E W T S A R K R Q P V M Y E V C F L Q D L D W W L W I S E A L D S A T S F Q R A L M N T
RANDOM_4705 T E T L N S M G P C K V H D K G G N K S A D T S Y L G S D I Y V K T G G Y I T G A A A R T N N T S G K A F Q N A
RANDOM_4706 T S A Q T V A R R D G D R Y Q R Q C Q D D V T S S R T R E S I R P Q N S R I Q T L S I V N V L K R V F G D Y V A
RANDOM_4707 N E H S G R I K Q F V A E S N T A N S Q V T A W D F V S V A I Q P V T T K A P G C D R K A S K H G A G T T I F S
RANDOM_4708 I E S E E Y G E K L D R A K G V E I A N G I L P G N D R D M P E S F A H L Y E K K T S V S N W K T Y I S G V I S
RANDOM_4709 F P K D T E Q L P I G F A G I G Q S R T G Q L D L E G T L Y P G L V S F I I T C Q K S R P R P V I A R S V M E M
RANDOM_4710 K A G E D A K D T A F Y L M G G Y L A D N V T D Q K E Y G G S G N V T S V R V Y E G I V F D A G S S S D V S T I
RANDOM_4711 V R A W I K W V W A A E E F S Y E S V M K V I K K S E N L N D R A K A R L Q L M R E T L N K T T V N E S R T S L
RANDOM_4712 S Y F G S I N D R G V N I T S G L I A Q Q S L M F K L P K D R K E S V M E V F A M S S F P G L Y V A A L L I K T
RANDOM_4713 I F M H L R E K R P L S T L E L L W T I G L E P R L V D I K L K P E E F P S F P A D N A L I G E D V K M G G D R
RANDOM_4714 P Q V N V H K Y S G I N T W T P H D S V K N A I M I V V T S N A S V I D F N S D V F Q S E K V I A A C V G R A R
RANDOM_4715 L Y G T V D F C A V E C Y E F R K I S R G G E E V K G Y K E V G I G T I I L C F L R Y S G V K Q W G I H E R E E
RANDOM_4716 A L A K N K A H A Q N P F L Y R P N N C P D K V M H E I L I S C N K K N I V S W F K E S T G N L V E E T Y K A L
RANDOM_4717 P E S K D P I L H Q A N N F C A D Q P K P A F K K E P L S A A T D Y A H A L D Y D F D V H A Q I L R Y P L S N G
RANDOM_4718 G R S L D K S L R L L A D A I S A L S A P G T S A I L T R A A R A A N L I G F A M V K K L L A T V S R S I K T L
RANDOM_4719 H E T R I Y S Y F I T P I A T V Q R R V L S G L T D N N R N M V R E L M G A G L N Q R R F E T E T Q Q T E S E Y
RANDOM_4720 E V G R V E H P F L T K E K I I D E F S L V R G T N D H R S A N I E G G A I F A P F A D T Y P A V T W C S G F T
RANDOM_4721 K F I P N L D L T G Y G W K R E T F Q K E C I W K N A T V T S K P S L V L L N A K P S D M S E D K N Y V Y T S R
RANDOM_4722 L L L N W K E H L A R M K E E N L P M L L C K H K T L E V S R K D S G I G Q T A N N G A H A Q C P R V I G R I P
RANDOM_4723 T N S S S E L R I M A A Y Q I V I C T V V T Q G I L D V A Y E L N V P S I S L R K L F G D P P S E L V E P H V I
RANDOM_4724 K E S G P A C S L R I T T G R Y I S H I Q L G S S T T K L F E T S Q S D Y V P R T V L S A N T V G G T D V N E E
RANDOM_4725 T A C I D Y N A S V I D E E T L T N D R R V Q K S P T K Y D E K L L E T E W S V C S A S V R G A V Q M G V S Y A
RANDOM_4726 D R V G T I T G V V L K G D E V E A A A Q P C A M R A S Q D E E E K P L T I A R S D K F T P E G K Q P V V V N E
RANDOM_4727 R P Q A Y D P L F W P N K G L S R V G D S E Y E K D A V T E V G I I A G A V S L S R R E K E P L Y G P S G V L G
RANDOM_4728 H V L Q A L T S H E K Y G H N Y E S K H K S M A S W V D A G N G F Y D I C V L Y H S N N G N L N E Q A D L I F G
RANDOM_4729 R G G T S A I L W K E S A E A H G Q T G L P P A R R Y A A S T D S G A R E I Y V A A G F S L Y P T M P I R V N R
RANDOM_4730 P N M G A I A S H S Y P P L D G Q A D P K E A P P F E R N L A A A R V T C A L Y D G I N F Y E N L V T K C P F D
RANDOM_4731 R N T E L K V G R V E L V L R H Q L L C A A E Q G A T T T G A E K V T N I D F L S F W R D G R I D T M I C Q S K
RANDOM_4732 V A T S D K N G E P S T A T N K M A G Y V E L V V R M G V F S R A K A G Y D K V V C F P T N T D E L N C Q A F V
RANDOM_4733 F S N V G F P G R I R E A R P Y K N A E D C L L T P P L D D V R K L S Q H P L I S H S P V K P E S Y G G T F A R
RANDOM_4734 Q G W V E E Y F T V S K P M V A D S F E I G P A T S M W V G I K A I L D D R G G A I P T L K E Y G K P K V S L D
RANDOM_4735 P D P V W I Q N P V C K S W Y A G L Q A N Y S M D F R F F T T P D A G L E F D A F P V N I P S K M G N A T L A F

RANDOM_4736 S Q M Q V Q W P P T G S L T Q V S I R I S E S A L N S C R T Q R I L T T L G G T G T P V K S V R C A L P C R K A
RANDOM_4737 H K S R V A S C S G P L P V K Q D L W E L P R S E S V Q R P L S V Y E L T C E E L V E V N M I A V I E L G A N E
RANDOM_4738 F C Q A V F D R K G N T D G S T L D A M A F L H E G H S A N P F T M K T I L S S S V K A L L P I V W P L R L A V
RANDOM_4739 Y L L P G Q R S A V D I A A M E E E V E Q G T L D S T N M I C G C N A N S S I F G S F I L F P A G Q L Y I V D D
RANDOM_4740 A M E P S K T R V A E W S H F Y D I F Q L F R A D S T V M V L Q V A I C G I L T N A F P K I P T G G D E F E G V
RANDOM_4741 R N P S A Q N E E E H L G Y Q M K N P L R W D P V N G D F A F R P S L S L S C N T E K A P S L L D T A T E M D T
RANDOM_4742 A Q S Q Y M I D Q G P R Y L W D F T T K A N L L T M S S K K E S K P L G R A E F K A L W S L G A T I L I V I A P
RANDOM_4743 L D G E L P S V E R S D L L K I G V E E L E C E E K N P Q T S G P S V K G L K R L I Y E T N K L C R Q N R Q Q A
RANDOM_4744 P R M S S D F N K P I V G V L Q G R K N T E L F P L E K N E T R S Q L I V I S L T F K F K P R K P V L F K L V E
RANDOM_4745 F E A H S N V Y N A G K S P S P R N K R V K D V H L S R H C S A S I P T G V N Y C K R G C F E V E G Q I Y I N S
RANDOM_4746 S A W A W C K F L H S Q S L S N R G D K Y A H E E Q S E H T L T Q E P C T A L G Q N F D K R P D M C A V H L T E
RANDOM_4747 T T R R G W V Q T D I F A A P K S E H D H Q I A D A Y F H N V N A G Y Q E L A Q L D Q R L S H Y F L A I S K F R
RANDOM_4748 C P D D Q L F P S P A T V I T T Y I P F D N N F A L N P D G I L Q M I S S P L L E N D A T H A R F I L A V N I L
RANDOM_4749 R E A P G M R G T E S S I M Y V I D G M S R L A G F D F A F V P I I M L E R I A E D V N V P F W P S D R G I D S
RANDOM_4750 A W L G N T W H E Q P A L R F E T L K S A M Y T I N D I S I E L A L S E V P D S G C W W T M T S A Q G E R T A I
RANDOM_4751 R R Q A T S E Y K K T R T T E N E E I S R A L T T S G K C D I A E A H A I R I N A R E D S V A F A W S C C G G T
RANDOM_4752 K G S L A G E R Q S L E G F P W E P L A E F S R S H S I E T W Q G K P P S L E R A L L N N P P R R N P Q R S H L
RANDOM_4753 Q G V T F G W D E F L E A D F A P N A G S L R A E Q Y Q A A S A N T K V E L S I E I G E D T L I M Y F C I L T A
RANDOM_4754 T H K Y R E R L L M N G Y L D H V T E N G T I S R T F D A P V D N I S G P V A V F I E G L L D S L D V T K G R P
RANDOM_4755 F C R L K L A F P V G H R F R F A D R Q E A M E G V M R H L V Y V R S T R L S S N D L T K G L E Q Q G Y G R Q L
RANDOM_4756 L M G K S T L R K E A Y P S V G L T G D L A S R G F A I C N G T P V C G K V I M M W P S Y Y M L A K E D R L S K
RANDOM_4757 K R I P E E V M G V H G I R F P N N G G K L A R L W Y P G V Y S E A L G W K L K D I M V A M A T L E D W G S G D
RANDOM_4758 R E G I E G S D A T V L T V Y D A I T E A R H Q S A K V A V W F E N N Y L G M P Q S P A P L K N V P A E K V N K
RANDOM_4759 S H F V T S N F L G H S E C L F A P E K V L M H A G K V L E K N Q W E D E S T T L P S Y V P L P G A F D V Q N E
RANDOM_4760 S Q V I E G A V A H P A Y T Y I A Q Q D T S I C Y A A V S V G F L F E A R V N F G E K W Q T N G Y R K V E V G S
RANDOM_4761 E L L G A E A G E R S L A K A N E F G C K K I D L D D A Y M D L L N D S D I V A G H A K H E F R E R V M G L P I
RANDOM_4762 E S F V D Y S V D A F P P N S I D Q V V R F E S R A H Q C E E P V I A S G L L Q L T S R G P V T K Y S L C N V G
RANDOM_4763 S K S A N C V V E T R R S T Y I M P T E I L L R E A V R G Q I T L I V T V V Y P A S G Q E Q E P F Y A V Y R K L
RANDOM_4764 G D M D I T E E T M R F L V Y R T I E L Q P H N Q C Q T F H A G I E L K G T Q L F V D G E D S R L E A I E L K A
RANDOM_4765 D G G L V E T E L E G G L A T S S S A Q R Q L C S I S E R F A Q L N V R Y A F T G E A D V L V T E Q N V T Y L K
RANDOM_4766 G T L S Y S Y E T K A F S A K P A S D H Y V K V A G P E R A F F T W E Q N K W S S P A G F P D E T I H K S A S A
RANDOM_4767 S V H C I L G Q C T V P D A S K H Q P E P L M R T S D L T S L Q C P R T R S I R L E N L T N L H N K P Y K L L N
RANDOM_4768 A G C T L A V E Y V R F C S M T T V A G Y F A V K L L L V F G F I M T R E I G E C T Y T F I D S R V Q N F S S V
RANDOM_4769 D R G F M V Q T G G V I T L T K C L K K K R E M Q Q R S Y L T E H Q A G R L S N T L V R K L R K H L N N S N K C
RANDOM_4770 S A A A M K N Y Q Y Y P S H V R V D A V D V Y L S D V A A V A A P R M S S Q R E Y S K I C L S Y K E C G R S T A
RANDOM_4771 N Q V K L V H V P N S E D T A E T L S E P Q D Q L Y V K T R E K E Q E K R H R V Q Q R L K Q F V R D V I F L W D
RANDOM_4772 W T K R P N W W P G D W N L N S I A D L V L R K L D S S T Q W H L L E L F K S A Y K V S V D L A C I L I Q K S R
RANDOM_4773 M S S K A H C I P K I T K L T P D T N P Y E E I S L L G E T T Y P D V Q S A H A T S K G D P P P R N E D L T E G
RANDOM_4774 D E A R A E L A G M G V N S V H M K C A A N L N G N L R R G D K V Y V R G S G F E Q K Q T L N Q Y F V E Y D T L
RANDOM_4775 V G E G R V L M S I F S S F F N D H V P R N Y M S A R A P L L G N R N T N V N A C A G L V R R D Q W K A G K K G
RANDOM_4776 F L D G A Q M D L G P A W L A R Y K G V A T H L V N A P A S L L V V Q M L I P A L I Q K N I T F L F I M K Y F K
RANDOM_4777 G A E A A K Y T W H K Y R Q Q T D V P G Y Y L L K E L T P I C K F F L A E T E P S R D T Y W I W R P E A E S S G
RANDOM_4778 L D G L M Y I I C N G N E L S Y G C E K R S L P A N V A R H E H G V I M T R D S S L T N G N T M L I F E L N L N
RANDOM_4779 P I S V A N H F Y H V H P R V A A A T K T R Q R R Y G F S D E L A D C A M C G D Y R I A E G V A L K E V F V T V
RANDOM_4780 T S G D D V T T V H H R T G E T N G R R L S D H I K D S F I Q A F Y Q V W H Y P Q A I E V I A K A C I T T H V E
RANDOM_4781 N M L G F G P Q V E V L I S R E D G G L T K Q P F L Q E S G A M S N N A S S F P G T C A L L V H K C S L G F D V
RANDOM_4782 D D L Q Q A L Y G E C G F E P N K V K S D M P C D Y F I A T E F K N D T K K G E P L D S F Q I A L W M I P W S L
RANDOM_4783 S I G N S I E T Q Q Y C A K Q N E G H A H Q T A C A S T S N K A I L K C H G F E D L G K L L P L R G P A Q Y T S
RANDOM_4784 K Q F E L P D N R A A D D P H S E V V W I G K A I P N S P G D H E A A G G A P V T A F L E D P T I E V N E F K A
RANDOM_4785 M G E S K D T G G E G H Y S D E S K Y T F R S S E T L K T V D A A R L H W R S F K L G A E E P A A S H V K Q M P
RANDOM_4786 N C L F C W R S L A I L T R K S T L F R K V N D G T E S Q V L K H I T K Y D L R Q H A E T I V S A T V T L L Q K
RANDOM_4787 H I C S E D C F N F H G S L Q C I S L S T F S R G D T E V E K N I D S S T G H K I S I I L R V R G Q V T L L P F
RANDOM_4788 A F N E N R V A T V T S L A P K T R T L D D L K E D Q K G S A T S A R A T R I I Q C S L W I G I L T Q L N E N Q
RANDOM_4789 M L R D L K R N M R T A L Q K Q H E G P L R S E T L N S S E G Y G F L Y G V Y G E V G L P A A K D K E R D A P H
RANDOM_4790 D Y V G T L S N L S N E L N D G C N D G W D A T I V G T N D T L E T A L M A R D T G I G F E L N L T Y T I A N S
RANDOM_4791 K V R L K S A G Q Q A G E F C I L K L P P K E A Q L L L T A L K T L S N Q D P P W A G S T G Q A T I V G D N V S
RANDOM_4792 M L T G E R E Q S I V A R E Q Q A F E K L E N R Y P W V A L F N D K K A H G D S P L R W I N Y S K L S T F D D E
RANDOM_4793 I L M P G S H R V P H V R H I P Y D G E R E V N M H I S M R I M P K L D T E T L K F A L A E D N N G P L Y V K A
RANDOM_4794 E L H P A A K G C Q A R A E E S A D F Q E P I Y D D K G T L N A R N L G K T G A P L I T Q K K M P H C Y F N V F
RANDOM_4795 I K Q V V G W T K R K G T R Q P Q N L A E H C G T A K K T K P E T L H T N M D L V V L M K G H Q F T V R A M R R
RANDOM_4796 G T R R E G L K N A F R Q V T D N H T K V Y N G R R Q G I V I S S T E D Q N S W H E M E Q K L D I N W A P W A S
RANDOM_4797 A I E V K I D A L V V R V F E S R Q Y C S L Y L E S A Q L V E L K G N S Q A R I S I A T S L P N S M N L E I P D
RANDOM_4798 S A P R T A N S A K D Q A V N Y Q Y A E V R V P S V T L Q V R H Q Q C P S S G Q K D A F F Q T V A V Q D K D Y E
RANDOM_4799 L D S D S V L T R A G K S I G R N E L C M C R F A Q L P L L V V S A N K T F G E T K N K P E H H D G R V P L S P
RANDOM_4800 F M P T A A K N Y D T D L Q S I E A L H D A L I V Q G E W I L G Y F R T F N V C R T A A H V A L S A G V T R I T
RANDOM_4801 V S R G C V L A S E Q S T I G T L V R S R W S C P F P I E R S V Y A Y R G G N V V I N T N L K L N G Y A G H A V
RANDOM_4802 I Q Q G S D I D N N Q N N R Y G A Q S A A A Y T E G G L L F P N D K P S F A S F K T Y G Y T H L G Q D R L C G I
RANDOM_4803 P F L E L D N N G N G V P K G S I R V P G N F M Q V F K A H D Q Y M T S E I V N R I G E S S T G V G D F D L K M
RANDOM_4804 K F G E N H D G W F T P L F Q Q N G V A T L L D W P I D G V L T L L N N N E I T P L N V V E S F A S F S I W D P
RANDOM_4805 E V P S K D I F A F V D T H I K T E R P A L E Y N K T C A I A S W F I R K E M M G T L W D I I S Y S D R I A D F
RANDOM_4806 Q L M G E L W V L C G L E L E Y S C A G A L I M P A N I V D L A S R A Q P F S H S E P R R E S L G D N I Q R N V
RANDOM_4807 S N V D G G E W G N M Y T T S P L A R K I P Y A K V F V R S D A G I S V L Q A N K F R I K T Q Q I Y E A L D L T
RANDOM_4808 N L C N H N P F N K A D V T K N W S S G V D K L S V S L T Q I H G A G Q V R G E G T K R D D R Y G G A A A D V K
RANDOM_4809 K A V K E S N L I E D A V I Y D K I K D P E V R K P I Y Q S K T G D Q K K H T W P R F K A L Q G T G N G S G I V
RANDOM_4810 K T P V S Y T W Y T Q Q S M K Q G Q T E K R S S G A R G G K K A L E V L H S R P T S A H D T S N V P T L V R D F
RANDOM_4811 M E P K L L A H W L L H L I G L R F P A P G D V R G C I R V L T A V A R C A Y K T V A S A K E D R W W K I E S A
RANDOM_4812 E G K G G F T L G I D S F N T P S N G Y G R Q L K P F Q L A D E E P G H H L T N E A L V G E A A N M E E V S S N
RANDOM_4813 N Y M Q C V T H E H H S A T P V T N Q P L E R P N A P Y V I C I T F K F S F L D C I G N N D S S G L Y P R A N L
RANDOM_4814 S S Q D A V N L Q G G F A A N N H F L Q A A T N T R L N P K Y P T T L H N E S K E S T G F K F L D V E L T A H V
RANDOM_4815 A V Q S W Y A C T T G S Y V E F G A A A E D P C P T E V E C A Q K F R P D E V E A P N P V Q S T A G I Q S A S Y
RANDOM_4816 K L A T L Q A P W I F A E V K A F T A A G Y P I N T Q I P N C P T G K I C R G V F K Y K L F H A G S G M I V R Q
RANDOM_4817 T E L E I M K E L E N K I P G A I L D L L A K M T L I Q G T R Q F K P H G A A C I N V N Q A V N L K Y E K T N P
RANDOM_4818 L E V D D M Q I Q L D S D K K K V V G C L W T A V V P R A L P L G Y K A P L H A I V G C R Q L G F Q T T I S D T
RANDOM_4819 H F G I K F L N S H N Y R D R W I C S L M V V G T A A V W W L P A T D P G C I C S A D I A A G L M R R D K W I Q
RANDOM_4820 G H Q Q V W K V Q G I A Y L E P V S D G L A E N G G A Q K V R G K S R D H A W D I A A P S G G V S L A T W D V S
RANDOM_4821 V L A C A V Q T V K S K A N R L N C V T P K S Y Y K K I F V S R H Y Y D T L D L L I D L I R L A V V E S Q A R I
RANDOM_4822 P Q V G L D E S E S V R W H H P E G S Q L K Q D N Y G Q K T A P V K I V Y S S G E H E S H T S F Y E W L S E F I
RANDOM_4823 P R R K L M L L F C L M A L M L V R P F N A I E K V F R Q D G I T K K T I R G D C R A R D A L H T D D G N S L E
RANDOM_4824 D F V A D L L G Y K H N M F P E Q F G P T A I S D T N E L A S S P E L T I Y K I D Y A Q Q M G N T K R R G L P A
RANDOM_4825 G V N D G E C T G W I R D L R P Y Y T K R D S P L D T K V G W G R I S L F V A E D V L G Q S R H Y F L G E C Q L
RANDOM_4826 S E E Q G V I R W G D K R W G Y S I V K S D S K V V S Y D M P N T D H K N L E T E Y E A S A K L Q S N G P T R F
RANDOM_4827 R D N L H G Q H P K H I A E S A C F R V A G L A E T P G D S Q C G Q S R V V P S N T F G G V D G L S R G A L A V
RANDOM_4828 E F L L Y Y S E I F F D D G I Y V Q Q A F E A C D N S T Q G Q R V Q W D I G P G N C K G L P I P E L I I Y N R A
RANDOM_4829 Y A K F Q E K I E L H M K T L A P T L Q E Y H G A E H I L C P R P C I Y A R P L R E V L E R E L I V P K R Q T V
RANDOM_4830 A R T Q T E R T F I C R Q V G A R R K K D V Y R L V P D S M F N W A T T S L R P R S E Q N A Y P N Y H D D T L F

RANDOM_4831 A E K R V S G V R P V K F A Y T T R G E Y E V L L G E K D V P H F N H A G S W Q K C N Y E N P G L R V F S R R V
RANDOM_4832 Y F S T F F Q L A G F V G H V R E H E P K A I G E P Y G R K Q G L V S S D L N T I D A I G T L D Q E G A T K L S
RANDOM_4833 L Q D I I L N A H E S K Q P G L G V L N T N L G N N K L V V L P G P T G W S P L H E L H T V P K I K I Y F Y S A
RANDOM_4834 M C F L N H S F V A R Q I P S G V R N I L T G K H Q C D R G N A E W S G F I P P T F K I T V N S Y L Y V G E T P
RANDOM_4835 K L I S I T L G G N V Y V I D L S P M A D V F L F S V G A K L F V C T S Q R R T L I R D L G F A H Y S I L T Y K
RANDOM_4836 L E L A F D L T N S G K V H L L D L Y W G R L R L G W Q S Q A Y Y S A Q T L S T N E G I K A D V S L T G T A A V
RANDOM_4837 A A T A H Y Q Y T V D N D C F R N I L W I T K C I A S L C L P D S A S D A E I F V R S G N A K V Y G E H G K K E
RANDOM_4838 N T I V I F D Q N L S W E N Q I D K M S A D T P L R L A A T S I N P L T H T L R V D K P R T E K G P K R L S T A
RANDOM_4839 D Q H A G E A H A G P R Y V P L D S T C Q G L D P T N G E Q I C Q Q I L P M A D R L F V Y T H F F A D G Q V Q A
RANDOM_4840 S G L C P Q P A E A S K E K G Q E G D V G P R G P W Q P L P A Y L P V A G Q G L K I L N V D I F Q G G K L K P F
RANDOM_4841 T I S P A G P E N N K E L A G H E K L K S L D L T Q N A C E R K G T A R I Q P E Y V L N V G Y I L A V T A F D G
RANDOM_4842 A K V S N L D M P R G K L G M Y K P V E A T E F G M P M S F V A Y I A D E M P D T V R L Q F C S L Y L I P A F R
RANDOM_4843 G P M Q G S I A I H T D F D L Y D I N L S V L Q G L G N A S D A V L F N K C V Q G T D S K M Y L V L Y E L L C A
RANDOM_4844 I A T V N A P F H P P G C D D D K M G T F Q L K L Q G G T L T I E P A S F Q L F G R I D D G E R H K W M T Y A I
RANDOM_4845 N S F K L Y D S I D T A A I S S A A F P M F V V T S L D Q V R R R I G D P F S Q G V W E G T R L L K A V F F H W
RANDOM_4846 T M I W V N N D H A S V A E L S K T W E R S G L I Q L E A D K Q E H R K E T D I S P G A K K Q N G P Y E F S G T
RANDOM_4847 R D E L D F T S S D D K N E I D F E D K F T C E F Q R E Q K L G I C L V P E C A I W R I E L V G D T G K K E A V
RANDOM_4848 P E L F S R E G L S G T V I Y L Q T I I I I P S M F M D R M G D F Y E D S M A G F K L T A P V K S D E K F D T C
RANDOM_4849 E N L H R Y L G P G P D F R V E G V R E Y I T H A P I P A P S D I V F R D K D Q S R M P F P R E D S G N P A A T
RANDOM_4850 Q S S H K W D S K L V V Q L I H G G I V D R H S A M M A A E V L F A R A P G K K L T Q Y K R E V E A P A T S E A
RANDOM_4851 Q K Q A Y R F W G E F V V V N Q V L T D K F Y A L N I L F C M G W G N V P P I E K L G V V E N L M R R G N V E Q
RANDOM_4852 L E K R A G A S G Q L Q F V S A P L C D A P R G K T E A N P Y P F D E R K D A N K G K S V M S E N S N H L N T M
RANDOM_4853 P E M A E F Q N G E K T M V G L K V V S T E A D S S A K G A H P L A L C K P Y P D Q G F L P V I H G L V S E L L
RANDOM_4854 S L V A L D P Y S E F F N T G L Q N G S Y L G A S W K Q V G S P N A W S S Y S L N V T M A L V V S F N Q Y K K L
RANDOM_4855 D E G P A T Q K R Y Y K E D R D N E A V G A S E C E N A P Q L S T W T K G K K V K Y T Q D K I T V S E E T L S E
RANDOM_4856 I Y A D T G T G T P L P V G P V A D A A R Q R T R V P Y A M L G T T V K I V S T Y L R T I S E S I G Q G W V F R
RANDOM_4857 D L P A W Q Q E P Q L Q Q C K G T A F A R T F L E D L L V E L D V G I D D F L T Q V R S G A F A S G K Q I V T F
RANDOM_4858 V M D S P D E F K A I N T V K T V M Y I G N E G S G H I D D D L L L G I A S V D K G A K Q S S N Q Q H V K R L Q
RANDOM_4859 T I N A M L L T T V S F L V L A I G G P P S Y K C N E P L Q I F I N G N C A D K P F G T N L T P D A A A C M G V
RANDOM_4860 D V N M C N L D M M S V E P E S G P G G L N A N R T R P I D G R E Q E T E K S G Q N P G P I P P Q E A D G G E K
RANDOM_4861 D R C F C E N A F I E D H E F P K K E K N F T Y E D K T E R C K G V I E A F S Y S S G F V Q A S P R T I V N A E
RANDOM_4862 S F P K M T G G P Q V K S A Y A M C Y P R S R R M L T R Y F H A A N S P K E W P L S G P K T S G R L L S P H R P
RANDOM_4863 G K E R L S H D S E V V E A D G F E P N F A N D K G G N G E N A R Y E S E V K L G F D K L I A R F K S L I I Y D
RANDOM_4864 E Y V E G G F L I A R F S S G G S N S S H K S T P L H L F K A Q A E V S I Q K H G N Q L D T F A E P W N A V S V
RANDOM_4865 K A T T C F A Q L E I S Y E S N F L L V A P A A E I W S E L P S M P D V E A V C Q A D A C F V M K Q V F S L Q E
RANDOM_4866 I E N D Y A A V H F E P P W E G L R R E G S W N H V N G S I K V G A Y S M I L I T F Y K F P V P V D Y L G P S V
RANDOM_4867 Q R A V S Y V D A R L S V E W F N G V T Y K E Q Y K S V G S T F Q H P D E T V F F E D G T L G A W I Q V E P R Q
RANDOM_4868 E R Q R I L D G V G K S L S V P L S S L H V I N E H C L C E L N Q T K S I H C S S D D L Q Q G K P N P T H N D V
RANDOM_4869 K H A M K G L G N N C K V S N R V P R K Y A I E L K N S T D K N N A V L S M C G K N R F A G Q T I G D D F T D H
RANDOM_4870 F A E H S F H L V K A A A G D T A H C H P D V A N Q V A S A R P G Y F R G D E R D S G Q S A A D K A P A L K S P
RANDOM_4871 V V L L V N L I S P R R H P K K A Q E Q Q I F I F G Y K F A M A K K T E L L G A I G V N P D A G C V S D A V S T
RANDOM_4872 T A G Q Q P L G E N H A V K I G G E L Y K N V G R I A L P L I N S E F S V P R K S L F H E M V Q A I I D D Q W V
RANDOM_4873 P Q D P S L S C G G Y E A Y D G L E E K R A L R E V E Q S S V L H K G E L Q N T N K S I V N T L L S W D D G A P
RANDOM_4874 D P L P M A T G R S A R T E P S L Q S F R R N Q V P Y G G K D P P P M Q L S L G Y R A P G K G V L E I E G P V N
RANDOM_4875 R Q E G E I M W Y D G L L S Q V T L L K Y D D L A R H A G I N G R A L C E M S K P V G K Y T M F K C E K A G F M
RANDOM_4876 N W S E V T M E P A D N E E D A S D Q Q P M A L I L T W R D H K P T C L A V A E E R D L V K A A L N F C Q A S E
RANDOM_4877 I R H S E F L S N K L I G T A K T Y I L D S K E S C E L K S L L I N V K A A R D K K F L P G L G T N D E V A S N
RANDOM_4878 H Y D V N L A S M T R T L N E G T H Q V V T R R L A A R L Q S Y D F V S G Y V L E L I V R Y Y R T G E R E R L Y
RANDOM_4879 Q S Y V S N K D K P K G S T A P Y F S D I S G C R G E D I T K D Y R N T R L D N H Y R R Q E I V A F I L S I E L
RANDOM_4880 A L T D E Y E A G P L K H I K C A C R V V Y K I K H E T G I G A C C Q L E T E H I E A H A P G H N Q E G S E C A
RANDOM_4881 K L Q I V K N P P S S L E R H L S K S K F S S K E T A I Q G N K L D T G T I F A G C E D H G G L R M F V S Y T D
RANDOM_4882 V Y P V S D I E C G L G E I K G L F N K Y R E Q E N V A I S G E L F L A F V R L L A Y R D F A T N P Y I T T I P
RANDOM_4883 F L Q A D L C V S L V V I L L D I Q R M C A Y Y N T I M L Q G S L T K F L L K T F E G S F D Q E L F I S G L A R
RANDOM_4884 R A K S L K A N D L A T F K Y V F D R R M G G I L S Y V A N E H P H E R E D P G I G L N L G F E G G K D L Y A W
RANDOM_4885 V I T L T W E H L L S S M S V A F F E C C V G Q L K N F M N N E T K L C S D L T G Y K Y M A E E H L T N Y R T W
RANDOM_4886 L S G R L S R C P E Q G Q A E G C V L L Q G L L M L V T I G A V T A V N E E K S S A V R L L R L L T K D Y S D L
RANDOM_4887 H L R T Y I K D E S C R I A Q A T R L R P Y E V D K L E R G F G A E S G P V G L Q Y C Q V H E L V H P N H M F I
RANDOM_4888 Y D V F K C T E G C L K D L L W H E C R K I P L A C K P N L E I R V S G P P D R A S I L E M P R K T C L S R L Q
RANDOM_4889 C A L Y S I F E I V N G A N I S F S Q M T R E M E K M H G F L V M S A G M A V K S L L S A I E N L L E A L K P V
RANDOM_4890 P V L L T D G W E C F S D G T L Q Y T E P Q P A L S G I G Q E L Y A E S S G G A T T D S T A V A E A M R V F G P
RANDOM_4891 D L L A A E A M P N L F P V A R H L F Y A L V A Y A V D A L A S S S I K S P W K C D P R A A V E T C E W F E F I
RANDOM_4892 D A V Y A G K A I T M L G S D T T C I A T S V S Q Y D M I N M S I R E V V G A I F S Y K K E K P G V G F L L K G
RANDOM_4893 S Q Y V Y C K F I I H G H N W L I L G E P A M W S K N H L Y T K S D I K L K G F V H F A C R P L V F S N L V G T
RANDOM_4894 E C K V K R D T L T A E P L A S A F C D Q Q T E K G F S E A Y C T T N L G H S F T Q L Y R S L R Y G G K F A G Q
RANDOM_4895 N G E D A E N H D V T M D L Y E G D F I P E Q A A Y L V E Y W G I K L A E L E N R E I G P G A D L G G S W E R D
RANDOM_4896 G E I R T G G F L R N A V R L R H Q W L L V F L G D K L N I T K F D N Q G A H R G S S H E V T D D A G P L D R G
RANDOM_4897 D P S E G I K S A K G T L M V P C L P T L H S N G D L R N K A Y I T I Q P P Q C D C R L T H Y S N L K Q D L P G
RANDOM_4898 W I S S E R R P R T R A R I F Q C T G K V G E N H V L D F R L M G L A Y C Y E H A S A K M A E A P S R A L G I G
RANDOM_4899 W S L G L H A D R T T S E V A N N P P E K L V A S T N P W S V N S T D G S K S I S I K N E S E N S E E T L D A G
RANDOM_4900 A N G I R P T I K N H T G T Y T G E A L T G V C R I A S Q A K D S S S L D E V V P D N Q V R S M Q I L A D L K Y
RANDOM_4901 L S Q S M S H I K V P S D C T R P N A G V A V L F E L S A Q T F L A T R T Q H K Q W K P G L P Q G I A R S V V G
RANDOM_4902 L T E N D V L T G I N M T F P N H A L V G Y A Q E G S Q Q G D T N G T L A H D I Y A L A V A A R S S V K G V S T
RANDOM_4903 A S M S P H V Y E P M D S V L D M Q L I L I V V L G Q W P R L E W A E S D D C N T A E R T E G G R P T I G G A P
RANDOM_4904 H G A M I K L Q R E V S A N S L I M S G F C T P P S T I V N E T V I K G Y Q P E L L P M K Y I K D F G D G L T D
RANDOM_4905 N M H P H P R H G F S L L F D L H I P Q L A V F E A S L H R L V S V G L G G T N A K D P T C F A N S R V N G Q F
RANDOM_4906 Y K P Y N L G L F T L M S W N G G I W K V L L L A P D D G T R R S G N S A R S D N I L G T T E T C N L S M Y D A
RANDOM_4907 R V K T Y E P E G C V P T S T P N R K E E Q Y D F Q N A N F A A N F T I K E A A H S L G G H R G M E H R K I P F
RANDOM_4908 L C L D G A L Q E F F A K M K W I D F E R F Q G L Y L G I K K V E N S K M L L L L Y E E K P F A A F S Y L D S N
RANDOM_4909 D V G K R C D G D V Y V K V V I I G A A I R T L M F G W D A E R A I M F S S S G T L S G V L I P C A P L A Y G I
RANDOM_4910 S N R K D A L G N D D Y I G Y I V E G L V T A Y D P S V Y V K L Q D R A S G A S Y Q Q G E Q H Y P K P R C M E L
RANDOM_4911 S L K T E G G S N M E W K H V K G E D P G N A F D G S G P G A S K S G S F V E G G E K L Y I A L L L G S T K G Y
RANDOM_4912 E Q N C R G C A P Y V S K L L A S D P V E P Q C G V E C S Y E E R E R A C V D S T E T G R S T I I R S Q K E Y A
RANDOM_4913 M N P H K E S Y D T N K A R D D K E N G A R T L W A V H K N Q G F T R S I A V T V A K E A E F M F P L K Y L V V
RANDOM_4914 N E I L K P D S R G P L W Q L D Y F C G N S G Y A G E C R L A I K V H V L V T P K E P H T G D Q L K A L N S D A
RANDOM_4915 V E I D D F I F W N R D Y G L G G P S E G G A M E T Y A E Q K S F P G V T V N L D Q F G L V S T G G A S E E Y G
RANDOM_4916 H D A R A C N L K D I F V V G Q I L C S S R R A P W D L D N N K T M R H A F L I K K L T I H V R M K A Y Q I V R
RANDOM_4917 S G E D I S R P K Q I N E K L P P N G F G K T T V E Q G P E W D T L Y R D A F L G R N P L M N A T S E A N I A E
RANDOM_4918 G A L R E Q A D I N V G T G A A G S G R Y S E F R G R E V M N L T N T G G F C S D F A I D A K L H S R A P V T N
RANDOM_4919 K R T K A Q Q P F G Y A P C N L W H D S D D F L C A L L K K T V P I F N P K Y N E T G F E E T D E A E G N G D R
RANDOM_4920 L T D I A A E V S P E P G N V C L I Y N M G L V A V K H M T Y A I Y A Q K Y T H D Q C N K F G P G Q Q R G G P S
RANDOM_4921 T G G E M S D L N G R Q V L Y W N P D Q G S H T Y I E K K Q S Q A K G V H M M G V G N V R R Y F W V S F M T H D
RANDOM_4922 K N L A H M I D C Q E Y C Y R A K L R T G D D A Y K E I S T H V K H C Q L A A C M L E S T K T R Q F K M M D T P
RANDOM_4923 N L H L Q R F D T Y I Q S L Q Q G K A E S S K H P S A V T A G F N V R H R V R S S F S A P A T A M F K R R P L T
RANDOM_4924 K R S E N V Y I I S F V F D V F C K T P L V A H A G A L F F V I V S I H R R L A E A K P I N E S S V P G H Y P L
RANDOM_4925 Y T G F V V S D Q F E T I I V R V H Q I R Y G R G P S L A N A L E S E H A T P N E R R A G L G A K I K N S N H Y

RANDOM_4926 G C A G N R I F R E F K T D R T Q D I L Q P D G N W V G V P G V I T G V L A L L V R M D Y L M A E V P E V Y I H
RANDOM_4927 G Y F F L L F L A S A S E G A H K L R T R D L V E L S T P R K D G D N V I E S E A E W Q N Q M G N D W R R A L Y
RANDOM_4928 F Q C F T A V R E V Q P P V K L F A V P N Q L F P L N P L T C F Y Q V Q S D H T L A N G G G W L E K P D M D K V
RANDOM_4929 S D L P E I T N S G P G V M N I A S H L G D E A E F Q D V Q I T T P R P L A C F H P F G S G V K I L R F I S S P
RANDOM_4930 T A L Q I K W L S N S L E P S F K E F G S L S C S V M E E L T K F A L G E I N Q D K Q E I I T G F L I Q L F V S
RANDOM_4931 A L A I L Q E M S Q E Q K H V V F T P S N D G A R L H A A I A D A L C D N L M F V P T S R D L S N L H Y S L L V
RANDOM_4932 A A L A G E D Y E G V D T I E K T Y S E E S H V S P V D F H K E V A Q C Q A D K E N Y P S A P S E Y A N K M L Y
RANDOM_4933 A E Q E G P K D K D P A Y Y V S R A E K T I Y P C A I E A I L M N D A R V S R R G E R C C I F D P L G D E P H T L
RANDOM_4934 L P K D H G F L K L M G F V W N N S G G L R S K A W C V S V D I S W R G S D K N K R P G L C T I F M M W H M R S
RANDOM_4935 K G K P I P I L T R E A T S R L R D A E N I P L T C P E F K Y F K N L A P K V F G G A S E G H A A H L V T E K G
RANDOM_4936 T E V K R Q D S C C T N R V Q F R D V I Q D P N H M K E E E V T R E F K M T A A E M H S N A E S G V N A Y P Q D
RANDOM_4937 C E R H K M L I N E T R I S L R V P Q M G G E S L I N R Q R K P S I V N R P W Y G R Q V P M K E H Q T G H I G Y
RANDOM_4938 A D L R F A K E G R I L K R I N L K F Y P T G K T L F Q Q I A Y G V A D M A T N P S L P L A C N Q Q N F S E P L
RANDOM_4939 I C G Q A A K G E E R I P L A K S T A V F C T T L D S A P K S L R Y E E L L A Y E N R D K A V M P S P D E E E V
RANDOM_4940 P E L E E N A S S A R D K D F V A R S G S G V F R A G A E I S L G A K Y G Y A Q E N K F K T F L G Q D N K G C F
RANDOM_4941 S P T H E K H T I A V H R G L C K R F H V S Y V L L K V G Q G L S Y L M N T R V A G L F Q K F A S E T S S P R N
RANDOM_4942 T A E R F S R A M A S I V S S L G L I E R R D F V L P G A S S N V G A F V P L Q N E T L G N K D K N L K R I T D
RANDOM_4943 M S I P A G L A T S D L Y S N P G W D A I R N V K G T D L R S V C E Y F L T K D N L Q G Y L D T L L L L Q Q D Y
RANDOM_4944 T A L V M T L P R V V E K K S S P A L K L L V Y V V T P G I K L H V R T L T E L S Q K G K G H R N S M I L E N R
RANDOM_4945 T D M S H A S K Q L G P L E L I D K T L G Q A L A E Q R P S T G N L K G I A M S H T S L V K I F L Y T E A G E L
RANDOM_4946 E Q K A Q T E E W K K T E A T K K L V K D A S F T K V W T N V C L G R G G P F Q G E A Y T T R P L G L P G M K A
RANDOM_4947 E A S V Q A I K T G Q A R S S T L N V E R D L A A I K D L K Q L N K M T C D L G K P F T D H M Y V I L K E I I K
RANDOM_4948 F S R P D Q R V A A I S I L K G K R V T L G N Q Q F F L D S I R Q E Q Q V G S S R D A A A S D A L A L G A V N Y
RANDOM_4949 I T S D L R D S A K S V I F K D P F S A S K G P G K E G S L L K R A A E R F F K E C F E S R A D L R L T L G T T
RANDOM_4950 V D V C S Q L A G P G E S M Q Q S Q G S D K Q D T S S I N L G C V V A I E Y I E T R E E F A R L H R Q E S V G G
RANDOM_4951 S V S C A V M A C E P F K R I N A F V A G Q E T Y S G N E A S Q P R M C G Q T S R T T G R K K T R T S P F C A L
RANDOM_4952 C R A L E S L T T E S D H N M L N S N S K D N L P I F V L S P Q K I E V H T D T P A T G A K Q E D F K G D D L E
RANDOM_4953 A R L I K T S R I V A L Y V A K T E N I F C N I P L E F Q A N P N T G L N P N R G Q D Q L P S G H A V A E V L I
RANDOM_4954 Y A Q L I G S M Q S N E T V I H S D L L E L S K N V F R G V R T M G E A D K E Y H V A N M E V R I K Q N F P T W
RANDOM_4955 Y D D D L V A M L P T L G L N A G K R P T N R D L T E G I S D V S V L A S Q V L V A K I A F M C M F T P D V A H
RANDOM_4956 R S V N K R H I E S D P L G A T R L A M T N H A P P S H S Q D I M L P H D T E D I L H Q S F H C S L I A L L V D
RANDOM_4957 E R G D F V I D G S S V T P R E I K T G S A P D S L E L T V K K Q C D V G K T V Q A Q L S A P S R T L W D L G S
RANDOM_4958 Y L I L T G Q P A S S A T H G M C E L L Y V G P S P T G V R L K N V A R T V I S S A Y S L P P L G S V I A T N F
RANDOM_4959 R T G K E E K L H Q M T E Y F N S Q K S S S H R A Y V N D R G F D S N S T V E L K V I V L A E V I S K E S C Y P
RANDOM_4960 G S F S I T Y F A N A E N D V G L L L G T F L Y F S G P L L A F D A A E L S K R V P L K S A G K A L K K L Y Q G
RANDOM_4961 K G A I S N L D V G A E G S S E G A I S A L L P G L H Q N W R A Y L I S M P C T P A S K G A A L S D S I K I F S
RANDOM_4962 A N Y L A G P V A L T H N G P S L K A R G R A H S G I I S G L S V Q I P L Y P N A L S D A L A S V K H Y Q E T L
RANDOM_4963 Q L E P V G C I K V R E R R N Q C D N F G T V W R E E V L A P H L I F S P A Q L K R L S H N G L E T T S T M P I
RANDOM_4964 E P V G R N F K V P T Y E N A A L D Q G P V I N T L P F V K F L K Q M K K Q P V G V D L A Y V E R E G R G Y T K
RANDOM_4965 V I S Q A E Q Q K S T A W D V L N K K S L T L H L K A E A P L S R F P G Y L E L S C I R T P R H A I L I S D C F
RANDOM_4966 I I E R T N G T Y D G Q S A M F E L A W E T T K F D D K P K T R Y R S R N M E N G L K C G N S F C G P Q F V L Q
RANDOM_4967 K E S V T T I Q Q F K I E L Y Y L K A R T Q M S R G S Q H S H R G P E G K R P L L K A P A M D M T V V L D G W V
RANDOM_4968 L Y C L Q Q V I S S A E E I T Y D A C P I R T F C K P Q G I D A G T D Q A S D D S G N N H Y A G E T T A K I T A
RANDOM_4969 C M C H L G E G E L Y R N M T T R Y H K I S R E R T K R M L G V R F E R N N V N K A T T T M A M D S R S L N Q N
RANDOM_4970 M T D C Q G A F A D R R R N Q R R D K V S C K A V E V K R M L N Y N F V D S K V E G E S P F S A N M G V S S A K
RANDOM_4971 G S K R F G K N V E D P L V N V E Y Q Y D G L F Q G V A A A K P T N S A R G W H E L D D W F S G S R Q L T F E D
RANDOM_4972 E K G N F R N L F D L I A G R G L F P L P L V I R N E K I S V G S A G G F L G G M K P S T N V L M P S L P Q L E
RANDOM_4973 Q P F I E T R G R V G D L E K A I D E A T K L K Y G L C H A K W R E C A M T H K P L V N L K A A S L R V G P S I
RANDOM_4974 R D V P A L S C Q K I V K D Q D Q A R P L E S A E A W E L A K E V T S G I V Q N D F L N A E V E T A L R V L N E
RANDOM_4975 F Q T K S T Q P S S V I K A P Q T F P S P S T R S A H E I G D C K A W D E V G A D A A M Y A T S A K N H L S E S
RANDOM_4976 C Y P D A Y K L H G K P F P L S R V E L W S R H E Y V E Q R L V G Q S K T L F D L E T N Y N R T A M N A N W D I
RANDOM_4977 K I Q N L K D R L S T Y G L E N A A A A H E K T D L W E Q Q E E R A G T D L K G L V P N A S R I K N F F N F L G
RANDOM_4978 N T E E P P G A P A Q N A N E V V E V C R L A Y P Q V T E K P T I A S Q P I S S P T F P C D E E V C L G T E E A
RANDOM_4979 K A E P L N E K D W S E H T V R I W T L G W P K M L T L K I D A T R W Q W I A H V S V G S D I P G F L P E V T S
RANDOM_4980 K G T F N R A S T S A M S A E S Q K G L K L G E V K D S I I P V E S D G P V A R N A D E A M A K A V S S V L Q I
RANDOM_4981 G D C L A A L R E C L Y D E P L G A E D P I C T D R Y A E Y Q D P V N P E T L G Y L A N E G L H F D A V A S L F
RANDOM_4982 K L R Q L L P Y S E Y G L I S C F L K V A P T F L G T Y I R R L A I K S K G A K R R V T A S Q R Y Y Y K V S T V
RANDOM_4983 S I Y T E G L T G S E C A F K G R L E N S V I R F A T G N A E E Q H S G A R T N Q P P C E K G L I N T P P A L G
RANDOM_4984 D V E I I N S E I N E S L A L V L K G N R E L A D L R K N F S N A K S A D T K T Y T G I N V P Q T I E A R T L I
RANDOM_4985 R Y L Y Y I S E S E G L I P I S G L T F T K G G G L P R A V A D V T M V Q G I L K L F K D I C C E D D A E G L F
RANDOM_4986 K V G E Q A N A F S K T Q K V H R R G T S N Q T A L D H R L R L K T L T L K Q V Y L L T C R V E A Y G K A A E L
RANDOM_4987 P A L W F V E V S I E P G E L V K Q S E K E H C A N N I A R G C Y I P G G P K D E C V A I G E L A S P A Q V P K
RANDOM_4988 N K T S F L G P S D L R L Q T K G A P R G L K L Q R A A H E A T H D E T T A A Q T D P T F Q F S K D L R R N S C
RANDOM_4989 L F P V N E E G Y F G P S N E A A E A L D L I I L A N Q I F Y S F T E P Y D S T P R D E V G S N L P A H A Q P L
RANDOM_4990 P N L I S V L A S L L A F D P D A Q F L G E G I G I V A F A T V M Y G M P G K M L P K L L K H G E S G G N R E Y
RANDOM_4991 P A G Q H Y A L A S D N S R S I I P R K E P Y A K R M F H L P Q L S S S F E C F E P D K R R G E H V D Q V G K L
RANDOM_4992 A C S Q Q R Y D R V N S G E D W K D K A L L I A L E M A I H V L Y Y V K V P Q V T R S T I R N Q V L D A V E F P
RANDOM_4993 I G K E R G E K P F L S A G R S P A T W P P K E A Q T L G E Q F Y M F A A D T N R Q A G I A G T Y S L N L G C K
RANDOM_4994 T V S S E L A A A S A I R I Q V A V N K N G Q D I C T K S P S P D N V K D T K I I G I G A R V P S L L A T K L E
RANDOM_4995 P N A P Q K F V K M A E V P R N S A S Q R Q W L I C M E I M Y G G R F P R I A L H S A L N I D I S K Q E I H Q L
RANDOM_4996 R G R A G V I R N N V K L A Y C L G R G I K F H K H T Q R M K G R N T K K D E H A V S L Y A I E H C T H G R K G
RANDOM_4997 P L H L L S K S G C E G T L Q K A P K D G S R M S I H P H D V C Q S A D F H K G N G R K L M D L A D E M G A N C
RANDOM_4998 P L N Q L N H R I E L D N D V S L P M V K G Q F K T T C Q G E A A R Q S H K N F L D K I P E R V I N H A T Q P L
RANDOM_4999 E S L S G L L Q A D P L V V R G K T T C Q N G V E L E L H L G L C R F H G E P S K A Y G T I N T E E L S A E S D
RANDOM_5000 R M W T Y G A G E T N M L A K L G I Q S S K A P A D H P A A E P K K C T S V A Q L N Q V I S A I S T T E V K R N
RANDOM_5001 E I C N G M I S G R T L L L S W T L I L A L P W L V M P L V T M T T G P E E Y N V N I L V P Q D V F S I L G P A
RANDOM_5002 K G T A L R D E N E A G Q N D E V G S V I D A P T F V T A L R E A W G R A L A Y G D F F S D N T S K A A R C K F
RANDOM_5003 S H L P S D K F V H S G L Q T Q S V A G V F F Y I G H R G T Y S P T I A E L A I G S Q Y E D V A V H I I V I V H
RANDOM_5004 K E E L D Q L N D A A I I T D G R Q A T N E G E P I L V A Y H Q I L K V P L N N V S A A K E S P I L R A L R L F
RANDOM_5005 L Q T P I Q P V I N E K V R A E R L E L L A N V L E E L Q Q I A G Q I D L V A Y F A D P A F E P E Y G I Q V P L
RANDOM_5006 E E G S Q E T E L K T N V S K T V Q A E I V F D T V T L S K F R R L D R L L K L F V S Q N V N C L N G V K D Q H
RANDOM_5007 R G F E E C L W P M G V I F G N N I A K T N D A S Y Y A N S E L T A G K N T S L P T V A T F F R S E L G D C E E
RANDOM_5008 Q K K I E R G V T V N M L N R I Y Q K G F F D F K R K S R E T G E Q G Q S M A P C V P A L I G T A A N Q T V L V
RANDOM_5009 L E D I A N R L N S D L I Q A V L S Q A I A L Q S A P G L G C S A M L T Y V Q V P P F S M A V R R I D I K T N I
RANDOM_5010 V A R K V E V Y E V S A L L A E T L H R Y Q G D T L L G T I N F V R R G I K K V L R V A S D L Q G L F P T Q F A
RANDOM_5011 E I A V Q I V I L V S Y R G Q Q L R D K G K D G F H Y Q D K S N T S P E V D D F V D V G Q P S A K P L A Q A N R
RANDOM_5012 Y H V R S L A A L M S N V S I F P L L S S D R T Q H I Q K T Q L L A L R Q Q T H T P S Y Q T G L A P L A T A A K
RANDOM_5013 K A G P P W P D K W A N S N R V I N S I R K N P A I L N E E K V A T I F T V R R A L L K P Q F I P S Y I L Y R R
RANDOM_5014 L H A P K T H E F T D S A N F M K F P A M D V P A G Y S T K E T S S E S P E P Q V C K L S S G L A I S L N F E W
RANDOM_5015 F E K L S G I A G E P V V V R L L T K T Q D Q S L D K K D S V N T L F E V P V F G V T H E E C A L T L I D D Q G
RANDOM_5016 G R T S K D T N K S L L R S I V E F V A N R G R G F L T F N N E V D V M P D N H R K S F M V Q S M D V W A L S K
RANDOM_5017 V I G T L I M W Y N G L L E P T R G N L L L E I V A C A I P I L H V H G K P D A T I G M E H G A G A K S R T N A
RANDOM_5018 N G T E D G T E N G V A P Q V G N F R R R Q A I C R A V A G T H N K P N K I K K N V E S S L D L G T G P V E L R
RANDOM_5019 K L G P D P N Q P T Q G D R S V G L W V I P V L G C L K V K D T Q N S K A S H E Q A N L F K Y L Q E M H D H M D
RANDOM_5020 D L E K N A G A H S A P T A C Y S I L A A P S S I H L K G Q Q V R N D R Q A R D D F N Y R G S C H N N H T D I G

RANDOM_5021 A K G T Y G A S L A A M D L Q G Y L A V P L P I C L M A R D V E A P H F E I M E V N V A G E L W G L H F F N F W
RANDOM_5022 H S K V N Q K A G N A L F V G W D A V N F L E T A R R A A T V D P V S L A D R E S H M D G I A F G M K F Q S A S
RANDOM_5023 P G A K D T W A L M T F M E T V P H P W D G R D M V Q G D C Q A K N L A S I D F L L T L T L C P L A V F P R E Q
RANDOM_5024 T A E N N F S S L V H M R T F L R P S Y F G F W L A G V A F C E I M M C P T V K P G G P F Q I V S S L T L T S S
RANDOM_5025 G V E A V S S D E V E N G A T M L T Y A A V L V H F T L N D S P L T N Y M G A P L G M A T G G F E T G G A N G V
RANDOM_5026 H E G K F G S A Q S N W K G N P Q A L I L A S V G I N G I R L T W G L S E S G S K F S Y G T M A A Y Q T Y D R G
RANDOM_5027 A S D A V G K V S R P K Y D S S Q N L L F G T P R E L D G V I N C I A D Q T Q S D E G M A P L N A C Y Y N N I R
RANDOM_5028 R H D H D G S H E T G K F K N S P D A N L G E L A F H L Y D A F Y N A Q G L I P D P L K K V D L V R M R P H L T
RANDOM_5029 E G L M P E E C S L L R N N T L R Q L T A N T D A S T P R V R G M L A H V W D L P E P L A I I Y I T A T M T D W
RANDOM_5030 F I D P I Q I S V W Y T F A S N L L L G G R V K A A A V E D L R A C S V E A L Y K P L L Q H N A A F L R F Q S K
RANDOM_5031 H H F R T V A T T W A K G V N S H E I Q E N D A R A K A G A R Y I S M V A L L D S T I V I L G V P K V L N V T P
RANDOM_5032 S G A I V Q P I G K A A W P T R K K I Y L L T E N A D P S G W T F F Q T N V I C S L L A R D K S G L L G S G L R
RANDOM_5033 P L T G F P G R R Q S L Q P E A I I F A R R Q G T P A P I P S T E I G I E D P V V M G I L C A S I Q D E S S S S
RANDOM_5034 Y E Q A N F P C L P T E A G L S L N L P T R G V D S Q N K L A F L R L V K D S K A A L M S W A R S S A I Q I R K
RANDOM_5035 Q G A L H Q G P F R S F H K H K A L D K Q N Q I G G A E F V V I V F I E L E M L G A W G A V I S L E G T E V E D
RANDOM_5036 E S Q M S L N L D Q V Y L E F H A T T N N L D T G Y N D V F E C S G T M D L E E D G L C N I N M A P G E M V P A
RANDOM_5037 P N R F K D G A G K A Q E T R N N R A T I Q V K N I D Q I N P A W L S A T Y K V D E H F T N A A L T R P Q Q V H
RANDOM_5038 K Q I P G G V W Y A E E V R G A E G M K L F S A V L V Y K V N A D T F I S Y Q E R G V K T V V E N L T L A F V T
RANDOM_5039 N D Q C S A T Y K S I L N L P Q Y N F C L F H L T S N Q I W S L W L P A D P E D V P T V Y Y N P K T F D Q F I V
RANDOM_5040 I T G D G I A T N I F A L A E R C P Q T T P N G L F A H F I A E V Y K V R N Q F K P E P Q L G R G A R F L P V R
RANDOM_5041 M E F D I A L M A A N S K H L D A I L E V T N H S P Y H I L K P G H N Q V V V E A Q L G E Y E A R A A I R E I G
RANDOM_5042 N V L T R P T H N R L V G M L K R F T M S Q H N I D L K Y L A I I A A P R Q W Q G V Y S L Y N A L H V D V A N A
RANDOM_5043 T E M A R S G L E T L I R H G P A S L S Y S I G V D Y P E E D K L G K M S N L K N A A Q D A S P A F G K S L S Q
RANDOM_5044 P Q A V C T D H Y V S E A S A R R E V A D D F C P R T S K I P V I K R H S T Q K S S Q N H V A K I A L V K G L V
RANDOM_5045 A S S K R T P A L F I E G V L T L G T A I P K D M L G K N T V V R V V I T L A A P K N S A A T I K E P N S V G S
RANDOM_5046 F V V G N S T A E D A S V K Y C E Y L G I H L E T R D R W I A E E R L D S Y E R P M S I G E N D V A T F C K V P
RANDOM_5047 L L M N M R T I A Q Q F F A I N S I V D C D L A P K K I G Q N T L R I G L K E H T F K T F K K M Q D Q T F T N G
RANDOM_5048 S L T Y L L G E E G I S G V N R A H R K T K N L Y C A H R R G V C F T A V F I R D V T V R K G G L S K E D P K R
RANDOM_5049 P P I G R T N V Y A M T V I V K V Y H E Q S V Y S Q Y R T E R M R R K W R S I S M P A V Y G D V P N K G V V M K
RANDOM_5050 F S F P A L F P A D Q F K Q L P A R N S K W T R L S R Y Y K G S N V S A F G G E G N L I R A K L L P M L Q V P G
RANDOM_5051 A L S Q A F A N A F N K I V V E E D V A F L C A A N E A K L N R D A K R Q Q L S K G F G S V A R L R N Q V A E C
RANDOM_5052 I A I E H H F V F E V G Q D R L A R S G S R D T R V G V N H R D S K G G G P D N G K T T G T T A S S K E K E P H
RANDOM_5053 A N T K A K L D E D E T G L I G K K S L A L L I G L A T I K A N V S E K A N S E P R K T I F R P A K S S A I T A
RANDOM_5054 V C A L A M T I L P P L A D N N T K S G Q P Q V Q M E P P G K N S L I I S R H K R L L A T E G S G I S S I S S A
RANDOM_5055 V R H L L I C D Y L K K Y Q A S K G C K G A R I V V G E I R H A C T V Q F G E P L Q G P G R H D F Q F L S V G G
RANDOM_5056 G Q D A I A R F R H H A P G K A M T T K L A G A T K D T P N P T Y P E L I L V D T V A Y R K I D Q S A Y A V Y F
RANDOM_5057 I C P N A A D K H L S A D K E I L S L L S G F Y Y D G P K R W M A E F Q F R R P E G N R V T V E A D A S W L F S
RANDOM_5058 P D T Q N V T E N S P S A Y L Y P L P Y N N G L Y G G Q A V C A D F D Y A E K T E I V G N H N K E A S D Y K Y A
RANDOM_5059 P A K C Y K T D Q A A V N Q R D K G S V F C A S D D T L L W Q Q P D T V E G C Q E D S P R T T I A V I P M G T I
RANDOM_5060 T A I V S P D N V S L C R S P N M T L T K P Q R R V A A V A G S A K R I N L F R G S F Y P I P Y I N I R G A C I
RANDOM_5061 L N C T I S W F F M P S P I W C Y L I G L T C E V F D S V D A D P V L A P P N A R T T T L A G G G A A F V I A R
RANDOM_5062 N K Y V F P F T A S G A E I Y F F N K A P T V K P S S N N S F A T A I S I N E Q T K A R G A D A Y R I P H N T D
RANDOM_5063 I V A E S K R G E I T I S I A S M W L R Y E A S E L T K S I V N N K I E K G P S R Q G V G I V T M F T K N E R H
RANDOM_5064 Q R Q G Y F R F D A D F S F K K E L L N H S D E C E R E G P M K I A L L H V S L R K V G F D K G R I S W A T I V
RANDOM_5065 G I L L F K V G N K I R D V E T A G A C T A K S A N A V M L K P K K P N L G L F A S R N K T H N A N Y L I F C G
RANDOM_5066 S N S P G I G I Y E A C P Q T Q V I A K N R L T K S N N L P T L A D T Q S K G A E H H T I G L V G P P C S I P K
RANDOM_5067 F K T R D N E G L Q A F L K E A V I T D A C K T G G T N C G L D I D S A Y R G I H P Q T G H I G A Y P G Q Q I A
RANDOM_5068 A L E G Q S G V G L T S F L V V I F L V T S E E S S E N G D L T Y T L A K D P W R S L H I I Y D L V G K N I G P
RANDOM_5069 G A N R I H S E C H P N L C E S D M E N A I K K L D T T L L E R Q Y A E S V M C G H E A P L A L S G K L F M Y E
RANDOM_5070 D K F Y M A N G L P A L L C P R P T S N V E D I R A Q G L W G E S N D R W A N F N H K A R V K K L C A G A Q A K
RANDOM_5071 H L R L V P Q I N V C A K H C S I A K F G M R T D Y G G S A S G G N D D L R A A H G S H A F L K L Y N V L K R L
RANDOM_5072 R G Q Q F A W M E F L T E G G E G A P A E E V T G D N R E F G V K E V E S N G N K R E P N G L P A R F L Y T N T
RANDOM_5073 K A N Q N V G S K A K I L G K W G A N T E L K L Y S D C V G V L G A D F A G D L G K G R S T P T S S A G R I L I
RANDOM_5074 F N T Q N C I S A E L I D F C K T R I M S I A P A K N G R M V T G V R I K T D I P S P T R V R M A P L K T A T Q
RANDOM_5075 Y K S E I G F G F P P I G F R R N R E T A I M L K V D L F A W A G L W K V S N R E I E K R K D I V Q E R V T V A
RANDOM_5076 L D R V I S T V L A R G L S V R F K R A M V E I K L L Y I M Q E F N N M D P D F N S L D Y N T M M F I I F T S K
RANDOM_5077 T D S L P V T I H F V A A D G K Q R I I V I Y T P L L K R A P T T G S G G D W P K P G K G V L I D V G R R W V N
RANDOM_5078 Y N D N H F H A R E G S R T K R T S P S S T Q P E I S K S L V L E P I N V M V V K I S V D S Y S G L Y Q T T D Q
RANDOM_5079 T L A E R Y N S S E L G E A G W E P N L V Q E K K K D R Y K V K S G S P R S R D D C I Q A I I D E G P S C P S S
RANDOM_5080 Q V Y Y D L R T A F F P D D L I R L F P V I Q H S N W M G V A S L A A H T I S S P E V D S G N A Y G E Y R L N A
RANDOM_5081 T L S Q V D I R G S L S E W V P S K A F L I T L L K I G V K S G A E I D D A G S I L P Q E A T H A D Q P E V D T
RANDOM_5082 S G G G T E M L E D N L A K I S F E D L L Q Y P F F W E G N L K E L A I K A L P L P E V I H A L M A Y L T G A N
RANDOM_5083 K C D K P I K P T I I P L E F T G A I C R F V T L A C S S T A D G Q Q L V E K E I V N G V V G L D H R G N F R L
RANDOM_5084 S A Q D T K F H V Y A L Y F F P E K G E G K I E A N S R E K D D P E A T P E T D P R V S I V Q A G L K F S E A E
RANDOM_5085 L E S L L N C P Y T I C P A S L E N Y G W T L V Q H I N K D I R A L E S A I L Y R H T C N D L E V K E I S C A D
RANDOM_5086 F V K G D A K Y S K E G T I P C H Y W Y R E V D T Q P L L M D K F D P K V G L A L V L G A R P E T Q L K L G R G
RANDOM_5087 I L T G S L Y A V E E K G S T H R L V D E D N S A Q M Q K T Y I V E A C S W M K Y L M K P L S Y W F L L V E G I
RANDOM_5088 C V P R Q K L T I L S S K P G L A N D E T D G R E I G G R I S L R I V D M L L G L G N N S G N G A T Y E T A H I
RANDOM_5089 M N G V G L G G I S A E M F L V I F V Q K K F H R L T N V F Q T K Q V A V M Y S T S I I P D P N I K I E L R Q N
RANDOM_5090 I F T A S V S D A W N R Q T F G A P A K K S L D D I L L K L R L M I L M R S T T D A D E D I N Y C Y V K Y S Q P
RANDOM_5091 C E D E S F G L Q H S L R L I F R C S D A F A E I I R F T K L A L Y F E V L D N T Q D S A T I G T V H M A H T D
RANDOM_5092 N Q I L A T T L L L K A R E V L K D V T V A C C F S A G S I P Q S T R N G D S K L A S R P L Y V N G T L I L Y N
RANDOM_5093 V N N S I K P N D K G Y Y Q E D I A A R K F G P G E L A E G N G I C A L L G L A Q A L V C D S S K L S F I E G P
RANDOM_5094 A G A R A G L H Y R M T S E F Q E R L K T M S S H L T L A Q L R K G Q V E I S C T N L P L I L S R R T Y T I E S
RANDOM_5095 D R A P F I G K S Q V D F H N L A P W G C D I N I F V G H L Q Q K K E A Q L N H A D E G R T H P G D V N G Y G K
RANDOM_5096 I R A Q I I T A V F L G V A S E K L I T Y N N K L Q R C S G F M C V N M T L V V S L E L A L A A E K A D P C H K
RANDOM_5097 L F L T E R P V E T P Q V S K K P L V K Y I G L G T G S S E P E A I P N A E E P A F I E V S N L L R T E S M Y D
RANDOM_5098 G I E S C G G L P T A T D A D N N D A C V E D N Q A I I L G V V S A R I N N R D S E S P P I I F N R Y A A S R S
RANDOM_5099 E A S L S E P D D T F I E G N P S A G V P S E S Q D K K S L F L C K E E S D I S E E I L G E E Q P R A T L S R K
RANDOM_5100 G I L M T E N P T S F A C G A Q F G R S G S P D Y G T H S L G V L L N R A M K I L I L P L G L V W A G S I L L M
RANDOM_5101 A E S K H Y L D K H G D V T P V L A I P Y P L M R H D S F V D L A E T F V I W L N Q A D V A T A N N L L G L R N
RANDOM_5102 V V D S G K D E L A E W G K L P F H G V E L L Q L G K F A A S G T T Q L N S C R L Q G R A R R N D H A D P S K G
RANDOM_5103 P L F S D D S P L S S V W Q L N Y R R A K P D Y A T Y T M P A T L L K P V S G C E P V R A I F A K L N N L I W K
RANDOM_5104 V A V F P E V S P R G T Q E Q D F G Q A S S E P P P L Q D S K Y W G E E A L T T Q I N P G P F D P R I K G M G S
RANDOM_5105 I E I K I D F P E D C M P S S C E D F N D K A C G R Y P E R I L G A L A I R G E N R L I P W G D G P H I W A V G
RANDOM_5106 P K H K S S R A S F G N A L Q L P L E A I I P F Y L Q L L S A D L A I A M E D L S G K V Y I T G G R R C L K W Y
RANDOM_5107 L P R S I S C E V V G T I V V E R V S A K L E D G Q E K K M A L E T A L H L Q Q V E S E D S A L D T S L N R Y T
RANDOM_5108 T R C S T T C R T P S T V F I V P V D S G K G T T A A L Q N P T R K E L D S I Q A E G H G Y T Q A G E E K Q R G
RANDOM_5109 I V E L K R F E S K V Y I P K V L E V S P Q K H L V F G V L R A L P G N I N Y L S I E A K Y H L A S T T L V Q L
RANDOM_5110 L S L G L C I V I T N V I S I M F G K M L L F L E A D G I V S T T K L T R Y A S A C K D L I E T V E Q E Q E V D
RANDOM_5111 L R A L M P L V N W L H A T L I T T V S V V G C N P A V Y T R S R F I L P I F G L A K I Q E L K K G P T Q H L N
RANDOM_5112 L G K E A R E I P Q T T V A G D D Y K H V S E I A L D T L Q T D N I T L I A K V M L D P F P A V N N I N Q P A L
RANDOM_5113 G V R R L Q K D G T R P L S C S E S S F L S A I E G P T K L C H S G I M C R V N D T K D F V F S S G A A D G A A
RANDOM_5114 H T L G L Q I L V T E E L Y G V A S A V V S A P G N W K W D K G G V K C S K R G I L P P G L P V G V D H A Q I T
RANDOM_5115 K S I G N R Y A K K D E S H V L I L M S Q T Q V T Q H G Y L D G G C V G M S T M E D P L T I A Q S V S N S G F A

RANDOM_5116 R Y V E Q T D Y N L A Q E S V H S A A N K D T T W A D N H E I V L V N L C V K G T G I R S T N S A I T T P V K K
RANDOM_5117 A E K R G Y K F T Y M Q Y L L D G L L Y A K L E P I R E F K F S F A K H T L R A S V C T Q K G W C S L K L P Y T
RANDOM_5118 Q V H E L A F D V G L N S L L P N T K V G F S Y L T G C G S W A L Y V E H W A S E A V A Y A M D F S P L T R R L
RANDOM_5119 Q S G N I N V D P K L H L G T G P I D L E L W E C R L N A R D I R Y E K T D A Q A V V R I I Y R K Y C L D H L L
RANDOM_5120 S T T L I S G E P A H K G I G L W N E Y N D T L G F R A R E G A T K V I S G S M G V V R W S A E P V K S Y T C W
RANDOM_5121 I D R G S P R T Q S V D E L L D V N L F A A L Q A H L M A I V F S F V I C Y A I P L D I D L S E I Y G R Q K V P
RANDOM_5122 A V G L D E I D D L I H S H S N G D Y I N T K F K W P T R I S T F P L A L L E N F H F Y L G D D A T G C L Y L E
RANDOM_5123 F Q L G I F P P Y K N M R K G L L G L W D C L L I T N D A A N A S I N E L L I G F R E M Q T M D E C N P V T L M
RANDOM_5124 A S V A A Y V F V G T D Q P A G I P D E Y M I K D S F A H T A F G K V R R R V S K G V E L T L C L L V G L I R I
RANDOM_5125 V F G L T I D N P T S L N H P P D L G Q M V T I G D A P L Q Y R I P V V A A N E E R S D C R P V G W L C R S G T
RANDOM_5126 E L L R A I V L L E V L P H N A A T T N P I L L L L Y A Y T K A M F R N S D G R Q M T T L E V C H E G T P V G V
RANDOM_5127 Y G H S L L L P N T S I G E Y T Y N L V G E P N S H L G A E S S T M V L N V I M V P A I G L A E P W K E C A S N
RANDOM_5128 E V A S E S K G A H M G G I Q S Q T V R E S H Q T Q G H L R K I D C Y G E Q L E R T S H A R E Y R S S A T I G L
RANDOM_5129 H E E T P P S E G C D G T V N S T S S Q N F D Q S F T H K G L T G W T K A F L T N L Q Q K I S V L T F N C H P W
RANDOM_5130 V F I K S K A W V F M H C P T P T H W I S D Q A K V S Y T E L W T K M S K A P P E C K M K T A R T P A G L E Y E
RANDOM_5131 N L A P D V K H E G F Y E E E D N K F I G I Q N V E E L H Q C R D A N D S K K I N K Q R N L A W K E G L H P C E
RANDOM_5132 R L V L G E S E A T G L V D A M Y F L Q I T R A L V R E D E D I L G L N A S V V D M D E A R L L T P I E F P E G
RANDOM_5133 P Q I R D L A L V K K I I D S V V I T N G S Q L S E K Y R I S L H I L P E L N I H L I V C L T T H F W G A A Y L
RANDOM_5134 I N K F N I R R Y P M K S P P V P Y N S L A V G S H N R P M L A K D S N R W A I K D G D V Y I V D M G V V P P Q
RANDOM_5135 L S N N P D S L K E L E A E T A E H F L Q I K S F G T K F A K C A F V L E I G H M V E L R P V C I G S I K L R L
RANDOM_5136 F A V D N T A P Q S D C I G G W K M E A S V S L S S S Q E R N G S N A Q P P P G L R P I K T Q M I R G T Q Q K F
RANDOM_5137 G E A M A S E G K Y T P F V G A E I T V L F L P E G S Y C F T E A A I A E T D S A E L N L I F R I L I D Y A R M
RANDOM_5138 A D W I V P E D T R F I W Q Q K T T E E V A L F D D A A K L A P I Q G V V C S K L E K K R S H L G N N D M G R Y
RANDOM_5139 Q P T R V E S P T V E A N T D S G H A R T K D K I G A L R G K L G H R T N S I A L K P G P Y Y C A A P K V E L P
RANDOM_5140 L L A F I K D S S S E E T M I T S F E E R A L A S G N G P E V D S C S Q M L E Y Y F T R W N L N G I S E E D K S
RANDOM_5141 G P L N S D V V E Y I H G L K R D S A R S N M V I T P A Y K I K S E M Q K R I S V K L V V S G Y F C E V Q G K I
RANDOM_5142 E A K A L P K T I R S L F I G V P L G T A P V F E A D R V R V L K L R R D A I T K S L R E E F N F P V V E I P D
RANDOM_5143 Y V S F K A T E A V G L A S G S T D G L G Y L E G I N R F G F G K I P Q E V C R K V L P E P S P A C S D Q R W S
RANDOM_5144 T E Q I Y S I N H K W S M Y F V L I R T L S Q S D Q E Y K G R F E G A P A G E Q S L N R W K P P P L Y N L G D S
RANDOM_5145 R N G D V D C L Q A S G G E V L T E Q T L T D S H P Q Y L Q K S K L A H C K A D G G V N F C P P A R R R H Q D P
RANDOM_5146 K G L L F R I H E P V R I A V D S G T D Y R T P D E E G G Q F V I A R I N K D L R G S T G A I T G K I L P L A R
RANDOM_5147 R T N L M L S P L L Q K V S I L A D H V K E G R A F P Q G C E G S K N S T L G Y E K K D L T G S W M R W E L L N
RANDOM_5148 S W S E T S I R V K E E G K Y M P I I W V N C V D I K T T T I P G V W K A N W S T V E A P F I D G V V L R A V A
RANDOM_5149 S Y I P Q K A E N N I T S A R L D A V C T S L Q A D K L P Y R P E V G H Q G G L I K T E V E L D P R T C E S L S
RANDOM_5150 L L R L A N A E V A K R A S N R Y T I D W T L K I H L E Y V Y V L W Y P F R T Y G P C H L Q T S R P G A D G V A
RANDOM_5151 V C G T Q T L N D T P Q K V K D E I G Q V E D N N E D N S Y E S P S Q E S H F L A L T L D A V D V L W I K S E V
RANDOM_5152 V E S M Y E K S G G A G C S L L A Q E G I E A R Y Y L S F F D A V V A L S Q I R P L C V Q T S N C K R P L Y R G
RANDOM_5153 A K F D D A A K I K V E V H Q T K Q G K T K S R A V L N Y I T K A V L F I A E F T L D P E M R E I G W L L I C S
RANDOM_5154 D D G I L R M E K L V L S V T V F V S L G I D C N H S A N Y L L H I K T L G L T Q I R G N F G L F G I I D M N S
RANDOM_5155 Q I M C V T E S E H P T I E Q S T D C R A A S S G Q C R V S I V T M L A N C D K T S L S Y D A S K R L I E K G N
RANDOM_5156 R L Q K S V A M Y P F G W K V L R W P S F L I K P V M E M S Q V V S L M T W I R G T A V E V N I H R V L D H E A
RANDOM_5157 I D Y G P P N S L A G T P L R W F L N T S L I T R K A E H N G D L V I Q L S R N A L Y G V Y S I E G L L E N L L
RANDOM_5158 R L R T Q V V G E L S L D P E E F T R K G D A W W L R T A S Y E Y W K Y Q E G Q T T S E L Y T I V Q R Q K R Q F
RANDOM_5159 P K M A M N K R R S H V G P D N L F T K A D L P S E E P D G I P P P E L A S H G E S S F A Y F I P Q T G L D L D
RANDOM_5160 D V L K F F G C A S Y D N N C T E Y T A A N Y P R R W Q Q G L P Q I T P T L G Y S F K E I V A L F S P E D C R K
RANDOM_5161 K L V D T K D I G E D V G T A K S L V V R S P L L C M E S L I R Q K R V L M N Y I A N N R N L D H R N N L W G S
RANDOM_5162 P V N N E S Q L E A K L T C R P Y L G T K D S T A L V A Q G T L D G K A E W V E N N V K P V L F L E R S A L E I
RANDOM_5163 I E K M K W P T S K C Y V L E E P N V K T P V V V L Y S M D A M R K A K D K D G R A S P L T H G S V L W G S W Q
RANDOM_5164 N N K N M M S L A M S Q K V T F G R Y K E S D I G H T D I F Q K L K P Q D G Y Q A Q V R G Q L N K S R R G Q E K
RANDOM_5165 W D L M Q S S E D D A T N A A I P L V K C E I R Q A T S T G A R R C C E I V W S A E T L K Q N V L A E R V A D H
RANDOM_5166 F C D P K L N T D Y M P C Y G S T S P L Q S L Y E G K L Q D R R S A T H S F R D A G A V K Q F G P G M E N L A E
RANDOM_5167 Q S T P S K G E G G A R W H S T K T L A M S S T G K R T V K T A D V K S P V G V I K W W V A E L S D A C Q G V V
RANDOM_5168 A S S K V G K P C G P E S L R G G L A K L R A E S S T R D S G F K Q F M E G P Q A Y S D G V V A T P C L D F L N
RANDOM_5169 T I A G K V V N K I L E E T G K K K I K F Q G L N C P Y A G Y S F N Q A D L A L E P H L V N V K P E K L S T E M
RANDOM_5170 E D S N W G E K E V D M K R Q Q P E N E W K R G L V P M M I G T L T V E N L I G G I A H R G G L Q V A P A E E S
RANDOM_5171 A L N R R A S Q R C T A L L G Q C P T F A G G N Q N Y V E K V C E P N V S L A A R Y Y F D S L A N T N D K I G D
RANDOM_5172 R G C R C G F E T D A F K E A R F L S E T G T L A L Q E N E E A Y E K V I T F Q K T D C G K F G M E A E G I T L
RANDOM_5173 K K T W A E G I K R Q Q E K E Y G I L R R E E L Q F D D H G A I W L A H A L S Y M W E P F G A T A Q I V F H L Q
RANDOM_5174 D G I L A Y T P P I R Q Y L A E P Q G E S I L Q H F A E K G L F F H T T P V N Y G G G F V N S E E K E P A L V E
RANDOM_5175 Q A D F A C D L V F A T Y G T P S A T P F E G T G T S I P S S P G S P P N R K V F W N D V L G P R G G E E L K N
RANDOM_5176 S T M Q R T G K P C Q R R T T F L R R D C L D L K V K G V R K V V E K M P Q K L G I T D D G Y D P T T E G P E T
RANDOM_5177 R C Q N D Q V Y L G Y P E T D P I K Y R G H S A G Y Q P K T H P K A P T R H C S K R I R R F A G M V P N V V L N
RANDOM_5178 K A L T T T A V F L A Q Q Q D F E K G Q S L P S G G I L A L C D A F Y L R G S D G G R S R A G Y S T L L A A R S
RANDOM_5179 I S Y H T T I L D E V T G S G I Q G Y A S M D L K M E M A S G L I R Y A G K F S K V L L F L V K L K F Y A N D R
RANDOM_5180 Q S G E L D E Q F S N E L E A I S A M I L P I T P H K S P G R W M E A V I W E G L T E I A Y G N N V T G R S Q I
RANDOM_5181 L P S I E P Q K S K A Q K S S A L H N Y K S E G G E G T P L V E I R N V Q I S P P R E E I D S A A I T P D Y T G
RANDOM_5182 K C F K D S V V L L N E V V I I K N Y G T L A K D P L Q L P Q E W A Y E K T I H Q I G S V I Q L D G N H V H Y F
RANDOM_5183 I I R V L T E P S Q P D R L A L S R E S E E L E E E S Y R S I D N N Y I P K K N G L S N F Y K I E F H L L L C R
RANDOM_5184 I Y Q A M K N S H N N F S A E M Y I P I K T Q R P C M P K I F A H S K N Q S K G T G F G T R S K N V Y G F P P D
RANDOM_5185 L V Q K T N I P E P T F L R H R V G A Q E R L I N Y G V L V K V G Q T S M S C I P E P N H E P E I N E K F A N M
RANDOM_5186 K T T D K L K P K C W S P T S V K N K E H A E K K A F A D V D D L C N G R V D S C D K E L K L T N E I Q V E N L
RANDOM_5187 L L P S R V E L F D R V D A D S V S L L P A P D W L T F V R C C I A P L I H P R L L A V A G Y I A G R G T L Q K
RANDOM_5188 K M V P Y K D K N L S M R Q C Q I N A D P K Y E L E D D Y P I L T T F D P I T R Y G R G T R T S D I D F E N M T
RANDOM_5189 V T G A P I T F G Q V V K F S E Y P R V Q L Q G K R V P A N S K G F V R A F G L P P F I R T W D Q L D P K D N H
RANDOM_5190 L R R H D T S W T S C R A F K S L H A K D N P I G K S I E M K F P W P N A T H Q S G V S G L M L E E D C V G S N
RANDOM_5191 A F P L R K I Q G P V S G K E L M W L N L F Y S E A L G A A R I M F Y K Q L P H R Q D A A L D K H Y A L N F A D
RANDOM_5192 W T K E V Q W Y E D L N P R N S T L N T D V R G T T L L F C F Y E S M R T C K F I L A W H F T V M G I P K G D Q
RANDOM_5193 G K I H Y C L E P F A R L D A A L K K L D A V L K M K H L T V L E L M G K V D L I M A H R I A G G A V Q G E M A
RANDOM_5194 T V G V D P I L A I L L L T N L S F G L R E I T E R H I P D M Y D L L E L I R N K T L K A A T F L P L E E I E T
RANDOM_5195 L N E I W E P L K A G P G K G K V H L F I I P Q V S T Q V K A A A S T E Q I I I L F G S S S I H H N G A R Q Y L
RANDOM_5196 S N G D E A L R S K G V P G S D F L G D D F Y H T C V G P N T H S E N G A K N L V G C Q C A L L D S K A Q V A H
RANDOM_5197 N M H N A Y N S L G Q S V A F P S T F N P K T I H K F L A T G T P Y G C E Q P L A H N Y A A T V N G K G I G T R
RANDOM_5198 E W Q K K L K R G T L L Q S K L A E R E F L L S I F T E R P I E P S T Q K S V N G E P H C L T R T V L L A V A F
RANDOM_5199 K P T V D A L T R S L K P F Y N G Y I A E I P Q E T A D K A L A R L D V G L E S A T K F S Y H I R P G I D G L D
RANDOM_5200 Q Q L A R D V L I L V F K E Y N S V R W P P E T D K V D K N R A I R G G D L Y N R V A L E L N A A F A K D G I F
RANDOM_5201 P R S D A W A G F S E Q I I P A S L Q V N N L D R R M D I E S S F A L P E L K E H A G G L C L R S P R L K E A A
RANDOM_5202 V R G K S F A D S A F S K H R K H A V I F L A L F W Y S T I N E Y T L V D C D V M H M I Q G C V L K E P Y F D K
RANDOM_5203 P N E R T M K K K R R H R N V E T Q Q E T P V Y N Q M D P K E V F F R T I G T V N M D I Q L I Q H S D C F V A V
RANDOM_5204 E F V G G S A L M G N A Q L E W L V M Y N K M D M V L S A C I T D I P V G T T K K L R L T L L L I T T T A A Y A
RANDOM_5205 I L C D E T S W G G S S K I F G G V A I I I Y S P P N E N C D E D L A L Y K L T K L V Y K G N L N P L G S P M G
RANDOM_5206 V V L E G H V V V M Y W K L K R D D I L V A G P P Y P G A Q Y V D I L D T T L E F Q P L Q H A L N T S G Y N A A
RANDOM_5207 D K L T L Q S M D E K R D M K G E R T E V V V F A Q Q Y L D V L L H Q K K Q G M L S R D S A I I V P Q V I V Q A
RANDOM_5208 L Q P T P D K T W F I N W G K P E D D N N N E R L P A F E Y E T A L Q D A G D V R K T L E G R A I M C E D F T A
RANDOM_5209 I I A I T Q Q H L P K N A K G E K W G N G V V R Y E A N A G G Y K N T M E C R P Y V W E L W F R N R A R Q A H C
RANDOM_5210 N S G D S S V I P L D G L F P R A A F G L P Q K F I A P S L K E C L T F Y L P P E T E H G W A N N Y Q L H G S G

RANDOM_5211 R A T V E F Q P Q I N A K G P F P T D E I A F S R P P C E E T L T D A G E V R Q S A S S G F Q K C M G T L Q D K
RANDOM_5212 D M L N E S D H N V C G L D W T G H I G G P A P S S G L K R V N L R A L S F D A A G K D V R H Q H Q T S R L L P
RANDOM_5213 V Y D E K L E D I G P E I R M F N S V M I L M R S A E D G K P I P A Y W A V G R Q A K L A A M T V T Y E S R K L
RANDOM_5214 P A A K E A R K L G I P C A S D A D K I E W P I C K F H A R K K P N L N L I Y R D Y Q N P H H Y Q E L A E T A S
RANDOM_5215 L Q R E Q E L T K G V T D Y L S N G T K L Y G R S L L A L R P Q E A V L P K G T V P D L C F G S D E P H A T S L
RANDOM_5216 A R F D T R S E Y M A E I I Q E N R F V L K E I P N K P T R Y N S L T S E P G L A R T V L F V R V L Q K N H A S
RANDOM_5217 R A D E G T E A S K K I F Q H A D P D N F Y S N F T S G Q L T L T S D I D W T G N T P K V R A A G R G C K F E G
RANDOM_5218 E Y R N S D E E N E Q S R K Y V G E P T A M Q V A G T S P G V L T R P T L Y S E L I G T E E A A S K V I V S S I
RANDOM_5219 P R C S N L F V G E V Y E K P I S P I K E T K L E T D Y G D G R L C E R L R I E K E A V Q N C N P E G A L I P I
RANDOM_5220 A S G Q P G N V E V K L P R A N G F K A G V K E Y I R Q W V F A G P T Y I P P Q R S Y T P V R T G E Q G G P L S
RANDOM_5221 D T T L L N L E V R V K G S S H K M P D P G M H F T G G K G G N L E V V G P K Q P Q G N D S Q G V K G E L G P G
RANDOM_5222 L S Y S I G P S I G I Y A L V G T P R N V F R I G H T F T I Y L N I P P Q E Y A G R Q N D N K F T T T I T G G N
RANDOM_5223 G Q L L G W L F P P L K T I F L W L L K A G K V D I I R E A I I M F V S I L A Y V A H R G I E I L G E G V T A R
RANDOM_5224 K A R V L G A L I P K A N G A S P C A N A D L S G C P L Q E V R K L L N A E P Q D V G A W L P Y F R Q M S L A S
RANDOM_5225 S Q E V S R G F A E S G A Q D L R A V F G Y R K E R E Q S A Y Q P V E A K D F I S P S E F R E L L A F K E I N V
RANDOM_5226 A H Q K K L I D E M S Y D T E A T F L G K R T R L E F I A R S V K R F W A W P I Q T S L A W S E L C K N Q T P A
RANDOM_5227 P F H S Y P A G V T K S T Q A F F R S G E S T D V I S N A G G R L A A I D L N V A K T F T S K Q E N L Y I L E I
RANDOM_5228 T G I N T N T V G G D P L P I P P T N Q H D I P P Y A A T T K F S T N V I L V S Y L D E V A K D K A L R I D N I
RANDOM_5229 A M K Q V S I I V N T P A I A N Y G D L L L M Q Q N T L V I K L L L Q T Q P T D E V V I G E G V H L S E G T E I
RANDOM_5230 C V F S P E R L M V D G S C Q T P E D I S N P I M G S L L W C K R I L D S L S A D E L V A S F K G K I G D P V Y
RANDOM_5231 K N Q P V R L S D Y C V E Y Y C P V Y N I F I V V Q L F G D V P N A G N K L N V Y T I G A G N D S V F M V E E G
RANDOM_5232 F L V H V F D P S E L P A A A E M W M F G D G Y I I N A P H Q G G F G G A K N Y K H H D C K G P A H I P A L N P
RANDOM_5233 T R Y D R N T N K L V W D C W L C P A V G N K E E L K D C E F R D G R D T F S L A V T M Y K A T G K V G L V I Q
RANDOM_5234 Y L K F S F P I A H D K G S L L L L S L P D K A N A N Y Q E G G R P N G A S D A T G G V L N L E I K D I A Q A V
RANDOM_5235 N P I V T S R M C G V K K A M K P N Y P Y A R S E N F D L R G T S A R K G L R A T G H T Y K Q H A S F E L T Y G
RANDOM_5236 A M R D L A S R E G L A A W T E E T T K N A P E K G D P S Q Q G P E S M L L S T D L Y N D T G V D A V Q D G E L
RANDOM_5237 M T K I A K F Q N E L S R Q M Q A K Q E S E N T V N Q R R A V L G K T L V G Y E V S D K M R A A T C N P F V S F
RANDOM_5238 F V I Q L D S R N G P R A N A S L P T M A N G S F W S R Y W W E A A A L F R S S V N N A H G T Q G T L I A T V P
RANDOM_5239 I L L F E W L D E S N P M M P L W S V K E A L R A H S D D G M Q T E V Q P L I R N E I D I K G A E M N F K L V A
RANDOM_5240 L A D S I A G I G F L T V D D T L L Y K C V L Q E N G V E F V T L G L A W F G L S A D G N A K E Q N D L S E L E
RANDOM_5241 N H E G S T L S A V N H L A K N Q I K L W G S T C G S L F H Q G E Q K R S K A G C I V K K G F L A L P A V V P H
RANDOM_5242 N G M R I R R L L G Q N F T P Y G L P V L T G P R N V Q L Q T P C P E E H I C G E G E L N E M I N E E H K V T F
RANDOM_5243 T D D E L E N D D S L T G R A I A L Q P E Q I R C G T K P E R F G M A P I T H G R M P K F T E Y V M R I S K P R
RANDOM_5244 S T L D E T T L K L K A Q K P N S L L A Q V R V I A K T L P P N Q A K I H T C L E G V N K G G P R Q C Q E N H V
RANDOM_5245 R G E A H N I F I E S A D L Q L R N N D M L S A L D M T T R A H W A L P G E P C P N L I C G K H P A V I Y A P V
RANDOM_5246 E W E H N D I L Y Q I N P T R G A W R R K G I S R A G N S M F I V S K L S K A D N E F K L I K D S S N P I V N Q
RANDOM_5247 C L V Y P E Q K L N M E P V K I G V F S D I Y G F V V P K D G N F T K S P V A L V D S I K F S S K L K K K K Y L
RANDOM_5248 V N K R V A N F A D D N E R A G G N I E L V L H F M A W D S F R L R V I E P D K D L R P I E L R V W V Y F S I E
RANDOM_5249 P G E A N G S F R L E K L R L S N S M T Q K G Q Y S M A V L Q I S E V L G R G S A N G P D E L I V T A S Y Q A Q
RANDOM_5250 W T V N G E Q G W P E K T Y V G K D N A G N S N G T S Y Y V Y G R A S N A V C D E F I S Q Q V C L T Q Y L A L V
RANDOM_5251 I S E C T M I F P R D E F I G L Y Y M A V H T K A A P Y I L P E H V S E F M N L Q R G L L E G W P A L T L K A N
RANDOM_5252 L D V Q A S T P R G R E C E A G Q Q L E P R E A Y G R V Q F A E L K T A S T M S L R E W L H H M L Q P L E T D A
RANDOM_5253 K G T H M G G F A R P R A L D M Q L V L M N F V N I A Y E I G A Y G Y A R L L I E A H I K P I I F F E E S H A Y
RANDOM_5254 G H V C V E A G K E L T P R S G Y T E R L H T K I P Q I N F R D A T G T G F T G G L V M N K D K E P P E Y T R Y
RANDOM_5255 T S A D A E I T P E R R N R H P S L F H I S S V Y S Q Q D T N D S P F K E Q E A C E P P L R E D A A Y A G A T A
RANDOM_5256 D V V N G G E V K N R P K K L I N G H H I T G L P V G G V S T K I E K Y A G G L L E G G K L N Y E D Y Q R V G R
RANDOM_5257 T V E A P T P L P E F G V F W R D A Q A P I A P L R N Q L P Q V P S L Q Q R G T N T P Q A N D L F G E P H D K G
RANDOM_5258 W A L Q K S F W D Y A W F A A L P S C T A R R S V G Q K G R T S Q L R S V F S L I I G E P R Q K V V Y Q A K I A
RANDOM_5259 A N E A P S A D T L V V I G N I E L E Q R V W S T V N Q L I R P A H K V V A Y D G E T V I D G Y Y D S G D S A D
RANDOM_5260 A S A T D T T I S P P E E I A H P F F G A R F L A V A P C R I A M L A H S G S C L N G S L D V A E Q T S V N M K
RANDOM_5261 S G H H F G I A V H V Q M Y A Y Y R D T A E N I F G T I R A R Q N V T P C S C G Q D L L S D D F P I D K Q K A R
RANDOM_5262 L A P A N I R G K H T K A G S D L S E C L F K L S D K P V S A C G G Q A G L K G K E Y I G R D W W D Y T W P V R
RANDOM_5263 V F V R Y V I E L R Q K P I A E S R Q L E I H E P K N G T P S N F D G D T G I L S V H P K M N G G C W Q N Q E S
RANDOM_5264 T D L I H E S W R I S C G N R S A G S D Q V V S C F P V Y A P M D L A Q F D R P E Q S T R N Y R R I E V P T S L
RANDOM_5265 L S E T G R K L I L M S M Q L M N N L D N R N C I A M D G L L G T G S D G H G G P R P S F S G Q A K G T D P T L
RANDOM_5266 N S E V V A L L F K T I V L V R S F Y A H V V K T T Y G V V Y G V T G S A D R E S I D V F L P L D L D N A T E L
RANDOM_5267 V F G P V K T Y A Y I I G P T C G I D P T V E L M N M K K L F I H P I V L P T L S L A A K F A V Q S R A A R G L
RANDOM_5268 D H V N S P D Q T K F D P K S F G R E E M T Q V I F G S Q C F A A L T W S V R R E A D Q G S L L M H L K E V D K
RANDOM_5269 M K I L D N G Y M N K E V S T Y I I F T R R K G G I L V V E M G D Q Q L V A I G V M E T P D R L T D Q E F G A K
RANDOM_5270 Y W C V H S K G Q G D L S F R F K Q H G S G N A V V I M P E S Q R T L S G V K A V L E G W A T S E Q I N H V E E
RANDOM_5271 I E D Q V G A A Y T S L R L L V A I R W V R N Y L R Q A Y F E Y L G D A Y Y G S L G I D A I T S S P S F E K A W
RANDOM_5272 T A V G K C N W P E C V I Q S P R S A L A G T S F L G R E I R V T T D V R R W G A R K G V G I S S D I A T T E A
RANDOM_5273 R M K H F I Y E E E A K M G E I V G G E L V V K I L G Q A L T G K K E F T T M F N E R R M N K R Q S I I N P E L
RANDOM_5274 C Q H I K V N A F F T K G G G E E G K E L A N E S F G N F P S A D Q Q E L P V G M K G L F A V G E E L A A I K Y
RANDOM_5275 G I A K S S K K D S N H F K T P A G I I Y L L D T R V S I R I S D A E I G R Q E P N P G G A K Q V V A N Y L T F
RANDOM_5276 E H E W T H T I C T K V L L M A V I A G S K I V V S E A H L N G K K R K H A G K D A E N Q V V K G Q H P I C R L
RANDOM_5277 L Y G K S I M K H E W A A A R V G D P D S V L N K A P A S L G T Y D C T K A A S D S S G W Y V I Y C E M G S Y L
RANDOM_5278 C V D A M S D R A N I T W A D R E H L T E A L Q S G K L S H I F N N Q A P R N K R A A R E S C G E I K V T V A P
RANDOM_5279 T G A V R L V E W T N S F D G K V M N L L E S R A T S S K D K D E Y I V G P W S L S G V A F I S S K R Y E V L A
RANDOM_5280 S G I T V D E V S D V T L G V L Y E G D T F V I F G Q G D L S Q R E I A S E D K S F T L V R Q N A S G P H P F G
RANDOM_5281 N S G D V R W V K Q E C G R V W T K Q F K F A F A P L F K I P S K L L V N K H I A S L L P L S F F V T I P Q K A
RANDOM_5282 P R Q G Y R H A D A D D I W M G A N G M P Q L E G D K D G V V A L Y G G D R S N Q H K A P G L V G T L N D L Q Q
RANDOM_5283 V R H S F L G L Q R Y P P A V E A F A Q K E S V N L V I Y T I R E E Y H C L G L S E I D Q L A M G V S Q H W F P
RANDOM_5284 S E G E M I A I K I T G Y V K G P S T F D R A D A A R D S N W A G F A T Y N A H L E K H G N Q G Q L I G S L K D
RANDOM_5285 A G R Q V R L D P S H H V A I S L H N A P A K L K T N P F S L S N L L K D V D N R K V E P A H I L Q T A L I T V
RANDOM_5286 P Q L A Q A A R A E D P E T G A Y I P I S P A N L Q K I P G K L Q T I P F A W T M L I V D P V D T T S D E H A A
RANDOM_5287 L E R F V P D S M G Y I A S N D L G T L S S S L I Y S E K G K Q V Y D L E C P R N G K W V R S A V P I A W A K R
RANDOM_5288 L G A N L A T Q S K Y S C P P K S K S Q T M P H P P C S M N K L R P V S S K A L N K T F E E M E T K L H G E T D
RANDOM_5289 G D D G A L T S V V M D A Y L V K A E T K S T Y E A I S I C S G N E F T H Y T H I M D V R I H E F G F S K S T P
RANDOM_5290 K V C R K L R L P E R V Y K V F I L V G D E S K W V R C C G G T V E D I A G G A I D S D G F F L S M G I G I K Q
RANDOM_5291 H E V D E S D E G L K T Y S G A S S K S K I Q V R L S R P R Q S N V G H V L E T N Y M Y N R H T E Q K A N Q Q I
RANDOM_5292 D Y K E P D Q I V N P I H I A P L L V N D V N C P T L D T D L T A F D L K D L V M P A K A D T D A S P D V S E F
RANDOM_5293 E L F N K V E G P R S F S S G Y A E G E E R D C P A M P S R S P S I N C H L W L E S A K S G E G T G P P R G Q L
RANDOM_5294 V G C A D G S C G S G V V M Y R D Y S K V D D M W L D A I V N G Y S M L R W L R K F V T K N L D W R L V H K S D
RANDOM_5295 F P T G T D T L Y H T S S I Y C D G Q K S A V C M K T F A R P W W E R E K K T Q N G A L M Y A A E V G A A N L I
RANDOM_5296 E N D M V Y I R K T S L K Q P C A S V H N V C L P K L P K F S Q A D D M F S G I A A R I S I P S G I G R T D G Q
RANDOM_5297 Y P G F R W L V E G F D L V D S W R D C K S Q S I S D D D T R L P L S L D Q S K S A A F K L Q I N F F K G G V A
RANDOM_5298 T Q P A I K T T L Y P A H A N N P A L E L K N A G L D D I V K Y A F A T F R L D R E D S S L R D I A V P G H K I
RANDOM_5299 G R D L F M S Y I L P E L I V S A R Q V G E N S G E R E T F E T V E G A G S I V D S L S F D V R V A P S V T R T
RANDOM_5300 L D T E I Y C S C T Y R Y Q F T Y E I D T A Q N R S L T H G K V R D I V M G T F G R Y S E T I N P D V K A S N L
RANDOM_5301 V R D R E E E C Y H Y G V G L A G A F P V Q A N V S E S H A Q T D Q Y L E V K D T N Q V S V S T Y Q S P G S E T
RANDOM_5302 K R E V A A E T L S V L L I W L T Q I V E P L R R L I D Q V R K N A V S I D G Q P V S V E P G F P N E R R W G Q
RANDOM_5303 V E I F C S P W D K V L A A A D M F H R T R V A H S V V R H D K A D A P A L F A A E S G M E H A A T S A G D H T
RANDOM_5304 T D R A P K E V V W D D R K S E Y T Q G D I G K W D R M R P E L Y D V F D N G L E I E V Q D C I V S D T L A L D
RANDOM_5305 N G S V H V T S K G D D L K N I L K N E S G P T T K K K S A T G G I L K G N G S S P A V Q N T K R D D T R Q K Q

RANDOM_5306 M P I L N C A K G L Y V S A T A E S G R K A W S N L L K V L E L G S V P T L L A L P F L E N G W T Q T K T I P L
RANDOM_5307 P P G D Y G M A D Q A L T Q T G V G Y T Y T A R G S Y T G A V T D K P L T F A V E S K E N V E T F C H D L L I D
RANDOM_5308 A E L D P S T A T Y G L T N D T S L N P G K M A V R V A K Q L S V T L A A V K V N L H D G K F S V D S P K F N K
RANDOM_5309 G I G L S M I V A V R Y F N E A K Q K M Y A A I C R R S I G D P R A S G H K M N R K L V E P A L N L R A L R W D
RANDOM_5310 E N A W S A W E R W V E K G K G D V A I D E K D I G V D A L V G H L G S P W K T Y R F S C W L L G G E N L A P M
RANDOM_5311 L G L Q A S I H N L F I I R Y P A R K R N V F S F D A V C E N R S T L D S E L A G D P H K I C V V N T N S A R K
RANDOM_5312 N F E R I L E N Q L Y A T I Y L L D N A G F I T T K L V K Y E F P T A V V C Q V A G T I M P F Y L I F S R D G E
RANDOM_5313 E T D A Y S P L A V V G T L I R T N V I G A A E P Y E V G N A N V R Y P C V C M A K G Q E Q F P G A I M N C I V
RANDOM_5314 K S E T I I T I H I L T D V G T E F N D D E V Y V T T G G L G A T I V T C R G G Q G L D P L I I D A P S P A L A
RANDOM_5315 R F S P K S R Y I S Q A N Q V S A G R G P N R K G L T K N K K H T S W N F V H R L R T W G Y V Q I T I I D E T D
RANDOM_5316 V K I E E V M H S E C R S G A F L S Q L L A T P D D G S G M D D V S I Y A F V E H K A A Q L R R R S C L T M K K
RANDOM_5317 A G I R E N I A G N I L E L E K K N N N D G L A L S A P P T K G L L Y K K A S T P W I I R G P K V S S P G L T R
RANDOM_5318 T E I T D K H F Y L L G D E R I D D C Y T L R I D W K E Q V N N Q A I F V T I T D K K L D V P N E Q Q T E T V N
RANDOM_5319 P Q F A T F D M F E Y G F S G K G K A R L K T M D I A Y Y D D Q H R Y L I S A T V A A G F N N I V N W R Y K L I
RANDOM_5320 G A V H G A A Q D D K K H V K V I S C R S E L P R S E K T G S M N V P L A S I Q P D A A G A I K V N D A Q M E P
RANDOM_5321 Y F T S A Q F R E L D R N E I H Y G E L L Q N E N D N Q P N I L K V W Q L Q I K T E K N R L T N M F G N S S D K
RANDOM_5322 K S A S V P R E L R F G R T N S A K T G E P A G C A E T P K G N R V G Y P H S V N I N E F S G T D E L I L L D V
RANDOM_5323 G K P S E E K E E T V L Q Q E Q C T C G G F E K A P L A L L L Y S Y R L D Y Y L G E E Y N D Y S H L L I Q G R Q
RANDOM_5324 Q L R D V P D T Y L E L F A S P Q S K A L S D E T Q K T Q R L L A Y Q Y K G Y F C D R D W N I S N Y S D G I V L
RANDOM_5325 I T E G A R A F S V M G S A A R G R S Q L S L R K Q A P L I I G D A G I D Q L R Q K I Y Q E D T I P S L P V R A
RANDOM_5326 T V S G G Y R T E G S F E S H L T Q Q L D A Q N L E R K H N H D K E L E T L M E D L C P L R G V G T G E E G R G
RANDOM_5327 R C F L E T S K Q L L E Q G K F Y L L G L A F G I G A T L T L L R I R P L N R E N T T V P V G I C Y T G T S E D
RANDOM_5328 H F E Y E R P R T P G D C A G L R A E K T E G A A E D N A L P A A S E V K Y K L S E L E I D D L R D I L V E E R
RANDOM_5329 Y M E S Q E A P G L N Q K S G F K V A C E G P G G A S F R D L N G G A K F A N H G V T T G K T Q I N Y D L A C V
RANDOM_5330 V E A G E I H P Q L A L L L E I E C S L E E S S I A L L Y L Q S K T K T E G D T V D A S M T W A A M S D R W L G
RANDOM_5331 A E R H L V S T S S Y S D Q D G H R D P L Q L T F L T G P E V A L K G G C A D L K R Q F P L G M I V A P E R Q V
RANDOM_5332 C V R D N F R I A Y F V I N E K L G V R E F S F L S D M A G E H F T I Y S L L A P S I Y R Q V V S V P V E R L A
RANDOM_5333 A M K I Q L F D K S D D S R T V E G T S A N G L D V A G V L I A S P W Q D G V K G F E D R I A T N L P V L P V K
RANDOM_5334 S T K Q V D T F S E A Y A V V Q S G V F T G R A K Q D T N S P P R V A Q I T F S W T W S D D M L E N I P F Q N E
RANDOM_5335 L L T L D D R M W T K I E R G N G E E E S G G F I S G Q V W E T A R N Y I P T G T E P E P R T E S K S V L I R Q
RANDOM_5336 P A R I V F L E G P T V F K A P V D A R T A S S A P E Y C R R Y Y L S S S R G S L P S A I G G H I F R W A L T P
RANDOM_5337 S A S K V D R P S P S Y L R D K P N L N K N Y A T D V T D G W G R L A N D G Q V R R L N D P P G A S A D S S L L
RANDOM_5338 D T I V N R L D K I V A I Q T N G S F P S S D I N Q A K M S A I T M D S F I F H V T K P G E C G S S F V R H I H
RANDOM_5339 A P V V K N R H S D F E G G I H K L E Y P D A M E E N A K F S E Y P R D L R E F K T S P L T I G V E E I N R R N
RANDOM_5340 T R I L M N H S Q V F E L S V T G Q I I P L P A T W A V E P Y V W H A I T K K D E R R I P S S K H T L R T D Y L
RANDOM_5341 S K T E E Y L R T Y Q L I G A N C H L M R R L Q G K T N L L V G Q N S S L Y K A S Q V F G V W D H T E I A A A P
RANDOM_5342 H S D T N V L I A A V E L G L R L A S F N A A D Q G F D I K M L R L S V Q L S T R S H K F L E Q F T L F G T D A
RANDOM_5343 D T C I V P E H R P G Y F D P S S M V L P S M E R V G I P N L L V S A T A S K T H L L R D E Y Y L L E I V M A A
RANDOM_5344 E E H Q L P P V F R V D G A S E E A V V E C I G D S T K Q L L E L N T Y R R P A S C C E L A E L H V M G S M M T
RANDOM_5345 F T M G S P Q R S G K S E L K L C C C C G Q H P E N K T N G A I G S R L F M L G D A C N G A N Y I I K P A D S W
RANDOM_5346 S D S A Q Q A Y F R P V F E D P F G R V S Y P A Q D M Y E G R P G S S T R R R P A G E A N N A M C G I A G R V R
RANDOM_5347 H R I E R E I K K N R S A A H T V P V V S A D S K Y R H F Q R L N K K Q N Q A P V P T L G E G T G V V L L S Y G
RANDOM_5348 K Q N S T P A E V S Y W F D I S W D L E L T A K G A Q M N L Y L V A H T Q A Q K L Y N P P L E E K K L G Q V A L
RANDOM_5349 S E V S H C R V F T I S S Y K S E Y N A R G G Q D K I V Y G L M T K K A V F T G A R S N G E F G S N K G G V V V
RANDOM_5350 K A S N I I L K G G D F V T H G Y S L S R A D D S R L V L L A T E N L N K L Y K D P L K A D R S L A V S L K R Q
RANDOM_5351 T G V P T P S T V S T A E K M E K V D E R K K K V A G G Y Q E L E H G G V L A R F N I K K S D T D Y A K Y A I E
RANDOM_5352 N R A I D W D I R I F G T E D A D K V P S A K N I A Y S E A Q T V Y S K V I I S I L E Q A A M K L L D A S A L A
RANDOM_5353 Q M D L I W E T C M L S N L C G D E K G N A R A Y Y E E F L G A G E S L S V D Q G L N H S K Q G R V G F I I T D
RANDOM_5354 Y E F S K K W D V A K F G S D F K A G D D D E W L D E E Q N D R S G N L F T I S Y A S V Q L G F A T C S Q H A L
RANDOM_5355 S F S K T T L S P Q S Q I N A I F F G N K S Q Q V L E K N N H G P G M Q T T P C V I G R D Q S I S D D H G I E L
RANDOM_5356 W C I D A I T V E Q A E T T R K F E A P D N I N I Q C E K Q F A D S V E A A V I S G V C S V P G P R F G V L N E
RANDOM_5357 R L S L T D L P A K E A K R K E V D G R T S M C A K G K Y P G E G A E R I A C H W P C A T Q K V Y V Y S M A E E
RANDOM_5358 G S T K P E W T N T A E E A V R Y Y G D P F V E W V Q L L F F F N A F V D G C S E E R I R D L R A H C V L G Q N
RANDOM_5359 L L C C S T V L Y G S E E D V R L I A V V A L D K I K S S W P Y M K L D I A G E G E E N L L D I T F T Y E L G V
RANDOM_5360 K A Q N Y S R A I L D N L G N A V W I Q I E R V M A L K D D G E N S E L M E V R V E E A R T I E V E E K Q N W F
RANDOM_5361 V V Y S G L L R A L N M R H P A Y T K Q F V L S E S G F A S Q S V E M A V V V I E Y G N W Y D H A I D Y K K N S
RANDOM_5362 A P S H V G Q E K S G G N A Q Y T K T I Q G A L A G Q A G T Y S E M V G T Y V S Q E P S K E R E S R R I V F V K
RANDOM_5363 K T L C R S E V S N W M Y D G E A M P F P D F P Y N N Q R D H P K S L D A L S Y Y P Q L Y S L T G D G M A D N G
RANDOM_5364 H E Q N F S S P D E A N V L K I Q T L Y H S S H R I S S G V L L R P K G A I Q Y R Q K A N P P F L P H N S C C F
RANDOM_5365 T A R P A K Q E I T H K Q L S Y N I G V L G V N E P R S V G R L N D C L A N Q K R L T H I E R S K P L Q L I W V
RANDOM_5366 G C S E L R Q G T M G K V E T S L G V P K V S E C W V R R N G V R L K L W L C R I Y D F T K L L L N K G A F Y S
RANDOM_5367 K N S A L T R Y G A A N W P C V Q M P Q Q S V A V I A I L A C M V M R R V Y A P L G K K P H D N G E A V L E A G
RANDOM_5368 Q L V Q A S Y S I G H I R F G R I P Y V M H T E R E Q P L H S L T R N L E G T K G L P G G V L L R A G S W G R G
RANDOM_5369 V G F T A Y S N R N A C K S D C C E D G Q E A E S M T M P K F I G G K E N T R S L Y T E S S N G E Q I Y E L K T
RANDOM_5370 I S S L S V P A I D D A K E Q S A R E Q A S K Q M L P Q E L G H F H F L I Q F T S F Q F K D L V W I E L P A V N
RANDOM_5371 L H T C S L D Q G S Q T E T L A Y A P I S S A L T Q L L Y P W I A A T N Y I V S D P H L V D V V E G H H G A N P
RANDOM_5372 A S E T R K P T V E E G R S W D P R Y Q K D V N L S E I Q H D Q Q S G K N L R T R L G G L L A I D F S Q R T T I
RANDOM_5373 N L N K S L F G F L F G T E C L Q H E G E A K P N T K T G I I S Q C I L Q S S D H I A E V M S D R H T T K L D P
RANDOM_5374 P I G T A G F K D T E V F A E K H T A L Y Q V A S H W H A T D E T W N R T Y G T T K R V L A C N E A S D F D S L
RANDOM_5375 R M P V A Y V G S D C R A N A E K I K Q K R F D K F I S V S S A G P T D G H A G V K C D V A Q T E S L R C V I R
RANDOM_5376 L Y E L P V T C W L K D E G N Q Y G C R I G S P R N G L D A T G S T P L S Y T E D M L L Y R N C R E D G T P A P
RANDOM_5377 L S A D C D I V P N I I L V D K V E F K H D Q R E I I L G H K A R A L N A L W K Q P L C C H N A V S V K V H D A
RANDOM_5378 I I T Y T F E T K L Q I E L N S R E G C I P D L A E C V L E P A G M E I H A D M S N D F N P S G T W L G M K D S
RANDOM_5379 I Y Q M Q V F E S T G N N A L G G P K E A A K Q L R E Q I D E R Q K L F V Y R N P Y E S N E F L N N I A F S E A
RANDOM_5380 N V V T L A W G R T T G D I T T S G A F E Y I T R R S V V T I V F G I V R L N D G S G G Q I L N E L L I G L K D
RANDOM_5381 E C V M E D I V E N A D P S S G A T N N K D P S D G Q R G I R W A L Y A I I G A L M K A F Q T S E E E E E H K R
RANDOM_5382 K K P V V I D G H Y S A T L E E S L A R Y P I W T T S K D T T V R Y A E L L N Q G S V G A R L K A N S I S E E H
RANDOM_5383 L G S K I K G Y F R L G S G L E Q L K L R V M I L D I W S D L E H C G W E N E I D V K H F L K V P D P L S G E K
RANDOM_5384 A Q Q D S G H Y T I I S S R K A M D Q S N L R K P R P G L S D L D I A G F L K T I D L L D Q M V G N E P E F D S
RANDOM_5385 L P S D G D E L E P S L V C N Q L N G H T K S F F V F L R L P M L E S Q V G R V V G V L G R E I I M W V I C Q Y
RANDOM_5386 S S D I F M L L G W Y R A N S G K T S A G P L H P W G S G T V N S D R L H I F S K R R P V T L P A G G I R A A R
RANDOM_5387 Y L P L L H N I T E Y D G L C Y G R S G L A L K G I E S E N L Q A R K D I I P Y K H D V L Y Y R W G L E E V V D
RANDOM_5388 A D A A P N R W F D I E V R M T D L I V G Y A K G T A A G A V A E G A R L Q V L G L L W Q C P S K L R D K M G Q
RANDOM_5389 E E E Q S I F L R D S G R F Q Y D R L E R R G P P N I L T C S S A G A V E N L L K D N Y L G G E H A C T D K P L
RANDOM_5390 L N V T M E A A G W L C M E E D A G Y F G R G D A E K G Q L F V K D K A D G S Y G M V F K S N L R K E L T T Y R
RANDOM_5391 G P G D I G Q D K K E S G T D N D C P A L E S Q R L I S R H P A L S A H S N L P V R E W K K V K D I E P N K N L
RANDOM_5392 E I S G L E A M V L Q E V R D L E K A D A R K E D I E W N I D P S C T N V L A R V D I K G K A E I I S A P I G I
RANDOM_5393 K K D S M L R A G A D P G P E T R L M M A I R N K Q G T T K T T T H V L D L I Y E P D A K Y G V L N Q N C V V I
RANDOM_5394 T T D T L L G S E V N R A Q V F A N A C L A I C I D S P S D D G L D F A L N C T L P S F Q L V M E W A M H R E Y
RANDOM_5395 I L I P S D I F T E N V G P T L R F G D T L K C H V P F T I G A L E N M G G G F Q C A L R A Y P R F I D G V V P
RANDOM_5396 R L T S S L S H F P V C E C A E T T A K Q H V I V D H D G V S S L R S T A S N Q I H T L E L S T M L T D D L P A
RANDOM_5397 Q C G S C L T Y L G G C C D V D P R N T G T K D L P S K V G P I S A A I I T I G P P I S C R P V P V E F T K F A
RANDOM_5398 L F R L N N D A V E E S L P Q P I E I D A S Y V S E P Y S G L L F E Q V Q W D A W I L F S H F G S C E L V A K R
RANDOM_5399 R L V H N L L G A G D V N E V D G N F E M P V Q Q Q Q K T E P F V Q E F I V G I K A V L E Q A E E I A D A G C M
RANDOM_5400 Q N I Y G S A M A M I L A L I T P S T F L T L I Y V P I C L I I G P T I Y M L A K K G S W F A V L E N G L G Y K

RANDOM_5401 V R K L C F P E V E R G I T D K N L S P L T A I V A M V I C T K K W R P F L L H K A R T V R K T S A D W Y L M Q
RANDOM_5402 L V F T T H G K L T L G Y A H T S A R R T V G I T T H K R M F A E E W A R H L H L F P T S F D R E K F R A G G Y
RANDOM_5403 Q I L Q K F S K V A I G A S G N P K E A L D L R M Y S A P L H Q Q K Q C P F K S N D N K E T P Y P S Q P T G R I
RANDOM_5404 V F D A N K F D A T G L L K L R L I S R K P K D N F T V E A T T L A M S A C A M M T V Y K L E Q P E V Q S S M A
RANDOM_5405 L G I K S Q Q S M S P I Y A P M I Q S L E Y N V L T A V L P R D F K A G D K R D L E T E R H R G E K A S R P G G
RANDOM_5406 E A D Q T N G N R A L P P Q T R R E V N W L T E E K R G F L N S G S S Y E P G F I S P S T N T G E F F T D K L E
RANDOM_5407 I K G V V F S H G M R Q L E I S Y L A A W E L T A S R E P C V A A P R T N N A L G V G M D A T V V E P K L T Y Y
RANDOM_5408 N T G Q V E D V P D W Y K A V A A P T K V R A T V L I I E G I L F C E S S V F G M D A P V A K V R L P E K C N D
RANDOM_5409 T P G F P N S L D S Q M A R F S E K L S L R K L S F L K E Y I D I K D R T V G N F L E K K A E N N K G P S T I D
RANDOM_5410 K V C R E A Q Q G P T L L K M Q H I V D Q S P T P D E L D H Q Y L R T Y P P S K V K T A D P K T P I V S S R N T
RANDOM_5411 P P C I N P A L K D E Y V N V Q A R L I L L F G T A H G T G I C V A A G R Y N L S K Y V A T L H A P V T T T V A
RANDOM_5412 S G L I I S L K F S L N I F W I P R T W K C I N G D P L S L V A R E C E F D R D L I I I Q A S L L N Q N A F Q A
RANDOM_5413 N N A T V T I L D I L T I F C G E R V I D V D P V V L H V K A H K N V V K R L P L R I R C Y G I L G S I R K I D
RANDOM_5414 S D Q T D S E G I L Y S A W F D R D F E N I G Y P A G D K M P R P T V D K V I I F F Y Q A K L S Q T N N I R W H
RANDOM_5415 R N G G S C R G T Q P P F D R E A T T S S A A Y F C K E F D G T M P Q P L L P T K L Q P S G H W V I I S N N L S
RANDOM_5416 N T N E Q V S R G V L R G G S M A Q A R M Q S D K Q C F S L P A G E R G L P A V P G S T E E L A V K N Q S E E P
RANDOM_5417 G L E T N Q S L W G D I Q R K N T L A Q L A V P A E L P S R S Q A C L G P Q Q R N R Y G S L K F P N T R W E D L
RANDOM_5418 V I K V V E K I D I A W E S T P T D A S T G T T L S V D E A K N E V G T P L L D P V G A E Y P P H Q E G Q R M S
RANDOM_5419 T F E R I Q S G D T K N S V N L A I T I N M M N M R M V L P N S H P K E A A R A C A N Q F E G K T W L F D V K R
RANDOM_5420 S T K Y R L D M A I C S A N A S T H I R V I K G G T L G E G K H A D D E S A P Y V G F G R V L A T H R K K D E L
RANDOM_5421 N E N Q S A S L T D Q S R N V Q L S A E R E T R G P D S A P E Y E L L L R G G E E K L Y I A V V Q T V T K E L R
RANDOM_5422 P E V S T G S H E I R C S G Q G K H T E C F E Q D L L N S T G D M V P I Q Q V F T E T T E K S M G E L K D V K S
RANDOM_5423 K A L F P G T V I T C K S Q S C V G E F G S K R N I L N V K D F N E S P Y L L I V V R Q L A M Q G Q E S P G S S
RANDOM_5424 T T S N P I R T E N D T F Q R A K I A D A P N G D G L K F N P G M T I Q D T Y L F T K Q H T R H L P H A G I L V
RANDOM_5425 S A E I Q Y A S V T P H L G L A Y C F R N T T I N F V P F S S D Q T L D P K C I R Y K N G L W P P I N A E D E D
RANDOM_5426 P D R N Q W K D S V T Y H E V D R D G P E S G Q N S W A H T N R E L A I R S L M S N L I I K W C A P D L A P F L
RANDOM_5427 F G P L G F G W R L P E A G I R V H R R D L D P K K T T K V K S V K A L C V V V N L Q Y Q Q S V Y K T V P E Q A
RANDOM_5428 P E K S Y H R I N K E R Q G F N Q S K A T F Y S L G E D D V A T E V L I T N F N S V T T R Y F I P H S L L K V K
RANDOM_5429 A L K R F F L C L T K P L T A L A C F M L K D P F A K P E L L T V A G T K G G F C Y I I S S V E R G N L S V I K
RANDOM_5430 L K I A P L L F R D E T K T Y F V V T Y T R P D E K A I K P N Y Q A M H V H F E D E F W K M K I A L D T K Q D C
RANDOM_5431 T T I E R V L K R A F A Y I G N M M S L V C A H N I P D Q K D F L V S G Q V D Q F V Y G L N V C V S L S G L F L
RANDOM_5432 L R S A L W Q E R L L R R K M T S A V A F T N N A R T G D Q G V L V G P A D A N V I L A K A R N V M G F Y F K R
RANDOM_5433 L L T L T R D I Q W V N Q G A N H F A E C Q N A L S L Q I K I Q Q S K V V F P L M C Y Q I T A V A V V R A R T V
RANDOM_5434 E K G A R L A Q K T N L R H A F L L A T Q K T P Q R V L D V L P N D W Q K D G L D F S T V L S N P S L Y V P T A
RANDOM_5435 S E V E N E D P V I V E V A L P Q T K D A R D A W V T K A F D P G D D V K R V S D D S E K W T W V P A W I C G E
RANDOM_5436 L A S K Y V G L W A L M E G S H R G A I V G L A S I W E E D R S K D V Y Y G F Y L W I K I I M V A L E A S E L Y
RANDOM_5437 H V L G K C Q E E S K E L T I L Y E R S Y L E V G I I E G A N L Q E L V A E F S I V S L S D Y S T I R L H T A S
RANDOM_5438 L Q I Q D T E G F N I A G Q S S I I S F L L V V T G P C V L A T R H Y T Q F I L A A G E V K C L C L V V D E L R
RANDOM_5439 I R E L E I S T V T L Q I S D V G I S P Q F E L T L L S D P K M R D Q E A S E R L P A K Y T A E W E P P Y N L A
RANDOM_5440 V K S C E L Q I T E V A A Q P A R K K N K N I G A P N E W C K G L L H L V E G A E D L I C I T T K Y H G Y N V D
RANDOM_5441 N T K L R N T Q G S D H Q A G F A F M V G L L R S M K D C P Q V A V I W V E S L A L K A H K K T S S S A Q T A M
RANDOM_5442 E L P E D P K V K T K V R S I I P E V L I F P L I R L M K I S P P C G E E A S P T L G C K I L A A D L S D Q K G
RANDOM_5443 T Y G D Q V V G L S V A M I L E E G L V T N E Q V I G L P G A G R T K G T H A G K S G F G G M L P D D I Q L L I
RANDOM_5444 A T I P L R L S K N A L A E T E I N V G H A L D P N A Q E A Y V L I E V S L L Q L C V E L R K E V W S T L R M P
RANDOM_5445 F S L A R S S D G M E P S G A N S R A A A A A Y D L I C R C V Q C A G N S G D G I S K D E F I R A L N T T V F L
RANDOM_5446 M T W T K P R G N N M G Q L I R K V Y H D V G P P K A H I R N K Q R L T S Y Y T D G A L C D V F E D V I S V R I
RANDOM_5447 M N T V P R A H A D G Y T V T S K V L T V R K A V V S F F R R N W R K I M P F S F E L A C T C L L K A G S A F N
RANDOM_5448 G E C E Y P N I M D W A A D G P L T I K F E R A G G Y E Y R T T E G L F D E S G Q G F Q L A D D M T E V D T W A
RANDOM_5449 V P E L A T S I T N G R K T V I P A G V M K A G A A S Y K V T S I R A G Q E R L K C M V Q S E I V A M S R R H N
RANDOM_5450 W I A G G S I T L I N A I V E F F L I R A C G K V T L I G G A K L G V S Q L A R R S N Q K L R K P N M S P N E Q
RANDOM_5451 F T K A A K A V G A V N R M F D S S E N L Y R C N N C S M S T G D E F F I T A L N E Y K F E E V C V N R N P T V
RANDOM_5452 P A C G P M L S H N K K G L L A T L S Q N S L S S L A L F S P N E G T V I G E K H C W G E A A S V K K G S R V G
RANDOM_5453 Y S P P T Y F S R E L V K S S E G M K W K L L M I V E I E S S V T P R R M A A I Q H L G T L S E K T I L G G K H
RANDOM_5454 K D Q D P V E V T L P T S D A Q K C F N P L P V W R M S G A T K A V I R P S N D H T E C E T P D C T G S K Y Q D
RANDOM_5455 P K T M H H N T N G H I K Q N G L H A S A E I T C G K H C L E A S E V T A L T N E W T V W R G L E G L P K A S L
RANDOM_5456 P C I D V E T Q S A H S L N D A G S Y A T D D A I D K P T N Q G I L V A Q E G V V Q G L E R K T G V V A D C G K
RANDOM_5457 K D G G R G D Y E Q T V F I E V I I N V L K R P I Q T L V T D T F Y F Q A R F S S D M L A W I F Q S Q T A I G M
RANDOM_5458 K L L D K R H A K V I G V G L L Q V S E F M V Q R V K Q L L H S P Q Y L A A R F G R D S A N P M K A F E V A C F
RANDOM_5459 K D D A S V R T S Y V L W P P R R T W N S L A A E G K I G I S M N C G E G C G A T E Q S N K Q H S A K T R L H I
RANDOM_5460 G S Q L A V I R G N E L K V H V F A V T S M N L G L R F E P H N L L Y C R D L N A N E I P T Q D N F C P V G V I
RANDOM_5461 V G D A L S E T P G A Q F N Y P T L P K R T G R R F A A K T A E T E P A E I C K A V R F L T R A E D C N S F G N
RANDOM_5462 R H V A V G G Q G H N M N V Q N H K P L C H C M D R M I S S P H G H P I K S D V A I R V K I Y H V M Y R S W D K
RANDOM_5463 E N M A N K N V N H T G T K T T I A Q R R S Q T Q T V E Q I A K R A L V G S Q L Q S W T F T L E I C W S M M C G
RANDOM_5464 N P R Y P S G R E A G T S M F L L N S V S S L S Q D E P C K E R W Q Q A K T V I E G W G T E P P A T E L F S Q E
RANDOM_5465 V V L P L T D S C G Y S F Y P L A M M D G F T P I E M F G N A K Q V N V S S N F M V I L S Y H T Y R L S F T P T
RANDOM_5466 N I E P G I E T R Q D E L T L Y G M L A N S P I L N V G L S G A P S S R L N K S Y D A A F A L Q D Q S G E G K R
RANDOM_5467 F L V M I G F P N G T S S N T T T F E T L S S V T T D A A E L T I L E D C F S G E A S W E G A G I G Y I R A G N
RANDOM_5468 A V R E E L T S I A T S F Q Y D S Q L S E L A T Y H S V P F L L L G S Q K H R P V S A L E G P K S T L E A S D L
RANDOM_5469 H K D V V L I Q F A V V V M Q G E K R G E D Y I Q S G E Q E V S E A D R T T G I V A L H E L D N T G V L C S D K
RANDOM_5470 R Q I Y G T A L Y G H L L Y P Q S V E G T P A G P Q S I P L N A I S G I D L Q K P L E L S S N E N R F D I G V T
RANDOM_5471 S V Y S R E L P A A S G N A S K A E L G K R E P A A I Y P F N I T I S G T D H E G L F E N R T D K I C P G I N L
RANDOM_5472 S T P S R R K N S V G I F T T S S Q A H G W S A G E D T A V R M L D Y E R R M T K S I D E K V A A S S N R S A M
RANDOM_5473 A Q P Y A T H T P M L E A R E I S P Y K A G M K E D D Y A L P V S N V V S G G V K G F I Y D P I G N S W L T E S
RANDOM_5474 L G S N D L T V V E K Y L D S F K L R K D P Y M A L P Y E P E M S G F F S Q R Q I I A Q E Q R I A S S Y D T N S
RANDOM_5475 S K F K L C V D P V Q V G N E N E K T P P S H G M Y L I Q V L E A R I V P D D K M P G M Y K Y F A S R T G F L P
RANDOM_5476 S T H S V A F S S I K E Y N E Q G P S G S K A V L D Q E G K A I F Y L F D R I V S D Y V F R L T I F I L F R S F
RANDOM_5477 E E I N T D P G Q L S D N R V T F P S E S I W L M M S V D P N D G T L L L L V F N P L F G T S E N T E F V C N N
RANDOM_5478 V G E H T S L V D G F T V N K A N L T T S A R A D S E N I I A C L G C L P T N L D G F N I G W I Q C E K S A T E
RANDOM_5479 D A I L P T S C M P D D V I K G C V K S N T M T P E A E K N T A P H Q K D F G K Y F D W P N E K C N A F E C N N
RANDOM_5480 Y A M A E T V V L Q E N E V I L G R N D Q K S K S I N F P C F L Q L Q S M A C S L V A L L A D E L C E R K L S A
RANDOM_5481 S V P A G D Y L Y Q E D Y K A A R A D Q P S P Q V S S N H V I I V G D T P T A C R D G N I L P G E A M A V T F C
RANDOM_5482 F P T E I A G L L Q R I G K S L S Q P M E F A V Q E K F Q Y I R A I G A M P D G F F G F D T S M Q D E E D L Q I
RANDOM_5483 S L L E C A W G G L L R E I V E K K V L A K L I Y I L A L D V D G R A E Y V R E F P H T E S D A V V L P S N S A
RANDOM_5484 E K L M Q G I A V G F C E Q Y S F R G V S T A Q S K E L M K Y L V G S V G E K N E H D M A I N G P G V Y A V L D
RANDOM_5485 A W G V V L A S Q A T K M L Y G A A I A L W G N E Y A A M D A A S R Y A N D S P S L I D Q P C K L F S K I G G E
RANDOM_5486 E G I A D I I Q A G I S G G N E E D V G E H V Y G R V I W V P T P Q K K L S F N Q G Y C K V K Q V F H R Y A R T
RANDOM_5487 V N F I P T C N K G T S T T I P A R C Y V R A F F R E K I A K A H L L G P K C I L Q K D R N Q C V I V D P E A Y
RANDOM_5488 P K R K S R A V A L V G P D D K L D D M Q Q E L D I Y K T P G R Y N T S P V N H I G A C G D N L L F V T L V K T
RANDOM_5489 R N L M A D Y Y W A T R F N K Q K V T F N V N Q G L T W H N K L T K Y L V E A K D V R T G L L V H S I D N E D D
RANDOM_5490 Q R S D A E C A I A V V L W L R I E K K A A N F I K P P G V G G G S G K L D L C S V T D L Q S S W V D K C F I T
RANDOM_5491 N V G Y A Y E I R V D M S M A V L N D P A H S N I S G G D Y H L M K Q P G G H I E Y F L T N S P D S L E Q A L R
RANDOM_5492 L G N R H K Y N I V E V D Q D G N L L Y G Q K A Y Y S P W S Q L E G H R V S D M I L E I E K A D L Q Y A P F R L
RANDOM_5493 V V E A A T L R T K Q S S Q G W K A Y M N G D S Q A N Q L P Y S Q A N A L P E A Q R S T T K K Q E E A F R L I R
RANDOM_5494 S V L P S G Y T T G T V L T V K K Q E R G Y S I E L A K F L T G Q Q V D K V L G R K E F S R M E F K L K I N V G
RANDOM_5495 E A A P L F T V L K A F D H C M T Y M V E G V Q W T T L R F F K L S T H R E S Y N L K E E G D E T C K D P S Q L

RANDOM_5496 I F D N P G S S V E K A L Q I L A E T G V F K Q G Q S L N S S R F C D K T T D T G G A K L D R V H I P L G P N P
RANDOM_5497 L T W A C W A A R N S T N S N S V S L I L I H Q P T A H N V I W G H L I H Q N E S A R D L V N I L G D T L P G S
RANDOM_5498 S L S I A V F K P G I C E A T S C G P L G G M E T V K D S A S P Q I H K L D R G P A K R T E K V P A M I P D V F
RANDOM_5499 Q L H S N I D E S T Q R F N H T C N D D Q T L D R D L V G L G I E T F I F K T R G P G L L G F D C N I W L G S L
RANDOM_5500 E N F S E Q T T E T K M D E E K C D D E V C E T E V S I L I E T Q D L D D I L D A H E K V D E I T L T L R E C D
RANDOM_5501 S T F S G I Y P V L T P S T I Q Y L P N D D F V E T T R G P V A V N A E S Q E R D I G E Q S Y T V E W A H Q E R
RANDOM_5502 E T G G Q D L L G T G E D F V P Y H R A L T M D S D F F G E N C A S E Y K V I M S Y F Q M N S L A E L V P V S D
RANDOM_5503 Y V Y L T V S E T T L P A A G E L E M I S S H P T G L G F P D G P N P R L A I Y I D G G S T I N S R K R R S P L
RANDOM_5504 A V T V E S L L L L V P I A D K L E H Q L A F A I N Q F Y L S K N E T A K P S S S R N Q M P S R N A S S G R E S
RANDOM_5505 I E A L A L V A L C L R M K V S E Y S F L A K P A M P W I N L P Q T V A R R C A W L S V E I H M N A K G S K K V
RANDOM_5506 R G R V I G N G R S N P H S W S S D G E C M R I T E G P A V Q E D D L A A L V A N Y F G Q Q R P Q F K R P K F A
RANDOM_5507 G L G P Q R Y R K H Q Q L E D I P P T K V A I L A D A Y L T L L L C E K W A G R I Y N R V N V N S Q K T T G V D
RANDOM_5508 A K M L S S Q M S R L R Q R E L E Y D G Q I E T Q Q E L G G P V G C L M H T D I E D N A S D I I S E G L V A Y E
RANDOM_5509 M A T W V K V A F L G K C V A Q L S L V A S G R V T A R F D I T G G E P F N N A E D L G R N P T H D D R L S R A
RANDOM_5510 L V G Y R L V Y I F N I H L G S Q I D L E A S Y Q D I L R L R N K I W T L G N F I T Y S N N I Q N N N K R G L A
RANDOM_5511 K I K V L A A P S A A V Y T D M K G E I C I D E Y E N P D F P V R I D M F K M D F A T V G N R A T A E N V C K L
RANDOM_5512 T C K L V P A L V F I G S E G S H N V S C H P S Y V P T N L F A L N S E F E G A Q Y R V W V P K R A L S L Q Q K
RANDOM_5513 I D G S A R R G K D V D D T L I H V F A D E T I D K P P T L N I E S K K S D G S N N R I N C R A G I S Q A I R A
RANDOM_5514 K G N V K Y K R R I T K V F G G K N E Q S M L K D D R S G I N V A L P A G L L R Q A Y D S T P Q S A R L S Q S G
RANDOM_5515 T A G L T G Q Y M Q G A E H T S Y H Q V K K Q D P A S L D W H L H I S L K R A P K Y L I S T E I C R S C A V S I
RANDOM_5516 A D Y D T V Y N A V N K F Y K Y E I L L V I S R L Q F S P C I T P P R K M T A E E P D T A H S P F G G S C D D M
RANDOM_5517 H K Q K T L D L Y L L P K Y V P I A T Q C R I I P R R N Q M E T S V G N T G D K V L I I P D T R V S D A D V F Q
RANDOM_5518 H K W M T P I N L L G A T L K R N T K Q S D G M G L S P Y N Y G S E E P V Q P P I V R T L I T D N L P I D T L T
RANDOM_5519 M I R D F L V I Q D P V K N K E K S Q V L T D A I E I S K N C L T N E Q H D N T N E P G P G C V L H F A E S N K
RANDOM_5520 N E R P S E R V A D D N S H R R M K S S N I W V T D E K F D S E S D G S V V H K C G F L V L S S L F L T S G E I
RANDOM_5521 T E R P L L N S F G L V C A I R K M H N T S V R A N S N V R I G V M P D Y G R E P S N K A G I N S H D I A I E T
RANDOM_5522 S V A R L W G E K W L K D Y G R T E D Y D S R I I R G G A L T A N M G E G H Q V Q A S Y R N T D V H S M G E A Q
RANDOM_5523 Q A G S N T L E V F M I L K S V N G F R H L G N K V F G S Y I C E T D P R D T L W E L Y P F L E D V P S A D R T
RANDOM_5524 D L E M L G R Q K H L R N P S S T F E Y T L E V D F P P E E K D K V N G G Y V R E C V Y W T Y V C E K G K E I I
RANDOM_5525 A N Q K K R G P Q V Q S Y P N G D T N Q K Q E K G L Q F L D P I V V P L S Q K F T A N S G A P D S E G I M K Y K
RANDOM_5526 Q E Y L V G S S K M D Y Y V A L S Y R S L G V H M K E F G S A P M A D P R F R V V A G G P T N T C V R E K W G R
RANDOM_5527 E A T A R T D R C K R Q N H L I R V S P W A F T Y N Q R T G L L G K E G D E P L R E F L R E P A F V L V D G I Q
RANDOM_5528 N I N L S L D S L E N Q H R V T P E Y C R S Y I Y V Q G P P T D R K L V L K G K K T I C A G R Y E D T T F G I P
RANDOM_5529 L R N A Q K A L V A K R V P H Q G V G G V L F S I A G E V G N S Y G V I G R L T I I Y N M G R D C A T E L V Y L
RANDOM_5530 T Y M N L A G C T Y I C V D D A D D C Y F C E N P D I Q W E F T L P R W K A V T G F P L Q G R C H L I D G Q A R
RANDOM_5531 L E L T K E Q G T L A V N P H V F V E A L Y T E I S K K L Y K S G V A K G D D F S S L P T K L S Y Q P A T W W D
RANDOM_5532 R F L D K F V V T A L F F S Y M N T A L L T E K G G Y C M D P S N V A K L Y Y S P V F H S E Y R A D T K N F Y S
RANDOM_5533 K E L K E V D S H V E L V H T K T E G N Q F A M R I I T L D S L L L A Y N G G S S F T K S K T H E V R R T I I S
RANDOM_5534 D I G G G I D T V Y L N R G I H G Y M L V Q T C G L N G T W S E L V H L T D C F D P E L N M K D D K A I V G I G
RANDOM_5535 I E G D E M S I Y P K R V L K R S A S T G V K E G A T A A I K G S D T S T L K Q G V A F D S Y I Q V T C E G A R
RANDOM_5536 R R G T V G N V I V L N S I G C S N P W D T I D A I H E D I L L H K G M R G G L S V V Q T C D S E V L K P T N T
RANDOM_5537 S L D T V G P V S H P D A S K F H N A W E L C I L G E T V L N K S K L L L G S L V V G D E S V I H A L F L I V V
RANDOM_5538 T V K L E V N E C V V E L D T K I Q F L K L L H L E E V V V L A V K P G V Y A Q W Q D L T A N Y T N E K A D I V
RANDOM_5539 L I Y Q D H G G V C S N L I R F V L P G V H G L M Q S C H P Q N F Q V L P C D C E D V I I L F Y L R Y K Q P N T
RANDOM_5540 E D S R L M Q Y W F L L S C W C L G R L T C A I A G C A P V E S V F L Q E L A G T F V M D P V E E R E D P W S K
RANDOM_5541 H E A T I A K L C G A C E M T R H T S P P R H D N A P A E F P D D N K N E E L D L T Y P S A L Q V L G E E K Q P
RANDOM_5542 T I Y V D H A G V Q C N H G T N G V K T A A R T E V T A P G S Q R E A A L L E N L V P T E S F S C M D V L G N H
RANDOM_5543 L Y E V D V P A F L L L N Y A V T R G L S P K L G G N A K L M F K G E S K N A K N F C R F G N R A Y Q I R N L L
RANDOM_5544 S S I A K Q D E L L A Q V A T L I P Y E A V C I L I Q L A A E A V G N E A D Y L F R V C S F R S E T L E V S D G
RANDOM_5545 E T Q P F G K Y G D V A L A V H D V E A N A V K A Q S A T S P G P T R S W I L S L T A M E R A R I P I E P I R D
RANDOM_5546 T E R I L P I A T R L L L R S G T K G Y I I H E L Y V S Y S A V Q V M I L G Q P G M A E A G W R M L V L V K L L
RANDOM_5547 V V T V L D Q N A A I D S K M Y S R Y K E S E I S I E P L G E C Y Q H T D Y Y D F L K G P R L E Q I L T V I K Q
RANDOM_5548 D V P L S V K E T S L V I R R K N H L Y A E M R I G G S W C L R N S A P L C K K T N F G E P N P L K L G I L T A
RANDOM_5549 S K T V L P K K I W Q L I K L K R Q S Q L Q S I A L V I M R N K N E T A K S Y K G Q L L V K Y I A F R D K Q V T
RANDOM_5550 N R R L I T S N K K Q P S N K C K V Q T F I E G A T Q T V T Q E K V R A S A M V I N R K F N F H G E Q T C M A S
RANDOM_5551 V G E A Q K G Y G A Q N L L A K S K Q S A R D E P T S I A V F S F T E G W S P R Q I L I A N L R P L I T S S H M
RANDOM_5552 S P L G A V G P D I G S R H D V G A E T E N G D P G G E S R K M K G T S S E V G Q L I N V R C Q L S T R T I T R
RANDOM_5553 V R K D T V P T R G F R S R L E K D P T L V L K N S S L K V G A R T G C A L V K Y D A Q G T L P V E A R N L I L
RANDOM_5554 S Q I K M N G Q A M P T D Y H L E G P T K W R L E A S V Q S S G D T C S C M Q K K A A Y W L F V M T R I L K L K
RANDOM_5555 G G L L E S A P G E G N T I R T A N P N K R E N R Y A I V I G Q S G T T H A Y E G H P L V N E H C V E N T F E N
RANDOM_5556 G D I D I S W D E H K E A I A S M S L N S G C T G R V Q D T E E W E C D L I A N S P G D M D N R A W G V P E L E
RANDOM_5557 G S R T Q V E W I R A P L V Y I P I G R Q D R L S F H N M M G P E G H E L E G D P A P M E D C Y R E P L N F A Y
RANDOM_5558 L L D F D P S Q L N C F L R D T V R D R N D K E V P Q E N M Q W I V L N I I S S S D R C N T A G E I I L S K P V
RANDOM_5559 S G R R E V A V Y A A D G V I E F D C N T D S V F A Y V L S G Y P L A G P L M V N L G M G N E P M A L C D I L E
RANDOM_5560 I T A I L S S E T R H A T A L Q R K E V D Q R T T E G G I Y V M V L N L E L D K C C A T L F I V R P A G A E E S
RANDOM_5561 N C N T P G V T E V Y E I I A R S A L V V G Y A D V E P S A G A I K Q F P I N E T K F E I L K V M K K R E G I Q
RANDOM_5562 F K L I E C K R Y I I Y R L I E E V R I R N F E D F G S I K Y R I A A T V S I R K P I N L H K F I V W K N W G Q
RANDOM_5563 V L A Q S I I G N F R N A P K G N R N I D S Q P V V S F G S F V G Q S A S L M E A V S H I L G N G E E L L C L A
RANDOM_5564 G N A Y V K L G L F A K E G T R V P V P R A G T L Y R Q I Y S A C A A R L H V K Q Q G T S L N I K G L Q Q L V Q
RANDOM_5565 R G R K V I P F D V P S K H T N I H L I Q V P D H S E L Q Q V A A Q Y F K I P E Y A C F T V I N L F V V T L I D
RANDOM_5566 S A W Q V S R L L W D Q K D V G N A F A N K L L Q L C E S T K S M I M D A D A F V R A D T V L R H N G L A G S L
RANDOM_5567 Q P T C K V G G P A G R V F D W Q N R L W S V V G G T D A K I E A L V K L A Q E V P T M A T C H Q M V Q R L Q W
RANDOM_5568 N Q Y V A V F T E N N A V S L W Y E T R V S V M T Y E S K S D N G C S E A G P E Q T L G S H S V L T G R E Y Q A
RANDOM_5569 S E D R I E T S C D P D V I A C F P L S Q L I K V D A G M L D V T H K H G N M D A K P R R G N K E S C I F T I G
RANDOM_5570 L P H Q A L K L L L Y H W I Q N K K A V A F S K C P M S I W S E H I F D P P A Q N V D K R A D G L G L L V I T W
RANDOM_5571 F A M T R T T N K D I V K A P A N G Y A G K Q N E D G Y Y R E Q Q A S L P F F A K L P K K E P W V A A A S I Q L
RANDOM_5572 G D N K T S E S Q H N V S I C S I T Y D R R F T V G D F E D A T V G G D Y P N E T K E V A T P G R E A E T D N S
RANDOM_5573 A P R E D E E A I G F Q I W L M I P M G E L A A M N T V L T Y P M D T Q N S Q F Y Y A V D G N T N K M G S I P R
RANDOM_5574 V Y T G G Y L A G D R S F S Q R R V S D P V M L Q A D D G M Q K I S I I V N F L N K P N P N R T K V E Q E V V L
RANDOM_5575 E G L V S T K S G E M K P V E F D N I E L P S E L Y S Q S P T N N Q S L V Q F I T P Q K T D S L T F P L V I A N
RANDOM_5576 G I L F S L D M P A M P N L D M E I V I N E L S L K C G F L K I S V I E G Y Q G I D L T N Q I Y A V V A S L L Y
RANDOM_5577 L Q Y A F Q D S D I D A E V T S A A D Q I N N F L S P C T A V D E W S D D K P S A Q I K A K Q V V W F N R V I L
RANDOM_5578 L N S S F S V M Q T K G S G S A L S Y Y S L Q L V C C D Q N W I L L P F V A V S R D P V L M A A D S R C V S R Q
RANDOM_5579 I A K D E S M V D K L F K M L P L S A Q D K K V G F A K Q P F Y S K S D I L R H M Y T E A P D P S V I S T K W V
RANDOM_5580 H A L L F A V T P P T S K F V C L P P H D C P V L V L Q T I S Y D A D P R A S A N G I A H P F H G F D H I C R G
RANDOM_5581 T N F S T A G K N E D G L E Y S M V T R E Q C I S L W A Y S L E V V G G E D Q E S N G G E R N A R E I N E Y R P
RANDOM_5582 P R I A A I G V D M Y K G E E D S L F N G A R A I A P T A T Q Y V Y N R N A N L M E P R Y L M G A R A D K V N W
RANDOM_5583 S G P D G V L G N G H G E S Y E L S Y S P V L I R A S K V A Q R S P A E S G D K K T A R V G D L T S Q V G P G I
RANDOM_5584 Y F I E D L G I T F E E Q Q P D N D Q T A P R I K V Q G N N A K M I D E E L V N S G P K Y V K E N E I A D K F V
RANDOM_5585 S M H P V N L S V L Q S A D R C T A R P D K P S A V G N P D S D L E L G N A A N Y R G T R T K M R D L D T L K M
RANDOM_5586 S E L H S A T L K K K I G A M P F G W T E L N S I P S S Q E K Q L E A A S Y P E K W V V W H F I I T W H P G T S
RANDOM_5587 L I T Y R T H C N N I V S T R R V Y S S L G D G L S P L D S T A R M Y G M Y L I G I M I F R G F D Q A L T L P I
RANDOM_5588 Q G L K I L A R L M L A Q V Q D L L Y E P L R V N P Q V G K R I N S S P L G I G D L E D R R S K F N M G N Q I Q
RANDOM_5589 Q I G L V K P D C D K E L S K N V V V L G Y P R D P L A T K R A I Q K L R S N K Q M A Y D E I R A P Q L G L V L
RANDOM_5590 A E L L R E N Q A C T S K K G C A E L S L L G E Q Q A D G E L S D A P K D A V G S V I H Q G L Y N Q C R A L S E

RANDOM_5591 L P N L L A G G S K V R G T A L R S D I E V F C C E Q N Y V S A S C E G Q L D N T M N Q G A M D H T Q S I R E G
RANDOM_5592 R T L I W S F T T D E L Q S I P Q I P F D L P N L N T E G E R L I D F M A P L T R K G T T P P R L A R W A K A G
RANDOM_5593 K R N P Y R S G I T G P R S F K L P R R T K S Q D C I G S V K L I S E R E L K K L E P G R V M N D V P L S D Y T
RANDOM_5594 T D V V N V H G L Q W V N V F A P G Y L S Y S I F P E G A S E E G D F I R W E L L Q I E C Q V Y I T V E I R L S
RANDOM_5595 F N Y E F R K P G Y E A S V L D L V G F T T G H R E D T S C H G T I N P H Y A S M V F G G A D E F Q I G E F E E
RANDOM_5596 E E S F D L L E L Y G T L A R T I A K G R R N G R G R V A R H L L G N Q N L P I V A Q K C E G S P K V S T Q L A
RANDOM_5597 F R T S R I L L E N R V T T Q H Y P F I Y K L F L A Y R E H A D N T Y P E L N S N E G D D N G N R E H P D E S N
RANDOM_5598 S P E D T T Q I V P K G K K T G T S Q P R P I T E T L K R T T A Q Q P I A G G Y P W F T L G F Q I F I S N L L C
RANDOM_5599 A Q V T V R L W H I C E G S N E L S Y V R G L A L E F W D L R Y D L A F E I N P E T E P G E V A V S T P T K R L
RANDOM_5600 D R S P Q N T E P I L F G S T K Y I Y I R W P R S P P L G N C Y G A W K E F V S N S L D N Q R E W K K G F A S P
RANDOM_5601 K T V E F S I M R A D E Q E L L L R K D F K D Y G S A Y E K P L I K S K T A I E T T N S A A K V T T T Y S F L W
RANDOM_5602 C G F G T V M S N V C D N G M E G D W P A T I R L Q K P Q L I L V T T Y D P C K K E E S Y L P E G Q V G H K A N
RANDOM_5603 M L K P G V Q T T N V S G S V A T D A D S L S N H L A M V R I V R G T Y T P A S W N L A T P M I P G R F D A E H
RANDOM_5604 H L I E A N L S V P S D G R G Y D D I E A A K L G P K P Y A H L K E S G V I L P L Y I N E E A A P T S P P K E M
RANDOM_5605 D Y D L G S S E A V L A S V A A K D V R I R D A G L R K E K I F L D Q H S L Q L A V H S M K E C G M E E A R W P
RANDOM_5606 K Y N A D T D I T Y G K P L L F K I A G D G V T V P P A I N T V D G H T G D V H D R A A D E S S Q I C V T M D A
RANDOM_5607 L L H E S L K Q I R A A N G L V N R M L T K T L T G I Y V L A L T N K E Y D W K K K D P S A S V G N D N K N E S
RANDOM_5608 K D D P L L T G F Q Y C N S I G G S A N Q A S S M E N T A A F K F N A P E A A S V C Q S G K A N L R S A A F S L
RANDOM_5609 E D E P C L E Y E G E R D N T D G S A S R S H N R S G A T S F V P L T G L K V D M V L L F R S Q G G S A E E S V
RANDOM_5610 K N V L S K A K E S F S T P E G S G D A S A L I K H R M M N R F L E R Q I V F L Y S A T D P G E Y P Y V T A V H
RANDOM_5611 L V M V V A W P A L L S D G V E A G D C S A E L L V S R S E E V Y H S V K S T Q M K W M N L G A V V L Q E S K K
RANDOM_5612 A A F P G S A T A S L D A C S L G E E P K E R A V P A L P D D T A S L P T M H Y R N D I Q T Q K F D S N Q A A F
RANDOM_5613 A W V A G N L E V G A L F C S V K A T T V P Q S S G N E I E R V A Q R I T V S K V C T G V F V K T Q S P R L Q D
RANDOM_5614 L Y R R R P F C D V C P L G L D V V L L L I Q G I D T E E H M Q A S Q I E D M S L T R E G V T L G L R I I G G L
RANDOM_5615 T S E L D R Y R Q W P S K L P Q H R K V E A P Q G Q M R R I F S R A G S N K P R S L E V V P I L T R P S L L I Q
RANDOM_5616 A A T S E G E A A V E P I V R E A R S L I D K K R P I K L S V S W N S T S H V Y D L T C T P P I K Q I N G R G Q
RANDOM_5617 V G T L L T T L L E R A V D A I T V A V R G A P E M S S G P E A A S P L S Y Y G V Y L P E I L S F H T P N Q N Y
RANDOM_5618 S V K Y R S S L E A T E G V T K W I V V Q S I P E T L L R S I P S K R K H Q T C V P V A S P M Q R K Y V G L L G
RANDOM_5619 K V L D R G E Y I R Y A K E M G S E R E H I G D Q N L N S A E R P L P I D P K S V T R S T G G L I I Q I F G L T
RANDOM_5620 L I R V Q F R Q V S G Y R Y S R V S A C L T D F A G N N G A F A L C D M K V G E L C G K L V K G P F V S H Y H T
RANDOM_5621 L T L G E Q S E P Y L A R P Q T V L A T K L G H L Y I S P K A V P L A S T A P K L V N E P P V T Q A M T L V E Y
RANDOM_5622 F S S T V T V I N Q A K M Q S D E K L K A S D P G S G K Y N G L P G Y A L F S T C I N N D A L W L A R T Q L Q S
RANDOM_5623 G I F W G G V E L R W A A N A V T D A V I P S P V L N L H I T D L Q H H E G E N G E C H H S A R I T C P P Q S K
RANDOM_5624 L Q K I E H Y P A P Y M G H F K F K V A V G I D G N V A A N A G D Q L T P P Q V R T E E K D P P G H T L K P G H
RANDOM_5625 A I C K K V V V P P P L Q N Y G L E D Y G G R L W A I A M Y G K R T T T R K P R N T G T L W D P P S S I V L K T
RANDOM_5626 A A R P G D D S L G E S N W E F V G K V E D L S F T L R A K G G L S Y Y N E A F Q R T E R A L V S A V N N C G N
RANDOM_5627 N M M G L S V Q A V P H T G F Y P A K G F I I Y D K P L T D D S N L I N D V S P K D V H D G T M L S K L R A P L
RANDOM_5628 R T E N S L E P H Y D T D W A I D I W K A Q A E G S N V M L H T A E A A G A Q F K A N I R D P Y R P N N L D L A
RANDOM_5629 Y G E V A E V D P F S I V Y T R V L F L I S D E T Q K R S S L N K T K Q T P R G P V F R I D V L T K D Y Y G F V
RANDOM_5630 L K Q V G V A N P N K N S G R H W H T I F R L E V E D V L L A A L L W R Y R Q H I T S L I T W L R V G N P Y E P
RANDOM_5631 E T L T L G D V Q P E G S V A L A L L Y T Y R S P F C T M E N N N K E Q C D C Q L S D T N V I I D T T L T I F M
RANDOM_5632 V V T E A K N E K E Q V E D V T R K G Y Y I G R C V E L L Q V D T E L G L T E R K S T P Y G K A V T G W T D R L
RANDOM_5633 I Q K M G P P S D Q Q N V D L W P T E N A C S L T F F S L I T R T N V L T S I A R A L Q Q R E G W H G D N Q P N
RANDOM_5634 A R A G S G E D A Q D K V A N K C N L L Q N P E L N S Y L A N N G E L H S Q G S P P G V K S V L E F K K I E L V
RANDOM_5635 T L K Q F T S E L R Y L R A L R H K D A C G G R K P Q N Y M A L V R P N C G E D D H Y N L Q K A K K H M S P S K
RANDOM_5636 M N D L K G M Y Q R V K G A M H D G V I F G S G F K G L G V Y E R L N V Q S A G S E A L W H V E Y V H G D F F H
RANDOM_5637 G E E K R W G P A E I L G I T L E R G M A G G K K H A V N A A E Y P I T G R A A E V I Y L Q L F G G L M T A A E
RANDOM_5638 V Y F A K S R S Q C N K R E D V I Y E K L K C I C Q M I E D A F R K F Q N E D K C V C T S T L N T R N S Q V V G
RANDOM_5639 L S M Y D F P T K D R R L Q F A R D K G K L T R L S P E I V E W T N S A K T P V L G V D L S R V I T T T W G D P
RANDOM_5640 S G G A S S E V G R G N A N H F V P A C D T S N E I S S D Q R P V N T D N D F F S G A P R Q S D D K G I K R D L
RANDOM_5641 G T R W Q R L I T P H H A R F S V F E L P S K I L K A P H H K L C E L D P V T S T I P G Q S L E S Q L Y G F H F
RANDOM_5642 W S Q L M V V L K A C Q F L R V D A A I S G N K T D P D G T S H L E I Y H V Q L V S R A L L N H Y N K K A K G N
RANDOM_5643 E V K R T R T A D I D F Y R Y G S S G L A E S S T S F N K A F T T R V H A F L Q L S S Q D I S R L V Y G S D W N
RANDOM_5644 R Q P I V Q S R N L L I E L S Q D V T L Q F D L V T Q H C N P D P R G G D S E Y F R T F S S D R P A P D L R L S
RANDOM_5645 N V A G Y V H L R F V G E R Q D G A I E A L I R N E Q R Q A T P G Q L S L V P H N K A K V T T N A V F G M V S Q
RANDOM_5646 V T G G W N E E L D Y F N G K A D T A R T N E A I L F N A P T E Q R A G R T A G I V V K I H E L L Y P E F R
RANDOM_5647 E S D S S P A G R A F V V G P A I V L T E L R K N L E R E A G R S P H D G E S A T D D H A I P Q G G Q L C E A C
RANDOM_5648 L E G H G T L V Q P E V A P A P I L K E I G W E R V L C Q S G Q N R Q M G E Q C C I K Y V K A N L I T R K I G G
RANDOM_5649 I G A N P N C G T G I V T L Y Q N M G V V H S K K G Q N A T K S D R Q P S V T S D N A E K K I V K I G Q G F V S
RANDOM_5650 Q R C A S A K Q G G E V V Y T P A L S N T G V D C N T T L L K I F V L P I A R V P N A N D R L L K N M Y K E K V
RANDOM_5651 F P R D K F K I R V Q F I T D T Y R C C L D E M F D A S C A P I Q P V I E K L D M F Q T M T D A T A L S K T T T
RANDOM_5652 E G F A N A A L E Y V Q A E E A P S E Y S T P R K A L E V S A Y K K R M V P L L S E S D V I V N I K N C L L V A
RANDOM_5653 V K P A I W Y E E I L H R Y H T G L Q R N Q Q V H W S G L G S T K Y E V D V I Q L Y Q E P V G Y D P I L S Q K T
RANDOM_5654 G L S D T F F Q L G E M E L P S Q M D A K G T C Y D E A N N W I L G D V S L W R G A A G T S L E G G G K H K K K
RANDOM_5655 Y G K A N Q A S G G I E L N G Q Y C E L C L A G I L E G I R K P V C A P C D M R G F A R T N D H S S G E R K K R
RANDOM_5656 A P F K R L K G R A I S A L N S E I G N L G L S G N L M C K M F N R I E T E G Y Q K S V E S H E R A L S V K L A
RANDOM_5657 W L S I S F P M L S A S R S I V D L I H H Q E S C N T D G C H M D H E V E A S A S L G G A K G K S L K T L T T P
RANDOM_5658 C G E T L T Q E S S L L A T D S I S N P P G M S P L Q T P T D D N R W L G G E Y A S A T L E L G Q S L V T K G E
RANDOM_5659 V D T E Q T D H E R I A R R Y M G T S Y V D P V A M K S P E E P Y E E M A D H W R L E G F G I P G A S V N K R N
RANDOM_5660 K S Q T H G A N D Q I D Q S K G N D V F A D V G K V S K N E A T N D E I S R L A V A V S N D G Q K E F P L R A H
RANDOM_5661 Y M H A S R K F F L V V N T I S V D F N R T R E S T I Y L R L K M T G I G I P C E R I F R I R K A L G R W L G M
RANDOM_5662 D Q T E R R G V L D H M V G L I E S A L T L G R M K A R N Q P N P P W P A K V K V S A D D Y F H E A E G L Y P S
RANDOM_5663 H R G A Y P I P K A L S E S G V S S I G G E A G T V Q V Q S P L G E Y E I L R T S Y R K V S S L Q E L L A L K P
RANDOM_5664 M V P V N V S C M C N G G Y I I T P H I T V F W K D L I I L F P R Q Q D F V D H Y S L N L L W V L R N D D R L A
RANDOM_5665 G C G S E A A P D Y F N L F A K L V L N R Q D I L G V Y V L W L R C P G K F K S L E L D R K N V G N G K F P N N
RANDOM_5666 Q P I E T L F R T S D P Y A E A L A K V L G T A E V S Y S L A P P T N D A S Y S S T K S L V A K I A V D F L G V
RANDOM_5667 Y A T E E W R K Q K S I R Q S S H E M T V S R E S L L V A S I M N G T T E Q L A P A S E E I P R Q F I G T F F P
RANDOM_5668 V H T Y G P A D R E M L D R D I L K M L N F I K R T V I D Q A R G K N C L N S D R D E C H S Q V S G D W R E A T
RANDOM_5669 E S F D K L K S K F N T K E W F A G A Y K Q D A A L Q Q I I P Q K C D V N G H V V K V Q M T Q P Y I N N I T R P
RANDOM_5670 Q A E S E V H D G A E M L Q Y L R W R S Q A I N I V D R I E A M S A D K L D D K G V F T I N D E S S N S D C G D
RANDOM_5671 P N G A D A M S G P F I S Y R G E G V N I E I T V M R P L G S P K N G L Y M R Y F H E W N V L H G P V V S T T S
RANDOM_5672 T N A V C E L D P G F G Q I T Q E Q A D K G Y C L K V G S D P S Q E L E R L A R G G Q S K A Q S R M R K E Q D V
RANDOM_5673 D P D F L V E K N Y A A G K G Q C Q S V L S A L F M T G K Q T S V G L K G M D S C I G I C L R V P R V L R P I S
RANDOM_5674 L E E A V R Y F R P E H S E A Q D V H N H F L G R T M V T F L I S G L C Q H Q A V P F T R T A V S K F A I P A E
RANDOM_5675 V Q T M R A D L Y K E T M F V I T T Q S D S T F D F A S K K R I Y F I D D E A R Q E D V E L S S P S S R H N I D
RANDOM_5676 V G G L I S Q S A L D E P G G A Y I N E L Q R S F L V A L V L K K Y G V V P G L D G A A F R K D Q Q P V R L E D
RANDOM_5677 E K I S R R R G R P D S A I N Q L G C Q P K Y P L F T W G L I D A V F S W F E K R I E L K A V V C L A R P V N D
RANDOM_5678 L F E I W G N Q L P T G P E S P N E Q L A P P R M K G T E S G W A R Q I P I E E L N C K I L R H N D S L F Q V G
RANDOM_5679 R L S Q D P A V T A T K Q T G D L N D F G K V H Y L H T K V V V T S I N I S L G L M K L D N H Y G N P D Y T L N
RANDOM_5680 V K S V A K V M D S G L K V K I L Q V A Q L Y P M Q S F N A V N L V V H I A L A K S P S M V I H Y Y D Y G E L A
RANDOM_5681 T P D N N N D G Q F V D A S A A N E H T G S L A L Q D E S M Y F T Y E V W D L G D E A L Y S C L S N D E K G V A
RANDOM_5682 Y D L D K C N K E R L A N R L V Q F G K S P E G H Y V A Q L R I N V D D R C S A C K Q F G K G K S G F P I K D I
RANDOM_5683 A D G N W P R D S R V E D G V E Y H I S R E L H A G D S W T V P T A Q R N T T D K A L K A L D I D G C L E A A A
RANDOM_5684 N A Y A A Y I N T I R Q T T I D C I L C L Q N P V E Y K N Y A R F C W H T L D V T T V K E L D E S V H V G K A K
RANDOM_5685 L P D P Y S E I Q L A V L K F E D K T G G T P I D P G Q A N R V M D D R Q F D L R K H Q G L A V I F G Q K L G L

RANDOM_5686 L L Y G G G P D P N K Q T L N K A G P N P N G M R R D N F A N Q T N S G F S G G E K D R V A E L G K F P I G V I
RANDOM_5687 D D P F L R W G R A V T F K Y F S G I V L E V Y V T K S N E K N L E N L D F V V L Q P E L F V E G Q K T G L V L
RANDOM_5688 W V I H V W V Q T S L I D V P I F G E L E S D L V I P A S L A V A G A D F M F I F L D R F N I G G S D A T G T H
RANDOM_5689 L V A H P E D G K P Q W N K N L N G Y K K G R N Y S F I F D F P D R F R A S S S P D L G D L E Y K T K Y M L S N
RANDOM_5690 S Q E V V L L I P G E E A K S R L R L Q E A G T I L Q G M Y G T I S S R K V P S T N P Q G V Q I Q R V P L D Q L
RANDOM_5691 G K E A A V Q E P I I L K T L S F D A V Q G V Q F V T G L G L T I P Y E K I G P P K S E A T P C G A R R Q C L Q
RANDOM_5692 H Q H D R M A D F E L L E I P H H K H H Q A L F Q Q L K A I F A D P V N V L F G K L T D N L K L T L T P A V A G
RANDOM_5693 T N A D L S L T G S T L S A D A G N S E L F Y Y G A Q L N G E H Q P V W H T R L A K E S D R V G L T E G L Y T L
RANDOM_5694 L T V R P V D L I F A S R S M E H D L F Q P E G D S D A L S G L T R N S A S D N E V S A L V S C A C I F V P G S
RANDOM_5695 V V Y T S K I D E L Y S F I E Q I P I F D H V L E A Q C Q S D E L Q P F P W E K G A E S L R I D G M S Q R R A V
RANDOM_5696 K L A V S V N G L D F F D F A G S T R H V A V L V E R I E G D L Y T Y F V I A L A E V A L C E V R D E P H K S T
RANDOM_5697 S A F T T F E I D G K R A D Y A V V L A N Y T S Y K R Y D P P K K L T I S V W T H A R L L G R R V P Y S E L K N
RANDOM_5698 E G G T Q S S Q T E F S V L F N V S M K A A D S P D H S I V Y I D Y A A K P S A A V V K K K E C R R Q K T G E Y
RANDOM_5699 N I L M P A Q L Y A T G G R T E K G A A I A A E E N W A Y F D P E G A H Q A L D H T A T D N K V S R W S E L K T
RANDOM_5700 E S I R H V L P I F D I I L G I K L P A Q L S L V V V E V R K Q S S K A T L A E H I A M K P G L R E S D E P V E
RANDOM_5701 G E T K N G L A N G D T S Y G N Y M L Y E I S L P Q L G F G E D T S K Q D A S Q E A T G E V S I L E Y L Y A S D
RANDOM_5702 E N C Y D E V L S L P F Q V I D L V T P L R V M R E T A Q L D D V G S V I A E K E K E K T G I R A K L D F A A A
RANDOM_5703 N Q F D I L R I T R R H T Y W I L L I A D P L A G T A E G S V S A L V L R A K E A I N N R T V P R G S H A D G L
RANDOM_5704 G T S L R T D M F P C G K P K L I L E P S Q E M P I S Q M D F M Q P K V G E L P P Q E T L P K T V L M D R T A K
RANDOM_5705 Y A P P A A S V E T V D P G V T P S E F C H A E A E R G S V T V L Q P I Q T R N E K L C F M I Y N L V H A T Q A
RANDOM_5706 R F N C H G A V V S K A R R E Y D H L A L S N M P F N V F S N F E Y C K Q L D Q Y F Q G E R A L V V E L K W K A
RANDOM_5707 S P G T E S E Y F K D G M F D T T G W D R G G I Y P A R S M L F A S P E A G Q F L H E G T H G C L P V H C M T A
RANDOM_5708 L N E G Q S I A T H G V E S A Y F K K K E S T V T Y A R D G P L I G L I F Q I E R V G C A D R G L I Y N A M R E
RANDOM_5709 L Y R S L H S L N V H T A D F E Y D I M H D H E P Y T W N L P I G Q K K I N K E F S A S E I F T V D A L Q P K G
RANDOM_5710 S L S R S S S I V E T T F Q R G R T L N K V C A S D R F G S Y R T G D F L S L D M L L D K L S S A T Y D P G K V
RANDOM_5711 Q G S L P W H L D A Y I K V L P D Y G V L G I R N T A A P S P N Q Q Q R Q D R D A I G G D S S K A R M R R G C G
RANDOM_5712 L G H N A G P V S V K S D P V L V P Y I Y G W W I V A Q K E I S A P Q A A K V L G D R A G K E I G F M A Y Q R I
RANDOM_5713 D C E P G N A K D G N V Y C Y M A G S K I V T E K A Y L R L I R S Q Q K K F A K E L A V T E R C P F E F L H N E
RANDOM_5714 Q F D H F R G K A S I S F K L A P T N D N T S S P H S Y L F F L T H F R P S Y G D L E K R T E R S Q N G V K E P
RANDOM_5715 N M Q L M H R K E V Q Y K Q S L D W V F R G S K L M G R V K H V A F S K G Y G W K S C P P G P Q A T G F G L N T
RANDOM_5716 C A N T Y P H V A A A P G A G P L A F E Y A L F Q V N R A P D K P F S D A V C G I L G K R V A T G G T E A P D D
RANDOM_5717 P T H F P I N R A R S C L L D G W R F L S N A C P D G P K K R L T R N E D L Q D Y M G A L G P D R G S T I I R V
RANDOM_5718 A V L P N L A S A Q E C G A D S V K G W V F Q Y H G D D N T E I S A P Y E T S L F Q I G S D G N W N R E D L T Q
RANDOM_5719 T K G R P L R E N V Y L A V F D P I Q H A G V T R H S T L L G Q N C K G G Y D N V C L A V S N A S T Y N H F F R
RANDOM_5720 D A D Y N L T H I I N S T N V R T P A Q L F L R T D E E V Y K F R C D V D L I S T S W A P P V L C L L S L N H R
RANDOM_5721 E V G F A K H A M L V L I V F L A G T G A P L S V L G A D P A R T H A Q K N R T K V Q E K E G G Q L A A V L I A
RANDOM_5722 S N L T S L R N Q G F Y Q R T V A D S L K W R Q W H R T P G I P M D K M L E G K S V Y E K A R L I V M Q Q P I G
RANDOM_5723 T R S F M Q E G H Q N W E S D G E V Q G N A S N F G F A N W E T E P V Y V N T N E V V V K C T L R G Q M S A D K
RANDOM_5724 R G Q F F V I A E V T S R I G T S L R T P I G Q C L L E F E N C P I L S K P K W S D H G S L G A V A P E G P V H
RANDOM_5725 A E V S E G A G A G P G V Q L E G G L W K W Q F T G K F T A E A T K A G A V K H M H A L Q K T T H R I P Q P D V
RANDOM_5726 I Y R L S Y P L V N E D V M A T K V S T V P E N M Y I E S G H V E A R L D P V R L K I P R L E K A V V K T I R A
RANDOM_5727 H K T N P F A Y I W T D L N L D L D I V E Y N M E G N A L D W K G G L Q Y V L R K Y K R D A N N G L R F S S S L
RANDOM_5728 S R V S S P K W D E Q A P D T D G Y D Q G E P C S R D K P L W N L R T D H V A S L N E N A N F L G G F T W G T P
RANDOM_5729 A R Y L T E F T N E I K V Y Q E W E T G M A S F N M G K E S V N T R F H K G P A V S M L E I S M K Y I F G L D R
RANDOM_5730 G T L G G S E A S V V L L K L L E N M A E E K G L R E P E A G Y D P V L V G A R F E L M V G H A H A L E T D K T
RANDOM_5731 E Y V G M A A V D V A T E R L N E L F G G N S K V L E D P G C D P F V T M S V L F L I L W R N N A A L L Q R T L
RANDOM_5732 A T V A F P N M D W A M A L G K L S L D I F A P E D Q K E R H Q S L I K T G R D E N A R K T V I A A V K S A P T
RANDOM_5733 H A R L L V A A A T N N R T L Y A N Q A S F D K T E D I G A I A I R F D D D R E P A K D R S I E L W L G C I S K
RANDOM_5734 W P S G P V G N C L Q E A Y L T D L D P L A K M H C L T R E D H F E N N L E C C C A A E E P V I Q G P Q D L G E
RANDOM_5735 K S I N T P Q R V D R C L L V W G V E C D F E Y R L H V F L T K E V M D T P C K S I S C R H E G P P S I W P K L
RANDOM_5736 T G G K H L P K D M N L P Y T Q H T S G F A D K A A S Q G K E T W A T K E Q P K A T D P P V M Q S F K A Q T K P
RANDOM_5737 Y L D A G V K L A L T V L E M S S Q P I W A G Y N L F I D N K S K I G D V W V N P R P P S M E I Q N E E K L I K
RANDOM_5738 R V S A S K E E I H A P Y L K N F K L S R S G N V L P P E G K G L K I T A S I S K P T G V G L V G N F Y E T E D
RANDOM_5739 M E M P F A N G E A I Q T Q S G K V Q L E N M S A G P N K I K A R S S L K G L V I K V N T A K D P V V P P V W V
RANDOM_5740 L K C L E V H A I T C Q Q V S G I Q E W N V V G Y A S I R R L R L E G S L D P S A V D I A R L S G H E S R N D K
RANDOM_5741 Q N G D F P A S F C M L L S D T L L F S A Q Q V V Y K L S D R L T R M A A A W K R P G G E A G A D H T V S D Q G
RANDOM_5742 A L F Q Q G L S H Q F G E V I P K V G Y Y A R E A A L N R I V K A G I E A G A K M P M N E A I K V M R V G F V L
RANDOM_5743 S S A S S R Y R N G V Y L E W A Q F I V S P D R C K G T E L I T V I S T E T H G S F E E R G V Q V G E A E I Y V
RANDOM_5744 P Y T D Y K S L K V E D I L G I R V R K L G G I T L S F L E S I V T K M R N D D M Q H W L E G D H S P M T D T G
RANDOM_5745 A K D F K N A S C N M G V L F K P C T W T D A A S D L L D Y G T Q D C K S P E V N K E S D N P R R E I G S G R T
RANDOM_5746 K H H T A I S D R L P D A I I E R K I V E D A T R L E I G G L T C L S S D I R S S Q G L P Q G Q I F N L A L K T
RANDOM_5747 R S G Y L S K L F G P S R W F S Y I I I T D G T L I F T S P A G K I D A Q Y A I K N D S G G A I K N H S L A A V
RANDOM_5748 A K E E E K G L L F G Y K R A L G P H N E A E F K Y L N K M S S Y A L L R P G N S S H G N Y P L S N G N V A K E
RANDOM_5749 G P T T C N G S R L M I L R E F K M A A Y L E V T N L K T A S H I V P M T E K M R R D S D L E I S V G S S F D I
RANDOM_5750 G V C L E G I P L A A A Q A T A V M G Q G S N Q S S S K R T L C V R V T L A D D K N G P K E V L D F T L H C N W
RANDOM_5751 Q E P Q N L N A T T F H R T I Q K D D C I D V I L P C G I F S A E D Q T Q L N S K E S T C R Q L D C T G H G D S
RANDOM_5752 C W S A P V E E A W Y Q L S M A L F G Y P L N N A C L A L K G P G K K W K E C T V L D I G C P V N I C Y P M P P
RANDOM_5753 V S L L P L E T F T A K S P A L M I A D D S V I T L Q N T T L T V L Q S N E T K G P D Y L T Y A G R S P C W N I
RANDOM_5754 H N Q P G E L K G L A W D E G F E W R K T T K L S N S G I F G L L A P P D T K G S Q K M D Q S S M P A T L G Y T
RANDOM_5755 T P R N K L L G N K N F L A V A E D R D S T L G G K A S S E H V W M Q D V E S L R L D A T H S L S L A K A P R Y
RANDOM_5756 E M S T T R D L L L E V E E Y N G W E Q V T T E P M E G A V L P N L T T L I D R F P S P D Q A S E T D G G D N S
RANDOM_5757 G W V L D L D F A P F G I I F I L K R R E P A N L G D Y A Q F N A S S C I T E V T C L W P D G K K A A C F N H D
RANDOM_5758 F T G L T D D K G L P Q S Q P Y K Q G N G L I N V I S I K R N A E P L K T A L T E I A G W A V Q S L H R F T S V
RANDOM_5759 G L Y D P F K L V A R Q A T V S L S L P S L S A P S P R D A V K N N W V T A G A P F Q S A V L S R C P I S A L F
RANDOM_5760 P F Q Q A L Q L T G N D N D L K Y I I R S S G M Y R N L Q D S V N F A V Q V I L P S L R F R K R S S D K L V L L
RANDOM_5761 E V T N N L K Q T A Y G P R W G S T W M L F L T S K A F I A R M L A R A S K A S C I L T E A E H L T D D A L A R
RANDOM_5762 R V K L V T R P F S A L K G G T F D I I L E T L P K L Q L H S R L Q N N R I Y L E N H I S E C G V T E T G A S P
RANDOM_5763 C S V I I L I Y I L K V R Q F L V L H F G T F L V M M G L V C K S F A V W I K T G N A R Q T Y V P L S L I L L F
RANDOM_5764 E D S V P Q D L V L I P A V S T R G D D E F P S N W P S E D R A A I I S L N T Y V T A G Y L R Y K V M I Q Y K Q
RANDOM_5765 Q Q N Q Q Y P M V N A Y A W K A G Y G G K K T E F Y I C D T P R F D V A N N I Y T N F K S G R P Y T M G F P G F
RANDOM_5766 T N A F G M Y T I L L E A I V A I Y D V F L S S N A L I G F T N G T V H T I V G R I V D I M V S R L F W V L D I
RANDOM_5767 G T E K F G R L F A A A I S G E S L N A M K Q L L L L S V S L R K E Q I G E G C A G C D V A T Q G K Q S Q H A L
RANDOM_5768 K I F M L L K L S H S S T N E D N R E H A V R V R F I Q D L A S P D F K P G K A K F L T L L A G D R G S G I Q A
RANDOM_5769 W N E V I F A T N P S L S I V Q Y Y N H L P V P L G R R D T M C E R C D F N S S T V G T G T L L K E I T D E I V
RANDOM_5770 Y L Y K K R K H N L V F L V G G K L Q T P S D V T L E L N T K Y T D E K P G E Q Q L N C E D K S M V C A V A P F
RANDOM_5771 M V N V A L C I C K P V W S K I V I T G P E E L D I R K H F R M M A P C T A V T D Y P S V L N Y C I P P A L A R
RANDOM_5772 R I R A F S C N I R W R N E L F K P C G R I L V L L Y R L F H K G G L L R T V S Q E W P P A L K K H E K Y Y R V
RANDOM_5773 A V N Y T E E L E G N E L G L S R P T E S V I I R V W L W V A Q K F K P G R D F V N R L G R F S H K T P G C V P
RANDOM_5774 T I Q C G V S C T S M W E S M A N L A I A K H H R G G V N N N K I A T E R K P L R E N K W C D K Y K L G Y D P I
RANDOM_5775 S C A G T K G F V K R P E F K T E M N A M G K L E V S L R P F E T H D H S M R F K K K T N L G L L A V I D M G L
RANDOM_5776 A F E I I Q K I A L K K E F G Q G G G I N I E S M N S D N S D E E E I S S E G F Q Y I F S D I V T C N N R Q A I
RANDOM_5777 L A A H G I T Q D G Y V D L P D K M W R G L H E N P Y A Q K D K G C D A L A A G V Y L T D R I Q N N D R T Q R Y
RANDOM_5778 F D K F I S L P D V Y L K D D R I C C K Q V N G D L H I E T P V T G Y L E G L L E N A A D P E G M T T S E T T Y
RANDOM_5779 N S S L L K E K L T L R Q V V D G I N H I L H K K V T L L K E Y F D Y L H R C G R S V A V T I A Q T D N T F W R
RANDOM_5780 N Q A R Q D Y L I I N S C Y S M P D C L L I E Q A T D E A G I L S Y N F N A Q P S P G S M D I S Y K G A V K L D

RANDOM_5781 K T L V V Q D D D N A Q D R F K M F S V R L R M V L S G K K A N I N G A M E L A A V I E G P L D L S K D G E C F
RANDOM_5782 A R S N D D S G V G A N A V L R A H D T R A L G V N V S S K M R P F S D Q A E G Y V E A F S I V A M S L A L P T
RANDOM_5783 D R A I M F L K F P L C S W K R T S K I R E E P T N T L R A T A C S K A Y A E G P A C G E N M I A V L L D P L K
RANDOM_5784 D P K Y D H Y D R L P Q N L A V T K T P D L P S D L L L G V Y G P P T A L L E N L E A E V Q S Q D P F R A E K M
RANDOM_5785 G E T K E P L L S L F L K Q L A R C T Y N P P E P L E D D D G A Q L E F A L S F P D V P L A P K L A G V Q L T F
RANDOM_5786 S Y A V A T T P L T T C K D N L L R I Q I T Q S R P R V H L P R S G K A N K I R N E L T W E K F A D G E L P E A
RANDOM_5787 F Q E L S I G N S D A Q F I P L P C N P E K G V V I T R F R I T F T Q F G D V Q V L E A K M V I G P F T D Q R R
RANDOM_5788 F D D L F Q G A E T Y G A E I S L N I T D R R W I F E S E C A N G L R H Q N C G Q D V I F T S P E S V F H D N I
RANDOM_5789 C G G L N R V V S D A L L G R L V Q Q S C V G S H R T D N L G Q S L L K F F R T T G I P R E M S D S V K Y Q G P
RANDOM_5790 D L Y A L P G T C D I F F E M P S E W L L E H L R G Y E L Q F A E I N L F Q L L T Q G K T L L P A S N T E I L K
RANDOM_5791 D G P Q T S C S P P E G A S H V F G E S V T S Y E A E L L A D R H F T Q V R S I S L A R S L A L I T T V A K K E
RANDOM_5792 F R V V L L K R Q G S S E M A H Y P P I T S K T F G D N V Y K A E L G L E Q E E P I I A A S N K R A S V I D A D
RANDOM_5793 I L P E T P K Y N H V I E A Q G I V G A N S G T L L Q Y W W N A I E Q N E S Y G R E K A G K V E S T A H A P L E
RANDOM_5794 I D Y L R S I G K S L L I E V V N G P S Q V A S L N L E E S A L Q A D S M P G C A I N L V E A D V E V P L E S I
RANDOM_5795 T G E T R D L E S K I S D L L Q V L W L F T T L Q T V C G S N M R L L C N R S R L D A T K S A M A N A S G T A F
RANDOM_5796 A K Y M D Y P A W Q R K G A A Y A N K L E S D R C A L S L C N W G A I L Q E T M Y I S A D A P T S L M E N S E P
RANDOM_5797 E R E A E C E G C E N A T D G E T A L Q S A I K D A K R L V A A H A Q T I G K L A V S R K C S G A N K G G V R F
RANDOM_5798 D T Y Y V V G P K R Q S I M K A D C R V L L L V F A E L A S V K T G V S L I A S L T P T R D S S T M N F S N Q V
RANDOM_5799 E P Y F E F G R L E T A S P F I A I L V L G L H P G W N I K K T P L L M S D A K P P L N H I S Y K I S L R Q V D
RANDOM_5800 L V T R D K G L A I E E Q V F T R M R Y W G Y H L V V T F V I D A Q Y E E L A E L M S Q Q K C E S K C K R K V E
RANDOM_5801 P E S K W T H S W A Y C S G L G A S I E G L S W N I I L L D G G Y V R S V F L R S L N L T E L I D L I S R E F V
RANDOM_5802 N C N S G V V E A R E T F P F A Y I K D E K C I G G L G E S F E A S V T Q A K V A W T V D K N S M I D V L E V P
RANDOM_5803 I N V S K P K D F I L K L K L H N L Y K F F I V D S T H V T Y D R E A D D P V A E E L D N S R F G E T L T I C G
RANDOM_5804 I S N A G E S Y I I S L T M E R A V G F V D P S A Q F L T H A T V F A A R T I R F I A N Q L F E A G C R L V R H
RANDOM_5805 R T P P D D K W A P G I G G S A A A E Y W V C K L Y E E Y A S I A D R E L G F N N E D G G N A P K P E M Q L L G
RANDOM_5806 V F K F A K Y L K S L I H K N T P V G N N V E I Q K A P V L N D E E G S C E E V M L V E G F I F V S K E L D L R
RANDOM_5807 L G T D P G D L T E V L M M A Q A G E H L E I L V G A Y E D T Q R D H S G G H P K L A A E A E C D E E F S T M T
RANDOM_5808 L S S P A L L R L R E L D L S S S A G G I S I V L C V G A R D S K F M T T F Y V G Y D C Q E D A N S R N N V N S
RANDOM_5809 M S A A S S V T D N I N P E S F T T L S R D A R L R R D A L A L V L D R P P F I K D V F T E V N V D Q V F Q L D
RANDOM_5810 I A G H C E S V E A G I K T F E V G N E C D A S L R E S F V M L L G L Y F R E I T S N M A F F V G V Q V Y L S R
RANDOM_5811 P P Y P F N K A G A V L A Y V L V I F F E F A K G P S K E V N Q A Q N N Q Q E V T A S D E Q Q I W S N L E V D A
RANDOM_5812 V V G G V P G S D I P Q V A F E L K D V S I V N G K Y I A S I F S Y L S R N S W D D N I H E L V M I V G I P A F
RANDOM_5813 G V D E A P I V M T G F A Y I V Y R I G F K D D S Y E A A S K N H A Y Y T D A G K Q F V S G E N C C I G F I K D
RANDOM_5814 G K K E F G A G D A E L A G T D Q K P P G T I S D N L K T D A A A A V F R L V Y L T A V V F L R L Q I Q G I D M
RANDOM_5815 P F S L A P V A V E M H N I N V D Q I D R T E K Y A G Y L E N N Y N A F D I E W N A V G D L I E C R V A D P N Y
RANDOM_5816 R M N G C P T I A F C K D K R K C H K V A G L V G D Q S Q E S D P A M P W I S L A I E S Q S L D N G W V A T R Q
RANDOM_5817 C L W D E P V H I A L A I A G Q Q F S Q C F D T F A Q L E D K E A I K D A L L Q N D A N V T N R L S S K R I P Y
RANDOM_5818 E F C L I T F G C T K N I T Y P P I T L H Y Y K D F P I L T T K V F V Y Q T G P P S A Y T R A D V K S Q C L A S
RANDOM_5819 V V K H C Y D R G K P K G R R S E N A S F T D S A F V V A R Q A G Y A E G Y M A L R Q K I P V L A Y G L L E S S
RANDOM_5820 I S V L R A V K E D Q L V T N Y F D E R A V N L P K R S E L L C E T V G S L K L N A Q W D F T Y N K T E D E R E
RANDOM_5821 K L L A N F G A Y A T T V K I N Y A A T G L A N K K R G V D F H W R Y E A S R T L Y G K N K L A A R S G N E S F
RANDOM_5822 K S R G K A L E P Y S D M A P R Q N H F H Y S G F W Q S E R I R N M G E F M G R V V C N M S K R G H P F V T D P
RANDOM_5823 D G A K S E E W M D L F A G D S L L I G S E K L I S D P D A L Y S H G M A M V A S T L L K S E F G E D F H P R T
RANDOM_5824 G T R A S M E S P T V L S V G A E N S N L A L G H P A G F Q E C Y L D Q C L Q I L L G G T M K E H K G E R Q L D
RANDOM_5825 L F N L T Q L V A V F I L A R P L K E D D I V D D Q S D R P I M R E T A Y A H A A G V L R G T R A R G T T G I A
RANDOM_5826 D D K K Y S A R P K E L L P G L D A S I K P P G P V A R A L H E P Q R P R N F S E E A A L N S V D D G F V I H E
RANDOM_5827 F G A K Q G F S W E P P K K E K L H H G I E Q A Y K D F A R E H V T F K A D L F P M R K K D K Y Y G Y A I P S E
RANDOM_5828 I P Q V Y Y L V Y E A H L P T T S P I E M G S W Y S S N V D S A D P P E V G Y A V Y C M H S Y S Q E D C G L N Y
RANDOM_5829 R T A P L E K C F M P E I L T Y D T L E Y V S Q D W R G P R G D V G V E E T Q L G E I L P L L C H Q L K N V A T
RANDOM_5830 D K V G P A V G D F K R M V K A P Q S R A L T R S P I P D P F R L R K A D Y G F D Y H G I S L W S T L I K S I V
RANDOM_5831 D N S S H V E N V V L H D P V P Q E A Q M K A K A K E L T T E S Q V G Q P E N L I E E K F G S W D F A T V P F V
RANDOM_5832 A C I W F L F I H A A G S I L K A F T Q L D I D M G Q K L E A A R D F I H S E A V T G Q S K G L M S N M P P F C
RANDOM_5833 Q L P F T E G P A E N L I Y Y Q A V H E D Q L P Q V F I N G E E E P E P C P C H E T E G V R A V A M L D A R S G
RANDOM_5834 A A A V V I K A P G L F V W T N K S T Q G F Y A V D I E I L K M P I A G K C P N L E L G P L E I K D V D P V V F
RANDOM_5835 L G S D V H G W S L V G E P P E P Y F V P D W Y A P L V R H R G I V R E L G F S S E R N Q L Q L Q T Q V P A D P
RANDOM_5836 N D A V Q Q T R W C D G Q H E K A R P G E V L L S Q V T G L S R G N Q A E P N G L G D E R H R T S Q L E T T L T
RANDOM_5837 E S E D P F S L K G D V H P L I A V K P T M E H G F P S D S H R R L S D Q E I P N A A R L V E W T M S F G G Q V
RANDOM_5838 L G S A R G T S D V S M L D P L G P Y T E I E S Y R V A L L G S V L E S G M V E Q F E D D I D D D T A P G L R K
RANDOM_5839 L Q E I F T V A A S T G T K L R E I L Q H G E I E A L R Y H V F K G A Q Q S S G D I D L I T A G M E I E G S R N
RANDOM_5840 Q A T I K D V L A R Y Q M C F P L H N V W V T H K Q L L E L Q I N M T S T V N T Q S C Q N L E V Y N L D T R Q V
RANDOM_5841 C A I I A E A F K W I K N C L L I T S D I V D I P S V G I E L K G H I A G R R P M L N M E A C N D G G L E S G L
RANDOM_5842 K L S L C G V L I Y D M E G Y E M N E V A E R G P P S N R D V I E M C S E T P G R T D V L G P G F L G L C N D M
RANDOM_5843 R S E F S I S I A K R K S M S N G L T S A V L T T A G L F I S I T G D T Q V V G N K P A C A D A T V P V C T T R
RANDOM_5844 N V L N I N A M A D K A V V V S D P N R L L P L S G G L N L P T N G M Y H N K E G F V A N P T A L E F G H K I G
RANDOM_5845 D L I G G Q V E M A T I S Q A D M S A I T L E R P W V I V V S D N T A Y G D L I R Y S P K I L E I L N E R K T L
RANDOM_5846 G L E G G K R A S T T P T S I G F G I R S A A G S G D R L A L G E Q R E N T A T T Q P V T R P E G G G G R K I I
RANDOM_5847 M Q N M P E F N Y P R K T K L M L D P P E Y N C L A L I K I Y S C K H Q L K W V G Y I N A E P T Q T G N D R M I
RANDOM_5848 H E Y A D D D M Q A A Y N S H K V E T I I A Q Q A H G G K Q D M N E G S Q L D S I L R G S Y S V F L Q A G S L R
RANDOM_5849 M H E A V I A K A E I I A E P Q F Q A Q L R S K F F S G S R C I R R L V E K M N G G D L T D G N A P Q P L K S A
RANDOM_5850 P S H S T R R Q Y A L K S S M N Q I H Y S T P P F N N R Q A T G L L R I E Y E R D G S L A R I K P P G E V T D R
RANDOM_5851 L N F A G W D H W N L R K L N Q Y S D F G I V E V Y Y V S I A T L K S F C H D R P G L V Q E E N V H A Q D D V G
RANDOM_5852 S V F P D V D G E Y E K I L D D G L T A G K R V R N P S V E G V I I L R L D E K C L Q G L A E S E A K L N S T R
RANDOM_5853 I R G L L V G F A G V G Y S Y C S R A C F S P A V A M V L S L W F A S R S K E S R S V G A I S V P Q I P N V L P
RANDOM_5854 L Q L R S M I L T G K N L D L Q M D M I P K G N W G Q L Y P R T A T R A T R Q K L W A N V G L G I C F A R I C F
RANDOM_5855 A A D H D S V N G P L R I R E T K Q L G S K G S V I L S E T A P I T Q I G A V Q I A V I G P N R Y E G V R D T L
RANDOM_5856 Q D N S H G L T P N L T V A W A G E H M N I T V Y N F V L P A M P R L H P E K Q L I E N E L A L G G Y A R V V R
RANDOM_5857 F K S S R S N A R P Y E S H E P P K G K C N D A S A R P L V D L A A D H A T S L K D F V L K I S P H L T L M F G
RANDOM_5858 S Q P H Q R K M D M A G A P K T D C I E V R Q T T G S G E M V S D M P D I R R T L Y G T Q V K G F V N S D E A F
RANDOM_5859 Q A T D S S M C M S R I A E R K G V V L G G T F P I L D Q K L K T P K Q P L D G A V A V E K A K Q I D Q G A K L
RANDOM_5860 L I S Q P M T P Y T S G K V G K L V K P V G L A L F N A L N E E N Q S L W R L G L G E I T E E V C V V L L V L E
RANDOM_5861 W Q T R P W Y Y F L A L L A R E F Y G A K A A D T E Y I H P V P P T S G E N R V V K D K N C S A V V L S G F V G
RANDOM_5862 Q G L Q E C K R L E H K C N E A L N A L Q F G E T A V H L V W V P F M S V D R V P C A I C Y R A P M K D G D P M
RANDOM_5863 S E P K Y A A S L V T P H V I T T S Y Q N R G R M I V K A M E G A Q G R W Q A L L D L N A R P D T Y C L Q G G E
RANDOM_5864 V E L L A E A P F L E S E K A K L S E Q L A V Q A Y V G K I N D R K R A P L Q R A Y Q I R S V P L A K L C Q R A
RANDOM_5865 I G T R L S D I S H P I R A V A L F G D V L K F T Q L D A R V S P M I E R T G L E L V L K D R L Q G H H Y T D L
RANDOM_5866 K S F P T P I P I P Y G V L D L G Y S G N I E S I Y D Y L G Q G R D S G N G D K E I R G Y W S N K A F S E V P R
RANDOM_5867 S L I Y E C Q I S S G L F Q D M K N A C L F A Y N F L Y T G G E S S A T L Q E I P I A R K P F S W R M Y I V M N
RANDOM_5868 R P R G L I L A R K D I F I S D Q M N Q R R I V M D Q D V A I H G Y A Y S R D D D I E E K L Q V A K Y A D F V L
RANDOM_5869 S E G V T M Y Q A P D V I K F S I S P L L A P I S E Y A D R L K A D T Q D E S A T L I L A D E V P I A G R L E K
RANDOM_5870 G T F G I Y G N E Q P N E A G K D N S P N S G V I P E Q Q H Q K R E Q E P I D V N N P G S I G V G L N P K R N T
RANDOM_5871 V S V G G I T K T I E G G R E G Y V P R S S S Q A L H I A R V W G N C S V V K E Y Q T P E S Y G D A M H L E F M
RANDOM_5872 L E F Q S T Y I K L R G D N V S K R K L F G P D F L P I A I S M E G W T R H I S T V R L A Y P K G S I Q H I D D
RANDOM_5873 P K T Y E T I H A V S T N A T P Q N E D S T L L P K F R E C A I K S R L R M C F G Q T I L Y T P M N S H A V I N
RANDOM_5874 N I E M A G K I D T R F N L S V A L E L S S E C A Y Y D H E G G Y G E G L S E V L I L P I M F T K E H R A E Y H
RANDOM_5875 L S Q R D S N Y E R Y Q I S K T D D A K V N D W A H G V F S A S T T R P K L H L E A V T N S R S G T N N I N R I

RANDOM_5876 A T K D S K L D G D M Q A L F M I C L R E S F Q V F G G V K A C R T R V Q S H V S I T Q W L G T Q I T R P L I S
RANDOM_5877 S T Y E M D G E K E G G L S G R S P A A V I S K N P M N E S H G A N T Y N D H V R T I V V S H Y P V S R N L M W
RANDOM_5878 N A K S S Q Y K T K C F D S S P R V L G R L L M K D Y A T P V T I I F I Q G K K M S G C V L Q W A P E L R T T N
RANDOM_5879 E N K Q K S P R R Q V R R A P S C H L E A A L D H G T Y A H A S H G V Q G G A A Y D I E P E V M E V Q K T A M F
RANDOM_5880 R R H I M L G P P F Y D V A E A N V I R N G D V S D M Q T A S V E S G G D V R Y V H V A L W N G R A K E S I T N
RANDOM_5881 G V G T R Y S V F T S A L K A A I P W G H R Y V Q R P L Y G K V L D V T G V I V V M T P A Y K A D A S V W V Y S
RANDOM_5882 L R T M K N Y L L L H R S L Q L I G V G N M A Q S S P E S T G V R F C N Q K S K K E S I F E A D N G A K Y A N R
RANDOM_5883 L G L E P R T V M K L I N R T L L S S S H Q E G N N R R F R E I L A G S N D I S K R P K L W I Y P S D T L G T S
RANDOM_5884 L T R T A Q L G A K F L F V D K I M S E A E S R E T R K L D A L R Y T S P E T L H G A A E T A R S Y E D S T I V
RANDOM_5885 S P D E L Q M A K V D V I I P I C E H I P L A T L A Q I L I M P L H L T T R R T D S K I S D K A R V G A S I I S
RANDOM_5886 L D L I I V E P L C A H I Y I A G H K D N D C N S I I T G Q H G G F G F V A E L G M E E I K C V E L M K P G T C
RANDOM_5887 A V T A I T T H G L A S F S D Q T S A F R W H V Q V A T G Q N V T L T N M S L L H E E C D Q P G R S G D T T P S
RANDOM_5888 K R Y L T L Y F S T Q Y V E P M P L A T N V N E L I P Y N A L V S K V R G E V L A C R S I I M Y R R D V Q K A L
RANDOM_5889 Y Q E M P P F P K V S L T E R Q D D T Q K T G D K L M C C I Y H A I A V W E R L S E C K V Y G E L L P D T P Q S
RANDOM_5890 A S L K K S R Q R T R N L I I G P Q G L E V L S I A P A P M F D E A R Y S L L G N G V I V W K P K T Q N G C L A
RANDOM_5891 L A H N L N D S P L V K L V T N N C A Y A K T V D N D A H A N Q D N T I T S L Q E R W E N M Q T I V L L D L A C
RANDOM_5892 H S L N R R S G A T L S L S R A F G S H G G E S Q A T V A K A L S C T V D I F Y G D K L G T Y S G H L S L N V A
RANDOM_5893 F M F Y D S S D N N D D F I Q K I L G F E E H K D K T K K V L L G E A W A D I L S P N V G D G S P I S A K R N K
RANDOM_5894 N E D A S F I S R G E L F Q E E M R K F Q H G K R S T L L P V V A G E S Q R R G K I K E D W A N E G V Y R A C L
RANDOM_5895 P V I R F L D T H V G D H L N H I K D K G K N H K E A Y G P K A A D V L T I V R K P G F R P S P C C G K T A R V
RANDOM_5896 A A S G G N D Y D L R P I V I A E D F I L V V G G Y H I L S G I L K A G G S R A L L I N T F E Y E H W P I S T I
RANDOM_5897 F K A K E Q G Y T L K N Q A F V L D A A T Q W Q Y K H E P R L F P V A E P H S A Q G G P K V T K R K Q D L E E V
RANDOM_5898 S A W T P I I A A Q L R A Q S Q V R E Q P R E G Q G Q Y E A H D P K T V E S S V G S E I Q A H Q I V V G R A Q E
RANDOM_5899 E E G A G I E A M H G S D D T F V Q R D F I D A A P S W W E F Y P I W T E E E V M F V K D S C M P G C G G Y H F
RANDOM_5900 P K Q V S N P L T L M S E L N D K P H D A E I W W Q E E M G A I V I R R A G V T F K D V M H A F A G A A I E Q L
RANDOM_5901 G E L V D V T S P S I Q M R Q R F S V R L W W I G A G R V A R Q S P D D S V V R P K Y P L V E S F C K V Q C L Q
RANDOM_5902 L D L F A C G V K N Q F G M E C G I I S A M K G I R P Q G A M G A R A D E L Y G S Y W T V A P N A D E V H L S F
RANDOM_5903 G G L C T F P E D L R F V R V Y A L G D V D L T A R A I E M Q V L Q M R L L Q M Y S G K Q V L A Q I V F Q K K F
RANDOM_5904 W L L G S P G V G H E S T K N N G K L Q F A T K N I A E R Q P I V N L L T T D Y L W I G E H F I C T A G M P T V
RANDOM_5905 W V C L T L D S D K W K W D K K Y I H N G Q I E T S V S E L K P R E S C T P K C R S S L K K F E T G I D R N A D
RANDOM_5906 D G A P R E A S T A V Y M S C G S N K Q P K S F S S D I T Q L M R F M S K T E T P D S N S L R I F E F S T F V N
RANDOM_5907 P L R A G T E A E K K G D A D L P V V A L G S R K G D E Q L E G R Q P P F N K K N N I P K R L V F G I F S E W G
RANDOM_5908 A F I E Y I V L R D P G S V R V F A T E S E P S N Q E F Y G I D N R N L T K M Y V D F A L L L M S G F A A S Q D
RANDOM_5909 I I P A G P F I R I I W D N S D N S T I V H F W T L V P V A S L E A A A V G L R A K L G T E S R L E D L F K I Q
RANDOM_5910 N A G E A V G D V D L R N Y A L A N V K G A E E S A I R S V H C N K L G V R G I K G F P G C R V K E F L K K V C
RANDOM_5911 A T Q S G Y R V Y R P V E D R C V Q Q A G Q A I S L S Q G L E I H V D C K I N E L V S V L P N P K D V M N N G G
RANDOM_5912 A A G I I W A K Q Y G L K I T L Y G S E Q R G S F N L N C T F S T I V S E E S Y L K A K S L Q K F G L N K K L T
RANDOM_5913 G G G R P R P Y T N G I S Q I I G K A N T E H A P N Q Q S I D T K V Q Y T I C A N V M P Q Q I D Y M R G S I A C
RANDOM_5914 A N E V N D T F K S G K K R P K F Y E T T F G N C D T Y D E K L G N D S A R L L C D I A G I C I R V I I D D E V
RANDOM_5915 S H N V I R N D N I V D A V R D A G R M S D L K W R K P G K K T K T N G L R A S N K S L I A L C L T F R E F S R
RANDOM_5916 E H S C V H L S F M A A A M G H C A V L Y M N N I S A I W S E G G L Y S G P R I N N D N K L T K A Y L A I A Y M
RANDOM_5917 F I K I D Q F E W L E G R N A S E R L G A Y L S G F D L S D G I R N L N D D E E R Q N L K L V R R L F S E E K Q
RANDOM_5918 E R V F S I E N L S A Y E R R T S Y V H T V T K L L G T R D I A A D R G R D L W E R Q S L V F P Y D A G R D Q V
RANDOM_5919 N E L W I L G S G K S L Q P P D K G S V V Q V E S S G S G I S G V D P W P W Q A V T K I L G E G C T V W T S E Y
RANDOM_5920 K L N V S L A G K K G E C E Q Q G V P Q R T S L E E D D L F I K T R S H G I H P A S N W H G L S Q F E E N Q F H
RANDOM_5921 Y E T D L S G A L A P F T M E E W I N L T A F D R D N A E K A K D L S D E I R L Y Q T V N V A R D T L P T P T Y
RANDOM_5922 M L D A E P L L E N H L F T D A Y S A D F C G H L D Q Y A D C L V I Q E A V C L C T Q I N E A A N L N D A H V K
RANDOM_5923 L C Y M L A D T S Y Y L L D V E E T P R D D E R L L L G L D L G N N M S M F Y L D Q N V N L S V I R Y T K D H M
RANDOM_5924 T S P T Y Y D P D N Q G P Q L S D D T L T Q L V L P L E E A K K E E A Q D L S E F A D G V N I K V I S F M T A V
RANDOM_5925 L R G Q L T P K E F L G H S N N Y V L D V S L G G H E I G Y A S E L H V L E V D I R K Q S E L E L S A L L R R S
RANDOM_5926 T T R G D G C D I E N L I H E D N V M L M D E P H P T P T N R P W S Q G C D E L N K K E N D F P N G L L A S L A
RANDOM_5927 V D P S A R P E P S A L T A L S S S D P G G K N A E T A F L A F V L K T K V E M R L S L L F D R C L N S A E V V
RANDOM_5928 T K Q E M D G T K H T D S L S T P T H P S T L A H S L A G M R D E M P V V N P P H V D A F G K H V K A Y V V I V
RANDOM_5929 N T I G E H E L K K T L F D G P R G V V S G C E E S F I S G Y P G E N K Y F E S Y F G C D Y G F E S N P G G L Q
RANDOM_5930 R R S I R R D T A A M E V A Y P P Q C K N A S G H A I R E S L S M Y A A K E P G V I I D V G I G A M C T D A T M
RANDOM_5931 D E A S L A S T S K D Y A P D N S K D L G L E T G G T E E H T K K D R E C L T N L F G G V R I F L N L N P G P A
RANDOM_5932 M Q S L A D N H Y S V A E H Y R P S T E G F I A R S A G K G Q C N S P V I P Q K Q H G A A V V S L A T K V A L A
RANDOM_5933 G T S Q N D N S P A V P D K M V T V A G W P V I N G P E M M G G G Q D M K E Y W T A K P I A K V G Y N S E L S E
RANDOM_5934 K E P V E V C P L A G I T T T Q R S K E N M M P L P W P S R H K T Q M Q N F E G Q D K L E Y L T P D L L N K R M
RANDOM_5935 V M K V H P A F D P G E I A K K L G Q K M Q S L L R P Y E N G R L M L Q P Q Q I S S P K I H E G P V Q A G A G I
RANDOM_5936 L A R T G R F E V L E Y L Y M L N A V G P R E F S S A P F N L C Y E W V R A T K I T V K K L D G L A N H I L I R
RANDOM_5937 Q L F L V S T N P K F Y G L V P L Y G A V M N S I Y G T D T V A W I L L F G V L Q T C R G E P G A G P R A V L D
RANDOM_5938 E R V C T E C T I G N L I G Q Q G H Q S E N E K A I G A Y L E C L T V K S L S F P H F T Q L E I V S D K T S S L
RANDOM_5939 L F L S V A P E K G L S Q A A V R Q L A Y Q V L D L W Q Q S V G Q L C Q V K V S Q R A Y F C Y R M T A G A G D Y
RANDOM_5940 G S T G L V D G T G N G T S S E L D F A R P S T L P C P K A R T H I V P G T Q S M N Q N A R D R D A G A V N N D
RANDOM_5941 L N T N V A A Q L K I L L A K V T N V R V K L R G L G S V K V F V G N D T K V A V T E M Y S S E L S N K N L N Q
RANDOM_5942 T R T N A L E A L C G M R I C S S S R D V T N Q M G T L K L Q G A V E A L L R A D D G G P G Y S Y M Y V L N L L
RANDOM_5943 S K M C K H S F A H A I G R K K S S E G L A R R A N F E T L I V N N K I K V P I S D K R K D D L F A K I C R R T
RANDOM_5944 S L P V N R N A K R G A R K G V Q H L Y D T I R A T T E S V S H A A R Q F G C P T G L M P V A I T A D G E V V S
RANDOM_5945 D P P V E F K V N P G L A C R P S M C E E G A P H N C D S F S F L L D E A Q I S L F L D C P G G L A T S P E F K
RANDOM_5946 V Q R K M F R D K A P Y S G P I D N P E R V G A A D V P V D N R L I I C T N T M I P Y G F S P V G I N K V Q C G
RANDOM_5947 I L I Q D S L Y N E I W G G R G A G A L S L Q E G G N H L E G D L S F V R E L V D A Y S L G A F G A P K D P R K
RANDOM_5948 T T N T C Q Y F L Y P Y D S G Y A K F S N V E G E S H L V Y Y P A L S G T R R K E P A E T D L L I Q Q I P T E F
RANDOM_5949 G V G A R R G P C G L M G L V C T V E D I T C P S L D K D I D S S K K A R G C A K R M A E R E R N P Y G Q C D D
RANDOM_5950 C L E D S L K K A N S G G G A F A Q L S E C I S Y D T P F I L Y V F Y I E L G H Y F L Y E L H R L K T A A P E L
RANDOM_5951 A P Q V L G G D L G L T A V E L Y I Q K V Q Q K A S L A T F I H G L A P Q I P A L L S S T R P G V S K P Y S N S
RANDOM_5952 G A A H G R E A R S T H E E C S V K T Q A S G I E Q M A M S K A Q K S I K Q T E N L Q T H G I G D D E S L K D G
RANDOM_5953 S A K K V G V Q L P D K I P G S D I G T E F E S G G R R I C A P K Q P D M K G I N T E F F L E S R T N G R H D L
RANDOM_5954 G Q P E E K Q F S H S I S D S V F N S L R P T F Q E G P L E L M D L G S P S L A H F S F P A T C I L R T G S F N
RANDOM_5955 D R Q K M V H I N F G V I E F D K L N E Y D L S Q R Q V F Q R L A Q P R E G F D M G C G K K G Q I R L K S F R T
RANDOM_5956 T W G I G P S L F G V S N A K E Q W Y F R P K H I M E A I G I K A K A G S L R T K P K A A D D S L M G Y K L E C
RANDOM_5957 G R Q G R T L I D T N D S I I I I Y K G Q V L T R P A M R K M K F F G L K L N I L H F S T L A K I D V C A E S G
RANDOM_5958 A N I E T L A W T N E E N Y A T E P D L R K E L R T P T G M K L G G E I T V V T V L T F P I W W T V L I Q A A A
RANDOM_5959 S P N N A S P A A I L V A N W T V I R F P V V E T Q T R F E V F E G S T S H G L V S G S E F W T K S G G H L F A
RANDOM_5960 T G S E V T L Q T V P R Y H K L T R V I Q A E F D I R S A G S S V L I D V G K I T S T A N S S M P T T W S L D M
RANDOM_5961 F V H S I Q H R L A M S N F S L G R T L S N G D Y T F K T N E T E K E S S Q A E K F P L R S T D V A L Y W Q P D
RANDOM_5962 T L P W S G H N V L L N K L W M I R H V P E D S I A T K G D S P G E M S F V V A L K R N R D E G K A K V E K G I
RANDOM_5963 D H H P L S L Q I S L S W V C P F E F E N M S R I T S I S S A E R Q V N D V V W G D D D A D S A A A P N A S G K
RANDOM_5964 A S N T A T F N R D H S G D Q W A R F L N G C R G S V K A S T D R V H E E E V V E E G F L E L Q G W S T P N K R
RANDOM_5965 P V Q G E G Y S D N K E G A A R Y N M V T T D G L A L K Q G P G D L D N L I N G N G I I S T T Q C Q C G S D L S
RANDOM_5966 I S L D A R P S G P I C K A S P R Y D A K K I L R D D I Q E Y T E K S G L N T S L F K L M V T D V A G W E I I E
RANDOM_5967 E C I G G N D F L D D V H I E Q G A D A N N R M Q G G S I Y L P S Q V A D T N S W F C D T A S P R L A P V H T E
RANDOM_5968 H Q E C I A S Y N I V K R F K R A L A E D R A V A K K A K N I K K V L Q W P D I K V Y S K T P E H R K V S I V A
RANDOM_5969 Q D L D M Q T L G P V D R A M E K F E I N A R V Q H G D I K V L D D I F D K I D A E D R T R R K Y G Y I A E P M
RANDOM_5970 G W Q V I E K L E V D L N V E T M K E Y R W N E M H S V N G A Q I K L T E G H D S R S K M G H T C V K L Y D Y L

RANDOM_5971 L I H L L R E T E F L P G K F D N Q T P M L I E Y L I E V N L N T R C T P H K G A V G E L N S T N P W A T I N Y
RANDOM_5972 I A S V W Y N R P E E K N V D E L T D S F V V S E L R Q F F P F T C V V A A A K G G L E P S M T A P F A K P H L
RANDOM_5973 K T H L L I G S E D V I V V E R F R K K C G V L A D Q F Y F T H Y H D K A G L P I W G T M L E L A V T G S V H V
RANDOM_5974 N K M R V R E Y T S Y V K V N I I N V A I V P E G S K A H M W I N S V E F S L V E N Q R T E R F G S L S K Y P Y
RANDOM_5975 D P L P Y I F S I T L G V N T K A V G E Q T E G G V G T R K R Q G F L N S G F P N E E A S Q P T S L P S E A A W
RANDOM_5976 V A G K D L N Q V L A D Q M H T K S W G S T V N R L T F E K S L L Q D R D P A E P P Q H Q D K E A N M S S Y L C
RANDOM_5977 A A A E Q I L I P P I S M A S K A C S S A T G R T L R E L A I G W W V A T L A A F S H L P G R T A A V F Q G L I
RANDOM_5978 K N P K L I L I Q L E R E W A Q D I V D T V E S C F P Q A G M R E Q C S I H V Y A P R L G A E D D W R Y L N H R
RANDOM_5979 P V S E I K T E L P S T R R L F P L L I I G E Q V E N A Q E R F N H L F G D K R Q Y Q A A L Y G E A R K R M G S
RANDOM_5980 P Q T I T Y E E K N S M F A M A N G T L Y C Q A L Y I P V L L V G F D T K L N N K E T L K P L I A R V P Y K S F
RANDOM_5981 C D W C T A L S D N T L N S Q R G G S A L D A G D E S L F L K S Q E V I N L E S R A D S N F P K L E L V A F K V
RANDOM_5982 L Y C K N E S V H V I D R G P R A N F V Q T T S G A E I Q T V E V E L L S T Q E M I Q F K L C L K V P S K Y S G
RANDOM_5983 I P F A S T F L D F G L K R S L G I A R A G N I C A N Q V S S P Y L I A V L A D L A N R S T I A R F I S R I E G
RANDOM_5984 R M A A M S C I I A V A L K G L D A M V V L Q I H N E D L L A Q L I T W C Y E T D Y L L V N E F I T G F I G E D
RANDOM_5985 R Y Y L A N K E K C V M S K Q G L F S A M T A D S D E P Q L A W T M A M A Q L R Q P N E G G S I I E A P H V P R
RANDOM_5986 Q L M L I L R N M V T R C S R D V T D R N E F G L L V R S D T D V A D W K S Q M V T V S H T W K T T A S R C N M
RANDOM_5987 E K T R H A V C N G V F R T L D A R K S S D T E W N K H A H L N N R V S P I P E G N T P N Q S L T A L E M A W Y
RANDOM_5988 A E V D G R I S V I Y T Y G T E A F I A Q V R F G F S F N R G D L P I D G G H L E T G I I D A K D N F E E M T S
RANDOM_5989 L G M Y P T V G V V P S L G L L V L F Y N E I D N I H I Q P A S Y T L N I M L K V E D H G V L V C I G T D V E S
RANDOM_5990 Q S L L R T A I E T M G P T S E A S K I T D G M T S V L I R T D W S S D N Y R V I L M Q D P P D L V D I D I I V
RANDOM_5991 K I A V V V G D K H P P G R V Y I A K Q W K G G K L S E H P W S A P G T C I E Y K K V D L L H D D C S A V V L Q
RANDOM_5992 S E P E A L G H T V L W S K Q A R E S V E D G Q I N E S V A V S Y T W A Q P K I D A F S G S P D Q E L R D S D W
RANDOM_5993 T N Q G P E S P N L S G F A Q C E Y G D A A G R N K R D C I P E G P H L A D R A F T P Q F V I R E A F N N T S E
RANDOM_5994 S I A R A L L E L E S Y K I G I P D P G M T F G E A Y D T F R T L G S M T R A P H N V P I I S S V D P S E M S A
RANDOM_5995 I F N N W E L M N K R V V V T P T K R C K E P L A D V D R A L Q L D R L N N H V I T L P T V G E L I A R D G E Y
RANDOM_5996 D E G F I G Q G K S C G A C M T T S L K K R T I N I K T K A T A H T K G H S D G A T K M D I A K G S N N I I E A
RANDOM_5997 A T F R N L T L R V E L I D Y M T T Y D P C G F E I F T V R N R M F T G W R P I S F A I E H H A H D K P N Y D T
RANDOM_5998 R A N N D N V L F G T S P L E G R T G F V R V V V I S I F D G S N D N C L L D P G E V D D D L G R D L G I R L S
RANDOM_5999 S T S G R P D S V W A G G F G L C H C K L F T N R R P P R Q V W E R S P S E C L L D Y G A A M K T P V C P S S Q
RANDOM_6000 D E A Y V G D N P F K K V N A Q L L G C E D D P E K G W Q K T K R S A S G R Q G D Y T W G E L V G H S D K R S L
RANDOM_6001 I A E K V H A E L A V L T R N G A T F T V A A Q S S I N D E L L Q D G S A R Q N A V G E I L T R L D S T L N S Y
RANDOM_6002 L Y N R T A T G W T N D D S N C D R V T G Y S C G S S V I A C T V G K K A N I R A N E Q A T T G T H M L H G K N
RANDOM_6003 A S D A D D S N G Q H R M I H R A S F D K A F G G T F P G F R S G S T S I R G F K S V L M H C N I P G I D V F D
RANDOM_6004 F V I D K N Q T R T M S H K K Q L A S L G L P N F P C W L L G M A T Y W V G Y P P H F I S Q A L T C K R D R H R
RANDOM_6005 A N E N L E S K M R S S R P G K L L K T L A R L A I I E Y E P A S D V L G A K L G A G I V F R D E F L L Q T L G
RANDOM_6006 M V K Q R L I I P P Q R V A A R D R L S F G V L C K A Q G A A L R G E G R E T L S S A G L L R T N G S R V L R M
RANDOM_6007 L Q K A G E Q P F A K F F V N G T Y S R P Q S V K K V S E A R E L Q K G L Y I D D C L E T L N P L D G S E V Q G
RANDOM_6008 W E F C C I D S V T G L S E R A W G A T H Q A Q V K K A I I E Y K I I V Q Q S E R H Q E L L Y K Q E N S E M A K
RANDOM_6009 H E Q A L N A K A G L D T I Y R L R K V T R N Y C R N A R T E L A T K S G V V G E T D G E E K L P K R T E G E R
RANDOM_6010 A M M A F R C I I L E R I G L K E V V I S H G K T T R K L F G T E G L L L V G V N T G K K P S T S K D G M Q F L
RANDOM_6011 S L L V V I H S I R Y G G L K G A D A H A L P I G I R G G R W D Q A P C P V R M K G V L A G N L D G L I V Y P K
RANDOM_6012 C C T F N S E A E R A L N G P N G Q E A K R R R A K I A E R L E G P G W S G T E L D F C V V F A Q L L M R W I P
RANDOM_6013 S P K F A L E K P G L V P C Y H A I K T A S A A A I N A R S H S P A D V E T S N Y V W R H T S L F V L I Q F L G
RANDOM_6014 T I G L R M F I T L F A R L A I A W D F L T R P T G S I P P V Q V M L F A K S D M A V H E V P G M A R G G S T V
RANDOM_6015 V Q G H F E G G R M L S P L N F S F I A R L S L P I W K R V C L T I C G N V T G Y F F E A K V Q W P A Q A E H N
RANDOM_6016 D T R A Q A Q D E E S A L N D R A Q C G I L L P R L S F V N P N A S T E V D T I E E S N L F Y A G L S K R G T E
RANDOM_6017 F P S D G V M W T S I V S L W E Q M I V M T P M A Y V I L V L V P A H Q T A V L F H N I A K G V G G V M D L K D
RANDOM_6018 S I S F S S Y H I L V P F I I K K V F V A Y Q Y V H V L V A N E E E V C A A Y V A R R G Q L M E D F F N E I T G
RANDOM_6019 I A A F H T A P C D M M K A I D P G E R V M R Q L S A Q M T N E S R D L Y R N Y Q V E D F A P Q P P N N E I E E
RANDOM_6020 D K M S L P E L G A N E S T Q T I R V C T S L A P N I W E V R V L N N P D K A T L D D P E L Q T Q G L K I A S F
RANDOM_6021 Q A R I P P L P S L N T D D Y L S C E Q S I E R D S C R I N Y I D N K K M C S D E V N I F Q T A Q D V G K P R E
RANDOM_6022 P A K N L V I Q L E Q T I E P E P S R T Q P G R P F N G A I S S D G F E A A P A R L K S E V L K Q K V L V V V D
RANDOM_6023 P L S K I Y P I K L V P K P D V I D I V Y G V A D Y V V T T H Q G F G I P S K E Q E L V S R V S S F I K R D L G
RANDOM_6024 T S P T T F G V R A P K G L L A D V A Q V M H S Y F P Q Q V S D Y A R G P A S L E G H M I R S R V E C N F A G P
RANDOM_6025 L E E K S D Q A Q D T S E P N G S D I S S E E D A V M D A E T L R E D W T K P S A P V S N L G A L E H S A S E T
RANDOM_6026 T E I N N E A I T P G D P K P F E N E K I E G A V G A M T A E I R E T V T Y N D K K Y A L E L A V F R V E Q V R
RANDOM_6027 Q E L A G T S C L L N K D I P L A Q Q E S C K C F I S K I T L F K I H G D E D L Q G S L C A G S L H A I Y A R C
RANDOM_6028 K R N M K M V E G Q G L Q I N R D K G K Q I D A S R V E D A I F M G S F L T E L I A E E D A Y Q E S I V R A A N
RANDOM_6029 S L V P S A L K E A Y V E Q F E Q S V I A P D L N G H F T L F V V R G I K P E L Y N L A E I K S M Q L E L A E I
RANDOM_6030 D P A S A A V P F L G L A P F A N Y T A K R D H W S Y R E L T N K N A S P R Q F Y E M G A Q L R R I M Q N F T M
RANDOM_6031 E Y Y F T N S K N D F Y T D N N P R E G K P E A V D R A A I A R I V P V R Q S V G G W W L L K N K K S A I A I V
RANDOM_6032 D G S S E I L F G E F I A K R N E Y S N G E S A P A T T D A Q L Q H E E G P E E T G N L P K K G S R Q E A V S S
RANDOM_6033 S I A S P I Y T R E Q S N D K S P E Q I R I I S H E L E Y S L T M E T I K K F L N G N S F W K E S K A G F V A K
RANDOM_6034 N S L P E K L Q S Q L N I L Y Q T A A Y T S R P G P E P N F K P K S S A V S N S A P V D I W E M A V D L D P G K
RANDOM_6035 T P I R R L T Q I K V V E I R L I A D W V Y G D G V G D H V P F M Y Y D G P S V E A V L I Q P K A Q P M K R N K
RANDOM_6036 R S A F D V T D F A I G L T E S D C E E R L G E R I E Y P E A L M M Q G M V V A I G S D G T P A H L S T R S K E
RANDOM_6037 K Q S T L L Q G M D D D A G V A C E E P N K H R K G G A L D S R H P V V L I E E Q V K Q G C I S P H R D L V N L
RANDOM_6038 Y E P P W G Q R M G F V L Y D D I K Q N L N G S A T R V A G H E G S D F S G M E I C A T G K P A V G A Y P D E A
RANDOM_6039 N V V T I D A A T N L S D V N I V R R C L V Y I I T H A D F N K R G R I S L E A Q E D F I F M L S N V D L A E F
RANDOM_6040 S G P T I C S D Y I I V T V H C K A K E A G G G A G Y L L L E T I Q D D R E Y D L S S E L S L V T S P S L Q A E
RANDOM_6041 L H S Q R K G R L V L R I H V I I Y S T S A L V P N Y D T S N I S S L A L D G T F R D T S P K S N G G T E A C A
RANDOM_6042 F C H I G D P V G T K R T G S V T N G W I C I E D A Q I I Y K W D T S F A N G K D H V L K F T A W W S G V P F P
RANDOM_6043 A P L C A Y S L R R M L G K A P M D A A T D S R D C T G Q R V A D I S M A G G G G L E K Y R H K D K I Y V N E V
RANDOM_6044 G A R L D R L V N Q F L V Q G A R K D G Y S D Q I L V E H D P P G A V G M S M V F Q T A L E T K G L Q T N Q D L
RANDOM_6045 V Y F F M L S N N A H I A E C T A R W L L K S G G D Q D L S T P V G V E L Q G M I V D R S E K L D A C D N P D K
RANDOM_6046 V D T E V A L K D K N K N G Q G L Y N V G A Q T V S L S I E L T Q N H R A L R A P P F N R C Q L Q S P H V Q D S
RANDOM_6047 S R L G L W R R T L P D F Y K K K G E S V D L Y N G S S N T G S L R T D K V M Y E V G G N H S G Q V F R I M V S
RANDOM_6048 A Q D L I Q P Q E D Q Q L Y S G F D I E K F S T R K A I G G I E A F N R L L A G I E L L G L R T V Q A R I G K H
RANDOM_6049 T E V S A V Y F K D V F E K N T S L A V A A P E A K R R P C K T L G A K V Y S G V F T A V N G Y S D V E L F V Q
RANDOM_6050 T Y D R E R Q R E G G E L I A P H E G N R R L V K S D P P L L G N A Q A K N K A A D V T T G M T E P N T T P L N
RANDOM_6051 A V N W R L N D G I M A P L G T L H I E M E G A L G L A A T K M V V E P G G D V V R V H L K S N F W G P L S L K
RANDOM_6052 R N G G S L L M F R H T V A S R E R V R E V S C Q E D V K K G V F N R R S A I Q Y G T A P E Q L I H V S N L I D
RANDOM_6053 P W D S N G I T S K K Q G A R L S S P S R N R F C R K L P H Y S T R S A S E N N Y S S K D K W L E G F R I T S M
RANDOM_6054 G G D S L S G V K T T H Q Q T L S T H L G N T S S V V L T R I M A N V F V F L T H T Q Q V Q N G Q F S E K D T L
RANDOM_6055 I K K E Y K P G V V A S P E L E G A D R G G A F V K C G N Q A R L A R K Q G Y F I F H K V T P K P G Q P T K L A
RANDOM_6056 R D E R S L L M S R V G H P V Y R A P K V G I H F Q S W R G D E A L N F T D G Q S S L V V C S T A L D N L M P K
RANDOM_6057 F W N K L K Q E A A A C L I P P P R K V E M V L V L D P L S T G V E K N D V D L A S P E Q A D L S L N L E W A K
RANDOM_6058 W R L T F P H A E L S S D W I E P K Y K V D A G E K L R T P S L L L G S G N N A Y L P V L L T G S K K K M K I K
RANDOM_6059 F K S R V A A E G T T A V S N G V G H S L P R S T V T V H K W R Q E N T T I F K C A L V R T F D S I E Y I V E C
RANDOM_6060 T S D G V S W Y K V T D V Y A R S Y V S S R T V S C L L I I V P N L Q D N E D L G A P G N G A V S L F A Y V L Y
RANDOM_6061 T V K A K S F G S Q C I K I G I E A D S L Q S M S S A C E F A Q D R G Q R G Q I P Q T F W L W V P R N L V A P V
RANDOM_6062 L D E D T E P A D K R Y S V D A S L D V V E D V S D K E A W T T E N S H K K N A I I C Q R G W K R D S H T A S A
RANDOM_6063 L G T V I T L S D L E R A F L R N Y D A Y V S V K T L A I I G Y G E N R Q A K D K A M A N N F K N R D L S K A V
RANDOM_6064 V A G L R D I R Y S Y A L E F S P I L L L L M H S K H Q C F I R V A V T N D P A S T T N E H I S R P L R V L R E
RANDOM_6065 N L G M N P A D G S L N D I N S V E V R G V P P R I M Y S R V P K E M R F Q G V P I A P P T L I L L P G Q N W L

RANDOM_6066 E P M S K N P L V V G P V I S L A N G Y R V E F V G P Y T A M I K I A C R F S A G G Q R E P Y Y K L A L Q Q R G
RANDOM_6067 S F L A I R R A Q S T S N E L L I Q K A V P F D K A L T R Y D H T L I K E N T G L A N N I A G S V F A M T R I R
RANDOM_6068 S Q K Y C Y S V G P Q T E S R K A S F E M T P D P E G K L N I L F G Y L S T K I C M S R G P F T K T I Q A M I A
RANDOM_6069 E T K S S K D A K L S V I E D S D L R Y P S L M D A E D T R E N Q P K Q Y K Y D S Q C H P H E T G S F A F R P L
RANDOM_6070 A G V Q V P M I K K T G R L P S D M M Y H Y T L G V V R I P I T E R V S S A G G L A P N F S D A P R L N C A T W
RANDOM_6071 L S K A R R S D G G D A S C I P S I Q E I N G V L N P S L P C Q G E S E I L I E N H M G D E V Q D H L E L G D Y
RANDOM_6072 K F N W T W I V V G V V R L A Q R L T V T P I A G Q T F G R H K A I F K Q Q A G G F D W W G P K V D K H V H F I
RANDOM_6073 Q I K G Y I A L R M M M Q G E A G E A S Q P N A L R S D C T T L E F K P C S L W K R M C R P G G G K S M L Q P D
RANDOM_6074 T R Q D C V P T D S V T Y I Y R R E H A R S G T M D R I R I S A D L G S S E S L L D K S D S L N Y S S E H V A I
RANDOM_6075 K K G P S T E E G T L P N L R Y V M I K D E K S Y K W E N L I V T N D D G K F S D H M P N L T N G E T D V V Q H
RANDOM_6076 V I T P H V C V T V V K G S G D S R N M E Q S D A R E L R V A F G L M V S H L M Q V L L E S D E A S F I G T Y L
RANDOM_6077 Q H P P V L C G V Y L V K V N T D F S L L G S T E D R D T S V D V K S D G V P K A Y H A V G S I S Y Q D E D D L
RANDOM_6078 L D G G T R T G P Q P R I N G E V N V A N A P D D Q D Y V T L H I A S K P E K E R K T L R Y M I S E K P T A Y V
RANDOM_6079 I A Q A D R A Y I T H Q T P A R S Q R T Q I I A I S S G G K F N T G S H W V I G A A Q M Y I N L Q E E V R A I G
RANDOM_6080 E V Q V D L G E R R A P S N L P S A R S T V T R R T L V D S T Y P K D K S G I A K Q S D L I K K N R K L E G M P
RANDOM_6081 R E P S R W G H P C S S G F G K T Y I G K L W T G Q V L Q N N G F K T G V G Q P R A L V C P Y Y E S F N E F T L
RANDOM_6082 C S A L T W P T Q V E H G N A T H N N A A G K I S T T N P H L S A P W A R V K V V L W G P Y L A N L L K V W R G
RANDOM_6083 Y Q G I A L F L V K I E L K I P I V L Y G Q K G L M Y P P W V D S L K P Q D N K K A L S I S G R W D W A V N Y G
RANDOM_6084 K K S V T I L G I K D R G K R S E A L K P I E D D S L E N G R T V L I S A K I D N R I F D A V V W F G A V R F I
RANDOM_6085 K G G L F P S R T V L D P L C V T K S W Y A L A F P E C Q E F M D C Q A S A V M C A M L K G K E R G A L C V F Q
RANDOM_6086 K R G D I A E K L N N G A G Y Q P S V S F K I D E G C W H R A A Q F V E A S Q F C A F T S V V V M G G I P I P A
RANDOM_6087 H E L A S M K H E G S P N E D Y E R P G S K G T E G I P A L R T E I K Q A A P R D K E T A G I A N S K K M E N M
RANDOM_6088 S D S K V E Y L D K E T V V Y G G Y L P L M P G N A D G F Y R T L K S A A A F G S D M D K I P D V L T P N E G K
RANDOM_6089 M L T A I R E C P D E G D Q M L G V S V N G G N P S P I C F G N A F K N S S T S L A D K R D G S Q E F N K S K F
RANDOM_6090 K K G T Q E L K R A E E N F D A L D E A P S K K F P A A I R E P C A T M A L V N L G A H F K D Y E P Q L A K I E
RANDOM_6091 K N K I D G V L H P Y Y T G T K A G E G D M T Y E G E F A C E R A H V S V A V S E D H I F P A D I Y H G E Q T L
RANDOM_6092 F V V D V C P L Q L G K L D F V P H S E D S Q S A T L P D V L I C T L L E Y T E M P L N L K L N A Q I V L T Q G
RANDOM_6093 T A D I D K T L S D V G F E D I Q L L D Q G G H D G R K L G K N C G L P A L R G V Y R K I E W E E E M A R S L P
RANDOM_6094 M L G N M G V G S E V S E V E T P P G A S C R E S T H G K H A V D K W A L P Q N S Y Y L S C L S S T N P F E Q N
RANDOM_6095 S Q A G A A I G E E C L G R E K L E T Q N S G N K E R Q G P T D I K R I G T V F I A S L K T A N A E K K L L V V
RANDOM_6096 F R P P S T S D T P K G S I P K G D I V V T V H R S Q S G K P I G S G R P C V Q S L V F P E S K S W V S S G S P
RANDOM_6097 I E L T W I G Q L Q L R S M L G N V T K D I L N T A R G L L E D V F S P S G T A I V F L R P A G K G E N D L V K
RANDOM_6098 V C I L A A R D Q C P S E L S L K R C A P I G K C P Q W T S H T V I V L K L P N G H I L I N K W K R F S S T Q N
RANDOM_6099 M T K C G A Q G K K M S G K I F K K T F I D T D Q D F I S R F S T Y T P S A H I M K S V G S G V A L N S V I D L
RANDOM_6100 D E A N L G L G N G Y N A T Q D T S A R K T F E R W A V L N V T D P Q T S Y G A G L K C F R Y T N G V R S S P P
RANDOM_6101 S K G A S T S S R I P S M D P V T Q E I S Q F I K E A D R G G T V P F E R G Q L M V F N S G P R S D I D N A T L
RANDOM_6102 L L L V P F F Y E L R L N L I G E L E G A C D T I K A V L E T H P A G R E P R F G V F L M S T R S L V S V F G A
RANDOM_6103 R G Q L A A C T H N P E I D L T K K R H D P P G I E I D T I N H S S P L Y P S F K N G L G A Q H L L V N L A G I
RANDOM_6104 L A E F A A P S L Q P V Q K G N E E N D A A K T M D N W G V E I G E E C D I K K A R L R P P S T R A F V V I I T
RANDOM_6105 A G R G T R A V V V T G P L E A D Q I F H V T Q S H E N P E I I T T M E P G D N L L C F Y I H P Q L T Q N G K F
RANDOM_6106 M R L G I N H L S K F I L Q I Q Q S P G I D C I T A F C R G V P A G A T E N I S L F L I C E C L V D A G E E N N
RANDOM_6107 Q Y F N K L E C Y Y S S E E E L K Y S E T S E K R L R P D H A Y Y Q V L N H V G A H V R T M K A G S N D K A Q A
RANDOM_6108 L G T A L R D V A D G N S M E S G T K E D R S V L P M E V M W A T N P N V L P E R A E L L D A I I C K L Q S A W
RANDOM_6109 E L K H S G D L Q G L T L V Q K K T S M A Q G K S Y K G W D A A Q D E L S I F K K S E D R M C L T W V Q D K D F
RANDOM_6110 L R D T L E R L V L T Y M R L A T E T D A A E E P C R D T N Q R R D G V R Q G G L M D N R G L A M F Y H I R K E
RANDOM_6111 L I S G D E S W D K K D L P Q Q T V K F A I D I T I R N Y A S K I Y E V A N D L G C G S T F Q S L L D D V K S S
RANDOM_6112 L S K Q F Q M R N H C S T T S K T S G C I D G A M F M A D G M A R R S I P Q F R L L Y N V Y T L F S R G P R E P
RANDOM_6113 L K Q E G E H H G K Y H A L L G N Y V Y G Q K L V L S I K E T K I G T R E V R P K P A R R S I V A L R F E A F K
RANDOM_6114 P G R K D T V I Q S I I S Y S E A V I P T D S P P I R E A Y P N A P H R M P C K T N A L P P P F G P E H P D A F
RANDOM_6115 P I G P E Y G L W K K S E E P K D H A I F V K N G L A Y Y S P L S P A T L A A S F A Y Y G G G V L M R K V M D I
RANDOM_6116 P A P G N D L G E A D N A M E Q R N K R L T D Y R D N P A F D Y A N V D E H L D S V M D S N G K N Y I V F G H E
RANDOM_6117 Q Y F P T D E K M G H K I Y N E P G A I A D L V P V S R E Q T K I Y Q S S V A S V N K P G P L L T V R T R L S A
RANDOM_6118 D I S N A E D V R K E S E C H F K L S S L K Q V A E Y A P E K R S S L N P G N A E N P K G W G G Y G L R G I M S
RANDOM_6119 K C T G P P D G K A I P N C D R D V H L S E R E S S P R I L L S A I E L W R F P S V L L S D G L T D L D E L W F
RANDOM_6120 T G F E T M P M A I L P F K A L F P K I A A D M C W G S I R A P L S N R G M P E L V V D L F Q G P G R V N N A R
RANDOM_6121 P G L T G E L G L S F K S M V S D N V F L H K V T E S C A S G L L C S G F K V Y A V T I L Y F L S A R E S E V I
RANDOM_6122 N N N L K Q P L P V N P R K V T V A V P N A M T L E R H Q E C D K S K T Q T K R S R D K F V G E Y T V R T K H P
RANDOM_6123 P G Y C I I L P V E I S S P K S A E L N Y N G N K P E S D L V A S Y R P L Q E T R L K T C A A D G V I D A E F A
RANDOM_6124 W I L D R I A E Y S D R M I Y P G V W V T E S F G T R T A A E G S E D S K A E G L V S L Y T A K K S V S T P D S
RANDOM_6125 S M I E V P V I P G E N A N E E Y V L T D I F S V S L A L T K G G G P L V R L R Q A Q G N G K L L D E L F E G D
RANDOM_6126 A K M P R E K E L A M Y T G T A Q C H T C L P V G R D T E Y D Y T A R F P A E E K K E S V V L F D S P K E V I Q
RANDOM_6127 K K S E F R A S W E I F F F A I G A Q D S A P G L C D A G H T R C A S V T I L E A R G L R L F I I W Q A R E G Y
RANDOM_6128 F V I S F L S N R G V K G K E S W S L S D N R E S I L Y T S K E W D E K Q P E I N S H K Y F K V E D A Q T H M I
RANDOM_6129 V I H S K S M T Y E I I L D P D R P I G L R K P L E K C L F K A Q E D T H A W C G H D A V Y L K M D V C C A A P
RANDOM_6130 H A A G V L A R G T H A L G A G F H L V E G F V N T P G E G T V Q K D V I A L L L E L G Q A E N P Y W S T C T T
RANDOM_6131 G T T K I S E L Y G G S R V K R D L C P K A E G Q A D R D C T R V A N E Q K G P Q Y R V D S F D I P N G C L A S
RANDOM_6132 P G D P D D L E T I A Q P E H G Q P V E P K S N L L R E N V I P A Y F L D H S P D P L T T Y A F W V I N A L A E
RANDOM_6133 S H A N F S L F K C M G E N F L G R F P T H R T D T Q H I A F L Y I P V D R A H E Q L V I Q S L L T E S G T T S
RANDOM_6134 M N P V A Q C I L I N F P M N E Y S L W G F A S Y I Y R L I A Y K L A Q F L A S G V E K A L N L N R L I Q E D D
RANDOM_6135 G L L V S Q N Q E Q G L I N S D T N G I Q G V F L A K E S A D G K E G M L F H R A Q A E L W H G E D P I L A F G
RANDOM_6136 I V Q T G S E P V E E R N K A L S V P E V H V P I A F I R Y V G L N F A Y G A K G G T G T D K L R L K S Y L K Y
RANDOM_6137 T I S D G K L A L A E R K N N E T N D R E G D G S P G Y S F W A M L L A P G P V R G N P L G K S H I Y S R V R N
RANDOM_6138 D G V S D A G H Q A D Q G N G G A V A R K N E S I F V E D M Q I T Q G S K W N A H S C G K L I L A V I S K S L K
RANDOM_6139 I Y L H F S Q V F S D M K L F P N G Q R K G P D D F I K D R K C T T I N V H E R I G V Q A Y P Y N V V G M C R A
RANDOM_6140 S A S S Q K K P D T V P A H T R T A V S F I K N A N L G L L E E L G P V N G P I D W H D S A K T R K I V N A A S
RANDOM_6141 N H Q Q A P R A G A P Y G G M A A E P N Q V A A S I K N S R L Q Q P E Q G R N V E Y R E Q E V G A L P P D A R M
RANDOM_6142 T D V S A E A M L I P Q S Q P G N F E K P G V F T E R T A G P N E A M A L A A K L T H V L T Y L I K P D S G Q D
RANDOM_6143 A G K G M F P D G D D P D Y W V N R T G L L Y E G K E N L V L L I Q T A V G F W I F V I E T T S P Q C A H V G A
RANDOM_6144 V Q D Y M V W R I A P K I I T Q R K L V T Q K S S E E T H G Q D K L R T P D K P P R V I S I R E K I D L D N W K
RANDOM_6145 L G F A A G K S V R Q C I E D R S T Y Q I D Q F T G F V K Y P T W L L S S E F L A K V R T L E V T D L P G K N P
RANDOM_6146 K S V P S W E T G L T E G V R N K G A I A P E A Q Q A S K G T D P G V G Y Q V S V G V A K T F L R G A L P V V T
RANDOM_6147 I R P C E C V V Q V Q R E T P E I V G V N T I C T Q P I D S I Y G Y R R E N V T F Q K T K Q Y S R C S N I P T I
RANDOM_6148 A D P S A A Q W V D P Q Y T F R Q T P T K G A K S I A R Q M C N T A D E G R T Y S K P D G V A M G G S Y P D E L
RANDOM_6149 G F Q P P T N G P E T F K R S I G A C Q I A M T R T E C T L L R P G K E Y R D M A K S A E N P D Y Q S S H T L L
RANDOM_6150 L T H N D K L L S R I R A G T E V V S G D P V P A L I Q E I K Q I A P N H A V D P P K V A V E L V I Y Q A Q N L
RANDOM_6151 Q E Q T L I Q D K N L I P L K K V G L N L V A V K D L R N S D H K A R A R K G S S L S E L S D G E V T I A I D G
RANDOM_6152 K I I M S V T T G I R R E A H N S D S L Y P L S A H I Y G Q T V L G G G M D V A R S T G G D C T P W F A R H R I
RANDOM_6153 F T D A F L S T E L V T A L K P K K S Y S Q R R D A D I I G E I H P D S R G N G H G L V V D K P P C F M P E T F
RANDOM_6154 E V P G R P L L L K T L R K M S E E K D A I G L Q Y Y Y W V S Q L R T S S V Q A L I A T D P N E V I H G W A T L
RANDOM_6155 E I A R V D D G L Y G A N I T M V E P E G L I E P A N V T G I R L T C A V H T H T S S D G L D S I E A I S L L T
RANDOM_6156 K A D V A L V A E R L P L N H Q G K E F K E E P E P S T Y L H L V N G H D A M M D M A F I V G K D L E I D S K A
RANDOM_6157 Q T E S G M E M K A S H K S E H F E S A T Y E T I I G S F N F V D L C R L Y Q S H F D L L S R K N I V A L C H M
RANDOM_6158 A D L E V N A G Q P F S G S S V F F I Q L G V G A M L D L G L A I E L G A S E S A L L L G R C H V K K I G P R P
RANDOM_6159 G T Y L E K A E P E T I E T S T A E P Y K G D E E K S R D A M S Y Y Y A R R I G R I S S S L A T R N P I A A Q V
RANDOM_6160 Q L T V K C D I F N S K L L I A D G G S P W P N N Q H I R W D L E M L M D P E K I R P W T R G M T R I W D F V C

RANDOM_6161 L G V E V G T M D K Y S V F G V R N P L T G P R V C K C I Y R D F L F T D I S V W N A A N L G F L K K P S G Y L
RANDOM_6162 T T V M G G P Q V D S F L V S N M F E G D T T L Y S E V E I P P R M Y D E P Q N S K D Q E V V L M N I S S P N F
RANDOM_6163 L S G H L G Q E S V V G Q T E A D R L V I E N S M R G G E F Y E A T G T P F G Q K L A E A A V G F G F S P A G E
RANDOM_6164 K W S R L N Q Q K W P W I T E R D L E L R I K S V T T R F A D P F A W A E V N Y V P F I L C L K Y L T C T S I D
RANDOM_6165 T V N F V N E E V G C G R A M L T S L K I E T R F F V D R S N Y S D D S M S F S I S M F Q S P G Y R M V E R L K
RANDOM_6166 L L N W Q A R G M N V C G L F P A C R Y K R F M K T K L D D E T K G R A G S G I L H G G L S T I T E P I K A D T
RANDOM_6167 T I S F A A T L E L A K N E K S H P S T V L A P P A S V K R T T F A E D N L S Q E F N S E A K S Q K S S K D T R
RANDOM_6168 G A S V L L N F K E V E F V W M L S V H I P A K L K E M E S G A L K S K H G S T T K N Q V Y D G P H P Q R P G V
RANDOM_6169 K Y G V Q D S K N Y R P P S A K L Y E E A D E L R T K A G V L K L D I H L V A W N V V H Q A G Y S W I T Q A T Y
RANDOM_6170 G T I K L G P I F T T S Y A T S S S G V L P A P L S S A A T T M S L L R E Q V D V E L N G T E M I L M E D D V A
RANDOM_6171 E L G E L V D Y S E R D F P F L A T G I Q S L H F L F K L R T K S V Q L E F P K W V A M R T H Y R P A L T R N E
RANDOM_6172 F K P D R I G A P L L T L V C S P I A I H Y P Q H E D N G P V H S K P S L S G G W P D E V R R Q Y A V F R F V V
RANDOM_6173 Q R V M K V D E T Q Q S V P V N N G P F T A N H I V A G S T G E G H H A K V P S M Q Y K D E R K S K G A L F W P
RANDOM_6174 Q N E M Y S R L K R E H N L F P A R R P H Y S G L D I E T L G R H S F P Y L L L I Y G P A E P R T N Y I L V V A
RANDOM_6175 K P A N E V T I M P N Q P I L L S S L L I G G V E A A A F D Q Q A C S V H E E T Q D N A I L N K H A K D S G L L
RANDOM_6176 E G H L P G N E E F A A I D P E T L I E A G L R R F Y F D E E E S N L G P G E N K R T S I D D L P G G A F R I I
RANDOM_6177 G F P T P N G K T S A G A R S P T L V I F N K Q Q G G Y R Y D R K F A E N T S G N P M Q I C T E F G S L M H N S
RANDOM_6178 I Y D S V V V F N E E W S Q L E E L F T G L Q C K T W L L T G T D L V P V E P A L G T M L E Q E I I Q H K V A N
RANDOM_6179 V P G K R Q F T L N G F E E H R Y Y E Y N P V Q E M K L P T E A L I F G V N I R L V F N K P E P N T D R Y V K M
RANDOM_6180 A L D D E I H I Q E K T L S S K L E S Y E A K W L G D D L E R G F L R E E T I E N F T Y E L I Q I E G D A V V K
RANDOM_6181 E H W Y A Y G W P A T V Y L G I C K H F H R E L Q R C A N L I H F Q S K F H T E V C Y E A I I F F N P V R T E G
RANDOM_6182 S C G R G F G N I H C V G T V G N T K V L G D V A A G M P S C A V S G W N D Y K S N A F R R P A S G Y L N H M V
RANDOM_6183 V D E S K A F Y G D T W K G Y G H V V T N L F F Y T S K P S P N I A A K D G E Q N R Q N A F R L A T I M G L A I
RANDOM_6184 Q D V I E L E E L V A L V G M T S H A S S R T N N T S K D M A C Y V L I A N K P G M Y E I D K M N S K P D E T P
RANDOM_6185 Q N G L S K H I I V L L E I E G H R C D E K L S I H A V S S A I S G Q N G V F L V K N G C A M P G E G L A I R A
RANDOM_6186 V A E A G K W P K W Q Y W A G L S V L R D I M R Y G L D G K M T I T F S T P F S D W V F A G G G L S R T A S G S
RANDOM_6187 V I V S D Y W V G A T C R T G V H G S R A E F G A L V L E K L W Y L E E A I R A R L M A I S L D K Q G I G K Q Q
RANDOM_6188 G D K I N N K R V P G F E I I D V Q L L M W C L E K N G G R V L K V W R F K A S F K G A T L Y V P D K L A G H V
RANDOM_6189 P Y M Q L A M E L P R A E W L V P E M Q L G N T R A L L Q L G S G A M V Y D K E Q L K W E D S I Y P D P H L I P
RANDOM_6190 Y V S Q S I G T L K F T A K L K Q G G K T S S V G V L Y L T T K L K T N F H S I D D G N G G P W A P G K M V N V
RANDOM_6191 S Y L I M H I E A K S A R L V I I R G A I L D G A G G K A R S S A S I I G A L Q L L E F P C Q K N F K R E T L E
RANDOM_6192 K L T I G A V G A K I E V P V V G Q A N K S G I T L A H A S V L Q L V P Q I K S M S A L P W V N P P A G V P A F
RANDOM_6193 A E P L L A N K T N I G G M P G H N A T C I A K L N N V M R R Q T R E S I R G K F M L N D I P E V N A E I D P K
RANDOM_6194 V L G L V S T G C V G D A D A P P P Y Y D F I N K T T T C S L A R A S A I T G L H V K K M A I G E Q S R K K F A
RANDOM_6195 P A T L E F N Y P N S S A I C L Q N I M G Y R T F A P A R A I S F Q Q N M P T T H L G S L M R Y L L D V S E A Q
RANDOM_6196 Q L G L T Q P S W G I L N R L N Y W A A G C K A L S D V A I L S T N T A V D P F E W E S N I G I V L E S V P L M
RANDOM_6197 R L F G G T G M H G E V P E G T A A S C Y A E Y G E Y A N R E A N A G W P V A E F Q M P F S E Q P T S N C R Q V
RANDOM_6198 R G S F I S A L R A I G V V G F H M T V P S D S P A S V P Y A Q F D M P F C V R E L D G L Y T T L V I H T K T V
RANDOM_6199 A V S S A D L Y V K G V E L E E G K E T R V N A F D E I G R G N V G S A P W V F N D A T A T L S G A V I L L I L
RANDOM_6200 V V A G S F Q T D R L R S P A F L T D R S V R L C K T G D C V E S F E L E P D I N G E A K F P A V P D L L K R A
RANDOM_6201 F A S E P L K G T C N K C K M P N D N E E F K F G S V Y V T T A F G A P N A P E D F F A K I K K F F F R G N P I
RANDOM_6202 Q Y A L R L N G T N S I I E M T I Q Q G M R L P G S P T P S L T P F I Q A D G S I V E K D I T I Y A T Y Q E Q D
RANDOM_6203 P G Q L A P F L A L V R A A K K A R I T H A G L E W D E S L A P M E N E V K V D M A T S P G L D L A L A T P R E
RANDOM_6204 H A N E Y P F I L M K L A L Q V K I I D A A L A F L H P H G N G G D T D D S R H E P K L Q F Q K A M P E P L F N
RANDOM_6205 Q E D M T H R E D S S E N I G A D P Q T A I T V L L L A V E E L R G L C Y S S V A I L S F P I V S P P L R A G M
RANDOM_6206 L S V G Y P A T K I D E H R L V M T E F Q F N R L D V Q C I S K S R A C F T N A D S P E V S N D A Q A V G C I S
RANDOM_6207 L R M V L G L Y D Q V T G L F R E E L V V Y A P K C L T W N K V K D M Y D A A G F P L K Q V L A Q V I N K D L A
RANDOM_6208 T V M V V G T Q Y P C S P D K G R Y L I N D G L I A Q E V V L R V P S N A V L L I L N P E A Q C D G R N S V Y E
RANDOM_6209 M A K K E T Y L K P L R Q R L Q A D H P A P N I I G T I S L I I L Y D A T I K E V D M S L D K C G A L I P Q W S
RANDOM_6210 G L M A L M H A F I L G Y L N N G G R P S Q K L R H I Y P L N V A V D P Q S S N V T S F E V P K L A G F V R V R
RANDOM_6211 Y A L H C E A E Y E S P K V S Q Y M K K T P M Q Y V L G G T V D E L K D K C G S E H R I N L A G N N Q T S N Q S
RANDOM_6212 H P Y Y L L E K K K V S L F L Q R A K K V D T R K G L P T A L A D P K L A M S G I D C D E D T W T V R K G G E L
RANDOM_6213 D D P F N Q D N A W M S L V V Q L E T Q Q K L Y N G A N D F R M D E P L P H Q I K I N T K G S G D A Q A I E L K
RANDOM_6214 G H T Y E Y P S D R G E M V Y G W G S Y F N F R H K I S G V M D L S P T V F S L G L S A K V D P S D D E E K T V
RANDOM_6215 G A A K F W P F Y D S C T T G I E K R P V E V R L L Q T M T L M L T P G N A S F A S N I M E L E L G I R I N A V
RANDOM_6216 G P G F D L D L P V P P L T C K H F T V P K V Q L K K A R A P L I N T T Y N Q T K E L D T E A H S P D R V L I L
RANDOM_6217 V V Q V P G A Y N E A A P S S F T A V P D K E T G P I S G D W E Q R R I R S E G T Q A G T Y E I A P E H K G L L
RANDOM_6218 L E S L P A L Y Q S V L L S T R S D A P E A I P R G S K L C E Q V P K P P N E F D S M K L W V P L K E E P A N F
RANDOM_6219 V Y R S N F T G Q G E K Q A T E R T G A P T I L Q V V A N G Y T S F Q M I A K I A I D N Q C T A K K G E K Y G Q
RANDOM_6220 F Y N C A S V V Y N V G V A Y V P K T D S G D Q D C E E L N T L R I A D D E A E K Q Q N I D P P G H F N N P I E
RANDOM_6221 T I G L I N M S C N Q D N D Q N W F W D V L E E T L S Q E Q M D V L P C N I D T L G N S V L Y E Y R G A P A V L
RANDOM_6222 A T F D N E S P K H A I G P S A T G T F L N P M K D A V E T F N K S I T S W N K K A K S P F L T R L D S S V S Q
RANDOM_6223 H A D M S E P Y S R I D F L A E P A S D A I Q A P Q V E T Q G W T Q K K G D S E E L L T H K S G R L E V Q D I G
RANDOM_6224 E A Q L L L T L A T Q T R M A I F S G A P S N N V G S A G Q P S G K H S Y D L Q S K T K H K L F L T M L A D F W
RANDOM_6225 T H E L Q P R S A E V R M N E T E L A H F E E G V A P L A V L N P S S N F F Q L E W L L M V E Q C G A F P A G V
RANDOM_6226 K S S K E C G I S W T S S I K Q P I G N S K E L E A G L N W W E S R D F H E K P L P S V A T I R A L E Y E S F R
RANDOM_6227 Q S P H E C S N E T T C L R R I L P F D R K G S D A L D E N L Y F K N I M V Q F L T P A G S Y S A V G Y I Q H G
RANDOM_6228 G P K V S N F S E S K Y V R L P S L Q D S V S I N T L K K Q Q S M N I G N Y L E Y A L A Y R E F M R E Q I V K G
RANDOM_6229 Q P H V C E P C R R I H V N K P D V I W W P R A G I E T D L A W A L T L W A T E R A G A K Q A E P R E S D G N N
RANDOM_6230 I P P L Q P S W G P K K R F L R K E I Q G V H R K D K V G R L H K N Q M A Y D D A G S R N Q Q A K Y T D P D A G
RANDOM_6231 L D P A D A D S K I N D I E V L D A Q K K E S G M G I V Q F V V Y I I Y Q A E G T D P L I F P I V G R G L I I F
RANDOM_6232 F N L F S P E D M P R V R A H E A G E Q P Q L T R P A L S A I T N V D P H E S R E R I L I L G A C H P V L E E R
RANDOM_6233 I G N K F A S G D P S L H K E Q R V L V N F K W V R Y S I N R V Q V L F I P D A N N W L S W T L N A P T R N G R
RANDOM_6234 L E L K K A K Q D W S A G E R A V A A G F Y L E I A L Q I I H F C T D T P L T A A H K L S E F D V L D G W Y S G
RANDOM_6235 K E L A C A D M G A R I L A S E N S K N L S L A V S M V G Q S E F A K G E H Y T T N A A G D G D A P W G I R R T
RANDOM_6236 A V T A P A I K L Y F Q G Y E A L M K V V V N D R G R K K P S S V A L T V A S A F N V F I E A E L L F S F K E E
RANDOM_6237 D F D I G E T K N L R Y R P E A P A K F K K L D R I D F L T A S L S F N D Q T Y A I T L G P N P G L A Y F G V L
RANDOM_6238 P C S L D D V P K I I S G I I G N A W L E G V E N Q P N S I G F S V H D D D I D P V I L V R A Q G L I A G Y R T
RANDOM_6239 W S I R V S T R L K T I S E S R Q L L P I F P A I N P D L I S V Y G L P P L A A V K L A R Q N S W L S S M A S P
RANDOM_6240 E A P L D A W D E C Y P C I E Q M K Y K E S A D M K R I A I W N R A N H Y N H E D E K V A G M K G D V G Q D K N
RANDOM_6241 L G L L K V T T V E Q G L G R A L G D N L L A H P I R K A I V Y E E P W M R R L V Q Q D F D C Q T F I P T I Q G
RANDOM_6242 L L A L S Q R K M F E V S H W Y G D P K P V N A H L D V G D T E G M R R G D K H P K T V P P L Y D S A N A S M A
RANDOM_6243 V P P A R S A N Q G D E S P I T I A Q Y F E V Y E T T S E L A S S E G I G I A T F V A V V D K R H L R T Q D S V
RANDOM_6244 A D N Y V S V L V P K L Y D E H K L A L A G K V Y G L S V E W A N D P G G G A G D C K K D L Q I S M D D L V A S
RANDOM_6245 L C I V L H E T Q P V P S P G D A T N S K N A E Q S E L V N C W P A S P L R A L A S A L A Q L E K I S L Q L K R
RANDOM_6246 H S K D Q P E F E K R K N V C R E P S S L G S L S Q Q R S A K A S L T I L E H R M P D D L F V P D V Q L V S R E
RANDOM_6247 E L Y G A L S L E S T E T C A D E G G K S A V A P N P V G V S K L E Q S V K C G K G V Q T K Y R T E F E M V V L
RANDOM_6248 V K D P I H P G D K R N C L F W Q V K D H F I L S L D K S T F R L S L I F V E R L D P E V K M E R Q A Y W T P D
RANDOM_6249 D V N F G E A Q T L F H L A Y F A K Q T K N L M E G H H Q T K Q P F R E D Q V R I A K D H K L E F G G F K A S D
RANDOM_6250 Y Y V R Q L A H F D I F A P R Y K A Q E L S V E P E L R Q D V L G F N L L K D K R V E V M S G C A P V D D F Y Y
RANDOM_6251 A N M G A A L L W Q F R S E V D L L E K A P G H V V M R L W V L T D F Q G N P A A G M V S E L F D S K Q I Q E H
RANDOM_6252 L W P E A F Y E L A P G W G T T P I P A P A D S L S A K I L K P K P A Q I R N I D R P P A D Y R I H N A V V S A
RANDOM_6253 A F V T G G F A I L T W Y T T K S L I S C I L A G W F P A W S L A L L A M G Y M Q G A F P T K D D D G G A R K G
RANDOM_6254 S F G L P I V Q M K V R R A M G H D V V Q T G V N L E G G M M Q N F C G F C N N I I Q V P L E K G Q I L N A F N
RANDOM_6255 R G E M E D M M L R L P A L G V M T S N E T T F L E A Q R W S A V S D K V S V P V L F V V D E A R L K A P S T L

RANDOM_6256 R D G P K P Q N K I L G V E A S S L A Y I N Y L P E Q A G K T H H D S E K S V A G Q A D L S N H R R I V E L I H
RANDOM_6257 Q L L V D R M A P E Q Q H F V S K G N K H I Y H S L K V P E R K M R E G K K Q S F K F D Y Y V Q M S E T N I K I
RANDOM_6258 N G L A D G Q V I Y K P G S R Y I D F A T L G R N E F F I M D G T Y I G D R L L S R D T T R M Y S G T C G A D V
RANDOM_6259 E A V I Q Y A D K Y A G C D N R L W D A D Q I Q H S D T L T H M N S P G Q F I T N V S L K P A N S D L D Y N S S
RANDOM_6260 I E R Y A A Q L D E K M T A P E K G I S E V F R D T A P I K A S K Y V D S L V N V V M S R D G A V Y M V E W G K
RANDOM_6261 P G A Q P P G M P S S K A S T E F L T N G V I H G S L G S E A Q E R G P D M A S R I G G N V L G A L Q T E P L M
RANDOM_6262 A G N C R D G I T G E D F F E T V C T S K S W G A R A W G T T V G E T T T P A R K S L S Q N K M S W E Y V P I S
RANDOM_6263 P L A I A N H L S V A G S V A T N V I T V P K N T Y I S E R D I S L S I V T L K A L C G Y N K V L A E H L R Q T
RANDOM_6264 A Y D D T S Y D S G F A R R A T A R H H A E E V R T D N D S T V F L K S I K R Y V M Q V E K H S S A S T P F E Y
RANDOM_6265 K N L P A L G V S Q K D L K D T P G I L L T L P E P I P F I D M D I F S G L L H S P N S W D Y Y G Q D Y G N P N
RANDOM_6266 F P C G S K D D L R T G Q D S H T T L P S T M D N N G D T S H S K L L A L E K L Y D H K E M E D P S S G N V A V
RANDOM_6267 P F W S E Y K V S A R N K C C E T Y G V E S S S V Q H A V A S K D R A V M A Y S L E Q E G R C S N I P F Y E A A
RANDOM_6268 M V G C D I M P E E V Q H I T D V E Y I T T N T A L A L R V S E R Y L G L T P E E L P Q S V F V L L D A S L P I
RANDOM_6269 A Q V V P R S G S T N P T A L K M N R S V S N R G S K R N I K E V K R G L N I A P D S E K Q L C F R L H I T Y S
RANDOM_6270 D N E N P V E P P K Q N C H T L Q M A T S D A A I A T K P R P Y F R Y A I Y V A L C I K S V D G V C V K L L A E
RANDOM_6271 S G G V N G F I I E D V Y G W T G T A T S Y E Q A C A N V A E Y V K P A I C L D Q V R F F A N T M L L L P L T Y
RANDOM_6272 Y S P L R D G L L A N D T N E M K A Q L A A S F Y S W A V V R Y E E C F N R T D L Y V N S R E T A V Y G G R V E
RANDOM_6273 I A T R V V D K A E C A E Y R T Q T I V V A S C H I E G E P D V T R W M S Y F D P I T K Q R K M G P L K G R R L
RANDOM_6274 Q T V D G I H Q C D K M L P W G L S Y D Q M A S R A K A A L R Q S D R I R P N E A D V S A D V A K A P W T V S Q
RANDOM_6275 L P P V S K L C G W E E S R C C S W A F V A E V L D P S L G C V S Q K E A S H P L G T M R D C L L N Q C F A D Q
RANDOM_6276 P A S S E P W N D T D I D A L L L G V D D I C D R L L G A A E N T K A L V A H L V F L N G N A N G N A D D R M P
RANDOM_6277 K F L N Q F L K E A Y R V K L S S N M A E H S K V I A E P V Q V N G P P P Y P Q G S G P E V S A K R N M E T T S
RANDOM_6278 F Q T A P V A V I D D C F T P G S C M G H N K Y P D K R T F R Y I P Y S F E S T N T F L I A E R A W W S L S I V
RANDOM_6279 Q V A I H W G V D R N D K Q N L A R Y E L I L P V R T S Y K E E F N C K S T R Q D L Y R G M N T K S T S D Y K L
RANDOM_6280 R G G Q M N D S K D R A R F L N E E Q V L T N V G D G A N D L E S D K V D P F Q G E Y S H N F E V D S E I R D D
RANDOM_6281 Q F Y T D W H E K I C L L E Q L E P V V Y G C P C C V L E L Q E G S T V Q K N A L P A A T F D R V E E T Y T N N
RANDOM_6282 Q E F V D A C I L Y I E F E H W R E R E W L H F S S G Y G M F F Q L V L A Y D A L E K F G H D Q S N K S S T V D
RANDOM_6283 A G T H I N S E K C V M V L R S I F E R T A N V K K E D C E P N T S M E A D Q A S F G S A L T L D E A K E H T A
RANDOM_6284 A Q V S V G A D M K R F F M L Q A D W V L T T L R S P Q K Q L G G Q D I E S G L S Q R R V L Q E E L H Y S A E I
RANDOM_6285 W S K S P N V W A Q P V V P L Q V E R L D V H Y Y T T R L K V Y Y V A M L K T A S L A D G Y S L V S P N A A L A N
RANDOM_6286 F V S A I V I E A G G K F N E C G R I P F M S P S E V S M S M P S R T G V D F L N G S V V G G K L Y Q A C M G P
RANDOM_6287 F S G N E V L R R S W V P P Q F I D G S V G H G T E W P A M V S K D F R S Y T K A Y G L C G L S R D L E N A L S
RANDOM_6288 L K E F S R L E L L Q A A T E P G F T V E G I F G A L A L T A K Q V D P L T V A T V R A L R F N E I G L E W C Y
RANDOM_6289 R P G T I L Q L L S I S N D P S G S V T P P L E S D H L A V G E L D D A S E F K G K K F H D L F P G I S N W Q E
RANDOM_6290 T A S G C D P H Q M K V K F I A S P W C V V R A P K R K E L V P W T V T Y S Y V S L E F E G L P V I V N A Q R V
RANDOM_6291 H G P G K I V P D Q G L V A D D S S G R D Q V E D H T A K A R F Q A N K R K D H L E D I L P K Y G R A G P S R P
RANDOM_6292 Y R I L D S D L A D A H G K T G L L G K R G G D V T L D A N K G K D R G A L K L S L M M Q V K N N E G S I N K P
RANDOM_6293 G L I V E V S L S T N P P E I Y E V V H L K A P P F K L L T G K T L R L G V R M D S G K P L I R F I F T A G I I
RANDOM_6294 M L F L S V N F E K V I I P M V S F R S M R L E F Q L I K Q N V G L A N T S I C C K L V S H R Y P G E L D Q L T
RANDOM_6295 A R C P E F R W P R P R Y A T Y D Y V L S A H L D K L F S K K A K T A I Y S E D V I S V V R A E R Q I V L V M V
RANDOM_6296 F W P K N T Q R S P V T V A K Q N A A N Y I I P Y E Q G R D V A L L D E G V K P A D L A E S L P V Y Q M R L T G
RANDOM_6297 S V Q S N P C Q R S R G A S N G D S G R V A T T G I Q L G G A H I T W M S A V A Y Y K F K Y G E N C R T E L D Q
RANDOM_6298 A N E S P R R I A K D D Y Q L D P K L K S P V A H G E A S A L A V R R K L E L V Q T K Q R C E A S L A E T E S I
RANDOM_6299 K D M L R C E L V L V S I C G D E V N Q S A Y E S R V S I P G D K G G E Y G G G L P A I G T N L A V T P S M C D
RANDOM_6300 L E T Q A G V K P A I D K C S T L D R N E V S G K G G G R S I D L V I T G I E E F L Q H G L C S T L G C K E A N
RANDOM_6301 V R Q I F V S S T K K L G D A R N N R N G H V H V W Y V N E E Q L N V L L I N R A F A E G I L L R L P Q L R Y L
RANDOM_6302 F E K G V D L K T N N F I V G V R Q K L K H R A D E E V N L N S R Q M L K G H Q R C G N V L E C H A I D Q K A W
RANDOM_6303 I K K G D A G V A R P T E V F F R V I E I Q A A V V M I Y T I M W T K L E K V V T S D L R F I K E Q V S A I N S
RANDOM_6304 A T N N F I M Y G K P Y F I F W K H A A F C G T E T T P I L K R M A E E A L G A N V K C G G E I S A N I N P I Q
RANDOM_6305 S G T L R F T G A Q D S Y V M N A D T D A Q E T I P A V R I L G G V L H G S A G I V R G H Q L L K Y R P L P S
RANDOM_6306 D K E S L L A R P H M Q T L E E F H P N E D F K K M G V A E I K T V R D S W S L N M V L L N S T S F P I K K F N
RANDOM_6307 L T L W L L L K L E G L I R C Y N C G I T Q P D H L G A L E T D M V T V G S L R E L F I H G M A V A P G L L A V
RANDOM_6308 V I P K G G A I L L E E K G L A T S A S W I H F P D L L I D F C R D S R C M S A Q E S D H G L H T R S F Y E A L
RANDOM_6309 V G T I C Q D V A F V G Q S K H T M W R S R E A H P C E Q P E K G L N S R Q G L T A K T G V W T P V T E S A T K
RANDOM_6310 K E H Q L F P Y L E S Y A T A D G E V K Q E A A G S N H S L D V D D D A T L R D V R S S K K D Q S G S G H K L A
RANDOM_6311 A I E L V P Q L L L G N T F L I L P G P R T W L N H N V L F R D K A R S I P I V D S V A S E G S L I R Q I E D P
RANDOM_6312 E G S Y P V L V A K A Q D I T D A E L F D M S V L G V H S E V M T A Q M T Y A I K Q M C N E S D V Y G P E A H D
RANDOM_6313 M G L R A F E M R R K E L S L G K D L R V I F V E Y W H G H F L K K F F P D K T K G V T K W Q Y Q D G T H V V T
RANDOM_6314 V V S G L G H Q M E W T Q S R G Q A K W G S E Q T K S L Y L I R Y I Q N Q Y A G E I T D K D K S Q R D P Y Q E D
RANDOM_6315 Y K Q L Y T S S D Y L L G H K P E H T R D L M C L I G E V A L R F L S E V M S V F E A L W L G D D E Q I L K F D
RANDOM_6316 M N L Q G Q I K T S A K D L P T M E E L V L D I K G I S K A V E P E T R L E S D S V N S I M P A Q V R Q V G P G
RANDOM_6317 N S R G K A A C H R A I T V I T V A P L V I E F D A V R N Q E R L L W N T P F T D Y R Q R S V A V A L Q T R T C
RANDOM_6318 V Y Q G G A S G D L S D G E Q T G I R I R E S P G S D M D L L S I I A W W C K S M S G A V S T T N K T S E L T D
RANDOM_6319 K T H P V E L E T I F N Q Y Q E A K G S R G E D Y K S W W G F A P L R R G M M H G I D R W F P D M Y P P N T A F
RANDOM_6320 N A Q F Q G E Y P Y G S R A R G Y D D A K M P N K L A Y M C N T D A H A R D V V G D S F P Q G S E I R I E G A S
RANDOM_6321 G R L P L G E D F G V P T Y N V A P S H N W Y N Y F G S G P Y G G P R D T Q A N M D H N P I K T P F Y T Y N H C
RANDOM_6322 N V V Q S G T V A L Q Q N A R P P V E D R G Y R D T S E E K A P Q L D M L L L A P S A Q A A M Y V Q V V E L P H
RANDOM_6323 Q L V E L Q T G I T S P E S V T I G E E S P T A L S W V A A H G G Y N C F V C M M I N V Q I S K P L Y T L S T P
RANDOM_6324 N N K K T P V M I T E A F A Q S D G N N S G A T E Q A Y D R K V A F E G C P D P R Q F D R N E V A A Q V G F N P
RANDOM_6325 V E L K I A C V A I V C L E F A A N Y S S H I D P T L L S D N F L H R D N T L H R I T Y K S A S E D M S A D D Q
RANDOM_6326 S Q T D S K I T D L S T L S E V L N I S N L P G A F L G A A K M S S Q T N P F A E L L G V A M D D G G K A Q Y K
RANDOM_6327 D G T V T W G D S F S I E F T R K T A T Y A Y Y P T V D L P S K L M P P M L M L E W D K L A A G H Y K P Q N S N
RANDOM_6328 D R M N E A V E K K T D V G L A E G C L L K Y F I R N A V T H G A K K V G A I P P V E V P A P K V S S R L A E V
RANDOM_6329 S I Y M A L R L S F M E P D Q G H S D A F I Q E E S G R N T S A E C R D E S F V V I Y Q R N R G F V Y N V E E Y
RANDOM_6330 Y D D S Q E T A G R R L K P I T L P G A K C N V Y L G K E F T M R A L E T V G H G K T Q G K L A E K L D C D P L
RANDOM_6331 F G A R T C A L F E H R E P A S Q G S T D Q T K R A N L D A V G P Y K G F E S T D N G F S S C A S K L G S T T P
RANDOM_6332 V L H T H F F N I T D C F F C R D D A G N K Y G E L A G D T M T F A L H K K R F K L M I T D G C E D S L Y R I A
RANDOM_6333 T A L G T A E T A R F R R Y E P Q E L Y F T L I P D N A F I I D S M D H F V A V R A C A P V V T T R G V L Q Y E
RANDOM_6334 G R L R Q E L Q E P L M L A E G D N T T G H L V S M F Y G N K M M N E V Q R G A R R L L M P K R F A E N N V V G
RANDOM_6335 K G E A L A N L T D S Y F G Q A F R M G G A I Q M Q R N M R P S L G N L R K M P L A R R F Y E N S H I V V P T Q
RANDOM_6336 H S R H E E V K A D T M L G G K A N E S K N L K W S G H D T T P K Y D C I V M F Q R R I P I G Y Q N K R V D E E
RANDOM_6337 P K C F R G E V V N L E G P Q S K D I Y T V Y P N S E A G E I T N T D D K M T K G R N L D G P L M I D M A L T L
RANDOM_6338 S A G R D R L G E E S P R G K S K G T P I F L W N V L G G T G V Y L I L Q I P E L L R S A S N V Q G I S I I A C
RANDOM_6339 A G C S T A L L Q V F E W D D L K Y H S S L G S V P E Y S A N Y L S L L N I R L P A T G A I N T S A S H G T N A
RANDOM_6340 R K T P V T R T D V A N E A P K L L Y S K G K D L R T E R L S N V S S D Q R K G N F V A G R E A I E L Y W D R K
RANDOM_6341 A R L R G S K L C N N L N I I L P N A L P E P W P M D P F A T I S R T P C A V V E D V Q K Y A I K T K L S I D L
RANDOM_6342 E L I D Y Q V N K C V I I E P A F T S A T L R F N Y G A N N G R R Q K R S P F D K W R K V Q G T T F L K T E D G
RANDOM_6343 T L S A Y D N R P G L K K N L A D G V K S K Q E R E L W V A H Q T G Q L L G S L Q T E K T D W Q F R N V Q I D L
RANDOM_6344 W L A K R E D E Y A E E L F I W E G K T R Y A Y R I T S E L H Q V L N V S L W T A F Y A T N E A E V P N C K A I
RANDOM_6345 A T R F G D E V A Q S L A N R N R C N S F G T W P S I V V A E Q Q R P R M T P Q T G I T A T H L H Q R Y F A M A
RANDOM_6346 L H E A L L V I P K K P S I G V R T S N L I L Q L G S K L V V E V V D H C N S R N R K L N S L T R Q S V I G L F
RANDOM_6347 T G D S D V H K I L Q D Y G Y F T A I H Q P V K S A P K S A H Y Y T E D V L N E T N G L T S K G V D E S K Y S N
RANDOM_6348 E N D A T K Q K E W S S V A G P G T G P Y E A D P A S R L R I S Q L T T R Y Q L K I L I E F L A K S A N S L A M
RANDOM_6349 V Q C V I F R V Y N T T H A R L E W S M T A I L W A V Q H G H L A G L E T L V Y R L W L A N W A T S T R S T R L
RANDOM_6350 E S T K H K Q S V P R D Q L K G S D D A A V Q A E V L Y P A K E G P S P R D D T K L Q V H E Y L S L T M T I V A

RANDOM_6351 L R P T V S W S R P G F Q L S D G K S Q G G C D G P E Q P P T A Q R V C E N P A P T Q V F T K S N Q K E L L Q L
RANDOM_6352 I Q Q K R L W C T L C K S F A Q F Q D H R E K R C S L W G E P M D V E Q F D G A K N N E H C A T D G N M E P L G
RANDOM_6353 E T L G L R K E G L G T Q P E L K S K F Y G P S V T I Q G I K Q S N G S V G P V A S L N E H T G P C I L L S K R
RANDOM_6354 V W T Y K D G S S D E G Q T S C S H L D Y E Q L L T G A S D K Y A I I F T K M F E L Y D S D Y N I L D L G K G V
RANDOM_6355 L T L R S K L A N R S L R E F A V L D T N D I R E T L I H F G G N E A D L Q G N L W K E L S G M E G S N L A L T
RANDOM_6356 P L I E S L Y N L S Y R T Y C A L E G S E E L T A E K Y R L H K S L N N K D N T K V L V S C F D T V E S V T P M
RANDOM_6357 A N V P Y G R D D A S N V G L S H K N S P P E W I I L N E E G D A F E D N F A G V F L I Q S Q S R M H L E A P F
RANDOM_6358 T I N V Q E E D T I L D M A L D M D L D A R E Q M G C M S H A K F A A N F P I D H A Y S I H D S S R D R N A K D
RANDOM_6359 V L E T L H I L S E E K G N A C L Y R T N P P E L H T D P N K A I G T D L A K E R Q E T E L P S I P R N T P N E
RANDOM_6360 S I D S S L C P K V D E Q T R G V S R Y T E S K S M S P A L S Q P V L E G K Q R T I I L I L V P H P Q Y A N D L
RANDOM_6361 L Q F D P F N P I E M A N P P M D H P E Y F R S H L G C K G V D D F V G N L R Y S I S V E D T A S R C D S G V V
RANDOM_6362 E L L H N C V A T Q W K L T R P L N R G R V L V G E L N C D E T Q S L G L I V E I F K H E K K S A N S S S D G R
RANDOM_6363 E P K T G P N T T Q S T P F T G K G T L I S H L D K Q I F R L R S Q I G N N F P N S S A A C D I R I S C Y L E G
RANDOM_6364 A G V S K F M I F S G G C L K G A K L E T C L D F S E T M A E A T P T D A S A P C R R D D G T N L F Q S S N G H
RANDOM_6365 N Q K G I E L G C N M V G C Y V D E S H P V I F Q R K L S H S M S A F K E P R G T V A G Y P R F M P I S N L P G
RANDOM_6366 T E S P G N I A P D D K M P K H K S K H L A Y P T T S L A E Q P V P D Q Q G N C D D Y K P M F Y R V D Q S V V A
RANDOM_6367 F K S N G S P K W Y V M Q L L G G V I V D M E T V S T S S M E R A D F S R H L E R E P I T N R P L K Q V T E T M
RANDOM_6368 P S V I S P M V E C W G N A I N K K G T T A V F G I S D P A N L A L F G P L K T G P S F S Y K G C A E I K G I I
RANDOM_6369 P E L N K G P M G E F C T A V A D N H L Y L G V C V S T Q V P G K F A I G A G I S A S R T G H P T L R T D L K A
RANDOM_6370 M V Y K V A Y S H E Y I L V L K V G N Q G R G R P G T S T K A K I R I N M D S D E V L A L A R N M E Y P G H N D
RANDOM_6371 F S V S T K K A D P N R R S P H A N H R A T A Q V R F M D G I H K S W G K D H S N K L D P A N N S G A T T K A L
RANDOM_6372 E S T Y S Q T A T T Y G G G L E W T E Y A G R N Q P S T K V S Q K P E V N T V C Q N P L K K A E G D D K E Q N A
RANDOM_6373 V G S N L V T A Q A E N Y G S I D G A V T V Y I G K G K C V G S E F V F H Q V K D R V L D N F G Y F K L I D S T
RANDOM_6374 G A Q F L K I S T N L A Q G L R M P T S T F G S G A T A N L N I N T V G V G V A D A A I K V D P L N L C V N S N
RANDOM_6375 M S I T Q R V A A H A M A D Y G A Y S N T Q I T L P N G N L K A D G V R K K S L I T A T R A S C S V Q K D V G D
RANDOM_6376 F K E L N F T S A N S A C R D E A R L S Y F H S Q T N G E A N S L P D D K D D R F L D P A D R G S I P A W L G N
RANDOM_6377 A L D E L L V I R E V S A H G K F L K N M W T A G A T D P T P R G H P T T R H K V G T E M M P L L L T A L I S V
RANDOM_6378 K S S A Y Q F D Q I G L S Y L R F G R E G S M V N S D S N R R K L Q F G E I L D N V L L L K A T A Q Q T S Q V R
RANDOM_6379 K L E V P I W K T T A I L E F V R M L I Y C M L V T G R C S E H S G V Y G G K A T Y A V Q F V G I R L R F Q S L
RANDOM_6380 Y R L N K Q L Y T C S R K D L R W K K V K G Q I F E I A S E L I G N P I I L Y L R F G R P N V S Q N F P I L L L
RANDOM_6381 G L N E N A H V A S L P Y T S E S K V P Y V D G A A P K P P P F K P G L V C L S Q R T V L P R P S T S G V R Q S
RANDOM_6382 R A C M P H L I M V P A D F L Q V S Y G V N D A A S S H C G L A P P V Q P H P N G Y R M M L D V S L M G K E N G
RANDOM_6383 V D R R E T Y I P Q I V N N K C S G G G N Q L N A N I Y N R V M Q N D Y F S I L F E S H P G D Q D G E G N Q L V
RANDOM_6384 N L I M N G P C A L I S F T T L L E V L D P P I Q E C F L R G P P N F G R G R G C C G A Q L G L M L A A K A G I
RANDOM_6385 E Q A A W I I E V E A F S L S T P C L D T S L S E I H Q K V P S Y L N E L N T Y S I F G G D M P S A R A W P L G
RANDOM_6386 S D T S N F L A S E T F Q G K K T Q F Y Q F S T P N S V I G I K L F P R V N Q K L L K V I F S P Y C K D Q V Y H
RANDOM_6387 H L G M R Y N L D T T A A H E G Q L Q S L T A C N L P E D T Q M D R C L R F N C D L K L I L I A H Q K Q K P K F
RANDOM_6388 N A V N M N E I K D A L P G W N R D N G Q P S S A E M R V A Q E S D D L A D L N M T L P F D L T Q A N L F P L R
RANDOM_6389 S F Q R G H A T H Y A G I R Q P G I H K L C T I Y Q I A S F G Y G R P I E Q G R P Q T F Q G P R P A F S F L M P
RANDOM_6390 H S M P N E P T A R L Q T G L D L A D Q N L W W D T A A P F A L S S S R A K E R S K P L G A K V F V A Q L A N Q
RANDOM_6391 P T K A G A M N H E N L Q R I K G T S M E F N L A R W A E G E M E A S S I I I S P Y D S G E G C Q D D R R P N K
RANDOM_6392 P A A L I D K V C K I F Y L A Q I Q D K E E S E R P H V L K C N C R V V R R G D L S V S N F S W L T D I G N G I
RANDOM_6393 T N P V T S S L T V L P E F M S L L A Y Y G K N P E R R L D R L K D A G G P T T C N T Y C H T R K G Y N R H Y R
RANDOM_6394 Y Y E N I L Y K Y G A A T M R S S C S I N Q L E I P V V H D S C I I G P F D I L H V G K G L T W K S Y M P K F V
RANDOM_6395 A M P R K P Q E G A A F A G K L Q V P S V E S G P V V D A D F P S A N L F S L L A E V Q C A G K T G A K R R T N
RANDOM_6396 L P S C G V E I K E V N A G V E T W G E Q K L K L V H D A A L D K K A T Q R E P Q A Q D V M A Q S T E T Q H R I
RANDOM_6397 G A K M G H I L W E Q D D L I G G F F T H G D H L Y P N S E T L F T R L E Q G H R S C M S T D S P E Q A E T G A
RANDOM_6398 R V P S N G E Y A Q L V S R S G Y N T R N Y Y R S A V I P S G D E E I K T D E R Y R Y V Y V T D T G D L A S L Q
RANDOM_6399 N I T D G S G G G G F L K E I I S P E L G W N V C G G W D G Q L D V D S E T S H L Q T Y L C A I S D I D D Y D A
RANDOM_6400 F G L S H E D C Y A N N S P N E V A V I V D E Q L V V G V V T I N G I A T A E D D D F E P P A Y D E C P A S L E
RANDOM_6401 L P Q T S L N E L V Q A T L C A S D V V A G L I Q S G Y D N H I I T P N A H G F S D V C K H C C L V R T E L Q G
RANDOM_6402 S L F N Q L I G K I L K A L G A P A Y F D S P G T T P A A S I E L Q I E A I V A G N D L A G Q S S N T I A L G A
RANDOM_6403 W F D V Y G Q H D T L T A T K C Q M Y A K N G D P K P K Q Y G Q E Q V L A H S A G R V T S G Q S G A E L S G K E
RANDOM_6404 S M Y S C I P V L S D P P E P P V D R D A T S C G Y K V T K T E I V L P L N D E E K R M P I Q Y Q A I A K W E L
RANDOM_6405 G C H T I G D E G R L A Q Q N V G G L V F T T P L T T K N F I D A G D L K N D C E N A N F A S D V A S L L V D L
RANDOM_6406 N T P Q Q R V H L P I A V S V P H S D T V L Q Y L L L M E V V L N A E V Q L L Q T V A E R G D V L V N E I I N T
RANDOM_6407 A T P R G K H H T L S G E L Q V S T S S H T V I T E I P V E P D F V F Q T K K Y N F F V E H E Q A L V I V K R K
RANDOM_6408 L K I P S R L Q A V P Q Q N N N L G K A T S G E T M D K R P T D K A S K A I S L K G A N A M D E T E L E G G K N
RANDOM_6409 P I E Y D R P N V L V G L N L A M Q L L N V H I G K V P T S G F E D D D V G A D I V N N D W G A E P V S N K K D
RANDOM_6410 D D L N Y Q A D R E T R T D P E D S G L G D Y D Y T G G A R A K P T E R L A F P I P A Y A Q K R E T I E L D V V
RANDOM_6411 N R T V Q C Q K S F F E D N D D P P S E S I S Q Q T C A A A S Q G R D S C T V Q V V N E K N L S V N K V T L V D
RANDOM_6412 V D T K G E N L N Q A I S P E N L E L C E G T E L P E G E L F M T N A W L A S V N K L N Y K P D E F V E T L S V
RANDOM_6413 R V K D N S V F S R S G Q P L V W V Q T E W D Q A P E T D P G M A K V G S K A T V A A V D T S I M G F D D A G N
RANDOM_6414 V R L V S I N A A F H K N R V V M S H T E V I Q G D E F A A F A E T S L L L L H T F V K E F T E T D R L F T L N
RANDOM_6415 E V E R A I R L R S L I V F K S P E N G I Q V R V K Q L N C G V N G K F Q S E I L A Y F N R A K G R A N E L N I
RANDOM_6416 P Q V Q T R S I K I K I G E A L P K I N G H E Q P A T T S Y L S N T F F T H W I V N A S E A Q E L E V E F N I V
RANDOM_6417 G F A N N S I D L V G G R P G Q L Q P T L Q A N K D S L N T F R F L R H G E Q A E L L T E P M N S P P E C A K L
RANDOM_6418 K D S L T R G P D T S R R D I L N E S E G T C K Y E A A R M V R K A V E G C I L Y T A G R G N D A V D L C L I V
RANDOM_6419 G A Q Y A N P K P T S R D E A G C P Y E L E L Y N E L A F A H L T R K F T G I G C A E A V F L V A S G R G V R S
RANDOM_6420 I A S P G G Y P K L R K N N R G A I R Q G V S M S C D F V K S E N N M T Q T K E T E W F R L V T C E I I Y F M C
RANDOM_6421 G L M R L T Q A N Q T E L N L I L A G T V V R I C E L R S P D A P E H N E I H A Q V K A W D P M E G R H L S K I
RANDOM_6422 Y L E D V R Y Q L M E E Q C L V W Y K I E Q V S T N G A C K D D A E Y G F T P E D W Q G P T T L D N D S A L L K
RANDOM_6423 C K N N K P K A A Q G G Q S S V D P P K F L P S T C A G A S Q R N Q C E Y M A T L Y H A E A G R F C F S R T L S
RANDOM_6424 S T W G S A M E N F I Q A L F M K K G M K E G G K L A A L D E A G N A L N K L E A E E T H N D E T G F M N P Y I
RANDOM_6425 Y K P V R Q F W Q N C N D R K D G N G A G A I T K T E R D K E Y E K W S I F A C R C A Q S R A F K S A H K L A F
RANDOM_6426 L S Y S P P I H P G N L P K G Q S Y G Q K S Q L S R Y A D P A K L G K M F M T A F V L G P V T L C S A I V L A K
RANDOM_6427 K A D V V S N I S I Q V V A V N S S K I E L I K G S P M S V R P C F T E C Q N A R F I L S I L R L T Y N G R S A
RANDOM_6428 N L G N K A I N Q G E F I L D M I D T L P A Y R P Q F K D I E Q S R R L K P A E E D L S E G K V P P F G F R K Q
RANDOM_6429 K G S P L D V N L K G T I H A T S H E R L A D D P T V S I E E K P E I N M A L S L C T I R G R D G A Y G S K T S
RANDOM_6430 V S V Q C K M N Q Q A K K L E S N M Y Q V V C M A L S I S P E V L I W A I S A P N T G K Q I P S C G P A D S P C
RANDOM_6431 L N T N K R H V M P S F E S F C W F I P E E T I C A P K S D R G A R R W T V L Y A D L K V E S D E T P Q A K I L
RANDOM_6432 L G F G D L G K P L P A A K I A P S Q L F N R L H V R I Y G R G Q S D E D G L E D P S Q D S A N L F R D R N P V
RANDOM_6433 T V E L V H A Q W W H V I A R A K T G R Y N T Q T S S S R Y D D D T W D R N T Q Y T N P P Q V I P I K R D F D Y
RANDOM_6434 I R M Y G P N V P Y S L P R M F N Q N E T P K F E W V A A V E P V T N K R D V N G D L L A D A T I E E I I T E T
RANDOM_6435 A C L A N P E T H G D F F S R P A A A K K D N A L H S P F P V Y F A S R F C P V A Y S I H Q C F L G G L K R P T
RANDOM_6436 A T Y N C M A D E R S L T D N S L S N E K P T N I A K Q I K F I F T S Q A P Q N I R E F T S Y E P V S L L I E K
RANDOM_6437 V D L T L K I N I L F E S G V P D F R G K S Q S G S G H S F A E L E Y K L V F I D R A E P L D R C A L T K E L I
RANDOM_6438 A T I E E I K Q L S D L Q F A D G W F Q T K A E P I K L S G M K M N I A L A L S V Y S N L S E Y I Y N V A V A I
RANDOM_6439 T T D H C Y N P Y N T G I G K L N G R V I R S L A L Y F C C Q Y F P P K G A S F V C H L D N S N T V D M A E S K
RANDOM_6440 I G T K N L E M G A L F Q A A D I L M E Q A L A F Q M N V Q P R S V A V A S E G D W F T E F Q I T L H H L T V E
RANDOM_6441 N I Q A C D C P K A M H A A G R F D K L V V V A D V I F K Y P S D Y I I S N A S Q N N Q E G V L V T A A V A F A
RANDOM_6442 R F E Y L V P E R Q V N W L L K R K A E R P N M S V I R V L Y E E V A T E G D E R T S A P L G A Y M G P K N W T
RANDOM_6443 L E P F K R S W E D K E C V A Q D L A D L D V R Q M L E L R V T R P E H R E R P C V V I A S I S N A Y G P K N L
RANDOM_6444 D M N P S T N R S P Q D D E L T K M G D F P Y Q N L T A P Q I A G R K V L P L L T K S L L I Y C P Y T D L K G L
RANDOM_6445 R E T G G Y A K N A G I A A T N Y N G C D F D K N Y L R S L L Q G R D W D S C A A I V N Q M T K P G I L S S S L

RANDOM_6446 G I A T D C P C G N N M P V R L V G V G F F W L G D I K G T P A L L W K I A P G H G H I D K V K C P C G P K L R
RANDOM_6447 S F M L D L P F Q P Q G D L A E K V Q E E A D T I E I W F D L L R T W S S V D I G R L Y E D L R D N D R A N H A
RANDOM_6448 A H Q F K A K S W W K K F H G E I S G E A V I L E A D T F C L S F M K A P Q M K Q Y S P P T A S Y S G G K G Q E
RANDOM_6449 D Y R E S A I E L T C G G S C N I G A S E L S P M Y E S V R D I S F I L V L C V R L R R V V G E K P N K E Q G R
RANDOM_6450 N G Q P F F N T Y E S G K A G M A L A N K P D F A E Q L L S P R G L T V T A M G A R P G T A N F F P Y E M A G L
RANDOM_6451 A Q V G G L Q V M P L V D I L L N W Y M V F D G K F S Q A G I G P S L H R M G A A P S Y L D R P V N H N K M T T
RANDOM_6452 N G N E T D L T L V S E K K G R V Q K E A A F Q Q V I A G E L A A E P N V G T P A E V N L R V V A H S V V R T P
RANDOM_6453 F G E S G R W G I L T K S S Q D E D L D L Y Q A A L P Y P C L E F G G T T E T T D I N F H E Y E L R A Q S V G V
RANDOM_6454 F N L S A L C E N F A T E N H L E L L A F E V E I L N R P I M Y G T Y K R K T S L H G S C I I H S L K A G E K A
RANDOM_6455 I E S G L I E K Q T E N Q K V Y C E N L R K I V P K C Q C R S Q G T K V P I P K T L T N R K Q S V G F I G S G Q
RANDOM_6456 V L T L C Q F M N T L P I A K D Q A C G C K H M M N V S W S L K H T A D R L P L I T P I H E K H L G L I G L M V
RANDOM_6457 D G L L E D A N V Q P M P H L E R C V G T P N A T F T E V A F Q G R K G T T D E G S T H A W L A E D G C E P T P
RANDOM_6458 Y R R A L E P G N F K Q T S A P P P P V A A D G R A I Y E D I S S N K G Y G R V L F A S V E S S D L L V E N E S
RANDOM_6459 R L D F I P G L N Y G G Q G N Y N T L R A V E R S L D E L G Q L D S L Q P N A H Y A H V A L F T A L Q V F I L E
RANDOM_6460 G R C K T E Q Q L R I T L A H A T N R E E N K S V S Q I A Q Q A K P A R N Y M G Y T R R W H D G Q T F S N N P L
RANDOM_6461 S K S L P I L D T D K Q L F V S A V G L S C A I F V Q N E V I M R K R Y K I S V S Q T D G A K S G D H G K D F D
RANDOM_6462 S L L I G K E S S I G D K C A D G S P A D T G G Q V C K N E W A L M D A W Q I D E K K P S S P M S T S K I D P F
RANDOM_6463 Q S V I F P L L G R V Q G K S T S N T S F V V Q A H G Y D S T K N L S G P N H E F E L D E D Y R H R L R W L S C
RANDOM_6464 V S E A H G F S R G L K N V V L T H H V H K R A E L L G R S R Q R C L A T D F K A T G D L I I P L L H T K T H F
RANDOM_6465 S Y Y L W I T I P W N S L A H G S E G Q G K S S E S G D R K L A I A L C K M A L I Y I R K I D K T L F S M S K G
RANDOM_6466 V L G E E G E K D P V R E R D S L A L S Y V Y G L Q G E K P N K I L L S K P A G T R G A V R L N Q P I F W K A A
RANDOM_6467 N R V L A E E Y L K E Q F N G D S N F V I K V S R Y H T A N V F Q G P K I G D V I G I P T R A P P Y V D N M F I
RANDOM_6468 F R A R I E R G P E M P D R L Q R M A N S Y R L L N G S L K V V G C T E F F A H A N S N E S T V Q H N R G I C E
RANDOM_6469 A L H E W Q W L L G S P E Q K G S P Q M G H L T T C P S E D Y K R A D Q T H P L I P Q D Q L P K E M R M T V L L
RANDOM_6470 I P P I H F A R G P T L A T D S S N L S G A N N L K E T S K E G S D A S K G L V E K H F P S S T Q C D E S W P A
RANDOM_6471 N T D T P I A E K E D P F I V R P K E D R N H S S L Y R K W M R W A A S G L Q L F R T S N R V K L R I V A L R W
RANDOM_6472 S G Y R R D T G E S T L H V F L D G C N F E V N Q D E Q F E E N A A D S P S Y T E H G N T V S A P A V T L V S N
RANDOM_6473 Y L D V T G H N V N P K E E S F M K P N D A Q Q T T Q A N S Q G I E K D L P R V Q L S V K A K I C M N K G N G K
RANDOM_6474 K T Y D L Q G D L N P I V I D A C L L K A T D C A I E D G R H R E R S G Y A P Y E L Y R G N S L V Y R P P H F G
RANDOM_6475 R G L G Y T C G M N Y I A S F Q L L N D T C D S C L P F G D C E A A M T G K D D S K R H K N Q P L A I N G T R V
RANDOM_6476 L K N S C A N N I E N L L K K K P A L N L W Q N S D W E N L S H A S F S D H R F Q N H N N Q I Q A G S T T V G P
RANDOM_6477 D E T M Y T C T H L A H I Q L T G R A R A Y M K K A L Q Q I S V S W G A D Y Q T D R S C I K V V W R S Y E I F L
RANDOM_6478 S G K G N Y K N N E H I N E R G P P Y I L D N P S R P G G I A L S G R D Q N C E Q V N A A Q F I I R A K E G A I
RANDOM_6479 M S Q V D G L F I L G F E T L P M E A E N Q L R M E D C P H V G K S K Q D R T I F I P M A P R C D L S L D L P G
RANDOM_6480 V L L Y V Q D E Y S Y G K T E N S V H F F Y R K P V R T I T R Q V R I I R E S D M S D I T P D C E S S L V V D N
RANDOM_6481 L N L T G Y R S M V N E L F L V K Y G D E G Y Q V E I A A A L L D N E D R D Y Y L W V H E K L S S Q P L T G Y H
RANDOM_6482 L A K F I A E N R D A D L N T Q S M K S D E V S M G S G M V V R D S L V L M K F E V L T A I C H S G P K S T D T
RANDOM_6483 H H H F G A L Q Y A D T Q L N Q N F D G L S T V Q I H S M V G I C L C P G E V K E A L H I G H I L L S D V D L C
RANDOM_6484 T K L V P I I M F F I A N L R Q I P I F H R K R L F E L C L Y G Q C P I W D N E L N E K I G T K P A H N P E P E
RANDOM_6485 K A L D T W M D P V L Y H F K E G C R S A N K K D V P Q P S V S D A H D V P L L L G R V E V C G E Y L T R H S T
RANDOM_6486 F T N Q S Q G G T I K R K P R V E V G S L G R E L R I E Q S L L E T K A S L R P A A M Q E R A L M V M E K L W L
RANDOM_6487 G G A S I S R N C T V T V G V L T L S P V K S L G Y T L I D Y D S Y M E A V S A N I G K V T T V P Q R C I K V V
RANDOM_6488 E A L Q K K I A E E D L R A A G T L E E L M I G E S T A W R I G T P N A L T E I F E N M Y W A G S P L F I K Y L
RANDOM_6489 E H P A D P T I L L K F N E L N T P E G T Q Y S H N V W D R A F V L T D E A Y E A K G L N D T V A R T E L M V R
RANDOM_6490 D E I S F A F Q T R N M S G E V A W S A H K E P C K I G D G G G P A K Y L K N A P E I S D L Q W G M C R H S F R
RANDOM_6491 F V K A A R Y Q S D L M E Q Y A T A D S R H G T W A S I D I A K M Y A R G T Y K A Y I F E P C P D L C V I F D S
RANDOM_6492 T V D V F F G S D H K I C G T F M G L H D P T A L F V Q E E T L P E F A L I G Q A A F T Q N P G V H G Y V S H S
RANDOM_6493 I Q A S C I P A Q R Y L P V Q W T T D T H P P L S E Q A K I Q S E K M A Q S D K I Y E G S S Y R D S L I G M Q R
RANDOM_6494 M D T P V D N W S T K D I C H S E V A H G Q Q R F T V N I V L F D E D Q S P D E N N P L V K S G R A H D F A M L
RANDOM_6495 S R E E N T A A T G A G E I M V P C K P L Q R F S A A Y R F Y Q R T G T K N A S E K P N T L G T A R V I M A G Q
RANDOM_6496 K K N N Y P S F I T I F Q T P A K G A I L K P E L S E T L Y F R K F G E P G F N M K W A I P I N R K K L Q T D M
RANDOM_6497 F A E A N R G Y A I D P T G T E M S G T V M S L V N R Q Y G T T T R D N N N Y C W D I L P P N K A G R F P P V M
RANDOM_6498 R M D N A G M W F T F A A E P V S L Y N G D T V R I M T T W K K T K V Y C V K H A S T F R K Y E K M T L Y S C A
RANDOM_6499 H G P S S C S D C R K I E T Y T L G N T Q E I N V T T L C R E N V R E A A V T K G F G P E A L T V L P F R T S E
RANDOM_6500 L S L I V S C D D T H P M V Q G D G F L G V G S E D K S Y T A C L D P Y C K N A F A G E S V T A K G N Q F L T E
RANDOM_6501 M P S A N T T N E A K V C S E N R V Y T S T K P R G T E D G N D L K G K A H T A Y I A A H G H A T R T N N M T Y
RANDOM_6502 R V Q V D M K G L P R R D C C P R R I C P Y H R R Y L P D M V D T T D E D N W T K M E T M N L L A H P A R N P V
RANDOM_6503 P L T P S P V M Q S F Y S T K L G S T N F V S A G N V L P I L S P T K G Y T D L Q D A K D S G F F V E F F R A H
RANDOM_6504 T I D T S V L A E H Q T H H E D P T L S Q H F A F D L Y V P E E T V V V F P V G F K S G L K E K T E D C L S S D
RANDOM_6505 G I E V V T P L A I S K K E R R D R C E A A V T S L Q R D G G I L S G L N S C S S F G L E N L G A K Q K T R S N
RANDOM_6506 E D N D E K L D E K D P G I Y P I N L I E D A L T Y L W L S F H R K P L A C R N V H T Q A V F E T G L R T N R L
RANDOM_6507 D S G D L L T I K G C E L C F H Y W A D S E L L A R S A K E R N D R D K G A P V E R T I G P D F Y L Q E S N R E
RANDOM_6508 D H K E G V Y G S T V S L T V F T L Y A I A G I A T K A Q P E G K T P N Q I L K T T T A S A Y Y A S Q Q V F C S
RANDOM_6509 P T L T T R F Y S A H D A Y D L G S A H D V A N W K T F L D N V T L S D K E P T L P E E E E M N D F P Q S E A S
RANDOM_6510 T L S M A L D D P P P R S L T G F L F N A I A Q W P Y Y G M F Q S N L K E E G D L V P E E V N A L G F A Y S N V
RANDOM_6511 L L R V N M S S E K P L N V L H T A D K T V D D P N T R V Q T V C Q S A G K N D Q P F L E A V F A E G E F H T L
RANDOM_6512 D G K R E A S D A N P D Y E G Y I K S N G L E L D Q N P N D H P L R S A A C F V V K I D L K T S L D Y L S R G S
RANDOM_6513 I K T C W L T Q A G H Q A S A L N D V N G S D D E F E G S D P W V L R N V T P S Y Q D K P K R H A C I S G S K K
RANDOM_6514 R Y A G E H P I L R T I I P V I N R A F S Y P S V G Y S R D M Y K A M G L P E I A I M K A S Y M L G Q A E A H N
RANDOM_6515 A N Q Q Q K G L A A V P K R A G A N Y F I I Q Q A S T I A E A M D N Y R L V D P M R R K A E N F T M A V G A S L
RANDOM_6516 R L D K E V I L Y N L G M V N L K T A G S P S G T I R L N N G T F D G Q K G T G P L H S S M M A Y H V R L L N N
RANDOM_6517 T A Q S L A D Y P S P G S G A E G K T T I I R K G G F R L L S P T S A K T K T P E A K P G C A I F K A T P T N D
RANDOM_6518 E Q G L F S Y W V P Q Q L E L V T R G E R F G N C Y P S N P C L E T K H G M I S V N S D K V I S S A T N E E H P
RANDOM_6519 E L V K A A A T C A P Y I Q H P S S D Y L G D G L R C A A T I A P A R P V N G T R C A A S I H N A D D R S Y M D
RANDOM_6520 E A N V V E G L M T D E V V K L T V N Q D G A G Y A A R D A Q K V Q S E F V G Y M L P L F S L G F M S Q T A S Y
RANDOM_6521 A H F L I L N I S R L N M W S E M D G K A Q T S S S A R L P A C S L K Y K P G R D I A V T Y M E P T L P M A A S
RANDOM_6522 K L V C P A A Q G E G F D I E A K V R P S N I I A Q W L C R Y G Q E D Q I K G R V A E V F A D S R L F Y Y G F A
RANDOM_6523 Q G H L K A A C I V V I A V W G L K I L H G D M P D Y Q S Y P A E A A S T Y D S R D D K K R H G V A G P N S S G
RANDOM_6524 T G H L G L L R L K A V L S N E S L F S L N N N S R R R E N V E E H K L G S R V L L T S N E A I K L Q P Y S Y S
RANDOM_6525 R A T K E Y T Q E V D A V A I N N A E M N D V L S G P M I G P Y K T R H N L E A Y K N W F K N E G T I V N L L K
RANDOM_6526 K I M K I N F T D V I S R G Y H G G A P L Y D A R T K V A G S K N S K L I G N D E A G S C P T Q N F R N Q Q R R
RANDOM_6527 V A L Q Y G P L E Q S Q L D G D T T T K Y C S I H P I Y G S I N H P V N K D S E A A K V S L V T A M S V P K F M
RANDOM_6528 A R H K E E L W E M A H R A T K W K C F P L T F R N L H V D A I A D R D L T V N D E C S N S I R T G F A A G Q S
RANDOM_6529 W G G E F D V C C P C T G R G G I F K T K D L D Q I Q P I Y H C L Y Y W Q G S L I I V S T R H F R F Y E R I T I
RANDOM_6530 Q S L L S D W I G L R A C H A D I G P A R Y T T G V S D L C I Q A A T D R S A W R A M Y D S V P E P K L G L T F
RANDOM_6531 V G R C A L L D V R D M A K N K E I V N R R R G A E L N A R K E W L G Q H S R E E D E C W N P P K N I N T S E I
RANDOM_6532 V Q P K E L C Y E F K K L A R A L Y H P A L T D S Q P V I A L R E A I L F D R V V R A I A W P I N T G L A G S L
RANDOM_6533 I T L V A P D R G E Y T P F S L G R R G G P A E L S G S R P A K E A G L L R Q I A D S I E K V H K A E G P N G M
RANDOM_6534 K K I V R S E K Q S T C I L D V S K V M P F Y A R P F S L R L E D S L T L N Q S T P G V K G T T K A H S M R L D
RANDOM_6535 S I S N V Y A F Q E M P F S Y M T F R Y I D V D M L I E L K A V G T Y G L K A S K F V Q Q E A M V D L G V Q I V
RANDOM_6536 S T E N V Q H I Q V Y T P K S G T S E S V M T Q L T Y P R L L D R V R S K K D W K K T K F S K Q S L N L G Q N Q
RANDOM_6537 K V A Q L T N E A T T S S D S K R A A S P E L K A Q N H N Q V R S S L I L R N N V G R S L M L N R I C M I S T E
RANDOM_6538 D S P S R I G V W T Q E P P E L P V N K I D S L I S M N T I Y D P T P A D G F N K M V L Q Y K G D V K V V N E C
RANDOM_6539 N G G C A A D Q G M S G T D E E E D I I A I S C A E E L L K L S R A G R K P I A I A W S V G A Q A V N L K D S V
RANDOM_6540 W T C T E A P F G K P K P L V S S F N L S Q T Q V Q S K L Q D S S D L C K D T K A D K W F R P A Y I I R E W M R

RANDOM_6541 Y S G S H N K D S G I T N I G N Y I K R I V T E P P L A Q K G L S S Y P A G S Y K R Q E S L C K A K L G P K T H
RANDOM_6542 H R A R L E N V P C R S Y R S D P G E L L H E F Q F D S L T Q V E D N P C N I V E E K P I Q A A S S A D Y H V D
RANDOM_6543 I S K G I N E T E N R G M D T A L A D F C E G G E Y E K Q G T I Y A T H K P P H D T L L Y W V D C K D T E C G V
RANDOM_6544 S S L T E N P H R A R G L S S D Q T D L P G S A T L V K S E A A I C V D L Y F E C G G T L E S S S K K D V D L Y
RANDOM_6545 N M H T T H F P K S P V A E K D N T E N K N S F T G W P Q L L I S N T T V V T K A M Q K K I C I G N A V I K P L
RANDOM_6546 C P D H L E F G I A A S T I L Q V P D H D T H R V N Q A V G S S S L A N H S R A R E G R M L A S R T L M T K K E
RANDOM_6547 K K N G T L E S F L S G N A V R M Q Q V T D S M I N S S E G D H G Q E V S Q T T I H A G R Y N E S A V A M I T I
RANDOM_6548 S S P M I Q M I R D Q P R F F Q M A T L V T G D A A Y C K T Q D Y S G D L I E T D A L K N I P D I R K A G L S R
RANDOM_6549 T K V E S L E T I E F P I Q T E N S K D A R F K R M Q Q G N E K G I L G I T P Q T Q M L Q Q T L L G K T L T G G
RANDOM_6550 S G T L A Y V D Y F D D I P Q R D K T H K N G T D A V Q N G T L N S A A I Y S Y V G P N G D E K T C F D S Y I N
RANDOM_6551 P G Q C V Q S E G S Y N S R S V K E M D T R V S D S L S L T H R G H L G M C A G H C I T V P N I I S G D R G S H
RANDOM_6552 E A V N F V G P E P L S N A N Y S I H A I K P T M F G L A H H L A V S R D N G L W V K P T K P F L G P G V F N S
RANDOM_6553 C F K D N I L D I V K Y A Q W V L G E L R G F A S R I R L N A W I A R V S G M A L T L Y A I A D E K P L L H E Q
RANDOM_6554 Y T A S A D V K S D A R G S D Q S T V N R G D L L L C T Q S K G E S T A V D T V G A V L R A Y K F P A K Q G L P
RANDOM_6555 L I G I L G T A T E A L N V S G K N A A A T T E D T L A A L Y R E V K W S V T I P V S M S V A V S E G T A V M I
RANDOM_6556 P G E N R L R V Q R R F T E N T L P S I N S K P R D P V Q N D G H N P L F D V F A K D A L K D Q N P M A S V G Y
RANDOM_6557 L L A V A G V E E E L E R L M F F T L E K K S Y K H H M A I D T K G T G I F L N Q I F L R L Q I I F R A L V A L
RANDOM_6558 E R I W V S Y S E R A T H G G S E Q N C T L L K P G V I P V L S V T K T V R Y A F E G A Y W G Q K F P R L L A S
RANDOM_6559 H A E E G H C S L P Y V P Y D G A D G T D A H A H E P T S Q L D F L L W L S A E L V A P W R S D A V G S S S P V
RANDOM_6560 P N K A K S G A L P W K Y N V G T A S I D C G G S Y V E D W I K Q S G V M K S E H G I L W Y E E Q L L T D L K L
RANDOM_6561 T L F T P K L T T D L V A M Y L P S P K N G I V D I S E T F S I G P Q L Y P S C A Y F C R R R G S Y E L P I S A
RANDOM_6562 E I Y P T A A L A F F R M N R G A T G E P P S E D A W Y S A E V T I D L K K G G H S V R E L D E L E K D N A P G
RANDOM_6563 S F Q G K I E K I G C L E G S E S N M V C I A T S N F P A Y A I L P T S V V P A T N G N V P F S R M P V G P Y Y
RANDOM_6564 A H K E K H Q D P V N G V H N E A A E I S Y C A H Y E R A Q V A F G D L F T Y Q A P P G R E R I D A C Y V E K L
RANDOM_6565 T G N D G S W T T M Q A A T T D D L E L H Y E I G G I R P E L S R D M K Q E G G M R A P I E V T K I G N R Y L R
RANDOM_6566 R G R I A P T L Y H Q Q F N I F Q L T P P V F M A S T R L A I D T F K G K K G A A I N N L M P I F N E T K C T V
RANDOM_6567 I I K G L R Y L Q F E I Y F A P A S A P E E A K T L N K G G G S V G V G K N P N L T Y T P G G M A F F L P K V D
RANDOM_6568 M Q A H T S R I H W Q L T S A Q G L A K M I L K Y A D P E I A K A V A P I T L H E P H V M F G A R E A F E H E L
RANDOM_6569 A G V R C Y S E I V R L F N H S P Q R K I N P V D A N N Y L R P K P K H S V R N L G S D E M P V K R L F D L L R
RANDOM_6570 I V Q S H L M E I L S A L S E V T C F I Y A Y C D F K I L G H K K L L L K F T C R R R R S P G A A T M I M A K I
RANDOM_6571 T N S V C T T D L T H D A D L D Q S K H S L H E E G E R P P L F A L L S L R S K K P A T N D K T S V E P Y H E Q
RANDOM_6572 V L L V G A T E S W P Q T P P G L V V T V A D I W E T G E A F S A R H E N I S Q G L Y Y F Q A S V D H I D Y V P
RANDOM_6573 L K G D R A F D E C F I R N P A R R D C V R P I C D V G K L Y V T A I T G G E T P T I S F E P Q P R N G T A S I
RANDOM_6574 Q R R R P Q W F V N S Y D W I M F G H G Q L A D N K Y Y T Q A I I E T R I D Y S V F P V V G L S R Q P K I T G K
RANDOM_6575 V S G F R Q A F A Q G C I I Y S Y A T L Q G E G L E M N N L N V E M L L L N P F R K S E E W H M H E G P R H Q G
RANDOM_6576 W K T R P A K L T L A G N R A V W Q G H D G D E V M D K D F G V A R D G F N L N L K H D T I Y A T F K V E A R L
RANDOM_6577 T C A E S R S M T N L E D T Y P V L K D E L A T E R G L A S G K V M D D E G V H R A K L E A D V R T Q K V G D A
RANDOM_6578 L S L Q L P V S Q G T L W A A C L P S G H V R S P E V L V L T E M H L S S K N W S K G Q Y C I S F E L M R E H S
RANDOM_6579 Q D Y I A N A P T F R I R S S F A S A S F V Q L F P T G F Y L D Y Y P L I A C W V A E H E E S R E S L S L L I G
RANDOM_6580 A D I P E I A I A A R I K K I S S W P F D K R S K S D G R A T S R N L E G V Q W R Y D E T L F L P S A A P Q T M
RANDOM_6581 G E K D Y M E V T M N T T I L S C A L M Q M F A G A E G S Y F V E G V F A A M D P K T K S A I F P K S M G P S Y
RANDOM_6582 K E N L V L K R Y L L C M L D S T K N I S F Q P T P G A E T Y A T R L E G G K F V E W I G R W F K L G T L E S K
RANDOM_6583 N G R N L C F R R T R G D V V E L F T N R E L I L L R S A Q V A S W L D P M C A G L V H S A Q V L I S K I W P S
RANDOM_6584 S C N A N F G S F N F M I L R F N E K S F R N S P G R L L R S R R T V V Q K S L G A Q R Q V P L I G I V L P M L
RANDOM_6585 K E D L N R D T F S V V L T M S R V L P R P H I F I I C K E G S G G K R F V F L P S L A M N R T L E R L C V L R
RANDOM_6586 G N I G S Q L K D R D S G R P L V G Q E I P L T L S D D E V D L E L D V L S N N E W N S I R K L D D P L L A F E
RANDOM_6587 R E R A P I A Q T N D S D L Q A E L Q N K S W L C G I S M P N I L E R F D E N Y S D G K Y D R K E A L I V C L R
RANDOM_6588 E L N S P Y S S Q N K Q N A E A I V L V K K E E R D R T S A Y I G L Q P G V T A P V Y I E T S G L N T K M P R I
RANDOM_6589 T A P L R I L R A G V H I S P D I S Y V K H S A L N K P V N W S C A A L N D T Q Y H S A S A S G N D G R R T K S
RANDOM_6590 I I P F V G N I F Y F I N E T E T L W E S H S M A R K P N S D L E N I R S L C K L L T P T F L Y V C I N V F A Y
RANDOM_6591 E F R D I D E E Q P L L K T L C C Y Q Q F P L A N I A I G L L E P S C S H A G F M L V N L R V L A H S Q P D D V
RANDOM_6592 T I E I D Q P Y M V G S V L T L I S P Q F D C R P K E R P R C S T T E K L G V L E M S A E L A G V Q C R K G D L
RANDOM_6593 C I G R A D G I R D I D T I C V R F D A K L F W K L W I K F E D S S K K M P A T T K V G S F I H H Y V A S Q A P
RANDOM_6594 F L G L R D E S N Y T S Q A K Y E M P V I T R A D V A T R P Q K T G S P S G F A G M I G K A G N D S G S R P R A
RANDOM_6595 Y E L S G E L E G K N I R N T E E D D G L P L A C T T A D F A I I H I L V A L Q V E G Y R A C F A L G L P L M M
RANDOM_6596 A I Q S G S P M V K P P T M A P R H R V A G T F D D M R E I L L L F G N H V Y R R M S L G V G D K D G Y N A P L
RANDOM_6597 E E A W G K R S I R S I A H G H S Q R P W V W T F S L L S S E Y E L A A A N A K T I F L I R P Q L P T K V Y S T
RANDOM_6598 I Y S E G K L A V E F A I L S P F K T A R E G E C I P S V I Y V K Q D D L T T Y K M T S E L E W L I L D V L P E
RANDOM_6599 G L D D C I S G T T L T A G C H M Y V I E T Y E F S W D N A Y I N Q I R G G I A T I L I T I T D N E E A L T H P
RANDOM_6600 K C G L G D M G S E P F F L T I I V E Y P K R I Q G I A E S C H R S R F G V A E T Y I A L G K A T N K S Y T Q T
RANDOM_6601 S A A R S A A L S G G I H G D A H D T D N V E E E L A A D S S A A P D R V Q D Y F E L F T M Q S I A F P S L R Q
RANDOM_6602 M P P G L D S G E L Q N D A K I Q S Q N S L V G V S W I A E E V T R S S K F S T G A Q E A A R P Q P T K N P V I
RANDOM_6603 Q M S T E V P Q V A T V P W S V Y T S K G N D G A T A E T H F S A I Y S K L L T S I S V T A F V R D D V L G V E
RANDOM_6604 K F Q N G M F R S V I H K P Q Q V L D V L A L T G P S L F Q D T T F G I L S T H L A G D H I L P G K D D T L H T
RANDOM_6605 A D I I A L A L G R D M S S N A I K N S A G I N P Q S T R D W L G T F K I I N D F D P E K Q S E G Y L R F A N Y
RANDOM_6606 K V T S A T Y K P L A N D I R L A K S F N I N T N Y E V C L S I L P M G K R V V S T G L N E L D S I A S E L N V
RANDOM_6607 E D E D N A D T T R E V H L H M R G Q D G T L E L D E L L L Y W D I G V E G Y I S D E D F G A F T S I V L D A S
RANDOM_6608 G E E Q K A R L R I L I V S L T L E H I N P P S F L K M S V P R L C F A G T V P P L L F I I I S Y P S I L K K D
RANDOM_6609 F G A D H E A C A F I F G L L C R R M Q S S S P I A W C I L H V P V N S S V E L M A E R A S P T G R G G D A L P
RANDOM_6610 R G L D G G E L F L P G K A K G F V P G F L R E K N A G L K Q V V K G L M I R E T N S Q F M V A E R E Q E D R Q
RANDOM_6611 T I Q L L Y E V D A T G S A A R K P V Q G W A Q A A G M F R S A G N K K N K S L I K V E A P E F D C L K K Y T F
RANDOM_6612 C V L E F S T T A S T T A S P T L C L D W G S P H T E Y L L H A K F L I S Y D L E A L G V A P M T W N V K N S A
RANDOM_6613 L G R I M G G E T S S L Y G R T I P K D L V E V L L A P G D S S L Q S A A L Y P S D T D A R V S P D G G G V E R
RANDOM_6614 E Y S R D L M P I S E D G V S K Y K A P F C S G E M G N K L N S E S P Q L N E Q T I D F E Y C T K M E D N R S Q
RANDOM_6615 K S F P V T V L R P S E A A W G Q H A D S D Q Y P I E S G F A G L A N H L W T T K L D I H V Q S R K R E I S P V
RANDOM_6616 F P L G V K S F I P S E P G G K P T G C I V I N I G T M M F P E K D A I Q Q E S I K A L Q P V R M G T K V F K V
RANDOM_6617 K E Y P D Q P E A P P V K G Y Y S G H Y H E L V M K K D E L P T L K A F L T V H V V E W V P N V Q S E T A V N Q
RANDOM_6618 L W V Y E K N E N P G F S L S L C L L L V N D E S I M T E R L L K D L E K R A Y K E K R Q S Y C P G K G V G S S
RANDOM_6619 T S N E S L D E V I H S A P D L V T G D E Y A V E M R A A D F F F R F D G R V S P S C T Q R K N R F T L T G L N
RANDOM_6620 T G V I G P Q Y G V L E D A M T G D R I P F Q K I E H P P A L R P V F I S T H E P N F R V E Y V K W K V I R K D
RANDOM_6621 G E L A D Q S T R L T T I Q G L K L R V M F L E I Y C A W G N A Q D D S T C L N A L R I G N I T L L G Q V A A R
RANDOM_6622 Y M N A Q S T K A A E R G T D K M L L I R T R C V A H E V S G G G H S A T V T F A P M E T D D L E T D T L C N M
RANDOM_6623 P A V E E A V T A P G G N D D S F T A T T P I L N V A M Q A L D M E Q G A T G Q R S A G V Q S G D S S F P D T W
RANDOM_6624 I L T S W E K F Q E F L L E G A I G Q M N S V I M L P T C R L E P L R K F C L F F I V E S S K Y A P A W T W I D
RANDOM_6625 K D T A N S A V S V I D I P G P R V N A Y E A G E L M D D L F I E N G D N V S T H D E T M E F T V R V Y K R T V
RANDOM_6626 P F N M D L S L N K E A T V S F D H I D F W D R K G G R S T C F Q T K V K L S S G A T L T P Y T C Q S S E V T A
RANDOM_6627 M I A Y M P D N K D L A T K T A R S V L A A I I T L L R A F I I G F F L S L V K F V G T L K A N L N Y A A G E E
RANDOM_6628 D T T T M F A A A E A C R T D D V G W F I H P S D I I G G I L E G H E N P P H L N Q N K T A H V S G L L K I L R
RANDOM_6629 K L N L K Q N L Y R V V Q V D R V P N L Q E K K S I T V V Q V E E I I A I N R L Y A G D R I K L A G A F S V K V
RANDOM_6630 L P Y S L E A I H M S R F K S N K R S R G W R T S A A C A V G T A Y E D N I C G L K V A L P S A T A W K S A L R
RANDOM_6631 M A E K A K S G I T K R L V T V P K T E K N S Q P H Q A S P M Y E Q S S D Q R I M Q G T S R F S L M L Y Y A H S
RANDOM_6632 N D L N E N H N A I R T W K D L L G D N K V V P E N P L S G A N Y A G S V N S S N R S Q D P N G E N D E I C S S
RANDOM_6633 G R N H F R T R A E L Q A S G D S Y S R T T P I E Q Q A R V N S M A R A S I A V A I R G P A Q A I E P T I L L V
RANDOM_6634 G V A S F S T L R P K V Q Q Y T G A H E Q D F F L K Q A E T G N T A R C A K H V L M I L E D Q Y E A A Q L E P A
RANDOM_6635 G Y T D Q Q L C G S S V Y N I T Y P V M D A P Q E V S Y T N N Q L I L I W Q Q V Q Y R I Q F Q I R G N V M S S Y

RANDOM_6636 P F L V N R N L L K R W K L R T K D N K K T L A D T F F M Q S A T T S F H Q V D K N A Y K M T S I A V Y E L S F
RANDOM_6637 F P D F K F Y R E I G Q L N F N R A H D D S V K D G S I V P T V R E L N A L G A S G Y S E G W I A T A K P R E R
RANDOM_6638 I S L L T G S I F K L I L G N Y T T I R P L A L G L T G T Y E L F Q D D V A L M E E R C E N S D A A V E Q R Q T
RANDOM_6639 E R F A V G V K G S P F V E D I P N G G E R A L T R Q L T D D V L L P S I P T H E H G T E M F L V A K E P L S P
RANDOM_6640 H R V E A N E F I L L Q L S A A A L V T A E Q L G M A P C Y G Q L Y V T A V L H R R N T L L S D L Y K I T R K D
RANDOM_6641 T F H G P E T L M S G T V G C G H M I K F K R S E Q Y G K R D L G P L L G Y Q P V S T M D N G S I G R L G A S P
RANDOM_6642 L A P V V V E V E S L A A P K L A E P Y E G A N S R C S T E Y V H K Q H V V S T F R N I E K A A A G G E G H M I
RANDOM_6643 V P L P C A S E N K R A N T G R N Y H G I G F I M A E P Q K K G L D G Q G I D S K R P P G D Y L T E D G V A L N
RANDOM_6644 R E P N T K K L F C V R A A L P E K Y Y L V G I A V H S T C W T Y D D A S Y F W L E I P T M V Q L N G T G S P G
RANDOM_6645 V T D G L K W P L V S Q D A Q K I I G N P F S S R N E A Y G T S I S E N H I S A G A G S G C C A P K A S K D T M
RANDOM_6646 D A S A S A N G V G R K F Y K V E Q N S A N S V V G N T P Y N L K L A E L T V A E T I D R T C A G V N N L S S T
RANDOM_6647 I L M N G K K L L F V R F L Y Q A D F L V G E K T A D T L L G G R P Y Y Q W A E E L A L L R V I M Y S Q N S A G
RANDOM_6648 K D G H N H P N C A A D D A I A R G T Y A G R G G P A S C I E D W C Q R D A T L V H F P Y S R L L N Q A Q C S A
RANDOM_6649 C I G G P S S A K E R R V V S V F Q Y A I I S V L P K A D T L K H N Y L S T Q Q N T L K K K G E V G S E K H V C
RANDOM_6650 Y R D R D G R P G Y G V G R L L I Y S F K I N H K D M P F D P E L L E E T F K A G S I K Y H Y D Q A L W F Q A A
RANDOM_6651 Y N P L S L S D E M K T T W E K W C L Y T L P E A S Y E Y A Q Y L A E V A L R C R C V A A P I N K D E I S R S K
RANDOM_6652 V Q A I W L L R H D E R D A L N R E T E S I K S L P E I Y Q F C I E A R G L S A R Y A A P E N V E L R R M T T D
RANDOM_6653 M T E R E G G E S V A Q A S S E M A T R F K I K D Q I G S H P A P I A H V A L G V A L I S D H F D D H W F Q C L
RANDOM_6654 A A G G R T I Y L G R A M V G V K V G H Q D R G T N E G I T Q L W K R L S G E P K I Q Q R I V I P G H I I N E V
RANDOM_6655 A E P L D K L G L I R N Q V N M G D V K F R R T C Y W P I A F A L P Y I P W G I C G I R L Y N Q P S T D Q R A G
RANDOM_6656 V D C D E V Q G A S E H C I F E T V E A A N Y G K K I V P V I C R F I Y S A E E S C A V K M A F G L P I V A I L
RANDOM_6657 A D R A I I A K R K C D R L G H T G I W D M L T H S K S Q I L L Q Q I Y Y Y L T K V E I Y M G F P K P G V E Y S
RANDOM_6658 L P S A A P D S A V I P K G V I E R Q A H D I E Q Q V F I Y K L S R S D S S G D V L D R N S A S R L V G D S V G
RANDOM_6659 S Y D D D S D T N L M S I N L G G A K Q S D A Y S H I V H N Q A V C A A L G C Y V K A A V K G T A M V E S K V E
RANDOM_6660 L V F I A L F N P V V V D L A L F F R Y P M Y K H I S S S R V F K G N S V P A E E A V G L G F G D Y T E P K A L
RANDOM_6661 K Y A A H L R A L N I L F M G P I T V R K W S Q L E L Y G L L Q H I A Y I I R G F E K S E T E H V T K H N K S E
RANDOM_6662 Y L N N R T Y N N H L G V W G Y D T G S V F S R T P G L V Y K I E I N S E V T R G S M T L Q A V T T N S Q G R Q
RANDOM_6663 E E A R Y V S P K E A N S T R P H Y K T G P V D L T E M S I A M I V L V E T L D S Q E P H L P R S K D H G T N K
RANDOM_6664 G C Q Q L V G N G T W E S R G E I R K E E E G S R L P D N A P S G F P R R F S N A V A Y L L G I R E D K L L P W
RANDOM_6665 D Q I Q C K R A R Y I V F P E T N L G I Q K L K K D S Q F S Q R N A D K L K F L H L L S P H K V C E N Y E A E Q
RANDOM_6666 Q L R I S T A T L K V F F D Y L G W F Q M R G L G E D Q M N D D W N K K V R P Y V Q S I V L D Q G A D C R V V F
RANDOM_6667 I I P E I F R Y F F P F I N R A R G R V D F T Q Q N R E K E V V G F G K K L K K L T G D S V K V R R G W P F A G
RANDOM_6668 I I D H Q A G L A H S V A P I D F V C T P K S W L M A R E K E N L T A H S P A N L A N T D A E P E T Y E L L C D
RANDOM_6669 G F M F V E S K S L T W S L D G P K E Q V G D D I L T A A L T L L V N G E G A Y L L S P A L D Q K P V D M V T G
RANDOM_6670 N Q K E T Q T E N Y I V V S R R A D E T D R A S T K I T Q P I L R E I G G Q N Y A H R A D E Q D H A R T G V G E
RANDOM_6671 E L K R E D A T Y E S A M D R Y F R V M T G Y L S A T A S T N C V E H F T N A K K S S P C C A G D M Q G E C K L
RANDOM_6672 P R H K S D P T Q R N A G K S E F T F I P A V N Q P K Q D C V N I P I L V E T R M L L I F G L M T Y D K D V K T
RANDOM_6673 S R K S D Q V E A S L G P Q R A S S L N D V T N L R L E V R T G K A V M N Q G P L I D Q L F T R I N Q G R N P V
RANDOM_6674 D S K N P R L G I N M G N L R H C K Q R V P A G E V E I Q K N S L Q K Q A C F V V K S S P K D Y R R D K R D E L
RANDOM_6675 Q W M P L I M L N V E Q N S T Q R E I S S L V F M N N G T A K I V Q F G T C E G A K V R P K Q E S L I P G N I G
RANDOM_6676 N L Q V G V S I T S G R G G A Y A D L A C C L L Q R E E E K G P A G F G F L K W L N V L V E G H A I M P L D D E
RANDOM_6677 W R K A I L F A L F P H A V G T D L T G G M E H T N Q T E P F F D K T D R E S Q F C A E G A K P R I H A L G M S
RANDOM_6678 T E L Q N H Q N R K V S V S G H T G T K V V S K Y H M Q D G E R A L S A D R M W C L L E G Q I C Y S S R T V L R
RANDOM_6679 S G F E N D H T Y K T M W N L A D P I E N G Q F G F T D V N Q T D V L D E S L M K A S E A P E A E A Y L H I E K
RANDOM_6680 T M Q Y V G Q D M L N K E R Q K P R V E A A V D D K F T D Q P R G R V G I S I M D G R T G G G K I E T Y D Y V F
RANDOM_6681 M A S K S R D V H T P K S G E N D I K V L C G C R E E T V D D C D V Q I Q Y L N E L T V R S L N E D Q V Q Y K R
RANDOM_6682 R S T G S Q I S I I I A A S S V T V K F A E T S N E M Q N A L G G S L I F A V G P G V G D E E A L R E A F E Q L
RANDOM_6683 L K G C P V R D G R I A A P S E A S I E A V V L L Y E D N A A L G R D C D T F L G M G A R T T A V K P I W A K I
RANDOM_6684 R G S K T I C S M G N P D E G P E G D D T T N S Q K L N D P H E E S M K I T V I E G E N Y L A K L C K G G L L S
RANDOM_6685 A R N L L A A I A H A M V P E F T T D Q S F N T K M E P P K V D L M T I A T A C Q I V N L V V F G M L P Y Y T L
RANDOM_6686 V T T S R A D P G L L D M F N S N T T I Q P T G K T D F R S M S L T P S E P D L T E T I E H K L N C L T K E E K
RANDOM_6687 S K S G Q A D K S P R S I T F T S S P T K K A I S A I L G D V L E C T G E F H E G P V P L H L D K T Y Y N G G S
RANDOM_6688 V I K T R L A V Q F R K A G S I E L D R E H R N N V I P L P T V T P I G H T A V D V S G I S L Y C S S G T Y Q Q
RANDOM_6689 G E M S N A N E I T A R Y A E S A K T C T A R S A Q G A L P R A A T E S R A I A V P A E T T A A P D A H T C S L
RANDOM_6690 I P P E R L P S N S L N V A S L D G K E T M S K I C A P Y D M T L S T G F T T H R L Y A Y L F S M G V K P D Y N
RANDOM_6691 K E S L D R A L L F A V S L W L G G G G S N T D N A L T V I A G M L F E E H N D K L V H T L P G T R Y I E M Q G
RANDOM_6692 A I Y G T N N V L S S F G V R M Q S G P L F K N G F A G V N V V D E F A L C G A D Y L F A S E E L T V Y L T S N
RANDOM_6693 R K A H L S P D W T S L N A L G N K G Q S Q N G H G D D H A Y E S L C E L E R Q K L T S G E C I E T M S T V P L
RANDOM_6694 L H N L D F F L D V T V F A W T P T N A S V R D Y P H V C Y V G W K A A E H E V G Q A V A S N V Y E M A F M K R
RANDOM_6695 K R N E A V G A I F A L T I V E L Y H L A K F D Y T S I S L Q E H E T E Q Q D L Q Q N V H A K S G E L K S G V L
RANDOM_6696 F A L D N D D H V D P S Y Y N I T M P S P T N C A M I D N H L E H E V K S L L N A L K Q L D V I I A M N V D E V
RANDOM_6697 L T I V W E D P S A V D K M K G L T L G T C T E F V C K N A N Q S P Y E S F E G F H F Q D S P V I E L L E A D T
RANDOM_6698 S E F D Y H L I L E C D T R K I V K V L K Y S A D A K P L R L V K P G S A A V D K T A E I I N T F F Y Q S D K V
RANDOM_6699 N T I K N I A P Q F R N A N H N D L D E G Y S L Q V K L R G G D F L N H V H F E R D G V P G C M E W E A T R M N
RANDOM_6700 G Y I L G P L R E C S P L E F R F G E F I A D T L I G F L H M K D C A P V S Y S L Y F N D L L G E A L Y R Y F T
RANDOM_6701 N C L D E A T M E F G L F F D T V G A I T A I H H I E L L N T L S G G K G A V S G A L P E Q L G L L T G Y A L V
RANDOM_6702 V S F N H F L A G D E P P H G Y Q D G L I I V F T S A R Q H P P K S C P R Y E G T P A G T A Q V Y G Q A V C R S
RANDOM_6703 H K N S I D D D V K A D S V I K Y N A R E T P I A M V Y A F G V N A L T P G E E F L R E L G K C Q D P E F A I V
RANDOM_6704 F K T S E I V K I Y V H I P P R D D G S K E T Y N I E F V R R G E T E G L G N T V L L G R S N G E H S G E I A V
RANDOM_6705 I I I N A L Y D I R Y N G E A D T L L D Q Y D V I E L L H C S D D T K K R F Q N Q S E I L S A E W Y E T P N E S
RANDOM_6706 G E M K S V L R E G G A R G V H G F T W D L V G A P R P R I N F I A D I N L Q R D S G I V R V G G T Q M T L A R
RANDOM_6707 P L L S N Y L A G E M R A P G V N R V G E R N L C A G A E R P L V A L C S N M E W I K N A K T R S F F Q R K D E
RANDOM_6708 V L V D Y N S G C Q L T P F A A K S N R L G K V V T N Q E I L K G T Q N S V P V K P K K E E N E A E L P Y V E S
RANDOM_6709 Y L S V Y K D E N V Q I E E F F N S A F R A W C S R K E S I N M L N K I S N A N L K T D L Q M V L K L R Y H S A
RANDOM_6710 P E H G D L L D P D R V D R K K I K A F T V E V V P D E N Q G E A I V D L N N K H E I N D N E T P T T T N S T G
RANDOM_6711 G S Q S P Q R L D T D G S G K F H P D R D S F G R P E V T Y D A T S V T L R Y A P V T M G Q D G M A I E F F E F
RANDOM_6712 F A D V A C I D A E G Q L K D D R K S S R S L Y L E P M V L L W G R L P D L K K K L Q V E T F L H A Y G E L T S
RANDOM_6713 R P G S D T E S T K Y V A A V Y V Q G L E I P T F N A N E A T I P T Q K C A A V W E S W A G N L D R N G T K W R
RANDOM_6714 I C I A L G P N P E P R I V I P N A I E D N R N L S M V A G L F S D E L V G N W A S L E V T C W H V T L D D D G
RANDOM_6715 W V Q S D V V H L S E W I F T D F N G A W G L D L F T A L N A V K T G L L K L T I E D Y S A L V K G M R V C S L
RANDOM_6716 R E G C G L S R K S F A H N I L E L G L G G K M S L W A K F E N Q G V A L L Q Y N L T L I V A V C T A F G N D L
RANDOM_6717 A C V S A T G S E A M M W N P I T T F P W G P R G I I I F G W L F Q L G V A S G L P K R V I Q L Q D Y A K P R G
RANDOM_6718 I F E I T H I Q W L V L S P D T D A T E A Y D V P Y P L I A G K V V G I S P T S R L R F D L E E G I A T I V S G
RANDOM_6719 R E P M G G V T E L Y L A I K N A I R R Q E Q Q A C S R S R S K Q K D P S I S H Y Q W K A L I T R G G Y G S E N
RANDOM_6720 S I M I V A S H S P V S G M P V V V F E A G G I F A E H S Y N S N D V Y Q M I L P Q I L K R T F I A N L H G P V
RANDOM_6721 E V A L G L R Q M T I K D L A G A P K R G E V I S K V G R D I L R D G T V E N R R G S T R L P Y I A Y Y I A K S
RANDOM_6722 N E K L R G V Y Q C V F R S T N K H T V K A E E A V P V A A I V V P D S A L Q Q R L V G L I S L Q G F I G G H P
RANDOM_6723 E D A I H Y K K R P T S R G P K E G N F P S N H A N V L Q L A G A A D Q Q N V Q G Q D H I L I R I M A R G L P K
RANDOM_6724 A W G A E K E L A L L T K P E R Y A L Q Q D G S A V F R E S E A N R L F L E D I L D N V R A P V L V T A H F Q P
RANDOM_6725 K E S F K A K K N S D R V K H E D K I S S P G R A I A V I C G G G N D L G A D I S E G P N K I E S C V D I W E D
RANDOM_6726 P W L P L S C P D I P E H P K G D Q V M I V S T R P T A M S Y Q K S A S L S K D Q E C K Q P D G L L E G M L Q D
RANDOM_6727 D T E P F F S G G T L E T F A V H D C D I A R L I I R L F A A T Y P S L V C E L E M I F P F R T H F E V V N L T
RANDOM_6728 Q I T P I L G S A N R S D N Q V K A Q P G C V G A I P T R A E V G G G L A A T L L P C R G L F K M S S F N A R F
RANDOM_6729 G Q D V N S V Q E N R G T Y D L A H E G S V C S T S D F E T N H T M T R R Q Y V S T T G A G F Q R S R K P P G P
RANDOM_6730 L N N N I I L M F L E T K S E I E P V P A E Y L Y Q L M S D A A P R D L S A V N R D T W Q F Q Y R D N I V G H V

RANDOM_6731 T P I S Q L R K K Q L L R E G A N V I N R V E K H N C K S E A A I P Q P L G N R Y N L Y A S P H W P Y Q L Y L L
RANDOM_6732 T I T K G T W T S T Y Y F T M L A L T G N G V E P A P A T C Q P A R G K K F G G P G S C S K N T V I D R E K C K
RANDOM_6733 S F N S K R S F P R K Q V N E F R T G K K F S R E L S G V L N E A L D E I L E E I F H K R K W G K G H H L P L V
RANDOM_6734 P Q D V A V E F A Y Q S S I T K G D E V P N V D E I R E V C C E N L V P D H G H A L D G K T W L R E C P L S T N
RANDOM_6735 T E P K E P D N R E T A N M E S P G M G Y H S F P G E T Y E S Y F A G N E A V G N L A S P W T L E I T V L V P L
RANDOM_6736 C V E P E N E L A N L S V I K F A I Y Y F S R Q N F P W F N I A Q T T S E E A C M K V A G G S P A Q N G I S Y F
RANDOM_6737 I I G D L A F F V S S S Q R M Y L K A D D R T D A G A G G T P R T G F L T T P G Y L N V S K S Q D R P G L V R A
RANDOM_6738 R S A G N G K F D A H P S T C T T I D E V L A M Y L R D R D A V N L A V L C P Q R G S T P Q K V G G K E S A I F
RANDOM_6739 Y P I V F E C K E Q D K W V N S R M E Y S E D P L A V L G D E K V G P F R E I T Q G K V P I F T N V L Y D P G D
RANDOM_6740 M Q Q Y Y C T L F P C V G W V D P E T M R N K R I L C E A R L L L K A I K N Y P S A G L N S I N G P I R F Y D G
RANDOM_6741 D V N M F E T G P Q F G Y F T T V P K L G Q M L V W A L N R T H T R S L N A Q L L S A F N K G S T P L T F E L R
RANDOM_6742 A T F V Q G F G A D S V E S S N W Q L Q T C Q K C V M V R H I K Q I F F Q Y M G G D C T R N M D V G S L S A L D
RANDOM_6743 G F R S Q F Y N M L N A G L D D T S R S S E E Q L D L S K T N T R T G F M Q D M E R V A Y P A G G I Y T L E V Y
RANDOM_6744 S P D A S G Y S N T C T N F G A S M L I K I W K E L A T G V M L E Y G D G E E K V Y I E V N S E L V E K C P Y R
RANDOM_6745 S N D M S D K N H A Y P Q D Y K Q V S F L G V A T S L L S P A V A V I P L V K D A E G I W E I F Y D K E K K D R
RANDOM_6746 I I E L V I K E K E A L V Q T S Q W A C S P E S D I N D P R S S D L E L A N R T E E D G F E Y T F F G T T A Q A
RANDOM_6747 Q V F T R K H I G I D L A D T H V A A N Q W P I A E D Y T E V V L Y K V V G L P I L N G Y S P F P N P L C D T N
RANDOM_6748 V E V K I F P S V N D I K G V S R E E D A S N A G T G G N Q Y D V F V E K V H N S D N G E P V A K E L S Q K E P
RANDOM_6749 G L T V A T H A A K G F R S E P A T N G G V S H P H Q K N T Q Q T A R Q N G L K L V Q L K N D Q L F H V A V V E
RANDOM_6750 Q H V S C H V K R A S T P Y Q V N A E I A Y S P S Y Y S P A H R D T G S Q L G M K R P P P R A D P V F G D H S N
RANDOM_6751 T I T C E A S C L R L P L K S M V K G L D C G N R M A M R T L A V S G D F S V T L A N P A L K Y S Q G N D T T S
RANDOM_6752 S L E V G L K G V G E Y S L L E K L D S V E A C D A S R S T P D Y H T V V E Q T Q P W A P G S L V M H S G R L E
RANDOM_6753 S K D R N D G N Y N Q P D N D I I K F L M V A C R P E M V A N A M R G L S D L E H I M A E E C Y H F P M N A R N
RANDOM_6754 F E L F E L S C V F L L A S P P F E W A Q M L G Q N W D L A L Q F I V S N T T Q G F F Q K C S A P V A L T P E V
RANDOM_6755 A P L Q A W V I L G F I S S I A F I L K S E R E Y S Y T N N S E P G D N K L L G A R N V T D A T D Q R L Q M I N
RANDOM_6756 P N A A L K V A R K Q P R K D L Q T H P C P V T W F D D A R A D C I T P V G L K E G G S Q N Y Q E S Y S F E E D
RANDOM_6757 A M Q F S R F S Q K N H P Q G E P H A N D T Y L G K F I W P T V G L D T P D L I D A S T A S K N Q V V D R Q I A
RANDOM_6758 I G F C K G D D C A K R D L T G F H L G L T S I I F L A Y K E V C I P V G K F C D G E P Q L D A G S I G A T L L
RANDOM_6759 Y Y L Y S S G E G S R R K G G G L F A I G D D Y Q L D V W K I L T R L A S Q T A I L L M L P L L T V D A P Q A P
RANDOM_6760 S H F Q T P Q F L Q I R L V G N G L A V F L P A R G A V D F A H T V P K R Q A G R Y N S P A Q M R L A C M M P W
RANDOM_6761 M R A G D G G I S S I I E E Y T W S P E K T S I N A L A T G L L Y L G L M A C E R L T L T E D R S R D D L H F G
RANDOM_6762 Q I Y E R D C G S N K N T S N R Y K K R K N G N K P G L Q T V T G N V C Q N V G I V S K A R E I F L V I V A F K
RANDOM_6763 H Q R N V N M L S Q D A G W K T D T S R G R T F N N S V A T T T L T F E R G C N F F K P E N G P T I A I E E E Y
RANDOM_6764 R E L L R I K T K K L E L L I P V Y F W F E I S Y R V S G I T V T P T C K I M D T D A P L T V H S K S E A R A S
RANDOM_6765 K T D Q M H Q A W I T G E V G S F S C C G D P P V S L I K R Q Q H E N T K E N T R R I Q Q V G K W T D L C T V C
RANDOM_6766 D T A G R S C T T C N A V R E A P N E I G Q Y G S K T N F E L R E Y F E I W A L T E D F G E E C P N V R D L D Y
RANDOM_6767 R N D S D K R G Q W R L G G T V G A T A V G F S S S G R P N G T M N E L P T K L F F A E L G F I N V M I I K I G
RANDOM_6768 K L F W G V C S N V P C V M S I S V A K E M A D Q D A R D P H G G K L K A K E S G Y V A P S G T V N D L E G V P
RANDOM_6769 R V E S C V D G R P V Q K Q L P E V A F M N D T A D L H N P T V N A C N R Q Q L S G R R E A G E G A L Y N S S V
RANDOM_6770 I R V N G E C I T P D L R S I A S C L I M N T D M F E V D G K A R E A G P L G F A G N K Q T T E S P M A P N Q V
RANDOM_6771 R Q E Y C W A S I G L T Y N S V W R P K S F V N S N T D K E G R E I S M I S L N Y H I K Q P R G L I L D C H A P
RANDOM_6772 M R E E P D L D L F E N G A V C A G R R V F T F E Q L K V A R K E D I F S R G E Q E I M Y G H H I F W L P D A N
RANDOM_6773 F K S V Y R G F T C G V V M L Y F A P M K A L F E G L A S N F V T H I G E Y M E R P K G I F S V E S L M A A T R
RANDOM_6774 V V K L C L A L S L A N D L S M A Q T I N E I D Q Y D A R L A I R L S C R A W S S T G N I D T F S V L Q R N L Q
RANDOM_6775 T R V A G C C I C C S E A D R S E P E Y T H N A Q D D D G V V Y A M F K A G R D D S K P E V Q L V E A A E G F D
RANDOM_6776 Q A G E D L P G V K S I M A V G M A A F F Q Q T C S G N L G L H L Y S S T T Q T I G S H I M V S P I R E L T T L
RANDOM_6777 Y K N N K V D A R P Q I F T T A K S K Y H H K N T Y M V S A E I Y A W A A A T L L T T L A A D F E Q M V N A Y R
RANDOM_6778 Y K P E V C I M G P V Y E V E F V Q V D S A I D Y C V T E L G F V V T R L T G S E M E V V V I Q L G V R P K I K
RANDOM_6779 R T L K K N I K G D L S T Y I I D R F R A R G F P T G Y E F G A Q I S D A T D F H S I K H L V P V N E R V I V Q
RANDOM_6780 P G C S A Q A R N V V K M V I S L N L F Y A D M L P A Q E A E L L F F N V A H A V L T L A Q L E L I I R P L C A
RANDOM_6781 P F F L N V L L L V V F S D Y A C F D Y T T D Q R A Y L L A S W M I D S L F P F K N D S L Y T T D S L A A N E Y
RANDOM_6782 P R Q L Q D P K W T L M V G T S R C L K I P A P V R A L P A S Y N E F P V M G N K I G S M E A A F L K N E R L S
RANDOM_6783 M D H I D S N S G Q T Y R G V L K P R A A P L T E E A I V L M R G T P I N T S Y Q S N T T L R L H V P D N V L Q
RANDOM_6784 P I G R A F L A M L V A L V M K P T S L V R N E E S P I E T A L R S L K S N N E I Q V L A Y Y S A Y G T C P L P
RANDOM_6785 R T P Y C S E L R A K Q S Y L V R A A E S G N G V H P V L M G H E Y E F E P K F D K V F V D G D Q P T A N F L L
RANDOM_6786 V T T E G E L S A E P E K W R T R L G D S V A G E G T E A T S Y E L I E T S A R I N S A A H G K V V A E A G Y E
RANDOM_6787 R C P D K D L D K S N T N D L T G S A A Q L Y Q Q L E T S I D M Y K K L D F Y L Y D F V R R T E C A Q E W E T D
RANDOM_6788 A H H L A L M H L R S V H L M K G Q Y K W S L A L E P N V S Y G P F M E G I A V F F N C F M H L K P I G Y S I E
RANDOM_6789 G D Y V T Q K N L A V R F A Y R R L L L R S A A I D L S S G D Y V T P R A L K T V R A N H N N L D N M S E Q T Y
RANDOM_6790 V P K R Q K V T E L A N L V Q S P H A E T E P W L Y L S I L I S T G L R A R M A F L V L N N G L E E P P I E T P
RANDOM_6791 T A G V S Y T L L L R E R S Q N P Q A E F Q G I T L S M M D T I G C W P G F T L D F K K A F E E A A I A P D C R
RANDOM_6792 I E N P S D S Y F L L P D N T Y S D G D Q R D G K K G R T E S T D L G P E E G K V G V H A E F F I G M T H G L D
RANDOM_6793 K A V V L A K A R S G L L N Q N M A F A H N V S I V T W G Q Q N K I K S A L F P D T R G S V G G G T D H R K G G
RANDOM_6794 R G T Q E R E S T D F C S D S A F P F V L L G A S G L T V I P Q S E K Q E A P Y L D S V E T S V L E C K I E A G
RANDOM_6795 T A C G A A A V L G F H N D Y K K T C I L S G G R A S K M A M S R V A V C V L A V I V K T V I G F L G V D L G K
RANDOM_6796 D A K S P G A V N R I G M I S N S E I Y S F H I K N P K E D K I D E A L Q V P V E L N P L E Q V D I T G L Y H D
RANDOM_6797 V M M K C A H Q W L A P V G L I N K D D R M R G A A T D Y A L A F L S P K P F Q W T H F E N G L E S N V L E I S
RANDOM_6798 R K I D Y N D D I D D A G Y C K T V K S R N K E V I G W N P D D P I L A L Q M I G C S I V H Q G Y K M I C L V L
RANDOM_6799 T P Y G T R K N C G E H N L D A P A L S E G F N F K G K S P L R E Q L L H G V N Q N D A Q R G N R S P T V P Y P
RANDOM_6800 G A I T T L S V V D C A D F Q L E E E S V V S R A A G G K S T C N Q N E S L V A E S Q D Q L E E T E G D D G P I
RANDOM_6801 D F W T T P G Q Y S S S I G C L M R T P Q T V N G D G H A S A V G F H P I D D M Y K S G L A I G G G S Y G C V V
RANDOM_6802 L P V D A D D D L V F T A S S Q P L F T K K L M R N Q C W T L V K V K S L R E A S K G W S C R A G P P A S V N P
RANDOM_6803 A T F T H H L A L Q N W D Q S R I G D S V R E V L V A P N R D A F F P G K V G T T V Q T T C S T Q I R V N K T E
RANDOM_6804 M D T N S V D T T E W E K Q T Q V L G A Y M I F L W L P T Y V Q Y L A L L T K L Q K L R Y N L N A F L A R R I I
RANDOM_6805 A I D D R E T V D S T P A D K I Y I K N K P T A L D G T G S R R K M K G I T G I L R T R M V S E I L G Y P H S D
RANDOM_6806 E V G L S S L L I G R V D K A L F P R F K C A A L S Q Y D V T P S Y D P T M Q D T L S M Q K T G H T E T G Q R M
RANDOM_6807 R G A C L F L D H V P A A S F E D Y L G K D A W R L L P F I H F C P F Y A K S V V I L A A E R Y T Q S A E L L V
RANDOM_6808 R S F Y K V F D Q S A S S M L L Q V S L V I D A Q S G C D M F M Q D L V D A V D P K V A V L K N L S N F D G F F
RANDOM_6809 E A I F S L H L V L Y K T E Q E D E T L Q A M A C W F G A Q G I I S S E H T T R P K Q R V Y N P E E S V I Q D R
RANDOM_6810 S S N A Y A V L F Y K L Q M R L P A R V S E S V P S N N L N I H L Q P V Q Q F T K T V L L A A V R V C E P G I G
RANDOM_6811 D G L A Q I A Y S S H D L C R R T K Y R I I K R I K P Q L A A T S E T A F T K L P C H T L I G S E K V K L P P P
RANDOM_6812 G V E F P L K W E W G T L F P D L P E R G L K L N E G E Y S A L R M K F V N L T S L S A S N V T L H V A P F K T
RANDOM_6813 T K S E Y D G M R E N R L Q F T I E L L M L K I Y S G K V A I L E A T L G D T L K V F A E K N F S N K L A H T V
RANDOM_6814 P E V L S R M L S D E F F K L L H R N D Y D L L Q N A D T Y A S E I V S E F T V I A F M E C T N W M Q E S V L K
RANDOM_6815 K D V L A E L M K A S S S V K V Q K L I T S A S D Q Y L P N I L E E G D L E L S K M L M E N V D D A Q L K E L L
RANDOM_6816 V V T S A D G F D F A S S P F Q I P H E L Q Q L L V G G L D W C R Q F D K I A V N G E T D P A L A E K A E S M H
RANDOM_6817 V A I R A A N D P Y T G G V D M L L S K K N W F R V A S E G K I G V F M V E W E F F D L T Y E K V E N S L N G G
RANDOM_6818 W S V R P P D V Q W I G A R N P F E Q A L H P M H Q V H S T S F L G D T T R I R T K F D T C C T D C G V Y Q D K
RANDOM_6819 T G H V R P D A L S L G C S Q G E D G H S I G S H D K K G H I A G Y R A P F Q I M T A A P A S W Y Q P D G L P V
RANDOM_6820 Q R G K R T D S F D V V R Q T E L P I Q K N L N F T S A D N K K C V Q S G V E H I G S F T T V S R K D N E A S K
RANDOM_6821 T A P W G A P V G C A G T L S N Q A A L E L R W V L A A G A L W T P H Y H K F I M K T N A H G H E I D L G L L V
RANDOM_6822 S I G K G L S E T R S G Q V M S L Q I S I E E S Y D S D S I D D V N L K N M G S H L A G R D S S T A P R K A H D
RANDOM_6823 D N S I A T G S N Q P H V D L E A E N A P V Q Y E S V L P D S I I E K K H D P G G E H I I Q S P I A R M E N T S
RANDOM_6824 D E T F E L R P S D L M V R H K A D A V R T A M I A S L V T S Y E S E H S E F L D L E V E P Y V P A S Q K F E Y
RANDOM_6825 C G I V I G G G S N F K I A G P E G I N L P P F D G T A Y L G P L V S A Y S T E P V L R R Y T I A S N N L Y G R

RANDOM_6826 S H Q P T N A P L P L V L L D D E V P K K Q I Q T T S A D P Q F D I H A P L Q K G R R D P R A F S G K V M I P G
RANDOM_6827 K R T G V D K F S E G G T A G E A Q I P I T A Y Q E A Q H L A G M W E E P T G S T T T D N R P H N M E K R P M A
RANDOM_6828 P G F C G S I D R L P K H P P L A R D L G G L Q S T P G E L R H Q V S S K P N L E I E S S M N T W D E E D P D D
RANDOM_6829 K E P Y W I A T A G L A E G F T N E P V E R V I A A R K F F R L I C D F R L K I N W E S A F R V P N K L I V H L
RANDOM_6830 T L S I V L A C S D V T V F Q E S L D H E N H R Q A Y A R T L Q L K L P A Q I T V I S L S G F R R Q L Q L R M D
RANDOM_6831 S N M G P S A G T T D E Q S I N T A E N C T P F S F Y V N L Q N V M W L A K G G Y E F E L Y R M K R L D G T K L
RANDOM_6832 W N L P E D S N L N K V I K L L L N L A K M A P N D G E Q R H T R F D N K L H L R T G A F F Q K R K K N N A E A
RANDOM_6833 A L Q S G C D A L T I I S Q I E G G R F R K N L A G V M M L I L E T V V P N K A V S D G S P N Q W K K K Q P Q K
RANDOM_6834 Q T T M C K M L M G L K L S V I E S N T L D L S V E T I A R C V T N L A G I V I T V K Y W G P L E W P K T L P K
RANDOM_6835 T K A T P V N V T R R L I R H G T T R D E Y V V V F M P S F N L T T Y I L R S K N R L E D E F V P I R G E Y N H
RANDOM_6836 S G D R M A N R I Q L L T E I S R L V I L L G E A D Q Y L I L G E G P P E S A H H C S Y S D Y E Q Q G M L S D Q
RANDOM_6837 D K L K R D R G F L L W N Y A G Q R S A V P D G R I L W G S G L Q I A A L P A S K R C S R S A D C G D N F H H N
RANDOM_6838 K D P E A G T E H G F Q F I A N E I Y K A T D V I G I P G L F D P V Y G L E V M D L D D R V A Y F Q P G D I T D
RANDOM_6839 Y E S V Q N S G V F E R K V Y N P R S E G P A N A S G R D L C S V L R V S Y W S D D P Y E P D S L Y C H A L N D
RANDOM_6840 H F I R V V A G L P D E M K I L E R K N L A L V Y S F R F L K L N K P N F A K L K S W G I Y Q T E A Y S V E G K
RANDOM_6841 S N S L V M G F H A D I A F K S S P Q G L N Y S P A K H G V G Y Q L V P V P P A S L L R A V L A K H L K V D Y G
RANDOM_6842 S D Q Q N I P G K L S S R K K F A L V A S Y V W S L T W L L T N N L N G L Q L T Q R E N H A A S V K E G A A R N
RANDOM_6843 A V T Y D S E S A I E I A G G F V S Q P I R S G R P G R F P I L Y S D D M K D Y K S H D F L I G E S G V I S T A
RANDOM_6844 S R S V A E Q A M N S F L L L R D I T L K W D D S I V N E D F G G D Y T D T I V G I P I P E C E L D E F R E S T
RANDOM_6845 Y C D E D G E D A L D G D I P T S V G R F D R T E A S G K H E R F E G T F D A A K S L G D I L L G K Q S V L W A
RANDOM_6846 Q R K W R I L S Q E K V S Q G M Y T K A G P V I T R L A C S N S K K K F I G G C A N L L A T F N R Q F S D L I I
RANDOM_6847 D V M T N R G S G D A K G V K S N S E C S I I I D A P L L Q S A Q T Q I S S D G R I V A E P K H K N N G E L E F
RANDOM_6848 D H K V C L G K A G T E R C N V L I I E L S M L Q R S L S K T A A S F S T H N S Q E F E L R S P T A L Y T R L Q
RANDOM_6849 L A Q K F N T P D V N V W C M S F E H E P L P V N I A S L I P V P G K P T E R W D N A M A F T F M T L R V S A G
RANDOM_6850 V V Q P K L G A I Q V N K K K P Y L S P K I G Y E A T N S Y K R S F L L R K R W A R M I N V D I N S T Q I T A K
RANDOM_6851 G I S S E G T V V L D T C Y N D Y K N K S A K V N T D K W L E D I P E K N A Q S G A I F S R G V S R L G M C Y L
RANDOM_6852 T S A F A K L P A I T S K K G M P T D T G C M G A S K I A M H A G N Q E L M V P L S G E I S E C D I V D A A E Q
RANDOM_6853 V F K S M A S S N S P P Y P S K Q L L Y V I F E V K V Q S E N C G G Q Y D S I P G Q S E S R T Q T D T Q M Q N D
RANDOM_6854 Y R E V S S K E G Q I E N R R V T P F E Y G D C A E S D N H F V R D T Q G E G P Q D R S R P G E I W E E E K V T
RANDOM_6855 R K G G H L Y F I K F C S N V C G V V L L I I Q T L F K K V N F E D K E P P A L Q T Q P V Q P W A A Q M A K A V
RANDOM_6856 M A E N G I R K A P M F A V E T L P S Y Y I T G L G Q P L R N Y R N F Q P T S I A G A G T S G I C D S V K G G P
RANDOM_6857 L I P D Y P L E P S Q S A G Y Y N P D A N T G S G V M C C K A R P F V G D A K G E D D G E A K Y I I N D K W A P
RANDOM_6858 L I V R P A P K G T L R R L G K E P R I V D R R F K G Q L R Y M Q W S L R V L K M R G T F A K S D R E Y L R M L
RANDOM_6859 A M L P T V P T S G H A K S F H D P S V F I N L D Q A V Q I A I V S I D R D N H Q F A I A Q H P I L E E N R R I
RANDOM_6860 C L K G N G M S L I A M F K T K R N L S P A M L P P R T R M E N K A V Q P E P T Y S S N P Q S L Y A K K Q C K E
RANDOM_6861 A A D G W I T V Y A L S P L V C G I N D G G P G S S M R F Q E Q A L R V I S C F L L V S T K S E I Q E T S L L I
RANDOM_6862 Y P A L G F G P L P S S Y L Q L A D P E I S V Q F F N A V S N L G K E P T L L N D G D K A S I E P W V G S G I E
RANDOM_6863 E N N Y C G P Y N C Q V V S V N D E W K I N L Y I I K C A N R R Q D N E S M D V E N D R P V D T G S S G D L V D
RANDOM_6864 K L G L K D K L M K R V D E V F L G W P A Y Q T G R K R D D V R E G V R I E Y A R L M P R P A N K K L I T S S F
RANDOM_6865 S S V G L K Y G G A P G C N H I A I F K M P C E M D L G N G Q K R V A A R A E M A R Q D I N R E E Q G L A A V I
RANDOM_6866 W A P A L L W N L V V V H P P V L E E G V T C F L F F K K Q A G F G I P L M S G R H C N R H L K Y C V L R F L R
RANDOM_6867 N R R E A S T S P T S G A F T F T Q L V I L V Y T Q E S G R L A T S P D G G K R E S F L R A P A R L K L T S R I
RANDOM_6868 K I S S E N R H V R V I I L A N L T R I G K L R G D V Q Y A R P P P L H V L Q E L Y V G T F A N C V D T M F L A
RANDOM_6869 K V P K F S C G L W P S A A N S D Q T M A D Q L Q K P K A Q L F F K V K A L E F I E Q H P V P V E S T K T I E Q
RANDOM_6870 I I G D V R P V V V G G G D G P F S S Y T G V L F K Q A A A G H A I W P R C L E L V G S E P E K Y E A D N S L D
RANDOM_6871 Q V G A V P I S L M S H D T L N A I I S S L V C L G P D R N V P A A N T S R N K E H Q T S E L S A D H V G K S K
RANDOM_6872 V A I K L D A G L E G A R G Y L T R A L Q V V P F S Q D Y T R H D D G D S S I A S K T H L E G C N W V A S V Y Q
RANDOM_6873 L E F G R Y E G R V C A P Q F G D D I D T S D G T S D K R G Q I I R W S D A N V T T A G S N T A D Y V K I I A H
RANDOM_6874 S L T L S A Q I D A N V N C K S I H V S E T L E R E L K E S K E Q K A H V L P Q A R V L A A K G P C L R S E K D
RANDOM_6875 W L Y I V V N N L F E R S F G N K C H P Q E M C F A E K S R V V T G I E E Y V F H D E G L E L L R V R E M P M E
RANDOM_6876 H S S F S V N Q R L E T E S Y V F A V E A A Q A I E I V V R L K Q T N Q P V E T E D F L Y G P V L L I K G G E F
RANDOM_6877 E Q V V A L R I D F R L L F L I A R M E T K P P N E T S V R V A R S T D D P V A L T N E E E A C Q S V C L N V T
RANDOM_6878 P G K K I T A F R R H V P L K V R I R G G Q L Y E K N L V K N D P Y S Q D N L A L D Q D T W A L H L P W E G Q S
RANDOM_6879 R W G A F L I E K Q Q A S L G A E M E L P G S T L E G S S T P M L A V M F A D N E E L K S A A G S G T V M L L D
RANDOM_6880 D V N A S T T N M D G L A F K K E A K C T V T E D S S F G K S H K Q E I Q F V N R A E L H V R D S Y D V G N T Y
RANDOM_6881 N M G E E N L N E A K A K A Q L I I L D I Q N G T K M A P F S L F L F N R H K N E A R M G F P T R V I N G F N G
RANDOM_6882 H P S N R L L A T Y T N G L I I M P G H P D Y Q P T A P D N W S L S S R T W L L Y P L A G A N D D P K A L I L H
RANDOM_6883 A R A Q R K F S K G S K Q D I R V S H N L P M I K R D N G R G I R K S S Q V C I Q V E A P T S E E V L L K K G A
RANDOM_6884 I A I F E N L V V E P A T G A W S H L Q Y T C G V N V D L Q A V D T S K L L V M E G L Y A I L D C K S D R L F E
RANDOM_6885 L G Q P G P E L V G A S A T F Q T V T G K G Q D V G E Q I Q V R D A N L G K P R P P V L M G S Y L P H D T L L D
RANDOM_6886 C S I I I P F V T G K D N L I L L C V A A S G H D L F F R R T Y V C L G Q I Q E E I H W F V P S R E R V L P S K
RANDOM_6887 R G E M I R K R V Y R N A H S T I M E L F Q T R S S C V H W L P N V R N A K G D L F M Q I R D R V S M G L D L L
RANDOM_6888 S L P M Y V E E D T D G K V D L Q P I R S E E T L V G L M L L L I K H E T A I G N S Y D D E I G P D Q I M S A E
RANDOM_6889 A L L K Y P A W W E P D L A L T G K I K K S G N L G H T S T L V F T Q R L Q S A Y A I G R L I I Q P T N T I V R
RANDOM_6890 K N V A F S K I F G H L G L D D G I G E G Q F L S I A I F Q F D K R D D S G T I C K G K T C Y S P K L E Y T P W
RANDOM_6891 I L P K H L I P L N E F Y F Q A L V P V E S E P E T K A P K A K S S C K L C T F D C V S L D P D M V A R A K E A
RANDOM_6892 Q K S E P R V V G L Y L S P S Y D G G G T L A L A E S P E F D H T V M L P D G P L P T G W P R S L S E G S K G R
RANDOM_6893 A T S A A R F L T S G G V W T A Y Y D I S P Y D C P M Y Q A L S T A I A Q R K L V N S P G N P I P R T P E G M Y
RANDOM_6894 A N S T T Q Y I V S P C L H D I Q A G K L G L Q F H R E G R S D A A I L G G M G P R R L I P K A A Q Q K A R W D
RANDOM_6895 I A R Q S I T L A P S M S T Q K S I G A R F L T I I N P P V C L T L A G S N A A V I V K E T G S D V T H T D T A
RANDOM_6896 V F G N S Y S I T M A R T S L V A A L C S S K Q R K M R S L I M K D G F K Q I L A G P M A E L L Y P Q K Q W S K
RANDOM_6897 S P I A R R M L S P F P L H L R G A V G S Y D C I R H L D D G R A E D R A G I V S H T E T C I D G V S H F S Q S
RANDOM_6898 F L I Q K I E D D A W P E Y S T E T C D G T W Y Q K R Q A D R V G A L T L Q K E S W L G P F M T W K H I I S M E
RANDOM_6899 N P A V G F M L T N R Y M G T L S L P F V V N L Q I Y V G G G V S A L T Y G D K I S N N E R Y R T S G D R G A E
RANDOM_6900 L G C D L K G E K K I L K Y L S S D V R V V Y L G T V F V G L R D K N Y N G E S A T I P G H V V S G F Y V R I Y
RANDOM_6901 I M I I R T E F M P L C E V V N P N M T W F V V R L N E A T Y P L G H T E A G A V P D L E L L G F T K S H F D K
RANDOM_6902 C D K K G S L G E A K T L G P A K F S E L P R A P P F I K G H S R A M G S Q S A L R S G Y S E A D Q S Y G L R L
RANDOM_6903 R K Y F G V G S N I G I T S Q A S S Y F S K A A R E A A S A S V S E K K H G L K P C K A R H Q I G A I I L V L Q
RANDOM_6904 M L D P Y V F G K W K Q M K S A L V P K S S L S P P A Q G W P C P L D L L S Y R S T G T I G L N P R L H Y D L S
RANDOM_6905 R P K D A T N S L T L D L S V T L V I I F Y N V C V T M D G F E H K G R A N V G S L V H N A C K K A N T E F L I
RANDOM_6906 D L H F H T Y P L S K D S T L T I S E K D A E A K T G E V A S T C K F I I S D L M Q A E T I A F K L S R Q T I T
RANDOM_6907 Y F T R V L C L P E N I G Y Y K L I R G Q E T G Q A G S C Q E C H F Q L D G E G G Q Y M L V K Y E I D P A Y L T
RANDOM_6908 N Q P T M E G H P P S V L V N I A D W K E M V P G N P D G G I K Q Y D S S M F W P P M S L G K R V Y T Q I Y Q L
RANDOM_6909 R L T V K K K Q A L G V Y F V Q P S V D I A P I S H D L A M I L T P M H G K M I F L D G Q L I R D V E A P Q L F
RANDOM_6910 V V G A E P V D I W G R D V S L E I E K R D L D M T L L H G T T D G G K L I R A N G V T N V D T S E E L E L S G
RANDOM_6911 Y T K G Q H T V L I A L I N N A F V D S Y V K N I K L V L K S S T F V H F R F P K Q E H Y A S P V L I F K S F S
RANDOM_6912 K P Y T R M M H G I G A T K E L E S L T L S L I P I M D M S V I E L K Q E D Y P P A T Y L K D L A S K L L G G L
RANDOM_6913 P K D I E G L E E T P S P M T I R G D G D L P C G I L D N D G P V E W A D S M P H T A L V T A E S V T Q I K W A
RANDOM_6914 K H G V S T I A P F V K G E L R T S G E V V F K A Y K L C Y Q D D Q T L I K L K D E T I V L V Q Q D I G S C T G
RANDOM_6915 A V G I D N M A A F A N E V R L D P K Q L V Q D Y Q I S L S P A Y D D N C G F D N S P P S V F G L G P F K R L S
RANDOM_6916 L M S R G P G D H E A V L L S E V C T R L V L P P Y T I R A E N C K L D A A I P P E Q K Q R R L L Y A Y L G S L
RANDOM_6917 E D L F T T L G V D I R G A G I S N Q N N L Q K Q A L Y L G D G I R I D L D K G S G R P Y I G G L E V K I S E E
RANDOM_6918 V N T F Q C F V G N L C D I P E L T P R S G R I K I V M L K D S E D E Y Q E H M T Q L P D L T V H L E A K I R S
RANDOM_6919 N R W S I K G V P K Q L T E A L S C D S S T I L I I A E F W P K A L Q E L K Y G R K L Q A I I T S E T W Q Y N G
RANDOM_6920 P C N K T H C G G I P F K D K S G I P N K T G G P G G V K N S H A Q E C T L E R L D G G P Y K I G T T Q I S P S

RANDOM_6921 H F N L V L E G S P S L R L A I Y V L F N A F S G Q L D L S A L E S I K Y G G G K I N M S A G L T E T Y L H I V
RANDOM_6922 R E N E A S V E G L R F Y D M A M P P G S R L L T K Y T N K G G Y L S S R R V V Y K E S T W K K P N K R N Q H L
RANDOM_6923 I T R V V P T R W V L S Y D S R A I S S R I S T I L A P L Y I L I Q D Q S K T N K L K A D E P S L S F H Q E H F
RANDOM_6924 D S A D K G V E K S F F F V S M L V A I G E K M L E K V T L I G V N E M L W S D G E Q Y A C Q L E G T E L G L K
RANDOM_6925 R N N Q R S T A A E T Y D M H K L Y G L L F R N L I D A S H E V D S Q V D N I T K N T A A L I R M L Y N S A R I
RANDOM_6926 F S V R W E L E P G R Q I S T G A K S L D D M G P S F T H A F T V S A S C P A K N K D V T S N Y L Q V N G G K V
RANDOM_6927 T S H L I S N E D H V K S G K S L D T K C L T Q R S M Q G M S I V F K A P A T P G K Y V A N T P A V N P R A Q I
RANDOM_6928 W Q Y I A D F V E K T A T V P R S P A A S K E V K R D Q V S Y P H V K N L G D I V F P Q S G N G A L H N R Q F Q
RANDOM_6929 T L G C P S L V L K V D T A Q L I P P G Y E W A S D G S L T Q E N L V Y D A K E E I R K V E G S R L T L Y G V F
RANDOM_6930 R S R N S V G H D R N G E L S N L F P S V E D T N L E S V G R L S I C P H N Q G S A F E L I F E K N V Y V G T T
RANDOM_6931 T W Q K Q L A S G Q R D L D E T E T N V P C A A S L D S E G R E Y F T G E A R E L E I A K I F T S L I K A E G Y
RANDOM_6932 E G A L S F E T N E S V L E P F C L D Q D L V L G N A E K R E I E F W D A L N V K D Y F E N Q L E K Y K K D N V
RANDOM_6933 V T G D K S F C T V M V A R N H G V C R T K E Y S Q T V K R G V E Q S A E E A V L V Q I W G G L S H Q K D T E L
RANDOM_6934 D R K G S F S V D I I K D H T E V A R H G M V P W I I S L Q F K L G Q G S R S Q Q D K E D A R G E D M P R N Q R
RANDOM_6935 K L Q Q F L W E S C P G P D P T D A S V A W L W D R N S I P M H V I D N N L S L R N A P T L Y S V M A A F K C V
RANDOM_6936 I Y Y T W A D F V S M S I G I G N Q R A H D A D D Y L D L L W A V I C L G Q T T L N K N F S S P S R E E E M V G
RANDOM_6937 P A D A S G K M C S R N I H A R N F A G S Q K I F P C G D L E S G C F S G R T K E K V Q E S L L R L K S C I P V
RANDOM_6938 A I K S S W Y F W K T L T A S A F L R V F E Q K V S H P S D D K A N T A F H T T D H S G A S Q Q A A M V D E R K
RANDOM_6939 A Q A K P A R M K G K K T Q M F N G E E N Q I A Y L C F A D Y E Y Y K G R L D D N G V S I N Q Q G Y Y C E P A G
RANDOM_6940 R L K S N Q F R Y Q H T F A C V T S R F E A S L K V V P L L T R T S D F I L K C T G D T I P S F A D D E Y Q H V
RANDOM_6941 Q T A G F I R L Q P R K E L V D I E H R T L Q C L L A E T S T P L V A G D T D S T S A K L T L A T L W H M T K A
RANDOM_6942 T L N P R P V Q A I A D A D E Y I I V N F G E M M I T Q C A E Y D E I K A L T L M G T Q L A K L P R S A D L Q T
RANDOM_6943 Q L L L I E S L E L H F G D E T R A G S L S P N D L S A Q A K V V Y M E K S R R P K V T S G S K R D K L G A L R
RANDOM_6944 Q L S L V T A L I L N G T A G A F L Y D E D A P L E H L F R D S D R K E G I D A R G I H I L N M F I F Q I V N P
RANDOM_6945 V K R L W C V Q L A F N S T S S E D F Q A N L L Y Y K P Y K S V K S I G L Y R G I R R F A L S G A Y K Y T A W P
RANDOM_6946 V S D I V V K D N T T G S K R V A V G H N N A A S M F N T K N G N P A L A S V T L Y F S L S P V V P A A M I I L
RANDOM_6947 E N T I T L D K T F A E T I N A L I C A S L G M F A P G N V W N V K G F A E S H D P H D M K G L H P N E R V Q N
RANDOM_6948 Y G P F W K S T N S E W S V E E S A V M D S M P F V Q D E E G T V N C N L N T D Q S W E V Q H K V T N T Q P V G
RANDOM_6949 K Q H N E I E V M D P F A Q K E P A N P L V S E I S G R V E Q L L T S V E I P V A N G L C D D D K V G L L A P N
RANDOM_6950 A I P P L P S A E F T I V E L S A A G R R F D H R S N F C G V W N Q L S E R D L M V E A N G S Q R V G K G P K R
RANDOM_6951 N Q E K T G I N S P R K L N A T A V V T A L S P L N I I K S V T K N G Y N G K M W F R S L A F F L T I L N G L L
RANDOM_6952 V E G L R L L D W A V G C D P R E L R T T P E E L T G T S G E S F N L E A T M E D Q A S G N Q Y M G S C E S N N
RANDOM_6953 Q Y L L A A A S Q P P V Y G V V R E R Y K G S M T A T T D V F A L P L D T L S K E L A H Q G V V V Q P D N N S E
RANDOM_6954 L N R R P R N T E G S P R H Q V G V V L N Q F E P L Q T R C P S P T A K V S W L M N R T R G S I L F S P V L V R
RANDOM_6955 I V G A K G E P A R F L C N M E I S K Y D E V I T N L I A A L E E A I E S M N W N Q Y N A P T H K A I E A A E C
RANDOM_6956 E Y E G T N S Y S K N L V S K A R A V K S T L L M L T V G A A L C T I Q H K F K D G K Q K S L R M T S F Q L R R
RANDOM_6957 E L Q G G T F L A K Q I S M G H L R N F C P A D A A R K S S S A L N H H A D C G K V L G F L P S M P A R L E R L
RANDOM_6958 G P E L G V F N P S H P K K A S I S F Y I M A S L I D G L T T N D N S L E N R S V D R R S G M D Q R L S S K D A
RANDOM_6959 F E K A E L N G G M I V G N E K A A Y I L H R L L V F H N G E S P A L G S D P R S I V Y T K N N P R D V L L E G
RANDOM_6960 Q K L R T L L L K F S F P Q T N K Q P N F L R F L R S N P I K L L P N P T Y G F F I R P N R R I E A S L V I A T
RANDOM_6961 K R I L Q I V G H L Q E T D E I A A G G M I L I N T L G S L V P G G T G E G I S N I L L Q Q P A K P I E P A E K
RANDOM_6962 E A S L D T A A G C L R L G A A V A T T S R S A Q L V S W G A S K C Y S N Q T A K I D G I E W R E T L L H E I K
RANDOM_6963 K N E R I G I F I G R R L M L I H S G I A R V H L A G G I T L K P I I A Y I P Q E T S D I A A L G D G G E L T E
RANDOM_6964 T Q Y V Y N D P W D E C V T S S L P V K G S K Q I T M C L T L P F E R S G K E T A A N C F E I T S M W T V M K L
RANDOM_6965 S R M V M I S P P G N A P S P G S D D W E G D N Q S S R Y A L S D L P K E E V L S D E H G K R Y L K I L I G L R
RANDOM_6966 F R D D N Q R T A Y R G I C R P E K R A D W F H Q Y N E E Q R T D I V R K I T A L K Q G S A A G Q I T S I R S V
RANDOM_6967 A D G D A R T K P R P A W E P L I I P S F S N E A G I E F A E A L L D R P L R S Q G V D T A N G C E F S S L T K
RANDOM_6968 E S D N S A F G V N E L T Q A V M V W I Y A D V A Q K A G Y A S K I G L E A P G D Q I K V L K A F T H S G S K T
RANDOM_6969 C F Y D T Y R W L P T A T S G K E S T P V G A K N I P G V G G R A C E S C E T D V T K G E V V H Y Q R T T K A S
RANDOM_6970 I S H S P G N A G T K L N A Q I R P G D G W E R P I D S L D A H Q E L P S S T E A Y E E P L P D S S Y V E S N D
RANDOM_6971 A F K L V G Q C L R A I Q V F K F D V L S I R F F K W D H D K M C V S V P K P S L V K G P N N Q D L I G G H G A
RANDOM_6972 K Q G T A R I M S L I Q L Q L C T T C M G D F T I W A W K S E S T Q T C E N R R E D C W V I T A V V S A L D V G
RANDOM_6973 L D K P A T A S T H F I M T F T E L Q D P D H L G D G I A G G L D D E G T Y S R K G Y A Q E R F R Q H R V A N D
RANDOM_6974 I G K Q I S C T N N K E T E I D T I V S Q D E P L K M M G M L V P M T L H Q I G H F S I Q L I L T N P M R P G E
RANDOM_6975 T E K N Y K G D T G G R G L V Q K R A R T Y R G Q I G L I S E V L H P F K L L M R S A S G L S D F D D E D Y E L
RANDOM_6976 G S L D A P T G P E P A N N H L L V E R Q V N F L A L D K E L T E E Y E K L N T I I V L G S N G M A A C M P I W
RANDOM_6977 L N R S W F A T M E T E E S I N M S T S E T L I L V T F P N V R R T A A A T D R K K D A P G T G L G D L G H V T
RANDOM_6978 I P E Y G Q R D D G N C T E N N S P L I K L A D T A S H V R L A V W G A V T I D A A V D D A A M Y I Q A A K N L
RANDOM_6979 Q I L P E L A N G T N E Y H R A M H E S K F V Y P P K R Q L E T L G G C P C V F S I R R A N D S V S C V P Q E V
RANDOM_6980 F S T H I L E G K L D M H T V I Q F T T G F T I C G F S T D I D Y V T V I S Q R A S Q D P Y Y K P R I Q D S H K
RANDOM_6981 T N D E S P S G N L V I L A G L C A Q T R A T H D E L L I R D A G I L G G T V Q L A L I S D R V I D I G K Y V S
RANDOM_6982 I S N A R P L I F N T R L E W M C L H D V S Q R S G G L H N P P V H L H P T V A N D M I V R L V L K G K T W Q N
RANDOM_6983 V S A N Q A N P Q L I P E G P H T K I C F G L N P D N G N N L L F E K Q S G Y P V D E P S M Q M W Q A K Y L T W
RANDOM_6984 A S H K V D V Q K T Q E G P V D M D A R R E A P F S Y S Q C N D A A G W A D E V M A D L F P W V L R G I Y K Y N
RANDOM_6985 S K A L V N T Y G P R E N D G I P L E P I G F S C D T N A S G G L K F C D T H A A M L F D L K P V S D S A K K M
RANDOM_6986 M T C E P P P C K Q A R K P F V V T V K L G D P Q R G G M G D C N A R S K D L L E S A H W L S F T L S N A N P V
RANDOM_6987 Y T D P V D C L K N V T P V G K A P P P D E L K R V L R K E F Q L A K G M L S Q T Y G R F L I N S T L G A N G L
RANDOM_6988 L Q Y M C Q K G C Y G N E D Y K Y P G S Q I S A P P V D L K D P R G L P A A L P S N Q L L S L R C D E I F V E V
RANDOM_6989 T Q A Y Q Q P G G S E E Y L E T Y K Q V G Q A P P G V P A R S G V K Q W K D T H D Y A E Q L F E D K A K R V A T
RANDOM_6990 S D L T S L A G R P E T V P S G R K P Q E D Q I T S A S H T T P Q D T L K S Q F K L D W T E E S R F I P C T G A
RANDOM_6991 A A T A S R P D P I N L K V D R H E I Q R A P R Q R P N L E I L L Q G H T C G V R Q A A K G D A S G N P G L C T
RANDOM_6992 C T A N E D I A E L N F W K L E T N P N A Q S T G I Y S Q N I N V A L L I K Q Q V E T L E K R K V G T L V S P S
RANDOM_6993 T P M E R A D K P E K G I K G K A F L N G H Y M I I D T D L P S E V L T I D L C A S I G F A E R K N L S I F G L
RANDOM_6994 L L A I P G R N G Q D G T S A L M C I A F V V G K N N R F L C C P A Q W V R V R E G P K P C T Y E V A S E K A M
RANDOM_6995 Q K D V E D L Q A M P G T K L A V G A A K A A L G A F A L S K Y I D K G H H M L A S S G M H T K D D K P T L S I
RANDOM_6996 T H F P D F T A S G F G L I G D L A K D D Y A G D S A G R N T C I K R L I K A T P L A A I N D I N Y E L F K I G
RANDOM_6997 L I D N K A L K Q I T Q S V C L A H G L M G R F F T W P E A L D K V T D Q G A A E A A L N E K Q I E T K I G G D
RANDOM_6998 A I R G T A I K A Y T P L V K D E F T G K S G E A K R R A A I A R Q F Q G M N P I M Q Q Q G A S R R Y C W L T S
RANDOM_6999 R L I P H Q K S S A N P N Q E W S T T T L L G S D I A Y T H S W K N Y K N I D G Q D V P N K E T L L K Y L R K Q
RANDOM_7000 M V L V G D G F Q L I P K A L L A S H L V Q K R V E K K A L I E G G Y M E F P V K T R A E T R L F W A P A G K D
RANDOM_7001 Y K G M I V Y Y M I G D L A I Q K F I F N F M E S F F V T E D D C L D L F D V G I N P T K A S P T N L C S T L A
RANDOM_7002 F N V N S Q I S Q E F C G N T V R I A E R A W A V V R P L V V L R F V K M P G T V A T A S L V A E V S G D A V Q
RANDOM_7003 L G I E H P D F E V G G F L R K F P G N M H A I C D E R D R Q D P G I I I F S N L H R T T I A D E A Q D A G H S
RANDOM_7004 N I I A L I A H M F N A A M Q T Q S M R S C G L V Q P E V F G I F S T E R D T K P K V L H Q V E D K S L I L S H
RANDOM_7005 L V L T V L S E L A G N S A T S D S A V I L D L I S M S V M Y L A P K V D D I Q K A G L F D E Q G V E Q A A D F
RANDOM_7006 L C N V C R P G W N A T S V L E L C A D G L D L D T R R V A G Q S Q N L G K G T S G K T G Y A A I L I F Q E K E
RANDOM_7007 Y I F K H G E L L Y A R A H L G P Q L D E F R H V N E G A C E Q S Q G C H G L M T M E V L D D Y T R V I K A G S
RANDOM_7008 G M Q T I V K G S T R R T F P V N S F G I K F V K K L P D G G S D T G S F V P E T P Q S R S N Y A W G I F K D S
RANDOM_7009 K T T R V Y I D V A E K T M Q D T R F G T N S A V W R C G P L L T P D P I D T D R S A S W L C A G S I D L A G R
RANDOM_7010 N L L G D W N H A T L P M V L K E N K L P K F K E K A D G Q D E F K G H K L Q L V E P G P A T D E T F K Y P S L
RANDOM_7011 V G E G T N G S T N L K R S I T T T G L E Q E G N P F K I Y T I I S A I Q S D P P F R Q S C D C A G V S C E P E
RANDOM_7012 L I M C S N G R S G A H A L P V S H C L G H P T N K G Q P E R G P D L E K V Y L A E T R W F G E V Q L G S P N E
RANDOM_7013 A G K F P I D L T T T I L Q L F T R E S S T P F T Q Y I T G A E R N N F G A A N D P R G L Q L T L V N M E G T L
RANDOM_7014 V R G K L N L L C N D P S T D A N A E P C T L Q C Y G L Y R E M D V L P A A S S G V L L L R E E R F M I A P D V
RANDOM_7015 G D V T I D K T E V I M E D L D R S Y Y V R F L L E E I Q L N L P I A R K I G W A S A D M K L C G V Q L L I I T

RANDOM_7016 P A F P F S S F F I L L A S L N Q S M G G Q P F E K R R R L V S G P L H R G L I C I L E P S E L G T I S S L S A
RANDOM_7017 P D T D D D S G Q N E M I A V A Q S L D V A H K P S K Q L C L P I A D L Y T V F D P R I T Q A Q H V I P I N V D
RANDOM_7018 P V E P L N R T G L V S R E G P A I P A A A S D K N S P V P L W R A W I T C G K K Y P M T L A Q Q L G L E I V S
RANDOM_7019 H K T V R A P S L P W N F S V T S K G A A H K D T P T F T H E K A Y P I Y E T I I Y E A P E H Y E P G E T D A S
RANDOM_7020 F A G K L Y F L R A Q L G A N S F S T P R Q S A F D T T G C N F A I G K A G G N S Q Y D T L G R S R V Q N I E D
RANDOM_7021 T P Y E A E V I Y K A L A I M G P Q L R I E V A I G G F I V E F G L F A T C P Y K G E S M I A G E P D G N P R I
RANDOM_7022 V I E I N K E S S K V H A H G R G N Q D K F G A M K S F G V A S A T E H H S N F M S P T F S S K D R N S K I R R
RANDOM_7023 E F A S G R T E L E K P G D S N T Q D G K L A A K T S T M S S K L E F Y L G G D E K I L L N E L K M P F L G L Q
RANDOM_7024 I P E G T G L E K E G D S R E I D Q A S S G Y M G K F K K I E A H E K V K K C I V V C K K A F H I Q Y H E I E T
RANDOM_7025 L S K A P T N L I A G G E P I V E S D V H S P L C W V Y G R K D R G Q D S V L D M R T V I Q M Y K S M I N E R I
RANDOM_7026 R K F T S P E I F V D T G A F H T T A A V Y T R D L A V V I I I F R S K D S L G G A P R D G S N E A V R M H D A
RANDOM_7027 C G A A G Q G L G I C F H I A A I H S V L G S R T E Y T D V A E K Q Q L H F T H T G D A G Y T Q S E K C V K A K
RANDOM_7028 E P F K V M A V V D M R N D Y A L R N I P Q K V R L K K T F E F T R I K N A P L A S R M G S I T Y Q D F Q A C T
RANDOM_7029 T D Q N K N I L Y A R S T D F T P F E I M Y Y S Y I T K F A F S Q Q P D T V P Q N T C A G D N P P G C K K N T L
RANDOM_7030 V L V C L P S Y S E E A L P V K P M T R K K H S A S K R K D K K P Y L L I M N G I L A M R Y E A S G Y I A G G L
RANDOM_7031 E A R Y S A L P Q G R Q G N M A T L D V L F L I T N T D V K H V S Q V V T E P Y Y I L Q T T E F G V N E N H E P
RANDOM_7032 A T E A C W K T E T T M Q N K N C R P T Y N Y R S C K P E L E N L I T R A T F I G A D Y K V G C N Y D N I I I G
RANDOM_7033 R T N Q N L V P A R E L N D N D L E L F G V I Q P G A A T N N Q L H V F R D N I S P I D I V H V N G T D L A V T
RANDOM_7034 C V S P A P G W G A H H P G S A G V T K D S S F L G E I K S I C E K Q G G G R A L W G V I N D P G I T P I I Y P
RANDOM_7035 T C L P M D P H S E S G A K P L Y I F N I C W R E V D R A V V C I L L D W R E Q S K Q V L A S Q S R P A E C C P
RANDOM_7036 P F T A F G V R C E H H C T S Y Q I D D G N D T K M G D L G F F P N N I P S S C V S K G V F P K S D I K G V F I
RANDOM_7037 V P F K P I N E P F L R N D P F S N K T F A Y C T L G G F E N A K L G N I G G G G F T K T S P E P A E L G E T S
RANDOM_7038 S Y V T E F L A D G V Y V A R G G G L L L A G K L I L A V D E H T I P T A D F S I W A K Q A N C E T A F E A F F
RANDOM_7039 K E Q P C S G L D K G G G Q Q V N E G P S D G Q R N P L P R Y E K N L Q I R N K E Q S R M F N Q H T T P G E E R
RANDOM_7040 S I T K C D N L I T K K V F Q G T L D V L D P H M Q G L P E T S Q T I T A L N K H L M R E A V L R G V F V E K Q
RANDOM_7041 G G Q V L P Q F A R E Q D H E E C T V S S N Q Q W Y E P S S G K C F M N H T K I N T F R A V Y M S V V W V L W I
RANDOM_7042 F N E N E E Y L K P V K R G V P K L K E G E Q Y M N Y Y S K F E D H R L L G L N V E R D A P K C I N A N I E E H
RANDOM_7043 I S V R A N N G G V L H F Q E V D S P V L S V Q G Y W E I A A P I E V P L F N P K L P A V I A Y H N D L L F D K
RANDOM_7044 F S K D T E I T I V Y T A C N M T P R E S H S G V D G W G T Y S W I R E W Q E S V R F A K Y L A E K V L H M L A
RANDOM_7045 C G N D A G E S S G V G G I R C F C P T F N L A V T R V H V D G Y V L F V L W L F Y Y A Q I K T T V K S T N L V
RANDOM_7046 R S K P A Q L E D K G V V P D D V S A F F V V V P A V K N V D L D E H K L P H G G P A S F L T R L S T G N L Y N
RANDOM_7047 I F G F L H S I C K Q K G V S S D T L M A T C F L A Y T V T D D T K N L F M P D N E V Q V P Q V R P V L C I Y Y
RANDOM_7048 A F K A D A K I D G C W A H Q L S A P R S A S K F V P G D I H I R D V R L F S N S H Y L N G N L R L G L K R W A
RANDOM_7049 K M N K F I N V T M G S T Y V V N S R I L G M R L R I P S A E S D T A L R R A Q D R G Q K G E L A G I P A P I E
RANDOM_7050 G E F A V I R N G C I I I A G V Q L S T Y N K T P Y S V F G K L G R V L V E I Q P L R R Q T S K V A Q P K D M N
RANDOM_7051 R P L T Q D P D E Q D Y D T I V Q K E C K I Q I V K G D R F A A S I K A A P G G Q A I V A T P P H P E D T W I R
RANDOM_7052 S W D T M A S L W Y Y E N K E K A C I E S E T S A Y R Y T G E D E A Q W F Y Q T C P D D D L N N L E V V V A K E
RANDOM_7053 M R P R Q P Q G N D I D A N A E Q V Q F V A H E W M F D K G N N Y V K G N L K L N P L T I I L I F M V Y N E F R
RANDOM_7054 N E R P F P S L A A F A V H I I I M V G K R G A L C R R A K Q R I L V I S F P E Q V V S D M I P A L A L V R S K
RANDOM_7055 I Q L Q Q E S L P Y K R Q I D D S G H T R K F G L L S E R T H K T K W E A V S L G L I F R L R M D H I R N D I G
RANDOM_7056 R I S L K F L L D D K L T P N A L W M D E P L A G T A N F S E V C S A E M M V K P N V E A K E H Y A L G L R K Y
RANDOM_7057 R V L C I K G H R P S K E V S Q T E N H G D L A R K K A A K A G S L D L P N H A P K G G Y V V N S I K E L A G Q
RANDOM_7058 S H T L P G S G H D N E Q L T A V Y L A D F Y D Q L S E R Y L K Y Y P G A T G S L D A F K G P C M L A V S P R L
RANDOM_7059 G E D G K P L L P Q P D V L L L T G S N Q K L L R V P E H L G V K V S K I K Y I S R G P E T D P A S L R I Y R L
RANDOM_7060 F C R Q G M V R L V D G L K C S A H G L A P A Q P V R T I D L M S Q R A R P A D P E C G T F D S E E K G D R I T
RANDOM_7061 R R K C R V L H V E L I N S V E L I I R Y T P E L L H C S S D R A A Y S D Q R L S T V E N K D T N L L C N P Y I
RANDOM_7062 E I L D V E V V Q V V Y S R S P T G L S G N L T D S R T T D N V S G D Q A R F P R P N Q A E T K K M L H Q S F I
RANDOM_7063 A G A D E F V S C A T G A V K T R P G I H N L N E V F L Q A G K N G N V G E I M V T D K Q L E I Y L A V Y S P Q
RANDOM_7064 M E Y P C G S L G A L P K S G D R H S T T T P Q L C F L N L I P C F A L P N A E L Q L E K T A Y T I F L F A Q C
RANDOM_7065 A V P P W P S L V L Q A F Q G A V M P K K F I L W S Q E F F N L D R Q S I V S K N P R N M D E Y A R E E A F N R
RANDOM_7066 T P K W P A C I A D Y A R L L L Q I F K A V D D E F D T A L Q L E A T Q D M Q S V D R S R D K I Y H L S S D G E
RANDOM_7067 M L Y A T K Q E A F G P F G I Q R K T I T I L W R H E S C S S Q S L R T L E H N I Q A S C T T S G H G A A L L T
RANDOM_7068 V T S N V L A Y I L R F F F W G G L A Q S P K E E G P E N V I E S L A F A V F V T Q I T A E P C G I T Y S I Y G
RANDOM_7069 V P A A M G L M A F L V Q V G I M W Q L G V P T V T L N P S D E T Y E D E L G R E N G G D D L A L K K P T Y N T
RANDOM_7070 E L I N T P N S M V V R K K E G E V D K A G S G T E L G Q A D Q S K S L K Y C S N G R I Y E L E L E P L I G V N
RANDOM_7071 R R F N P A R S L L T Y D N A T T Q N Y A G T N I E R S F R N S H E V S I V R N A L V G V S N E H V G P L F W G
RANDOM_7072 Y T V F Q E E P L L T I L Q F N S R Q D I M T T I K E A A V G P V L V S T L L F Y I G E G V Q M L G D F H E A H
RANDOM_7073 R L E A S A E A F N V G L D G R G D A H P K I E H N R Y P T Q M P T P Y R I L R T N G C G S A L G A D V G R S Q
RANDOM_7074 V E A L S S W M L R Q I H W V V V S L G K S L H Q T Y N G A R H G Y G A A C E L V S L N K R I D E F I W G G L G
RANDOM_7075 P S L C I V L N A M E L D V L P F T E H I E V I V A I E Y E M A D L R I Q T N W M D T V S V V K S S T Q L G E R
RANDOM_7076 V S Q P F F P L R H A T L D L C R N N F L G D L T Y L I I Q P Y V K L H A P T I Q T D S I D I P G K D F D V T K Y
RANDOM_7077 D P C T K L D I I M L S G L V P E I N C T I K F N W Q K C E V A K L L P M E A C E M C R I Q L A N L A E S W M I
RANDOM_7078 T E I D P K Q V A L I D V S R V Y P N M G V L D G G C K T Q Q P T K S D D K P N E I L G G S T P K H N V G Y M P
RANDOM_7079 M P V H N D H S E R G G G I H A M L P G I Y C F G V V G L T I W K H F P E P E S G N A R A L M H M N A F D T T Q
RANDOM_7080 R V E M V R P S R Y L E W A P G V A P P R F L T Y G C L N H H D G V A Q L C F T V N G S V F P P P N V L K K A P
RANDOM_7081 K D I V Q G S A Q C I S M T Q P D G I I G D S P V E V G G S S R A F F N K S K A S P A G E V A L E R N D S N G G
RANDOM_7082 I R Y G A V G P F K F A L S P I G L T S K S G Y Y K L G S L D R P E F P Q K G H D E E A L D L D T L V L L G V D
RANDOM_7083 G T S S Q T T L G T S A T T L P L A L E I G Q A L P N G G K G I H L N G G E S K L I P K H K K V T E C P A A P Q
RANDOM_7084 L K L L K Q P H F H K T I A S I M P T R D L A C T S G P R Q G N V N R P G V D W S Q I L T H A Q P A S S A F L D
RANDOM_7085 L P L L G I G E Y A V T K T V I D N S G E A P A T T S P L L A L A M P E G G A G I M L D N E E K P N G G G I G S
RANDOM_7086 H E Y V I D L R L N F K T V E S A G S G V A V Q Y A F S K E A G L H F S W R D D N K K S Y P C V F T A D D P M H
RANDOM_7087 Q S H A G Y Q G T P P E Y V L I T L L S E R E V L N K T T I P I S P L C T E V G P Y I E I N V K A E F N A L E R
RANDOM_7088 A F S T G E K P A P D S L Q A A A C D G Q W P D V I P L V L L M A T D S P E L I R N L I D W L S K V T Y E Q L G
RANDOM_7089 L S V Y V A V W R G M V K N S D L I L R M T G G E I G M I R L H Q E E P V P Y S C G S Y I I E P V V Y E S S Q V
RANDOM_7090 N L N A I L G E Y P D G H P F A R D F S L D P V N L P V T G S T S L S E T P Y R I A D K I V K K A K R A P P D V
RANDOM_7091 L E H F A F K S H Y R S H S K S R G A P L M R V D V E E K F Q E M F E E D R V Q I A E G M N A D D T T R D M A L
RANDOM_7092 S V T P A V L G L S T L K A P C D Q E G I E I D V V L W L T E E A A S Q R L Y V V M A G A V G R V D N Q L I I F
RANDOM_7093 R H V P T K G E L R L E K S K K K M F V N A S S P E L E K G T F T L S N F R G A H E M L P K D R A G D I N P A Y
RANDOM_7094 G Q R K H F V E L T A G M L R G D D L E N R V Q V A I T E C G R A V F C V P D K A W V R K I W E R H L N L V L G
RANDOM_7095 V T I Q I C A R H A S S K A G E G T M D M S L I L L R N T P R D T R K S T Y E A W L A G T A D D A A V E E H A K
RANDOM_7096 E G G L R G N Y E C S S V L S H N N R F A A C A T P E V E A R R M A Y G D I T Y K T V P P S G T S T G D T E S F
RANDOM_7097 S A R S V A S I E S D V D C A Y F I H D P M V V W N T Y A E F E E S W E H G D V A P E M T F A P Q Q D G H L V D
RANDOM_7098 N I K V I D T D A Y G V K M S R P R S S I L G L K I P K Q E V S Q H V F F C T T N T S L F L H A T L A V A G A V
RANDOM_7099 N R Q G T V K D H G L L V T W D Q L R A C R E G K S A G K A S T L K S R G A N P A T P H P T G S G N E H T Y E A
RANDOM_7100 V K V E W S Q A I T P T R E E G M T Y V R L N I F S K R G L N G F I G G E D W L F C E C A S E V E I S S R P H M
RANDOM_7101 I V I P T Q A G A S S R S L D I L D G H V S D Q Q P C T T H K G R Q W I N G N P E T Q K Y N G D G E F G T I T G
RANDOM_7102 V V Y Y P S V P P P T L D T L A N L E I V E V E K R K G F H H L R I V D C S L M V P L R G F I R T I A T D A E H
RANDOM_7103 T A T G Y P T W G S Y N S D V S E Y P I G E T A L G I P M Q C L E E P T K D S A T M T K V T D G E L V L Q A G R
RANDOM_7104 P V S M N Q P S G G P P W Q M K I A L E E T D L Y N V L T F S G S G G V T T C L H Q A A D N L T E Y L V D K A R
RANDOM_7105 C W A D S K G Y V Q V L Y G P S W Y C L P Q F T V S G A R S V A Y I T K A V V W T F G E C V R V P Y L C N Q D L
RANDOM_7106 F F P E Y G P E V A Y I D K T N G F F R L I N Q R A F L V P Y L W S D D E N T K R N N G D P K G G A P T Y F G Q
RANDOM_7107 S E F F T E M P C H F P E A L K R F T T G L Y R I K Y P V Q P A T V E H R A F H V A V K F S G G F N F L E L I Y
RANDOM_7108 E E T A L L Q A G L R G V S D E A V N R Y T K K P D G L V Y Y N T D V D F V G N I I N G L K D L I N I N K P L T
RANDOM_7109 S V H K P S Y P G S R G G P I S H K R L R L F Q W F N S A W W Y P W V D E G Q L V G V M E D Q P A E A T K L F S
RANDOM_7110 A Y K C L V I Q S N L D L K E M T N Y I F I S F R D Y R H Y L P E N L Y L C W D S M K H M K G L R A F A G R S H

RANDOM_7111 P F P C N G S G R E A Q T N N P A S E N A S F T H E T E P A R A E A G H D Y V I D I R F N L V F R G D A A V E G
RANDOM_7112 F R T S C Q I G Y S N N D I R S C A L S D W P L L S M D W V H I F S D A T L P S N H G P P G D P L M R D G A P P
RANDOM_7113 K I I V L F V A D K E G T I K Q T S R M R D S N P R D G Q S D D T E V Y L Q T E Y C Y S L I T L V D Y G L K I K
RANDOM_7114 G G L T R G I M E K A R D A L A N R K Y A G H R F G R Y H F S S L G P I L C V A G E E D S Q I S S R F A L P R T
RANDOM_7115 L D H Y A L Y Q R N K D P D R Y N R S V S S T D T I E L D S E T Q N T S K I I K Q H N N M T H I C R L Q R G Q G
RANDOM_7116 R F I V I N S S Q L K N G G S A A A C A T G V S E R A T A P G T K T N S L E K D F Q P A Q E D G F V V W N Q G R
RANDOM_7117 S C T L D I I L I S E N D G Q D L R M D K K F A N M S V F P Q E S P K R G E E I T S I R P V E Q Q V E L E K F G
RANDOM_7118 S E P R L F F W M G P N N R P N V E Y I T K K V E L G C D A T P L Q A I E T F P A L A H L A F P F S S K D K K I
RANDOM_7119 V L T T I H G R V V L A V K L A I E E A I G D Q S G M N M G K A E P F L L L A V M F A R Y F G H N E I A S V N E
RANDOM_7120 F S V A M I N T S D N W A P L E I S S K Q H T G L C Q A Q R N Y A D F T S H F K E T M L G T W C K G S I K P Q D
RANDOM_7121 G K I V R A N Y C H V E M S F N I E Y T F D S I S R S M R V H A D M G P V L S F E G T C L L V I E A K V D H H P
RANDOM_7122 G Y I E G S R D L F E Q E D M E R A P N D T G S H N Y N L S L Y S S D Y L D E A V K Q G R G V A K D L G E P K S
RANDOM_7123 K V K T V N V N M K K K D T R Y S S T Q D F L Q D V K Q V A D F T S E N R S Y V V E K P Q L S A R K R N W L I R
RANDOM_7124 D Y L R T M T V N E Q A D T V H E T R S V E A R R L E S S A T F R Y K A T L T I R A E D V E S N K E L L E P T A
RANDOM_7125 W L L I L K I A T A K E K E Y P D E S F L E M D Y S A S R K L L L A T H V T S P Y P L L V R E V L I K Y A P E S
RANDOM_7126 H I P T H D R Y I L T Y E E V K V P L M L G E A I V N A L K Q S E L R T I E S L A R V N W S P G C G I A F P G V
RANDOM_7127 A D M L S Q D G Y F T T E K R R W E A A E G L I T V I S P K A R D S A K V S E K C N L W G K S T P L C L G Q E L
RANDOM_7128 I E V G Y A N L E D S T D S R S V K Q T G A A L V L G S Q G A F K H V R Y L S C L L Y G C S T Q Q E E A W A L I
RANDOM_7129 L R R Q S I R S L L G R T Y K A Q V A E V V L Q S V C A H D E C N A E Y A R R G E T G Q V C G D A K A R D P A A
RANDOM_7130 S M I A M I R Y S V M N I A V F L Q V M I P M A V E T Q K T L M N V P N V I L G G S H G H G D D M D D L C K K A
RANDOM_7131 L S W T I F K P F N L G Q V P P A D N K A D K A A G R F Q E V S P G G A E D I V E Y I W M A R K L A L F L S A E
RANDOM_7132 C I A K T L R F R E D I D L D K I Y L N D L P L T C F G Y S L V T Y A V L L R T M F H W G S A F L A L P L I L E
RANDOM_7133 T T I N E G L I T I Q D T L S V Q G E F P K T A D C Q D G H F V F P D R Q P K L T M E Y D I V I E E W L W T I T
RANDOM_7134 Q T G R V G E D V T Y G P I H A V D L V S G A L T S Y P V T K S A L P T H Y G P G L D M M V E S L C P S E C N P
RANDOM_7135 S Q N T L I N E M C S G L F R S V E V G L Q D L D L F N T A L A K T M E L H F E Y A K L A R P G G E H V L V S D
RANDOM_7136 S A S K K D S S N D G G E M T C G L R K V F L Q A G K V D A L S A G A S L A S E K K K A F G I A R D S A E S I H
RANDOM_7137 S K P V N E S N K M G G N D A N H M Q G G E S Y P I Q G A G H K D L G K T Q E R P S D H N A I E D L Q G L R W S
RANDOM_7138 V A Q K V M E F R V L I P R F D G A M I Y K R P P M L P V L S C E A N D T E M F V V Q R Q T S G S N A W Q D N G
RANDOM_7139 P A N P C N R H A L L K A R Y L I E K T Q L G D K K H P T Q E R V N F N H N F E G F L C D P D P P S Q E P E G S
RANDOM_7140 E I L P P Q S I C L A L I C A K E V N V L A S P C E C A G N F R H I T Y S E Q F A K P Y A A G T D A A D A S E T
RANDOM_7141 I N F T L R S V P R F N W I C P K I I E P L V L Q F L V R R R I I L E L V I A I C Y P V N D N R R I T F V D Q G
RANDOM_7142 T G E D V W G M V E K G S L E K L K K Q N D T V S V V D K M V K K E C M P W K Q E Q T T D V M K R N F I P V L G
RANDOM_7143 S N A A K L I H G M A L I P S G S G P R L R V E G T T R V V R I P C I T S G H E G G K D L Q H A G P P T S M C H
RANDOM_7144 W P A R A D H S K G H V A L M D I E E H N T L R I A F Q F G S P H L I Y Y V T E T P S E S G G A K T G Y P L F Y
RANDOM_7145 H I A C Y I K R V K V L I Y A N Y G H F D A L S M H Q H T Y R Q Y C E T V L E G L P K V K F P M N L C E E K G L
RANDOM_7146 L S D H A Q Y Q I I Q K K T Y E D I E I I V V W I H R V L G Q F A S N E L Y D A P S K S N G D G G L K Y A I F L
RANDOM_7147 T I S D F E L K D T D D N R N L Q C R Q D V Y Q G C L M A V A D T W E V L I K I K I E Q E N T Q A M E Q S A L F
RANDOM_7148 M L D P D R M G L Q S V I L A K R P A D N F Y F S P G N D D Y E K Q R Q S E G A V S V D I N L M L S T Q Q E H V
RANDOM_7149 A Q L G K P K R C M E S I S K Q M Q E P G K S V Y K G T V S L G F R P K D S T D S A S H F N N G V S Q Q M K S G
RANDOM_7150 I P W G K R L S I S R P G N K K R M D P A P M H S G K Q L K P S Q E T S Q Q N F Y Y K Y F G K E R R A D V L I K
RANDOM_7151 L Y G T V L P L P F V A Q K E H H V A D R A G D R T S E Y L P G N L F F E W K D Y L A D S G D G F E L L R M S G
RANDOM_7152 T K E V L E R S V R Y T F E K L E Y E P K D T M I P R D L I L I Q D G S P G R E F I K A M K N C D Q C A R K S R
RANDOM_7153 A D A T D F E C K R S A I P C T T D R S G I K Y I V N P Y S V R I F K G Q N N E W Q I L M C N G R K P K F K P E
RANDOM_7154 G I D S P W T D L G I D L A C T P K S N L G G Y N V V N R A G Q G P T P Y E K N D L A L A R K S N P K A D H T I
RANDOM_7155 K V E S S S G H F P W E E S M F S T L T S V K P P N G G D E A R V L L D N D D L G N T R Q V N C M Q V Q I G R N
RANDOM_7156 C N H V R R Y G L C A E P R N E G F F V N G D L L E V L F S A F Y A L P D S Y N V Y F P R K T D G H H R V E V C
RANDOM_7157 W L S I S L Q I L Y A S Q Q S Q Y A G G Y Q Q D P S W L T I P V L A K E E L V R T W S D P V E A R V L A A S T S
RANDOM_7158 G L K A L R V K K R A V F F E Y V T V V E Q G A T G V L T A P Y A S G E M L H A E K P K A A I I Y V S H I V Q A
RANDOM_7159 E L P Y P D E C P A G I E P E A N G I G A E Q I T T D D A K C T P R S Y D P P I D L S Y G A M Q A C T F Y A G S
RANDOM_7160 L M L K D A V V N E M G C P I K A M I T T T A N E N T L V P A A G I H S G P V Q K L I V N C P K R G A P P V E V
RANDOM_7161 M L P E I Y P Q N E K R F A L Y S N L L I A L R F R N E P I K K M S N V E H S V L I S G G Q M K V G V I T L A G
RANDOM_7162 I E Q T K K S M P S N I S S K G S P H S I Y W Q G P K N E S G S V A V V S C V A V G V E N F L I K E Q A D R D P
RANDOM_7163 E I M P L L L R D S P P D R Y M D Y K R N I L N M N V S Q D V K T M G L Q S S V P L V C R V E Y R P K K Y V I H
RANDOM_7164 K L S D V R L P D K G W V S V V N S I Q N N R A F F T S D E C D V E K L L G V D T I P N L H A P W S D A P R K K
RANDOM_7165 F E G V V F V Y L L R S R Q F A Y E I L G R G G D A L I V P E L S T L I K N V A G W R K I G A A R S A A P A I N
RANDOM_7166 R I M P V H R Q L E G W T G Y I K K K A S I E K H N S L R T L P S E R T L F Q Y K P W S G G T E P T S W Q S P V
RANDOM_7167 E T I G R K G G L P L S T G G S A A P F N S R N T T S C V A L M F P V G I G H L S F K R V H L Y M L E A F Y Q D
RANDOM_7168 K V P H K N L N G V Y Q L A A L L A A R A N H C M V V N T N I T S K S I A P C S H I E R I L P S N T E A Q S L L
RANDOM_7169 L D A K I S G L V Q H Q Q A G I G E E V Q K N K R I Q A T V N S S K N Y C N L T R S C K L F P S A S N R A G E E
RANDOM_7170 D T Q I G H I N V Y S K R D Q Y T H G A N V K N V G E Y T K P R A E K T A F R Y L K L G P L D S K G A Y H L F R
RANDOM_7171 V H E F M A V G R G P C Q D P G R Y K Q L D Q G E E K D Q F L A N T V S L V D G R V L S I Y L A I G F Y A V V Q
RANDOM_7172 E A W V E K D K H T P E E A V G P E H H I C A M W K F V I S S V H L T G Q I Y L P E V K D E Y F G D G G E L Q K
RANDOM_7173 E A F R G A T A G E E K T F F D N E L H E T V L S H D A K I E E G E F A H S S E T N G N P L G N L S E P N L G K
RANDOM_7174 V G I K P W N G S S T L L K H K H I R D G D A I T L D E P A T L D N F D Y N I K A S P S V E T T I E Q V L I V T
RANDOM_7175 G I A R L H D E G D V K P G R K T L Q S A A R G K A M E R V V G K S A G E G R P F V V Y F Y S V K N K P L C A P
RANDOM_7176 E E L I P A E F A T V I G K L S G P A Y L M N D V G N V R R Y G G Q A P F N R E P T S S I K E A F P Y L M Q F F
RANDOM_7177 F K R W E A F K G L E Y N R F L G I E A T R T S F D R N G R E V S R G G S M Y T N S V R V I H E D L L V N N R D
RANDOM_7178 T V C G R P P K A L G P A G D V L Q S M V I H P R E I P G V H I L Q C D R T I G A P H H M G P P T P V L K R G I
RANDOM_7179 T A A K P D Y E I L V A A C N N E G G P V A P E R P E L P V P F N H Y G I G Y L P H L A S V L R M R D A S Q Y V
RANDOM_7180 E D L A M S R R D A T H N Q E N C L S K Y G N L P I S K T L V R R C V V D R R L L V K G F G S D F L L A I R P R
RANDOM_7181 Q L A I L K Q D Y E E L P E G V P D E Y Q G I A T S G L A H D T R L N L V L A P L L Y L L Q S V I E G S D S V R
RANDOM_7182 E Y E Y W V T L Q K H P S N A V E F E D V I K N A H M L G T R T S E G L I S K G S P A R A T L K I Y I K E H R V
RANDOM_7183 L L F D A S Y E Y K L A I T T K F F D I L T L K L G V S I V M F E K A L T L V P S M I S I E M K G S L E D K F T
RANDOM_7184 L L G D D P Q S D D P L R S A L P K N A G L V W H W N G E I V V F G S Q Q R H L S L R F R N S A K D Q F G R Y G
RANDOM_7185 K N L W E C P L K L Q T S F E V E S H S S L G A K P V S T N H S V K H T E L P A P P V A L R A H E G D Q T R E E
RANDOM_7186 I V K I A N A A S L Y S D S V T K D A Y A K K L D C R I P M I D A Q D F N G K M A L W E A G F K N D A A S R A G
RANDOM_7187 V R K F L P V D R A Q F E Q P V N T P A P G S E S H I S W R P D S Q G S S Q W F G H A G P E Y T M D M E I E A V
RANDOM_7188 L I F E G A K C L L L Q I S S G L T L N N Y S L S A N T N A G S Q K L V V E I A L Y D I G L F T T D I A R F L D
RANDOM_7189 G V N Q T R Y L E A L N K L N Q T F F P T N N E G Y G E T L L Y E T K Q L Q K T V I S M E I I S A L L M I M S D
RANDOM_7190 R H Q T S L T T A E L V R K I N G P V P N Q L E K H V A R S A H L S A V V E T R G L A Q G D F D T G K K R A T V
RANDOM_7191 I S D T G Q Y Y P G R R V S K K T E Q P W P E K I S C A E P V V Y C L D I I K S E F G N N L R S A D G G S L S Q
RANDOM_7192 A T A I T D C K A K R E G Q T C A Q G L E P L L L L F M S N N D E L N S A P M G E G V F S D M L K A N D G S V L
RANDOM_7193 N Y K A A A D P G R V F S R Y N N C V E M L V N N R F R K P R C T E S E L L C A V I D L E L G S E S P H G R Y D
RANDOM_7194 R R A V V V A N T S K A L I R K S G G S G L L L I G S G V T L Y T D F N L R Q L E K L A T M P V Q L R L I V S V
RANDOM_7195 N A G E Y E V N M A P I A E T D D E S R L P R T I S T E P K G G P T S Q E Y L Q N F W R M D V G I L I K G N I Q
RANDOM_7196 D A S G T Q F G F E Q E F S G D S S H E K V A E E P G F N D L E M T D I I T K L D D K K Q F G D T P K C E Q M K
RANDOM_7197 D T A F S P K L N E N F P N E S P L E Y G Q L T D K W C G I A A T L L R P R Y M A L R K P G P P E L K M G D D V
RANDOM_7198 R K S Y S P F P G R L C F G S S L F K D Y D K N K G F N K A N S L V E W V W D S T H A H I M G L K V A C L T S Q
RANDOM_7199 H K E V P P Q K K K E R F M V T A L T Y L D N L M P F I Q R W A A Y E P V E V I E L L T N R P I A P A L P K Q G
RANDOM_7200 K C R P V I M M D M H F T Q A S N F C L F F A G L E E I E G N E K T S A F E F A Y R V V G D V S D T T M F A I G
RANDOM_7201 K E R E G I R S G T L K N R C N P G A K A P A L T R I P F K G G D G V A M S I A F I G G E Q R N A R E P Q I N R
RANDOM_7202 T R Q G M H I G H W S P Y K F N S L R L K K T A S G L E Y T M Q L G P A Q A V A D L N K G S S G M C G G C R P L
RANDOM_7203 T S V T T T L P T M E K E N S E V E S S F L E A E N Q A R V L Q G K F C I R I L A A I F N T T D L S A H A E N G
RANDOM_7204 A P Y P P Y N D S G L E E R K V E A R S L L H S M M D G L V G R A R L Q I R P D V T W L V P A D H V Y T G V G D
RANDOM_7205 R A I S D L Q R T K S P N T E P Q S R G P V G L Q N I I C K N L Y S K R K L K T M L E L N I T A P P W S W G Q K

RANDOM_7206 T A A R D Q D C Y F S T G P S S P A N V H L A H E R L H L A A R N A K Y I F S G L V N A V N A Y P D K K I D A K
RANDOM_7207 S E P E N D K Q I E V G G E S T G L A K T N Q H G A T Q A G M E G V L F L L R F M G L E R L R R F L E K M A A N
RANDOM_7208 M V F E A G V V K Q M N E S M K L R N V S K G S K W A A E G V E A Q L P Q S Q A T E G T K G K Q P A S T D G I L
RANDOM_7209 I M D A I D T V N Y D K L V L F L V A L M G V K R R T I I G S V Y K A M G Q Q T D V L T A I K P A L A A G R W K
RANDOM_7210 L G H E N T W N R E S K A V Y G A L T V H A L F F Y F V A G L Q A Q V Y L H P E S S P T F A S T A T I I H L N N
RANDOM_7211 F R V A H R K S D L V T Y A I K P S R P H L A G P V P C S I Y R K T A N L A G S L A N Q D I T T Q H Y E Q E N G
RANDOM_7212 A T H P V V I G T F A S N A L P I G M N L R S I V R A M A F T E W V S F G M K A V A T L A A K C W R R D N G A N
RANDOM_7213 V G G A P I E M K G K G D P L T H T V T K Q H M F E R V V S I H L T D T F N N E D N P T V R F A N N I G E N L V
RANDOM_7214 L Q K D R V D K K G I D C E Y S R L I Y L T V R D L D A K S G A D E N D N D Y I N K D V N R T I S G T L E R A H
RANDOM_7215 Q T Q E R N Q V N D D Q F A A M P G N I Q P R I I G K V S R N K L Y D K T L I T D V S S I E L K R D R S N D V G
RANDOM_7216 L V D G R C S K A N I R T R G V T A P D E L T L A G G K K G S G D H G L P I L K M P K W S I S K W I K L D E Q Q
RANDOM_7217 A T D P A K E V I T K G D P I T E E E R Y E Y A N Q M Y Y Y A A M T P E S C G S Y Q Y L L L H K G W L L L I A K
RANDOM_7218 C K A G G L H K G L H I G Q R E V G I Q K Q P L I P E K K P A T M D M F G N S D E P A A P K K R Q L F P D Q S N
RANDOM_7219 D V L G N Q G I V A F Q G I Y E I G L S L A C Y L L D P V L C V A E S E Y S E E E R S Q M A E E K S G K L L I N
RANDOM_7220 P N Y T T L G W V I E A V R Y G F L P L E E I E K P G A K H V V V Q N Q K A L G R Q I R K N V K A K S S E D N N
RANDOM_7221 V L E T A I V G I R N G K I P L A K E S V S V R L R S A Q T K F Q E V T L I Q L E T P G I L G V L F Q L Q S E P
RANDOM_7222 V W S P L A K N L V A R G P H Q P C G V G K P Q N T V S Q G M Q P M S G E G I V L Y D S V L S T V D I P K M P K
RANDOM_7223 L G V Q P A K H K S L T G A P Q D A V K V I D V Y Y S V R S E T P N Q K E R R F P L T T Y T N H Q S P A W S K A
RANDOM_7224 W C G V N E Y D L Y A G W L G S Q L I Y Q L N F C G P K T S T N R G E G S A P V G S P K D P F S I I R G I K F V
RANDOM_7225 L N P L V R V G N K Y R E H N L T H H R S T L T V A M Q E N M Y F P G E K K N R V S P D N L Q V E G W F Q L A S
RANDOM_7226 F D S W S L L G S D I R G I R L V C Q Q V C M T A D R A A M L L F G V K D Q I D S I E R T N S P V R H I I D S W
RANDOM_7227 G T H H W N Q N F D I T M T P N A L E T Y E G L S L F N H N G L E C D G I D K M A L L R H E N E Q Y K L G G S Q
RANDOM_7228 L A E H R G F N K T G H S W L Y G E Y M E Q K E K P V A S D A T V A T G N G E V S H E T R S P A I M R E V C S H
RANDOM_7229 I E F R Y F V K D K N A G R Q D G W S W I Q L S V N I R G I E S G V K L P P E K G S G V K E L G P Y M K W L N C
RANDOM_7230 R G R R L D V Q I A L H N H G S M P Y R C E E V V D S E E I L R A L A V Q A H R P Q P W P S M W A Q I Q E K F V
RANDOM_7231 P D D R A Q G G C R V L D L D I G L D G S G K R I S N R A N N D K T S Q V S K R P M N Q T P L F A S K R A V C N
RANDOM_7232 E Q P F Y T L L E P V E A S S I I D T G C D V V P V G E P I Q V P Q S S A H A F H S R S N I P K Y T S L V C G G
RANDOM_7233 M Y E L A S I G E G G K Q M M E G V F G E S K R N V R A S D P E W E D A V P S E H N L I L D A D I Q E G I Y L G
RANDOM_7234 T G A V S D C A H A F G E V N I N D L I D E S F Q Q V M T S C A K A V A K T L S M G P E G I T F N T M Y Q V G Q
RANDOM_7235 P T D G L F N V S V D K R A Q Y S G A P M S Y D C D G G A S I R C P A C N D I V Y S K L P V D S L S Y L S E Q S
RANDOM_7236 A G K L N H R H W E C A H Q T S V L P D N L Q K E N Q F L N E L F N N F P L Y G N H Q A K E K I I V C P L Q Y M
RANDOM_7237 H P I P F E L M D N T E F V L E P S F W E K D A P Y P P K T K P E M G D N Q Q Y P M L L C G D N D S I I N C T C
RANDOM_7238 V S I M Q L F D G T K P S V A E T G F R S E H H T S N H F G R G N A S S P I A S K F K D D F M I D E A S L G Y I
RANDOM_7239 S F L L R H Y N L H G E S R N G L M R S M P G I D I S R L I S Q S L D G K P N E D S S V E F I I A F I Y S S N G
RANDOM_7240 K T Y A S A K E T P L T P P S V H Q I K S T R A D G P E S A T A S L A A L K G K S W P I Y T L S T S L I N F F M
RANDOM_7241 Q N V P Q S G N R R K L Y C K L R R Y T S G E L L E C V E L S P V Q D H V V K V M H G E A C N G R N T G L K K Y
RANDOM_7242 R S L L N L R K G L V D L S S G F R W L L D S A K N D P L C E I R K S S Y N Y D V E A M L A I D D M L Q L P A L
RANDOM_7243 L W G E I I L A S R R I I P E D L P M E I T V I L P E S K K C S L D F D R G T A E D D I A M A F L T T L C G M Q
RANDOM_7244 H G Y S F S A F D S D A A G T G T T G F N N Q V F D L G K A E T L G N D K K N R P M D V A R Y E R Q I M E F P G
RANDOM_7245 L E K N K H N V M I L D E Q V S C G T W L N G D A V K C D S R K K D T I E A L L F I F G L G L E L M M P P S C K
RANDOM_7246 P L E R T C P D I H Y V E A L L S E S H E G P I Q I T R K V A Y T L V W C L F F T S V I A N V R W T L A A L M T
RANDOM_7247 Q N A T K S A Y F S K S Y N T L P N F Q S E V E D K S K E L K D S Q L A Y P H A D T C N V D E Q G F K K I H V F
RANDOM_7248 P W A D S Q T T H A T S G T R D S H N F S H E L A A I A M M Q D A S A D G G E R A K L S V A N A K P A G A V S R
RANDOM_7249 K G C G N Y S P A S E V V D L K A P L C A M G L T V G Q G P A D P S V R Y R L E Q E G V R L V I V S A T G E I D
RANDOM_7250 T V G A Y Y T Y F N H A L E V A V A Q M K E G Q L W T T P R Q S L V T A I E E Y R G L G H P S W Y F N L G V L N
RANDOM_7251 A E E N R I A C E E E K I D G F A D A R R K S F R E K Y T Y G V K L S V S A T E F D N Y R S G S L E E T S E Q T
RANDOM_7252 T G S A L H C T T L Q G I D E W Q C H E S T E Q N K L G V W R R P Q V R G H P N E R R Y T W V D F Q Q Y E G L R
RANDOM_7253 A A P K E S V F L S N P G G V A F D A E I T F S R M I G G V L C A W S H E L M I A H A A D K V G L R M Y S R K L
RANDOM_7254 L F S N D C Q M R F V H N C S R T R H G T S L K S L L Y V S V I E Y R K T P L N G N A G I P P V P D A R G T L M
RANDOM_7255 S G V P Y D A V P S K T Y E S N T S G G E E C I M M R K S Y C G Q T N N D A R V M D R F C M D G T Q V Y S S L P
RANDOM_7256 L Q N H L D T D Y M Q L S A N Q E Y L K P M Q L I I E T L I F Q P L G T A A N T A I K N E Y Y L G G T E P N M P
RANDOM_7257 G E P C I Y H P F T E L L C M L D Y P G C A R D G K P S V G T F L A E N V A H E I K D A K P L E G P N G T L G K
RANDOM_7258 N E N V L D G N S H K S L A A G G G D D I W V P N V K G L V I P T F M P S E T E H S Q I S S P G S L N T V N A K
RANDOM_7259 A G A R K N V T S A N R L D L M K S S A A A D S K R V N A K W I A V Q G V K T E I I I V E G N F L C P D S V I A
RANDOM_7260 R P T A R T K I N G R I N G K L H A E L A A F K I A T G V G R A Y L Q A A M L A P E S A I D K L M K S G S Y A K
RANDOM_7261 L P R I M M D P T L S D L I L Y T I A T Q W I Y A A L F E A G L M F D T Y V N K I Q F F T I I T E I C K V S A S
RANDOM_7262 V F D I T R S N K S V L H F A Y L G N V S S E T T V D R G K C S I A L R A G S A K Q E Q S F Q S V A R L R A G Y
RANDOM_7263 T S K E H E I N S G V V K N I A S S P Q F P K Y Y K E S A F C I S Y E D E E M I G I W T Q P L T Q H Y L T L A M
RANDOM_7264 Q F A R K D V G L T T E K V V C T V E V W K S G G A V P Q P K E D Q D A A L E D T K R I W D E A E T L A Q D S G
RANDOM_7265 Y V D G N L T T D V A L N P T D V F D T P I G E V P L K D T T S L G M D F A D A I G E I W Q P R G I L M K N P G
RANDOM_7266 R E N L S R P L S V K S V F S H I Q S C R L Q W N A K N S E M H K C V K L A V S T G D K I G Q I V G R Q W N W I
RANDOM_7267 P L L Q N N V T F V P Q T C Q D S S K L V T G N Q L P P P N T D V S E N A V A E P G Y E L G D S H T P A L V Q V
RANDOM_7268 K P K P P E P G L T N G I E G C C K L M W L K M K P P A K Y H C A V T K C F N E K T V T G E R K A S L L C G H V
RANDOM_7269 A R Q F K D A A G V S I A G R L G H D K A N L T T W N Q V E A Q I V V K H V G G G A F K C Y K E P R N L N S L N
RANDOM_7270 G P N R A D V K E Q L I L K F F G P E A E K E I I T T I A D T C V L R M T L G G D M V F T N H I C P I G G Q G L
RANDOM_7271 Q Y L G F A H A N A R V P N T I K P D S Q N Q I Q D V N T P I I P Q P C R E L V I A T S L G S D R E T E V D W T
RANDOM_7272 A R A T L D C K R L M P G M A R P S S T S A Q P L G G I A Q V G G S A T V V E A Y T A L G G L I P I A P S L P D
RANDOM_7273 V L T T V K V L I T V L P I C N Q P A E Q Q Q N S A E A G H T Y A W A I S L L P D S F E T E C I F L R A T R N T
RANDOM_7274 H D C I E E P L I T A G D V L S K S Y D D P L S S K F M S C D K C G E R V L E Y Q K A L G Q E N Q R N E V S T A
RANDOM_7275 L I T T F P S V S G E L T V I G L A E M F E C P I A L Q E T V W D Q Y L L A T G D I Q K V A W A P P D S D K A Q
RANDOM_7276 P E Y P P K G G M T C A G T D F G A H F D E I Y F M L L S K D Q D T L T S T Y Q S T H K G E P N L S T S A I Y N
RANDOM_7277 G D L Q C L E N S K E R G K A G E S P V E P Y S S R E P L T E P A H G K F A E R L D Y P E L L L G R Y V G A G P
RANDOM_7278 L L C A L V S V F G S V G C R L R V E V Y F P I D W G N E G T I S T P F P I E E V C R K G A I D I D L G Y L E T
RANDOM_7279 S L S K L V L M N V A E S V A C S Q I D P T S N L S C T P G G I A S E P D P V W N N F Q W P A R W H F E K K F S
RANDOM_7280 V K G M D Y K L I L T A E T Y I G L Q A Y L A V M C D L S R Y R I G F F K Y G G E R N W T E R T K M T K M T L H
RANDOM_7281 I G Q F D T V A A A R A R K K R L T Y E K D V S L I S S G Q G T A P G V Y I A R S E W P H Q E Y P A V V D R K S
RANDOM_7282 G V S S F I V L V P D D S L Y I E S E H Q L L G L A D E E L G T E Y E C F T K G T Q G K P F V K A T E G V A T A
RANDOM_7283 Q K Y E G T L T N T C I T S A G K H L M M S T D Y L C K T F I L A S Q Y R G R E T K T T R Y Y S I T M L S V R E
RANDOM_7284 L T V S I P N E E E Y L D Y E N G S G N D N M K A T A L L K A K G V M Q A G V T K K D T K P T Y H L L H T V N F
RANDOM_7285 D C S Q S I R G A R I T H S A G T E M H K T E G S A P R V R V L Q L P E Q T I I N G A G N V T M M G I F K I T R
RANDOM_7286 S G L F A M Y A L K L S C L A K D A L R T T M A C V K L T G Q R P E R L Q K I P I A L Q A F K P G S N M Y S R S
RANDOM_7287 K N Q E E E K H A G S Q Y S F L V R K A P V E F E K Q A R G K E Q E R Y L E M Q L P D R A S T T C E E R R F S N
RANDOM_7288 M S N P G S G I G V G M T T T L H H Q L N A A Q P K T K D Q S I R I D C E F Q E S A T V D K R K I E T W F Q A I
RANDOM_7289 E A G Q P G D V P N N L Q R F D S A I V N V A R V G L Q A G V F F H K G K Q I N L K Y R M P M L I Y T C A I S N
RANDOM_7290 R L C R L F M P Q H L H L L I K S V G Q F S N G R P P P R A E S T G N H E V P Y D F L S S S I G L A F A G D T P
RANDOM_7291 A L I K S L I L L G E L P A A I Q I P D I I D L H I Y N S N Q E K N L W R I T S S L M G H L I R L R G L G F Y I
RANDOM_7292 V H F Q L Q A V I N E I N M I K W N G A P E P K L K S I M A S H R E I I R V I N C L I T P S G A A L K R L G G K
RANDOM_7293 S A E G Q S D L K F C A A Q G Y T L P T F T E E T N A Y G Y Q Q Y A R F A G N E N R N V C P K D L L K N L S D V
RANDOM_7294 S E L V I E E A D G L G Q Q R P T K T V I N L R Y L K K L L A Y Q A P G V G L I G N N G V R F R N I M R G T G L
RANDOM_7295 E E R P P V S G G T M E R K M P S K M A L Q E D K A V R D K Q A T S F Y V R I K T L A A T S C E T D P L M T K P
RANDOM_7296 S N N M R S L I P P F C I I S E L T R P T R Q T L A G I R K G P K Q I P A H A M V D L T A G D D P F G T N Y G N
RANDOM_7297 L R I L Y L I E R D E G L N M R V S D T L W R R V T V L C N H K S L S S D N T A G H K L A L Y N T D R A I S A S
RANDOM_7298 T V V G K A C H I D N A G H L A Y F A V M M D F I E G H E R G L L V T A R F L G I E L R M T I A G S K G D F S N
RANDOM_7299 A G V M P S G R D K R N D V E I Y E T K D D T K T W I L F P K Q H P K L W R L E D V S I D G T V R T R R S F F L
RANDOM_7300 T S K V Q R E S Q A N W E D N N Q C G P F L A K T K G G S P P R E L T T F L I D N I P G T K L N H A P E D T G N

RANDOM_7301 K A H F Q G A L G H S G C I S K Q N Q K C V Y S R E M H D F I E L H W T L R T L T C R G I G A V G S E Q S T L T
RANDOM_7302 D S A G L T K S E Q M I L S A K K D V R R Q M A V A S S S T N M P P Y K L G A P V D N E T H W K Y P N N N G L C
RANDOM_7303 K T T R F A R K N P R L V K S G K L R E Y E K V V I K C L C L F S Y T L V A P I L L C L P L T M D Y T E K K S E
RANDOM_7304 P L E V L L D Q E P K H E I R I A A H F L D V D V F A H D N M Q L V Y R N S K Y K K D S D N G S T R L K K D V R
RANDOM_7305 Y G I P V K S S V N R L R C T T R I P R A D Q I L S D V N D L W A F D G A T L I G N S A C P L T F V K Q T V D V
RANDOM_7306 T L R P E S R Q L L A E K F T E K E V D V T H P P G G A L G Y A T P R C Y E L K S A R S G L H I Q K A N R A P E
RANDOM_7307 G A I A K A T A K L L E E N Y R H S Y V L T M V L L C G V V R A M Y N S G P M Q V A H V W L S F K G K Y T A A E
RANDOM_7308 A G S R G D F L T D K P L I S S N D G K K P S L N A D N Q L A E F S R N K V N C G S E L K F D P V T D R Q N L M
RANDOM_7309 W W I Q S L L A K F K R M P L F Q I I K G D P F D Q P E V C T L G G A T A S D D P Y A E L D V S T K F F S M S T
RANDOM_7310 N K I G Y W C A E K G D T L L K L P A K H W D E G N K T Q G K P E A S N Y Q V H S D P E R S S R A C V R C E P P
RANDOM_7311 V N Q G P C D S L K C K P T M P S A I H L K R H V K P E S R T T S E L R P A A R F E V G V S E Q P E K T S V K V
RANDOM_7312 Y M L N V M S D E D G D D R R K K A M H Y G M A P I S S S I K L E T T W E K I D L V S S G S D C N L L D A F I V
RANDOM_7313 W H Q K T I S F L L T I D E P T W N A S N K K T S K T M E P G V L T V V C P Y D A A E L G F L H F D L G L Y N K
RANDOM_7314 D L I M D T T E G D F L Y S H P T Y S A M I E K K W E V W P D R R A M W L D G P Q S T Q N N S D R Q H R S P Q S
RANDOM_7315 S I I E M Y N V V T M N E A V E S T G F S S N S D A R P S T T G K G L G D G F D Q P M F N N I K A D N N T T G H
RANDOM_7316 K T K F L K R M A T N E F T E N R D D P L V S Y A D Q F F A F G P H L Y I V K K Y Q L R H S G Q P Y G V A L W F
RANDOM_7317 A M F I K S K F P V S I L R L V I K Q R P H S G F G S E G L C Y H C A A G T Q L L K Q T P E T R E L N K N D L E
RANDOM_7318 R E H T E G I F L A C G G L V G Q T G L G F D D I G L P V N K C F D A G A D K L R D K R V D L R I I E V E E V P
RANDOM_7319 S L D S E Q N C T L F G S P P V V F E G A G Q F I T I R R V L M I S R V F T P Q P A P E S T A P N I I P H K V P
RANDOM_7320 L L S V Y N V D L L L T T R T Q L A L D P S I A P K P P G S D T A N V K H E T C L P E W D P A K A L A R T Q W G
RANDOM_7321 V K E L T A P E G G S G K L S E L S G P Q I H G D Y N A D Y E C P E R N T L R Q K D L S M T K S L K T D Y K Q V
RANDOM_7322 A R A N S D I E G H R P K Q E N E K S Q S C L D R S K W K E C N N G S E G L K R V A V C A M K D V P I V E T G D
RANDOM_7323 F W Y K A E I L S P T V T V P I Y G T F D G F F D Y G M L A L V Y V P A Y N P L D V A L K P G S N K C P Q G N L
RANDOM_7324 F G I P V L I V C D N L I E Y K L H V P L C A E D L S D V A S I V A A Y S E S K N S P P P E F T W V S G Q E N F
RANDOM_7325 Y K A C A S V S D P F Q L P D E G C G V P I T A I H L F Y T L N F L D R N S G N W R G W P E L Q R D V K P L S L
RANDOM_7326 S Q K I A C I I A P H S K A E L N N D T A D A G V L T T R D I Y T D S S I P N G D T F L V D I D G Q V P R E K V
RANDOM_7327 M P E D K Q L I D G L I Q L Q I V K D G E N L K P K A G A K S D D L K K S S P K S P E K Y Y I F V A D R L N C E
RANDOM_7328 A D Y Y P T G I E A Q Y I P F V I K K C F L K P D K P R G A T Q G S Q A T V G G K E E L S R G G K D R K I M S R
RANDOM_7329 G H A F D V A S K Q N N V V L R T Y L D M A R G S E V G H Y A P R H A L G L S R R T M P T E A F C A I I G V W R
RANDOM_7330 H N K A P N L I K R C A A W D H A F V A Y A L A R I L G C P V P E A I S K D L E Q V I S F A L Q D I Q M V G Q H
RANDOM_7331 V V K N G H L S P G N G R A S P F L N L L K V I A I R I D T R I F F N V T N L I V G F T R S M L T V S F E R V K
RANDOM_7332 Q A P G Y D P P L S A N T V N G K A E P A M I G I S A V Q K G I S S S V L F P L S S F D P A Q F Y D A G L S S E
RANDOM_7333 P I E L P P G V W A C H F T A F T V L N A I D N E A I S K E E F S G W K L M Q D Q K T L F E L Q I L L A Q L P L
RANDOM_7334 Y L D G V L E D L N L I D Q S D M P A Q T C A L A W T M P D G L W L D T I L G D L L G L Q L M N R Q P C L Q V M
RANDOM_7335 L T L K D E V G F V Q T E V W L Q T Y V V L Q P I F G P I P R N D E C P D N V A A I I A I K E A V T A V D Y G S
RANDOM_7336 S I S Q K L Y L L G S N H K S I A G I A S C I G L P S E S F Q S A H L T R D A P F A K N T L E Y L E A N D K D A
RANDOM_7337 Q E G L D S S Y H T A N L A G F W C R H N S N T H R S V V L N S G S F M G Y V S V E E S V I G G V D K A R C N I
RANDOM_7338 N A I R G K L Q E H K D A Y N R L K S P D R R S G D E D N A I A W L N Y M M L R W G R D E Q I E Y L F F M P M C
RANDOM_7339 G S H R E A L D P P V D F G V G T T D E V W K D L V G A G S K S L K G P I P T T K S S A R C I G R Y L F A M K F
RANDOM_7340 N I K Q V E P V D R V A K A M A K T Q L I T Y D N M S G L Y L L T M N P D I D A Q E A G P E I S P G M E D K Q F
RANDOM_7341 S T P E G H P P E G V F M Y R I L L R Q K A H I H F W M Q V C P K A F E V I D I L T N M M S T Q D M E L V E S F
RANDOM_7342 N H P F R D I W A D V K Y K Y G A G S L V F G L M A I P K G R D G F L V K E M T K W T V A V T P T Q H K G Q L L
RANDOM_7343 L G N A F G L I H A A V L I H P D F A S Y S D K I E V A Q L G S D E L G V L L Y D K P G K G Q R D V T E E G E A
RANDOM_7344 E G M E R D R L L G K K V T R K V P C K C L D E M T Y V N E Q P S G P Y P A N S A S I N N E R V Q K A L D R H G
RANDOM_7345 I A D S P A L Q R Q V S F T S R D L A E G I E T G D V Q G S H V V D M H D E A I S S N S D K I K V Q G H V V E I
RANDOM_7346 C V T Y G V Q T A F T R L V F A R V G M C C E F S A L S A Q R S K T L P F S L K G K E S E S Q P V C Y Q I A A K
RANDOM_7347 T V C E G G P A A A G L L H S A P G F E C S C I G G F N H K G M D R V K E V V M E K K F L K F I E A P I A H S G
RANDOM_7348 V E T S S V P R G E A V L K D T C A K W Q L V L L L K V P G D F S I I Q N Q T G N P R E E S P L T K A P A H Q N
RANDOM_7349 T L M A E L I S A Q E V Q K L V D N E E V K Q G Q K L L I A H S Y V I E K T R F E N N D R Y A G L L D P R L K D
RANDOM_7350 R Q Q S L V G G K E L V P T T R L H S R A F E R H F G A R A Q I R C S Q K L P W A V M A F N E V T A N L I G S E
RANDOM_7351 G E V E T L T S N C P P I T I Q L V V L P P L D D E P E E H H T R K T T T Y Q K C D D R P T L M G D C G P E D R
RANDOM_7352 E V I F R R M A E A T Y F L N T N E P T G K P L D V Y D G D C R N Q G I P A D Q I P I T L R R I A P N G L K V G
RANDOM_7353 E A V Y Q I E E S F A A M E G S Y V R E S F P I A K L S M G S M E R V G L L L D D D M T T S I P T S E F Y L G K
RANDOM_7354 P M N S D D P L I L R L Y E A A D M I L L E M G G Q A M Q T V K L K E T P G N N L R G G G A V Q R V V L I Y C F
RANDOM_7355 N R Q N I D S L G V D F P D E G E L G G A K K T W L D S C Y E H L P C S A M L L I L K D T S D G S A A L A G F L
RANDOM_7356 G F L D L R L E M V A A D F I Q K L F L R G A H L V T G R S E S H T M G C N L L A V P E V E M T Y P K L G F I T
RANDOM_7357 V S L L D Q K M V G S T D Q M E L K I V N E K G K E L K D S T D P L K W V K D V R N H L Q G K G Y S L V E N L D
RANDOM_7358 I A V C Q Y L T W P G E K K S Q D T E N T L R Y A N Y D K Q T V I F V A L S L N D S K S R T L N D D L S E F F L
RANDOM_7359 I G S A K L H P V I I A N P P D R D D K S E A S P Q L A R L I L R V N T N Y P F D V S T H R Q L R I K S P A S E
RANDOM_7360 Q I I T P E E K S A D V T Q R P P K S E S S L I E N L N F A I D G L L E A V A D N I E T D C Q R A L I G A L S K
RANDOM_7361 T F C K R A A T S E E A M D V R G D A E A C I L L L K I L L L I Y M P R A W A A P M K G A G S C R F Q V P K H R
RANDOM_7362 E N L R I V E A H S P N V K A V N I R L G D Q F W M M P P V F R T R P L T T A V V G L S S Y A W H S F D I P D A
RANDOM_7363 H K I V S E C N N A M D Q R M G N P L T I G A E F F H S Q K Q C S E Q V Y S A T E G R G H A V C K C D P T V V V
RANDOM_7364 A D S L F T T K V K R F D N G T Q R S E Y I T S R S S E P P L N R T K K R D A E D G G A E G R H E T M I S G G P
RANDOM_7365 N A W I N P A N Y C P A L V S C K L D D T P A W K N Q V K A T H F F P T E V T I H P V A D G M A N M C T D A Q D
RANDOM_7366 K E L P V T G R F C L V K R Q L L K T D D A K R A K E F K I Y Q K I I Y E L A N I I S G Y Q F V A V A S V Q L R
RANDOM_7367 D N Q S F W S V E C S D R K T H G S S E V L V T G L K P S S F A I V I K S V M Y L H E E Q V S A F I V Y I S Q L
RANDOM_7368 V R Q M R K E R Q C T D H S S R S Q V H N N K W S G A M T A I A F Q D K A K T L N K A A E W T T Q L C M D K H R
RANDOM_7369 C L M I V D G T P V L S T L H V D V T Q G S T P K R T A I Q G G R L A T R L I M E H L P Q S G R R S P L A Y P T
RANDOM_7370 A G Y I F Q L L T L P V L S A L P A S L L T Q V R F A R H S F Y I K E D T T Y K L I Q M Y N S V E S P S I F Y I
RANDOM_7371 K S T Q D R P G N V L A L K W K T A L P A S F Y P T E L G A T V D L P S Q H D K G G T G M R G P P L K C M L Y Y
RANDOM_7372 P T G R R G G S G I F Y L Q P R S A M F D E E W E I E P Q P P N Q K L S F S C K G A N Y S N W V G C F A F R F E
RANDOM_7373 E E I D A F Y L L N L R S Y L R D L I H A S E G L Y V V D G I A L Y T I N L G E R A V V I D Y R S L M D I R Y S
RANDOM_7374 I S E G S I R G L Q F L N L V I Q D R P V S I T L G D T S R Q I F S R R F I R N G R Q K A A V V L Y S P F T L D
RANDOM_7375 V N E G F S T R I E V E A D I D T I G K E T P F A R I A C Q L T P V S V C R S R S T A L V N G Q Q G R E V C T K
RANDOM_7376 Q A H N P A T S Y K I G I G A S Q T V D P G R R E L R A V I T S K E T T F D A P F L S K L S I A T P F A A A D S
RANDOM_7377 L P P I L H N V D P P G L N T D A K V A F L L L G E L A Y K P E Y L Y R H W G K V A K D H F I N G T F G D W G D
RANDOM_7378 L I E L S S P E L A E T I Y S K V C K I A Q V G P Q K V S L V T L G C D S P L E R I Y T P L I A D V P T N L P D
RANDOM_7379 K E L F M V K E D R A R T S Q A S H T D H A I P A S N I W S Y V L S K A A K P P M A S H Q W M I P R E I L D A E
RANDOM_7380 V D L G N S D P C D S T T N G C Q S G E T S K Q I L F V Y S T N K I N G L G A F L Y L D L F V A W V E D P K I A
RANDOM_7381 A I F C D S L K P S R G Q T E S A L F K A I S L Y V T I T K P Y E V C T T R P P A I A A N E Q M K A A Y R V L P
RANDOM_7382 K N Y Q L S K K T I N K L V N Q F A V L Y L C G K L Q R T D P E Q C A A R V N G Q P G C V Q Q A C L V R I P I S
RANDOM_7383 T N Y L D A N G Y E L A P L P F G S Q L V G G K G A T N R C K K A Y A A Q N A A V P A R A E K I G V Y T G L H L
RANDOM_7384 S W T V D P H V A D Q G K R L I S L N Y E L E A V E F Q P A L T T S R E Y G W R Y G R E S I C T A D E V K A P A
RANDOM_7385 A L S A G I D G L Y V L V I G Y F I K A V G D D P Y A R T G D N Y K T C S R D C G Q P Y S D G T V L G V V G K Q
RANDOM_7386 D T C Q Y S E K F L K Y V W I G Q S L T E M S P P G A S D G P G P D G V L C T N F G E S G G D A Y K S L V R I V
RANDOM_7387 L D A S L P M R D S L L W L K F G T P Y F D H M N Y P P N K I W Y R L K G T A N P G Q L L N K P Y T T P V Y G G
RANDOM_7388 V V S K G L C E K E L T D E I K A A P T G R V S D I A S Q A C N C V Q S K P Q S E G T L D K A D G C A D V A D P
RANDOM_7389 A G Q L L S L V D E A A F D L Y R I Q P G R P G Y M I S S D Q A Y F K G G K E D A I G T F F L N G S K T S A L Y
RANDOM_7390 G L L K L I P I T V T F E R Q E Y Q S A V A A I P Q K A L L W M F A E G A Q S P T Y L L T F L M N N V M F G Y I
RANDOM_7391 Y L Q D V D Y S E Y G V W V H G R S V S V A A T K M A R D I M K I T P R F E A K K A Q S V T T C R L N I T T A T
RANDOM_7392 I G L R G A R I V G F G Y F R P T G V I E P A D D A A T A E N C C W L T F E Q R L F G G D E V D K S V S R R R Y
RANDOM_7393 R N V R G A T T E N F P L S G V K V E T S M L M S Y L T S R I A A I K D Y P N G H L C I Q K K S L G F A V R A P
RANDOM_7394 R G N L A L E E L Q P H I E L E K A N L E I E A S A T D V H D K F H S L Q Y G H Q I E R L S Q N V S Y N R L G I
RANDOM_7395 I G H A K E P R T T Y G A V S N P L K P F L D K I L G P I Y I A G A A L S I F G H N A N F Y L A I L N M I S R Q

RANDOM_7396 P P L F R I V L D T E P S G A L E L V S K R M A I P A R S K F E Q S Y F D I T R T C V Q P I R I D G S L T K V A
RANDOM_7397 P L L R D L G A R N G A E Y H L K E R M C R P I L A P N K R A M N G N R G V S Q L T R K K G D P A G A A L N A C
RANDOM_7398 L D P F I G P R H P G S E L G K P S L E R L G P M K S L M S E G L T W S L F E A F Y Q S T K K L V C G G A K D L
RANDOM_7399 I K E V V K V P C A M F K L K F E L E S P V F K Q Q K R Q F L E T A G T G H R T E V D A H N G K D M I N H D P A
RANDOM_7400 E L I E E V C T G P R T V D Y Y E R R L E V L P G Q E L L Y T V V Q D E F M D A R N T A P K G L S Y F R I L L G
RANDOM_7401 S G P A K G D Y T T C G E S E T A M S A T N V M L T Y Q D G V E M S M A Y A M M V A E A I V K V W Q E L Q D W D
RANDOM_7402 E K V S L L S G G L L N N I S L A A A A K V R D F G K N R T N L S E D F I P W A Q N L A N N V D N G E E R W K Y
RANDOM_7403 E V I V L V S T G K I D G V A E F Y A G V T G T R E E C V V V A T Q V Y L L A P S Q L N C L D N A G E N S V V V
RANDOM_7404 F F K C D F C E K G F V I C A R R S T I V L Q N A N D A K R K V C N L E E T I A A V Q Y L V I T F A A R I M I S
RANDOM_7405 K G G V F L F V Y R A G N P Q R Y Y W T F I T R N A V D L V S I P A N V K I M A Q I P D G K E E Y K P A A I N G
RANDOM_7406 F Y T T A L A E L V E A G I E V A F C P L L V G Q C H E K A V Y C G D L H E L L F M V N K L K K C T D F S K W E
RANDOM_7407 L F N T P E N F G L Y P G L L I R F E I Q K L G Y Y R F N I C K G A L K A G S A Q H R H A Y N D Y Q Y M Q D G S
RANDOM_7408 T V Q A Q L Q W P G N I L T E Y D K A P S I H A G E H G V A T D A K F H R F R S F P Y D C A E T V Q S E S V R S
RANDOM_7409 Y I K V T S E G I L E D S A G H T L E N P T E S F P R R G A Y I T Y A W L D E N S K Q N P I D L L Q R S Q R Y K
RANDOM_7410 H T L G D K P I S L M G W H S S E G F V L R A L L T G E E N A S G R S M R V N K V V Y P P G K D W D I S N M W S
RANDOM_7411 L E G D S S M E K A T M F A D V N S A I G N S N L K V G P K E T V S D P L M E Y S V V S T V P A A D G C S F G T
RANDOM_7412 Q L R G Q L D V R T N M K I T D A G I A K H E K E I D R G F N M F R E R W A V G G Q G G Y L S R E L P A V E T L
RANDOM_7413 V I Q A K V G I D G P S E K R V P K W A Y A Y Q L E R V S V G G R I G I Q P Q M R N K T K G L Y L G K N K I D S
RANDOM_7414 S A G Q K S L S N S L F K L I A S S G Y L G Q Y T E V I L E M E L T E K K N K E K P S L L Q A E V D R L L S G L
RANDOM_7415 I S E P T L V R Q C G Q G K F R E G E L L G T G S S I S G E S L K I A W T V A Y R E L I G K Q Y V R D E L P C E
RANDOM_7416 C L A L F G R L I D N S F C P G K S A N L V P E I L D P D H A A A S P A I D T A L R H A K A G L R P T F L P A P
RANDOM_7417 D Q T L V D N A G I T F T P I P A Q I T G A K V E K A V L K A P S Q H S G I K E M D I A M Q V A A L C L F H T I
RANDOM_7418 V L Y V T W S V D T G M E G T C S K I T V S S S E A G A S H L K E E F T G M C F G Q T V F V L D R V S E G A Y S
RANDOM_7419 K A G I S E E K Y P P K P E V V S A S R V S N K R D G A T T Q L H Y Y C N Q W N N I Q N S V R P Y G A E E V D V
RANDOM_7420 E T V A L N Q L S L F I Q K V N S C S C V I N K K D T H A K A I N I L Q S M T K I T K S A L R Q R L D K A S D M
RANDOM_7421 D A Q K V A Q M Y S N L R K S H Y E T Y Q Y D V G E R D W F S L L P P E G T R F D R I Y I V D S Q L A Y V A Y D
RANDOM_7422 S A E M R W M S T S S G L G T T E H V I L I A L S R E T Q I T T F R P M T Q Q L S P V Q Y N Y M Q A A R C Q A E
RANDOM_7423 G F C Y H F S T E L L I R D I V G A G K L V Y P V Y L H A T L N T T F Q E D P S V H L F L P M E L T I P L Y I E
RANDOM_7424 A E L D R S V D D E A E K K Q V T G D V G L D C M P L S P Q S V A T I P L G E L L G P T A S G K S A K P N T N S
RANDOM_7425 S K E M Y G A E S K I K E A Q S S P G K K L V T S G L L I R H E A G Y E E G V T T P A T Y A I W A G G P F Y D I
RANDOM_7426 L E E K M R P R Q K R E P F P G G I Q S D Y L T S S V C G F V Q L K E N A E F V G G H M S T L H G M A E V G A G
RANDOM_7427 S L R N P G F D V P V G V G K P I P E L K Q V N L S F I T R M E G P T P G W Y I Y T T A Q I Q L Y W C I A K A N
RANDOM_7428 P R G H E K I D F P K I L T A K W C R G G N F T Y V L L G L F I H L N L D F N A T F K S D V V F L N G P N S D P
RANDOM_7429 R T F Y G V I T A T S E K G Q S V K A T M N S M P A T C F T V Q G L A R D K Q E S S N T K L L K V A L S E T S Y
RANDOM_7430 K E E R N K P K P G D E K H F Q F W Q P N L D L D L F G V N K P D D L L A R D I V T L H V R P L E N K S P L T S
RANDOM_7431 S Y I E F D L H L G P P I D P T L E G T V V M Q K V G A T V E Q H P G E G A E N P A I S F P A S P P V V E T I I
RANDOM_7432 F P M S P A D P Y N G V T L E A Y Q C K G Q C L L T L S D P Y G G A S I W S G A A A S V L F N F P L D V Y L H S
RANDOM_7433 A G Q L A G C E A E L R V G Y A M F W E V A M A N H A A G T R S A G R F C V G A L T M M P G V L T I S C S N E T
RANDOM_7434 K Q K F K E A V V S W L Q P G N R A I T A Y A F L A L L S L D G S V H L I D G E E G G P E S V K I R L K I T V M
RANDOM_7435 K K T S G S S R I D L Y I Y N K L D N A K M D A P I P P D N L S E A T S A E C C T A K A A P A I E D E E G S I N
RANDOM_7436 V Q H I K S M W K L L I I P C S E H H G E M R A A N L V A G R N L F A M P V S A C A L L D Q Q A N A L K K A A P
RANDOM_7437 A A P Q E N E S W E T R Y H N R Y V G E V I N E D P N A G G Q R I G E A L V K A I E K P V D H L M I T R Y L A P
RANDOM_7438 N G C N T Q R N L C F E S S I E G P L H W D C T K N L S M P V K S A E A L L A V I Q A G G K R H E G F D G P G L
RANDOM_7439 N L E H F D T V D P A E V S V K R G V E V A A K C V F A R C K W G T M V D G I Q D Q H R G Q I G T L L P K G Y G
RANDOM_7440 L W N E I E M D L Q V D Q E D Y F L P C Q L S S K G A H A Q D K M V P P V G F R N I E Y G A M S K P G I A N S I
RANDOM_7441 S T G V H K S T G K T R V I M S V L C L F D E L K L T E P N I A N V M G M T L R W A T E V V E T K G A K L L S L
RANDOM_7442 S Y N V E K D V R A F Q H G N G Q K N M C P L M G I N D G G D K A V C A I K C T R T K A N K A A L A Y E L I L P
RANDOM_7443 A I R A Q V S K P P V V G I R L Y G E S A R V I P A E A G I A E P D D T G F Q V T E L L L K R I Q V Q N E R F Q
RANDOM_7444 V P L L A I G A N K S N V D K N L V P F I S F C A T A G L K D G D Q T S S G T E S V Q L S G V L A A G V I G T P
RANDOM_7445 G W E T G Y G A D L P Q L N M S L S G T C F K V Y E K D K P V T L K G A A Y T M G N S Y K N D C P D E L T V S S
RANDOM_7446 L V D S C L I T K S S D A E F R A D E G L N T A G F T Y I N D N D E S G S K A T L N T Q A P R Q N R I N K V G T
RANDOM_7447 L S A S C S Q L V G T K V E S A F E A N D D D V C R I P D Y D F L R K L M G N A A I G E H H V L F Q R C K E I N
RANDOM_7448 E N D G Q Q Q H P S S Q A Q E T T I K A S I L Q L E K P D F F T L L L Y K A S W L Q D S A I T N V R H Y L S C R
RANDOM_7449 Y T M M T E V E V A L I L A S Y I N N T D T R G L L V F Q V H A T M M Q Q V G K T V P E L V G L A H H R L G E K
RANDOM_7450 W C W T S A S L D L E E L Y K V R L C L I C N C G T P A L V Y V S K C L G D Y D T M E R P S A R F S W H M L A L
RANDOM_7451 V A T N K F P F N W A M K P R G D N I R L A I S T L Q L F A Q R W C A W G E V A G D D L F L H R E T V T G L L L
RANDOM_7452 Q I Q T V L G Y P I P M K N P I V R S F N K W R D V E R Y L K G D Y D V T L F L K G A A H L A C V C V W P V E K
RANDOM_7453 I M P G W G G Y E K F Y N E K L L K L N D S L I C E I K I I A T L L G Y Q I D C E I L V K I A M I L A S E M R L
RANDOM_7454 L E G F V V G K R G S Y G L A E A D P L S Y N D E G L A P C A G I E F G K A G T A S N K G S A N M R E V L L E Y
RANDOM_7455 G F G L Q S A G L T L E N A F G G H G L N C T L R V P L Q Y T Q T P L D F I M N D L E Y K A V T R L G V E Q P Q
RANDOM_7456 G L S V A L G G L P T R D S Y K C A G C C A W C G Y K F T R V A D F Q A V D T D I Q P D P F C T A S S R I P L V
RANDOM_7457 T N S D V F V E L I Y L L V A K P C K V V E T A E N G S G Y K S L N G C L Q G I N F E V K S Y A R V L I M G G Y
RANDOM_7458 I V L R R H P I N Y G R V P L P D F H E N N R M T D G R Q G Q L D S T M E I G E G R S I S D I A W P R P D Y I S
RANDOM_7459 K G C F Y A Q G N E M D A G I F T L G L D S L S S V E I G H S S P A F D R R V L P S V S G V K D V S Y L A V E T
RANDOM_7460 K Q G R D K G Q R H A A H A E K L M D L S I V A E V I L S E L A L L Q P G G P A S Y Q S V S C T K T N F D K Q R
RANDOM_7461 K I D T S N D S S V I G E D S C D A C I K E F S S R L P F G L Y Y V C A V G T G S F G T S P G Q K L D D G F D V
RANDOM_7462 Y M Q M N D G S L G G E G S G V P S K L F I W P S E Q A L D N A N R W C Y S I A G G A H A K A K T E N L F L V D
RANDOM_7463 D K Y D P S Q S Q S C C T P P K G Q L E A T S T R S T A T I R A T E N L M N I K K M F S D L S G A R E D A T E V
RANDOM_7464 P R G A Y S N K S S S D G R D V D K I A I N L E G P T R Q I K L N G G K Q L S A N D D Y T M L E W Y Q R P R G
RANDOM_7465 L F V K S G N H R I K V P N C K T A T H A T I G N G F C M T S T V S T Y G N I S P E F I I C I I C V V F H P I H
RANDOM_7466 P K A K Q R R G C V R A Y A D K D P A E A D F Y P S V Q C M L V P A I L A N V G S F C R F Q I R N I S L G F L R
RANDOM_7467 P V K K G Y P N L T V L R S G N M A P Y V I H R Q F S H R R T L D Y Q G I G S G E Q T G K V H D L R A G H G F
RANDOM_7468 G R N G F A V L G V I E N A S K V C G G D V R V G P Y V E K L I L A P P S S Q Q G K Y D P L A C V C E A W A S P
RANDOM_7469 K E A V L Q I W T N R E V G S S G L K S L L I A N V A L T P A M Q K H Q Q F F S K A N E T C W P L E N Q M G Y F
RANDOM_7470 V V D S H A Y G L S V K A R I Y K G K L L P F Q H K N I L F L N V A C A L K S T V D R L G G G G L Q S Q A Q V R
RANDOM_7471 T L F Q E K T T G E D F A C T T I L F T P S G K L T P M N Y V L P D S P Q G L F L T L I L T K G R R G G E D V F
RANDOM_7472 Y G G L T L D R I H A F L V L R A L G S S E K S A E V A K T L H T P H K E T G R N V I A N A T D A T G L Q G S E
RANDOM_7473 N Q N L T K N K A P I I L I R A P V F T D M H E G F A S R H Q V I E G G F Q L L L K D R N A S F R N L N S L T S
RANDOM_7474 A Q A C C A Y T S Y K S N S G R H C C L L S Q I T K S T I A L E I R S E L D Y K L V M Y L M V D E Y E G D A R G
RANDOM_7475 Q Y D V E G I D V A E D G R K K D T Y Q A V Q K L R N E D S A A R S K K G K M R D T M E A T G R G V A A V H I D
RANDOM_7476 L S V F T I P L S F V G S T G S S V I R E L R L R C D S K A D G G L N I A W K L I K G P E I V H K T T R A D K E
RANDOM_7477 Q G T M S P T R L L L Q S N L L P R F E S L P T P G L A T D H R D N S F P H V S C L T T M D P R F G D R L L P Q
RANDOM_7478 K L A D G A P N E R V A S N W N K P D F V I E Q L P P L L H G D A A E A Y K T L F F M F L D G E F F K L E N G V
RANDOM_7479 Y T R L Y T G I L T L K C M S I E I P T K Q E S L G L S A D N R L S A A I I W R Y Q A G T K L L S G D T R V V H
RANDOM_7480 R A G H Q A W N N K K L Q Q G F V V E K E L R S K L H P P K D D E N Q L N M V I Q L I N R G P V F N G Q C D G W
RANDOM_7481 K Y L V F Q P S R N D E R A W V G C Q D N H K S S Q M V D E R N S W Q K V R Y A E F K D D S Q E A A F A K A S K
RANDOM_7482 S K L F R K K Y G R L G K C Q K E C T V G R K I L M A H K P R H D L S I V S G D S W N G L K P N L F I G L D D S
RANDOM_7483 R F E I L N A S M A A T M W G E S M C E V N Y T G E P T T A P E Y G K D S G V D A E L H D G A T T K V L P R A S
RANDOM_7484 I A A D Q L G K F R T K W K P K I D R P K G S E R E G T L Q V Q Q P G D A G N T L R A E M A V L I A A R L I A L
RANDOM_7485 E L Q R E G T K P L W H Y K C L N W D L L D I F R N A S E D L A T A P T S E C N N R E K M F K P Y F T T G Q G F
RANDOM_7486 S G I K L N D E V S G L N I A L E D I E S C L N Q L S L N M Q G V L R D M I F T T I R I Y S K P S L G T S A L E
RANDOM_7487 D H V P F G F K K S G E P C Q N D G A A I C L Y S R W L D L E P A T W G A D S H N T T A W F D N F F S L G I A V
RANDOM_7488 S L G K V E D M E T L E P T H E T Q A C K Y E V T S V D Q P L D D T K I K L V G K A V F K D H K C A T I L Q G K
RANDOM_7489 V L V G V N N V S R S A N K N Q L V A K H G I T D E K V L Y D V E E G L E D L P L S L E P F M E I N E A L E E A
RANDOM_7490 I R R S A W G K R S Q Q E P I W W M Q K F D Q V K V S G D Q F A V V L P S I E P L S I R G K S S G T W W A R H K

RANDOM_7491 G F V C N A S E G E P Q K A F L Q Q K K A R F P L K I V S E H Y T V K F A Y V A E C V I G D R F V Y E E P A H S
RANDOM_7492 S A D S G N I R K P A N I T S F M E F H L A G N G V V A P Q F R K A G D T T P M R Y D V R Q K K E D H L F K A E
RANDOM_7493 E F D L I I P L S R A I P I G I Y D R L L D T L A E N D N A E N D S A V P M E G S L T D N D K V G G N L I R N R
RANDOM_7494 N G D K E L S W L T Y S S C G R A L S R V F H Q F F A S P V Y L V E T L Q S V V L S R S E R T G A Q L S K L V T
RANDOM_7495 C I W D D T L S R E A R Q E Q P L N V A I W S R G K Y D K M L P P L T L T R Q E R I Q G L S K G D K Q T R F K E
RANDOM_7496 V D L H Q A P R F N P Q P V G I K A L N S E L P T L H K E T S E V A H V T K Y I H L P K V A R P I A V Y H I Q N
RANDOM_7497 K Q A I G I E P A T G I S P T K D I Y G D Q K S H E P D N I K T G D I Y K V D W W M T E L L V C A I G V Y S V D
RANDOM_7498 H V H G G D N L Y L S D L D D A G L I Y G L L A G Y K D L F F L P S N R D L L Q K Q L I L N G Y F D Q L V S S P
RANDOM_7499 Q V D K M G A S Q N V R L F V R R E A Y A I E G G E N E G P T D V T A F A S G F N T L S S P I M T P G C T G G T
RANDOM_7500 T I G V I R Q E R T M T S A I N E G I L K L Q S A N R D T Y R L L A G A H K A L P G P L A M Q A F S I M R M I A
RANDOM_7501 D A L E A A E L N G L L E E V Q P F V P V S D S Q S H S A T W G L K C S Q I W K A Q N A P D L E E G L E P N R V
RANDOM_7502 N I T Q V Q T K Y F A K A Q L P V H D I K I L M A K A K I Y K A K M R T A T L D I Q K I S A K A F P A G E L D T
RANDOM_7503 Q V V T Q T G G A G G I A E Q F T N D P D K A M Q W I D F L N V D I S L A Y G G P P V E F Y R K D K A T L S I S
RANDOM_7504 R S G L I V A W F S S A D E S P Q R L A E G T L K Y S L K G T D I E P H E V K W Y L P Q M R L V Y P L G A V F S
RANDOM_7505 L Y Q S P G L N A W V A A R L I M S G D Y V T T F T Y F S R A E S K C R V A Q D K F I R R G S S N P K K S E D S
RANDOM_7506 L T I R N L E S Q L E F R K M I M Q N L K L Y Q E V L P I Q G G E Q K A G C E P P E W N L A D F L K L P T I S S
RANDOM_7507 V N S L G I R V P A Q A T C R L Q R Y C A A S R R G E L R P K P L T T L S N L M V N D M D C Q E G Q G R E Q S P
RANDOM_7508 P A E A S Q T G N L I Q V P S S R L G M C N M R R R A P T P E A Q E T Y V S S T S G A D A R T G I I E A S G M Y
RANDOM_7509 A T M A E S L L S R L T G L H S A P T E Q E L Y F Q S T D L S E K T N K S K E G E T R M W P H V N A G V A T K M
RANDOM_7510 Y T S N M A V D S T I N F K V Q G I V L S F R D Q K Y K K A A T D I D V K T K E Q H G W I I R F P V I V G S S N
RANDOM_7511 V A L R D V R S V G A R T L V Q V D P G Q S D R N Y C Q Q T G I L M F C K G S G S N S L A K K F D I G S Q Y G E
RANDOM_7512 S R T P P V S L E E V S T R F E E Q M K Y R N A W F L G V L K L K D P P H L I H H H G R V T D T T V L V S A P D
RANDOM_7513 V G H W L V T I R P D D D S V A N Q V I P L I D V N Y V T N S D M G L W S S P L E Y N D V G K L P R F R V K E V
RANDOM_7514 N V I I S V I T G R S E V Q G L F S S V M L T V K P L T L T V V P V L R M K P R T V I E A N K E L K V K Q V P L
RANDOM_7515 A R A S K Q R A D Y K S L I I S V P D P D A S I L K C A L P G N K S Y S G D A A G Y D S P S M V Q I I H D N I S
RANDOM_7516 F H P E N K A G P A N S L G D S T I R G I L E V E Y N R P S N R Q I P M T T A Q A V D S L L Y S A N N G H R W F
RANDOM_7517 F H T N N T M L L V A I K Y W S T D V D G Q A K G A L F G D T E S P Q F N T M A Q R N S L I F E I S E T E P E E
RANDOM_7518 H P T Y L Q L Q R L S D T G S P P A Q A N L P I V P Q L K K G S N T A S Q R V T L A E A T I G L S V N A C G T K
RANDOM_7519 G Y Y G R E L R R P V I N V D E M S L V K Q I D P Q N V V F E A D D Y L L P F R I Q L E K G Q G E R K D P I D P
RANDOM_7520 K I T E K S D D T F E R A K L I V P K T S S A S L V R L T T M D Q Y L R H E L E N R A A P I G E G S A G D T A K
RANDOM_7521 K P F N L M I A G T P A K Q P P P N E S D R K T H A Q L E R V N I A S L C V L R N V T T E D S L Y S A Y L R F Y
RANDOM_7522 R P T P Y H V H G I N C P R K C G R S V L I D D H L N A K T N T F D I L E N Y K E S A A F L R D V V A S Q I I P
RANDOM_7523 S I S T F A R S L E I P S E V P R I F S R V A S D V P I L L T T P Q A F A E N D Y V K D L A S Q G S N L Y Q W F
RANDOM_7524 S R Q Q C A L K G M T F L V L K E V N A A L N T L T E A Y T L P K A V K P L V H E N V L Y S Q S Y I V N E Q K T
RANDOM_7525 Q Q G Q T E A E S A K E V R D K S V A G F H R M S I D T N A E A D S N K T A P G F G N K P K L I V Y V T H H P I
RANDOM_7526 L L I K V D Y A G T R N G G V V Q V T H K G F A P E L V C P A G P H L R M F V M D S N S W Y T V S T D E G G V P
RANDOM_7527 E E Y V E Q L A Q P F I I F K K A Q Q C L L L H V E R A A T P W E N T V A V L I Q P S V S V S G V A L D P K N I
RANDOM_7528 L W G P K F A V I F E G N H G A R Q C A L E R D C Q A S D Q F K E T V T I N G A G A N F S T D M C F E S E V P I
RANDOM_7529 M A N Y F A H E P G P E L P L K D E L V E E E V N F P E T S T Q G K Y T R E K S S V P G Y T G Q W P P P E F Q N
RANDOM_7530 T N I A F V F D L I Q A A F T S S V E S F Y I S L H M Y S I L L Q L K R V G W P D S G L G I R V K M R N T R N G
RANDOM_7531 F A K S N T L K I V L K P R R Y N E D L Y E W P D V F Q I A A R K E T T A E Q L H S F T H N Q K I D H G L Y T G
RANDOM_7532 Q N G E V L G G V V Q A T R N L R W M N S L T V H H P L F H I G T A E I L Y K R G S K W A V V Q Q L V G G V V E
RANDOM_7533 C E K I P V N S C G D E K P D Q I K C L Y Y P E A T R S M G Q Q H K T L D G P L S S A A K S Y K L E I A S S Y S
RANDOM_7534 A V V D E D P H L V L L Y Q E D I T I Q E S I R M N V C A G N F D G K I E P Y R N E Y K G Q R Q V E V L A C L H
RANDOM_7535 D R T V F T A P N I S L D Y Q N V P N W E R D A P D V E L Q V K T G R Y W T E K A V F G D G V E Q I I M E I A R
RANDOM_7536 E K M S D A D I C L I S S G I C A Y G E G Y S G N L D K F N H S A Q M D I H S L A V Y H Y K D D P E K Y L E G T
RANDOM_7537 C L K K N E S T G R L I V V Q Y F E F Y A L A K E M P L A D D S G V M P R G L Y S Y C I L D H V S S L S L G D P
RANDOM_7538 M E L A S I M Y H T Y K G I D D Y Q G A Q E L Q R A Y L K A G I R N T I I N L D P G G I F I S L R E G C T C P P
RANDOM_7539 R V N S L L E S Q A S S A M G A A L T T N E Q T R E D A L K V I L P Y F L R G A H D Q H P F F E L L Q I D P S V
RANDOM_7540 Y S T L I V K D L L E A E T V L G G K T N E M Q C L V P N L P K S D G T Q K V L Y M S P E K G F K P G H N K M S
RANDOM_7541 I A M C R F V L E G S A T Q V H E P D K K Y T K F T L V A G V C G Y T T A F Q A P R E N P W L K A W E L R S G D
RANDOM_7542 E L P L V I K P V R A V D S M F E D K V Y T G R H D S R P A I G Q K S C D D T V D K G A F G E A I P A C F N K L
RANDOM_7543 L R A G W C R N C V Y D Q Q A I I Y E Y L E I S T S C V P Q S T L K I F I T S Y L H D F F L I Y N V R A L S E L
RANDOM_7544 V L N A I G S I S L I D E S N I K K S N E H T I D G M S R I T L E Y Q R T Q L L T D P F I L F S S E T S P N S N
RANDOM_7545 D R G L P Q F A I V A F Q V S Y C R T V C L A G T D G L I T W V Q A V Y D A G V T E F V Q N L V S D Y T V T E A
RANDOM_7546 T N G D L C V Q T S R G G W P C R M A T G G K N E A R E N S T S S A D E K F F D L F G S M R L Y G P R L G I R G
RANDOM_7547 Q Y E N E T A D V T N L K A L V G A S G A E M L L P L K Y V S L E G G S P P Q S T N Y D W I L Y N P P A I E A L
RANDOM_7548 A S V L I H M V Y E K L S A P L D K G K G K D M S G T M E L G K L N E F V L R T N F D S D P L E L S P L L A I C
RANDOM_7549 T A D S D Y Q I S H R P S E Q N N C P L S P R D L S D C H K K R D Y L N A I G A E A E Q E A G S V A G D D N F Q
RANDOM_7550 E C E T D S G T P M G I G L A R S A E P E N D A D T L E T N I L D K I C L R V K G I D K D S E K L H I A L L I R
RANDOM_7551 P H S K Q L Y D M G G A G G S P E A W T M N P S D Y C S E A Q G A W Y K S D E T S V A R A Q M H K E F G Y A A T
RANDOM_7552 C V L V E K Q T L K K A K A F T G E C Q L P K I I N E I T I T D Q V L S D D T K D K N E R A G E G L D M A D R S
RANDOM_7553 S T M V E V G A Y T Q S F H S V V C N I S E E V T I G R G A E L L N V A P Y S A S M P T S V D A L G F E Y I G T
RANDOM_7554 P P M E E D E L F I L G T E F N R I N Q I L P K V D F D T Q D I T W L L L V Q A L D T L R L P T K K R N K S L H
RANDOM_7555 P L G D V V M I R K S S G N I G S Q C Q F L H Y I V K L L S M I A S D A P E E I T H Q C F G Y P K K S T L E Y K
RANDOM_7556 I E L A L D M T I I R R S H L S A A D S Y Q E R D Q D I T E Q E C Y R P F T V Y L P L Q A R P M S M T E D D A S
RANDOM_7557 W A T I H T S T V V K P G S N H K W T P N H G L K P L R S R E L N P N N A E C K T K L G Y L R S S L L H E I N N
RANDOM_7558 D H I K T K I P L L S S K V R C S E R D L G H G D E A N Y P S L D I P L R Q T A E F A N K Y K P E T P G A S G E
RANDOM_7559 C V S H P L K K A F F P I M S R N H V R R L K L I Y T E R P K M V G V G T L S W D G P A K R K E N L G I L Y S V
RANDOM_7560 A H L P G N F N L T F W E G Y D G H A V A G L C S S F Y Q Q G L L M T S Y N Y Y C A A F G S R R Y L K K I H K R
RANDOM_7561 G I N N L P S V F M V S F L R T Y V G L A V G R L A P V I L A A L R S V V A W D K Y T V K C A S K S A L E A Y I
RANDOM_7562 P S L K A E V M I H S R S P I E M I N N Y V E K I G G P E N A S V G S A M T A K A S A F D N A E K S E A S A R S
RANDOM_7563 K L A L S F H V V Y G R K T I M A I G G T A L V A C S S E A R A V T D T P T M G D R L Q Q W F L C W Y S I A M I
RANDOM_7564 K T G G P V C K S V F R C D S N Q L G L I G V Q I R H L K A C K A K G Q T M Q V A V N K E N V E G W A K V E Y T
RANDOM_7565 G A D D Q I R W L Q D G Q G S I V K T R N R E K W V W P D T I S E R F L R A E K R C G C F V T D I D K A A S E T
RANDOM_7566 T D R M L E L I G G Y E G N I S Y V Q N V A G N R W A L T P E C F D P K N R I I L L E R K N S H R N V E S E K F
RANDOM_7567 H H L N E S S R I G L V P S V I D S C S F L Y A G L R A D E M K V A E G V L I D P A Q E V G G V G T S P D I P L
RANDOM_7568 V W V A S F T I Y K N L T L E N T K A L N P V G M K F I P Y Q Y A R Q G I R A R A V V L G G D K H K Q F C N N V
RANDOM_7569 Q D Y L T A C L A A A S P E L Q V P Q T L K T D V K F L K T E V R N G N Y S P V G G G P M T P F Q D L T M T G P
RANDOM_7570 A E E T S R P G K F D I Q E A L A E Q G H R D P E G F S V G V N S M W V E D G C E Q C Q A A E A F A L L D V Y E
RANDOM_7571 G A S V A P N L N S A T L E I G R G V A D S D G L K A A A A D S N G R P R K R P A W V F K D R L V R A V D D L D
RANDOM_7572 A V I A I N F D L F A P L P L M T M D W S R Q V T Y M P L G T S G L V Y Q F W G G P E H Q N W L L C G T R G R D
RANDOM_7573 E E Y N G R V V C K E V P A V L C Q P L L L P V Q D A G V P R C E D N K Q P W N Q P T S S K V A Y E E Y S D I S
RANDOM_7574 L N Y Y K I E P R S E Q R N K P Q S K V A K C G K G H L V T A E I Y P L P K K Q V G S E A Y N D L V E I L V P A
RANDOM_7575 C Y N A S I Y K H A T N L K E K D A I I G R L C S D V V I P V E Y I G L E S D W D G D A L M L D G M E V P S A K
RANDOM_7576 K S F E Q D A K F G S Q L H D D V K I L G H C F L F L P T V I S S I Q R K M S D F I A K S P Q F S Y G D F S Y D
RANDOM_7577 L T Y R E Q N G W D G P T Q E S D Q L E A D P E F G P L G I E S K H E H T N A V E Y N D S T V A H S G K N H F S
RANDOM_7578 C G K T G G A P K A R C R E S R P P Q D K Q S I N S A R S A F L I I T Q K L K K K N C I T N S F A P G R E D D V
RANDOM_7579 D T K Y E H A C V P E I L P D R I G Q L L Q G R E I D M L K V L I A K P N Q F K M F K L G V N L K P P F D Q T I
RANDOM_7580 R M D P N K C P D P W G K G T Q T F C L T A G G L R F G N T D L A G E K A E T A S Q A E G L G I I R K V Q F E G
RANDOM_7581 T T A S D T D A T A N F Q I A F Y G R I D R G Q G E H D Y D L K I I R K Y R V L I P L E S A S R D K M K E T G V
RANDOM_7582 Q L P T I P Y Y G G D T C A A E V Y T A D G F E Y K S T F L I S Q R L H F S P D Q V L A K N D F R K S T D G E I
RANDOM_7583 E I A Q R Y Y K F Q D Q W G E V A N E T T A Q W R E P V D A Y M L K K K T G K A V L D A M L A I S M K N Q M Q S
RANDOM_7584 S L D L W L K P D A R V I E N A S M V L P S L K S M A V T N G A L D K V P R D K A A D S K R D L P D D P Y G N D
RANDOM_7585 R Q G Y E V K E Y V R G S Q F C T A C D T N L N L W T L C G V V R K H C A P Q F L I T Q E W N W P D L P I I A Q

RANDOM_7586 E R R E N S A A T L V Q G F V P G K Q A N K K D E F G M S E Y T T P S N V P L S G A F P G L E G F A I N E F E G
RANDOM_7587 P V K Y E P E N K V H P E S S Y T G V S G S F D Q I K I W S S R L R G L T K R A V N T V G Q G E E F D G N H K G
RANDOM_7588 H A G E Q V G C D A T D Y V K E Q Y L H K A G D S R D V Y Y R S G T A C A Y W M S Y G G Q P I E I H F G H E V N
RANDOM_7589 F Q L P N P K G R V F F P I E I A Y K S V I A I Y S E A E K L A M M T F N T A M K Y E L L G M E F K L L L Y Y I
RANDOM_7590 S Y N E D T Y S N D V G F H M H Q D D V Y K Q G T R D S K V F K I G E L I F E Y Q G Q F F L D Q L A L K Q T C L
RANDOM_7591 R T P R G M S L F P I K Q E H A C I G V F P K Y M D Q L T V P E D R L A S W A S W P L L A S W R Y G T V A N F D
RANDOM_7592 K L G P K E M S D D L M K L P A D S H A D G A D E P D T E R R M H A A W I P R P T A E S K P F E K E E F L A S N
RANDOM_7593 A K V L L L Q K T P V S D S A P A E A E A F A Y L L C G G P Q S V E T Q G Q G F Y K V M D I L G N A I N S D P L
RANDOM_7594 E R S V K K G R T G P S R E M L D R D M V G D D Q G I A N L Y L I N K A R F A E G A G M L G M P V M L V F P H P
RANDOM_7595 I H P A T P T N L G L M G K L A K T M P K V A S M I V Q Y K G Q G Q K K F P N L A F P L S S L F V A D P T L V A
RANDOM_7596 C T S A D Y S Y Q M K G T Q K F C D G R D A E C N E P V A L P I I A K D G E S A F D P A F T M E N A L A F N Q F
RANDOM_7597 A A P R M T Q S G R D N K Y T S R D V K Y L E V L L S Q F L R H C Y L S R A L A V M N P C R A A P K P E N S S Q
RANDOM_7598 F L L G D C L F C A G A G C F K N G Q D E Q G Y V F E Q R G Q Y G E G I R V G A N F T D V P Q G R K G L P R R G
RANDOM_7599 T Y G F S I Y Y H Y E R E S G D R Y F I R K L E V P G A Q D E S P A Q R M R G S T Q L G T Q H L A D V N K E S I
RANDOM_7600 F G Q Y Y R T A G A R N A E E L H D K T L C G D A E V Q A K D C P T E T R G I E C Q G V S A N Y L V L P R V L C
RANDOM_7601 E D M Q N Q V W I Q I R Q A S D D E V L K L A C A N L T A I V A F A T S D G C T L R M S S A Y Q S G P A K V Y D
RANDOM_7602 I L I Q E S G V P G Q T H A S T V V H P I P R A D F S A I R F G F P S D H L D V L F F I E G D Y I R G G C Y P A
RANDOM_7603 S A S L G M D G T D K K L E Q T A K Q F N E P S K T W P L S K V Q S E D V K E E C F S K I A N V K V H G W L K M
RANDOM_7604 N T N P E S N Q A G N T L K R R E H G L L K R N P S I E E F F L K L E D C K V G P R K A A T E Y E T L S G Y I V
RANDOM_7605 I L L A T K W K R F S G V A Y T K P T E T P I S W L V M L L K G M T N D K G L D A I L P S P K G D D K G C P G A
RANDOM_7606 L D E E L P I L L N P L G N G Y N E I V Y K M D Y F S N C L P M Y L A P Q P G Q Q R F H P G N V K D T K E V A E
RANDOM_7607 K N A V E P A F D I D T V L H N L A S N I G E L V R I V Q M L R D D W S N L N Q L R D A V Q Y E E L E A Y H I P
RANDOM_7608 I Y I A S F V Y Q I T Q Q T F P A I A F P A G D G I P W R S A E W Y W E N S R R T E G M S Q G H P V Y P L Q L P
RANDOM_7609 Y P L K D A Q T W K N W K S Y F D P L L V E S S C R T A D K P R R M A D G S E A N D D F I N G Y Q E A L D P L M
RANDOM_7610 L T G G Q P V T E R K T N S I E A F S S V V S T T V N S V P E E K K D S V N V R S R K E N Q V V S Y D N V E L G
RANDOM_7611 E A R L F E R Q A E Q V D V A K I K T F F P G L G A T F N T P T D Q V L L I D Y V Q L R L F A Q L G R A S N L V
RANDOM_7612 E E R G V L G V S K E Q V I N S M R E I I M Q S W A Q A A P I I R F V K I L Q N G A R T L G L P D E F E K F Y D
RANDOM_7613 G L R K D M V W V M D D L L A Q K Q E I A D V G T L T W F I E D I L V K G A D E A R A L R V A G R E R L G A I E
RANDOM_7614 D G F I A F P G D G A V F A S A D F C R S P A K V K E L T G D S A F I D A V H D T R F V S K A L L L P W P K D F
RANDOM_7615 P E V L K S V Q N T S M N S E S E L S G G S E L S T K Q A K S A Y A Y K T S K F T I W L K V Q S P S T S T R L E
RANDOM_7616 A A L Q H K Q R Y L K R P W G A M W E Q Y L E D G L Q G G I D A K K V V H M P I S R N L A Y A K V K G P K Y A V
RANDOM_7617 L M K I S P G R L Y L S A T L C K E T T A T N G D Q P L N K L L I I T K P L Y Q F R A E S G Y M I E S L I I K E
RANDOM_7618 A G R G T Y M R H L Y S T T F R G V A L T M S S M G Y L L T D A Y S A K P G S I T V N M G K L P R P W L T D I A
RANDOM_7619 A H I D R L L R L Q S R P L T C N R V A A S D T I L Q R A E R P Q K V G V G Y S C V P F Q H N I R K R R D E R A
RANDOM_7620 G L D K A R S W V Q K E Q Q G A Y T D K I T G I V T R S R D T Q M V Q R L T D G N K E S N F R T L V S H I R W G
RANDOM_7621 V Y V T N E T T V N E D D I G P K E S L K K G T H E R Y S R L S L E L T Y G G F L R R R L N E P N Y L E V I E L
RANDOM_7622 H P C F P R Y L R T G A T G L T A K G D I V S V R A L L R N N N A F N K Y L N E D V K A G D I V E N A R G L V E
RANDOM_7623 L I R K L T S I A D A Q G R D M A R A G S P E D T G T L S A V K Q A N K T C R A A L N L A L K N S T S Y E E T A
RANDOM_7624 P L V L T G Q G I Y E I I V E R P E R E F S V A S A T I K A D P N V D S T L A S G A H S I G I R K A N I A L G V
RANDOM_7625 P E R A E N T G S V M L L Q K T T P V K L T G G K A Q L A T Y L R G D S S R G F G C M Y T G N F P V L N D V Y P
RANDOM_7626 K Q E R T E T P L Q T N D D A I S G S A Y A L N A N T D P L R D G G L T Y D G M K A K L V G Y A D Y D S N I R E
RANDOM_7627 N K D W G A Y R N G D E Y Q K R G G T G G E D S I L T F C P S F G R V V F Y V F N A K L S S K T T E A A S L I D
RANDOM_7628 F T A K T A I I T E T S K S E E N N R A S K D A D E L D S P F S V W L V I C L E S V R L G G S D P I S N P Q P T
RANDOM_7629 V S L T G T N K L I L Q A T N A K L G C D F W S V T L A M N D I A I D A R R G V T G V L L V Y E D Y Q D L F K D
RANDOM_7630 A A L L D A I S V L F M L D L F G A T V Y F T R D K S F G D P K E S L K V A P K L R T C Y E N V T T T G I Y R D
RANDOM_7631 F N W N S T K D G G L M D N E E K T L K T N D I P L V S I D L Q K K S V G E A R P Y T Q N A M D K T Q Y C T L A
RANDOM_7632 A L K A G R L G E E V P S P I K F P A L G L N T Y Q I E L V S S I M I F D S A K L S L D G K L L I E S Q Q F G F
RANDOM_7633 S I P Y F S P I M F A A K A V S N E K I N V E M V T S I V R R Y D E I P R A G R T R Q K L A C I N K G G M R S S
RANDOM_7634 E V H G F E D V A L L T C V R Q I E T F F C V S V I A P A M K I N E I P L G V V V W D E A I P E T R G A T H N L
RANDOM_7635 K F S R K G G A D D N I C Q C L I V L Y I Q D I S A N K E Y Q E W S N A L R R E S M T F D K K A G I A V K N V Q
RANDOM_7636 F P F L N K R V I L T V F P L G P L R T G Q Q T V T V Y G K A M A C G R N L L H Q K P F E K I S A G S I V G G S
RANDOM_7637 F G A R I T T F K S L L T I Q D D E E V R S F C Q L E A D F C D G A T A S S A T Y V H H D I S V T Q V A V D L P
RANDOM_7638 S L T R E A P R Q V A P I L G A S Y M N P E L Y E N L G R A E T A K I P D K E Q Y G V Y P L R Y T Y P L S T A V
RANDOM_7639 L T G N I G S S D E L N E K K K S V I I K R I R C A Y V N V S M W Q P Q S T S F I V H N N T K A I G G S Q G G A
RANDOM_7640 G F A E Y L F I R A L L A I S Q T S A K L L A R L V Q L R K E N S P I H S D L L N Q A L D W Q Q N T I G Y L A R
RANDOM_7641 L S L I L I T Y I K P I D H K S L R Y L E M L I T V L F S Y C D F L E P P E L A R G E S F Y E E L V L Y I V I T
RANDOM_7642 P I V G S C Y G Q L Q V E V L K S V H L A R S I T H H S H F R G R T L D K F R R V D L A A S W V L L G Y F N A S
RANDOM_7643 F I F L R E S I V G S L N I S N G S L G L S S T Y K F C A D G V P K F G N Y Q N R D S R D K E I K L N R L I K S
RANDOM_7644 Q K G F A L L K G S L L K Q L S S G L G L A P V C Y V P R Q M R P E S D L S K G Q W R I C D K Q V Y G S S R H V
RANDOM_7645 F S T V V R E L H E V T F Q L V N T G N E N D L I L V R K M K Q H E T K F G G H V C S L I P R G S H G K D V K I
RANDOM_7646 T R Y D W E Q S R L T A Q V H A V Y L A V L R R K G G F D P G I G N G Y T L N G D W T G P D V L S A C K T S V A
RANDOM_7647 P L V A P W S L A V P G Q S L D L T I L P I F L K A P E P Q P G G T H S R E S I R N T E V S K I S D V G L V D G
RANDOM_7648 Y K I Q A I V K S A I I G K D I Q P E S V G L R D I Q Y A I A S N F Q P E D Y I G T K S N D P D T G L P R Q G G
RANDOM_7649 P P A E G A S S I I R E L N V V S V K L G G G V Q R C H D L I I R L R L R R R S W G D A T V S Y G K I G D C G D
RANDOM_7650 D G A N D V A G S P Q Q A K Q T R G K E F A A T V D G A K S R K M C R L R L K V K R L L T L Q S L N P P F G S V
RANDOM_7651 L E C Y K Q F G M L G S Q N P W W S I P Y T T E I V S D R P T T V L E E T P I G Q I L V G T K H E A M E I S M S
RANDOM_7652 H T R S D L Q R P T E I S Q H K G Y C H N K D V K T R I L A I E A P T D T N F S Q D L D D G E F P V N E V A K Q
RANDOM_7653 A L A V N S K A T D Q T H Y C G K N R Q H E V G Q E D K I A F D Q R A M D D S A K S I F I R T S L V I V P A S F
RANDOM_7654 G T E E R E I W K S L D L C K E Q P V G T G P V K Q H N Q G S E A R I N S E W E A H K S G L E Q Q C A N A Q P S
RANDOM_7655 N L T V K N L K F T I K R N W P R L R D T R L F P K T R Y H P K K C S L D Y L F G P I F E L T R Q K Y I G A T I
RANDOM_7656 L P S G P K E L C P T P E V Y M N A R A S D T G A A A E W L T H Q G N N Q I Y A V V A G D F L R R V G I L K L D
RANDOM_7657 I I E L T S I R G E N L T L A S Y P T I S E S E E E T S V G I T R F P S T Y A T Y G A M A A L I S A T R D V A S
RANDOM_7658 D N I K T W T A D A Q P N C V R R D I T G S L E I V R G A Y K K Q H L S I A D M A H V F F N W R I C N A E V I R
RANDOM_7659 A I V T V D G K I N W V F I R Y S A E T F C Y L G Q S T G Q N H Y L G A A H I P F L S Q N Q K A N D K S A C F I
RANDOM_7660 H K T Q R P G G S L E F K G S P I G A R A V K T V A R G A C Y A G F Q S S P N P I Q Y Y G F E T A H K L V W S G
RANDOM_7661 V L S S Q A C L C A N H G Q G E L S M Y L A A I Y L Y S D K G G L T G K S G C N G F T L N K A V W E G L P N G P
RANDOM_7662 D V P W I N P L P D S G H N K W F V V E N C T V N L K E K R K A T N D N T G K I K N D Q H A L S L V E S D N F I
RANDOM_7663 R L P V N S N I T T P T S F N V C S N V A Y V V V Q F D E V G G T V V D S E S I R Q G K K Y H E D P H V L C T G
RANDOM_7664 E A L N R V M R I T G V D L T V R H Q L E R A Q S R I G V P S A D V Y T K I F F S S D Q L S V E S D L L A L S I
RANDOM_7665 F G K K E L W D C S F G V S D G R I Q M K A Q F E L E L T Q T D E E C R A T A V Q C A K M G Q V D S L I G L A M
RANDOM_7666 S L P D S V L F E K E Q C H S L V Y P K F Y S D K A S A T Q A E K M L D Q Q A T M F V Q R G L G S G I P T S E G
RANDOM_7667 Y L D Q L S G K F L K L K T P D R S E F L D P R S Y D T A N F T S L G D L A Y A S E K L Q S K T T P A D S E L N
RANDOM_7668 I A G S N E A T T A G E A G S L P D T T N P A L R D E A Y D I V K S A P S V K K A R L A I V R A I C G G F A E E
RANDOM_7669 T R L P R G D M V K K T G Y P A T A I Y G M G Q G A S G G S G S V L Q Q T P V K R G E S D I I S G L S P A S I P
RANDOM_7670 S S A Y K K L H K K V T Y A E L N P N V E P P S P A H M G L L P M T K E P W D L Q G T Q Q P L L A D F Q A T A G
RANDOM_7671 A S D L S D F S A V D Q E I R S G Y V D P Y G E A I S R W R C D C V V G G Q P D L D S N N A K T D N K I V G V V
RANDOM_7672 V F L E L E S Q I E S Q A L M L A D G M L H S A V I D A K V G L Q C L K K L V G F V M L S V F P L G L L F E A P
RANDOM_7673 K H S L P V E Y K L M H P P T T G I N F T G M P S I T I W G P P P A I A P N S E G N Q V G K S E A P Y R Y M A S
RANDOM_7674 Y L T Q M F L I V K L N R S E R I G A G Y K L P T E G L H V I Q C K S S V T D I Y P E T V H R A L L T K F A I G
RANDOM_7675 W Q R V R E P A Q G Q Y A K I G A G W T A G G D N H R I I K K G N T N L M K N N G V L P I H A V L E E I D H K M
RANDOM_7676 H Q Y Q A S N L K I L L L Q L A H H N G E N C V A L L N V I H P H V S E Y R K E S L R T G L D A R R N M Y E S V
RANDOM_7677 V V D A V S D Y L I N L K Y F S E T G S L L V P S L Y L L Q N D G G L D P Q F G L E V K H P W D D V P A K Q M K
RANDOM_7678 N R L A P A S L K E P V G Q H A H A Y D K K R G W K I L Q N K A T L C N A I S I C A G H N F E G H Y R K G W L L
RANDOM_7679 R I N L I S K L R L V F G F A K I H T I S N P L Y E A D L S V A A H V G S K G G D D K V H R P I P L Y I E T G D
RANDOM_7680 Y R V I D V N F T K E A Y A I R L E S S R I P S E L I A P W L I L L E P G P A E Q P Q T Q P T A P A A S N N A Q

RANDOM_7681 S A L A L S V S L K A L M V L I V G P V E W R G F N I M S S Y V A V E L Y F A L A V S N S I E E T K T Y H E L D
RANDOM_7682 D H K V T I D L S C P I V L L N R L T V L D P V A G S P Q S K A R Q L A A A E N D A D P K P N L L S L P Q F V Q
RANDOM_7683 N P Q T F Q K T G A A G K N V A R C A A V S C K R G R T N A D G D V A S G R L K A M I L E G A W L K L E Q E F K
RANDOM_7684 G G M I N L E S K E V Q G V K A L Y D L Q A N K A A V K C L G R R V P V F D K A P W R L I V L M H L I T S N K S
RANDOM_7685 H A K D N V L A T T L R A F F F C I F K Y L I S I S K P S P E R G L E S W F K E S A F T L Y Q P Q T S A T D A E
RANDOM_7686 V F G I S D M T A Q R P T N A C N P D A I K L V R M I K S V R V R D H V V P D P N P K A P P P C E N V S L N I K
RANDOM_7687 S D A A L T K F E L H E A V Y P L Y S A V F S L P G S G A Y G H N T T M V E R P E E V T I P E L Y A E I S T D S
RANDOM_7688 Q A S C L A Y R G E I K F L E A E Y T P L D E L I I F Y N E D Q I H P S I Y K S R Q C A K A R D Q Y V L P T M R
RANDOM_7689 M I Q V I L E R L I L D S Y D S V E V I E V D L G K P N S D L P G P K G R V L N P I L P G K T Y V T A F I K H K
RANDOM_7690 V N V G V V I F E G N S H D I G D E W N I H L G F S D I N E S A K S N S N D E S I W G F D N I L K H W L K T D P
RANDOM_7691 A E F S P L G G S P L G Q V G E A Q E S F F M S N N T V S P I P D H S N D G T S R I G M A P P N P A G D G S S W
RANDOM_7692 F V G Y A L T N N G E P G G K F P N E V L L N S L D A Q A P I A M K G M P I L E Q N E G D A P R R G R A L G A H
RANDOM_7693 V V D N T Q S C G S K E S D S V G T S Q L L A F L D P A S Q I V A Q V M A V T P T R I R Y V Q R G R G N R K E E
RANDOM_7694 L A S I N H A A N T V R L I V A D L K E P Q R G A F E F P D T T L K M S V V V A G R I R F W S F K K K G W M N L
RANDOM_7695 C L D T L Y V H R D F D V A T A M T V R I L T L S N A I G Q Y A V Q G R L P L H K S E V N G L Q L L S P L E H K
RANDOM_7696 V G C S D A Q E A M N L I D L G P V V N A N N N R S A R S D V E G I K L L L G P V Y D Q P D I N Q Y A V T K I N
RANDOM_7697 C Q S Y T A L P I V D R F T G A T A V Q E P R V I D L G L E F T H T K T Q K L S L K P T T K S N G T L N N I L G
RANDOM_7698 R T T R P E Q L E I V T G T D V G V R D A D N C R S N F F L D L D N G A F V S V S K V F D M T D P T T A Q L K Q
RANDOM_7699 V G K S G G G P A S G P N E V P G Y Q S Q S S C G V V D G T Y E A I A H T N N E E P C L T P G Q V D H M V I A V
RANDOM_7700 S G L R K I P L I N S Q E L E E Y P V G C I F M A E E L P A T V Q E G N K L D G G A L V A R G L F A E T L K A S
RANDOM_7701 T E F D D L G G A N K L Q L N A G A S P R Q A P P L D E N P Q D Y P E T E V W P Q R L Q C K T M G C C C E C A S
RANDOM_7702 N D P V T Q G V I S L H D G Q K V R Q L Y P G P L D T L A T G V A K G G P T A T T D D M I Q K D Q V N I G G I L
RANDOM_7703 G S A F A D D H T F I Y T I S T W W P M Q S L V K G Q P T S W P G E R F K L L L A L S A C A E F G L G G V G K P
RANDOM_7704 G T N N A R W F D K T V N C L N L T T G S G H I K R L L K H G H A E S E V C Y G A A D G M V H D S A V F G L R A
RANDOM_7705 F F L P H K S L Y K V T E D G V E I I G R Q I L D G P M Q K S Y I D L V R E P K M S N E V L I D I Q T N F Y K A
RANDOM_7706 I A W T S M P K C S P T T K A I P K R K D L A S D I D I D D A L A V T K L D E R W D E K A S A Q F L F S L G N G
RANDOM_7707 L C T C Y F L T S F L S D T I P R K G P Y I K T F T V M E C I L E D S R Q F P K L N H A T H L G S T D R A V E A
RANDOM_7708 E R V N H R L S R G C R F T G F D E L T Q L F E G G P Q L Y L G K S S I P A E A H V D G F I H I E V L P L A L G
RANDOM_7709 D V T A K L L T N R C E R C W I K T S V L S L S S F I V K L L I R R T G P L P I N A L D L Y D D D R G K F E C Y
RANDOM_7710 F G K M L N E I Q N K I K E V G E P N G S F Q W E T W M K A C S R D G E K S S M S M T P G A D V F L S S R F E Y
RANDOM_7711 Q M P Y K V K P T G I A F A R I N P T I C A D L G I A H H A N F G T C E P G V V N R P S G L G E V K G R L R K L
RANDOM_7712 S Q D H R I L T M R Q V N L A V S G F Y W G G V K V L N V L N P R Y R A D P D R A R S P Y G G N P V S T W D V H
RANDOM_7713 L D Q L N M L N K G K S S P T R F L V Q W S R T Q S N V N R T G L Y K S N K F A I R L K E L K S I Y K V Q G L N
RANDOM_7714 V A R D A A W R P R H S R T G K F K F I K T E E L C E R E G Y I W V K N P S Q Q S R S G N S Y L A V F Q T E P E
RANDOM_7715 Q K A P P H S A A R I G L G I S H L L D R N R F K R E A E T G G A V T K M G T A E T D D G A W N I K R G Y R A M
RANDOM_7716 S V A A P N T T N G G V T N K V C L P T P S I A C S D S S A C D G L G W S E E A L L N Y V T V Q N G Q R V H E V
RANDOM_7717 T P K K D Y P A G R I D E I A V L F A G S D V E I D L D E R S A S L Y I T D T N V F L I P W Y D F L Q F H T I K
RANDOM_7718 A A A R N N S C D P D S I A T G H L S A E I F P I E L Y K T S T V E S C G I L Y K C I A R K V L N K E Y T Q Q H
RANDOM_7719 L M E A N A G G R N V G D Q V E P F K T R D T I G A F E T S G T P D Y D A D P M N L A S M T I S G P F W H S T G
RANDOM_7720 S I L V L H N A A Q T Y V D R K E H D C Y P R P I Q A A S L V G Q A A V A G P I Q A T T K E L A G P K A W K I S
RANDOM_7721 V I E A V N L R K I K Q N T Q T K Q T T I E Y V A V Q S C V V K S V E H P Q L P Q G T V K A A R C K M K T Y Q A
RANDOM_7722 H L Q S H Y I T C W L Q A A T I S T F V E E R A K I R R E I T F D E A N I S E E P S R N V I Q G I P R P K F G Q
RANDOM_7723 R C T L S V L A P D W I T S F N C R D E N F Y E G R L R A F A Q R L I F H V E K K G Q C L A S A S D G A G M K L
RANDOM_7724 G T T P L I R V K T S N N R S V N E C V P P T R P K L N R I S V G S K V R I A I A P S P Y S I G P L I R D L K D
RANDOM_7725 K S G F S G A E E F L T R D A Y R A Q K S D R K I I D T Q F K G I N H L E S I H A K M L S L Q L Q R V G H Q R D
RANDOM_7726 Y M A S E T G S C C G H P A A Q P E G L V E T R E M S N K E T L G N D N W K H F T K T I S E K I S V L V V S V V
RANDOM_7727 L S D T R A D T F M N K I H N N I T E E G G G A V Q L Y V D F N A K V N D T S P V C A S G S E I G F H I Q D A E
RANDOM_7728 M E V S Y S N D D G G D Y V Q N L F A H N L P T A I C I E S E L T A E E G D L T F D Q K D K A E E T Q D S G F R
RANDOM_7729 L E E P F N H W A G F N Q K C L N P D D I L R T F K A N D C L G I I D N D G K N I Q P E T W E Y F N C H A S A D
RANDOM_7730 L S N S G G G L P P T V E P I H S P F L N G L Y P V D N V L Q I E V G G S S M E E V R E V K H H Q R L P P F S P
RANDOM_7731 E F T D I L H P K A E A V A Q C S I N Q V M F E R N P D P D V G A K T D F Y G Q N H L G C S Q K K C A S D E F V
RANDOM_7732 T M T R F V D L V Y I N V D G E L S L A F S A T P C K I D M S L Q Y T A N Q D D D A M L E V D E N L I S N P S A
RANDOM_7733 P G R E D N A H D S A A Q L N S E P R G A A D V L E A K T R A E N N K L R Q A S E A K G T T L F S A N I Y Q Y F
RANDOM_7734 N G D Q T R E W P A D Q E A P V F N T P W E S R V N L F H V P L A G G N R S P A K Q A A P L E L Y N I P Q L D I
RANDOM_7735 G A E E K I M P L Q P G G V G K E Y S L E C A D N Q D S G D G L T S R E H Y F S S T N L E L C P N F K E Y E S G
RANDOM_7736 E L S A L L T R C A T R F E F D Y S P D V A P L E F E P L A H D L R S L A V L F T H Y Q L E N P L L D V K G K R
RANDOM_7737 H D T T K L I A T I D Q A V P Y P K G N M Q Q I P L G E A W V F A E D T E N K N E S V L R G K V N N K R L A D R
RANDOM_7738 F A L A R I V A L G T V E G L Y S E G Y R Q W C I E T T L V R L Y T P D R L M K H G K Q F K D D G N I S K L L I
RANDOM_7739 A T L A L K T V D S H N S R A V P M L V M I L Q H I A H A H F S D F V T V F I T D Q S D V V E N I S W T T F P G
RANDOM_7740 D A P N A G F A Q L I M H I N R S V A D F Y E V I I G Q E S V P Q V T D D A S K S A V F T T K G L A L E M D D K
RANDOM_7741 L P A P I S N E D H K F R L R E R F A V V V Q L T P T T L M E K G F A S N F A A R C Q S A L E G Y F Y N Q R G I
RANDOM_7742 S G A L H D Q T G M T N G A S G I A K S E S D E M W T M G R K G Q T G D R L V R E G L D G F A Q D K L Q D L Q T
RANDOM_7743 L A P C T E L K R S D M E V G T K L K G P A P K Q C Y R L L E P I S T K K Q P A V S T G Y C C V V D S V P N N E
RANDOM_7744 H N G S K R S F E L D F F Y K G L Y P N H A F G P T R V V Y D E E T V N K R G S E F Q L A V Q E V M F S C L P R
RANDOM_7745 I N V F P R F E E H H E W S S I S N P D I T P G D D A S K S L A I S L A Y S V S A D A M H R I A W N P T D F S S
RANDOM_7746 R M D E E K E H E K I N Q L R Q I K R V C Y N L L E W V D T K N K C L A S A E P S R T S L T E P I W V H L S Q G
RANDOM_7747 Q R T D C R D G K A T Q S M R N N L S G V A I H E A E L S Q V T K G L S S A K G L L V N N F K E L D I Q D L R S
RANDOM_7748 L L V H S L E W G L Y A L Q T G P K V A Q K M G Q S K R P A L A G S N W N D E A H D K E D E Q Y F E A L C F R K
RANDOM_7749 V L G K A G E M H G Y Q L T E S K R L L T G V L F Y I C S G L D V K A M L E G V I Q Q G C S S A V N Y G G S E W
RANDOM_7750 V A P G Q F G S P A M G Q R N T N E T F L S S N A T T G G N D G I K D Q E Q A L P S F D L Q Q I M S T L V Q K E
RANDOM_7751 Y G Q S P L G R G I G F H Q T D V A E T V T G R R G A V A G R T A S E N H H C K Y V A I T H T G K G I V Y A K A
RANDOM_7752 E V L T I G Q M I A E L M S D L F N K L Y K A G V Y I D T P Y A R M D Q V E I K R P T Y T L E G W P T Q R V L K
RANDOM_7753 G K L K N A S G S V N L V G P P A L C Q T K Q A S S Y L T F W H E V D S G N W A I R T R V V S I K E Q S I R E T
RANDOM_7754 V E P Q G W Y V A T P S E S D K R D Q G G K L G Q K Y H P E W P R L T G F L A A S A N D E A L S E I E Q F H H V
RANDOM_7755 T D P P C V A A Q L S A V C P S S L E E V V C V A S D V F S P E P P L N L K V E K L T D Y V V K Y P M T I S D A
RANDOM_7756 L W R N E L W T Y L A S P L N D R I P Q L F S T Q N G L G P K E F T F K L P T Q F A S M L V G T R Y I E R I N L
RANDOM_7757 R L P H T K S G G W F A A M E A K R A L Y I P L F T W N H K A D G R T T P Q T E L E V F V I E V S T S Y V G S G
RANDOM_7758 Q G F G S D S E A Q P E Q V D R G E T L K A Y I V K A H I A W L T T D I W R K Y H N A L S L D V E F Y Y S T S S
RANDOM_7759 P T D T N D K T G K V L V A Q A A T Q L S G L H T S V G Q Y V R R L L V G G D V M V D W D V L Q K G E A E G C T
RANDOM_7760 E C C H R Y L D D V N I V S D Q V M V E P R T L R R R E L A T D C G R P E Q A L F I L V N R C V K F A A I S V I
RANDOM_7761 Y A G L A P N T L N I D Y F C S G P V T V Y I Q A V S T G N Q V L N T G K T L I R G L C H L T F A E R S P L S I
RANDOM_7762 Q P I R N V Q E Q H A I L R S T G A P G W E P G I P W G I L E L P Q V R G Q K S K E G G R L T K G L S G V N D Q
RANDOM_7763 F R G A N D D D S W M P A G E K H P G C E V D N D Q G A P I I G V H S N F A G Y P L T V N A G E G P S L I W V E
RANDOM_7764 K E I G D A V E R D W E E V L R I R M V V K D I A K V K E D K A S D K G G A D E H N K R A Q S P L I I E Q R V K
RANDOM_7765 V L S A A P M M L K H P A L K D S V I C T F R G V T R T G I Q L L G G A G G S T A E P M G P M Q I E I K C G E S
RANDOM_7766 L C I K Y R Q K E N K I R E T T Q Q F I V V E D V T L R D L G A L V N L G G T Y Q Y I S S K R A G N Q R A I Q D
RANDOM_7767 N S K E P I G T T V P M I V R F T N S E K I N F I D A Q T Q V Q M A R V K I A D L G D G E D A F E R I K Q P Q H
RANDOM_7768 C F Y T G I Q P I D A R T K A I A R D T V D A V E F V S I T G A E E A V G N K T Q P P Y I P R Y G A F W L C P V
RANDOM_7769 K H T R S M I A K R K K A S I S L A A A I A P T G C L T I K D T R R V M R L W G G G T K A A V G A Q N N F V P E
RANDOM_7770 P Q W P L F Y E K F V E K Q A G L T E P M A D Q K L P S D E V R V E C V S I T V V N L K H Q E G V R L D R I A L
RANDOM_7771 A G L N N N I E Y E K N L A Y M F L T E R S D F Q S N K L K I F T N E S N M K N K G E T K D I E W M V D G R A G
RANDOM_7772 V P F E P V C G K G A L E I L G K Q A T R A I V S N W R S C M D A G N W E P D D N C S S F A L G K K G E E Q T V
RANDOM_7773 T S F S F D S G G R L Q P S G R S T E T R V V F V D C A L H L G K S A I E L M S L G P S S R V V S A C R N L V D
RANDOM_7774 N G F T A A G Q H A T E F P G E I F V N T G T E K T Q R Q L L R R T L D L D A V N Q K E N L T S C V F A I V N W
RANDOM_7775 L A W M M S K K Y K D V L G P E G S T D G N G G H N T R A H V T S W P D L W K L L S G S N E E N P P V I T F K V

RANDOM_7776 Q E I E N E L K T T D P L L R T V D L Q F P A E R T W T I G E E V I Y S L Q R V H A D L F T R A K L P D V G K L
RANDOM_7777 D G R D K V K P Y A S M L S S A A C T D I N V C F D Q I K V R W E K Q P E L L N G S T E G S P L Q D F D N A Q G
RANDOM_7778 Q F P N S R L E V T L E C W V N Q I T F A Y N C E Q I E N D K T S V D P E T Y N N Q F P N F K T L G S D S A G M
RANDOM_7779 C S S Q M E T G I K P T F Q A A V G K T E E R I G R F V S F G N H F L Y A K N F E T G S R V G I K F E D G S T Y
RANDOM_7780 K S L Y L G E A S T R A Y G Y G H C T F D T T A V H Y G L L P N V N S D L F L V V A F T D L F I T P D K A S S M
RANDOM_7781 H I P D D D A N M E S R W R Q G S Q E Q E I Q R Y Q I Q S T I K L T K N I V K A K A S A Q D G T V D A K D T V F
RANDOM_7782 K G D K R E P N K P E G V L A Q I Q P E V H M R L C H D Q R G C M H L D Y G Q Q A A E G S S G S I S E L H E I Y
RANDOM_7783 H E A A A L S D E C G S I Q D L L A K D H G T Q Q K S R R Y A M N C S R V C I K R L F G L D L A E C I A A V V R
RANDOM_7784 F R E R P G R S I D F V Y L N K I G N A S F R E F Q A A E W I S R K L C K I L T I T P N V P N E T Y Q P L D M N
RANDOM_7785 S D Y G K A D K F S A S P L C S T L N V Y N W N L M Q P G A V G I F E P R T R D K Y R G Q V R S D C E M S T D K
RANDOM_7786 A H N V F V V P L A Y C R A A E V V L V R E R F S E V G Y R A P E E C G K H G K V D M V K C R D P L L W S V P L
RANDOM_7787 D I K A H H R V K E S G T T S F S P V G L M R E T I E L R E N E A E Q P H F S R R T V A Y A R T T E G A T T E H
RANDOM_7788 Y V E I H S V V Y A L L L Q G D K L C S C A A T A M E P S L T V I P G L T T K A F W T N R I P T L P A A I N D D
RANDOM_7789 P Y S M E S R L L S R E S R F V N G S G N T W A A K V P G E G Y L V L S L Q G T L D L I V Y G D L G D P V R F K
RANDOM_7790 L T S G V S Y E V S V I E L Q D A E R K G Q S A T P G D L L T G F S I V T M N P D D F L N P H T V E K S K I T S
RANDOM_7791 Y G K F T I S A P A G M R L E D V I R S K A D L N L D Y N G G K S S E I A P A E V L P Y T R C F M E T A D N H I
RANDOM_7792 T F H Q V S Q N T T D Y Q H A H A V G D Y I L Q A L L D P T T L V P F A T Y D N W P T F G G I T E L R E V G S T
RANDOM_7793 K C P T E D S G P A Q P F T K I R M V V L M L V T M A L E R N R D D I Q E D F D V C K K M P V I E N L P L Y A E
RANDOM_7794 V S T A L Q E D T R D C S K G Y T I Y I E L A I R T A A L D R L Q I T R G S Q A L I A K A Q Q G F E P L S P L A
RANDOM_7795 E A K E L R K L V T G P G N P R C E P D I A K T R K G L V K N V G N I M P I L L P L G N T T R L S L I P L A E S
RANDOM_7796 T L E E A D M I G G S N E S A C V F L M H L L A T K Q F E Q G L A G K D Y T L L L A I H V Q V M N K V I R H T M
RANDOM_7797 P I N S S D V V K C G I K A F R M E C A T R A E D A N V L S N R T P M E G T S I D H E V G S A I I V G L A S K V
RANDOM_7798 Q G L Q K L S K K I G L G A T L T S E M A F P I E D R H S T R Y L W D K N V S L F H F D I V I H E E G M L L D Q
RANDOM_7799 Q L I P L S I A H K A D K I Q F D K D F T Y R E K R S S M S Q A I S A E R K R G A Q L E I D L R E V K S V C P D
RANDOM_7800 L Q S S K E L A F L E D K L R F Y Y H K G V D D M Y V K L E G I P D H A R P Q E Y F C D D Q W G S V F V E T D F
RANDOM_7801 S E L T E S P G M L F P G R P V G A G T V V L P V L V G A W I S C A I A N N S T D F E G F G E I M R L Q N A G N
RANDOM_7802 G Y P Q D R K S V K K R N A T L A F E I M L E L I S D P C N Q T L L G R C G L P N F K P A T V E N S G V A W L Q
RANDOM_7803 A L A R K T Y N T I V L G I A V D K S H G G A A W P K L Y F Q D T Y V E A E F T E R A H T V G K Y G S V L N I N
RANDOM_7804 P H K F K C K A R V F D R E D P Q R A I G L C D T S H S S M M T K V R K G Y E V V D F Q K S S L M T K R Q G E V
RANDOM_7805 G L W T M V F C G M Q E T G I M L V Y G E S Y K V N A N F P N S P A L S N E Q I I V T S Y E T G A I W T G K Q T
RANDOM_7806 T K H T L R E Q S K E A V L F E K Y I K C G T G W I C H D S N D L T E P E L K P W N E G I C N S H G D T G D A T
RANDOM_7807 R V F E S K T H P E A L L T S N I A L Y S L T A Y L S I G F S L V G M I L S F L F D V S N K N A A T E V Q K R R
RANDOM_7808 K D A P Y A D W A M A T G Y L R A E G K A D E V Q L L E F H T L L G L K Q N A G E A K V E L L Q E L V P R T S K
RANDOM_7809 A V G I I S D A F E I A Q Q F E R K F I G S M F V E R N S N I N Q Q Q G K R I D N L L E D C S D G T V L A E S V
RANDOM_7810 G Q N N S Q L F V P V Q P K T V L I Y F T A W T L V L V T Q Y I R D Q I L I I T L Q I R L I L P T S Y L S A I Y
RANDOM_7811 K H E G P T V G E I K A P C Y K D Q E R C L V P G L E A T K L P L W A P D K K N S G I P S R T N P D P V A V T E
RANDOM_7812 V K S G R Y M D I Y Y S K A M K E G H T T E T L T G N N V E L Q L I H I A E H K V V L P Q K R P N N L S K A E I
RANDOM_7813 A V Q G D S Q A D V D R V K S T S G Q V P K E S I C P Q T W P L S G E P K L I R F H L N L V R K L A C V Y N H A
RANDOM_7814 R S S P R S G G K K V S C L H A V G A W E N P H V D P L F A G L E N F E Y A E T S G L I H E P N N D S K R V A L
RANDOM_7815 K T A R D A I K S N T A F L G T P G K A D K I S P A I W Y F C Y I E L Y V Q K D R G E T S T G F P N V L Y G T C
RANDOM_7816 P L M Y Y T S Y P N D I R V R V T F R S G N A A Y G T D E Q F A E G G W M I P Q W S T L C G T G I A V L L R K E
RANDOM_7817 V V N V L N N L Q L K Y H D S L A H I S I G A C G F N P N D L R A T V V V S G A G I V G D F P Y T L G G V S R R
RANDOM_7818 T E T Q A C L Y L I F M K P Q F G D Q V D F G E A L Q V T P G I M S K E I F E L A F F S Q L F T T G P I P G S E
RANDOM_7819 A G M N V Q S N V N K Y V Q G K L V T V T Y S Y P L V L A P A M L S S S I Y A D F D L V E L F K A V I E D R D E
RANDOM_7820 V A W D F S A L A H L G A A G I G N F M D Y G P V A M E S S M N K D D E R P A E G F R S G A S N Q Q G K G N S V
RANDOM_7821 K E S C Q T T E C S E E P K E R R N L M C G E S L D S V A L L Y E Y L G I N T L A Q V Q V D H L D L I A G T V Q
RANDOM_7822 D D N R A Q M S M T Q N R V T G P K K K D V R V S G E N G S D C T A C S Q D L Q A P R M I P F D D I G A D Q V E
RANDOM_7823 L D R P G G L E S D G L Y Y E N V Y D E R V R K T L R S V S V C R A M G D R K A M T C G S S K G K V A Q N N R A
RANDOM_7824 S R Q C L T P N H E G V M K K L V M A D C I S A D A M G T N K L A M M A S L R P K G S L N T F L L C K V I Q Y E
RANDOM_7825 R K G D L I V G Y F F V S Q I V I S M T V C E S T W I P K C E K K R A L Q C A K P G Y T N R E L D D N F E M I Q
RANDOM_7826 A G K T V I G S Q G Q R L Q K L P L H N F S I A N S Q G S P H S V I G V L Q Y Q N P A Q P Q S R N L L R I S H T
RANDOM_7827 A S L P H E F K W A L A K V A L Y R V L R Q S G A E P V I R Q S S I K N F Y V Q K L N V T H R V F L R L N Q F S
RANDOM_7828 D S Y G L K K N A L G F K T S Y R L D P D R A P L D P G D T V L Y H G M M W G E N W S D E N P F T P L K N H A K
RANDOM_7829 A V I I T L A A E Q T F Q T P Y R Q T R K I E G L I Q F Q L C W G W H E L M N A G T L D L G C G D E V Y T Q S S
RANDOM_7830 Q C M R A F E S G M Y I N A A I E P Y G A F L Y N T G D Q L W T P A A P T Q G L S D R Q A L T R L A K V L E P I
RANDOM_7831 G D L Y S S L L R L K Q K A F T I A T D V E C L D P R N I G G G F L M F G G T D E Q F R N K I D P L Q N V N G G
RANDOM_7832 A G G P I T H L M D L I Q G W N V L R W E R S P V V V V G Q K D L I Q A N D T L R N D N T E A F S Q F R R Y Y L
RANDOM_7833 A L I R P E V I L A D E K I S E D Q P E F P S F Y R T P Q R P P H S L I L G D Q N E E G L H L I I E E V N K D T
RANDOM_7834 A N M Q E V A E E V A R Q P K R I S G L R E L L F L I L K G G Y I S L G G P F A A V V K E K L R W M P T A G M I
RANDOM_7835 V T D A R A S M L N Q A K W E V T Q E L L H V S K F I S T Y I V A P E N R T N P A E S P L F H A S D L S K I A S
RANDOM_7836 F N A T P S R S D E Y N A H P K K F N P T M G K P A Q A T S W P W K P S E G T A L Y E M S L V D P K A A S E P M
RANDOM_7837 T G L P V A T M V G S F L V G S E P D G H V A K T D T S S S A E F R N T F S G F L E D S D P Q P D V E K V G Y Y
RANDOM_7838 H Y E V G Q S V T V L V P E D L T A R R T L C M Y P G R W A P S W G S G D E R Q E K L F S L D L D W Y E R G L L
RANDOM_7839 S E I K G S L R L S H L V A G E A D G R A V Y Y V N V T P P S L W T P S K A G G R S F L K L I T C S D L K E S A I
RANDOM_7840 A K V E Q A V H N S L L S E Q E A D P S Q I I Q T L L T N L Y E R A P K K T T D L Q F G L D V R P V E M T K A A
RANDOM_7841 A R N T L N G C V P V L F R C S S I S K D R L V I P K P G A L L Q T A E K E K P I L E A V C L I L T S Y D L G E
RANDOM_7842 A A L H D S R V G T A G D I F A N A I N R F I L M A N N G L E N Y S L S K K N Y F C L I F E I F H H F W G A S V
RANDOM_7843 H V P G K G G P I I P G G K Q D T P F L L D K K K V G K C S Y T H K A H A I K K D P F T V H P D P A F D H D L V
RANDOM_7844 P H V D D I G T K F F I Y V D N K V S I N A S K C E L W L T G T V S G Q K L W G R L A E G A L V P F V R F I P W
RANDOM_7845 D C D I Y S W C M Y L Y R L D E T M G E P I T Y T L G E Q R P N H E E A T R T I Y L F R E R S G D N I F L L V D
RANDOM_7846 Q A S D D G V F S T A W W I A V G A A S P D N P R P S L P Y R L E I Y I D I G D G E T S C F L G A A D Q H Q S P
RANDOM_7847 G G Q L A R H K S V R A A R R I L L H H D V G C K C E R L G S Y L G L G V V D T G D D K I G S N D D R A F H C E
RANDOM_7848 M G H F A L K T L S C A A F P E Q G D F K F V P N A G T W G Y Q I G L T V L W F A Q G E R D L R I S R S K I V V
RANDOM_7849 G T Q T H G A P V A K R A A V A P V N V D L A K S F V A M A R Q N G G D Y A S K T T L L K Q L S K P V E G A A F
RANDOM_7850 A S S R V A K K V R V F L L G I L M G S I K I Q L A Q I P C G M M S F S S A S G V V D A P S L W L C S L H E G A
RANDOM_7851 L V S E W C G T A L L G P L K K D G Y R K G H K M A H Q C T A D I D K Q L T P A S T P S D L N S V L H R R L I E
RANDOM_7852 Q C T L D T R A N L T P A Q G H R E K A A T L H A R A E D K K F Q L D A K Q L E C Q D E Q R E E Q R S A A G G V
RANDOM_7853 A A L V S N V I T L P N I P Q A L A S H R D R H L G F G G G H S N F P L N S G R Q G T W T L P R F A L G S V S L
RANDOM_7854 K Q V P G E G G Q S K N L R I I S G M A Y I I I L E I R L E Q T L T H G T Q I A G E Y D P K L E C E E I L C F A
RANDOM_7855 T P L K P E L V K V S S M G E V D P M K A F K T R Q R P K D F K S P G I P H G A G N T N F L G R E Y T D H F H L
RANDOM_7856 D G N K G A R Q A Y R A K E F W H H G R A Q R Q F F A R P P Q C W E T R P N P T V I W V T V M L V V T D N I K G
RANDOM_7857 A C Y Q V M Q G K E T Q L V L R P L T M H G A Y N T A I S G M W G T N L R G K L V M L Q E E C L R P S T K L G H
RANDOM_7858 P T N E V G W F A K T S H L L G G F I V F L P P K D K N T M N R G A N L Y D W N F I V K V A I F E I S R S F E D
RANDOM_7859 V L C W L G I T G M D S M T L K G L K L A F R S V Q K N P K P S L Q R A D W F N R R R T D I Q E V K Y P L P E V
RANDOM_7860 E Y R P F P T K V L S F R D S M L F L I Y G A Y Q N E N Q G D G T Y L M I S E T A N G R L K H G A S A I V L S P
RANDOM_7861 P N L T T R N R A I E T M D F E M V D A E A V S E R S D Q E G F C F V V M T D S L V T G I D T K L T T E N V D V
RANDOM_7862 E Y M Q L F R P D T A Q F K E R L K T R T Q T V R W D A Y R V Y Q K D S D C C S Q S D I Q P G H M F Y K K I T G
RANDOM_7863 W L F D R Y R G L Q S Q S V L L D Y T K D N H V R P L R V Y E S T E P S L A T F S F L S S T K G M L Q I V S H P
RANDOM_7864 V G I K I L E K L Y P L D Q V R R N K A V G L S I G K P Y W S T T G V L D Q G I E K T Q T W A N Q I R N T N H A
RANDOM_7865 A V K D S A K Y S K R L L K G H L N K V L E N N R L R R D G L P F L I I W R I S K G N L A R H Q A S I V R F L V
RANDOM_7866 T E K C T A Y L P R G E A K I K L Y A S E P G K S T V F G H A I T G G E R R L V P G N S P V P P H V M A D L Q F
RANDOM_7867 N P Q F I F D G Y Y Y G K G K A T K P A I E T D D V N L A E P K D V V D P E G Y K E N Q V G I C H N A S G G C R
RANDOM_7868 K R L C F N L K Q T E N I E N Y I N F D F I P I S L K D Y W T P R S A A L W K F T W T Q Q C I E K P K M D F F G
RANDOM_7869 W P Q K F P S C A L F Q N D A M R W P T I R G P Q I T K K S R F A E K L D N Q A G A N A D E L E G Q L K G W I A
RANDOM_7870 V N R D I H V Q G H H A Q A R A Y F D F E L E S P P E T A I D Q E E L V F P F E P P T Q N C V C I V C Q V S K T

RANDOM_7871 A Q R S Q S T L A D L K E T P N T Q Y T K S L E A V G N L A L E V D P P A T P E Q P V D E A D L N A E E I K K Q
RANDOM_7872 G G D T S T T A Y K S M E M H E K C F G S I G I L G A L P K K R A K D G A D E A M I V N T Q R K K K N V V G H L
RANDOM_7873 E K I Y T I Y E A L E K Y A S D L V Y R R N K P T F N R G M I P I D T S Q R A K D G M S N K T L L H A T A A V V
RANDOM_7874 M S L V L A K H L G L P E A L A P T T G L G G L I L I S R D A T A A I E E A E D D V A A T A E S G D V I R A G A
RANDOM_7875 L E F R A S H I Q M A S S V S D L N H A A I A N R S L K I T T I A A I E T E P L Y A G V V F C R A A K S V N D Q
RANDOM_7876 A D I A V T L E M S C L G G E L N L L N G R F E I R A Q C V T V G M G F V S T D S V E S L H S Y D A P Y E P H V
RANDOM_7877 V I P L S A Q C N L C Y R D M V D K M P Q R N L P V S A T E S I D T Q R N L I R E E G I E A I A G I W C A Q T A
RANDOM_7878 K G R I L K N L P R V M G L A Q L I Q T G L G A F Q Y A Q E N F L W V F C G I K E A E P D A L A S T D A L R T E
RANDOM_7879 I W R T V K L E D V H D K F L E S F N G A L C A D L N K K N E G L I G E R Q K P H S S M P G K G A G Y I I Y E L
RANDOM_7880 V T V F E E E W A D D E G Q C I N N R S R F S E L N D K G E H S E T P C K S T L F D K I P E A K Q A V L P T L Y
RANDOM_7881 G L G G I T P H Y S F F A G S T I C N P Y I T R P Y P Q D L I G E E Y H K A Q P E M K I D T L P P I A S G S E I
RANDOM_7882 S S P G V C D G I T Q N A T I A G E D G V L E Y R A N V M L H F L N I P P L G W N A I A I F T T L V V V L R E L
RANDOM_7883 M I L G D A Y A C T W K V T T K L E G S E Q A L G T M P V A K P Q D I F P L S A I C R P K N E P D L G H N A T L
RANDOM_7884 A M H I L L S R V L S E E E A L P P T E G I L E R A L E T F Y K N R I V D L N D M L E F Y G F R G A A P S M S L
RANDOM_7885 P N K H D E E R P F M D L G C P A W L S T H P R P S A F D T D G M V R S K P A W W A L K S H S H E T D A L G G N
RANDOM_7886 N G W T S K Y G L T G E Y P Y T K L S D G P G N Q Q T R E G A Q V T V D H K N L A G C V S F F R L L Q L Y A A T
RANDOM_7887 S S P G R Q Q A E C K G K I S L S T L A Q G W S F D V R T E K M S P M M A V V K H A D Y R E P Q D I S K T E G L
RANDOM_7888 L L L P R S W I P E N I N T V S A N G A A G S F G N V D A T N I G H V L V N I P H E V T N R F V G W T T R G L Q
RANDOM_7889 S L P D A P N E K A N T L V I L F S I K T R G N Q I K R N A A G H S R K S Q Q E A A P F N L A E Q L M L P N E A
RANDOM_7890 T D R N K L T S H L A A D I R K Q V N Y N M N E A V D F A E A T V N T A L N D L S S A K R K N Q P K L Q S T G Q
RANDOM_7891 T G Q E F L T S A F W Q F T I D E P S L S F Y N R Y D R W K G S S T Y V T R E P C D P D D L I E G Y G G A L R C
RANDOM_7892 T L V D Y Y A A K A L L D D Q V E I T S E Q V S F K W S K R Q K Q L A G R T R G E K V W M K G R E K Q Y P H D I
RANDOM_7893 V W I V E E N L R G P P V G G P L V Y G Q E E S Y V D S F T R G I R K S L I A A A F I G S M E G A Y S I T L I S
RANDOM_7894 L V A N P S V R L A M R S V I A V A E A Q I L S A M C L G A D S M R S G S L G F R K E R T A A L V P D A N Q G S
RANDOM_7895 D Y I R A Y V E Q T A W E K T T V N Q Q K N P P L K A E G L Q T Q S T A S E Y Q F D M Q T I T R E R I P P E A H
RANDOM_7896 A W R G D Y A V S E I E S P N R R E E T R W C S I L D K V R D S L Y I A N P K V A R A A K C A E E H D L R R D T
RANDOM_7897 K T P C L A A D P R E P I D Y A S Q A A Y F N N L F L E K K T V L S V V Q V Q A P Y E T E A H Q N I E Y R I A T
RANDOM_7898 V S F S T C L L T M S I R V T T A N T F I S F S I A I G R K V S P S A I K G G Q W F R S S T Q C L N D E A V F A
RANDOM_7899 E S H K V V F K A T F G I A T F P E I H Y V L Q G H H N S Q L C L Q P T G E Q N F M L I G A V P I A E K L A C V
RANDOM_7900 E T K K A F F A D D I D G E S I V R F L P F A V K N K Q G T P E I A D G Y K I R A S K D S S E V I L K D L K A A
RANDOM_7901 L I T S L P L T G A F A W E K S K K T L V L P S E D A G C A P N G T F K N S P L R M N T F D N I A A Q E F E D I
RANDOM_7902 D P M L L I E Y A I L E T A S T N S S R G G S D F L G K L G R M Q A E F K T D C A G W L G F V C W D A R G Y N P
RANDOM_7903 G L F S R L D G R S I P F F P N P G Q L T I G P F Q E F K A G V L T S V L P F C I G P C L V L H I M E L R R L K
RANDOM_7904 T F Q E I P G I P L L L P K E G P E V S P P D K I E F K D V I S E V S E V Q L L Q S G F P E A V G L N C K L D E
RANDOM_7905 G T C L E I S I F D A D P H E R K T G A R E T C E Y S E L P A I Q T G M P S F T H Y D S I G F P E N G Q S E L L
RANDOM_7906 M L T N M P N S H Q N D G A H I P Q P G E F Y L R S S I N L P Y V L Y K D G S K S E V N N T P A N A K D G E P N
RANDOM_7907 L A D N R S T R G D G T A Y V Q G L W A E A V G N V V G A N D T S V N S V V G G S Q P A Y V G P P S F E N A L V
RANDOM_7908 D T Y G D P T N S R I Y R G C K T P S K S A T E E R D Y G D P K T T A A E N G L F I G R L T A G K D Q Q E E T F
RANDOM_7909 L K M G S S Q F N G H V V L K A P V P P M E T I D Q K K V E I N E V K V K T S D V R V A D G N L K T W P N L L T
RANDOM_7910 K P P W V V S S A A R P T K A A V T E G M S L A F R P K G G D I V L A L M R L A N Q N M T I I G K L R N L E G S
RANDOM_7911 K K E H N S E G Y A Q N A Y L R F T R Q T E F S T I L N L G D L K E E N P D I A V A E T P S V L G T R S H I G K
RANDOM_7912 L E L A A R S Y A L P T Y R A H K Q L F D E Y S A I L K C G A R P F L L A V G G V G V G N V N G Y L D P F Q G Y
RANDOM_7913 A R T F E K N H V K F N N G N V R I N Q T D D E Q I K K Y D K Q C G R G E L Q V L T Y R K S T L T F C L Q K K P
RANDOM_7914 S T Y A R C S Y V L F T A V S P A T Q I G Y M A K K I D L R S I T Y R R S F T V T H T E Y L C Y Y G P W Y V S R
RANDOM_7915 G H F T P D A E I P R T K S T G D G A R T F F D I L L E W V Y G T T A L W V F K P S Q F G D L A H Q R S C L V K
RANDOM_7916 M A E N T A A P D P T P K S G V A T Q R E A A A V C V E K V D K S S D L T V Q R Q L G S N S R S Q A C T F R P A
RANDOM_7917 I V T V I G V T L N P W I V E G I L K E P P P S L K A I C G T P F G T R L I A Q E G A K S L A N C C M Q T K L F
RANDOM_7918 P F L N M S L L T T N N L G E K L F T G L Y F I W A G V V D A H P R G F G L A G A S Q Q G H Q S C S A G Y S K K
RANDOM_7919 Y P E I L K L W V L V I L E C I L V D T N E S L Y T G K F F M P K A L K A Y V Q V R F D S R Y D K D F L L S S S
RANDOM_7920 L D A F G G L L A F D I T K I E W A R L H D A V F R W L T E D P E S A A T S R N R H L S L Q N I Q R P K C L E D
RANDOM_7921 P E I F K L A F S T E E V C D D R S A L K N F G Q N E S Y T T T T A N T K W H I N E L P Y P S G V H S R A A G L
RANDOM_7922 M R D W D V E L G L T R L D E P V L F A L E F V D I D C T S I I T G R Y T E K C K A E F N Q S Y I I T G W K S Y
RANDOM_7923 Q I N D D Y H K K N D P D T M A V N L L D Q V P S L V V L H L I S A N N Q P T R G S Q E S D S Q R T V F T T N V
RANDOM_7924 D E C Y N A N P A Q Q I D W T Q K E R F G A P P E N C G T K L E F G V T Q I D G P H L L T L H Q D H Y T V D L L
RANDOM_7925 T E S Q L A P F M L M S W I P L K A S T S A Y C Q H I D K L Y A R L Y I A L A T L M I Y P L D L D Q C S Y G E D
RANDOM_7926 Y R G G A A G V D A E Q T K D T V G E K S L V R T S K S V A L G L I V T Y H A K V T L C T T A L L L V Q S T N L
RANDOM_7927 G I E S S G V E G K S I T P L N E P M K K K E A D P I T L L S A V S N K T H K N S A E A D M Y S G A I E Y W A N
RANDOM_7928 Y I V R A R K E V R R G S L T L Y V L N P A N G E K A L Q R G P Y S I V L G Q H A D P V I R N L A P A D V E P G
RANDOM_7929 G I L A I L T S L Y S I G G I E D M E R L K P E L A V I A D R L K P R W A A R D K H Y R G I K R T K V E S P G S
RANDOM_7930 P L E S K G A K I V T G V S R S P I E K S A K W K S E G S F R E F A C M Q S L K S N P K I Q V V R A D V N T A A
RANDOM_7931 T C F K V D I K L A I A I N A T T N C L H F E T E V T E L F R I Q L V S G V A L G E I I Q E A V M T A G K H E Q
RANDOM_7932 D V T V P F T R D Y P E L T V M I K Q I S V H Y M G E L K D S W S L T Q M N N N R V E T C E R V K A P K L F F I
RANDOM_7933 E A E S D G R I G G K E L V Y D N E F E S N Q E T K A N L S V N K T K W D A D E D Q E R G Q S F G V N V E L D E
RANDOM_7934 A F P P R A G H L W E A V H R S C R N L A S N H Q S A E A L T G S A D P G A E E L L E E I G A R P S G A A M S V
RANDOM_7935 K A V G T S A S Y S C Y I P W Y Q V F S I V G N D M W L A P L Y V M F G G V V W K I H N R R V D G K F V A V V P
RANDOM_7936 F V C I A L N G M E Y W G F K G K G M K I K N A Q P D T Y H V A V E P L A T R T L Y E S G Q N V P F Q T P D P E
RANDOM_7937 N T T I V H V C L V K I E S I E E S H S S M L R S E A R T S I I I D R S K S C L E K E L I G Q D D D S L G T I Q
RANDOM_7938 I F T E E I Q D K K K S G Q V P N F F L T N L G G E Y K L L T K T R K L D G E G V D F I H K Q K K K S Q F P T I
RANDOM_7939 P F L Y P K D E A L A W L K S Y V L A N C H Y R A S M P I T S S F S I C V L G H R C A N L N R E F A D V I E F F
RANDOM_7940 R E I R P R Q M I Q S N H I F S V E G P E N D L G S P F F F L E A N E C E A T Q Y A L D G H P Y L N A I Q K A V
RANDOM_7941 T A D L I M G L D T R S I L G A K T R L G A G T A S A L K V A I A Y W S E M M F G K Q H F D I S S S Q H Q N D L
RANDOM_7942 L G I L N Y H K I L C V C K V I D V M Q L P I D I K G G P T M A A G L T V G G S K S F K G Q S A T E I P L S E E
RANDOM_7943 L N S G S F D K H F R I D G G T K A K R L M V E Y I M G S E V K L T L F Q V A T G A S F T E Q C A Y W L S A G D
RANDOM_7944 G Y S T R W S V V A D L A P F M G T S D I T A L A E I P E A A G P T S D G K I G M A E P T L L K T N Y N S E T V
RANDOM_7945 A V N L R P I V K F M T N A P K I Q I V L L E G G K N D T E V C L P M G N A D G H N T L L N D A A Q T H T P K A
RANDOM_7946 E G A P P G A T T V S G T K G V K R Q A Y K Y S Y P L L R R F L T P G F I G T L I A T L Q E V E A P A P N T A L
RANDOM_7947 G A G T V K K T L P T A H T P T L Y G H S Q F P I C L L S K D G G C S L L T G F G V S A H I K G I T D S Y D N D
RANDOM_7948 R G S S V Q K T L A N L A A I F V S L L T L Y L F K F T L E Y A E P M H N H K V P Y D Q F T I P E L E K V L A K
RANDOM_7949 K E N V F M S S S Y V G C L G R V K D F P N I H M G K Q D L S T R A V N N T G G Q T N D K L K L L C M I E L G E
RANDOM_7950 D F A L G C Y F L V T H V V S A P S K Y G T R D A F F T V E F L H S P G K L S G G A S P G R L D A Q V A A P E S
RANDOM_7951 A S I E G A I P A Y V P P L K E Q I M T A Q I S K L I G R T V G D V C Y D H R S I T H F P K W G P S G D K A S A
RANDOM_7952 Q A V G S D L F R I P V S I R H V P T G R K A H V V A P G E N K G Q V R S L G M F G V R P L L L E W G P K T G A
RANDOM_7953 K I Y V H G P L E F G N C Q E G G C S A N F T I G G H G S G T S E M T S R D N G T N Q P S F L I A S F S H G Y I
RANDOM_7954 K L V Y Q K M A G N T D K N A K Y P A F L A E K G N G S L Q P S S F L F S S V S D S C T L Y I I R T E E N N E R
RANDOM_7955 G L A P T K T D S K L P I I F N G Y V K G V E L R I A S V G C T E R L L V L C K E E S A E N E A C P A S G R L T
RANDOM_7956 D K P R S V S S P D H A A N G L P S T Q A P G G S R T K E L A R I S K L F L H E T W G M M R P D C P I E E P E I
RANDOM_7957 E D G D W V C M G Q A S T I L A N G T Q W T P S Q I S T L A Q D E T F Q V E A A L A Q N I L R I T D S I I T W R
RANDOM_7958 S E L K D Q K S P Q I E E Q W A G N V L N N W R T A P R N R A S V M F D N D L R A N I M M R P C L H V Y E D S A
RANDOM_7959 R I K Y D G E V E T P N L A L S A V W E R E E G S D P S R P P L H Y H T G Y D R C Q P T C S P L T A S Q D F T G
RANDOM_7960 M V V L L V A N G T V N A S S T Y Y G N I N H A R W K F S E A K A D L V L F I Y D L V T K G D L V G S R M I K L
RANDOM_7961 W P L T E E K S F S V L K F M P Y V E T N P N G D K V V Q H K V K G G A T C G G V S N G P R M N K R Y M G K S N
RANDOM_7962 K A T G L G T Q L K A A A H P K K D K L D M K E L T P D H S E L A E A L G V V L L A M V V T E N P K Q C M V A V
RANDOM_7963 A I A N F R N D Q R A D F S A G V G R P V R Y V S T S P K S I M V N L V N A R Y N S T G I L Q T K G F Q Q V N C
RANDOM_7964 V S G R P K I K S W N W A A V T A Q I V N A N E A S P L P I R D K K Y E A A K P Y A I E P E N T Q R A N G M R D
RANDOM_7965 L A V L T D I P Q Q V I T L V G E G E G M Q A K M D A L E T S T Q L Y T K V K V S F Y Y D L E V T V S T N V L L

RANDOM_7966 A G A K T Q S G M E C V D S D R A Y G G K A E K L Y P A L H T I A I Q R N L P V N H N Q S E S T E A V F A A Y V
RANDOM_7967 I R E G I D E V A V C F N R N K P K P S K P M S K L D E G V P N N A W Y K M K F K G L N E P V L Y P S I G R A Y
RANDOM_7968 K V I D A T K K A K T G Q C E G F D D S E R R R T V A G M E H N W V D D L C V Y A G C N D G Q E K A Y G T A P M
RANDOM_7969 D G D M W S L G Q A L N Y G V Q L I Q G G H G P V F F N L K E D Y V Y Y V F G W S A L N A P P E N L G N S T C G
RANDOM_7970 R K Q A K V R L Q T K S P C H V S H K P F V W Q Q N L E S A T D Y P I A G M K S K Q V G V S K R S L L I L G N D
RANDOM_7971 G R S W A V K Q Y K G K Y G L Y R K L C A I S P P P E Q T F P L D K R H A V W V K F N G R L R K F L E T Q I G F
RANDOM_7972 L S T E P V N L F R V C S L E D P R T F L L N H K L L N T D Y L V N D L S T D D K K P L I R V D Y D P I G L A A
RANDOM_7973 F S L N K N G E Y K Q G R D K M I V L H N V L R N T M E P R I V V R F L P R L L A Q P K T Y L I A S Q L S P S V
RANDOM_7974 E E D I S P G E Q A P T S E R E L M I Q Q D E T V L Y P L T G Q Y I Y E G A L E S N E I L L E I N T G S G E N F
RANDOM_7975 E K Y K K A W G S N E C A G A C Q L E S S C R A H A D E L V C I G V V G D I G G L S N G A R E P T V V F V Y S D
RANDOM_7976 G P S A T V D K G D T E P G H F P I E R D V V T Q E P L N D L E I P K G L A E T L K V I K E M I S H G W K L T N
RANDOM_7977 V F D D L R I E I I S D Y K W V L K G A L V S G G W G V L A P G F E R Q Q N P T K R G L L P K E I A T A L F R L
RANDOM_7978 W E G A A L Q K R V E I A P I G S C Q E L A R M S C A Y P E S E I A E E L V N D C Y A K V G S P K K F N G P P G
RANDOM_7979 T L V F P K D A F S L W S I L D E A N E N D K G D R N Q L A E E D V Q G G T G K K I L G S S P Q H V V S A K K K
RANDOM_7980 Q R E H H K G W D S V Y L I A L R A I C L T Y S M L E V T S G L A F R P E A S E N I G L T G A F G L K G L L S L
RANDOM_7981 L A D G L E A S S Q P T V T R E C N G A I R P G P Y D T L H F C S S P S D A I G Y V L T T P G M Q L S V L Y E K
RANDOM_7982 G L F Q V I S E L L L M V P F P S V Q F N P E V M L K G T M T M S I N Q E A L R M V S F L E A C G D E T K W K D
RANDOM_7983 I D R F P A K T F A V G T E S N A K E P G F E K I W V F E D V P G K G F I A H V E F V S M G P S A S S G G A L T
RANDOM_7984 K Y T S I M A L E V N N H L Y Q E T I V V F P L V T R M E Q F D P A M V I G R P V V P V D R S L T A G S K L Y L
RANDOM_7985 L A M R L T A K A T R G T D F P D Y D E T V T S Q L C D A G V N N N M E G A S D P D C S F P T T A L A R G V D T
RANDOM_7986 Y S A P P T A D H D R T S Q S V E K F A K A G F E R A S T K S S S V I G F W D I Q A D E E G H S L A S Y T L I A
RANDOM_7987 Y L I R P V H V L I A R L N L W Q Q N N I K F M G T G N I G I R Q A K G Q R K M P L W R S F L R R S Q H K D F C
RANDOM_7988 N T R T R S L I T V H L I F F Q S K I D F L T S L S K L K L K T A G R A I I L T W A Y H I N I P E D L K S G N Q
RANDOM_7989 I T T N T I A T A S N P D R C N K C W P E G L N T D G G N D W A A E S D G E T A E F E F L Y L N R F T K M Y V K
RANDOM_7990 I R E A Q G I I Y M D E C K T S A M G H C E G I L N I L Q D E V L R M G N L F G L F V A H I P G I G A N T R D N
RANDOM_7991 G E N A I D L F W F S F F L Q G L I K C V T F S H L D C I G F V L L S K R P R T P F A D P N D V A L Y R I W G E
RANDOM_7992 L V N S M S A I K I Y V A R E I S R D A H R G M Q A V S T D E G T I K I I Q P A N M S S Y N E G L H L Q K P L L
RANDOM_7993 D P L S L N L L A F V G E E I L Q P A Y T G A T D L I M E F P L I L E K D V N T L C V G F V I M L D D L I E G C
RANDOM_7994 I K L T K A F I L K A S G Q I M D Y R N D R T N R S G N F T P H L I V I I M E V E A I Y D T L V P N L I Q G R L
RANDOM_7995 V A V T L V C P A S E T G T S M Q G V L A Q L Y G P G L S L N D D Q D K S S G G L C C L A W L L P R L N N S I T
RANDOM_7996 G K R K G D I L F C I G K L M Q G V E I L Q G F D K H K E D L I L D L R S L D E L S I I I F S K F A I W S S T I
RANDOM_7997 G P L E P I L E T D P L K T L L G F T G N S P T V K D Y H I R V P P S C I A E L A K R T R E A A K A G I L P Y Q
RANDOM_7998 I Q D T I P F T I A L A E E L S T S D G A S L E L F K P P W W T V L L A L L L S M A S P N R Q N G H G V E Y D G
RANDOM_7999 G G Q G S S S R K F L F E M L G K S D P S N V P V A S V V T A S D C L F E S A E L Q V F L C F L I Q P Y R T L V
RANDOM_8000 G F E R F L W R K A N L R N W G M D V V N D G I E P E F N L A A E R S E G F Q G F L G T K E K H F I W I Q A K T
RANDOM_8001 G A K S D F P H D K K V K V N K Q G A T I A G E L R Q V K T N G D P S G E L R H W V F Q E V L Y A L A P T T D P
RANDOM_8002 L A L S Y L L I P K F R S Q A F F E R D T K T V G V D I I G L A D G I Q L K A Q S A N V E L P I K E Q A H F D N
RANDOM_8003 G P N F L P E I L E S T A F G Q D L L A V G P C T V Q T A P Q H Y N P T T K P G I G V L R D D K V Q D L L I P I
RANDOM_8004 M T N G T Q R N F Y H T K R G E R T I Q V L L L G Y F R K D S Y E S A L A Q R N I S A M R I K G I K S R V S H P
RANDOM_8005 F S F L A V Y I L F A R S N E G L F A L T A C L V T G Q H F D T A G L V E V V D I E K G I R R I P E D L L G M A
RANDOM_8006 Y S T N Q C K G S S A H V T T Y S K R P L E R K E F G V Q K E Y T S W S A T S Q P V I V I P N N L T L T L F T A
RANDOM_8007 L L L N V T G A F V L A H R F E L T V L M N Q A S Y E P Q P M K R G L M A N E H E A S S F T S N Q T W M K S M F
RANDOM_8008 L D L S S V S A F Q A N E L V N D E T T L V N D K Q T L R S Y E G Q Q E M P T L D S S A S F T E Q N A V I K L V
RANDOM_8009 G A G T R R Q G Q Q L R K D L Q E D N I Y D V R G V A V E D G R E N E V I K M I H P S Y H S Q Q I F S V D E P D
RANDOM_8010 V N R Q P P G S D K K K D K R Y N P V V G G L V P A R G G V A G T A H K N E N E F K S A I P C L D G E A F K A F
RANDOM_8011 G H L F N N G T A S A A E F L Y M K K P L G S G S P V Q A P V G E P E I W E K R G Y F P H A D D Q S P L S Y C T
RANDOM_8012 G Q C S Q L E N C S N F H Q S E D L R A T Q T M G V S R W I K I E A K L Q T D I G S D G R D T C N T T H C I S E
RANDOM_8013 M I G F A Q E Q S D R M G K L K V V Y F H W T L T W C M I M A R G K I V F V L V Y Q N K R N K T L Y P P A K V A
RANDOM_8014 I V W T Q I E A E E T K L H Y S D G T L A V D K G I G E S D D V V G A V T C I N S D F W S A G K I I N N T L D L
RANDOM_8015 L W A S Q L E I I G S R L D Y R F R I D Q K L G V A S L R L P E A V N D A Y Q P Q S N P L C E D P S A A K P R Y
RANDOM_8016 G V Q V E H P M N S G I Y E H F P T S D T N Q D E C E T G I P K I D K H N R I A K E T I P T S K L D R L W L K A
RANDOM_8017 A A H K L I H I V S N S K S Q A V G E Q T I A V L C R G K L D N M L E A G R G L V G K D G V P T V A K A R R V F
RANDOM_8018 I S K A C A I L L E L S I T Q G V H P R K L K Q S E S Y G S M L I I N K S W E E T E R V N A T D E H G F N S G R
RANDOM_8019 W L G K S E Q L P L E A V P S E A A N T V G V Y E V G N S S D R M L E G A E L M L Y T T P P P K H G I N S H S V
RANDOM_8020 G D I V K T F A R G D A A Y T F A R Y L V E S E E D G G V P R Q K R N D S G K L T A L F A D P K S C D F G D I D
RANDOM_8021 V R A G S I G A G Q S F M K V I K G I K T K K D N D Q F R W G G M M R C R F G Y K R N R L F G C L L I A R E R D
RANDOM_8022 T L D L A Y D E L I Y H Y R I Y D I V L E R L G Q Y S I R D Q P A L N M R G A L L S R P A L R D S I N A S C A D
RANDOM_8023 P D D S S D P K E T L G Y F N G A V Q Q T T L R A R A G S F A V T G F G G Y C L G R E L K S T V Q W L L H I N F
RANDOM_8024 I L G A S A Y L V G A L G P S N V K A K S I A N H D C V A I R G I E Q Y W L D D V K V L S I I G L G R A N E I Q
RANDOM_8025 E V T L A T F L A E Y H F V T F Q I T D D N I Q L V V N R E R L L G R E F C L R A G Q L G A A R Q D L N A G S A
RANDOM_8026 S D V A T K S F F L V G L A P A A L G T Q I E E T G A G S E M K N N T S V E K C F I G T G L V A S A K W S T G F
RANDOM_8027 V E T K R D S G A I L K N R S H P A S Y K F L K E Q G G I E D L G G P Y R G E I D L S K E I S M V A L L F I I S
RANDOM_8028 D T V I P G L L T Y Y G R N R K G L K T L S G A L K T W P G A Q V C F R G V M A M Y L E I Q N K A S P P G T W Y
RANDOM_8029 T R K T Y E V L R Y C L V N R Q T L N C Q G A G A K I D L L E I K H F S T D A A I N Q D P S F T M V P P T Y S T
RANDOM_8030 T S L G T N E E Y A L H T V K S M C L K F W N R K N D N S I V G R G L L K A H P S G R P P A K P N N G S G H V D
RANDOM_8031 R L P L P A K E V A Y Y R K L G L S A P F I I S G Q S F Q L D G E R G F L F F E N K E P S Y K I G H L D N E G E
RANDOM_8032 A L V G P A A A R Q G D E K L G L T Y I A C S K T R G K H H A A F R L K H V S G F V R Q V V N N L K F E G P L N
RANDOM_8033 G V K D A E G A P N K S L F I G F S G F S G G F S I V T P L K A A H S F N T L E G L S G E T L V Q T V I S A A A
RANDOM_8034 P L A Y H F A S P G A K S G D K I N L R N V Y Y L C P M R I G G K F G C F G L G D A C A F S S D G P M A K M K M
RANDOM_8035 G K Q V I G A T K A T K Q Y N E G V I E K A N T A D L E P R I L I E A S K P L K P A A C P S S S G I F M A P S A
RANDOM_8036 N F D T A K V S Q S P I N N D V A D A Y S Y T F T C R G S S K R I L W A A V N R P T L G P D M Y D C R W S V T I
RANDOM_8037 M Q A E I E D N L R D G S G N G D S T G K N G A Y R F Y D I E K H D L D S D K Y P E Q H C H G S K R E K S L M A
RANDOM_8038 L G H D L K N L M S S Y Q A I I V R F K I W A S E T A I R K F P N A I A T E P T C W I K T K I K F I S F P L C G
RANDOM_8039 D R C A I K G D D D V L E D A R T A D I I P C G R M I K I S G L L L N D P R F S D G T N M G V Q E G Q H I L D A
RANDOM_8040 L F M G D L P G G K A A N I C R T A R L E N A Q T C E D I T P E R Q V S D A K T D G L G D K E G A A F S A R Q Y
RANDOM_8041 P E K M K I A H V V N F Y M E L R L K L E G R Q S K V H L S E F S K S Y N K K A G N E I L G A V V E S V S F I V
RANDOM_8042 K L I G K E L F A L K D N T V G Q S I Y I N G L L A F G R G P S N E V E S W Q A G F T T G H A C A S V F P K H T
RANDOM_8043 Q E Q R S H S K F E L H I V P M C T V L D S L Q N L V A L E D H T W G E K H N G L L R P S S G D F A L L D K K L
RANDOM_8044 K Y S E S N L M I G V P S A A A Y G K K F P K G V D T L P E V V A D R A C Q F K I K E R Q A L D W E T G D N E K
RANDOM_8045 Q R L I T S Q V D K R Y I I N K V S L R P L L Q G T G Y G A K K V S R L T K G I E H N F D R G P D Q A A V D T K
RANDOM_8046 A E D C K F R T E E G I A F L K T L F R K I S A T R M G V N L N N G Y L A T L K L H R L K G G L G L R D C K Q P
RANDOM_8047 F T R H M N S E Q S A S E S N R G F D L Q K P D L N V L G I A F V S I T V I Y K L P E A A G A Q K I G W K F L V
RANDOM_8048 L S K I K K E I N V G L E L V E V G Q K L T P V L T L K A T P L Q E S V R A T E F F I D S Y K Y K L L H L D E L
RANDOM_8049 I T I L S R A G N K L K L E M R L Y S D T Q K T P E C I P R R P D K V V T V E P E T K I L A L A D S P L Y P N S
RANDOM_8050 A Q K W C K H A K E L A L E P G R I S K A M T A E N S E D S T V R L S R S Y R L V K L A S C T C G I D Y M L E Y
RANDOM_8051 P I N N M G E L Y R V A Y T V K M V T H E A K Y R T D T D E T S C S T C D G L L G G D I T V I G K G E I V A F F
RANDOM_8052 N N F R P L E P M D G M E Q S I K L S T H Q P C V L S D L E A D D L A P P K V N N L P S T Q G R I D L L Y G H K
RANDOM_8053 V T L V S H P F M L L I P T H N K T G K L A P N V G V K S C K A W P A G K I V E M L G A M Q Y R T G L E K K F R
RANDOM_8054 R A R R S V I A L I P Y T Q F A N V A A F F R V G E S E S V C I P G R E D I A S L F T N V V Y W I A G L A A N L
RANDOM_8055 H T F E I M L V H C E Q V V D F V E V K G I G P I F T I E M V F G A S Y G L K V F G N A N M T G G S I G R L F V
RANDOM_8056 I K A I Y W D C A L V S V Q D K S K W D V T K T V K K Q G D A V G P Q A P Y Q S K P D L K P L D E G V Q K E G C
RANDOM_8057 A D F G S S R L H D D A T L Q F E T L P Y W S L L I T L L N A L L K M N K D L L N G C L H T D S E D Q I S L L Q
RANDOM_8058 G C L G P F A G M T I Q G E A R Y P G L K R N P G L D E L Q D C C P R N E A D I R H I Q M G V A Y A E A T Q G R
RANDOM_8059 Q S L K A E V E L P Q S E V C V P G N G A T K V W A A D Y R E Y E F Q E E C I G M C S L T Y A T L T Q K V A E V
RANDOM_8060 A A S E R S G H N F N A L Y T E S A G C D G I H R D L E Q P G W A Q T G M R L L V R G R R R F M W V P N A G N V

RANDOM_8061 P G R A L R L E G I N T W E Q P G N A V N A F Y L Q K I G N F P Q V E R F C N S K S T I I D H G E L T R A V A G
RANDOM_8062 L Q Q S I F K G S S K N K A D V F P Y E I I S V G E Y K V I R P A R T E A H R A H K A N S I F R C N D S H K D K
RANDOM_8063 H L A H E G W L V Q S V V T V S D G S S R G I H S R S E C L M G T T K S G S A A T M S G K L L T R E N I G A P P
RANDOM_8064 V R Q D P L T T P K E E S R L G F P I M S H W L F Y S D E V R S G V A A A S N A K G I T L E H Q K T R Q L L W W
RANDOM_8065 H N A Q K W A D I V L V T T W V R I Q L E R I P D I C A E D L R L P E G M A I G I I G E I V R S F T V S P G L K
RANDOM_8066 Y L V Q V N R E R V A D A R G Q S G N H Q D V T W I Y D S V I I F V V A F Q K L A N A N F H G R V S I T T Q G E
RANDOM_8067 D F C C L L S D P K S G A D G R Q A S W G V N E A L V G C P S I P V G K E T K V F S G L V D E R Y K M G K A L V
RANDOM_8068 K D S M C I R E A F T A L R T N T N E R V Q H D P T A K T K Y Y N Q A L G F Q E R V A F I L L D P T Y Q V F L L
RANDOM_8069 E E I Y G A S D S S Q I G P E A D R Y P T L P K A G T Q D G M Q F R K L A F E I S V L P L S P P L N K R L E D G
RANDOM_8070 F V D A S K L Y T D A L D E N M H W T E D Q D D I Y A L T S R Q N Q G V L A A R Y L L G E I D D E M D P P I T S
RANDOM_8071 I N S K L T M P K N K E P P F M F A T Y G R G W G T V M R L Q Y L L E E H V D F L D G H C Y S V F S C T S L K A
RANDOM_8072 P E P Y S G G R E I G A S C A C G E V A T E L L S K R H A M L G R T P L G V R K E P C D P I K N K F Q T R S L F
RANDOM_8073 Q T Y K T A S L A G V G A D R V F M G L E H E L R S P H S D K L G I I L D S M D F A E Q K V P H A L Q I K L V D
RANDOM_8074 R Y L S Q F Y F S K N W Q D I V A I M V S V P D G G L V S G H A A Q T S A V T T A E Q F I E D M F G H L L P Y L
RANDOM_8075 C P M G A V L H S I W M E Q A E Y P D V L G K L A K P S A E V L E F P V V V R D D N Q S T C W N D I N L P L Q L
RANDOM_8076 V T L N K G Q A H D G P K Q D C E G D A L N V T L L V L A C C S H G C P W S R T L N V C H L Y Y C A V S K K P I
RANDOM_8077 K G G H A L R C M T G D S N K T V P C E D K S K G L G S E A N G I A H S L F I R A A E V A M K F K N E I V R N W
RANDOM_8078 R A V A T V A T T M A T F Q L H I V S V K V S R G K I G M V L A L E Q R A C R L A I A G E A T R E W I Q T G T V
RANDOM_8079 V E G Y L D T L A A A F T G I S A A S S I K K M D T A R L S W Y D D Y K W L G G E L F S E F P R T F R V F N A F
RANDOM_8080 A E L F L A K C E T T L N I D K A K P K F K Y Q L R P L T T G T V P F K A D L S T G A Q S V S S K F Y N Q M E A
RANDOM_8081 A P V D G H Q H R K N V S I E L Q L T P N K T T L L W A E A N H V N R T P A E Q A Q A G K L E L V G E S I Q T L
RANDOM_8082 P G M L Y I V R E K G R T M G G F Q P K G R C A Q G P T Y I D M E N Q H G E R R D L Q G N L Q C K A K L R T L S
RANDOM_8083 S S M L E L S I G K G E K I F G P N G I Q S P A N G I Y S S Y V T E V N G I D S R D E S G M L Q V N L K V Q H L
RANDOM_8084 Q S F V G S Y T N I G K Y Q P D T Y Y E D E L F R T G A H A E D L N D E K G V E Q Y K L K E H T T L A V K V Q D
RANDOM_8085 Y A Q V N A G E D T S L T L G S R V H N E D I K C R G L L A R K L P D C L D D Q G G L P F A A N A W N V S H N R
RANDOM_8086 R G V S T N F T A T S I P L L K A R E V N D F Y R F L N K K E S A T E K A V R C Q L M H I V L E Q Q I D V S Y G
RANDOM_8087 T R I K A F N K V I R F F A V D G E Q V E F S Q V G T P T G L A L P P L T Q L R A F N R V I V A Q G P E A C I I
RANDOM_8088 I S V S A R F G P P L Y W S G I M Y G Q F A V T Y A M E T S G C M L I T F L L D I E A I S R K N C K F S H K V A
RANDOM_8089 I E V L L Q A V W R V S A C T S T F A P K V S S L D G S G N D N Q S T T F K R V E G F D R N T G E V V N A T L S
RANDOM_8090 T V E G D L E G Q T S K G N V S T L S F M G S Q E T N G G E P R A E A Y F G C P G G K A V G S G F S F I L N K N
RANDOM_8091 S S V T V V T N I C E T G G D N I L S F L E A Y I V M N M F A P S Q S S E N Q V P N A V E R Q E W L E F K V S V
RANDOM_8092 S A Y S K K D G H A L P S L S M T I Y P K L E D V V E L Q V G K N P L E A L V S N T D R V C I G A I V S L I N L
RANDOM_8093 D D P S H I S S P K A G Q G P V P K S G V M S I V R G C Q N G P A E T Y D A T K P G A L Q S G V K T Q P T A G S
RANDOM_8094 R V K L A Y V F Q K L M N R G Q A G N K T G L R P F R K Q K N Y G T H R F T K G G L R N D P I Q F L Y F V E A T
RANDOM_8095 N W L D T S S F I Q Q F V Q V E S L L S H G D A Y Y E D Q A W W G R V I A I E T K I L C M G V P K Q S M V N S L
RANDOM_8096 K A T S Y V L N I D T R R T H S F G V M N T V Q P E K F K P T P G H F Q S T V C A F V Q E M V S N I G L K E P C
RANDOM_8097 E D S Q F N T T A G Y L I Q V E E F L G S M E F H N T E G I G H S V L T E E Q S I R H D E V N R P S R F Y A M K
RANDOM_8098 G S V D A P W S R P P P H N F T A V A T F A C D L G N L R A K P P F A D E Q S E Y F I D S M M N E L P I T G G G
RANDOM_8099 L M L Q M R Y N D K E G S A T L S V L T L G S D N L A P G L L I I G S I G Y W S P K A G S C T A G Q G R T L H T
RANDOM_8100 A V Q V R P I F N C S D A G S C G T N T D E E P Q A I L V A Q E L P A I K D D L N K G D S G N Q K L A Q L Y K L
RANDOM_8101 Y P L M L F Y L N R R R G L L Q D H Q Y L I A P P A R L C I V V T G F P G K G K I S E L C S A G Y V A P L E D L
RANDOM_8102 Q D L L L D S A D I L G S R I D E A S P A S S F L I Q L S Y V E G S D N F L R L V R A P K P F C A F P G I T E N
RANDOM_8103 Q Y M V I T F R N A R G F G G Q Y G P P E T G M N E Q T N D L W L N T I T V L V N Y D G I E D D S L V R V L T A
RANDOM_8104 F A C K Y L L T T D L D I E K Q A G A F H R E N L R L L E S E K D H R D A L S S M N L T T S I L V V T K F K I I
RANDOM_8105 N K L T N R E L G A A M A T M K V G R G T K C I E K K C L A A D P V Q G A H D Q P V Y I C L Q L L A E W K S P D
RANDOM_8106 K A N I G I N N M P R L V L T G I S E T G G G E E M Q T K M C K P A R V S N G G D H L S V E D T Q S A R N D S I
RANDOM_8107 C Q A T L G N L I G R I S R H R C I S K E T Q M G P E S S V V R Y P A E Q S P E F G I E Y A S E G L L L L T T T
RANDOM_8108 M L Y A N P Q V V H T S E N S P K V G N R K D D T A T R I T M S D T L G R P C F Q K C S L G V L V A A K E P L V
RANDOM_8109 T R S S V N A L L A L K N G L A H G E V I G S T R P Q V L W I L F L E K R K A D A A E S P E A V T A A P F I E N
RANDOM_8110 T G F L A L R P S Y I N E D V F V N D D L P Q P K V T N S S Q Q W N V E T E S R S L M E H C F P V E E F D P I H
RANDOM_8111 E N N D L D F L N K A I Q D F A K G V G A K L K S P L P F W C C R N R K Y K V G L Y K R G M L V L N H T N S L D
RANDOM_8112 Q K F V D A M F R G V C R R G D M G E D L M T H V V A A P S D L K I T N L D I A I A D I C G E P F M V N S F P T
RANDOM_8113 H Q T R D R T P L A N S C A P E A Q R L I G K T Q R V I E Y Y A Y V M Q H T N T A K K A D K A E S L L D T T W P
RANDOM_8114 N S Y K T R R S C Y V C F R F G R D A K A L G M H L I K R V P E S S L Y N I K M Y G G W V A M E Q K L S G N E S
RANDOM_8115 T S A N K W L N S I A G V V T G E I A V T C G S L Y V P E N L P F S K E A I H P I R K L V V Q T L L V S I S T I
RANDOM_8116 N R L H L A R K P F V Q S E N H S P Q A Y T Q K L S D E R Q Y L P K Q R V C N E N L L P G S F E G G T S G Q Q K
RANDOM_8117 K L E E S P F S K Q E D G T V I K D E L T Q K Y D M G A E L R Y D S V V V E L S N D R D V S Y F F N D T A L S L
RANDOM_8118 S T A C V L D K D K E K T K E G A T G K T G R A G S N M Q H T T N G P I G P G Y T G F L Y T F H E Q N V T Q S P
RANDOM_8119 E G E Y D A T G K E V K E L T G V Q G E D S D F Q E A Q T N E P E Q A L F T I L R G E Q V T L Y D S L S E R F D
RANDOM_8120 G W K D L P P H D E A Y C L K L W F Y A N A D R H M V E H L L D H K D K V G T P D A I K A S E R S T G N P V I L
RANDOM_8121 T A I L V E L Y L G I A L Y G E V N P T R A S T S G M F A D R K A I L I L Y S I S D S T E K L L N A P N A G F S
RANDOM_8122 D R K L Q D E K P P I C T A A V C T K D D S F I M C E P V H M L K R L N Y W K D A P L M S K N R F D G Q R Q I I
RANDOM_8123 E G P N M Y A I R W E L E A I T T P G N E A N N G V K C S D T T K D P S N K N F R R V N K S T A S K S L R T N P
RANDOM_8124 T Y A K Y R N V C L Y P M P S D L A R V T K T T L H K E V A T H Q P P K R L Y G Q D E G H S V A P K D I A S T G
RANDOM_8125 L A E L W S G T G F A E G H F L V N I P G T V A P D E M I Q Y V P A F P H L D L V E T E P E E H L S S D V P S K
RANDOM_8126 Y A V L K L A M T A N Y R P C E Y K A V G M S I Q L P R T A N V M L A F L L M P A A K D F Q G C S K E K Q S T E
RANDOM_8127 N V T S Q E S V K E S W I S E P F E I Y G L P E G F V V F L T G N V K Y E K N R L T K A H K Q A N V A Y R A E G
RANDOM_8128 E I T Y Q L T L R K A T P S A G I L G R E Y S E A D G A S R V A Q A R A E V A F T Q Q R S A V A T P E S F D G K
RANDOM_8129 F G L V G I G I G S Q L S V T R D Y D G V I N D R K Y A A S C G D W I T F G L Q L E S D L E G L T T N K G V L A
RANDOM_8130 D G L M M S L A N V V R L R A I T S N E E K L A A I K A G L S Q L R I S D L A I L A G E A L T R A G S Q I K I F
RANDOM_8131 S L N L Q K A N P T R T T E A D G A D R E N L Q L H N G A L W Y D N L N L L K I Y V M E L S L P D C G K Q D T H
RANDOM_8132 G G Y D Y I Y K V D S L I P S H K D N T G G Q A A Y V C D V P N P V F H S K A G A F T P S Y S V V L T I P F I R
RANDOM_8133 P N Q V P K H S K P E N L A F L A T F R G R G Q T F V T F A E G W T E D K S P L E C L T T A V M L K G G H E H A
RANDOM_8134 L K D G S V D S W F W T F D R E P I A A C I I A K F Q S T L K P A E E T G H L E S S V F Q G R N D K I V S Q S Y
RANDOM_8135 G A P V T F G R A K E L L Q N N R P L H L N E G A D S L K Q K E W D R G Y T A V I K D D I I L V L K F G C L K S
RANDOM_8136 L R S Q V N K Q F E P G M D M A R A L A G V A H Y K K L I I S N L H S A A T Q T G D G R P G G V H S N F P K F S
RANDOM_8137 G K V I N V L H D M C I F D L I K F I L L I L A S Y L L H E T T Y Q P Y A D D P V L R E T R R Y A F A T S L N W
RANDOM_8138 T T R N E W L A I A Y R N S A V V F E A M G G L V K P Y K H G Y H T P A E S L R R A A G E D W N V V G V F R L E
RANDOM_8139 K D A S A D D R P D I P H R G A S L L G Q Y G C K I S V Q L Q C S S N G R C R F E I A I C F G G L Y P T N E A R
RANDOM_8140 G K E I A C P G D T L Y N L S W L F N I E K L G T I N R K T T I L L H M Q Y A G L D L F N Q G Q A A S G L Q L S
RANDOM_8141 K R L K K S I T G K V T I V A R A H N E K D A I L I D E N V N W R S L A N V G G N M D A I G T P S I A N S L L A
RANDOM_8142 N A V F L K S E D D L G T L L L F R S T F T A I R Q G G G G S V M V E Q K A V L L N E Q A L M P E R A K G L Q A
RANDOM_8143 L K N E N F K A C F E L A R K T H K H L E T S C K V Y G A V C R Q E I D Y S I E E F E A T A K V S S P P P C N T
RANDOM_8144 I L F P I I N M P L A P D V W I F I K P W D D K T T N L P Q S P K S L Y E M L V K T A L A S K G D Q Y W K T S L
RANDOM_8145 R I R L I T Q V F A T V A F I P A T V G A Q R A P H K G G S M P A K I I L G T D G K F W I P D G T H E A E G T T
RANDOM_8146 F L T H D L R V R V I E S Q C S N T Q A I V D L M K D N P Y D R R F S L V T A E A A Y T E S A H Q S P N H L E I
RANDOM_8147 Y S Q P G C A K D K L A L D G N F A R D E K A Q P A N E R Q E V D A W Q F G E R D A W K T T H Y V S Q V V I L P
RANDOM_8148 G R C L V S G G P E F S V P G W L L S Q K L A L A R Q A S F L K E E T Q G E D G A E Q L S G R L L P W M G L G V
RANDOM_8149 F E D E E P E K L E E A D E Q A Q Q N E N S E H F E L N Q R M M G T C E G L V I V E R Q V T E W D F R F E S S D
RANDOM_8150 T P D S E V Y Y G A G S L S N I I W S N R D G T I P E A G G A L V I N D V F Y I W S G I A T G R M Q R Y D R P L
RANDOM_8151 V Q P F S A G R P G G E P R V Q A S L E G P A M V D R E D I H E G E I A G R M A A R G C V K G S V R R V Y S V L
RANDOM_8152 V W P P N L F Q S D P P E D V S N M V L G L G I S I P L F R C D M R E T V H I S C R L P L D K S H T I P F P Y W
RANDOM_8153 V K S R I S A S V D H L E S V I R F S D I F H R G T K P V S I D K D T D L D A T S Y Q F A D D A T Q L V R T P L
RANDOM_8154 S G E P K P D T V Y F D P E Y V P S L V W D Q N P W S F G K V P Y M K Q L R T T V Q Y L T S L D E E P P Y E D C
RANDOM_8155 N S D E F D R S E T H F D I Q C E W T L L P A G Q P G A G A D V L V V W S G P Y W A T K E L H S L S G A K N A I

RANDOM_8156 D L I D S N L D G E G K G L G L T K L T V S V W F E G H Y T E V H G K T K E V D G A S A H R P T N S N I D T V L
RANDOM_8157 S P G D S T L V K A K A G E R H P D N G V V F L P A T K M M F A T E E K P Y T V Y E T G L L G N T R L D I L A L
RANDOM_8158 K N V N K V A S W S P K S L I Y I V E D N K C G K K F S M S Y E Q G D W L G E G T K I H D I L A K V S P Q E Q L
RANDOM_8159 L L N K S A D T S S R L G L S I G L A L L A A L G Y P K A S M W R H L L H P P T V T S Q N D T A G G S N S D K G
RANDOM_8160 L R T D G F V C S V H V N M D L G R C Q S C A E F K L G N D A D S V T Y R K G F W V G E E S L W A T V Q A A T Y
RANDOM_8161 D I R R R Y A S Y G S Q W K G S I T D I D I T V R E W N I Q L L K L F Q V V A T T P G A S R Y Y D T D V K T I G
RANDOM_8162 R A Y G R D R K I T G F L L N R K M K A A W G G L I R A T G R R R V T R L P V P V S W A G P D F V Y D T T P I T
RANDOM_8163 Q Q T Y M D A Y L R V G V V G G T D Q F A G G E M K T T H L D G G F Y C G I T K I A I H L S R E Y A G R R L I N
RANDOM_8164 F R A G Q G P V I L E E V N G T M K T K V Q G P E A S L Q M D Y G T K D S N Y A Q E K R T L Q E I E F H H N S S
RANDOM_8165 L K D R K A A V G D W C I L T T K L R F L W V Q P T E S H F I Y T V S L D V Q R G R E E H G R Q V A Q S Y T C A
RANDOM_8166 L G W G L E D F S R G A K F P V D R I P F G S A L R L P Q P N E G G S D S D Y I R A A T Y K D F Q I I I P G T D
RANDOM_8167 L A R T I L V K A G T K G I E G I G F K D Y E A R A G E E G S L D R K N P D E T L R G P A A N E A A T L S K I L
RANDOM_8168 E T V L T L L Q T M G F L R A G R T K D R A G L F L D K K R T D E A G E L Q K W A A D N T C F E P Q M A D D T V
RANDOM_8169 S R R I P K G C T L C T Q T Q K N H L I E S T K C L W V L W S I C S I M R A I P L A S V S P S T S Q F Y I A L A
RANDOM_8170 L P L L A G S T R D S K E M S I G E L Q Q P K D A H V N L L E N L Q I V Q I R N T A A D A D H P I V I F T V K A
RANDOM_8171 A L T G E S L W S L Q R D S P E F G D S P P G T G F M G L N K A S V S A A I I G A A F E P Q F V S E A S Q L D S
RANDOM_8172 Y S L G F H G M L A N T T L R D E S W C S V G T F R I G D G S V T A I T L G V S D N L N G D V V K D G L S L W N
RANDOM_8173 Q I G M Q K E T K V S E Q V G Q T L T N R E T S N I D T A G R R G Q Q D D A R G D L Y V L Y V K P Q W K G S F A
RANDOM_8174 V A T T G V E K Q E G S K Q K L K G V V P L T L M G S H L K A K G G D L A T T I E T M A A A F Q S I K A N V P D
RANDOM_8175 I R S L V G S S F Q F A R P A G E M D T S G I N K S S A D A S K A K V Y R F Q K A N M L A M H D Q L W V Y F I R
RANDOM_8176 V R A L H E D I K Q R T I A Q R R L E L F D V L S D G W A V V K G E N E S R V L L G Y R G V N D S V G K A G T A
RANDOM_8177 E R S E T Y R D K G Q N Q N R G F Y I V P R K E W D V K T S Q Q L G Q L N F L H N K T R S F G K V T D L D L A M
RANDOM_8178 S F M D L L Y S M L S I L R T I V R G H R L V L G Q K V I G T D Q A T A G V Q A E K Q T R W G R A Q E E S G Q D
RANDOM_8179 Y L R K I P E D T R V P Q A V E D L E D Q D N D M V K E D A A E V P E A S P A G H F S R T A K Q Q N G L K E A T
RANDOM_8180 G L E L Q K S A N P S L L I L L R V M L E C T Q R W G V S Q T E G A C A P E L Y A F L H V N K N R D K E Q S V D
RANDOM_8181 D G N R P A P L S G L R E T I S V T N D R V W V S K W A L Q R A D I E P L V G K L P I F N E R P Y A G H N G V R
RANDOM_8182 N A S W V K G V M L R V S G V T L D P M L I Q K N A P P A T N I Y F C Y K F P P D H M K F D W N E T L L K L D I
RANDOM_8183 Q C T N P S I F M G V L L R G S A L T T L G A Q L N K C F A V D F E H A Y Q T L Q F D L G E G G D E N V V E R R
RANDOM_8184 L C G V M Y Q A W Y P C V E P K C K L I Y D Q G P T A S F K A L R V L I L P A V F S Q V I V S G R E G P H K S T
RANDOM_8185 V C T V C F T Q P P G Q L T V W G M W E F I L A A M G M A E E S K V W K I A L A Q V Y R E F K V L Q F V S R G P
RANDOM_8186 T L T T N T I P K K S S A E N W S S L T R Q G T G P L N K A T A R T Q P P M C I N I L D G E H L P S A V N G T R
RANDOM_8187 L P F S F A R E S G Y A E L K S R G K K S Q V S D L L T D N I H N A K F I D N R L A Q A I E R G S V D P A M G T
RANDOM_8188 L R A G I G Q Q F D R P P E K N A T V L Q L K L A H I G S E G K D K V H S V R E I G L D F I P L K S G R E E P Q
RANDOM_8189 N V L Q L T P E H E C S G H T T R S A S A R R R I M G T D S S N L R F T R G V V K S P L K Q I T A P D E L P S L
RANDOM_8190 F K G Q D G A G P D I N A C A P E E A T N A R S P Q N P R K G G D K D Q E K Y T T Q D T S R L T V E R F D T R V
RANDOM_8191 T T A K P T T S R N A G C T G L E R Q Y V G L W L L D K S V V E S E M G Y N G E D Q P I P I A H M Q R A Y I T K
RANDOM_8192 L T E D S D T E S G S P K S V I W I P L K N K L Y K V R D S P C E H G V R T T I E T Y A S A Q K H Q I V D G G A
